# Shared Neural Dynamics of Facial Expression Processing

**DOI:** 10.1101/2024.08.05.606517

**Authors:** Madeline Molly Ely, Géza Gergely Ambrus

**Author notes:** **Corresponding Author**: GGA. **Conflict of Interest** : The authors declare that they have no known competing financial interests or personal relationships that could have appeared to influence the work reported in this paper. **Funding information**: This research received no specific grant from any funding agency in the public, commercial, or not-for-profit sectors.

## Abstract

The ability to recognize and interpret facial expressions is fundamental to human social cognition, enabling navigation of complex interpersonal interactions and understanding of others’ emotional states. The extent to which neural patterns associated with facial expression processing are shared between observers remains unexplored, and no study has yet examined the neural dynamics specific to different emotional expressions. Additionally, the neural processing dynamics of facial attributes such as sex and identity in relation to facial expressions have not been thoroughly investigated.

In this study, we investigated the shared neural dynamics of emotional face processing using an explicit facial emotion recognition task, where participants made two-alternative forced choice (2AFC) decisions on the displayed emotion. Our data-driven approach employed cross-participant multivariate classification and representational dissimilarity analysis on EEG data. The results demonstrate that EEG signals can effectively decode the sex, emotional expression, and identity of face stimuli across different stimuli and participants, indicating shared neural codes for facial expression processing.

Multivariate classification analyses revealed that sex is decoded first, followed by identity, and then emotion. Emotional expressions (angry, happy, sad) were decoded earlier when contrasted with neutral expressions. While identity and sex information were modulated by image-level stimulus features, the effects of emotion were independent of visual image properties. Importantly, our findings suggest enhanced processing of face identity and sex for emotional expressions, particularly for angry faces and, to a lesser extent, happy faces.

## Introduction

The ability to recognize and interpret facial expressions is a fundamental aspect of human social cognition, enabling us to navigate complex interpersonal interactions and understand the emotional states of others. While the neural mechanisms underlying facial expression recognition have been extensively studied, our understanding of how the brain simultaneously processes facial expressions along with other forms of information, such as gender and identity, remains limited. This is particularly crucial given that the integration of identity and facial expression information is essential for social judgments and interactions.

The primary objective of this study is to characterize the shared processing dynamics related to the perception of facial emotions and their interaction with face sex and identity information. Using data-driven multivariate cross-classification and representational similarity analyses of electroencephalographic data, we explored the stimulus- and participant-independent neural dynamics associated with these attributes as observers evaluated expressive faces of unfamiliar identities.

In recent decades, neuroimaging studies have significantly advanced our understanding of the neural mechanisms underlying facial emotion processing. Much of the research in face perception has concentrated on distinguishing between the changeable and dynamic aspects versus the stable and invariant aspects of face perception (Bernstein et al., 2018). Identity and sex characteristics are embedded in the consistent features of the face, defined by the underlying bone structure (Mello-Gentil & Souza-Mello, 2022), proportions, and skin texture (González-Álvarez & Sos-Peña, 2022), which remain relatively stable over time. In contrast, emotional expression is changeable, manifested through the dynamic movements of facial muscles, and can vary with an individual’s emotional state and reactions to external stimuli (Zebrowitz, 2011).

While early models propose distinct pathways for face-emotion processing (Bruce & Young, 1986), a substantial body of research indicates that the processing of facial identity and facial expression interact at various stages of the information processing sequence. Indeed, through the examination of the statistical properties of face images themselves using principal component analysis (PCA), certain principal components appear to encode identity, some encode expression, while others encode both dimensions concurrently. Notably, shape changes within the facial features may serve as a coding mechanism for either characteristic (Calder et al., 2001). This suggests that the dissociation between identity and expression at the image level might be partial rather than absolute (Calder & Young, 2005; Young & Bruce, 2011).

### Functional neuroanatomy

Haxby, Hoffman, and Gobbini (2000) proposed that the dissociation between facial identity and facial expression originates from the utilization of distinct brain structures within a core face network. These structures are involved in processing the visuo-structural properties of these two facial characteristics. According to this account, subsequent to receiving input from the inferior occipital gyrus, the fusiform gyrus handles the processing of the invariant (non-changeable) properties crucial for coding facial identity. Simultaneously, the superior temporal sulcus (STS) is responsible for processing changeable aspects of the face, such as facial expressions. Following this initial processing stage, further cognitive processing unfolds within the extended face network. For example, facial identity information undergoes additional processing in structures located in the anterior and medial temporal lobe, while facial expression information is channeled into the amygdala and limbic system for further analysis.

It has become increasingly evident that the distinction between facial identity and facial expression processing at the neural level is more nuanced than initially perceived. Cumulative evidence suggests that both the fusiform face area (FFA) and the STS structures play reciprocal roles in expression processing. Notably, these two areas may extract different types of information concerning facial expressions. The FFA’s response to facial expression appears to encompass a broad sensitivity to shape information, while the posterior STS-face area (pSTS-FA) may specifically respond to face shapes conveying emotional information (Duchaine & Yovel, 2015). For instance, the FFA, traditionally associated with face identity computation, also demonstrates involvement in expression processing. Conversely, the STS, typically associated with biological motion processing, may also play a role in identity processing, particularly for dynamic stimuli. In this capacity, it is thought to integrate identity information transmitted by form and expression information transmitted by motion (Dobs et al., 2018).

This interaction challenges the traditional separation and hierarchy between facial identity and expression processing, indicating more integrated and interconnected neural processes governing the perception of facial information. This perspective aligns with the demands of everyday social interactions, where the ability to attribute social meaning relies on tracking changes in expression and gaze direction across individuals, even as their invariant features (identity) remain constant.

### Behavioral studies

From an evolutionary perspective, prioritizing the detection, processing, and interpretation of emotional cues likely conferred adaptive advantages, such as anticipating threats through the recognition of angry expressions or signaling prosocial intentions and fostering trust with happy expressions (Feldmann-Wüstefeld et al., 2011; Hager & Ekman, 1979). However, ongoing scientific discourse surrounds how these emotional expressions are most effectively processed and prioritized, and how their processing interacts with other cognitive mechanisms such as attention and memory. Employing similar experimental paradigms and stimuli, previous studies have produced conflicting outcomes.

A wide variety of investigations propose that angry faces are prioritized for processing, more readily capture attention, and are remembered more, indicative of an anger superiority effect. There also exists a growing body of evidence that aligns with the presence of a happiness superiority effect, positing that happy faces are attended to longer, their processing is prioritized, and more accurately recalled at a later time point. Zsidó et al. (2021) found that both children and adults identified happy faces faster than angry and fearful ones, regardless of the age of the faces. Similarly, Halamová and colleagues (2023) observed longer fixation durations for happy faces, followed by angry faces, in a faces-in-a-crowd study. Fixations on contemptuous and sad faces were similar to those on neutral faces. Švegar et al. (2013) reported higher accuracy and faster reaction times for happy expressions in a change detection paradigm. Additionally, several studies have shown a happy face advantage for future memory (D’Argembeau & Van der Linden, 2007; Liu et al., 2014; Shimamura et al., 2006). Conversely, angry face superiority effects were observed in visual search tasks (Dixson et al., 2022; Horstmann & Bauland, 2006), Changes in angry faces are detected more efficiently than those in happy faces (Lyyra et al., 2014). Additionally, research exists that shows an angry face advantage for visual short-term memory (Jackson et al., 2009, 2014).

### Neural dynamics

The temporal dynamics of the processing of these features is also a matter of interest, and Multivariate Pattern Analysis (MVPA) using M/EEG continues to contribute significantly to the understanding of the representational dynamics of face processing. The preliminary phases of visual category-specific discrimination, distinguishing between faces and non-face stimuli, unfold remarkably early, manifesting before or at approximately 100 milliseconds after stimulus onset (T. A. Carlson et al., 2013; Kaneshiro et al., 2015; Klink et al., 2023). Face-sex information has been reported to emerge at ca. 70 ms after stimulus onset, and identity at around 70 - 90 ms (Dobs et al., 2019; Nemrodov et al., 2016). Face-familiarity information has been observed to emerge in the ca. 200 to 400 ms (Ambrus, 2024; Dalski et al., 2023; Dalski, Kovács, & Ambrus, 2022) and 400 to 600 ms (Dobs et al., 2019; C. Li et al., 2022) time windows.

Recently, multivariate pattern analysis studies have begun to investigate the temporal dynamics and the network of brain regions involved in the perception and processing of facial expressions.

Smith and Smith (2019), using EEG multivariate classification, investigated how task (implicit and explicit categorization, i.e., task congruency) modulates the neural representations of face-identity and expression. The study was conducted on 15 participants who were asked to categorize facial expressions or identities (happy, sad, fearful, disgusted, angry, and surprised) displayed by 3 male and 3 female models. The study found that task context affects the neural processing of face identity more than that of expression, with identity being better decoded under explicit conditions. Both the effects of facial expression and identity peaked within time-windows centered around 90-170 ms over posterior electrodes. The task context affected the decoding of identity at early (pre-200 ms) stages but not the decoding of expression. At later (>350 ms) stages, both face categories were better decoded under explicit conditions. The authors interpreted the independence of early effects in emotion decoding with Muukkonen et al. (2020) presented color images of four identities, two male and two female, with angry, happy, fearful, and neutral expressions, to 17 participants, who underwent separate EEG and fMRI measurements. Both the original stimuli and morphs of identity and expression between stimuli were presented, with participants performing a gender identification task. Decoding and representational similarity analysis results revealed that emotion information is available already at around 100 ms, spreading from occipital to temporal areas within the first 100 to 250 ms. Signals associated with happy faces peaked earlier than those linked to angry or fearful faces.

In Li et al. (2022), EEG representational similarity analysis was conducted with 20 participants viewing images of eight identities (half young, half old, half male, half female) displaying fearful, happy, and neutral expressions. The images were grayscale, normalized for contrast, brightness, and spatial frequency, and cropped to exclude external features using an oval mask. Participants performed a 1-back task during the experiment. The processing of emotion began before the extraction of identity, starting at approximately 120 ms. Identity decoding occurred considerably later, around 235 ms. Importantly, the study found no significant effect of sex. The authors attributed this to the preprocessing of the stimuli, where the ears and hair were cropped from the facial images, leading to the loss of gender-specific cues.

In Zhang et al. (2023), 20 participants viewed luminance and contrast-adjusted grayscale images, cropped to the inner features, of eight male and female faces displaying happy, sad, angry, disgusted, fearful, and neutral expressions while performing an orthogonal fixation-cross color change detection task. Source-localized MEG patterns from the lateral occipital cortex (LO), fusiform gyrus (FG), inferior partial cortex (IP), and posterior superior temporal sulcus (pSTS) were subjected to time-resolved representational similarity analysis. Their results revealed that all regions they investigated differentiated between neutral and expressive facial expressions in the ca. 100 to 150 ms time window, with the LO and IP representing categorical, rather than image-level information as early as ca. 100 ms.

### The current study

While previous studies have explored the temporal dynamics of facial expression processing using M/EEG MVPA (Y. Li et al., 2022; Muukkonen et al., 2020; Smith & Smith, 2019), the extent to which neural patterns associated with facial expression processing are shared between observers remains uninvestigated. In past studies we explored the generalizability of the neural signals for familiarity for faces (Dalski et al., 2023; Dalski, Kovács, & Ambrus, 2022; Dalski, Kovács, Wiese, et al., 2022; C. Li et al., 2022) as well as other visual object categories (Ambrus, 2024; Klink et al., 2023; Ozdemir & Ambrus, submitted). Furthermore, although the time course of expressive versus neutral face processing has been described (Zhang et al., 2023), no study has yet examined the neural dynamics specific to different emotional expressions. Finally, the neural processing dynamics of facial attributes such as sex and identity in relation to facial expression, have yet to be explored. Characterizing the generalizable neural signals of facial expression processing provide insights into the neural architecture and processes involved in social cognition and can serve as a steppingstone for investigating how these processes are affected in conditions such as autism, schizophrenia, or social anxiety.

Here, we investigated the shared neural dynamics of emotional face processing employing an explicit facial emotion recognition task, where participants made two-alternative forced choice (2AFC) decisions on the displayed emotion. Our approach involved cross-participant multivariate classification and representational dissimilarity analysis on EEG data. We investigated the effects of stimulus identity, sex, and emotion, including pairs of emotions, while also considering the impact of visual image properties on the observed neural dynamics.

Our results are in line with previous findings demonstrating that EEG signals can effectively be used to decode the sex, emotional expression, and identity of face stimuli. Moreover, these can be decoded across stimuli and participants, indicating shared neural codes for facial expression processing. In terms of the onset of these effects, multivariate classification analyses revealed that sex is decoded first, followed by identity, and then emotion. When analyzing pairs of expressions, emotional expressions (angry, happy, sad) were decoded earlier against neutral expressions. Representational similarity analyses indicated that visual image properties reduced the effect of identity and sex decoding while they had minimal to no impact on emotion classification. Identity and sex representations are strongest in angry faces, followed by happy and sad faces, compared to neutral faces.

We demonstrated the feasibility of cross-participant multivariate classification in decoding emotional face processing, revealing the shared neural dynamics across individuals. Our analyses unveiled distinct patterns for stimulus identity, sex, and emotion processing. Notably, the effects of emotion were found to be independent of visual image properties. Our investigation into pairs of emotions sheds light on the temporal sequence of processing, providing insights into the prioritization and interaction of different emotional expressions. Importantly, our results suggest enhanced face-identity and sex processing for emotional expressions, particularly for angry faces and to some extent for happy faces.

## Methods

### Participants

Our sample consisted of 24 healthy university students (8 males and 16 females). Participants took part in the study for partial course credits, and all provided written informed consent before the experiment. Volunteers were recruited through the SONA research participation system or personal contacts. Participants disclosed no history of neurological conditions, had normal or corrected-to-normal vision, and were right-handed. The experiment was conducted in accordance with the guidelines of the Declaration of Helsinki, and with the approval of the ethics committee of Bournemouth University [Ethics ID: #52261]. Written informed consent was acquired from all participants.

### Stimuli

The stimuli consisted of frontal color photographs featuring eight individuals sourced from the KDEF database (Lundqvist et al., 1998), including four males (AM10, AM17, AM24, AM31) and four females (AF07, AF15, AF26, AF28). These images depicted four distinct (posed) facial expressions— happy, angry, sad, and neutral—amounting to a total of 32 images. The images were presented in a randomized order across runs.

The decision to focus exclusively on happy, angry, sad, and neutral facial expressions in this present investigation stems from a deliberate choice based on several considerations. Firstly, the selected emotional expressions represent a well-established and widely studied core set that captures fundamental affective states. Anger and disgust have consistently emerged as the least quickly and accurately recognized facial expressions across existing literature (Hendel et al., 2023), while fear and surprise are also easily confused (Zhao et al., 2017). To maintain methodological clarity and enhance the reliability of our findings, we chose to exclude surprise, fear and disgust from the current investigation.

### Experimental Design

In the context of a 2AFC (Two-Alternative Forced Choice) design, preceded by the presentation of a fixation cross for 200 ms, each face image was displayed for a duration of 1000 ms, followed by the appearance of a choice screen featuring the correct emotion and an incorrect emotion and an interstimulus interval between 500 and 1000 ms (**Figure 1**). Participants were instructed to select the correct emotion displayed by the presented face using the left or right key, with no time limit for making their choice. Each image was repeated 12 times. For each image, the veridical facial expression displayed was paired with an incorrect choice 4 times, with the response key assignment balanced. To ensure a balanced number of correct trials, in instances where participants provided an incorrect response, the trial was reintroduced into the trial sequence, effectively rescheduled for a subsequent time-point. As such, the dataset for each participant encompassed 12 repetitions of an image, 96 presentations of a facial expression, and 48 presentations of an identity, totaling 384 trials. The experiment was programmed in and was carried out using PsychoPy (Peirce, 2007; Peirce et al., 2019).

**Figure 1.**
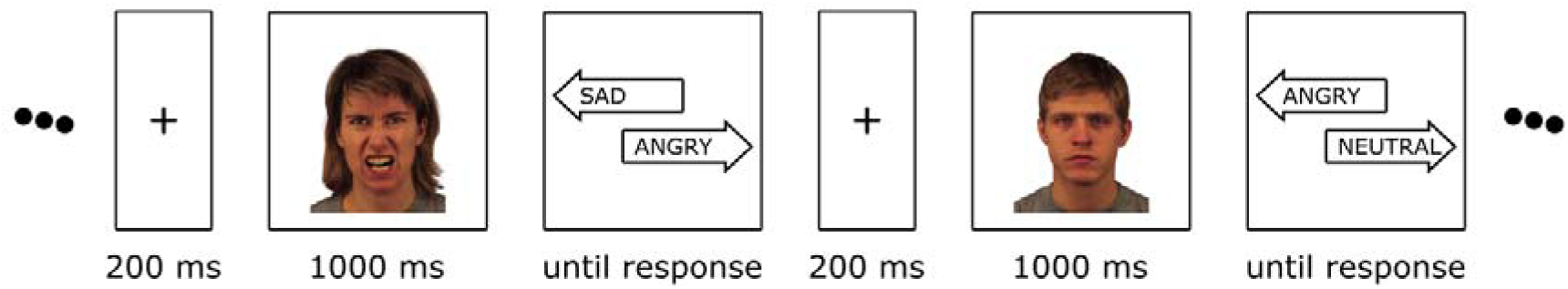
Experimental design. The stimuli comprised randomized 32 frontal color photographs of eight individuals (four males and four females) from the KDEF database, displaying posed happy, angry, sad, and neutral facial expressions. In a 2AFC design, each face image appeared for 1000 ms after a 200 ms fixation cross, followed by a choice screen with the correct and an incorrect emotion. Participants, with no time limit, were asked to select the correct emotion using the left or right key. Images were repeated 12 times, totaling 384 trials with 96 facial expression presentations and 48 identity presentations.

### EEG recording

EEG was recorded using a 64-channel BioSemi Active-Two system (https://www.biosemi.com/products.htm) from electrode sites based on the 10-20 international system, in the EEG laboratory of the Department of Psychology at Bournemouth University. EEG data was processed in MNE-Python (Gramfort et al., 2013, 2014); it was bandpass filtered between 0.1 and 40 Hz, segmented between −200 and 1200 ms and baseline-corrected to the 200 ms preceding the stimulus presentation and downsampled to 200 Hz. To enhance the signal-to-noise ratio and expedite computation (Grootswagers et al., 2017), for each participant, evoked responses for trials presenting the same image were randomly grouped into 3-trial bins and averaged. No further processing had been performed (T. A. Carlson et al., 2020; Delorme, 2023; Grootswagers et al., 2017), 2022; Grootswagers et al., 2017). Data handling was carried out using the numpy and scipy packages (Harris et al., 2020; Virtanen et al., 2020).

### Analysis pipeline

#### Classification analyses

Classification analyses were carried out using scikit-learn (Pedregosa et al., 2011). Linear discriminant analysis (LDA) classifiers were systematically trained across participants to categorize the identity, sex, and facial expression, as well as the combinations of facial expressions of the presented faces. The procedure followed a leave-one participant-out scheme. For identity classification, classifiers underwent iterative training on three emotion categories, successively tested on the emotion category left out. In the case of sex and emotion classifiers, training was conducted on six distinct identities (three male and three female) and subsequently tested on one identity left out. The classification of pairs of emotional expressions followed a similar logic, with the distinction that only two emotional expressions were included at a time in the classification procedure. The training involved six identities (three male and three female) and was subsequently tested on one identity that was omitted during training. Classification accuracies were tested against chance levels (identity: 0.125, sex: 0.5, emotion: 0.25, pairs of facial expressions: 0.5, see **Supplementary Information Figure S1**). For a similar approach, see Klink et al. (2023). To investigate the time-course of face identity and sex processing across various emotional expressions, cross-classification was employed (Kaplan et al., 2015). This involved training on data from trials with neutral expressions and testing the classification performance on trials featuring one of the emotional expressions, and vice versa, following the previously described methodology. The resulting classification accuracies in both directions (e.g. neutral-to-angry and angry-to-neutral) were averaged within each participant (Man et al., 2012; Oosterhof et al., 2012) and tested statistically against chance (see **Supplementary Information Figure S2**).

Time-resolved classification was conducted across all electrodes and pre-defined regions of interest (ROIs), encompassing six scalp locations along the median (left and right) and coronal (anterior, center, and posterior) planes. The spatio-temporal searchlight procedure systematically tested each channel by training and testing on data from that channel and its neighboring electrodes, applying the same time-resolved analysis logic as previously described (Ambrus, 2024; Dalski, Kovács, & Ambrus, 2022).

#### Representational similarity analyses

Representational similarity analyses (Kriegeskorte, 2008) entailed the creation of participant-level empirical neural representational dissimilarity matrices (RDMs) for each time-point. This was achieved through leave-one-subject-out pairwise classification of stimulus pairs, resulting in matrices of dimensions 280 by 32 by 32. The classification was performed following the previously outlined procedure (for a similar approach, see Ozdemir & Ambrus, *submitted*). Subsequently, these matrices were compared to predictor representational dissimilarity matrices modeling identity, sex, and facial expression (32 by 32 matrices), as well as pairs of expressions (16 by 16 matrices), utilizing Spearman rank correlations (see **Supplementary Information Figure S3**). The resulting correlation values were then Fisher-transformed. Partial correlations were used to account for the effects of visual image properties. Model RDMs were constructed using the Euclidean distances of maximum cross-correlation (Collin et al., 2022) and face-descriptors obtained using the dlib package (King, 2009).

#### Statistical testing

A moving average of 35 ms (spanning 7 consecutive time points) was applied to all participant-level classification accuracy data. (Ambrus, 2024; Ambrus et al., 2019, 2021; Dalski, Kovács, & Ambrus, 2022; Klink et al., 2023) The statistical evaluation of the results employed both cluster permutation tests and Bayesian statistical analyses. Classification accuracies underwent two-sided, one-sample cluster permutation tests (10,000 iterations) against chance, using MNE-Python. The Bayesian tests (Teichmann et al., 2022) also employed a two-sided approach, utilizing a non-directional whole-Cauchy prior with a medium width (*r* = 0.707) and excluding an interval from *δ* = -0.5 to +0.5. The resulting Bayes factors were then thresholded, with values exceeding 10 considered as strong evidence (Moerel et al., 2022; Wetzels et al., 2011). Bayesian statistical analyses were performed using the BayesFactor R package (Morey et al., 2015).

## Results

### Behavioral results

Behavioral results are shown in **Figure 2** . As mentioned above, incorrect trials were repeated; this analysis does not account for those repeated trials. The mean classification accuracy across all participants was 96.6%, with a standard deviation of 2.86%. The average of the median reaction times across participants was 645 ms (±105.13). In terms of reaction times, no difference was seen between neutral (0.641 ± 0.1144) and angry faces (0.664 ± 0.0962), neutral and sad faces (0.673 ± 0.1891), or angry and sad faces. Reaction times for angry faces were significantly higher than those for happy faces (0.635 ± 0.0955). These results align with previous studies that report faster choice reaction times for faces displaying positive expressions compared to negative or neutral ones, indicating a happy-face advantage that is theorized to primarily occur at premotor processing stages (Leppänen et al., 2003).

**Figure 2.**
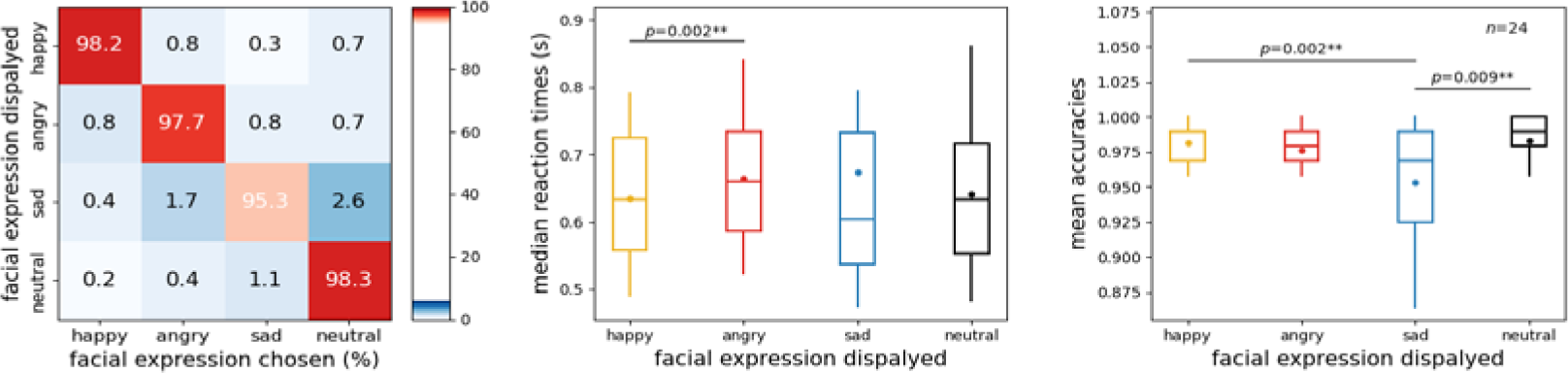
Behavioral results. Left: confusion matrix. The rows correspond to expression categories, while columns denote the responses selected by participants (happy, angry, sad, neutral). The diagonal elements signify accurate responses, while off-diagonal elements indicate errors. The color scale and numerical values within cells denote the average percentage of occurrences for specific facial expression and response pairings across participants. Middle and Right: reaction times and accuracy scores; first, second (median), and third quartiles; whiskers with Q1 − 1.5 IQR and Q3 + 1.5 IQR. The dot denotes the mean. Wilcoxon signed rank test, puncorrected < 0.05.

**Figure 3.**
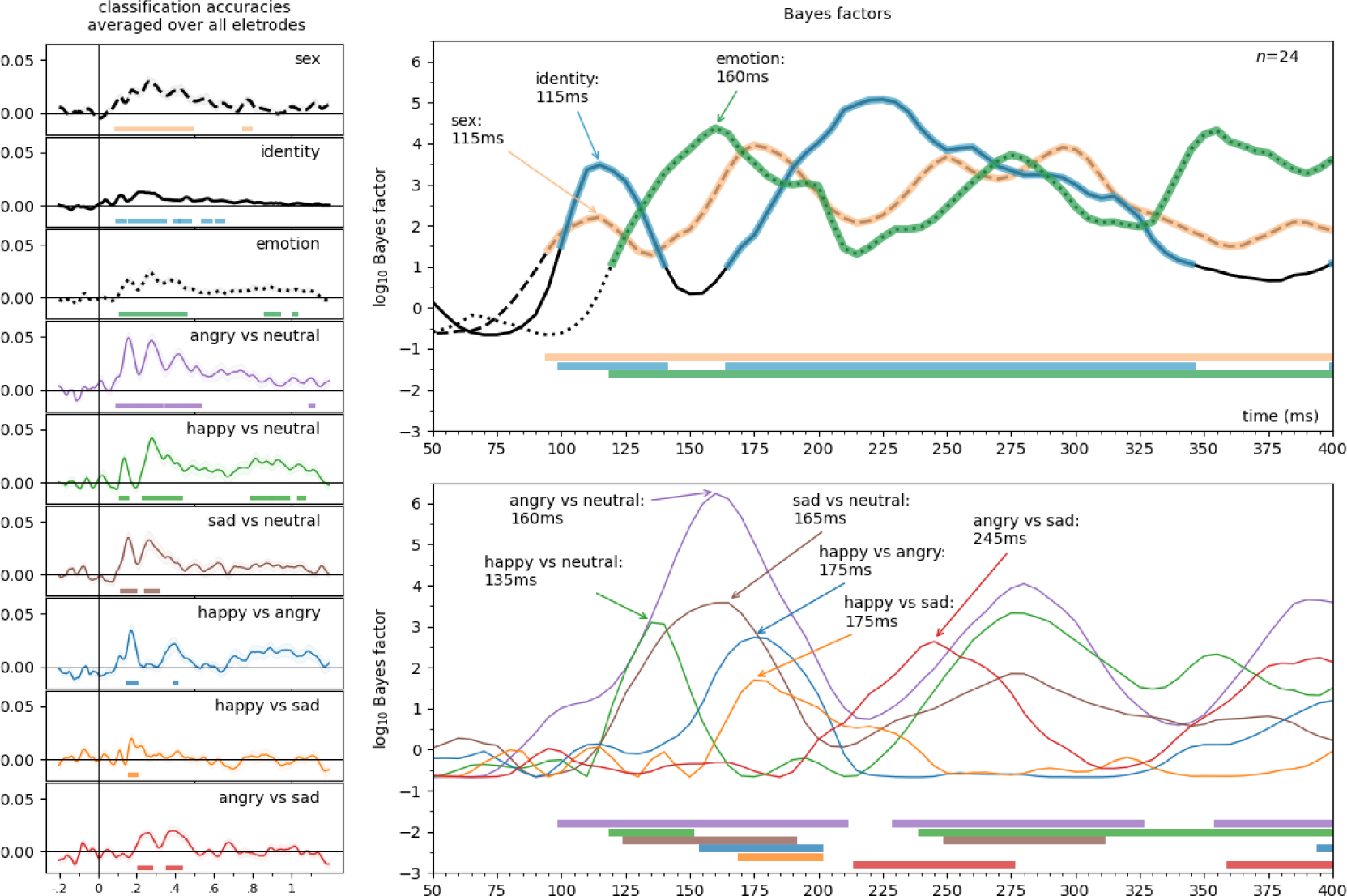
The onsets of the effects of identity, sex, emotion displayed, and emotion discrimination. Time-resolved searchlight decoding was systematically performed on all channels and their neighboring electrodes. The classifier performance was averaged across all channels to emphasize the onset and duration of these effects. Onsets were determined by Bayes factors against chance, with values exceeding 10 considered indicative of strong evidence (horizontal significance markers). The left panel shows the entire time course of the effects in terms of classification accuracy averaged over all channels; the top right panel displays the effects of identity, sex, and emotional expression, while the bottom right panel displays the emotional expression discrimination in log10 Bayes factors, between 50 and 400 milliseconds. The first peak in each effect is indicated.

In terms of accuracy, the highest accuracies were seen for neutral (0.982 ± 0.0299) and happy faces (0.982 ± 0.0160), followed by angry faces (0.976 ± 0.0238) and finally sad faces (0.953 ± 0.040). While accuracy rates for neutral, happy and angry faces were not different, sad expressions were harder to recognize, most often confused with neutral (2.6%) and angry (1.7%) faces. Accuracy results also align with prior literature, with the sad expression being most confused with neutral and angry emotional displays (Du & Martinez, 2011; Goren & Wilson, 2006).

### Classification analyses

### Effect onsets

To estimate the onset of effects, various approaches were considered. Inspecting decoding accuracies or model correlations across all electrodes might overlook weaker effects due to the curse of dimensionality. Another option, focusing on *a priori* regions of interest, could miss effects not manifesting at those specific scalp locations. Therefore, we chose to average decoding accuracies in the searchlight analyses across all electrodes and timepoints, thus providing a more comprehensive assessment of the temporal dynamics and mitigating the limitations associated with other methods. For a visualization of first peak latencies so calculated, see **Figure 4** . Note that the other previously mentioned analyses yielded similar findings.

**Figure 4.**
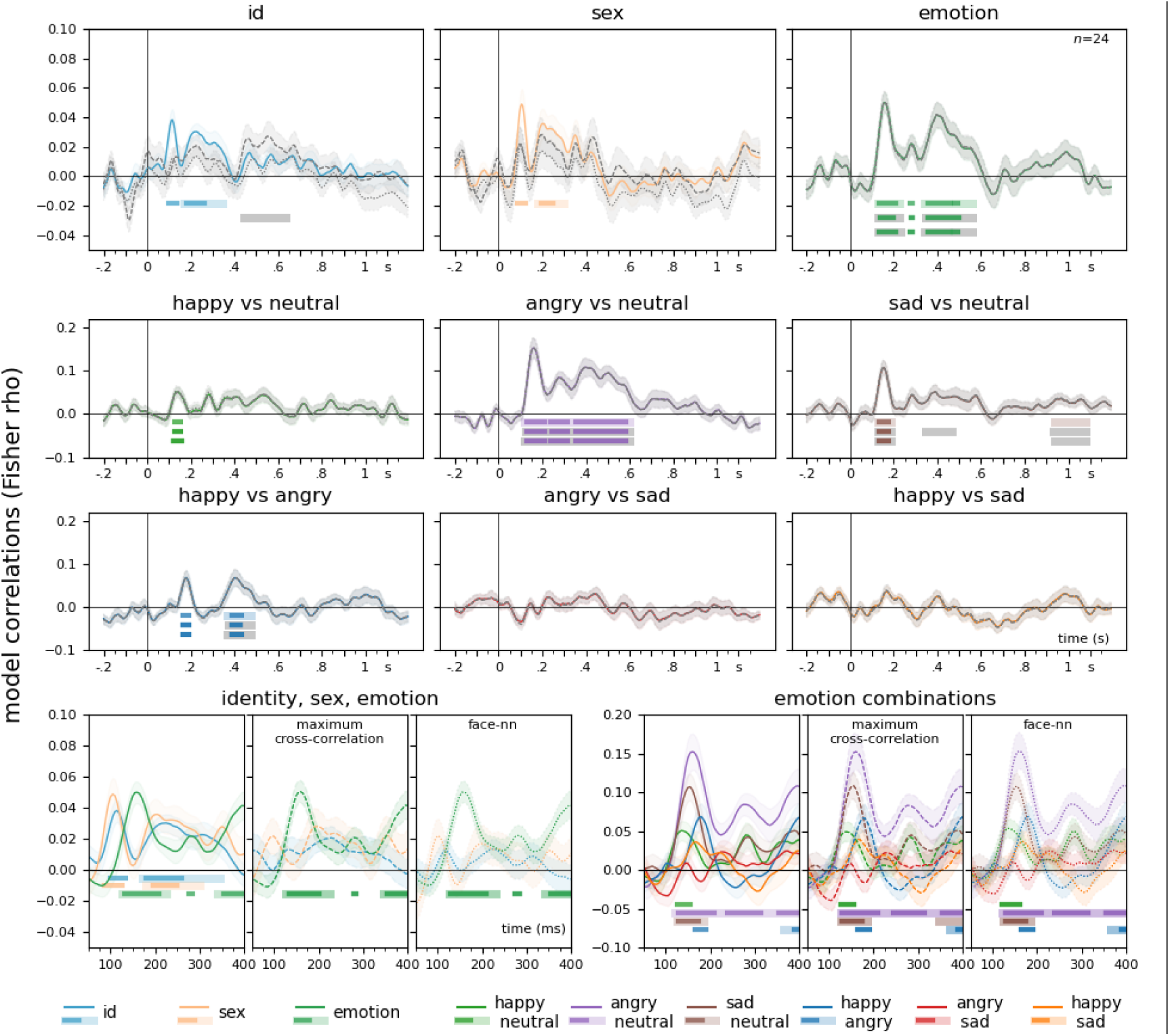
Representational Similarity Analysis. Results for all electrodes are presented. Time-resolved neural representational dissimilarity matrices were formed through pairwise decoding of EEG data, adopting a leave-one-participant-out approach. These matrices were then assessed against model RDMs for identity, sex, and facial expression, along with pairs of facial expressions, utilizing rank correlations. To control for visual image properties, partial correlations were employed. Solid lines represent model correlations, dashed lines illustrate results when the effects of maximum cross-correlation are controlled for, and dotted lines depict results with the effects of neural network feature distance partialled out. Visual image properties diminished the effect of identity and sex, but not emotion. The figure shows the analysis conducted over all electrodes. Light lines denote significant clusters revealed by the two-sided cluster permutation tests, *p* < 0.05; dark lines denote results of the Bayesian statistical analyses, two-sided one-sample Bayesian t-tests, bf >10. Error ranges denote ± SEM. For detailed statistics, see Supplementary Table 1.

The results pertaining to the first peaks and peak latencies of the effects are as follows. Sex: the effect spanned from 95 to 485 ms, with a peak at 115 ms and a peak Bayes Factor of 161.63. Identity: observed between 100 to 140 ms, with a peak at 115 ms and a substantial peak Bayes Factor of 3024.61. Emotion: extended from 120 to 450 ms, peaking at 160 ms with a remarkably high peak Bayes Factor of 23860.78. Angry vs. neutral: spanning from 100 to 210 ms, the peak occurred at 160 ms, accompanied by an exceptionally high peak Bayes Factor of 1736495.2. Happy vs. neutral: Manifesting between 120 to 150 ms, the peak emerged at 135 ms, with a peak Bayes Factor of 1251.35. Sad vs. Neutral: present from 125 to 190 ms, the peak latency was at 165 ms, yielding a peak Bayes Factor of 3784.43. Happy vs. angry: taking place between 155 to 200 ms, the peak was observed at 175 ms, accompanied by a peak Bayes Factor of 543.72. Happy vs. sad: ranging from 170 to 200 ms, with the peak occurring at 175 ms and a peak Bayes Factor of 49.46. Angry vs. sad: first peak between 215 to 275 ms, the peak was noted at 245 ms, with a peak Bayes Factor of 430.02.

### Representational similarity analysis

Time-resolved neural representational dissimilarity matrices were generated by pairwise decoding of EEG data at each time-point, employing a leave-one-participant-out strategy. This process involved pairwise classification of all stimulus pairs, resulting in matrices with one empty off-diagonal. Subsequently, these matrices underwent evaluation against model representational dissimilarity matrices (model RDMs) for identity, sex, and facial expression, as well as pairs of facial expressions, using rank correlations. The neural representational dissimilarity matrices were compared to model RDMs, which featured zeros for within-class comparisons and ones for cross-class comparisons in cells, employing Spearman rank correlations.

To account for the potential impact of image-level stimulus properties, two visual image similarity metrics were used. Maximum cross-correlation served as the metric for assessing lower-level image similarity, whereas higher-level and face-specific visual features were gauged using neural network descriptors extracted from a dedicated face identification model. Maximum cross-correlation resembles pixelwise correlation similarity but is equivalent to calculating the highest correlation achievable by translating one image in a pair relative to the other through all possible positions. This process involves convolving the two images, generating a cross-correlation map where the peak indicates the maximum potential correlation had the images been optimally co-registered (Collin et al., 2022). The neural network feature similarity RDM was calculated by computing pairwise Euclidean distances between the 128 feature descriptors derived from processing the images through a neural network specifically trained for face identification (dlib, King, 2017).

Results of the representational similarity analysis over all electrodes are presented in **Figure 4** . Model correlations over all electrodes revealed similar patterns of effects described for the previous, decoding searchlight-based analysis. For face identity, Bayesian analyses indicated evidence for an effect between 100 and 135 ms (peak at 115 ms, bf = 1291.34) that was not picked up by the cluster permutation tests. Cluster permutation tests identified an effect at 170 to 345 ms (cluster *p* = 0.004). Sex-related effects were evidenced by Bayesian analyses between 85 and 125 ms (peak at 105 ms, bf = 270.66) and exhibited a cluster from 180 to 300 ms (cluster *p* = 0.034). Emotional face processing showed clusters from 125 to 225 ms (cluster *p* = 0.02) and 340 to 560 ms (cluster *p* = 0.005). No significant clusters were found for angry vs sad or happy vs sad. Bayesian analyses indicated an early effect for happy vs neutral between 125 and 155 ms (peak at 135 ms, bf = 49.71). For angry vs neutral, a significant cluster emerged at 120 to 600 ms (cluster *p* = 0.0002), and for sad vs neutral, a cluster was observed from 125 to 185 ms (cluster *p* = 0.047) and 935 to 1080 ms (cluster *p* = 0.023). Happy vs angry exhibited a cluster at 365 to 475 ms (cluster *p* = 0.035).

When controlling for low-level and face-specific image properties, in the case of identity, all effects before ca. 400 ms were attenuated. Partialling out low-level image properties yielded a ca. 400 to 600 ms identity effect, in line with results of previous investigations indicating an image-independent representation of identity in this time range (Ambrus et al., 2019). Controlling for face-specific image properties reduced this effect as well. The effects of face sex were also reduced when controlling for both low-level and face-specific image properties. The effects seen for emotion, as well as for pairs of emotions, were essentially unaffected by controlling for either low-level, or face-specific image properties, indicating that these effects capture image-independent processing of these factors.

For the results of the representational similarity analysis in the pre-defined regions of interest, see **Supplementary Information Figures S5 and S6**, for detailed statistics, including Bayesian analyses, see **Supplementary Table 1**.

### Multivariate cross-classification

Results of the multivariate cross-classification analyses for identity, sex, and emotion are presented in **Figure 5**. For detailed statistics, including Bayesian analyses, see **Supplementary Table 2** .

**Figure 5.**
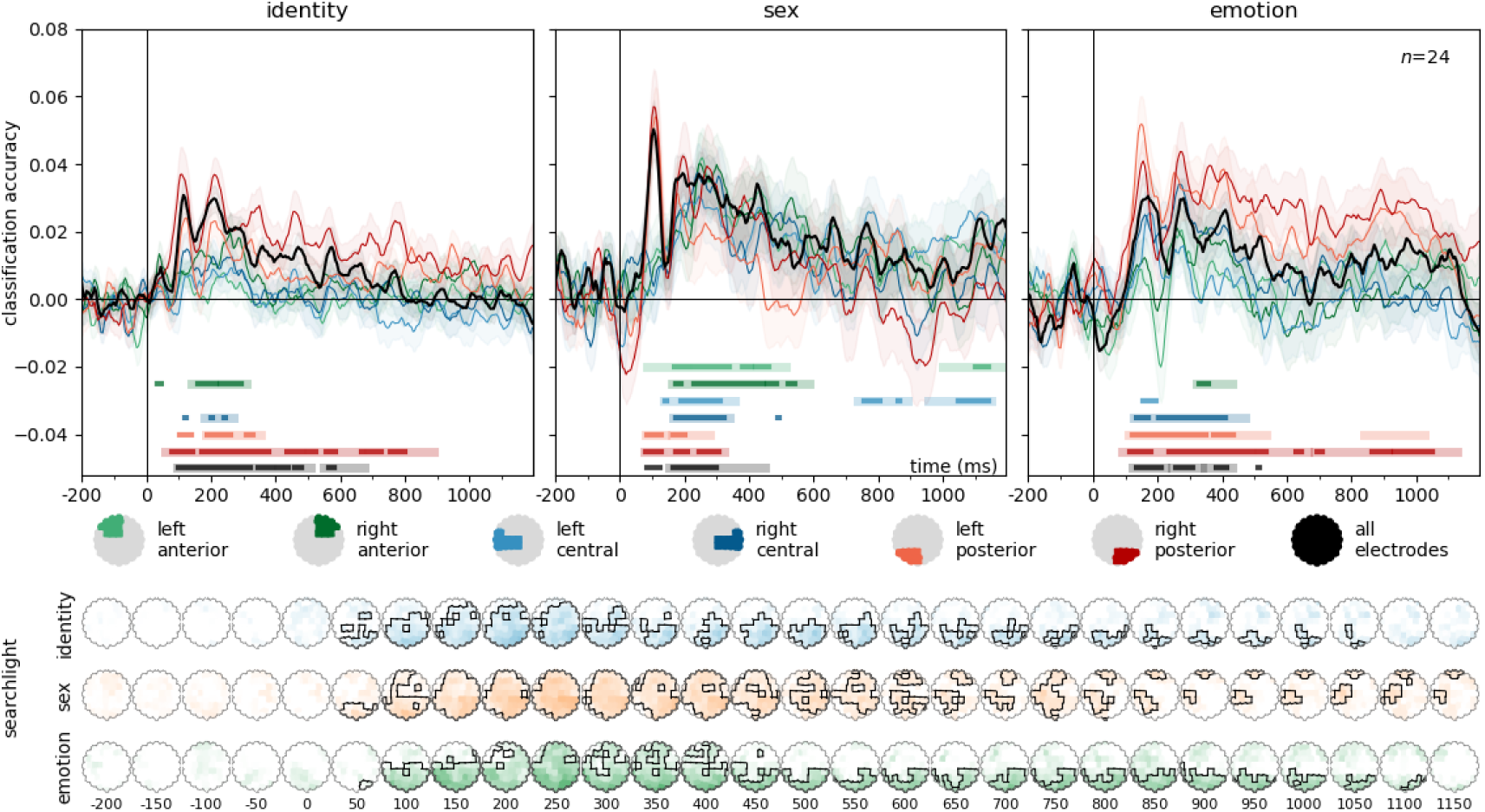
Time-resolved, leave-one-participant-out classification of identity, sex, and emotion. . Classifiers were trained, in a leave-one-participant-out scheme, to categorize the identity, sex, and facial expression of the stimulus faces. For identity, classifiers were iteratively trained on three emotion categories, and tested on the emotion category left out. For sex and emotion, training was performed on six identities (3 male and 3 female) and tested on one left out. Classification accuracies are plotted against chance (identity: 0.125, sex: 0.5, emotion: 0.25). Error ranges denote ± SEM. Color lines represent regions of interests. Light lines denote significant clusters revealed by the two-sided cluster permutation tests, *p* < 0.05; dark lines denote results of the Bayesian statistical analyses, two-sided one-sample Bayesian *t*-tests, bf >10. Spatio-temporal searchlight results are shown as scalp maps, with classification accuracy scores averaged in 50 ms steps. Sensors and time points belonging to significant clusters are marked. (Two-sided spatio-temporal cluster permutation tests, *p* < 0.05). For detailed statistics, see **Supplementary Table 2A-C** .

#### Identity

Over all electrodes, two significant clusters were identified. The first cluster spanned the time window from 95 to 510 ms, peaking at 115 ms, with a highly significant cluster (p<0.0001, peak Cohen’s *d* = 1.6927). The second cluster, from 550 to 675 ms, peaked at 575 ms (cluster *p* = 0.0194, peak Cohen’s *d* = 0.7775). Further significant clusters were observed in the right and left posterior, right central, and right anterior regions of interest. The searchlight analysis yielded a single significant cluster was identified, spanning from 75 to 1075 ms, with a peak at 225 ms over PO8 (cluster *p* < 0.0001, peak Cohen’s *d* = 1.5945).

#### Sex

For all electrodes, a single significant cluster emerged in the time window from 155 to 450 ms, peaking at 195 ms (cluster *p* = 0.008, peak Cohen’s *d* = 1.0624). Significant clusters were observed in all pre-defined regions of interest as well. In the searchlight analysis, a significant cluster was found in the time window from 70 to 1195 ms, peaking at 105 ms over POz (cluster *p* < 0.0001, peak Cohen’s *d* = 1.8928).

#### Emotion

Three significant clusters were identified for emotion over all electrodes. The first cluster spanned the time window from 120 to 225 ms, with a peak at 170 ms (cluster *p* = 0.0149, peak Cohen’s *d* = 1.1992). The second cluster, from 245 to 335 ms, peaked at 280 ms (cluster *p* = 0.0326, peak Cohen’s *d* = 0.917). The third cluster, covering 345 to 430 ms, peaked at 410 ms (cluster *p* = 0.0437, peak Cohen’s *d* = 0.7775). The pre-defined regions of interest also yielded significant clusters, with the exception of the left anterior and left central regions. The searchlight analysis yielded a significant cluster from 80 to 1145 ms, with a peak at 150 ms over PO7 (cluster *p* < 0.0001, peak Cohen’s *d* = 1.6457).

Results of the multivariate cross-classification analyses for the differential decoding of facial expressions are presented in **Figure 6**. For detailed statistics, including Bayesian analyses, see **Supplementary Table 2**.

**Figure 6.**
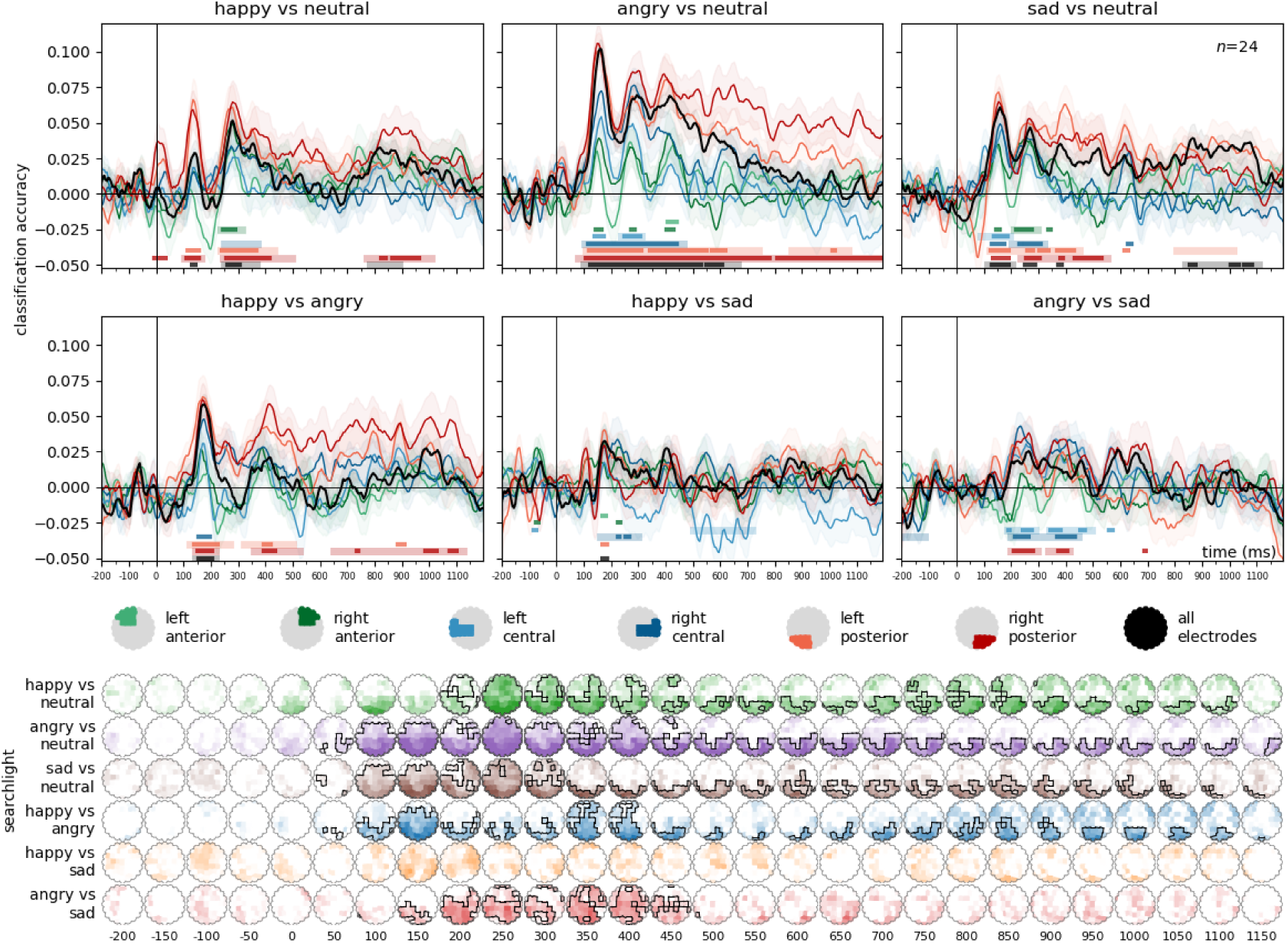
Differential decoding of emotional expressions. For the classification of emotion pairs, training was iteratively performed on six identities (3 male and 3 female) and tested on one left out. Classification accuracies are plotted against chance (0.5). Bottom panels: spatio-temporal searchlight results are shown as scalp maps, with classification accuracy scores averaged in 50 ms steps. Sensors and time points forming part of the significant clusters are marked (Two-sided spatio-temporal cluster permutation tests, *p* < 0.05). For detailed statistics, see Supplementary Table 2D-I.

#### Happy vs Neutral

Two significant clusters were found for all electrodes. The first cluster spanned the time window from 250 to 365 ms, with a peak at 280 ms (cluster *p* = 0.0271, peak Cohen’s *d* = 0.8811). The second cluster, from 785 to 890 ms, peaked at 820 ms (cluster *p* = 0.0427, peak Cohen’s *d* = 0.5888). In the searchlight analysis, a significant cluster was found between 215 to 1145 ms, peaking at 290 ms over channel P8 (cluster *p* = 0.0005, peak Cohen’s *d* = 1.1647).

#### Angry vs Neutral

Over all electrodes, a single highly significant cluster was observed across the entire time window from 100 to 665 ms, with a peak at 160 ms (cluster *p* < 0.0001, peak Cohen’s *d* = 2.106). The searchlight analysis identified a significant cluster between 80 to 1195 ms, peaking at 150 ms over POz (cluster *p* <0.0001, peak Cohen’s *d* = 1.6915).

#### Sad vs Neutral

Two significant clusters were identified over all electrodes. The first cluster spanned the time window from 115 to 205 ms, with a peak at 160 ms (cluster *p* = 0.0313, peak Cohen’s *d* = 1.0432). The second cluster, covering 840 to 1105 ms, peaked at 865 ms (cluster *p* = 0.0062, peak Cohen’s *d* = 0.7572). The searchlight procedure yielded a significant cluster from 75 to 1195 ms, with a peak at 155 ms over POz (cluster *p* = 0.0003, peak Cohen’s *d* = 1.6955).

#### Happy vs Angry

For all electrodes, a significant cluster was found in the time window from 145 to 215 ms, peaking at 175 ms (cluster *p* = 0.0267, peak Cohen’s *d* = 1.2307). Two significant clusters were found in the searchlight analysis. The first cluster spanned from 95 to 260 ms, peaking at 170 ms over channel PO7, (cluster *p* = 0.0437, peak Cohen’s *d* = 1.1389). The second cluster extended from 225 to 1155 ms, peaking at 425 ms over Oz (cluster *p* = 0.0098, peak Cohen’s *d* = 1.1444).

#### Angry vs Sad

No significant clusters were observed in this comparison for all electrodes. Significant clusters were seen in the central and right posterior regions of interest. In the searchlight analysis, two significant clusters were identified. The first cluster spanned from 140 to 500 ms, peaking at 260 ms over channel CP2 (cluster *p* = 0.0002, peak Cohen’s *d* = 1.1469). The second cluster covered the time window from -200 to -80 ms, with a peak at -120 ms over CP1 (cluster *p* = 0.0278, peak Cohen’s *d* = -0.948).

#### Happy vs Sad

No significant clusters were found in this comparison for all electrodes. Significant clusters were seen in the central regions of interest. The searchlight analysis yielded no significant clusters either.

### The effects of expression on face identity and sex processing

Results of the multivariate cross-classification analyses for face-identity and sex decoding in the different emotion conditions are presented in **Figure 7**. For detailed statistics, including Bayesian analyses, see **Supplementary Table 3** .

**Figure 7.**
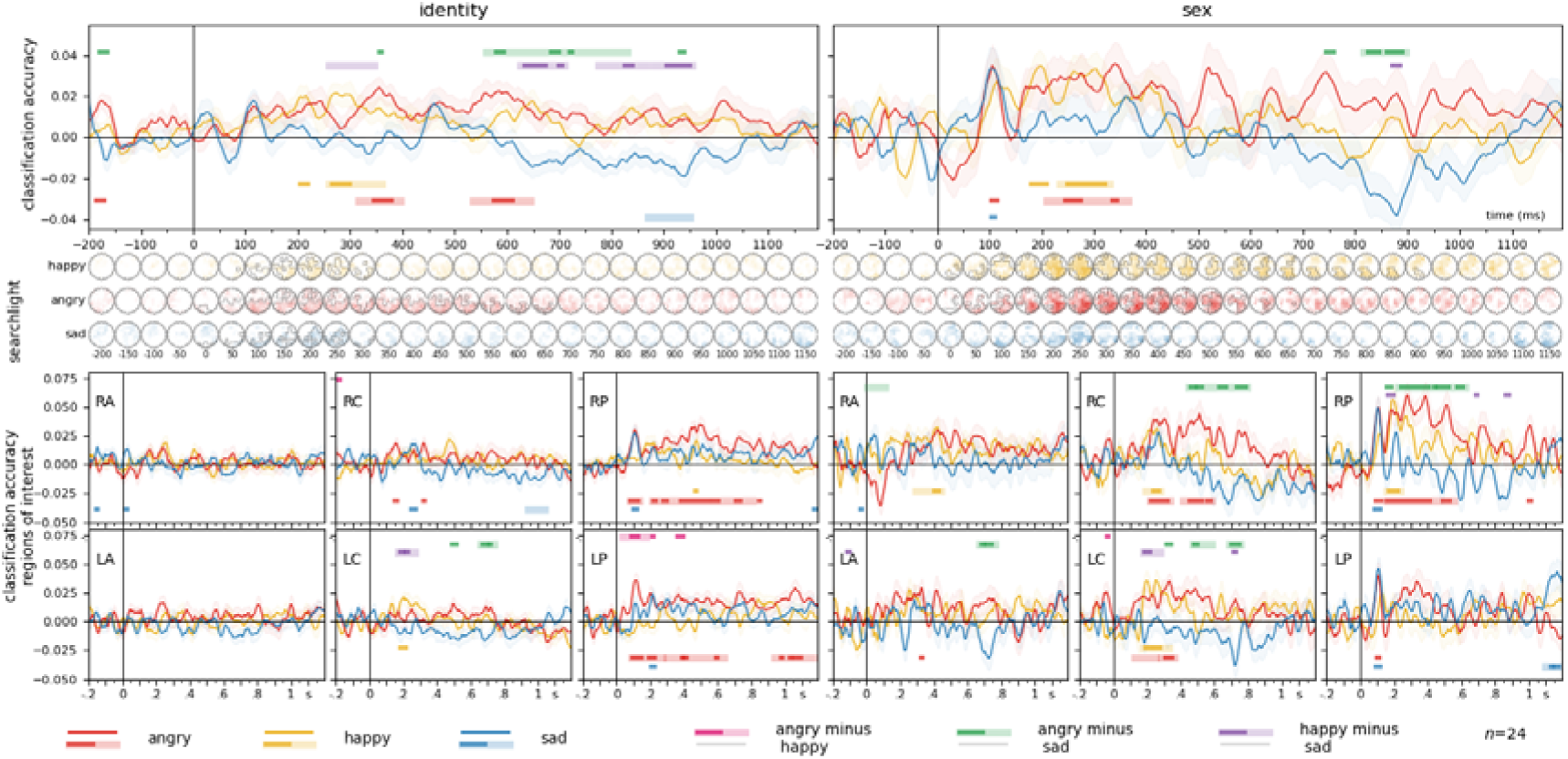
Identity and sex decoding for emotional expressions vs neutral (classification accuracies for the directions emotion-to-neutral and neutral-to-emotion averaged). Middle Panels: spatio-temporal searchlight results are shown as scalp maps, with classification accuracy scores averaged in 50 ms steps. Sensors and time points forming part of the significant cluster when tested on faces are shown in the top row, sensors and time points belonging to the significant cluster when tested on scenes are shown in the bottom row. (Two-sided spatio-temporal cluster permutation tests, *p* < .05). Bottom panels: ROI analyses. The same analysis as in the top panel but repeated for six pre-defined electrode clusters separately. RA/LA: right/left anterior, RC/LC: right/left central, RP/LP: right/left posterior. For detailed statistics, see Supplementary Table 3.

### Identity

#### Happy

Over all electrodes, a significant cluster was observed in the time window from 260 to 360 ms, peaking at 275 ms (cluster *p* = 0.0193, peak Cohen’s *d* = 0.8727). In the searchlight analysis, a significant cluster was observed from 140 to 335 ms, with a peak at 190 ms over P6 (cluster *p* = 0.0044, peak Cohen’s *d* = 0.8283).

#### Angry

Over all electrodes, two significant clusters were identified. The first cluster spanned from 315 to 395 ms, with a peak at 355 ms (cluster *p* = 0.0315, peak Cohen’s *d* = 0.9274). The second cluster covered the time window from 535 to 645 ms, peaking at 585 ms (cluster *p* = 0.013, peak Cohen’s *d* = 0.9170). Large significant clusters were observed over the left and right posterior regions of interest. The searchlight analysis yielded a significant cluster from 85 to 675 ms, with a peak at 110 ms over O1 (cluster *p* < 0.0001, peak Cohen’s *d* = 1.1285).

#### Sad

A significant cluster was found over all electrodes in the time window from 870 to 950 ms, peaking at 870 ms (cluster *p* = 0.0283, peak Cohen’s *d* = -0.4318). The searchlight analysis identified a significant cluster between 80 to 325 ms, peaking at 275 ms over P9 (cluster *p* = 0.0104, peak Cohen’s *d* = 0.7277).

#### Difference, Angry-Happy

No significant clusters were found over all electrodes or in the searchlight analysis.

#### Difference, Angry-Sad

A significant cluster for all electrodes was observed from 560 to 830 ms, with a peak at 645 ms (cluster *p* = 0.001, peak Cohen’s *d* = 0.6286). No significant clusters were found in the searchlight analysis.

#### Difference, Happy-Sad

Four significant clusters were identified over all electrodes. The first cluster spanned from 260 to 345 ms, peaking at 300 ms (cluster *p* = 0.0345, peak Cohen’s *d* = 0.5879). The second cluster covered the time window from 625 to 710 ms, peaking at 650 ms (cluster *p* = 0.0215, peak Cohen’s *d* = 0.7639). The third cluster extended from 775 to 865 ms, with a peak at 830 ms (cluster *p* = 0.0286, peak Cohen’s *d* = 0.7068). The fourth cluster spanned from 880 to 955 ms, peaking at 940 ms (cluster *p* = 0.0162, peak Cohen’s *d* = 1.0248). In the searchlight analysis, no significant clusters were found.

### Sex

#### Happy

A significant cluster was found in the time window from 235 to 330 ms, peaking at 305 ms (cluster *p* = 0.0229, peak Cohen’s *d* = 0.7960). Significant clusters were observed in the central, and right anterior and posterior regions of interest. The searchlight analysis revealed a significant cluster between 75 and 905 ms, with a peak at 205 ms over channel P4 (cluster *p* = 0.0005, peak Cohen’s *d* = 0.9495).

#### Angry

A significant cluster was identified from 210 to 365 ms, with a peak at 340 ms (cluster *p* = 0.0211, peak Cohen’s *d* = 0.7002). Significant clusters were observed in the bilateral central and right posterior regions of interest. The searchlight analysis identified a significant cluster from 75 to 650 ms, with a peak at 390 ms over channel P4 (cluster *p* = 0.0002, peak Cohen’s *d* = 0.9530).

#### Sad

No significant clusters were found over all electrodes, and no significant clusters were seen in the searchlight analysis.

#### Difference, Angry-Happy

No significant clusters were found over all electrodes, and no significant clusters were seen in the searchlight analysis.

#### Difference, Angry-Sad

A significant cluster was observed from 815 to 895 ms, with a peak at 880 ms (cluster *p* = 0.038, peak Cohen’s *d* = 0.8669). The searchlight analysis yielded a significant cluster from 225 to 340 ms, with a peak at 300 ms over O2 (cluster *p* = 0.0167, peak Cohen’s *d* = 0.9104).

#### Difference, Happy-Sad

No significant clusters were found over all electrodes, and no significant clusters were seen in the searchlight analysis.

To better visualize these results, we plotted identity and sex classification accuracies averaged within time-windows of interest over all electrodes and the right posterior area (**Figure 8**). The most prominent effects were consistently observed for the angry facial expression.

**Figure 8.**
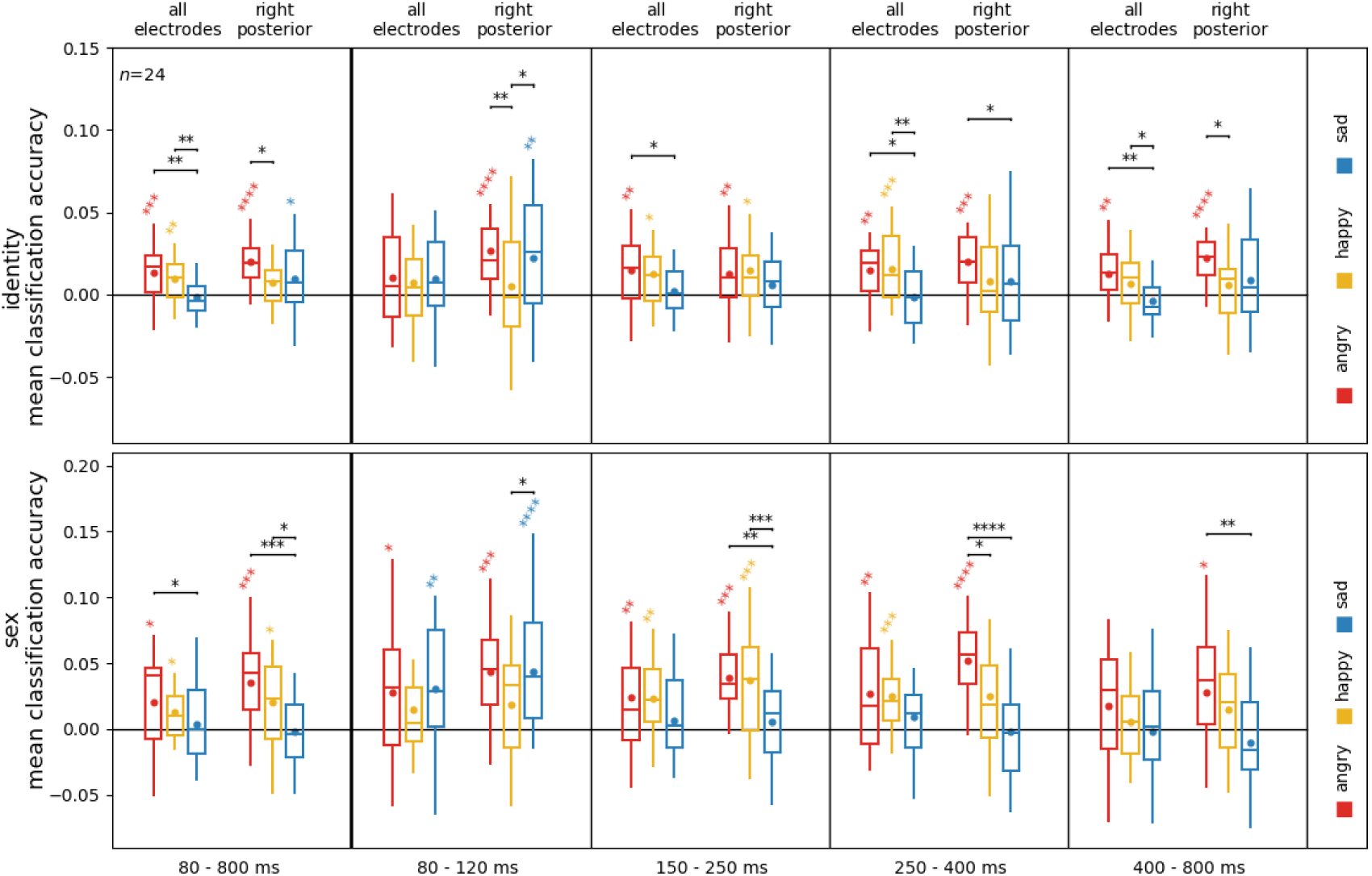
Sex and identity processing in different emotional expressions for all electrodes and the right posterior region of interest. The same data as in Figure 7, but time-resolved data averaged in different time bins for better visualization. Boxplots depict first, second (median), and third quartiles; whiskers with Q1 − 1.5 IQR and Q3 + 1.5 IQR. The dot denotes the mean. Asterisks represent uncorrected results of one-sample t-tests against chance, and paired sample t-test between emotion conditions. \**p* <0.05, \*\**p* <0.01, \*\*\**p*<0.001, \*\*\*\**p*<0.0001.

## Discussion

Our study investigated the neural dynamics of emotional face processing through an explicit facial emotion recognition task employing a two-alternative forced choice decision on the displayed emotion. We employed cross-participant multivariate classification and representational dissimilarity analysis on EEG data to probe the effects of stimulus identity, sex, and emotion, including pairs of emotions. Additionally, we explored the impact of visual image properties on these processing dynamics. The key findings of our study were the following: 1) The onset of sex and identity processing occurs early, approximately around 100 ms, but these effects are largely influenced by image properties. 2) In contrast, facial expression effects, commencing at approximately 120 ms and peaking between 200 and 250 ms, are not significantly influenced by visual image properties. 3) This independence from visual image properties also applies to analyses involving pairs of emotional expressions, with peaks occurring between 100 and 275 ms, suggesting image-independent processing of emotional expressions. 4) The temporal sequence of emotional expression processing follows the order of happy–neutral, angry–neutral, sad–neutral, followed by happy–angry, happy–sad, and finally angry–sad and 5) Identity and sex processing are enhanced for angry and happy expressions compared to sad and neutral faces.

### The processing of facial identity and sex occurs early and is image-dependent

Our findings align with prior research regarding the onset of face identity and sex effects, indicating a pattern where sex-related effects precede those related to identity in alignment with the coarse-to-fine account of face perception (Goffaux et al., 2011). Using cross-participant multivariate patten analysis, we also extended this research by examining the neural patterns associated with these attributes shared across participants.

The earliest effects of face identity have been shown to unfold before or around 100 ms. These early effects of face-identity tend to be highly image-dependent (Ambrus et al., 2019), as these effects exhibit a strong correlation with low-level image properties and can be diminished by implementing controls to account for these factors. Previous studies have shown that these identity effects can be seen even for previously unknown identities (Ambrus et al., 2021; Dobs et al., 2019; Vida et al., 2017). Image-independent representations tend to emerge around 200 ms, and especially 400 ms following stimulus onset (Ambrus et al., 2019; Vida et al., 2017).

Our current findings are in agreement with this pattern. After accounting for face-independent low-level visual image properties (maximum cross-correlation), only a residual identity effect persisted within the time frame of approximately 400 to 600 ms. Additionally, this remaining effect was further diminished when controlling for higher-level, face-dependent image properties, specifically those extracted from a neural network trained for face recognition. The effects of stimulus sex were also greatly reduced by controlling for image properties.

### Effects of facial expression follow that of sex and identity, and are not image-dependent

In our observations, we identified significant and robust effects related to emotional expressions in general. This effect was evident in both our time-resolved decoding and representational dissimilarity analyses, initiating around 120 ms and reaching its peak at approximately 160 ms. Importantly, the presence of this effect remained unaffected even after implementing controls for image properties, implying that this emotional expression effect is independent of visual image characteristics. This finding is concordant with those of Zhang et al. (2023), who found that face-selective regions in the lateral occipital and inferior parietal cortices represented emotional face stimuli in a categorical, rather than image-dependent manner.

In a prior study by Smith and Smith (Smith & Smith, 2019), the authors reported even earlier onsets, 31 to 90 ms, for similar effects, with a comparable peak at around 150 ms. However, their methodological description of the train-test procedures did not provide clear indications that they left an identity out when decoding facial expressions. Consequently, it is plausible that the earlier onsets they observed could be attributed to image-dependent effects. Our results are more in line with a recent EEG-RSA investigation by Li and collaborators (Y. Li et al., 2022) who reported the first emotional expression effects between 120 and 800 ms with a similar peak around 170 ms.

Evidence for image independence in the behavioral domain comes from Alais and collaborators (2021), who in a recent study found that illusory faces displayed positive serial dependence for perceived facial expression, similar to human faces, suggesting shared mechanisms for temporal continuity. Furthermore, cross-domain serial dependence of perceived expression was robust between illusory and human faces when interleaved, leading the authors to suggest a common mechanism for facial expression processing beyond human facial features. It would be interesting to see how these effects manifest on a neural level, for example by employing cross-dataset classification (Ambrus, 2024; Dalski, Kovács, & Ambrus, 2022).

### The discrimination of emotional expressions follows a distinct time course

Our findings unveiled a sequential pattern in the processing of emotional expressions. Initially, discrimination between emotional and neutral expressions took precedence. Subsequently, distinctions emerged between different emotional expressions. The temporal sequence of emotional expression processing adhered to the order of angry–neutral and happy–neutral starting around 100-120 ms, with an earlier onset but later peak for angry expressions, and sad–neutral starting at 125 ms and peaking at 165 ms. Following this, discriminations unfolded between happy–angry, happy–sad, and ultimately, angry–sad, with onsets at 155, 170, and 215 ms, respectively.

In a recent MEG study conducted by Zhang and colleagues (2023), their exploration of emotional facial expression discrimination revealed a strikingly similar temporal pattern when compared to neutral expressions. They observed distinctions between neutral and happy expressions commencing at 90 ms and peaking at 120 ms, neutral and angry expressions initiating at 80 ms and reaching a peak at 150 ms, and neutral and sad expressions initiating at 120 ms. Our present results not only replicate but also expand upon these observations by including the distinctions between different emotional expressions. In both our representational similarity and cross-classification analyses, these effects unfolded following those seen between neutral and emotional expressions. These findings are in line with the proposal that stimuli carrying emotionally significance are encoded more rapidly than neutral ones as a result of emotion-facilitated sensory processing (Schupp et al., 2003). It is noteworthy that the effects associated with sad expressions were the last to manifest in the temporal sequence when compared to neutral faces, as well as with other emotional expressions (see **Figures 3** and **4**).

### Face-identity and sex information processing is enhanced by angry and happy facial expressions

Our findings suggest that the processing of face-identity and sex information is enhanced when participants are presented with facial expressions conveying anger, and to a lesser extent, happiness. In other words, the presence of these emotional expressions appears to enhance the brain’s ability to process and distinguish information related to the identity and sex of faces, offering further evidence that these facial attributes and facial expressions are not processed entirely independently.

D’Argembeau and Van der Linden (2011) suggest that anger, and in general, negative stimuli have a tendency to narrow attention towards indicators of threat, while positive stimuli tend to expand attentional breadth (Derryberry & Tucker, 1994). Thus, they propose that angry faces tend to automatically direct attention toward facial expressions, thereby disrupting the process of elaborative processing (Vuilleumier et al., 2001). This line of argumentation suggests that the influence of facial expressions on memory for facial identity may stem from the disruptive effects of angry expressions on the processing of facial identity. In other words, negative facial expressions draw attention to the specific features conveying the emotional signals, interfering with the processing of local features more than positive expressions. Consequently, this disruption interferes with the identification of a person.

Our results appear to directly contradict this notion. Although both happy and angry expressions elicited stronger shared identity and sex-related responses compared to sad expressions, the most stable shared representations were observed for angry expressions, particularly in the more posterior regions of interest (**Figure 7**). Our findings suggest that the brain is more sensitive to face-identity and sex information when it is presented in the context of an emotional expression, particularly that of anger. This enhanced processing would predict (Craik & Lockhart, 1972) better subsequent memory for angry faces, which is at odds with studies that find a happy face advantage for future memory (D’Argembeau & Van der Linden, 2007; Liu et al., 2014; Shimamura et al., 2006). However, our findings align well with research that shows an angry face advantage for visual short-term memory (Jackson et al., 2009). In our study, subsequent memory was not tested, thus the nature of this enhanced processing, and how it relates to memory-related processing, needs to be the subject of future research.

## Summary

This study provides insights into the neural mechanisms underlying the processing of facial expressions, identity, and sex. Through the use of cross-participant multivariate classification and representational dissimilarity analysis on EEG data, we demonstrated that sex, identity, and emotional expression can be decoded from EEG signals across participants. Our findings reveal that shared neural patterns associated with facial expression processing exist, indicating a common neural code for these processes. Our results also showed that the processing of emotional versus neutral expressions occurs earlier than that of pairs of expressive stimuli, and that the effects of emotion are independent of the visual image properties, suggesting a robust mechanism for emotion recognition that transcends image-level features. The temporal sequence of decoding revealed that sex information is processed first, followed by identity, and then emotion. Furthermore, emotional expressions, particularly angry and to a lesser extent, happy faces, enhanced the processing of identity and sex, highlighting the influence of emotion on these facial attributes. These findings underscore the complexity and integration of facial expression processing within the neural architecture, underscoring the interconnected nature of affective information processing.

## CREDIT author statement

**MME**: Conceptualization, Investigation, Formal analysis, Writing - Original Draft, Writing - Review & Editing; **GGA**: Conceptualization, Methodology, Software, Data Curation, Investigation, Formal analysis, Writing - Original Draft, Writing - Review & Editing, Visualization, Project administration, Supervision

## Supplementary Materials

**Supplementary Table 1.** Representational Similarity Analysis.

**Supplementary Table 2.** Classification analyses; identity, sex, emotion; emotion pairs.

**Supplementary Table 3.** Classification analyses, identity and sex decoding for different emotion conditions.

## Supplementary Information

### Classification pipelines

The figures below illustrate the classification pipelines used in this study. They detail the processes for multivariate cross-classification and representational similarity analyses applied to the electroencephalographic data.

**Supplementary Information Figure S1.**
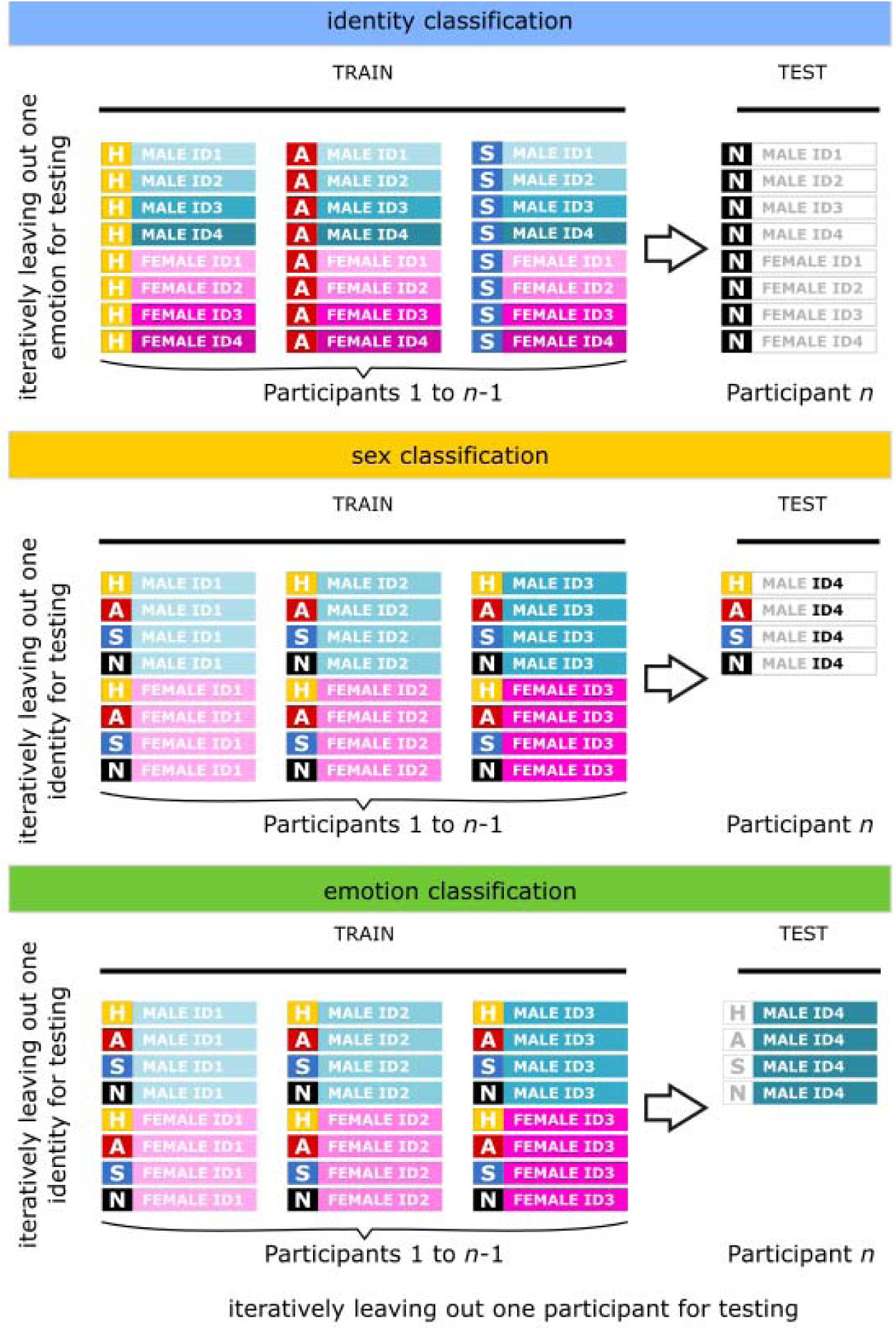
In identity classification, classifiers were iteratively trained on three emotion categories and then tested on the emotion category excluded from training. For sex and emotion classifiers, training involved six distinct identities (three male and three female) and subsequent testing on one identity omitted during training. Classification accuracies were compared against chance levels (identity: 0.125, sex: 0.5, emotion: 0.25). H: happy, A: angry, S: sad, N: neutral.

**Supplementary Information Figure S2.**
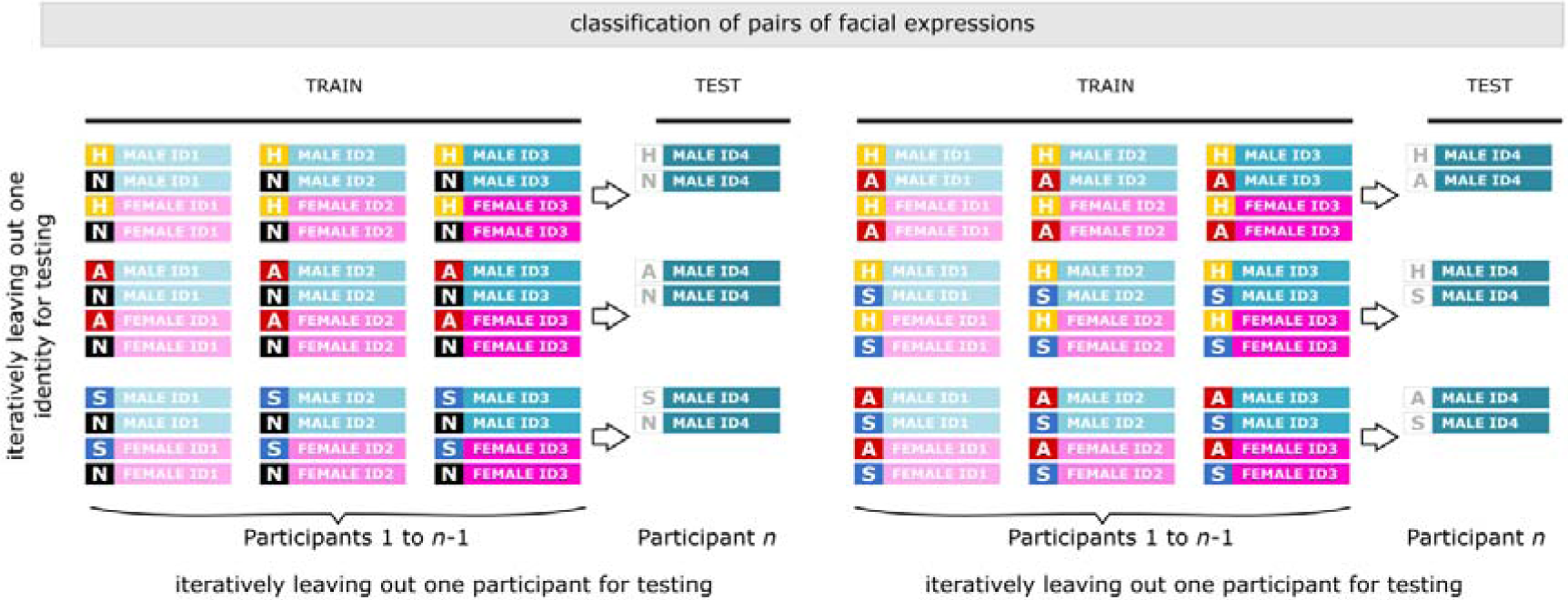
The classification of pairs of emotional expressions followed a similar approach to that of emotion in general, but with two emotional expressions included at a time. Training involved six distinct identities (three male and three female) and subsequent testing on one identity omitted during training. H: happy, A: angry, S: sad, N: neutral.

**Supplementary Information Figure S3.**
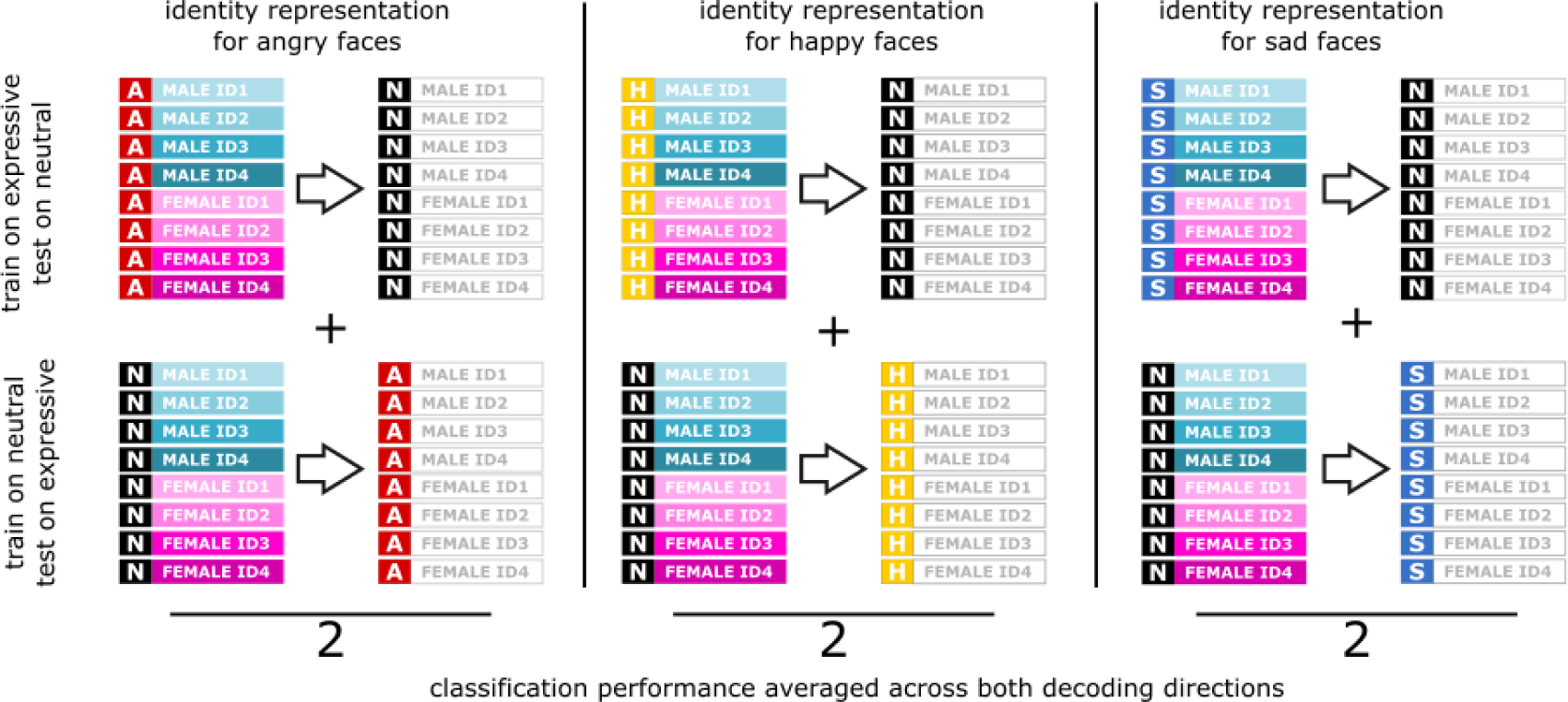
To explore the temporal dynamics of face identity and sex processing across different emotional expressions, cross-classification was implemented. This method entailed training the classifier using data from trials with neutral expressions and evaluating its performance on trials featuring one of the emotional expressions, and vice versa. The obtained classification accuracies in both directions (e.g., neutral-to-angry and angry-to-neutral) were then averaged for each participant and subjected to statistical testing against chance. H: happy, A: angry, S: sad, N: neutral.

**Supplementary Information Figure S4.**
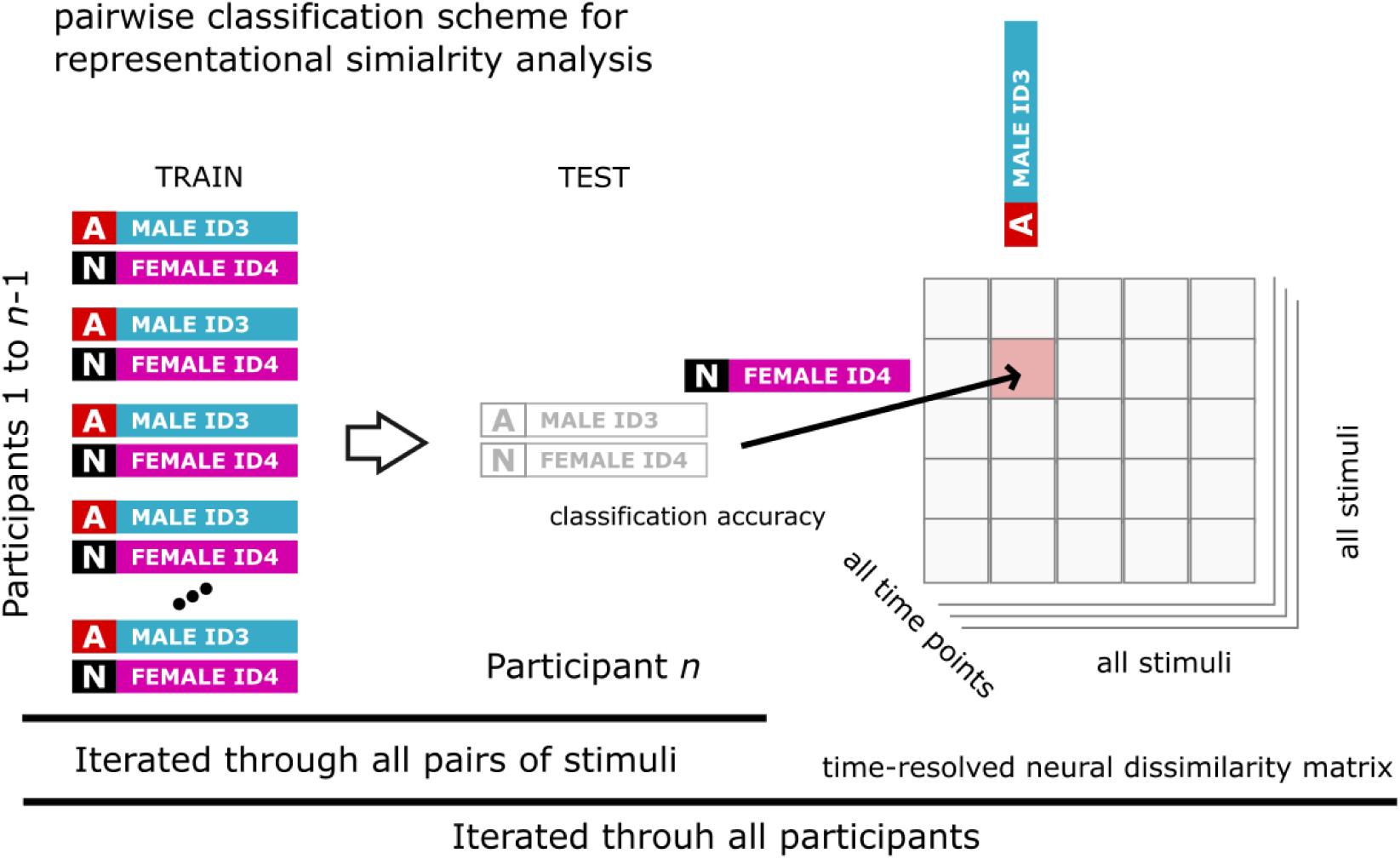
Representational similarity analyses. Representational similarity analyses involved constructing participant-level empirical neural representational dissimilarity matrices (RDMs) in each time-point for each participant. This process was achieved through the pairwise classification of stimulus pairs, resulting in matrices with dimensions 280 by 32 by 32. The classification followed a leave-one-subject-out scheme. Subsequently, these matrices were compared to predictor representational dissimilarity matrices that modeled identity, sex, and facial expression (32 by 32 matrices), as well as pairs of expressions (16 by 16 matrices), using Spearman rank correlations. The resulting correlation values were then Fishertransformed. H: happy, A: angry, S: sad, N: neutral.

### Supplementary Results

**Supplementary Figure S5.**
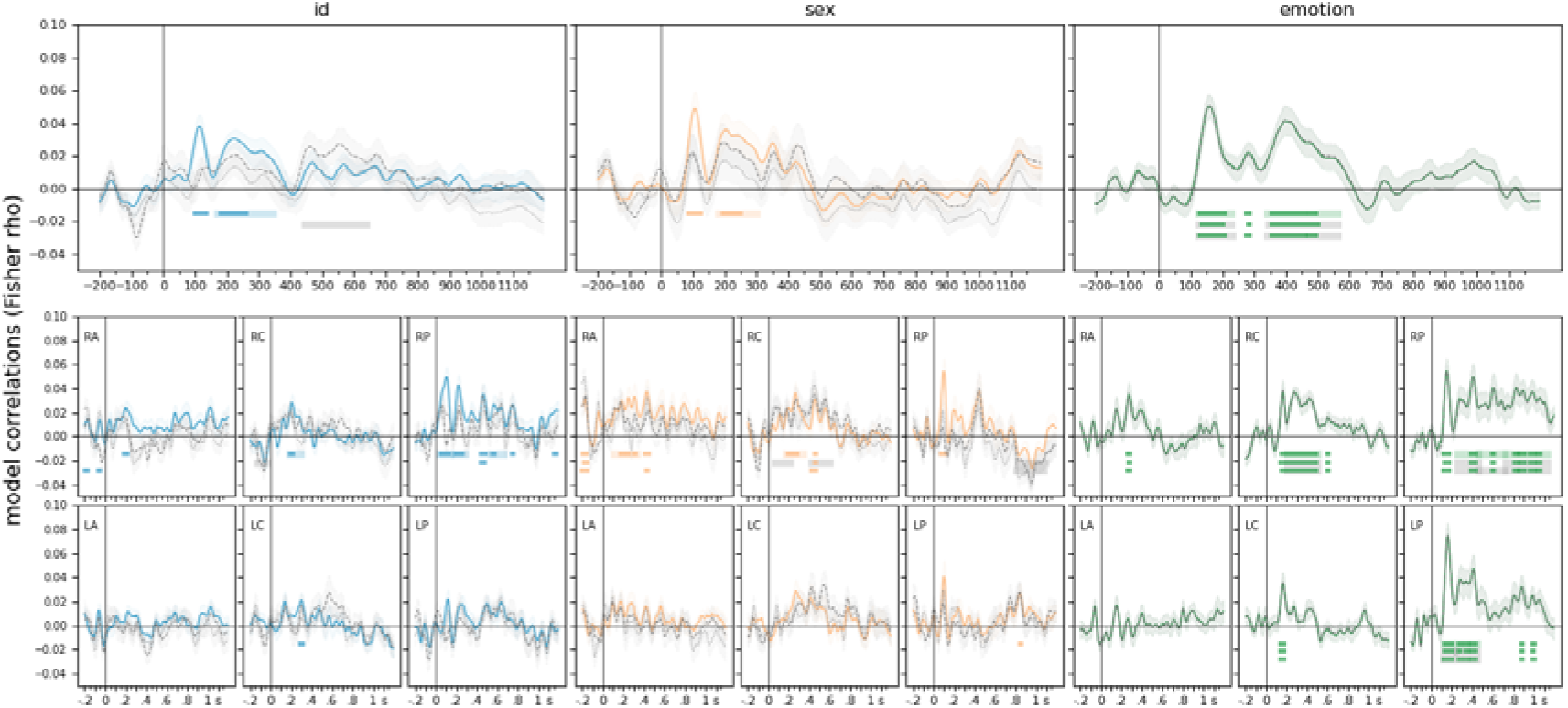
Representational Similarity Analysis for identity, sex, and facial expressions. Results for all electrodes and pre-defined regions of interest. Neural RDMs were calculated using time-resolved, leave-one-participant-out pairwise decoding of EEG data. These matrices were then assessed against model RDMs for identity, sex, and facial expression. Solid lines represent model correlations, dashed lines illustrate results when the effects of maximum cross-correlation are controlled for, and dotted lines depict results with the effects of neural network feature distance partialled out. Light lines denote significant clusters revealed by the two-sided cluster permutation tests, *p* < 0.05; dark lines denote results of the Bayesian statistical analyses, two-sided one-sample Bayesian t-tests, bf >10. Error ranges denote ± SEM. Top panels: results for all electrodes. Bottom panels: results in the six pre-defined electrode clusters separately. RA/LA: right/left anterior, RC/LC: right/left central, RP/LP: right/left posterior. For detailed statistics, see **Supplementary Table 1**.

**Supplementary Figure S6.**
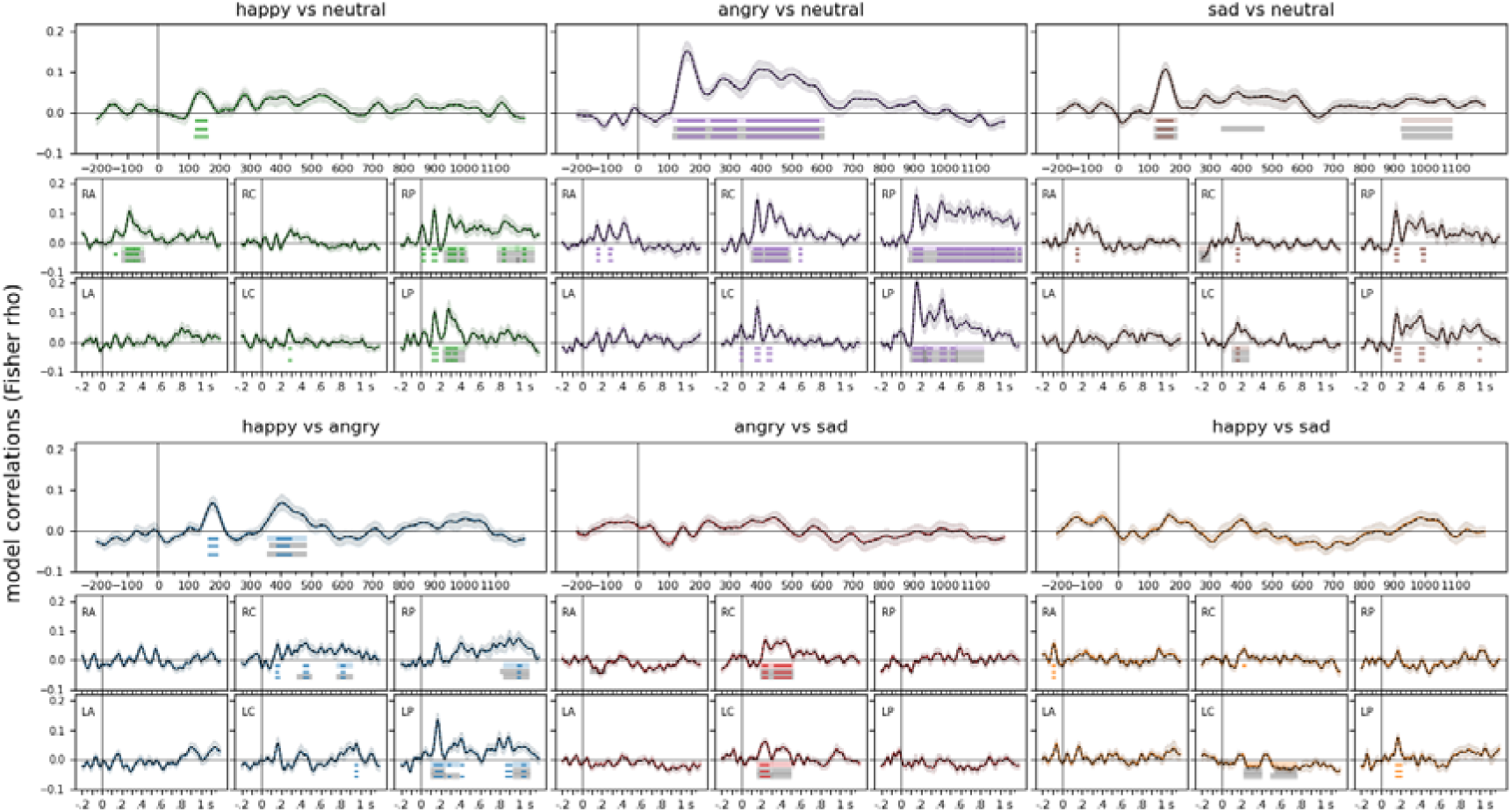
Representational Similarity Analysis for pairs of facial expressions. Results for all electrodes and pre-defined regions of interest. Neural RDMs were calculated using time-resolved, leaveone-participant-out pairwise decoding of EEG data. These matrices were then assessed against model RDMs for pairs of facial expressions. Solid lines represent model correlations, dashed lines illustrate results when the effects of maximum cross-correlation are controlled for, and dotted lines depict results with the effects of neural network feature distance partialled out. Light lines denote significant clusters revealed by the twosided cluster permutation tests, *p* < 0.05; dark lines denote results of the Bayesian statistical analyses, twosided one-sample Bayesian t-tests, bf >10. Error ranges denote ± SEM. Top panels: results for all electrodes. Bottom panels: results in the six pre-defined electrode clusters separately. RA/LA: right/left anterior, RC/LC: right/left central, RP/LP: right/left posterior. For detailed statistics, see **Supplementary Table 1**.

## Notes

### Competing Interest Statement

The authors have declared no competing interest.

