## Supplementary Table for "Shared Neural Dynamics of Facial Expression Processing": SupplementaryTable_1.html

  


### Supplementary Table 1.

  

Representational dissimilarity analysis

  

**Representational Similarity Analysis.** Time-resolved neural representational dissimilarity matrices were formed through pairwise decoding of EEG data, adopting a leave-one-participant-out approach.
These matrices were then assessed against model RDMs for identity, sex, and facial expression, along with pairs of facial expressions, utilizing rank correlations.
To control for visual image properties, partial correlations were employed. The effects of maximum cross-correlation and neural network (dlib) feature distances were partialled out.

  


**A** identity

**B** sex

**C** emotion

**D** neutral vs happy

**E** neutral vs angry

**F** neutral vs sad

**G** happy vs angry

**H** angry vs sad

**I** happy vs sad

**J** maximum cross-correlation, identity

**K** maximum cross-correlation, sex

**L** maximum cross-correlation, emotion

**M** maximum cross-correlation, neutral vs happy

**N** maximum cross-correlation, neutral vs angry

**O** maximum cross-correlation, neutral vs sad

**P** maximum cross-correlation, happy vs angry

**Q** maximum cross-correlation, angry vs sad

**R** maximum cross-correlation, happy vs sad

**S** neural network features, identity

**T** neural network features, sex

**U** neural network features, emotion

**V** neural network features, neutral vs happy

**W** neural network features, neutral vs angry

**X** neural network features, neutral vs sad

**Y** neural network features, happy vs angry

**Z** neural network features, angry vs sad

**AA** neural network features, happy vs sad

A) identity

  
|  | time window | peak latency | cluster *p* | peak Cohen's *d* |  | | | |
| **all electrodes** | 170 - 345 ms | 225 ms | 0.0039 | 1.0967 |  | | | |
|  | | | | | | | | |

Model correlations, cluster permutation tests

|  | **left hemisphere** | | | | **right hemisphere** | | | |
|  | time window | peak latency | cluster *p* | peak Cohen's *d* | time window | peak latency | cluster *p* | peak Cohen's *d* |
| **anterior** |  | | | |  | | | |
| **central** |  | | | | 180 - 290 ms | 205 ms | 0.0376 | 0.8111 |
| **posterior** |  | | | | 30 - 140 ms | 105 ms | 0.0191 | 1.2088 |
  | | | | 185 - 285 ms | 220 ms | 0.0295 | 0.9876 |  | | | | 535 - 660 ms | 570 ms | 0.0352 | 0.7086 |

  

Model correlations, Bayesian statistics

|  | -200 | -195 | -190 | -185 | -180 | -175 | -170 | -165 | -160 | -155 | -150 | -145 | -140 | -135 | -130 | -125 | -120 | -115 | -110 | -105 | -100 | -95 | -90 | -85 | -80 | -75 | -70 | -65 | -60 | -55 | -50 | -45 | -40 | -35 | -30 | -25 | -20 | -15 | -10 | -5 | 0 | 5 | 10 | 15 | 20 | 25 | 30 | 35 | 40 | 45 | 50 | 55 | 60 | 65 | 70 | 75 | 80 | 85 | 90 | 95 | 100 | 105 | 110 | 115 | 120 | 125 | 130 | 135 | 140 | 145 | 150 | 155 | 160 | 165 | 170 | 175 | 180 | 185 | 190 | 195 | 200 | 205 | 210 | 215 | 220 | 225 | 230 | 235 | 240 | 245 | 250 | 255 | 260 | 265 | 270 | 275 | 280 | 285 | 290 | 295 | 300 | 305 | 310 | 315 | 320 | 325 | 330 | 335 | 340 | 345 | 350 | 355 | 360 | 365 | 370 | 375 | 380 | 385 | 390 | 395 | 400 | 405 | 410 | 415 | 420 | 425 | 430 | 435 | 440 | 445 | 450 | 455 | 460 | 465 | 470 | 475 | 480 | 485 | 490 | 495 | 500 | 505 | 510 | 515 | 520 | 525 | 530 | 535 | 540 | 545 | 550 | 555 | 560 | 565 | 570 | 575 | 580 | 585 | 590 | 595 | 600 | 605 | 610 | 615 | 620 | 625 | 630 | 635 | 640 | 645 | 650 | 655 | 660 | 665 | 670 | 675 | 680 | 685 | 690 | 695 | 700 | 705 | 710 | 715 | 720 | 725 | 730 | 735 | 740 | 745 | 750 | 755 | 760 | 765 | 770 | 775 | 780 | 785 | 790 | 795 | 800 | 805 | 810 | 815 | 820 | 825 | 830 | 835 | 840 | 845 | 850 | 855 | 860 | 865 | 870 | 875 | 880 | 885 | 890 | 895 | 900 | 905 | 910 | 915 | 920 | 925 | 930 | 935 | 940 | 945 | 950 | 955 | 960 | 965 | 970 | 975 | 980 | 985 | 990 | 995 | 1000 | 1005 | 1010 | 1015 | 1020 | 1025 | 1030 | 1035 | 1040 | 1045 | 1050 | 1055 | 1060 | 1065 | 1070 | 1075 | 1080 | 1085 | 1090 | 1095 | 1100 | 1105 | 1110 | 1115 | 1120 | 1125 | 1130 | 1135 | 1140 | 1145 | 1150 | 1155 | 1160 | 1165 | 1170 | 1175 | 1180 | 1185 | 1190 | 1195 |
| --- | --- | --- | --- | --- | --- | --- | --- | --- | --- | --- | --- | --- | --- | --- | --- | --- | --- | --- | --- | --- | --- | --- | --- | --- | --- | --- | --- | --- | --- | --- | --- | --- | --- | --- | --- | --- | --- | --- | --- | --- | --- | --- | --- | --- | --- | --- | --- | --- | --- | --- | --- | --- | --- | --- | --- | --- | --- | --- | --- | --- | --- | --- | --- | --- | --- | --- | --- | --- | --- | --- | --- | --- | --- | --- | --- | --- | --- | --- | --- | --- | --- | --- | --- | --- | --- | --- | --- | --- | --- | --- | --- | --- | --- | --- | --- | --- | --- | --- | --- | --- | --- | --- | --- | --- | --- | --- | --- | --- | --- | --- | --- | --- | --- | --- | --- | --- | --- | --- | --- | --- | --- | --- | --- | --- | --- | --- | --- | --- | --- | --- | --- | --- | --- | --- | --- | --- | --- | --- | --- | --- | --- | --- | --- | --- | --- | --- | --- | --- | --- | --- | --- | --- | --- | --- | --- | --- | --- | --- | --- | --- | --- | --- | --- | --- | --- | --- | --- | --- | --- | --- | --- | --- | --- | --- | --- | --- | --- | --- | --- | --- | --- | --- | --- | --- | --- | --- | --- | --- | --- | --- | --- | --- | --- | --- | --- | --- | --- | --- | --- | --- | --- | --- | --- | --- | --- | --- | --- | --- | --- | --- | --- | --- | --- | --- | --- | --- | --- | --- | --- | --- | --- | --- | --- | --- | --- | --- | --- | --- | --- | --- | --- | --- | --- | --- | --- | --- | --- | --- | --- | --- | --- | --- | --- | --- | --- | --- | --- | --- | --- | --- | --- | --- | --- | --- | --- | --- | --- | --- | --- | --- | --- | --- | --- | --- | --- | --- | --- | --- | --- | --- | --- | --- | --- | --- | --- | --- | --- | --- | --- | --- |
| left anterior | 0.695754 | 0.625959 | 0.540020 | 0.411051 | 0.282831 | 0.226619 | 0.214739 | 0.235525 | 0.275586 | 0.311862 | 0.333221 | 0.394339 | 0.442304 | 0.496071 | 0.601861 | 0.648954 | 0.620681 | 0.590023 | 0.531707 | 0.461624 | 0.513158 | 0.404642 | 0.281578 | 0.217905 | 0.242045 | 0.532972 | 1.096819 | 1.546705 | 1.317050 | 1.047181 | 0.665872 | 0.330870 | 0.216482 | 0.409139 | 1.277159 | 3.130017 | 7.480395 | 8.455732 | 6.218774 | 3.142240 | 1.601099 | 0.771155 | 0.463975 | 0.358447 | 0.307715 | 0.282393 | 0.252666 | 0.233078 | 0.222411 | 0.220184 | 0.219515 | 0.223057 | 0.223261 | 0.222532 | 0.220844 | 0.218022 | 0.214787 | 0.217674 | 0.237779 | 0.276942 | 0.324448 | 0.394023 | 0.454846 | 0.455826 | 0.387592 | 0.315741 | 0.263000 | 0.235754 | 0.225496 | 0.225190 | 0.226897 | 0.232294 | 0.238322 | 0.236941 | 0.231354 | 0.227215 | 0.220421 | 0.218576 | 0.224171 | 0.243821 | 0.287349 | 0.401189 | 0.658686 | 1.093421 | 1.719499 | 2.312060 | 2.317720 | 1.591530 | 0.934491 | 0.552841 | 0.471017 | 0.465286 | 0.440192 | 0.396119 | 0.381128 | 0.357922 | 0.364779 | 0.396657 | 0.457239 | 0.513450 | 0.601687 | 0.576768 | 0.518452 | 0.465077 | 0.393891 | 0.302297 | 0.248999 | 0.218212 | 0.216413 | 0.233458 | 0.264509 | 0.314377 | 0.349949 | 0.382660 | 0.392124 | 0.392862 | 0.393790 | 0.403488 | 0.414832 | 0.420940 | 0.401448 | 0.378328 | 0.361506 | 0.333813 | 0.336187 | 0.370222 | 0.426278 | 0.547670 | 0.711738 | 0.786109 | 0.733456 | 0.594026 | 0.425435 | 0.325822 | 0.253982 | 0.220121 | 0.216494 | 0.234205 | 0.262158 | 0.273240 | 0.272815 | 0.260906 | 0.243024 | 0.224650 | 0.216294 | 0.214894 | 0.217619 | 0.218948 | 0.218879 | 0.217304 | 0.214827 | 0.224579 | 0.249172 | 0.283066 | 0.280226 | 0.252949 | 0.233426 | 0.221417 | 0.217588 | 0.219607 | 0.225367 | 0.235558 | 0.254207 | 0.275703 | 0.292710 | 0.303550 | 0.299180 | 0.293431 | 0.284089 | 0.277913 | 0.282130 | 0.309344 | 0.357400 | 0.449823 | 0.586245 | 0.758081 | 0.799774 | 0.676717 | 0.499860 | 0.359774 | 0.275796 | 0.236224 | 0.223198 | 0.221813 | 0.233490 | 0.261475 | 0.307135 | 0.355177 | 0.365736 | 0.341135 | 0.328839 | 0.334707 | 0.337057 | 0.359705 | 0.366966 | 0.348439 | 0.287097 | 0.246166 | 0.220722 | 0.214646 | 0.219369 | 0.221145 | 0.218696 | 0.214930 | 0.214741 | 0.215243 | 0.214628 | 0.215979 | 0.221498 | 0.225713 | 0.222908 | 0.216116 | 0.214628 | 0.214686 | 0.214814 | 0.218385 | 0.229858 | 0.241617 | 0.245522 | 0.237703 | 0.221012 | 0.215057 | 0.228928 | 0.266245 | 0.330134 | 0.395055 | 0.432008 | 0.459405 | 0.459362 | 0.454639 | 0.457232 | 0.451791 | 0.451734 | 0.453448 | 0.408458 | 0.355270 | 0.331706 | 0.319608 | 0.331589 | 0.382223 | 0.483979 | 0.630914 | 0.839815 | 1.114614 | 1.310700 | 1.461810 | 1.457820 | 1.407643 | 1.277082 | 1.169682 | 0.862759 | 0.692966 | 0.565777 | 0.481600 | 0.425000 | 0.424964 | 0.403818 | 0.409447 | 0.383007 | 0.350071 | 0.329229 | 0.325753 | 0.304928 | 0.292180 | 0.277793 | 0.259945 | 0.236343 | 0.220151 | 0.214635 | 0.215979 | 0.218468 | 0.218605 | 0.216031 | 0.214639 | 0.214965 | 0.215363 | 0.214979 | 0.214641 | 0.214899 | 0.215751 |
| right anterior | 0.465714 | 0.794441 | 1.401896 | 3.079983 | 4.871103 | 5.445120 | 4.481452 | 2.204145 | 1.036112 | 0.583182 | 0.388399 | 0.259080 | 0.218160 | 0.214850 | 0.215299 | 0.214960 | 0.214723 | 0.214657 | 0.219043 | 0.233718 | 0.253464 | 0.279430 | 0.284840 | 0.249414 | 0.215510 | 0.278052 | 0.852912 | 3.249592 | 5.672031 | 4.117535 | 1.648776 | 0.809235 | 0.469644 | 0.320510 | 0.242532 | 0.216599 | 0.226180 | 0.309209 | 0.409354 | 0.386564 | 0.341807 | 0.268585 | 0.226431 | 0.214627 | 0.234582 | 0.319565 | 0.507161 | 0.933268 | 1.525940 | 1.495682 | 1.020381 | 0.550856 | 0.330968 | 0.260715 | 0.230446 | 0.220983 | 0.221985 | 0.228369 | 0.240811 | 0.275264 | 0.327170 | 0.424795 | 0.605529 | 0.886447 | 1.335654 | 1.790287 | 2.248117 | 2.243172 | 1.702406 | 1.248492 | 1.002329 | 0.864372 | 0.898127 | 1.037016 | 1.373956 | 2.287102 | 5.452268 | 15.733572 | 46.896862 | 92.037624 | 103.581248 | 84.983611 | 41.767775 | 19.188824 | 9.084250 | 5.216836 | 3.247988 | 2.539164 | 2.021976 | 1.709090 | 1.425447 | 1.092458 | 0.743932 | 0.516323 | 0.391979 | 0.331492 | 0.328035 | 0.340007 | 0.380129 | 0.412561 | 0.396218 | 0.352036 | 0.333069 | 0.304988 | 0.276699 | 0.260846 | 0.247669 | 0.237967 | 0.236297 | 0.244909 | 0.257645 | 0.278331 | 0.297236 | 0.311818 | 0.354425 | 0.402442 | 0.415873 | 0.430499 | 0.473158 | 0.444491 | 0.386590 | 0.332445 | 0.306581 | 0.296779 | 0.299104 | 0.314426 | 0.342690 | 0.359867 | 0.365862 | 0.356808 | 0.334209 | 0.303864 | 0.263254 | 0.233608 | 0.219666 | 0.216833 | 0.220919 | 0.243781 | 0.291470 | 0.362297 | 0.412887 | 0.419445 | 0.388846 | 0.339966 | 0.306254 | 0.286471 | 0.288181 | 0.296658 | 0.316103 | 0.317555 | 0.322478 | 0.301759 | 0.273426 | 0.250326 | 0.245773 | 0.244519 | 0.258989 | 0.275657 | 0.312135 | 0.348593 | 0.367033 | 0.355519 | 0.337300 | 0.299713 | 0.272814 | 0.249109 | 0.233126 | 0.227289 | 0.231291 | 0.249600 | 0.313082 | 0.450363 | 0.774138 | 1.515568 | 2.908288 | 4.502417 | 5.929350 | 5.185140 | 3.162339 | 1.660600 | 0.934361 | 0.602931 | 0.497798 | 0.485317 | 0.524046 | 0.589154 | 0.724973 | 0.879636 | 0.951669 | 0.986872 | 0.968238 | 0.973193 | 1.005667 | 1.021083 | 1.011296 | 1.162564 | 1.261353 | 1.301144 | 1.452898 | 1.644420 | 1.859750 | 2.201197 | 2.243358 | 1.899247 | 1.496421 | 0.970588 | 0.622610 | 0.453365 | 0.357781 | 0.304290 | 0.282962 | 0.277535 | 0.285644 | 0.298982 | 0.314399 | 0.334248 | 0.365156 | 0.393536 | 0.416907 | 0.433640 | 0.462413 | 0.476580 | 0.511133 | 0.562174 | 0.715435 | 0.935633 | 1.124845 | 1.084204 | 0.919679 | 0.674964 | 0.545434 | 0.456281 | 0.414343 | 0.423969 | 0.471621 | 0.518826 | 0.654978 | 0.826542 | 1.114533 | 1.552587 | 2.054821 | 2.703921 | 3.832773 | 4.529102 | 4.918945 | 4.899701 | 3.960876 | 2.892331 | 2.143445 | 1.531706 | 1.047298 | 0.735330 | 0.536661 | 0.428440 | 0.364957 | 0.326200 | 0.313456 | 0.329911 | 0.347362 | 0.359472 | 0.368605 | 0.375996 | 0.372946 | 0.398356 | 0.469931 | 0.672038 | 1.148119 | 2.095126 | 3.012780 | 2.766565 | 1.730953 | 0.985984 | 0.667654 | 0.572527 | 0.537491 | 0.572863 | 0.647173 | 0.750100 | 0.760171 | 0.796357 |
| left central | 0.215206 | 0.215784 | 0.216034 | 0.216489 | 0.222175 | 0.256679 | 0.342888 | 0.543697 | 0.757024 | 0.661093 | 0.472812 | 0.408414 | 0.340568 | 0.294552 | 0.290789 | 0.282210 | 0.273438 | 0.267421 | 0.256946 | 0.255305 | 0.257166 | 0.239319 | 0.246920 | 0.262610 | 0.269847 | 0.267435 | 0.261577 | 0.242558 | 0.238463 | 0.224124 | 0.216954 | 0.214714 | 0.223603 | 0.242723 | 0.258145 | 0.255854 | 0.226289 | 0.214642 | 0.223113 | 0.241349 | 0.240410 | 0.223918 | 0.215516 | 0.216271 | 0.223706 | 0.227203 | 0.232868 | 0.240513 | 0.250373 | 0.261225 | 0.268493 | 0.275810 | 0.283553 | 0.277334 | 0.260342 | 0.244396 | 0.226568 | 0.221401 | 0.217883 | 0.214690 | 0.222959 | 0.247036 | 0.336585 | 0.557955 | 0.932654 | 1.429234 | 1.976680 | 2.304899 | 2.406434 | 1.920914 | 1.399875 | 0.835265 | 0.538992 | 0.398470 | 0.351534 | 0.352915 | 0.387837 | 0.439973 | 0.488877 | 0.476427 | 0.430828 | 0.415169 | 0.382526 | 0.335570 | 0.318483 | 0.300214 | 0.297779 | 0.305504 | 0.322749 | 0.364784 | 0.479760 | 0.591482 | 0.766804 | 1.032462 | 1.384314 | 1.779380 | 2.858746 | 4.703138 | 8.060807 | 11.850104 | 18.806375 | 23.789251 | 18.197447 | 9.052456 | 3.472834 | 1.161341 | 0.538574 | 0.338154 | 0.260345 | 0.231618 | 0.217604 | 0.214860 | 0.219388 | 0.222089 | 0.221917 | 0.215533 | 0.218359 | 0.252143 | 0.326056 | 0.408822 | 0.433348 | 0.426807 | 0.383610 | 0.330795 | 0.283850 | 0.255621 | 0.245693 | 0.247222 | 0.259720 | 0.294682 | 0.364204 | 0.465788 | 0.589628 | 0.699308 | 0.741811 | 0.668751 | 0.539202 | 0.435168 | 0.358676 | 0.307099 | 0.275310 | 0.250072 | 0.231231 | 0.219388 | 0.214765 | 0.215915 | 0.217651 | 0.217649 | 0.215126 | 0.215531 | 0.222779 | 0.241774 | 0.274823 | 0.304085 | 0.310965 | 0.287930 | 0.253589 | 0.228176 | 0.215794 | 0.215872 | 0.220645 | 0.222506 | 0.220060 | 0.215996 | 0.214930 | 0.222400 | 0.232881 | 0.242781 | 0.248318 | 0.244814 | 0.231703 | 0.220519 | 0.214825 | 0.215495 | 0.216307 | 0.214784 | 0.216601 | 0.228761 | 0.244493 | 0.257266 | 0.259032 | 0.243634 | 0.221066 | 0.215154 | 0.235669 | 0.289573 | 0.345369 | 0.349185 | 0.316101 | 0.266061 | 0.234669 | 0.220484 | 0.216337 | 0.217861 | 0.228103 | 0.261077 | 0.335475 | 0.451804 | 0.485653 | 0.459515 | 0.396454 | 0.352714 | 0.336005 | 0.331536 | 0.329848 | 0.368128 | 0.447633 | 0.512142 | 0.631154 | 0.759787 | 0.760226 | 0.548666 | 0.361338 | 0.260041 | 0.227428 | 0.216074 | 0.214882 | 0.219067 | 0.243101 | 0.319185 | 0.484074 | 0.866592 | 1.553790 | 2.839374 | 4.429032 | 5.791580 | 5.377652 | 4.363190 | 3.088050 | 2.411751 | 2.011899 | 1.864922 | 1.979114 | 2.254110 | 2.649319 | 3.591303 | 4.334204 | 3.929885 | 2.591588 | 1.254429 | 0.596381 | 0.339607 | 0.250979 | 0.220839 | 0.214998 | 0.214675 | 0.214765 | 0.214668 | 0.214686 | 0.214724 | 0.215166 | 0.216060 | 0.217518 | 0.218255 | 0.219213 | 0.219516 | 0.221289 | 0.224916 | 0.235051 | 0.252800 | 0.278077 | 0.308538 | 0.342074 | 0.349586 | 0.349741 | 0.344908 | 0.338025 | 0.335738 | 0.370925 | 0.438389 | 0.616920 | 1.186851 | 3.050248 | 7.287948 | 12.159360 | 13.297670 | 9.680092 | 5.896219 | 4.630313 | 3.689868 |
| right central | 0.216478 | 0.227045 | 0.242388 | 0.294340 | 0.336342 | 0.340470 | 0.325657 | 0.307505 | 0.293957 | 0.335262 | 0.460147 | 0.789154 | 1.381073 | 1.397483 | 0.714585 | 0.411746 | 0.306908 | 0.250784 | 0.240659 | 0.255994 | 0.298300 | 0.372224 | 0.603899 | 1.007064 | 1.761270 | 3.432918 | 9.902245 | 34.069944 | 65.505980 | 46.751871 | 18.330473 | 8.840649 | 4.534332 | 2.053212 | 1.091895 | 0.575904 | 0.331204 | 0.232471 | 0.215621 | 0.237741 | 0.269147 | 0.331731 | 0.384105 | 0.374199 | 0.361105 | 0.365016 | 0.370213 | 0.333010 | 0.249907 | 0.215566 | 0.226762 | 0.283902 | 0.379890 | 0.482496 | 0.484372 | 0.396860 | 0.295215 | 0.239271 | 0.215042 | 0.222700 | 0.265870 | 0.370628 | 0.659713 | 1.321823 | 2.452796 | 2.561903 | 1.620770 | 0.796591 | 0.422381 | 0.288456 | 0.254171 | 0.253110 | 0.287229 | 0.366023 | 0.531156 | 0.914112 | 1.938999 | 4.207216 | 11.805176 | 34.792283 | 73.396231 | 72.266640 | 48.227829 | 22.685315 | 11.324399 | 5.713666 | 3.288708 | 2.210312 | 1.837105 | 1.635506 | 1.679420 | 2.127506 | 2.997838 | 3.931808 | 4.921497 | 5.527695 | 4.835563 | 3.393123 | 2.028946 | 1.060138 | 0.552998 | 0.335374 | 0.259076 | 0.231188 | 0.222991 | 0.225203 | 0.233933 | 0.237148 | 0.239719 | 0.244197 | 0.243335 | 0.241384 | 0.242296 | 0.249381 | 0.277087 | 0.323333 | 0.373151 | 0.423618 | 0.411584 | 0.300530 | 0.223369 | 0.234584 | 0.393678 | 0.671857 | 0.822000 | 0.723355 | 0.501481 | 0.336853 | 0.250769 | 0.215929 | 0.224156 | 0.258702 | 0.323106 | 0.390699 | 0.397925 | 0.332577 | 0.266252 | 0.224989 | 0.214629 | 0.217589 | 0.215858 | 0.214767 | 0.220655 | 0.232141 | 0.235592 | 0.236958 | 0.232485 | 0.223314 | 0.220757 | 0.225031 | 0.232512 | 0.247541 | 0.271953 | 0.288050 | 0.302258 | 0.305201 | 0.295680 | 0.301314 | 0.317409 | 0.317566 | 0.316835 | 0.317457 | 0.306919 | 0.293760 | 0.275860 | 0.253610 | 0.237040 | 0.222022 | 0.214640 | 0.222246 | 0.237495 | 0.255537 | 0.269438 | 0.270588 | 0.253572 | 0.243179 | 0.234937 | 0.232256 | 0.235649 | 0.247858 | 0.262682 | 0.290307 | 0.310889 | 0.295668 | 0.292137 | 0.293114 | 0.280976 | 0.278074 | 0.270007 | 0.245794 | 0.224590 | 0.214920 | 0.219577 | 0.222883 | 0.218562 | 0.214626 | 0.222479 | 0.249386 | 0.290373 | 0.335707 | 0.368319 | 0.350641 | 0.301181 | 0.258982 | 0.228778 | 0.216656 | 0.214725 | 0.215714 | 0.217045 | 0.216439 | 0.215167 | 0.214660 | 0.216387 | 0.222568 | 0.231829 | 0.245683 | 0.257932 | 0.253384 | 0.244069 | 0.233926 | 0.222526 | 0.215414 | 0.215380 | 0.222512 | 0.235060 | 0.251131 | 0.263130 | 0.255330 | 0.231014 | 0.216686 | 0.216753 | 0.232517 | 0.255099 | 0.270471 | 0.268847 | 0.251653 | 0.237415 | 0.226437 | 0.217590 | 0.214649 | 0.215950 | 0.221968 | 0.233247 | 0.245486 | 0.253993 | 0.251247 | 0.242971 | 0.234283 | 0.224679 | 0.216325 | 0.215441 | 0.226596 | 0.257453 | 0.300899 | 0.369178 | 0.460452 | 0.641686 | 0.948686 | 1.461288 | 2.096157 | 2.789981 | 3.088245 | 3.327276 | 3.838793 | 3.605131 | 3.106258 | 2.805401 | 2.629689 | 2.547468 | 2.603426 | 2.301114 | 1.812383 | 1.309318 | 0.961071 | 0.795315 | 0.683925 | 0.596308 | 0.539832 | 0.507640 | 0.457732 |
| left posterior | 0.228479 | 0.224651 | 0.223605 | 0.221167 | 0.218106 | 0.217403 | 0.234476 | 0.308869 | 0.532911 | 1.123574 | 2.526381 | 2.696120 | 1.470561 | 0.653086 | 0.347383 | 0.243285 | 0.217024 | 0.216532 | 0.219619 | 0.215750 | 0.214997 | 0.223320 | 0.248476 | 0.303498 | 0.412242 | 0.662221 | 1.208232 | 2.810826 | 5.407012 | 6.134376 | 4.036413 | 1.766439 | 0.642226 | 0.317683 | 0.232090 | 0.214654 | 0.220072 | 0.225974 | 0.227795 | 0.225561 | 0.220393 | 0.218437 | 0.215517 | 0.214975 | 0.220146 | 0.236792 | 0.262698 | 0.293012 | 0.317858 | 0.333509 | 0.307558 | 0.301512 | 0.298057 | 0.266044 | 0.234090 | 0.217203 | 0.219897 | 0.251747 | 0.338119 | 0.607127 | 1.209022 | 2.481218 | 4.509636 | 5.264522 | 4.367577 | 2.797724 | 1.221158 | 0.474579 | 0.233922 | 0.248638 | 0.535697 | 1.583991 | 4.515414 | 7.112613 | 5.173756 | 2.136123 | 0.709912 | 0.322706 | 0.220923 | 0.230596 | 0.320340 | 0.505187 | 0.786572 | 1.165484 | 1.615096 | 2.216963 | 2.413302 | 2.769112 | 2.782805 | 2.846438 | 2.744073 | 2.996644 | 3.037620 | 2.871217 | 2.030417 | 1.299842 | 0.849200 | 0.605729 | 0.443603 | 0.359452 | 0.326841 | 0.326486 | 0.321212 | 0.317384 | 0.321499 | 0.310406 | 0.278677 | 0.254125 | 0.241913 | 0.241020 | 0.237843 | 0.230532 | 0.224924 | 0.217980 | 0.214725 | 0.221720 | 0.234745 | 0.245345 | 0.248703 | 0.243595 | 0.228117 | 0.218254 | 0.214985 | 0.214636 | 0.214672 | 0.216182 | 0.224749 | 0.242625 | 0.291178 | 0.390523 | 0.479989 | 0.527288 | 0.597130 | 0.716311 | 0.891477 | 1.179188 | 1.703147 | 2.641048 | 3.795985 | 5.053331 | 5.689410 | 5.344789 | 3.641321 | 2.203785 | 1.306627 | 0.789239 | 0.521684 | 0.401482 | 0.349590 | 0.338643 | 0.356841 | 0.408064 | 0.501443 | 0.575196 | 0.545952 | 0.461775 | 0.364761 | 0.296744 | 0.263070 | 0.256769 | 0.272572 | 0.324860 | 0.456480 | 0.792584 | 1.480252 | 2.287562 | 2.984119 | 3.190640 | 2.906619 | 2.508191 | 2.434300 | 2.257173 | 1.840512 | 1.145570 | 0.696658 | 0.448028 | 0.332837 | 0.280059 | 0.256742 | 0.249502 | 0.252390 | 0.256916 | 0.263162 | 0.265728 | 0.264267 | 0.260171 | 0.253107 | 0.245885 | 0.245111 | 0.241360 | 0.233919 | 0.230616 | 0.224313 | 0.217523 | 0.214733 | 0.216554 | 0.227522 | 0.242123 | 0.253212 | 0.252292 | 0.231830 | 0.217364 | 0.215139 | 0.222567 | 0.235689 | 0.248676 | 0.262173 | 0.265277 | 0.262692 | 0.246381 | 0.230183 | 0.219110 | 0.214723 | 0.218126 | 0.227827 | 0.242384 | 0.261404 | 0.275054 | 0.281462 | 0.279054 | 0.268130 | 0.250669 | 0.239534 | 0.227418 | 0.221287 | 0.217991 | 0.216692 | 0.214743 | 0.214784 | 0.214810 | 0.214800 | 0.218666 | 0.234452 | 0.274981 | 0.356143 | 0.491138 | 0.607870 | 0.719501 | 0.731373 | 0.668282 | 0.561194 | 0.479749 | 0.387284 | 0.324091 | 0.274519 | 0.243786 | 0.231449 | 0.231540 | 0.239172 | 0.257267 | 0.299929 | 0.384061 | 0.561082 | 0.859192 | 1.337377 | 1.956672 | 2.486642 | 2.143449 | 1.501308 | 0.861587 | 0.453252 | 0.283044 | 0.226226 | 0.214652 | 0.217791 | 0.218296 | 0.216631 | 0.214647 | 0.218160 | 0.231892 | 0.253236 | 0.278683 | 0.291203 | 0.294904 | 0.287463 | 0.277377 | 0.263108 | 0.252347 | 0.246099 | 0.239562 |
| right posterior | 0.270943 | 0.273559 | 0.283385 | 0.277713 | 0.257720 | 0.232189 | 0.224768 | 0.220253 | 0.227052 | 0.252679 | 0.311390 | 0.331333 | 0.316109 | 0.252520 | 0.214756 | 0.246004 | 0.326654 | 0.401912 | 0.377497 | 0.293451 | 0.228334 | 0.226051 | 0.307448 | 0.320450 | 0.274020 | 0.234748 | 0.220402 | 0.217075 | 0.215867 | 0.214668 | 0.218163 | 0.228770 | 0.243775 | 0.262307 | 0.270384 | 0.279195 | 0.269910 | 0.251272 | 0.232194 | 0.228121 | 0.229149 | 0.266513 | 0.343309 | 0.456090 | 0.599070 | 0.873647 | 1.399601 | 2.566638 | 5.185325 | 9.941544 | 15.881831 | 23.186937 | 34.115961 | 55.757606 | 108.837592 | 244.619038 | 381.904486 | 581.846422 | 925.834009 | 1595.515072 | 2705.145435 | 4509.803143 | 4759.658550 | 2276.719540 | 644.241141 | 119.928108 | 22.424180 | 4.772160 | 1.346543 | 0.530617 | 0.305222 | 0.237362 | 0.222696 | 0.225185 | 0.247905 | 0.344484 | 0.730926 | 2.396665 | 10.934780 | 57.548811 | 194.298012 | 394.224951 | 550.171067 | 504.802710 | 343.850150 | 222.998955 | 148.393334 | 94.694869 | 64.486566 | 42.859084 | 28.630857 | 16.116990 | 7.991001 | 4.332953 | 3.042395 | 2.613806 | 2.201333 | 1.730535 | 1.322134 | 0.883536 | 0.620773 | 0.543012 | 0.546641 | 0.656970 | 0.910775 | 1.327905 | 1.948135 | 2.520131 | 2.669163 | 2.989329 | 3.218943 | 3.181971 | 2.760167 | 2.214891 | 1.633706 | 1.266219 | 1.053171 | 0.972415 | 0.947704 | 0.897608 | 0.775257 | 0.651354 | 0.577377 | 0.536537 | 0.551360 | 0.676766 | 1.082302 | 2.129289 | 4.711747 | 10.645987 | 21.849711 | 35.145523 | 46.186245 | 45.579151 | 34.724389 | 22.360493 | 11.557281 | 5.319790 | 2.583732 | 1.471444 | 0.883377 | 0.616590 | 0.499178 | 0.459700 | 0.460086 | 0.528431 | 0.713221 | 1.244980 | 2.962106 | 8.954820 | 24.252745 | 38.668340 | 37.029281 | 29.161543 | 19.441756 | 11.028183 | 6.470491 | 3.810351 | 2.481848 | 1.810296 | 1.633328 | 1.592527 | 1.719999 | 1.919115 | 2.526002 | 3.323607 | 4.517780 | 5.264802 | 6.023440 | 5.401908 | 3.931295 | 2.366590 | 1.494558 | 0.865478 | 0.537956 | 0.386785 | 0.330605 | 0.313940 | 0.351676 | 0.446417 | 0.667822 | 0.985126 | 1.335949 | 1.615893 | 2.019136 | 2.609618 | 3.644508 | 5.322680 | 7.466674 | 10.421556 | 13.057132 | 14.370409 | 11.837884 | 7.763189 | 4.054274 | 2.094611 | 1.195444 | 0.739832 | 0.518840 | 0.418237 | 0.362531 | 0.321915 | 0.295471 | 0.282127 | 0.276586 | 0.276289 | 0.285520 | 0.308118 | 0.320644 | 0.321084 | 0.302828 | 0.283723 | 0.272539 | 0.271963 | 0.277456 | 0.289410 | 0.291588 | 0.284359 | 0.277341 | 0.262024 | 0.251483 | 0.249514 | 0.257056 | 0.275370 | 0.305265 | 0.345065 | 0.375997 | 0.368293 | 0.328340 | 0.271997 | 0.227215 | 0.215238 | 0.251807 | 0.363668 | 0.525990 | 0.640383 | 0.644853 | 0.502802 | 0.373010 | 0.268129 | 0.215890 | 0.249063 | 0.454118 | 1.227405 | 3.415215 | 7.359283 | 10.131662 | 8.128682 | 4.579944 | 2.238439 | 1.057703 | 0.609328 | 0.420677 | 0.347033 | 0.303503 | 0.293778 | 0.300763 | 0.348062 | 0.438975 | 0.721314 | 1.273077 | 2.350001 | 3.810843 | 5.253172 | 5.097570 | 4.982396 | 4.990091 | 5.214736 | 6.044631 | 8.706419 | 12.335450 | 17.147449 | 19.798283 | 18.733167 | 14.314714 | 12.046042 | 8.887014 | 5.807138 | 4.422007 | 3.738462 |
| all electrodes | 0.514194 | 0.424800 | 0.330807 | 0.260219 | 0.216896 | 0.230790 | 0.309119 | 0.373871 | 0.379521 | 0.285019 | 0.217597 | 0.267040 | 0.464125 | 0.643685 | 0.659459 | 0.685191 | 0.536980 | 0.495773 | 0.529331 | 0.661178 | 0.745578 | 0.876800 | 0.713642 | 0.636737 | 0.409270 | 0.261948 | 0.218060 | 0.215124 | 0.224159 | 0.227156 | 0.219468 | 0.215110 | 0.214659 | 0.219183 | 0.224964 | 0.226446 | 0.218683 | 0.214774 | 0.230776 | 0.254809 | 0.289919 | 0.304462 | 0.292097 | 0.261908 | 0.249240 | 0.243742 | 0.259144 | 0.300535 | 0.382004 | 0.454025 | 0.438829 | 0.364857 | 0.288390 | 0.245468 | 0.229985 | 0.234365 | 0.283326 | 0.492179 | 1.349827 | 5.755933 | 32.392327 | 156.355558 | 596.525238 | 1291.342627 | 1201.862825 | 565.554073 | 142.746001 | 16.749791 | 1.676380 | 0.460068 | 0.285687 | 0.265030 | 0.334271 | 0.574959 | 1.373825 | 4.767455 | 14.825173 | 36.892542 | 115.418565 | 267.385828 | 523.663065 | 1192.663527 | 1884.381228 | 1929.553511 | 1316.265269 | 743.655124 | 373.854426 | 245.877555 | 239.003409 | 219.924804 | 154.376705 | 66.158741 | 20.945830 | 7.776628 | 4.337044 | 2.868020 | 2.480554 | 2.452628 | 2.977735 | 3.763476 | 4.193685 | 4.330148 | 4.026772 | 3.599061 | 3.197385 | 2.941271 | 2.407943 | 2.450030 | 2.071657 | 1.711667 | 1.204007 | 0.844792 | 0.568465 | 0.415107 | 0.306042 | 0.258984 | 0.230634 | 0.216799 | 0.215905 | 0.227137 | 0.244524 | 0.260037 | 0.265338 | 0.248125 | 0.222632 | 0.214729 | 0.223884 | 0.252616 | 0.321505 | 0.456679 | 0.698386 | 1.075499 | 1.509687 | 1.822293 | 1.767145 | 1.330793 | 0.927894 | 0.648914 | 0.517688 | 0.460515 | 0.428004 | 0.401699 | 0.380588 | 0.339565 | 0.291926 | 0.272000 | 0.271878 | 0.292207 | 0.349006 | 0.462160 | 0.659583 | 1.005239 | 1.288945 | 1.405070 | 1.372895 | 1.080123 | 0.788797 | 0.637646 | 0.502598 | 0.400908 | 0.374441 | 0.365107 | 0.389572 | 0.451718 | 0.569172 | 0.723748 | 0.924752 | 0.969952 | 1.104352 | 1.289107 | 1.751376 | 2.667073 | 3.545023 | 3.149106 | 2.334741 | 1.370718 | 0.796090 | 0.462150 | 0.323876 | 0.264755 | 0.253040 | 0.261977 | 0.301091 | 0.373331 | 0.492963 | 0.640500 | 0.854336 | 1.050660 | 1.310074 | 1.199237 | 0.953784 | 0.838709 | 0.668889 | 0.472149 | 0.364813 | 0.300452 | 0.247807 | 0.223109 | 0.215358 | 0.214732 | 0.214628 | 0.215420 | 0.217320 | 0.222091 | 0.229896 | 0.232335 | 0.231486 | 0.232496 | 0.235977 | 0.244871 | 0.256227 | 0.276364 | 0.299706 | 0.330054 | 0.329879 | 0.302288 | 0.264890 | 0.240309 | 0.224061 | 0.218102 | 0.217457 | 0.224241 | 0.254836 | 0.312191 | 0.383644 | 0.465078 | 0.462790 | 0.392556 | 0.335529 | 0.284541 | 0.248089 | 0.232327 | 0.218910 | 0.214656 | 0.215876 | 0.219054 | 0.221206 | 0.218998 | 0.216665 | 0.214731 | 0.216420 | 0.220418 | 0.222356 | 0.223454 | 0.223143 | 0.220183 | 0.217200 | 0.214811 | 0.214630 | 0.214627 | 0.214636 | 0.214650 | 0.214673 | 0.214762 | 0.215193 | 0.215814 | 0.217555 | 0.219364 | 0.221221 | 0.219451 | 0.218591 | 0.218456 | 0.218671 | 0.221078 | 0.225795 | 0.230497 | 0.232843 | 0.229485 | 0.221694 | 0.217195 | 0.214734 | 0.215103 | 0.216164 | 0.217811 | 0.224293 | 0.232759 | 0.254725 | 0.302986 | 0.336344 | 0.370810 |

B) sex

  
|  | time window | peak latency | cluster *p* | peak Cohen's *d* |  | | | |
| **all electrodes** | 180 - 300 ms | 205 ms | 0.0337 | 0.7515 |  | | | |
|  | | | | | | | | |

Model correlations, cluster permutation tests

|  | **left hemisphere** | | | | **right hemisphere** | | | |
|  | time window | peak latency | cluster *p* | peak Cohen's *d* | time window | peak latency | cluster *p* | peak Cohen's *d* |
| **anterior** |  | | | | 110 - 330 ms | 305 ms | 0.0185 | 0.6209 |
| **central** |  | | | | 175 - 340 ms | 265 ms | 0.01 | 0.8236 |
| **posterior** |  | | | |  | | | |

  

Model correlations, Bayesian statistics

|  | -200 | -195 | -190 | -185 | -180 | -175 | -170 | -165 | -160 | -155 | -150 | -145 | -140 | -135 | -130 | -125 | -120 | -115 | -110 | -105 | -100 | -95 | -90 | -85 | -80 | -75 | -70 | -65 | -60 | -55 | -50 | -45 | -40 | -35 | -30 | -25 | -20 | -15 | -10 | -5 | 0 | 5 | 10 | 15 | 20 | 25 | 30 | 35 | 40 | 45 | 50 | 55 | 60 | 65 | 70 | 75 | 80 | 85 | 90 | 95 | 100 | 105 | 110 | 115 | 120 | 125 | 130 | 135 | 140 | 145 | 150 | 155 | 160 | 165 | 170 | 175 | 180 | 185 | 190 | 195 | 200 | 205 | 210 | 215 | 220 | 225 | 230 | 235 | 240 | 245 | 250 | 255 | 260 | 265 | 270 | 275 | 280 | 285 | 290 | 295 | 300 | 305 | 310 | 315 | 320 | 325 | 330 | 335 | 340 | 345 | 350 | 355 | 360 | 365 | 370 | 375 | 380 | 385 | 390 | 395 | 400 | 405 | 410 | 415 | 420 | 425 | 430 | 435 | 440 | 445 | 450 | 455 | 460 | 465 | 470 | 475 | 480 | 485 | 490 | 495 | 500 | 505 | 510 | 515 | 520 | 525 | 530 | 535 | 540 | 545 | 550 | 555 | 560 | 565 | 570 | 575 | 580 | 585 | 590 | 595 | 600 | 605 | 610 | 615 | 620 | 625 | 630 | 635 | 640 | 645 | 650 | 655 | 660 | 665 | 670 | 675 | 680 | 685 | 690 | 695 | 700 | 705 | 710 | 715 | 720 | 725 | 730 | 735 | 740 | 745 | 750 | 755 | 760 | 765 | 770 | 775 | 780 | 785 | 790 | 795 | 800 | 805 | 810 | 815 | 820 | 825 | 830 | 835 | 840 | 845 | 850 | 855 | 860 | 865 | 870 | 875 | 880 | 885 | 890 | 895 | 900 | 905 | 910 | 915 | 920 | 925 | 930 | 935 | 940 | 945 | 950 | 955 | 960 | 965 | 970 | 975 | 980 | 985 | 990 | 995 | 1000 | 1005 | 1010 | 1015 | 1020 | 1025 | 1030 | 1035 | 1040 | 1045 | 1050 | 1055 | 1060 | 1065 | 1070 | 1075 | 1080 | 1085 | 1090 | 1095 | 1100 | 1105 | 1110 | 1115 | 1120 | 1125 | 1130 | 1135 | 1140 | 1145 | 1150 | 1155 | 1160 | 1165 | 1170 | 1175 | 1180 | 1185 | 1190 | 1195 |
| --- | --- | --- | --- | --- | --- | --- | --- | --- | --- | --- | --- | --- | --- | --- | --- | --- | --- | --- | --- | --- | --- | --- | --- | --- | --- | --- | --- | --- | --- | --- | --- | --- | --- | --- | --- | --- | --- | --- | --- | --- | --- | --- | --- | --- | --- | --- | --- | --- | --- | --- | --- | --- | --- | --- | --- | --- | --- | --- | --- | --- | --- | --- | --- | --- | --- | --- | --- | --- | --- | --- | --- | --- | --- | --- | --- | --- | --- | --- | --- | --- | --- | --- | --- | --- | --- | --- | --- | --- | --- | --- | --- | --- | --- | --- | --- | --- | --- | --- | --- | --- | --- | --- | --- | --- | --- | --- | --- | --- | --- | --- | --- | --- | --- | --- | --- | --- | --- | --- | --- | --- | --- | --- | --- | --- | --- | --- | --- | --- | --- | --- | --- | --- | --- | --- | --- | --- | --- | --- | --- | --- | --- | --- | --- | --- | --- | --- | --- | --- | --- | --- | --- | --- | --- | --- | --- | --- | --- | --- | --- | --- | --- | --- | --- | --- | --- | --- | --- | --- | --- | --- | --- | --- | --- | --- | --- | --- | --- | --- | --- | --- | --- | --- | --- | --- | --- | --- | --- | --- | --- | --- | --- | --- | --- | --- | --- | --- | --- | --- | --- | --- | --- | --- | --- | --- | --- | --- | --- | --- | --- | --- | --- | --- | --- | --- | --- | --- | --- | --- | --- | --- | --- | --- | --- | --- | --- | --- | --- | --- | --- | --- | --- | --- | --- | --- | --- | --- | --- | --- | --- | --- | --- | --- | --- | --- | --- | --- | --- | --- | --- | --- | --- | --- | --- | --- | --- | --- | --- | --- | --- | --- | --- | --- | --- | --- | --- | --- | --- | --- | --- | --- | --- | --- | --- | --- | --- | --- | --- | --- | --- | --- |
| left anterior | 0.945998 | 0.865148 | 0.865693 | 0.723664 | 0.549853 | 0.365848 | 0.259668 | 0.215042 | 0.240729 | 0.313308 | 0.369479 | 0.403602 | 0.414924 | 0.441764 | 0.445625 | 0.374090 | 0.299071 | 0.233567 | 0.214643 | 0.221639 | 0.218398 | 0.216758 | 0.214640 | 0.220250 | 0.238822 | 0.249473 | 0.246103 | 0.238591 | 0.226504 | 0.221950 | 0.220398 | 0.230345 | 0.278896 | 0.513963 | 1.232011 | 2.553689 | 1.900873 | 0.732519 | 0.335696 | 0.231293 | 0.217436 | 0.257405 | 0.302994 | 0.312410 | 0.280715 | 0.236883 | 0.216993 | 0.215292 | 0.218541 | 0.225965 | 0.230240 | 0.232289 | 0.224928 | 0.215685 | 0.220939 | 0.255502 | 0.339881 | 0.515582 | 0.847354 | 1.412952 | 2.208236 | 2.674431 | 2.501424 | 1.839236 | 0.953523 | 0.536224 | 0.398378 | 0.384833 | 0.415729 | 0.528520 | 0.681372 | 0.869143 | 1.028898 | 1.168292 | 1.252151 | 1.365138 | 1.304910 | 1.046661 | 0.764531 | 0.540384 | 0.399580 | 0.315008 | 0.272993 | 0.252096 | 0.253451 | 0.274001 | 0.324319 | 0.414147 | 0.557460 | 0.712290 | 0.809665 | 0.877170 | 0.886403 | 0.834296 | 0.860811 | 0.988084 | 1.109729 | 1.096765 | 0.960356 | 0.738343 | 0.550972 | 0.385602 | 0.307327 | 0.269993 | 0.258901 | 0.248737 | 0.252016 | 0.248114 | 0.243438 | 0.232482 | 0.221153 | 0.214638 | 0.215310 | 0.215265 | 0.214681 | 0.221005 | 0.245882 | 0.300905 | 0.406234 | 0.515716 | 0.601851 | 0.770273 | 0.916112 | 0.965518 | 0.939719 | 0.820384 | 0.640948 | 0.477650 | 0.364666 | 0.294144 | 0.249671 | 0.223428 | 0.214645 | 0.221837 | 0.247043 | 0.276646 | 0.287693 | 0.264615 | 0.232527 | 0.214905 | 0.222016 | 0.265308 | 0.349309 | 0.419415 | 0.435276 | 0.442027 | 0.389837 | 0.391969 | 0.411568 | 0.444413 | 0.558242 | 0.831399 | 1.224637 | 1.595306 | 1.482452 | 0.925747 | 0.513797 | 0.294062 | 0.227242 | 0.215831 | 0.214641 | 0.214697 | 0.216237 | 0.224029 | 0.234426 | 0.246154 | 0.238221 | 0.224289 | 0.217415 | 0.215245 | 0.215284 | 0.218370 | 0.225530 | 0.243635 | 0.276664 | 0.307022 | 0.309235 | 0.282520 | 0.251161 | 0.224638 | 0.214669 | 0.219563 | 0.229722 | 0.245942 | 0.260233 | 0.275361 | 0.276923 | 0.257097 | 0.231443 | 0.216596 | 0.217147 | 0.236737 | 0.273652 | 0.326031 | 0.352864 | 0.359303 | 0.289578 | 0.234231 | 0.214989 | 0.231919 | 0.253461 | 0.243596 | 0.230089 | 0.216504 | 0.214748 | 0.214670 | 0.216479 | 0.222623 | 0.235752 | 0.242212 | 0.244126 | 0.236744 | 0.231262 | 0.232038 | 0.242808 | 0.262748 | 0.296069 | 0.313212 | 0.302233 | 0.265319 | 0.226951 | 0.214675 | 0.221239 | 0.228094 | 0.227682 | 0.222702 | 0.222421 | 0.227289 | 0.244532 | 0.295899 | 0.393780 | 0.467483 | 0.558608 | 0.565188 | 0.479378 | 0.361090 | 0.302062 | 0.269867 | 0.277519 | 0.305155 | 0.369821 | 0.411785 | 0.442207 | 0.443366 | 0.393961 | 0.342963 | 0.316717 | 0.311579 | 0.343939 | 0.460524 | 0.625514 | 0.901815 | 0.968868 | 0.830561 | 0.637672 | 0.502820 | 0.396540 | 0.373285 | 0.362475 | 0.389322 | 0.422700 | 0.481727 | 0.476989 | 0.420176 | 0.341878 | 0.307798 | 0.281868 | 0.264162 | 0.252209 | 0.253502 | 0.253128 | 0.252001 | 0.243678 | 0.233633 | 0.221008 | 0.215204 | 0.215513 | 0.219752 | 0.222854 | 0.224218 |
| right anterior | 5.677794 | 11.016558 | 19.480311 | 53.353673 | 129.755665 | 246.046429 | 251.836903 | 104.584276 | 33.719467 | 7.213036 | 1.668134 | 0.616981 | 0.302820 | 0.220433 | 0.218763 | 0.250156 | 0.300865 | 0.362120 | 0.383403 | 0.404781 | 0.391962 | 0.415087 | 0.401241 | 0.337870 | 0.277792 | 0.238483 | 0.215318 | 0.311224 | 0.782153 | 1.756133 | 3.254948 | 6.628922 | 9.292236 | 6.042143 | 3.306794 | 1.778863 | 0.704527 | 0.396101 | 0.352788 | 0.383944 | 0.460511 | 0.537449 | 0.558111 | 0.502353 | 0.449109 | 0.380755 | 0.379581 | 0.418499 | 0.499973 | 0.628283 | 0.916778 | 1.176778 | 1.349463 | 1.505724 | 1.331580 | 0.773157 | 0.503208 | 0.401929 | 0.390899 | 0.473248 | 0.706342 | 1.037635 | 1.903134 | 3.430367 | 5.355646 | 5.466661 | 6.385117 | 5.322791 | 3.717523 | 2.375519 | 1.884927 | 1.750966 | 2.285763 | 3.631302 | 7.754850 | 19.477277 | 42.271788 | 53.740597 | 42.130659 | 20.058952 | 8.427100 | 3.862006 | 1.971857 | 1.329518 | 1.242367 | 1.590838 | 2.622642 | 5.353309 | 12.393969 | 22.885961 | 26.533879 | 17.747997 | 9.088004 | 3.818705 | 2.047338 | 1.301483 | 1.257444 | 1.602759 | 2.602584 | 4.637811 | 7.916788 | 10.552201 | 10.560958 | 7.609902 | 4.221995 | 2.479929 | 1.499170 | 1.051144 | 0.861233 | 0.732166 | 0.672412 | 0.666501 | 0.645592 | 0.662295 | 0.754010 | 0.895387 | 1.126044 | 1.728498 | 2.612800 | 3.465664 | 4.134673 | 5.398687 | 7.408790 | 11.691273 | 17.450382 | 23.501970 | 28.525997 | 24.154887 | 13.019806 | 6.604930 | 3.358520 | 1.777244 | 1.120064 | 0.845353 | 0.707934 | 0.660844 | 0.658311 | 0.640344 | 0.622721 | 0.604215 | 0.572492 | 0.537986 | 0.549006 | 0.563596 | 0.581617 | 0.602088 | 0.626872 | 0.651509 | 0.713539 | 0.759931 | 0.819827 | 0.930540 | 1.023060 | 1.071712 | 1.122776 | 1.112624 | 1.062489 | 0.923557 | 0.720819 | 0.548703 | 0.439892 | 0.348984 | 0.292276 | 0.262842 | 0.250315 | 0.247029 | 0.257894 | 0.277632 | 0.322942 | 0.419800 | 0.587514 | 0.831855 | 1.218371 | 1.680256 | 1.986425 | 2.098860 | 2.033910 | 1.936472 | 1.709173 | 1.482978 | 1.187952 | 0.907299 | 0.644454 | 0.495061 | 0.389117 | 0.355709 | 0.374626 | 0.459253 | 0.591168 | 0.874434 | 1.247416 | 1.596928 | 1.945568 | 2.425154 | 3.110845 | 4.593098 | 6.753608 | 8.471800 | 9.489135 | 9.476081 | 7.997029 | 5.946063 | 4.159377 | 2.711642 | 1.870245 | 1.371045 | 1.183654 | 1.119655 | 1.103865 | 1.020625 | 0.918349 | 0.727735 | 0.560999 | 0.440508 | 0.373573 | 0.353808 | 0.391613 | 0.468947 | 0.609065 | 0.842666 | 1.253547 | 1.764337 | 2.666796 | 3.953745 | 6.547355 | 9.809815 | 10.898578 | 8.274858 | 5.257492 | 2.341810 | 1.051261 | 0.615437 | 0.460277 | 0.421179 | 0.465150 | 0.531895 | 0.666704 | 0.818742 | 0.898638 | 0.883073 | 0.858830 | 0.796919 | 0.874687 | 1.037140 | 1.220482 | 1.344559 | 1.266780 | 0.894578 | 0.571290 | 0.390932 | 0.291704 | 0.244455 | 0.222017 | 0.214765 | 0.216769 | 0.220987 | 0.221994 | 0.215853 | 0.217988 | 0.252026 | 0.362958 | 0.688986 | 1.415053 | 2.702676 | 4.480015 | 5.998794 | 5.756354 | 4.312168 | 2.841931 | 1.930986 | 1.355607 | 0.931265 | 0.670797 | 0.553439 | 0.494552 | 0.502682 | 0.570536 | 0.692633 | 0.736939 | 0.810184 |
| left central | 0.214901 | 0.218578 | 0.226202 | 0.242403 | 0.257770 | 0.250380 | 0.229902 | 0.214860 | 0.245789 | 0.309337 | 0.349519 | 0.372303 | 0.308514 | 0.258242 | 0.236690 | 0.226763 | 0.229933 | 0.244366 | 0.254746 | 0.255910 | 0.260292 | 0.245901 | 0.245893 | 0.244503 | 0.247243 | 0.258173 | 0.305722 | 0.318840 | 0.323446 | 0.278364 | 0.226139 | 0.220375 | 0.277331 | 0.399702 | 0.419914 | 0.359070 | 0.305183 | 0.277622 | 0.292502 | 0.375293 | 0.670163 | 1.523180 | 2.390021 | 2.481391 | 1.967421 | 1.149441 | 0.752394 | 0.624473 | 0.513336 | 0.488564 | 0.474029 | 0.478344 | 0.430916 | 0.332714 | 0.253923 | 0.221991 | 0.218847 | 0.246027 | 0.255926 | 0.250319 | 0.239042 | 0.228583 | 0.225641 | 0.223418 | 0.226241 | 0.242673 | 0.262574 | 0.285943 | 0.326378 | 0.328600 | 0.291925 | 0.262442 | 0.239362 | 0.231653 | 0.234650 | 0.242489 | 0.259202 | 0.290972 | 0.328824 | 0.375687 | 0.442246 | 0.501346 | 0.561093 | 0.612586 | 0.687987 | 0.724172 | 0.853514 | 1.099720 | 1.579707 | 2.310127 | 3.800102 | 5.273848 | 7.188400 | 8.323485 | 8.163512 | 7.195288 | 7.227502 | 7.602679 | 7.972009 | 8.154823 | 8.923851 | 8.637606 | 7.656323 | 6.688581 | 5.236833 | 3.497046 | 2.085561 | 1.310963 | 0.847378 | 0.585323 | 0.443500 | 0.362519 | 0.321091 | 0.302496 | 0.304458 | 0.324475 | 0.367567 | 0.450699 | 0.605168 | 0.840583 | 1.115674 | 1.334396 | 1.474440 | 1.378877 | 1.102523 | 0.857441 | 0.697176 | 0.593497 | 0.530527 | 0.485579 | 0.461436 | 0.446593 | 0.440901 | 0.439056 | 0.470189 | 0.486135 | 0.495757 | 0.517299 | 0.546152 | 0.562797 | 0.614134 | 0.677072 | 0.709466 | 0.669992 | 0.574205 | 0.485114 | 0.429345 | 0.395699 | 0.390818 | 0.411736 | 0.437634 | 0.440991 | 0.413152 | 0.366148 | 0.305993 | 0.246049 | 0.217100 | 0.219286 | 0.244973 | 0.281458 | 0.306920 | 0.316596 | 0.295792 | 0.266365 | 0.237909 | 0.219418 | 0.214777 | 0.219144 | 0.223584 | 0.222358 | 0.217617 | 0.214655 | 0.224029 | 0.248931 | 0.275202 | 0.291884 | 0.287475 | 0.260315 | 0.241256 | 0.227447 | 0.219894 | 0.218205 | 0.218854 | 0.223948 | 0.236377 | 0.252228 | 0.262440 | 0.262381 | 0.234304 | 0.214965 | 0.234852 | 0.317771 | 0.481376 | 0.697078 | 0.927051 | 0.913463 | 0.750184 | 0.584239 | 0.482732 | 0.406369 | 0.354480 | 0.318913 | 0.290585 | 0.268628 | 0.254310 | 0.245220 | 0.242003 | 0.250229 | 0.261445 | 0.273247 | 0.287674 | 0.298187 | 0.290433 | 0.272777 | 0.241657 | 0.219138 | 0.215107 | 0.226466 | 0.258971 | 0.294478 | 0.322996 | 0.349441 | 0.366473 | 0.388277 | 0.412550 | 0.448842 | 0.489869 | 0.539085 | 0.524253 | 0.506833 | 0.476595 | 0.432539 | 0.364859 | 0.302841 | 0.254361 | 0.227551 | 0.216966 | 0.214735 | 0.218205 | 0.226401 | 0.240623 | 0.261662 | 0.275536 | 0.278310 | 0.265129 | 0.242692 | 0.224139 | 0.215770 | 0.214763 | 0.215901 | 0.215138 | 0.214730 | 0.217712 | 0.223326 | 0.232098 | 0.235030 | 0.236542 | 0.235312 | 0.233365 | 0.229741 | 0.230183 | 0.228524 | 0.228582 | 0.229890 | 0.229230 | 0.225191 | 0.222789 | 0.219731 | 0.215875 | 0.214701 | 0.217931 | 0.228059 | 0.243882 | 0.258235 | 0.274048 | 0.293205 | 0.316003 | 0.337128 | 0.359545 | 0.384296 |
| right central | 0.578170 | 0.528753 | 0.421948 | 0.292960 | 0.231056 | 0.214987 | 0.239376 | 0.328210 | 0.566116 | 1.051962 | 1.993772 | 3.645157 | 4.818834 | 2.851451 | 1.307570 | 0.709979 | 0.511009 | 0.451665 | 0.473983 | 0.694005 | 1.456374 | 3.180902 | 3.417766 | 2.265975 | 0.829513 | 0.345833 | 0.227009 | 0.218112 | 0.252428 | 0.295933 | 0.296686 | 0.268204 | 0.232312 | 0.215094 | 0.216881 | 0.221871 | 0.230448 | 0.237744 | 0.245154 | 0.255018 | 0.280208 | 0.280503 | 0.259812 | 0.233547 | 0.217044 | 0.229329 | 0.311168 | 0.455257 | 0.573339 | 0.598776 | 0.511858 | 0.400550 | 0.312907 | 0.259052 | 0.239919 | 0.242978 | 0.264975 | 0.305711 | 0.419154 | 0.599402 | 0.909640 | 1.375383 | 1.992323 | 2.119251 | 1.921062 | 1.313128 | 0.826852 | 0.500675 | 0.350808 | 0.274866 | 0.271114 | 0.310998 | 0.438436 | 0.660983 | 1.132698 | 1.980219 | 4.162818 | 8.749971 | 19.784286 | 30.881244 | 30.801098 | 17.958922 | 10.320450 | 5.741264 | 4.018432 | 3.243855 | 3.203108 | 3.635478 | 4.922277 | 7.400611 | 12.718718 | 26.252878 | 54.764244 | 86.478399 | 88.466935 | 54.216159 | 22.804673 | 9.838935 | 5.030309 | 3.239919 | 2.565579 | 2.320936 | 2.473574 | 2.865940 | 3.123399 | 3.049953 | 2.482380 | 1.887712 | 1.508066 | 1.108558 | 0.816794 | 0.680573 | 0.616548 | 0.594606 | 0.579905 | 0.514096 | 0.413265 | 0.350826 | 0.298486 | 0.267462 | 0.256199 | 0.258620 | 0.262045 | 0.287787 | 0.357278 | 0.518283 | 0.848612 | 1.472875 | 2.451819 | 4.568853 | 8.401517 | 13.777780 | 17.024048 | 14.858991 | 8.530221 | 4.628815 | 2.568025 | 1.620876 | 1.015531 | 0.693761 | 0.504647 | 0.418813 | 0.413248 | 0.463075 | 0.527923 | 0.690098 | 0.964707 | 1.234545 | 1.611382 | 1.894059 | 1.901234 | 1.838738 | 1.682672 | 1.466140 | 1.371824 | 1.266000 | 1.107594 | 0.935089 | 0.684899 | 0.442013 | 0.304642 | 0.252190 | 0.238157 | 0.243716 | 0.266730 | 0.325787 | 0.441205 | 0.588756 | 0.678465 | 0.631629 | 0.508961 | 0.364843 | 0.267068 | 0.221945 | 0.215749 | 0.234238 | 0.271234 | 0.351611 | 0.457552 | 0.552001 | 0.630573 | 0.585424 | 0.480523 | 0.400686 | 0.352244 | 0.296092 | 0.257957 | 0.228092 | 0.214932 | 0.221583 | 0.246146 | 0.295139 | 0.363320 | 0.444534 | 0.541066 | 0.607943 | 0.563222 | 0.464372 | 0.377849 | 0.303004 | 0.255514 | 0.234009 | 0.223695 | 0.217886 | 0.216855 | 0.218663 | 0.223819 | 0.226915 | 0.225241 | 0.220449 | 0.215381 | 0.218387 | 0.240239 | 0.288334 | 0.366837 | 0.475315 | 0.555257 | 0.558103 | 0.488873 | 0.407906 | 0.336028 | 0.277280 | 0.239129 | 0.220440 | 0.214682 | 0.217960 | 0.222915 | 0.222704 | 0.216182 | 0.216517 | 0.243087 | 0.301103 | 0.369109 | 0.399061 | 0.376719 | 0.328700 | 0.281611 | 0.245270 | 0.222259 | 0.214750 | 0.218037 | 0.229254 | 0.250563 | 0.272332 | 0.284748 | 0.279149 | 0.264953 | 0.246720 | 0.235545 | 0.228433 | 0.227060 | 0.228404 | 0.230086 | 0.232597 | 0.235704 | 0.237332 | 0.233841 | 0.225407 | 0.216009 | 0.215102 | 0.217744 | 0.220920 | 0.224534 | 0.226460 | 0.226946 | 0.224953 | 0.220316 | 0.216023 | 0.214674 | 0.216193 | 0.217090 | 0.216585 | 0.215414 | 0.214934 | 0.220176 | 0.231666 | 0.248517 | 0.275583 | 0.289608 | 0.294315 |
| left posterior | 0.570781 | 0.583264 | 0.591679 | 0.644454 | 0.644753 | 0.523888 | 0.339480 | 0.247976 | 0.214668 | 0.269435 | 0.465207 | 0.730579 | 0.779995 | 0.502434 | 0.291853 | 0.224597 | 0.215289 | 0.238244 | 0.261372 | 0.256602 | 0.252844 | 0.252222 | 0.246650 | 0.232380 | 0.219121 | 0.214706 | 0.216371 | 0.224266 | 0.237089 | 0.257333 | 0.304158 | 0.388605 | 0.480284 | 0.415141 | 0.299675 | 0.227996 | 0.222802 | 0.314065 | 0.449793 | 0.552866 | 0.512747 | 0.362185 | 0.253196 | 0.214835 | 0.242592 | 0.327593 | 0.458910 | 0.585286 | 0.633056 | 0.647574 | 0.591529 | 0.474173 | 0.325241 | 0.223315 | 0.254622 | 0.542848 | 1.478020 | 3.747861 | 6.546546 | 7.970770 | 7.037070 | 5.288535 | 3.435286 | 1.992041 | 1.014709 | 0.503219 | 0.261605 | 0.221457 | 0.453070 | 1.299848 | 1.736822 | 1.196446 | 0.702767 | 0.358470 | 0.234037 | 0.219780 | 0.270622 | 0.339413 | 0.344195 | 0.318450 | 0.283717 | 0.268797 | 0.269483 | 0.294931 | 0.358157 | 0.480194 | 0.661492 | 0.853139 | 0.887792 | 0.710222 | 0.547363 | 0.428219 | 0.356519 | 0.318572 | 0.293850 | 0.266673 | 0.247555 | 0.229352 | 0.216448 | 0.215207 | 0.217473 | 0.216720 | 0.214637 | 0.219901 | 0.240706 | 0.275402 | 0.310983 | 0.340778 | 0.349373 | 0.354360 | 0.329066 | 0.293553 | 0.260121 | 0.235681 | 0.219066 | 0.214742 | 0.215505 | 0.216968 | 0.219563 | 0.221972 | 0.222390 | 0.219466 | 0.215638 | 0.215104 | 0.223146 | 0.243756 | 0.267496 | 0.265635 | 0.251972 | 0.229771 | 0.215387 | 0.221761 | 0.249946 | 0.294908 | 0.328268 | 0.352773 | 0.357746 | 0.358760 | 0.372358 | 0.397235 | 0.413811 | 0.423359 | 0.385071 | 0.313691 | 0.262327 | 0.236436 | 0.224853 | 0.218544 | 0.216383 | 0.216645 | 0.217546 | 0.219055 | 0.220759 | 0.221709 | 0.222940 | 0.221389 | 0.217761 | 0.216452 | 0.215235 | 0.214701 | 0.214627 | 0.214686 | 0.215255 | 0.216316 | 0.220675 | 0.229164 | 0.240907 | 0.258966 | 0.286476 | 0.329089 | 0.389467 | 0.458761 | 0.459409 | 0.392421 | 0.304028 | 0.246127 | 0.220639 | 0.214748 | 0.215329 | 0.214959 | 0.215827 | 0.225676 | 0.257384 | 0.325224 | 0.418496 | 0.522666 | 0.679117 | 0.896449 | 1.185513 | 1.393551 | 1.251205 | 0.928048 | 0.625266 | 0.418642 | 0.321606 | 0.275026 | 0.254273 | 0.251688 | 0.268018 | 0.305692 | 0.408826 | 0.597680 | 0.882498 | 1.256889 | 1.852022 | 2.630623 | 4.005672 | 5.856127 | 8.305985 | 10.618318 | 12.691795 | 10.462568 | 5.980826 | 2.473665 | 0.998684 | 0.504609 | 0.345392 | 0.286842 | 0.265450 | 0.268047 | 0.281443 | 0.317740 | 0.378518 | 0.499543 | 0.661901 | 0.805109 | 0.829585 | 0.855981 | 0.807606 | 0.703929 | 0.648101 | 0.669950 | 0.769000 | 0.941633 | 1.164940 | 1.202759 | 1.116842 | 0.877374 | 0.680370 | 0.535189 | 0.434491 | 0.356833 | 0.329265 | 0.306004 | 0.291937 | 0.284835 | 0.281066 | 0.261975 | 0.246193 | 0.228802 | 0.219994 | 0.214891 | 0.217835 | 0.234777 | 0.253603 | 0.283942 | 0.285152 | 0.253626 | 0.222677 | 0.214970 | 0.228858 | 0.252185 | 0.284564 | 0.305340 | 0.304504 | 0.293467 | 0.279007 | 0.259478 | 0.243896 | 0.237097 | 0.237766 | 0.244003 | 0.257114 | 0.286055 | 0.340911 | 0.435586 | 0.563134 | 0.668531 | 0.713331 | 0.745038 |
| right posterior | 0.313050 | 0.305029 | 0.283104 | 0.245658 | 0.223503 | 0.215427 | 0.215894 | 0.220230 | 0.225739 | 0.231616 | 0.237409 | 0.228926 | 0.215438 | 0.226167 | 0.291101 | 0.356737 | 0.329308 | 0.258930 | 0.216346 | 0.234079 | 0.283008 | 0.282683 | 0.233541 | 0.225876 | 0.445517 | 1.382890 | 2.835264 | 3.156064 | 2.263411 | 1.183396 | 0.618607 | 0.326213 | 0.215654 | 0.296436 | 0.574650 | 0.869966 | 0.828248 | 0.623192 | 0.486254 | 0.369927 | 0.339247 | 0.345618 | 0.368836 | 0.393009 | 0.361275 | 0.284193 | 0.242937 | 0.218457 | 0.217568 | 0.237686 | 0.272893 | 0.345011 | 0.573600 | 1.797312 | 16.292616 | 169.831134 | 761.560102 | 1289.111360 | 1111.992379 | 981.599769 | 700.273917 | 324.894534 | 97.348413 | 21.826779 | 3.983173 | 0.881958 | 0.304538 | 0.214762 | 0.295206 | 0.437184 | 0.408354 | 0.282219 | 0.219866 | 0.227350 | 0.296676 | 0.414212 | 0.551572 | 0.686276 | 0.810931 | 0.911934 | 0.921152 | 0.885151 | 0.835671 | 0.771532 | 0.689770 | 0.610370 | 0.530914 | 0.473479 | 0.432309 | 0.445353 | 0.520349 | 0.642872 | 0.763113 | 0.850889 | 0.839903 | 0.752735 | 0.570082 | 0.388214 | 0.276907 | 0.231611 | 0.216629 | 0.214733 | 0.214743 | 0.215892 | 0.218675 | 0.223437 | 0.228549 | 0.236427 | 0.240730 | 0.242744 | 0.244310 | 0.249677 | 0.255610 | 0.266347 | 0.288186 | 0.325235 | 0.379745 | 0.461009 | 0.592437 | 0.749918 | 0.913912 | 1.045557 | 1.182166 | 1.292371 | 1.466853 | 1.736501 | 2.240773 | 2.884811 | 3.564876 | 3.899073 | 3.816621 | 3.193854 | 2.326575 | 1.525086 | 0.995169 | 0.662381 | 0.468667 | 0.360946 | 0.305260 | 0.269176 | 0.247021 | 0.234990 | 0.232962 | 0.238783 | 0.255903 | 0.294736 | 0.362172 | 0.439162 | 0.497302 | 0.535031 | 0.523977 | 0.487791 | 0.447027 | 0.399187 | 0.355088 | 0.322210 | 0.293114 | 0.262990 | 0.244444 | 0.228937 | 0.219114 | 0.215071 | 0.214649 | 0.214629 | 0.215498 | 0.218795 | 0.225878 | 0.233780 | 0.242418 | 0.244326 | 0.244326 | 0.240296 | 0.238342 | 0.232990 | 0.228198 | 0.222464 | 0.217214 | 0.214662 | 0.219129 | 0.232778 | 0.248844 | 0.260513 | 0.259450 | 0.242996 | 0.224000 | 0.214829 | 0.219555 | 0.236004 | 0.255701 | 0.274742 | 0.284495 | 0.288321 | 0.286566 | 0.279240 | 0.267856 | 0.258278 | 0.248898 | 0.234765 | 0.221204 | 0.214682 | 0.232506 | 0.317639 | 0.585926 | 1.243453 | 2.337219 | 3.361103 | 3.719931 | 3.334322 | 3.028534 | 2.655562 | 2.540364 | 2.497758 | 2.342839 | 1.844683 | 1.394920 | 0.941154 | 0.678142 | 0.550108 | 0.503833 | 0.485405 | 0.512453 | 0.570904 | 0.680932 | 0.856274 | 1.310193 | 2.381054 | 5.381403 | 14.671037 | 38.965038 | 83.401123 | 143.867468 | 197.833223 | 226.350712 | 247.322217 | 170.310542 | 71.453893 | 20.025284 | 6.343827 | 2.331603 | 1.174995 | 0.765046 | 0.630726 | 0.575233 | 0.599266 | 0.646329 | 0.689365 | 0.762611 | 0.874499 | 1.026954 | 1.226503 | 1.606216 | 2.000498 | 2.383732 | 2.357251 | 2.006229 | 1.321756 | 0.768113 | 0.430139 | 0.285161 | 0.225111 | 0.214834 | 0.224053 | 0.236210 | 0.248267 | 0.255852 | 0.265388 | 0.270849 | 0.275790 | 0.283979 | 0.296076 | 0.310380 | 0.333887 | 0.353778 | 0.359869 | 0.350664 | 0.340202 | 0.324449 | 0.292527 | 0.281019 | 0.266834 |
| all electrodes | 0.248314 | 0.291553 | 0.323980 | 0.421056 | 0.648427 | 1.141517 | 1.740083 | 2.024881 | 1.477650 | 0.567232 | 0.244405 | 0.220700 | 0.254175 | 0.285187 | 0.284940 | 0.301100 | 0.302998 | 0.313607 | 0.326499 | 0.385291 | 0.328657 | 0.266552 | 0.222510 | 0.215483 | 0.245202 | 0.286447 | 0.298344 | 0.268514 | 0.250455 | 0.226953 | 0.216753 | 0.215786 | 0.216251 | 0.216125 | 0.218263 | 0.219587 | 0.223894 | 0.225578 | 0.228323 | 0.228422 | 0.224955 | 0.214744 | 0.216834 | 0.245541 | 0.316725 | 0.396957 | 0.362623 | 0.342172 | 0.311988 | 0.309509 | 0.293053 | 0.296296 | 0.304819 | 0.270875 | 0.215442 | 0.363587 | 2.480632 | 20.878789 | 76.175426 | 172.001255 | 226.800923 | 270.660161 | 249.737467 | 181.979228 | 79.940255 | 20.614419 | 3.235465 | 0.623057 | 0.262580 | 0.216910 | 0.214660 | 0.218289 | 0.246414 | 0.350187 | 0.632184 | 1.280050 | 2.413182 | 4.757425 | 8.543091 | 13.769826 | 19.123142 | 25.263301 | 27.871634 | 30.237483 | 24.762893 | 21.876307 | 18.262290 | 16.532163 | 14.230690 | 12.724018 | 11.284055 | 8.436748 | 5.866602 | 4.826322 | 5.003461 | 6.100530 | 7.080055 | 6.318404 | 4.248085 | 2.557841 | 1.408446 | 0.908906 | 0.648070 | 0.539382 | 0.517161 | 0.632878 | 0.919674 | 1.765933 | 3.133404 | 4.590102 | 4.427939 | 3.332619 | 1.943058 | 1.102580 | 0.640620 | 0.427693 | 0.326033 | 0.283151 | 0.271858 | 0.279058 | 0.294036 | 0.316845 | 0.353955 | 0.402620 | 0.476859 | 0.546321 | 0.550605 | 0.493008 | 0.404014 | 0.331940 | 0.292559 | 0.263839 | 0.242935 | 0.232303 | 0.221474 | 0.214736 | 0.221856 | 0.249135 | 0.297270 | 0.378899 | 0.493288 | 0.526960 | 0.511331 | 0.458387 | 0.410312 | 0.368368 | 0.310488 | 0.270552 | 0.254497 | 0.241234 | 0.229703 | 0.225225 | 0.222443 | 0.221428 | 0.221498 | 0.224531 | 0.232302 | 0.245987 | 0.260793 | 0.287509 | 0.319809 | 0.353609 | 0.377647 | 0.386361 | 0.379151 | 0.376219 | 0.354824 | 0.339988 | 0.318053 | 0.284128 | 0.254841 | 0.237521 | 0.228352 | 0.227076 | 0.230760 | 0.234985 | 0.247374 | 0.263018 | 0.282047 | 0.283903 | 0.274072 | 0.260157 | 0.251566 | 0.243878 | 0.246365 | 0.248408 | 0.240663 | 0.226516 | 0.215225 | 0.218488 | 0.234152 | 0.262210 | 0.290332 | 0.273646 | 0.247154 | 0.224255 | 0.214634 | 0.225830 | 0.241328 | 0.244807 | 0.231221 | 0.218441 | 0.214739 | 0.220545 | 0.227550 | 0.225188 | 0.217298 | 0.214635 | 0.218065 | 0.224457 | 0.229921 | 0.228847 | 0.226039 | 0.222063 | 0.219588 | 0.220710 | 0.226207 | 0.234190 | 0.243826 | 0.254282 | 0.253687 | 0.251794 | 0.240641 | 0.228703 | 0.220698 | 0.218083 | 0.215784 | 0.217405 | 0.222789 | 0.233760 | 0.246757 | 0.261768 | 0.267417 | 0.270555 | 0.255588 | 0.232645 | 0.216679 | 0.216254 | 0.228580 | 0.248973 | 0.270627 | 0.279183 | 0.276226 | 0.266841 | 0.245288 | 0.222261 | 0.214693 | 0.225073 | 0.239652 | 0.246372 | 0.241864 | 0.229759 | 0.216582 | 0.217502 | 0.239120 | 0.274887 | 0.324282 | 0.366481 | 0.404750 | 0.453079 | 0.563638 | 0.730563 | 1.086643 | 1.726320 | 2.660293 | 3.286640 | 3.467504 | 2.777300 | 2.008959 | 1.419330 | 1.081337 | 0.954720 | 0.926866 | 0.872494 | 0.846887 | 0.820691 | 0.731167 | 0.605454 | 0.572801 | 0.513761 |

C) emotion

  
|  | time window | peak latency | cluster *p* | peak Cohen's *d* |  | | | |  |
| **all electrodes** | 125 - 225 ms | 160 ms | 0.0197 | 1.4671 |  | | | |  |
 340 - 560 ms | 395 ms | 0.0046 | 0.9563 |  | | | ||  |  | | | | | | | | |

Model correlations, cluster permutation tests

|  | **left hemisphere** | | | | **right hemisphere** | | | |  |
|  | time window | peak latency | cluster *p* | peak Cohen's *d* | time window | peak latency | cluster *p* | peak Cohen's *d* |  |
| **anterior** |  | | | |  | | | |  |
| **central** |  | | | | 210 - 495 ms | 275 ms | 0.0002 | 1.832 |  |
| **posterior** | 120 - 440 ms | 160 ms | 0.0025 | 1.4723 | 255 - 455 ms | 405 ms | 0.0233 | 0.9189 |  |
  | | | | 470 - 700 ms | 605 ms | 0.0242 | 0.6597 |  | | | | 725 - 1125 ms | 830 ms | 0.0047 | 0.6765 |

  

Model correlations, Bayesian statistics

|  | -200 | -195 | -190 | -185 | -180 | -175 | -170 | -165 | -160 | -155 | -150 | -145 | -140 | -135 | -130 | -125 | -120 | -115 | -110 | -105 | -100 | -95 | -90 | -85 | -80 | -75 | -70 | -65 | -60 | -55 | -50 | -45 | -40 | -35 | -30 | -25 | -20 | -15 | -10 | -5 | 0 | 5 | 10 | 15 | 20 | 25 | 30 | 35 | 40 | 45 | 50 | 55 | 60 | 65 | 70 | 75 | 80 | 85 | 90 | 95 | 100 | 105 | 110 | 115 | 120 | 125 | 130 | 135 | 140 | 145 | 150 | 155 | 160 | 165 | 170 | 175 | 180 | 185 | 190 | 195 | 200 | 205 | 210 | 215 | 220 | 225 | 230 | 235 | 240 | 245 | 250 | 255 | 260 | 265 | 270 | 275 | 280 | 285 | 290 | 295 | 300 | 305 | 310 | 315 | 320 | 325 | 330 | 335 | 340 | 345 | 350 | 355 | 360 | 365 | 370 | 375 | 380 | 385 | 390 | 395 | 400 | 405 | 410 | 415 | 420 | 425 | 430 | 435 | 440 | 445 | 450 | 455 | 460 | 465 | 470 | 475 | 480 | 485 | 490 | 495 | 500 | 505 | 510 | 515 | 520 | 525 | 530 | 535 | 540 | 545 | 550 | 555 | 560 | 565 | 570 | 575 | 580 | 585 | 590 | 595 | 600 | 605 | 610 | 615 | 620 | 625 | 630 | 635 | 640 | 645 | 650 | 655 | 660 | 665 | 670 | 675 | 680 | 685 | 690 | 695 | 700 | 705 | 710 | 715 | 720 | 725 | 730 | 735 | 740 | 745 | 750 | 755 | 760 | 765 | 770 | 775 | 780 | 785 | 790 | 795 | 800 | 805 | 810 | 815 | 820 | 825 | 830 | 835 | 840 | 845 | 850 | 855 | 860 | 865 | 870 | 875 | 880 | 885 | 890 | 895 | 900 | 905 | 910 | 915 | 920 | 925 | 930 | 935 | 940 | 945 | 950 | 955 | 960 | 965 | 970 | 975 | 980 | 985 | 990 | 995 | 1000 | 1005 | 1010 | 1015 | 1020 | 1025 | 1030 | 1035 | 1040 | 1045 | 1050 | 1055 | 1060 | 1065 | 1070 | 1075 | 1080 | 1085 | 1090 | 1095 | 1100 | 1105 | 1110 | 1115 | 1120 | 1125 | 1130 | 1135 | 1140 | 1145 | 1150 | 1155 | 1160 | 1165 | 1170 | 1175 | 1180 | 1185 | 1190 | 1195 |
| --- | --- | --- | --- | --- | --- | --- | --- | --- | --- | --- | --- | --- | --- | --- | --- | --- | --- | --- | --- | --- | --- | --- | --- | --- | --- | --- | --- | --- | --- | --- | --- | --- | --- | --- | --- | --- | --- | --- | --- | --- | --- | --- | --- | --- | --- | --- | --- | --- | --- | --- | --- | --- | --- | --- | --- | --- | --- | --- | --- | --- | --- | --- | --- | --- | --- | --- | --- | --- | --- | --- | --- | --- | --- | --- | --- | --- | --- | --- | --- | --- | --- | --- | --- | --- | --- | --- | --- | --- | --- | --- | --- | --- | --- | --- | --- | --- | --- | --- | --- | --- | --- | --- | --- | --- | --- | --- | --- | --- | --- | --- | --- | --- | --- | --- | --- | --- | --- | --- | --- | --- | --- | --- | --- | --- | --- | --- | --- | --- | --- | --- | --- | --- | --- | --- | --- | --- | --- | --- | --- | --- | --- | --- | --- | --- | --- | --- | --- | --- | --- | --- | --- | --- | --- | --- | --- | --- | --- | --- | --- | --- | --- | --- | --- | --- | --- | --- | --- | --- | --- | --- | --- | --- | --- | --- | --- | --- | --- | --- | --- | --- | --- | --- | --- | --- | --- | --- | --- | --- | --- | --- | --- | --- | --- | --- | --- | --- | --- | --- | --- | --- | --- | --- | --- | --- | --- | --- | --- | --- | --- | --- | --- | --- | --- | --- | --- | --- | --- | --- | --- | --- | --- | --- | --- | --- | --- | --- | --- | --- | --- | --- | --- | --- | --- | --- | --- | --- | --- | --- | --- | --- | --- | --- | --- | --- | --- | --- | --- | --- | --- | --- | --- | --- | --- | --- | --- | --- | --- | --- | --- | --- | --- | --- | --- | --- | --- | --- | --- | --- | --- | --- | --- | --- | --- | --- | --- | --- | --- | --- | --- | --- |
| left anterior | 0.228279 | 0.241114 | 0.242453 | 0.252804 | 0.255122 | 0.253886 | 0.245924 | 0.244347 | 0.299178 | 0.389186 | 0.430318 | 0.490204 | 0.547255 | 0.460040 | 0.376453 | 0.298889 | 0.265891 | 0.264194 | 0.263568 | 0.267469 | 0.269299 | 0.268245 | 0.246406 | 0.215136 | 0.261470 | 0.408004 | 0.892191 | 2.000508 | 3.391354 | 3.535802 | 1.746643 | 0.460949 | 0.216787 | 0.446068 | 2.058720 | 5.168993 | 6.382655 | 5.087490 | 3.434367 | 2.498832 | 2.167214 | 2.376169 | 5.159039 | 11.598611 | 33.322414 | 30.316695 | 9.207604 | 1.889380 | 0.746607 | 0.391900 | 0.310943 | 0.294673 | 0.386499 | 0.571895 | 0.889141 | 1.035755 | 1.020062 | 0.714204 | 0.566373 | 0.470828 | 0.363610 | 0.289033 | 0.248855 | 0.217015 | 0.228964 | 0.297711 | 0.453582 | 0.707165 | 1.164901 | 1.394986 | 1.516702 | 1.339706 | 1.274106 | 1.040486 | 0.778873 | 0.458803 | 0.290958 | 0.217996 | 0.243954 | 0.419815 | 0.801630 | 1.013298 | 0.861829 | 0.659444 | 0.447585 | 0.307129 | 0.247619 | 0.221133 | 0.214643 | 0.220519 | 0.232404 | 0.248243 | 0.260311 | 0.283083 | 0.291688 | 2.675294e-01 | 0.235050 | 0.217234 | 0.221178 | 0.264515 | 0.352642 | 0.459941 | 0.526877 | 0.514578 | 0.451391 | 0.362739 | 0.279689 | 0.235795 | 0.218216 | 0.214934 | 0.214703 | 0.214727 | 0.214905 | 0.215170 | 0.216676 | 0.222054 | 0.242815 | 0.283769 | 0.350936 | 0.433391 | 0.470694 | 0.458361 | 0.399267 | 0.318852 | 0.263137 | 0.234648 | 0.219394 | 0.214805 | 0.214720 | 0.214875 | 0.214653 | 0.215099 | 0.215575 | 0.216801 | 0.219148 | 0.217856 | 0.215831 | 0.214821 | 0.214706 | 0.215397 | 0.216680 | 0.217718 | 0.218488 | 0.217738 | 0.217041 | 0.214856 | 0.215852 | 0.217797 | 0.216456 | 0.214844 | 0.214804 | 0.217476 | 0.222566 | 0.229579 | 0.230769 | 0.229455 | 0.223331 | 0.216686 | 0.214640 | 0.217092 | 0.221305 | 0.221487 | 0.218538 | 0.216265 | 0.214923 | 0.214692 | 0.214913 | 0.216889 | 0.225271 | 0.236172 | 0.240277 | 0.229462 | 0.219018 | 0.214811 | 0.221793 | 0.233562 | 0.230344 | 0.221019 | 0.214627 | 0.220492 | 0.240645 | 0.258414 | 0.264702 | 0.257296 | 0.251959 | 0.241296 | 0.229965 | 0.220712 | 0.217488 | 0.215095 | 0.216016 | 0.226316 | 0.240534 | 0.250272 | 0.242017 | 0.219032 | 0.227848 | 0.339724 | 0.609350 | 0.989262 | 1.417969 | 1.502128 | 1.054538 | 0.548122 | 0.327131 | 0.245659 | 0.221976 | 0.218365 | 0.225234 | 0.248570 | 0.295154 | 0.352421 | 0.431046 | 0.556189 | 0.720158 | 0.885147 | 1.052759 | 1.104765 | 1.062200 | 0.981444 | 0.830329 | 0.667587 | 0.557099 | 0.481179 | 0.412207 | 0.355956 | 0.325908 | 0.306038 | 0.297394 | 0.284242 | 0.271394 | 0.259676 | 0.249807 | 0.234797 | 0.231432 | 0.231411 | 0.229699 | 0.227128 | 0.224158 | 0.219213 | 0.216621 | 0.215102 | 0.215298 | 0.218797 | 0.219681 | 0.218295 | 0.215504 | 0.215188 | 0.220699 | 0.229194 | 0.240785 | 0.249260 | 0.257343 | 0.269313 | 0.274050 | 0.264357 | 0.257861 | 0.250820 | 0.241137 | 0.235800 | 0.236615 | 0.243627 | 0.263467 | 0.298124 | 0.339488 | 0.390524 | 0.439835 | 0.454787 | 0.440257 | 0.422900 | 0.393305 | 0.357252 | 0.332306 | 0.315010 | 0.309303 | 0.313508 | 0.337852 | 0.386921 | 0.415715 | 0.458604 |
| right anterior | 1.242715 | 1.069604 | 0.707512 | 0.436186 | 0.302630 | 0.235074 | 0.219982 | 0.215581 | 0.214627 | 0.217638 | 0.250377 | 0.426087 | 0.746599 | 1.497872 | 1.938160 | 1.581413 | 1.040053 | 0.543721 | 0.269308 | 0.214904 | 0.236544 | 0.320887 | 0.472618 | 0.683198 | 0.988533 | 1.459780 | 1.739355 | 1.158698 | 0.525792 | 0.287094 | 0.215598 | 0.272026 | 0.425596 | 0.534445 | 0.501096 | 0.407789 | 0.307201 | 0.263470 | 0.228067 | 0.222924 | 0.231227 | 0.252528 | 0.280442 | 0.339260 | 0.356832 | 0.354678 | 0.306179 | 0.249365 | 0.218311 | 0.215898 | 0.227850 | 0.239594 | 0.252149 | 0.258456 | 0.270778 | 0.269790 | 0.267513 | 0.246392 | 0.231186 | 0.218533 | 0.214803 | 0.216015 | 0.214648 | 0.231569 | 0.340958 | 0.712817 | 1.718562 | 2.566486 | 2.603859 | 2.070544 | 1.670760 | 1.314330 | 1.037862 | 0.780562 | 0.619459 | 0.545072 | 0.490338 | 0.431064 | 0.380392 | 0.314662 | 0.256180 | 0.233942 | 0.231687 | 0.238772 | 0.259865 | 0.300886 | 0.361485 | 0.429680 | 0.556258 | 0.819324 | 1.390102 | 2.523473 | 5.142756 | 10.378725 | 18.639949 | 2.210882e+01 | 17.818987 | 12.082235 | 7.991189 | 4.418819 | 2.763873 | 1.851000 | 1.243913 | 0.916587 | 0.750372 | 0.651431 | 0.621448 | 0.644940 | 0.697470 | 0.792788 | 0.944585 | 1.165488 | 1.308486 | 1.498379 | 1.781787 | 1.948099 | 2.160672 | 2.405218 | 2.376339 | 2.477473 | 2.651131 | 2.536783 | 2.257288 | 1.766269 | 1.291866 | 0.983707 | 0.755607 | 0.587408 | 0.491552 | 0.439957 | 0.380236 | 0.329256 | 0.298424 | 0.291796 | 0.285856 | 0.277352 | 0.264792 | 0.248079 | 0.227111 | 0.215099 | 0.218223 | 0.232903 | 0.246971 | 0.241440 | 0.224856 | 0.214819 | 0.220810 | 0.239888 | 0.261917 | 0.298176 | 0.321007 | 0.334063 | 0.330505 | 0.308621 | 0.270692 | 0.243260 | 0.221526 | 0.214627 | 0.223524 | 0.252910 | 0.298982 | 0.337676 | 0.390708 | 0.456026 | 0.518942 | 0.588697 | 0.687496 | 0.773376 | 0.842042 | 0.862358 | 0.865347 | 0.779144 | 0.610486 | 0.474173 | 0.398066 | 0.351161 | 0.316303 | 0.305152 | 0.310533 | 0.324549 | 0.347094 | 0.421667 | 0.594543 | 0.858462 | 1.151582 | 1.263490 | 1.103673 | 0.831542 | 0.620613 | 0.471406 | 0.424901 | 0.395606 | 0.369405 | 0.352207 | 0.337276 | 0.305274 | 0.273234 | 0.249070 | 0.239737 | 0.246713 | 0.270295 | 0.314631 | 0.388403 | 0.455192 | 0.461665 | 0.403722 | 0.338129 | 0.290049 | 0.257442 | 0.236390 | 0.228625 | 0.227712 | 0.227411 | 0.226019 | 0.222602 | 0.221235 | 0.222139 | 0.220502 | 0.219141 | 0.219416 | 0.217380 | 0.214926 | 0.214796 | 0.217133 | 0.217369 | 0.215006 | 0.215086 | 0.217364 | 0.218897 | 0.214766 | 0.221557 | 0.258206 | 0.335610 | 0.444536 | 0.499579 | 0.479144 | 0.368111 | 0.273851 | 0.227479 | 0.217384 | 0.217188 | 0.225714 | 0.253569 | 0.298907 | 0.345626 | 0.369435 | 0.373231 | 0.334952 | 0.299235 | 0.263424 | 0.241675 | 0.228718 | 0.225606 | 0.223148 | 0.230474 | 0.250841 | 0.288914 | 0.337783 | 0.407583 | 0.461657 | 0.508367 | 0.544547 | 0.575928 | 0.586232 | 0.556656 | 0.488802 | 0.384891 | 0.283459 | 0.225813 | 0.216148 | 0.243882 | 0.284275 | 0.320741 | 0.345059 | 0.352860 | 0.354232 | 0.340601 | 0.326587 | 0.317957 | 0.299728 |
| left central | 0.284199 | 0.293956 | 0.306663 | 0.338711 | 0.362116 | 0.366250 | 0.328387 | 0.284739 | 0.259775 | 0.239819 | 0.224278 | 0.216870 | 0.215998 | 0.242846 | 0.331767 | 0.419637 | 0.483115 | 0.409727 | 0.288894 | 0.248536 | 0.228600 | 0.214844 | 0.226536 | 0.258159 | 0.351139 | 0.459169 | 0.575908 | 0.521571 | 0.383194 | 0.259043 | 0.216714 | 0.225167 | 0.241381 | 0.232898 | 0.215885 | 0.219416 | 0.243762 | 0.277437 | 0.311293 | 0.333058 | 0.316855 | 0.272522 | 0.230160 | 0.214683 | 0.223268 | 0.238178 | 0.245545 | 0.234467 | 0.223805 | 0.217327 | 0.214814 | 0.214795 | 0.214972 | 0.216204 | 0.216826 | 0.220058 | 0.225765 | 0.231112 | 0.229035 | 0.227166 | 0.221751 | 0.218766 | 0.216770 | 0.217261 | 0.224902 | 0.283376 | 0.541661 | 1.855302 | 8.240368 | 29.208042 | 65.974834 | 93.017245 | 81.533696 | 54.721592 | 32.099455 | 14.974627 | 7.251159 | 3.911038 | 2.310800 | 1.585886 | 1.250481 | 0.949993 | 0.811053 | 0.779741 | 0.752666 | 0.746179 | 0.744100 | 0.682816 | 0.693941 | 0.711789 | 0.934115 | 1.497041 | 2.590838 | 3.792759 | 5.422651 | 5.420495e+00 | 3.839003 | 2.014713 | 0.892878 | 0.440316 | 0.291547 | 0.240093 | 0.223181 | 0.221652 | 0.223690 | 0.230917 | 0.248135 | 0.278118 | 0.291345 | 0.284628 | 0.263911 | 0.246416 | 0.235842 | 0.234148 | 0.249076 | 0.289752 | 0.332019 | 0.382205 | 0.464791 | 0.565753 | 0.634819 | 0.663908 | 0.637368 | 0.615078 | 0.607461 | 0.642529 | 0.683163 | 0.871759 | 1.156881 | 1.429950 | 1.665148 | 1.904083 | 1.458932 | 0.908166 | 0.564588 | 0.387839 | 0.321063 | 0.288768 | 0.259543 | 0.237711 | 0.217654 | 0.223248 | 0.295771 | 0.514883 | 1.066524 | 1.795739 | 1.901433 | 1.469912 | 0.968353 | 0.617154 | 0.425574 | 0.335037 | 0.292497 | 0.278408 | 0.276827 | 0.283934 | 0.294262 | 0.290366 | 0.277233 | 0.259216 | 0.242041 | 0.233983 | 0.235699 | 0.247559 | 0.279257 | 0.321407 | 0.352561 | 0.382794 | 0.386414 | 0.373813 | 0.365743 | 0.380297 | 0.398278 | 0.439559 | 0.499858 | 0.610914 | 0.714038 | 0.797261 | 0.662782 | 0.525591 | 0.421175 | 0.348560 | 0.287327 | 0.263151 | 0.249387 | 0.254358 | 0.264524 | 0.289647 | 0.320816 | 0.368250 | 0.389785 | 0.423306 | 0.491692 | 0.579052 | 0.600827 | 0.547683 | 0.438311 | 0.310444 | 0.255911 | 0.232339 | 0.223398 | 0.220934 | 0.224336 | 0.235708 | 0.263320 | 0.306369 | 0.343947 | 0.350181 | 0.321819 | 0.285080 | 0.254628 | 0.239948 | 0.233999 | 0.229926 | 0.229078 | 0.230413 | 0.228457 | 0.227636 | 0.227900 | 0.223954 | 0.217774 | 0.214707 | 0.216802 | 0.223849 | 0.237303 | 0.260530 | 0.291124 | 0.337441 | 0.433882 | 0.630376 | 1.007182 | 1.411093 | 1.566162 | 1.424943 | 1.009291 | 0.607370 | 0.370759 | 0.268948 | 0.235177 | 0.224053 | 0.220883 | 0.221814 | 0.223207 | 0.224350 | 0.219066 | 0.214756 | 0.217051 | 0.228489 | 0.258402 | 0.310426 | 0.393235 | 0.482115 | 0.587607 | 0.712740 | 0.738511 | 0.640561 | 0.513990 | 0.397558 | 0.337124 | 0.303933 | 0.294137 | 0.307296 | 0.342753 | 0.393848 | 0.476893 | 0.596923 | 0.736691 | 0.893176 | 0.992632 | 0.960248 | 0.856576 | 0.788654 | 0.730674 | 0.772838 | 0.804310 | 0.846088 | 0.905625 | 1.018334 | 0.963623 | 0.989190 |
| right central | 3.773983 | 4.256309 | 5.026474 | 6.093438 | 5.709908 | 3.848157 | 2.670290 | 1.872124 | 1.510793 | 1.274570 | 0.936939 | 0.556752 | 0.325465 | 0.237044 | 0.215636 | 0.214645 | 0.216646 | 0.225165 | 0.247723 | 0.286340 | 0.344585 | 0.401076 | 0.436562 | 0.406263 | 0.328219 | 0.257227 | 0.217729 | 0.218804 | 0.232867 | 0.229366 | 0.216352 | 0.227225 | 0.287044 | 0.392752 | 0.484086 | 0.432109 | 0.290798 | 0.220524 | 0.228129 | 0.262857 | 0.268131 | 0.242650 | 0.219398 | 0.222907 | 0.283787 | 0.465738 | 0.723164 | 0.903592 | 0.706038 | 0.498740 | 0.331093 | 0.247443 | 0.215870 | 0.226599 | 0.315044 | 0.596765 | 1.398905 | 2.640973 | 2.521250 | 1.121135 | 0.493618 | 0.293434 | 0.233438 | 0.215081 | 0.219140 | 0.248001 | 0.322808 | 0.593180 | 1.817475 | 9.499464 | 45.544443 | 161.866613 | 254.155918 | 145.205407 | 52.294068 | 14.798783 | 4.576437 | 1.783001 | 0.841454 | 0.509490 | 0.481795 | 0.649456 | 1.657673 | 6.630729 | 26.821965 | 74.608252 | 101.498762 | 73.467462 | 58.942670 | 62.912627 | 113.422224 | 364.902151 | 2284.888447 | 27664.384212 | 478927.039729 | 1.434484e+06 | 730004.661826 | 97300.540311 | 11975.636525 | 2593.073594 | 1172.503081 | 1055.688483 | 932.362450 | 695.271538 | 358.278212 | 155.843412 | 57.598698 | 30.212998 | 20.048583 | 17.812964 | 14.449228 | 14.075816 | 16.508004 | 24.766377 | 29.297755 | 32.290074 | 28.332887 | 26.202004 | 23.723479 | 25.805113 | 33.868916 | 65.372440 | 114.744897 | 211.507474 | 402.609551 | 557.365196 | 454.269892 | 308.083919 | 205.333478 | 176.313209 | 193.079829 | 266.190864 | 341.643395 | 335.441809 | 193.164781 | 94.056055 | 37.209726 | 12.554993 | 3.842909 | 1.622327 | 0.893365 | 0.639545 | 0.544817 | 0.511523 | 0.472830 | 0.432954 | 0.411431 | 0.430011 | 0.497923 | 0.583223 | 0.651530 | 0.694822 | 0.657495 | 0.630034 | 0.609422 | 0.680560 | 0.954018 | 1.613788 | 3.385936 | 8.635722 | 18.989022 | 23.106034 | 14.551576 | 5.400881 | 2.355507 | 1.269462 | 0.997111 | 0.961168 | 1.108043 | 1.107961 | 1.066814 | 0.870781 | 0.748016 | 0.626161 | 0.547857 | 0.468582 | 0.461297 | 0.469283 | 0.469986 | 0.454427 | 0.430642 | 0.395376 | 0.377325 | 0.367004 | 0.369287 | 0.391678 | 0.410644 | 0.431337 | 0.468318 | 0.498660 | 0.532164 | 0.587963 | 0.645076 | 0.673419 | 0.672346 | 0.595395 | 0.474355 | 0.390496 | 0.356497 | 0.349794 | 0.372172 | 0.415441 | 0.470810 | 0.521312 | 0.559071 | 0.490761 | 0.390633 | 0.296541 | 0.238620 | 0.217485 | 0.214892 | 0.216051 | 0.232008 | 0.278412 | 0.332421 | 0.417332 | 0.467439 | 0.431299 | 0.356046 | 0.302004 | 0.258921 | 0.245967 | 0.246012 | 0.270131 | 0.327343 | 0.424492 | 0.502641 | 0.562227 | 0.576650 | 0.528630 | 0.441363 | 0.353836 | 0.289839 | 0.250401 | 0.235740 | 0.230135 | 0.227440 | 0.223136 | 0.217555 | 0.215496 | 0.231908 | 0.278624 | 0.373950 | 0.517503 | 0.654564 | 0.735371 | 0.661415 | 0.498997 | 0.370259 | 0.278066 | 0.244472 | 0.232566 | 0.222907 | 0.218393 | 0.216132 | 0.214633 | 0.216030 | 0.226577 | 0.253720 | 0.282272 | 0.318903 | 0.363451 | 0.345409 | 0.320453 | 0.295676 | 0.271880 | 0.251780 | 0.233125 | 0.220208 | 0.214979 | 0.216233 | 0.228206 | 0.257390 | 0.311055 | 0.380766 | 0.486122 | 0.563713 | 0.562750 | 0.540829 | 0.510151 |
| left posterior | 1.273682 | 1.822367 | 2.674107 | 6.210895 | 11.803936 | 12.492592 | 5.420023 | 1.811058 | 0.544854 | 0.255974 | 0.214741 | 0.246694 | 0.302786 | 0.303823 | 0.255561 | 0.214708 | 0.377757 | 2.844146 | 42.166362 | 121.475703 | 31.087612 | 3.755847 | 0.492931 | 0.214801 | 0.490021 | 1.859605 | 3.050814 | 1.282904 | 0.512980 | 0.243873 | 0.219406 | 0.252312 | 0.231069 | 0.215020 | 0.230940 | 0.276143 | 0.336448 | 0.378384 | 0.394295 | 0.365502 | 0.314326 | 0.275181 | 0.268821 | 0.293109 | 0.337357 | 0.378592 | 0.380806 | 0.368049 | 0.337921 | 0.297230 | 0.258972 | 0.229294 | 0.214859 | 0.223914 | 0.256503 | 0.302937 | 0.346066 | 0.385609 | 0.431593 | 0.410338 | 0.302954 | 0.227454 | 0.226511 | 0.409313 | 1.550593 | 9.717910 | 69.316260 | 371.974200 | 1537.118888 | 5615.157911 | 16117.771992 | 37860.923790 | 65982.837654 | 86410.441953 | 74791.560214 | 50150.895511 | 21136.722559 | 9729.725453 | 5238.356619 | 2189.705457 | 511.153836 | 74.625053 | 14.086831 | 4.626089 | 2.370937 | 1.648476 | 1.388779 | 1.151967 | 1.042873 | 1.206620 | 1.623697 | 2.388929 | 3.999377 | 6.434169 | 9.925437 | 1.475710e+01 | 16.981149 | 13.607278 | 9.121393 | 5.803062 | 4.084019 | 3.221370 | 3.032862 | 3.418136 | 4.416482 | 6.455030 | 10.211242 | 16.019435 | 21.717470 | 22.888920 | 17.588513 | 11.699366 | 7.090543 | 4.807037 | 3.637065 | 3.287275 | 3.695810 | 5.202441 | 8.628071 | 16.144055 | 31.716067 | 56.667716 | 80.741701 | 88.583798 | 74.546313 | 39.869944 | 17.068744 | 6.028772 | 2.221228 | 0.998881 | 0.631196 | 0.485330 | 0.455911 | 0.464884 | 0.503292 | 0.566106 | 0.676061 | 0.809634 | 0.920922 | 1.019947 | 1.073509 | 1.039300 | 1.055215 | 1.107015 | 1.139115 | 1.043687 | 0.826806 | 0.596486 | 0.448449 | 0.348008 | 0.291844 | 0.259415 | 0.240223 | 0.231188 | 0.227200 | 0.228075 | 0.235295 | 0.250682 | 0.269412 | 0.294727 | 0.317731 | 0.343343 | 0.358360 | 0.358197 | 0.345258 | 0.328285 | 0.307794 | 0.291275 | 0.277837 | 0.262972 | 0.250629 | 0.244075 | 0.241359 | 0.244943 | 0.259237 | 0.281945 | 0.312761 | 0.358985 | 0.410449 | 0.461620 | 0.514002 | 0.554237 | 0.566030 | 0.587227 | 0.591924 | 0.564305 | 0.514698 | 0.461675 | 0.400359 | 0.351555 | 0.324811 | 0.305702 | 0.297060 | 0.300616 | 0.326227 | 0.373762 | 0.477565 | 0.646845 | 0.915818 | 1.211506 | 1.507125 | 1.595660 | 1.573787 | 1.413258 | 1.232477 | 1.073841 | 0.956810 | 0.884580 | 0.874147 | 0.970746 | 1.181401 | 1.626316 | 2.395110 | 3.882624 | 5.869225 | 8.767978 | 11.923471 | 13.924426 | 12.100602 | 9.372928 | 5.812413 | 3.489395 | 2.091179 | 1.358988 | 0.991167 | 0.857873 | 0.751901 | 0.702458 | 0.674032 | 0.663123 | 0.657806 | 0.744921 | 0.965131 | 1.520536 | 2.860996 | 6.228702 | 11.746938 | 20.162288 | 25.338757 | 23.127167 | 17.060298 | 10.556851 | 5.067068 | 2.811080 | 1.899371 | 1.649227 | 1.916431 | 2.577597 | 3.131673 | 3.316436 | 2.920103 | 2.257439 | 1.702084 | 1.292893 | 0.994981 | 0.838212 | 0.686037 | 0.588748 | 0.522055 | 0.453839 | 0.376772 | 0.317542 | 0.273250 | 0.247401 | 0.232511 | 0.224588 | 0.220974 | 0.219521 | 0.218485 | 0.217273 | 0.215253 | 0.214632 | 0.216976 | 0.222313 | 0.227695 | 0.227999 | 0.226967 | 0.223365 | 0.223098 | 0.220447 |
| right posterior | 1.431169 | 1.657403 | 1.991867 | 2.342420 | 2.385374 | 2.091420 | 1.577576 | 1.002552 | 0.539448 | 0.326844 | 0.254570 | 0.226912 | 0.217730 | 0.217850 | 0.234072 | 0.258600 | 0.300223 | 0.316703 | 0.289601 | 0.240673 | 0.216495 | 0.230587 | 0.263946 | 0.263752 | 0.238103 | 0.214651 | 0.253767 | 0.396429 | 0.758792 | 1.519199 | 2.598032 | 2.052872 | 0.966786 | 0.410683 | 0.228091 | 0.227997 | 0.312470 | 0.379458 | 0.386062 | 0.324382 | 0.285397 | 0.254221 | 0.243877 | 0.239512 | 0.239968 | 0.235469 | 0.221224 | 0.219240 | 0.314990 | 0.602648 | 1.022456 | 1.132622 | 0.938368 | 0.653875 | 0.418344 | 0.311672 | 0.280813 | 0.275129 | 0.278085 | 0.281107 | 0.248748 | 0.215011 | 0.278141 | 0.775224 | 3.701865 | 20.863881 | 100.784566 | 332.693546 | 730.299010 | 1364.516027 | 1676.128868 | 1401.092388 | 772.496558 | 269.791111 | 78.689108 | 25.954267 | 8.750455 | 3.190488 | 1.371055 | 0.625352 | 0.345709 | 0.260239 | 0.239871 | 0.240975 | 0.255239 | 0.291321 | 0.334981 | 0.401416 | 0.513617 | 0.721340 | 1.056191 | 1.768610 | 2.873823 | 4.358731 | 6.116615 | 7.173999e+00 | 6.947501 | 5.985182 | 4.933619 | 3.920647 | 3.571363 | 3.384327 | 3.400181 | 3.680688 | 3.977833 | 3.888237 | 3.834789 | 3.766979 | 3.367970 | 2.864887 | 2.360178 | 1.921684 | 1.734617 | 1.896523 | 2.252963 | 3.235256 | 5.857851 | 12.116873 | 28.674853 | 70.977562 | 149.994384 | 239.775809 | 248.813081 | 170.058455 | 88.887005 | 38.889050 | 15.691131 | 7.010141 | 3.560286 | 2.142130 | 1.494100 | 1.241179 | 1.148329 | 1.125682 | 1.189249 | 1.383126 | 1.595994 | 1.861219 | 2.118029 | 2.445365 | 2.772996 | 2.954725 | 3.185378 | 3.837829 | 4.763072 | 5.376333 | 5.045141 | 4.251745 | 3.265898 | 2.381239 | 1.714243 | 1.364074 | 1.156844 | 1.126890 | 1.281118 | 1.799331 | 2.827469 | 4.893266 | 8.059230 | 11.664339 | 13.087051 | 13.040052 | 10.955030 | 8.300521 | 5.830495 | 4.121870 | 2.866709 | 2.250513 | 1.822092 | 1.630850 | 1.634129 | 1.834794 | 2.094422 | 2.718249 | 3.444567 | 4.359254 | 4.721316 | 4.366072 | 3.446712 | 2.459184 | 1.591718 | 1.125836 | 0.879574 | 0.792158 | 0.861817 | 1.121440 | 1.721701 | 2.859396 | 4.359133 | 5.001722 | 4.522673 | 3.386497 | 2.697873 | 2.464483 | 2.454725 | 2.695647 | 2.950124 | 3.028822 | 3.195219 | 3.636388 | 4.446112 | 5.813628 | 7.660675 | 10.048043 | 12.285989 | 13.375711 | 13.427615 | 12.679444 | 11.525170 | 10.151150 | 9.255252 | 9.120497 | 10.354306 | 13.820401 | 20.619441 | 35.233159 | 58.474022 | 74.968379 | 78.107144 | 68.684661 | 43.860827 | 25.047253 | 14.773862 | 9.174236 | 6.684341 | 5.850473 | 5.522359 | 5.608258 | 6.179943 | 6.626613 | 7.345048 | 8.729608 | 11.602297 | 15.565675 | 19.419426 | 23.013622 | 24.993360 | 25.293133 | 23.899475 | 22.308683 | 17.669865 | 13.452981 | 10.138298 | 8.239612 | 7.489755 | 8.059931 | 9.780776 | 14.553631 | 20.739015 | 25.234937 | 25.671304 | 21.136450 | 13.679692 | 8.000758 | 4.764151 | 3.253309 | 2.589060 | 2.315583 | 2.336629 | 2.673293 | 3.085386 | 3.296655 | 3.193729 | 2.602181 | 1.938230 | 1.312714 | 0.892203 | 0.656931 | 0.554765 | 0.475861 | 0.446676 | 0.445835 | 0.459405 | 0.502218 | 0.617677 | 0.765614 | 1.003134 | 1.320038 | 1.478114 | 1.613044 |
| all electrodes | 0.429001 | 0.520781 | 0.528270 | 0.470150 | 0.402684 | 0.358644 | 0.266566 | 0.227831 | 0.216054 | 0.283378 | 0.434926 | 0.477455 | 0.494756 | 0.392590 | 0.307818 | 0.252656 | 0.224134 | 0.214739 | 0.223723 | 0.257002 | 0.271221 | 0.235664 | 0.215173 | 0.251168 | 0.408463 | 0.662091 | 1.221244 | 1.173005 | 0.886246 | 0.563138 | 0.424717 | 0.285304 | 0.253134 | 0.258545 | 0.302021 | 0.352941 | 0.418692 | 0.382025 | 0.311799 | 0.236276 | 0.217400 | 0.293001 | 0.552395 | 1.254826 | 1.677678 | 1.222887 | 0.589764 | 0.321420 | 0.262345 | 0.247126 | 0.248417 | 0.267253 | 0.321178 | 0.389518 | 0.509045 | 0.593657 | 0.567243 | 0.464203 | 0.387636 | 0.327794 | 0.266953 | 0.221806 | 0.222843 | 0.311810 | 0.688050 | 2.189114 | 11.199383 | 78.887652 | 592.607773 | 3073.914409 | 12172.318240 | 34972.456876 | 75801.712663 | 82158.734554 | 50808.740296 | 13461.678548 | 1934.357303 | 279.294686 | 59.852139 | 23.692901 | 15.083949 | 13.316038 | 11.507469 | 7.261958 | 3.407250 | 1.863084 | 1.142484 | 0.758921 | 0.593942 | 0.519468 | 0.556001 | 0.679114 | 1.055196 | 2.150974 | 5.974028 | 1.297355e+01 | 20.994792 | 15.028873 | 7.608720 | 3.328924 | 1.634209 | 0.861674 | 0.583210 | 0.483388 | 0.458683 | 0.502787 | 0.584900 | 0.775413 | 1.208393 | 2.314468 | 5.144889 | 13.459832 | 37.954129 | 84.540301 | 174.861685 | 234.179322 | 190.401105 | 151.358394 | 152.337631 | 147.738194 | 199.457944 | 305.196365 | 461.185771 | 475.307204 | 346.158774 | 184.580733 | 115.650420 | 67.433420 | 41.040793 | 23.873738 | 15.204920 | 11.979568 | 10.806844 | 9.589316 | 9.931348 | 11.426370 | 12.291682 | 11.822589 | 11.788174 | 10.407936 | 8.159630 | 5.487884 | 3.379248 | 2.460594 | 2.127276 | 1.975792 | 1.890494 | 2.022221 | 2.065036 | 2.048689 | 1.932025 | 1.722458 | 1.446729 | 1.104478 | 0.781823 | 0.550801 | 0.421628 | 0.347378 | 0.309490 | 0.272573 | 0.244366 | 0.224648 | 0.215448 | 0.216279 | 0.225383 | 0.246288 | 0.276639 | 0.323741 | 0.380743 | 0.429553 | 0.445894 | 0.465540 | 0.447609 | 0.433900 | 0.406409 | 0.349954 | 0.283887 | 0.239353 | 0.217782 | 0.216596 | 0.231839 | 0.253329 | 0.253457 | 0.241655 | 0.226851 | 0.218697 | 0.214644 | 0.217747 | 0.226772 | 0.231329 | 0.236122 | 0.233558 | 0.224518 | 0.217958 | 0.215111 | 0.215178 | 0.217215 | 0.220043 | 0.226360 | 0.233202 | 0.238593 | 0.247884 | 0.257762 | 0.264229 | 0.262469 | 0.253762 | 0.251376 | 0.264496 | 0.289470 | 0.329908 | 0.384078 | 0.445096 | 0.453587 | 0.407321 | 0.339672 | 0.295109 | 0.269710 | 0.270476 | 0.290048 | 0.318915 | 0.332563 | 0.324249 | 0.305396 | 0.294258 | 0.294500 | 0.309462 | 0.362008 | 0.439328 | 0.524244 | 0.620800 | 0.686702 | 0.619469 | 0.574529 | 0.551451 | 0.579777 | 0.672734 | 0.809297 | 1.005300 | 1.321900 | 1.558800 | 1.486678 | 1.223115 | 0.924589 | 0.688293 | 0.531224 | 0.479877 | 0.477591 | 0.472139 | 0.480421 | 0.511745 | 0.465978 | 0.404063 | 0.333358 | 0.268369 | 0.227004 | 0.214893 | 0.241217 | 0.286838 | 0.318531 | 0.319186 | 0.277901 | 0.232003 | 0.215760 | 0.215943 | 0.219419 | 0.219051 | 0.214741 | 0.223246 | 0.266451 | 0.348162 | 0.438150 | 0.534910 | 0.601984 | 0.669836 | 0.677825 | 0.706529 | 0.696671 | 0.678247 | 0.590508 | 0.585751 |

D) neutral vs happy

  
|  | time window | peak latency | cluster *p* | peak Cohen's *d* |  | | | |
| **all electrodes** |  | | | |  | | | |
|  | | | | | | | | |

Model correlations, cluster permutation tests

|  | **left hemisphere** | | | | **right hemisphere** | | | |
|  | time window | peak latency | cluster *p* | peak Cohen's *d* | time window | peak latency | cluster *p* | peak Cohen's *d* |
| **anterior** |  | | | | 230 - 405 ms | 280 ms | 0.0075 | 0.9909 |
| **central** |  | | | |  | | | |
| **posterior** | 245 - 415 ms | 280 ms | 0.0131 | 0.8676 | 260 - 445 ms | 290 ms | 0.0135 | 0.7171 |
  | | | | 800 - 975 ms | 845 ms | 0.0251 | 0.6384 |  | | | | 985 - 1120 ms | 1050 ms | 0.0411 | 0.644 |

  

Model correlations, Bayesian statistics

|  | -200 | -195 | -190 | -185 | -180 | -175 | -170 | -165 | -160 | -155 | -150 | -145 | -140 | -135 | -130 | -125 | -120 | -115 | -110 | -105 | -100 | -95 | -90 | -85 | -80 | -75 | -70 | -65 | -60 | -55 | -50 | -45 | -40 | -35 | -30 | -25 | -20 | -15 | -10 | -5 | 0 | 5 | 10 | 15 | 20 | 25 | 30 | 35 | 40 | 45 | 50 | 55 | 60 | 65 | 70 | 75 | 80 | 85 | 90 | 95 | 100 | 105 | 110 | 115 | 120 | 125 | 130 | 135 | 140 | 145 | 150 | 155 | 160 | 165 | 170 | 175 | 180 | 185 | 190 | 195 | 200 | 205 | 210 | 215 | 220 | 225 | 230 | 235 | 240 | 245 | 250 | 255 | 260 | 265 | 270 | 275 | 280 | 285 | 290 | 295 | 300 | 305 | 310 | 315 | 320 | 325 | 330 | 335 | 340 | 345 | 350 | 355 | 360 | 365 | 370 | 375 | 380 | 385 | 390 | 395 | 400 | 405 | 410 | 415 | 420 | 425 | 430 | 435 | 440 | 445 | 450 | 455 | 460 | 465 | 470 | 475 | 480 | 485 | 490 | 495 | 500 | 505 | 510 | 515 | 520 | 525 | 530 | 535 | 540 | 545 | 550 | 555 | 560 | 565 | 570 | 575 | 580 | 585 | 590 | 595 | 600 | 605 | 610 | 615 | 620 | 625 | 630 | 635 | 640 | 645 | 650 | 655 | 660 | 665 | 670 | 675 | 680 | 685 | 690 | 695 | 700 | 705 | 710 | 715 | 720 | 725 | 730 | 735 | 740 | 745 | 750 | 755 | 760 | 765 | 770 | 775 | 780 | 785 | 790 | 795 | 800 | 805 | 810 | 815 | 820 | 825 | 830 | 835 | 840 | 845 | 850 | 855 | 860 | 865 | 870 | 875 | 880 | 885 | 890 | 895 | 900 | 905 | 910 | 915 | 920 | 925 | 930 | 935 | 940 | 945 | 950 | 955 | 960 | 965 | 970 | 975 | 980 | 985 | 990 | 995 | 1000 | 1005 | 1010 | 1015 | 1020 | 1025 | 1030 | 1035 | 1040 | 1045 | 1050 | 1055 | 1060 | 1065 | 1070 | 1075 | 1080 | 1085 | 1090 | 1095 | 1100 | 1105 | 1110 | 1115 | 1120 | 1125 | 1130 | 1135 | 1140 | 1145 | 1150 | 1155 | 1160 | 1165 | 1170 | 1175 | 1180 | 1185 | 1190 | 1195 |
| --- | --- | --- | --- | --- | --- | --- | --- | --- | --- | --- | --- | --- | --- | --- | --- | --- | --- | --- | --- | --- | --- | --- | --- | --- | --- | --- | --- | --- | --- | --- | --- | --- | --- | --- | --- | --- | --- | --- | --- | --- | --- | --- | --- | --- | --- | --- | --- | --- | --- | --- | --- | --- | --- | --- | --- | --- | --- | --- | --- | --- | --- | --- | --- | --- | --- | --- | --- | --- | --- | --- | --- | --- | --- | --- | --- | --- | --- | --- | --- | --- | --- | --- | --- | --- | --- | --- | --- | --- | --- | --- | --- | --- | --- | --- | --- | --- | --- | --- | --- | --- | --- | --- | --- | --- | --- | --- | --- | --- | --- | --- | --- | --- | --- | --- | --- | --- | --- | --- | --- | --- | --- | --- | --- | --- | --- | --- | --- | --- | --- | --- | --- | --- | --- | --- | --- | --- | --- | --- | --- | --- | --- | --- | --- | --- | --- | --- | --- | --- | --- | --- | --- | --- | --- | --- | --- | --- | --- | --- | --- | --- | --- | --- | --- | --- | --- | --- | --- | --- | --- | --- | --- | --- | --- | --- | --- | --- | --- | --- | --- | --- | --- | --- | --- | --- | --- | --- | --- | --- | --- | --- | --- | --- | --- | --- | --- | --- | --- | --- | --- | --- | --- | --- | --- | --- | --- | --- | --- | --- | --- | --- | --- | --- | --- | --- | --- | --- | --- | --- | --- | --- | --- | --- | --- | --- | --- | --- | --- | --- | --- | --- | --- | --- | --- | --- | --- | --- | --- | --- | --- | --- | --- | --- | --- | --- | --- | --- | --- | --- | --- | --- | --- | --- | --- | --- | --- | --- | --- | --- | --- | --- | --- | --- | --- | --- | --- | --- | --- | --- | --- | --- | --- | --- | --- | --- | --- | --- | --- | --- | --- | --- |
| left anterior | 0.249005 | 0.256427 | 0.247153 | 0.237094 | 0.222855 | 0.214633 | 0.227149 | 0.247964 | 0.249419 | 0.233834 | 0.229589 | 0.221082 | 0.219810 | 0.226515 | 0.263315 | 0.332791 | 0.432845 | 0.437299 | 0.378608 | 0.310742 | 0.247139 | 0.214677 | 0.238512 | 0.261625 | 0.240154 | 0.216630 | 0.243831 | 0.330044 | 0.394116 | 0.404937 | 0.325694 | 0.218298 | 0.278845 | 0.737805 | 2.760643 | 4.244555 | 3.627336 | 2.037219 | 1.296153 | 0.900353 | 0.841255 | 0.688935 | 0.646012 | 0.558138 | 0.548068 | 0.515616 | 0.432702 | 0.300563 | 0.244870 | 0.222251 | 0.215788 | 0.215859 | 0.228747 | 0.273076 | 0.392182 | 0.627030 | 0.918367 | 0.944228 | 0.751873 | 0.461265 | 0.272244 | 0.215339 | 0.239464 | 0.377511 | 0.625542 | 0.812260 | 0.855156 | 0.781165 | 0.734383 | 0.669050 | 0.578552 | 0.463462 | 0.353667 | 0.253131 | 0.215107 | 0.231320 | 0.292643 | 0.392786 | 0.475796 | 0.535740 | 0.613645 | 0.584163 | 0.451694 | 0.335632 | 0.255916 | 0.223663 | 0.214628 | 0.225099 | 0.254402 | 0.289056 | 0.336498 | 0.373261 | 0.365527 | 0.338589 | 0.320875 | 0.290469 | 0.267654 | 0.249549 | 0.232234 | 0.219901 | 0.214908 | 0.219684 | 0.243516 | 0.290188 | 0.336219 | 0.353252 | 0.298809 | 0.243970 | 0.219745 | 0.214898 | 0.214681 | 0.214640 | 0.215304 | 0.216886 | 0.221909 | 0.227466 | 0.226816 | 0.224586 | 0.221575 | 0.216583 | 0.215071 | 0.215005 | 0.216287 | 0.221374 | 0.230896 | 0.236423 | 0.237560 | 0.234125 | 0.221001 | 0.214731 | 0.227099 | 0.262242 | 0.322043 | 0.396147 | 0.465618 | 0.499754 | 0.451313 | 0.398066 | 0.339939 | 0.302193 | 0.280703 | 0.280290 | 0.289329 | 0.320183 | 0.369643 | 0.409631 | 0.417136 | 0.357674 | 0.287956 | 0.237163 | 0.216080 | 0.223054 | 0.253689 | 0.289026 | 0.302665 | 0.295424 | 0.266214 | 0.236038 | 0.218803 | 0.214648 | 0.217395 | 0.219382 | 0.216007 | 0.215361 | 0.227416 | 0.259109 | 0.303971 | 0.352136 | 0.357678 | 0.330887 | 0.285822 | 0.258272 | 0.234364 | 0.226453 | 0.221403 | 0.218400 | 0.215099 | 0.214653 | 0.217227 | 0.221880 | 0.228470 | 0.227278 | 0.227319 | 0.232218 | 0.251726 | 0.281366 | 0.358545 | 0.491592 | 0.648466 | 0.733995 | 0.721021 | 0.591835 | 0.495911 | 0.415357 | 0.359697 | 0.363115 | 0.464066 | 0.681141 | 1.063590 | 1.497279 | 1.935648 | 2.142598 | 1.853358 | 1.248788 | 0.904837 | 0.696724 | 0.581439 | 0.541302 | 0.539311 | 0.578507 | 0.653447 | 0.723271 | 0.820110 | 1.060454 | 1.333101 | 1.389840 | 1.249071 | 0.944592 | 0.694432 | 0.524075 | 0.409399 | 0.336270 | 0.307809 | 0.294642 | 0.295002 | 0.304348 | 0.340529 | 0.383049 | 0.414085 | 0.412111 | 0.396313 | 0.373628 | 0.357212 | 0.341839 | 0.358415 | 0.408558 | 0.443046 | 0.461376 | 0.449205 | 0.374173 | 0.298487 | 0.244880 | 0.219861 | 0.214819 | 0.214664 | 0.214649 | 0.217845 | 0.226044 | 0.237732 | 0.247159 | 0.265556 | 0.288408 | 0.347018 | 0.466925 | 0.773883 | 1.069756 | 1.186986 | 0.950480 | 0.658260 | 0.418379 | 0.314824 | 0.271299 | 0.271934 | 0.303076 | 0.374779 | 0.473120 | 0.556966 | 0.565247 | 0.510368 | 0.459285 | 0.420340 | 0.380107 | 0.361504 | 0.361959 | 0.350948 | 0.341354 | 0.340146 | 0.334777 | 0.321050 | 0.320985 |
| right anterior | 3.221912 | 4.007240 | 4.332450 | 5.446765 | 3.389579 | 1.398419 | 0.867838 | 0.669280 | 0.531612 | 0.364796 | 0.248067 | 0.215447 | 0.273474 | 0.555569 | 0.857440 | 0.949924 | 0.834840 | 0.569780 | 0.333205 | 0.251556 | 0.223206 | 0.220654 | 0.223397 | 0.223530 | 0.222449 | 0.216768 | 0.220134 | 0.254560 | 0.291688 | 0.321276 | 0.320308 | 0.264143 | 0.228408 | 0.214773 | 0.219670 | 0.235181 | 0.233726 | 0.224815 | 0.214766 | 0.215629 | 0.216933 | 0.215504 | 0.214631 | 0.218285 | 0.224464 | 0.230668 | 0.228126 | 0.229562 | 0.227585 | 0.225750 | 0.221340 | 0.219434 | 0.216070 | 0.214627 | 0.216604 | 0.220567 | 0.222052 | 0.220624 | 0.219045 | 0.217518 | 0.217411 | 0.224680 | 0.279785 | 0.569350 | 1.480424 | 3.624053 | 9.207970 | 12.230435 | 7.969425 | 4.360513 | 2.382628 | 1.619421 | 1.256670 | 0.992416 | 0.878952 | 0.951582 | 0.916764 | 0.891371 | 0.900785 | 0.806360 | 0.626170 | 0.592967 | 0.609756 | 0.629438 | 0.764974 | 1.057015 | 1.600535 | 2.330584 | 4.068085 | 9.466340 | 27.565096 | 72.693121 | 180.076198 | 378.181935 | 645.154446 | 616.758363 | 393.421244 | 229.214142 | 135.627285 | 65.637085 | 33.206158 | 18.656753 | 10.831273 | 7.447597 | 6.124523 | 6.253808 | 7.216791 | 9.432337 | 11.252993 | 12.426557 | 12.383980 | 12.600611 | 11.660146 | 10.136324 | 8.684649 | 6.563977 | 5.291062 | 4.301002 | 3.226527 | 2.203904 | 1.848119 | 1.482306 | 1.274462 | 1.069617 | 0.965136 | 0.882179 | 0.862872 | 0.829501 | 0.807047 | 0.819135 | 0.746068 | 0.732508 | 0.794790 | 1.004368 | 1.353989 | 2.165703 | 3.247600 | 4.131094 | 3.539912 | 2.749980 | 2.081659 | 1.476683 | 1.048416 | 0.745280 | 0.502769 | 0.385100 | 0.332830 | 0.311085 | 0.307878 | 0.311571 | 0.293306 | 0.276161 | 0.267697 | 0.259429 | 0.252625 | 0.257901 | 0.270819 | 0.290029 | 0.313050 | 0.329699 | 0.344022 | 0.352124 | 0.335985 | 0.310189 | 0.286760 | 0.257094 | 0.232763 | 0.219080 | 0.214680 | 0.215486 | 0.217058 | 0.219349 | 0.219021 | 0.219588 | 0.218113 | 0.215394 | 0.214630 | 0.215376 | 0.218606 | 0.220412 | 0.222617 | 0.230919 | 0.239988 | 0.253343 | 0.273172 | 0.295348 | 0.312656 | 0.334915 | 0.340998 | 0.352985 | 0.372835 | 0.393361 | 0.404570 | 0.430289 | 0.488039 | 0.587100 | 0.712678 | 0.896025 | 1.034673 | 1.026353 | 0.917459 | 0.794233 | 0.661544 | 0.599095 | 0.538151 | 0.520547 | 0.517743 | 0.506210 | 0.460793 | 0.423354 | 0.354795 | 0.287035 | 0.244944 | 0.230496 | 0.231181 | 0.246907 | 0.279644 | 0.335132 | 0.424557 | 0.538200 | 0.672398 | 0.754620 | 0.725675 | 0.681621 | 0.549878 | 0.436969 | 0.422869 | 0.492214 | 0.590271 | 0.988994 | 2.081894 | 3.660267 | 5.364293 | 5.559605 | 4.306243 | 2.768983 | 1.525828 | 0.797567 | 0.493247 | 0.365897 | 0.324418 | 0.322247 | 0.350014 | 0.388470 | 0.432968 | 0.476430 | 0.545764 | 0.535421 | 0.509536 | 0.438546 | 0.382849 | 0.323538 | 0.295139 | 0.279976 | 0.292177 | 0.325416 | 0.398738 | 0.540806 | 0.783142 | 1.041466 | 1.192505 | 1.212480 | 1.123423 | 0.974193 | 0.819171 | 0.670643 | 0.492425 | 0.344191 | 0.247329 | 0.214915 | 0.234078 | 0.282526 | 0.342249 | 0.388344 | 0.396765 | 0.383437 | 0.344819 | 0.284460 | 0.265400 | 0.247882 |
| left central | 0.328939 | 0.365151 | 0.385470 | 0.436507 | 0.475612 | 0.453161 | 0.337951 | 0.271770 | 0.234821 | 0.216189 | 0.216466 | 0.218142 | 0.219045 | 0.223699 | 0.249771 | 0.300810 | 0.457140 | 0.767832 | 0.849978 | 0.647043 | 0.535355 | 0.408220 | 0.338587 | 0.276659 | 0.223199 | 0.220241 | 0.295574 | 0.502285 | 0.942010 | 1.456190 | 1.708938 | 1.534368 | 0.992293 | 0.460737 | 0.270701 | 0.225335 | 0.216681 | 0.219248 | 0.224161 | 0.233990 | 0.254448 | 0.263209 | 0.249327 | 0.251266 | 0.248884 | 0.244788 | 0.241986 | 0.240778 | 0.238759 | 0.234136 | 0.221572 | 0.214641 | 0.224345 | 0.241403 | 0.265233 | 0.286692 | 0.312812 | 0.343222 | 0.476223 | 0.623297 | 0.812826 | 0.725019 | 0.550278 | 0.354860 | 0.240009 | 0.216213 | 0.248597 | 0.325082 | 0.411421 | 0.388318 | 0.315244 | 0.250499 | 0.218680 | 0.214716 | 0.217344 | 0.226257 | 0.230532 | 0.228167 | 0.228826 | 0.229146 | 0.230113 | 0.234931 | 0.244794 | 0.263325 | 0.292535 | 0.325519 | 0.337051 | 0.323338 | 0.277932 | 0.234268 | 0.216092 | 0.295029 | 0.614201 | 1.644950 | 4.551854 | 10.045095 | 14.962433 | 17.175197 | 15.279050 | 9.246702 | 4.487937 | 1.692509 | 0.800397 | 0.519514 | 0.378550 | 0.292897 | 0.255625 | 0.232979 | 0.221482 | 0.217151 | 0.215645 | 0.215045 | 0.214849 | 0.214626 | 0.214876 | 0.216814 | 0.217389 | 0.215733 | 0.215225 | 0.214705 | 0.217851 | 0.222982 | 0.224994 | 0.225368 | 0.218913 | 0.214688 | 0.216545 | 0.226515 | 0.243743 | 0.260286 | 0.267039 | 0.270082 | 0.255883 | 0.243561 | 0.231387 | 0.224497 | 0.220541 | 0.220014 | 0.220714 | 0.222826 | 0.224860 | 0.223539 | 0.219357 | 0.215073 | 0.216916 | 0.231929 | 0.253682 | 0.284801 | 0.320492 | 0.377411 | 0.494146 | 0.686535 | 0.674858 | 0.626992 | 0.626938 | 0.525340 | 0.392690 | 0.301963 | 0.244372 | 0.215563 | 0.221221 | 0.240754 | 0.248115 | 0.242603 | 0.229146 | 0.223084 | 0.218414 | 0.215510 | 0.215268 | 0.215189 | 0.214983 | 0.220186 | 0.233522 | 0.257246 | 0.287105 | 0.316293 | 0.344522 | 0.360113 | 0.335222 | 0.299256 | 0.261523 | 0.233433 | 0.215899 | 0.215662 | 0.224293 | 0.229642 | 0.228346 | 0.221489 | 0.217060 | 0.214661 | 0.214737 | 0.214945 | 0.215631 | 0.216358 | 0.216100 | 0.215542 | 0.214979 | 0.214718 | 0.215615 | 0.218314 | 0.222103 | 0.226575 | 0.229013 | 0.229103 | 0.226788 | 0.223682 | 0.218922 | 0.216421 | 0.215478 | 0.215047 | 0.214870 | 0.215247 | 0.215528 | 0.215555 | 0.215454 | 0.214872 | 0.214687 | 0.216691 | 0.223562 | 0.235341 | 0.239041 | 0.235661 | 0.225384 | 0.218697 | 0.215751 | 0.215420 | 0.215664 | 0.216278 | 0.215600 | 0.215199 | 0.214634 | 0.214784 | 0.215312 | 0.215220 | 0.214696 | 0.215358 | 0.218992 | 0.222432 | 0.218971 | 0.214838 | 0.217197 | 0.227398 | 0.239597 | 0.249223 | 0.247605 | 0.236923 | 0.226031 | 0.216919 | 0.215427 | 0.226266 | 0.252341 | 0.281192 | 0.308414 | 0.331689 | 0.348895 | 0.342047 | 0.337125 | 0.336298 | 0.352096 | 0.361585 | 0.376469 | 0.402887 | 0.437179 | 0.497605 | 0.589536 | 0.716924 | 0.999003 | 1.272871 | 1.377588 | 1.411434 | 1.203523 | 0.937857 | 0.797630 | 0.697295 | 0.592653 | 0.524368 | 0.439023 | 0.386039 | 0.358818 | 0.331898 |
| right central | 0.222389 | 0.224464 | 0.232364 | 0.256838 | 0.273095 | 0.302459 | 0.291513 | 0.275884 | 0.242190 | 0.217419 | 0.222422 | 0.253556 | 0.312544 | 0.334426 | 0.296945 | 0.248047 | 0.219590 | 0.215972 | 0.216153 | 0.214627 | 0.218100 | 0.238194 | 0.277723 | 0.276565 | 0.261656 | 0.235301 | 0.218742 | 0.215301 | 0.215423 | 0.217542 | 0.225444 | 0.236999 | 0.232333 | 0.215969 | 0.225468 | 0.297300 | 0.450184 | 0.601151 | 0.621889 | 0.478030 | 0.319094 | 0.232251 | 0.214867 | 0.234956 | 0.278074 | 0.351810 | 0.454312 | 0.506550 | 0.467262 | 0.406598 | 0.318061 | 0.254218 | 0.221042 | 0.217146 | 0.256833 | 0.347230 | 0.522765 | 0.759235 | 1.108219 | 1.498344 | 1.973387 | 2.316737 | 2.145260 | 1.130654 | 0.521823 | 0.262038 | 0.215313 | 0.285164 | 0.558336 | 1.238526 | 1.751979 | 1.581809 | 0.936376 | 0.474555 | 0.276793 | 0.215710 | 0.248340 | 0.387839 | 0.736958 | 1.228185 | 1.114342 | 0.625898 | 0.339810 | 0.235549 | 0.214645 | 0.230858 | 0.264289 | 0.283856 | 0.330860 | 0.423364 | 0.622932 | 0.913767 | 1.409367 | 1.913875 | 2.552376 | 2.672341 | 2.925966 | 3.135475 | 2.981674 | 2.561841 | 2.366162 | 2.266279 | 1.895467 | 1.395420 | 1.046697 | 0.769321 | 0.537361 | 0.472039 | 0.493645 | 0.620053 | 0.775904 | 0.975400 | 1.206715 | 1.472459 | 1.474490 | 1.263730 | 0.815027 | 0.519416 | 0.361063 | 0.288750 | 0.262092 | 0.280888 | 0.310142 | 0.351809 | 0.381509 | 0.427402 | 0.457337 | 0.487729 | 0.506695 | 0.548527 | 0.566500 | 0.578649 | 0.536513 | 0.505177 | 0.473540 | 0.446325 | 0.413842 | 0.371678 | 0.309278 | 0.267061 | 0.234844 | 0.218927 | 0.214705 | 0.214687 | 0.216121 | 0.219879 | 0.229265 | 0.246514 | 0.249547 | 0.232461 | 0.217398 | 0.217842 | 0.249147 | 0.291739 | 0.325015 | 0.314704 | 0.271817 | 0.241237 | 0.223386 | 0.215657 | 0.214626 | 0.214900 | 0.217177 | 0.227041 | 0.239103 | 0.252593 | 0.250451 | 0.250385 | 0.252448 | 0.268113 | 0.284218 | 0.316657 | 0.348020 | 0.418909 | 0.483135 | 0.588528 | 0.620659 | 0.597265 | 0.508778 | 0.434906 | 0.363552 | 0.330986 | 0.298017 | 0.285234 | 0.285891 | 0.307971 | 0.353419 | 0.446606 | 0.566406 | 0.672160 | 0.635455 | 0.484566 | 0.392315 | 0.336519 | 0.310198 | 0.302436 | 0.301390 | 0.302907 | 0.302085 | 0.288421 | 0.268167 | 0.258222 | 0.243900 | 0.231522 | 0.222924 | 0.219485 | 0.218450 | 0.222990 | 0.233641 | 0.251507 | 0.279493 | 0.322628 | 0.325867 | 0.310070 | 0.300362 | 0.283758 | 0.277084 | 0.309436 | 0.369155 | 0.434308 | 0.489115 | 0.461998 | 0.386342 | 0.304127 | 0.241378 | 0.216302 | 0.220651 | 0.243572 | 0.260751 | 0.246798 | 0.224723 | 0.215317 | 0.238142 | 0.324617 | 0.500420 | 0.823369 | 1.300394 | 1.916550 | 1.899894 | 1.878253 | 1.405766 | 0.939425 | 0.626739 | 0.465012 | 0.378857 | 0.378627 | 0.435218 | 0.577270 | 0.852940 | 1.152021 | 1.356299 | 1.304157 | 0.954022 | 0.738659 | 0.599626 | 0.506374 | 0.446499 | 0.381252 | 0.327184 | 0.298892 | 0.264979 | 0.241557 | 0.229239 | 0.220799 | 0.217543 | 0.217785 | 0.219143 | 0.222925 | 0.231840 | 0.243023 | 0.260509 | 0.286865 | 0.382685 | 0.592632 | 0.845430 | 1.152203 | 1.355284 | 1.139281 | 1.021622 | 0.943383 |
| left posterior | 0.416210 | 0.514106 | 0.693033 | 1.174084 | 2.559619 | 3.398618 | 2.176660 | 0.791328 | 0.293190 | 0.214977 | 0.278676 | 0.448976 | 0.648011 | 0.677784 | 0.554799 | 0.336014 | 0.224995 | 0.231967 | 0.328709 | 0.427738 | 0.321901 | 0.221089 | 0.242756 | 0.390427 | 0.734898 | 1.392381 | 1.762485 | 1.238461 | 0.742150 | 0.396116 | 0.233660 | 0.225946 | 0.273643 | 0.289591 | 0.268177 | 0.246955 | 0.230231 | 0.221516 | 0.215089 | 0.219057 | 0.241211 | 0.295004 | 0.416888 | 0.647483 | 1.051534 | 1.530675 | 1.774099 | 1.984368 | 2.143305 | 2.552224 | 3.310768 | 3.037746 | 1.718429 | 0.732436 | 0.306046 | 0.215185 | 0.260508 | 0.430140 | 0.684139 | 0.649710 | 0.393439 | 0.220491 | 0.412729 | 3.466341 | 28.971601 | 148.219908 | 542.820107 | 1302.098233 | 1718.290397 | 1021.777173 | 387.581810 | 95.536834 | 23.142179 | 6.625720 | 2.426596 | 1.154696 | 0.658302 | 0.436023 | 0.336822 | 0.294231 | 0.271813 | 0.267105 | 0.273232 | 0.293050 | 0.333189 | 0.384914 | 0.455488 | 0.551876 | 0.726730 | 1.463475 | 4.923059 | 20.045204 | 57.930078 | 101.422583 | 123.759393 | 132.948256 | 111.852876 | 71.521280 | 36.792413 | 18.964729 | 10.524844 | 6.798716 | 5.350901 | 5.671724 | 7.011919 | 8.963693 | 10.757208 | 12.347827 | 12.832836 | 12.645128 | 11.941224 | 10.752662 | 7.915590 | 5.415178 | 3.303038 | 2.135449 | 1.601561 | 1.399656 | 1.329614 | 1.462605 | 1.815051 | 2.071164 | 2.120504 | 1.893114 | 1.367155 | 0.793276 | 0.433496 | 0.267603 | 0.216996 | 0.219278 | 0.233828 | 0.238609 | 0.234390 | 0.224671 | 0.220284 | 0.215864 | 0.214755 | 0.220758 | 0.233713 | 0.259555 | 0.292299 | 0.339168 | 0.418681 | 0.504637 | 0.578880 | 0.625402 | 0.649546 | 0.588792 | 0.532192 | 0.471497 | 0.435605 | 0.398836 | 0.370281 | 0.352511 | 0.331343 | 0.304686 | 0.274282 | 0.255223 | 0.239003 | 0.236600 | 0.241789 | 0.261575 | 0.288150 | 0.314174 | 0.325793 | 0.325497 | 0.308187 | 0.277886 | 0.246886 | 0.223601 | 0.214627 | 0.219132 | 0.222210 | 0.220527 | 0.217442 | 0.214649 | 0.216897 | 0.225240 | 0.242992 | 0.270449 | 0.297188 | 0.308615 | 0.311362 | 0.305254 | 0.296964 | 0.288945 | 0.273331 | 0.251064 | 0.229155 | 0.217084 | 0.214705 | 0.217170 | 0.220911 | 0.218526 | 0.214667 | 0.218818 | 0.237261 | 0.273393 | 0.330639 | 0.408696 | 0.495963 | 0.589721 | 0.667273 | 0.698113 | 0.664567 | 0.621329 | 0.550676 | 0.486219 | 0.439017 | 0.440696 | 0.474518 | 0.562940 | 0.724333 | 1.032932 | 1.378267 | 1.630587 | 1.579328 | 1.402734 | 1.105263 | 0.874537 | 0.664726 | 0.574567 | 0.515552 | 0.462819 | 0.404240 | 0.355065 | 0.300310 | 0.262915 | 0.235223 | 0.224497 | 0.221378 | 0.223550 | 0.234978 | 0.267753 | 0.327594 | 0.431149 | 0.546180 | 0.628172 | 0.604196 | 0.510010 | 0.424113 | 0.365452 | 0.314571 | 0.292226 | 0.274667 | 0.270076 | 0.282093 | 0.304407 | 0.323809 | 0.341514 | 0.333284 | 0.308674 | 0.288893 | 0.266026 | 0.247462 | 0.236260 | 0.228668 | 0.223281 | 0.222154 | 0.220438 | 0.219734 | 0.218446 | 0.216102 | 0.214775 | 0.214888 | 0.219564 | 0.232012 | 0.252287 | 0.281658 | 0.320214 | 0.384166 | 0.527146 | 0.778309 | 1.171032 | 1.663633 | 2.017988 | 1.871498 | 1.592471 | 1.513603 | 1.447340 |
| right posterior | 0.263200 | 0.263549 | 0.275434 | 0.275852 | 0.293226 | 0.347542 | 0.374904 | 0.338232 | 0.259539 | 0.215298 | 0.229629 | 0.270386 | 0.331067 | 0.373591 | 0.352905 | 0.322556 | 0.268468 | 0.228577 | 0.215556 | 0.214969 | 0.216294 | 0.214675 | 0.214725 | 0.215312 | 0.215120 | 0.217221 | 0.223709 | 0.227067 | 0.233380 | 0.246049 | 0.261704 | 0.273696 | 0.251397 | 0.222172 | 0.219187 | 0.294236 | 0.550428 | 1.060384 | 2.155568 | 4.286350 | 8.361681 | 16.056231 | 29.009953 | 46.471950 | 76.017864 | 93.420294 | 45.650509 | 9.910726 | 1.416456 | 0.393373 | 0.232184 | 0.214629 | 0.221068 | 0.224190 | 0.224834 | 0.222147 | 0.224437 | 0.227125 | 0.229672 | 0.220635 | 0.220125 | 0.395757 | 2.527222 | 45.824103 | 776.451193 | 5012.485895 | 9573.159442 | 5715.563966 | 1478.115358 | 241.797909 | 38.306047 | 7.336342 | 1.860257 | 0.575930 | 0.269815 | 0.215415 | 0.311160 | 0.611182 | 1.131820 | 1.784029 | 1.730303 | 1.225520 | 0.720930 | 0.414246 | 0.276682 | 0.225340 | 0.215401 | 0.239431 | 0.302425 | 0.430771 | 0.627738 | 1.073977 | 1.838731 | 3.368513 | 6.331164 | 11.528604 | 16.934242 | 21.219891 | 21.638297 | 19.144954 | 18.224346 | 20.364617 | 28.151339 | 49.809512 | 101.273418 | 152.632087 | 123.684296 | 59.237810 | 23.602685 | 10.652972 | 6.463872 | 4.331626 | 3.069117 | 2.602164 | 2.365625 | 2.516119 | 3.102784 | 4.073042 | 5.857007 | 9.454171 | 14.161189 | 18.108639 | 19.462724 | 17.105874 | 12.761799 | 8.539310 | 5.108951 | 3.041312 | 1.846608 | 1.359529 | 1.055572 | 0.952676 | 0.935025 | 0.934631 | 0.964963 | 1.156251 | 1.382576 | 1.536639 | 1.619080 | 1.545684 | 1.471600 | 1.359077 | 1.306314 | 1.511119 | 2.358290 | 3.851449 | 5.638199 | 6.359909 | 5.016381 | 3.561448 | 2.455901 | 1.695591 | 1.189853 | 0.927852 | 0.816667 | 0.841723 | 0.943450 | 1.196295 | 1.611155 | 2.191065 | 2.630570 | 3.270124 | 3.597977 | 3.521155 | 2.919549 | 2.420280 | 1.948441 | 1.756627 | 1.519234 | 1.362451 | 1.279250 | 1.363845 | 1.669232 | 2.576814 | 3.995282 | 5.804932 | 6.178196 | 4.993053 | 3.537228 | 2.387808 | 1.547620 | 1.022701 | 0.722028 | 0.557207 | 0.488243 | 0.504796 | 0.572473 | 0.662793 | 0.787183 | 0.919915 | 0.970259 | 0.944966 | 0.885197 | 0.852744 | 0.861535 | 0.943447 | 1.008283 | 1.024313 | 1.036890 | 1.121868 | 1.298186 | 1.660791 | 2.319890 | 3.402034 | 5.089219 | 7.538155 | 9.922522 | 11.486352 | 10.749578 | 8.165551 | 6.291953 | 5.228727 | 4.932000 | 5.924229 | 7.128456 | 7.822491 | 8.063169 | 7.252072 | 5.364021 | 3.774968 | 2.480474 | 1.747732 | 1.422099 | 1.326789 | 1.410052 | 1.730615 | 2.065182 | 2.398100 | 2.567092 | 2.564884 | 2.337585 | 2.069906 | 1.818108 | 1.585156 | 1.399038 | 1.278194 | 1.178275 | 1.171929 | 1.247825 | 1.406964 | 1.570112 | 1.775520 | 1.978179 | 2.283854 | 2.787110 | 3.875405 | 6.975592 | 13.287298 | 19.020944 | 16.187181 | 9.691325 | 5.743855 | 3.465794 | 2.455670 | 2.176732 | 2.510245 | 3.180423 | 4.411835 | 5.836959 | 7.105300 | 6.431305 | 4.919164 | 3.545156 | 2.434865 | 1.689721 | 1.138537 | 0.794903 | 0.620448 | 0.603588 | 0.624535 | 0.742049 | 0.996047 | 1.294151 | 1.438298 | 1.653855 | 1.548784 | 1.333861 | 1.259613 | 1.189897 | 1.128201 |
| all electrodes | 0.413794 | 0.407702 | 0.365503 | 0.257875 | 0.216799 | 0.218884 | 0.270239 | 0.385468 | 0.513044 | 0.681450 | 0.820204 | 0.726979 | 0.759811 | 0.646460 | 0.421495 | 0.288688 | 0.236977 | 0.216738 | 0.214752 | 0.220504 | 0.229297 | 0.231151 | 0.218572 | 0.214686 | 0.230933 | 0.313643 | 0.510553 | 0.665234 | 0.833129 | 0.742785 | 0.577943 | 0.399676 | 0.285018 | 0.240856 | 0.240910 | 0.239086 | 0.241286 | 0.248421 | 0.256764 | 0.244714 | 0.227886 | 0.222998 | 0.215424 | 0.215140 | 0.216361 | 0.215705 | 0.223024 | 0.220297 | 0.231287 | 0.241462 | 0.251847 | 0.265120 | 0.291257 | 0.322843 | 0.363230 | 0.399232 | 0.377644 | 0.350114 | 0.307911 | 0.262118 | 0.217029 | 0.236094 | 0.436297 | 1.399166 | 6.477279 | 29.449168 | 101.832902 | 103.903613 | 49.710698 | 18.867140 | 11.876587 | 10.212857 | 9.187683 | 5.678124 | 2.685680 | 0.934982 | 0.407294 | 0.253132 | 0.217242 | 0.214765 | 0.214753 | 0.220550 | 0.244881 | 0.269919 | 0.267312 | 0.260157 | 0.249582 | 0.234116 | 0.233345 | 0.248277 | 0.290683 | 0.399461 | 0.622047 | 0.995616 | 1.821419 | 2.888007 | 4.396781 | 5.865631 | 5.401032 | 3.139978 | 1.599556 | 0.615900 | 0.317611 | 0.248976 | 0.230350 | 0.236343 | 0.293073 | 0.415839 | 0.692145 | 1.172540 | 1.887408 | 2.150164 | 2.226088 | 1.630178 | 1.328598 | 1.038206 | 0.921656 | 0.767854 | 0.771228 | 0.848698 | 1.112668 | 1.525348 | 2.520671 | 2.996663 | 2.372855 | 1.437736 | 0.911605 | 0.571924 | 0.412090 | 0.327625 | 0.292039 | 0.287594 | 0.322227 | 0.378853 | 0.463465 | 0.581720 | 0.744078 | 0.949517 | 1.452496 | 2.164037 | 3.076317 | 3.381919 | 2.738077 | 2.520715 | 2.827350 | 2.988671 | 2.841609 | 2.663380 | 2.615266 | 2.657458 | 2.719139 | 2.640010 | 2.655084 | 2.030276 | 1.275614 | 0.779787 | 0.561347 | 0.442106 | 0.429843 | 0.406493 | 0.370498 | 0.315243 | 0.268537 | 0.234049 | 0.219430 | 0.214648 | 0.218052 | 0.221757 | 0.222269 | 0.221316 | 0.221335 | 0.223080 | 0.225262 | 0.231031 | 0.237901 | 0.232911 | 0.222326 | 0.214899 | 0.221044 | 0.252159 | 0.323644 | 0.466161 | 0.603825 | 0.722562 | 0.755463 | 0.690062 | 0.518927 | 0.359648 | 0.251769 | 0.218409 | 0.216250 | 0.219464 | 0.216896 | 0.214661 | 0.220609 | 0.239923 | 0.259216 | 0.265382 | 0.258773 | 0.250043 | 0.245595 | 0.250328 | 0.280202 | 0.358621 | 0.451099 | 0.574992 | 0.741272 | 0.917775 | 1.038847 | 1.186602 | 1.111026 | 0.924922 | 0.636795 | 0.433915 | 0.318968 | 0.279271 | 0.268718 | 0.288718 | 0.352400 | 0.429015 | 0.426614 | 0.360029 | 0.298690 | 0.255534 | 0.241403 | 0.248673 | 0.300544 | 0.410107 | 0.611549 | 0.895249 | 1.146248 | 0.950599 | 0.807578 | 0.674256 | 0.610821 | 0.592431 | 0.566125 | 0.498141 | 0.425535 | 0.355759 | 0.295668 | 0.261997 | 0.249854 | 0.251960 | 0.259223 | 0.283091 | 0.304883 | 0.311678 | 0.307340 | 0.283889 | 0.244779 | 0.220801 | 0.214800 | 0.224325 | 0.236634 | 0.255446 | 0.261484 | 0.242264 | 0.222384 | 0.214636 | 0.231467 | 0.292097 | 0.384275 | 0.515609 | 0.597295 | 0.578300 | 0.473783 | 0.343289 | 0.255301 | 0.221080 | 0.215070 | 0.228534 | 0.249183 | 0.271234 | 0.284917 | 0.307130 | 0.329996 | 0.356816 | 0.354734 | 0.383101 |

E) neutral vs angry

  
|  | time window | peak latency | cluster *p* | peak Cohen's *d* |  | | | |
| **all electrodes** | 120 - 600 ms | 160 ms | 0.0002 | 1.3357 |  | | | |
|  | | | | | | | | |

Model correlations, cluster permutation tests

|  | **left hemisphere** | | | | **right hemisphere** | | | |
|  | time window | peak latency | cluster *p* | peak Cohen's *d* | time window | peak latency | cluster *p* | peak Cohen's *d* |
| **anterior** |  | | | |  | | | |
| **central** |  | | | | 120 - 475 ms | 160 ms | 0.0002 | 1.3139 |
| **posterior** | 120 - 235 ms | 160 ms | 0.0183 | 1.1281 | 95 - 1195 ms | 155 ms | 0.0002 | 1.2863 |
 250 - 540 ms | 415 ms | 0.0043 | 0.8329 |  | | | | 580 - 815 ms | 660 ms | 0.0113 | 0.5732 |  | | | |

  

Model correlations, Bayesian statistics

|  | -200 | -195 | -190 | -185 | -180 | -175 | -170 | -165 | -160 | -155 | -150 | -145 | -140 | -135 | -130 | -125 | -120 | -115 | -110 | -105 | -100 | -95 | -90 | -85 | -80 | -75 | -70 | -65 | -60 | -55 | -50 | -45 | -40 | -35 | -30 | -25 | -20 | -15 | -10 | -5 | 0 | 5 | 10 | 15 | 20 | 25 | 30 | 35 | 40 | 45 | 50 | 55 | 60 | 65 | 70 | 75 | 80 | 85 | 90 | 95 | 100 | 105 | 110 | 115 | 120 | 125 | 130 | 135 | 140 | 145 | 150 | 155 | 160 | 165 | 170 | 175 | 180 | 185 | 190 | 195 | 200 | 205 | 210 | 215 | 220 | 225 | 230 | 235 | 240 | 245 | 250 | 255 | 260 | 265 | 270 | 275 | 280 | 285 | 290 | 295 | 300 | 305 | 310 | 315 | 320 | 325 | 330 | 335 | 340 | 345 | 350 | 355 | 360 | 365 | 370 | 375 | 380 | 385 | 390 | 395 | 400 | 405 | 410 | 415 | 420 | 425 | 430 | 435 | 440 | 445 | 450 | 455 | 460 | 465 | 470 | 475 | 480 | 485 | 490 | 495 | 500 | 505 | 510 | 515 | 520 | 525 | 530 | 535 | 540 | 545 | 550 | 555 | 560 | 565 | 570 | 575 | 580 | 585 | 590 | 595 | 600 | 605 | 610 | 615 | 620 | 625 | 630 | 635 | 640 | 645 | 650 | 655 | 660 | 665 | 670 | 675 | 680 | 685 | 690 | 695 | 700 | 705 | 710 | 715 | 720 | 725 | 730 | 735 | 740 | 745 | 750 | 755 | 760 | 765 | 770 | 775 | 780 | 785 | 790 | 795 | 800 | 805 | 810 | 815 | 820 | 825 | 830 | 835 | 840 | 845 | 850 | 855 | 860 | 865 | 870 | 875 | 880 | 885 | 890 | 895 | 900 | 905 | 910 | 915 | 920 | 925 | 930 | 935 | 940 | 945 | 950 | 955 | 960 | 965 | 970 | 975 | 980 | 985 | 990 | 995 | 1000 | 1005 | 1010 | 1015 | 1020 | 1025 | 1030 | 1035 | 1040 | 1045 | 1050 | 1055 | 1060 | 1065 | 1070 | 1075 | 1080 | 1085 | 1090 | 1095 | 1100 | 1105 | 1110 | 1115 | 1120 | 1125 | 1130 | 1135 | 1140 | 1145 | 1150 | 1155 | 1160 | 1165 | 1170 | 1175 | 1180 | 1185 | 1190 | 1195 |
| --- | --- | --- | --- | --- | --- | --- | --- | --- | --- | --- | --- | --- | --- | --- | --- | --- | --- | --- | --- | --- | --- | --- | --- | --- | --- | --- | --- | --- | --- | --- | --- | --- | --- | --- | --- | --- | --- | --- | --- | --- | --- | --- | --- | --- | --- | --- | --- | --- | --- | --- | --- | --- | --- | --- | --- | --- | --- | --- | --- | --- | --- | --- | --- | --- | --- | --- | --- | --- | --- | --- | --- | --- | --- | --- | --- | --- | --- | --- | --- | --- | --- | --- | --- | --- | --- | --- | --- | --- | --- | --- | --- | --- | --- | --- | --- | --- | --- | --- | --- | --- | --- | --- | --- | --- | --- | --- | --- | --- | --- | --- | --- | --- | --- | --- | --- | --- | --- | --- | --- | --- | --- | --- | --- | --- | --- | --- | --- | --- | --- | --- | --- | --- | --- | --- | --- | --- | --- | --- | --- | --- | --- | --- | --- | --- | --- | --- | --- | --- | --- | --- | --- | --- | --- | --- | --- | --- | --- | --- | --- | --- | --- | --- | --- | --- | --- | --- | --- | --- | --- | --- | --- | --- | --- | --- | --- | --- | --- | --- | --- | --- | --- | --- | --- | --- | --- | --- | --- | --- | --- | --- | --- | --- | --- | --- | --- | --- | --- | --- | --- | --- | --- | --- | --- | --- | --- | --- | --- | --- | --- | --- | --- | --- | --- | --- | --- | --- | --- | --- | --- | --- | --- | --- | --- | --- | --- | --- | --- | --- | --- | --- | --- | --- | --- | --- | --- | --- | --- | --- | --- | --- | --- | --- | --- | --- | --- | --- | --- | --- | --- | --- | --- | --- | --- | --- | --- | --- | --- | --- | --- | --- | --- | --- | --- | --- | --- | --- | --- | --- | --- | --- | --- | --- | --- | --- | --- | --- | --- | --- | --- | --- |
| left anterior | 0.314901 | 0.338487 | 0.379532 | 0.500486 | 0.724065 | 1.678487 | 3.742264 | 9.715261 | 33.270690 | 47.645767 | 17.101599 | 4.849047 | 1.525277 | 0.581526 | 0.326085 | 0.276366 | 0.293130 | 0.388877 | 0.732535 | 1.719756 | 3.675966 | 5.576814 | 3.455727 | 1.021546 | 0.383554 | 0.233954 | 0.224408 | 0.325327 | 0.539224 | 0.630913 | 0.573198 | 0.326268 | 0.229912 | 0.218804 | 0.251244 | 0.259133 | 0.243485 | 0.239592 | 0.239540 | 0.236222 | 0.237752 | 0.252918 | 0.302292 | 0.366736 | 0.459516 | 0.824760 | 1.727560 | 2.031966 | 1.692474 | 0.897608 | 0.474516 | 0.321276 | 0.248556 | 0.221827 | 0.215794 | 0.214665 | 0.218131 | 0.227772 | 0.239538 | 0.246520 | 0.260288 | 0.248924 | 0.237783 | 0.240503 | 0.285246 | 0.371825 | 0.601241 | 0.886591 | 1.202371 | 1.190573 | 1.053103 | 0.739880 | 0.512099 | 0.351382 | 0.250572 | 0.215452 | 0.272114 | 0.494969 | 1.020657 | 1.591220 | 1.540523 | 1.127105 | 0.751213 | 0.516007 | 0.340466 | 0.259181 | 0.226896 | 0.214751 | 0.223104 | 0.255464 | 0.323951 | 0.429512 | 0.558325 | 0.711373 | 7.199846e-01 | 6.297123e-01 | 4.906631e-01 | 3.616241e-01 | 2.693282e-01 | 0.228315 | 0.215073 | 0.216491 | 0.220860 | 0.221221 | 0.220907 | 0.218573 | 0.215536 | 0.214906 | 0.218087 | 0.225278 | 0.244522 | 0.271409 | 0.309006 | 0.342684 | 0.392925 | 0.445690 | 0.544674 | 0.789945 | 1.446649 | 2.889452 | 4.836862 | 6.634484 | 5.933912 | 5.089072 | 4.130013 | 3.056804 | 2.101933 | 1.412464 | 0.863964 | 0.532274 | 0.372672 | 0.290810 | 0.246202 | 0.225567 | 0.217646 | 0.214645 | 0.217432 | 0.216794 | 0.214705 | 0.218874 | 0.228110 | 0.235926 | 0.239555 | 0.242520 | 0.230811 | 0.221878 | 0.215649 | 0.218344 | 0.249035 | 0.314214 | 0.390393 | 0.451198 | 0.465577 | 0.465378 | 0.434307 | 0.364499 | 0.290820 | 0.251621 | 0.228645 | 0.216145 | 0.215578 | 0.219002 | 0.224519 | 0.232847 | 0.246412 | 0.274938 | 0.338113 | 0.446901 | 0.618045 | 0.787810 | 0.857300 | 0.716897 | 0.533259 | 0.370713 | 0.280699 | 0.241242 | 0.233082 | 0.238387 | 0.263220 | 0.291645 | 0.319415 | 0.330297 | 0.328722 | 0.331525 | 0.357259 | 0.400712 | 0.446449 | 0.510619 | 0.578645 | 0.592322 | 0.475312 | 0.376221 | 0.312292 | 0.275148 | 0.260644 | 0.265735 | 0.300506 | 0.354839 | 0.394939 | 0.388233 | 0.355653 | 0.301622 | 0.248863 | 0.216642 | 0.222062 | 0.248200 | 0.262108 | 0.240084 | 0.216430 | 0.226051 | 0.284388 | 0.363803 | 0.419159 | 0.449958 | 0.395942 | 0.311800 | 0.260550 | 0.234987 | 0.226798 | 0.226365 | 0.222978 | 0.216169 | 0.215125 | 0.226012 | 0.271327 | 0.381484 | 0.558170 | 0.743351 | 0.758441 | 0.656551 | 0.553290 | 0.540757 | 0.629765 | 0.904914 | 1.287555 | 1.698369 | 1.967388 | 1.543698 | 0.949736 | 0.572482 | 0.366413 | 0.267224 | 0.231399 | 0.217954 | 0.215042 | 0.214629 | 0.214634 | 0.214628 | 0.215323 | 0.217711 | 0.222203 | 0.223449 | 0.221475 | 0.217131 | 0.214836 | 0.214900 | 0.215784 | 0.217439 | 0.217045 | 0.216826 | 0.216964 | 0.216854 | 0.217326 | 0.220028 | 0.223420 | 0.230147 | 0.246050 | 0.260953 | 0.270631 | 0.285345 | 0.292829 | 0.295869 | 0.305940 | 0.322384 | 0.345437 | 0.392413 | 0.461487 | 0.556431 | 0.614321 | 0.711569 |
| right anterior | 0.219687 | 0.218767 | 0.214737 | 0.224329 | 0.252311 | 0.304511 | 0.359357 | 0.382385 | 0.410982 | 0.447518 | 0.550489 | 1.018583 | 1.308918 | 0.949236 | 0.704209 | 0.505544 | 0.375328 | 0.327693 | 0.294106 | 0.271574 | 0.267212 | 0.249255 | 0.245278 | 0.268680 | 0.274975 | 0.276873 | 0.280390 | 0.300913 | 0.320513 | 0.317305 | 0.297710 | 0.355539 | 0.375100 | 0.336427 | 0.287860 | 0.236864 | 0.214950 | 0.240093 | 0.322636 | 0.345205 | 0.299280 | 0.267748 | 0.228114 | 0.215537 | 0.231311 | 0.257612 | 0.267931 | 0.242761 | 0.215311 | 0.222975 | 0.251969 | 0.271611 | 0.293079 | 0.295739 | 0.314065 | 0.315306 | 0.327049 | 0.339681 | 0.349261 | 0.325006 | 0.301414 | 0.257115 | 0.236471 | 0.235794 | 0.260799 | 0.340460 | 0.611552 | 1.360737 | 3.263837 | 6.710260 | 11.725462 | 15.552841 | 13.052507 | 7.684642 | 3.747407 | 1.974717 | 1.143742 | 0.738858 | 0.521870 | 0.425929 | 0.340734 | 0.296163 | 0.270461 | 0.243848 | 0.227275 | 0.221870 | 0.221818 | 0.226226 | 0.248167 | 0.327274 | 0.559599 | 1.148145 | 2.480444 | 5.555463 | 1.068477e+01 | 1.558766e+01 | 1.668623e+01 | 1.481514e+01 | 1.082047e+01 | 7.060683 | 3.889963 | 2.041394 | 1.048828 | 0.684299 | 0.528591 | 0.481201 | 0.434554 | 0.395068 | 0.352834 | 0.313227 | 0.281512 | 0.270715 | 0.265502 | 0.283388 | 0.320872 | 0.363712 | 0.455703 | 0.629289 | 0.882124 | 1.396187 | 2.407539 | 4.225226 | 6.968796 | 9.230026 | 10.175147 | 10.616226 | 9.219605 | 7.704380 | 6.215479 | 4.851956 | 3.583053 | 2.627707 | 1.763737 | 1.269217 | 0.810744 | 0.538374 | 0.384027 | 0.290997 | 0.237362 | 0.217512 | 0.214975 | 0.218182 | 0.218646 | 0.219454 | 0.218280 | 0.217742 | 0.219453 | 0.220666 | 0.218476 | 0.214783 | 0.218920 | 0.240385 | 0.291472 | 0.396065 | 0.498799 | 0.483278 | 0.394279 | 0.288798 | 0.222828 | 0.223275 | 0.287116 | 0.400774 | 0.567272 | 0.696029 | 0.758511 | 0.784787 | 0.880584 | 0.904490 | 0.905019 | 0.879604 | 0.702496 | 0.489192 | 0.367747 | 0.309779 | 0.283117 | 0.282743 | 0.282749 | 0.288398 | 0.301362 | 0.304329 | 0.297236 | 0.308718 | 0.323928 | 0.331739 | 0.339351 | 0.353039 | 0.372609 | 0.383624 | 0.378848 | 0.358388 | 0.344546 | 0.311573 | 0.286697 | 0.271027 | 0.260839 | 0.252902 | 0.246558 | 0.241866 | 0.250630 | 0.277159 | 0.306705 | 0.340562 | 0.358685 | 0.356141 | 0.351113 | 0.350277 | 0.355207 | 0.365202 | 0.345156 | 0.308901 | 0.275071 | 0.252295 | 0.237636 | 0.232776 | 0.232123 | 0.237483 | 0.248743 | 0.269413 | 0.296862 | 0.342453 | 0.378683 | 0.405632 | 0.415928 | 0.447637 | 0.499462 | 0.610032 | 0.747628 | 0.847106 | 0.848352 | 0.750473 | 0.601090 | 0.501958 | 0.454422 | 0.417141 | 0.419145 | 0.436136 | 0.473561 | 0.514978 | 0.552793 | 0.535023 | 0.491145 | 0.402506 | 0.319218 | 0.268910 | 0.244479 | 0.233695 | 0.229682 | 0.236106 | 0.250135 | 0.268630 | 0.288738 | 0.323609 | 0.354725 | 0.390341 | 0.400285 | 0.387769 | 0.355168 | 0.316280 | 0.268525 | 0.238050 | 0.221834 | 0.215354 | 0.215146 | 0.216111 | 0.215378 | 0.214715 | 0.220817 | 0.242745 | 0.282072 | 0.356976 | 0.437191 | 0.486097 | 0.525754 | 0.530037 | 0.473881 | 0.431864 | 0.378656 | 0.345678 | 0.325985 | 0.308897 |
| left central | 0.218558 | 0.230882 | 0.239825 | 0.252566 | 0.260074 | 0.238488 | 0.218979 | 0.215204 | 0.225092 | 0.243574 | 0.264856 | 0.265399 | 0.241149 | 0.221338 | 0.215076 | 0.219000 | 0.225306 | 0.236435 | 0.248532 | 0.255985 | 0.254017 | 0.224709 | 0.214646 | 0.228260 | 0.304336 | 0.461336 | 0.618569 | 0.718947 | 0.565245 | 0.366618 | 0.272501 | 0.236617 | 0.243709 | 0.315531 | 0.590832 | 1.345319 | 3.523359 | 7.834248 | 21.881702 | 41.535980 | 48.951249 | 21.776724 | 8.197877 | 3.065741 | 1.342097 | 0.672826 | 0.448601 | 0.350352 | 0.285837 | 0.259474 | 0.259197 | 0.259029 | 0.264319 | 0.279862 | 0.291893 | 0.317305 | 0.378150 | 0.504115 | 0.720045 | 1.016537 | 1.054368 | 0.683442 | 0.340731 | 0.269569 | 0.286703 | 0.464908 | 1.287719 | 5.791280 | 23.170580 | 71.939117 | 166.689831 | 304.306246 | 301.449199 | 200.845339 | 96.621945 | 34.213784 | 9.107824 | 2.407731 | 0.772441 | 0.403207 | 0.278860 | 0.231226 | 0.221670 | 0.225256 | 0.226873 | 0.241582 | 0.271227 | 0.308944 | 0.371150 | 0.513903 | 0.883569 | 2.294685 | 8.496764 | 34.055908 | 1.020773e+02 | 1.950358e+02 | 2.284292e+02 | 1.376217e+02 | 5.573996e+01 | 18.978209 | 7.612357 | 3.701804 | 1.800609 | 0.916465 | 0.565733 | 0.427203 | 0.353058 | 0.319607 | 0.316035 | 0.318396 | 0.321241 | 0.319569 | 0.328498 | 0.341443 | 0.384128 | 0.440054 | 0.507134 | 0.587064 | 0.745638 | 0.936253 | 1.097841 | 1.117585 | 1.111726 | 1.025989 | 0.889442 | 0.742847 | 0.681427 | 0.670552 | 0.611983 | 0.496993 | 0.384952 | 0.294438 | 0.232980 | 0.215377 | 0.214899 | 0.215093 | 0.226173 | 0.255205 | 0.318802 | 0.428965 | 0.490654 | 0.411500 | 0.301285 | 0.224748 | 0.220253 | 0.260466 | 0.289502 | 0.274193 | 0.241355 | 0.216979 | 0.219012 | 0.249526 | 0.286127 | 0.310632 | 0.308636 | 0.293672 | 0.265427 | 0.248372 | 0.231930 | 0.220928 | 0.217138 | 0.215044 | 0.214768 | 0.216279 | 0.222125 | 0.229886 | 0.235411 | 0.243996 | 0.252722 | 0.253250 | 0.246240 | 0.239376 | 0.228746 | 0.217710 | 0.215118 | 0.220704 | 0.230520 | 0.238088 | 0.243568 | 0.244802 | 0.238546 | 0.231657 | 0.230021 | 0.226599 | 0.222502 | 0.219452 | 0.218093 | 0.216746 | 0.215552 | 0.214829 | 0.215112 | 0.216058 | 0.219250 | 0.229773 | 0.255917 | 0.305054 | 0.362347 | 0.387384 | 0.377757 | 0.340953 | 0.278524 | 0.230408 | 0.214664 | 0.230563 | 0.280655 | 0.342563 | 0.361511 | 0.324220 | 0.271592 | 0.235796 | 0.222319 | 0.220614 | 0.223627 | 0.229303 | 0.240519 | 0.247123 | 0.244528 | 0.238772 | 0.239201 | 0.241910 | 0.252163 | 0.261180 | 0.275833 | 0.287724 | 0.258390 | 0.230596 | 0.220246 | 0.215154 | 0.215648 | 0.221752 | 0.244440 | 0.280064 | 0.330034 | 0.395152 | 0.445205 | 0.423318 | 0.353014 | 0.274261 | 0.233591 | 0.215014 | 0.222360 | 0.243438 | 0.277141 | 0.329542 | 0.422568 | 0.593400 | 0.785710 | 0.989053 | 1.294520 | 1.704917 | 2.012158 | 2.080690 | 1.823624 | 1.313237 | 0.703256 | 0.388443 | 0.275325 | 0.234624 | 0.223100 | 0.222367 | 0.232762 | 0.260720 | 0.311159 | 0.369961 | 0.404779 | 0.420050 | 0.420771 | 0.375994 | 0.331296 | 0.284101 | 0.242802 | 0.220334 | 0.214698 | 0.215638 | 0.216447 | 0.215403 | 0.214713 | 0.218741 | 0.222103 | 0.226954 |
| right central | 0.508316 | 0.552230 | 0.617687 | 0.648460 | 0.673773 | 0.852570 | 1.343168 | 2.000268 | 3.471963 | 4.934034 | 4.094859 | 1.802743 | 0.717546 | 0.292331 | 0.221060 | 0.217590 | 0.245871 | 0.285635 | 0.310444 | 0.330636 | 0.314249 | 0.279728 | 0.243461 | 0.214815 | 0.243372 | 0.421755 | 1.079108 | 3.375807 | 7.096576 | 9.283930 | 3.537189 | 1.027327 | 0.423309 | 0.275654 | 0.230858 | 0.214627 | 0.231164 | 0.283361 | 0.403056 | 0.582043 | 0.656986 | 0.736322 | 0.765896 | 0.846003 | 0.832584 | 0.744278 | 0.526901 | 0.367441 | 0.263742 | 0.226169 | 0.215226 | 0.215194 | 0.221313 | 0.231142 | 0.238674 | 0.238493 | 0.223476 | 0.214821 | 0.231531 | 0.280048 | 0.360997 | 0.439928 | 0.539993 | 0.731848 | 1.184865 | 2.790864 | 9.979353 | 61.081099 | 473.935571 | 3068.153539 | 9722.684170 | 19401.006430 | 17717.646695 | 8237.708616 | 2459.283656 | 577.769380 | 133.043527 | 38.368339 | 11.554907 | 4.485158 | 2.616717 | 1.898077 | 1.732416 | 1.783661 | 2.072359 | 2.636279 | 3.953510 | 6.353130 | 13.075872 | 44.049439 | 188.899273 | 1329.891996 | 14779.031501 | 157464.329225 | 1.028311e+06 | 3.350700e+06 | 5.014438e+06 | 3.998505e+06 | 1.916739e+06 | 473882.397405 | 75501.383307 | 13989.692686 | 2748.953091 | 688.154305 | 235.121856 | 104.756341 | 50.342924 | 26.865888 | 17.386239 | 15.154502 | 14.843638 | 19.480781 | 39.094043 | 100.961053 | 249.153970 | 615.297174 | 1178.529534 | 1916.479554 | 2588.560554 | 3106.521243 | 3392.441502 | 3776.246118 | 2641.706314 | 1296.306711 | 634.047636 | 283.129149 | 116.049830 | 55.179494 | 31.160516 | 17.079601 | 10.169553 | 5.792229 | 3.097246 | 1.879013 | 1.206893 | 0.983349 | 0.973228 | 1.154644 | 1.302695 | 1.409386 | 1.308522 | 1.167263 | 1.100001 | 1.074132 | 0.997701 | 0.878642 | 0.624617 | 0.457656 | 0.365099 | 0.328207 | 0.316928 | 0.344916 | 0.409454 | 0.575873 | 1.029346 | 2.468974 | 6.553179 | 15.498371 | 23.366165 | 23.406823 | 16.402678 | 9.445203 | 3.753929 | 1.539961 | 0.820013 | 0.541016 | 0.431222 | 0.421377 | 0.490038 | 0.630626 | 0.878133 | 1.205665 | 1.634674 | 1.933422 | 2.060684 | 1.993003 | 2.121736 | 2.031191 | 1.901949 | 1.735959 | 1.571124 | 1.333675 | 1.099722 | 0.889950 | 0.731166 | 0.578998 | 0.460730 | 0.398915 | 0.363948 | 0.329358 | 0.329463 | 0.334414 | 0.341366 | 0.359505 | 0.408026 | 0.422535 | 0.443597 | 0.492538 | 0.606347 | 0.637479 | 0.598965 | 0.540495 | 0.483839 | 0.397877 | 0.332897 | 0.287563 | 0.263838 | 0.249181 | 0.242436 | 0.249342 | 0.286457 | 0.384761 | 0.644565 | 1.303405 | 2.468927 | 4.516221 | 5.658611 | 5.144049 | 3.776772 | 2.444361 | 1.356290 | 0.961689 | 0.808442 | 0.880104 | 1.164619 | 1.857104 | 3.008907 | 4.341010 | 4.229839 | 2.734313 | 1.470692 | 0.767879 | 0.468068 | 0.328652 | 0.290459 | 0.291866 | 0.314702 | 0.371196 | 0.445915 | 0.442915 | 0.361210 | 0.264202 | 0.216634 | 0.225736 | 0.262281 | 0.293039 | 0.274192 | 0.240133 | 0.216814 | 0.216049 | 0.220031 | 0.222260 | 0.228524 | 0.236043 | 0.257581 | 0.318623 | 0.417697 | 0.653280 | 1.080293 | 1.610609 | 1.963081 | 2.130483 | 1.693215 | 1.313241 | 0.899983 | 0.542384 | 0.371282 | 0.301453 | 0.265616 | 0.255507 | 0.267792 | 0.280066 | 0.303215 | 0.322029 | 0.351352 | 0.382006 | 0.445207 | 0.521030 | 0.580775 | 0.638662 |
| left posterior | 0.218741 | 0.214732 | 0.216163 | 0.231314 | 0.274141 | 0.356026 | 0.488768 | 0.590976 | 0.494346 | 0.323386 | 0.240976 | 0.217259 | 0.214728 | 0.215801 | 0.225591 | 0.292019 | 0.483590 | 0.845416 | 2.098795 | 4.677281 | 3.757387 | 1.060273 | 0.321479 | 0.215846 | 0.325616 | 0.691404 | 1.138684 | 1.062804 | 0.756675 | 0.515940 | 0.371480 | 0.335661 | 0.394483 | 0.623166 | 1.157865 | 1.881932 | 2.732822 | 3.201907 | 2.962997 | 2.497049 | 2.091571 | 1.724867 | 1.654677 | 1.703760 | 1.382448 | 0.945226 | 0.637394 | 0.421053 | 0.296458 | 0.237060 | 0.215445 | 0.221295 | 0.244731 | 0.277107 | 0.287298 | 0.282039 | 0.282117 | 0.284557 | 0.299370 | 0.332906 | 0.305477 | 0.237660 | 0.218128 | 0.367253 | 1.290171 | 5.892291 | 24.751493 | 82.219536 | 215.580639 | 466.500643 | 847.359499 | 1222.221832 | 1446.210141 | 1559.924838 | 1442.408102 | 1132.327521 | 811.714978 | 689.763028 | 657.581040 | 609.742967 | 415.411916 | 163.436840 | 42.649243 | 10.923850 | 3.657548 | 1.941371 | 1.428643 | 1.108537 | 0.975092 | 1.030267 | 1.298196 | 1.903774 | 3.592793 | 7.700674 | 1.870239e+01 | 4.187138e+01 | 5.928100e+01 | 4.752611e+01 | 3.035901e+01 | 16.704038 | 10.126492 | 6.637967 | 4.545053 | 3.181411 | 2.431852 | 1.923723 | 1.775466 | 1.841766 | 2.195508 | 2.766780 | 3.632213 | 4.262225 | 4.745348 | 4.836247 | 4.972671 | 5.105896 | 6.048307 | 7.296619 | 9.781872 | 13.422521 | 19.527662 | 28.902355 | 45.446865 | 68.794512 | 96.721290 | 103.561006 | 81.754867 | 46.773826 | 22.388877 | 9.559805 | 4.693103 | 2.666897 | 1.854028 | 1.502522 | 1.439312 | 1.583369 | 2.019441 | 2.780388 | 3.982157 | 6.005311 | 8.611375 | 11.798653 | 15.462297 | 19.214058 | 19.236260 | 13.913033 | 7.085063 | 3.192078 | 1.620784 | 0.992821 | 0.738488 | 0.636059 | 0.626996 | 0.691086 | 0.841917 | 1.109830 | 1.620752 | 2.516618 | 3.373416 | 3.966888 | 4.539477 | 4.939761 | 4.949222 | 4.554372 | 3.973020 | 3.402169 | 3.156677 | 3.236591 | 3.834859 | 4.581544 | 4.977103 | 4.913544 | 4.511386 | 4.245345 | 4.309355 | 4.025490 | 3.598379 | 3.493496 | 3.222526 | 3.018049 | 3.049126 | 3.070378 | 2.822625 | 2.750578 | 2.405919 | 2.221152 | 2.043205 | 1.892594 | 1.657312 | 1.525910 | 1.429492 | 1.369086 | 1.432704 | 1.666536 | 2.102355 | 2.566953 | 3.541764 | 4.831922 | 6.042853 | 5.950010 | 5.056628 | 3.394468 | 2.282253 | 1.514098 | 1.027555 | 0.716702 | 0.529339 | 0.426841 | 0.383741 | 0.379017 | 0.418275 | 0.526379 | 0.774337 | 1.257940 | 1.954125 | 2.570482 | 2.630723 | 2.051372 | 1.243995 | 0.759128 | 0.478724 | 0.331501 | 0.265377 | 0.241728 | 0.232688 | 0.236073 | 0.249437 | 0.266769 | 0.289496 | 0.322043 | 0.373247 | 0.441230 | 0.544886 | 0.719329 | 0.999204 | 1.284423 | 1.478573 | 1.614121 | 1.623379 | 1.427381 | 1.182747 | 0.997285 | 0.869568 | 0.818336 | 0.845632 | 1.044906 | 1.668212 | 2.973597 | 4.149799 | 4.356951 | 3.256345 | 2.020870 | 1.305032 | 0.888615 | 0.648051 | 0.562952 | 0.525292 | 0.520220 | 0.567889 | 0.632106 | 0.613220 | 0.529877 | 0.425616 | 0.327439 | 0.258075 | 0.224265 | 0.214629 | 0.222800 | 0.246015 | 0.279893 | 0.334401 | 0.386513 | 0.435664 | 0.428253 | 0.391533 | 0.340987 | 0.302406 | 0.275469 | 0.264375 | 0.250356 |
| right posterior | 1.000577 | 0.623676 | 0.410528 | 0.233641 | 0.223901 | 0.284551 | 0.368450 | 0.389611 | 0.303439 | 0.238938 | 0.219382 | 0.215476 | 0.214636 | 0.215271 | 0.214862 | 0.223474 | 0.251826 | 0.288683 | 0.304081 | 0.305813 | 0.263309 | 0.216157 | 0.226961 | 0.248511 | 0.273648 | 0.256768 | 0.228270 | 0.214627 | 0.232480 | 0.304930 | 0.374382 | 0.379467 | 0.301238 | 0.225521 | 0.235393 | 0.459390 | 1.364788 | 2.343945 | 1.353725 | 0.688852 | 0.484081 | 0.450081 | 0.510233 | 0.640771 | 0.728493 | 0.736606 | 0.580471 | 0.383975 | 0.240327 | 0.215997 | 0.240937 | 0.247084 | 0.231458 | 0.214630 | 0.258111 | 0.442150 | 0.750347 | 1.164380 | 1.456679 | 1.320490 | 1.241991 | 1.268707 | 1.511266 | 2.344554 | 5.119888 | 14.091245 | 43.224775 | 138.731918 | 516.517750 | 1858.863246 | 5088.690222 | 11910.340890 | 15816.849649 | 13241.359495 | 7225.388172 | 4019.047855 | 1880.288885 | 744.538644 | 257.139013 | 71.474255 | 18.204775 | 5.971655 | 2.866492 | 1.819763 | 1.571619 | 1.675753 | 1.900269 | 2.211076 | 2.758494 | 3.608885 | 4.434355 | 5.467653 | 6.685800 | 7.791291 | 8.569078e+00 | 9.188434e+00 | 9.163238e+00 | 8.633205e+00 | 7.740024e+00 | 6.414504 | 5.284009 | 4.430988 | 3.973956 | 3.657026 | 3.618226 | 3.630169 | 3.645549 | 3.526481 | 3.476474 | 3.370107 | 3.426600 | 3.710933 | 4.590995 | 6.586795 | 10.804822 | 18.662384 | 35.649102 | 77.708414 | 170.572479 | 309.988167 | 469.822874 | 612.635159 | 621.976659 | 468.793277 | 278.924430 | 150.370457 | 76.318962 | 42.266058 | 27.879368 | 20.201552 | 15.672361 | 13.990831 | 14.157004 | 15.207208 | 17.869256 | 21.656019 | 23.215548 | 23.741420 | 23.435076 | 22.000915 | 20.032932 | 19.236871 | 18.339605 | 20.180544 | 26.338707 | 38.703512 | 52.838803 | 63.961548 | 67.444668 | 60.640191 | 46.416419 | 36.049473 | 29.850842 | 27.528651 | 31.356106 | 43.918660 | 70.032150 | 125.767660 | 193.330599 | 219.172806 | 167.168185 | 96.599193 | 45.187585 | 20.829505 | 11.022432 | 7.444526 | 6.022773 | 6.322096 | 7.404285 | 9.615743 | 13.595981 | 19.837276 | 25.904105 | 40.052143 | 64.763365 | 102.372262 | 166.032245 | 240.458722 | 269.303598 | 225.811599 | 149.219181 | 79.384379 | 45.779913 | 26.591424 | 19.882646 | 20.698156 | 30.002333 | 46.701780 | 79.277000 | 135.752978 | 209.175474 | 277.526991 | 369.319816 | 469.933283 | 468.354443 | 405.934889 | 299.457524 | 223.807407 | 175.140420 | 165.751682 | 194.452027 | 264.933016 | 327.415347 | 348.571452 | 305.603633 | 219.123572 | 143.338448 | 100.703642 | 79.125790 | 65.725510 | 68.530862 | 82.580429 | 116.640509 | 184.626685 | 293.785033 | 453.503547 | 881.523296 | 1210.812653 | 1280.755252 | 1026.371227 | 489.540517 | 139.353379 | 45.549646 | 19.298592 | 11.708502 | 9.868572 | 11.035166 | 15.544872 | 26.136890 | 42.329209 | 55.930941 | 67.694342 | 73.469381 | 79.881428 | 79.487046 | 72.282624 | 64.298707 | 61.498173 | 52.599894 | 49.180829 | 46.653095 | 44.837221 | 47.382876 | 50.568869 | 42.477588 | 34.784507 | 28.709711 | 26.255791 | 28.570475 | 31.067519 | 27.894560 | 21.718597 | 15.742428 | 11.289246 | 8.848164 | 8.202222 | 9.008845 | 11.111954 | 14.495229 | 19.856348 | 26.016625 | 34.032052 | 45.752896 | 60.214034 | 75.224366 | 73.246276 | 51.487850 | 26.364679 | 11.695615 | 4.997810 | 2.455692 | 1.542222 | 1.235778 | 1.248931 | 1.637385 | 2.585193 | 4.726644 | 9.018839 | 12.559712 | 16.696864 |
| all electrodes | 0.231619 | 0.238963 | 0.239659 | 0.241145 | 0.235041 | 0.235641 | 0.234218 | 0.241811 | 0.253282 | 0.266872 | 0.264038 | 0.304240 | 0.364046 | 0.501707 | 0.785995 | 1.398740 | 2.103410 | 3.325408 | 5.061906 | 6.521746 | 4.313967 | 1.278301 | 0.396035 | 0.231295 | 0.219721 | 0.221058 | 0.222444 | 0.346365 | 0.794308 | 1.614531 | 1.769175 | 1.563422 | 0.873046 | 0.395957 | 0.229202 | 0.222456 | 0.327361 | 0.476153 | 0.417274 | 0.306289 | 0.242748 | 0.214630 | 0.228477 | 0.252349 | 0.260588 | 0.261615 | 0.257778 | 0.270462 | 0.339529 | 0.452791 | 0.545471 | 0.596089 | 0.615271 | 0.462462 | 0.345528 | 0.261303 | 0.225032 | 0.217312 | 0.218570 | 0.225039 | 0.237497 | 0.225142 | 0.218926 | 0.304739 | 0.780808 | 3.882130 | 25.859756 | 150.332651 | 657.276013 | 1780.300780 | 4606.100690 | 8775.644049 | 15283.310202 | 23077.660287 | 29233.632859 | 19402.285796 | 7381.027283 | 2194.614706 | 745.375620 | 238.384156 | 72.648592 | 30.068709 | 14.568127 | 9.027374 | 6.478007 | 5.125452 | 5.025665 | 7.222179 | 10.901120 | 28.566431 | 131.546624 | 512.251149 | 1533.162325 | 4681.236984 | 6.943213e+03 | 6.866914e+03 | 5.109504e+03 | 1.756447e+03 | 6.556570e+02 | 312.401292 | 165.222747 | 90.888963 | 45.757950 | 18.906510 | 9.530877 | 5.184713 | 3.175429 | 2.706018 | 3.241597 | 5.029481 | 9.585951 | 21.279279 | 46.703601 | 109.655668 | 214.882515 | 285.396631 | 287.851069 | 285.491092 | 282.723981 | 253.066135 | 343.930259 | 584.470648 | 975.830931 | 1374.486104 | 1738.917343 | 1583.628982 | 1572.536908 | 1101.163016 | 684.976504 | 345.821409 | 156.236162 | 78.302455 | 51.765454 | 33.551319 | 30.483006 | 39.059175 | 64.176587 | 129.578804 | 312.940470 | 833.988735 | 1662.981228 | 1436.356338 | 621.478863 | 317.101107 | 171.377207 | 103.648766 | 77.243638 | 64.663301 | 65.052053 | 80.572209 | 84.904689 | 79.537789 | 76.077374 | 60.934594 | 47.812702 | 34.135922 | 23.144439 | 13.805359 | 6.546487 | 2.964686 | 1.535898 | 0.944389 | 0.659203 | 0.552111 | 0.448013 | 0.365564 | 0.320570 | 0.291056 | 0.266604 | 0.256980 | 0.250678 | 0.241091 | 0.243129 | 0.245323 | 0.252978 | 0.278544 | 0.337637 | 0.402909 | 0.515750 | 0.659842 | 0.773644 | 0.789169 | 0.770108 | 0.734378 | 0.692407 | 0.754080 | 0.787119 | 0.776566 | 0.739938 | 0.727170 | 0.673766 | 0.716091 | 0.794203 | 0.968113 | 1.121116 | 1.212291 | 1.189458 | 1.059000 | 0.794832 | 0.546498 | 0.392778 | 0.327334 | 0.301944 | 0.290691 | 0.295183 | 0.299370 | 0.318564 | 0.347216 | 0.372387 | 0.404642 | 0.454830 | 0.455260 | 0.427949 | 0.387110 | 0.362810 | 0.360191 | 0.400716 | 0.489441 | 0.649882 | 0.831163 | 0.916755 | 0.734188 | 0.545442 | 0.403579 | 0.312932 | 0.254972 | 0.234211 | 0.222870 | 0.220222 | 0.218253 | 0.216228 | 0.215070 | 0.218180 | 0.222718 | 0.222343 | 0.219477 | 0.215617 | 0.215339 | 0.227015 | 0.252341 | 0.277318 | 0.290371 | 0.278538 | 0.244424 | 0.218767 | 0.214879 | 0.222193 | 0.244180 | 0.273480 | 0.283973 | 0.308421 | 0.320107 | 0.329835 | 0.364876 | 0.433643 | 0.531319 | 0.651225 | 0.646748 | 0.504283 | 0.386958 | 0.282739 | 0.229837 | 0.216126 | 0.216903 | 0.238719 | 0.361444 | 0.845212 | 3.300568 | 9.999113 | 13.236905 | 7.901300 | 3.377063 | 1.628375 | 1.126160 | 0.863300 | 0.749508 | 0.682401 | 0.589152 | 0.510904 | 0.467094 |

F) neutral vs sad

  
|  | time window | peak latency | cluster *p* | peak Cohen's *d* |  | | | |  |
| **all electrodes** | 125 - 185 ms | 155 ms | 0.047 | 1.2823 |  | | | |  |
 935 - 1080 ms | 970 ms | 0.0232 | 0.5591 |  | | | ||  |  | | | | | | | | |

Model correlations, cluster permutation tests

|  | **left hemisphere** | | | | **right hemisphere** | | | |  |
|  | time window | peak latency | cluster *p* | peak Cohen's *d* | time window | peak latency | cluster *p* | peak Cohen's *d* |  |
| **anterior** |  | | | |  | | | |  |
| **central** | 135 - 245 ms | 160 ms | 0.0466 | 0.6459 | -200 - -135 ms | -195 ms | 0.0428 | -0.7566 |  |
| **posterior** |  | | | |  | | | |  |

  

Model correlations, Bayesian statistics

|  | -200 | -195 | -190 | -185 | -180 | -175 | -170 | -165 | -160 | -155 | -150 | -145 | -140 | -135 | -130 | -125 | -120 | -115 | -110 | -105 | -100 | -95 | -90 | -85 | -80 | -75 | -70 | -65 | -60 | -55 | -50 | -45 | -40 | -35 | -30 | -25 | -20 | -15 | -10 | -5 | 0 | 5 | 10 | 15 | 20 | 25 | 30 | 35 | 40 | 45 | 50 | 55 | 60 | 65 | 70 | 75 | 80 | 85 | 90 | 95 | 100 | 105 | 110 | 115 | 120 | 125 | 130 | 135 | 140 | 145 | 150 | 155 | 160 | 165 | 170 | 175 | 180 | 185 | 190 | 195 | 200 | 205 | 210 | 215 | 220 | 225 | 230 | 235 | 240 | 245 | 250 | 255 | 260 | 265 | 270 | 275 | 280 | 285 | 290 | 295 | 300 | 305 | 310 | 315 | 320 | 325 | 330 | 335 | 340 | 345 | 350 | 355 | 360 | 365 | 370 | 375 | 380 | 385 | 390 | 395 | 400 | 405 | 410 | 415 | 420 | 425 | 430 | 435 | 440 | 445 | 450 | 455 | 460 | 465 | 470 | 475 | 480 | 485 | 490 | 495 | 500 | 505 | 510 | 515 | 520 | 525 | 530 | 535 | 540 | 545 | 550 | 555 | 560 | 565 | 570 | 575 | 580 | 585 | 590 | 595 | 600 | 605 | 610 | 615 | 620 | 625 | 630 | 635 | 640 | 645 | 650 | 655 | 660 | 665 | 670 | 675 | 680 | 685 | 690 | 695 | 700 | 705 | 710 | 715 | 720 | 725 | 730 | 735 | 740 | 745 | 750 | 755 | 760 | 765 | 770 | 775 | 780 | 785 | 790 | 795 | 800 | 805 | 810 | 815 | 820 | 825 | 830 | 835 | 840 | 845 | 850 | 855 | 860 | 865 | 870 | 875 | 880 | 885 | 890 | 895 | 900 | 905 | 910 | 915 | 920 | 925 | 930 | 935 | 940 | 945 | 950 | 955 | 960 | 965 | 970 | 975 | 980 | 985 | 990 | 995 | 1000 | 1005 | 1010 | 1015 | 1020 | 1025 | 1030 | 1035 | 1040 | 1045 | 1050 | 1055 | 1060 | 1065 | 1070 | 1075 | 1080 | 1085 | 1090 | 1095 | 1100 | 1105 | 1110 | 1115 | 1120 | 1125 | 1130 | 1135 | 1140 | 1145 | 1150 | 1155 | 1160 | 1165 | 1170 | 1175 | 1180 | 1185 | 1190 | 1195 |
| --- | --- | --- | --- | --- | --- | --- | --- | --- | --- | --- | --- | --- | --- | --- | --- | --- | --- | --- | --- | --- | --- | --- | --- | --- | --- | --- | --- | --- | --- | --- | --- | --- | --- | --- | --- | --- | --- | --- | --- | --- | --- | --- | --- | --- | --- | --- | --- | --- | --- | --- | --- | --- | --- | --- | --- | --- | --- | --- | --- | --- | --- | --- | --- | --- | --- | --- | --- | --- | --- | --- | --- | --- | --- | --- | --- | --- | --- | --- | --- | --- | --- | --- | --- | --- | --- | --- | --- | --- | --- | --- | --- | --- | --- | --- | --- | --- | --- | --- | --- | --- | --- | --- | --- | --- | --- | --- | --- | --- | --- | --- | --- | --- | --- | --- | --- | --- | --- | --- | --- | --- | --- | --- | --- | --- | --- | --- | --- | --- | --- | --- | --- | --- | --- | --- | --- | --- | --- | --- | --- | --- | --- | --- | --- | --- | --- | --- | --- | --- | --- | --- | --- | --- | --- | --- | --- | --- | --- | --- | --- | --- | --- | --- | --- | --- | --- | --- | --- | --- | --- | --- | --- | --- | --- | --- | --- | --- | --- | --- | --- | --- | --- | --- | --- | --- | --- | --- | --- | --- | --- | --- | --- | --- | --- | --- | --- | --- | --- | --- | --- | --- | --- | --- | --- | --- | --- | --- | --- | --- | --- | --- | --- | --- | --- | --- | --- | --- | --- | --- | --- | --- | --- | --- | --- | --- | --- | --- | --- | --- | --- | --- | --- | --- | --- | --- | --- | --- | --- | --- | --- | --- | --- | --- | --- | --- | --- | --- | --- | --- | --- | --- | --- | --- | --- | --- | --- | --- | --- | --- | --- | --- | --- | --- | --- | --- | --- | --- | --- | --- | --- | --- | --- | --- | --- | --- | --- | --- | --- | --- | --- | --- |
| left anterior | 0.527579 | 0.469429 | 0.454648 | 0.349553 | 0.262967 | 0.233885 | 0.217677 | 0.214643 | 0.232167 | 0.245683 | 0.246014 | 0.232342 | 0.228542 | 0.216028 | 0.216379 | 0.239540 | 0.256719 | 0.251891 | 0.241733 | 0.239760 | 0.234107 | 0.234634 | 0.251779 | 0.293318 | 0.366407 | 0.406733 | 0.450199 | 0.448265 | 0.394954 | 0.279206 | 0.227225 | 0.216326 | 0.255465 | 0.434894 | 0.873216 | 1.334575 | 1.319020 | 1.491415 | 1.481519 | 1.643040 | 2.138715 | 4.534657 | 12.062185 | 36.765637 | 101.960531 | 131.406449 | 107.872723 | 56.552454 | 28.622397 | 13.250211 | 6.412206 | 2.035444 | 1.018511 | 0.578511 | 0.363896 | 0.267029 | 0.234327 | 0.218748 | 0.215687 | 0.215994 | 0.216546 | 0.215338 | 0.214644 | 0.220247 | 0.250254 | 0.323252 | 0.483301 | 0.746503 | 1.084639 | 1.418789 | 1.715465 | 1.654587 | 1.531239 | 1.269833 | 0.945340 | 0.583661 | 0.386131 | 0.274645 | 0.225660 | 0.214677 | 0.221732 | 0.224805 | 0.220385 | 0.215587 | 0.214716 | 0.219355 | 0.224706 | 0.225521 | 0.224600 | 0.226829 | 0.232612 | 0.247632 | 0.271866 | 0.312588 | 0.352461 | 0.359751 | 0.350761 | 0.348245 | 0.319536 | 0.284858 | 0.253918 | 0.234415 | 0.232295 | 0.238881 | 0.249297 | 0.272319 | 0.308001 | 0.325760 | 0.341768 | 0.335384 | 0.322249 | 0.296254 | 0.270786 | 0.251196 | 0.241899 | 0.233746 | 0.233217 | 0.231815 | 0.228869 | 0.225007 | 0.221103 | 0.217809 | 0.216631 | 0.215191 | 0.214653 | 0.214651 | 0.215874 | 0.225039 | 0.254197 | 0.329099 | 0.479844 | 0.693066 | 0.783322 | 0.663285 | 0.457983 | 0.354682 | 0.296886 | 0.263541 | 0.243361 | 0.233548 | 0.227491 | 0.222725 | 0.218950 | 0.215879 | 0.215003 | 0.214894 | 0.215112 | 0.215390 | 0.216137 | 0.215339 | 0.215912 | 0.228220 | 0.253024 | 0.296076 | 0.353988 | 0.430846 | 0.539447 | 0.630653 | 0.701534 | 0.801874 | 0.833654 | 0.873022 | 0.896915 | 0.889192 | 0.881654 | 0.914542 | 0.880152 | 0.907900 | 0.922715 | 0.881340 | 0.770977 | 0.638925 | 0.492835 | 0.392789 | 0.331939 | 0.307510 | 0.310112 | 0.344378 | 0.411874 | 0.531197 | 0.712805 | 0.867232 | 0.919236 | 0.799917 | 0.619096 | 0.440296 | 0.330679 | 0.263837 | 0.234434 | 0.221438 | 0.215350 | 0.214880 | 0.215657 | 0.215047 | 0.214644 | 0.219037 | 0.244009 | 0.313476 | 0.442644 | 0.652747 | 0.914860 | 1.024489 | 0.775712 | 0.453435 | 0.282051 | 0.230317 | 0.219507 | 0.223070 | 0.250683 | 0.333608 | 0.482190 | 0.692945 | 0.885147 | 1.012152 | 1.068140 | 1.088037 | 1.010107 | 0.960430 | 0.926347 | 0.942863 | 0.913687 | 0.869562 | 0.822342 | 0.769055 | 0.725210 | 0.720308 | 0.792691 | 0.898641 | 1.023364 | 0.945485 | 0.751281 | 0.555525 | 0.447079 | 0.370144 | 0.353003 | 0.375498 | 0.453297 | 0.514545 | 0.565986 | 0.581229 | 0.561813 | 0.489957 | 0.415647 | 0.337442 | 0.288813 | 0.250040 | 0.225574 | 0.218103 | 0.216393 | 0.216330 | 0.218431 | 0.220138 | 0.219637 | 0.217149 | 0.214742 | 0.216934 | 0.225629 | 0.240970 | 0.252373 | 0.257656 | 0.257321 | 0.247214 | 0.231553 | 0.221715 | 0.217207 | 0.215396 | 0.214942 | 0.214628 | 0.214890 | 0.216254 | 0.219814 | 0.224908 | 0.228177 | 0.230796 | 0.230996 | 0.230343 | 0.235424 | 0.250229 | 0.255375 | 0.268152 |
| right anterior | 0.229295 | 0.223120 | 0.215584 | 0.216467 | 0.218331 | 0.217084 | 0.214803 | 0.218999 | 0.252769 | 0.283767 | 0.270830 | 0.245078 | 0.231031 | 0.215423 | 0.223264 | 0.271458 | 0.387794 | 0.471305 | 0.447044 | 0.369600 | 0.283798 | 0.229715 | 0.214704 | 0.234205 | 0.287505 | 0.359496 | 0.395574 | 0.336665 | 0.266536 | 0.222256 | 0.217297 | 0.256808 | 0.346975 | 0.426575 | 0.366468 | 0.275636 | 0.228298 | 0.217954 | 0.214879 | 0.214626 | 0.215383 | 0.217166 | 0.217068 | 0.218082 | 0.217478 | 0.216489 | 0.217386 | 0.215716 | 0.215082 | 0.226185 | 0.269844 | 0.374636 | 0.567739 | 0.928192 | 1.499009 | 2.265143 | 3.565919 | 3.972786 | 3.340005 | 1.907346 | 0.958463 | 0.562221 | 0.455605 | 0.480157 | 0.725440 | 1.518068 | 3.652158 | 6.666305 | 9.586143 | 11.643865 | 12.140772 | 10.211648 | 7.093003 | 4.510038 | 3.132774 | 2.478949 | 2.107008 | 1.905635 | 1.684115 | 1.277558 | 0.882484 | 0.683440 | 0.625921 | 0.675168 | 0.842567 | 1.126399 | 1.538062 | 2.042850 | 2.347111 | 2.814854 | 3.265904 | 3.297845 | 3.415200 | 3.825479 | 4.172744 | 4.491054 | 5.225372 | 5.832003 | 5.873180 | 3.884749 | 2.363253 | 1.475245 | 0.966906 | 0.662144 | 0.603919 | 0.650250 | 0.817967 | 1.047629 | 1.302794 | 1.368983 | 1.342166 | 1.164981 | 0.961520 | 0.787617 | 0.653237 | 0.477531 | 0.352707 | 0.271378 | 0.231781 | 0.216909 | 0.214712 | 0.217625 | 0.222892 | 0.235398 | 0.255740 | 0.295198 | 0.367741 | 0.471011 | 0.542539 | 0.537047 | 0.480172 | 0.423864 | 0.356300 | 0.300474 | 0.269545 | 0.261485 | 0.267663 | 0.291256 | 0.342235 | 0.448668 | 0.624757 | 0.867710 | 1.208582 | 1.489344 | 1.463568 | 1.276579 | 1.063228 | 0.726698 | 0.495685 | 0.345772 | 0.267121 | 0.231015 | 0.216931 | 0.215289 | 0.222417 | 0.234963 | 0.240845 | 0.233453 | 0.221034 | 0.215167 | 0.236116 | 0.275309 | 0.331353 | 0.370033 | 0.360167 | 0.346071 | 0.321211 | 0.290854 | 0.277822 | 0.274886 | 0.279070 | 0.289687 | 0.291033 | 0.285632 | 0.281041 | 0.268776 | 0.255399 | 0.244215 | 0.242040 | 0.253100 | 0.270392 | 0.299912 | 0.348925 | 0.403392 | 0.457604 | 0.490640 | 0.521738 | 0.593001 | 0.662791 | 0.662691 | 0.672282 | 0.633171 | 0.553962 | 0.497633 | 0.425163 | 0.339256 | 0.292871 | 0.266027 | 0.256654 | 0.269819 | 0.310397 | 0.356008 | 0.410511 | 0.421780 | 0.396178 | 0.330154 | 0.276996 | 0.239136 | 0.220136 | 0.214678 | 0.218395 | 0.224522 | 0.234020 | 0.250324 | 0.269467 | 0.277250 | 0.285283 | 0.292181 | 0.291955 | 0.277319 | 0.279703 | 0.279241 | 0.276003 | 0.269647 | 0.260936 | 0.241703 | 0.226575 | 0.215777 | 0.215683 | 0.215288 | 0.214828 | 0.218420 | 0.228109 | 0.238121 | 0.225167 | 0.216447 | 0.215635 | 0.224921 | 0.240987 | 0.234618 | 0.220734 | 0.215861 | 0.244389 | 0.298376 | 0.327317 | 0.300137 | 0.253629 | 0.224352 | 0.214884 | 0.217925 | 0.222282 | 0.221256 | 0.216456 | 0.214671 | 0.216301 | 0.221981 | 0.231125 | 0.234867 | 0.244422 | 0.248264 | 0.248854 | 0.249213 | 0.255770 | 0.256299 | 0.255565 | 0.254976 | 0.252419 | 0.243116 | 0.230742 | 0.221452 | 0.217103 | 0.214656 | 0.216519 | 0.225966 | 0.241744 | 0.263249 | 0.278987 | 0.288327 | 0.287514 | 0.274185 |
| left central | 0.977165 | 0.963285 | 1.179758 | 1.311170 | 1.322662 | 0.996551 | 0.727222 | 0.435056 | 0.302610 | 0.226941 | 0.214995 | 0.235655 | 0.284040 | 0.330224 | 0.297370 | 0.251269 | 0.222889 | 0.214905 | 0.244570 | 0.291236 | 0.318485 | 0.289310 | 0.277657 | 0.258242 | 0.254507 | 0.237496 | 0.231773 | 0.226198 | 0.218957 | 0.215013 | 0.221134 | 0.234589 | 0.244943 | 0.257686 | 0.247215 | 0.235468 | 0.229263 | 0.227491 | 0.223235 | 0.225610 | 0.236211 | 0.259055 | 0.289249 | 0.341503 | 0.395179 | 0.426391 | 0.384036 | 0.319627 | 0.259706 | 0.223904 | 0.214638 | 0.221373 | 0.234306 | 0.258792 | 0.281926 | 0.319642 | 0.358652 | 0.384616 | 0.396617 | 0.406757 | 0.400664 | 0.398153 | 0.414828 | 0.432652 | 0.475212 | 0.589819 | 0.818598 | 1.319522 | 2.279500 | 3.633191 | 5.886314 | 10.076858 | 11.618534 | 10.274249 | 7.454793 | 4.480389 | 2.688984 | 1.851503 | 1.446087 | 1.457068 | 1.772651 | 2.096327 | 2.567365 | 3.475091 | 4.023805 | 3.959550 | 3.003600 | 2.107936 | 1.621733 | 1.187076 | 0.960810 | 0.821439 | 0.693086 | 0.598014 | 0.541366 | 0.476675 | 0.435840 | 0.385162 | 0.329472 | 0.290202 | 0.270904 | 0.272426 | 0.293563 | 0.326587 | 0.346814 | 0.371170 | 0.370416 | 0.330665 | 0.258471 | 0.223936 | 0.215249 | 0.227897 | 0.246522 | 0.248322 | 0.240232 | 0.225946 | 0.222100 | 0.220492 | 0.217330 | 0.214855 | 0.214738 | 0.215846 | 0.219966 | 0.230049 | 0.245195 | 0.255903 | 0.256077 | 0.242075 | 0.222521 | 0.215558 | 0.215924 | 0.221943 | 0.226405 | 0.225464 | 0.226153 | 0.227602 | 0.240856 | 0.249146 | 0.250103 | 0.256091 | 0.259099 | 0.246275 | 0.239130 | 0.230397 | 0.223445 | 0.219314 | 0.217473 | 0.214638 | 0.219102 | 0.236727 | 0.259990 | 0.281257 | 0.280781 | 0.269751 | 0.264396 | 0.259454 | 0.253791 | 0.234352 | 0.219099 | 0.215307 | 0.236824 | 0.315715 | 0.470446 | 0.740993 | 0.999746 | 1.161940 | 1.192283 | 1.039629 | 0.788338 | 0.596336 | 0.429868 | 0.287527 | 0.222276 | 0.228311 | 0.338289 | 0.562647 | 0.716851 | 0.813007 | 0.735903 | 0.723545 | 0.672945 | 0.547438 | 0.443571 | 0.399584 | 0.329775 | 0.307686 | 0.295879 | 0.281417 | 0.277054 | 0.294697 | 0.312931 | 0.388925 | 0.583165 | 0.935286 | 1.431663 | 1.755151 | 1.453423 | 0.984140 | 0.663575 | 0.468584 | 0.394857 | 0.384061 | 0.418832 | 0.521196 | 0.768712 | 1.191308 | 1.769538 | 2.117239 | 1.868874 | 1.360321 | 0.942354 | 0.719945 | 0.615734 | 0.544605 | 0.550899 | 0.610952 | 0.624501 | 0.661343 | 0.801018 | 0.814710 | 0.632517 | 0.415339 | 0.277226 | 0.228129 | 0.215448 | 0.216789 | 0.220299 | 0.217972 | 0.215849 | 0.214634 | 0.214877 | 0.214627 | 0.215259 | 0.220723 | 0.238304 | 0.238175 | 0.227191 | 0.220369 | 0.218137 | 0.216431 | 0.215825 | 0.215826 | 0.217048 | 0.218430 | 0.221678 | 0.231637 | 0.253168 | 0.288758 | 0.320061 | 0.329402 | 0.294181 | 0.245106 | 0.216431 | 0.224221 | 0.267529 | 0.325853 | 0.376512 | 0.401677 | 0.405420 | 0.410766 | 0.400761 | 0.390569 | 0.378551 | 0.393958 | 0.422481 | 0.459706 | 0.470529 | 0.482761 | 0.428376 | 0.353475 | 0.288249 | 0.251520 | 0.232501 | 0.230875 | 0.229316 | 0.232628 | 0.241925 | 0.254057 | 0.253048 | 0.259390 |
| right central | 18.094848 | 22.836997 | 30.734896 | 52.069043 | 87.680648 | 110.901787 | 118.357353 | 221.063396 | 937.900962 | 1069.297526 | 385.606766 | 53.261482 | 7.992221 | 1.587190 | 0.539697 | 0.314735 | 0.251360 | 0.218845 | 0.215097 | 0.224550 | 0.238935 | 0.247013 | 0.239013 | 0.224807 | 0.216834 | 0.257536 | 0.392993 | 0.721490 | 1.074226 | 0.678607 | 0.309177 | 0.214628 | 0.251720 | 0.285056 | 0.271497 | 0.262961 | 0.238199 | 0.221442 | 0.214630 | 0.221281 | 0.245157 | 0.255815 | 0.259179 | 0.245154 | 0.225242 | 0.214853 | 0.219429 | 0.251211 | 0.302305 | 0.367327 | 0.434670 | 0.456689 | 0.397852 | 0.320546 | 0.246296 | 0.215039 | 0.220722 | 0.228604 | 0.228883 | 0.217912 | 0.214700 | 0.217094 | 0.220838 | 0.221821 | 0.222704 | 0.245135 | 0.318624 | 0.552769 | 1.472322 | 4.368886 | 9.012013 | 13.899220 | 15.205912 | 10.606088 | 6.080390 | 3.044250 | 1.524641 | 0.844986 | 0.513543 | 0.328918 | 0.265405 | 0.235973 | 0.234952 | 0.247686 | 0.269422 | 0.281269 | 0.296992 | 0.292925 | 0.294179 | 0.290417 | 0.287414 | 0.283087 | 0.283555 | 0.276297 | 0.278191 | 0.285252 | 0.289594 | 0.289642 | 0.300547 | 0.320730 | 0.367814 | 0.453002 | 0.494349 | 0.456902 | 0.337034 | 0.247138 | 0.214631 | 0.237079 | 0.301820 | 0.330565 | 0.324690 | 0.293268 | 0.272316 | 0.250463 | 0.242265 | 0.242111 | 0.247605 | 0.243364 | 0.231454 | 0.219695 | 0.214754 | 0.224579 | 0.239748 | 0.247563 | 0.253792 | 0.250099 | 0.233215 | 0.220505 | 0.216702 | 0.216426 | 0.216943 | 0.218756 | 0.220645 | 0.221496 | 0.219648 | 0.217083 | 0.215237 | 0.214828 | 0.214759 | 0.217620 | 0.224766 | 0.238270 | 0.268363 | 0.311469 | 0.374499 | 0.463451 | 0.523571 | 0.515401 | 0.440844 | 0.356632 | 0.288134 | 0.252530 | 0.241369 | 0.239777 | 0.250047 | 0.277570 | 0.300084 | 0.299053 | 0.277818 | 0.242128 | 0.219527 | 0.214722 | 0.219544 | 0.222180 | 0.224484 | 0.227732 | 0.230259 | 0.228176 | 0.225960 | 0.217614 | 0.216494 | 0.254282 | 0.353299 | 0.467840 | 0.550623 | 0.560907 | 0.468028 | 0.392691 | 0.364337 | 0.342627 | 0.335102 | 0.337800 | 0.336712 | 0.336046 | 0.322941 | 0.293088 | 0.264728 | 0.235206 | 0.216249 | 0.219236 | 0.239222 | 0.279426 | 0.317405 | 0.318661 | 0.289603 | 0.257617 | 0.229845 | 0.216551 | 0.215753 | 0.223457 | 0.229562 | 0.238934 | 0.256553 | 0.266737 | 0.259662 | 0.262854 | 0.261455 | 0.269594 | 0.286610 | 0.305728 | 0.301912 | 0.293869 | 0.264616 | 0.242877 | 0.235116 | 0.228900 | 0.225165 | 0.229653 | 0.235919 | 0.241850 | 0.249427 | 0.249643 | 0.236154 | 0.219240 | 0.215486 | 0.227859 | 0.244831 | 0.259054 | 0.265730 | 0.265165 | 0.259500 | 0.254463 | 0.249706 | 0.240436 | 0.225349 | 0.215740 | 0.216046 | 0.230007 | 0.268080 | 0.321341 | 0.376799 | 0.425318 | 0.465604 | 0.498076 | 0.535369 | 0.560642 | 0.633664 | 0.698587 | 0.780881 | 0.714703 | 0.715634 | 0.767256 | 0.839907 | 0.781252 | 0.635305 | 0.427856 | 0.289187 | 0.220869 | 0.227254 | 0.278534 | 0.330655 | 0.369183 | 0.318225 | 0.253435 | 0.217950 | 0.221357 | 0.258637 | 0.292571 | 0.326123 | 0.346059 | 0.365913 | 0.388172 | 0.421207 | 0.470067 | 0.535216 | 0.607226 | 0.657296 | 0.696503 | 0.721396 | 0.699876 |
| left posterior | 0.871302 | 1.093416 | 1.415745 | 2.329371 | 2.095102 | 1.659912 | 0.977798 | 0.597455 | 0.373694 | 0.247095 | 0.215130 | 0.224589 | 0.255313 | 0.273844 | 0.279665 | 0.269950 | 0.242604 | 0.223659 | 0.216444 | 0.215823 | 0.220324 | 0.231977 | 0.244123 | 0.241169 | 0.220588 | 0.214679 | 0.226193 | 0.301691 | 0.509286 | 1.092077 | 2.673599 | 4.426606 | 5.312822 | 3.716525 | 1.261256 | 0.512147 | 0.287799 | 0.218136 | 0.222545 | 0.232184 | 0.224132 | 0.214941 | 0.217024 | 0.223223 | 0.229739 | 0.233744 | 0.234307 | 0.237950 | 0.255006 | 0.301386 | 0.393760 | 0.623898 | 1.035312 | 1.758592 | 2.345846 | 3.239449 | 4.346623 | 4.857655 | 3.937834 | 2.881457 | 1.643411 | 0.780001 | 0.296594 | 0.216746 | 0.341525 | 0.728271 | 1.795896 | 4.601636 | 11.596317 | 26.783610 | 57.567160 | 95.978057 | 124.225637 | 110.605095 | 75.271675 | 45.247743 | 20.959589 | 8.117700 | 3.238938 | 1.418770 | 0.733087 | 0.467820 | 0.393952 | 0.419773 | 0.507089 | 0.617604 | 0.768794 | 0.854357 | 0.822971 | 0.784022 | 0.766163 | 0.744885 | 0.759874 | 0.781086 | 0.800544 | 0.784715 | 0.723113 | 0.647962 | 0.583430 | 0.540485 | 0.549625 | 0.610465 | 0.769637 | 1.084275 | 1.577565 | 2.148992 | 2.700332 | 3.032657 | 3.250370 | 3.452096 | 3.918495 | 4.644032 | 5.159308 | 5.641620 | 5.959508 | 6.498733 | 7.447994 | 9.126740 | 11.397086 | 13.814827 | 15.514511 | 17.464787 | 19.140130 | 17.648148 | 14.581739 | 9.460908 | 4.995084 | 2.499166 | 1.358310 | 0.797706 | 0.572016 | 0.459665 | 0.427041 | 0.407149 | 0.397524 | 0.386478 | 0.376410 | 0.352742 | 0.331031 | 0.311702 | 0.293734 | 0.278424 | 0.271347 | 0.270309 | 0.274280 | 0.275331 | 0.267521 | 0.256765 | 0.247438 | 0.239783 | 0.234097 | 0.229582 | 0.225406 | 0.222512 | 0.219720 | 0.220570 | 0.225591 | 0.238983 | 0.264459 | 0.315442 | 0.407955 | 0.579645 | 0.828859 | 1.093293 | 1.222662 | 1.109224 | 0.811670 | 0.554073 | 0.386672 | 0.288064 | 0.234435 | 0.215335 | 0.220229 | 0.236306 | 0.239895 | 0.228444 | 0.215466 | 0.225079 | 0.280422 | 0.377021 | 0.507373 | 0.609596 | 0.610119 | 0.561805 | 0.473710 | 0.381141 | 0.325111 | 0.294543 | 0.270445 | 0.257391 | 0.256368 | 0.253013 | 0.249569 | 0.246747 | 0.245188 | 0.242549 | 0.251488 | 0.273114 | 0.320599 | 0.405103 | 0.563875 | 0.769159 | 1.025939 | 1.201918 | 1.330304 | 1.437879 | 1.488649 | 1.400755 | 1.350514 | 1.354920 | 1.344601 | 1.369826 | 1.387617 | 1.382417 | 1.218261 | 1.029115 | 0.866535 | 0.737341 | 0.636655 | 0.605120 | 0.592182 | 0.610518 | 0.709614 | 0.890942 | 1.119786 | 1.389504 | 1.563617 | 1.572010 | 1.469476 | 1.350332 | 1.255730 | 1.323593 | 1.566545 | 2.127358 | 3.134261 | 4.906755 | 7.650091 | 10.217526 | 10.023183 | 7.930412 | 5.469701 | 3.424712 | 2.086384 | 1.370117 | 0.931255 | 0.666699 | 0.552677 | 0.501399 | 0.462238 | 0.437432 | 0.411567 | 0.381283 | 0.353704 | 0.334759 | 0.327262 | 0.326687 | 0.314475 | 0.306344 | 0.291774 | 0.273305 | 0.256753 | 0.244915 | 0.235105 | 0.231222 | 0.230095 | 0.234310 | 0.245088 | 0.266049 | 0.291016 | 0.306893 | 0.301028 | 0.290859 | 0.262765 | 0.238607 | 0.223407 | 0.217492 | 0.215526 | 0.215335 | 0.214994 | 0.215782 |
| right posterior | 0.344404 | 0.419647 | 0.615151 | 0.990249 | 1.408258 | 1.205255 | 0.888039 | 0.476052 | 0.323542 | 0.238388 | 0.218352 | 0.214705 | 0.217663 | 0.228429 | 0.229799 | 0.236008 | 0.234955 | 0.261996 | 0.354964 | 0.433763 | 0.373494 | 0.324404 | 0.251937 | 0.214762 | 0.251821 | 0.428252 | 0.877245 | 1.589399 | 2.107798 | 1.597377 | 0.745790 | 0.409442 | 0.355173 | 0.385332 | 0.412054 | 0.404453 | 0.399009 | 0.384059 | 0.406583 | 0.480648 | 0.595318 | 0.775151 | 1.004775 | 1.065244 | 0.888543 | 0.714210 | 0.598425 | 0.515528 | 0.522186 | 0.709603 | 1.095326 | 1.508237 | 1.561600 | 1.616094 | 1.171797 | 0.796879 | 0.519152 | 0.342120 | 0.259281 | 0.233611 | 0.217478 | 0.216890 | 0.251011 | 0.387711 | 0.765029 | 1.581575 | 3.414069 | 6.629666 | 10.982254 | 17.994367 | 26.070250 | 28.356481 | 22.078335 | 13.965333 | 7.842931 | 4.276178 | 2.166441 | 1.039244 | 0.534894 | 0.317338 | 0.233180 | 0.215361 | 0.214627 | 0.216490 | 0.228509 | 0.268251 | 0.339303 | 0.492463 | 0.798642 | 1.424353 | 2.703399 | 5.320184 | 7.750480 | 9.574922 | 10.016555 | 9.084601 | 7.966960 | 6.994749 | 5.580471 | 4.922656 | 4.281303 | 3.777175 | 3.087354 | 2.589512 | 2.093012 | 1.736987 | 1.411451 | 1.252719 | 1.129242 | 1.073167 | 1.053799 | 1.134704 | 1.290640 | 1.582326 | 1.937661 | 2.513754 | 3.141963 | 3.700496 | 4.588986 | 6.056214 | 7.788796 | 10.012741 | 12.103305 | 13.717556 | 15.014185 | 15.144391 | 12.670240 | 9.182586 | 5.683604 | 2.941329 | 1.534742 | 0.931592 | 0.689093 | 0.621788 | 0.673695 | 0.798488 | 0.935537 | 1.087642 | 1.100503 | 1.054113 | 0.967450 | 0.839896 | 0.774360 | 0.834000 | 0.946521 | 1.008108 | 1.027275 | 0.900924 | 0.687731 | 0.484248 | 0.362029 | 0.298494 | 0.270308 | 0.263029 | 0.271974 | 0.296702 | 0.350473 | 0.460301 | 0.669318 | 1.058373 | 1.681671 | 2.475197 | 3.108101 | 3.185632 | 2.549405 | 1.663804 | 0.908707 | 0.522095 | 0.350634 | 0.279665 | 0.250150 | 0.241859 | 0.242086 | 0.253827 | 0.277590 | 0.322285 | 0.387108 | 0.458919 | 0.506808 | 0.512813 | 0.464029 | 0.392322 | 0.332752 | 0.303481 | 0.292592 | 0.294822 | 0.316223 | 0.350090 | 0.380517 | 0.386764 | 0.383316 | 0.368416 | 0.365720 | 0.372273 | 0.403994 | 0.462647 | 0.535440 | 0.569238 | 0.565696 | 0.554902 | 0.593354 | 0.687843 | 0.853540 | 1.069377 | 1.184270 | 1.135605 | 0.965274 | 0.746139 | 0.575377 | 0.472961 | 0.408602 | 0.375619 | 0.364615 | 0.377865 | 0.417644 | 0.457241 | 0.497703 | 0.537184 | 0.542628 | 0.508140 | 0.444408 | 0.385709 | 0.342406 | 0.316189 | 0.313556 | 0.339719 | 0.391379 | 0.470007 | 0.591326 | 0.717029 | 0.831169 | 0.984531 | 1.122599 | 1.122045 | 1.030187 | 0.977782 | 0.902827 | 0.770727 | 0.604860 | 0.461040 | 0.334291 | 0.259203 | 0.229406 | 0.220884 | 0.219774 | 0.224493 | 0.239308 | 0.283043 | 0.361523 | 0.461736 | 0.587473 | 0.699213 | 0.706148 | 0.645331 | 0.562972 | 0.486420 | 0.435127 | 0.409150 | 0.403643 | 0.418685 | 0.432258 | 0.425988 | 0.362417 | 0.293488 | 0.245147 | 0.220545 | 0.214736 | 0.216668 | 0.215728 | 0.214660 | 0.220021 | 0.240105 | 0.268119 | 0.296554 | 0.350627 | 0.408609 | 0.497980 | 0.611670 | 0.713775 | 0.772813 |
| all electrodes | 0.216518 | 0.217217 | 0.215331 | 0.215256 | 0.221726 | 0.226135 | 0.257383 | 0.291063 | 0.372092 | 0.543137 | 0.823044 | 0.870015 | 1.165731 | 1.141983 | 0.987379 | 0.683872 | 0.447480 | 0.277803 | 0.228504 | 0.216878 | 0.215553 | 0.214630 | 0.214803 | 0.216818 | 0.226990 | 0.260719 | 0.398126 | 0.645457 | 0.740403 | 0.807015 | 0.745942 | 0.516320 | 0.389163 | 0.350695 | 0.317105 | 0.331067 | 0.316188 | 0.251743 | 0.216651 | 0.233866 | 0.367928 | 0.679790 | 1.216417 | 1.850526 | 1.607329 | 0.960672 | 0.438835 | 0.262115 | 0.225827 | 0.215310 | 0.214769 | 0.217207 | 0.221388 | 0.229120 | 0.224729 | 0.220617 | 0.216016 | 0.214719 | 0.215612 | 0.219910 | 0.220882 | 0.215464 | 0.225479 | 0.332879 | 0.893065 | 4.595031 | 39.795916 | 287.615680 | 1422.510342 | 4418.117289 | 8445.214743 | 10304.613439 | 6634.246753 | 1709.759191 | 264.135619 | 44.500478 | 7.647998 | 1.606420 | 0.595460 | 0.336838 | 0.263992 | 0.250294 | 0.260839 | 0.279472 | 0.308476 | 0.309164 | 0.288142 | 0.256004 | 0.241067 | 0.234842 | 0.244016 | 0.275019 | 0.372222 | 0.615152 | 1.205092 | 2.135182 | 3.237650 | 3.224181 | 2.329249 | 1.444951 | 0.982084 | 0.711748 | 0.602974 | 0.557486 | 0.601918 | 0.729607 | 0.861990 | 0.890727 | 0.926294 | 1.015432 | 1.242265 | 1.736539 | 2.587707 | 3.895048 | 6.463370 | 7.809735 | 7.654431 | 7.130163 | 5.758105 | 3.900476 | 3.043788 | 2.375583 | 1.982003 | 1.631650 | 1.311037 | 1.177815 | 1.367514 | 1.546156 | 1.792416 | 2.055908 | 1.945644 | 1.494712 | 1.190598 | 0.991418 | 0.897148 | 0.936615 | 0.935885 | 0.933240 | 0.916945 | 0.849799 | 0.693640 | 0.617506 | 0.578063 | 0.565114 | 0.525232 | 0.499045 | 0.459823 | 0.419582 | 0.393121 | 0.398036 | 0.437432 | 0.582242 | 0.921483 | 1.454672 | 2.053265 | 2.421230 | 1.947708 | 1.378079 | 0.948437 | 0.636939 | 0.453285 | 0.350176 | 0.281413 | 0.245246 | 0.227901 | 0.215404 | 0.218699 | 0.241037 | 0.268775 | 0.305236 | 0.308816 | 0.300951 | 0.271347 | 0.259467 | 0.238737 | 0.224013 | 0.214732 | 0.219995 | 0.239183 | 0.262727 | 0.299032 | 0.352865 | 0.377456 | 0.378806 | 0.367903 | 0.365956 | 0.343810 | 0.317468 | 0.307981 | 0.307084 | 0.309245 | 0.335701 | 0.383887 | 0.412906 | 0.422941 | 0.395236 | 0.330229 | 0.298811 | 0.303234 | 0.317053 | 0.319712 | 0.333306 | 0.332555 | 0.299339 | 0.264809 | 0.244372 | 0.232105 | 0.235266 | 0.259894 | 0.314169 | 0.436431 | 0.640760 | 0.799017 | 0.815306 | 0.649560 | 0.476092 | 0.356767 | 0.315150 | 0.301699 | 0.325335 | 0.355874 | 0.392960 | 0.416170 | 0.490223 | 0.616043 | 0.831443 | 1.152295 | 1.617824 | 2.292801 | 3.505379 | 4.817455 | 5.525865 | 4.408370 | 3.334247 | 3.116184 | 3.214131 | 2.927340 | 2.873846 | 2.652778 | 2.502420 | 2.363108 | 2.505827 | 2.873074 | 3.094820 | 1.888171 | 1.448771 | 1.427776 | 1.494857 | 1.720066 | 2.622957 | 3.391475 | 4.873656 | 6.575170 | 7.648165 | 7.032700 | 5.187940 | 2.061484 | 0.953738 | 0.551835 | 0.390981 | 0.323168 | 0.305598 | 0.303202 | 0.319304 | 0.358242 | 0.420299 | 0.498594 | 0.581979 | 0.684390 | 0.800178 | 1.057153 | 1.504408 | 2.140064 | 2.508176 | 2.799076 | 2.420549 | 1.768126 | 1.021970 | 0.861967 | 0.647000 |

G) happy vs angry

  
|  | time window | peak latency | cluster *p* | peak Cohen's *d* |  | | | |
| **all electrodes** | 365 - 475 ms | 405 ms | 0.0336 | 0.7268 |  | | | |
|  | | | | | | | | |

Model correlations, cluster permutation tests

|  | **left hemisphere** | | | | **right hemisphere** | | | |
|  | time window | peak latency | cluster *p* | peak Cohen's *d* | time window | peak latency | cluster *p* | peak Cohen's *d* |
| **anterior** |  | | | |  | | | |
| **central** |  | | | | 785 - 895 ms | 820 ms | 0.0434 | 0.8076 |
| **posterior** | 130 - 215 ms | 170 ms | 0.0289 | 1.1229 | 860 - 1065 ms | 895 ms | 0.021 | 0.5456 |
 265 - 365 ms | 340 ms | 0.049 | 0.5897 |  | | | | 965 - 1080 ms | 990 ms | 0.0401 | 0.4992 |  | | | |

  

Model correlations, Bayesian statistics

|  | -200 | -195 | -190 | -185 | -180 | -175 | -170 | -165 | -160 | -155 | -150 | -145 | -140 | -135 | -130 | -125 | -120 | -115 | -110 | -105 | -100 | -95 | -90 | -85 | -80 | -75 | -70 | -65 | -60 | -55 | -50 | -45 | -40 | -35 | -30 | -25 | -20 | -15 | -10 | -5 | 0 | 5 | 10 | 15 | 20 | 25 | 30 | 35 | 40 | 45 | 50 | 55 | 60 | 65 | 70 | 75 | 80 | 85 | 90 | 95 | 100 | 105 | 110 | 115 | 120 | 125 | 130 | 135 | 140 | 145 | 150 | 155 | 160 | 165 | 170 | 175 | 180 | 185 | 190 | 195 | 200 | 205 | 210 | 215 | 220 | 225 | 230 | 235 | 240 | 245 | 250 | 255 | 260 | 265 | 270 | 275 | 280 | 285 | 290 | 295 | 300 | 305 | 310 | 315 | 320 | 325 | 330 | 335 | 340 | 345 | 350 | 355 | 360 | 365 | 370 | 375 | 380 | 385 | 390 | 395 | 400 | 405 | 410 | 415 | 420 | 425 | 430 | 435 | 440 | 445 | 450 | 455 | 460 | 465 | 470 | 475 | 480 | 485 | 490 | 495 | 500 | 505 | 510 | 515 | 520 | 525 | 530 | 535 | 540 | 545 | 550 | 555 | 560 | 565 | 570 | 575 | 580 | 585 | 590 | 595 | 600 | 605 | 610 | 615 | 620 | 625 | 630 | 635 | 640 | 645 | 650 | 655 | 660 | 665 | 670 | 675 | 680 | 685 | 690 | 695 | 700 | 705 | 710 | 715 | 720 | 725 | 730 | 735 | 740 | 745 | 750 | 755 | 760 | 765 | 770 | 775 | 780 | 785 | 790 | 795 | 800 | 805 | 810 | 815 | 820 | 825 | 830 | 835 | 840 | 845 | 850 | 855 | 860 | 865 | 870 | 875 | 880 | 885 | 890 | 895 | 900 | 905 | 910 | 915 | 920 | 925 | 930 | 935 | 940 | 945 | 950 | 955 | 960 | 965 | 970 | 975 | 980 | 985 | 990 | 995 | 1000 | 1005 | 1010 | 1015 | 1020 | 1025 | 1030 | 1035 | 1040 | 1045 | 1050 | 1055 | 1060 | 1065 | 1070 | 1075 | 1080 | 1085 | 1090 | 1095 | 1100 | 1105 | 1110 | 1115 | 1120 | 1125 | 1130 | 1135 | 1140 | 1145 | 1150 | 1155 | 1160 | 1165 | 1170 | 1175 | 1180 | 1185 | 1190 | 1195 |
| --- | --- | --- | --- | --- | --- | --- | --- | --- | --- | --- | --- | --- | --- | --- | --- | --- | --- | --- | --- | --- | --- | --- | --- | --- | --- | --- | --- | --- | --- | --- | --- | --- | --- | --- | --- | --- | --- | --- | --- | --- | --- | --- | --- | --- | --- | --- | --- | --- | --- | --- | --- | --- | --- | --- | --- | --- | --- | --- | --- | --- | --- | --- | --- | --- | --- | --- | --- | --- | --- | --- | --- | --- | --- | --- | --- | --- | --- | --- | --- | --- | --- | --- | --- | --- | --- | --- | --- | --- | --- | --- | --- | --- | --- | --- | --- | --- | --- | --- | --- | --- | --- | --- | --- | --- | --- | --- | --- | --- | --- | --- | --- | --- | --- | --- | --- | --- | --- | --- | --- | --- | --- | --- | --- | --- | --- | --- | --- | --- | --- | --- | --- | --- | --- | --- | --- | --- | --- | --- | --- | --- | --- | --- | --- | --- | --- | --- | --- | --- | --- | --- | --- | --- | --- | --- | --- | --- | --- | --- | --- | --- | --- | --- | --- | --- | --- | --- | --- | --- | --- | --- | --- | --- | --- | --- | --- | --- | --- | --- | --- | --- | --- | --- | --- | --- | --- | --- | --- | --- | --- | --- | --- | --- | --- | --- | --- | --- | --- | --- | --- | --- | --- | --- | --- | --- | --- | --- | --- | --- | --- | --- | --- | --- | --- | --- | --- | --- | --- | --- | --- | --- | --- | --- | --- | --- | --- | --- | --- | --- | --- | --- | --- | --- | --- | --- | --- | --- | --- | --- | --- | --- | --- | --- | --- | --- | --- | --- | --- | --- | --- | --- | --- | --- | --- | --- | --- | --- | --- | --- | --- | --- | --- | --- | --- | --- | --- | --- | --- | --- | --- | --- | --- | --- | --- | --- | --- | --- | --- | --- | --- | --- |
| left anterior | 0.450992 | 0.611555 | 0.756650 | 1.499930 | 1.988714 | 1.616249 | 0.759661 | 0.408608 | 0.285134 | 0.244371 | 0.225155 | 0.251082 | 0.372012 | 0.904024 | 2.066914 | 2.036172 | 1.536416 | 1.317496 | 0.999937 | 0.869410 | 0.723332 | 0.543262 | 0.389887 | 0.278186 | 0.217213 | 0.222620 | 0.258544 | 0.303887 | 0.322411 | 0.295919 | 0.247285 | 0.216315 | 0.229089 | 0.321255 | 0.655693 | 1.810554 | 4.850635 | 4.621129 | 2.096289 | 0.729095 | 0.354891 | 0.246596 | 0.227135 | 0.221711 | 0.239478 | 0.302423 | 0.457081 | 0.462098 | 0.489950 | 0.436414 | 0.451066 | 0.573045 | 1.145249 | 2.098177 | 3.671607 | 4.444681 | 3.537845 | 1.790481 | 1.024789 | 0.610314 | 0.380009 | 0.288386 | 0.244202 | 0.220912 | 0.214751 | 0.221359 | 0.236126 | 0.254569 | 0.303089 | 0.342615 | 0.389183 | 0.434628 | 0.518317 | 0.535548 | 0.488181 | 0.412048 | 0.333180 | 0.268608 | 0.225880 | 0.217673 | 0.305668 | 0.684716 | 1.559102 | 2.299881 | 2.031948 | 1.456998 | 1.031691 | 0.652509 | 0.475200 | 0.410836 | 0.413152 | 0.506071 | 0.944606 | 2.225544 | 6.465297 | 17.360419 | 44.258179 | 77.967191 | 86.279983 | 65.240024 | 39.546400 | 18.996210 | 11.162063 | 6.968729 | 4.315910 | 2.630152 | 1.647126 | 0.803978 | 0.475626 | 0.358550 | 0.302629 | 0.268064 | 0.245417 | 0.227633 | 0.218711 | 0.215144 | 0.214640 | 0.216443 | 0.217238 | 0.217808 | 0.216254 | 0.214902 | 0.216622 | 0.228053 | 0.258865 | 0.312502 | 0.398739 | 0.472249 | 0.486134 | 0.428338 | 0.344773 | 0.282228 | 0.248289 | 0.229212 | 0.219358 | 0.216071 | 0.215183 | 0.215158 | 0.217605 | 0.226643 | 0.254094 | 0.318014 | 0.462385 | 0.654470 | 0.825015 | 0.643992 | 0.429431 | 0.306755 | 0.251552 | 0.227482 | 0.220860 | 0.220669 | 0.230723 | 0.255484 | 0.300441 | 0.419775 | 0.646659 | 0.827864 | 1.235891 | 2.124372 | 3.075896 | 4.848439 | 7.129967 | 8.212402 | 8.885151 | 7.796814 | 4.787285 | 2.776295 | 1.449042 | 0.759443 | 0.462185 | 0.351544 | 0.302518 | 0.300590 | 0.323642 | 0.376450 | 0.431325 | 0.456422 | 0.424106 | 0.377080 | 0.319661 | 0.287965 | 0.274448 | 0.287308 | 0.329564 | 0.435213 | 0.647063 | 0.881772 | 0.791894 | 0.680555 | 0.569812 | 0.475870 | 0.394772 | 0.342267 | 0.284188 | 0.245985 | 0.219174 | 0.215037 | 0.219750 | 0.230518 | 0.251255 | 0.271988 | 0.292669 | 0.306629 | 0.320246 | 0.327274 | 0.331031 | 0.348236 | 0.401580 | 0.437370 | 0.472484 | 0.489415 | 0.554979 | 0.725164 | 0.993667 | 1.310711 | 1.813758 | 2.417951 | 2.929932 | 3.062537 | 2.994280 | 2.641615 | 2.441468 | 2.150436 | 1.706674 | 1.349372 | 1.085159 | 0.832447 | 0.674846 | 0.599850 | 0.528923 | 0.485734 | 0.439544 | 0.393986 | 0.375649 | 0.363780 | 0.316272 | 0.283188 | 0.253337 | 0.226919 | 0.216595 | 0.214793 | 0.215787 | 0.217893 | 0.216289 | 0.214713 | 0.215952 | 0.221910 | 0.227062 | 0.230051 | 0.231005 | 0.232450 | 0.242542 | 0.268099 | 0.308367 | 0.364933 | 0.445958 | 0.531292 | 0.568271 | 0.543241 | 0.522108 | 0.503072 | 0.526954 | 0.585800 | 0.664218 | 0.762208 | 0.864967 | 0.895972 | 0.895413 | 0.948536 | 0.982886 | 0.969827 | 0.912135 | 0.821849 | 0.769770 | 0.700746 | 0.624874 | 0.585700 | 0.578618 | 0.538576 |
| right anterior | 0.442506 | 0.351417 | 0.295759 | 0.235521 | 0.214794 | 0.250499 | 0.356918 | 0.563088 | 0.927310 | 1.229188 | 1.263752 | 1.113001 | 1.045976 | 0.891715 | 0.627578 | 0.427950 | 0.306227 | 0.235040 | 0.215117 | 0.231357 | 0.249090 | 0.268260 | 0.343980 | 0.439831 | 0.559815 | 0.620301 | 0.606807 | 0.455361 | 0.288381 | 0.218557 | 0.237983 | 0.426260 | 0.891702 | 1.620087 | 1.908672 | 1.392597 | 0.828215 | 0.553878 | 0.370012 | 0.314490 | 0.312259 | 0.341889 | 0.377256 | 0.452424 | 0.654385 | 1.147206 | 1.257022 | 1.046638 | 0.600669 | 0.382534 | 0.320478 | 0.310458 | 0.317640 | 0.362629 | 0.369337 | 0.388347 | 0.405934 | 0.475032 | 0.466817 | 0.449071 | 0.354406 | 0.300683 | 0.235564 | 0.215609 | 0.218572 | 0.225011 | 0.236881 | 0.236195 | 0.235946 | 0.247523 | 0.302981 | 0.360615 | 0.412374 | 0.470456 | 0.490969 | 0.477892 | 0.437740 | 0.362914 | 0.273428 | 0.220254 | 0.234592 | 0.339150 | 0.546565 | 0.773138 | 0.972748 | 0.859620 | 0.622176 | 0.482511 | 0.345095 | 0.254500 | 0.220769 | 0.216304 | 0.238776 | 0.278931 | 0.326065 | 0.354725 | 0.316756 | 0.286427 | 0.255298 | 0.234461 | 0.219976 | 0.214901 | 0.217026 | 0.220162 | 0.225475 | 0.224855 | 0.220182 | 0.214741 | 0.216210 | 0.224599 | 0.240882 | 0.274403 | 0.310888 | 0.378293 | 0.500632 | 0.731253 | 1.116168 | 1.869484 | 2.849566 | 4.558801 | 7.012323 | 9.363409 | 11.387386 | 11.802296 | 7.087258 | 2.904040 | 0.997115 | 0.464506 | 0.276893 | 0.225507 | 0.215976 | 0.233694 | 0.267115 | 0.274122 | 0.266978 | 0.247334 | 0.245347 | 0.261040 | 0.299056 | 0.365161 | 0.377768 | 0.345115 | 0.252206 | 0.218392 | 0.348296 | 0.755640 | 1.702255 | 3.090923 | 3.690867 | 3.928010 | 2.842157 | 1.959556 | 1.228646 | 0.653672 | 0.343310 | 0.249077 | 0.214766 | 0.234930 | 0.300986 | 0.394676 | 0.495049 | 0.569649 | 0.626727 | 0.641230 | 0.719113 | 0.749267 | 0.656208 | 0.533119 | 0.405087 | 0.296911 | 0.245088 | 0.220627 | 0.214709 | 0.214680 | 0.215278 | 0.223089 | 0.240375 | 0.269836 | 0.292460 | 0.281592 | 0.267937 | 0.286443 | 0.377539 | 0.705529 | 1.574312 | 2.699789 | 3.260002 | 2.577206 | 1.772688 | 1.303491 | 1.213873 | 1.354584 | 1.572524 | 1.878134 | 2.064520 | 1.778542 | 1.291723 | 0.871782 | 0.704165 | 0.696000 | 0.906951 | 1.324489 | 2.318831 | 2.460816 | 1.825622 | 0.938772 | 0.497407 | 0.313826 | 0.258997 | 0.236243 | 0.235632 | 0.239641 | 0.249689 | 0.257352 | 0.258936 | 0.254927 | 0.257701 | 0.251511 | 0.244819 | 0.238520 | 0.227457 | 0.216154 | 0.215430 | 0.228719 | 0.249498 | 0.265186 | 0.261875 | 0.257668 | 0.251379 | 0.259470 | 0.274703 | 0.307055 | 0.375998 | 0.479488 | 0.529165 | 0.495434 | 0.392042 | 0.280173 | 0.222711 | 0.216100 | 0.228123 | 0.231931 | 0.223971 | 0.216774 | 0.215327 | 0.227155 | 0.249975 | 0.265212 | 0.274839 | 0.262850 | 0.248639 | 0.242673 | 0.242847 | 0.247309 | 0.275517 | 0.332755 | 0.416088 | 0.540833 | 0.673853 | 0.759716 | 0.791006 | 0.770394 | 0.699874 | 0.646638 | 0.606575 | 0.563675 | 0.478660 | 0.374628 | 0.290408 | 0.237380 | 0.216864 | 0.214796 | 0.215535 | 0.215704 | 0.216504 | 0.218966 | 0.220377 | 0.221695 | 0.223765 | 0.223298 |
| left central | 0.540560 | 0.502034 | 0.517198 | 0.485797 | 0.411017 | 0.351024 | 0.356838 | 0.359110 | 0.360732 | 0.370421 | 0.409812 | 0.429506 | 0.499472 | 0.652332 | 1.026155 | 1.667064 | 2.479991 | 2.647620 | 2.171060 | 1.571601 | 1.178615 | 0.810456 | 0.582697 | 0.514396 | 0.448899 | 0.382173 | 0.355712 | 0.384788 | 0.398636 | 0.426432 | 0.553252 | 0.689370 | 0.663006 | 0.436663 | 0.305842 | 0.246075 | 0.224228 | 0.216347 | 0.214932 | 0.215176 | 0.215364 | 0.214659 | 0.215301 | 0.218577 | 0.227928 | 0.232479 | 0.229528 | 0.219094 | 0.214655 | 0.216423 | 0.222077 | 0.230940 | 0.238367 | 0.232693 | 0.225780 | 0.219293 | 0.217175 | 0.214647 | 0.216573 | 0.225033 | 0.233838 | 0.247489 | 0.261515 | 0.274259 | 0.263172 | 0.240840 | 0.214974 | 0.245138 | 0.442339 | 1.019149 | 1.893959 | 2.393029 | 2.752479 | 2.737599 | 2.349824 | 1.706852 | 1.115675 | 0.706337 | 0.448107 | 0.273698 | 0.214711 | 0.291677 | 0.639027 | 1.576449 | 3.338057 | 5.515039 | 5.854622 | 4.638901 | 2.701161 | 1.410548 | 0.734436 | 0.435668 | 0.332266 | 0.295909 | 0.276483 | 0.281907 | 0.335868 | 0.430812 | 0.624201 | 0.936113 | 1.296431 | 1.388696 | 1.175648 | 0.865907 | 0.597042 | 0.406240 | 0.297262 | 0.243566 | 0.221307 | 0.215590 | 0.214871 | 0.217568 | 0.221486 | 0.229520 | 0.245146 | 0.271999 | 0.308521 | 0.358485 | 0.404160 | 0.441214 | 0.455153 | 0.443810 | 0.395086 | 0.350646 | 0.311388 | 0.282707 | 0.268967 | 0.272861 | 0.282017 | 0.300044 | 0.323413 | 0.345486 | 0.334932 | 0.309235 | 0.268030 | 0.231409 | 0.214950 | 0.221928 | 0.259509 | 0.331506 | 0.484980 | 0.802381 | 1.338412 | 1.968801 | 3.292965 | 4.941091 | 5.350881 | 4.180163 | 2.361875 | 1.175032 | 0.673365 | 0.458394 | 0.374259 | 0.372848 | 0.385382 | 0.422891 | 0.469091 | 0.538582 | 0.528816 | 0.473989 | 0.393567 | 0.382861 | 0.388044 | 0.483691 | 0.703191 | 1.048761 | 1.218856 | 1.084917 | 0.741967 | 0.579082 | 0.470340 | 0.407096 | 0.356480 | 0.328778 | 0.299506 | 0.273922 | 0.243998 | 0.229144 | 0.217507 | 0.214846 | 0.219217 | 0.221512 | 0.223097 | 0.219470 | 0.216256 | 0.214719 | 0.217292 | 0.225920 | 0.229610 | 0.235643 | 0.237196 | 0.234900 | 0.222004 | 0.215519 | 0.219736 | 0.246736 | 0.296224 | 0.347212 | 0.374293 | 0.386310 | 0.414122 | 0.435100 | 0.413644 | 0.371923 | 0.322710 | 0.284083 | 0.268763 | 0.268511 | 0.286413 | 0.331850 | 0.406698 | 0.485386 | 0.563070 | 0.610676 | 0.639137 | 0.668080 | 0.729057 | 0.852734 | 1.067551 | 1.398563 | 1.848773 | 2.298463 | 2.218417 | 1.679081 | 1.178995 | 0.922384 | 0.910446 | 1.205701 | 2.274925 | 5.884223 | 13.415177 | 17.299224 | 13.995935 | 7.871785 | 3.582725 | 1.641982 | 0.826403 | 0.473460 | 0.342398 | 0.287027 | 0.264370 | 0.248509 | 0.238144 | 0.229992 | 0.218137 | 0.215535 | 0.225998 | 0.239341 | 0.250293 | 0.261367 | 0.269302 | 0.259978 | 0.245139 | 0.232813 | 0.220323 | 0.214646 | 0.218689 | 0.231378 | 0.254506 | 0.287764 | 0.319440 | 0.337624 | 0.339233 | 0.334960 | 0.317587 | 0.296008 | 0.276504 | 0.249329 | 0.225916 | 0.215323 | 0.216916 | 0.234677 | 0.267067 | 0.317425 | 0.372063 | 0.422922 | 0.450531 | 0.456454 | 0.443799 | 0.434953 |
| right central | 0.244518 | 0.269193 | 0.281668 | 0.331339 | 0.388341 | 0.415303 | 0.401329 | 0.390977 | 0.344619 | 0.331517 | 0.288905 | 0.240643 | 0.218002 | 0.214679 | 0.219672 | 0.221614 | 0.231058 | 0.240517 | 0.242928 | 0.242018 | 0.246336 | 0.242895 | 0.267837 | 0.340164 | 0.462146 | 0.567079 | 0.529313 | 0.468301 | 0.438124 | 0.430827 | 0.442188 | 0.584739 | 0.805882 | 1.087922 | 1.038618 | 0.546982 | 0.275826 | 0.214741 | 0.315846 | 0.736092 | 1.803988 | 4.056918 | 6.378675 | 3.614227 | 1.309183 | 0.418045 | 0.243383 | 0.215605 | 0.215237 | 0.215206 | 0.222769 | 0.250291 | 0.308430 | 0.406248 | 0.709086 | 1.366398 | 3.323305 | 5.413831 | 3.025678 | 1.065636 | 0.499473 | 0.290929 | 0.229906 | 0.214628 | 0.224989 | 0.238066 | 0.254738 | 0.310566 | 0.463552 | 1.109967 | 3.855714 | 11.480493 | 19.841340 | 20.129131 | 10.712090 | 4.367466 | 1.800074 | 0.791868 | 0.407722 | 0.274889 | 0.230991 | 0.222628 | 0.230198 | 0.247496 | 0.277138 | 0.342593 | 0.387835 | 0.415419 | 0.519697 | 0.770083 | 1.309299 | 2.533237 | 5.006953 | 8.200147 | 11.198086 | 9.451636 | 6.681510 | 3.450614 | 1.492869 | 0.693594 | 0.445010 | 0.367636 | 0.384406 | 0.494894 | 0.673596 | 1.017579 | 1.456301 | 1.548842 | 1.171128 | 0.771222 | 0.533510 | 0.419867 | 0.382607 | 0.397716 | 0.455821 | 0.588573 | 0.853004 | 1.279891 | 1.943754 | 2.746558 | 3.175249 | 3.258871 | 3.151663 | 3.163263 | 3.769134 | 4.943654 | 7.359772 | 12.223825 | 18.345671 | 24.282214 | 31.279281 | 26.841936 | 17.521080 | 10.536788 | 6.397201 | 3.817683 | 2.396546 | 1.409451 | 0.933516 | 0.699164 | 0.602537 | 0.595625 | 0.673607 | 0.726560 | 0.731921 | 0.677060 | 0.581322 | 0.489161 | 0.502279 | 0.527346 | 0.558170 | 0.590759 | 0.630987 | 0.611794 | 0.607974 | 0.647708 | 0.727091 | 0.827698 | 1.121635 | 1.684866 | 2.521211 | 3.302964 | 3.476921 | 2.539725 | 1.637302 | 0.994720 | 0.711458 | 0.554580 | 0.515998 | 0.561715 | 0.665448 | 0.776277 | 0.963976 | 1.087814 | 1.131701 | 1.119721 | 1.142298 | 1.294265 | 1.482633 | 1.579279 | 1.681697 | 1.726399 | 1.561774 | 1.234975 | 0.922986 | 0.684503 | 0.514227 | 0.393395 | 0.346592 | 0.331296 | 0.332975 | 0.365328 | 0.434507 | 0.574541 | 0.852604 | 1.211152 | 1.329638 | 1.253239 | 1.207649 | 1.338591 | 1.867035 | 3.700550 | 11.175779 | 39.338818 | 88.739892 | 92.209296 | 45.144341 | 13.168708 | 3.318278 | 1.311192 | 0.869790 | 0.900107 | 1.223191 | 2.025106 | 2.956810 | 3.781192 | 3.593363 | 2.886999 | 1.968480 | 1.358560 | 0.933363 | 0.818748 | 0.768658 | 0.827682 | 0.954130 | 1.069441 | 0.974226 | 0.891295 | 0.808670 | 0.733738 | 0.635376 | 0.553440 | 0.502364 | 0.517033 | 0.573866 | 0.660227 | 0.730921 | 0.760527 | 0.730936 | 0.586487 | 0.442724 | 0.337464 | 0.267716 | 0.228722 | 0.219442 | 0.218329 | 0.226006 | 0.248675 | 0.297214 | 0.374510 | 0.485199 | 0.603649 | 0.719651 | 0.729275 | 0.627313 | 0.488837 | 0.345047 | 0.275296 | 0.237773 | 0.218583 | 0.214656 | 0.214632 | 0.214647 | 0.215580 | 0.228176 | 0.264415 | 0.310346 | 0.336617 | 0.345547 | 0.338070 | 0.309620 | 0.272492 | 0.244799 | 0.224521 | 0.215951 | 0.215160 | 0.216772 | 0.215379 | 0.214678 | 0.214724 |
| left posterior | 0.440518 | 0.516416 | 0.611458 | 1.015059 | 1.316237 | 1.218730 | 0.819478 | 0.604092 | 0.414452 | 0.290537 | 0.247439 | 0.232853 | 0.240012 | 0.279703 | 0.530954 | 3.526811 | 134.668068 | 4399.163618 | 27542.910264 | 10433.941966 | 380.624772 | 15.834129 | 1.869739 | 0.407122 | 0.220207 | 0.224136 | 0.258795 | 0.282172 | 0.297913 | 0.243671 | 0.219477 | 0.216076 | 0.224576 | 0.229268 | 0.234620 | 0.233563 | 0.234183 | 0.222002 | 0.217326 | 0.215846 | 0.215420 | 0.215765 | 0.226191 | 0.254459 | 0.307096 | 0.399028 | 0.506284 | 0.601806 | 0.676851 | 0.754355 | 0.711123 | 0.620791 | 0.526481 | 0.472343 | 0.396310 | 0.318914 | 0.287346 | 0.289294 | 0.284516 | 0.280843 | 0.288034 | 0.289888 | 0.305425 | 0.349994 | 0.440310 | 0.702400 | 1.445107 | 3.578419 | 10.233720 | 30.760568 | 79.588787 | 193.479215 | 460.446000 | 1010.382296 | 1853.694926 | 2403.978025 | 1889.608765 | 1151.066847 | 486.607841 | 161.752980 | 55.790207 | 19.845540 | 5.490299 | 1.473370 | 0.568789 | 0.360691 | 0.287431 | 0.251319 | 0.248914 | 0.269359 | 0.314623 | 0.451675 | 0.931770 | 2.406829 | 7.262561 | 16.364587 | 21.313016 | 16.853148 | 10.972483 | 7.716034 | 5.407856 | 3.458368 | 2.523228 | 2.415128 | 2.633722 | 3.556583 | 4.602158 | 6.169424 | 6.871709 | 5.612828 | 3.629040 | 2.262608 | 1.408244 | 1.060854 | 0.944943 | 0.951706 | 1.119593 | 1.537394 | 2.193892 | 3.667199 | 6.840700 | 11.101135 | 12.424707 | 10.999831 | 7.160739 | 3.798632 | 1.965113 | 1.118150 | 0.710392 | 0.523033 | 0.436872 | 0.395860 | 0.386295 | 0.395389 | 0.410284 | 0.425512 | 0.468051 | 0.537808 | 0.587498 | 0.649429 | 0.743020 | 0.839690 | 0.942102 | 1.119285 | 1.229647 | 1.293823 | 1.075392 | 0.856072 | 0.739571 | 0.682504 | 0.649264 | 0.675668 | 0.677362 | 0.608206 | 0.509779 | 0.393782 | 0.315930 | 0.259949 | 0.228618 | 0.214890 | 0.221319 | 0.244201 | 0.264513 | 0.268366 | 0.258539 | 0.247239 | 0.225202 | 0.215144 | 0.244209 | 0.334126 | 0.547437 | 0.890016 | 1.221089 | 1.480089 | 1.584175 | 1.473328 | 1.319643 | 1.082954 | 0.846150 | 0.762180 | 0.781779 | 0.831933 | 0.956679 | 1.103690 | 1.150016 | 1.048591 | 0.874318 | 0.704664 | 0.580495 | 0.497201 | 0.454331 | 0.468034 | 0.545306 | 0.705404 | 1.059041 | 1.796579 | 2.923769 | 4.279136 | 5.432254 | 5.152030 | 4.004343 | 2.722470 | 1.807732 | 1.261462 | 0.996454 | 0.814539 | 0.745918 | 0.752656 | 0.853684 | 1.077543 | 1.742923 | 3.349741 | 6.852864 | 14.558084 | 29.924230 | 50.175044 | 73.749804 | 101.513574 | 89.876602 | 56.610383 | 28.724537 | 12.223661 | 4.866898 | 2.191959 | 1.205142 | 0.864949 | 0.733241 | 0.763029 | 0.905301 | 1.132602 | 1.185054 | 1.222141 | 1.158571 | 1.122081 | 1.173177 | 1.334635 | 1.540542 | 1.981032 | 2.410784 | 2.791453 | 3.397449 | 4.242989 | 5.011421 | 6.615901 | 8.634948 | 12.446935 | 16.359256 | 19.337267 | 17.565914 | 14.481701 | 9.295053 | 5.666713 | 3.612604 | 2.529530 | 1.784542 | 1.458425 | 1.212869 | 1.039691 | 0.889984 | 0.754876 | 0.595647 | 0.468512 | 0.385403 | 0.341924 | 0.318628 | 0.307307 | 0.310514 | 0.315470 | 0.323419 | 0.326390 | 0.320570 | 0.306493 | 0.284021 | 0.258161 | 0.241232 | 0.231398 | 0.225026 | 0.222276 | 0.220378 | 0.219327 |
| right posterior | 0.375225 | 0.340690 | 0.306167 | 0.276858 | 0.270443 | 0.296385 | 0.345834 | 0.501383 | 0.817657 | 1.135474 | 1.153473 | 0.798591 | 0.445643 | 0.332118 | 0.302497 | 0.290095 | 0.320615 | 0.412262 | 0.544130 | 0.558077 | 0.510680 | 0.367168 | 0.269225 | 0.222728 | 0.215272 | 0.216898 | 0.238860 | 0.285682 | 0.386291 | 0.645573 | 1.266763 | 1.586109 | 0.880712 | 0.403356 | 0.245383 | 0.214653 | 0.236334 | 0.268624 | 0.302143 | 0.294433 | 0.252120 | 0.215415 | 0.232946 | 0.327758 | 0.499125 | 0.697672 | 0.851814 | 1.109017 | 1.227406 | 1.195769 | 0.901277 | 0.576135 | 0.356611 | 0.256193 | 0.224917 | 0.217269 | 0.217633 | 0.225629 | 0.241883 | 0.268371 | 0.305870 | 0.310331 | 0.274834 | 0.245787 | 0.225880 | 0.214820 | 0.224249 | 0.287489 | 0.511688 | 1.269119 | 2.936922 | 5.958604 | 9.202302 | 9.756640 | 9.129813 | 7.544872 | 4.970315 | 2.948418 | 1.490328 | 0.651164 | 0.322967 | 0.229446 | 0.214673 | 0.220602 | 0.227537 | 0.227610 | 0.225516 | 0.222206 | 0.218984 | 0.215707 | 0.214690 | 0.214627 | 0.214878 | 0.217086 | 0.222168 | 0.230317 | 0.235459 | 0.231799 | 0.224145 | 0.219181 | 0.214710 | 0.220099 | 0.238901 | 0.281409 | 0.343930 | 0.400326 | 0.459178 | 0.504063 | 0.497812 | 0.496587 | 0.471730 | 0.419178 | 0.383603 | 0.366358 | 0.360384 | 0.382826 | 0.443455 | 0.570379 | 0.821743 | 1.288771 | 1.963903 | 3.017871 | 4.053051 | 4.244887 | 3.238898 | 2.283854 | 1.473690 | 1.035329 | 0.794680 | 0.669366 | 0.595975 | 0.574520 | 0.548891 | 0.535392 | 0.527588 | 0.529101 | 0.514549 | 0.507749 | 0.539479 | 0.651175 | 0.851410 | 1.169459 | 1.738137 | 2.407764 | 2.543681 | 2.199340 | 1.478368 | 0.920044 | 0.590088 | 0.436773 | 0.343537 | 0.301201 | 0.269700 | 0.256221 | 0.252938 | 0.270729 | 0.304491 | 0.377682 | 0.522524 | 0.731875 | 0.912692 | 1.131733 | 1.286523 | 1.333062 | 1.378666 | 1.441686 | 1.436747 | 1.508105 | 1.342001 | 1.087132 | 0.854945 | 0.703590 | 0.582898 | 0.508473 | 0.458208 | 0.446544 | 0.430360 | 0.419543 | 0.408310 | 0.400172 | 0.384337 | 0.390990 | 0.422032 | 0.511955 | 0.701133 | 1.114867 | 1.772947 | 2.494600 | 2.683757 | 2.342882 | 1.671211 | 1.189130 | 0.943791 | 0.862925 | 0.872447 | 0.997142 | 1.189584 | 1.385021 | 1.515498 | 1.557188 | 1.471686 | 1.364750 | 1.232738 | 1.174847 | 1.178664 | 1.192424 | 1.199948 | 1.263474 | 1.351985 | 1.324693 | 1.207622 | 1.110655 | 1.145633 | 1.247357 | 1.473851 | 2.049455 | 2.782240 | 3.324892 | 3.609368 | 3.781126 | 3.664354 | 3.560877 | 3.319178 | 2.928861 | 2.644902 | 2.430667 | 2.286427 | 2.241583 | 2.285629 | 2.469020 | 2.967235 | 3.681626 | 4.506722 | 6.024940 | 8.117293 | 9.735426 | 11.231494 | 12.605569 | 12.707349 | 11.996140 | 10.126272 | 8.391622 | 6.576238 | 5.064964 | 3.975765 | 3.214989 | 2.755968 | 2.575982 | 2.450965 | 2.417998 | 2.396281 | 2.251671 | 1.904161 | 1.452259 | 1.040332 | 0.783310 | 0.651497 | 0.601770 | 0.616778 | 0.684982 | 0.781906 | 0.885774 | 0.996317 | 1.073158 | 1.132490 | 1.095854 | 1.025878 | 0.909452 | 0.735967 | 0.525095 | 0.384479 | 0.288807 | 0.239795 | 0.221837 | 0.217590 | 0.217345 | 0.225020 | 0.245504 | 0.262485 | 0.277064 |
| all electrodes | 0.869712 | 1.892428 | 3.751912 | 12.529950 | 20.821092 | 12.675677 | 3.967611 | 1.798133 | 0.930645 | 0.527613 | 0.361369 | 0.307109 | 0.287190 | 0.321124 | 0.362582 | 0.404593 | 0.488589 | 0.619807 | 0.699681 | 1.361018 | 2.519777 | 2.462698 | 0.878618 | 0.351185 | 0.225324 | 0.214649 | 0.220786 | 0.215222 | 0.217466 | 0.235023 | 0.255072 | 0.299908 | 0.293311 | 0.270086 | 0.232816 | 0.219421 | 0.214774 | 0.218198 | 0.224023 | 0.219628 | 0.215073 | 0.249080 | 0.409543 | 1.231164 | 4.072188 | 6.968618 | 6.737027 | 4.260373 | 2.029241 | 1.204432 | 0.622605 | 0.427635 | 0.389824 | 0.441956 | 0.509492 | 0.518616 | 0.385121 | 0.265353 | 0.215875 | 0.226584 | 0.254606 | 0.281830 | 0.269853 | 0.240138 | 0.219797 | 0.215747 | 0.223950 | 0.217228 | 0.220498 | 0.293914 | 0.619050 | 1.999713 | 8.420256 | 31.310184 | 67.611490 | 71.884671 | 53.384296 | 31.628421 | 15.847407 | 8.313477 | 3.790782 | 1.233454 | 0.443815 | 0.238986 | 0.225101 | 0.306307 | 0.445105 | 0.574372 | 0.620987 | 0.742871 | 0.791701 | 0.769460 | 0.710353 | 0.642846 | 0.485678 | 0.372290 | 0.298363 | 0.263455 | 0.243730 | 0.242266 | 0.242603 | 0.246938 | 0.255770 | 0.253418 | 0.242361 | 0.228928 | 0.218083 | 0.215585 | 0.229833 | 0.271198 | 0.339509 | 0.487586 | 0.826190 | 1.359848 | 2.189106 | 3.413653 | 5.308962 | 7.480545 | 12.161161 | 15.373021 | 19.805765 | 21.212982 | 21.471593 | 18.459592 | 17.203054 | 13.500054 | 10.427690 | 7.547813 | 5.434061 | 3.969878 | 3.430620 | 3.845092 | 4.560011 | 4.208331 | 3.297854 | 2.006346 | 1.074366 | 0.586598 | 0.380846 | 0.296646 | 0.262862 | 0.243631 | 0.234174 | 0.234357 | 0.244121 | 0.252027 | 0.261480 | 0.272281 | 0.283692 | 0.275351 | 0.255306 | 0.227402 | 0.215353 | 0.220761 | 0.250602 | 0.313532 | 0.356316 | 0.389771 | 0.374513 | 0.356199 | 0.338580 | 0.338607 | 0.324146 | 0.318685 | 0.322694 | 0.316543 | 0.312623 | 0.330139 | 0.365714 | 0.385099 | 0.387881 | 0.374144 | 0.353577 | 0.313648 | 0.279900 | 0.252781 | 0.234418 | 0.220863 | 0.215323 | 0.215422 | 0.220295 | 0.225346 | 0.226528 | 0.222307 | 0.215636 | 0.216565 | 0.231485 | 0.274476 | 0.348421 | 0.423183 | 0.488187 | 0.486998 | 0.397625 | 0.323426 | 0.268552 | 0.234108 | 0.219136 | 0.215012 | 0.215274 | 0.219579 | 0.232890 | 0.258312 | 0.281813 | 0.304767 | 0.303459 | 0.290012 | 0.276680 | 0.283887 | 0.301183 | 0.347049 | 0.386742 | 0.461643 | 0.567679 | 0.694354 | 0.833041 | 0.882521 | 0.759451 | 0.583198 | 0.433762 | 0.326615 | 0.268423 | 0.230134 | 0.219808 | 0.219990 | 0.232994 | 0.269180 | 0.371215 | 0.456070 | 0.506357 | 0.500545 | 0.476238 | 0.408292 | 0.366500 | 0.329747 | 0.309546 | 0.314669 | 0.347578 | 0.398182 | 0.486411 | 0.584266 | 0.659033 | 0.675116 | 0.623775 | 0.552475 | 0.514623 | 0.493910 | 0.494171 | 0.510158 | 0.529356 | 0.559938 | 0.533631 | 0.497073 | 0.434530 | 0.357980 | 0.278702 | 0.235269 | 0.215661 | 0.220153 | 0.245598 | 0.277245 | 0.302049 | 0.284120 | 0.257985 | 0.243276 | 0.241821 | 0.246797 | 0.271921 | 0.328965 | 0.434826 | 0.572684 | 0.697257 | 0.887962 | 1.029420 | 1.071124 | 1.023285 | 0.995057 | 0.875541 | 0.716125 | 0.591761 | 0.516443 |

H) angry vs sad

  
|  | time window | peak latency | cluster *p* | peak Cohen's *d* |  | | | |  |
| **all electrodes** |  | | | |  | | | |  |
|  | | | | | | | | |  |

Model correlations, cluster permutation tests

|  | **left hemisphere** | | | | **right hemisphere** | | | |  |
|  | time window | peak latency | cluster *p* | peak Cohen's *d* | time window | peak latency | cluster *p* | peak Cohen's *d* |  |
| **anterior** |  | | | |  | | | |  |
| **central** | 185 - 280 ms | 240 ms | 0.0198 | 0.986 | 215 - 495 ms | 240 ms | 0.0007 | 0.8443 |  |
 335 - 475 ms | 420 ms | 0.015 | 0.6436 |  | | | ||  | **posterior** |  | | | |  | | | |

  

Model correlations, Bayesian statistics

|  | -200 | -195 | -190 | -185 | -180 | -175 | -170 | -165 | -160 | -155 | -150 | -145 | -140 | -135 | -130 | -125 | -120 | -115 | -110 | -105 | -100 | -95 | -90 | -85 | -80 | -75 | -70 | -65 | -60 | -55 | -50 | -45 | -40 | -35 | -30 | -25 | -20 | -15 | -10 | -5 | 0 | 5 | 10 | 15 | 20 | 25 | 30 | 35 | 40 | 45 | 50 | 55 | 60 | 65 | 70 | 75 | 80 | 85 | 90 | 95 | 100 | 105 | 110 | 115 | 120 | 125 | 130 | 135 | 140 | 145 | 150 | 155 | 160 | 165 | 170 | 175 | 180 | 185 | 190 | 195 | 200 | 205 | 210 | 215 | 220 | 225 | 230 | 235 | 240 | 245 | 250 | 255 | 260 | 265 | 270 | 275 | 280 | 285 | 290 | 295 | 300 | 305 | 310 | 315 | 320 | 325 | 330 | 335 | 340 | 345 | 350 | 355 | 360 | 365 | 370 | 375 | 380 | 385 | 390 | 395 | 400 | 405 | 410 | 415 | 420 | 425 | 430 | 435 | 440 | 445 | 450 | 455 | 460 | 465 | 470 | 475 | 480 | 485 | 490 | 495 | 500 | 505 | 510 | 515 | 520 | 525 | 530 | 535 | 540 | 545 | 550 | 555 | 560 | 565 | 570 | 575 | 580 | 585 | 590 | 595 | 600 | 605 | 610 | 615 | 620 | 625 | 630 | 635 | 640 | 645 | 650 | 655 | 660 | 665 | 670 | 675 | 680 | 685 | 690 | 695 | 700 | 705 | 710 | 715 | 720 | 725 | 730 | 735 | 740 | 745 | 750 | 755 | 760 | 765 | 770 | 775 | 780 | 785 | 790 | 795 | 800 | 805 | 810 | 815 | 820 | 825 | 830 | 835 | 840 | 845 | 850 | 855 | 860 | 865 | 870 | 875 | 880 | 885 | 890 | 895 | 900 | 905 | 910 | 915 | 920 | 925 | 930 | 935 | 940 | 945 | 950 | 955 | 960 | 965 | 970 | 975 | 980 | 985 | 990 | 995 | 1000 | 1005 | 1010 | 1015 | 1020 | 1025 | 1030 | 1035 | 1040 | 1045 | 1050 | 1055 | 1060 | 1065 | 1070 | 1075 | 1080 | 1085 | 1090 | 1095 | 1100 | 1105 | 1110 | 1115 | 1120 | 1125 | 1130 | 1135 | 1140 | 1145 | 1150 | 1155 | 1160 | 1165 | 1170 | 1175 | 1180 | 1185 | 1190 | 1195 |
| --- | --- | --- | --- | --- | --- | --- | --- | --- | --- | --- | --- | --- | --- | --- | --- | --- | --- | --- | --- | --- | --- | --- | --- | --- | --- | --- | --- | --- | --- | --- | --- | --- | --- | --- | --- | --- | --- | --- | --- | --- | --- | --- | --- | --- | --- | --- | --- | --- | --- | --- | --- | --- | --- | --- | --- | --- | --- | --- | --- | --- | --- | --- | --- | --- | --- | --- | --- | --- | --- | --- | --- | --- | --- | --- | --- | --- | --- | --- | --- | --- | --- | --- | --- | --- | --- | --- | --- | --- | --- | --- | --- | --- | --- | --- | --- | --- | --- | --- | --- | --- | --- | --- | --- | --- | --- | --- | --- | --- | --- | --- | --- | --- | --- | --- | --- | --- | --- | --- | --- | --- | --- | --- | --- | --- | --- | --- | --- | --- | --- | --- | --- | --- | --- | --- | --- | --- | --- | --- | --- | --- | --- | --- | --- | --- | --- | --- | --- | --- | --- | --- | --- | --- | --- | --- | --- | --- | --- | --- | --- | --- | --- | --- | --- | --- | --- | --- | --- | --- | --- | --- | --- | --- | --- | --- | --- | --- | --- | --- | --- | --- | --- | --- | --- | --- | --- | --- | --- | --- | --- | --- | --- | --- | --- | --- | --- | --- | --- | --- | --- | --- | --- | --- | --- | --- | --- | --- | --- | --- | --- | --- | --- | --- | --- | --- | --- | --- | --- | --- | --- | --- | --- | --- | --- | --- | --- | --- | --- | --- | --- | --- | --- | --- | --- | --- | --- | --- | --- | --- | --- | --- | --- | --- | --- | --- | --- | --- | --- | --- | --- | --- | --- | --- | --- | --- | --- | --- | --- | --- | --- | --- | --- | --- | --- | --- | --- | --- | --- | --- | --- | --- | --- | --- | --- | --- | --- | --- | --- | --- | --- | --- |
| left anterior | 0.229075 | 0.238616 | 0.238677 | 0.231521 | 0.217165 | 0.220035 | 0.253915 | 0.272056 | 0.251024 | 0.232048 | 0.216515 | 0.225372 | 0.292751 | 0.481443 | 0.867296 | 1.032334 | 0.952872 | 0.715437 | 0.424263 | 0.308692 | 0.254167 | 0.216862 | 0.215737 | 0.241119 | 0.305048 | 0.302599 | 0.334544 | 0.411669 | 0.515512 | 0.618529 | 0.544719 | 0.291291 | 0.215435 | 0.273585 | 0.405942 | 0.527741 | 0.560681 | 0.507568 | 0.398455 | 0.361079 | 0.319668 | 0.310490 | 0.327281 | 0.376031 | 0.394723 | 0.360616 | 0.269438 | 0.222176 | 0.221468 | 0.290160 | 0.336285 | 0.305291 | 0.227074 | 0.244644 | 0.567350 | 1.454725 | 3.135467 | 3.563257 | 3.721637 | 3.489197 | 2.270580 | 1.712703 | 1.738760 | 1.395744 | 0.858492 | 0.612543 | 0.423634 | 0.340816 | 0.317537 | 0.356002 | 0.472841 | 0.845332 | 1.440038 | 2.128931 | 1.829010 | 1.044461 | 0.534512 | 0.394567 | 0.382245 | 0.527244 | 0.984482 | 1.648308 | 1.886596 | 1.384563 | 0.805198 | 0.481276 | 0.370860 | 0.314638 | 0.275708 | 0.240714 | 0.233917 | 0.230132 | 0.227845 | 0.223379 | 0.223495 | 0.228951 | 0.244115 | 0.265611 | 0.316762 | 0.463427 | 0.745103 | 1.329812 | 2.423843 | 3.436547 | 3.153283 | 2.606105 | 1.463661 | 0.796551 | 0.491288 | 0.437958 | 0.412480 | 0.421786 | 0.408663 | 0.408732 | 0.322582 | 0.249242 | 0.214867 | 0.229179 | 0.276464 | 0.327263 | 0.352126 | 0.343526 | 0.324624 | 0.289287 | 0.259394 | 0.241887 | 0.236303 | 0.238972 | 0.243811 | 0.253183 | 0.259934 | 0.248513 | 0.221948 | 0.215158 | 0.240171 | 0.309302 | 0.429376 | 0.572510 | 0.599478 | 0.600846 | 0.577579 | 0.563161 | 0.571763 | 0.651632 | 0.682879 | 0.724322 | 0.641901 | 0.530681 | 0.424998 | 0.344024 | 0.275241 | 0.243418 | 0.226104 | 0.217840 | 0.215156 | 0.216321 | 0.216456 | 0.216207 | 0.217283 | 0.218402 | 0.217058 | 0.215808 | 0.215082 | 0.214827 | 0.214724 | 0.214739 | 0.215149 | 0.216664 | 0.220990 | 0.223143 | 0.217649 | 0.215604 | 0.237407 | 0.279057 | 0.322436 | 0.343214 | 0.311359 | 0.268571 | 0.238096 | 0.220249 | 0.214698 | 0.216021 | 0.217823 | 0.215107 | 0.216472 | 0.234040 | 0.304331 | 0.493389 | 0.682328 | 0.871323 | 1.162928 | 1.756238 | 2.902210 | 5.821761 | 10.470498 | 11.935713 | 6.810493 | 2.898280 | 1.189563 | 0.641730 | 0.430049 | 0.375774 | 0.407801 | 0.563075 | 0.790866 | 1.205714 | 1.705229 | 2.509542 | 3.781474 | 6.945355 | 16.178423 | 48.402552 | 90.294527 | 109.645900 | 70.911186 | 35.739709 | 15.396645 | 6.984263 | 3.390775 | 1.922323 | 1.299901 | 0.947810 | 0.781664 | 0.663687 | 0.652531 | 0.697332 | 0.859246 | 1.000249 | 1.115984 | 1.124449 | 1.122624 | 1.057394 | 1.010191 | 0.959032 | 0.949445 | 0.946804 | 0.959484 | 1.043826 | 1.063394 | 0.973240 | 0.835609 | 0.679075 | 0.539185 | 0.500121 | 0.480734 | 0.494329 | 0.544678 | 0.588083 | 0.606717 | 0.612946 | 0.626205 | 0.663740 | 0.777636 | 0.915768 | 1.206820 | 1.616586 | 2.081209 | 2.303412 | 2.178331 | 1.538980 | 1.012826 | 0.638004 | 0.459499 | 0.359630 | 0.322431 | 0.306973 | 0.310351 | 0.334548 | 0.385452 | 0.498936 | 0.711524 | 0.967300 | 1.170077 | 1.322170 | 1.126148 | 0.844768 | 0.585172 | 0.403883 | 0.335860 | 0.291822 |
| right anterior | 0.262060 | 0.239128 | 0.221422 | 0.214803 | 0.223067 | 0.244077 | 0.260812 | 0.266506 | 0.249816 | 0.237431 | 0.242112 | 0.256273 | 0.255603 | 0.284578 | 0.365939 | 0.414628 | 0.409395 | 0.344671 | 0.268903 | 0.223740 | 0.214859 | 0.229063 | 0.255724 | 0.298216 | 0.313385 | 0.341759 | 0.335281 | 0.289986 | 0.238409 | 0.222615 | 0.215080 | 0.223447 | 0.229956 | 0.231764 | 0.228389 | 0.218444 | 0.214879 | 0.214643 | 0.217535 | 0.217455 | 0.214955 | 0.214868 | 0.216978 | 0.220647 | 0.215726 | 0.216120 | 0.233668 | 0.295928 | 0.417149 | 0.531436 | 0.493204 | 0.364239 | 0.265488 | 0.223142 | 0.214779 | 0.217921 | 0.223301 | 0.244348 | 0.298204 | 0.454353 | 0.760575 | 1.618705 | 3.108511 | 6.074944 | 4.177613 | 1.896516 | 0.925198 | 0.612529 | 0.499292 | 0.648648 | 0.993217 | 1.875625 | 3.008161 | 4.285444 | 4.498405 | 5.369033 | 6.293540 | 9.022842 | 14.566129 | 17.313888 | 16.579416 | 14.108436 | 7.552091 | 2.879681 | 1.239637 | 0.594803 | 0.365473 | 0.290450 | 0.258356 | 0.247995 | 0.249231 | 0.240853 | 0.228380 | 0.222855 | 0.217248 | 0.214661 | 0.214707 | 0.215831 | 0.220469 | 0.239693 | 0.270896 | 0.305344 | 0.325546 | 0.292917 | 0.256589 | 0.246629 | 0.235403 | 0.230604 | 0.232060 | 0.231957 | 0.223928 | 0.214850 | 0.220463 | 0.245029 | 0.286450 | 0.338359 | 0.353509 | 0.336282 | 0.303220 | 0.292346 | 0.282102 | 0.276382 | 0.275601 | 0.274377 | 0.274428 | 0.274949 | 0.267405 | 0.266029 | 0.269744 | 0.269241 | 0.257999 | 0.236158 | 0.216350 | 0.216480 | 0.231495 | 0.261008 | 0.288118 | 0.299625 | 0.349176 | 0.455394 | 0.611408 | 0.865965 | 1.161255 | 1.047128 | 0.939197 | 0.752676 | 0.558610 | 0.466959 | 0.400890 | 0.317904 | 0.285498 | 0.269980 | 0.257448 | 0.256143 | 0.267891 | 0.295256 | 0.349022 | 0.470323 | 0.764813 | 1.271994 | 1.732596 | 2.401108 | 4.191871 | 8.739681 | 18.749274 | 25.901438 | 29.758718 | 32.943376 | 44.905680 | 36.344382 | 25.672320 | 10.847259 | 4.389617 | 1.797314 | 1.055630 | 0.721177 | 0.601568 | 0.587090 | 0.696162 | 0.867798 | 1.174743 | 1.839431 | 3.139618 | 4.207078 | 5.228670 | 4.334279 | 2.915657 | 1.896592 | 1.393318 | 0.986340 | 0.844462 | 0.714184 | 0.599209 | 0.495581 | 0.426744 | 0.366099 | 0.319456 | 0.301144 | 0.316367 | 0.378210 | 0.554703 | 1.102571 | 2.398900 | 4.633875 | 5.608532 | 5.019282 | 3.575220 | 2.408602 | 1.552634 | 0.928805 | 0.607051 | 0.441427 | 0.348942 | 0.305882 | 0.297518 | 0.317362 | 0.413676 | 0.651851 | 1.130223 | 1.565372 | 1.549504 | 1.052281 | 0.659205 | 0.437192 | 0.359497 | 0.358236 | 0.380722 | 0.413372 | 0.435312 | 0.427536 | 0.374024 | 0.303684 | 0.261084 | 0.233488 | 0.220659 | 0.215744 | 0.214933 | 0.214997 | 0.215174 | 0.215748 | 0.226574 | 0.256911 | 0.317414 | 0.419462 | 0.500415 | 0.512836 | 0.443433 | 0.342953 | 0.263054 | 0.221798 | 0.215210 | 0.221194 | 0.224930 | 0.229204 | 0.227599 | 0.224478 | 0.224089 | 0.224269 | 0.225054 | 0.230633 | 0.227878 | 0.223327 | 0.217205 | 0.215036 | 0.214699 | 0.215078 | 0.217065 | 0.223730 | 0.236346 | 0.255610 | 0.283477 | 0.321977 | 0.375585 | 0.418469 | 0.434903 | 0.430660 | 0.382525 | 0.350241 | 0.331261 | 0.311196 |
| left central | 0.234985 | 0.231680 | 0.229125 | 0.223137 | 0.221732 | 0.228227 | 0.244514 | 0.255032 | 0.294070 | 0.380528 | 0.439675 | 0.456757 | 0.452010 | 0.384358 | 0.329529 | 0.300645 | 0.294201 | 0.331123 | 0.388700 | 0.415237 | 0.531237 | 0.848548 | 1.724611 | 2.304340 | 2.675816 | 2.215661 | 1.646836 | 0.653789 | 0.292996 | 0.215144 | 0.266770 | 0.359934 | 0.368647 | 0.276860 | 0.220851 | 0.219954 | 0.243497 | 0.258222 | 0.262660 | 0.267315 | 0.239372 | 0.218606 | 0.220758 | 0.293398 | 0.545119 | 0.870125 | 0.962267 | 0.667800 | 0.508787 | 0.425829 | 0.358655 | 0.303385 | 0.251252 | 0.217422 | 0.218520 | 0.239207 | 0.272765 | 0.296496 | 0.303983 | 0.292344 | 0.271071 | 0.255722 | 0.251504 | 0.249871 | 0.246245 | 0.247947 | 0.231299 | 0.218257 | 0.215084 | 0.215046 | 0.216152 | 0.229610 | 0.250924 | 0.289846 | 0.356973 | 0.497056 | 0.886392 | 2.334553 | 8.196729 | 22.151973 | 30.621772 | 27.032119 | 23.632655 | 25.671785 | 36.586798 | 69.268068 | 140.095144 | 265.670903 | 305.222244 | 178.724762 | 98.827250 | 54.821804 | 34.679318 | 18.603646 | 9.157395 | 3.863664 | 1.589498 | 0.643206 | 0.352668 | 0.249814 | 0.219830 | 0.214796 | 0.214631 | 0.216758 | 0.242698 | 0.344486 | 0.636990 | 1.581184 | 3.278659 | 4.306946 | 4.728604 | 4.263490 | 3.301566 | 2.930939 | 2.585399 | 2.750020 | 3.065491 | 3.945155 | 4.822926 | 6.571451 | 7.649972 | 7.819570 | 7.588539 | 7.188631 | 6.743415 | 6.285078 | 5.099527 | 4.049542 | 3.280641 | 2.615301 | 2.847734 | 3.784991 | 5.150234 | 5.766795 | 3.955549 | 1.859401 | 1.062676 | 0.696074 | 0.460450 | 0.324672 | 0.243749 | 0.214703 | 0.252531 | 0.379339 | 0.600926 | 0.743624 | 0.559757 | 0.356221 | 0.242614 | 0.214676 | 0.241072 | 0.294687 | 0.337736 | 0.363306 | 0.392198 | 0.392978 | 0.376639 | 0.362012 | 0.330275 | 0.282942 | 0.238750 | 0.216216 | 0.223182 | 0.268040 | 0.341747 | 0.421117 | 0.479356 | 0.456604 | 0.356558 | 0.260381 | 0.216396 | 0.226989 | 0.268814 | 0.296476 | 0.270559 | 0.234559 | 0.216966 | 0.216873 | 0.226904 | 0.234602 | 0.234905 | 0.227104 | 0.218402 | 0.215579 | 0.215202 | 0.216822 | 0.222566 | 0.238382 | 0.257816 | 0.263585 | 0.253548 | 0.243462 | 0.239414 | 0.241242 | 0.245104 | 0.256928 | 0.288552 | 0.333705 | 0.391938 | 0.449175 | 0.500763 | 0.527603 | 0.554120 | 0.593636 | 0.651970 | 0.707799 | 0.711989 | 0.654044 | 0.561797 | 0.467278 | 0.375562 | 0.329636 | 0.304688 | 0.291641 | 0.274100 | 0.258762 | 0.238717 | 0.224041 | 0.214846 | 0.219281 | 0.235498 | 0.247722 | 0.253442 | 0.256462 | 0.260779 | 0.274121 | 0.314073 | 0.394677 | 0.521564 | 0.706500 | 0.906502 | 1.038858 | 1.029584 | 0.932745 | 0.778341 | 0.635926 | 0.523053 | 0.479853 | 0.469727 | 0.485638 | 0.522622 | 0.576381 | 0.586302 | 0.604395 | 0.542633 | 0.441578 | 0.347422 | 0.280502 | 0.236017 | 0.218550 | 0.214696 | 0.214627 | 0.214661 | 0.214646 | 0.214642 | 0.214628 | 0.214643 | 0.214628 | 0.214646 | 0.214694 | 0.214816 | 0.214900 | 0.216955 | 0.221146 | 0.231273 | 0.243051 | 0.260585 | 0.276222 | 0.279768 | 0.260241 | 0.245109 | 0.242775 | 0.237696 | 0.253746 | 0.268829 | 0.287074 | 0.312045 | 0.346035 | 0.331540 | 0.346242 |
| right central | 2.196799 | 1.818113 | 1.650384 | 1.537046 | 1.490423 | 1.416406 | 1.242413 | 1.230829 | 0.836961 | 0.790272 | 0.843903 | 0.681450 | 0.382934 | 0.272737 | 0.227977 | 0.220427 | 0.219392 | 0.215833 | 0.215046 | 0.223770 | 0.259801 | 0.360797 | 0.453929 | 0.517048 | 0.458745 | 0.416180 | 0.342862 | 0.286191 | 0.234978 | 0.216049 | 0.214654 | 0.220807 | 0.236154 | 0.266080 | 0.316073 | 0.355614 | 0.375121 | 0.375211 | 0.326404 | 0.296350 | 0.291392 | 0.289252 | 0.299607 | 0.345520 | 0.366577 | 0.352854 | 0.321580 | 0.309496 | 0.291323 | 0.282423 | 0.257085 | 0.226205 | 0.214989 | 0.235734 | 0.327416 | 0.624660 | 1.567352 | 4.399370 | 6.906840 | 2.092946 | 0.603722 | 0.295919 | 0.236517 | 0.219825 | 0.223246 | 0.236745 | 0.286810 | 0.378334 | 0.521447 | 0.643018 | 0.651042 | 0.595250 | 0.506733 | 0.427659 | 0.332029 | 0.287303 | 0.249275 | 0.235174 | 0.225958 | 0.218070 | 0.219103 | 0.299148 | 0.918888 | 4.897276 | 27.799813 | 128.555343 | 237.764883 | 176.325065 | 84.996281 | 35.931069 | 16.838919 | 9.934577 | 7.145586 | 5.413565 | 4.017963 | 2.914028 | 2.121906 | 1.458304 | 1.194422 | 1.076129 | 1.184202 | 1.587950 | 2.390892 | 3.379405 | 4.589563 | 5.295430 | 5.653619 | 6.711309 | 10.099133 | 18.937451 | 32.157639 | 47.296953 | 57.774307 | 77.631147 | 70.908709 | 55.586872 | 31.614407 | 20.068532 | 13.342709 | 12.690089 | 15.738557 | 26.928792 | 41.886651 | 58.116249 | 77.538274 | 106.260615 | 113.019065 | 100.596310 | 91.048025 | 86.826388 | 94.125370 | 177.964050 | 526.007286 | 1651.822430 | 2633.394427 | 989.987653 | 160.705226 | 22.711781 | 4.880305 | 1.613526 | 0.621060 | 0.326859 | 0.252998 | 0.229522 | 0.223453 | 0.229125 | 0.249603 | 0.312077 | 0.448158 | 0.650050 | 0.874504 | 1.133515 | 1.094228 | 0.974658 | 0.884006 | 0.843306 | 0.927859 | 1.244229 | 1.778684 | 2.356318 | 2.305561 | 1.604769 | 0.942610 | 0.532204 | 0.362197 | 0.301661 | 0.293350 | 0.338990 | 0.489410 | 0.739367 | 1.088611 | 1.278064 | 1.209704 | 0.998956 | 0.817669 | 0.690237 | 0.707258 | 0.708042 | 0.670576 | 0.665792 | 0.604062 | 0.530850 | 0.516819 | 0.541964 | 0.604824 | 0.757379 | 0.876471 | 0.949549 | 0.967397 | 0.885500 | 0.671510 | 0.542283 | 0.450865 | 0.347431 | 0.282143 | 0.244034 | 0.217487 | 0.218030 | 0.232953 | 0.235459 | 0.217430 | 0.225265 | 0.292460 | 0.442672 | 0.622362 | 0.589435 | 0.401723 | 0.277065 | 0.222886 | 0.218162 | 0.249889 | 0.284122 | 0.288965 | 0.278679 | 0.270721 | 0.253663 | 0.233820 | 0.223122 | 0.217886 | 0.215181 | 0.214641 | 0.214708 | 0.214732 | 0.214951 | 0.216379 | 0.220530 | 0.227296 | 0.240234 | 0.258323 | 0.272719 | 0.278333 | 0.272438 | 0.258598 | 0.255470 | 0.264397 | 0.275833 | 0.297560 | 0.323744 | 0.333987 | 0.320119 | 0.297926 | 0.259446 | 0.227269 | 0.214718 | 0.225353 | 0.257600 | 0.288458 | 0.297287 | 0.273653 | 0.236554 | 0.221909 | 0.215550 | 0.215031 | 0.218447 | 0.224150 | 0.237845 | 0.251681 | 0.276865 | 0.305172 | 0.321449 | 0.339556 | 0.357460 | 0.330908 | 0.303102 | 0.292637 | 0.290149 | 0.282655 | 0.276709 | 0.265658 | 0.248750 | 0.235021 | 0.223240 | 0.216510 | 0.214745 | 0.219550 | 0.242077 | 0.274291 | 0.312355 | 0.334788 | 0.362504 |
| left posterior | 1.911067 | 2.066497 | 1.741830 | 1.214298 | 0.703139 | 0.480420 | 0.378989 | 0.345729 | 0.301463 | 0.260248 | 0.226532 | 0.217071 | 0.256551 | 0.302793 | 0.318231 | 0.273642 | 0.215963 | 0.265453 | 0.478282 | 0.683621 | 0.515706 | 0.290439 | 0.217107 | 0.492799 | 1.944375 | 3.384253 | 2.414671 | 0.963303 | 0.398837 | 0.261003 | 0.232965 | 0.234105 | 0.276099 | 0.386426 | 0.683647 | 1.298699 | 2.000739 | 2.376315 | 2.244135 | 1.263629 | 0.407211 | 0.215720 | 0.389020 | 0.752334 | 0.981113 | 1.007194 | 0.933949 | 0.790116 | 0.635932 | 0.548057 | 0.482002 | 0.441061 | 0.396948 | 0.356904 | 0.303918 | 0.264473 | 0.247056 | 0.264015 | 0.364869 | 0.714599 | 1.240689 | 1.444744 | 1.149506 | 0.706917 | 0.432753 | 0.301670 | 0.243354 | 0.221574 | 0.215325 | 0.214659 | 0.214691 | 0.214828 | 0.220221 | 0.243642 | 0.312692 | 0.433839 | 0.479195 | 0.366253 | 0.258615 | 0.217586 | 0.219781 | 0.234167 | 0.237331 | 0.227392 | 0.217358 | 0.215961 | 0.231462 | 0.268172 | 0.317132 | 0.358519 | 0.411412 | 0.471237 | 0.528793 | 0.600724 | 0.672949 | 0.676055 | 0.676436 | 0.636404 | 0.543582 | 0.484985 | 0.463656 | 0.484851 | 0.551287 | 0.628455 | 0.653137 | 0.629443 | 0.530534 | 0.427892 | 0.362882 | 0.307247 | 0.271844 | 0.247576 | 0.228380 | 0.217301 | 0.214660 | 0.215345 | 0.215716 | 0.215934 | 0.216421 | 0.215707 | 0.216079 | 0.216678 | 0.215296 | 0.214653 | 0.216741 | 0.237351 | 0.262217 | 0.275110 | 0.276053 | 0.255320 | 0.229644 | 0.218159 | 0.214628 | 0.215600 | 0.215834 | 0.215382 | 0.214628 | 0.216529 | 0.223137 | 0.236320 | 0.258093 | 0.296395 | 0.330004 | 0.345052 | 0.330007 | 0.318706 | 0.311152 | 0.323187 | 0.366209 | 0.467888 | 0.686859 | 1.096042 | 1.681822 | 2.269610 | 2.656698 | 2.246492 | 1.511578 | 0.990619 | 0.668756 | 0.508354 | 0.503684 | 0.666138 | 1.078234 | 2.032708 | 3.098004 | 3.402138 | 3.304267 | 2.738238 | 1.944788 | 1.495823 | 1.085810 | 0.784791 | 0.623553 | 0.529923 | 0.447672 | 0.421929 | 0.429101 | 0.475192 | 0.585241 | 0.806379 | 1.087327 | 1.496572 | 2.046670 | 2.573862 | 2.909869 | 3.028613 | 2.671755 | 2.122335 | 1.647774 | 1.499003 | 1.593014 | 2.139538 | 3.090440 | 4.766171 | 5.836469 | 4.746215 | 2.964676 | 1.713136 | 0.926334 | 0.581903 | 0.457267 | 0.414442 | 0.417827 | 0.430224 | 0.450776 | 0.444782 | 0.429572 | 0.391492 | 0.364681 | 0.345098 | 0.342172 | 0.324105 | 0.295229 | 0.264330 | 0.241361 | 0.217563 | 0.217436 | 0.237007 | 0.265094 | 0.293605 | 0.284289 | 0.257674 | 0.228248 | 0.214762 | 0.225759 | 0.265298 | 0.328112 | 0.394200 | 0.414387 | 0.394623 | 0.369077 | 0.342958 | 0.313710 | 0.289386 | 0.261710 | 0.236521 | 0.221234 | 0.215063 | 0.214810 | 0.215724 | 0.214780 | 0.215348 | 0.219211 | 0.221034 | 0.223566 | 0.222328 | 0.218096 | 0.214936 | 0.214964 | 0.216686 | 0.219097 | 0.221579 | 0.225829 | 0.231436 | 0.233752 | 0.236307 | 0.230443 | 0.222639 | 0.216722 | 0.214671 | 0.221156 | 0.232993 | 0.248876 | 0.266679 | 0.278137 | 0.285875 | 0.293630 | 0.302709 | 0.322486 | 0.369261 | 0.416380 | 0.454594 | 0.542008 | 0.602250 | 0.634974 | 0.648567 | 0.666086 | 0.655731 | 0.665814 | 0.627379 |
| right posterior | 7.223216 | 8.503882 | 9.384345 | 10.494381 | 6.574725 | 3.812279 | 1.923723 | 0.965891 | 0.505291 | 0.385470 | 0.368473 | 0.364825 | 0.415639 | 0.604715 | 0.847893 | 0.986339 | 0.916020 | 0.604682 | 0.390341 | 0.253968 | 0.215458 | 0.268222 | 0.344066 | 0.406045 | 0.394351 | 0.325882 | 0.249225 | 0.217790 | 0.215411 | 0.221937 | 0.229990 | 0.238305 | 0.242742 | 0.236432 | 0.234831 | 0.256035 | 0.290541 | 0.319178 | 0.370364 | 0.449626 | 0.544099 | 0.630863 | 0.761455 | 0.807764 | 0.894474 | 0.754002 | 0.656281 | 0.696346 | 0.955981 | 1.101605 | 0.974741 | 0.565043 | 0.394978 | 0.351587 | 0.376264 | 0.464870 | 0.711168 | 1.377330 | 3.007380 | 5.215951 | 6.366282 | 7.682425 | 6.386935 | 3.292366 | 1.305922 | 0.541736 | 0.293097 | 0.228891 | 0.215887 | 0.214706 | 0.214794 | 0.221471 | 0.233803 | 0.251658 | 0.286091 | 0.305248 | 0.289431 | 0.256192 | 0.223288 | 0.214974 | 0.223352 | 0.241022 | 0.255019 | 0.263707 | 0.267110 | 0.267844 | 0.254051 | 0.242759 | 0.238399 | 0.245980 | 0.263675 | 0.289672 | 0.315295 | 0.311451 | 0.287068 | 0.250510 | 0.225081 | 0.214630 | 0.219493 | 0.223898 | 0.219578 | 0.215216 | 0.216689 | 0.231239 | 0.257228 | 0.290837 | 0.331842 | 0.377069 | 0.406464 | 0.417107 | 0.413349 | 0.393423 | 0.370891 | 0.365533 | 0.355951 | 0.363974 | 0.390962 | 0.453000 | 0.534358 | 0.649433 | 0.724745 | 0.675914 | 0.492568 | 0.318331 | 0.231777 | 0.216118 | 0.236212 | 0.247487 | 0.235362 | 0.218800 | 0.215570 | 0.235845 | 0.277270 | 0.311043 | 0.336341 | 0.342522 | 0.323817 | 0.302834 | 0.279871 | 0.257629 | 0.237976 | 0.224395 | 0.217896 | 0.215818 | 0.214643 | 0.215342 | 0.217994 | 0.220174 | 0.219785 | 0.216404 | 0.214634 | 0.221623 | 0.252758 | 0.331835 | 0.493702 | 0.850755 | 1.476083 | 2.407535 | 3.540926 | 4.228494 | 3.688660 | 2.647238 | 1.504678 | 0.773207 | 0.473396 | 0.349658 | 0.292858 | 0.276714 | 0.288385 | 0.311051 | 0.342280 | 0.369642 | 0.393802 | 0.410880 | 0.403749 | 0.387552 | 0.355904 | 0.311153 | 0.267026 | 0.235849 | 0.217509 | 0.215136 | 0.223466 | 0.238609 | 0.249369 | 0.253067 | 0.248760 | 0.238392 | 0.230832 | 0.226819 | 0.227997 | 0.239678 | 0.252948 | 0.265748 | 0.263350 | 0.242798 | 0.219351 | 0.215225 | 0.230919 | 0.249403 | 0.263224 | 0.263345 | 0.254120 | 0.237303 | 0.225558 | 0.215986 | 0.215316 | 0.222918 | 0.241177 | 0.279758 | 0.355599 | 0.469382 | 0.542209 | 0.498557 | 0.387367 | 0.270477 | 0.216435 | 0.231869 | 0.297965 | 0.400042 | 0.461779 | 0.426090 | 0.335859 | 0.258268 | 0.222767 | 0.214644 | 0.220867 | 0.227123 | 0.227529 | 0.225155 | 0.219641 | 0.214644 | 0.220360 | 0.252469 | 0.324494 | 0.439610 | 0.557082 | 0.620825 | 0.583702 | 0.530829 | 0.470622 | 0.408727 | 0.348015 | 0.293471 | 0.248894 | 0.223535 | 0.214630 | 0.216834 | 0.216524 | 0.214731 | 0.216536 | 0.228140 | 0.249067 | 0.269473 | 0.280393 | 0.279620 | 0.283457 | 0.290549 | 0.310671 | 0.346414 | 0.386596 | 0.393176 | 0.384845 | 0.349458 | 0.323776 | 0.305190 | 0.295190 | 0.284261 | 0.280235 | 0.267106 | 0.254867 | 0.243678 | 0.230947 | 0.222932 | 0.218285 | 0.215538 | 0.214687 | 0.214943 | 0.215924 | 0.217138 |
| all electrodes | 0.214658 | 0.222013 | 0.250137 | 0.334540 | 0.443083 | 0.417620 | 0.331771 | 0.271577 | 0.227338 | 0.214742 | 0.218034 | 0.222633 | 0.227450 | 0.231065 | 0.231473 | 0.237101 | 0.249479 | 0.301879 | 0.370802 | 0.427672 | 0.539892 | 0.738126 | 0.670067 | 0.630699 | 0.593000 | 0.599847 | 0.749904 | 0.816966 | 0.694301 | 0.597127 | 0.501655 | 0.387701 | 0.380670 | 0.386023 | 0.396295 | 0.411498 | 0.425134 | 0.388799 | 0.355741 | 0.327231 | 0.280532 | 0.247908 | 0.230733 | 0.225558 | 0.224353 | 0.240940 | 0.282586 | 0.334980 | 0.313249 | 0.279986 | 0.236793 | 0.215424 | 0.231631 | 0.290791 | 0.395163 | 0.520145 | 0.808963 | 1.309323 | 2.266581 | 3.160270 | 3.459905 | 2.479308 | 1.874532 | 1.173035 | 0.668702 | 0.342484 | 0.226464 | 0.225159 | 0.287288 | 0.336772 | 0.314893 | 0.247576 | 0.214659 | 0.262625 | 0.398673 | 0.521736 | 0.509473 | 0.375789 | 0.264006 | 0.217425 | 0.224893 | 0.274296 | 0.364212 | 0.475015 | 0.601826 | 0.643209 | 0.611304 | 0.560173 | 0.501768 | 0.389469 | 0.341334 | 0.287903 | 0.248817 | 0.232512 | 0.232985 | 0.230918 | 0.233457 | 0.235524 | 0.239350 | 0.242096 | 0.237480 | 0.237690 | 0.240513 | 0.239338 | 0.237362 | 0.238225 | 0.236711 | 0.244254 | 0.255444 | 0.277917 | 0.335288 | 0.428190 | 0.545860 | 0.676137 | 0.816039 | 0.835582 | 0.796785 | 0.666370 | 0.577141 | 0.549777 | 0.550208 | 0.545879 | 0.625525 | 0.776739 | 1.001618 | 1.329720 | 1.659305 | 1.589125 | 1.374797 | 1.046521 | 0.777194 | 0.617483 | 0.474806 | 0.358986 | 0.300536 | 0.271402 | 0.244459 | 0.227963 | 0.217963 | 0.214847 | 0.228178 | 0.260236 | 0.319967 | 0.342570 | 0.341338 | 0.321584 | 0.297019 | 0.259113 | 0.239347 | 0.222074 | 0.214668 | 0.219394 | 0.230738 | 0.246685 | 0.248204 | 0.242247 | 0.232437 | 0.224399 | 0.218663 | 0.215225 | 0.215084 | 0.217931 | 0.225197 | 0.240251 | 0.267599 | 0.311992 | 0.392960 | 0.515642 | 0.618279 | 0.630191 | 0.589613 | 0.515707 | 0.435202 | 0.369708 | 0.323600 | 0.286463 | 0.257339 | 0.236876 | 0.226813 | 0.223906 | 0.228106 | 0.239890 | 0.276921 | 0.344214 | 0.437493 | 0.505857 | 0.561836 | 0.580234 | 0.581887 | 0.517794 | 0.485236 | 0.457293 | 0.423698 | 0.406193 | 0.419520 | 0.413533 | 0.424440 | 0.407230 | 0.350900 | 0.307153 | 0.278732 | 0.262535 | 0.262634 | 0.268581 | 0.280721 | 0.323015 | 0.364621 | 0.371478 | 0.363619 | 0.351774 | 0.321374 | 0.292708 | 0.280866 | 0.284795 | 0.282940 | 0.273717 | 0.263149 | 0.252301 | 0.235462 | 0.221961 | 0.216476 | 0.214997 | 0.215049 | 0.215918 | 0.218158 | 0.222678 | 0.229742 | 0.235117 | 0.246250 | 0.261642 | 0.266594 | 0.255326 | 0.232473 | 0.218448 | 0.214638 | 0.218026 | 0.224873 | 0.227663 | 0.228679 | 0.226068 | 0.220355 | 0.217032 | 0.215232 | 0.214626 | 0.215490 | 0.216270 | 0.216491 | 0.215370 | 0.214689 | 0.215455 | 0.217220 | 0.217678 | 0.215033 | 0.215722 | 0.227916 | 0.260551 | 0.331420 | 0.428103 | 0.532049 | 0.607257 | 0.615759 | 0.492421 | 0.405808 | 0.345875 | 0.314615 | 0.310949 | 0.360557 | 0.425493 | 0.594602 | 0.872992 | 1.074083 | 1.222118 | 1.509359 | 1.468873 | 1.492742 | 1.441232 | 1.285433 | 0.944366 | 0.822211 | 0.692153 |

I) happy vs sad

  
|  | time window | peak latency | cluster *p* | peak Cohen's *d* |  | | | |  |
| **all electrodes** |  | | | |  | | | |  |
|  | | | | | | | | |  |

Model correlations, cluster permutation tests

|  | **left hemisphere** | | | | **right hemisphere** | | | |  |
|  | time window | peak latency | cluster *p* | peak Cohen's *d* | time window | peak latency | cluster *p* | peak Cohen's *d* |  |
| **anterior** |  | | | |  | | | |  |
| **central** | 250 - 385 ms | 360 ms | 0.0287 | -0.5323 |  | | | |  |
 520 - 730 ms | 690 ms | 0.0047 | -0.8102 |  | | | ||  | **posterior** |  | | | |  | | | |

  

Model correlations, Bayesian statistics

|  | -200 | -195 | -190 | -185 | -180 | -175 | -170 | -165 | -160 | -155 | -150 | -145 | -140 | -135 | -130 | -125 | -120 | -115 | -110 | -105 | -100 | -95 | -90 | -85 | -80 | -75 | -70 | -65 | -60 | -55 | -50 | -45 | -40 | -35 | -30 | -25 | -20 | -15 | -10 | -5 | 0 | 5 | 10 | 15 | 20 | 25 | 30 | 35 | 40 | 45 | 50 | 55 | 60 | 65 | 70 | 75 | 80 | 85 | 90 | 95 | 100 | 105 | 110 | 115 | 120 | 125 | 130 | 135 | 140 | 145 | 150 | 155 | 160 | 165 | 170 | 175 | 180 | 185 | 190 | 195 | 200 | 205 | 210 | 215 | 220 | 225 | 230 | 235 | 240 | 245 | 250 | 255 | 260 | 265 | 270 | 275 | 280 | 285 | 290 | 295 | 300 | 305 | 310 | 315 | 320 | 325 | 330 | 335 | 340 | 345 | 350 | 355 | 360 | 365 | 370 | 375 | 380 | 385 | 390 | 395 | 400 | 405 | 410 | 415 | 420 | 425 | 430 | 435 | 440 | 445 | 450 | 455 | 460 | 465 | 470 | 475 | 480 | 485 | 490 | 495 | 500 | 505 | 510 | 515 | 520 | 525 | 530 | 535 | 540 | 545 | 550 | 555 | 560 | 565 | 570 | 575 | 580 | 585 | 590 | 595 | 600 | 605 | 610 | 615 | 620 | 625 | 630 | 635 | 640 | 645 | 650 | 655 | 660 | 665 | 670 | 675 | 680 | 685 | 690 | 695 | 700 | 705 | 710 | 715 | 720 | 725 | 730 | 735 | 740 | 745 | 750 | 755 | 760 | 765 | 770 | 775 | 780 | 785 | 790 | 795 | 800 | 805 | 810 | 815 | 820 | 825 | 830 | 835 | 840 | 845 | 850 | 855 | 860 | 865 | 870 | 875 | 880 | 885 | 890 | 895 | 900 | 905 | 910 | 915 | 920 | 925 | 930 | 935 | 940 | 945 | 950 | 955 | 960 | 965 | 970 | 975 | 980 | 985 | 990 | 995 | 1000 | 1005 | 1010 | 1015 | 1020 | 1025 | 1030 | 1035 | 1040 | 1045 | 1050 | 1055 | 1060 | 1065 | 1070 | 1075 | 1080 | 1085 | 1090 | 1095 | 1100 | 1105 | 1110 | 1115 | 1120 | 1125 | 1130 | 1135 | 1140 | 1145 | 1150 | 1155 | 1160 | 1165 | 1170 | 1175 | 1180 | 1185 | 1190 | 1195 |
| --- | --- | --- | --- | --- | --- | --- | --- | --- | --- | --- | --- | --- | --- | --- | --- | --- | --- | --- | --- | --- | --- | --- | --- | --- | --- | --- | --- | --- | --- | --- | --- | --- | --- | --- | --- | --- | --- | --- | --- | --- | --- | --- | --- | --- | --- | --- | --- | --- | --- | --- | --- | --- | --- | --- | --- | --- | --- | --- | --- | --- | --- | --- | --- | --- | --- | --- | --- | --- | --- | --- | --- | --- | --- | --- | --- | --- | --- | --- | --- | --- | --- | --- | --- | --- | --- | --- | --- | --- | --- | --- | --- | --- | --- | --- | --- | --- | --- | --- | --- | --- | --- | --- | --- | --- | --- | --- | --- | --- | --- | --- | --- | --- | --- | --- | --- | --- | --- | --- | --- | --- | --- | --- | --- | --- | --- | --- | --- | --- | --- | --- | --- | --- | --- | --- | --- | --- | --- | --- | --- | --- | --- | --- | --- | --- | --- | --- | --- | --- | --- | --- | --- | --- | --- | --- | --- | --- | --- | --- | --- | --- | --- | --- | --- | --- | --- | --- | --- | --- | --- | --- | --- | --- | --- | --- | --- | --- | --- | --- | --- | --- | --- | --- | --- | --- | --- | --- | --- | --- | --- | --- | --- | --- | --- | --- | --- | --- | --- | --- | --- | --- | --- | --- | --- | --- | --- | --- | --- | --- | --- | --- | --- | --- | --- | --- | --- | --- | --- | --- | --- | --- | --- | --- | --- | --- | --- | --- | --- | --- | --- | --- | --- | --- | --- | --- | --- | --- | --- | --- | --- | --- | --- | --- | --- | --- | --- | --- | --- | --- | --- | --- | --- | --- | --- | --- | --- | --- | --- | --- | --- | --- | --- | --- | --- | --- | --- | --- | --- | --- | --- | --- | --- | --- | --- | --- | --- | --- | --- | --- | --- | --- |
| left anterior | 0.214637 | 0.214910 | 0.219864 | 0.248417 | 0.284563 | 0.300470 | 0.278078 | 0.244268 | 0.214629 | 0.241756 | 0.293051 | 0.325151 | 0.333474 | 0.276664 | 0.232791 | 0.214809 | 0.228225 | 0.244457 | 0.259492 | 0.285248 | 0.294201 | 0.304104 | 0.332147 | 0.449370 | 0.697106 | 1.094444 | 2.122907 | 4.147863 | 6.597803 | 10.706508 | 10.669460 | 4.813691 | 1.653123 | 0.442574 | 0.222836 | 0.230729 | 0.303160 | 0.311900 | 0.282551 | 0.292120 | 0.324354 | 0.364427 | 0.433747 | 0.427486 | 0.356784 | 0.249954 | 0.216173 | 0.300154 | 0.513379 | 0.914348 | 1.138132 | 1.103050 | 0.635236 | 0.321123 | 0.215519 | 0.241498 | 0.313946 | 0.346499 | 0.387793 | 0.415502 | 0.400959 | 0.319124 | 0.262335 | 0.222208 | 0.218039 | 0.245899 | 0.285217 | 0.353007 | 0.505440 | 0.636393 | 0.954606 | 1.684761 | 3.183182 | 5.186260 | 7.377491 | 5.496910 | 3.126324 | 1.563556 | 0.691049 | 0.372702 | 0.260000 | 0.223234 | 0.216012 | 0.218024 | 0.223736 | 0.237518 | 0.253562 | 0.268273 | 0.272280 | 0.274057 | 0.258192 | 0.236545 | 0.218779 | 0.215307 | 0.214756 | 0.215066 | 0.218878 | 0.227716 | 0.262873 | 0.337437 | 0.438167 | 0.525080 | 0.552419 | 0.480352 | 0.350225 | 0.269409 | 0.222830 | 0.214782 | 0.221097 | 0.224548 | 0.223568 | 0.216988 | 0.215003 | 0.214773 | 0.215361 | 0.221280 | 0.255604 | 0.323235 | 0.419289 | 0.501340 | 0.468650 | 0.345951 | 0.249439 | 0.215487 | 0.282046 | 0.446845 | 0.619425 | 0.611482 | 0.350367 | 0.225644 | 0.234681 | 0.306033 | 0.389735 | 0.472994 | 0.548236 | 0.550142 | 0.506468 | 0.411454 | 0.334008 | 0.281671 | 0.257528 | 0.252247 | 0.258700 | 0.263192 | 0.283284 | 0.326411 | 0.397749 | 0.461385 | 0.487845 | 0.458426 | 0.418982 | 0.358040 | 0.298340 | 0.250599 | 0.226218 | 0.216029 | 0.214643 | 0.214627 | 0.216259 | 0.223529 | 0.231039 | 0.226150 | 0.218902 | 0.215367 | 0.215028 | 0.219915 | 0.230035 | 0.237051 | 0.244064 | 0.262741 | 0.296857 | 0.341943 | 0.394860 | 0.433251 | 0.454249 | 0.450259 | 0.395488 | 0.355734 | 0.329238 | 0.294893 | 0.255787 | 0.236151 | 0.228016 | 0.228122 | 0.235632 | 0.256432 | 0.287917 | 0.338128 | 0.394858 | 0.466052 | 0.517784 | 0.607756 | 0.587827 | 0.478462 | 0.323799 | 0.230578 | 0.231449 | 0.378289 | 0.599922 | 0.693307 | 0.674798 | 0.528840 | 0.394874 | 0.310397 | 0.258870 | 0.228879 | 0.218009 | 0.214685 | 0.215342 | 0.216738 | 0.216643 | 0.215552 | 0.214878 | 0.222806 | 0.253800 | 0.329539 | 0.472772 | 0.652418 | 0.774575 | 0.720358 | 0.588443 | 0.494609 | 0.428316 | 0.381402 | 0.366387 | 0.354028 | 0.335323 | 0.306056 | 0.281283 | 0.261105 | 0.255881 | 0.256181 | 0.262039 | 0.271970 | 0.287458 | 0.280007 | 0.261202 | 0.245210 | 0.227813 | 0.217254 | 0.215043 | 0.214822 | 0.214707 | 0.215436 | 0.221303 | 0.233979 | 0.255988 | 0.289280 | 0.336693 | 0.376184 | 0.424391 | 0.445300 | 0.456877 | 0.456741 | 0.437783 | 0.388085 | 0.374011 | 0.363987 | 0.353984 | 0.349366 | 0.361660 | 0.379977 | 0.414457 | 0.454488 | 0.487111 | 0.515827 | 0.531098 | 0.519873 | 0.498703 | 0.466164 | 0.424387 | 0.372991 | 0.334232 | 0.304598 | 0.295711 | 0.291260 | 0.298024 | 0.315044 | 0.325641 | 0.330108 |
| right anterior | 0.459236 | 0.595599 | 0.572916 | 0.582460 | 0.570302 | 0.473320 | 0.431664 | 0.401508 | 0.376210 | 0.325202 | 0.240101 | 0.215922 | 0.238414 | 0.305034 | 0.349650 | 0.282186 | 0.219399 | 0.264478 | 0.904756 | 4.946408 | 24.575032 | 68.458179 | 67.953132 | 40.136788 | 16.378045 | 7.492007 | 3.838664 | 1.672544 | 0.746304 | 0.420181 | 0.245162 | 0.215996 | 0.240681 | 0.264839 | 0.272352 | 0.272467 | 0.287425 | 0.323768 | 0.359660 | 0.453657 | 0.614692 | 0.619182 | 0.491038 | 0.394100 | 0.297618 | 0.253062 | 0.228595 | 0.217456 | 0.214684 | 0.218566 | 0.227362 | 0.232409 | 0.232506 | 0.229748 | 0.220348 | 0.215919 | 0.214628 | 0.217191 | 0.225297 | 0.229311 | 0.230272 | 0.224500 | 0.215728 | 0.219201 | 0.235425 | 0.253389 | 0.272730 | 0.277170 | 0.258334 | 0.245790 | 0.240759 | 0.247651 | 0.265911 | 0.291353 | 0.321818 | 0.357613 | 0.373784 | 0.365637 | 0.348024 | 0.329224 | 0.322568 | 0.323993 | 0.355864 | 0.448314 | 0.623919 | 0.881328 | 1.127272 | 1.131808 | 1.017517 | 0.899953 | 0.798923 | 0.685657 | 0.628691 | 0.561492 | 0.500751 | 0.411322 | 0.345615 | 0.293913 | 0.276070 | 0.260957 | 0.267835 | 0.282979 | 0.306993 | 0.316152 | 0.297558 | 0.246789 | 0.219434 | 0.214958 | 0.220309 | 0.220509 | 0.215367 | 0.217129 | 0.234073 | 0.270632 | 0.312119 | 0.366398 | 0.421850 | 0.452891 | 0.418799 | 0.382505 | 0.328185 | 0.271353 | 0.227394 | 0.214686 | 0.218851 | 0.231333 | 0.249806 | 0.280577 | 0.302587 | 0.339393 | 0.376547 | 0.406523 | 0.344913 | 0.271443 | 0.222525 | 0.215930 | 0.234501 | 0.263604 | 0.293729 | 0.292064 | 0.284418 | 0.283995 | 0.286253 | 0.287489 | 0.301353 | 0.320691 | 0.346546 | 0.393083 | 0.425696 | 0.476155 | 0.504859 | 0.482289 | 0.370026 | 0.283693 | 0.228548 | 0.214692 | 0.229210 | 0.243008 | 0.245146 | 0.238428 | 0.227028 | 0.217809 | 0.215469 | 0.216407 | 0.221108 | 0.232765 | 0.260597 | 0.305838 | 0.372772 | 0.443079 | 0.490039 | 0.478479 | 0.388445 | 0.291061 | 0.240569 | 0.218382 | 0.214897 | 0.218097 | 0.218102 | 0.216256 | 0.215069 | 0.226420 | 0.278554 | 0.346235 | 0.385241 | 0.338178 | 0.280307 | 0.234379 | 0.217563 | 0.214723 | 0.215239 | 0.219722 | 0.233192 | 0.260070 | 0.306179 | 0.354989 | 0.358356 | 0.318887 | 0.280665 | 0.257924 | 0.238342 | 0.228524 | 0.226858 | 0.229174 | 0.221747 | 0.215718 | 0.214826 | 0.216354 | 0.220102 | 0.220754 | 0.216575 | 0.214754 | 0.217661 | 0.221564 | 0.223085 | 0.221546 | 0.219403 | 0.215482 | 0.214627 | 0.215026 | 0.216141 | 0.218882 | 0.221923 | 0.226577 | 0.226517 | 0.222860 | 0.220700 | 0.221090 | 0.230278 | 0.278757 | 0.446428 | 0.982810 | 2.329657 | 5.216292 | 9.004412 | 10.079413 | 7.233657 | 4.125562 | 2.043890 | 1.143002 | 0.747867 | 0.532436 | 0.416540 | 0.345199 | 0.307269 | 0.286402 | 0.281571 | 0.287708 | 0.309359 | 0.333627 | 0.366441 | 0.405415 | 0.462469 | 0.508007 | 0.509746 | 0.494279 | 0.484204 | 0.454079 | 0.399884 | 0.368956 | 0.345512 | 0.328777 | 0.317881 | 0.328996 | 0.351800 | 0.386694 | 0.405938 | 0.392029 | 0.350851 | 0.299453 | 0.250266 | 0.227792 | 0.220992 | 0.216597 | 0.214638 | 0.214830 | 0.215817 | 0.218421 | 0.220071 | 0.219101 |
| left central | 0.244769 | 0.269764 | 0.296961 | 0.352708 | 0.360862 | 0.383304 | 0.365089 | 0.319098 | 0.279542 | 0.261167 | 0.248111 | 0.249064 | 0.263432 | 0.280683 | 0.287722 | 0.274257 | 0.264948 | 0.264376 | 0.282092 | 0.275494 | 0.279062 | 0.273279 | 0.267660 | 0.260233 | 0.275894 | 0.266622 | 0.254823 | 0.234132 | 0.219938 | 0.215874 | 0.228837 | 0.292818 | 0.412381 | 0.543079 | 0.504685 | 0.413186 | 0.290020 | 0.243806 | 0.234911 | 0.236641 | 0.247838 | 0.276016 | 0.315126 | 0.338916 | 0.354512 | 0.347103 | 0.320215 | 0.279190 | 0.249767 | 0.233893 | 0.239148 | 0.267947 | 0.316892 | 0.378486 | 0.446828 | 0.471808 | 0.475945 | 0.419101 | 0.360227 | 0.311715 | 0.282895 | 0.264567 | 0.266327 | 0.276235 | 0.306007 | 0.329443 | 0.342212 | 0.344131 | 0.302646 | 0.241968 | 0.215126 | 0.230379 | 0.274958 | 0.315507 | 0.327741 | 0.302309 | 0.276249 | 0.274584 | 0.299474 | 0.367019 | 0.530248 | 0.682456 | 0.808545 | 0.770444 | 0.617179 | 0.381529 | 0.243348 | 0.222723 | 0.328401 | 0.715228 | 1.572870 | 3.443229 | 4.731312 | 5.233220 | 3.660744 | 2.048859 | 1.519674 | 1.344835 | 1.630050 | 2.285456 | 3.255288 | 4.617635 | 6.796272 | 6.209514 | 5.633449 | 4.947627 | 2.742063 | 1.863117 | 1.776959 | 1.642958 | 1.665148 | 1.952997 | 2.080693 | 3.339841 | 5.803727 | 6.564450 | 4.430615 | 1.928472 | 0.610527 | 0.308309 | 0.232341 | 0.214767 | 0.221758 | 0.250479 | 0.284333 | 0.329495 | 0.354398 | 0.379262 | 0.402170 | 0.413719 | 0.379500 | 0.373176 | 0.353746 | 0.324310 | 0.296831 | 0.265810 | 0.237474 | 0.221867 | 0.214642 | 0.226952 | 0.277368 | 0.393081 | 0.655020 | 1.109167 | 1.683417 | 1.960915 | 1.686998 | 1.321041 | 1.183811 | 1.200711 | 1.238416 | 1.667495 | 2.614521 | 4.137375 | 6.459939 | 11.235443 | 22.308336 | 35.834468 | 39.820707 | 29.435201 | 17.694269 | 10.665074 | 7.601357 | 5.095398 | 4.077824 | 3.728925 | 3.155802 | 2.873457 | 2.749818 | 2.932562 | 3.576345 | 4.486583 | 5.851630 | 8.530760 | 10.672757 | 12.410973 | 17.925686 | 27.685555 | 41.652236 | 52.720600 | 46.802461 | 28.676407 | 15.091937 | 8.024472 | 4.030863 | 2.349462 | 1.400815 | 0.971637 | 0.777327 | 0.671726 | 0.620267 | 0.624208 | 0.745393 | 0.967031 | 1.440555 | 2.445009 | 4.397478 | 6.000976 | 5.369261 | 3.630687 | 1.800100 | 0.887009 | 0.540128 | 0.429489 | 0.370848 | 0.350922 | 0.352223 | 0.345821 | 0.332824 | 0.325067 | 0.315601 | 0.307588 | 0.314273 | 0.311246 | 0.319778 | 0.339781 | 0.373430 | 0.416602 | 0.480791 | 0.496229 | 0.493000 | 0.442296 | 0.383218 | 0.324234 | 0.290818 | 0.266671 | 0.257790 | 0.253488 | 0.255348 | 0.254242 | 0.246790 | 0.238653 | 0.236569 | 0.237880 | 0.252507 | 0.302586 | 0.433332 | 0.654384 | 0.886573 | 1.004688 | 0.993311 | 0.855710 | 0.687078 | 0.524505 | 0.428081 | 0.372790 | 0.347413 | 0.350516 | 0.391651 | 0.467762 | 0.606284 | 0.842707 | 1.206994 | 1.653807 | 1.869633 | 1.695220 | 1.371518 | 0.939777 | 0.767224 | 0.712707 | 0.745809 | 0.886660 | 1.275587 | 1.849871 | 2.719827 | 3.450840 | 3.721946 | 4.439438 | 5.462446 | 7.886875 | 14.271756 | 29.640580 | 44.856284 | 54.857692 | 44.107664 | 26.111520 | 13.047803 | 6.494234 | 4.317614 | 3.182931 |
| right central | 0.473078 | 0.426442 | 0.372495 | 0.303377 | 0.254119 | 0.219322 | 0.215318 | 0.220449 | 0.221671 | 0.218537 | 0.215414 | 0.214886 | 0.215457 | 0.217156 | 0.219767 | 0.221146 | 0.230510 | 0.257341 | 0.300130 | 0.302315 | 0.302882 | 0.303706 | 0.298010 | 0.294388 | 0.296006 | 0.301332 | 0.281629 | 0.253954 | 0.230007 | 0.226091 | 0.224901 | 0.233692 | 0.231676 | 0.246638 | 0.258424 | 0.236811 | 0.216846 | 0.217584 | 0.263996 | 0.328465 | 0.376091 | 0.450087 | 0.398069 | 0.274368 | 0.220133 | 0.222059 | 0.275631 | 0.340343 | 0.392130 | 0.355193 | 0.299943 | 0.253110 | 0.227154 | 0.214629 | 0.234095 | 0.332621 | 0.591642 | 1.090486 | 1.484604 | 1.247207 | 0.862469 | 0.565578 | 0.428739 | 0.377332 | 0.408966 | 0.534522 | 0.864522 | 1.559244 | 2.290632 | 1.682059 | 0.654493 | 0.260933 | 0.220038 | 0.282782 | 0.364182 | 0.412278 | 0.437217 | 0.481559 | 0.558438 | 0.690116 | 1.005660 | 1.477214 | 2.595558 | 5.329894 | 10.683500 | 13.772613 | 9.552640 | 3.899125 | 1.807046 | 0.902245 | 0.596657 | 0.447395 | 0.356348 | 0.291796 | 0.275570 | 0.260743 | 0.248434 | 0.254074 | 0.277577 | 0.302705 | 0.364202 | 0.451438 | 0.492350 | 0.462469 | 0.362324 | 0.276312 | 0.235808 | 0.223569 | 0.222194 | 0.231281 | 0.243538 | 0.263623 | 0.269494 | 0.256893 | 0.230288 | 0.215463 | 0.220316 | 0.238899 | 0.247076 | 0.244071 | 0.229820 | 0.218329 | 0.214629 | 0.218493 | 0.228842 | 0.241111 | 0.255161 | 0.268181 | 0.261028 | 0.251547 | 0.242520 | 0.242163 | 0.235372 | 0.227220 | 0.222748 | 0.221610 | 0.217196 | 0.216070 | 0.215545 | 0.215061 | 0.214809 | 0.215186 | 0.215795 | 0.218907 | 0.224492 | 0.233365 | 0.242165 | 0.252818 | 0.268894 | 0.288290 | 0.297971 | 0.288146 | 0.256313 | 0.224120 | 0.216338 | 0.256672 | 0.339886 | 0.421895 | 0.423538 | 0.345013 | 0.254116 | 0.216173 | 0.229840 | 0.281384 | 0.349285 | 0.391065 | 0.409987 | 0.393015 | 0.340239 | 0.279861 | 0.241296 | 0.220884 | 0.215336 | 0.214631 | 0.214664 | 0.214709 | 0.215031 | 0.214637 | 0.216078 | 0.219818 | 0.222204 | 0.223118 | 0.217062 | 0.214633 | 0.217879 | 0.230060 | 0.253264 | 0.286664 | 0.329480 | 0.363306 | 0.383047 | 0.380504 | 0.349212 | 0.320910 | 0.285426 | 0.260690 | 0.247775 | 0.237744 | 0.232710 | 0.227953 | 0.221438 | 0.216042 | 0.214626 | 0.215915 | 0.217865 | 0.222477 | 0.229010 | 0.236402 | 0.238363 | 0.235615 | 0.226150 | 0.218518 | 0.215433 | 0.214848 | 0.215564 | 0.217439 | 0.224542 | 0.238081 | 0.263983 | 0.273634 | 0.272763 | 0.261119 | 0.249295 | 0.228442 | 0.219565 | 0.215538 | 0.215030 | 0.214742 | 0.215336 | 0.217749 | 0.224779 | 0.233348 | 0.253082 | 0.282248 | 0.320008 | 0.373934 | 0.447270 | 0.562725 | 0.759803 | 1.085060 | 1.407521 | 1.409178 | 1.245401 | 1.004831 | 0.792694 | 0.576352 | 0.421426 | 0.332967 | 0.288292 | 0.251838 | 0.238557 | 0.240240 | 0.239921 | 0.241074 | 0.244913 | 0.245236 | 0.239721 | 0.228545 | 0.218224 | 0.214808 | 0.215456 | 0.217782 | 0.216957 | 0.215973 | 0.215093 | 0.214668 | 0.214830 | 0.220159 | 0.241939 | 0.295389 | 0.422015 | 0.709965 | 1.204830 | 1.948420 | 2.851834 | 3.666843 | 3.945218 | 3.833160 | 3.457732 | 3.081191 |
| left posterior | 0.924559 | 1.139986 | 1.565848 | 2.559900 | 2.732489 | 1.817452 | 0.838934 | 0.380085 | 0.230088 | 0.218605 | 0.248882 | 0.299164 | 0.343222 | 0.322092 | 0.271385 | 0.218184 | 0.268380 | 1.062866 | 3.935725 | 8.379383 | 9.869583 | 9.058096 | 6.084536 | 4.632473 | 1.677974 | 0.725505 | 0.398953 | 0.304745 | 0.245439 | 0.228481 | 0.219485 | 0.218261 | 0.216712 | 0.219884 | 0.220492 | 0.221133 | 0.217928 | 0.217334 | 0.220366 | 0.232239 | 0.251205 | 0.275880 | 0.292091 | 0.251135 | 0.216100 | 0.228396 | 0.264100 | 0.326003 | 0.370776 | 0.357255 | 0.292218 | 0.237660 | 0.214748 | 0.236958 | 0.286696 | 0.279409 | 0.238378 | 0.214704 | 0.255068 | 0.397984 | 0.464987 | 0.486138 | 0.501885 | 0.465365 | 0.423733 | 0.387324 | 0.380268 | 0.393120 | 0.570960 | 1.320987 | 4.909035 | 25.191763 | 124.819979 | 379.892464 | 622.656286 | 571.523910 | 238.294477 | 88.756445 | 21.577758 | 4.373594 | 1.299524 | 0.602647 | 0.393187 | 0.321626 | 0.274772 | 0.243078 | 0.229964 | 0.219585 | 0.216004 | 0.216574 | 0.219374 | 0.224905 | 0.237326 | 0.247255 | 0.248747 | 0.252097 | 0.249398 | 0.239864 | 0.231132 | 0.230422 | 0.228320 | 0.229892 | 0.239412 | 0.265530 | 0.301266 | 0.334509 | 0.334906 | 0.304529 | 0.248441 | 0.216279 | 0.222586 | 0.260710 | 0.330511 | 0.411475 | 0.460050 | 0.436612 | 0.352357 | 0.269755 | 0.221522 | 0.219011 | 0.255394 | 0.313573 | 0.358034 | 0.359799 | 0.317251 | 0.258964 | 0.225009 | 0.214674 | 0.226047 | 0.249507 | 0.250829 | 0.236440 | 0.220130 | 0.214628 | 0.222146 | 0.236157 | 0.247577 | 0.248082 | 0.237864 | 0.223204 | 0.215467 | 0.215255 | 0.217098 | 0.217107 | 0.215545 | 0.214626 | 0.215335 | 0.214806 | 0.216176 | 0.234399 | 0.280956 | 0.374032 | 0.513379 | 0.635198 | 0.735873 | 0.785103 | 0.748565 | 0.639712 | 0.543109 | 0.449864 | 0.405639 | 0.373449 | 0.348663 | 0.330688 | 0.322882 | 0.314576 | 0.332370 | 0.366510 | 0.415243 | 0.458661 | 0.525865 | 0.553156 | 0.536950 | 0.456007 | 0.375085 | 0.309497 | 0.270245 | 0.242257 | 0.227557 | 0.219214 | 0.215180 | 0.215480 | 0.221785 | 0.235049 | 0.255986 | 0.279940 | 0.295521 | 0.304225 | 0.305378 | 0.297224 | 0.287243 | 0.273618 | 0.265546 | 0.264699 | 0.268640 | 0.278985 | 0.307237 | 0.339108 | 0.382823 | 0.451777 | 0.537089 | 0.609003 | 0.652727 | 0.664595 | 0.614037 | 0.546977 | 0.469994 | 0.427951 | 0.396054 | 0.381687 | 0.374457 | 0.383084 | 0.401517 | 0.452661 | 0.516828 | 0.636341 | 0.814045 | 0.963385 | 0.934547 | 0.810640 | 0.637675 | 0.548544 | 0.502148 | 0.467125 | 0.460223 | 0.472190 | 0.446401 | 0.420710 | 0.381281 | 0.342701 | 0.307693 | 0.293056 | 0.288026 | 0.307735 | 0.353797 | 0.439311 | 0.540595 | 0.593842 | 0.523608 | 0.407726 | 0.325499 | 0.283902 | 0.270156 | 0.276786 | 0.304024 | 0.361498 | 0.465244 | 0.640099 | 0.883013 | 1.120782 | 1.331417 | 1.424132 | 1.229921 | 0.916070 | 0.656756 | 0.464189 | 0.338694 | 0.285271 | 0.258608 | 0.243297 | 0.232687 | 0.227817 | 0.224422 | 0.224192 | 0.225411 | 0.230642 | 0.242488 | 0.261948 | 0.289508 | 0.326850 | 0.368007 | 0.410207 | 0.455070 | 0.530639 | 0.674354 | 0.890505 | 1.123269 | 1.404765 | 1.520556 | 1.588258 |
| right posterior | 4.563489 | 2.829250 | 1.766252 | 0.808528 | 0.392076 | 0.259848 | 0.218097 | 0.216404 | 0.239722 | 0.261571 | 0.262341 | 0.242094 | 0.220123 | 0.217075 | 0.268417 | 0.364840 | 0.441490 | 0.395510 | 0.315572 | 0.226610 | 0.220234 | 0.288579 | 0.364862 | 0.361035 | 0.247918 | 0.216197 | 0.301272 | 0.502680 | 0.546147 | 0.347459 | 0.246375 | 0.215496 | 0.220121 | 0.235782 | 0.264847 | 0.280083 | 0.264854 | 0.237946 | 0.224662 | 0.215204 | 0.214737 | 0.216319 | 0.216684 | 0.216660 | 0.215345 | 0.215059 | 0.214638 | 0.217509 | 0.237912 | 0.286186 | 0.364767 | 0.460377 | 0.507184 | 0.517314 | 0.470315 | 0.440595 | 0.441319 | 0.499659 | 0.522031 | 0.454070 | 0.327293 | 0.248331 | 0.217707 | 0.216298 | 0.220195 | 0.218160 | 0.214840 | 0.214665 | 0.214910 | 0.231751 | 0.317490 | 0.514171 | 0.789949 | 1.007739 | 0.983763 | 0.785410 | 0.566975 | 0.383774 | 0.270362 | 0.220632 | 0.216479 | 0.228427 | 0.242419 | 0.255111 | 0.267645 | 0.297618 | 0.379513 | 0.499744 | 0.598519 | 0.634658 | 0.515501 | 0.396680 | 0.334464 | 0.324657 | 0.326885 | 0.370582 | 0.438192 | 0.550010 | 0.664110 | 0.742088 | 0.707608 | 0.631272 | 0.518211 | 0.398457 | 0.327816 | 0.306861 | 0.324295 | 0.369467 | 0.493931 | 0.768312 | 1.264068 | 2.011749 | 2.454381 | 1.858308 | 1.248234 | 0.779326 | 0.475156 | 0.331278 | 0.256015 | 0.223934 | 0.215139 | 0.214847 | 0.215706 | 0.214852 | 0.214636 | 0.216718 | 0.226398 | 0.246344 | 0.297176 | 0.436423 | 0.786281 | 1.989367 | 7.681900 | 26.133948 | 69.957204 | 106.205446 | 87.980795 | 53.133179 | 25.560484 | 11.141854 | 5.148893 | 2.561383 | 1.367600 | 0.843753 | 0.563387 | 0.407701 | 0.338194 | 0.309785 | 0.289176 | 0.287137 | 0.292435 | 0.322677 | 0.362791 | 0.425342 | 0.488358 | 0.524235 | 0.430087 | 0.341763 | 0.272138 | 0.236160 | 0.223769 | 0.222223 | 0.225240 | 0.233299 | 0.243425 | 0.253642 | 0.264880 | 0.283703 | 0.335525 | 0.440171 | 0.602975 | 0.861435 | 1.091857 | 1.063735 | 0.897676 | 0.719200 | 0.573914 | 0.525159 | 0.518699 | 0.538745 | 0.590027 | 0.529136 | 0.418667 | 0.325692 | 0.262869 | 0.234422 | 0.224239 | 0.218857 | 0.221359 | 0.231951 | 0.243574 | 0.261601 | 0.275991 | 0.268773 | 0.255292 | 0.244030 | 0.231460 | 0.224833 | 0.218565 | 0.214982 | 0.215738 | 0.221452 | 0.232774 | 0.245371 | 0.262969 | 0.287289 | 0.331358 | 0.400983 | 0.511059 | 0.631852 | 0.696226 | 0.630360 | 0.484621 | 0.361008 | 0.279567 | 0.235509 | 0.218313 | 0.214742 | 0.214626 | 0.216218 | 0.227414 | 0.252387 | 0.279430 | 0.286367 | 0.274686 | 0.252861 | 0.230851 | 0.219460 | 0.215055 | 0.215308 | 0.216575 | 0.215431 | 0.215603 | 0.226433 | 0.253907 | 0.322604 | 0.451979 | 0.590864 | 0.732068 | 0.831378 | 0.824782 | 0.802738 | 0.752604 | 0.743662 | 0.817659 | 0.844393 | 0.913704 | 1.098736 | 1.149376 | 1.025722 | 0.813405 | 0.562688 | 0.385842 | 0.287389 | 0.239943 | 0.222827 | 0.215873 | 0.215068 | 0.219198 | 0.225201 | 0.231685 | 0.238775 | 0.245439 | 0.257651 | 0.274887 | 0.308533 | 0.359264 | 0.402824 | 0.413469 | 0.405961 | 0.358678 | 0.305687 | 0.263151 | 0.231066 | 0.216522 | 0.214725 | 0.216904 | 0.219084 | 0.219664 | 0.219631 |
| all electrodes | 0.252279 | 0.244828 | 0.227114 | 0.214971 | 0.224891 | 0.242200 | 0.287032 | 0.322888 | 0.385041 | 0.601498 | 1.019263 | 1.641599 | 2.175977 | 2.376359 | 1.896676 | 1.669314 | 1.504211 | 1.318289 | 1.149764 | 0.868845 | 0.576460 | 0.484668 | 0.504133 | 0.511266 | 0.651400 | 1.169201 | 2.626959 | 3.344570 | 3.356530 | 2.864401 | 2.583769 | 1.893765 | 1.370748 | 0.741734 | 0.459071 | 0.316889 | 0.257628 | 0.230851 | 0.217645 | 0.219450 | 0.257982 | 0.354626 | 0.537574 | 0.823759 | 0.787472 | 0.548692 | 0.324255 | 0.233512 | 0.214633 | 0.217132 | 0.215818 | 0.216234 | 0.250879 | 0.350608 | 0.516080 | 0.649732 | 0.582950 | 0.401555 | 0.296349 | 0.234168 | 0.215480 | 0.217080 | 0.224211 | 0.227864 | 0.224534 | 0.218239 | 0.217514 | 0.224830 | 0.261231 | 0.374578 | 0.749586 | 1.659779 | 3.154886 | 4.019400 | 3.772565 | 2.276055 | 1.284554 | 0.839618 | 0.623957 | 0.616899 | 0.795034 | 1.109514 | 1.373678 | 1.445055 | 1.367841 | 1.290583 | 1.018077 | 0.601629 | 0.346470 | 0.250854 | 0.219750 | 0.214671 | 0.220193 | 0.224989 | 0.229436 | 0.233006 | 0.248058 | 0.288044 | 0.370261 | 0.501112 | 0.703040 | 0.798370 | 0.679530 | 0.497415 | 0.370329 | 0.284292 | 0.243513 | 0.225178 | 0.217837 | 0.215601 | 0.215312 | 0.214739 | 0.214815 | 0.215926 | 0.227876 | 0.255251 | 0.295205 | 0.364114 | 0.531291 | 0.682135 | 0.891544 | 0.981452 | 0.884895 | 0.590048 | 0.398407 | 0.277630 | 0.226008 | 0.215450 | 0.229819 | 0.251184 | 0.250367 | 0.235477 | 0.220808 | 0.215043 | 0.216136 | 0.219405 | 0.219006 | 0.216070 | 0.214632 | 0.221578 | 0.246467 | 0.288979 | 0.359419 | 0.466537 | 0.562385 | 0.593648 | 0.515120 | 0.425956 | 0.372357 | 0.352073 | 0.372867 | 0.483782 | 0.780080 | 1.385395 | 2.097670 | 2.541085 | 2.441972 | 1.898330 | 1.345136 | 1.139227 | 1.069369 | 1.112444 | 1.098506 | 1.065080 | 0.939939 | 0.780969 | 0.646972 | 0.584521 | 0.599636 | 0.720122 | 1.056839 | 1.922301 | 4.424926 | 9.791966 | 16.845956 | 21.055734 | 20.700586 | 13.809531 | 9.583661 | 5.221908 | 2.427846 | 1.113349 | 0.752921 | 0.605773 | 0.573448 | 0.603104 | 0.724412 | 0.903757 | 1.064275 | 1.163790 | 1.251912 | 1.237018 | 1.066767 | 0.922873 | 0.736329 | 0.559681 | 0.448968 | 0.360374 | 0.285782 | 0.255218 | 0.243014 | 0.235074 | 0.227938 | 0.222986 | 0.220548 | 0.220793 | 0.219936 | 0.216859 | 0.215183 | 0.214633 | 0.215158 | 0.215259 | 0.214656 | 0.217362 | 0.227775 | 0.241610 | 0.252528 | 0.250472 | 0.239731 | 0.223879 | 0.216486 | 0.214842 | 0.217442 | 0.221723 | 0.227734 | 0.242082 | 0.257286 | 0.282256 | 0.303432 | 0.323724 | 0.348345 | 0.384970 | 0.433452 | 0.532808 | 0.672934 | 0.836270 | 0.932352 | 0.865612 | 0.771187 | 0.674173 | 0.600401 | 0.548510 | 0.520234 | 0.495603 | 0.493718 | 0.492848 | 0.518377 | 0.529639 | 0.528045 | 0.492713 | 0.415533 | 0.325000 | 0.265073 | 0.227691 | 0.215525 | 0.216731 | 0.233703 | 0.261555 | 0.299389 | 0.342270 | 0.367285 | 0.333488 | 0.282157 | 0.239731 | 0.219458 | 0.214683 | 0.217951 | 0.221283 | 0.219578 | 0.215786 | 0.215221 | 0.222625 | 0.238770 | 0.250221 | 0.243502 | 0.233530 | 0.224081 | 0.221465 | 0.220437 | 0.222414 |

J) maximum cross-correlation, identity

  
|  | time window | peak latency | cluster *p* | peak Cohen's *d* |  | | | |
| **all electrodes** | 445 - 635 ms | 550 ms | 0.021 | 0.5689 |  | | | |
|  | | | | | | | | |

Model correlations, cluster permutation tests

|  | **left hemisphere** | | | | **right hemisphere** | | | |
|  | time window | peak latency | cluster *p* | peak Cohen's *d* | time window | peak latency | cluster *p* | peak Cohen's *d* |
| **anterior** |  | | | |  | | | |
| **central** |  | | | | -125 - -40 ms | -65 ms | 0.045 | -0.9028 |
| **posterior** |  | | | |  | | | |

  

Model correlations, Bayesian statistics

|  | -200 | -195 | -190 | -185 | -180 | -175 | -170 | -165 | -160 | -155 | -150 | -145 | -140 | -135 | -130 | -125 | -120 | -115 | -110 | -105 | -100 | -95 | -90 | -85 | -80 | -75 | -70 | -65 | -60 | -55 | -50 | -45 | -40 | -35 | -30 | -25 | -20 | -15 | -10 | -5 | 0 | 5 | 10 | 15 | 20 | 25 | 30 | 35 | 40 | 45 | 50 | 55 | 60 | 65 | 70 | 75 | 80 | 85 | 90 | 95 | 100 | 105 | 110 | 115 | 120 | 125 | 130 | 135 | 140 | 145 | 150 | 155 | 160 | 165 | 170 | 175 | 180 | 185 | 190 | 195 | 200 | 205 | 210 | 215 | 220 | 225 | 230 | 235 | 240 | 245 | 250 | 255 | 260 | 265 | 270 | 275 | 280 | 285 | 290 | 295 | 300 | 305 | 310 | 315 | 320 | 325 | 330 | 335 | 340 | 345 | 350 | 355 | 360 | 365 | 370 | 375 | 380 | 385 | 390 | 395 | 400 | 405 | 410 | 415 | 420 | 425 | 430 | 435 | 440 | 445 | 450 | 455 | 460 | 465 | 470 | 475 | 480 | 485 | 490 | 495 | 500 | 505 | 510 | 515 | 520 | 525 | 530 | 535 | 540 | 545 | 550 | 555 | 560 | 565 | 570 | 575 | 580 | 585 | 590 | 595 | 600 | 605 | 610 | 615 | 620 | 625 | 630 | 635 | 640 | 645 | 650 | 655 | 660 | 665 | 670 | 675 | 680 | 685 | 690 | 695 | 700 | 705 | 710 | 715 | 720 | 725 | 730 | 735 | 740 | 745 | 750 | 755 | 760 | 765 | 770 | 775 | 780 | 785 | 790 | 795 | 800 | 805 | 810 | 815 | 820 | 825 | 830 | 835 | 840 | 845 | 850 | 855 | 860 | 865 | 870 | 875 | 880 | 885 | 890 | 895 | 900 | 905 | 910 | 915 | 920 | 925 | 930 | 935 | 940 | 945 | 950 | 955 | 960 | 965 | 970 | 975 | 980 | 985 | 990 | 995 | 1000 | 1005 | 1010 | 1015 | 1020 | 1025 | 1030 | 1035 | 1040 | 1045 | 1050 | 1055 | 1060 | 1065 | 1070 | 1075 | 1080 | 1085 | 1090 | 1095 | 1100 | 1105 | 1110 | 1115 | 1120 | 1125 | 1130 | 1135 | 1140 | 1145 | 1150 | 1155 | 1160 | 1165 | 1170 | 1175 | 1180 | 1185 | 1190 | 1195 |
| --- | --- | --- | --- | --- | --- | --- | --- | --- | --- | --- | --- | --- | --- | --- | --- | --- | --- | --- | --- | --- | --- | --- | --- | --- | --- | --- | --- | --- | --- | --- | --- | --- | --- | --- | --- | --- | --- | --- | --- | --- | --- | --- | --- | --- | --- | --- | --- | --- | --- | --- | --- | --- | --- | --- | --- | --- | --- | --- | --- | --- | --- | --- | --- | --- | --- | --- | --- | --- | --- | --- | --- | --- | --- | --- | --- | --- | --- | --- | --- | --- | --- | --- | --- | --- | --- | --- | --- | --- | --- | --- | --- | --- | --- | --- | --- | --- | --- | --- | --- | --- | --- | --- | --- | --- | --- | --- | --- | --- | --- | --- | --- | --- | --- | --- | --- | --- | --- | --- | --- | --- | --- | --- | --- | --- | --- | --- | --- | --- | --- | --- | --- | --- | --- | --- | --- | --- | --- | --- | --- | --- | --- | --- | --- | --- | --- | --- | --- | --- | --- | --- | --- | --- | --- | --- | --- | --- | --- | --- | --- | --- | --- | --- | --- | --- | --- | --- | --- | --- | --- | --- | --- | --- | --- | --- | --- | --- | --- | --- | --- | --- | --- | --- | --- | --- | --- | --- | --- | --- | --- | --- | --- | --- | --- | --- | --- | --- | --- | --- | --- | --- | --- | --- | --- | --- | --- | --- | --- | --- | --- | --- | --- | --- | --- | --- | --- | --- | --- | --- | --- | --- | --- | --- | --- | --- | --- | --- | --- | --- | --- | --- | --- | --- | --- | --- | --- | --- | --- | --- | --- | --- | --- | --- | --- | --- | --- | --- | --- | --- | --- | --- | --- | --- | --- | --- | --- | --- | --- | --- | --- | --- | --- | --- | --- | --- | --- | --- | --- | --- | --- | --- | --- | --- | --- | --- | --- | --- | --- | --- | --- | --- |
| left anterior | 0.245088 | 0.253959 | 0.254084 | 0.264466 | 0.262671 | 0.271850 | 0.289106 | 0.332760 | 0.399902 | 0.351859 | 0.253853 | 0.214631 | 0.251848 | 0.344276 | 0.467130 | 0.570249 | 0.551179 | 0.487394 | 0.434252 | 0.402051 | 0.464016 | 0.481342 | 0.449236 | 0.344472 | 0.243815 | 0.223024 | 0.308605 | 0.444639 | 0.434777 | 0.364383 | 0.252837 | 0.218078 | 0.348710 | 0.619704 | 1.049945 | 1.927872 | 5.230370 | 13.171243 | 25.930936 | 24.584155 | 7.981610 | 1.337845 | 0.411459 | 0.256665 | 0.218716 | 0.214895 | 0.219010 | 0.224871 | 0.229265 | 0.233222 | 0.247323 | 0.262460 | 0.272817 | 0.284024 | 0.289829 | 0.284845 | 0.282724 | 0.282395 | 0.275182 | 0.258086 | 0.237418 | 0.225369 | 0.218247 | 0.214766 | 0.215767 | 0.219542 | 0.226522 | 0.232893 | 0.234422 | 0.232612 | 0.229987 | 0.229942 | 0.233170 | 0.242007 | 0.253502 | 0.266708 | 0.275547 | 0.274902 | 0.255936 | 0.232703 | 0.217773 | 0.215656 | 0.227417 | 0.237596 | 0.239589 | 0.235614 | 0.227350 | 0.218995 | 0.214663 | 0.218133 | 0.225135 | 0.232849 | 0.246241 | 0.268546 | 0.296232 | 0.324478 | 0.335227 | 0.316990 | 0.270712 | 0.230105 | 0.215024 | 0.236781 | 0.276162 | 0.304395 | 0.285076 | 0.234822 | 0.214676 | 0.231421 | 0.274424 | 0.326808 | 0.383549 | 0.457948 | 0.526921 | 0.642191 | 0.884211 | 1.337137 | 2.269168 | 4.527609 | 8.856353 | 13.746542 | 15.707276 | 15.432701 | 14.510993 | 12.240439 | 9.500441 | 7.240433 | 5.277181 | 4.264157 | 3.429886 | 2.646664 | 1.881232 | 1.235494 | 0.723138 | 0.461690 | 0.310202 | 0.240465 | 0.216835 | 0.215121 | 0.217668 | 0.216814 | 0.215358 | 0.215014 | 0.219367 | 0.229946 | 0.248102 | 0.289777 | 0.343637 | 0.406078 | 0.487370 | 0.574738 | 0.602538 | 0.587238 | 0.546015 | 0.505889 | 0.497790 | 0.468453 | 0.412232 | 0.387380 | 0.366623 | 0.331570 | 0.300329 | 0.277358 | 0.254932 | 0.235451 | 0.221175 | 0.215413 | 0.214678 | 0.215950 | 0.217818 | 0.217479 | 0.216968 | 0.217889 | 0.220842 | 0.226326 | 0.236231 | 0.247317 | 0.254006 | 0.245922 | 0.230292 | 0.218195 | 0.214646 | 0.219369 | 0.226598 | 0.229612 | 0.225888 | 0.220647 | 0.216418 | 0.214633 | 0.215815 | 0.218670 | 0.223915 | 0.234893 | 0.247496 | 0.264141 | 0.278967 | 0.287779 | 0.272388 | 0.258619 | 0.245652 | 0.237709 | 0.232038 | 0.236586 | 0.241754 | 0.254080 | 0.260875 | 0.252934 | 0.231771 | 0.217252 | 0.216818 | 0.233365 | 0.259508 | 0.276105 | 0.288799 | 0.304822 | 0.317378 | 0.323878 | 0.334941 | 0.331935 | 0.325215 | 0.315014 | 0.288848 | 0.258248 | 0.239841 | 0.225652 | 0.217032 | 0.214769 | 0.214629 | 0.214729 | 0.214707 | 0.214643 | 0.214641 | 0.214863 | 0.215338 | 0.215027 | 0.214939 | 0.214675 | 0.215274 | 0.220160 | 0.230357 | 0.247625 | 0.266237 | 0.283355 | 0.297798 | 0.309683 | 0.315233 | 0.330315 | 0.340994 | 0.345646 | 0.339131 | 0.330992 | 0.298889 | 0.274597 | 0.250362 | 0.235396 | 0.226425 | 0.224697 | 0.221280 | 0.220521 | 0.218863 | 0.216755 | 0.214953 | 0.214734 | 0.218623 | 0.228853 | 0.244550 | 0.261845 | 0.280129 | 0.295689 | 0.308795 | 0.316732 | 0.324509 | 0.326152 | 0.310010 | 0.289066 | 0.270565 | 0.260546 | 0.254152 | 0.250697 | 0.248724 | 0.248789 |
| right anterior | 0.271116 | 0.309469 | 0.376456 | 0.478559 | 0.648342 | 0.932510 | 1.434887 | 2.027229 | 1.833037 | 1.102334 | 0.472591 | 0.253729 | 0.215585 | 0.214688 | 0.215177 | 0.219923 | 0.232141 | 0.239080 | 0.218545 | 0.217600 | 0.242073 | 0.297026 | 0.385797 | 0.472076 | 0.479227 | 0.353181 | 0.289760 | 0.284546 | 0.343842 | 0.606222 | 1.262481 | 1.905501 | 2.427825 | 2.740302 | 2.694422 | 2.292094 | 1.929010 | 1.764885 | 1.228349 | 0.676935 | 0.444273 | 0.300573 | 0.232437 | 0.215351 | 0.219823 | 0.238258 | 0.252358 | 0.264118 | 0.268219 | 0.248407 | 0.230982 | 0.221162 | 0.217257 | 0.217603 | 0.218893 | 0.224634 | 0.247019 | 0.299788 | 0.375654 | 0.482180 | 0.549989 | 0.618562 | 0.691842 | 0.733140 | 0.718022 | 0.660935 | 0.598367 | 0.538474 | 0.510443 | 0.533887 | 0.570123 | 0.603053 | 0.660205 | 0.662034 | 0.676063 | 0.704599 | 0.758005 | 0.836614 | 1.014693 | 1.211294 | 1.371148 | 1.127279 | 0.705443 | 0.473174 | 0.342897 | 0.276472 | 0.242852 | 0.226444 | 0.215808 | 0.216361 | 0.230623 | 0.257417 | 0.301036 | 0.344628 | 0.384642 | 0.407823 | 0.419145 | 0.425595 | 0.433474 | 0.437901 | 0.458597 | 0.484974 | 0.466339 | 0.419932 | 0.410746 | 0.401724 | 0.395125 | 0.399128 | 0.415029 | 0.398754 | 0.398057 | 0.376837 | 0.364235 | 0.373885 | 0.366352 | 0.336673 | 0.359988 | 0.411171 | 0.480292 | 0.779926 | 1.763223 | 4.672297 | 11.242041 | 17.468698 | 15.081519 | 6.286087 | 1.658620 | 0.564535 | 0.297586 | 0.227862 | 0.215235 | 0.214721 | 0.219336 | 0.249688 | 0.325547 | 0.438462 | 0.487883 | 0.401851 | 0.298155 | 0.245117 | 0.230784 | 0.238977 | 0.272638 | 0.356037 | 0.512257 | 0.710861 | 0.750169 | 0.690378 | 0.604563 | 0.581693 | 0.564300 | 0.599807 | 0.617124 | 0.565946 | 0.450057 | 0.349785 | 0.267901 | 0.230883 | 0.215519 | 0.216556 | 0.220146 | 0.218995 | 0.216611 | 0.214703 | 0.217423 | 0.225262 | 0.237104 | 0.243760 | 0.235826 | 0.223445 | 0.215431 | 0.232005 | 0.262297 | 0.298996 | 0.351180 | 0.388798 | 0.455792 | 0.477328 | 0.454930 | 0.385214 | 0.327174 | 0.288188 | 0.290791 | 0.318327 | 0.384966 | 0.473399 | 0.602585 | 0.636502 | 0.516082 | 0.384282 | 0.314828 | 0.283009 | 0.275189 | 0.271211 | 0.268492 | 0.278022 | 0.272047 | 0.254862 | 0.245399 | 0.242458 | 0.246637 | 0.268886 | 0.296360 | 0.330159 | 0.355380 | 0.331985 | 0.284127 | 0.247906 | 0.222240 | 0.214652 | 0.217268 | 0.221198 | 0.222817 | 0.221687 | 0.221635 | 0.222475 | 0.223937 | 0.227815 | 0.233610 | 0.240873 | 0.247542 | 0.258056 | 0.272740 | 0.292355 | 0.284446 | 0.259590 | 0.235293 | 0.219612 | 0.214685 | 0.214991 | 0.216204 | 0.215953 | 0.216048 | 0.220270 | 0.235379 | 0.272136 | 0.374082 | 0.547702 | 0.836985 | 1.177369 | 1.296819 | 1.366901 | 1.386442 | 1.191995 | 1.016291 | 0.942358 | 0.853788 | 0.795119 | 0.731151 | 0.652648 | 0.536430 | 0.430836 | 0.342596 | 0.298928 | 0.273888 | 0.261404 | 0.259157 | 0.270892 | 0.283242 | 0.293615 | 0.285000 | 0.258023 | 0.229857 | 0.217585 | 0.214717 | 0.215260 | 0.223242 | 0.254114 | 0.317723 | 0.419240 | 0.492935 | 0.502252 | 0.476591 | 0.460895 | 0.425384 | 0.403672 | 0.375601 | 0.352345 | 0.331299 | 0.315391 |
| left central | 0.253510 | 0.248484 | 0.246120 | 0.239372 | 0.220686 | 0.222438 | 0.301530 | 0.521934 | 0.797516 | 0.802422 | 0.529303 | 0.342936 | 0.246378 | 0.217543 | 0.215194 | 0.220151 | 0.222166 | 0.218298 | 0.215151 | 0.215200 | 0.214640 | 0.223423 | 0.254807 | 0.336286 | 0.623174 | 1.082688 | 1.441720 | 1.588346 | 1.525984 | 1.324866 | 1.038588 | 0.740611 | 0.616885 | 0.485831 | 0.386843 | 0.303576 | 0.235349 | 0.215020 | 0.237359 | 0.277089 | 0.289100 | 0.264645 | 0.244520 | 0.230261 | 0.222078 | 0.219030 | 0.217715 | 0.217769 | 0.218035 | 0.220091 | 0.225198 | 0.232198 | 0.235642 | 0.240651 | 0.242788 | 0.238646 | 0.230635 | 0.220406 | 0.215682 | 0.215314 | 0.215943 | 0.217174 | 0.228388 | 0.264126 | 0.333199 | 0.442329 | 0.572720 | 0.650034 | 0.637345 | 0.542979 | 0.467543 | 0.396242 | 0.365446 | 0.351784 | 0.339281 | 0.314078 | 0.277256 | 0.244251 | 0.223085 | 0.214627 | 0.222499 | 0.235136 | 0.252336 | 0.274886 | 0.279641 | 0.276717 | 0.270890 | 0.269293 | 0.274967 | 0.290722 | 0.304834 | 0.341110 | 0.352299 | 0.320682 | 0.261218 | 0.218073 | 0.236198 | 0.358409 | 0.648963 | 1.025112 | 1.350173 | 1.286383 | 0.889906 | 0.531502 | 0.359342 | 0.276928 | 0.244741 | 0.234903 | 0.228818 | 0.223836 | 0.217679 | 0.214842 | 0.222896 | 0.232304 | 0.238641 | 0.229763 | 0.217167 | 0.216860 | 0.238794 | 0.289496 | 0.351157 | 0.421050 | 0.451148 | 0.445342 | 0.431424 | 0.431662 | 0.443353 | 0.475232 | 0.493730 | 0.526058 | 0.576524 | 0.633975 | 0.691335 | 0.731071 | 0.671581 | 0.565654 | 0.460758 | 0.379487 | 0.324157 | 0.296072 | 0.286041 | 0.287300 | 0.300720 | 0.324566 | 0.349175 | 0.376774 | 0.415410 | 0.466960 | 0.554755 | 0.683519 | 0.797781 | 0.952347 | 1.105482 | 1.193565 | 1.191089 | 1.136096 | 0.990551 | 0.857457 | 0.733046 | 0.649286 | 0.610700 | 0.615704 | 0.624961 | 0.667135 | 0.748248 | 0.835331 | 0.848711 | 0.836826 | 0.781731 | 0.708019 | 0.606432 | 0.526011 | 0.470599 | 0.445602 | 0.442492 | 0.475850 | 0.537642 | 0.609889 | 0.674060 | 0.727473 | 0.738728 | 0.679159 | 0.539997 | 0.413851 | 0.329345 | 0.282871 | 0.262154 | 0.260267 | 0.264582 | 0.274648 | 0.281620 | 0.284861 | 0.282187 | 0.258573 | 0.231002 | 0.215452 | 0.220312 | 0.244153 | 0.258281 | 0.259575 | 0.244650 | 0.228708 | 0.217813 | 0.214750 | 0.221786 | 0.228252 | 0.227238 | 0.223659 | 0.217007 | 0.214661 | 0.216224 | 0.215299 | 0.215005 | 0.220359 | 0.231312 | 0.247198 | 0.254014 | 0.245199 | 0.228654 | 0.215598 | 0.217697 | 0.240373 | 0.292359 | 0.393383 | 0.550011 | 0.785638 | 0.996850 | 1.186126 | 1.191679 | 1.111444 | 0.937662 | 0.771799 | 0.639197 | 0.537713 | 0.461743 | 0.437975 | 0.391242 | 0.330081 | 0.289755 | 0.260087 | 0.239844 | 0.228458 | 0.221129 | 0.216290 | 0.214775 | 0.214748 | 0.215169 | 0.215197 | 0.214841 | 0.214803 | 0.214639 | 0.215522 | 0.218296 | 0.222372 | 0.230424 | 0.237339 | 0.244251 | 0.250838 | 0.264105 | 0.290791 | 0.346732 | 0.434649 | 0.573419 | 0.673440 | 0.686754 | 0.584712 | 0.465869 | 0.374036 | 0.339194 | 0.325082 | 0.344314 | 0.406645 | 0.534551 | 0.696874 | 0.854105 | 0.984262 | 1.000461 | 0.999145 | 0.987663 | 0.971040 |
| right central | 0.215928 | 0.214668 | 0.214642 | 0.214914 | 0.214799 | 0.224497 | 0.258170 | 0.300428 | 0.338310 | 0.300246 | 0.231056 | 0.219511 | 0.299924 | 0.534851 | 0.966578 | 1.681981 | 2.474478 | 3.086820 | 4.219834 | 5.376418 | 8.080615 | 13.542862 | 30.093142 | 47.131364 | 38.995986 | 37.836482 | 70.617378 | 109.887126 | 105.156650 | 64.429330 | 24.144699 | 8.420435 | 2.785187 | 0.806005 | 0.340883 | 0.226335 | 0.224656 | 0.299474 | 0.396635 | 0.441973 | 0.420842 | 0.396335 | 0.375005 | 0.366963 | 0.391747 | 0.433934 | 0.472123 | 0.457440 | 0.348602 | 0.271739 | 0.242685 | 0.233708 | 0.242340 | 0.283203 | 0.399962 | 0.617141 | 0.799432 | 0.761220 | 0.690911 | 0.561856 | 0.487204 | 0.442060 | 0.461660 | 0.570011 | 0.841351 | 1.185252 | 1.492844 | 1.455025 | 1.147152 | 0.880984 | 0.708136 | 0.631892 | 0.657617 | 0.735914 | 0.835808 | 0.988077 | 1.121852 | 1.210790 | 1.402810 | 1.813306 | 2.166994 | 2.225757 | 1.774947 | 1.082305 | 0.633335 | 0.418560 | 0.323568 | 0.288352 | 0.271574 | 0.264128 | 0.264309 | 0.275960 | 0.295817 | 0.326843 | 0.374319 | 0.452921 | 0.553646 | 0.681006 | 0.772961 | 0.681464 | 0.465375 | 0.317374 | 0.257297 | 0.235731 | 0.229967 | 0.233423 | 0.243571 | 0.250167 | 0.261200 | 0.287686 | 0.334030 | 0.424493 | 0.618842 | 0.989951 | 1.821740 | 3.275846 | 4.773785 | 5.536389 | 5.198439 | 2.879189 | 1.371249 | 0.679135 | 0.380338 | 0.260008 | 0.221522 | 0.214753 | 0.218519 | 0.219356 | 0.216265 | 0.215159 | 0.224176 | 0.239175 | 0.260411 | 0.273527 | 0.265273 | 0.238860 | 0.222615 | 0.216484 | 0.216144 | 0.221270 | 0.245328 | 0.295541 | 0.365232 | 0.434685 | 0.450536 | 0.434582 | 0.399205 | 0.349183 | 0.310770 | 0.291596 | 0.277782 | 0.274648 | 0.283265 | 0.290646 | 0.309206 | 0.339204 | 0.382338 | 0.480908 | 0.648728 | 0.802668 | 0.963271 | 1.020882 | 0.897669 | 0.719540 | 0.535186 | 0.394695 | 0.331796 | 0.298128 | 0.285558 | 0.297271 | 0.349122 | 0.431552 | 0.562082 | 0.698776 | 0.862775 | 0.933807 | 0.918067 | 0.890437 | 0.899153 | 0.881783 | 0.868257 | 0.733702 | 0.569305 | 0.466859 | 0.377370 | 0.310477 | 0.289543 | 0.276238 | 0.271382 | 0.284356 | 0.306526 | 0.317237 | 0.332088 | 0.318623 | 0.291940 | 0.263025 | 0.240491 | 0.223295 | 0.215355 | 0.216644 | 0.227301 | 0.239707 | 0.240422 | 0.233345 | 0.220049 | 0.214648 | 0.222224 | 0.241860 | 0.281597 | 0.330818 | 0.381169 | 0.401214 | 0.355832 | 0.283853 | 0.235589 | 0.215448 | 0.218772 | 0.227815 | 0.236362 | 0.240438 | 0.235376 | 0.227544 | 0.221872 | 0.217466 | 0.215546 | 0.214850 | 0.214641 | 0.214658 | 0.214843 | 0.215018 | 0.214962 | 0.214766 | 0.214800 | 0.217438 | 0.224369 | 0.235965 | 0.245474 | 0.249620 | 0.255456 | 0.260453 | 0.255945 | 0.247666 | 0.241516 | 0.236188 | 0.231657 | 0.228147 | 0.225815 | 0.223421 | 0.220169 | 0.215736 | 0.214817 | 0.218416 | 0.226926 | 0.236523 | 0.245180 | 0.249678 | 0.257515 | 0.272716 | 0.291272 | 0.311909 | 0.354740 | 0.403771 | 0.453830 | 0.506827 | 0.507227 | 0.471330 | 0.431766 | 0.383606 | 0.354782 | 0.341388 | 0.320687 | 0.297260 | 0.275360 | 0.252221 | 0.236570 | 0.224968 | 0.217727 | 0.214978 | 0.214628 | 0.215181 |
| left posterior | 0.250008 | 0.272634 | 0.308375 | 0.401216 | 0.536089 | 0.659959 | 0.615602 | 0.468863 | 0.334297 | 0.245198 | 0.214900 | 0.258082 | 0.341576 | 0.398684 | 0.401193 | 0.350242 | 0.299136 | 0.270503 | 0.278857 | 0.335306 | 0.440448 | 0.662029 | 1.032912 | 1.800708 | 3.135172 | 4.854095 | 6.833189 | 8.813036 | 8.126005 | 5.560409 | 2.890128 | 1.268559 | 0.549095 | 0.304283 | 0.233219 | 0.214901 | 0.217943 | 0.231199 | 0.256630 | 0.299965 | 0.362890 | 0.491881 | 0.663940 | 0.879430 | 1.113202 | 1.198046 | 0.982182 | 0.625216 | 0.394262 | 0.289852 | 0.258068 | 0.233397 | 0.219058 | 0.214676 | 0.219151 | 0.245932 | 0.289513 | 0.410267 | 0.622882 | 0.858569 | 0.996427 | 0.839160 | 0.487980 | 0.307503 | 0.237807 | 0.217584 | 0.214935 | 0.216024 | 0.234092 | 0.320355 | 0.569469 | 1.322950 | 2.892743 | 3.625206 | 2.436441 | 1.212288 | 0.595338 | 0.371748 | 0.266063 | 0.224141 | 0.214680 | 0.221773 | 0.235575 | 0.251147 | 0.268953 | 0.283313 | 0.276045 | 0.271594 | 0.259562 | 0.248673 | 0.236819 | 0.233476 | 0.228603 | 0.225444 | 0.219412 | 0.215450 | 0.214629 | 0.214861 | 0.215496 | 0.216010 | 0.215947 | 0.215743 | 0.217602 | 0.224411 | 0.237503 | 0.261640 | 0.301437 | 0.337767 | 0.335951 | 0.296482 | 0.256447 | 0.233599 | 0.222121 | 0.220362 | 0.227568 | 0.253607 | 0.318022 | 0.471266 | 0.815204 | 1.435993 | 1.792667 | 1.540752 | 1.023396 | 0.686380 | 0.485954 | 0.346149 | 0.260350 | 0.222351 | 0.216730 | 0.250020 | 0.307463 | 0.383829 | 0.517192 | 0.711674 | 0.969765 | 1.268022 | 1.476644 | 1.613254 | 1.571802 | 1.318822 | 1.024363 | 0.754950 | 0.518355 | 0.384496 | 0.307376 | 0.264004 | 0.243662 | 0.237356 | 0.241829 | 0.262246 | 0.305492 | 0.389137 | 0.565203 | 0.839353 | 1.060803 | 1.093772 | 0.896600 | 0.653477 | 0.498327 | 0.438282 | 0.436858 | 0.482381 | 0.572468 | 0.720155 | 0.847107 | 0.801441 | 0.671047 | 0.521944 | 0.410639 | 0.336019 | 0.297328 | 0.273977 | 0.256424 | 0.237396 | 0.223599 | 0.216529 | 0.214690 | 0.214627 | 0.214924 | 0.218804 | 0.232653 | 0.258072 | 0.285468 | 0.306707 | 0.318257 | 0.322367 | 0.314595 | 0.305694 | 0.304213 | 0.303608 | 0.301709 | 0.305119 | 0.293912 | 0.269362 | 0.244031 | 0.224194 | 0.214993 | 0.216241 | 0.220698 | 0.222263 | 0.218948 | 0.215460 | 0.214908 | 0.218803 | 0.225881 | 0.232484 | 0.235658 | 0.233285 | 0.228093 | 0.219502 | 0.214638 | 0.219385 | 0.237381 | 0.277667 | 0.331345 | 0.390777 | 0.468868 | 0.533623 | 0.587045 | 0.626195 | 0.640548 | 0.635720 | 0.651959 | 0.641862 | 0.660836 | 0.691443 | 0.662418 | 0.532222 | 0.409773 | 0.330899 | 0.296963 | 0.288836 | 0.304452 | 0.348500 | 0.432828 | 0.577694 | 0.773067 | 1.054032 | 1.343784 | 1.546432 | 1.544435 | 1.366879 | 0.999007 | 0.677432 | 0.443530 | 0.316894 | 0.262416 | 0.243793 | 0.241868 | 0.253876 | 0.292050 | 0.375975 | 0.569317 | 0.963679 | 1.852920 | 3.419153 | 5.517257 | 5.577134 | 3.771994 | 1.763140 | 0.734710 | 0.363174 | 0.243278 | 0.214698 | 0.225229 | 0.244465 | 0.257455 | 0.258613 | 0.244623 | 0.226380 | 0.215459 | 0.217130 | 0.228129 | 0.242322 | 0.254691 | 0.258703 | 0.255933 | 0.248377 | 0.245820 | 0.240900 |
| right posterior | 0.281880 | 0.277865 | 0.252566 | 0.219760 | 0.216572 | 0.241336 | 0.267155 | 0.278226 | 0.248501 | 0.216013 | 0.256682 | 0.369301 | 0.513998 | 0.522571 | 0.433746 | 0.324874 | 0.262277 | 0.239996 | 0.266267 | 0.379107 | 0.932301 | 4.716264 | 18.761971 | 33.191568 | 31.574982 | 11.254636 | 2.965023 | 1.070451 | 0.540003 | 0.323601 | 0.244555 | 0.217133 | 0.218430 | 0.245172 | 0.285643 | 0.364963 | 0.477550 | 0.671583 | 0.969919 | 1.311966 | 0.912308 | 0.640479 | 0.536135 | 0.538203 | 0.613490 | 0.799368 | 1.137684 | 1.521773 | 1.699347 | 1.791497 | 2.069710 | 2.612894 | 3.157489 | 3.107166 | 2.337614 | 1.277914 | 0.497009 | 0.249782 | 0.221790 | 0.373223 | 0.652410 | 0.587415 | 0.357764 | 0.235722 | 0.218389 | 0.268523 | 0.344254 | 0.364947 | 0.325801 | 0.263952 | 0.225387 | 0.214693 | 0.222807 | 0.232950 | 0.231969 | 0.222832 | 0.214701 | 0.224244 | 0.268179 | 0.399190 | 0.674935 | 1.149543 | 1.912500 | 2.617097 | 3.000406 | 2.966142 | 2.718072 | 2.182338 | 1.555863 | 0.927141 | 0.554448 | 0.345617 | 0.247851 | 0.217514 | 0.215292 | 0.216911 | 0.216443 | 0.214943 | 0.214725 | 0.214810 | 0.214629 | 0.214629 | 0.214644 | 0.215854 | 0.222234 | 0.239400 | 0.276408 | 0.332891 | 0.407909 | 0.539099 | 0.700185 | 0.824377 | 0.804027 | 0.644876 | 0.460600 | 0.341132 | 0.274252 | 0.246071 | 0.234966 | 0.232264 | 0.232426 | 0.233787 | 0.239860 | 0.254204 | 0.292421 | 0.402042 | 0.786783 | 2.352404 | 8.052900 | 21.704001 | 37.109278 | 40.124314 | 34.478374 | 23.349112 | 12.140594 | 6.078620 | 2.964465 | 1.512600 | 0.892073 | 0.632169 | 0.455275 | 0.362853 | 0.310993 | 0.280436 | 0.260881 | 0.255230 | 0.259038 | 0.290561 | 0.377661 | 0.564412 | 0.867832 | 1.200659 | 1.453408 | 1.570331 | 1.585688 | 1.393883 | 1.180109 | 0.964245 | 0.829287 | 0.741887 | 0.781344 | 0.914078 | 1.202344 | 1.725760 | 2.950117 | 4.844302 | 7.147081 | 7.633774 | 6.629095 | 4.093090 | 2.075678 | 0.954437 | 0.522766 | 0.322592 | 0.245936 | 0.220851 | 0.215401 | 0.214627 | 0.214791 | 0.217462 | 0.231921 | 0.265727 | 0.328090 | 0.440565 | 0.683589 | 1.205739 | 2.422521 | 5.060366 | 8.163400 | 9.714292 | 8.374246 | 5.729489 | 3.309250 | 2.005911 | 1.180174 | 0.741444 | 0.511754 | 0.385626 | 0.315199 | 0.278171 | 0.255703 | 0.242728 | 0.235749 | 0.232879 | 0.233302 | 0.238112 | 0.247349 | 0.262248 | 0.270269 | 0.265876 | 0.246004 | 0.226700 | 0.215684 | 0.215643 | 0.222782 | 0.235148 | 0.254656 | 0.280491 | 0.315264 | 0.360256 | 0.399674 | 0.403527 | 0.358799 | 0.301441 | 0.258381 | 0.232391 | 0.221065 | 0.218136 | 0.219662 | 0.229874 | 0.267980 | 0.364094 | 0.604052 | 1.041576 | 1.512531 | 1.715152 | 1.600437 | 1.153226 | 0.782413 | 0.495695 | 0.314536 | 0.234339 | 0.214707 | 0.233727 | 0.274473 | 0.323570 | 0.355721 | 0.340028 | 0.303262 | 0.266930 | 0.238548 | 0.225262 | 0.219212 | 0.217277 | 0.215960 | 0.215804 | 0.215761 | 0.217029 | 0.218495 | 0.222567 | 0.223322 | 0.222201 | 0.220281 | 0.218003 | 0.214781 | 0.215746 | 0.219168 | 0.221962 | 0.222077 | 0.217260 | 0.214645 | 0.221324 | 0.242178 | 0.274826 | 0.299322 | 0.326607 | 0.325162 | 0.304566 | 0.290380 | 0.284562 |
| all electrodes | 0.274426 | 0.270445 | 0.236688 | 0.217491 | 0.227072 | 0.336235 | 0.622481 | 0.903689 | 0.958304 | 0.512427 | 0.240288 | 0.235062 | 0.377804 | 0.525762 | 0.594368 | 0.608147 | 0.441177 | 0.377487 | 0.507767 | 1.117018 | 3.021292 | 9.900554 | 22.431678 | 52.414957 | 50.082277 | 17.768117 | 5.681325 | 1.680494 | 0.561638 | 0.327845 | 0.277663 | 0.260205 | 0.251110 | 0.240765 | 0.225195 | 0.214674 | 0.229096 | 0.300789 | 0.444351 | 0.604053 | 0.783614 | 0.856702 | 0.705288 | 0.548975 | 0.428056 | 0.348032 | 0.324423 | 0.322917 | 0.341329 | 0.363284 | 0.375147 | 0.385889 | 0.396134 | 0.379683 | 0.337793 | 0.285254 | 0.242616 | 0.218887 | 0.214829 | 0.217012 | 0.215622 | 0.215349 | 0.238913 | 0.337322 | 0.580080 | 0.897277 | 1.058579 | 0.837868 | 0.571625 | 0.424458 | 0.352545 | 0.326586 | 0.354140 | 0.406265 | 0.493957 | 0.709659 | 1.032199 | 1.460846 | 2.554362 | 3.442079 | 3.807802 | 4.432728 | 4.732609 | 4.462009 | 4.411346 | 4.556974 | 4.820722 | 4.702073 | 4.059359 | 3.085324 | 2.125961 | 1.376762 | 0.878123 | 0.581684 | 0.455524 | 0.391684 | 0.371630 | 0.374286 | 0.385893 | 0.406776 | 0.401026 | 0.380963 | 0.365592 | 0.367745 | 0.363157 | 0.358379 | 0.341324 | 0.347888 | 0.334961 | 0.326837 | 0.305670 | 0.275673 | 0.250270 | 0.236384 | 0.223694 | 0.218531 | 0.215776 | 0.214659 | 0.216653 | 0.220431 | 0.224073 | 0.223119 | 0.218486 | 0.214662 | 0.228453 | 0.282182 | 0.392393 | 0.615706 | 1.034123 | 1.723543 | 2.713310 | 3.776746 | 4.224249 | 4.141805 | 3.678330 | 2.842551 | 2.263088 | 1.755561 | 1.532584 | 1.603298 | 1.783554 | 1.929565 | 2.410121 | 3.225133 | 4.105899 | 5.165475 | 5.345810 | 5.142613 | 4.800995 | 4.498231 | 3.954719 | 3.750344 | 3.187021 | 2.633244 | 2.109476 | 1.715840 | 1.449701 | 1.386818 | 1.296902 | 1.233935 | 1.299019 | 1.403789 | 1.629774 | 1.811903 | 1.820880 | 1.614177 | 1.406991 | 1.150758 | 1.075489 | 1.105053 | 1.406081 | 2.063921 | 2.755444 | 2.972169 | 2.571484 | 1.823368 | 1.219687 | 0.790329 | 0.561302 | 0.448325 | 0.393764 | 0.370341 | 0.380871 | 0.395233 | 0.415564 | 0.408736 | 0.385269 | 0.362851 | 0.349177 | 0.324115 | 0.310941 | 0.313952 | 0.307182 | 0.292109 | 0.279714 | 0.268671 | 0.254061 | 0.246048 | 0.242603 | 0.244085 | 0.256519 | 0.272824 | 0.285240 | 0.299219 | 0.303815 | 0.282976 | 0.263027 | 0.249729 | 0.245372 | 0.248395 | 0.254977 | 0.263905 | 0.273760 | 0.278612 | 0.268531 | 0.249445 | 0.229657 | 0.217535 | 0.215018 | 0.221333 | 0.229748 | 0.229606 | 0.221503 | 0.216697 | 0.214791 | 0.215239 | 0.215857 | 0.215732 | 0.215484 | 0.214730 | 0.214905 | 0.215526 | 0.218647 | 0.223232 | 0.225386 | 0.228552 | 0.230216 | 0.227480 | 0.226447 | 0.224483 | 0.220834 | 0.219946 | 0.221001 | 0.220977 | 0.219645 | 0.217881 | 0.216058 | 0.214991 | 0.214806 | 0.217419 | 0.224057 | 0.229578 | 0.234266 | 0.231528 | 0.228412 | 0.223178 | 0.219897 | 0.217501 | 0.216405 | 0.214724 | 0.215134 | 0.217734 | 0.221762 | 0.224785 | 0.224155 | 0.220741 | 0.216821 | 0.214834 | 0.214640 | 0.214892 | 0.214777 | 0.214678 | 0.214651 | 0.214638 | 0.216529 | 0.221466 | 0.233737 | 0.261806 | 0.286567 | 0.303497 |

K) maximum cross-correlation, sex

  
|  | time window | peak latency | cluster *p* | peak Cohen's *d* |  | | | |  |
| **all electrodes** |  | | | |  | | | |  |
|  | | | | | | | | |  |

Model correlations, cluster permutation tests

|  | **left hemisphere** | | | | **right hemisphere** | | | |  |
|  | time window | peak latency | cluster *p* | peak Cohen's *d* | time window | peak latency | cluster *p* | peak Cohen's *d* |  |
| **anterior** |  | | | |  | | | |  |
| **central** |  | | | | 65 - 205 ms | 85 ms | 0.04 | 0.5606 |  |
  | | | | 420 - 600 ms | 450 ms | 0.0205 | 0.6243 ||  | **posterior** |  | | | | 820 - 1085 ms | 955 ms | 0.0059 | -1.1155 |

  

Model correlations, Bayesian statistics

|  | -200 | -195 | -190 | -185 | -180 | -175 | -170 | -165 | -160 | -155 | -150 | -145 | -140 | -135 | -130 | -125 | -120 | -115 | -110 | -105 | -100 | -95 | -90 | -85 | -80 | -75 | -70 | -65 | -60 | -55 | -50 | -45 | -40 | -35 | -30 | -25 | -20 | -15 | -10 | -5 | 0 | 5 | 10 | 15 | 20 | 25 | 30 | 35 | 40 | 45 | 50 | 55 | 60 | 65 | 70 | 75 | 80 | 85 | 90 | 95 | 100 | 105 | 110 | 115 | 120 | 125 | 130 | 135 | 140 | 145 | 150 | 155 | 160 | 165 | 170 | 175 | 180 | 185 | 190 | 195 | 200 | 205 | 210 | 215 | 220 | 225 | 230 | 235 | 240 | 245 | 250 | 255 | 260 | 265 | 270 | 275 | 280 | 285 | 290 | 295 | 300 | 305 | 310 | 315 | 320 | 325 | 330 | 335 | 340 | 345 | 350 | 355 | 360 | 365 | 370 | 375 | 380 | 385 | 390 | 395 | 400 | 405 | 410 | 415 | 420 | 425 | 430 | 435 | 440 | 445 | 450 | 455 | 460 | 465 | 470 | 475 | 480 | 485 | 490 | 495 | 500 | 505 | 510 | 515 | 520 | 525 | 530 | 535 | 540 | 545 | 550 | 555 | 560 | 565 | 570 | 575 | 580 | 585 | 590 | 595 | 600 | 605 | 610 | 615 | 620 | 625 | 630 | 635 | 640 | 645 | 650 | 655 | 660 | 665 | 670 | 675 | 680 | 685 | 690 | 695 | 700 | 705 | 710 | 715 | 720 | 725 | 730 | 735 | 740 | 745 | 750 | 755 | 760 | 765 | 770 | 775 | 780 | 785 | 790 | 795 | 800 | 805 | 810 | 815 | 820 | 825 | 830 | 835 | 840 | 845 | 850 | 855 | 860 | 865 | 870 | 875 | 880 | 885 | 890 | 895 | 900 | 905 | 910 | 915 | 920 | 925 | 930 | 935 | 940 | 945 | 950 | 955 | 960 | 965 | 970 | 975 | 980 | 985 | 990 | 995 | 1000 | 1005 | 1010 | 1015 | 1020 | 1025 | 1030 | 1035 | 1040 | 1045 | 1050 | 1055 | 1060 | 1065 | 1070 | 1075 | 1080 | 1085 | 1090 | 1095 | 1100 | 1105 | 1110 | 1115 | 1120 | 1125 | 1130 | 1135 | 1140 | 1145 | 1150 | 1155 | 1160 | 1165 | 1170 | 1175 | 1180 | 1185 | 1190 | 1195 |
| --- | --- | --- | --- | --- | --- | --- | --- | --- | --- | --- | --- | --- | --- | --- | --- | --- | --- | --- | --- | --- | --- | --- | --- | --- | --- | --- | --- | --- | --- | --- | --- | --- | --- | --- | --- | --- | --- | --- | --- | --- | --- | --- | --- | --- | --- | --- | --- | --- | --- | --- | --- | --- | --- | --- | --- | --- | --- | --- | --- | --- | --- | --- | --- | --- | --- | --- | --- | --- | --- | --- | --- | --- | --- | --- | --- | --- | --- | --- | --- | --- | --- | --- | --- | --- | --- | --- | --- | --- | --- | --- | --- | --- | --- | --- | --- | --- | --- | --- | --- | --- | --- | --- | --- | --- | --- | --- | --- | --- | --- | --- | --- | --- | --- | --- | --- | --- | --- | --- | --- | --- | --- | --- | --- | --- | --- | --- | --- | --- | --- | --- | --- | --- | --- | --- | --- | --- | --- | --- | --- | --- | --- | --- | --- | --- | --- | --- | --- | --- | --- | --- | --- | --- | --- | --- | --- | --- | --- | --- | --- | --- | --- | --- | --- | --- | --- | --- | --- | --- | --- | --- | --- | --- | --- | --- | --- | --- | --- | --- | --- | --- | --- | --- | --- | --- | --- | --- | --- | --- | --- | --- | --- | --- | --- | --- | --- | --- | --- | --- | --- | --- | --- | --- | --- | --- | --- | --- | --- | --- | --- | --- | --- | --- | --- | --- | --- | --- | --- | --- | --- | --- | --- | --- | --- | --- | --- | --- | --- | --- | --- | --- | --- | --- | --- | --- | --- | --- | --- | --- | --- | --- | --- | --- | --- | --- | --- | --- | --- | --- | --- | --- | --- | --- | --- | --- | --- | --- | --- | --- | --- | --- | --- | --- | --- | --- | --- | --- | --- | --- | --- | --- | --- | --- | --- | --- | --- | --- | --- | --- | --- | --- |
| left anterior | 0.402145 | 0.419149 | 0.441763 | 0.460684 | 0.481905 | 0.473640 | 0.428994 | 0.378265 | 0.310485 | 0.240162 | 0.214959 | 0.227639 | 0.282677 | 0.401704 | 0.525998 | 0.555247 | 0.447952 | 0.281817 | 0.216224 | 0.220479 | 0.220474 | 0.214814 | 0.228131 | 0.285753 | 0.402533 | 0.530824 | 0.556804 | 0.538161 | 0.479153 | 0.445511 | 0.445729 | 0.571445 | 0.869420 | 1.605066 | 2.196393 | 2.394903 | 1.774418 | 0.934671 | 0.475724 | 0.295468 | 0.219443 | 0.233083 | 0.320085 | 0.406316 | 0.420605 | 0.357495 | 0.290777 | 0.253739 | 0.236893 | 0.228147 | 0.232236 | 0.244248 | 0.262308 | 0.303522 | 0.387703 | 0.504421 | 0.705941 | 1.049193 | 1.424001 | 1.797720 | 1.861185 | 1.330668 | 0.736822 | 0.409309 | 0.263380 | 0.222673 | 0.216524 | 0.220792 | 0.235991 | 0.272610 | 0.330006 | 0.384425 | 0.418134 | 0.442369 | 0.461458 | 0.480747 | 0.493518 | 0.448275 | 0.375067 | 0.300432 | 0.248437 | 0.219872 | 0.214850 | 0.220615 | 0.222728 | 0.218094 | 0.214717 | 0.223267 | 0.245087 | 0.270812 | 0.277920 | 0.271989 | 0.259652 | 0.246977 | 0.240084 | 0.246255 | 0.259212 | 0.273417 | 0.288188 | 0.294370 | 0.289581 | 0.267618 | 0.250358 | 0.238108 | 0.232087 | 0.222633 | 0.218414 | 0.215771 | 0.214809 | 0.214708 | 0.216169 | 0.222765 | 0.227620 | 0.230585 | 0.235509 | 0.236529 | 0.235799 | 0.234688 | 0.230224 | 0.223667 | 0.217946 | 0.214843 | 0.219331 | 0.223492 | 0.227086 | 0.232794 | 0.233482 | 0.232140 | 0.228025 | 0.222734 | 0.216902 | 0.214735 | 0.221658 | 0.239869 | 0.273277 | 0.304425 | 0.320285 | 0.313120 | 0.288417 | 0.250526 | 0.227508 | 0.214925 | 0.221047 | 0.239533 | 0.250032 | 0.248773 | 0.231581 | 0.225131 | 0.221185 | 0.218779 | 0.220760 | 0.231548 | 0.246874 | 0.255158 | 0.243744 | 0.224102 | 0.214641 | 0.229768 | 0.273858 | 0.310850 | 0.338939 | 0.346355 | 0.312407 | 0.273904 | 0.243374 | 0.225085 | 0.220972 | 0.221961 | 0.222628 | 0.224967 | 0.225599 | 0.223307 | 0.220836 | 0.218428 | 0.216101 | 0.215658 | 0.215863 | 0.217715 | 0.222539 | 0.233818 | 0.250166 | 0.271981 | 0.292346 | 0.321079 | 0.358202 | 0.409111 | 0.438756 | 0.415499 | 0.340951 | 0.269150 | 0.227671 | 0.214630 | 0.225040 | 0.248653 | 0.267711 | 0.284069 | 0.268130 | 0.244468 | 0.223370 | 0.216625 | 0.215997 | 0.220748 | 0.226833 | 0.241903 | 0.247262 | 0.237885 | 0.222161 | 0.214648 | 0.224011 | 0.248287 | 0.292510 | 0.340820 | 0.387596 | 0.430186 | 0.459805 | 0.457727 | 0.464764 | 0.439203 | 0.403109 | 0.361057 | 0.315756 | 0.283700 | 0.279388 | 0.285225 | 0.301441 | 0.315187 | 0.313887 | 0.298633 | 0.274232 | 0.246369 | 0.229956 | 0.223692 | 0.219498 | 0.217991 | 0.217639 | 0.218251 | 0.216908 | 0.214993 | 0.215413 | 0.219296 | 0.224765 | 0.225265 | 0.224144 | 0.220392 | 0.215729 | 0.214643 | 0.214818 | 0.214634 | 0.217043 | 0.229243 | 0.247496 | 0.267640 | 0.266904 | 0.257188 | 0.242671 | 0.232363 | 0.223132 | 0.220322 | 0.219041 | 0.220909 | 0.220993 | 0.220371 | 0.216469 | 0.214962 | 0.222563 | 0.228619 | 0.228337 | 0.224878 | 0.220881 | 0.217183 | 0.215815 | 0.215583 | 0.216735 | 0.219779 | 0.227348 | 0.238299 | 0.249360 | 0.254927 | 0.256090 | 0.252877 |
| right anterior | 3.757664 | 5.684207 | 9.180563 | 19.310916 | 56.049318 | 182.702196 | 636.648894 | 1421.336881 | 1420.146859 | 228.173956 | 12.143137 | 1.127479 | 0.310191 | 0.217522 | 0.218369 | 0.230559 | 0.238915 | 0.246894 | 0.257197 | 0.279500 | 0.305470 | 0.385360 | 0.520123 | 0.701401 | 0.856154 | 0.930467 | 0.868599 | 0.589253 | 0.367793 | 0.285671 | 0.261058 | 0.236776 | 0.240283 | 0.264427 | 0.287619 | 0.279573 | 0.268254 | 0.228364 | 0.218545 | 0.284038 | 0.429099 | 0.554563 | 0.564212 | 0.457257 | 0.348759 | 0.266894 | 0.235131 | 0.223310 | 0.222113 | 0.226945 | 0.244665 | 0.270113 | 0.298220 | 0.340811 | 0.401648 | 0.456583 | 0.564358 | 0.758379 | 0.983923 | 1.309834 | 1.784803 | 1.994333 | 2.498859 | 2.912058 | 2.974817 | 2.543348 | 2.712463 | 2.435966 | 2.137076 | 1.841324 | 1.669972 | 1.610992 | 1.954899 | 2.379525 | 3.382048 | 4.769166 | 5.364475 | 4.742176 | 3.601013 | 2.093232 | 1.198088 | 0.703830 | 0.455557 | 0.353095 | 0.318366 | 0.322045 | 0.363456 | 0.436705 | 0.533557 | 0.563248 | 0.511694 | 0.404708 | 0.304998 | 0.245088 | 0.224999 | 0.218468 | 0.221350 | 0.239731 | 0.293154 | 0.419340 | 0.632873 | 0.830751 | 0.906119 | 0.830025 | 0.629649 | 0.495357 | 0.383149 | 0.316242 | 0.279509 | 0.254113 | 0.238712 | 0.235043 | 0.232200 | 0.233063 | 0.240754 | 0.254360 | 0.270038 | 0.308624 | 0.338039 | 0.347842 | 0.352917 | 0.367441 | 0.387901 | 0.460804 | 0.574502 | 0.735463 | 1.041295 | 1.290518 | 1.248944 | 1.117858 | 0.927577 | 0.676461 | 0.498470 | 0.383007 | 0.317927 | 0.280972 | 0.260767 | 0.244866 | 0.236533 | 0.230931 | 0.225849 | 0.220276 | 0.218334 | 0.217137 | 0.215827 | 0.215218 | 0.215364 | 0.215869 | 0.218476 | 0.223617 | 0.234382 | 0.263588 | 0.315627 | 0.384989 | 0.466514 | 0.533221 | 0.568342 | 0.556413 | 0.484955 | 0.401301 | 0.330048 | 0.265384 | 0.228736 | 0.215951 | 0.214718 | 0.215296 | 0.214673 | 0.218894 | 0.239433 | 0.284241 | 0.353066 | 0.409223 | 0.452510 | 0.461744 | 0.450074 | 0.426615 | 0.417602 | 0.427944 | 0.445978 | 0.455794 | 0.453160 | 0.435059 | 0.388248 | 0.345474 | 0.305836 | 0.288934 | 0.291239 | 0.309112 | 0.322949 | 0.355923 | 0.394534 | 0.410646 | 0.432384 | 0.480763 | 0.552621 | 0.655706 | 0.713731 | 0.650265 | 0.573971 | 0.501337 | 0.445352 | 0.420339 | 0.433297 | 0.484210 | 0.559758 | 0.617311 | 0.690758 | 0.726044 | 0.671554 | 0.560925 | 0.442051 | 0.333694 | 0.265389 | 0.231069 | 0.216767 | 0.214633 | 0.214627 | 0.215010 | 0.218094 | 0.227528 | 0.249709 | 0.281484 | 0.335661 | 0.388367 | 0.476102 | 0.570266 | 0.605421 | 0.573068 | 0.510561 | 0.377626 | 0.290420 | 0.250177 | 0.234129 | 0.232068 | 0.243409 | 0.265673 | 0.306656 | 0.348568 | 0.369215 | 0.359547 | 0.333182 | 0.297258 | 0.282368 | 0.276354 | 0.273430 | 0.273450 | 0.272938 | 0.262122 | 0.244224 | 0.227618 | 0.216628 | 0.214964 | 0.222357 | 0.234767 | 0.245672 | 0.246354 | 0.240086 | 0.223346 | 0.214627 | 0.228131 | 0.272647 | 0.356348 | 0.466555 | 0.584456 | 0.658019 | 0.680131 | 0.667046 | 0.620701 | 0.585182 | 0.581233 | 0.566468 | 0.505622 | 0.433876 | 0.371780 | 0.330578 | 0.312291 | 0.311333 | 0.324600 | 0.326149 | 0.328952 |
| left central | 0.300575 | 0.340959 | 0.382932 | 0.414183 | 0.389766 | 0.290893 | 0.224685 | 0.227269 | 0.322211 | 0.479737 | 0.526439 | 0.402620 | 0.247732 | 0.215475 | 0.263805 | 0.331019 | 0.298272 | 0.228989 | 0.214632 | 0.218760 | 0.216737 | 0.218912 | 0.258901 | 0.384278 | 0.581799 | 0.640227 | 0.556289 | 0.525486 | 0.534960 | 0.613338 | 0.819414 | 1.075116 | 1.307346 | 1.232418 | 0.772223 | 0.445523 | 0.310283 | 0.250091 | 0.239404 | 0.264848 | 0.371410 | 0.612300 | 0.686709 | 0.554321 | 0.434867 | 0.332745 | 0.273368 | 0.247334 | 0.228764 | 0.219881 | 0.215518 | 0.214632 | 0.215494 | 0.221705 | 0.239589 | 0.260926 | 0.303714 | 0.337925 | 0.309090 | 0.257235 | 0.223646 | 0.214638 | 0.220625 | 0.232322 | 0.235774 | 0.225840 | 0.221545 | 0.217642 | 0.214628 | 0.218222 | 0.221264 | 0.224834 | 0.226367 | 0.229644 | 0.234310 | 0.235269 | 0.234024 | 0.230801 | 0.222391 | 0.215730 | 0.214810 | 0.217023 | 0.218946 | 0.216779 | 0.214636 | 0.217326 | 0.227151 | 0.246222 | 0.277623 | 0.310820 | 0.353417 | 0.387730 | 0.440428 | 0.486100 | 0.534818 | 0.620989 | 0.830954 | 1.139583 | 1.484145 | 1.618449 | 1.570599 | 1.354635 | 1.087249 | 0.877892 | 0.774017 | 0.704550 | 0.657348 | 0.653638 | 0.611288 | 0.521345 | 0.444966 | 0.373581 | 0.322250 | 0.296172 | 0.292105 | 0.305172 | 0.351581 | 0.447763 | 0.641299 | 1.009873 | 1.564063 | 2.223234 | 2.799353 | 2.901007 | 2.542691 | 2.146299 | 1.714897 | 1.407372 | 1.141525 | 0.913663 | 0.749113 | 0.629995 | 0.542365 | 0.491615 | 0.480543 | 0.478890 | 0.514916 | 0.591605 | 0.711597 | 0.883485 | 1.222770 | 1.856510 | 3.071578 | 5.046985 | 7.255182 | 9.033631 | 9.764174 | 9.312369 | 8.492836 | 7.767137 | 6.767969 | 5.146153 | 3.632301 | 2.590279 | 1.860613 | 1.260438 | 0.834592 | 0.584719 | 0.460987 | 0.397061 | 0.372681 | 0.362390 | 0.367238 | 0.385323 | 0.416649 | 0.453997 | 0.494260 | 0.517297 | 0.503095 | 0.473551 | 0.436356 | 0.397004 | 0.344767 | 0.307374 | 0.284265 | 0.271247 | 0.266897 | 0.272541 | 0.281203 | 0.298470 | 0.323948 | 0.340617 | 0.343129 | 0.316549 | 0.282217 | 0.257999 | 0.246400 | 0.243294 | 0.266967 | 0.329897 | 0.488129 | 0.819130 | 1.465207 | 2.287192 | 3.116217 | 2.795475 | 1.921205 | 1.222907 | 0.836169 | 0.598316 | 0.483645 | 0.439023 | 0.447262 | 0.492127 | 0.584832 | 0.710396 | 0.855976 | 0.977529 | 1.013670 | 0.925644 | 0.818759 | 0.710233 | 0.581549 | 0.466814 | 0.364812 | 0.289058 | 0.247593 | 0.227971 | 0.216558 | 0.214674 | 0.215991 | 0.219621 | 0.226515 | 0.237978 | 0.259641 | 0.300330 | 0.372039 | 0.501317 | 0.587487 | 0.593946 | 0.514146 | 0.390237 | 0.282680 | 0.226658 | 0.215692 | 0.238529 | 0.275828 | 0.317683 | 0.336362 | 0.342781 | 0.341699 | 0.337356 | 0.318755 | 0.296662 | 0.270121 | 0.245343 | 0.226640 | 0.217009 | 0.214656 | 0.214923 | 0.214726 | 0.214698 | 0.215888 | 0.217472 | 0.219175 | 0.218990 | 0.219669 | 0.220590 | 0.223237 | 0.224568 | 0.226245 | 0.225115 | 0.224480 | 0.223725 | 0.223050 | 0.221830 | 0.221717 | 0.222570 | 0.222801 | 0.222233 | 0.220954 | 0.218420 | 0.215578 | 0.214632 | 0.216184 | 0.221273 | 0.230214 | 0.243652 | 0.255672 | 0.268582 |
| right central | 1.626283 | 1.862435 | 2.024793 | 1.564736 | 0.999569 | 0.600568 | 0.384901 | 0.260575 | 0.217089 | 0.218884 | 0.255617 | 0.363106 | 0.578746 | 0.987537 | 2.219319 | 5.347563 | 10.316806 | 23.181143 | 41.349781 | 68.637487 | 103.275563 | 115.974339 | 66.359587 | 26.763931 | 4.882274 | 0.979592 | 0.354315 | 0.224549 | 0.226224 | 0.272859 | 0.313986 | 0.337417 | 0.322690 | 0.291765 | 0.280324 | 0.272945 | 0.263868 | 0.251844 | 0.233715 | 0.221059 | 0.214759 | 0.220891 | 0.224033 | 0.217341 | 0.215217 | 0.238784 | 0.297891 | 0.392405 | 0.501805 | 0.608623 | 0.740547 | 0.945692 | 1.248205 | 1.602076 | 2.014510 | 2.738339 | 3.907933 | 4.417780 | 4.830272 | 4.713316 | 4.256918 | 3.492341 | 2.670164 | 1.888883 | 1.606940 | 1.492664 | 1.633612 | 1.966786 | 2.624380 | 2.707977 | 2.620426 | 2.553920 | 3.089050 | 3.071499 | 3.056215 | 2.921774 | 2.953595 | 2.928000 | 3.360936 | 3.615892 | 3.116615 | 1.711023 | 0.732676 | 0.386822 | 0.295592 | 0.279226 | 0.311706 | 0.402019 | 0.572028 | 0.920676 | 1.616468 | 3.035787 | 5.305054 | 7.329885 | 7.756176 | 6.464950 | 4.520324 | 3.183308 | 2.408763 | 1.919157 | 1.576313 | 1.334442 | 1.283699 | 1.358348 | 1.471598 | 1.505177 | 1.408887 | 1.253956 | 1.160462 | 1.023755 | 0.941374 | 0.957318 | 1.059803 | 1.248224 | 1.483795 | 1.519888 | 1.336371 | 1.099871 | 0.879944 | 0.756778 | 0.724286 | 0.791974 | 0.869244 | 1.026965 | 1.308581 | 1.766854 | 2.373796 | 3.149227 | 3.951687 | 5.610342 | 8.162200 | 10.923472 | 11.788230 | 10.529318 | 7.574276 | 5.434722 | 3.849672 | 3.116003 | 2.310767 | 1.807449 | 1.522568 | 1.537768 | 1.787476 | 2.341450 | 2.798205 | 3.778517 | 5.409304 | 6.649757 | 7.770608 | 7.807674 | 6.591320 | 5.590730 | 4.599147 | 3.686539 | 3.226260 | 2.787887 | 2.488284 | 2.486707 | 2.451358 | 2.118369 | 1.473632 | 0.913541 | 0.638773 | 0.555551 | 0.570042 | 0.683907 | 0.973786 | 1.476715 | 2.282209 | 3.383278 | 4.676772 | 5.213589 | 4.598585 | 3.180842 | 2.090945 | 1.410756 | 1.006050 | 0.705698 | 0.557148 | 0.475633 | 0.392852 | 0.352067 | 0.322309 | 0.298082 | 0.279221 | 0.278235 | 0.281643 | 0.305282 | 0.348160 | 0.411731 | 0.480110 | 0.562604 | 0.638109 | 0.716060 | 0.804029 | 0.860671 | 0.808012 | 0.672741 | 0.524641 | 0.385430 | 0.294949 | 0.251057 | 0.230871 | 0.220830 | 0.219126 | 0.223981 | 0.241487 | 0.274503 | 0.318192 | 0.361215 | 0.366585 | 0.307556 | 0.251467 | 0.220384 | 0.216287 | 0.237248 | 0.269308 | 0.297489 | 0.307881 | 0.302851 | 0.290340 | 0.275303 | 0.258755 | 0.244572 | 0.233369 | 0.224542 | 0.220491 | 0.219199 | 0.220767 | 0.226617 | 0.240982 | 0.250602 | 0.245004 | 0.231739 | 0.221103 | 0.215265 | 0.215717 | 0.222487 | 0.234949 | 0.251901 | 0.265027 | 0.266650 | 0.267582 | 0.260165 | 0.248664 | 0.238872 | 0.232659 | 0.226556 | 0.224369 | 0.223146 | 0.226163 | 0.235221 | 0.251746 | 0.274220 | 0.312742 | 0.361328 | 0.404487 | 0.401393 | 0.353060 | 0.302730 | 0.262389 | 0.233907 | 0.220236 | 0.215726 | 0.214829 | 0.215007 | 0.216818 | 0.223547 | 0.237430 | 0.256706 | 0.278669 | 0.301317 | 0.317845 | 0.310995 | 0.295459 | 0.281943 | 0.270419 | 0.254159 | 0.249190 | 0.244530 |
| left posterior | 0.892845 | 0.949558 | 1.081104 | 1.504607 | 2.309948 | 3.619741 | 4.690062 | 4.211065 | 2.065773 | 0.775422 | 0.364698 | 0.231591 | 0.219185 | 0.254544 | 0.274856 | 0.283052 | 0.267873 | 0.243799 | 0.231546 | 0.240331 | 0.261513 | 0.296029 | 0.370625 | 0.546518 | 0.723120 | 0.708047 | 0.574172 | 0.444182 | 0.344896 | 0.298261 | 0.308270 | 0.354627 | 0.432868 | 0.400569 | 0.304461 | 0.230736 | 0.220886 | 0.313776 | 0.512492 | 0.856100 | 1.335286 | 1.604799 | 1.648558 | 1.322310 | 0.831329 | 0.521667 | 0.343534 | 0.266992 | 0.241056 | 0.225310 | 0.218396 | 0.216868 | 0.219851 | 0.238436 | 0.320917 | 0.490777 | 0.756793 | 1.051132 | 1.188842 | 1.013088 | 0.658305 | 0.394503 | 0.258824 | 0.215161 | 0.242328 | 0.355663 | 0.622469 | 1.051763 | 1.836558 | 2.878103 | 2.649100 | 1.533669 | 0.756959 | 0.362568 | 0.238115 | 0.214964 | 0.232211 | 0.251094 | 0.246871 | 0.231289 | 0.217618 | 0.214720 | 0.216719 | 0.215938 | 0.214788 | 0.220688 | 0.232886 | 0.243061 | 0.238424 | 0.222816 | 0.215145 | 0.216139 | 0.223956 | 0.234448 | 0.244230 | 0.259917 | 0.275755 | 0.294676 | 0.324453 | 0.350576 | 0.357206 | 0.353630 | 0.336436 | 0.315390 | 0.296698 | 0.273961 | 0.253394 | 0.236138 | 0.225527 | 0.217855 | 0.215086 | 0.214664 | 0.214817 | 0.214693 | 0.215416 | 0.220249 | 0.234632 | 0.261392 | 0.314373 | 0.393256 | 0.483087 | 0.523023 | 0.474091 | 0.369874 | 0.277725 | 0.228286 | 0.214739 | 0.216185 | 0.217179 | 0.214662 | 0.221208 | 0.250764 | 0.299443 | 0.362046 | 0.388107 | 0.396137 | 0.398749 | 0.413913 | 0.453610 | 0.525526 | 0.608102 | 0.703759 | 0.715236 | 0.617215 | 0.494124 | 0.398053 | 0.328644 | 0.277937 | 0.246722 | 0.231859 | 0.224116 | 0.220909 | 0.218918 | 0.216788 | 0.215238 | 0.214673 | 0.216946 | 0.219738 | 0.223393 | 0.225303 | 0.223526 | 0.219276 | 0.216023 | 0.214706 | 0.216707 | 0.219670 | 0.221864 | 0.220356 | 0.216698 | 0.214742 | 0.215148 | 0.217168 | 0.215894 | 0.214730 | 0.221715 | 0.241270 | 0.266275 | 0.273445 | 0.260925 | 0.231399 | 0.214699 | 0.229297 | 0.287602 | 0.423605 | 0.656679 | 0.982041 | 1.398387 | 1.897082 | 2.261689 | 2.339164 | 1.941191 | 1.408803 | 0.933016 | 0.602207 | 0.423984 | 0.333335 | 0.289584 | 0.273882 | 0.280320 | 0.305409 | 0.374884 | 0.509863 | 0.753734 | 1.219285 | 2.175940 | 3.655281 | 6.022584 | 8.480954 | 9.543700 | 8.325136 | 5.616842 | 2.855969 | 1.304321 | 0.618567 | 0.357708 | 0.263281 | 0.228723 | 0.217063 | 0.214626 | 0.215058 | 0.215596 | 0.215052 | 0.214654 | 0.217315 | 0.224100 | 0.234032 | 0.242030 | 0.252455 | 0.258715 | 0.260911 | 0.268468 | 0.290087 | 0.326172 | 0.374260 | 0.422218 | 0.423528 | 0.377465 | 0.309983 | 0.263181 | 0.234958 | 0.221686 | 0.216490 | 0.216183 | 0.216934 | 0.220066 | 0.225210 | 0.232946 | 0.235599 | 0.235112 | 0.227844 | 0.222957 | 0.217830 | 0.214676 | 0.218572 | 0.232014 | 0.264177 | 0.294205 | 0.291025 | 0.253557 | 0.224520 | 0.214637 | 0.221321 | 0.245428 | 0.278996 | 0.315755 | 0.352235 | 0.375657 | 0.379526 | 0.367092 | 0.349366 | 0.332269 | 0.321187 | 0.311164 | 0.316287 | 0.329372 | 0.352771 | 0.375754 | 0.393074 | 0.393221 | 0.396571 |
| right posterior | 0.393062 | 0.411939 | 0.470359 | 0.488748 | 0.416362 | 0.340135 | 0.284631 | 0.248354 | 0.228116 | 0.217793 | 0.214719 | 0.218641 | 0.229252 | 0.248052 | 0.260405 | 0.271698 | 0.311550 | 0.448922 | 0.982869 | 2.506662 | 4.952887 | 4.918762 | 2.367038 | 0.716814 | 0.300689 | 0.216051 | 0.235008 | 0.302424 | 0.365945 | 0.339855 | 0.275758 | 0.223344 | 0.220240 | 0.265312 | 0.311309 | 0.299174 | 0.245795 | 0.219801 | 0.214685 | 0.215444 | 0.214674 | 0.217902 | 0.236930 | 0.279940 | 0.334563 | 0.366550 | 0.391394 | 0.384627 | 0.358757 | 0.342898 | 0.338288 | 0.311199 | 0.256096 | 0.217886 | 0.228354 | 0.280569 | 0.338403 | 0.342469 | 0.294521 | 0.240670 | 0.215932 | 0.226337 | 0.282170 | 0.417825 | 0.695900 | 1.099320 | 1.267221 | 1.161362 | 0.927108 | 0.676015 | 0.461727 | 0.323072 | 0.251237 | 0.218673 | 0.216544 | 0.227560 | 0.236923 | 0.240968 | 0.238563 | 0.231541 | 0.220445 | 0.214627 | 0.221302 | 0.239170 | 0.268688 | 0.299927 | 0.317384 | 0.314340 | 0.303679 | 0.275116 | 0.242558 | 0.222928 | 0.215797 | 0.214658 | 0.214750 | 0.214635 | 0.216785 | 0.229249 | 0.258658 | 0.295797 | 0.341712 | 0.372903 | 0.377965 | 0.350886 | 0.314412 | 0.280231 | 0.256762 | 0.236896 | 0.225714 | 0.219718 | 0.216341 | 0.214672 | 0.215183 | 0.217337 | 0.223028 | 0.233789 | 0.250185 | 0.271892 | 0.304654 | 0.342250 | 0.388275 | 0.448488 | 0.558177 | 0.735616 | 1.083082 | 1.689513 | 2.701582 | 3.980074 | 5.073885 | 5.108253 | 4.410026 | 3.148452 | 1.985282 | 1.145665 | 0.654962 | 0.402586 | 0.292266 | 0.244786 | 0.226807 | 0.218793 | 0.215429 | 0.214643 | 0.214672 | 0.214676 | 0.214707 | 0.216964 | 0.224546 | 0.237398 | 0.250220 | 0.259161 | 0.255143 | 0.244612 | 0.234875 | 0.225626 | 0.220382 | 0.218078 | 0.216159 | 0.214703 | 0.214742 | 0.215918 | 0.218322 | 0.219836 | 0.217296 | 0.214911 | 0.216200 | 0.223787 | 0.234099 | 0.238749 | 0.237303 | 0.228078 | 0.220606 | 0.215727 | 0.214627 | 0.215709 | 0.218090 | 0.222630 | 0.231242 | 0.252455 | 0.292830 | 0.356206 | 0.414604 | 0.443075 | 0.424109 | 0.357829 | 0.282555 | 0.232542 | 0.214970 | 0.220054 | 0.231901 | 0.239588 | 0.238515 | 0.232518 | 0.226261 | 0.221102 | 0.217666 | 0.216197 | 0.215588 | 0.214769 | 0.215133 | 0.222496 | 0.251344 | 0.330006 | 0.513794 | 0.875332 | 1.465519 | 2.281417 | 3.285130 | 4.087365 | 4.289118 | 3.834216 | 3.406917 | 3.178266 | 3.064582 | 2.907983 | 2.929777 | 2.998494 | 3.140618 | 3.566143 | 4.632677 | 6.153910 | 8.775571 | 12.453920 | 16.184708 | 19.566630 | 25.461665 | 35.138399 | 59.477966 | 116.848300 | 237.832514 | 472.042146 | 874.374900 | 1450.321401 | 1847.232772 | 1789.522504 | 1092.292863 | 507.338869 | 148.028194 | 42.675856 | 13.014914 | 5.761174 | 3.300883 | 2.436672 | 1.956537 | 1.906471 | 1.958277 | 2.049322 | 2.241173 | 2.686083 | 3.243972 | 3.944132 | 4.689950 | 5.019463 | 5.057034 | 4.906756 | 4.447762 | 3.649344 | 2.634849 | 1.567376 | 0.935354 | 0.607186 | 0.449242 | 0.374192 | 0.342554 | 0.326636 | 0.324591 | 0.332775 | 0.352510 | 0.374315 | 0.397879 | 0.408764 | 0.400307 | 0.361969 | 0.321950 | 0.289554 | 0.277248 | 0.271701 | 0.275566 | 0.295928 | 0.307699 | 0.321333 |
| all electrodes | 0.351455 | 0.388159 | 0.444615 | 0.620127 | 1.075399 | 2.451176 | 4.755412 | 6.754617 | 5.982962 | 1.830567 | 0.319892 | 0.217171 | 0.286324 | 0.412199 | 0.456439 | 0.452430 | 0.360717 | 0.320098 | 0.392040 | 0.828368 | 1.258142 | 1.443150 | 1.549558 | 1.625094 | 1.369803 | 1.117181 | 1.178844 | 0.954285 | 0.625967 | 0.428206 | 0.359311 | 0.295876 | 0.266362 | 0.223756 | 0.219309 | 0.274290 | 0.408732 | 0.632328 | 0.762099 | 0.782967 | 0.683629 | 0.461215 | 0.343100 | 0.251257 | 0.215516 | 0.225587 | 0.241238 | 0.262122 | 0.270162 | 0.269969 | 0.246860 | 0.227958 | 0.216189 | 0.217431 | 0.247806 | 0.339133 | 0.543944 | 0.872663 | 1.113571 | 1.203754 | 1.025552 | 0.807585 | 0.603696 | 0.453961 | 0.337955 | 0.273697 | 0.238920 | 0.228958 | 0.225264 | 0.224157 | 0.226791 | 0.245266 | 0.278584 | 0.350004 | 0.477940 | 0.686836 | 0.979727 | 1.505601 | 2.161422 | 2.981479 | 3.547328 | 3.952603 | 3.471564 | 2.830323 | 2.089543 | 1.790162 | 1.754983 | 1.843175 | 1.702710 | 1.600480 | 1.458972 | 1.144020 | 0.878572 | 0.759569 | 0.751245 | 0.874780 | 0.994188 | 1.014761 | 0.813051 | 0.570557 | 0.363405 | 0.271066 | 0.229880 | 0.219895 | 0.219325 | 0.232723 | 0.277789 | 0.404468 | 0.620072 | 0.913754 | 1.082489 | 0.993265 | 0.772895 | 0.586462 | 0.438005 | 0.346097 | 0.298428 | 0.276224 | 0.280782 | 0.308971 | 0.350475 | 0.420281 | 0.539919 | 0.693561 | 0.918379 | 1.195144 | 1.398211 | 1.439373 | 1.260142 | 1.011288 | 0.842391 | 0.651062 | 0.469880 | 0.359739 | 0.289155 | 0.243742 | 0.221050 | 0.214646 | 0.217964 | 0.227479 | 0.241525 | 0.243557 | 0.235177 | 0.222032 | 0.215428 | 0.215286 | 0.223362 | 0.239909 | 0.248481 | 0.252800 | 0.253164 | 0.246154 | 0.239685 | 0.234841 | 0.228680 | 0.224556 | 0.220053 | 0.216702 | 0.215510 | 0.214750 | 0.214896 | 0.216575 | 0.219726 | 0.224803 | 0.232717 | 0.248051 | 0.261826 | 0.274623 | 0.275474 | 0.253952 | 0.230786 | 0.218811 | 0.214953 | 0.214790 | 0.215221 | 0.216183 | 0.215915 | 0.215344 | 0.214784 | 0.214784 | 0.214632 | 0.215264 | 0.218313 | 0.225380 | 0.241616 | 0.260913 | 0.265839 | 0.250293 | 0.227461 | 0.215598 | 0.217084 | 0.230107 | 0.246293 | 0.245222 | 0.235378 | 0.223686 | 0.216497 | 0.214642 | 0.214950 | 0.214630 | 0.217302 | 0.229351 | 0.255425 | 0.288983 | 0.296405 | 0.267766 | 0.235020 | 0.217897 | 0.214859 | 0.218429 | 0.221863 | 0.222460 | 0.222464 | 0.223850 | 0.226495 | 0.235322 | 0.256777 | 0.292271 | 0.334604 | 0.381007 | 0.397565 | 0.422937 | 0.430582 | 0.424987 | 0.413384 | 0.415291 | 0.375675 | 0.360895 | 0.357230 | 0.366830 | 0.371065 | 0.380796 | 0.360096 | 0.333110 | 0.284216 | 0.244754 | 0.220464 | 0.214646 | 0.218794 | 0.225517 | 0.228903 | 0.225734 | 0.220268 | 0.216514 | 0.214630 | 0.217497 | 0.224450 | 0.231777 | 0.232664 | 0.227777 | 0.218754 | 0.214629 | 0.220130 | 0.237230 | 0.265666 | 0.293425 | 0.318789 | 0.332266 | 0.345688 | 0.368527 | 0.424044 | 0.501895 | 0.666049 | 0.924801 | 1.296308 | 1.594010 | 1.871968 | 1.885787 | 1.840249 | 1.705900 | 1.690950 | 1.778702 | 1.732152 | 1.465205 | 1.211726 | 0.987646 | 0.808279 | 0.640712 | 0.578593 | 0.528383 |

L) maximum cross-correlation, emotion

  
|  | time window | peak latency | cluster *p* | peak Cohen's *d* |  | | | |  |
| **all electrodes** | 125 - 225 ms | 160 ms | 0.0212 | 1.4434 |  | | | |  |
 345 - 560 ms | 395 ms | 0.0042 | 0.9521 |  | | | ||  |  | | | | | | | | |

Model correlations, cluster permutation tests

|  | **left hemisphere** | | | | **right hemisphere** | | | |  |
|  | time window | peak latency | cluster *p* | peak Cohen's *d* | time window | peak latency | cluster *p* | peak Cohen's *d* |  |
| **anterior** |  | | | |  | | | |  |
| **central** |  | | | | 210 - 495 ms | 275 ms | 0.0001 | 1.7929 |  |
| **posterior** | 120 - 225 ms | 160 ms | 0.0212 | 1.4779 | 260 - 450 ms | 410 ms | 0.0255 | 0.9059 |  |
 250 - 440 ms | 410 ms | 0.0185 | 0.8164 | 730 - 1120 ms | 830 ms | 0.0048 | 0.6782 |

  

Model correlations, Bayesian statistics

|  | -200 | -195 | -190 | -185 | -180 | -175 | -170 | -165 | -160 | -155 | -150 | -145 | -140 | -135 | -130 | -125 | -120 | -115 | -110 | -105 | -100 | -95 | -90 | -85 | -80 | -75 | -70 | -65 | -60 | -55 | -50 | -45 | -40 | -35 | -30 | -25 | -20 | -15 | -10 | -5 | 0 | 5 | 10 | 15 | 20 | 25 | 30 | 35 | 40 | 45 | 50 | 55 | 60 | 65 | 70 | 75 | 80 | 85 | 90 | 95 | 100 | 105 | 110 | 115 | 120 | 125 | 130 | 135 | 140 | 145 | 150 | 155 | 160 | 165 | 170 | 175 | 180 | 185 | 190 | 195 | 200 | 205 | 210 | 215 | 220 | 225 | 230 | 235 | 240 | 245 | 250 | 255 | 260 | 265 | 270 | 275 | 280 | 285 | 290 | 295 | 300 | 305 | 310 | 315 | 320 | 325 | 330 | 335 | 340 | 345 | 350 | 355 | 360 | 365 | 370 | 375 | 380 | 385 | 390 | 395 | 400 | 405 | 410 | 415 | 420 | 425 | 430 | 435 | 440 | 445 | 450 | 455 | 460 | 465 | 470 | 475 | 480 | 485 | 490 | 495 | 500 | 505 | 510 | 515 | 520 | 525 | 530 | 535 | 540 | 545 | 550 | 555 | 560 | 565 | 570 | 575 | 580 | 585 | 590 | 595 | 600 | 605 | 610 | 615 | 620 | 625 | 630 | 635 | 640 | 645 | 650 | 655 | 660 | 665 | 670 | 675 | 680 | 685 | 690 | 695 | 700 | 705 | 710 | 715 | 720 | 725 | 730 | 735 | 740 | 745 | 750 | 755 | 760 | 765 | 770 | 775 | 780 | 785 | 790 | 795 | 800 | 805 | 810 | 815 | 820 | 825 | 830 | 835 | 840 | 845 | 850 | 855 | 860 | 865 | 870 | 875 | 880 | 885 | 890 | 895 | 900 | 905 | 910 | 915 | 920 | 925 | 930 | 935 | 940 | 945 | 950 | 955 | 960 | 965 | 970 | 975 | 980 | 985 | 990 | 995 | 1000 | 1005 | 1010 | 1015 | 1020 | 1025 | 1030 | 1035 | 1040 | 1045 | 1050 | 1055 | 1060 | 1065 | 1070 | 1075 | 1080 | 1085 | 1090 | 1095 | 1100 | 1105 | 1110 | 1115 | 1120 | 1125 | 1130 | 1135 | 1140 | 1145 | 1150 | 1155 | 1160 | 1165 | 1170 | 1175 | 1180 | 1185 | 1190 | 1195 |
| --- | --- | --- | --- | --- | --- | --- | --- | --- | --- | --- | --- | --- | --- | --- | --- | --- | --- | --- | --- | --- | --- | --- | --- | --- | --- | --- | --- | --- | --- | --- | --- | --- | --- | --- | --- | --- | --- | --- | --- | --- | --- | --- | --- | --- | --- | --- | --- | --- | --- | --- | --- | --- | --- | --- | --- | --- | --- | --- | --- | --- | --- | --- | --- | --- | --- | --- | --- | --- | --- | --- | --- | --- | --- | --- | --- | --- | --- | --- | --- | --- | --- | --- | --- | --- | --- | --- | --- | --- | --- | --- | --- | --- | --- | --- | --- | --- | --- | --- | --- | --- | --- | --- | --- | --- | --- | --- | --- | --- | --- | --- | --- | --- | --- | --- | --- | --- | --- | --- | --- | --- | --- | --- | --- | --- | --- | --- | --- | --- | --- | --- | --- | --- | --- | --- | --- | --- | --- | --- | --- | --- | --- | --- | --- | --- | --- | --- | --- | --- | --- | --- | --- | --- | --- | --- | --- | --- | --- | --- | --- | --- | --- | --- | --- | --- | --- | --- | --- | --- | --- | --- | --- | --- | --- | --- | --- | --- | --- | --- | --- | --- | --- | --- | --- | --- | --- | --- | --- | --- | --- | --- | --- | --- | --- | --- | --- | --- | --- | --- | --- | --- | --- | --- | --- | --- | --- | --- | --- | --- | --- | --- | --- | --- | --- | --- | --- | --- | --- | --- | --- | --- | --- | --- | --- | --- | --- | --- | --- | --- | --- | --- | --- | --- | --- | --- | --- | --- | --- | --- | --- | --- | --- | --- | --- | --- | --- | --- | --- | --- | --- | --- | --- | --- | --- | --- | --- | --- | --- | --- | --- | --- | --- | --- | --- | --- | --- | --- | --- | --- | --- | --- | --- | --- | --- | --- | --- | --- | --- | --- | --- | --- |
| left anterior | 0.234029 | 0.248078 | 0.248626 | 0.256199 | 0.253734 | 0.247302 | 0.236353 | 0.231172 | 0.265926 | 0.327860 | 0.365165 | 0.427959 | 0.512179 | 0.467549 | 0.401666 | 0.320668 | 0.278867 | 0.271051 | 0.266314 | 0.268593 | 0.269029 | 0.269844 | 0.250872 | 0.216449 | 0.249758 | 0.372287 | 0.774860 | 1.628084 | 2.541600 | 2.480390 | 1.268254 | 0.388691 | 0.214679 | 0.511091 | 2.559791 | 6.211966 | 7.073866 | 5.391490 | 3.624020 | 2.664327 | 2.290933 | 2.399429 | 4.724738 | 9.350467 | 22.646958 | 19.507329 | 6.347745 | 1.453664 | 0.625949 | 0.349661 | 0.282981 | 0.267363 | 0.334638 | 0.473902 | 0.733370 | 0.889004 | 0.916265 | 0.670656 | 0.554673 | 0.477790 | 0.379646 | 0.305868 | 0.263075 | 0.221371 | 0.222289 | 0.278466 | 0.413040 | 0.633824 | 1.029415 | 1.228879 | 1.327622 | 1.166112 | 1.089162 | 0.877771 | 0.656380 | 0.399892 | 0.268746 | 0.215340 | 0.255086 | 0.454353 | 0.877152 | 1.123284 | 0.967338 | 0.745599 | 0.501702 | 0.334398 | 0.261627 | 0.227163 | 0.215292 | 0.216814 | 0.224629 | 0.235442 | 0.242845 | 0.258175 | 0.262961 | 2.454442e-01 | 0.223528 | 0.214682 | 0.229616 | 0.284620 | 0.384720 | 0.493419 | 0.550908 | 0.528407 | 0.462232 | 0.373170 | 0.288390 | 0.240715 | 0.220185 | 0.215525 | 0.214629 | 0.214628 | 0.214695 | 0.214817 | 0.215616 | 0.219039 | 0.234541 | 0.266015 | 0.317950 | 0.380395 | 0.404914 | 0.390505 | 0.342587 | 0.281130 | 0.241518 | 0.223318 | 0.215417 | 0.215044 | 0.215961 | 0.215938 | 0.214720 | 0.214751 | 0.215131 | 0.216222 | 0.218370 | 0.217224 | 0.215415 | 0.214653 | 0.215065 | 0.216410 | 0.218373 | 0.220001 | 0.221091 | 0.220092 | 0.219321 | 0.216177 | 0.214646 | 0.214954 | 0.214630 | 0.215929 | 0.219777 | 0.228458 | 0.238927 | 0.250555 | 0.250981 | 0.246290 | 0.235091 | 0.222844 | 0.216167 | 0.214685 | 0.216280 | 0.216526 | 0.215281 | 0.214668 | 0.214790 | 0.214968 | 0.214666 | 0.215381 | 0.221954 | 0.231456 | 0.234947 | 0.225242 | 0.216787 | 0.215939 | 0.226881 | 0.242848 | 0.238925 | 0.226333 | 0.215303 | 0.217630 | 0.235212 | 0.251929 | 0.257698 | 0.250299 | 0.244043 | 0.233396 | 0.223465 | 0.216810 | 0.215210 | 0.214633 | 0.217907 | 0.230263 | 0.245135 | 0.254702 | 0.244783 | 0.219652 | 0.227710 | 0.343377 | 0.633776 | 1.061297 | 1.574072 | 1.697438 | 1.179006 | 0.594584 | 0.344473 | 0.251953 | 0.224268 | 0.219391 | 0.225756 | 0.246919 | 0.287125 | 0.333363 | 0.395598 | 0.494666 | 0.627620 | 0.771965 | 0.939720 | 1.018499 | 1.000059 | 0.926862 | 0.777585 | 0.618083 | 0.510701 | 0.438268 | 0.374440 | 0.324173 | 0.297388 | 0.280040 | 0.272802 | 0.262912 | 0.252740 | 0.243362 | 0.235470 | 0.224757 | 0.222886 | 0.223865 | 0.223627 | 0.222717 | 0.220927 | 0.217124 | 0.215195 | 0.214637 | 0.217845 | 0.224779 | 0.226566 | 0.223986 | 0.218309 | 0.214630 | 0.217895 | 0.225382 | 0.236388 | 0.244475 | 0.251806 | 0.262580 | 0.266299 | 0.256669 | 0.250159 | 0.243288 | 0.234767 | 0.230191 | 0.230644 | 0.236054 | 0.252260 | 0.280896 | 0.315760 | 0.360305 | 0.406561 | 0.424338 | 0.415715 | 0.401930 | 0.375098 | 0.341340 | 0.317848 | 0.301216 | 0.295970 | 0.299925 | 0.322834 | 0.370074 | 0.398672 | 0.440622 |
| right anterior | 1.156718 | 0.965260 | 0.630452 | 0.394184 | 0.286083 | 0.232272 | 0.220263 | 0.216318 | 0.214751 | 0.216786 | 0.251047 | 0.451020 | 0.844065 | 1.775605 | 2.223408 | 1.650589 | 0.993155 | 0.501784 | 0.260103 | 0.214676 | 0.236819 | 0.316632 | 0.453666 | 0.625457 | 0.845163 | 1.152387 | 1.241447 | 0.776676 | 0.370832 | 0.234246 | 0.222778 | 0.376056 | 0.701998 | 0.901046 | 0.769773 | 0.553798 | 0.363376 | 0.286296 | 0.234759 | 0.225644 | 0.232636 | 0.253111 | 0.279958 | 0.339050 | 0.361915 | 0.369706 | 0.328072 | 0.267548 | 0.225880 | 0.214733 | 0.219743 | 0.229479 | 0.243066 | 0.252964 | 0.269507 | 0.272980 | 0.274076 | 0.253415 | 0.237026 | 0.221525 | 0.215647 | 0.215138 | 0.214806 | 0.232659 | 0.343700 | 0.736309 | 1.843066 | 2.786264 | 2.821421 | 2.234991 | 1.799957 | 1.407491 | 1.095053 | 0.806365 | 0.627733 | 0.542324 | 0.477533 | 0.410289 | 0.354390 | 0.290766 | 0.240809 | 0.223790 | 0.222395 | 0.227918 | 0.244599 | 0.277750 | 0.327738 | 0.382883 | 0.483966 | 0.692805 | 1.142318 | 2.044831 | 4.165911 | 8.569343 | 15.849795 | 1.934804e+01 | 15.683359 | 10.570696 | 6.802807 | 3.635155 | 2.215238 | 1.460220 | 0.985342 | 0.746248 | 0.628969 | 0.559609 | 0.542151 | 0.565560 | 0.609626 | 0.688060 | 0.814358 | 1.001098 | 1.121692 | 1.268974 | 1.484801 | 1.603244 | 1.761157 | 1.947680 | 1.924115 | 1.978935 | 2.065892 | 1.910870 | 1.635907 | 1.244835 | 0.905764 | 0.698800 | 0.557306 | 0.456225 | 0.402772 | 0.378240 | 0.341693 | 0.304152 | 0.278929 | 0.271642 | 0.264856 | 0.256587 | 0.246134 | 0.233472 | 0.219499 | 0.214950 | 0.224892 | 0.248372 | 0.271996 | 0.268885 | 0.245521 | 0.222969 | 0.214627 | 0.220425 | 0.231830 | 0.253622 | 0.269199 | 0.280468 | 0.282222 | 0.271817 | 0.247695 | 0.230118 | 0.216788 | 0.215526 | 0.229262 | 0.263080 | 0.313877 | 0.356053 | 0.412634 | 0.481310 | 0.545254 | 0.615499 | 0.714319 | 0.798145 | 0.865889 | 0.895417 | 0.912442 | 0.847289 | 0.689146 | 0.549951 | 0.466227 | 0.410006 | 0.362029 | 0.343638 | 0.344888 | 0.357203 | 0.379482 | 0.464809 | 0.667471 | 0.987789 | 1.334839 | 1.454526 | 1.249312 | 0.928569 | 0.691004 | 0.526925 | 0.478258 | 0.447039 | 0.416655 | 0.395812 | 0.377412 | 0.338179 | 0.299234 | 0.269785 | 0.259024 | 0.269988 | 0.302996 | 0.361649 | 0.454502 | 0.530300 | 0.526150 | 0.447115 | 0.365447 | 0.308218 | 0.270548 | 0.245839 | 0.236681 | 0.235948 | 0.236131 | 0.234604 | 0.230468 | 0.229179 | 0.231501 | 0.229808 | 0.228254 | 0.229386 | 0.226220 | 0.220117 | 0.216599 | 0.214677 | 0.214639 | 0.215814 | 0.219909 | 0.224428 | 0.226099 | 0.216914 | 0.217233 | 0.245750 | 0.311362 | 0.404550 | 0.450126 | 0.433723 | 0.339744 | 0.260152 | 0.222267 | 0.215353 | 0.215114 | 0.219855 | 0.238786 | 0.271328 | 0.304986 | 0.323603 | 0.329494 | 0.304302 | 0.278583 | 0.250826 | 0.233762 | 0.223874 | 0.221767 | 0.220335 | 0.227335 | 0.247522 | 0.285646 | 0.334466 | 0.403214 | 0.455533 | 0.495947 | 0.521395 | 0.539317 | 0.536652 | 0.498586 | 0.431674 | 0.340722 | 0.259615 | 0.218781 | 0.219876 | 0.253291 | 0.293605 | 0.328077 | 0.351771 | 0.360665 | 0.365508 | 0.356432 | 0.346668 | 0.339128 | 0.320895 |
| left central | 0.271348 | 0.280247 | 0.292198 | 0.324293 | 0.352293 | 0.365197 | 0.333407 | 0.290952 | 0.265028 | 0.243713 | 0.226175 | 0.217022 | 0.216796 | 0.251084 | 0.362403 | 0.475133 | 0.550577 | 0.450180 | 0.303977 | 0.255041 | 0.233363 | 0.216082 | 0.220460 | 0.241436 | 0.307062 | 0.382347 | 0.456915 | 0.411955 | 0.312490 | 0.232171 | 0.215024 | 0.240821 | 0.260742 | 0.243949 | 0.217816 | 0.218049 | 0.242690 | 0.279735 | 0.318262 | 0.343437 | 0.326849 | 0.279784 | 0.233910 | 0.215154 | 0.220400 | 0.233072 | 0.238979 | 0.228690 | 0.219684 | 0.215195 | 0.214985 | 0.217119 | 0.217973 | 0.220837 | 0.221576 | 0.225869 | 0.231960 | 0.236402 | 0.231912 | 0.227928 | 0.221003 | 0.217433 | 0.215399 | 0.215325 | 0.219608 | 0.263492 | 0.470029 | 1.528118 | 6.796391 | 25.070310 | 58.648763 | 82.461014 | 70.583092 | 46.281929 | 26.599941 | 12.316082 | 5.915306 | 3.133860 | 1.790761 | 1.175850 | 0.889311 | 0.672046 | 0.582494 | 0.569093 | 0.562943 | 0.566221 | 0.563520 | 0.506963 | 0.495323 | 0.486007 | 0.583299 | 0.835736 | 1.326287 | 1.912600 | 2.868256 | 3.192746e+00 | 2.560622 | 1.505636 | 0.739196 | 0.393997 | 0.273073 | 0.231200 | 0.218526 | 0.217404 | 0.218715 | 0.224109 | 0.238545 | 0.266315 | 0.281391 | 0.278470 | 0.261823 | 0.245811 | 0.235083 | 0.232866 | 0.246845 | 0.286826 | 0.330159 | 0.380837 | 0.464494 | 0.568728 | 0.643312 | 0.679685 | 0.661637 | 0.651761 | 0.659929 | 0.716163 | 0.776852 | 1.010957 | 1.342096 | 1.638361 | 1.869790 | 2.085264 | 1.542784 | 0.926545 | 0.563022 | 0.384663 | 0.319878 | 0.288512 | 0.260120 | 0.239115 | 0.218683 | 0.220662 | 0.279962 | 0.451102 | 0.843176 | 1.282428 | 1.279578 | 0.983348 | 0.672973 | 0.456739 | 0.338232 | 0.281165 | 0.254781 | 0.246319 | 0.245310 | 0.248740 | 0.253422 | 0.248333 | 0.238673 | 0.228174 | 0.220134 | 0.217159 | 0.218161 | 0.224225 | 0.243665 | 0.273992 | 0.300353 | 0.326158 | 0.332346 | 0.324057 | 0.317151 | 0.325366 | 0.334885 | 0.360762 | 0.400596 | 0.477972 | 0.552473 | 0.619893 | 0.530312 | 0.434637 | 0.360252 | 0.306881 | 0.261067 | 0.243329 | 0.233246 | 0.236459 | 0.243969 | 0.262728 | 0.286817 | 0.326064 | 0.347540 | 0.379044 | 0.438477 | 0.510460 | 0.522070 | 0.473695 | 0.388659 | 0.290315 | 0.247750 | 0.228803 | 0.221380 | 0.219112 | 0.221276 | 0.229222 | 0.248981 | 0.279733 | 0.306378 | 0.311022 | 0.290824 | 0.264683 | 0.242798 | 0.232548 | 0.228668 | 0.225809 | 0.225102 | 0.226055 | 0.224298 | 0.223243 | 0.222922 | 0.219537 | 0.215364 | 0.215098 | 0.220602 | 0.230506 | 0.246313 | 0.271124 | 0.301699 | 0.347081 | 0.443644 | 0.646697 | 1.048376 | 1.514904 | 1.727875 | 1.610297 | 1.151304 | 0.688894 | 0.410721 | 0.288504 | 0.245311 | 0.229520 | 0.224051 | 0.223872 | 0.224410 | 0.224948 | 0.219173 | 0.214762 | 0.217010 | 0.228352 | 0.257943 | 0.308769 | 0.389723 | 0.477426 | 0.584332 | 0.715835 | 0.757473 | 0.668126 | 0.538111 | 0.411793 | 0.343992 | 0.306202 | 0.294705 | 0.306060 | 0.339104 | 0.388074 | 0.466589 | 0.576122 | 0.704233 | 0.840541 | 0.914541 | 0.872620 | 0.772285 | 0.705501 | 0.655246 | 0.696074 | 0.728894 | 0.772775 | 0.832608 | 0.940233 | 0.897093 | 0.924069 |
| right central | 3.601892 | 3.916981 | 4.369142 | 4.776175 | 4.158477 | 2.776355 | 1.972099 | 1.463178 | 1.229311 | 1.055438 | 0.791814 | 0.492978 | 0.307815 | 0.235854 | 0.217085 | 0.215658 | 0.214636 | 0.216544 | 0.227718 | 0.252849 | 0.296178 | 0.341779 | 0.375279 | 0.357639 | 0.300269 | 0.245432 | 0.215619 | 0.221742 | 0.236507 | 0.231370 | 0.216561 | 0.228696 | 0.300310 | 0.437974 | 0.571305 | 0.512558 | 0.335161 | 0.233372 | 0.218673 | 0.245489 | 0.254899 | 0.237253 | 0.218138 | 0.225001 | 0.292484 | 0.487602 | 0.749530 | 0.922573 | 0.720608 | 0.515463 | 0.347726 | 0.260218 | 0.220548 | 0.217544 | 0.269371 | 0.438874 | 0.888034 | 1.596622 | 1.703705 | 0.903365 | 0.453240 | 0.288148 | 0.234629 | 0.215563 | 0.218001 | 0.246549 | 0.326483 | 0.625995 | 2.037065 | 11.158631 | 53.496239 | 185.242318 | 279.749210 | 152.002844 | 52.546377 | 14.452007 | 4.347075 | 1.655453 | 0.758893 | 0.442579 | 0.391109 | 0.470511 | 0.991222 | 3.346287 | 13.072434 | 38.767898 | 59.383324 | 47.023594 | 39.610844 | 42.382816 | 73.333944 | 217.884334 | 1247.896050 | 13975.773687 | 254799.289956 | 1.016336e+06 | 669630.583647 | 97961.297455 | 12316.261428 | 2679.089782 | 1226.876607 | 1109.278005 | 996.854748 | 759.212485 | 408.933546 | 185.786217 | 71.297232 | 39.019577 | 26.583280 | 23.848102 | 18.899253 | 17.893755 | 20.420745 | 30.121285 | 34.669833 | 37.817979 | 33.791715 | 32.846146 | 31.340432 | 36.121489 | 49.939016 | 103.731007 | 195.233579 | 387.477722 | 780.581648 | 1085.189334 | 804.665914 | 479.078037 | 282.065050 | 219.524320 | 225.754202 | 300.643468 | 372.986841 | 348.387528 | 191.751848 | 88.338177 | 33.657887 | 11.363779 | 3.631423 | 1.602795 | 0.921324 | 0.681902 | 0.593731 | 0.568232 | 0.533709 | 0.490490 | 0.465887 | 0.485121 | 0.550274 | 0.619855 | 0.659510 | 0.668972 | 0.613224 | 0.580871 | 0.565137 | 0.641601 | 0.923639 | 1.613169 | 3.467332 | 8.927859 | 19.350844 | 23.788000 | 14.947377 | 5.494280 | 2.350447 | 1.242973 | 0.959433 | 0.925319 | 1.085855 | 1.153323 | 1.198470 | 1.042346 | 0.938917 | 0.808664 | 0.711767 | 0.597774 | 0.581825 | 0.587922 | 0.586604 | 0.564796 | 0.531470 | 0.481804 | 0.453955 | 0.435424 | 0.433363 | 0.457132 | 0.475788 | 0.495419 | 0.532397 | 0.559540 | 0.589675 | 0.642980 | 0.699138 | 0.726327 | 0.723419 | 0.638028 | 0.507723 | 0.417404 | 0.380737 | 0.372508 | 0.394852 | 0.437839 | 0.493301 | 0.542235 | 0.578151 | 0.504626 | 0.400639 | 0.304463 | 0.244644 | 0.220913 | 0.217120 | 0.220328 | 0.243374 | 0.298274 | 0.358109 | 0.450875 | 0.500873 | 0.451203 | 0.362892 | 0.302706 | 0.257433 | 0.243549 | 0.242675 | 0.264290 | 0.316557 | 0.406515 | 0.482483 | 0.547487 | 0.574544 | 0.539009 | 0.458850 | 0.370950 | 0.303906 | 0.261044 | 0.244619 | 0.237957 | 0.234471 | 0.228393 | 0.220277 | 0.214749 | 0.227755 | 0.271140 | 0.362847 | 0.502732 | 0.636315 | 0.709640 | 0.635189 | 0.482477 | 0.361365 | 0.274778 | 0.242721 | 0.230929 | 0.221446 | 0.217153 | 0.215158 | 0.215182 | 0.218785 | 0.234558 | 0.270081 | 0.305568 | 0.346510 | 0.391897 | 0.365513 | 0.333584 | 0.304181 | 0.277913 | 0.256615 | 0.237071 | 0.222873 | 0.216049 | 0.214932 | 0.222641 | 0.245261 | 0.288827 | 0.345357 | 0.429298 | 0.486902 | 0.480980 | 0.460205 | 0.434331 |
| left posterior | 1.100674 | 1.530368 | 2.143446 | 4.409477 | 7.305935 | 7.113946 | 3.235460 | 1.209834 | 0.428768 | 0.235806 | 0.217563 | 0.261334 | 0.320442 | 0.311250 | 0.254465 | 0.215093 | 0.404373 | 3.684988 | 74.820785 | 260.401123 | 63.752963 | 6.838070 | 0.708719 | 0.220196 | 0.359995 | 1.119767 | 1.854083 | 0.945009 | 0.451105 | 0.239228 | 0.220113 | 0.253895 | 0.232382 | 0.215207 | 0.230015 | 0.274010 | 0.333270 | 0.377717 | 0.402035 | 0.385415 | 0.339460 | 0.298696 | 0.294358 | 0.328228 | 0.387143 | 0.439689 | 0.440901 | 0.422325 | 0.381927 | 0.328432 | 0.278628 | 0.239396 | 0.216750 | 0.220156 | 0.253546 | 0.313393 | 0.388691 | 0.491367 | 0.647236 | 0.687213 | 0.474628 | 0.288819 | 0.215079 | 0.298152 | 0.990441 | 6.365598 | 51.316550 | 316.648450 | 1466.970491 | 5758.382734 | 17046.290332 | 40094.933045 | 69611.809626 | 90264.581663 | 78301.663426 | 53292.386592 | 22881.637950 | 10404.690185 | 5227.966248 | 1873.270609 | 357.041250 | 46.636047 | 8.914038 | 3.117127 | 1.685274 | 1.203539 | 1.033188 | 0.878341 | 0.808229 | 0.925488 | 1.216319 | 1.751669 | 2.871522 | 4.604742 | 7.232287 | 1.103212e+01 | 13.117432 | 10.984527 | 7.672394 | 5.046748 | 3.642319 | 2.906498 | 2.741243 | 3.059198 | 3.877150 | 5.544853 | 8.630340 | 13.474093 | 18.605751 | 20.173771 | 16.001546 | 10.924005 | 6.722822 | 4.591626 | 3.491448 | 3.155868 | 3.529180 | 4.913593 | 8.013516 | 14.636858 | 27.786760 | 47.743980 | 65.669228 | 70.682243 | 59.331275 | 32.091134 | 14.031230 | 5.134373 | 1.967839 | 0.920640 | 0.597433 | 0.468644 | 0.444951 | 0.456611 | 0.496650 | 0.560597 | 0.666753 | 0.789680 | 0.882402 | 0.953853 | 0.978463 | 0.929629 | 0.930140 | 0.968268 | 1.000284 | 0.932251 | 0.757685 | 0.560772 | 0.430955 | 0.340374 | 0.288798 | 0.258428 | 0.240163 | 0.231595 | 0.228024 | 0.229342 | 0.237326 | 0.253788 | 0.273460 | 0.299078 | 0.320866 | 0.343149 | 0.353188 | 0.348107 | 0.331741 | 0.313090 | 0.292628 | 0.276826 | 0.264524 | 0.251404 | 0.240796 | 0.235344 | 0.233245 | 0.236408 | 0.248712 | 0.268929 | 0.297475 | 0.342091 | 0.394581 | 0.450343 | 0.510881 | 0.562044 | 0.583245 | 0.612819 | 0.623873 | 0.598306 | 0.545765 | 0.487856 | 0.419923 | 0.366729 | 0.338508 | 0.318524 | 0.309632 | 0.313509 | 0.340080 | 0.388716 | 0.495273 | 0.666589 | 0.938267 | 1.234975 | 1.525830 | 1.606369 | 1.577845 | 1.410288 | 1.223003 | 1.058039 | 0.933085 | 0.854599 | 0.834563 | 0.912701 | 1.087233 | 1.465565 | 2.120529 | 3.419005 | 5.183360 | 7.785038 | 10.619101 | 12.424138 | 10.750753 | 8.263647 | 5.051125 | 2.969550 | 1.756442 | 1.135020 | 0.828598 | 0.719413 | 0.638302 | 0.602184 | 0.586900 | 0.590661 | 0.601291 | 0.693590 | 0.908168 | 1.432250 | 2.668405 | 5.690854 | 10.480839 | 17.688504 | 21.964885 | 19.850633 | 14.417449 | 8.847595 | 4.304654 | 2.447754 | 1.698941 | 1.509591 | 1.779490 | 2.416375 | 2.956637 | 3.155682 | 2.796325 | 2.181370 | 1.655352 | 1.260406 | 0.967541 | 0.812156 | 0.662608 | 0.566934 | 0.502844 | 0.438648 | 0.366570 | 0.311573 | 0.270385 | 0.246442 | 0.232839 | 0.225681 | 0.222473 | 0.221470 | 0.220782 | 0.219563 | 0.216565 | 0.214853 | 0.215863 | 0.220946 | 0.227190 | 0.228444 | 0.227886 | 0.224157 | 0.223970 | 0.221153 |
| right posterior | 1.415190 | 1.640346 | 1.904937 | 2.106519 | 2.049221 | 1.735434 | 1.288669 | 0.839896 | 0.479932 | 0.308181 | 0.248745 | 0.226003 | 0.218603 | 0.220557 | 0.245492 | 0.282108 | 0.341006 | 0.366996 | 0.334626 | 0.265648 | 0.225282 | 0.218649 | 0.237523 | 0.236603 | 0.221049 | 0.218674 | 0.285032 | 0.464498 | 0.887146 | 1.687566 | 2.715473 | 2.097668 | 0.994294 | 0.410151 | 0.225350 | 0.235739 | 0.347100 | 0.436856 | 0.445243 | 0.362907 | 0.308394 | 0.264681 | 0.247761 | 0.239473 | 0.236933 | 0.230250 | 0.217437 | 0.225783 | 0.355630 | 0.734556 | 1.282774 | 1.430012 | 1.195020 | 0.851705 | 0.552505 | 0.421028 | 0.406400 | 0.446726 | 0.528275 | 0.642373 | 0.565826 | 0.332999 | 0.214890 | 0.359718 | 1.520048 | 9.642371 | 57.540275 | 235.749792 | 616.256329 | 1294.926307 | 1685.287276 | 1410.700164 | 753.410617 | 250.452138 | 69.732667 | 21.694189 | 6.891118 | 2.381856 | 0.984489 | 0.453990 | 0.272143 | 0.224916 | 0.217234 | 0.218209 | 0.225310 | 0.246475 | 0.276132 | 0.322986 | 0.401316 | 0.541166 | 0.760532 | 1.207787 | 1.874947 | 2.753367 | 3.834148 | 4.627738e+00 | 4.745156 | 4.354279 | 3.780813 | 3.133145 | 2.928998 | 2.807559 | 2.825931 | 3.046021 | 3.267226 | 3.168246 | 3.096926 | 3.028406 | 2.727022 | 2.361214 | 1.991094 | 1.650981 | 1.509076 | 1.651353 | 1.946579 | 2.738113 | 4.822729 | 9.669421 | 22.417454 | 55.150047 | 118.017459 | 192.965339 | 204.472494 | 140.894052 | 74.281327 | 32.897973 | 13.460745 | 6.115106 | 3.151847 | 1.912293 | 1.339491 | 1.106663 | 1.009149 | 0.972045 | 1.005110 | 1.141871 | 1.298959 | 1.502943 | 1.711880 | 1.985735 | 2.272452 | 2.460455 | 2.672900 | 3.182013 | 3.836258 | 4.169420 | 3.809630 | 3.179506 | 2.469087 | 1.842710 | 1.370520 | 1.118304 | 0.967460 | 0.953371 | 1.089085 | 1.524435 | 2.391043 | 4.150966 | 6.909245 | 10.193202 | 11.700795 | 11.944737 | 10.283753 | 7.973214 | 5.701954 | 4.080175 | 2.840012 | 2.209665 | 1.763013 | 1.550627 | 1.527868 | 1.692805 | 1.915736 | 2.473752 | 3.136636 | 3.969481 | 4.306614 | 3.976506 | 3.120147 | 2.219517 | 1.440033 | 1.026356 | 0.810540 | 0.737229 | 0.805801 | 1.047821 | 1.595101 | 2.609566 | 3.917745 | 4.457389 | 4.018419 | 2.998108 | 2.371581 | 2.160797 | 2.168019 | 2.419148 | 2.712400 | 2.867723 | 3.093293 | 3.575435 | 4.409143 | 5.816050 | 7.719589 | 10.195368 | 12.511514 | 13.696494 | 13.728315 | 12.866068 | 11.572682 | 10.019438 | 8.902533 | 8.498763 | 9.285374 | 11.889991 | 16.987479 | 27.690196 | 44.185665 | 55.087060 | 56.225114 | 49.227494 | 32.170975 | 19.027880 | 11.586302 | 7.362413 | 5.457829 | 4.837622 | 4.603479 | 4.712536 | 5.226061 | 5.646771 | 6.295426 | 7.497491 | 9.972609 | 13.470030 | 16.983133 | 20.277706 | 22.095830 | 22.250258 | 20.888578 | 19.310728 | 15.161024 | 11.425120 | 8.529495 | 6.844080 | 6.130482 | 6.471489 | 7.719081 | 11.317245 | 16.134615 | 20.013104 | 21.128867 | 18.236026 | 12.339645 | 7.444800 | 4.532473 | 3.118800 | 2.467673 | 2.166217 | 2.126079 | 2.331312 | 2.571201 | 2.629478 | 2.469629 | 1.985258 | 1.481428 | 1.015634 | 0.702453 | 0.526504 | 0.448514 | 0.389110 | 0.366161 | 0.364910 | 0.374403 | 0.406187 | 0.491231 | 0.601766 | 0.781109 | 1.031767 | 1.158111 | 1.271417 |
| all electrodes | 0.415990 | 0.512827 | 0.520368 | 0.460661 | 0.393658 | 0.347534 | 0.258708 | 0.223523 | 0.218222 | 0.297603 | 0.458945 | 0.493165 | 0.497999 | 0.387346 | 0.301132 | 0.249170 | 0.223515 | 0.214705 | 0.224382 | 0.265039 | 0.293088 | 0.256986 | 0.217270 | 0.223253 | 0.304199 | 0.461970 | 0.830108 | 0.861801 | 0.696830 | 0.479178 | 0.381288 | 0.270832 | 0.245212 | 0.253622 | 0.301059 | 0.362554 | 0.446495 | 0.420408 | 0.340991 | 0.249230 | 0.214829 | 0.267161 | 0.466547 | 0.976954 | 1.303995 | 1.025813 | 0.545279 | 0.314820 | 0.262194 | 0.247849 | 0.248165 | 0.263362 | 0.307072 | 0.359885 | 0.459301 | 0.545102 | 0.562245 | 0.515294 | 0.478199 | 0.438939 | 0.361914 | 0.269466 | 0.216969 | 0.242277 | 0.460791 | 1.451414 | 8.031553 | 65.761658 | 579.172183 | 3229.354713 | 12407.010331 | 32500.756036 | 59634.100001 | 55433.837992 | 30697.508423 | 8102.625147 | 1241.226134 | 193.687431 | 43.082776 | 16.836436 | 10.045159 | 8.036709 | 6.612456 | 4.347463 | 2.272421 | 1.351107 | 0.884996 | 0.614720 | 0.491103 | 0.431345 | 0.454982 | 0.540871 | 0.801924 | 1.537155 | 3.993069 | 8.430441e+00 | 13.486299 | 10.062347 | 5.366880 | 2.465099 | 1.257068 | 0.690556 | 0.482524 | 0.410604 | 0.396042 | 0.436265 | 0.510694 | 0.679401 | 1.058675 | 1.999351 | 4.337629 | 10.859265 | 29.541386 | 65.256824 | 138.290523 | 193.480897 | 167.345648 | 139.262503 | 144.464677 | 143.325495 | 195.554379 | 300.992245 | 454.218851 | 472.013535 | 355.616419 | 198.979754 | 130.325931 | 78.657948 | 48.919665 | 28.790757 | 18.708330 | 14.987368 | 13.480135 | 11.814477 | 12.186431 | 13.986715 | 15.189575 | 14.762410 | 15.052087 | 13.785779 | 11.236443 | 7.678212 | 4.752342 | 3.495638 | 3.074820 | 2.885061 | 2.736627 | 2.928231 | 2.949145 | 2.858158 | 2.604892 | 2.253567 | 1.829496 | 1.348571 | 0.914816 | 0.622706 | 0.464864 | 0.376999 | 0.332729 | 0.290141 | 0.256471 | 0.231648 | 0.217861 | 0.214844 | 0.221256 | 0.240126 | 0.269933 | 0.317287 | 0.376041 | 0.427754 | 0.445630 | 0.464664 | 0.444783 | 0.427259 | 0.396518 | 0.338547 | 0.274321 | 0.233290 | 0.215803 | 0.219122 | 0.237768 | 0.260444 | 0.258354 | 0.243664 | 0.227187 | 0.218467 | 0.214627 | 0.218634 | 0.229086 | 0.234273 | 0.239206 | 0.236056 | 0.225991 | 0.218477 | 0.215196 | 0.215213 | 0.217703 | 0.221526 | 0.229829 | 0.239120 | 0.246497 | 0.257894 | 0.269292 | 0.275276 | 0.270837 | 0.258907 | 0.254899 | 0.267270 | 0.291772 | 0.331937 | 0.386304 | 0.447598 | 0.455902 | 0.406952 | 0.337058 | 0.291764 | 0.266331 | 0.266983 | 0.286443 | 0.315108 | 0.327474 | 0.316672 | 0.294896 | 0.281354 | 0.279210 | 0.290734 | 0.336315 | 0.406678 | 0.487666 | 0.583287 | 0.655884 | 0.599656 | 0.562761 | 0.545462 | 0.578699 | 0.675655 | 0.816020 | 1.012598 | 1.324601 | 1.542210 | 1.444773 | 1.173269 | 0.882007 | 0.657178 | 0.510254 | 0.465784 | 0.469430 | 0.472505 | 0.490248 | 0.534330 | 0.494840 | 0.431232 | 0.350952 | 0.276346 | 0.229285 | 0.214765 | 0.241199 | 0.288891 | 0.322750 | 0.325046 | 0.283402 | 0.235353 | 0.216869 | 0.215033 | 0.217187 | 0.216775 | 0.214717 | 0.227067 | 0.274095 | 0.358722 | 0.451092 | 0.551220 | 0.619273 | 0.686492 | 0.691977 | 0.720451 | 0.704490 | 0.679764 | 0.592006 | 0.582382 |

M) maximum cross-correlation, neutral vs happy

  
|  | time window | peak latency | cluster *p* | peak Cohen's *d* |  | | | |
| **all electrodes** |  | | | |  | | | |
|  | | | | | | | | |

Model correlations, cluster permutation tests

|  | **left hemisphere** | | | | **right hemisphere** | | | |
|  | time window | peak latency | cluster *p* | peak Cohen's *d* | time window | peak latency | cluster *p* | peak Cohen's *d* |
| **anterior** |  | | | | 230 - 400 ms | 280 ms | 0.009 | 0.9598 |
| **central** |  | | | |  | | | |
| **posterior** | 245 - 415 ms | 280 ms | 0.0142 | 0.8487 | 260 - 440 ms | 290 ms | 0.0158 | 0.7016 |
  | | | | 800 - 1120 ms | 845 ms | 0.0063 | 0.6343 |

  

Model correlations, Bayesian statistics

|  | -200 | -195 | -190 | -185 | -180 | -175 | -170 | -165 | -160 | -155 | -150 | -145 | -140 | -135 | -130 | -125 | -120 | -115 | -110 | -105 | -100 | -95 | -90 | -85 | -80 | -75 | -70 | -65 | -60 | -55 | -50 | -45 | -40 | -35 | -30 | -25 | -20 | -15 | -10 | -5 | 0 | 5 | 10 | 15 | 20 | 25 | 30 | 35 | 40 | 45 | 50 | 55 | 60 | 65 | 70 | 75 | 80 | 85 | 90 | 95 | 100 | 105 | 110 | 115 | 120 | 125 | 130 | 135 | 140 | 145 | 150 | 155 | 160 | 165 | 170 | 175 | 180 | 185 | 190 | 195 | 200 | 205 | 210 | 215 | 220 | 225 | 230 | 235 | 240 | 245 | 250 | 255 | 260 | 265 | 270 | 275 | 280 | 285 | 290 | 295 | 300 | 305 | 310 | 315 | 320 | 325 | 330 | 335 | 340 | 345 | 350 | 355 | 360 | 365 | 370 | 375 | 380 | 385 | 390 | 395 | 400 | 405 | 410 | 415 | 420 | 425 | 430 | 435 | 440 | 445 | 450 | 455 | 460 | 465 | 470 | 475 | 480 | 485 | 490 | 495 | 500 | 505 | 510 | 515 | 520 | 525 | 530 | 535 | 540 | 545 | 550 | 555 | 560 | 565 | 570 | 575 | 580 | 585 | 590 | 595 | 600 | 605 | 610 | 615 | 620 | 625 | 630 | 635 | 640 | 645 | 650 | 655 | 660 | 665 | 670 | 675 | 680 | 685 | 690 | 695 | 700 | 705 | 710 | 715 | 720 | 725 | 730 | 735 | 740 | 745 | 750 | 755 | 760 | 765 | 770 | 775 | 780 | 785 | 790 | 795 | 800 | 805 | 810 | 815 | 820 | 825 | 830 | 835 | 840 | 845 | 850 | 855 | 860 | 865 | 870 | 875 | 880 | 885 | 890 | 895 | 900 | 905 | 910 | 915 | 920 | 925 | 930 | 935 | 940 | 945 | 950 | 955 | 960 | 965 | 970 | 975 | 980 | 985 | 990 | 995 | 1000 | 1005 | 1010 | 1015 | 1020 | 1025 | 1030 | 1035 | 1040 | 1045 | 1050 | 1055 | 1060 | 1065 | 1070 | 1075 | 1080 | 1085 | 1090 | 1095 | 1100 | 1105 | 1110 | 1115 | 1120 | 1125 | 1130 | 1135 | 1140 | 1145 | 1150 | 1155 | 1160 | 1165 | 1170 | 1175 | 1180 | 1185 | 1190 | 1195 |
| --- | --- | --- | --- | --- | --- | --- | --- | --- | --- | --- | --- | --- | --- | --- | --- | --- | --- | --- | --- | --- | --- | --- | --- | --- | --- | --- | --- | --- | --- | --- | --- | --- | --- | --- | --- | --- | --- | --- | --- | --- | --- | --- | --- | --- | --- | --- | --- | --- | --- | --- | --- | --- | --- | --- | --- | --- | --- | --- | --- | --- | --- | --- | --- | --- | --- | --- | --- | --- | --- | --- | --- | --- | --- | --- | --- | --- | --- | --- | --- | --- | --- | --- | --- | --- | --- | --- | --- | --- | --- | --- | --- | --- | --- | --- | --- | --- | --- | --- | --- | --- | --- | --- | --- | --- | --- | --- | --- | --- | --- | --- | --- | --- | --- | --- | --- | --- | --- | --- | --- | --- | --- | --- | --- | --- | --- | --- | --- | --- | --- | --- | --- | --- | --- | --- | --- | --- | --- | --- | --- | --- | --- | --- | --- | --- | --- | --- | --- | --- | --- | --- | --- | --- | --- | --- | --- | --- | --- | --- | --- | --- | --- | --- | --- | --- | --- | --- | --- | --- | --- | --- | --- | --- | --- | --- | --- | --- | --- | --- | --- | --- | --- | --- | --- | --- | --- | --- | --- | --- | --- | --- | --- | --- | --- | --- | --- | --- | --- | --- | --- | --- | --- | --- | --- | --- | --- | --- | --- | --- | --- | --- | --- | --- | --- | --- | --- | --- | --- | --- | --- | --- | --- | --- | --- | --- | --- | --- | --- | --- | --- | --- | --- | --- | --- | --- | --- | --- | --- | --- | --- | --- | --- | --- | --- | --- | --- | --- | --- | --- | --- | --- | --- | --- | --- | --- | --- | --- | --- | --- | --- | --- | --- | --- | --- | --- | --- | --- | --- | --- | --- | --- | --- | --- | --- | --- | --- | --- | --- | --- | --- | --- |
| left anterior | 0.263596 | 0.273655 | 0.261449 | 0.248221 | 0.228283 | 0.214670 | 0.229355 | 0.259450 | 0.268684 | 0.250830 | 0.245304 | 0.231094 | 0.226390 | 0.232453 | 0.269315 | 0.331629 | 0.417776 | 0.412976 | 0.352267 | 0.287873 | 0.234280 | 0.215442 | 0.248890 | 0.273460 | 0.246768 | 0.218435 | 0.237180 | 0.313878 | 0.371431 | 0.379861 | 0.308797 | 0.216067 | 0.293587 | 0.800712 | 2.938035 | 4.448278 | 3.846319 | 2.256959 | 1.533688 | 1.141622 | 1.155654 | 0.988280 | 0.943577 | 0.806759 | 0.810402 | 0.735045 | 0.563636 | 0.345655 | 0.262410 | 0.227567 | 0.216589 | 0.215712 | 0.227388 | 0.266726 | 0.372713 | 0.577777 | 0.828986 | 0.865972 | 0.738744 | 0.489640 | 0.298219 | 0.222132 | 0.221920 | 0.312495 | 0.502441 | 0.672499 | 0.741885 | 0.709309 | 0.689208 | 0.646476 | 0.569396 | 0.465077 | 0.360386 | 0.259154 | 0.216328 | 0.225071 | 0.273129 | 0.352614 | 0.417191 | 0.463452 | 0.528992 | 0.517949 | 0.424712 | 0.331386 | 0.258642 | 0.225809 | 0.214778 | 0.222282 | 0.247634 | 0.277960 | 0.318017 | 0.346325 | 0.335429 | 0.310783 | 0.296444 | 0.272908 | 0.255985 | 0.242114 | 0.228383 | 0.218580 | 0.214785 | 0.219334 | 0.240078 | 0.279486 | 0.318532 | 0.336878 | 0.292425 | 0.242546 | 0.219962 | 0.215049 | 0.214630 | 0.214708 | 0.215532 | 0.217178 | 0.222690 | 0.229360 | 0.230102 | 0.229134 | 0.226306 | 0.219572 | 0.216793 | 0.216422 | 0.218522 | 0.225093 | 0.235850 | 0.241149 | 0.241120 | 0.235185 | 0.220341 | 0.215101 | 0.231120 | 0.271389 | 0.338258 | 0.421461 | 0.500021 | 0.542973 | 0.490887 | 0.428555 | 0.362630 | 0.320956 | 0.298119 | 0.299616 | 0.312821 | 0.353011 | 0.413343 | 0.454352 | 0.453538 | 0.378925 | 0.298131 | 0.241930 | 0.217592 | 0.219768 | 0.244357 | 0.274489 | 0.287083 | 0.281325 | 0.255658 | 0.229889 | 0.216360 | 0.215468 | 0.220830 | 0.223630 | 0.218432 | 0.214641 | 0.222630 | 0.249468 | 0.289546 | 0.334380 | 0.341175 | 0.319299 | 0.278870 | 0.253940 | 0.231816 | 0.224531 | 0.219676 | 0.216748 | 0.214627 | 0.215985 | 0.222729 | 0.231133 | 0.240607 | 0.237309 | 0.235760 | 0.240717 | 0.262193 | 0.294130 | 0.378805 | 0.526458 | 0.706535 | 0.811074 | 0.801187 | 0.657267 | 0.548585 | 0.452592 | 0.384471 | 0.382269 | 0.484826 | 0.707514 | 1.114766 | 1.600269 | 2.135358 | 2.412751 | 2.099540 | 1.399956 | 1.006208 | 0.766638 | 0.636964 | 0.592668 | 0.593285 | 0.636569 | 0.717630 | 0.783205 | 0.870720 | 1.098836 | 1.353558 | 1.394410 | 1.263489 | 0.971231 | 0.720918 | 0.543815 | 0.422086 | 0.343874 | 0.313041 | 0.297945 | 0.297057 | 0.306047 | 0.342686 | 0.385610 | 0.416199 | 0.411864 | 0.390746 | 0.362793 | 0.342170 | 0.324879 | 0.338387 | 0.385797 | 0.423550 | 0.451122 | 0.448188 | 0.376116 | 0.298954 | 0.244649 | 0.219642 | 0.214813 | 0.214636 | 0.214781 | 0.219657 | 0.231055 | 0.246367 | 0.258036 | 0.278897 | 0.302229 | 0.362026 | 0.482593 | 0.787251 | 1.070689 | 1.178549 | 0.948583 | 0.666931 | 0.428182 | 0.322288 | 0.276297 | 0.275646 | 0.304028 | 0.369942 | 0.458453 | 0.535126 | 0.545110 | 0.498294 | 0.453606 | 0.419603 | 0.381924 | 0.366481 | 0.371448 | 0.365693 | 0.360875 | 0.366619 | 0.367155 | 0.352618 | 0.354109 |
| right anterior | 3.070529 | 3.714918 | 4.037464 | 5.275251 | 3.543295 | 1.514171 | 0.952376 | 0.742285 | 0.589606 | 0.397281 | 0.258803 | 0.214744 | 0.270186 | 0.572054 | 0.944940 | 1.110310 | 1.018521 | 0.691219 | 0.382995 | 0.271917 | 0.230326 | 0.223690 | 0.224143 | 0.221804 | 0.219141 | 0.214915 | 0.224005 | 0.257862 | 0.285406 | 0.301290 | 0.288629 | 0.240386 | 0.217449 | 0.217247 | 0.232228 | 0.254849 | 0.244752 | 0.227236 | 0.214656 | 0.216963 | 0.219471 | 0.217036 | 0.214803 | 0.217466 | 0.224815 | 0.233972 | 0.233757 | 0.238185 | 0.236481 | 0.234556 | 0.227965 | 0.224097 | 0.217987 | 0.214796 | 0.216080 | 0.220252 | 0.222217 | 0.220742 | 0.218719 | 0.216887 | 0.216407 | 0.221589 | 0.266924 | 0.519842 | 1.381156 | 3.779202 | 11.251689 | 16.978261 | 11.488349 | 6.085634 | 3.127864 | 2.017448 | 1.504749 | 1.146392 | 0.989318 | 1.053031 | 0.981333 | 0.915542 | 0.886926 | 0.762066 | 0.575701 | 0.535027 | 0.541865 | 0.551227 | 0.651047 | 0.859847 | 1.235548 | 1.713008 | 2.870276 | 6.467260 | 18.705966 | 50.266464 | 128.627023 | 277.131254 | 476.779430 | 453.036427 | 282.539172 | 161.646649 | 94.698277 | 45.984318 | 23.678283 | 13.608980 | 8.116215 | 5.816510 | 4.974882 | 5.244723 | 6.274049 | 8.435506 | 10.105279 | 11.145325 | 11.142276 | 11.434922 | 10.712655 | 9.304585 | 7.885527 | 5.837214 | 4.614390 | 3.703853 | 2.747264 | 1.855419 | 1.531688 | 1.215833 | 1.035969 | 0.869118 | 0.781576 | 0.714445 | 0.699134 | 0.675093 | 0.661703 | 0.676391 | 0.626937 | 0.615527 | 0.656990 | 0.801175 | 1.042160 | 1.608044 | 2.393683 | 3.076221 | 2.726255 | 2.181408 | 1.668197 | 1.179129 | 0.821620 | 0.573983 | 0.394549 | 0.314388 | 0.279880 | 0.267124 | 0.265875 | 0.267694 | 0.255178 | 0.245056 | 0.241511 | 0.238646 | 0.236421 | 0.242853 | 0.255274 | 0.273189 | 0.294823 | 0.309759 | 0.321099 | 0.327349 | 0.313179 | 0.290853 | 0.270327 | 0.244447 | 0.224508 | 0.215429 | 0.215884 | 0.220568 | 0.224760 | 0.229877 | 0.230632 | 0.233458 | 0.230977 | 0.222893 | 0.217813 | 0.214979 | 0.215273 | 0.216725 | 0.218918 | 0.226168 | 0.234751 | 0.247457 | 0.266977 | 0.289820 | 0.309526 | 0.333400 | 0.338621 | 0.347115 | 0.360836 | 0.372839 | 0.375816 | 0.393810 | 0.439890 | 0.520560 | 0.618001 | 0.760556 | 0.868153 | 0.865113 | 0.781366 | 0.686551 | 0.585444 | 0.544842 | 0.500691 | 0.489240 | 0.487377 | 0.472477 | 0.423189 | 0.381825 | 0.318984 | 0.262637 | 0.231020 | 0.221774 | 0.222980 | 0.235914 | 0.263791 | 0.310420 | 0.381723 | 0.463533 | 0.547042 | 0.585872 | 0.552281 | 0.522085 | 0.440553 | 0.369218 | 0.365513 | 0.428192 | 0.516933 | 0.859350 | 1.784913 | 3.098107 | 4.409032 | 4.435547 | 3.355695 | 2.151745 | 1.204089 | 0.654356 | 0.422779 | 0.326289 | 0.296149 | 0.296661 | 0.320850 | 0.353598 | 0.390911 | 0.426377 | 0.483412 | 0.477439 | 0.457680 | 0.398427 | 0.351686 | 0.301795 | 0.278703 | 0.267678 | 0.281447 | 0.315556 | 0.389961 | 0.533013 | 0.775951 | 1.029710 | 1.165627 | 1.164877 | 1.062676 | 0.907572 | 0.755024 | 0.614570 | 0.451723 | 0.320888 | 0.238732 | 0.214631 | 0.237812 | 0.286154 | 0.343905 | 0.387743 | 0.394739 | 0.382275 | 0.345339 | 0.285628 | 0.266481 | 0.249114 |
| left central | 0.347491 | 0.385451 | 0.404937 | 0.451914 | 0.484732 | 0.457470 | 0.339768 | 0.274026 | 0.237073 | 0.216960 | 0.215862 | 0.217784 | 0.219242 | 0.225236 | 0.255433 | 0.314798 | 0.498964 | 0.877920 | 0.973209 | 0.728216 | 0.594506 | 0.446135 | 0.372165 | 0.306462 | 0.236866 | 0.215092 | 0.267686 | 0.447395 | 0.855599 | 1.362495 | 1.682994 | 1.616613 | 1.103885 | 0.522715 | 0.295865 | 0.235698 | 0.221585 | 0.225374 | 0.232062 | 0.244003 | 0.266347 | 0.272303 | 0.252501 | 0.251232 | 0.246602 | 0.241516 | 0.239129 | 0.239557 | 0.239551 | 0.236898 | 0.224510 | 0.215346 | 0.219393 | 0.231185 | 0.248370 | 0.262834 | 0.280465 | 0.299611 | 0.393723 | 0.501926 | 0.655392 | 0.618052 | 0.500824 | 0.341200 | 0.238904 | 0.216119 | 0.249154 | 0.332857 | 0.439253 | 0.431148 | 0.351463 | 0.269997 | 0.224310 | 0.214814 | 0.216352 | 0.226164 | 0.233065 | 0.232407 | 0.235072 | 0.236846 | 0.238688 | 0.244247 | 0.256377 | 0.278144 | 0.311903 | 0.351350 | 0.371095 | 0.361982 | 0.312285 | 0.256794 | 0.215716 | 0.250059 | 0.437484 | 1.023301 | 2.663830 | 5.869872 | 9.009492 | 10.671495 | 9.459487 | 5.747305 | 2.829808 | 1.118736 | 0.564031 | 0.389122 | 0.302036 | 0.252395 | 0.233333 | 0.222549 | 0.217563 | 0.215914 | 0.215276 | 0.214859 | 0.214706 | 0.214713 | 0.214633 | 0.215491 | 0.215696 | 0.214800 | 0.214667 | 0.215212 | 0.219331 | 0.224201 | 0.225213 | 0.224324 | 0.217504 | 0.214719 | 0.218954 | 0.232776 | 0.253919 | 0.272666 | 0.278435 | 0.279210 | 0.261576 | 0.246834 | 0.232984 | 0.225207 | 0.220799 | 0.219903 | 0.220163 | 0.221806 | 0.223497 | 0.222462 | 0.219400 | 0.215687 | 0.215187 | 0.223262 | 0.235604 | 0.254126 | 0.274959 | 0.307439 | 0.373232 | 0.471583 | 0.456549 | 0.433690 | 0.440748 | 0.384649 | 0.307382 | 0.252631 | 0.221432 | 0.217541 | 0.241464 | 0.272844 | 0.277422 | 0.263710 | 0.240508 | 0.229184 | 0.220794 | 0.216066 | 0.215356 | 0.215161 | 0.214970 | 0.219642 | 0.230559 | 0.248683 | 0.270187 | 0.289455 | 0.307980 | 0.321337 | 0.305910 | 0.279044 | 0.249213 | 0.226813 | 0.214761 | 0.217517 | 0.228367 | 0.233632 | 0.231228 | 0.222862 | 0.217611 | 0.214691 | 0.214755 | 0.215089 | 0.215979 | 0.217059 | 0.216884 | 0.216228 | 0.215467 | 0.214628 | 0.215095 | 0.217341 | 0.221006 | 0.225660 | 0.228610 | 0.229215 | 0.227514 | 0.224728 | 0.219966 | 0.217381 | 0.216347 | 0.215714 | 0.215397 | 0.215827 | 0.216000 | 0.215917 | 0.215783 | 0.215062 | 0.214628 | 0.216108 | 0.222308 | 0.233793 | 0.238268 | 0.236243 | 0.226679 | 0.220066 | 0.216797 | 0.216538 | 0.217058 | 0.218048 | 0.216899 | 0.216052 | 0.214744 | 0.214760 | 0.215701 | 0.215965 | 0.215220 | 0.214716 | 0.216908 | 0.219686 | 0.217088 | 0.214632 | 0.219431 | 0.232321 | 0.247245 | 0.258886 | 0.256948 | 0.244129 | 0.230569 | 0.218546 | 0.214963 | 0.225364 | 0.253388 | 0.286811 | 0.319082 | 0.347566 | 0.369044 | 0.361739 | 0.354822 | 0.352347 | 0.366532 | 0.373986 | 0.388408 | 0.415881 | 0.453953 | 0.522183 | 0.622060 | 0.756758 | 1.047104 | 1.307670 | 1.382227 | 1.384809 | 1.173321 | 0.928610 | 0.815201 | 0.735238 | 0.633920 | 0.561114 | 0.464451 | 0.401915 | 0.370748 | 0.341036 |
| right central | 0.219500 | 0.220662 | 0.225431 | 0.241466 | 0.250728 | 0.268021 | 0.256584 | 0.245280 | 0.225046 | 0.214648 | 0.234853 | 0.273966 | 0.331895 | 0.336636 | 0.285838 | 0.236890 | 0.215515 | 0.221548 | 0.222292 | 0.216476 | 0.215600 | 0.235390 | 0.281475 | 0.285481 | 0.270106 | 0.239913 | 0.220278 | 0.215906 | 0.216210 | 0.219226 | 0.229297 | 0.244501 | 0.241446 | 0.220270 | 0.218136 | 0.268939 | 0.394545 | 0.530731 | 0.570045 | 0.456314 | 0.315560 | 0.232997 | 0.214710 | 0.233359 | 0.277172 | 0.353526 | 0.461973 | 0.521167 | 0.483600 | 0.422777 | 0.331480 | 0.264062 | 0.226092 | 0.215008 | 0.243629 | 0.319801 | 0.473581 | 0.687969 | 1.027077 | 1.443310 | 1.988764 | 2.440561 | 2.348382 | 1.259653 | 0.560613 | 0.266604 | 0.215457 | 0.295746 | 0.628353 | 1.555154 | 2.320131 | 2.033458 | 1.094021 | 0.500828 | 0.274976 | 0.214912 | 0.259172 | 0.423363 | 0.825726 | 1.373171 | 1.236988 | 0.694036 | 0.373861 | 0.249829 | 0.216419 | 0.219926 | 0.240163 | 0.253148 | 0.284283 | 0.345514 | 0.471527 | 0.642120 | 0.924450 | 1.229166 | 1.626018 | 1.779668 | 2.050679 | 2.318152 | 2.310811 | 2.051071 | 1.916497 | 1.844101 | 1.547321 | 1.145401 | 0.869900 | 0.648908 | 0.463922 | 0.412976 | 0.435031 | 0.546831 | 0.689247 | 0.877966 | 1.100647 | 1.362985 | 1.393313 | 1.228821 | 0.812192 | 0.525828 | 0.365893 | 0.291517 | 0.263244 | 0.281423 | 0.309396 | 0.349074 | 0.375062 | 0.414879 | 0.437163 | 0.461145 | 0.474640 | 0.510929 | 0.526348 | 0.537292 | 0.497559 | 0.469444 | 0.443636 | 0.423231 | 0.398106 | 0.363222 | 0.306719 | 0.266951 | 0.235625 | 0.219452 | 0.214799 | 0.214776 | 0.216495 | 0.220608 | 0.231274 | 0.251038 | 0.255379 | 0.237094 | 0.219507 | 0.215971 | 0.241076 | 0.281109 | 0.320022 | 0.318459 | 0.279331 | 0.248427 | 0.229936 | 0.220078 | 0.217504 | 0.221209 | 0.229098 | 0.248720 | 0.267394 | 0.284604 | 0.276713 | 0.271370 | 0.268203 | 0.280892 | 0.294503 | 0.326649 | 0.358307 | 0.434754 | 0.506682 | 0.622635 | 0.652200 | 0.621264 | 0.521982 | 0.443202 | 0.370477 | 0.338777 | 0.305666 | 0.293746 | 0.295548 | 0.321190 | 0.375395 | 0.486012 | 0.628622 | 0.757259 | 0.713652 | 0.529112 | 0.414517 | 0.345312 | 0.312448 | 0.300987 | 0.296983 | 0.296348 | 0.295901 | 0.285123 | 0.268790 | 0.262898 | 0.250961 | 0.238488 | 0.228764 | 0.224466 | 0.222896 | 0.229145 | 0.242673 | 0.262992 | 0.293092 | 0.339756 | 0.345719 | 0.332114 | 0.326054 | 0.308987 | 0.302085 | 0.342724 | 0.413211 | 0.483635 | 0.539693 | 0.503339 | 0.415828 | 0.322725 | 0.250385 | 0.218735 | 0.217810 | 0.236395 | 0.251301 | 0.239409 | 0.221413 | 0.216322 | 0.241774 | 0.330940 | 0.505508 | 0.816180 | 1.260260 | 1.826211 | 1.817485 | 1.819209 | 1.397167 | 0.962672 | 0.657809 | 0.494465 | 0.406109 | 0.409727 | 0.477364 | 0.637232 | 0.930987 | 1.213757 | 1.372435 | 1.264291 | 0.908217 | 0.699673 | 0.569878 | 0.483750 | 0.430706 | 0.374508 | 0.325991 | 0.301413 | 0.269962 | 0.247075 | 0.233925 | 0.224104 | 0.219727 | 0.219651 | 0.220777 | 0.224740 | 0.234435 | 0.246872 | 0.265942 | 0.293647 | 0.390388 | 0.589982 | 0.811743 | 1.060419 | 1.207347 | 1.007850 | 0.904569 | 0.835682 |
| left posterior | 0.447815 | 0.560868 | 0.754554 | 1.263173 | 2.651758 | 3.388521 | 2.133159 | 0.766814 | 0.284142 | 0.216030 | 0.294052 | 0.493094 | 0.723950 | 0.745378 | 0.589131 | 0.345135 | 0.227427 | 0.227373 | 0.305014 | 0.373793 | 0.286964 | 0.215935 | 0.257707 | 0.424740 | 0.780737 | 1.404734 | 1.706733 | 1.204464 | 0.737219 | 0.400414 | 0.234187 | 0.226934 | 0.282197 | 0.308523 | 0.288689 | 0.264558 | 0.242374 | 0.229300 | 0.217551 | 0.216300 | 0.235230 | 0.284485 | 0.395145 | 0.600449 | 0.948641 | 1.367282 | 1.615225 | 1.865627 | 2.074747 | 2.516888 | 3.244149 | 2.876657 | 1.597012 | 0.687354 | 0.294214 | 0.214655 | 0.268931 | 0.450337 | 0.727417 | 0.708803 | 0.428569 | 0.226043 | 0.374977 | 2.946129 | 24.978228 | 134.892602 | 528.696772 | 1342.039664 | 1890.035247 | 1189.315523 | 470.173300 | 116.104135 | 27.184552 | 7.288460 | 2.502808 | 1.133097 | 0.633373 | 0.418832 | 0.326334 | 0.287323 | 0.267141 | 0.262875 | 0.268340 | 0.286544 | 0.323925 | 0.371031 | 0.431748 | 0.512429 | 0.652356 | 1.228319 | 3.767398 | 14.777153 | 43.845282 | 80.478075 | 101.791712 | 111.248427 | 92.821120 | 58.289282 | 29.369302 | 14.902196 | 8.253790 | 5.354935 | 4.254406 | 4.525647 | 5.573584 | 7.109383 | 8.636988 | 10.118535 | 10.870215 | 11.140617 | 10.915920 | 10.171908 | 7.644894 | 5.253551 | 3.186889 | 2.050098 | 1.534489 | 1.339302 | 1.272592 | 1.410162 | 1.766972 | 2.037396 | 2.101775 | 1.895929 | 1.365595 | 0.781769 | 0.419234 | 0.259221 | 0.215308 | 0.223297 | 0.241788 | 0.246642 | 0.240348 | 0.227697 | 0.221657 | 0.216070 | 0.214809 | 0.221747 | 0.235881 | 0.262559 | 0.294891 | 0.340631 | 0.420181 | 0.509387 | 0.593150 | 0.656201 | 0.701997 | 0.648971 | 0.594118 | 0.524729 | 0.481764 | 0.437282 | 0.402852 | 0.380916 | 0.355907 | 0.323693 | 0.287430 | 0.264358 | 0.244735 | 0.241027 | 0.246131 | 0.266908 | 0.293770 | 0.318851 | 0.327729 | 0.323653 | 0.302181 | 0.270505 | 0.240641 | 0.220274 | 0.215055 | 0.221936 | 0.225370 | 0.222931 | 0.219080 | 0.214944 | 0.215815 | 0.222875 | 0.239398 | 0.266761 | 0.294221 | 0.306292 | 0.309159 | 0.304150 | 0.297373 | 0.291932 | 0.278904 | 0.257943 | 0.234496 | 0.219932 | 0.214927 | 0.215061 | 0.216964 | 0.215669 | 0.215023 | 0.222082 | 0.242943 | 0.281824 | 0.342743 | 0.426943 | 0.522582 | 0.624813 | 0.707443 | 0.739521 | 0.699831 | 0.650450 | 0.573043 | 0.503210 | 0.450816 | 0.448568 | 0.475643 | 0.553316 | 0.694066 | 0.963785 | 1.259903 | 1.479596 | 1.440126 | 1.297777 | 1.044891 | 0.844091 | 0.651171 | 0.564004 | 0.500627 | 0.441423 | 0.379386 | 0.330846 | 0.282537 | 0.251274 | 0.229278 | 0.221537 | 0.219836 | 0.222708 | 0.235171 | 0.270373 | 0.333553 | 0.438857 | 0.548254 | 0.614402 | 0.575503 | 0.478590 | 0.395498 | 0.341809 | 0.298078 | 0.280214 | 0.267501 | 0.266495 | 0.280718 | 0.305113 | 0.324681 | 0.338746 | 0.325864 | 0.298601 | 0.276732 | 0.254125 | 0.237105 | 0.227452 | 0.221292 | 0.217502 | 0.216625 | 0.215809 | 0.215592 | 0.215324 | 0.214753 | 0.214647 | 0.215148 | 0.219087 | 0.228948 | 0.244851 | 0.268483 | 0.301246 | 0.359737 | 0.492802 | 0.737850 | 1.144155 | 1.694298 | 2.119212 | 1.985729 | 1.664203 | 1.571712 | 1.482108 |
| right posterior | 0.314147 | 0.307652 | 0.315281 | 0.300033 | 0.305946 | 0.348133 | 0.360676 | 0.322901 | 0.252627 | 0.214814 | 0.231531 | 0.270914 | 0.326564 | 0.362054 | 0.334772 | 0.302031 | 0.253875 | 0.222500 | 0.214686 | 0.215725 | 0.217286 | 0.214957 | 0.215132 | 0.216559 | 0.216780 | 0.220875 | 0.229274 | 0.231323 | 0.234677 | 0.244255 | 0.255753 | 0.266902 | 0.248418 | 0.222930 | 0.217308 | 0.278123 | 0.494741 | 0.924292 | 1.865292 | 3.769511 | 7.540660 | 14.418022 | 25.682966 | 41.321581 | 68.526300 | 84.021576 | 40.843411 | 8.867734 | 1.278183 | 0.364672 | 0.225199 | 0.215635 | 0.226659 | 0.229789 | 0.229929 | 0.227434 | 0.232926 | 0.241441 | 0.252778 | 0.244000 | 0.215788 | 0.283450 | 1.272790 | 20.411818 | 385.359978 | 3091.000362 | 7220.605660 | 5015.726770 | 1438.865007 | 248.077721 | 39.238959 | 7.277733 | 1.770356 | 0.530039 | 0.252056 | 0.220970 | 0.371457 | 0.828980 | 1.673736 | 2.777154 | 2.695686 | 1.860739 | 1.039455 | 0.548719 | 0.333444 | 0.247933 | 0.216516 | 0.222288 | 0.264340 | 0.359383 | 0.512756 | 0.861686 | 1.481547 | 2.739232 | 5.219437 | 9.649374 | 14.492209 | 18.413998 | 18.779631 | 16.390051 | 15.176789 | 16.249221 | 21.262920 | 35.506816 | 67.075731 | 92.387752 | 68.255428 | 31.332980 | 12.744341 | 6.070326 | 3.887412 | 2.782955 | 2.102281 | 1.871274 | 1.761413 | 1.884760 | 2.281998 | 2.917830 | 4.073160 | 6.409658 | 9.550245 | 12.427452 | 13.721411 | 12.445522 | 9.580672 | 6.630466 | 4.125725 | 2.534775 | 1.569226 | 1.159660 | 0.903727 | 0.811205 | 0.795110 | 0.800065 | 0.833948 | 0.998004 | 1.195220 | 1.332084 | 1.413197 | 1.365461 | 1.305810 | 1.199337 | 1.138246 | 1.281499 | 1.905752 | 2.977052 | 4.233349 | 4.824114 | 4.006746 | 3.011455 | 2.180938 | 1.567530 | 1.126548 | 0.886328 | 0.780552 | 0.799624 | 0.887130 | 1.112608 | 1.476607 | 1.981244 | 2.361479 | 2.928537 | 3.229849 | 3.185012 | 2.659249 | 2.213600 | 1.770705 | 1.571365 | 1.337475 | 1.182345 | 1.097978 | 1.169779 | 1.436054 | 2.231881 | 3.500147 | 5.129415 | 5.431460 | 4.375641 | 3.086304 | 2.107603 | 1.378081 | 0.922450 | 0.659599 | 0.514275 | 0.452457 | 0.466116 | 0.522448 | 0.597594 | 0.700125 | 0.810516 | 0.855069 | 0.840497 | 0.794494 | 0.776634 | 0.793354 | 0.874163 | 0.940654 | 0.964847 | 0.983960 | 1.072784 | 1.248295 | 1.611160 | 2.282416 | 3.386204 | 5.085099 | 7.500605 | 9.782265 | 11.115547 | 10.298135 | 7.790008 | 6.015948 | 5.001239 | 4.728125 | 5.672410 | 6.811529 | 7.440283 | 7.664683 | 6.895157 | 5.089380 | 3.582564 | 2.362336 | 1.672742 | 1.373968 | 1.292763 | 1.383318 | 1.715392 | 2.069444 | 2.415840 | 2.593625 | 2.604850 | 2.378710 | 2.107384 | 1.855910 | 1.627690 | 1.448121 | 1.331503 | 1.234343 | 1.237086 | 1.330245 | 1.509881 | 1.688452 | 1.911405 | 2.124889 | 2.417606 | 2.890226 | 3.939657 | 6.931361 | 12.958103 | 18.370978 | 15.450256 | 9.168423 | 5.409744 | 3.274177 | 2.341846 | 2.103820 | 2.460075 | 3.167922 | 4.443853 | 5.821640 | 6.826577 | 5.998779 | 4.503722 | 3.271360 | 2.286719 | 1.607092 | 1.085673 | 0.760976 | 0.593593 | 0.576348 | 0.599293 | 0.721029 | 0.990295 | 1.316490 | 1.478538 | 1.696356 | 1.565485 | 1.317543 | 1.229518 | 1.148956 | 1.085520 |
| all electrodes | 0.423870 | 0.421077 | 0.372900 | 0.260544 | 0.216980 | 0.219542 | 0.277640 | 0.410498 | 0.577172 | 0.804918 | 0.966878 | 0.834463 | 0.846635 | 0.684298 | 0.431294 | 0.291858 | 0.237167 | 0.216484 | 0.214988 | 0.223025 | 0.235704 | 0.239564 | 0.223231 | 0.215947 | 0.224817 | 0.299202 | 0.488754 | 0.655651 | 0.847338 | 0.773905 | 0.609875 | 0.417006 | 0.289212 | 0.240452 | 0.238269 | 0.235044 | 0.235894 | 0.242112 | 0.248829 | 0.238184 | 0.223357 | 0.219047 | 0.214661 | 0.219677 | 0.227053 | 0.229033 | 0.248813 | 0.236730 | 0.250650 | 0.258173 | 0.264067 | 0.272061 | 0.291087 | 0.316192 | 0.360220 | 0.410439 | 0.401931 | 0.385154 | 0.344411 | 0.290681 | 0.225965 | 0.220771 | 0.344231 | 0.923665 | 3.739782 | 15.681105 | 54.893613 | 68.270077 | 42.384338 | 19.043924 | 12.674167 | 10.987815 | 9.460335 | 5.400893 | 2.342025 | 0.795309 | 0.357139 | 0.236091 | 0.214683 | 0.217306 | 0.215433 | 0.215783 | 0.229958 | 0.246337 | 0.243703 | 0.237468 | 0.228380 | 0.218317 | 0.216891 | 0.221254 | 0.241686 | 0.310662 | 0.463126 | 0.729881 | 1.313507 | 2.065180 | 3.094512 | 4.068894 | 3.734617 | 2.219521 | 1.174195 | 0.479896 | 0.268550 | 0.225318 | 0.216680 | 0.219560 | 0.256489 | 0.352245 | 0.579515 | 0.984699 | 1.597924 | 1.852254 | 1.958196 | 1.456113 | 1.203168 | 0.951831 | 0.857083 | 0.722021 | 0.729931 | 0.807586 | 1.063567 | 1.456985 | 2.393119 | 2.832462 | 2.311566 | 1.460366 | 0.955729 | 0.609730 | 0.440598 | 0.345619 | 0.304477 | 0.297845 | 0.333842 | 0.394285 | 0.489385 | 0.628535 | 0.837175 | 1.119228 | 1.811934 | 2.834453 | 4.160280 | 4.546602 | 3.563034 | 3.250292 | 3.672235 | 3.865232 | 3.633326 | 3.379627 | 3.282701 | 3.291075 | 3.321990 | 3.145645 | 3.099420 | 2.346530 | 1.449184 | 0.863011 | 0.606609 | 0.468918 | 0.451016 | 0.423866 | 0.383676 | 0.325924 | 0.277032 | 0.239441 | 0.222240 | 0.214780 | 0.216679 | 0.220511 | 0.221743 | 0.221316 | 0.221467 | 0.222769 | 0.224009 | 0.227951 | 0.232067 | 0.226145 | 0.217600 | 0.215078 | 0.229592 | 0.272700 | 0.360975 | 0.526766 | 0.673316 | 0.783917 | 0.794520 | 0.706746 | 0.523880 | 0.361408 | 0.252761 | 0.218954 | 0.215737 | 0.218493 | 0.216392 | 0.214747 | 0.221354 | 0.241706 | 0.263373 | 0.272696 | 0.267393 | 0.258041 | 0.252412 | 0.256743 | 0.287704 | 0.369201 | 0.465178 | 0.594642 | 0.766620 | 0.950414 | 1.077552 | 1.229716 | 1.142805 | 0.932693 | 0.624488 | 0.416765 | 0.303811 | 0.265543 | 0.256618 | 0.276309 | 0.340526 | 0.425048 | 0.431742 | 0.365726 | 0.301331 | 0.255889 | 0.240737 | 0.247347 | 0.298258 | 0.406702 | 0.607035 | 0.890508 | 1.128180 | 0.931183 | 0.793135 | 0.666449 | 0.608705 | 0.596045 | 0.573424 | 0.508046 | 0.436044 | 0.364138 | 0.301317 | 0.266704 | 0.254379 | 0.257067 | 0.265222 | 0.292167 | 0.316934 | 0.327318 | 0.325417 | 0.300839 | 0.255085 | 0.225182 | 0.214665 | 0.221468 | 0.233353 | 0.252578 | 0.260706 | 0.243598 | 0.223719 | 0.214658 | 0.229012 | 0.286113 | 0.376244 | 0.507418 | 0.597267 | 0.588299 | 0.488291 | 0.354108 | 0.261087 | 0.223329 | 0.214693 | 0.225792 | 0.244617 | 0.264005 | 0.274426 | 0.291209 | 0.306375 | 0.325053 | 0.321090 | 0.343393 |

N) maximum cross-correlation, neutral vs angry

  
|  | time window | peak latency | cluster *p* | peak Cohen's *d* |  | | | |
| **all electrodes** | 125 - 600 ms | 160 ms | 0.0002 | 1.3091 |  | | | |
|  | | | | | | | | |

Model correlations, cluster permutation tests

|  | **left hemisphere** | | | | **right hemisphere** | | | |
|  | time window | peak latency | cluster *p* | peak Cohen's *d* | time window | peak latency | cluster *p* | peak Cohen's *d* |
| **anterior** |  | | | |  | | | |
| **central** |  | | | | 120 - 470 ms | 160 ms | 0.0002 | 1.2993 |
| **posterior** | 120 - 230 ms | 160 ms | 0.0196 | 1.1274 | 110 - 1195 ms | 155 ms | 0.0002 | 1.2785 |
 250 - 540 ms | 415 ms | 0.0043 | 0.8301 |  | | | | 580 - 815 ms | 660 ms | 0.0115 | 0.5764 |  | | | |

  

Model correlations, Bayesian statistics

|  | -200 | -195 | -190 | -185 | -180 | -175 | -170 | -165 | -160 | -155 | -150 | -145 | -140 | -135 | -130 | -125 | -120 | -115 | -110 | -105 | -100 | -95 | -90 | -85 | -80 | -75 | -70 | -65 | -60 | -55 | -50 | -45 | -40 | -35 | -30 | -25 | -20 | -15 | -10 | -5 | 0 | 5 | 10 | 15 | 20 | 25 | 30 | 35 | 40 | 45 | 50 | 55 | 60 | 65 | 70 | 75 | 80 | 85 | 90 | 95 | 100 | 105 | 110 | 115 | 120 | 125 | 130 | 135 | 140 | 145 | 150 | 155 | 160 | 165 | 170 | 175 | 180 | 185 | 190 | 195 | 200 | 205 | 210 | 215 | 220 | 225 | 230 | 235 | 240 | 245 | 250 | 255 | 260 | 265 | 270 | 275 | 280 | 285 | 290 | 295 | 300 | 305 | 310 | 315 | 320 | 325 | 330 | 335 | 340 | 345 | 350 | 355 | 360 | 365 | 370 | 375 | 380 | 385 | 390 | 395 | 400 | 405 | 410 | 415 | 420 | 425 | 430 | 435 | 440 | 445 | 450 | 455 | 460 | 465 | 470 | 475 | 480 | 485 | 490 | 495 | 500 | 505 | 510 | 515 | 520 | 525 | 530 | 535 | 540 | 545 | 550 | 555 | 560 | 565 | 570 | 575 | 580 | 585 | 590 | 595 | 600 | 605 | 610 | 615 | 620 | 625 | 630 | 635 | 640 | 645 | 650 | 655 | 660 | 665 | 670 | 675 | 680 | 685 | 690 | 695 | 700 | 705 | 710 | 715 | 720 | 725 | 730 | 735 | 740 | 745 | 750 | 755 | 760 | 765 | 770 | 775 | 780 | 785 | 790 | 795 | 800 | 805 | 810 | 815 | 820 | 825 | 830 | 835 | 840 | 845 | 850 | 855 | 860 | 865 | 870 | 875 | 880 | 885 | 890 | 895 | 900 | 905 | 910 | 915 | 920 | 925 | 930 | 935 | 940 | 945 | 950 | 955 | 960 | 965 | 970 | 975 | 980 | 985 | 990 | 995 | 1000 | 1005 | 1010 | 1015 | 1020 | 1025 | 1030 | 1035 | 1040 | 1045 | 1050 | 1055 | 1060 | 1065 | 1070 | 1075 | 1080 | 1085 | 1090 | 1095 | 1100 | 1105 | 1110 | 1115 | 1120 | 1125 | 1130 | 1135 | 1140 | 1145 | 1150 | 1155 | 1160 | 1165 | 1170 | 1175 | 1180 | 1185 | 1190 | 1195 |
| --- | --- | --- | --- | --- | --- | --- | --- | --- | --- | --- | --- | --- | --- | --- | --- | --- | --- | --- | --- | --- | --- | --- | --- | --- | --- | --- | --- | --- | --- | --- | --- | --- | --- | --- | --- | --- | --- | --- | --- | --- | --- | --- | --- | --- | --- | --- | --- | --- | --- | --- | --- | --- | --- | --- | --- | --- | --- | --- | --- | --- | --- | --- | --- | --- | --- | --- | --- | --- | --- | --- | --- | --- | --- | --- | --- | --- | --- | --- | --- | --- | --- | --- | --- | --- | --- | --- | --- | --- | --- | --- | --- | --- | --- | --- | --- | --- | --- | --- | --- | --- | --- | --- | --- | --- | --- | --- | --- | --- | --- | --- | --- | --- | --- | --- | --- | --- | --- | --- | --- | --- | --- | --- | --- | --- | --- | --- | --- | --- | --- | --- | --- | --- | --- | --- | --- | --- | --- | --- | --- | --- | --- | --- | --- | --- | --- | --- | --- | --- | --- | --- | --- | --- | --- | --- | --- | --- | --- | --- | --- | --- | --- | --- | --- | --- | --- | --- | --- | --- | --- | --- | --- | --- | --- | --- | --- | --- | --- | --- | --- | --- | --- | --- | --- | --- | --- | --- | --- | --- | --- | --- | --- | --- | --- | --- | --- | --- | --- | --- | --- | --- | --- | --- | --- | --- | --- | --- | --- | --- | --- | --- | --- | --- | --- | --- | --- | --- | --- | --- | --- | --- | --- | --- | --- | --- | --- | --- | --- | --- | --- | --- | --- | --- | --- | --- | --- | --- | --- | --- | --- | --- | --- | --- | --- | --- | --- | --- | --- | --- | --- | --- | --- | --- | --- | --- | --- | --- | --- | --- | --- | --- | --- | --- | --- | --- | --- | --- | --- | --- | --- | --- | --- | --- | --- | --- | --- | --- | --- | --- | --- | --- |
| left anterior | 0.312903 | 0.337507 | 0.379526 | 0.506190 | 0.736406 | 1.696644 | 3.651648 | 8.911334 | 27.574472 | 37.147036 | 13.050861 | 3.817099 | 1.278038 | 0.523060 | 0.309215 | 0.269876 | 0.293115 | 0.404859 | 0.817953 | 2.094183 | 4.809566 | 7.598192 | 4.569053 | 1.204100 | 0.411462 | 0.238188 | 0.222904 | 0.323159 | 0.540142 | 0.625964 | 0.560073 | 0.315818 | 0.226114 | 0.221108 | 0.257133 | 0.264300 | 0.247016 | 0.243173 | 0.243605 | 0.240138 | 0.241580 | 0.255687 | 0.301936 | 0.358399 | 0.434339 | 0.740390 | 1.483203 | 1.742634 | 1.540251 | 0.877507 | 0.480827 | 0.327303 | 0.251378 | 0.222618 | 0.215878 | 0.214687 | 0.218558 | 0.228588 | 0.239837 | 0.245242 | 0.255911 | 0.243273 | 0.231796 | 0.233404 | 0.271877 | 0.350420 | 0.564515 | 0.840830 | 1.152464 | 1.155312 | 1.027371 | 0.718385 | 0.496180 | 0.341340 | 0.245958 | 0.216117 | 0.276020 | 0.503103 | 1.038132 | 1.618274 | 1.570436 | 1.153670 | 0.775142 | 0.536841 | 0.354748 | 0.268959 | 0.233081 | 0.216281 | 0.217779 | 0.239827 | 0.292803 | 0.379712 | 0.491992 | 0.629822 | 0.646463 | 5.749651e-01 | 4.557892e-01 | 3.419325e-01 | 2.595167e-01 | 0.223933 | 0.214626 | 0.218894 | 0.225295 | 0.226291 | 0.226666 | 0.224020 | 0.219384 | 0.215304 | 0.214776 | 0.217690 | 0.230144 | 0.251448 | 0.283693 | 0.314519 | 0.356987 | 0.396022 | 0.466410 | 0.636631 | 1.080458 | 2.008181 | 3.195388 | 4.253502 | 3.810165 | 3.291832 | 2.702017 | 2.028962 | 1.426485 | 0.994712 | 0.643949 | 0.423496 | 0.315186 | 0.258869 | 0.229075 | 0.217342 | 0.214658 | 0.217368 | 0.224706 | 0.223599 | 0.217949 | 0.214924 | 0.219358 | 0.224193 | 0.226649 | 0.228458 | 0.220502 | 0.215621 | 0.215322 | 0.231328 | 0.288718 | 0.398488 | 0.526810 | 0.625492 | 0.645360 | 0.641603 | 0.587981 | 0.469933 | 0.349939 | 0.286588 | 0.248295 | 0.224422 | 0.215434 | 0.214662 | 0.216155 | 0.220292 | 0.228799 | 0.248871 | 0.296437 | 0.380185 | 0.510732 | 0.639246 | 0.691398 | 0.587820 | 0.449864 | 0.325901 | 0.257851 | 0.229461 | 0.224356 | 0.228919 | 0.249738 | 0.274794 | 0.299789 | 0.309242 | 0.307024 | 0.308065 | 0.328305 | 0.361971 | 0.398269 | 0.452317 | 0.511332 | 0.527759 | 0.432341 | 0.349008 | 0.294572 | 0.262975 | 0.250447 | 0.255162 | 0.285941 | 0.334906 | 0.371560 | 0.365990 | 0.336172 | 0.287343 | 0.240179 | 0.215021 | 0.227046 | 0.257789 | 0.271840 | 0.244976 | 0.217375 | 0.224341 | 0.279259 | 0.353563 | 0.403030 | 0.428496 | 0.375998 | 0.297155 | 0.250277 | 0.228538 | 0.222035 | 0.221974 | 0.219500 | 0.214952 | 0.216631 | 0.231951 | 0.287791 | 0.422497 | 0.648800 | 0.896970 | 0.919817 | 0.783905 | 0.650406 | 0.630700 | 0.740263 | 1.088800 | 1.578934 | 2.094632 | 2.402205 | 1.833634 | 1.103376 | 0.654516 | 0.409003 | 0.289440 | 0.243444 | 0.223812 | 0.217826 | 0.215663 | 0.215450 | 0.215581 | 0.217500 | 0.221248 | 0.227321 | 0.229084 | 0.226694 | 0.220651 | 0.216433 | 0.214743 | 0.214697 | 0.215456 | 0.215336 | 0.215287 | 0.215440 | 0.215456 | 0.215796 | 0.217661 | 0.220232 | 0.225571 | 0.238851 | 0.251744 | 0.260386 | 0.273437 | 0.279679 | 0.281751 | 0.289834 | 0.303667 | 0.323362 | 0.365385 | 0.427834 | 0.516496 | 0.571751 | 0.663845 |
| right anterior | 0.222534 | 0.220786 | 0.215039 | 0.223731 | 0.253534 | 0.308395 | 0.364584 | 0.385204 | 0.412583 | 0.444018 | 0.540324 | 0.998370 | 1.292860 | 0.949518 | 0.718572 | 0.523707 | 0.389901 | 0.338871 | 0.300875 | 0.273929 | 0.266373 | 0.246966 | 0.241616 | 0.260875 | 0.264820 | 0.264525 | 0.264224 | 0.276774 | 0.290230 | 0.290933 | 0.279510 | 0.340971 | 0.373936 | 0.345927 | 0.298702 | 0.242759 | 0.214682 | 0.239853 | 0.331520 | 0.363465 | 0.318206 | 0.284702 | 0.236581 | 0.214628 | 0.225680 | 0.251740 | 0.268553 | 0.247844 | 0.216615 | 0.219959 | 0.246339 | 0.268281 | 0.295241 | 0.303711 | 0.328850 | 0.334177 | 0.349353 | 0.364377 | 0.373244 | 0.341848 | 0.311325 | 0.260124 | 0.235876 | 0.233814 | 0.256943 | 0.333683 | 0.597608 | 1.331519 | 3.184237 | 6.560816 | 11.541875 | 15.514125 | 13.266080 | 7.938307 | 3.904824 | 2.067547 | 1.202972 | 0.776118 | 0.545791 | 0.445181 | 0.354836 | 0.306085 | 0.277783 | 0.247731 | 0.228615 | 0.221987 | 0.221117 | 0.224158 | 0.242529 | 0.311225 | 0.512089 | 1.019997 | 2.169778 | 4.824168 | 9.302744 | 1.366447e+01 | 1.457458e+01 | 1.276400e+01 | 9.109066e+00 | 5.794092 | 3.164316 | 1.685868 | 0.901895 | 0.609810 | 0.481350 | 0.440033 | 0.397560 | 0.362559 | 0.327090 | 0.294516 | 0.268979 | 0.261789 | 0.259320 | 0.277685 | 0.314127 | 0.353869 | 0.437141 | 0.591334 | 0.806904 | 1.232320 | 2.041025 | 3.423637 | 5.372886 | 6.877820 | 7.467516 | 7.777976 | 6.874363 | 5.874839 | 4.818026 | 3.848833 | 2.907513 | 2.164575 | 1.477421 | 1.081616 | 0.708958 | 0.484501 | 0.355452 | 0.276330 | 0.231100 | 0.215726 | 0.216196 | 0.221351 | 0.222154 | 0.223426 | 0.221960 | 0.221426 | 0.224352 | 0.226647 | 0.223901 | 0.216905 | 0.215497 | 0.229748 | 0.270403 | 0.358935 | 0.447413 | 0.435123 | 0.359630 | 0.270407 | 0.218242 | 0.229781 | 0.307706 | 0.442408 | 0.635970 | 0.775143 | 0.832489 | 0.847316 | 0.939726 | 0.958090 | 0.956766 | 0.931629 | 0.745156 | 0.515862 | 0.385286 | 0.322895 | 0.293162 | 0.291674 | 0.290513 | 0.295086 | 0.307442 | 0.309883 | 0.302382 | 0.314805 | 0.331483 | 0.340165 | 0.348347 | 0.362477 | 0.382465 | 0.393507 | 0.388407 | 0.367326 | 0.353710 | 0.319642 | 0.294136 | 0.277958 | 0.267377 | 0.259192 | 0.252964 | 0.248491 | 0.259389 | 0.290228 | 0.323743 | 0.362009 | 0.381546 | 0.377388 | 0.370449 | 0.368160 | 0.372330 | 0.381372 | 0.357252 | 0.316668 | 0.279786 | 0.255383 | 0.240010 | 0.235081 | 0.234724 | 0.240897 | 0.253287 | 0.275512 | 0.304477 | 0.352602 | 0.390185 | 0.417412 | 0.427212 | 0.460734 | 0.514600 | 0.630157 | 0.771057 | 0.869198 | 0.860604 | 0.751134 | 0.594376 | 0.493826 | 0.445921 | 0.409981 | 0.412817 | 0.431471 | 0.471191 | 0.516533 | 0.559650 | 0.548035 | 0.508219 | 0.418869 | 0.332078 | 0.278636 | 0.251540 | 0.238653 | 0.232891 | 0.238253 | 0.250570 | 0.266605 | 0.284160 | 0.316735 | 0.346369 | 0.380733 | 0.390646 | 0.378970 | 0.347263 | 0.309428 | 0.263485 | 0.234779 | 0.220076 | 0.214883 | 0.215786 | 0.217125 | 0.216147 | 0.214635 | 0.218894 | 0.238118 | 0.273911 | 0.342874 | 0.417492 | 0.463806 | 0.503269 | 0.510711 | 0.460480 | 0.422950 | 0.373625 | 0.342514 | 0.323465 | 0.307185 |
| left central | 0.218811 | 0.232447 | 0.243062 | 0.260215 | 0.271868 | 0.247507 | 0.222772 | 0.214631 | 0.221629 | 0.238725 | 0.261392 | 0.265089 | 0.242647 | 0.222336 | 0.214932 | 0.218971 | 0.225657 | 0.237774 | 0.250928 | 0.258591 | 0.257204 | 0.226281 | 0.214635 | 0.226507 | 0.297758 | 0.444878 | 0.589784 | 0.680851 | 0.531725 | 0.347836 | 0.262341 | 0.230911 | 0.236680 | 0.298481 | 0.539030 | 1.204562 | 3.115405 | 6.933875 | 19.729372 | 38.847483 | 47.992484 | 22.453998 | 8.734413 | 3.327216 | 1.462229 | 0.722870 | 0.474293 | 0.366516 | 0.295048 | 0.265798 | 0.265341 | 0.265127 | 0.271266 | 0.289910 | 0.304964 | 0.335242 | 0.407166 | 0.555279 | 0.812608 | 1.174474 | 1.235019 | 0.770981 | 0.357379 | 0.272855 | 0.285215 | 0.451093 | 1.208583 | 5.311588 | 21.439524 | 68.689100 | 164.534259 | 306.005515 | 303.514279 | 200.795792 | 94.948870 | 33.142179 | 8.774862 | 2.336848 | 0.757996 | 0.398090 | 0.276223 | 0.230059 | 0.220839 | 0.223630 | 0.224148 | 0.235694 | 0.259166 | 0.287599 | 0.334628 | 0.443950 | 0.725750 | 1.779391 | 6.390702 | 26.409289 | 85.393794 | 1.738592e+02 | 2.152015e+02 | 1.373420e+02 | 5.782425e+01 | 20.018789 | 8.047370 | 3.889637 | 1.872708 | 0.940205 | 0.574819 | 0.432205 | 0.356798 | 0.323002 | 0.319995 | 0.323682 | 0.328143 | 0.326775 | 0.336306 | 0.349142 | 0.391799 | 0.447626 | 0.515689 | 0.594923 | 0.753458 | 0.941383 | 1.098793 | 1.114631 | 1.106747 | 1.019135 | 0.883273 | 0.736211 | 0.674219 | 0.663223 | 0.606074 | 0.493464 | 0.383399 | 0.293447 | 0.232425 | 0.215265 | 0.215009 | 0.214894 | 0.224725 | 0.251884 | 0.312203 | 0.417922 | 0.481918 | 0.412268 | 0.306554 | 0.227482 | 0.218007 | 0.250837 | 0.272848 | 0.258499 | 0.232141 | 0.215158 | 0.222063 | 0.255005 | 0.290606 | 0.312497 | 0.308263 | 0.292229 | 0.264321 | 0.248382 | 0.233093 | 0.222437 | 0.218665 | 0.216004 | 0.214669 | 0.215071 | 0.218854 | 0.224842 | 0.229572 | 0.236767 | 0.244349 | 0.245245 | 0.239776 | 0.234226 | 0.225328 | 0.216337 | 0.215897 | 0.222984 | 0.233856 | 0.241297 | 0.245951 | 0.245862 | 0.238598 | 0.231298 | 0.229510 | 0.226137 | 0.222294 | 0.219448 | 0.218209 | 0.216881 | 0.215626 | 0.214831 | 0.215031 | 0.215773 | 0.218437 | 0.227694 | 0.250981 | 0.295109 | 0.347043 | 0.370287 | 0.361343 | 0.327890 | 0.271092 | 0.227743 | 0.214794 | 0.231222 | 0.279201 | 0.337298 | 0.354872 | 0.320681 | 0.270629 | 0.236119 | 0.222967 | 0.221595 | 0.225156 | 0.231516 | 0.243482 | 0.250323 | 0.247204 | 0.240466 | 0.239701 | 0.240723 | 0.247919 | 0.253058 | 0.262629 | 0.270150 | 0.246514 | 0.225376 | 0.218049 | 0.214765 | 0.216121 | 0.222501 | 0.245603 | 0.281953 | 0.333442 | 0.400467 | 0.450732 | 0.425708 | 0.352986 | 0.272920 | 0.232270 | 0.214833 | 0.223253 | 0.245085 | 0.279051 | 0.330784 | 0.422518 | 0.591261 | 0.787773 | 1.007051 | 1.346258 | 1.826077 | 2.239643 | 2.382420 | 2.107319 | 1.491950 | 0.765155 | 0.404557 | 0.279991 | 0.235604 | 0.223082 | 0.221947 | 0.232009 | 0.259590 | 0.310592 | 0.370974 | 0.408057 | 0.422905 | 0.422289 | 0.374190 | 0.327428 | 0.279533 | 0.239484 | 0.218864 | 0.214627 | 0.216182 | 0.216942 | 0.215612 | 0.214678 | 0.218562 | 0.221887 | 0.226660 |
| right central | 0.590682 | 0.638452 | 0.711035 | 0.727208 | 0.734662 | 0.907336 | 1.389492 | 2.002696 | 3.291757 | 4.344309 | 3.440295 | 1.569085 | 0.668970 | 0.294025 | 0.224718 | 0.215161 | 0.233371 | 0.263380 | 0.284331 | 0.306378 | 0.300467 | 0.275868 | 0.244853 | 0.215316 | 0.236466 | 0.390354 | 0.974137 | 3.130997 | 7.321813 | 10.957484 | 4.191017 | 1.143092 | 0.448540 | 0.283320 | 0.233480 | 0.214671 | 0.229818 | 0.279217 | 0.390824 | 0.563036 | 0.643880 | 0.728341 | 0.769523 | 0.867196 | 0.873232 | 0.793498 | 0.564123 | 0.391594 | 0.275219 | 0.230736 | 0.216279 | 0.214695 | 0.218987 | 0.227038 | 0.233203 | 0.232937 | 0.220559 | 0.215359 | 0.233744 | 0.281894 | 0.360734 | 0.434842 | 0.529075 | 0.711318 | 1.135257 | 2.597097 | 8.885017 | 51.354159 | 382.642644 | 2462.199073 | 8024.671497 | 16514.021309 | 15323.338110 | 7175.675452 | 2166.741994 | 519.985008 | 123.130162 | 36.556476 | 11.252579 | 4.363232 | 2.464680 | 1.685004 | 1.448540 | 1.414292 | 1.560708 | 1.888990 | 2.688065 | 4.130207 | 8.224409 | 27.064568 | 117.743262 | 836.975126 | 9466.552169 | 101623.786095 | 676206.618663 | 2.257175e+06 | 3.471040e+06 | 2.877229e+06 | 1.440025e+06 | 364080.301370 | 58556.804020 | 10752.610551 | 2077.587827 | 512.122346 | 174.290660 | 77.207233 | 37.189256 | 20.149489 | 13.408031 | 12.102118 | 12.300323 | 16.574399 | 33.508597 | 86.945873 | 216.768352 | 552.719603 | 1102.395815 | 1921.971773 | 2786.720307 | 3575.898011 | 4134.325133 | 4833.651730 | 3395.816907 | 1610.865160 | 739.417931 | 306.852592 | 117.130286 | 52.406752 | 28.192997 | 14.815177 | 8.474925 | 4.662348 | 2.454083 | 1.487941 | 0.975149 | 0.811177 | 0.818677 | 0.988793 | 1.137417 | 1.242907 | 1.161927 | 1.039059 | 0.969402 | 0.927028 | 0.844297 | 0.740837 | 0.547874 | 0.417390 | 0.339930 | 0.306465 | 0.293486 | 0.310896 | 0.353696 | 0.468827 | 0.785817 | 1.799897 | 4.707776 | 11.459494 | 18.341787 | 19.458133 | 14.133193 | 8.380786 | 3.383890 | 1.394326 | 0.741858 | 0.490655 | 0.390877 | 0.379080 | 0.433091 | 0.549081 | 0.758462 | 1.047644 | 1.442467 | 1.747212 | 1.909682 | 1.882710 | 2.033252 | 1.971545 | 1.860631 | 1.702126 | 1.540331 | 1.298259 | 1.058612 | 0.843461 | 0.684600 | 0.537708 | 0.427110 | 0.369500 | 0.337163 | 0.306064 | 0.305154 | 0.308300 | 0.313111 | 0.327397 | 0.365586 | 0.375912 | 0.394150 | 0.438319 | 0.542342 | 0.588887 | 0.573006 | 0.527155 | 0.473881 | 0.389023 | 0.323923 | 0.279424 | 0.256686 | 0.243470 | 0.237745 | 0.244884 | 0.280414 | 0.375553 | 0.633251 | 1.314868 | 2.558335 | 4.750719 | 5.993126 | 5.464776 | 4.027457 | 2.605077 | 1.434046 | 1.011546 | 0.846860 | 0.917098 | 1.209382 | 1.919623 | 3.094109 | 4.507586 | 4.428837 | 2.833749 | 1.519111 | 0.796187 | 0.484587 | 0.337682 | 0.296091 | 0.295822 | 0.316977 | 0.371086 | 0.442560 | 0.440422 | 0.359235 | 0.262023 | 0.215938 | 0.228615 | 0.270626 | 0.306769 | 0.287099 | 0.247917 | 0.218817 | 0.215158 | 0.218383 | 0.220776 | 0.227382 | 0.235661 | 0.258574 | 0.323173 | 0.431318 | 0.689044 | 1.161206 | 1.744624 | 2.095813 | 2.245556 | 1.783595 | 1.377589 | 0.946972 | 0.572429 | 0.388348 | 0.313019 | 0.273710 | 0.263182 | 0.277931 | 0.292982 | 0.320446 | 0.343357 | 0.378682 | 0.416324 | 0.493363 | 0.587261 | 0.660193 | 0.730089 |
| left posterior | 0.216657 | 0.214681 | 0.217753 | 0.235348 | 0.279921 | 0.361384 | 0.485369 | 0.569713 | 0.466320 | 0.305576 | 0.231925 | 0.214969 | 0.215181 | 0.214638 | 0.220840 | 0.279859 | 0.454307 | 0.783265 | 1.915159 | 4.284848 | 3.531349 | 1.013448 | 0.312518 | 0.216597 | 0.329974 | 0.691434 | 1.124138 | 1.040935 | 0.738435 | 0.502149 | 0.361851 | 0.327421 | 0.383896 | 0.603910 | 1.121462 | 1.821817 | 2.658969 | 3.185567 | 3.054294 | 2.685462 | 2.332627 | 1.958102 | 1.887513 | 1.920046 | 1.519483 | 1.011262 | 0.667327 | 0.431311 | 0.298838 | 0.236929 | 0.215249 | 0.222220 | 0.247379 | 0.282755 | 0.295744 | 0.293829 | 0.299578 | 0.310514 | 0.339583 | 0.396171 | 0.362782 | 0.262551 | 0.214654 | 0.323263 | 1.068517 | 4.826420 | 20.678192 | 71.366852 | 195.283308 | 438.176911 | 820.026175 | 1204.760405 | 1438.433555 | 1560.991508 | 1452.770798 | 1147.794428 | 827.925420 | 703.334355 | 665.895397 | 605.302011 | 392.594694 | 143.075763 | 35.737276 | 9.154363 | 3.133206 | 1.683774 | 1.244854 | 0.971086 | 0.850957 | 0.883817 | 1.084192 | 1.535414 | 2.754390 | 5.631086 | 13.264300 | 2.943498e+01 | 4.312725e+01 | 3.694849e+01 | 2.509566e+01 | 14.387490 | 8.969014 | 5.973702 | 4.123912 | 2.889883 | 2.193648 | 1.719808 | 1.565004 | 1.594617 | 1.864616 | 2.317888 | 3.013899 | 3.536540 | 3.967120 | 4.116969 | 4.310848 | 4.510435 | 5.446899 | 6.692653 | 9.070139 | 12.568604 | 18.433864 | 27.524071 | 43.714615 | 67.012605 | 95.304451 | 102.454313 | 80.582033 | 45.814723 | 21.867597 | 9.382769 | 4.643256 | 2.654339 | 1.854200 | 1.504800 | 1.443261 | 1.591757 | 2.035490 | 2.805573 | 4.027202 | 6.074007 | 8.686234 | 11.861636 | 15.522921 | 19.272328 | 19.278628 | 13.972375 | 7.147143 | 3.227213 | 1.638216 | 1.000735 | 0.740960 | 0.633971 | 0.620567 | 0.678850 | 0.820251 | 1.068929 | 1.538477 | 2.344284 | 3.099807 | 3.609997 | 4.112156 | 4.458671 | 4.480720 | 4.167622 | 3.700932 | 3.229214 | 3.054866 | 3.181089 | 3.818184 | 4.623704 | 5.083684 | 5.059819 | 4.663101 | 4.384561 | 4.433897 | 4.117468 | 3.655626 | 3.533074 | 3.263228 | 3.078317 | 3.146842 | 3.216576 | 2.992433 | 2.931341 | 2.551656 | 2.337339 | 2.127255 | 1.944968 | 1.676346 | 1.520957 | 1.407775 | 1.335235 | 1.386774 | 1.603711 | 2.015445 | 2.456517 | 3.385539 | 4.605206 | 5.734272 | 5.626393 | 4.778209 | 3.211517 | 2.169745 | 1.450629 | 0.992825 | 0.696268 | 0.516087 | 0.416921 | 0.374808 | 0.369721 | 0.406770 | 0.511033 | 0.752522 | 1.233175 | 1.945773 | 2.609391 | 2.701933 | 2.125076 | 1.285306 | 0.779217 | 0.487370 | 0.334464 | 0.265963 | 0.241120 | 0.231184 | 0.232996 | 0.242993 | 0.255084 | 0.271440 | 0.296260 | 0.337288 | 0.393325 | 0.480873 | 0.629688 | 0.867222 | 1.113569 | 1.287303 | 1.408341 | 1.411806 | 1.238659 | 1.022189 | 0.857925 | 0.743962 | 0.697670 | 0.718395 | 0.876655 | 1.363003 | 2.357259 | 3.212566 | 3.324271 | 2.507503 | 1.608080 | 1.077455 | 0.754713 | 0.564277 | 0.498857 | 0.472213 | 0.473484 | 0.523107 | 0.590495 | 0.584080 | 0.515434 | 0.421505 | 0.328101 | 0.260079 | 0.225778 | 0.214742 | 0.221025 | 0.242218 | 0.275394 | 0.331994 | 0.390509 | 0.447573 | 0.443193 | 0.402994 | 0.347521 | 0.305104 | 0.275578 | 0.263736 | 0.249445 |
| right posterior | 1.005271 | 0.624422 | 0.411461 | 0.233963 | 0.223092 | 0.279410 | 0.356552 | 0.378648 | 0.299696 | 0.237863 | 0.219258 | 0.215729 | 0.214841 | 0.214665 | 0.216425 | 0.231264 | 0.269693 | 0.316587 | 0.334150 | 0.328964 | 0.272274 | 0.216815 | 0.226554 | 0.249297 | 0.274927 | 0.258176 | 0.229400 | 0.214659 | 0.230441 | 0.297527 | 0.361979 | 0.368555 | 0.296298 | 0.224261 | 0.237727 | 0.475855 | 1.401894 | 2.265546 | 1.287713 | 0.664020 | 0.470903 | 0.438237 | 0.493842 | 0.611923 | 0.685060 | 0.679225 | 0.528455 | 0.351989 | 0.230731 | 0.219153 | 0.251129 | 0.256626 | 0.237763 | 0.215102 | 0.245065 | 0.385306 | 0.593975 | 0.836558 | 0.980109 | 0.877627 | 0.828791 | 0.867828 | 1.069703 | 1.703763 | 3.811216 | 10.872788 | 34.722914 | 116.005946 | 449.605460 | 1679.334474 | 4739.142629 | 11183.988569 | 14531.806510 | 11556.096846 | 5943.377147 | 3099.285724 | 1376.684051 | 529.752866 | 184.959442 | 53.556234 | 14.341154 | 4.809556 | 2.322501 | 1.465322 | 1.236069 | 1.268152 | 1.390042 | 1.571974 | 1.904975 | 2.447547 | 2.981917 | 3.699504 | 4.576937 | 5.450082 | 6.154798 | 6.789877e+00 | 6.880823e+00 | 6.556981e+00 | 5.930808e+00 | 4.944037 | 4.095423 | 3.455183 | 3.118677 | 2.883253 | 2.854860 | 2.853482 | 2.846856 | 2.717214 | 2.625579 | 2.501093 | 2.514630 | 2.688182 | 3.283254 | 4.627454 | 7.413298 | 12.475668 | 23.362018 | 50.486842 | 113.378306 | 216.916907 | 352.882648 | 494.046330 | 532.220976 | 421.037723 | 258.267816 | 141.252764 | 71.731849 | 39.573362 | 25.774305 | 18.473339 | 14.146857 | 12.433974 | 12.386039 | 13.199520 | 15.480644 | 18.856206 | 20.498571 | 21.272811 | 21.230364 | 20.051843 | 18.304507 | 17.493082 | 16.477786 | 17.756893 | 22.364763 | 31.576493 | 41.795600 | 49.952914 | 53.036557 | 49.035034 | 38.829719 | 31.086261 | 26.234164 | 24.414366 | 27.877210 | 39.162030 | 62.657542 | 113.325805 | 176.577391 | 203.799048 | 157.674054 | 92.623189 | 43.984218 | 20.461760 | 10.905587 | 7.381395 | 5.956483 | 6.212744 | 7.211967 | 9.280224 | 13.068541 | 19.047545 | 24.952875 | 38.885610 | 63.327996 | 100.212882 | 161.228928 | 230.584452 | 254.135168 | 209.447801 | 136.298068 | 71.734300 | 40.991430 | 23.587413 | 17.425694 | 17.839350 | 25.363517 | 38.570668 | 64.126084 | 108.273429 | 166.222930 | 221.296738 | 297.030021 | 384.818246 | 395.836847 | 356.907325 | 274.017668 | 212.716199 | 172.975499 | 169.021059 | 202.824216 | 277.928705 | 339.462857 | 350.305942 | 294.914898 | 202.461812 | 127.590513 | 86.552917 | 65.859055 | 53.103511 | 54.110899 | 63.966434 | 88.820911 | 139.877259 | 222.905583 | 344.345045 | 669.925425 | 932.156660 | 1001.835554 | 826.125997 | 410.092169 | 121.363059 | 40.385883 | 17.182208 | 10.367942 | 8.614339 | 9.421309 | 12.942924 | 21.247014 | 33.988125 | 45.223150 | 55.820230 | 62.372020 | 69.821610 | 71.380615 | 66.312997 | 59.645264 | 57.096320 | 48.378801 | 44.157446 | 40.485336 | 37.271809 | 37.435069 | 38.076313 | 30.669806 | 24.172447 | 19.492054 | 17.797456 | 19.607500 | 22.088088 | 20.986157 | 17.435367 | 13.390439 | 9.949723 | 7.963196 | 7.426145 | 8.100921 | 9.862280 | 12.722100 | 17.143227 | 22.164391 | 28.679480 | 38.123583 | 49.738057 | 61.193585 | 58.566283 | 40.467916 | 20.604720 | 9.220505 | 4.074135 | 2.089515 | 1.367132 | 1.128941 | 1.166104 | 1.548591 | 2.469908 | 4.544875 | 8.709738 | 12.172620 | 16.268285 |
| all electrodes | 0.227134 | 0.234877 | 0.236883 | 0.240665 | 0.236045 | 0.237791 | 0.237163 | 0.245472 | 0.257345 | 0.268505 | 0.260386 | 0.292256 | 0.340510 | 0.458889 | 0.728616 | 1.352241 | 2.135898 | 3.544247 | 5.558868 | 7.257362 | 4.939271 | 1.469853 | 0.434430 | 0.238128 | 0.217637 | 0.219280 | 0.224280 | 0.349511 | 0.794957 | 1.633055 | 1.801310 | 1.614065 | 0.912245 | 0.409596 | 0.232315 | 0.220240 | 0.316523 | 0.464285 | 0.419588 | 0.313748 | 0.248001 | 0.214842 | 0.224670 | 0.245449 | 0.254170 | 0.257851 | 0.258248 | 0.276407 | 0.359399 | 0.494014 | 0.594888 | 0.636444 | 0.638325 | 0.465366 | 0.341051 | 0.257623 | 0.223885 | 0.217679 | 0.221234 | 0.233700 | 0.255638 | 0.239922 | 0.214767 | 0.273811 | 0.662856 | 3.265718 | 22.293418 | 132.007335 | 581.454900 | 1548.819628 | 3869.414202 | 6998.473144 | 11588.875093 | 16801.859850 | 21094.331584 | 14162.732683 | 5602.273571 | 1730.983353 | 602.337565 | 194.716742 | 59.748485 | 24.564532 | 11.928224 | 7.359003 | 5.237760 | 4.065945 | 3.917093 | 5.449640 | 7.982610 | 19.978252 | 88.678355 | 340.802856 | 1047.205451 | 3266.991518 | 4945.722736 | 4.931533e+03 | 3.602433e+03 | 1.236567e+03 | 4.699701e+02 | 227.962906 | 121.670842 | 67.059677 | 33.551766 | 13.987253 | 7.196708 | 4.006453 | 2.517801 | 2.183011 | 2.605908 | 3.967476 | 7.448168 | 16.352651 | 35.911526 | 84.455117 | 166.541243 | 225.191432 | 233.241550 | 236.012518 | 236.186559 | 212.236052 | 285.567428 | 479.200886 | 795.622576 | 1115.807662 | 1423.526322 | 1311.914490 | 1312.304470 | 925.555512 | 577.656121 | 292.620452 | 133.976945 | 68.344740 | 46.148298 | 30.652051 | 28.333868 | 36.684999 | 60.941394 | 124.094160 | 306.180917 | 847.226225 | 1753.502466 | 1518.482143 | 645.358755 | 322.667645 | 172.118132 | 102.765313 | 75.528252 | 62.927002 | 62.976531 | 77.505225 | 80.659140 | 74.633927 | 70.393063 | 56.084809 | 43.417144 | 30.879184 | 20.861184 | 12.428125 | 5.843588 | 2.628983 | 1.355390 | 0.835494 | 0.586533 | 0.494342 | 0.406995 | 0.338770 | 0.301879 | 0.278107 | 0.257925 | 0.250686 | 0.246027 | 0.237934 | 0.240491 | 0.243382 | 0.251388 | 0.277546 | 0.338211 | 0.404970 | 0.519063 | 0.661991 | 0.766473 | 0.771708 | 0.743487 | 0.698611 | 0.651332 | 0.699762 | 0.720490 | 0.703154 | 0.665429 | 0.649605 | 0.600726 | 0.632046 | 0.694145 | 0.839772 | 0.977374 | 1.072792 | 1.079835 | 0.986757 | 0.755516 | 0.524368 | 0.378973 | 0.317248 | 0.293464 | 0.282684 | 0.286391 | 0.289471 | 0.305885 | 0.329413 | 0.348228 | 0.373022 | 0.413316 | 0.412355 | 0.389811 | 0.356221 | 0.335717 | 0.334623 | 0.371819 | 0.453924 | 0.602644 | 0.777663 | 0.867740 | 0.705572 | 0.527449 | 0.390882 | 0.302617 | 0.247386 | 0.228104 | 0.218800 | 0.216875 | 0.215732 | 0.214841 | 0.216640 | 0.221528 | 0.227724 | 0.227721 | 0.224479 | 0.218899 | 0.214785 | 0.218975 | 0.234839 | 0.252343 | 0.262681 | 0.255178 | 0.231662 | 0.215543 | 0.216673 | 0.227466 | 0.252610 | 0.283259 | 0.293307 | 0.318918 | 0.331763 | 0.344093 | 0.385933 | 0.467007 | 0.581442 | 0.717439 | 0.709169 | 0.544446 | 0.411456 | 0.294786 | 0.235008 | 0.218034 | 0.219290 | 0.244975 | 0.372524 | 0.848679 | 3.159835 | 9.443194 | 12.995353 | 8.040015 | 3.490434 | 1.675165 | 1.144637 | 0.870425 | 0.752170 | 0.681105 | 0.586256 | 0.508778 | 0.464779 |

O) maximum cross-correlation, neutral vs sad

  
|  | time window | peak latency | cluster *p* | peak Cohen's *d* |  | | | |  |
| **all electrodes** | 125 - 185 ms | 155 ms | 0.0479 | 1.2977 |  | | | |  |
 345 - 465 ms | 390 ms | 0.0453 | 0.6015 |  | | | | 930 - 1080 ms | 965 ms | 0.0198 | 0.5836 |  | | | ||  |  | | | | | | | | |

Model correlations, cluster permutation tests

|  |  | **left hemisphere** | | | | **right hemisphere** | | | |
|  | time window | peak latency | cluster *p* | peak Cohen's *d* | time window | peak latency | cluster *p* | peak Cohen's *d* |  |
| **anterior** |  | | | |  | | | |  |
| **central** | 135 - 245 ms | 160 ms | 0.0442 | 0.6484 | -200 - -135 ms | -200 ms | 0.0454 | -0.7227 |  |
| **posterior** |  | | | |  | | | |  |

  

Model correlations, Bayesian statistics

|  | -200 | -195 | -190 | -185 | -180 | -175 | -170 | -165 | -160 | -155 | -150 | -145 | -140 | -135 | -130 | -125 | -120 | -115 | -110 | -105 | -100 | -95 | -90 | -85 | -80 | -75 | -70 | -65 | -60 | -55 | -50 | -45 | -40 | -35 | -30 | -25 | -20 | -15 | -10 | -5 | 0 | 5 | 10 | 15 | 20 | 25 | 30 | 35 | 40 | 45 | 50 | 55 | 60 | 65 | 70 | 75 | 80 | 85 | 90 | 95 | 100 | 105 | 110 | 115 | 120 | 125 | 130 | 135 | 140 | 145 | 150 | 155 | 160 | 165 | 170 | 175 | 180 | 185 | 190 | 195 | 200 | 205 | 210 | 215 | 220 | 225 | 230 | 235 | 240 | 245 | 250 | 255 | 260 | 265 | 270 | 275 | 280 | 285 | 290 | 295 | 300 | 305 | 310 | 315 | 320 | 325 | 330 | 335 | 340 | 345 | 350 | 355 | 360 | 365 | 370 | 375 | 380 | 385 | 390 | 395 | 400 | 405 | 410 | 415 | 420 | 425 | 430 | 435 | 440 | 445 | 450 | 455 | 460 | 465 | 470 | 475 | 480 | 485 | 490 | 495 | 500 | 505 | 510 | 515 | 520 | 525 | 530 | 535 | 540 | 545 | 550 | 555 | 560 | 565 | 570 | 575 | 580 | 585 | 590 | 595 | 600 | 605 | 610 | 615 | 620 | 625 | 630 | 635 | 640 | 645 | 650 | 655 | 660 | 665 | 670 | 675 | 680 | 685 | 690 | 695 | 700 | 705 | 710 | 715 | 720 | 725 | 730 | 735 | 740 | 745 | 750 | 755 | 760 | 765 | 770 | 775 | 780 | 785 | 790 | 795 | 800 | 805 | 810 | 815 | 820 | 825 | 830 | 835 | 840 | 845 | 850 | 855 | 860 | 865 | 870 | 875 | 880 | 885 | 890 | 895 | 900 | 905 | 910 | 915 | 920 | 925 | 930 | 935 | 940 | 945 | 950 | 955 | 960 | 965 | 970 | 975 | 980 | 985 | 990 | 995 | 1000 | 1005 | 1010 | 1015 | 1020 | 1025 | 1030 | 1035 | 1040 | 1045 | 1050 | 1055 | 1060 | 1065 | 1070 | 1075 | 1080 | 1085 | 1090 | 1095 | 1100 | 1105 | 1110 | 1115 | 1120 | 1125 | 1130 | 1135 | 1140 | 1145 | 1150 | 1155 | 1160 | 1165 | 1170 | 1175 | 1180 | 1185 | 1190 | 1195 |
| --- | --- | --- | --- | --- | --- | --- | --- | --- | --- | --- | --- | --- | --- | --- | --- | --- | --- | --- | --- | --- | --- | --- | --- | --- | --- | --- | --- | --- | --- | --- | --- | --- | --- | --- | --- | --- | --- | --- | --- | --- | --- | --- | --- | --- | --- | --- | --- | --- | --- | --- | --- | --- | --- | --- | --- | --- | --- | --- | --- | --- | --- | --- | --- | --- | --- | --- | --- | --- | --- | --- | --- | --- | --- | --- | --- | --- | --- | --- | --- | --- | --- | --- | --- | --- | --- | --- | --- | --- | --- | --- | --- | --- | --- | --- | --- | --- | --- | --- | --- | --- | --- | --- | --- | --- | --- | --- | --- | --- | --- | --- | --- | --- | --- | --- | --- | --- | --- | --- | --- | --- | --- | --- | --- | --- | --- | --- | --- | --- | --- | --- | --- | --- | --- | --- | --- | --- | --- | --- | --- | --- | --- | --- | --- | --- | --- | --- | --- | --- | --- | --- | --- | --- | --- | --- | --- | --- | --- | --- | --- | --- | --- | --- | --- | --- | --- | --- | --- | --- | --- | --- | --- | --- | --- | --- | --- | --- | --- | --- | --- | --- | --- | --- | --- | --- | --- | --- | --- | --- | --- | --- | --- | --- | --- | --- | --- | --- | --- | --- | --- | --- | --- | --- | --- | --- | --- | --- | --- | --- | --- | --- | --- | --- | --- | --- | --- | --- | --- | --- | --- | --- | --- | --- | --- | --- | --- | --- | --- | --- | --- | --- | --- | --- | --- | --- | --- | --- | --- | --- | --- | --- | --- | --- | --- | --- | --- | --- | --- | --- | --- | --- | --- | --- | --- | --- | --- | --- | --- | --- | --- | --- | --- | --- | --- | --- | --- | --- | --- | --- | --- | --- | --- | --- | --- | --- | --- | --- | --- | --- | --- | --- |
| left anterior | 0.532489 | 0.471854 | 0.456191 | 0.352083 | 0.265766 | 0.236815 | 0.219328 | 0.214819 | 0.226795 | 0.238591 | 0.240621 | 0.229361 | 0.226966 | 0.215837 | 0.216281 | 0.238609 | 0.256036 | 0.253261 | 0.244657 | 0.243862 | 0.237759 | 0.237014 | 0.252432 | 0.290719 | 0.358164 | 0.392853 | 0.430885 | 0.430006 | 0.383875 | 0.275863 | 0.226087 | 0.216665 | 0.256128 | 0.434155 | 0.858877 | 1.275386 | 1.230271 | 1.376126 | 1.379762 | 1.588576 | 2.161843 | 4.764429 | 12.872276 | 37.831884 | 91.706829 | 103.054244 | 79.369758 | 40.664116 | 20.409360 | 9.545529 | 4.774750 | 1.634986 | 0.871334 | 0.519328 | 0.339728 | 0.256722 | 0.229058 | 0.216704 | 0.214860 | 0.215074 | 0.215467 | 0.214787 | 0.214693 | 0.221919 | 0.254052 | 0.329810 | 0.494205 | 0.764611 | 1.109649 | 1.444587 | 1.743467 | 1.676696 | 1.543188 | 1.279756 | 0.958492 | 0.595861 | 0.396567 | 0.281842 | 0.229401 | 0.214778 | 0.218483 | 0.220967 | 0.217948 | 0.214868 | 0.215186 | 0.221499 | 0.228092 | 0.229690 | 0.229263 | 0.232484 | 0.239532 | 0.256330 | 0.282284 | 0.325029 | 0.364504 | 0.369764 | 0.359052 | 0.355739 | 0.326235 | 0.290564 | 0.257940 | 0.237282 | 0.235273 | 0.242993 | 0.255023 | 0.280333 | 0.318502 | 0.337682 | 0.353713 | 0.345358 | 0.330436 | 0.302706 | 0.275193 | 0.253713 | 0.242800 | 0.233505 | 0.232155 | 0.230023 | 0.226706 | 0.222877 | 0.219306 | 0.216421 | 0.215474 | 0.214679 | 0.214728 | 0.214941 | 0.216590 | 0.226229 | 0.255605 | 0.332254 | 0.490517 | 0.721241 | 0.825710 | 0.701295 | 0.480255 | 0.367561 | 0.303206 | 0.265955 | 0.243894 | 0.233347 | 0.227197 | 0.222818 | 0.219424 | 0.216468 | 0.215647 | 0.215764 | 0.216532 | 0.217454 | 0.219045 | 0.217651 | 0.214676 | 0.222449 | 0.243063 | 0.280791 | 0.332594 | 0.402006 | 0.498000 | 0.578666 | 0.643797 | 0.737372 | 0.770214 | 0.809929 | 0.835230 | 0.831693 | 0.828957 | 0.866170 | 0.840635 | 0.874694 | 0.896501 | 0.863149 | 0.759169 | 0.630652 | 0.485814 | 0.386868 | 0.326679 | 0.301847 | 0.303337 | 0.334530 | 0.395563 | 0.504015 | 0.667308 | 0.803732 | 0.848512 | 0.742247 | 0.579554 | 0.418462 | 0.319059 | 0.258923 | 0.232672 | 0.220966 | 0.215321 | 0.214838 | 0.215534 | 0.215009 | 0.214641 | 0.218898 | 0.243881 | 0.314887 | 0.450283 | 0.677501 | 0.973436 | 1.112358 | 0.840521 | 0.481640 | 0.292038 | 0.234013 | 0.221075 | 0.224267 | 0.251117 | 0.330130 | 0.467948 | 0.659018 | 0.831261 | 0.947428 | 1.005453 | 1.037099 | 0.977915 | 0.945671 | 0.924479 | 0.942587 | 0.910327 | 0.861719 | 0.811235 | 0.760388 | 0.719822 | 0.718831 | 0.796445 | 0.911829 | 1.044052 | 0.967906 | 0.767265 | 0.567040 | 0.456032 | 0.377702 | 0.361530 | 0.387838 | 0.473376 | 0.543586 | 0.605703 | 0.629545 | 0.611553 | 0.534426 | 0.452612 | 0.364305 | 0.308940 | 0.263320 | 0.232676 | 0.222184 | 0.219152 | 0.218536 | 0.220924 | 0.222439 | 0.221251 | 0.218029 | 0.214915 | 0.216426 | 0.224474 | 0.239174 | 0.250278 | 0.255478 | 0.255538 | 0.246281 | 0.231257 | 0.221689 | 0.217205 | 0.215370 | 0.214885 | 0.214629 | 0.215028 | 0.216592 | 0.220321 | 0.225351 | 0.228248 | 0.230414 | 0.230601 | 0.230081 | 0.235513 | 0.251347 | 0.257316 | 0.270725 |
| right anterior | 0.230149 | 0.223678 | 0.215755 | 0.216199 | 0.217801 | 0.216596 | 0.214718 | 0.218924 | 0.250439 | 0.278579 | 0.265886 | 0.241927 | 0.229458 | 0.215260 | 0.223372 | 0.271071 | 0.385970 | 0.468923 | 0.451977 | 0.381913 | 0.298276 | 0.239253 | 0.215466 | 0.224017 | 0.261779 | 0.317225 | 0.344610 | 0.298989 | 0.246181 | 0.216388 | 0.224843 | 0.286182 | 0.417693 | 0.531698 | 0.431380 | 0.297190 | 0.232788 | 0.218681 | 0.214801 | 0.214682 | 0.215056 | 0.216838 | 0.216993 | 0.218374 | 0.218075 | 0.217328 | 0.218922 | 0.216996 | 0.214644 | 0.222585 | 0.261820 | 0.363284 | 0.558158 | 0.935132 | 1.555977 | 2.420793 | 3.923247 | 4.458072 | 3.769609 | 2.137526 | 1.055501 | 0.606739 | 0.485196 | 0.508197 | 0.763410 | 1.584389 | 3.815252 | 7.009810 | 10.143685 | 12.385677 | 12.984648 | 10.959527 | 7.610826 | 4.838845 | 3.375953 | 2.670207 | 2.269139 | 2.056969 | 1.821226 | 1.385517 | 0.964379 | 0.750045 | 0.690685 | 0.752859 | 0.950337 | 1.286153 | 1.785220 | 2.407529 | 2.791666 | 3.379735 | 3.916097 | 3.907087 | 3.993503 | 4.450108 | 4.850335 | 5.272229 | 6.268843 | 7.210693 | 7.418871 | 4.898403 | 2.951697 | 1.825106 | 1.169919 | 0.781580 | 0.706872 | 0.764878 | 0.974273 | 1.263741 | 1.574639 | 1.634220 | 1.566411 | 1.326122 | 1.072087 | 0.860533 | 0.701056 | 0.502256 | 0.364345 | 0.275968 | 0.233270 | 0.217092 | 0.214750 | 0.218367 | 0.225186 | 0.241043 | 0.267148 | 0.317652 | 0.409781 | 0.536683 | 0.616420 | 0.593442 | 0.510714 | 0.435198 | 0.357298 | 0.298210 | 0.266388 | 0.258366 | 0.264351 | 0.287574 | 0.337584 | 0.441446 | 0.610951 | 0.842853 | 1.166707 | 1.425467 | 1.404762 | 1.235667 | 1.040398 | 0.720907 | 0.498258 | 0.349363 | 0.269954 | 0.232451 | 0.217260 | 0.215288 | 0.222980 | 0.236960 | 0.244499 | 0.237751 | 0.224259 | 0.214643 | 0.230955 | 0.266848 | 0.320429 | 0.359671 | 0.352353 | 0.339653 | 0.316224 | 0.287290 | 0.274674 | 0.271896 | 0.275904 | 0.286487 | 0.288816 | 0.284743 | 0.281046 | 0.269938 | 0.257610 | 0.247128 | 0.245386 | 0.257468 | 0.275744 | 0.305853 | 0.354304 | 0.406684 | 0.456480 | 0.486298 | 0.517294 | 0.592229 | 0.669444 | 0.681205 | 0.700219 | 0.663027 | 0.579217 | 0.517962 | 0.439624 | 0.348730 | 0.299811 | 0.271628 | 0.261972 | 0.275620 | 0.316840 | 0.361507 | 0.414677 | 0.424440 | 0.398746 | 0.333378 | 0.280323 | 0.241766 | 0.221850 | 0.214652 | 0.217048 | 0.222393 | 0.231263 | 0.247031 | 0.265304 | 0.271826 | 0.277304 | 0.280505 | 0.276809 | 0.261776 | 0.261721 | 0.260121 | 0.256819 | 0.252011 | 0.245078 | 0.230465 | 0.220080 | 0.214639 | 0.218059 | 0.216819 | 0.214630 | 0.217095 | 0.226222 | 0.236571 | 0.224714 | 0.216375 | 0.215672 | 0.225056 | 0.241153 | 0.234680 | 0.220861 | 0.215771 | 0.243593 | 0.296599 | 0.325597 | 0.300190 | 0.254582 | 0.225003 | 0.214977 | 0.217720 | 0.222190 | 0.221367 | 0.216587 | 0.214688 | 0.216246 | 0.222107 | 0.231571 | 0.235520 | 0.245289 | 0.249020 | 0.248707 | 0.247763 | 0.252332 | 0.250951 | 0.248720 | 0.246790 | 0.243317 | 0.235212 | 0.225475 | 0.218592 | 0.215785 | 0.214674 | 0.217472 | 0.227613 | 0.243763 | 0.265519 | 0.281989 | 0.293308 | 0.293496 | 0.280644 |
| left central | 1.002566 | 0.997144 | 1.232611 | 1.405245 | 1.443155 | 1.090108 | 0.792141 | 0.468211 | 0.320256 | 0.234013 | 0.214749 | 0.228654 | 0.276831 | 0.331443 | 0.305758 | 0.258517 | 0.226076 | 0.214665 | 0.243552 | 0.294954 | 0.325686 | 0.291869 | 0.274229 | 0.251505 | 0.245575 | 0.230235 | 0.224969 | 0.220394 | 0.215593 | 0.217832 | 0.228338 | 0.246565 | 0.259930 | 0.275136 | 0.261377 | 0.246413 | 0.237544 | 0.234169 | 0.227891 | 0.229858 | 0.240960 | 0.263799 | 0.292472 | 0.341322 | 0.390429 | 0.416464 | 0.372000 | 0.309592 | 0.254600 | 0.222291 | 0.214724 | 0.222293 | 0.236278 | 0.263749 | 0.292231 | 0.337372 | 0.382859 | 0.410045 | 0.418964 | 0.423902 | 0.411968 | 0.403770 | 0.415457 | 0.429269 | 0.468277 | 0.577325 | 0.800520 | 1.296248 | 2.261159 | 3.656119 | 5.988308 | 10.258823 | 11.901867 | 10.581031 | 7.740228 | 4.698697 | 2.849206 | 1.965739 | 1.531955 | 1.537805 | 1.873069 | 2.241457 | 2.762056 | 3.726895 | 4.294305 | 4.217336 | 3.211524 | 2.272310 | 1.765957 | 1.293889 | 1.039642 | 0.870495 | 0.717150 | 0.603043 | 0.536644 | 0.469852 | 0.431944 | 0.385648 | 0.334902 | 0.297941 | 0.280059 | 0.283735 | 0.308290 | 0.343957 | 0.363717 | 0.387232 | 0.384545 | 0.343305 | 0.266444 | 0.227890 | 0.214652 | 0.223321 | 0.239056 | 0.241507 | 0.235971 | 0.224212 | 0.221446 | 0.220550 | 0.217644 | 0.214958 | 0.214767 | 0.215661 | 0.218886 | 0.226730 | 0.237970 | 0.245084 | 0.243671 | 0.230843 | 0.216815 | 0.214903 | 0.222283 | 0.234091 | 0.240178 | 0.237582 | 0.237492 | 0.239262 | 0.257306 | 0.268439 | 0.268821 | 0.275857 | 0.278207 | 0.261318 | 0.252428 | 0.241776 | 0.232855 | 0.227769 | 0.225659 | 0.218132 | 0.214692 | 0.221772 | 0.235561 | 0.248808 | 0.248949 | 0.242226 | 0.239125 | 0.236038 | 0.232860 | 0.221108 | 0.214744 | 0.220969 | 0.256341 | 0.361766 | 0.557593 | 0.892663 | 1.195760 | 1.379239 | 1.400486 | 1.213882 | 0.907286 | 0.676449 | 0.477204 | 0.309318 | 0.228739 | 0.222028 | 0.312809 | 0.501505 | 0.626732 | 0.701169 | 0.626282 | 0.604118 | 0.556937 | 0.455865 | 0.373418 | 0.337300 | 0.283429 | 0.266122 | 0.257612 | 0.247985 | 0.245570 | 0.259027 | 0.274899 | 0.336309 | 0.497371 | 0.803869 | 1.267967 | 1.608991 | 1.359816 | 0.923113 | 0.618639 | 0.434270 | 0.362745 | 0.347697 | 0.370303 | 0.449853 | 0.650489 | 1.004781 | 1.492183 | 1.800147 | 1.599512 | 1.177289 | 0.820395 | 0.633472 | 0.545455 | 0.484536 | 0.485618 | 0.533838 | 0.543358 | 0.575890 | 0.691378 | 0.710485 | 0.568153 | 0.389702 | 0.268102 | 0.224730 | 0.214797 | 0.218508 | 0.222887 | 0.219705 | 0.216661 | 0.214706 | 0.214816 | 0.214633 | 0.215055 | 0.219905 | 0.236840 | 0.237264 | 0.227095 | 0.220770 | 0.218725 | 0.216954 | 0.216220 | 0.216207 | 0.217583 | 0.219318 | 0.223453 | 0.235808 | 0.261928 | 0.304755 | 0.341322 | 0.350154 | 0.306751 | 0.249858 | 0.217133 | 0.223411 | 0.266220 | 0.323833 | 0.372174 | 0.394014 | 0.394719 | 0.398987 | 0.391526 | 0.385746 | 0.378125 | 0.397950 | 0.430197 | 0.470133 | 0.480797 | 0.490407 | 0.431611 | 0.354734 | 0.288994 | 0.252339 | 0.233470 | 0.232418 | 0.231330 | 0.235399 | 0.245806 | 0.259198 | 0.258008 | 0.264646 |
| right central | 17.539824 | 21.553039 | 27.729322 | 42.824607 | 66.335904 | 78.666429 | 81.462897 | 154.604626 | 704.805064 | 948.407433 | 396.893239 | 58.322449 | 9.165988 | 1.821662 | 0.594043 | 0.332857 | 0.257970 | 0.220188 | 0.214872 | 0.223695 | 0.237102 | 0.243210 | 0.234714 | 0.221801 | 0.218937 | 0.268483 | 0.429525 | 0.838382 | 1.286727 | 0.788138 | 0.336600 | 0.215228 | 0.244805 | 0.277088 | 0.265528 | 0.258803 | 0.236842 | 0.221402 | 0.214628 | 0.220268 | 0.242258 | 0.252164 | 0.255728 | 0.242408 | 0.223971 | 0.214740 | 0.219729 | 0.252068 | 0.304352 | 0.370893 | 0.442082 | 0.473636 | 0.422624 | 0.346040 | 0.262559 | 0.218672 | 0.215891 | 0.221505 | 0.223496 | 0.216283 | 0.214977 | 0.217629 | 0.221197 | 0.222086 | 0.223320 | 0.248240 | 0.331038 | 0.599993 | 1.667435 | 5.012064 | 10.092107 | 15.038745 | 16.070655 | 11.179625 | 6.434677 | 3.239315 | 1.621000 | 0.887035 | 0.525229 | 0.328156 | 0.261773 | 0.232856 | 0.231809 | 0.244731 | 0.268663 | 0.284280 | 0.304373 | 0.302474 | 0.305308 | 0.302167 | 0.299489 | 0.294872 | 0.296377 | 0.289735 | 0.294345 | 0.306078 | 0.315060 | 0.317019 | 0.331864 | 0.356840 | 0.417559 | 0.533691 | 0.606171 | 0.580035 | 0.426068 | 0.293148 | 0.224397 | 0.217073 | 0.251247 | 0.275083 | 0.277099 | 0.258310 | 0.244898 | 0.231024 | 0.226183 | 0.226396 | 0.230477 | 0.228655 | 0.221841 | 0.215631 | 0.216901 | 0.233059 | 0.252500 | 0.261932 | 0.268629 | 0.262797 | 0.240813 | 0.224211 | 0.218899 | 0.218595 | 0.219585 | 0.222382 | 0.225172 | 0.226483 | 0.224141 | 0.220569 | 0.217515 | 0.216644 | 0.215071 | 0.214983 | 0.218307 | 0.225985 | 0.245174 | 0.274032 | 0.318688 | 0.385334 | 0.438209 | 0.442482 | 0.392594 | 0.327970 | 0.272398 | 0.242692 | 0.233054 | 0.231019 | 0.237730 | 0.256513 | 0.270431 | 0.266768 | 0.249482 | 0.225902 | 0.214913 | 0.217867 | 0.226998 | 0.229210 | 0.230640 | 0.233245 | 0.235139 | 0.232075 | 0.229532 | 0.220087 | 0.214968 | 0.241710 | 0.319193 | 0.409953 | 0.477191 | 0.484932 | 0.410716 | 0.349596 | 0.326328 | 0.307898 | 0.301169 | 0.301559 | 0.298589 | 0.295243 | 0.283511 | 0.260806 | 0.240702 | 0.221903 | 0.214793 | 0.228713 | 0.257477 | 0.307095 | 0.349646 | 0.347014 | 0.310543 | 0.271621 | 0.237968 | 0.220011 | 0.214635 | 0.218529 | 0.222653 | 0.229476 | 0.242931 | 0.250468 | 0.244275 | 0.246586 | 0.245750 | 0.252136 | 0.265225 | 0.279328 | 0.274985 | 0.267229 | 0.244628 | 0.229007 | 0.224160 | 0.220785 | 0.219037 | 0.222928 | 0.229000 | 0.235483 | 0.243644 | 0.245156 | 0.233340 | 0.218064 | 0.216270 | 0.231506 | 0.251785 | 0.269744 | 0.279548 | 0.279880 | 0.273706 | 0.267647 | 0.261548 | 0.250058 | 0.231396 | 0.217997 | 0.214784 | 0.223905 | 0.253983 | 0.297070 | 0.341718 | 0.380232 | 0.410501 | 0.431836 | 0.455884 | 0.468939 | 0.515055 | 0.555391 | 0.606718 | 0.556424 | 0.553600 | 0.588342 | 0.640779 | 0.613043 | 0.521677 | 0.371348 | 0.264634 | 0.215925 | 0.240620 | 0.313082 | 0.384148 | 0.437779 | 0.369903 | 0.281036 | 0.226872 | 0.215575 | 0.239444 | 0.265524 | 0.292711 | 0.310008 | 0.327828 | 0.348076 | 0.379457 | 0.424293 | 0.481238 | 0.541694 | 0.582803 | 0.613216 | 0.630635 | 0.611403 |
| left posterior | 0.797751 | 0.995053 | 1.281632 | 2.100884 | 1.855980 | 1.414936 | 0.826587 | 0.512866 | 0.333536 | 0.235265 | 0.214639 | 0.228481 | 0.259181 | 0.274576 | 0.277167 | 0.265446 | 0.237708 | 0.219992 | 0.214848 | 0.214626 | 0.215911 | 0.222009 | 0.228941 | 0.226341 | 0.215350 | 0.217261 | 0.234033 | 0.314351 | 0.520699 | 1.091648 | 2.616222 | 4.301674 | 5.191610 | 3.676500 | 1.252898 | 0.504647 | 0.282809 | 0.216986 | 0.224862 | 0.235900 | 0.227132 | 0.215747 | 0.215555 | 0.219485 | 0.223492 | 0.225215 | 0.224545 | 0.226021 | 0.236359 | 0.266911 | 0.330856 | 0.492644 | 0.793454 | 1.336870 | 1.882145 | 2.811139 | 4.192801 | 5.329678 | 4.871114 | 3.931134 | 2.431948 | 1.165299 | 0.374815 | 0.215318 | 0.296369 | 0.618961 | 1.573414 | 4.210880 | 11.028410 | 26.142244 | 57.033731 | 95.177306 | 123.060136 | 109.111154 | 73.954034 | 44.490209 | 20.689530 | 7.985182 | 3.187911 | 1.401942 | 0.726650 | 0.467776 | 0.395923 | 0.423244 | 0.511925 | 0.625433 | 0.780439 | 0.872099 | 0.845571 | 0.810452 | 0.794333 | 0.772888 | 0.786903 | 0.806774 | 0.825395 | 0.809578 | 0.748911 | 0.677397 | 0.615291 | 0.574531 | 0.587947 | 0.654385 | 0.823925 | 1.150860 | 1.644851 | 2.198411 | 2.717795 | 3.005595 | 3.189926 | 3.363181 | 3.796185 | 4.480097 | 4.961347 | 5.446252 | 5.817943 | 6.391317 | 7.348950 | 8.998878 | 11.137163 | 13.297299 | 14.616009 | 16.071376 | 17.200633 | 15.600945 | 12.724798 | 8.266922 | 4.428020 | 2.275351 | 1.271616 | 0.767865 | 0.561760 | 0.457604 | 0.428334 | 0.410032 | 0.401134 | 0.389847 | 0.378988 | 0.354348 | 0.331961 | 0.312017 | 0.293480 | 0.277531 | 0.269818 | 0.268137 | 0.271531 | 0.272609 | 0.265585 | 0.255903 | 0.247491 | 0.240448 | 0.235093 | 0.230738 | 0.226593 | 0.223692 | 0.220888 | 0.222130 | 0.228195 | 0.243394 | 0.271411 | 0.325774 | 0.421457 | 0.593561 | 0.834827 | 1.079345 | 1.186947 | 1.063352 | 0.774186 | 0.529353 | 0.371416 | 0.279030 | 0.230187 | 0.214742 | 0.222956 | 0.241679 | 0.245642 | 0.232233 | 0.216305 | 0.223333 | 0.277314 | 0.374775 | 0.506878 | 0.608857 | 0.608280 | 0.559580 | 0.472247 | 0.381000 | 0.325735 | 0.295198 | 0.270985 | 0.257574 | 0.256301 | 0.252748 | 0.249029 | 0.245705 | 0.243896 | 0.241080 | 0.249632 | 0.271034 | 0.318595 | 0.403906 | 0.566017 | 0.781281 | 1.054787 | 1.244280 | 1.377806 | 1.478190 | 1.507441 | 1.390808 | 1.314578 | 1.291816 | 1.256086 | 1.257487 | 1.262419 | 1.256072 | 1.121362 | 0.965585 | 0.825985 | 0.709495 | 0.615932 | 0.584645 | 0.570234 | 0.584786 | 0.673504 | 0.836009 | 1.038706 | 1.275073 | 1.429365 | 1.435736 | 1.343966 | 1.241032 | 1.159712 | 1.222354 | 1.444135 | 1.955939 | 2.876998 | 4.518937 | 7.073961 | 9.514452 | 9.442244 | 7.581582 | 5.285000 | 3.341158 | 2.052859 | 1.352910 | 0.921062 | 0.660262 | 0.547976 | 0.497932 | 0.459891 | 0.435711 | 0.410684 | 0.381419 | 0.354468 | 0.335875 | 0.328610 | 0.328459 | 0.316323 | 0.308218 | 0.293884 | 0.275470 | 0.258444 | 0.245930 | 0.235565 | 0.231227 | 0.229684 | 0.233457 | 0.243756 | 0.263958 | 0.287926 | 0.302904 | 0.296655 | 0.286555 | 0.259803 | 0.236984 | 0.222706 | 0.217276 | 0.215495 | 0.215353 | 0.215022 | 0.215888 |
| right posterior | 0.352526 | 0.427445 | 0.616126 | 0.929596 | 1.225530 | 1.007692 | 0.737888 | 0.414873 | 0.299074 | 0.233038 | 0.217977 | 0.214626 | 0.215849 | 0.222519 | 0.222418 | 0.225896 | 0.224500 | 0.243664 | 0.312804 | 0.373752 | 0.329199 | 0.293465 | 0.238337 | 0.215013 | 0.264815 | 0.466810 | 0.969390 | 1.736455 | 2.212977 | 1.594049 | 0.718092 | 0.389802 | 0.336447 | 0.362076 | 0.386067 | 0.380085 | 0.377532 | 0.366620 | 0.391530 | 0.468109 | 0.589239 | 0.778889 | 1.022696 | 1.089577 | 0.906099 | 0.723520 | 0.601497 | 0.516481 | 0.522797 | 0.710681 | 1.097516 | 1.518841 | 1.598092 | 1.711777 | 1.309317 | 0.943846 | 0.643724 | 0.429435 | 0.316751 | 0.278910 | 0.244460 | 0.219560 | 0.220578 | 0.302852 | 0.584969 | 1.274355 | 2.940554 | 6.065361 | 10.517601 | 17.688822 | 26.021202 | 28.472256 | 22.126263 | 13.888073 | 7.743833 | 4.177963 | 2.093732 | 1.001226 | 0.516744 | 0.308470 | 0.229518 | 0.214759 | 0.214917 | 0.215498 | 0.226151 | 0.264543 | 0.335326 | 0.487909 | 0.789216 | 1.392957 | 2.587546 | 4.940964 | 7.098947 | 8.798732 | 9.398363 | 8.839379 | 8.051803 | 7.287657 | 5.949460 | 5.325771 | 4.676975 | 4.144822 | 3.382503 | 2.819667 | 2.262492 | 1.867019 | 1.514133 | 1.344760 | 1.218426 | 1.163201 | 1.144338 | 1.228057 | 1.384872 | 1.674069 | 2.016259 | 2.554431 | 3.119182 | 3.589707 | 4.343243 | 5.578230 | 7.006334 | 8.777367 | 10.359550 | 11.493988 | 12.445570 | 12.524837 | 10.554290 | 7.779378 | 4.949369 | 2.666295 | 1.455475 | 0.913274 | 0.688998 | 0.626125 | 0.677856 | 0.801254 | 0.938897 | 1.091270 | 1.109192 | 1.068717 | 0.982613 | 0.853152 | 0.784187 | 0.837136 | 0.939218 | 0.990206 | 1.001135 | 0.878692 | 0.675968 | 0.480931 | 0.362649 | 0.301070 | 0.273820 | 0.267483 | 0.278188 | 0.305931 | 0.365030 | 0.484763 | 0.711276 | 1.130626 | 1.796925 | 2.626767 | 3.245925 | 3.270165 | 2.574672 | 1.664841 | 0.907853 | 0.522773 | 0.350796 | 0.279751 | 0.250044 | 0.241543 | 0.241525 | 0.253060 | 0.276479 | 0.320924 | 0.385588 | 0.457554 | 0.505073 | 0.509358 | 0.458750 | 0.387076 | 0.328977 | 0.301213 | 0.291757 | 0.295625 | 0.318857 | 0.354747 | 0.386621 | 0.392529 | 0.386848 | 0.368906 | 0.362828 | 0.366095 | 0.393809 | 0.447684 | 0.516706 | 0.552900 | 0.555799 | 0.549470 | 0.589700 | 0.685360 | 0.851386 | 1.063204 | 1.174142 | 1.123680 | 0.952558 | 0.734521 | 0.566321 | 0.464643 | 0.401162 | 0.368347 | 0.357011 | 0.368655 | 0.405540 | 0.441613 | 0.478593 | 0.513353 | 0.516955 | 0.484536 | 0.425784 | 0.370895 | 0.330243 | 0.305636 | 0.303181 | 0.327537 | 0.376342 | 0.451487 | 0.567242 | 0.687228 | 0.794741 | 0.933809 | 1.055159 | 1.048041 | 0.959005 | 0.909283 | 0.841328 | 0.722521 | 0.573418 | 0.443460 | 0.326460 | 0.256444 | 0.228431 | 0.220285 | 0.219050 | 0.222906 | 0.235038 | 0.271455 | 0.336864 | 0.421380 | 0.531432 | 0.635583 | 0.650696 | 0.602751 | 0.531448 | 0.462019 | 0.414646 | 0.388818 | 0.380207 | 0.387945 | 0.393063 | 0.380724 | 0.324834 | 0.268143 | 0.231248 | 0.215703 | 0.217511 | 0.222307 | 0.220124 | 0.215773 | 0.215740 | 0.228203 | 0.248632 | 0.270054 | 0.310019 | 0.352910 | 0.420289 | 0.510874 | 0.590747 | 0.642656 |
| all electrodes | 0.216751 | 0.217642 | 0.215481 | 0.215350 | 0.223015 | 0.229243 | 0.268054 | 0.310870 | 0.411876 | 0.632261 | 0.996692 | 1.043184 | 1.413880 | 1.362836 | 1.144596 | 0.775135 | 0.491196 | 0.288188 | 0.229472 | 0.216049 | 0.214703 | 0.215561 | 0.215409 | 0.214627 | 0.218341 | 0.240551 | 0.349065 | 0.554641 | 0.634111 | 0.691362 | 0.639941 | 0.452828 | 0.354207 | 0.331582 | 0.310799 | 0.335346 | 0.329025 | 0.262353 | 0.220008 | 0.225737 | 0.333009 | 0.589945 | 1.030069 | 1.502460 | 1.260793 | 0.756549 | 0.375803 | 0.245657 | 0.220212 | 0.214626 | 0.216050 | 0.220871 | 0.227854 | 0.239058 | 0.233530 | 0.227172 | 0.218918 | 0.215346 | 0.215354 | 0.221029 | 0.223963 | 0.217461 | 0.220714 | 0.314646 | 0.844757 | 4.591207 | 43.260336 | 336.166194 | 1753.487902 | 5557.432726 | 10464.086226 | 12071.503618 | 7223.250532 | 1733.675337 | 254.186267 | 42.013060 | 7.314427 | 1.587735 | 0.602635 | 0.344432 | 0.270398 | 0.257101 | 0.269981 | 0.292165 | 0.326695 | 0.328789 | 0.305531 | 0.268478 | 0.250498 | 0.242785 | 0.253943 | 0.291219 | 0.405554 | 0.692231 | 1.389402 | 2.489827 | 3.805820 | 3.795210 | 2.715294 | 1.662399 | 1.118883 | 0.801251 | 0.672551 | 0.616028 | 0.662047 | 0.800418 | 0.944166 | 0.970276 | 1.009164 | 1.105586 | 1.354737 | 1.910177 | 2.878092 | 4.379713 | 7.387163 | 8.898666 | 8.558696 | 7.839190 | 6.193532 | 4.094952 | 3.141968 | 2.419613 | 2.001587 | 1.643096 | 1.323242 | 1.201138 | 1.418923 | 1.632747 | 1.935454 | 2.283858 | 2.225878 | 1.725325 | 1.361362 | 1.114258 | 0.997283 | 1.037697 | 1.039509 | 1.042107 | 1.026038 | 0.949079 | 0.768900 | 0.681785 | 0.638392 | 0.627359 | 0.584916 | 0.557220 | 0.510293 | 0.462585 | 0.431483 | 0.438914 | 0.488472 | 0.670793 | 1.108534 | 1.822058 | 2.628601 | 3.135326 | 2.502938 | 1.757324 | 1.196024 | 0.785485 | 0.540844 | 0.404373 | 0.312125 | 0.262884 | 0.238060 | 0.218242 | 0.215878 | 0.233206 | 0.258318 | 0.291793 | 0.295213 | 0.287401 | 0.259802 | 0.248003 | 0.230238 | 0.218959 | 0.215022 | 0.225201 | 0.249641 | 0.277462 | 0.318447 | 0.378317 | 0.404213 | 0.404668 | 0.392337 | 0.390197 | 0.366217 | 0.337308 | 0.327047 | 0.327318 | 0.332018 | 0.365864 | 0.425751 | 0.461753 | 0.473545 | 0.440546 | 0.363498 | 0.326894 | 0.335471 | 0.355671 | 0.361405 | 0.379784 | 0.378371 | 0.333800 | 0.287744 | 0.259820 | 0.242901 | 0.245993 | 0.275316 | 0.338994 | 0.482991 | 0.728176 | 0.932267 | 0.976330 | 0.790921 | 0.576783 | 0.418712 | 0.361325 | 0.340811 | 0.368643 | 0.403625 | 0.444431 | 0.466275 | 0.549336 | 0.689795 | 0.933471 | 1.300493 | 1.843287 | 2.642668 | 4.053212 | 5.566564 | 6.389408 | 5.119705 | 3.877532 | 3.670612 | 3.839888 | 3.516776 | 3.441824 | 3.150115 | 2.922583 | 2.752987 | 2.937231 | 3.411823 | 3.684748 | 2.224238 | 1.682361 | 1.646507 | 1.730766 | 2.012521 | 3.152558 | 4.229842 | 6.326669 | 8.779728 | 10.419761 | 9.449679 | 6.556084 | 2.403809 | 1.045940 | 0.579820 | 0.400317 | 0.325005 | 0.303447 | 0.298625 | 0.312584 | 0.349338 | 0.410214 | 0.488299 | 0.573460 | 0.682156 | 0.808763 | 1.084510 | 1.558976 | 2.224109 | 2.587973 | 2.834913 | 2.393496 | 1.716624 | 0.989143 | 0.831513 | 0.627484 |

P) maximum cross-correlation, happy vs angry

  
|  | time window | peak latency | cluster *p* | peak Cohen's *d* |  | | | |
| **all electrodes** | 370 - 475 ms | 405 ms | 0.0359 | 0.7266 |  | | | |
|  | | | | | | | | |

Model correlations, cluster permutation tests

|  | **left hemisphere** | | | | **right hemisphere** | | | |
|  | time window | peak latency | cluster *p* | peak Cohen's *d* | time window | peak latency | cluster *p* | peak Cohen's *d* |
| **anterior** |  | | | |  | | | |
| **central** |  | | | |  | | | |
| **posterior** | 130 - 210 ms | 170 ms | 0.0342 | 1.1056 | 830 - 1065 ms | 895 ms | 0.017 | 0.535 |
 965 - 1075 ms | 1050 ms | 0.0453 | 0.6248 |  | | | |

  

Model correlations, Bayesian statistics

|  | -200 | -195 | -190 | -185 | -180 | -175 | -170 | -165 | -160 | -155 | -150 | -145 | -140 | -135 | -130 | -125 | -120 | -115 | -110 | -105 | -100 | -95 | -90 | -85 | -80 | -75 | -70 | -65 | -60 | -55 | -50 | -45 | -40 | -35 | -30 | -25 | -20 | -15 | -10 | -5 | 0 | 5 | 10 | 15 | 20 | 25 | 30 | 35 | 40 | 45 | 50 | 55 | 60 | 65 | 70 | 75 | 80 | 85 | 90 | 95 | 100 | 105 | 110 | 115 | 120 | 125 | 130 | 135 | 140 | 145 | 150 | 155 | 160 | 165 | 170 | 175 | 180 | 185 | 190 | 195 | 200 | 205 | 210 | 215 | 220 | 225 | 230 | 235 | 240 | 245 | 250 | 255 | 260 | 265 | 270 | 275 | 280 | 285 | 290 | 295 | 300 | 305 | 310 | 315 | 320 | 325 | 330 | 335 | 340 | 345 | 350 | 355 | 360 | 365 | 370 | 375 | 380 | 385 | 390 | 395 | 400 | 405 | 410 | 415 | 420 | 425 | 430 | 435 | 440 | 445 | 450 | 455 | 460 | 465 | 470 | 475 | 480 | 485 | 490 | 495 | 500 | 505 | 510 | 515 | 520 | 525 | 530 | 535 | 540 | 545 | 550 | 555 | 560 | 565 | 570 | 575 | 580 | 585 | 590 | 595 | 600 | 605 | 610 | 615 | 620 | 625 | 630 | 635 | 640 | 645 | 650 | 655 | 660 | 665 | 670 | 675 | 680 | 685 | 690 | 695 | 700 | 705 | 710 | 715 | 720 | 725 | 730 | 735 | 740 | 745 | 750 | 755 | 760 | 765 | 770 | 775 | 780 | 785 | 790 | 795 | 800 | 805 | 810 | 815 | 820 | 825 | 830 | 835 | 840 | 845 | 850 | 855 | 860 | 865 | 870 | 875 | 880 | 885 | 890 | 895 | 900 | 905 | 910 | 915 | 920 | 925 | 930 | 935 | 940 | 945 | 950 | 955 | 960 | 965 | 970 | 975 | 980 | 985 | 990 | 995 | 1000 | 1005 | 1010 | 1015 | 1020 | 1025 | 1030 | 1035 | 1040 | 1045 | 1050 | 1055 | 1060 | 1065 | 1070 | 1075 | 1080 | 1085 | 1090 | 1095 | 1100 | 1105 | 1110 | 1115 | 1120 | 1125 | 1130 | 1135 | 1140 | 1145 | 1150 | 1155 | 1160 | 1165 | 1170 | 1175 | 1180 | 1185 | 1190 | 1195 |
| --- | --- | --- | --- | --- | --- | --- | --- | --- | --- | --- | --- | --- | --- | --- | --- | --- | --- | --- | --- | --- | --- | --- | --- | --- | --- | --- | --- | --- | --- | --- | --- | --- | --- | --- | --- | --- | --- | --- | --- | --- | --- | --- | --- | --- | --- | --- | --- | --- | --- | --- | --- | --- | --- | --- | --- | --- | --- | --- | --- | --- | --- | --- | --- | --- | --- | --- | --- | --- | --- | --- | --- | --- | --- | --- | --- | --- | --- | --- | --- | --- | --- | --- | --- | --- | --- | --- | --- | --- | --- | --- | --- | --- | --- | --- | --- | --- | --- | --- | --- | --- | --- | --- | --- | --- | --- | --- | --- | --- | --- | --- | --- | --- | --- | --- | --- | --- | --- | --- | --- | --- | --- | --- | --- | --- | --- | --- | --- | --- | --- | --- | --- | --- | --- | --- | --- | --- | --- | --- | --- | --- | --- | --- | --- | --- | --- | --- | --- | --- | --- | --- | --- | --- | --- | --- | --- | --- | --- | --- | --- | --- | --- | --- | --- | --- | --- | --- | --- | --- | --- | --- | --- | --- | --- | --- | --- | --- | --- | --- | --- | --- | --- | --- | --- | --- | --- | --- | --- | --- | --- | --- | --- | --- | --- | --- | --- | --- | --- | --- | --- | --- | --- | --- | --- | --- | --- | --- | --- | --- | --- | --- | --- | --- | --- | --- | --- | --- | --- | --- | --- | --- | --- | --- | --- | --- | --- | --- | --- | --- | --- | --- | --- | --- | --- | --- | --- | --- | --- | --- | --- | --- | --- | --- | --- | --- | --- | --- | --- | --- | --- | --- | --- | --- | --- | --- | --- | --- | --- | --- | --- | --- | --- | --- | --- | --- | --- | --- | --- | --- | --- | --- | --- | --- | --- | --- | --- | --- | --- | --- | --- | --- |
| left anterior | 0.444690 | 0.586261 | 0.712174 | 1.352468 | 1.781044 | 1.427597 | 0.694466 | 0.383041 | 0.274702 | 0.239085 | 0.223624 | 0.248413 | 0.373249 | 0.923227 | 2.106164 | 2.060227 | 1.543647 | 1.253746 | 0.925293 | 0.790627 | 0.634241 | 0.474002 | 0.350937 | 0.262456 | 0.215668 | 0.223475 | 0.255243 | 0.291459 | 0.302170 | 0.277261 | 0.237417 | 0.214980 | 0.233328 | 0.330710 | 0.684489 | 1.991996 | 5.511789 | 5.354569 | 2.390034 | 0.796604 | 0.370703 | 0.248838 | 0.226013 | 0.219335 | 0.230084 | 0.270273 | 0.373619 | 0.370606 | 0.394425 | 0.363946 | 0.377108 | 0.465048 | 0.884510 | 1.598611 | 2.942832 | 3.946352 | 3.497231 | 1.904922 | 1.131239 | 0.678263 | 0.417671 | 0.312398 | 0.260074 | 0.229112 | 0.215555 | 0.216173 | 0.225035 | 0.238536 | 0.276567 | 0.312837 | 0.362082 | 0.414499 | 0.497687 | 0.514828 | 0.464643 | 0.384289 | 0.307557 | 0.249907 | 0.217498 | 0.230270 | 0.388972 | 1.070796 | 2.621017 | 3.669933 | 2.997781 | 2.020765 | 1.367949 | 0.835812 | 0.594124 | 0.509988 | 0.531292 | 0.716522 | 1.571734 | 4.347297 | 14.279848 | 39.414180 | 97.714938 | 155.480777 | 148.107209 | 96.629218 | 50.339152 | 20.594022 | 11.193083 | 6.864267 | 4.386318 | 2.784098 | 1.808976 | 0.838935 | 0.473643 | 0.353610 | 0.301133 | 0.269711 | 0.248816 | 0.230776 | 0.220960 | 0.216304 | 0.214916 | 0.214948 | 0.215031 | 0.214906 | 0.214670 | 0.215928 | 0.225200 | 0.246742 | 0.291616 | 0.359148 | 0.458476 | 0.525188 | 0.509744 | 0.424392 | 0.330126 | 0.267406 | 0.236289 | 0.221125 | 0.215319 | 0.214634 | 0.214767 | 0.214654 | 0.215848 | 0.224573 | 0.252941 | 0.320948 | 0.472301 | 0.659014 | 0.793322 | 0.598292 | 0.393934 | 0.284346 | 0.238755 | 0.221137 | 0.217306 | 0.217962 | 0.226908 | 0.250028 | 0.292548 | 0.402881 | 0.613725 | 0.801839 | 1.230239 | 2.162724 | 3.203595 | 5.061597 | 7.176623 | 7.896518 | 8.201551 | 6.978311 | 4.190506 | 2.392169 | 1.245710 | 0.656555 | 0.410075 | 0.319006 | 0.278411 | 0.275850 | 0.291513 | 0.326062 | 0.358712 | 0.369173 | 0.341608 | 0.308979 | 0.269703 | 0.248718 | 0.240066 | 0.246431 | 0.267915 | 0.323769 | 0.434233 | 0.576350 | 0.553639 | 0.504905 | 0.445156 | 0.389475 | 0.333628 | 0.296364 | 0.254200 | 0.227714 | 0.214652 | 0.221844 | 0.234726 | 0.255168 | 0.290186 | 0.324575 | 0.360926 | 0.390471 | 0.421052 | 0.434935 | 0.443568 | 0.474802 | 0.561547 | 0.611129 | 0.650394 | 0.650373 | 0.717244 | 0.904206 | 1.187898 | 1.500546 | 2.018899 | 2.626938 | 3.134353 | 3.230018 | 3.137472 | 2.725908 | 2.498260 | 2.167336 | 1.697067 | 1.326005 | 1.060534 | 0.806130 | 0.654975 | 0.583855 | 0.514829 | 0.469265 | 0.422117 | 0.377044 | 0.358551 | 0.347438 | 0.304957 | 0.275876 | 0.248479 | 0.224398 | 0.215699 | 0.214626 | 0.217099 | 0.220730 | 0.219193 | 0.216301 | 0.214631 | 0.217110 | 0.220848 | 0.223514 | 0.224900 | 0.226341 | 0.234806 | 0.256866 | 0.292256 | 0.340256 | 0.409349 | 0.483081 | 0.518437 | 0.505023 | 0.495390 | 0.487327 | 0.518423 | 0.581300 | 0.662556 | 0.767072 | 0.878753 | 0.922010 | 0.934308 | 0.999215 | 1.049750 | 1.049935 | 1.003859 | 0.919665 | 0.877826 | 0.802573 | 0.721569 | 0.679847 | 0.672000 | 0.621063 |
| right anterior | 0.533819 | 0.414861 | 0.339242 | 0.254834 | 0.216489 | 0.228628 | 0.293918 | 0.427788 | 0.681434 | 0.957082 | 1.125546 | 1.144924 | 1.189077 | 1.069438 | 0.742323 | 0.478446 | 0.320106 | 0.233061 | 0.217633 | 0.253656 | 0.302435 | 0.360106 | 0.548535 | 0.757631 | 0.885532 | 0.825925 | 0.683062 | 0.449062 | 0.274226 | 0.215625 | 0.249563 | 0.470393 | 0.991153 | 1.769072 | 2.030425 | 1.410856 | 0.785106 | 0.503432 | 0.333613 | 0.281504 | 0.273737 | 0.291968 | 0.312822 | 0.358126 | 0.477484 | 0.735977 | 0.767364 | 0.656789 | 0.450727 | 0.331348 | 0.296595 | 0.298264 | 0.314668 | 0.366441 | 0.379636 | 0.403148 | 0.419989 | 0.483008 | 0.463585 | 0.432314 | 0.336693 | 0.283742 | 0.227512 | 0.214627 | 0.223218 | 0.230857 | 0.243796 | 0.242968 | 0.244981 | 0.264392 | 0.340682 | 0.411137 | 0.462681 | 0.512175 | 0.504917 | 0.458519 | 0.389217 | 0.306414 | 0.236981 | 0.215144 | 0.276987 | 0.450326 | 0.768125 | 1.156606 | 1.637646 | 1.666122 | 1.303601 | 1.034768 | 0.676022 | 0.411159 | 0.298235 | 0.236834 | 0.215387 | 0.219284 | 0.235566 | 0.249371 | 0.236532 | 0.227403 | 0.217878 | 0.214639 | 0.218327 | 0.228098 | 0.249228 | 0.259003 | 0.272877 | 0.268614 | 0.253707 | 0.229822 | 0.219051 | 0.214627 | 0.219007 | 0.236846 | 0.260442 | 0.305484 | 0.388620 | 0.551257 | 0.837875 | 1.410851 | 2.194878 | 3.577274 | 5.529549 | 7.270584 | 8.701024 | 8.851190 | 5.274874 | 2.239688 | 0.836071 | 0.427487 | 0.272290 | 0.226769 | 0.215169 | 0.229439 | 0.258680 | 0.266522 | 0.262431 | 0.246998 | 0.248292 | 0.269669 | 0.319182 | 0.411150 | 0.443568 | 0.416307 | 0.290766 | 0.214967 | 0.281324 | 0.538368 | 1.156951 | 2.099095 | 2.596012 | 2.847335 | 2.130538 | 1.506274 | 0.976131 | 0.546970 | 0.307103 | 0.234531 | 0.215571 | 0.251549 | 0.339832 | 0.449167 | 0.551948 | 0.609497 | 0.638124 | 0.630389 | 0.690308 | 0.713435 | 0.628436 | 0.510382 | 0.386879 | 0.288781 | 0.241922 | 0.220279 | 0.214812 | 0.214630 | 0.215936 | 0.225221 | 0.243082 | 0.271432 | 0.291894 | 0.280867 | 0.268583 | 0.290512 | 0.392860 | 0.781275 | 1.894305 | 3.432698 | 4.131313 | 3.198048 | 2.141797 | 1.548103 | 1.432073 | 1.608007 | 1.859661 | 2.196273 | 2.376086 | 2.004976 | 1.452441 | 0.994368 | 0.819991 | 0.823146 | 1.088951 | 1.587367 | 2.753644 | 2.800368 | 2.015040 | 1.030947 | 0.542802 | 0.335543 | 0.272836 | 0.245874 | 0.244703 | 0.248827 | 0.260214 | 0.268611 | 0.269831 | 0.265089 | 0.267940 | 0.260768 | 0.252807 | 0.245354 | 0.232756 | 0.218695 | 0.214626 | 0.222033 | 0.237197 | 0.249584 | 0.248444 | 0.247148 | 0.244310 | 0.254018 | 0.271275 | 0.304303 | 0.368919 | 0.456896 | 0.488180 | 0.446383 | 0.351689 | 0.260884 | 0.218014 | 0.219015 | 0.235452 | 0.241279 | 0.232757 | 0.222833 | 0.215278 | 0.217257 | 0.228907 | 0.238568 | 0.245991 | 0.239240 | 0.230795 | 0.227557 | 0.227840 | 0.231154 | 0.251932 | 0.295963 | 0.362656 | 0.463280 | 0.568950 | 0.639472 | 0.666909 | 0.650931 | 0.594409 | 0.550577 | 0.513159 | 0.473176 | 0.401522 | 0.319689 | 0.256768 | 0.222120 | 0.214779 | 0.219403 | 0.221767 | 0.222296 | 0.224445 | 0.229895 | 0.232829 | 0.235155 | 0.238360 | 0.237151 |
| left central | 0.483587 | 0.451999 | 0.466937 | 0.450410 | 0.386976 | 0.336539 | 0.347778 | 0.353754 | 0.359585 | 0.373045 | 0.419572 | 0.453742 | 0.541624 | 0.724321 | 1.148740 | 1.806311 | 2.493032 | 2.557649 | 2.086067 | 1.525141 | 1.167414 | 0.803063 | 0.578945 | 0.515189 | 0.454438 | 0.391912 | 0.373338 | 0.416829 | 0.445669 | 0.488735 | 0.640347 | 0.788837 | 0.739860 | 0.476653 | 0.328293 | 0.259341 | 0.231303 | 0.219571 | 0.216705 | 0.214659 | 0.214626 | 0.214886 | 0.216090 | 0.218254 | 0.223530 | 0.223684 | 0.219724 | 0.214744 | 0.218599 | 0.225308 | 0.237149 | 0.254088 | 0.265656 | 0.253012 | 0.237716 | 0.224851 | 0.219684 | 0.214933 | 0.215968 | 0.224086 | 0.232491 | 0.244623 | 0.255923 | 0.265075 | 0.253732 | 0.233251 | 0.214802 | 0.264542 | 0.544529 | 1.396341 | 2.725920 | 3.490841 | 4.025818 | 3.947533 | 3.299425 | 2.285199 | 1.384042 | 0.798820 | 0.461678 | 0.264830 | 0.215519 | 0.335985 | 0.845352 | 2.282264 | 5.080577 | 8.802877 | 9.746130 | 7.863908 | 4.594909 | 2.361326 | 1.153155 | 0.617431 | 0.432532 | 0.363349 | 0.323068 | 0.318808 | 0.373962 | 0.463393 | 0.639909 | 0.912932 | 1.229091 | 1.343946 | 1.206267 | 0.930869 | 0.662413 | 0.456264 | 0.328894 | 0.261463 | 0.230631 | 0.220211 | 0.215322 | 0.214781 | 0.216235 | 0.220691 | 0.231612 | 0.253385 | 0.287386 | 0.338072 | 0.389589 | 0.433793 | 0.450987 | 0.438410 | 0.387020 | 0.340903 | 0.300986 | 0.272137 | 0.256917 | 0.256492 | 0.259013 | 0.267259 | 0.279075 | 0.289986 | 0.281425 | 0.264957 | 0.239434 | 0.219190 | 0.215730 | 0.232084 | 0.281416 | 0.368020 | 0.545958 | 0.904834 | 1.487179 | 2.122028 | 3.475299 | 5.227912 | 5.738974 | 4.499886 | 2.575675 | 1.304842 | 0.754831 | 0.513543 | 0.418093 | 0.415451 | 0.423068 | 0.448883 | 0.475195 | 0.522933 | 0.495659 | 0.433963 | 0.360283 | 0.346185 | 0.346535 | 0.424357 | 0.605308 | 0.905325 | 1.085445 | 1.001568 | 0.703752 | 0.556059 | 0.450755 | 0.386870 | 0.336917 | 0.308910 | 0.281725 | 0.259900 | 0.235834 | 0.224831 | 0.216223 | 0.215227 | 0.220255 | 0.222596 | 0.224380 | 0.220377 | 0.216678 | 0.214674 | 0.217157 | 0.225756 | 0.229201 | 0.234546 | 0.235217 | 0.231926 | 0.219765 | 0.214895 | 0.221302 | 0.247693 | 0.290586 | 0.332801 | 0.356315 | 0.367032 | 0.395187 | 0.419668 | 0.404958 | 0.370372 | 0.325809 | 0.287458 | 0.270869 | 0.268992 | 0.284779 | 0.326231 | 0.393262 | 0.462140 | 0.529321 | 0.568133 | 0.589330 | 0.609020 | 0.655647 | 0.757495 | 0.937287 | 1.221287 | 1.633413 | 2.072052 | 2.036223 | 1.555444 | 1.086285 | 0.844374 | 0.824203 | 1.070437 | 1.979567 | 5.143926 | 12.105778 | 16.696020 | 14.385548 | 8.452945 | 3.932967 | 1.807959 | 0.897169 | 0.500983 | 0.351758 | 0.286993 | 0.258492 | 0.239250 | 0.227855 | 0.220672 | 0.214722 | 0.220851 | 0.237494 | 0.252996 | 0.264252 | 0.274747 | 0.281660 | 0.269866 | 0.251937 | 0.237025 | 0.222680 | 0.214935 | 0.217291 | 0.228461 | 0.249917 | 0.281249 | 0.310165 | 0.325896 | 0.325597 | 0.319190 | 0.301658 | 0.280550 | 0.261730 | 0.238384 | 0.220707 | 0.214642 | 0.218619 | 0.236787 | 0.266214 | 0.310165 | 0.356996 | 0.400084 | 0.422117 | 0.427780 | 0.416760 | 0.408135 |
| right central | 0.240313 | 0.262561 | 0.271772 | 0.312653 | 0.355627 | 0.370403 | 0.350609 | 0.337364 | 0.297578 | 0.288029 | 0.259236 | 0.228099 | 0.215377 | 0.215143 | 0.220635 | 0.221589 | 0.231077 | 0.241418 | 0.244948 | 0.247127 | 0.257139 | 0.256809 | 0.290803 | 0.376645 | 0.505945 | 0.616350 | 0.568211 | 0.501177 | 0.464573 | 0.454879 | 0.469005 | 0.657376 | 0.973917 | 1.455821 | 1.452031 | 0.723876 | 0.320228 | 0.216976 | 0.277703 | 0.596669 | 1.450565 | 3.313752 | 5.320582 | 2.812843 | 0.973875 | 0.343580 | 0.227036 | 0.214808 | 0.214674 | 0.214639 | 0.220237 | 0.245065 | 0.296470 | 0.380151 | 0.626404 | 1.110002 | 2.403228 | 3.761056 | 2.220050 | 0.873184 | 0.447052 | 0.279336 | 0.228232 | 0.214626 | 0.223570 | 0.234574 | 0.248816 | 0.299779 | 0.441965 | 1.054198 | 3.618205 | 10.463216 | 17.283414 | 16.890202 | 8.741021 | 3.542880 | 1.458369 | 0.651144 | 0.345166 | 0.243229 | 0.216441 | 0.214695 | 0.214855 | 0.218444 | 0.228729 | 0.257821 | 0.285426 | 0.307684 | 0.372756 | 0.516900 | 0.806549 | 1.408425 | 2.571404 | 4.077538 | 5.737433 | 5.333074 | 4.086836 | 2.300424 | 1.082425 | 0.546070 | 0.369651 | 0.312631 | 0.321768 | 0.396992 | 0.519169 | 0.753942 | 1.077450 | 1.169421 | 0.911974 | 0.624760 | 0.449577 | 0.364177 | 0.336693 | 0.349016 | 0.393884 | 0.497286 | 0.704562 | 1.050681 | 1.612191 | 2.348003 | 2.808206 | 2.997363 | 2.961432 | 2.979380 | 3.519863 | 4.568837 | 6.742674 | 11.139351 | 16.619647 | 21.245665 | 26.379627 | 21.268891 | 13.329337 | 7.745074 | 4.665147 | 2.792636 | 1.797607 | 1.108596 | 0.778820 | 0.611865 | 0.550173 | 0.559306 | 0.645961 | 0.707011 | 0.724201 | 0.681808 | 0.594001 | 0.505418 | 0.523571 | 0.549758 | 0.574852 | 0.593743 | 0.610870 | 0.572419 | 0.551881 | 0.572420 | 0.632436 | 0.726769 | 1.006095 | 1.560302 | 2.446078 | 3.399634 | 3.774871 | 2.852773 | 1.837449 | 1.084986 | 0.744244 | 0.554245 | 0.497948 | 0.531288 | 0.624059 | 0.729153 | 0.909485 | 1.029618 | 1.069907 | 1.046484 | 1.044159 | 1.158585 | 1.297308 | 1.353153 | 1.413152 | 1.431553 | 1.287212 | 1.024203 | 0.777182 | 0.588000 | 0.452978 | 0.354816 | 0.317185 | 0.305117 | 0.306989 | 0.333392 | 0.392738 | 0.510338 | 0.739233 | 1.027026 | 1.127651 | 1.065167 | 1.022133 | 1.102311 | 1.452476 | 2.595400 | 6.771915 | 20.691492 | 45.026465 | 50.054309 | 28.125615 | 9.460714 | 2.702725 | 1.161178 | 0.816897 | 0.866749 | 1.188574 | 1.960611 | 2.886593 | 3.734719 | 3.555452 | 2.794918 | 1.859065 | 1.257070 | 0.849547 | 0.733790 | 0.677927 | 0.716258 | 0.807580 | 0.889542 | 0.810467 | 0.748824 | 0.692288 | 0.651747 | 0.588897 | 0.530325 | 0.494962 | 0.520167 | 0.580878 | 0.668177 | 0.732576 | 0.749148 | 0.704003 | 0.555558 | 0.415326 | 0.316604 | 0.253607 | 0.221532 | 0.215524 | 0.215060 | 0.219026 | 0.235409 | 0.274915 | 0.340773 | 0.436670 | 0.540773 | 0.643465 | 0.658735 | 0.575031 | 0.455694 | 0.328194 | 0.265760 | 0.232137 | 0.216235 | 0.215243 | 0.215948 | 0.217366 | 0.215138 | 0.218028 | 0.238913 | 0.267060 | 0.282686 | 0.288224 | 0.283749 | 0.266623 | 0.243548 | 0.226827 | 0.216282 | 0.214976 | 0.220367 | 0.223775 | 0.219972 | 0.217178 | 0.215686 |
| left posterior | 0.672952 | 0.843586 | 1.089188 | 2.160750 | 2.830040 | 2.266253 | 1.181654 | 0.715496 | 0.437075 | 0.291062 | 0.242670 | 0.226926 | 0.229269 | 0.253149 | 0.412178 | 2.056070 | 63.618434 | 2949.553428 | 32832.710349 | 15529.726135 | 590.389914 | 20.986372 | 2.170339 | 0.422500 | 0.220293 | 0.226062 | 0.265583 | 0.294494 | 0.314390 | 0.251456 | 0.222349 | 0.217831 | 0.228160 | 0.232142 | 0.235633 | 0.232067 | 0.230752 | 0.219210 | 0.215580 | 0.214839 | 0.214736 | 0.215060 | 0.225114 | 0.255644 | 0.311310 | 0.403394 | 0.501630 | 0.576536 | 0.627742 | 0.676555 | 0.624516 | 0.542663 | 0.462764 | 0.420581 | 0.362962 | 0.298200 | 0.271095 | 0.271122 | 0.263324 | 0.256654 | 0.261472 | 0.263297 | 0.276585 | 0.314466 | 0.391781 | 0.613296 | 1.232680 | 3.003622 | 8.571524 | 25.853608 | 66.811089 | 162.504970 | 390.134331 | 851.642772 | 1567.760305 | 2030.592613 | 1557.722804 | 887.316787 | 342.717731 | 102.233502 | 30.661945 | 9.044463 | 2.236171 | 0.655955 | 0.318725 | 0.243387 | 0.222150 | 0.215523 | 0.215604 | 0.221224 | 0.240068 | 0.306306 | 0.538236 | 1.254056 | 3.824346 | 9.579274 | 13.977048 | 11.924723 | 8.039361 | 5.785225 | 4.116417 | 2.645154 | 1.918677 | 1.805483 | 1.934702 | 2.539486 | 3.221084 | 4.227235 | 4.717075 | 3.929282 | 2.652500 | 1.746419 | 1.149345 | 0.899616 | 0.823511 | 0.845716 | 1.005805 | 1.397853 | 2.032911 | 3.419615 | 6.331354 | 9.868386 | 10.358856 | 8.597502 | 5.392310 | 2.817465 | 1.478644 | 0.862812 | 0.565505 | 0.427935 | 0.365296 | 0.337903 | 0.335723 | 0.347801 | 0.365223 | 0.383942 | 0.424268 | 0.485378 | 0.527263 | 0.575496 | 0.644136 | 0.712106 | 0.783734 | 0.907434 | 0.975758 | 1.005078 | 0.831185 | 0.665378 | 0.580773 | 0.546217 | 0.533057 | 0.563516 | 0.568266 | 0.514624 | 0.437882 | 0.347839 | 0.287570 | 0.244545 | 0.221597 | 0.214841 | 0.226436 | 0.252464 | 0.271960 | 0.272767 | 0.259912 | 0.246575 | 0.224361 | 0.215341 | 0.245269 | 0.331788 | 0.528239 | 0.838780 | 1.140471 | 1.372316 | 1.464860 | 1.367448 | 1.241679 | 1.037945 | 0.829392 | 0.763163 | 0.805580 | 0.891474 | 1.072903 | 1.290303 | 1.381130 | 1.271934 | 1.048280 | 0.830556 | 0.667282 | 0.560838 | 0.507826 | 0.523412 | 0.615109 | 0.808319 | 1.226533 | 2.087590 | 3.376967 | 4.889668 | 6.163108 | 5.843656 | 4.523146 | 3.024323 | 1.959054 | 1.323571 | 1.015323 | 0.809137 | 0.729422 | 0.731883 | 0.829890 | 1.048754 | 1.695537 | 3.278667 | 6.770636 | 14.634878 | 30.887475 | 52.972054 | 79.497448 | 112.507199 | 100.687060 | 61.896220 | 29.667821 | 11.575233 | 4.287287 | 1.853091 | 1.007404 | 0.725931 | 0.624376 | 0.659781 | 0.794210 | 1.019543 | 1.116472 | 1.199447 | 1.164662 | 1.135790 | 1.173162 | 1.293927 | 1.429321 | 1.751593 | 2.043329 | 2.322499 | 2.833322 | 3.618736 | 4.436999 | 6.071740 | 8.131196 | 11.832876 | 15.584525 | 18.061398 | 15.994813 | 12.755940 | 7.989321 | 4.820009 | 3.095999 | 2.164641 | 1.524418 | 1.249254 | 1.047313 | 0.909184 | 0.798200 | 0.696931 | 0.566878 | 0.459412 | 0.386562 | 0.349330 | 0.331243 | 0.324939 | 0.334145 | 0.346500 | 0.362523 | 0.371368 | 0.365376 | 0.346190 | 0.312599 | 0.275073 | 0.249672 | 0.235540 | 0.226722 | 0.223065 | 0.220602 | 0.219396 |
| right posterior | 0.413773 | 0.378414 | 0.338838 | 0.305530 | 0.297734 | 0.328763 | 0.387258 | 0.568504 | 0.919525 | 1.241157 | 1.220147 | 0.828301 | 0.455401 | 0.337707 | 0.307672 | 0.293799 | 0.321802 | 0.410846 | 0.536148 | 0.546390 | 0.500419 | 0.365759 | 0.270797 | 0.225076 | 0.217039 | 0.221195 | 0.251233 | 0.307992 | 0.423539 | 0.692738 | 1.274184 | 1.553662 | 0.905710 | 0.418084 | 0.250638 | 0.214859 | 0.234380 | 0.265187 | 0.295316 | 0.286705 | 0.247296 | 0.215043 | 0.232768 | 0.315261 | 0.435636 | 0.529945 | 0.573541 | 0.684253 | 0.776705 | 0.849779 | 0.760969 | 0.575443 | 0.402094 | 0.303040 | 0.267350 | 0.259984 | 0.277641 | 0.329061 | 0.417343 | 0.546221 | 0.690266 | 0.627884 | 0.427743 | 0.306396 | 0.245063 | 0.216265 | 0.224994 | 0.303207 | 0.572637 | 1.427728 | 3.104585 | 5.798482 | 8.363432 | 8.431360 | 7.536545 | 5.901147 | 3.685571 | 2.083518 | 1.031946 | 0.479031 | 0.270809 | 0.217455 | 0.218662 | 0.229175 | 0.235250 | 0.232403 | 0.228716 | 0.225056 | 0.222221 | 0.218675 | 0.217324 | 0.217656 | 0.221225 | 0.229438 | 0.241203 | 0.254764 | 0.259734 | 0.249418 | 0.234235 | 0.224287 | 0.215612 | 0.217869 | 0.234579 | 0.274405 | 0.335354 | 0.394648 | 0.458984 | 0.510782 | 0.508655 | 0.507218 | 0.479245 | 0.423075 | 0.383706 | 0.362846 | 0.354010 | 0.373279 | 0.429214 | 0.546539 | 0.780342 | 1.216630 | 1.846334 | 2.810071 | 3.719011 | 3.820189 | 2.865079 | 2.002080 | 1.283075 | 0.900771 | 0.689667 | 0.579042 | 0.511873 | 0.490255 | 0.467175 | 0.455597 | 0.448312 | 0.451246 | 0.443405 | 0.443615 | 0.475704 | 0.577446 | 0.763224 | 1.072738 | 1.636064 | 2.346673 | 2.573093 | 2.273907 | 1.522485 | 0.930736 | 0.578814 | 0.414019 | 0.317675 | 0.273780 | 0.244458 | 0.231829 | 0.227980 | 0.237657 | 0.259719 | 0.312207 | 0.421881 | 0.596930 | 0.779283 | 1.022881 | 1.244662 | 1.382242 | 1.514541 | 1.666068 | 1.706941 | 1.791805 | 1.566323 | 1.241183 | 0.960145 | 0.782048 | 0.639818 | 0.554872 | 0.498238 | 0.483458 | 0.463847 | 0.450674 | 0.435788 | 0.424511 | 0.406164 | 0.413190 | 0.447232 | 0.547879 | 0.760668 | 1.225367 | 1.963273 | 2.762485 | 2.952336 | 2.591914 | 1.886702 | 1.370754 | 1.120775 | 1.058375 | 1.098777 | 1.276751 | 1.523854 | 1.734405 | 1.832095 | 1.787310 | 1.604343 | 1.435428 | 1.268867 | 1.190238 | 1.191672 | 1.220884 | 1.260607 | 1.377947 | 1.539207 | 1.564749 | 1.447838 | 1.319089 | 1.327068 | 1.399010 | 1.590344 | 2.122643 | 2.761479 | 3.177106 | 3.345255 | 3.425198 | 3.265349 | 3.109083 | 2.823481 | 2.422424 | 2.127509 | 1.916623 | 1.798995 | 1.797622 | 1.900632 | 2.143712 | 2.684794 | 3.462003 | 4.394329 | 6.023512 | 8.135813 | 9.493393 | 10.361839 | 10.875569 | 10.320570 | 9.331468 | 7.776369 | 6.469478 | 5.141865 | 4.022461 | 3.222900 | 2.645290 | 2.297845 | 2.172985 | 2.098269 | 2.099708 | 2.122523 | 2.050202 | 1.797830 | 1.422754 | 1.051469 | 0.806131 | 0.673384 | 0.619030 | 0.626874 | 0.681248 | 0.756851 | 0.833215 | 0.916807 | 0.978104 | 1.035954 | 1.009230 | 0.950672 | 0.845235 | 0.684432 | 0.490284 | 0.362308 | 0.276207 | 0.233419 | 0.219112 | 0.216374 | 0.216663 | 0.224718 | 0.246585 | 0.264796 | 0.280811 |
| all electrodes | 1.206335 | 2.635085 | 5.101085 | 14.442401 | 20.773379 | 12.803682 | 4.116907 | 1.938433 | 1.042897 | 0.601296 | 0.415911 | 0.352911 | 0.325614 | 0.367465 | 0.412101 | 0.447447 | 0.523397 | 0.632169 | 0.676541 | 1.243080 | 2.264034 | 2.407780 | 0.978122 | 0.398893 | 0.238435 | 0.216365 | 0.216206 | 0.214627 | 0.217880 | 0.229763 | 0.241249 | 0.270952 | 0.265275 | 0.248785 | 0.223848 | 0.216115 | 0.215628 | 0.219947 | 0.225614 | 0.220238 | 0.214968 | 0.244746 | 0.383935 | 1.058219 | 3.557793 | 6.675942 | 7.049962 | 4.626586 | 2.421166 | 1.518253 | 0.784939 | 0.515248 | 0.446423 | 0.484676 | 0.549440 | 0.566632 | 0.446576 | 0.308583 | 0.228549 | 0.215163 | 0.226641 | 0.243704 | 0.240728 | 0.225750 | 0.215840 | 0.217494 | 0.224999 | 0.216362 | 0.224732 | 0.317109 | 0.703118 | 2.271787 | 9.024069 | 29.943768 | 57.412957 | 55.022082 | 38.752250 | 22.735247 | 11.219004 | 5.737471 | 2.588293 | 0.843738 | 0.329779 | 0.216912 | 0.262290 | 0.411875 | 0.645098 | 0.850669 | 0.918852 | 1.105198 | 1.165169 | 1.090626 | 0.972718 | 0.847906 | 0.613195 | 0.450933 | 0.350798 | 0.303929 | 0.276949 | 0.275837 | 0.274249 | 0.276205 | 0.281973 | 0.272176 | 0.253959 | 0.236295 | 0.221847 | 0.214633 | 0.222927 | 0.253988 | 0.307022 | 0.419844 | 0.675792 | 1.070200 | 1.688585 | 2.690442 | 4.382661 | 6.561414 | 11.316777 | 15.153060 | 20.203664 | 21.927724 | 21.645462 | 17.879440 | 16.020182 | 12.259382 | 9.446286 | 7.022309 | 5.184007 | 3.891502 | 3.398966 | 3.801738 | 4.411720 | 3.915478 | 2.912124 | 1.727187 | 0.930673 | 0.524945 | 0.354603 | 0.285950 | 0.260656 | 0.246988 | 0.241733 | 0.246789 | 0.263917 | 0.275641 | 0.286348 | 0.296947 | 0.305516 | 0.290385 | 0.262803 | 0.229334 | 0.215395 | 0.221665 | 0.256376 | 0.329465 | 0.380629 | 0.416871 | 0.395083 | 0.364710 | 0.336479 | 0.326275 | 0.306279 | 0.296504 | 0.297189 | 0.290962 | 0.288286 | 0.302471 | 0.332671 | 0.352069 | 0.357082 | 0.346591 | 0.330509 | 0.297194 | 0.268925 | 0.245948 | 0.229961 | 0.218395 | 0.214665 | 0.217414 | 0.225141 | 0.231608 | 0.232463 | 0.226575 | 0.217195 | 0.215382 | 0.228001 | 0.268895 | 0.346037 | 0.432043 | 0.510463 | 0.514243 | 0.420283 | 0.337905 | 0.277141 | 0.238755 | 0.221326 | 0.215666 | 0.214877 | 0.219155 | 0.234308 | 0.264995 | 0.296768 | 0.329464 | 0.329642 | 0.313796 | 0.296942 | 0.303676 | 0.322922 | 0.377628 | 0.428293 | 0.524620 | 0.659817 | 0.817571 | 0.978493 | 0.991015 | 0.786599 | 0.570086 | 0.412104 | 0.310328 | 0.258669 | 0.226212 | 0.217900 | 0.218176 | 0.230264 | 0.267862 | 0.379757 | 0.478409 | 0.536992 | 0.531054 | 0.505074 | 0.427483 | 0.379237 | 0.338135 | 0.314558 | 0.315586 | 0.342976 | 0.384377 | 0.455110 | 0.528750 | 0.579493 | 0.583641 | 0.537466 | 0.477950 | 0.445049 | 0.426598 | 0.426266 | 0.438291 | 0.454154 | 0.479663 | 0.465101 | 0.442529 | 0.396569 | 0.335332 | 0.268746 | 0.231027 | 0.214965 | 0.222640 | 0.252811 | 0.290747 | 0.323287 | 0.307286 | 0.277847 | 0.260088 | 0.257873 | 0.261933 | 0.288399 | 0.346140 | 0.449213 | 0.579868 | 0.697744 | 0.877525 | 1.015274 | 1.038433 | 0.974042 | 0.932009 | 0.804343 | 0.648191 | 0.537915 | 0.470303 |

Q) maximum cross-correlation, angry vs sad

  
|  | time window | peak latency | cluster *p* | peak Cohen's *d* |  | | | |  |
| **all electrodes** |  | | | |  | | | |  |
|  | | | | | | | | |  |

Model correlations, cluster permutation tests

|  | **left hemisphere** | | | | **right hemisphere** | | | |  |
|  | time window | peak latency | cluster *p* | peak Cohen's *d* | time window | peak latency | cluster *p* | peak Cohen's *d* |  |
| **anterior** |  | | | | 105 - 220 ms | 190 ms | 0.0303 | -0.7383 |  |
| **central** | 185 - 275 ms | 240 ms | 0.0239 | 0.9508 | 215 - 495 ms | 240 ms | 0.0008 | 0.8191 |  |
 335 - 475 ms | 420 ms | 0.0157 | 0.6426 |  | | | ||  | **posterior** |  | | | |  | | | |

  

Model correlations, Bayesian statistics

|  | -200 | -195 | -190 | -185 | -180 | -175 | -170 | -165 | -160 | -155 | -150 | -145 | -140 | -135 | -130 | -125 | -120 | -115 | -110 | -105 | -100 | -95 | -90 | -85 | -80 | -75 | -70 | -65 | -60 | -55 | -50 | -45 | -40 | -35 | -30 | -25 | -20 | -15 | -10 | -5 | 0 | 5 | 10 | 15 | 20 | 25 | 30 | 35 | 40 | 45 | 50 | 55 | 60 | 65 | 70 | 75 | 80 | 85 | 90 | 95 | 100 | 105 | 110 | 115 | 120 | 125 | 130 | 135 | 140 | 145 | 150 | 155 | 160 | 165 | 170 | 175 | 180 | 185 | 190 | 195 | 200 | 205 | 210 | 215 | 220 | 225 | 230 | 235 | 240 | 245 | 250 | 255 | 260 | 265 | 270 | 275 | 280 | 285 | 290 | 295 | 300 | 305 | 310 | 315 | 320 | 325 | 330 | 335 | 340 | 345 | 350 | 355 | 360 | 365 | 370 | 375 | 380 | 385 | 390 | 395 | 400 | 405 | 410 | 415 | 420 | 425 | 430 | 435 | 440 | 445 | 450 | 455 | 460 | 465 | 470 | 475 | 480 | 485 | 490 | 495 | 500 | 505 | 510 | 515 | 520 | 525 | 530 | 535 | 540 | 545 | 550 | 555 | 560 | 565 | 570 | 575 | 580 | 585 | 590 | 595 | 600 | 605 | 610 | 615 | 620 | 625 | 630 | 635 | 640 | 645 | 650 | 655 | 660 | 665 | 670 | 675 | 680 | 685 | 690 | 695 | 700 | 705 | 710 | 715 | 720 | 725 | 730 | 735 | 740 | 745 | 750 | 755 | 760 | 765 | 770 | 775 | 780 | 785 | 790 | 795 | 800 | 805 | 810 | 815 | 820 | 825 | 830 | 835 | 840 | 845 | 850 | 855 | 860 | 865 | 870 | 875 | 880 | 885 | 890 | 895 | 900 | 905 | 910 | 915 | 920 | 925 | 930 | 935 | 940 | 945 | 950 | 955 | 960 | 965 | 970 | 975 | 980 | 985 | 990 | 995 | 1000 | 1005 | 1010 | 1015 | 1020 | 1025 | 1030 | 1035 | 1040 | 1045 | 1050 | 1055 | 1060 | 1065 | 1070 | 1075 | 1080 | 1085 | 1090 | 1095 | 1100 | 1105 | 1110 | 1115 | 1120 | 1125 | 1130 | 1135 | 1140 | 1145 | 1150 | 1155 | 1160 | 1165 | 1170 | 1175 | 1180 | 1185 | 1190 | 1195 |
| --- | --- | --- | --- | --- | --- | --- | --- | --- | --- | --- | --- | --- | --- | --- | --- | --- | --- | --- | --- | --- | --- | --- | --- | --- | --- | --- | --- | --- | --- | --- | --- | --- | --- | --- | --- | --- | --- | --- | --- | --- | --- | --- | --- | --- | --- | --- | --- | --- | --- | --- | --- | --- | --- | --- | --- | --- | --- | --- | --- | --- | --- | --- | --- | --- | --- | --- | --- | --- | --- | --- | --- | --- | --- | --- | --- | --- | --- | --- | --- | --- | --- | --- | --- | --- | --- | --- | --- | --- | --- | --- | --- | --- | --- | --- | --- | --- | --- | --- | --- | --- | --- | --- | --- | --- | --- | --- | --- | --- | --- | --- | --- | --- | --- | --- | --- | --- | --- | --- | --- | --- | --- | --- | --- | --- | --- | --- | --- | --- | --- | --- | --- | --- | --- | --- | --- | --- | --- | --- | --- | --- | --- | --- | --- | --- | --- | --- | --- | --- | --- | --- | --- | --- | --- | --- | --- | --- | --- | --- | --- | --- | --- | --- | --- | --- | --- | --- | --- | --- | --- | --- | --- | --- | --- | --- | --- | --- | --- | --- | --- | --- | --- | --- | --- | --- | --- | --- | --- | --- | --- | --- | --- | --- | --- | --- | --- | --- | --- | --- | --- | --- | --- | --- | --- | --- | --- | --- | --- | --- | --- | --- | --- | --- | --- | --- | --- | --- | --- | --- | --- | --- | --- | --- | --- | --- | --- | --- | --- | --- | --- | --- | --- | --- | --- | --- | --- | --- | --- | --- | --- | --- | --- | --- | --- | --- | --- | --- | --- | --- | --- | --- | --- | --- | --- | --- | --- | --- | --- | --- | --- | --- | --- | --- | --- | --- | --- | --- | --- | --- | --- | --- | --- | --- | --- | --- | --- | --- | --- | --- | --- | --- |
| left anterior | 0.222913 | 0.228387 | 0.226435 | 0.219041 | 0.215301 | 0.238350 | 0.294628 | 0.316837 | 0.280637 | 0.248188 | 0.221660 | 0.219187 | 0.273246 | 0.437460 | 0.792958 | 0.968703 | 0.901460 | 0.667373 | 0.400418 | 0.298703 | 0.251283 | 0.217759 | 0.214637 | 0.226737 | 0.266983 | 0.268948 | 0.299145 | 0.369171 | 0.465482 | 0.571819 | 0.533745 | 0.301432 | 0.217188 | 0.267256 | 0.403060 | 0.532381 | 0.568145 | 0.512837 | 0.395892 | 0.353272 | 0.305706 | 0.288733 | 0.293975 | 0.323810 | 0.334445 | 0.311056 | 0.247776 | 0.216945 | 0.227900 | 0.309892 | 0.362753 | 0.331824 | 0.238252 | 0.229723 | 0.466390 | 1.117637 | 2.415679 | 2.819154 | 3.129711 | 3.171044 | 2.181310 | 1.738988 | 1.890494 | 1.585260 | 0.989128 | 0.712931 | 0.482189 | 0.375758 | 0.342786 | 0.381504 | 0.505323 | 0.911481 | 1.586594 | 2.375252 | 2.011169 | 1.113206 | 0.565172 | 0.420937 | 0.416946 | 0.597933 | 1.182341 | 2.073754 | 2.401000 | 1.683617 | 0.931644 | 0.529498 | 0.396704 | 0.331774 | 0.289116 | 0.249283 | 0.242671 | 0.239328 | 0.237730 | 0.232541 | 0.234225 | 0.244023 | 0.267924 | 0.298716 | 0.367463 | 0.555779 | 0.894873 | 1.539772 | 2.626215 | 3.556694 | 3.255407 | 2.720536 | 1.542647 | 0.838302 | 0.512610 | 0.453454 | 0.425450 | 0.435822 | 0.424630 | 0.432692 | 0.348056 | 0.265465 | 0.217536 | 0.221162 | 0.256637 | 0.297214 | 0.317325 | 0.308370 | 0.290872 | 0.260586 | 0.238072 | 0.227318 | 0.224891 | 0.227939 | 0.232987 | 0.241282 | 0.246538 | 0.237047 | 0.217025 | 0.218676 | 0.256439 | 0.342352 | 0.481935 | 0.645026 | 0.673254 | 0.681287 | 0.656612 | 0.641900 | 0.644670 | 0.725030 | 0.750098 | 0.796856 | 0.704662 | 0.582885 | 0.469018 | 0.381852 | 0.304649 | 0.266795 | 0.242948 | 0.228790 | 0.221709 | 0.224128 | 0.223372 | 0.221936 | 0.223296 | 0.224650 | 0.221973 | 0.219371 | 0.217606 | 0.216867 | 0.216433 | 0.215075 | 0.214700 | 0.214803 | 0.216859 | 0.217922 | 0.214919 | 0.220126 | 0.254519 | 0.310296 | 0.365669 | 0.393084 | 0.351527 | 0.296090 | 0.255066 | 0.229028 | 0.217326 | 0.214672 | 0.214749 | 0.215488 | 0.225338 | 0.260787 | 0.376864 | 0.685281 | 0.991435 | 1.298484 | 1.783254 | 2.813172 | 4.777146 | 9.697047 | 16.130346 | 15.715742 | 7.717860 | 3.032015 | 1.212695 | 0.652839 | 0.438301 | 0.383856 | 0.417758 | 0.576428 | 0.804469 | 1.209793 | 1.694736 | 2.506445 | 3.845146 | 7.213106 | 17.391474 | 55.892319 | 115.300286 | 148.412306 | 98.907880 | 49.089552 | 20.229872 | 8.828391 | 4.192234 | 2.366465 | 1.601813 | 1.173615 | 0.963764 | 0.803303 | 0.776648 | 0.821642 | 1.006303 | 1.159036 | 1.279836 | 1.274143 | 1.258096 | 1.168616 | 1.102693 | 1.029217 | 0.997223 | 0.973038 | 0.970810 | 1.047658 | 1.086826 | 1.033794 | 0.937617 | 0.796442 | 0.647258 | 0.613857 | 0.596342 | 0.610194 | 0.660179 | 0.691456 | 0.686128 | 0.667701 | 0.658509 | 0.679905 | 0.784136 | 0.916640 | 1.205222 | 1.629194 | 2.138412 | 2.449262 | 2.396265 | 1.733298 | 1.144960 | 0.712265 | 0.505420 | 0.391488 | 0.350260 | 0.335140 | 0.342685 | 0.375155 | 0.439194 | 0.581671 | 0.846135 | 1.160358 | 1.397598 | 1.553011 | 1.279560 | 0.929441 | 0.624034 | 0.420557 | 0.346182 | 0.298708 |
| right anterior | 0.257311 | 0.235771 | 0.219858 | 0.215089 | 0.223301 | 0.240220 | 0.251320 | 0.254541 | 0.242161 | 0.233530 | 0.240051 | 0.254984 | 0.253331 | 0.277080 | 0.344653 | 0.373190 | 0.354950 | 0.303171 | 0.252424 | 0.221487 | 0.215266 | 0.222438 | 0.237530 | 0.261703 | 0.269077 | 0.287762 | 0.283073 | 0.251404 | 0.221022 | 0.214857 | 0.218116 | 0.249212 | 0.261293 | 0.261214 | 0.249591 | 0.228699 | 0.218636 | 0.215903 | 0.215028 | 0.215115 | 0.214822 | 0.217211 | 0.222058 | 0.228324 | 0.219238 | 0.214682 | 0.225901 | 0.276264 | 0.376390 | 0.464710 | 0.425362 | 0.321139 | 0.245808 | 0.217362 | 0.215132 | 0.222177 | 0.228928 | 0.253035 | 0.310131 | 0.473338 | 0.806221 | 1.812383 | 3.847243 | 8.728652 | 6.761173 | 3.270360 | 1.570851 | 0.983164 | 0.746187 | 0.975478 | 1.498136 | 2.845800 | 4.460578 | 6.226683 | 6.256455 | 7.228357 | 8.393831 | 12.158052 | 20.203500 | 25.192460 | 25.311152 | 22.651217 | 12.474149 | 4.540225 | 1.795913 | 0.779420 | 0.439073 | 0.332132 | 0.288596 | 0.275727 | 0.280801 | 0.271046 | 0.252148 | 0.242606 | 0.230813 | 0.220948 | 0.221361 | 0.226903 | 0.240337 | 0.282216 | 0.346321 | 0.419584 | 0.459142 | 0.387439 | 0.310044 | 0.282809 | 0.259138 | 0.248581 | 0.249106 | 0.248641 | 0.236729 | 0.218486 | 0.215759 | 0.232397 | 0.265699 | 0.307790 | 0.317570 | 0.301586 | 0.274467 | 0.263994 | 0.254037 | 0.247121 | 0.242817 | 0.238068 | 0.234657 | 0.232953 | 0.229103 | 0.229696 | 0.233237 | 0.234797 | 0.229555 | 0.218919 | 0.216119 | 0.230472 | 0.260431 | 0.305832 | 0.342094 | 0.354737 | 0.424511 | 0.582024 | 0.821705 | 1.227274 | 1.680097 | 1.467766 | 1.297417 | 1.025311 | 0.748343 | 0.631222 | 0.541951 | 0.407083 | 0.349880 | 0.320384 | 0.297328 | 0.292958 | 0.311867 | 0.356705 | 0.443734 | 0.649931 | 1.166873 | 2.048257 | 2.757912 | 3.835079 | 6.724930 | 14.653546 | 32.952906 | 47.136065 | 53.641426 | 58.442685 | 76.187497 | 58.028781 | 39.780657 | 16.539351 | 6.558355 | 2.606616 | 1.494012 | 0.973052 | 0.785724 | 0.748668 | 0.880509 | 1.081413 | 1.458014 | 2.278170 | 3.942850 | 5.372902 | 6.937689 | 5.918052 | 4.038943 | 2.600155 | 1.894964 | 1.321675 | 1.119628 | 0.925948 | 0.756638 | 0.605902 | 0.506796 | 0.423498 | 0.362398 | 0.341150 | 0.368214 | 0.463052 | 0.728890 | 1.560989 | 3.504976 | 6.749092 | 8.131243 | 7.050789 | 4.868988 | 3.228687 | 2.039585 | 1.177283 | 0.741310 | 0.518304 | 0.396168 | 0.339660 | 0.330747 | 0.361972 | 0.504575 | 0.872225 | 1.614941 | 2.195142 | 2.046847 | 1.315977 | 0.789861 | 0.505513 | 0.407307 | 0.406809 | 0.435705 | 0.475669 | 0.501577 | 0.487969 | 0.416310 | 0.326454 | 0.272581 | 0.238740 | 0.222933 | 0.216493 | 0.215309 | 0.215507 | 0.216045 | 0.214827 | 0.221589 | 0.244554 | 0.292846 | 0.377968 | 0.446367 | 0.459321 | 0.403326 | 0.319789 | 0.252653 | 0.219121 | 0.215937 | 0.221883 | 0.224413 | 0.227490 | 0.225488 | 0.222785 | 0.223329 | 0.224661 | 0.226884 | 0.234591 | 0.233141 | 0.229119 | 0.221421 | 0.217569 | 0.216557 | 0.217744 | 0.221376 | 0.230816 | 0.246557 | 0.268475 | 0.298609 | 0.336599 | 0.388065 | 0.423787 | 0.431538 | 0.421868 | 0.379459 | 0.352651 | 0.335523 | 0.317411 |
| left central | 0.232607 | 0.231449 | 0.230436 | 0.226842 | 0.228356 | 0.241534 | 0.268824 | 0.285325 | 0.335194 | 0.435806 | 0.488746 | 0.488311 | 0.466324 | 0.389812 | 0.331715 | 0.303367 | 0.299046 | 0.342203 | 0.411308 | 0.448551 | 0.570195 | 0.889403 | 1.756784 | 2.244809 | 2.455437 | 1.926416 | 1.380682 | 0.559147 | 0.269678 | 0.217732 | 0.282045 | 0.387073 | 0.392241 | 0.285589 | 0.221990 | 0.220155 | 0.246823 | 0.264850 | 0.272353 | 0.279437 | 0.248192 | 0.222677 | 0.216800 | 0.270985 | 0.463002 | 0.701121 | 0.768032 | 0.552421 | 0.433163 | 0.371102 | 0.319792 | 0.278553 | 0.241100 | 0.216408 | 0.217762 | 0.232286 | 0.254585 | 0.268062 | 0.271889 | 0.264157 | 0.250109 | 0.239821 | 0.237068 | 0.235100 | 0.231073 | 0.230326 | 0.219149 | 0.214639 | 0.216280 | 0.216033 | 0.214859 | 0.219091 | 0.231858 | 0.259044 | 0.309707 | 0.417959 | 0.720084 | 1.841094 | 6.438681 | 18.194842 | 26.313459 | 23.609657 | 20.961305 | 22.872236 | 32.525199 | 60.675870 | 117.031662 | 198.273471 | 205.880088 | 117.723747 | 65.169158 | 36.081378 | 22.913803 | 12.272557 | 6.000438 | 2.545697 | 1.096015 | 0.493461 | 0.301120 | 0.232709 | 0.215554 | 0.215128 | 0.215478 | 0.214912 | 0.233802 | 0.322316 | 0.584768 | 1.424673 | 2.863095 | 3.617855 | 3.861752 | 3.529034 | 2.823563 | 2.563758 | 2.286337 | 2.438418 | 2.694139 | 3.409037 | 4.138747 | 5.716540 | 6.769164 | 7.068218 | 7.043797 | 6.842667 | 6.602606 | 6.290288 | 5.105775 | 4.044310 | 3.309455 | 2.695446 | 3.027906 | 4.250850 | 6.155866 | 7.051849 | 4.596893 | 2.064817 | 1.147894 | 0.738983 | 0.480419 | 0.332742 | 0.245487 | 0.214691 | 0.253684 | 0.385377 | 0.610560 | 0.733510 | 0.539337 | 0.342092 | 0.236852 | 0.215447 | 0.251222 | 0.317131 | 0.368415 | 0.400422 | 0.434464 | 0.433956 | 0.412711 | 0.394945 | 0.356063 | 0.300588 | 0.247725 | 0.218597 | 0.219894 | 0.260307 | 0.332138 | 0.412434 | 0.471443 | 0.446556 | 0.345065 | 0.251944 | 0.214883 | 0.234898 | 0.286455 | 0.316608 | 0.281923 | 0.238188 | 0.217336 | 0.217398 | 0.229977 | 0.240378 | 0.241626 | 0.232029 | 0.220506 | 0.216337 | 0.215480 | 0.216843 | 0.222247 | 0.238012 | 0.258500 | 0.266450 | 0.258238 | 0.249057 | 0.246189 | 0.249490 | 0.255164 | 0.270606 | 0.309905 | 0.364423 | 0.431347 | 0.492462 | 0.543578 | 0.563868 | 0.580111 | 0.608117 | 0.652141 | 0.696906 | 0.699531 | 0.649340 | 0.566631 | 0.478515 | 0.387438 | 0.340115 | 0.313756 | 0.298859 | 0.278167 | 0.260353 | 0.239224 | 0.223988 | 0.214820 | 0.219243 | 0.234975 | 0.246301 | 0.250671 | 0.252113 | 0.253954 | 0.263133 | 0.295914 | 0.365273 | 0.476389 | 0.643017 | 0.831274 | 0.962242 | 0.959258 | 0.872391 | 0.728492 | 0.593965 | 0.488519 | 0.447445 | 0.435345 | 0.445221 | 0.470402 | 0.505691 | 0.503812 | 0.507106 | 0.451230 | 0.371168 | 0.299968 | 0.252026 | 0.223231 | 0.214898 | 0.215518 | 0.215681 | 0.215044 | 0.214890 | 0.214834 | 0.215059 | 0.215339 | 0.215357 | 0.215933 | 0.216858 | 0.218515 | 0.220304 | 0.227836 | 0.238031 | 0.255551 | 0.270566 | 0.290706 | 0.305683 | 0.301305 | 0.271820 | 0.249518 | 0.242686 | 0.235696 | 0.250371 | 0.266892 | 0.289695 | 0.321984 | 0.367410 | 0.355984 | 0.376454 |
| right central | 2.176847 | 1.782800 | 1.601183 | 1.448655 | 1.355179 | 1.238409 | 1.075172 | 1.067883 | 0.749330 | 0.705028 | 0.735567 | 0.586969 | 0.343484 | 0.257506 | 0.224116 | 0.219357 | 0.219424 | 0.216403 | 0.214691 | 0.221311 | 0.253748 | 0.346235 | 0.431974 | 0.487832 | 0.427837 | 0.382033 | 0.314563 | 0.265906 | 0.226471 | 0.214857 | 0.214639 | 0.221318 | 0.240672 | 0.277328 | 0.336646 | 0.381876 | 0.404128 | 0.405743 | 0.352442 | 0.318626 | 0.313300 | 0.309804 | 0.319989 | 0.373233 | 0.398717 | 0.382711 | 0.345698 | 0.333478 | 0.312101 | 0.303029 | 0.274706 | 0.236921 | 0.215365 | 0.222645 | 0.281235 | 0.471709 | 1.036906 | 2.671358 | 4.295642 | 1.528640 | 0.519249 | 0.285993 | 0.237478 | 0.222494 | 0.228948 | 0.248740 | 0.315547 | 0.436945 | 0.622409 | 0.775743 | 0.782942 | 0.701493 | 0.583807 | 0.484881 | 0.366610 | 0.310627 | 0.264465 | 0.246673 | 0.234614 | 0.223440 | 0.215652 | 0.276178 | 0.771889 | 3.813214 | 20.737231 | 92.459783 | 168.738351 | 129.286365 | 66.077387 | 29.371471 | 14.207907 | 8.476933 | 6.022921 | 4.498172 | 3.337669 | 2.442056 | 1.806224 | 1.274938 | 1.076009 | 1.000517 | 1.130563 | 1.550632 | 2.396818 | 3.488866 | 4.835720 | 5.622293 | 6.026034 | 7.247846 | 11.279638 | 22.396151 | 40.469028 | 62.836338 | 79.050425 | 107.606708 | 98.639064 | 76.652846 | 42.227512 | 25.914451 | 16.548305 | 15.283531 | 18.780366 | 32.492896 | 51.121768 | 72.412877 | 97.502140 | 130.748793 | 130.768932 | 106.012074 | 87.381014 | 77.202374 | 79.581740 | 145.418436 | 431.339369 | 1401.139175 | 2379.757459 | 887.419786 | 138.751824 | 19.045231 | 4.218586 | 1.466957 | 0.596920 | 0.325767 | 0.256151 | 0.232677 | 0.226391 | 0.233284 | 0.256145 | 0.321981 | 0.461375 | 0.661130 | 0.876268 | 1.123596 | 1.080186 | 0.971797 | 0.901615 | 0.888643 | 1.021698 | 1.447185 | 2.196362 | 3.057316 | 3.033393 | 2.062973 | 1.152767 | 0.614476 | 0.400816 | 0.326237 | 0.316110 | 0.372690 | 0.557353 | 0.872241 | 1.324310 | 1.589344 | 1.509250 | 1.233164 | 0.990630 | 0.822283 | 0.836404 | 0.835067 | 0.790004 | 0.789298 | 0.713276 | 0.621305 | 0.599763 | 0.627432 | 0.700221 | 0.877955 | 1.010814 | 1.094467 | 1.116240 | 1.025434 | 0.774521 | 0.620079 | 0.509276 | 0.383343 | 0.302770 | 0.255532 | 0.221173 | 0.215693 | 0.227186 | 0.230741 | 0.216392 | 0.226981 | 0.298646 | 0.459481 | 0.656188 | 0.618889 | 0.414184 | 0.280609 | 0.223667 | 0.217788 | 0.248331 | 0.281489 | 0.287677 | 0.280915 | 0.276388 | 0.261738 | 0.241896 | 0.229682 | 0.222734 | 0.217976 | 0.216193 | 0.215400 | 0.215367 | 0.215031 | 0.214632 | 0.215935 | 0.219740 | 0.228838 | 0.243452 | 0.256592 | 0.263492 | 0.261194 | 0.251153 | 0.249526 | 0.257814 | 0.267317 | 0.284343 | 0.303229 | 0.307615 | 0.293543 | 0.274550 | 0.243990 | 0.220795 | 0.214972 | 0.231181 | 0.268553 | 0.303528 | 0.315246 | 0.291398 | 0.249116 | 0.230135 | 0.219289 | 0.214858 | 0.215503 | 0.219548 | 0.231417 | 0.245176 | 0.269995 | 0.297526 | 0.311613 | 0.324285 | 0.333887 | 0.306601 | 0.280914 | 0.271858 | 0.270833 | 0.266370 | 0.262897 | 0.255210 | 0.241979 | 0.230984 | 0.221323 | 0.215827 | 0.214973 | 0.220572 | 0.243888 | 0.276144 | 0.314117 | 0.334981 | 0.361777 |
| left posterior | 1.350254 | 1.379829 | 1.121802 | 0.753775 | 0.466822 | 0.350111 | 0.297516 | 0.282854 | 0.259307 | 0.236212 | 0.217940 | 0.222357 | 0.268961 | 0.315179 | 0.330324 | 0.282665 | 0.217200 | 0.263022 | 0.500888 | 0.796301 | 0.637332 | 0.352163 | 0.216611 | 0.331334 | 0.906526 | 1.436147 | 1.165632 | 0.587467 | 0.313612 | 0.238459 | 0.223970 | 0.226854 | 0.264581 | 0.372823 | 0.678963 | 1.318832 | 2.029416 | 2.396343 | 2.295417 | 1.372446 | 0.478964 | 0.215764 | 0.321441 | 0.578238 | 0.730992 | 0.753962 | 0.708843 | 0.619418 | 0.516486 | 0.458623 | 0.415020 | 0.390296 | 0.362754 | 0.339133 | 0.301895 | 0.271753 | 0.261688 | 0.298221 | 0.469926 | 1.096857 | 2.176953 | 2.647737 | 2.012924 | 1.101545 | 0.584676 | 0.357366 | 0.262380 | 0.227476 | 0.216775 | 0.214715 | 0.214669 | 0.215579 | 0.223629 | 0.252642 | 0.332534 | 0.469345 | 0.523720 | 0.395986 | 0.270448 | 0.220440 | 0.217052 | 0.227846 | 0.230307 | 0.222621 | 0.215628 | 0.217597 | 0.235787 | 0.274975 | 0.326703 | 0.372826 | 0.432044 | 0.499278 | 0.565638 | 0.651258 | 0.736958 | 0.746757 | 0.746602 | 0.694400 | 0.578347 | 0.504314 | 0.474118 | 0.490445 | 0.550915 | 0.622606 | 0.642612 | 0.618965 | 0.521482 | 0.419890 | 0.354540 | 0.300273 | 0.265540 | 0.242331 | 0.224938 | 0.215921 | 0.214749 | 0.216595 | 0.217150 | 0.217193 | 0.217461 | 0.216128 | 0.216116 | 0.216280 | 0.214917 | 0.214955 | 0.218503 | 0.243225 | 0.267815 | 0.275244 | 0.269697 | 0.246887 | 0.223647 | 0.215363 | 0.216070 | 0.219750 | 0.220751 | 0.219959 | 0.216759 | 0.214627 | 0.217265 | 0.226546 | 0.245026 | 0.281571 | 0.318193 | 0.340081 | 0.332790 | 0.328575 | 0.324889 | 0.340832 | 0.390716 | 0.506406 | 0.759872 | 1.250141 | 1.970931 | 2.697545 | 3.173756 | 2.640553 | 1.724728 | 1.096977 | 0.723145 | 0.544697 | 0.543234 | 0.732753 | 1.212887 | 2.351149 | 3.686797 | 4.105849 | 4.025395 | 3.301316 | 2.286651 | 1.714282 | 1.218189 | 0.864766 | 0.685323 | 0.582609 | 0.489468 | 0.460955 | 0.469229 | 0.519707 | 0.634943 | 0.856717 | 1.114192 | 1.481491 | 1.974686 | 2.442185 | 2.756226 | 2.893742 | 2.574724 | 2.040936 | 1.565941 | 1.406976 | 1.469525 | 1.938548 | 2.759678 | 4.293955 | 5.451203 | 4.665856 | 3.055904 | 1.853598 | 1.032095 | 0.650175 | 0.506739 | 0.452868 | 0.448417 | 0.453856 | 0.468559 | 0.457664 | 0.440109 | 0.400502 | 0.373342 | 0.353989 | 0.352772 | 0.335256 | 0.306760 | 0.274976 | 0.250688 | 0.221627 | 0.214992 | 0.226908 | 0.247259 | 0.269031 | 0.261538 | 0.241602 | 0.221106 | 0.214979 | 0.232238 | 0.276990 | 0.343779 | 0.413310 | 0.433719 | 0.411580 | 0.384338 | 0.356519 | 0.325952 | 0.300199 | 0.270384 | 0.242638 | 0.225189 | 0.216683 | 0.214830 | 0.214636 | 0.214957 | 0.218006 | 0.224385 | 0.226861 | 0.230494 | 0.228967 | 0.222920 | 0.217133 | 0.214817 | 0.214814 | 0.215847 | 0.217357 | 0.220246 | 0.224205 | 0.225738 | 0.227499 | 0.223005 | 0.217654 | 0.214774 | 0.216105 | 0.226751 | 0.241059 | 0.258276 | 0.275798 | 0.284928 | 0.289960 | 0.294182 | 0.299537 | 0.314604 | 0.354156 | 0.393223 | 0.424747 | 0.500608 | 0.554866 | 0.593033 | 0.617219 | 0.648615 | 0.653136 | 0.673615 | 0.641894 |
| right posterior | 4.252694 | 5.056882 | 5.478752 | 6.065012 | 3.997466 | 2.463840 | 1.349799 | 0.745471 | 0.425406 | 0.339431 | 0.332402 | 0.339122 | 0.404640 | 0.632528 | 0.957128 | 1.174560 | 1.122032 | 0.737358 | 0.465772 | 0.285581 | 0.216119 | 0.237687 | 0.285409 | 0.322346 | 0.310835 | 0.270230 | 0.227330 | 0.214626 | 0.219823 | 0.227973 | 0.235636 | 0.242450 | 0.243652 | 0.232760 | 0.228186 | 0.241104 | 0.262700 | 0.281094 | 0.318368 | 0.381237 | 0.465164 | 0.553611 | 0.687784 | 0.765231 | 0.893836 | 0.793753 | 0.721288 | 0.786911 | 1.107683 | 1.291928 | 1.134351 | 0.646236 | 0.444935 | 0.397440 | 0.441774 | 0.599757 | 1.083340 | 2.631500 | 7.273295 | 14.578284 | 19.099988 | 23.708265 | 18.829582 | 8.556316 | 2.774669 | 0.889710 | 0.383168 | 0.257344 | 0.224838 | 0.216537 | 0.217548 | 0.227626 | 0.241892 | 0.262478 | 0.301602 | 0.326715 | 0.314070 | 0.277487 | 0.235197 | 0.215973 | 0.216062 | 0.226719 | 0.238132 | 0.245362 | 0.247596 | 0.246692 | 0.235086 | 0.225830 | 0.221924 | 0.224628 | 0.234010 | 0.249870 | 0.266850 | 0.264648 | 0.248878 | 0.226831 | 0.215289 | 0.220116 | 0.236599 | 0.245000 | 0.234929 | 0.222734 | 0.214872 | 0.219386 | 0.235578 | 0.259167 | 0.290245 | 0.327965 | 0.357392 | 0.372978 | 0.375261 | 0.361189 | 0.343013 | 0.339197 | 0.331023 | 0.336612 | 0.357746 | 0.406252 | 0.467957 | 0.556695 | 0.621625 | 0.593127 | 0.446766 | 0.298022 | 0.225105 | 0.219113 | 0.246250 | 0.259813 | 0.244997 | 0.223247 | 0.214642 | 0.229008 | 0.265062 | 0.296112 | 0.318495 | 0.323025 | 0.305913 | 0.287316 | 0.267446 | 0.248836 | 0.232427 | 0.221184 | 0.216130 | 0.214774 | 0.215184 | 0.218048 | 0.222923 | 0.226057 | 0.224953 | 0.219675 | 0.215305 | 0.217479 | 0.240902 | 0.307091 | 0.449319 | 0.765161 | 1.322074 | 2.149268 | 3.139177 | 3.729563 | 3.262318 | 2.372916 | 1.387495 | 0.736037 | 0.460767 | 0.345282 | 0.290546 | 0.273928 | 0.283464 | 0.303191 | 0.330231 | 0.353380 | 0.372774 | 0.384855 | 0.375122 | 0.358199 | 0.328631 | 0.288657 | 0.250598 | 0.225817 | 0.214822 | 0.218372 | 0.231253 | 0.248154 | 0.256804 | 0.256829 | 0.249680 | 0.238428 | 0.231363 | 0.228907 | 0.232955 | 0.251111 | 0.273024 | 0.294794 | 0.293160 | 0.263741 | 0.227365 | 0.214737 | 0.224580 | 0.241102 | 0.254302 | 0.254500 | 0.245826 | 0.230963 | 0.221182 | 0.214772 | 0.217430 | 0.229309 | 0.254088 | 0.305077 | 0.407616 | 0.563465 | 0.667933 | 0.613649 | 0.468467 | 0.310885 | 0.226525 | 0.218267 | 0.256365 | 0.323159 | 0.365828 | 0.344660 | 0.286021 | 0.235423 | 0.215829 | 0.217817 | 0.232021 | 0.241997 | 0.242839 | 0.239865 | 0.231151 | 0.218504 | 0.214865 | 0.231028 | 0.279049 | 0.361177 | 0.451108 | 0.505836 | 0.488506 | 0.453519 | 0.410008 | 0.362780 | 0.315444 | 0.272430 | 0.237721 | 0.218882 | 0.215234 | 0.219958 | 0.219668 | 0.216140 | 0.214808 | 0.221476 | 0.236548 | 0.252845 | 0.262976 | 0.263446 | 0.266783 | 0.272021 | 0.287043 | 0.312922 | 0.341367 | 0.343322 | 0.334998 | 0.306910 | 0.286260 | 0.271294 | 0.263327 | 0.255052 | 0.251744 | 0.242909 | 0.234931 | 0.228119 | 0.220792 | 0.216833 | 0.215096 | 0.214639 | 0.215154 | 0.216792 | 0.218681 | 0.220608 |
| all electrodes | 0.221835 | 0.214653 | 0.225080 | 0.273338 | 0.343371 | 0.332301 | 0.284910 | 0.248052 | 0.220334 | 0.216029 | 0.220390 | 0.224776 | 0.228143 | 0.230710 | 0.230631 | 0.236171 | 0.249797 | 0.304178 | 0.368149 | 0.410474 | 0.489301 | 0.609094 | 0.520823 | 0.473551 | 0.444256 | 0.454532 | 0.564345 | 0.643498 | 0.600551 | 0.558140 | 0.489212 | 0.386218 | 0.382131 | 0.390697 | 0.407100 | 0.431222 | 0.455270 | 0.430054 | 0.404682 | 0.376665 | 0.320573 | 0.277895 | 0.251947 | 0.243382 | 0.240626 | 0.264633 | 0.317521 | 0.378910 | 0.341686 | 0.295521 | 0.242421 | 0.216360 | 0.229376 | 0.288860 | 0.401039 | 0.548923 | 0.921891 | 1.700703 | 3.538950 | 5.987340 | 7.337505 | 5.319213 | 3.766561 | 2.078736 | 1.017963 | 0.435230 | 0.242212 | 0.219272 | 0.277728 | 0.331724 | 0.312554 | 0.245203 | 0.214862 | 0.269249 | 0.420111 | 0.567293 | 0.568180 | 0.422261 | 0.290646 | 0.226027 | 0.216633 | 0.247881 | 0.317915 | 0.414130 | 0.532197 | 0.579175 | 0.559414 | 0.516330 | 0.463206 | 0.361229 | 0.315798 | 0.268979 | 0.236528 | 0.223473 | 0.222459 | 0.220185 | 0.220548 | 0.221006 | 0.223244 | 0.225566 | 0.224267 | 0.225859 | 0.229058 | 0.229679 | 0.229914 | 0.232261 | 0.232728 | 0.240712 | 0.251390 | 0.271633 | 0.323973 | 0.407877 | 0.516403 | 0.640255 | 0.779419 | 0.807337 | 0.776003 | 0.656973 | 0.576227 | 0.559540 | 0.575342 | 0.587150 | 0.694726 | 0.896353 | 1.206850 | 1.658812 | 2.077964 | 1.935535 | 1.618455 | 1.198824 | 0.877929 | 0.692045 | 0.522538 | 0.386094 | 0.318065 | 0.284734 | 0.254603 | 0.235847 | 0.223113 | 0.215215 | 0.219722 | 0.241754 | 0.289309 | 0.311058 | 0.313750 | 0.299874 | 0.280625 | 0.248370 | 0.232283 | 0.218790 | 0.214795 | 0.222199 | 0.235010 | 0.252250 | 0.253248 | 0.246574 | 0.235670 | 0.226587 | 0.219916 | 0.215716 | 0.214794 | 0.217087 | 0.223720 | 0.237747 | 0.263449 | 0.305649 | 0.381767 | 0.497434 | 0.597032 | 0.613762 | 0.580226 | 0.513067 | 0.436813 | 0.372388 | 0.326751 | 0.289527 | 0.259943 | 0.238913 | 0.228362 | 0.225100 | 0.229425 | 0.241384 | 0.278947 | 0.348520 | 0.445954 | 0.520023 | 0.589202 | 0.623727 | 0.642607 | 0.581376 | 0.551736 | 0.522815 | 0.484067 | 0.461899 | 0.476015 | 0.465385 | 0.472196 | 0.441186 | 0.367960 | 0.314791 | 0.280751 | 0.262302 | 0.261634 | 0.267781 | 0.281053 | 0.326906 | 0.372688 | 0.383796 | 0.377137 | 0.365090 | 0.332227 | 0.300916 | 0.286789 | 0.289848 | 0.286185 | 0.275022 | 0.263065 | 0.251708 | 0.235134 | 0.222032 | 0.216803 | 0.215428 | 0.215997 | 0.218225 | 0.222507 | 0.229597 | 0.239503 | 0.246125 | 0.259116 | 0.277081 | 0.282049 | 0.267860 | 0.239373 | 0.220917 | 0.214694 | 0.217229 | 0.223981 | 0.226754 | 0.227121 | 0.223730 | 0.217988 | 0.215293 | 0.214640 | 0.215687 | 0.218748 | 0.220220 | 0.220094 | 0.217513 | 0.215532 | 0.214749 | 0.216123 | 0.216893 | 0.214881 | 0.215941 | 0.228648 | 0.263167 | 0.338419 | 0.442207 | 0.550629 | 0.627553 | 0.632618 | 0.503858 | 0.415346 | 0.355754 | 0.326448 | 0.325174 | 0.383081 | 0.457898 | 0.650909 | 0.956965 | 1.153914 | 1.263215 | 1.517563 | 1.457999 | 1.509844 | 1.509468 | 1.412264 | 1.082732 | 0.961130 | 0.806941 |

R) maximum cross-correlation, happy vs sad

  
|  | time window | peak latency | cluster *p* | peak Cohen's *d* |  | | | |  |
| **all electrodes** |  | | | |  | | | |  |
|  | | | | | | | | |  |

Model correlations, cluster permutation tests

|  | **left hemisphere** | | | | **right hemisphere** | | | |  |
|  | time window | peak latency | cluster *p* | peak Cohen's *d* | time window | peak latency | cluster *p* | peak Cohen's *d* |  |
| **anterior** |  | | | |  | | | |  |
| **central** | 250 - 385 ms | 270 ms | 0.0234 | -0.7067 |  | | | |  |
 560 - 725 ms | 690 ms | 0.015 | -0.7141 |  | | | ||  | **posterior** |  | | | |  | | | |

  

Model correlations, Bayesian statistics

|  | -200 | -195 | -190 | -185 | -180 | -175 | -170 | -165 | -160 | -155 | -150 | -145 | -140 | -135 | -130 | -125 | -120 | -115 | -110 | -105 | -100 | -95 | -90 | -85 | -80 | -75 | -70 | -65 | -60 | -55 | -50 | -45 | -40 | -35 | -30 | -25 | -20 | -15 | -10 | -5 | 0 | 5 | 10 | 15 | 20 | 25 | 30 | 35 | 40 | 45 | 50 | 55 | 60 | 65 | 70 | 75 | 80 | 85 | 90 | 95 | 100 | 105 | 110 | 115 | 120 | 125 | 130 | 135 | 140 | 145 | 150 | 155 | 160 | 165 | 170 | 175 | 180 | 185 | 190 | 195 | 200 | 205 | 210 | 215 | 220 | 225 | 230 | 235 | 240 | 245 | 250 | 255 | 260 | 265 | 270 | 275 | 280 | 285 | 290 | 295 | 300 | 305 | 310 | 315 | 320 | 325 | 330 | 335 | 340 | 345 | 350 | 355 | 360 | 365 | 370 | 375 | 380 | 385 | 390 | 395 | 400 | 405 | 410 | 415 | 420 | 425 | 430 | 435 | 440 | 445 | 450 | 455 | 460 | 465 | 470 | 475 | 480 | 485 | 490 | 495 | 500 | 505 | 510 | 515 | 520 | 525 | 530 | 535 | 540 | 545 | 550 | 555 | 560 | 565 | 570 | 575 | 580 | 585 | 590 | 595 | 600 | 605 | 610 | 615 | 620 | 625 | 630 | 635 | 640 | 645 | 650 | 655 | 660 | 665 | 670 | 675 | 680 | 685 | 690 | 695 | 700 | 705 | 710 | 715 | 720 | 725 | 730 | 735 | 740 | 745 | 750 | 755 | 760 | 765 | 770 | 775 | 780 | 785 | 790 | 795 | 800 | 805 | 810 | 815 | 820 | 825 | 830 | 835 | 840 | 845 | 850 | 855 | 860 | 865 | 870 | 875 | 880 | 885 | 890 | 895 | 900 | 905 | 910 | 915 | 920 | 925 | 930 | 935 | 940 | 945 | 950 | 955 | 960 | 965 | 970 | 975 | 980 | 985 | 990 | 995 | 1000 | 1005 | 1010 | 1015 | 1020 | 1025 | 1030 | 1035 | 1040 | 1045 | 1050 | 1055 | 1060 | 1065 | 1070 | 1075 | 1080 | 1085 | 1090 | 1095 | 1100 | 1105 | 1110 | 1115 | 1120 | 1125 | 1130 | 1135 | 1140 | 1145 | 1150 | 1155 | 1160 | 1165 | 1170 | 1175 | 1180 | 1185 | 1190 | 1195 |
| --- | --- | --- | --- | --- | --- | --- | --- | --- | --- | --- | --- | --- | --- | --- | --- | --- | --- | --- | --- | --- | --- | --- | --- | --- | --- | --- | --- | --- | --- | --- | --- | --- | --- | --- | --- | --- | --- | --- | --- | --- | --- | --- | --- | --- | --- | --- | --- | --- | --- | --- | --- | --- | --- | --- | --- | --- | --- | --- | --- | --- | --- | --- | --- | --- | --- | --- | --- | --- | --- | --- | --- | --- | --- | --- | --- | --- | --- | --- | --- | --- | --- | --- | --- | --- | --- | --- | --- | --- | --- | --- | --- | --- | --- | --- | --- | --- | --- | --- | --- | --- | --- | --- | --- | --- | --- | --- | --- | --- | --- | --- | --- | --- | --- | --- | --- | --- | --- | --- | --- | --- | --- | --- | --- | --- | --- | --- | --- | --- | --- | --- | --- | --- | --- | --- | --- | --- | --- | --- | --- | --- | --- | --- | --- | --- | --- | --- | --- | --- | --- | --- | --- | --- | --- | --- | --- | --- | --- | --- | --- | --- | --- | --- | --- | --- | --- | --- | --- | --- | --- | --- | --- | --- | --- | --- | --- | --- | --- | --- | --- | --- | --- | --- | --- | --- | --- | --- | --- | --- | --- | --- | --- | --- | --- | --- | --- | --- | --- | --- | --- | --- | --- | --- | --- | --- | --- | --- | --- | --- | --- | --- | --- | --- | --- | --- | --- | --- | --- | --- | --- | --- | --- | --- | --- | --- | --- | --- | --- | --- | --- | --- | --- | --- | --- | --- | --- | --- | --- | --- | --- | --- | --- | --- | --- | --- | --- | --- | --- | --- | --- | --- | --- | --- | --- | --- | --- | --- | --- | --- | --- | --- | --- | --- | --- | --- | --- | --- | --- | --- | --- | --- | --- | --- | --- | --- | --- | --- | --- | --- | --- | --- |
| left anterior | 0.215764 | 0.214939 | 0.216330 | 0.240947 | 0.280084 | 0.306647 | 0.294299 | 0.261112 | 0.217506 | 0.226075 | 0.264031 | 0.293079 | 0.310157 | 0.269945 | 0.231401 | 0.215184 | 0.235643 | 0.266953 | 0.303034 | 0.356594 | 0.364696 | 0.354220 | 0.360480 | 0.448828 | 0.630918 | 0.889710 | 1.504519 | 2.527615 | 3.442485 | 4.706268 | 4.269887 | 2.082885 | 0.894452 | 0.336276 | 0.215505 | 0.248476 | 0.357100 | 0.374868 | 0.325987 | 0.331917 | 0.362111 | 0.387765 | 0.441519 | 0.424209 | 0.344604 | 0.241935 | 0.218701 | 0.320202 | 0.571150 | 1.050963 | 1.349447 | 1.372650 | 0.820136 | 0.406025 | 0.228781 | 0.222337 | 0.277184 | 0.315091 | 0.364438 | 0.404518 | 0.404786 | 0.335693 | 0.277716 | 0.228365 | 0.215675 | 0.237143 | 0.270495 | 0.326589 | 0.451308 | 0.553462 | 0.809729 | 1.403233 | 2.563914 | 3.978336 | 5.410050 | 3.935448 | 2.321710 | 1.235632 | 0.580243 | 0.330197 | 0.242350 | 0.216814 | 0.214699 | 0.214760 | 0.216960 | 0.226204 | 0.239123 | 0.251349 | 0.254588 | 0.255854 | 0.241930 | 0.225496 | 0.215121 | 0.214749 | 0.215291 | 0.217666 | 0.223950 | 0.234135 | 0.273317 | 0.349667 | 0.447661 | 0.522151 | 0.538643 | 0.468316 | 0.348252 | 0.271338 | 0.224307 | 0.214650 | 0.220112 | 0.223752 | 0.222890 | 0.216468 | 0.214714 | 0.214633 | 0.214634 | 0.216776 | 0.239716 | 0.292111 | 0.370556 | 0.446693 | 0.426149 | 0.320633 | 0.237868 | 0.220274 | 0.329225 | 0.575500 | 0.804632 | 0.779956 | 0.436153 | 0.251960 | 0.218153 | 0.269286 | 0.345739 | 0.426704 | 0.502636 | 0.512578 | 0.472391 | 0.384767 | 0.314007 | 0.267771 | 0.247479 | 0.243346 | 0.249418 | 0.253133 | 0.268532 | 0.300513 | 0.352377 | 0.393667 | 0.408498 | 0.381027 | 0.347122 | 0.300889 | 0.258543 | 0.227698 | 0.215818 | 0.215780 | 0.220167 | 0.219757 | 0.215748 | 0.214885 | 0.216722 | 0.215485 | 0.214644 | 0.215678 | 0.218988 | 0.225794 | 0.235967 | 0.241197 | 0.246369 | 0.264101 | 0.296987 | 0.343383 | 0.400727 | 0.451111 | 0.486780 | 0.500846 | 0.450210 | 0.409500 | 0.383539 | 0.343489 | 0.289669 | 0.260875 | 0.248047 | 0.249241 | 0.265731 | 0.306396 | 0.365918 | 0.458540 | 0.553235 | 0.660886 | 0.740588 | 0.878009 | 0.819237 | 0.628211 | 0.392979 | 0.255475 | 0.216803 | 0.300375 | 0.452847 | 0.528796 | 0.525411 | 0.422134 | 0.322362 | 0.261517 | 0.228958 | 0.215558 | 0.215378 | 0.220348 | 0.226350 | 0.231230 | 0.231614 | 0.228431 | 0.219707 | 0.214628 | 0.224992 | 0.266895 | 0.354022 | 0.469985 | 0.552964 | 0.526168 | 0.445248 | 0.384498 | 0.339447 | 0.308947 | 0.299768 | 0.291173 | 0.276994 | 0.257284 | 0.240910 | 0.228744 | 0.224802 | 0.223764 | 0.225197 | 0.229760 | 0.238818 | 0.238098 | 0.231138 | 0.225120 | 0.217492 | 0.214644 | 0.215687 | 0.216150 | 0.216670 | 0.215125 | 0.216099 | 0.225802 | 0.247680 | 0.282805 | 0.332123 | 0.373016 | 0.419595 | 0.438052 | 0.448590 | 0.449170 | 0.431797 | 0.385838 | 0.374578 | 0.365407 | 0.355243 | 0.349368 | 0.357980 | 0.370778 | 0.398138 | 0.427045 | 0.446580 | 0.465223 | 0.476728 | 0.470690 | 0.461363 | 0.443204 | 0.413347 | 0.369854 | 0.334242 | 0.304309 | 0.294214 | 0.287748 | 0.292637 | 0.307637 | 0.317398 | 0.322869 |
| right anterior | 0.388622 | 0.466345 | 0.440371 | 0.437438 | 0.446744 | 0.391956 | 0.377090 | 0.367231 | 0.364454 | 0.334196 | 0.253583 | 0.215037 | 0.220743 | 0.257719 | 0.284040 | 0.243357 | 0.214627 | 0.292272 | 0.936790 | 4.041301 | 15.938862 | 39.471329 | 39.028531 | 21.390674 | 8.146330 | 3.522212 | 1.668231 | 0.694835 | 0.337127 | 0.230432 | 0.227077 | 0.323779 | 0.415958 | 0.423558 | 0.388037 | 0.360464 | 0.368064 | 0.414654 | 0.469704 | 0.621811 | 0.863714 | 0.857139 | 0.635899 | 0.471656 | 0.331436 | 0.271322 | 0.239512 | 0.224003 | 0.215865 | 0.215299 | 0.221728 | 0.227773 | 0.231613 | 0.232596 | 0.224056 | 0.218743 | 0.215987 | 0.214678 | 0.217315 | 0.219318 | 0.220317 | 0.217549 | 0.214627 | 0.221557 | 0.235265 | 0.248310 | 0.262526 | 0.266452 | 0.251578 | 0.242239 | 0.239125 | 0.246573 | 0.263805 | 0.284328 | 0.307172 | 0.331128 | 0.333539 | 0.317267 | 0.297345 | 0.277948 | 0.269232 | 0.265991 | 0.279557 | 0.327767 | 0.424324 | 0.570841 | 0.717820 | 0.734395 | 0.670659 | 0.588589 | 0.510259 | 0.428141 | 0.377562 | 0.334910 | 0.303473 | 0.266447 | 0.244362 | 0.228499 | 0.223656 | 0.219803 | 0.222491 | 0.229409 | 0.244545 | 0.256444 | 0.253118 | 0.227209 | 0.214790 | 0.220845 | 0.236112 | 0.240025 | 0.227620 | 0.216038 | 0.217082 | 0.233629 | 0.256718 | 0.288950 | 0.323826 | 0.342372 | 0.318744 | 0.289350 | 0.253229 | 0.224029 | 0.215079 | 0.229947 | 0.251877 | 0.275471 | 0.293075 | 0.318505 | 0.323274 | 0.341686 | 0.363897 | 0.387647 | 0.334420 | 0.272994 | 0.226266 | 0.214638 | 0.223532 | 0.241213 | 0.261064 | 0.258472 | 0.249946 | 0.244963 | 0.242867 | 0.241679 | 0.248136 | 0.258381 | 0.274742 | 0.301674 | 0.321402 | 0.348266 | 0.364844 | 0.353416 | 0.294290 | 0.247543 | 0.218453 | 0.217068 | 0.237681 | 0.252730 | 0.254658 | 0.247453 | 0.235600 | 0.224309 | 0.220697 | 0.223776 | 0.233003 | 0.249775 | 0.283581 | 0.332726 | 0.404838 | 0.491593 | 0.566588 | 0.597348 | 0.521641 | 0.390018 | 0.303453 | 0.252791 | 0.228158 | 0.219250 | 0.218599 | 0.220275 | 0.231589 | 0.260581 | 0.344818 | 0.432417 | 0.474423 | 0.403217 | 0.326205 | 0.263709 | 0.235258 | 0.226398 | 0.232706 | 0.252658 | 0.292211 | 0.358196 | 0.462301 | 0.568118 | 0.579089 | 0.497088 | 0.419365 | 0.372691 | 0.328981 | 0.301929 | 0.294155 | 0.290216 | 0.256333 | 0.229354 | 0.217434 | 0.214755 | 0.215343 | 0.215541 | 0.214696 | 0.219571 | 0.229525 | 0.238936 | 0.242803 | 0.240949 | 0.238492 | 0.229325 | 0.224896 | 0.222807 | 0.221426 | 0.219064 | 0.217975 | 0.216264 | 0.216293 | 0.217604 | 0.218355 | 0.217217 | 0.214661 | 0.226142 | 0.295279 | 0.524312 | 1.062330 | 2.135876 | 3.470726 | 4.059807 | 3.236733 | 2.111978 | 1.197042 | 0.756960 | 0.534571 | 0.401015 | 0.324243 | 0.278088 | 0.253976 | 0.242010 | 0.239735 | 0.243853 | 0.257167 | 0.270938 | 0.286869 | 0.304250 | 0.330105 | 0.352152 | 0.354747 | 0.353059 | 0.358862 | 0.353872 | 0.328165 | 0.313241 | 0.298690 | 0.285570 | 0.274256 | 0.276152 | 0.283875 | 0.297318 | 0.300692 | 0.287962 | 0.265564 | 0.242286 | 0.222734 | 0.215906 | 0.214798 | 0.214873 | 0.217803 | 0.220905 | 0.226269 | 0.235865 | 0.241161 | 0.240402 |
| left central | 0.230464 | 0.250912 | 0.276354 | 0.332786 | 0.352132 | 0.392420 | 0.391975 | 0.351974 | 0.309357 | 0.288488 | 0.269709 | 0.265835 | 0.272513 | 0.275959 | 0.268041 | 0.250068 | 0.240813 | 0.242988 | 0.259856 | 0.259527 | 0.260212 | 0.251201 | 0.241620 | 0.233958 | 0.239603 | 0.232696 | 0.224947 | 0.216543 | 0.215224 | 0.233240 | 0.265345 | 0.375016 | 0.558812 | 0.723541 | 0.601250 | 0.437255 | 0.280924 | 0.231593 | 0.222354 | 0.222194 | 0.229729 | 0.254228 | 0.296135 | 0.332539 | 0.366041 | 0.373171 | 0.346141 | 0.296632 | 0.256527 | 0.233088 | 0.233893 | 0.255331 | 0.295621 | 0.353308 | 0.426540 | 0.472151 | 0.508280 | 0.457034 | 0.393221 | 0.339481 | 0.303291 | 0.277263 | 0.276968 | 0.285778 | 0.314673 | 0.333625 | 0.341335 | 0.337929 | 0.293670 | 0.234882 | 0.214657 | 0.239226 | 0.293890 | 0.343530 | 0.360246 | 0.328369 | 0.292599 | 0.281718 | 0.290384 | 0.318215 | 0.390169 | 0.440241 | 0.482208 | 0.465172 | 0.401920 | 0.286525 | 0.219906 | 0.245831 | 0.416212 | 1.026539 | 2.582512 | 6.785427 | 11.836902 | 18.366300 | 17.064692 | 9.081885 | 5.680064 | 4.212330 | 4.329236 | 5.226193 | 6.739589 | 9.093517 | 13.164420 | 12.680084 | 12.147142 | 10.414656 | 4.817840 | 2.631903 | 2.132995 | 1.777099 | 1.668530 | 1.868863 | 1.967085 | 3.032245 | 4.988201 | 5.934249 | 4.427665 | 2.083733 | 0.654909 | 0.316390 | 0.232995 | 0.214713 | 0.223240 | 0.257558 | 0.301298 | 0.360435 | 0.396835 | 0.430501 | 0.449407 | 0.449381 | 0.399479 | 0.384627 | 0.359341 | 0.326525 | 0.294015 | 0.259230 | 0.230575 | 0.216947 | 0.217361 | 0.242240 | 0.307382 | 0.429633 | 0.661933 | 1.003378 | 1.373048 | 1.523977 | 1.313820 | 1.065117 | 0.952303 | 0.946454 | 0.931189 | 1.116854 | 1.489544 | 2.020182 | 2.769131 | 4.342679 | 8.166994 | 13.000062 | 15.203991 | 11.992834 | 7.611315 | 4.778186 | 3.664711 | 2.650802 | 2.241210 | 2.198544 | 2.071193 | 1.974920 | 1.969494 | 2.111965 | 2.408968 | 2.696017 | 3.090208 | 3.836943 | 4.414327 | 5.176117 | 7.524391 | 11.806606 | 17.412446 | 21.008699 | 18.424129 | 12.138836 | 7.177810 | 4.255773 | 2.483613 | 1.605851 | 1.081782 | 0.823579 | 0.715955 | 0.671324 | 0.666675 | 0.701329 | 0.859569 | 1.118109 | 1.636669 | 2.684684 | 4.604642 | 6.047576 | 5.298646 | 3.520599 | 1.717341 | 0.842809 | 0.509362 | 0.399945 | 0.341339 | 0.319289 | 0.318747 | 0.315286 | 0.308606 | 0.308300 | 0.306785 | 0.305175 | 0.317852 | 0.320986 | 0.333101 | 0.355973 | 0.394392 | 0.443519 | 0.512147 | 0.529369 | 0.526296 | 0.473037 | 0.408754 | 0.343754 | 0.305145 | 0.278059 | 0.266628 | 0.260533 | 0.261253 | 0.259050 | 0.249915 | 0.240248 | 0.236159 | 0.235244 | 0.245226 | 0.281324 | 0.374191 | 0.529090 | 0.691927 | 0.772125 | 0.777523 | 0.691778 | 0.576819 | 0.458000 | 0.386243 | 0.340360 | 0.317768 | 0.315730 | 0.341626 | 0.389988 | 0.479826 | 0.629373 | 0.875144 | 1.208890 | 1.434894 | 1.396803 | 1.208124 | 0.866233 | 0.720777 | 0.674672 | 0.697494 | 0.792782 | 1.058921 | 1.415364 | 1.936357 | 2.387168 | 2.614920 | 3.158807 | 3.855302 | 5.388334 | 9.286735 | 18.220397 | 27.142389 | 33.511125 | 28.016159 | 17.703803 | 9.475443 | 4.984100 | 3.440119 | 2.594156 |
| right central | 0.430766 | 0.381785 | 0.326979 | 0.264008 | 0.228223 | 0.214960 | 0.228409 | 0.241852 | 0.240275 | 0.229533 | 0.220065 | 0.217141 | 0.217150 | 0.217935 | 0.218688 | 0.217694 | 0.222462 | 0.237584 | 0.264115 | 0.263163 | 0.265007 | 0.263854 | 0.257561 | 0.250606 | 0.248897 | 0.250491 | 0.240107 | 0.227145 | 0.217242 | 0.216673 | 0.217550 | 0.226485 | 0.230671 | 0.258176 | 0.285457 | 0.257640 | 0.224919 | 0.214632 | 0.239351 | 0.280877 | 0.309800 | 0.363115 | 0.340235 | 0.260868 | 0.220357 | 0.217824 | 0.250895 | 0.292119 | 0.330929 | 0.313274 | 0.282593 | 0.251598 | 0.232560 | 0.216448 | 0.221664 | 0.290212 | 0.494404 | 0.930087 | 1.381250 | 1.285895 | 0.987275 | 0.696518 | 0.546449 | 0.480696 | 0.521012 | 0.674288 | 1.049929 | 1.718921 | 2.108836 | 1.233725 | 0.448717 | 0.225882 | 0.237949 | 0.318532 | 0.398542 | 0.431278 | 0.432762 | 0.444668 | 0.462678 | 0.489803 | 0.602009 | 0.803732 | 1.339452 | 2.722400 | 5.818232 | 8.692901 | 7.069724 | 3.154048 | 1.497659 | 0.751344 | 0.502628 | 0.385221 | 0.316983 | 0.269891 | 0.260819 | 0.252567 | 0.244553 | 0.251039 | 0.274792 | 0.299286 | 0.358083 | 0.442649 | 0.489169 | 0.465865 | 0.371470 | 0.286123 | 0.244681 | 0.232442 | 0.233774 | 0.252086 | 0.275471 | 0.309384 | 0.318194 | 0.298209 | 0.255404 | 0.225365 | 0.214646 | 0.218986 | 0.222020 | 0.220239 | 0.215513 | 0.215647 | 0.223369 | 0.238996 | 0.260704 | 0.282872 | 0.305013 | 0.323865 | 0.305121 | 0.285180 | 0.266671 | 0.261230 | 0.246560 | 0.232262 | 0.224322 | 0.221821 | 0.217002 | 0.216082 | 0.215901 | 0.215774 | 0.215920 | 0.217833 | 0.219977 | 0.226529 | 0.235195 | 0.246080 | 0.254891 | 0.265900 | 0.282242 | 0.303245 | 0.314611 | 0.305980 | 0.274096 | 0.237908 | 0.215638 | 0.225885 | 0.267118 | 0.315353 | 0.322428 | 0.279695 | 0.227890 | 0.215705 | 0.250272 | 0.311466 | 0.373325 | 0.398377 | 0.404760 | 0.386301 | 0.346071 | 0.298600 | 0.264510 | 0.240925 | 0.230474 | 0.226502 | 0.226922 | 0.225478 | 0.224668 | 0.219046 | 0.215381 | 0.214657 | 0.214849 | 0.214727 | 0.216301 | 0.223513 | 0.239876 | 0.267755 | 0.310399 | 0.367319 | 0.432924 | 0.478049 | 0.496965 | 0.481593 | 0.427840 | 0.384093 | 0.333312 | 0.299776 | 0.280190 | 0.262501 | 0.250472 | 0.239318 | 0.226631 | 0.217421 | 0.214666 | 0.215654 | 0.217135 | 0.220413 | 0.224237 | 0.227430 | 0.226563 | 0.223954 | 0.218459 | 0.215227 | 0.214628 | 0.214656 | 0.214993 | 0.217045 | 0.225108 | 0.241648 | 0.273579 | 0.288654 | 0.292104 | 0.281157 | 0.267380 | 0.238541 | 0.224618 | 0.217496 | 0.216154 | 0.215425 | 0.216721 | 0.220369 | 0.228917 | 0.237417 | 0.254878 | 0.275860 | 0.300137 | 0.335688 | 0.388072 | 0.486051 | 0.676006 | 1.010684 | 1.378964 | 1.444111 | 1.292510 | 1.025465 | 0.778740 | 0.539978 | 0.384681 | 0.303747 | 0.266004 | 0.239188 | 0.230787 | 0.234086 | 0.234879 | 0.236845 | 0.240117 | 0.239100 | 0.232715 | 0.223105 | 0.215630 | 0.214861 | 0.217432 | 0.219702 | 0.217184 | 0.215181 | 0.214667 | 0.215739 | 0.218279 | 0.229177 | 0.258126 | 0.317889 | 0.443806 | 0.701261 | 1.081788 | 1.568082 | 2.070383 | 2.457007 | 2.519686 | 2.419234 | 2.198944 | 1.989687 |
| left posterior | 0.547671 | 0.666886 | 0.870227 | 1.342945 | 1.362424 | 0.918849 | 0.492017 | 0.282097 | 0.215357 | 0.236182 | 0.282129 | 0.337174 | 0.369050 | 0.324947 | 0.261281 | 0.214929 | 0.311625 | 1.723961 | 7.655528 | 16.958650 | 20.130444 | 18.819091 | 13.247606 | 12.024453 | 4.657125 | 1.676358 | 0.691259 | 0.426322 | 0.289667 | 0.246685 | 0.225908 | 0.220995 | 0.217029 | 0.218645 | 0.217651 | 0.216564 | 0.214683 | 0.214688 | 0.214684 | 0.218353 | 0.229280 | 0.247999 | 0.266055 | 0.237218 | 0.214629 | 0.239766 | 0.290654 | 0.383143 | 0.454361 | 0.440853 | 0.348909 | 0.264668 | 0.217152 | 0.223459 | 0.262650 | 0.267581 | 0.239770 | 0.215367 | 0.232092 | 0.313021 | 0.354221 | 0.366007 | 0.372894 | 0.347960 | 0.321653 | 0.304092 | 0.310005 | 0.334592 | 0.480498 | 1.030364 | 3.379271 | 15.347421 | 71.120358 | 213.361049 | 364.436211 | 344.119083 | 151.133452 | 56.653793 | 13.890110 | 2.943389 | 0.990956 | 0.513255 | 0.360143 | 0.304361 | 0.264914 | 0.235484 | 0.223238 | 0.215604 | 0.214768 | 0.215035 | 0.214815 | 0.214638 | 0.215190 | 0.215939 | 0.215632 | 0.215889 | 0.215413 | 0.214717 | 0.214661 | 0.214671 | 0.214882 | 0.216272 | 0.222888 | 0.243653 | 0.275110 | 0.304487 | 0.306635 | 0.281507 | 0.236840 | 0.214771 | 0.227218 | 0.268997 | 0.339400 | 0.416959 | 0.462832 | 0.440857 | 0.361018 | 0.279082 | 0.226494 | 0.215761 | 0.240084 | 0.283178 | 0.316983 | 0.321608 | 0.293754 | 0.250855 | 0.224157 | 0.214626 | 0.223479 | 0.244082 | 0.246594 | 0.234220 | 0.219301 | 0.214702 | 0.224215 | 0.240930 | 0.253177 | 0.252177 | 0.239305 | 0.222227 | 0.214825 | 0.217015 | 0.221189 | 0.221885 | 0.218960 | 0.215802 | 0.214630 | 0.214646 | 0.216397 | 0.230185 | 0.263681 | 0.334673 | 0.445065 | 0.550524 | 0.643533 | 0.704203 | 0.693396 | 0.609333 | 0.524447 | 0.438478 | 0.400126 | 0.374612 | 0.355941 | 0.342595 | 0.340975 | 0.338319 | 0.367393 | 0.418029 | 0.492934 | 0.568310 | 0.688883 | 0.756909 | 0.762749 | 0.651412 | 0.519564 | 0.400825 | 0.326110 | 0.272235 | 0.242681 | 0.225689 | 0.217154 | 0.214692 | 0.218746 | 0.228630 | 0.244409 | 0.262303 | 0.273702 | 0.280227 | 0.280335 | 0.273563 | 0.266927 | 0.257739 | 0.252932 | 0.254144 | 0.258551 | 0.266994 | 0.288859 | 0.310600 | 0.338654 | 0.382196 | 0.435072 | 0.482911 | 0.521596 | 0.547648 | 0.530114 | 0.489065 | 0.430574 | 0.396626 | 0.370384 | 0.360065 | 0.357476 | 0.369580 | 0.391722 | 0.444691 | 0.506796 | 0.611562 | 0.745040 | 0.828751 | 0.765346 | 0.642388 | 0.503379 | 0.435942 | 0.403293 | 0.379901 | 0.378870 | 0.393936 | 0.385036 | 0.372934 | 0.346707 | 0.318538 | 0.291687 | 0.281280 | 0.279359 | 0.300615 | 0.347943 | 0.433507 | 0.525546 | 0.565713 | 0.490072 | 0.380592 | 0.306340 | 0.271454 | 0.261972 | 0.272149 | 0.304488 | 0.370785 | 0.489447 | 0.690356 | 0.974217 | 1.261456 | 1.533970 | 1.687882 | 1.473583 | 1.087956 | 0.754543 | 0.509586 | 0.354292 | 0.288528 | 0.256119 | 0.238202 | 0.226791 | 0.221972 | 0.219087 | 0.218822 | 0.219985 | 0.224549 | 0.235722 | 0.256097 | 0.287465 | 0.331313 | 0.380741 | 0.427627 | 0.471556 | 0.540164 | 0.662650 | 0.836317 | 1.014069 | 1.233574 | 1.319310 | 1.368937 |
| right posterior | 5.641869 | 3.521452 | 2.070757 | 0.853141 | 0.383223 | 0.249215 | 0.215109 | 0.222096 | 0.259078 | 0.288263 | 0.287022 | 0.258725 | 0.225627 | 0.216214 | 0.279299 | 0.421384 | 0.555642 | 0.501418 | 0.389768 | 0.249675 | 0.214626 | 0.250182 | 0.297818 | 0.286803 | 0.218983 | 0.247537 | 0.480405 | 0.996968 | 1.092196 | 0.550871 | 0.307663 | 0.226744 | 0.214817 | 0.225326 | 0.255213 | 0.286193 | 0.285694 | 0.260940 | 0.244533 | 0.224648 | 0.218629 | 0.215309 | 0.214711 | 0.214692 | 0.214742 | 0.215527 | 0.216249 | 0.227991 | 0.271422 | 0.359805 | 0.500477 | 0.665307 | 0.750861 | 0.785561 | 0.733545 | 0.738227 | 0.843197 | 1.204131 | 1.619643 | 1.679644 | 1.042593 | 0.527598 | 0.316249 | 0.243347 | 0.225926 | 0.224214 | 0.228875 | 0.226186 | 0.216453 | 0.224931 | 0.303431 | 0.481965 | 0.718681 | 0.899560 | 0.866396 | 0.681445 | 0.482274 | 0.325406 | 0.237523 | 0.215089 | 0.243099 | 0.286292 | 0.325005 | 0.349886 | 0.371048 | 0.422604 | 0.569099 | 0.773560 | 0.931981 | 0.982259 | 0.760320 | 0.546563 | 0.438450 | 0.415297 | 0.411939 | 0.466516 | 0.548172 | 0.674907 | 0.793316 | 0.856278 | 0.796369 | 0.698173 | 0.562869 | 0.419766 | 0.336673 | 0.308258 | 0.320151 | 0.359714 | 0.476663 | 0.750740 | 1.316660 | 2.285490 | 3.031620 | 2.407515 | 1.606646 | 0.960612 | 0.558122 | 0.371967 | 0.275569 | 0.232489 | 0.217945 | 0.214996 | 0.214626 | 0.214909 | 0.215913 | 0.220830 | 0.234388 | 0.258252 | 0.313554 | 0.456365 | 0.789825 | 1.904994 | 7.158101 | 27.085936 | 83.933105 | 144.708584 | 120.890938 | 75.927537 | 37.390905 | 16.425015 | 7.495004 | 3.638663 | 1.874239 | 1.136409 | 0.746402 | 0.527102 | 0.428691 | 0.388530 | 0.357833 | 0.352709 | 0.357143 | 0.396990 | 0.448788 | 0.526706 | 0.606987 | 0.657924 | 0.532319 | 0.403697 | 0.303269 | 0.251365 | 0.232329 | 0.229373 | 0.233210 | 0.244439 | 0.259466 | 0.275870 | 0.294734 | 0.323969 | 0.394063 | 0.532257 | 0.746889 | 1.076477 | 1.375639 | 1.365201 | 1.154930 | 0.921235 | 0.727994 | 0.657426 | 0.635785 | 0.643379 | 0.668777 | 0.558817 | 0.419675 | 0.319714 | 0.258056 | 0.232186 | 0.224026 | 0.220206 | 0.225317 | 0.241382 | 0.259663 | 0.285910 | 0.305312 | 0.295024 | 0.275771 | 0.259294 | 0.241406 | 0.232152 | 0.223412 | 0.216971 | 0.214671 | 0.218220 | 0.228640 | 0.243284 | 0.264488 | 0.294692 | 0.348151 | 0.431020 | 0.555509 | 0.680844 | 0.722255 | 0.624168 | 0.462774 | 0.337938 | 0.261338 | 0.224605 | 0.214723 | 0.216821 | 0.218477 | 0.215305 | 0.216731 | 0.229913 | 0.245911 | 0.249319 | 0.243253 | 0.231229 | 0.219857 | 0.215180 | 0.215019 | 0.218680 | 0.221261 | 0.219265 | 0.214750 | 0.218622 | 0.236687 | 0.287629 | 0.386257 | 0.489159 | 0.591900 | 0.662763 | 0.653751 | 0.630248 | 0.589888 | 0.581545 | 0.631823 | 0.656025 | 0.717056 | 0.873229 | 0.962422 | 0.931364 | 0.806081 | 0.595515 | 0.417080 | 0.305559 | 0.248697 | 0.226803 | 0.217116 | 0.214787 | 0.218916 | 0.226571 | 0.236601 | 0.249324 | 0.263101 | 0.285612 | 0.315300 | 0.369227 | 0.449303 | 0.517923 | 0.536169 | 0.529201 | 0.462520 | 0.386293 | 0.322037 | 0.267316 | 0.233920 | 0.221006 | 0.215742 | 0.214645 | 0.214643 | 0.214686 |
| all electrodes | 0.241112 | 0.235945 | 0.220870 | 0.217911 | 0.235957 | 0.263706 | 0.326945 | 0.370710 | 0.439243 | 0.656780 | 1.032861 | 1.498570 | 1.779324 | 1.775635 | 1.374415 | 1.174917 | 1.088209 | 0.987257 | 0.836010 | 0.611249 | 0.414447 | 0.340954 | 0.321749 | 0.300468 | 0.327384 | 0.468135 | 0.915692 | 1.307746 | 1.460893 | 1.371646 | 1.299853 | 1.041515 | 0.861683 | 0.592094 | 0.435897 | 0.326561 | 0.271994 | 0.244026 | 0.225517 | 0.215067 | 0.240041 | 0.318243 | 0.469709 | 0.694553 | 0.661888 | 0.489189 | 0.309182 | 0.229440 | 0.215037 | 0.221755 | 0.220685 | 0.214704 | 0.233185 | 0.313331 | 0.479135 | 0.670970 | 0.689412 | 0.515974 | 0.391741 | 0.292708 | 0.245429 | 0.224586 | 0.217014 | 0.214965 | 0.214743 | 0.214972 | 0.214699 | 0.223478 | 0.277738 | 0.452167 | 1.037596 | 2.458283 | 4.577039 | 5.427384 | 4.610118 | 2.490376 | 1.278799 | 0.763567 | 0.521484 | 0.451114 | 0.471123 | 0.528543 | 0.573589 | 0.627896 | 0.687160 | 0.756553 | 0.711374 | 0.492018 | 0.312320 | 0.238085 | 0.216014 | 0.216429 | 0.226661 | 0.232955 | 0.238437 | 0.242423 | 0.261029 | 0.308206 | 0.405968 | 0.563902 | 0.812079 | 0.929696 | 0.776942 | 0.547030 | 0.393048 | 0.292020 | 0.245553 | 0.225174 | 0.217356 | 0.215172 | 0.214916 | 0.214645 | 0.214889 | 0.215853 | 0.226762 | 0.251500 | 0.289092 | 0.357587 | 0.533070 | 0.727088 | 1.039481 | 1.273325 | 1.233414 | 0.837127 | 0.535850 | 0.345525 | 0.257152 | 0.218914 | 0.215697 | 0.225991 | 0.228607 | 0.222819 | 0.216352 | 0.214652 | 0.218129 | 0.221966 | 0.221285 | 0.217642 | 0.214906 | 0.218503 | 0.239108 | 0.276182 | 0.334794 | 0.415646 | 0.477289 | 0.480572 | 0.408710 | 0.331551 | 0.288922 | 0.272955 | 0.281633 | 0.337843 | 0.494892 | 0.803493 | 1.162356 | 1.398856 | 1.388941 | 1.133024 | 0.845855 | 0.737229 | 0.706565 | 0.743634 | 0.763784 | 0.782943 | 0.739799 | 0.657428 | 0.572176 | 0.535173 | 0.557038 | 0.666540 | 0.966608 | 1.730072 | 3.884496 | 8.495663 | 15.275377 | 19.800493 | 19.481922 | 12.673780 | 8.593296 | 4.684031 | 2.250864 | 1.077448 | 0.754265 | 0.631623 | 0.614914 | 0.666925 | 0.823264 | 1.035371 | 1.198878 | 1.288143 | 1.344227 | 1.304821 | 1.112933 | 0.951471 | 0.746208 | 0.565182 | 0.453414 | 0.366100 | 0.291082 | 0.259702 | 0.247443 | 0.240011 | 0.233280 | 0.228770 | 0.227157 | 0.228363 | 0.227314 | 0.222242 | 0.218521 | 0.215758 | 0.214717 | 0.214629 | 0.215384 | 0.219691 | 0.231770 | 0.246428 | 0.258024 | 0.255695 | 0.244702 | 0.227729 | 0.219099 | 0.214861 | 0.214951 | 0.216789 | 0.220509 | 0.230279 | 0.240747 | 0.257878 | 0.270490 | 0.278984 | 0.289914 | 0.309393 | 0.338044 | 0.407405 | 0.522699 | 0.669741 | 0.787247 | 0.775190 | 0.715767 | 0.638745 | 0.577621 | 0.534267 | 0.516885 | 0.503635 | 0.513446 | 0.525639 | 0.566831 | 0.595355 | 0.610177 | 0.579286 | 0.490253 | 0.375266 | 0.293060 | 0.239282 | 0.219088 | 0.214755 | 0.225200 | 0.246444 | 0.275484 | 0.308506 | 0.330618 | 0.308972 | 0.271990 | 0.238095 | 0.220764 | 0.214776 | 0.215648 | 0.217910 | 0.217603 | 0.215652 | 0.214695 | 0.217381 | 0.224082 | 0.227845 | 0.223882 | 0.219966 | 0.216977 | 0.216995 | 0.217389 | 0.219602 |

S) neural network features, identity

  
|  | time window | peak latency | cluster *p* | peak Cohen's *d* |  | | | |
| **all electrodes** |  | | | |  | | | |
|  | | | | | | | | |

Model correlations, cluster permutation tests

|  | **left hemisphere** | | | | **right hemisphere** | | | |
|  | time window | peak latency | cluster *p* | peak Cohen's *d* | time window | peak latency | cluster *p* | peak Cohen's *d* |
| **anterior** |  | | | |  | | | |
| **central** |  | | | |  | | | |
| **posterior** |  | | | |  | | | |

  

Model correlations, Bayesian statistics

|  | -200 | -195 | -190 | -185 | -180 | -175 | -170 | -165 | -160 | -155 | -150 | -145 | -140 | -135 | -130 | -125 | -120 | -115 | -110 | -105 | -100 | -95 | -90 | -85 | -80 | -75 | -70 | -65 | -60 | -55 | -50 | -45 | -40 | -35 | -30 | -25 | -20 | -15 | -10 | -5 | 0 | 5 | 10 | 15 | 20 | 25 | 30 | 35 | 40 | 45 | 50 | 55 | 60 | 65 | 70 | 75 | 80 | 85 | 90 | 95 | 100 | 105 | 110 | 115 | 120 | 125 | 130 | 135 | 140 | 145 | 150 | 155 | 160 | 165 | 170 | 175 | 180 | 185 | 190 | 195 | 200 | 205 | 210 | 215 | 220 | 225 | 230 | 235 | 240 | 245 | 250 | 255 | 260 | 265 | 270 | 275 | 280 | 285 | 290 | 295 | 300 | 305 | 310 | 315 | 320 | 325 | 330 | 335 | 340 | 345 | 350 | 355 | 360 | 365 | 370 | 375 | 380 | 385 | 390 | 395 | 400 | 405 | 410 | 415 | 420 | 425 | 430 | 435 | 440 | 445 | 450 | 455 | 460 | 465 | 470 | 475 | 480 | 485 | 490 | 495 | 500 | 505 | 510 | 515 | 520 | 525 | 530 | 535 | 540 | 545 | 550 | 555 | 560 | 565 | 570 | 575 | 580 | 585 | 590 | 595 | 600 | 605 | 610 | 615 | 620 | 625 | 630 | 635 | 640 | 645 | 650 | 655 | 660 | 665 | 670 | 675 | 680 | 685 | 690 | 695 | 700 | 705 | 710 | 715 | 720 | 725 | 730 | 735 | 740 | 745 | 750 | 755 | 760 | 765 | 770 | 775 | 780 | 785 | 790 | 795 | 800 | 805 | 810 | 815 | 820 | 825 | 830 | 835 | 840 | 845 | 850 | 855 | 860 | 865 | 870 | 875 | 880 | 885 | 890 | 895 | 900 | 905 | 910 | 915 | 920 | 925 | 930 | 935 | 940 | 945 | 950 | 955 | 960 | 965 | 970 | 975 | 980 | 985 | 990 | 995 | 1000 | 1005 | 1010 | 1015 | 1020 | 1025 | 1030 | 1035 | 1040 | 1045 | 1050 | 1055 | 1060 | 1065 | 1070 | 1075 | 1080 | 1085 | 1090 | 1095 | 1100 | 1105 | 1110 | 1115 | 1120 | 1125 | 1130 | 1135 | 1140 | 1145 | 1150 | 1155 | 1160 | 1165 | 1170 | 1175 | 1180 | 1185 | 1190 | 1195 |
| --- | --- | --- | --- | --- | --- | --- | --- | --- | --- | --- | --- | --- | --- | --- | --- | --- | --- | --- | --- | --- | --- | --- | --- | --- | --- | --- | --- | --- | --- | --- | --- | --- | --- | --- | --- | --- | --- | --- | --- | --- | --- | --- | --- | --- | --- | --- | --- | --- | --- | --- | --- | --- | --- | --- | --- | --- | --- | --- | --- | --- | --- | --- | --- | --- | --- | --- | --- | --- | --- | --- | --- | --- | --- | --- | --- | --- | --- | --- | --- | --- | --- | --- | --- | --- | --- | --- | --- | --- | --- | --- | --- | --- | --- | --- | --- | --- | --- | --- | --- | --- | --- | --- | --- | --- | --- | --- | --- | --- | --- | --- | --- | --- | --- | --- | --- | --- | --- | --- | --- | --- | --- | --- | --- | --- | --- | --- | --- | --- | --- | --- | --- | --- | --- | --- | --- | --- | --- | --- | --- | --- | --- | --- | --- | --- | --- | --- | --- | --- | --- | --- | --- | --- | --- | --- | --- | --- | --- | --- | --- | --- | --- | --- | --- | --- | --- | --- | --- | --- | --- | --- | --- | --- | --- | --- | --- | --- | --- | --- | --- | --- | --- | --- | --- | --- | --- | --- | --- | --- | --- | --- | --- | --- | --- | --- | --- | --- | --- | --- | --- | --- | --- | --- | --- | --- | --- | --- | --- | --- | --- | --- | --- | --- | --- | --- | --- | --- | --- | --- | --- | --- | --- | --- | --- | --- | --- | --- | --- | --- | --- | --- | --- | --- | --- | --- | --- | --- | --- | --- | --- | --- | --- | --- | --- | --- | --- | --- | --- | --- | --- | --- | --- | --- | --- | --- | --- | --- | --- | --- | --- | --- | --- | --- | --- | --- | --- | --- | --- | --- | --- | --- | --- | --- | --- | --- | --- | --- | --- | --- | --- | --- |
| left anterior | 0.363823 | 0.347279 | 0.324974 | 0.281679 | 0.245819 | 0.228853 | 0.224898 | 0.226640 | 0.236636 | 0.248397 | 0.261580 | 0.247365 | 0.226932 | 0.217125 | 0.214998 | 0.221128 | 0.235093 | 0.274693 | 0.419791 | 0.989148 | 4.472206 | 9.744545 | 9.825655 | 4.851041 | 1.694897 | 0.375334 | 0.220192 | 0.227414 | 0.251019 | 0.264781 | 0.236216 | 0.214766 | 0.271482 | 0.370738 | 0.497099 | 0.597684 | 0.805308 | 0.840656 | 0.734871 | 0.514016 | 0.357938 | 0.248230 | 0.216364 | 0.218463 | 0.241234 | 0.270664 | 0.289780 | 0.292643 | 0.271631 | 0.244124 | 0.230631 | 0.222163 | 0.220686 | 0.224697 | 0.236928 | 0.254715 | 0.277459 | 0.292217 | 0.300673 | 0.290967 | 0.274170 | 0.262768 | 0.258910 | 0.254562 | 0.249180 | 0.239465 | 0.228493 | 0.218280 | 0.214827 | 0.215125 | 0.218153 | 0.224182 | 0.234199 | 0.253746 | 0.281868 | 0.310404 | 0.340113 | 0.337859 | 0.298503 | 0.249929 | 0.221430 | 0.215085 | 0.222697 | 0.224769 | 0.218663 | 0.214666 | 0.221563 | 0.238497 | 0.258136 | 0.278910 | 0.281903 | 0.275677 | 0.274947 | 0.292277 | 0.311569 | 0.329653 | 0.332311 | 0.318037 | 0.290853 | 0.265459 | 0.234475 | 0.222001 | 0.218180 | 0.217581 | 0.221586 | 0.246168 | 0.298585 | 0.407683 | 0.540442 | 0.678665 | 0.783641 | 0.857358 | 0.840570 | 0.790175 | 0.729063 | 0.705232 | 0.736302 | 0.800157 | 0.915518 | 0.936506 | 0.862306 | 0.787838 | 0.734233 | 0.620853 | 0.590469 | 0.633090 | 0.736725 | 1.010832 | 1.427399 | 1.594083 | 1.477430 | 1.124624 | 0.726967 | 0.467600 | 0.302718 | 0.229743 | 0.215173 | 0.233277 | 0.254228 | 0.253227 | 0.240381 | 0.224190 | 0.214824 | 0.222220 | 0.251356 | 0.301593 | 0.353826 | 0.381230 | 0.386767 | 0.371570 | 0.333776 | 0.296830 | 0.270660 | 0.243750 | 0.228414 | 0.217344 | 0.214977 | 0.225348 | 0.249909 | 0.290848 | 0.329943 | 0.337681 | 0.307946 | 0.274849 | 0.249116 | 0.234243 | 0.228878 | 0.229013 | 0.232258 | 0.235642 | 0.237284 | 0.241817 | 0.252782 | 0.270223 | 0.300423 | 0.344943 | 0.379605 | 0.384512 | 0.359723 | 0.317816 | 0.282928 | 0.258007 | 0.245884 | 0.246690 | 0.268740 | 0.304635 | 0.347886 | 0.365059 | 0.331552 | 0.289212 | 0.268104 | 0.260881 | 0.255932 | 0.261197 | 0.261666 | 0.258217 | 0.245433 | 0.235672 | 0.226513 | 0.221833 | 0.217752 | 0.217561 | 0.219536 | 0.225842 | 0.233674 | 0.238049 | 0.230275 | 0.220598 | 0.214629 | 0.226651 | 0.268359 | 0.341759 | 0.478351 | 0.700512 | 0.955500 | 1.122635 | 1.135710 | 0.997922 | 0.817431 | 0.668660 | 0.555672 | 0.466226 | 0.419055 | 0.373832 | 0.321375 | 0.277886 | 0.252051 | 0.231694 | 0.222287 | 0.218571 | 0.217865 | 0.218984 | 0.220911 | 0.221017 | 0.222355 | 0.222227 | 0.219203 | 0.216187 | 0.214724 | 0.214919 | 0.215691 | 0.216218 | 0.215853 | 0.215169 | 0.214647 | 0.214837 | 0.217997 | 0.226677 | 0.244000 | 0.264447 | 0.298799 | 0.325869 | 0.352085 | 0.354149 | 0.341218 | 0.298073 | 0.267414 | 0.242355 | 0.229867 | 0.225950 | 0.227017 | 0.228840 | 0.236418 | 0.247462 | 0.262048 | 0.291477 | 0.346987 | 0.427872 | 0.532859 | 0.645658 | 0.738436 | 0.818491 | 0.808284 | 0.739717 | 0.634053 | 0.543857 | 0.489659 | 0.487502 | 0.482851 | 0.500962 |
| right anterior | 3.485302 | 6.248610 | 11.328095 | 18.586572 | 19.841760 | 18.748013 | 12.735012 | 6.674682 | 2.414558 | 0.762701 | 0.303230 | 0.215073 | 0.244405 | 0.275068 | 0.296279 | 0.332257 | 0.368112 | 0.388972 | 0.426044 | 0.370006 | 0.318319 | 0.267492 | 0.226096 | 0.220252 | 0.352931 | 1.848538 | 12.865827 | 42.266679 | 40.516876 | 21.959265 | 6.229406 | 1.883662 | 0.654425 | 0.347519 | 0.242506 | 0.217043 | 0.220715 | 0.249930 | 0.281982 | 0.288586 | 0.309206 | 0.283306 | 0.250126 | 0.224967 | 0.214644 | 0.224649 | 0.250210 | 0.281071 | 0.289300 | 0.257159 | 0.222355 | 0.216851 | 0.246257 | 0.295382 | 0.360585 | 0.397524 | 0.385091 | 0.325156 | 0.265103 | 0.224676 | 0.214772 | 0.218283 | 0.226777 | 0.239717 | 0.256866 | 0.277025 | 0.292284 | 0.302941 | 0.309667 | 0.324009 | 0.341706 | 0.367955 | 0.410777 | 0.436936 | 0.454440 | 0.464741 | 0.509338 | 0.532876 | 0.534927 | 0.519870 | 0.516220 | 0.465352 | 0.395466 | 0.334570 | 0.283025 | 0.241649 | 0.217171 | 0.217198 | 0.237221 | 0.277454 | 0.362063 | 0.507417 | 0.791728 | 1.257305 | 1.921387 | 2.374304 | 2.674642 | 2.830969 | 2.622989 | 2.333708 | 2.168326 | 2.158160 | 1.971394 | 1.978635 | 2.082162 | 2.240024 | 2.060034 | 1.824682 | 1.466654 | 1.091689 | 0.800141 | 0.589422 | 0.434763 | 0.349906 | 0.282059 | 0.243417 | 0.237009 | 0.244840 | 0.252980 | 0.289113 | 0.348504 | 0.404133 | 0.417713 | 0.387893 | 0.328312 | 0.270350 | 0.233647 | 0.217776 | 0.214659 | 0.216199 | 0.215862 | 0.214626 | 0.220576 | 0.242683 | 0.277002 | 0.296639 | 0.273538 | 0.228970 | 0.214935 | 0.240467 | 0.290099 | 0.315301 | 0.285978 | 0.243734 | 0.220217 | 0.214711 | 0.216864 | 0.218380 | 0.217837 | 0.220608 | 0.223969 | 0.242167 | 0.287824 | 0.374059 | 0.450391 | 0.488980 | 0.427360 | 0.356760 | 0.269112 | 0.222601 | 0.215021 | 0.220410 | 0.225208 | 0.219138 | 0.214630 | 0.228175 | 0.281038 | 0.378255 | 0.478528 | 0.504545 | 0.373073 | 0.267755 | 0.219565 | 0.218982 | 0.248594 | 0.283559 | 0.319647 | 0.327068 | 0.306051 | 0.275492 | 0.253913 | 0.241144 | 0.247615 | 0.269005 | 0.301042 | 0.326209 | 0.345904 | 0.342410 | 0.323866 | 0.300550 | 0.282688 | 0.275525 | 0.275755 | 0.274605 | 0.268978 | 0.268539 | 0.258583 | 0.244116 | 0.234734 | 0.229074 | 0.226136 | 0.226303 | 0.223681 | 0.219489 | 0.216509 | 0.214698 | 0.220999 | 0.241934 | 0.281960 | 0.345829 | 0.419883 | 0.471228 | 0.459870 | 0.419353 | 0.362968 | 0.313592 | 0.272486 | 0.244286 | 0.229015 | 0.222705 | 0.219462 | 0.220470 | 0.226321 | 0.237822 | 0.244204 | 0.244598 | 0.240462 | 0.239402 | 0.240263 | 0.249900 | 0.263106 | 0.285308 | 0.292619 | 0.270179 | 0.234010 | 0.215624 | 0.222652 | 0.246249 | 0.274118 | 0.288803 | 0.279992 | 0.271061 | 0.268850 | 0.260671 | 0.254799 | 0.254893 | 0.246692 | 0.234982 | 0.224682 | 0.217442 | 0.214737 | 0.220300 | 0.233705 | 0.244736 | 0.255539 | 0.267866 | 0.276990 | 0.272391 | 0.272432 | 0.275326 | 0.288334 | 0.310291 | 0.346316 | 0.369500 | 0.371011 | 0.325574 | 0.272855 | 0.235150 | 0.219120 | 0.214687 | 0.214837 | 0.214800 | 0.214630 | 0.214742 | 0.215087 | 0.216466 | 0.218365 | 0.220878 | 0.220838 | 0.220492 |
| left central | 0.225767 | 0.222574 | 0.218854 | 0.215110 | 0.215373 | 0.221984 | 0.244095 | 0.314538 | 0.503297 | 0.797298 | 1.049374 | 1.209424 | 1.059364 | 0.746687 | 0.528829 | 0.355992 | 0.256858 | 0.215551 | 0.248227 | 0.383284 | 0.526562 | 0.629150 | 0.552839 | 0.445957 | 0.372084 | 0.332956 | 0.311756 | 0.327950 | 0.332677 | 0.376966 | 0.404894 | 0.412670 | 0.409821 | 0.387534 | 0.324971 | 0.260178 | 0.214977 | 0.240977 | 0.313995 | 0.372791 | 0.388793 | 0.344163 | 0.311959 | 0.281134 | 0.264683 | 0.248108 | 0.233157 | 0.223364 | 0.218042 | 0.216609 | 0.217846 | 0.223896 | 0.240276 | 0.281282 | 0.351866 | 0.430131 | 0.475520 | 0.437601 | 0.350329 | 0.293249 | 0.257092 | 0.234391 | 0.234431 | 0.260161 | 0.328161 | 0.490039 | 0.760036 | 1.058512 | 1.310240 | 1.402136 | 1.415233 | 1.229932 | 1.116520 | 1.084848 | 1.067464 | 1.081303 | 1.015148 | 0.834698 | 0.587122 | 0.370425 | 0.258339 | 0.224700 | 0.214750 | 0.232003 | 0.254772 | 0.270194 | 0.269329 | 0.258836 | 0.246303 | 0.234182 | 0.222083 | 0.219144 | 0.218150 | 0.217585 | 0.217049 | 0.216717 | 0.214718 | 0.223655 | 0.252846 | 0.291844 | 0.338738 | 0.340678 | 0.301130 | 0.254553 | 0.225094 | 0.214745 | 0.217243 | 0.223274 | 0.229476 | 0.230569 | 0.236614 | 0.246757 | 0.257677 | 0.260123 | 0.261107 | 0.243131 | 0.223729 | 0.214629 | 0.223241 | 0.239568 | 0.245750 | 0.244209 | 0.230357 | 0.218571 | 0.214695 | 0.215349 | 0.215238 | 0.214638 | 0.216675 | 0.231741 | 0.280077 | 0.369399 | 0.498904 | 0.694399 | 0.862783 | 0.926283 | 0.826246 | 0.744779 | 0.599596 | 0.450041 | 0.331960 | 0.257585 | 0.221804 | 0.214706 | 0.221276 | 0.229677 | 0.227188 | 0.219895 | 0.214680 | 0.219513 | 0.234312 | 0.258447 | 0.286291 | 0.297651 | 0.291888 | 0.279672 | 0.267252 | 0.260669 | 0.258250 | 0.264394 | 0.279513 | 0.306754 | 0.334633 | 0.355941 | 0.359498 | 0.349313 | 0.313170 | 0.286509 | 0.270131 | 0.258534 | 0.247878 | 0.243822 | 0.243648 | 0.252380 | 0.273667 | 0.328260 | 0.410120 | 0.486958 | 0.515130 | 0.497310 | 0.422849 | 0.332219 | 0.248895 | 0.215310 | 0.237275 | 0.359743 | 0.681654 | 1.206727 | 1.874524 | 1.906820 | 1.335652 | 0.822556 | 0.518292 | 0.396135 | 0.368176 | 0.376678 | 0.425081 | 0.499484 | 0.499365 | 0.464824 | 0.406421 | 0.359453 | 0.320441 | 0.286790 | 0.258649 | 0.255087 | 0.264557 | 0.276832 | 0.296473 | 0.310494 | 0.304599 | 0.269348 | 0.231482 | 0.215744 | 0.215984 | 0.224262 | 0.227470 | 0.216800 | 0.216946 | 0.242399 | 0.291241 | 0.355674 | 0.410873 | 0.504552 | 0.571242 | 0.603307 | 0.569223 | 0.495045 | 0.417477 | 0.380895 | 0.350467 | 0.333412 | 0.328791 | 0.324605 | 0.322184 | 0.339025 | 0.341226 | 0.322983 | 0.302653 | 0.277862 | 0.253601 | 0.235654 | 0.221926 | 0.214915 | 0.216924 | 0.226602 | 0.237115 | 0.242829 | 0.237366 | 0.227047 | 0.216278 | 0.215460 | 0.223303 | 0.231194 | 0.238294 | 0.241651 | 0.246617 | 0.254485 | 0.276986 | 0.312091 | 0.365408 | 0.426258 | 0.492609 | 0.511256 | 0.501041 | 0.450352 | 0.392149 | 0.335260 | 0.303236 | 0.279827 | 0.276553 | 0.293035 | 0.343486 | 0.429757 | 0.565196 | 0.747731 | 0.973426 | 1.189759 | 1.247499 | 1.308603 |
| right central | 0.281662 | 0.364469 | 0.453933 | 0.598164 | 0.564495 | 0.388887 | 0.246894 | 0.217468 | 0.318040 | 0.532708 | 0.616784 | 0.563683 | 0.373712 | 0.236536 | 0.216255 | 0.243480 | 0.297759 | 0.351578 | 0.443374 | 0.501057 | 0.498841 | 0.409975 | 0.411325 | 0.487916 | 0.950661 | 4.176239 | 64.880167 | 1218.276558 | 4554.149296 | 2105.746223 | 236.772438 | 20.265395 | 1.780020 | 0.356405 | 0.220242 | 0.226873 | 0.285817 | 0.369454 | 0.450919 | 0.454848 | 0.396904 | 0.371679 | 0.369869 | 0.374703 | 0.447954 | 0.563341 | 0.681244 | 0.680354 | 0.517566 | 0.379024 | 0.310365 | 0.263750 | 0.243995 | 0.242993 | 0.259523 | 0.286614 | 0.306778 | 0.310310 | 0.312424 | 0.296841 | 0.287475 | 0.286173 | 0.297629 | 0.326106 | 0.367109 | 0.379110 | 0.354110 | 0.308192 | 0.270427 | 0.253818 | 0.251343 | 0.269023 | 0.322780 | 0.423938 | 0.578879 | 0.864087 | 1.285115 | 1.820536 | 2.802426 | 5.658335 | 11.437090 | 16.448231 | 11.404234 | 4.370123 | 1.548023 | 0.695861 | 0.398715 | 0.296261 | 0.250770 | 0.226881 | 0.216275 | 0.214836 | 0.218006 | 0.223565 | 0.229673 | 0.234072 | 0.236467 | 0.236644 | 0.237008 | 0.243983 | 0.258611 | 0.280817 | 0.292936 | 0.284044 | 0.260903 | 0.233841 | 0.217029 | 0.214992 | 0.220720 | 0.235860 | 0.257337 | 0.280684 | 0.307248 | 0.344667 | 0.422692 | 0.554786 | 0.740080 | 0.945169 | 1.091128 | 0.955635 | 0.718153 | 0.491338 | 0.369787 | 0.297968 | 0.263337 | 0.241208 | 0.232777 | 0.230295 | 0.236553 | 0.252159 | 0.282253 | 0.315650 | 0.366882 | 0.406830 | 0.412434 | 0.370662 | 0.328709 | 0.288010 | 0.269998 | 0.269956 | 0.275327 | 0.271389 | 0.259324 | 0.241542 | 0.224420 | 0.217157 | 0.214968 | 0.214701 | 0.215934 | 0.227240 | 0.256618 | 0.308856 | 0.370446 | 0.390345 | 0.386379 | 0.376034 | 0.361938 | 0.375811 | 0.419946 | 0.446008 | 0.448390 | 0.422712 | 0.360207 | 0.303429 | 0.258007 | 0.233103 | 0.223106 | 0.217664 | 0.214927 | 0.214735 | 0.216305 | 0.220511 | 0.226160 | 0.229425 | 0.229384 | 0.229875 | 0.229093 | 0.225439 | 0.224720 | 0.229696 | 0.237353 | 0.258985 | 0.292067 | 0.305482 | 0.310545 | 0.320774 | 0.306649 | 0.302927 | 0.300770 | 0.286979 | 0.270862 | 0.258508 | 0.244211 | 0.244403 | 0.262597 | 0.303481 | 0.397051 | 0.597739 | 0.945042 | 1.290006 | 1.371118 | 0.980696 | 0.620136 | 0.423313 | 0.320947 | 0.273039 | 0.253973 | 0.247694 | 0.239781 | 0.235543 | 0.233561 | 0.233076 | 0.235530 | 0.243364 | 0.252411 | 0.269792 | 0.287160 | 0.285280 | 0.276228 | 0.265404 | 0.251454 | 0.242184 | 0.240693 | 0.240891 | 0.246163 | 0.255752 | 0.266426 | 0.278153 | 0.288769 | 0.281498 | 0.264363 | 0.247223 | 0.232693 | 0.224277 | 0.220228 | 0.217521 | 0.216976 | 0.218446 | 0.220933 | 0.226139 | 0.240014 | 0.262847 | 0.287986 | 0.306220 | 0.306389 | 0.294257 | 0.284551 | 0.277128 | 0.278338 | 0.290516 | 0.314289 | 0.343080 | 0.394654 | 0.459486 | 0.560174 | 0.685023 | 0.898714 | 1.173044 | 1.490460 | 1.782601 | 2.036200 | 2.116739 | 2.121252 | 2.142690 | 2.018989 | 1.858405 | 1.701555 | 1.608903 | 1.552498 | 1.453187 | 1.255505 | 1.041302 | 0.806753 | 0.608485 | 0.493022 | 0.416468 | 0.362566 | 0.321031 | 0.305723 | 0.288639 |
| left posterior | 0.462435 | 0.384656 | 0.324960 | 0.253414 | 0.219187 | 0.216198 | 0.221262 | 0.217102 | 0.215575 | 0.252235 | 0.405321 | 0.539518 | 0.503026 | 0.387300 | 0.281663 | 0.229598 | 0.214731 | 0.222380 | 0.236209 | 0.243548 | 0.250037 | 0.232690 | 0.216014 | 0.222123 | 0.269125 | 0.399808 | 0.648077 | 1.406843 | 3.343193 | 6.294286 | 7.978292 | 5.328368 | 2.084518 | 0.885542 | 0.446766 | 0.287797 | 0.245290 | 0.222692 | 0.214631 | 0.217766 | 0.223620 | 0.237274 | 0.247401 | 0.240070 | 0.225012 | 0.214694 | 0.224244 | 0.259156 | 0.314778 | 0.361648 | 0.350247 | 0.332686 | 0.312714 | 0.276356 | 0.244574 | 0.228441 | 0.219107 | 0.218293 | 0.219253 | 0.218380 | 0.215296 | 0.217842 | 0.254074 | 0.333813 | 0.452919 | 0.543378 | 0.542997 | 0.492170 | 0.377254 | 0.278846 | 0.230798 | 0.214627 | 0.228995 | 0.252396 | 0.261550 | 0.254842 | 0.230371 | 0.215306 | 0.227101 | 0.309083 | 0.550502 | 1.048918 | 1.639665 | 1.894837 | 1.821062 | 1.703936 | 1.383694 | 1.139965 | 0.865357 | 0.675598 | 0.539630 | 0.484573 | 0.451887 | 0.436038 | 0.385416 | 0.328709 | 0.282716 | 0.254142 | 0.231573 | 0.219399 | 0.215325 | 0.214629 | 0.216341 | 0.221489 | 0.229420 | 0.250840 | 0.299774 | 0.363147 | 0.364624 | 0.306878 | 0.256798 | 0.232060 | 0.218141 | 0.215177 | 0.215236 | 0.216298 | 0.217445 | 0.218410 | 0.219307 | 0.219069 | 0.216154 | 0.214651 | 0.214760 | 0.214729 | 0.214687 | 0.214933 | 0.214924 | 0.214745 | 0.214841 | 0.218922 | 0.224142 | 0.230129 | 0.245598 | 0.276386 | 0.319133 | 0.382111 | 0.460455 | 0.557102 | 0.652328 | 0.733271 | 0.798345 | 0.837394 | 0.757008 | 0.626577 | 0.484509 | 0.369856 | 0.294448 | 0.255038 | 0.237959 | 0.235060 | 0.240192 | 0.255393 | 0.288566 | 0.339942 | 0.393714 | 0.445631 | 0.471113 | 0.474879 | 0.467126 | 0.501473 | 0.604352 | 0.844840 | 1.369219 | 2.558934 | 4.665519 | 6.567613 | 7.312760 | 5.894333 | 3.713007 | 2.117368 | 1.426628 | 1.112640 | 0.945270 | 0.759790 | 0.599823 | 0.473061 | 0.381689 | 0.329187 | 0.295513 | 0.277914 | 0.268392 | 0.259510 | 0.249347 | 0.239697 | 0.232180 | 0.225644 | 0.219549 | 0.215787 | 0.215137 | 0.215071 | 0.215161 | 0.216143 | 0.217230 | 0.216870 | 0.216417 | 0.215484 | 0.214650 | 0.214749 | 0.215055 | 0.214984 | 0.214638 | 0.216032 | 0.219852 | 0.225944 | 0.230069 | 0.231233 | 0.227813 | 0.221101 | 0.216190 | 0.214821 | 0.220233 | 0.231222 | 0.247224 | 0.270962 | 0.291715 | 0.312710 | 0.343247 | 0.364051 | 0.381643 | 0.401766 | 0.424266 | 0.447726 | 0.503234 | 0.553772 | 0.629690 | 0.686068 | 0.664959 | 0.538367 | 0.425480 | 0.346118 | 0.316764 | 0.325848 | 0.378870 | 0.511006 | 0.789246 | 1.276195 | 1.788890 | 2.207491 | 2.304980 | 2.142702 | 1.785401 | 1.383210 | 0.938592 | 0.606778 | 0.398675 | 0.294662 | 0.252435 | 0.237308 | 0.235552 | 0.242176 | 0.267497 | 0.324433 | 0.444572 | 0.658187 | 1.107991 | 1.841331 | 2.839203 | 3.249820 | 3.142749 | 2.292081 | 1.299599 | 0.661492 | 0.388508 | 0.267451 | 0.231675 | 0.222749 | 0.221211 | 0.224683 | 0.236636 | 0.256571 | 0.285451 | 0.327816 | 0.362573 | 0.386142 | 0.386368 | 0.372839 | 0.349382 | 0.319173 | 0.305854 | 0.293618 |
| right posterior | 0.215778 | 0.215591 | 0.215359 | 0.215646 | 0.219861 | 0.233402 | 0.246746 | 0.260793 | 0.254908 | 0.232434 | 0.214693 | 0.232431 | 0.296018 | 0.341207 | 0.319298 | 0.279162 | 0.247181 | 0.231580 | 0.231373 | 0.250620 | 0.324668 | 0.568839 | 0.701619 | 0.485922 | 0.345506 | 0.275117 | 0.246620 | 0.245525 | 0.255255 | 0.240537 | 0.222204 | 0.214627 | 0.222106 | 0.239113 | 0.249039 | 0.260617 | 0.260599 | 0.253361 | 0.246227 | 0.259598 | 0.290845 | 0.425573 | 0.720939 | 1.283934 | 2.379548 | 4.070071 | 4.533614 | 4.154692 | 3.551224 | 2.924759 | 2.637058 | 3.030560 | 3.926819 | 5.327777 | 7.005831 | 8.841229 | 6.362791 | 3.527605 | 1.976077 | 1.196404 | 0.954910 | 1.065687 | 1.358938 | 1.805339 | 2.312864 | 2.259654 | 1.813906 | 1.156627 | 0.670530 | 0.395264 | 0.268285 | 0.220104 | 0.216925 | 0.239035 | 0.271174 | 0.285045 | 0.259678 | 0.225428 | 0.217383 | 0.288093 | 0.522387 | 1.030549 | 2.067494 | 3.634611 | 5.370218 | 7.168775 | 8.295460 | 6.764339 | 4.248761 | 2.057716 | 0.984156 | 0.528543 | 0.329668 | 0.245467 | 0.217928 | 0.215834 | 0.229607 | 0.250695 | 0.271365 | 0.301005 | 0.342773 | 0.365599 | 0.359063 | 0.311359 | 0.256855 | 0.220166 | 0.219073 | 0.253959 | 0.299701 | 0.343574 | 0.353791 | 0.321564 | 0.266768 | 0.231821 | 0.217079 | 0.214654 | 0.216260 | 0.215767 | 0.214653 | 0.215567 | 0.216889 | 0.216847 | 0.215878 | 0.215070 | 0.214661 | 0.214789 | 0.219318 | 0.241263 | 0.294101 | 0.403626 | 0.598077 | 0.870886 | 1.251827 | 1.648136 | 1.772908 | 1.694135 | 1.298086 | 0.837588 | 0.546780 | 0.398922 | 0.306259 | 0.259654 | 0.235198 | 0.222862 | 0.216958 | 0.215015 | 0.214647 | 0.215067 | 0.219374 | 0.239585 | 0.291856 | 0.367332 | 0.431192 | 0.460716 | 0.440188 | 0.368375 | 0.305002 | 0.262475 | 0.240314 | 0.232030 | 0.237452 | 0.251150 | 0.276370 | 0.318334 | 0.384933 | 0.466455 | 0.548508 | 0.573737 | 0.548751 | 0.463660 | 0.352221 | 0.265094 | 0.222190 | 0.219186 | 0.272290 | 0.386070 | 0.493276 | 0.488439 | 0.368075 | 0.272057 | 0.221632 | 0.217171 | 0.241287 | 0.273125 | 0.303220 | 0.320336 | 0.332906 | 0.349309 | 0.374556 | 0.418844 | 0.452441 | 0.483651 | 0.484309 | 0.472334 | 0.427676 | 0.384873 | 0.330321 | 0.282211 | 0.241460 | 0.219308 | 0.215304 | 0.226358 | 0.244887 | 0.258234 | 0.260101 | 0.252036 | 0.239066 | 0.228556 | 0.225590 | 0.227736 | 0.235403 | 0.251822 | 0.277274 | 0.308161 | 0.336313 | 0.361009 | 0.378006 | 0.376872 | 0.354109 | 0.334051 | 0.308263 | 0.279747 | 0.251598 | 0.229457 | 0.218125 | 0.214840 | 0.214656 | 0.214695 | 0.217292 | 0.227447 | 0.255888 | 0.312443 | 0.404378 | 0.510063 | 0.587801 | 0.623261 | 0.622011 | 0.548782 | 0.470594 | 0.384778 | 0.292704 | 0.232838 | 0.214779 | 0.241383 | 0.306861 | 0.386512 | 0.439559 | 0.417450 | 0.359221 | 0.303617 | 0.266711 | 0.248451 | 0.239430 | 0.236868 | 0.232937 | 0.229562 | 0.225294 | 0.223714 | 0.222853 | 0.226527 | 0.229101 | 0.233916 | 0.239970 | 0.243879 | 0.236541 | 0.229397 | 0.224580 | 0.221584 | 0.220841 | 0.225769 | 0.236528 | 0.257978 | 0.290467 | 0.335122 | 0.377902 | 0.434875 | 0.454076 | 0.439325 | 0.424206 | 0.416338 |
| all electrodes | 0.439504 | 0.386131 | 0.314641 | 0.255405 | 0.218462 | 0.221136 | 0.259774 | 0.301866 | 0.311021 | 0.273050 | 0.233884 | 0.214628 | 0.231025 | 0.247839 | 0.275943 | 0.338746 | 0.402203 | 0.561267 | 1.180073 | 2.085375 | 2.776256 | 3.123138 | 2.399195 | 2.067282 | 1.277384 | 0.585756 | 0.354636 | 0.286217 | 0.244627 | 0.233931 | 0.240940 | 0.257436 | 0.273415 | 0.308498 | 0.331135 | 0.336413 | 0.292515 | 0.238834 | 0.214802 | 0.223259 | 0.259051 | 0.300979 | 0.298750 | 0.269349 | 0.245573 | 0.227088 | 0.219235 | 0.217580 | 0.217897 | 0.217565 | 0.215393 | 0.215100 | 0.223310 | 0.239845 | 0.259140 | 0.275962 | 0.270230 | 0.249394 | 0.225661 | 0.214913 | 0.247958 | 0.368601 | 0.681633 | 1.019904 | 0.960354 | 0.653203 | 0.408757 | 0.266951 | 0.217442 | 0.216306 | 0.218898 | 0.217060 | 0.215262 | 0.224221 | 0.238394 | 0.263117 | 0.285027 | 0.289191 | 0.340568 | 0.412619 | 0.514113 | 0.787867 | 1.385365 | 1.812790 | 1.802407 | 1.520729 | 1.200087 | 0.948757 | 0.762019 | 0.551782 | 0.414902 | 0.334100 | 0.277371 | 0.243316 | 0.229450 | 0.221126 | 0.219569 | 0.222062 | 0.230612 | 0.250569 | 0.286156 | 0.324973 | 0.351618 | 0.367819 | 0.355289 | 0.312250 | 0.266161 | 0.242713 | 0.223632 | 0.216241 | 0.214677 | 0.217035 | 0.221502 | 0.225292 | 0.231693 | 0.233057 | 0.232339 | 0.233023 | 0.238100 | 0.247936 | 0.259962 | 0.269857 | 0.282272 | 0.284317 | 0.268618 | 0.251310 | 0.241469 | 0.230428 | 0.218365 | 0.214872 | 0.223815 | 0.249621 | 0.301111 | 0.387719 | 0.513406 | 0.610001 | 0.687412 | 0.665090 | 0.615535 | 0.565237 | 0.511311 | 0.415135 | 0.331480 | 0.268056 | 0.232344 | 0.219967 | 0.217009 | 0.219240 | 0.231201 | 0.259114 | 0.314769 | 0.425309 | 0.531441 | 0.565805 | 0.541096 | 0.453143 | 0.378243 | 0.349208 | 0.328213 | 0.311573 | 0.323049 | 0.340130 | 0.377865 | 0.427544 | 0.481465 | 0.521425 | 0.558004 | 0.554594 | 0.578236 | 0.611670 | 0.681397 | 0.755500 | 0.861507 | 0.863391 | 0.780992 | 0.628730 | 0.479494 | 0.343594 | 0.276692 | 0.239101 | 0.226399 | 0.225641 | 0.237768 | 0.261046 | 0.301157 | 0.315289 | 0.297102 | 0.274681 | 0.261579 | 0.242507 | 0.234389 | 0.237026 | 0.232444 | 0.222312 | 0.216424 | 0.214957 | 0.227398 | 0.262428 | 0.330200 | 0.415281 | 0.427050 | 0.403136 | 0.371773 | 0.321387 | 0.272105 | 0.244661 | 0.224128 | 0.215452 | 0.215376 | 0.220216 | 0.225003 | 0.227048 | 0.225680 | 0.220822 | 0.214718 | 0.223386 | 0.271753 | 0.391739 | 0.618044 | 0.886956 | 1.048087 | 0.866051 | 0.545228 | 0.373836 | 0.290722 | 0.248815 | 0.239971 | 0.237627 | 0.236800 | 0.241934 | 0.260380 | 0.281074 | 0.339643 | 0.439510 | 0.569408 | 0.710753 | 0.828311 | 0.880725 | 0.928754 | 0.915660 | 0.795329 | 0.700640 | 0.670702 | 0.647484 | 0.637257 | 0.674943 | 0.753509 | 0.875399 | 0.923895 | 0.854227 | 0.762215 | 0.731604 | 0.656871 | 0.655309 | 0.689303 | 0.736259 | 0.705582 | 0.650038 | 0.551827 | 0.500120 | 0.464422 | 0.436929 | 0.415306 | 0.393041 | 0.372206 | 0.356796 | 0.350149 | 0.356658 | 0.390376 | 0.437463 | 0.514655 | 0.591946 | 0.677573 | 0.803815 | 1.159808 | 1.733237 | 3.072308 | 6.218524 | 9.196179 | 10.726106 |

T) neural network features, sex

  
|  | time window | peak latency | cluster *p* | peak Cohen's *d* |  | | | |
| **all electrodes** |  | | | |  | | | |
|  | | | | | | | | |

Model correlations, cluster permutation tests

|  | **left hemisphere** | | | | **right hemisphere** | | | |
|  | time window | peak latency | cluster *p* | peak Cohen's *d* | time window | peak latency | cluster *p* | peak Cohen's *d* |
| **anterior** |  | | | |  | | | |
| **central** |  | | | |  | | | |
| **posterior** |  | | | | 805 - 1070 ms | 955 ms | 0.004 | -1.0297 |

  

Model correlations, Bayesian statistics

|  | -200 | -195 | -190 | -185 | -180 | -175 | -170 | -165 | -160 | -155 | -150 | -145 | -140 | -135 | -130 | -125 | -120 | -115 | -110 | -105 | -100 | -95 | -90 | -85 | -80 | -75 | -70 | -65 | -60 | -55 | -50 | -45 | -40 | -35 | -30 | -25 | -20 | -15 | -10 | -5 | 0 | 5 | 10 | 15 | 20 | 25 | 30 | 35 | 40 | 45 | 50 | 55 | 60 | 65 | 70 | 75 | 80 | 85 | 90 | 95 | 100 | 105 | 110 | 115 | 120 | 125 | 130 | 135 | 140 | 145 | 150 | 155 | 160 | 165 | 170 | 175 | 180 | 185 | 190 | 195 | 200 | 205 | 210 | 215 | 220 | 225 | 230 | 235 | 240 | 245 | 250 | 255 | 260 | 265 | 270 | 275 | 280 | 285 | 290 | 295 | 300 | 305 | 310 | 315 | 320 | 325 | 330 | 335 | 340 | 345 | 350 | 355 | 360 | 365 | 370 | 375 | 380 | 385 | 390 | 395 | 400 | 405 | 410 | 415 | 420 | 425 | 430 | 435 | 440 | 445 | 450 | 455 | 460 | 465 | 470 | 475 | 480 | 485 | 490 | 495 | 500 | 505 | 510 | 515 | 520 | 525 | 530 | 535 | 540 | 545 | 550 | 555 | 560 | 565 | 570 | 575 | 580 | 585 | 590 | 595 | 600 | 605 | 610 | 615 | 620 | 625 | 630 | 635 | 640 | 645 | 650 | 655 | 660 | 665 | 670 | 675 | 680 | 685 | 690 | 695 | 700 | 705 | 710 | 715 | 720 | 725 | 730 | 735 | 740 | 745 | 750 | 755 | 760 | 765 | 770 | 775 | 780 | 785 | 790 | 795 | 800 | 805 | 810 | 815 | 820 | 825 | 830 | 835 | 840 | 845 | 850 | 855 | 860 | 865 | 870 | 875 | 880 | 885 | 890 | 895 | 900 | 905 | 910 | 915 | 920 | 925 | 930 | 935 | 940 | 945 | 950 | 955 | 960 | 965 | 970 | 975 | 980 | 985 | 990 | 995 | 1000 | 1005 | 1010 | 1015 | 1020 | 1025 | 1030 | 1035 | 1040 | 1045 | 1050 | 1055 | 1060 | 1065 | 1070 | 1075 | 1080 | 1085 | 1090 | 1095 | 1100 | 1105 | 1110 | 1115 | 1120 | 1125 | 1130 | 1135 | 1140 | 1145 | 1150 | 1155 | 1160 | 1165 | 1170 | 1175 | 1180 | 1185 | 1190 | 1195 |
| --- | --- | --- | --- | --- | --- | --- | --- | --- | --- | --- | --- | --- | --- | --- | --- | --- | --- | --- | --- | --- | --- | --- | --- | --- | --- | --- | --- | --- | --- | --- | --- | --- | --- | --- | --- | --- | --- | --- | --- | --- | --- | --- | --- | --- | --- | --- | --- | --- | --- | --- | --- | --- | --- | --- | --- | --- | --- | --- | --- | --- | --- | --- | --- | --- | --- | --- | --- | --- | --- | --- | --- | --- | --- | --- | --- | --- | --- | --- | --- | --- | --- | --- | --- | --- | --- | --- | --- | --- | --- | --- | --- | --- | --- | --- | --- | --- | --- | --- | --- | --- | --- | --- | --- | --- | --- | --- | --- | --- | --- | --- | --- | --- | --- | --- | --- | --- | --- | --- | --- | --- | --- | --- | --- | --- | --- | --- | --- | --- | --- | --- | --- | --- | --- | --- | --- | --- | --- | --- | --- | --- | --- | --- | --- | --- | --- | --- | --- | --- | --- | --- | --- | --- | --- | --- | --- | --- | --- | --- | --- | --- | --- | --- | --- | --- | --- | --- | --- | --- | --- | --- | --- | --- | --- | --- | --- | --- | --- | --- | --- | --- | --- | --- | --- | --- | --- | --- | --- | --- | --- | --- | --- | --- | --- | --- | --- | --- | --- | --- | --- | --- | --- | --- | --- | --- | --- | --- | --- | --- | --- | --- | --- | --- | --- | --- | --- | --- | --- | --- | --- | --- | --- | --- | --- | --- | --- | --- | --- | --- | --- | --- | --- | --- | --- | --- | --- | --- | --- | --- | --- | --- | --- | --- | --- | --- | --- | --- | --- | --- | --- | --- | --- | --- | --- | --- | --- | --- | --- | --- | --- | --- | --- | --- | --- | --- | --- | --- | --- | --- | --- | --- | --- | --- | --- | --- | --- | --- | --- | --- | --- | --- |
| left anterior | 1.181835 | 1.106891 | 1.103781 | 0.858427 | 0.710467 | 0.536711 | 0.434911 | 0.343872 | 0.284837 | 0.245314 | 0.234296 | 0.227373 | 0.222731 | 0.221724 | 0.220382 | 0.220560 | 0.218134 | 0.215469 | 0.217252 | 0.256872 | 0.482330 | 1.024799 | 2.209801 | 4.449232 | 6.844791 | 4.473092 | 2.550111 | 1.545685 | 0.922988 | 0.612124 | 0.503310 | 0.495915 | 0.538364 | 0.553414 | 0.449324 | 0.349363 | 0.281960 | 0.227179 | 0.215552 | 0.240852 | 0.326853 | 0.595469 | 1.024696 | 1.370730 | 1.575223 | 1.332891 | 0.859539 | 0.553555 | 0.366971 | 0.264663 | 0.232558 | 0.223226 | 0.225252 | 0.248418 | 0.343508 | 0.584095 | 1.188563 | 2.669013 | 5.862953 | 11.027159 | 13.829012 | 7.940187 | 2.790446 | 1.047265 | 0.497871 | 0.327157 | 0.289576 | 0.292959 | 0.314306 | 0.352872 | 0.389651 | 0.409213 | 0.412762 | 0.399936 | 0.382511 | 0.373320 | 0.362347 | 0.333782 | 0.295055 | 0.256007 | 0.228165 | 0.214964 | 0.219874 | 0.235970 | 0.250566 | 0.256559 | 0.245133 | 0.227974 | 0.216309 | 0.215253 | 0.218357 | 0.220440 | 0.220148 | 0.216869 | 0.215493 | 0.216890 | 0.221367 | 0.227819 | 0.233583 | 0.232514 | 0.226948 | 0.217096 | 0.214819 | 0.219679 | 0.226706 | 0.240196 | 0.247458 | 0.253522 | 0.256534 | 0.267451 | 0.284975 | 0.318482 | 0.323181 | 0.293533 | 0.251921 | 0.221572 | 0.214992 | 0.227945 | 0.256542 | 0.300327 | 0.349685 | 0.462578 | 0.629303 | 0.839521 | 0.964361 | 0.945601 | 0.670913 | 0.433703 | 0.302297 | 0.242277 | 0.217072 | 0.217745 | 0.240562 | 0.273858 | 0.315404 | 0.335641 | 0.320480 | 0.275623 | 0.238535 | 0.216884 | 0.214993 | 0.221644 | 0.226497 | 0.224031 | 0.218934 | 0.217959 | 0.216407 | 0.218526 | 0.223931 | 0.235840 | 0.257235 | 0.301433 | 0.349650 | 0.389632 | 0.390150 | 0.366337 | 0.333759 | 0.303928 | 0.280285 | 0.273727 | 0.262131 | 0.245240 | 0.233452 | 0.224277 | 0.217368 | 0.215027 | 0.214903 | 0.217722 | 0.219267 | 0.219422 | 0.218776 | 0.217071 | 0.215468 | 0.214641 | 0.215856 | 0.219150 | 0.222284 | 0.222635 | 0.221843 | 0.218088 | 0.215615 | 0.214646 | 0.215194 | 0.219266 | 0.226208 | 0.239503 | 0.247682 | 0.243216 | 0.233783 | 0.222831 | 0.215484 | 0.215801 | 0.224460 | 0.239033 | 0.245498 | 0.248444 | 0.237931 | 0.225510 | 0.215989 | 0.214633 | 0.214815 | 0.214791 | 0.216821 | 0.224960 | 0.231594 | 0.232047 | 0.224185 | 0.216598 | 0.215981 | 0.235822 | 0.307859 | 0.489567 | 0.871310 | 1.431741 | 1.962540 | 2.088114 | 1.930471 | 1.558930 | 1.207211 | 0.901335 | 0.724022 | 0.648188 | 0.697551 | 0.800560 | 0.958447 | 1.001701 | 0.902026 | 0.716885 | 0.523513 | 0.375505 | 0.308918 | 0.285449 | 0.269234 | 0.265679 | 0.265828 | 0.267522 | 0.260375 | 0.250233 | 0.237108 | 0.231900 | 0.230687 | 0.237816 | 0.253413 | 0.287513 | 0.354977 | 0.473938 | 0.633881 | 0.769828 | 0.800783 | 0.703810 | 0.569992 | 0.455272 | 0.402773 | 0.359900 | 0.337292 | 0.311835 | 0.290803 | 0.267821 | 0.250075 | 0.233604 | 0.227038 | 0.223262 | 0.223619 | 0.231790 | 0.248109 | 0.263546 | 0.274851 | 0.285058 | 0.297462 | 0.307379 | 0.316744 | 0.332030 | 0.353636 | 0.380121 | 0.421767 | 0.459999 | 0.506877 | 0.564108 | 0.579670 | 0.586606 |
| right anterior | 350.822016 | 857.766805 | 1669.401493 | 5568.575640 | 17343.003697 | 49325.601225 | 33846.747097 | 8522.310653 | 908.873734 | 33.474329 | 1.760787 | 0.362200 | 0.214720 | 0.273326 | 0.432823 | 0.722861 | 1.104694 | 1.386938 | 1.466324 | 1.055625 | 0.634027 | 0.394096 | 0.264162 | 0.216312 | 0.228038 | 0.296294 | 0.476211 | 1.021194 | 2.534879 | 5.551999 | 8.773567 | 15.125010 | 13.311616 | 5.693396 | 2.158573 | 1.029600 | 0.506990 | 0.417116 | 0.493496 | 0.690281 | 1.005469 | 1.135466 | 0.783611 | 0.494827 | 0.322888 | 0.244049 | 0.226049 | 0.219544 | 0.217527 | 0.218707 | 0.220127 | 0.216697 | 0.215039 | 0.225048 | 0.244325 | 0.272743 | 0.290620 | 0.262208 | 0.225358 | 0.216662 | 0.258295 | 0.313747 | 0.386154 | 0.487191 | 0.573698 | 0.618734 | 0.816191 | 0.941262 | 0.952487 | 0.891080 | 0.897326 | 0.964084 | 1.307068 | 1.754143 | 2.471593 | 3.097217 | 3.152102 | 2.165814 | 1.279367 | 0.693928 | 0.427532 | 0.298792 | 0.248755 | 0.228153 | 0.224822 | 0.228216 | 0.237404 | 0.252029 | 0.279692 | 0.285824 | 0.263428 | 0.225515 | 0.219944 | 0.305250 | 0.484067 | 0.662081 | 0.618058 | 0.413612 | 0.257186 | 0.215017 | 0.258573 | 0.326746 | 0.356186 | 0.310947 | 0.254350 | 0.222923 | 0.214684 | 0.215992 | 0.216420 | 0.216561 | 0.215144 | 0.214946 | 0.219114 | 0.230977 | 0.259675 | 0.301598 | 0.343905 | 0.404779 | 0.463012 | 0.524277 | 0.655284 | 0.999746 | 1.731427 | 3.464340 | 6.550120 | 10.470322 | 13.716538 | 12.067301 | 6.637115 | 3.304110 | 1.664507 | 0.904741 | 0.602710 | 0.476848 | 0.409228 | 0.377565 | 0.365732 | 0.349697 | 0.341310 | 0.341228 | 0.350061 | 0.357832 | 0.368866 | 0.366901 | 0.354929 | 0.338906 | 0.327510 | 0.321930 | 0.327606 | 0.331514 | 0.339988 | 0.361038 | 0.376532 | 0.373101 | 0.362567 | 0.342356 | 0.322390 | 0.304886 | 0.291225 | 0.282179 | 0.275029 | 0.252502 | 0.232117 | 0.219415 | 0.214632 | 0.219981 | 0.228740 | 0.234917 | 0.229498 | 0.218297 | 0.215219 | 0.226356 | 0.249956 | 0.272873 | 0.284926 | 0.289331 | 0.289051 | 0.292478 | 0.291063 | 0.287086 | 0.279163 | 0.271863 | 0.256916 | 0.246165 | 0.234673 | 0.232152 | 0.235589 | 0.252862 | 0.278325 | 0.325093 | 0.362551 | 0.373294 | 0.375935 | 0.394505 | 0.423922 | 0.463777 | 0.501957 | 0.495250 | 0.453951 | 0.386276 | 0.321500 | 0.271945 | 0.244446 | 0.228825 | 0.222872 | 0.219497 | 0.220397 | 0.219358 | 0.217266 | 0.214690 | 0.218234 | 0.243158 | 0.292859 | 0.364880 | 0.402068 | 0.372849 | 0.292669 | 0.236961 | 0.215643 | 0.219376 | 0.243491 | 0.273681 | 0.301012 | 0.307970 | 0.304932 | 0.286738 | 0.262339 | 0.236317 | 0.221210 | 0.214672 | 0.223340 | 0.239603 | 0.247914 | 0.242232 | 0.225672 | 0.215724 | 0.216156 | 0.220882 | 0.219912 | 0.214864 | 0.220206 | 0.249953 | 0.290738 | 0.336216 | 0.382277 | 0.419632 | 0.459633 | 0.529616 | 0.606161 | 0.673844 | 0.734732 | 0.802166 | 0.908627 | 0.978142 | 0.987321 | 0.940976 | 0.850033 | 0.663059 | 0.487812 | 0.353759 | 0.273807 | 0.231493 | 0.216426 | 0.214886 | 0.215898 | 0.215786 | 0.215147 | 0.214664 | 0.214812 | 0.215474 | 0.217221 | 0.221769 | 0.228150 | 0.232679 | 0.232277 | 0.226763 | 0.219405 | 0.214772 | 0.214785 | 0.215800 |
| left central | 0.218038 | 0.214772 | 0.220924 | 0.247984 | 0.283025 | 0.289899 | 0.252969 | 0.215185 | 0.267142 | 0.601777 | 1.995556 | 5.257306 | 4.580964 | 1.906770 | 0.684566 | 0.308317 | 0.225106 | 0.216175 | 0.252776 | 0.371588 | 0.555261 | 0.781021 | 0.761495 | 0.633905 | 0.442477 | 0.333257 | 0.261740 | 0.253239 | 0.258634 | 0.303488 | 0.394487 | 0.511388 | 0.545821 | 0.573976 | 0.432373 | 0.300582 | 0.234976 | 0.218057 | 0.216005 | 0.228305 | 0.277526 | 0.400341 | 0.460388 | 0.409492 | 0.339190 | 0.288029 | 0.258501 | 0.251095 | 0.236018 | 0.224759 | 0.216317 | 0.215717 | 0.238210 | 0.326490 | 0.577857 | 1.113131 | 2.483693 | 4.265782 | 2.515463 | 0.841773 | 0.331648 | 0.220153 | 0.231462 | 0.315797 | 0.412864 | 0.407636 | 0.386682 | 0.310505 | 0.231801 | 0.217110 | 0.249053 | 0.314459 | 0.394230 | 0.509354 | 0.623408 | 0.640983 | 0.580545 | 0.497463 | 0.383354 | 0.297216 | 0.254501 | 0.232238 | 0.221240 | 0.218078 | 0.218829 | 0.220511 | 0.229366 | 0.254145 | 0.311777 | 0.401915 | 0.534721 | 0.642707 | 0.701824 | 0.673573 | 0.583860 | 0.459381 | 0.376282 | 0.316829 | 0.277973 | 0.259202 | 0.260300 | 0.263282 | 0.268518 | 0.279589 | 0.299226 | 0.323905 | 0.346218 | 0.359020 | 0.353627 | 0.338410 | 0.317781 | 0.298132 | 0.287793 | 0.284551 | 0.290667 | 0.316227 | 0.370605 | 0.471747 | 0.635835 | 0.820584 | 0.955964 | 1.022008 | 0.991866 | 0.827173 | 0.653535 | 0.528301 | 0.452850 | 0.403246 | 0.362758 | 0.328157 | 0.302303 | 0.281631 | 0.270043 | 0.272781 | 0.303751 | 0.368709 | 0.473915 | 0.652881 | 0.858952 | 0.954012 | 0.930909 | 0.832545 | 0.697263 | 0.595775 | 0.513801 | 0.483710 | 0.516624 | 0.598569 | 0.707814 | 0.857445 | 0.897618 | 0.717465 | 0.483626 | 0.335386 | 0.258474 | 0.224698 | 0.215507 | 0.214650 | 0.214651 | 0.215025 | 0.217979 | 0.222378 | 0.229604 | 0.236393 | 0.241799 | 0.244725 | 0.247881 | 0.246210 | 0.241163 | 0.233873 | 0.228791 | 0.223992 | 0.218854 | 0.216811 | 0.216810 | 0.219021 | 0.223906 | 0.232610 | 0.240214 | 0.244850 | 0.239399 | 0.227632 | 0.217108 | 0.216777 | 0.241640 | 0.304388 | 0.413772 | 0.547915 | 0.572281 | 0.461283 | 0.296887 | 0.218026 | 0.243502 | 0.375259 | 0.594764 | 0.704258 | 0.642806 | 0.511605 | 0.397263 | 0.322984 | 0.283352 | 0.263473 | 0.258645 | 0.265286 | 0.282512 | 0.304418 | 0.330411 | 0.360738 | 0.396196 | 0.417986 | 0.419845 | 0.403992 | 0.377028 | 0.341393 | 0.292568 | 0.250716 | 0.225276 | 0.214655 | 0.222984 | 0.239630 | 0.248947 | 0.243726 | 0.231094 | 0.224107 | 0.219150 | 0.217519 | 0.219213 | 0.224957 | 0.228864 | 0.232347 | 0.230869 | 0.222724 | 0.215101 | 0.218867 | 0.240238 | 0.273316 | 0.302118 | 0.319806 | 0.319373 | 0.314252 | 0.310938 | 0.311665 | 0.309371 | 0.305139 | 0.299419 | 0.286900 | 0.263264 | 0.242090 | 0.227260 | 0.218024 | 0.215379 | 0.214841 | 0.214802 | 0.215014 | 0.216023 | 0.216320 | 0.216922 | 0.216949 | 0.216345 | 0.215480 | 0.215282 | 0.214950 | 0.215038 | 0.215278 | 0.215509 | 0.215853 | 0.217076 | 0.219795 | 0.223643 | 0.228813 | 0.235024 | 0.239361 | 0.237622 | 0.232335 | 0.224402 | 0.217190 | 0.214783 | 0.221371 | 0.229354 | 0.240962 |
| right central | 0.391820 | 0.359413 | 0.306617 | 0.250046 | 0.224131 | 0.216332 | 0.216788 | 0.217765 | 0.221303 | 0.240191 | 0.253768 | 0.252846 | 0.239537 | 0.218739 | 0.222416 | 0.268672 | 0.393487 | 0.688228 | 1.283227 | 2.157376 | 2.874975 | 2.397778 | 1.221935 | 0.667934 | 0.350435 | 0.237626 | 0.214971 | 0.218612 | 0.246618 | 0.292422 | 0.347060 | 0.409898 | 0.482498 | 0.501855 | 0.542210 | 0.560525 | 0.509947 | 0.392236 | 0.294426 | 0.232907 | 0.215238 | 0.229389 | 0.231611 | 0.219137 | 0.217474 | 0.265381 | 0.385105 | 0.578326 | 0.861946 | 1.183587 | 1.585948 | 2.092151 | 2.677462 | 2.853419 | 2.947610 | 3.376274 | 4.055067 | 4.405844 | 4.178572 | 2.790173 | 1.657245 | 0.978792 | 0.625024 | 0.436573 | 0.346672 | 0.288267 | 0.253211 | 0.234763 | 0.233192 | 0.240426 | 0.275091 | 0.381971 | 0.649472 | 1.025845 | 1.564025 | 2.073749 | 2.425764 | 2.270813 | 1.957485 | 1.362061 | 0.851394 | 0.522075 | 0.363691 | 0.290474 | 0.258030 | 0.246952 | 0.256864 | 0.290014 | 0.361061 | 0.492377 | 0.683507 | 0.860796 | 0.932462 | 0.778934 | 0.565364 | 0.394432 | 0.297874 | 0.254446 | 0.241055 | 0.239079 | 0.247142 | 0.261360 | 0.292115 | 0.338315 | 0.397946 | 0.455021 | 0.520198 | 0.605247 | 0.706579 | 0.786069 | 0.858125 | 0.928262 | 0.935049 | 0.901590 | 0.860843 | 0.816582 | 0.760857 | 0.716779 | 0.697861 | 0.728443 | 0.853542 | 1.151963 | 1.795911 | 3.186253 | 6.304011 | 11.978874 | 19.146115 | 22.837026 | 21.786857 | 20.722008 | 20.524771 | 18.670294 | 16.048565 | 13.644670 | 10.601118 | 8.727552 | 7.294164 | 5.896034 | 4.064608 | 2.675013 | 1.600125 | 1.054816 | 0.802741 | 0.703677 | 0.656762 | 0.750375 | 1.013617 | 1.454563 | 2.329906 | 3.652321 | 4.918018 | 6.151492 | 6.652795 | 6.071706 | 5.114478 | 3.774037 | 2.502469 | 1.698606 | 1.107599 | 0.708549 | 0.450907 | 0.324007 | 0.270224 | 0.255578 | 0.261909 | 0.301702 | 0.389767 | 0.511843 | 0.622531 | 0.675815 | 0.589499 | 0.421783 | 0.292889 | 0.230149 | 0.214862 | 0.233675 | 0.272778 | 0.319786 | 0.347697 | 0.360748 | 0.383989 | 0.398086 | 0.405192 | 0.401270 | 0.372877 | 0.322690 | 0.279810 | 0.242472 | 0.221984 | 0.214943 | 0.215847 | 0.221735 | 0.228875 | 0.236301 | 0.242381 | 0.246366 | 0.241735 | 0.233094 | 0.224796 | 0.218068 | 0.214896 | 0.214703 | 0.215484 | 0.217197 | 0.218853 | 0.218788 | 0.218055 | 0.218597 | 0.220549 | 0.223433 | 0.229527 | 0.245965 | 0.269391 | 0.300103 | 0.340516 | 0.399991 | 0.455071 | 0.489096 | 0.480526 | 0.453909 | 0.421672 | 0.395757 | 0.380237 | 0.370582 | 0.368298 | 0.367198 | 0.371693 | 0.371978 | 0.376316 | 0.367857 | 0.363802 | 0.333499 | 0.294239 | 0.265489 | 0.250693 | 0.240054 | 0.232779 | 0.230066 | 0.229774 | 0.232583 | 0.241106 | 0.256979 | 0.269836 | 0.285205 | 0.300844 | 0.311466 | 0.320973 | 0.328317 | 0.325972 | 0.322641 | 0.307650 | 0.278686 | 0.255398 | 0.240538 | 0.230729 | 0.226690 | 0.227389 | 0.232915 | 0.247592 | 0.268372 | 0.289840 | 0.319629 | 0.361502 | 0.397480 | 0.416823 | 0.416719 | 0.382843 | 0.333475 | 0.287604 | 0.254418 | 0.235747 | 0.226867 | 0.221099 | 0.219525 | 0.219688 | 0.220288 | 0.221267 | 0.223723 | 0.225637 | 0.226879 |
| left posterior | 0.261067 | 0.270715 | 0.283166 | 0.323539 | 0.408851 | 0.540578 | 0.668971 | 0.745839 | 0.569910 | 0.305957 | 0.224642 | 0.215625 | 0.231413 | 0.239095 | 0.221755 | 0.214935 | 0.219597 | 0.254299 | 0.338977 | 0.469481 | 0.720362 | 0.882428 | 0.766565 | 0.550664 | 0.374784 | 0.285970 | 0.246350 | 0.225046 | 0.215510 | 0.221409 | 0.300890 | 0.528387 | 0.836066 | 0.901112 | 0.611195 | 0.368202 | 0.244165 | 0.216130 | 0.269761 | 0.394530 | 0.574700 | 0.625833 | 0.462078 | 0.280577 | 0.216743 | 0.229731 | 0.304632 | 0.466602 | 0.678115 | 0.867725 | 0.924655 | 0.695868 | 0.408832 | 0.244141 | 0.227164 | 0.402748 | 0.968551 | 2.276749 | 3.652260 | 3.391349 | 2.233524 | 1.293875 | 0.702669 | 0.410922 | 0.279625 | 0.228319 | 0.214830 | 0.215349 | 0.216168 | 0.214946 | 0.216359 | 0.232529 | 0.263936 | 0.341860 | 0.465046 | 0.641958 | 0.769189 | 0.835125 | 0.724199 | 0.624708 | 0.529982 | 0.494658 | 0.547207 | 0.731505 | 1.166764 | 1.990607 | 2.951773 | 2.596853 | 1.453060 | 0.646017 | 0.361230 | 0.258566 | 0.223322 | 0.214715 | 0.217080 | 0.226527 | 0.240445 | 0.262731 | 0.308713 | 0.375265 | 0.432289 | 0.470296 | 0.459107 | 0.387707 | 0.315525 | 0.266287 | 0.239500 | 0.225923 | 0.219854 | 0.216106 | 0.214806 | 0.214706 | 0.215465 | 0.216763 | 0.216875 | 0.217552 | 0.217628 | 0.217503 | 0.215721 | 0.214715 | 0.214797 | 0.214803 | 0.214626 | 0.216062 | 0.221725 | 0.230133 | 0.231535 | 0.221071 | 0.214679 | 0.233247 | 0.311265 | 0.510251 | 0.884527 | 1.453900 | 1.903483 | 2.281091 | 2.424779 | 2.762068 | 3.306871 | 4.204977 | 4.576090 | 4.194336 | 2.471274 | 1.251021 | 0.668725 | 0.446301 | 0.358200 | 0.314351 | 0.294227 | 0.292986 | 0.295899 | 0.300149 | 0.299339 | 0.281568 | 0.255513 | 0.228689 | 0.215006 | 0.217814 | 0.230420 | 0.247418 | 0.258954 | 0.263212 | 0.267374 | 0.264013 | 0.265648 | 0.270833 | 0.274079 | 0.281380 | 0.291531 | 0.307304 | 0.329272 | 0.364307 | 0.365129 | 0.340658 | 0.290140 | 0.248880 | 0.224039 | 0.216279 | 0.214672 | 0.214801 | 0.217376 | 0.224920 | 0.242424 | 0.277465 | 0.324610 | 0.366034 | 0.409304 | 0.445624 | 0.486688 | 0.525327 | 0.534862 | 0.507330 | 0.470122 | 0.411386 | 0.376203 | 0.363994 | 0.375653 | 0.412913 | 0.500591 | 0.626400 | 0.879927 | 1.290794 | 1.815609 | 2.491830 | 3.467477 | 4.482304 | 5.648288 | 6.260478 | 6.119788 | 5.321837 | 4.093404 | 2.607990 | 1.475550 | 0.793477 | 0.454864 | 0.305347 | 0.243026 | 0.220897 | 0.214923 | 0.214734 | 0.215312 | 0.215054 | 0.214657 | 0.215682 | 0.219104 | 0.224497 | 0.228103 | 0.234151 | 0.237863 | 0.241319 | 0.246287 | 0.253373 | 0.258778 | 0.259729 | 0.252959 | 0.238862 | 0.224799 | 0.216076 | 0.214955 | 0.218934 | 0.223981 | 0.226228 | 0.222216 | 0.217051 | 0.214638 | 0.217802 | 0.225682 | 0.232027 | 0.234223 | 0.228933 | 0.222856 | 0.216226 | 0.215368 | 0.227299 | 0.258026 | 0.319351 | 0.378595 | 0.392626 | 0.358527 | 0.308789 | 0.261366 | 0.228648 | 0.215112 | 0.217928 | 0.226840 | 0.235098 | 0.239206 | 0.237325 | 0.233823 | 0.235630 | 0.242055 | 0.250917 | 0.260674 | 0.276070 | 0.291461 | 0.302370 | 0.303092 | 0.301721 | 0.299492 | 0.295863 |
| right posterior | 4.018863 | 3.839326 | 3.094702 | 1.251816 | 0.636702 | 0.411297 | 0.320019 | 0.283109 | 0.267021 | 0.254504 | 0.240933 | 0.223723 | 0.215663 | 0.215964 | 0.225726 | 0.255593 | 0.319569 | 0.461575 | 0.702351 | 0.969494 | 0.880371 | 0.512886 | 0.277927 | 0.215426 | 0.328756 | 0.792102 | 1.744842 | 2.567508 | 2.116603 | 1.040156 | 0.472900 | 0.247678 | 0.224177 | 0.348061 | 0.531916 | 0.616379 | 0.517421 | 0.409138 | 0.323862 | 0.257977 | 0.229678 | 0.217949 | 0.214844 | 0.214649 | 0.214629 | 0.214781 | 0.220834 | 0.236887 | 0.264615 | 0.313921 | 0.379623 | 0.400590 | 0.317446 | 0.228195 | 0.236276 | 0.513723 | 1.964258 | 4.800950 | 4.842460 | 2.805137 | 1.282389 | 0.564329 | 0.301432 | 0.221717 | 0.227052 | 0.317174 | 0.504262 | 0.758576 | 0.895699 | 0.720507 | 0.483236 | 0.325268 | 0.253934 | 0.224336 | 0.215627 | 0.214651 | 0.214735 | 0.215706 | 0.218082 | 0.222315 | 0.232161 | 0.253351 | 0.281564 | 0.311693 | 0.346449 | 0.346480 | 0.312707 | 0.276224 | 0.248284 | 0.225867 | 0.215541 | 0.215562 | 0.217876 | 0.216146 | 0.215104 | 0.229482 | 0.299255 | 0.542887 | 1.224231 | 2.424396 | 4.191558 | 5.486241 | 5.260449 | 3.730642 | 2.223269 | 1.166199 | 0.701972 | 0.466195 | 0.385432 | 0.375444 | 0.387525 | 0.397706 | 0.399278 | 0.361445 | 0.295488 | 0.241478 | 0.216282 | 0.220958 | 0.254608 | 0.304422 | 0.353010 | 0.394126 | 0.436511 | 0.476686 | 0.517467 | 0.548612 | 0.587736 | 0.603877 | 0.578235 | 0.519554 | 0.453851 | 0.380179 | 0.321819 | 0.277800 | 0.249695 | 0.229752 | 0.218516 | 0.214633 | 0.216311 | 0.222085 | 0.231512 | 0.242869 | 0.252583 | 0.258772 | 0.255196 | 0.243614 | 0.231586 | 0.224271 | 0.221779 | 0.221262 | 0.223605 | 0.229155 | 0.239166 | 0.259802 | 0.292093 | 0.338931 | 0.412778 | 0.525181 | 0.639884 | 0.772405 | 0.883483 | 0.913547 | 0.780215 | 0.582409 | 0.421724 | 0.326100 | 0.278463 | 0.260007 | 0.258942 | 0.271488 | 0.295389 | 0.334877 | 0.390697 | 0.472064 | 0.566507 | 0.691487 | 0.865269 | 1.107053 | 1.400679 | 1.816180 | 2.173017 | 2.340378 | 2.206387 | 1.885596 | 1.548700 | 1.263437 | 1.003288 | 0.824823 | 0.689979 | 0.555827 | 0.450831 | 0.375542 | 0.317633 | 0.278798 | 0.254373 | 0.237301 | 0.230313 | 0.233843 | 0.257865 | 0.352256 | 0.747968 | 2.873358 | 16.919287 | 80.768303 | 187.940019 | 253.080336 | 226.545731 | 187.136675 | 176.370543 | 183.530319 | 215.862753 | 303.234616 | 397.346596 | 381.618448 | 311.676281 | 204.630439 | 106.445942 | 54.340909 | 30.104871 | 18.327830 | 12.939766 | 10.832293 | 10.909818 | 13.182866 | 20.339333 | 37.981408 | 85.536672 | 197.064801 | 407.558072 | 638.213508 | 780.172738 | 680.793444 | 371.180368 | 151.503351 | 55.452982 | 22.430278 | 9.536866 | 5.234688 | 3.333072 | 2.752249 | 2.630320 | 2.855289 | 2.970089 | 3.442237 | 3.788252 | 3.863184 | 3.728512 | 3.839012 | 3.823524 | 3.685137 | 3.261228 | 2.667662 | 2.121811 | 1.637682 | 1.259768 | 0.998881 | 0.841174 | 0.719170 | 0.639571 | 0.587454 | 0.539656 | 0.488380 | 0.449487 | 0.416620 | 0.395557 | 0.397661 | 0.422866 | 0.458053 | 0.486441 | 0.465978 | 0.411893 | 0.347466 | 0.301378 | 0.271764 | 0.260754 | 0.256114 | 0.259001 | 0.276583 | 0.286130 | 0.300129 |
| all electrodes | 0.259489 | 0.287996 | 0.305491 | 0.388207 | 0.541733 | 0.844117 | 1.172016 | 1.362116 | 1.130230 | 0.624474 | 0.310498 | 0.227612 | 0.215007 | 0.215997 | 0.222116 | 0.241860 | 0.268384 | 0.308255 | 0.386252 | 0.496405 | 0.522294 | 0.446735 | 0.328375 | 0.252359 | 0.220562 | 0.215057 | 0.218229 | 0.235681 | 0.258021 | 0.283411 | 0.305073 | 0.310195 | 0.322377 | 0.300439 | 0.262463 | 0.248662 | 0.238602 | 0.228703 | 0.220105 | 0.214904 | 0.215880 | 0.214734 | 0.215420 | 0.235582 | 0.306946 | 0.469085 | 0.604420 | 0.792675 | 0.958792 | 1.183965 | 1.179366 | 1.122873 | 1.023354 | 0.767359 | 0.440353 | 0.253148 | 0.218273 | 0.327987 | 0.619590 | 1.115769 | 1.440940 | 1.510738 | 1.206302 | 0.709047 | 0.337001 | 0.218755 | 0.252853 | 0.370509 | 0.502842 | 0.493387 | 0.370698 | 0.255121 | 0.217827 | 0.221423 | 0.258705 | 0.324135 | 0.391744 | 0.466973 | 0.504271 | 0.543433 | 0.531508 | 0.511857 | 0.472335 | 0.457509 | 0.422506 | 0.408244 | 0.405442 | 0.423613 | 0.414797 | 0.388225 | 0.347071 | 0.299407 | 0.258771 | 0.238420 | 0.232803 | 0.238400 | 0.244389 | 0.250550 | 0.249051 | 0.240448 | 0.226816 | 0.217589 | 0.214937 | 0.218430 | 0.219528 | 0.215704 | 0.217355 | 0.244278 | 0.300876 | 0.386077 | 0.449321 | 0.444016 | 0.386810 | 0.319927 | 0.266558 | 0.239327 | 0.230374 | 0.233677 | 0.250554 | 0.284325 | 0.327975 | 0.380949 | 0.430499 | 0.446004 | 0.445900 | 0.412374 | 0.340205 | 0.271921 | 0.228711 | 0.214643 | 0.223814 | 0.250802 | 0.282875 | 0.287035 | 0.276378 | 0.273880 | 0.286345 | 0.313711 | 0.363247 | 0.461435 | 0.615023 | 0.723707 | 0.791610 | 0.763670 | 0.697251 | 0.636920 | 0.543641 | 0.477595 | 0.481133 | 0.471550 | 0.430706 | 0.401423 | 0.365796 | 0.342968 | 0.328581 | 0.320966 | 0.327161 | 0.342222 | 0.344893 | 0.359624 | 0.377639 | 0.395282 | 0.406255 | 0.405721 | 0.392544 | 0.386547 | 0.363868 | 0.344246 | 0.322139 | 0.287905 | 0.265283 | 0.257642 | 0.257791 | 0.265103 | 0.287507 | 0.306670 | 0.335839 | 0.357403 | 0.365525 | 0.343307 | 0.332957 | 0.325720 | 0.334429 | 0.357285 | 0.424211 | 0.517883 | 0.592400 | 0.573414 | 0.464263 | 0.353614 | 0.277161 | 0.234843 | 0.220063 | 0.221943 | 0.234243 | 0.272955 | 0.369648 | 0.605220 | 1.023980 | 1.511617 | 1.432589 | 0.934893 | 0.564527 | 0.351163 | 0.267632 | 0.240796 | 0.235031 | 0.236489 | 0.246962 | 0.263154 | 0.289113 | 0.323473 | 0.381911 | 0.483119 | 0.630792 | 0.894307 | 1.469088 | 2.581513 | 3.909206 | 4.860562 | 4.487284 | 4.426301 | 4.016101 | 3.651350 | 3.224046 | 3.436992 | 3.138392 | 3.017350 | 2.894026 | 2.989290 | 3.096573 | 3.502528 | 3.664443 | 4.103480 | 3.812322 | 2.619705 | 1.546548 | 0.988587 | 0.714058 | 0.564239 | 0.485233 | 0.484354 | 0.552289 | 0.666705 | 0.928623 | 1.500495 | 2.571350 | 4.238980 | 5.684757 | 5.977799 | 5.111220 | 3.542615 | 2.047830 | 1.109657 | 0.666126 | 0.477604 | 0.371013 | 0.308965 | 0.265013 | 0.231452 | 0.215108 | 0.222893 | 0.263988 | 0.336972 | 0.421216 | 0.459676 | 0.436980 | 0.350388 | 0.279352 | 0.233883 | 0.218860 | 0.215749 | 0.214884 | 0.214663 | 0.214801 | 0.214963 | 0.214654 | 0.215129 | 0.215642 | 0.216881 |

U) neural network features, emotion

  
|  | time window | peak latency | cluster *p* | peak Cohen's *d* |  | | | |  |
| **all electrodes** | 125 - 230 ms | 160 ms | 0.0184 | 1.4788 |  | | | |  |
 340 - 560 ms | 395 ms | 0.0045 | 0.9584 |  | | | ||  |  | | | | | | | | |

Model correlations, cluster permutation tests

|  | **left hemisphere** | | | | **right hemisphere** | | | |  |
|  | time window | peak latency | cluster *p* | peak Cohen's *d* | time window | peak latency | cluster *p* | peak Cohen's *d* |  |
| **anterior** |  | | | |  | | | |  |
| **central** |  | | | | 205 - 495 ms | 275 ms | 0.0002 | 1.849 |  |
| **posterior** | 120 - 445 ms | 160 ms | 0.0027 | 1.4698 | 255 - 460 ms | 405 ms | 0.0223 | 0.9238 |  |
  | | | | 470 - 700 ms | 605 ms | 0.0238 | 0.6639 |  | | | | 725 - 1125 ms | 830 ms | 0.0046 | 0.6784 |

  

Model correlations, Bayesian statistics

|  | -200 | -195 | -190 | -185 | -180 | -175 | -170 | -165 | -160 | -155 | -150 | -145 | -140 | -135 | -130 | -125 | -120 | -115 | -110 | -105 | -100 | -95 | -90 | -85 | -80 | -75 | -70 | -65 | -60 | -55 | -50 | -45 | -40 | -35 | -30 | -25 | -20 | -15 | -10 | -5 | 0 | 5 | 10 | 15 | 20 | 25 | 30 | 35 | 40 | 45 | 50 | 55 | 60 | 65 | 70 | 75 | 80 | 85 | 90 | 95 | 100 | 105 | 110 | 115 | 120 | 125 | 130 | 135 | 140 | 145 | 150 | 155 | 160 | 165 | 170 | 175 | 180 | 185 | 190 | 195 | 200 | 205 | 210 | 215 | 220 | 225 | 230 | 235 | 240 | 245 | 250 | 255 | 260 | 265 | 270 | 275 | 280 | 285 | 290 | 295 | 300 | 305 | 310 | 315 | 320 | 325 | 330 | 335 | 340 | 345 | 350 | 355 | 360 | 365 | 370 | 375 | 380 | 385 | 390 | 395 | 400 | 405 | 410 | 415 | 420 | 425 | 430 | 435 | 440 | 445 | 450 | 455 | 460 | 465 | 470 | 475 | 480 | 485 | 490 | 495 | 500 | 505 | 510 | 515 | 520 | 525 | 530 | 535 | 540 | 545 | 550 | 555 | 560 | 565 | 570 | 575 | 580 | 585 | 590 | 595 | 600 | 605 | 610 | 615 | 620 | 625 | 630 | 635 | 640 | 645 | 650 | 655 | 660 | 665 | 670 | 675 | 680 | 685 | 690 | 695 | 700 | 705 | 710 | 715 | 720 | 725 | 730 | 735 | 740 | 745 | 750 | 755 | 760 | 765 | 770 | 775 | 780 | 785 | 790 | 795 | 800 | 805 | 810 | 815 | 820 | 825 | 830 | 835 | 840 | 845 | 850 | 855 | 860 | 865 | 870 | 875 | 880 | 885 | 890 | 895 | 900 | 905 | 910 | 915 | 920 | 925 | 930 | 935 | 940 | 945 | 950 | 955 | 960 | 965 | 970 | 975 | 980 | 985 | 990 | 995 | 1000 | 1005 | 1010 | 1015 | 1020 | 1025 | 1030 | 1035 | 1040 | 1045 | 1050 | 1055 | 1060 | 1065 | 1070 | 1075 | 1080 | 1085 | 1090 | 1095 | 1100 | 1105 | 1110 | 1115 | 1120 | 1125 | 1130 | 1135 | 1140 | 1145 | 1150 | 1155 | 1160 | 1165 | 1170 | 1175 | 1180 | 1185 | 1190 | 1195 |
| --- | --- | --- | --- | --- | --- | --- | --- | --- | --- | --- | --- | --- | --- | --- | --- | --- | --- | --- | --- | --- | --- | --- | --- | --- | --- | --- | --- | --- | --- | --- | --- | --- | --- | --- | --- | --- | --- | --- | --- | --- | --- | --- | --- | --- | --- | --- | --- | --- | --- | --- | --- | --- | --- | --- | --- | --- | --- | --- | --- | --- | --- | --- | --- | --- | --- | --- | --- | --- | --- | --- | --- | --- | --- | --- | --- | --- | --- | --- | --- | --- | --- | --- | --- | --- | --- | --- | --- | --- | --- | --- | --- | --- | --- | --- | --- | --- | --- | --- | --- | --- | --- | --- | --- | --- | --- | --- | --- | --- | --- | --- | --- | --- | --- | --- | --- | --- | --- | --- | --- | --- | --- | --- | --- | --- | --- | --- | --- | --- | --- | --- | --- | --- | --- | --- | --- | --- | --- | --- | --- | --- | --- | --- | --- | --- | --- | --- | --- | --- | --- | --- | --- | --- | --- | --- | --- | --- | --- | --- | --- | --- | --- | --- | --- | --- | --- | --- | --- | --- | --- | --- | --- | --- | --- | --- | --- | --- | --- | --- | --- | --- | --- | --- | --- | --- | --- | --- | --- | --- | --- | --- | --- | --- | --- | --- | --- | --- | --- | --- | --- | --- | --- | --- | --- | --- | --- | --- | --- | --- | --- | --- | --- | --- | --- | --- | --- | --- | --- | --- | --- | --- | --- | --- | --- | --- | --- | --- | --- | --- | --- | --- | --- | --- | --- | --- | --- | --- | --- | --- | --- | --- | --- | --- | --- | --- | --- | --- | --- | --- | --- | --- | --- | --- | --- | --- | --- | --- | --- | --- | --- | --- | --- | --- | --- | --- | --- | --- | --- | --- | --- | --- | --- | --- | --- | --- | --- | --- | --- | --- | --- | --- |
| left anterior | 0.228262 | 0.241107 | 0.242472 | 0.252805 | 0.255303 | 0.254441 | 0.246952 | 0.246025 | 0.303971 | 0.399541 | 0.443491 | 0.505990 | 0.564095 | 0.471270 | 0.383402 | 0.302542 | 0.267769 | 0.265150 | 0.263435 | 0.266113 | 0.266485 | 0.263927 | 0.242272 | 0.214709 | 0.268506 | 0.424323 | 0.934606 | 2.096433 | 3.550726 | 3.678976 | 1.803227 | 0.468961 | 0.217136 | 0.443652 | 2.070001 | 5.302887 | 6.640892 | 5.297336 | 3.549094 | 2.557044 | 2.199107 | 2.409976 | 5.264337 | 11.999935 | 35.026007 | 32.496629 | 9.829875 | 1.988482 | 0.771527 | 0.398804 | 0.314200 | 0.297147 | 0.390863 | 0.580787 | 0.906272 | 1.056085 | 1.037710 | 0.723026 | 0.570656 | 0.472017 | 0.363027 | 0.287797 | 0.247702 | 0.216718 | 0.229709 | 0.299876 | 0.458661 | 0.718708 | 1.190377 | 1.430060 | 1.560482 | 1.382951 | 1.322206 | 1.083653 | 0.811811 | 0.474019 | 0.296683 | 0.218827 | 0.242020 | 0.414202 | 0.789424 | 0.993796 | 0.841747 | 0.642632 | 0.436851 | 0.301481 | 0.244612 | 0.219903 | 0.214772 | 0.221852 | 0.234872 | 0.251989 | 0.265061 | 0.289562 | 0.298778 | 2.728385e-01 | 0.237917 | 0.218178 | 0.219944 | 0.260910 | 0.346110 | 0.450809 | 0.516438 | 0.504571 | 0.442556 | 0.356386 | 0.276092 | 0.234127 | 0.217644 | 0.214803 | 0.214782 | 0.214799 | 0.215006 | 0.215293 | 0.216889 | 0.222427 | 0.243575 | 0.285152 | 0.353482 | 0.437652 | 0.476328 | 0.464372 | 0.404279 | 0.322086 | 0.264953 | 0.235591 | 0.219775 | 0.214874 | 0.214681 | 0.214809 | 0.214680 | 0.215189 | 0.215688 | 0.216933 | 0.219280 | 0.217917 | 0.215848 | 0.214824 | 0.214701 | 0.215367 | 0.216598 | 0.217561 | 0.218233 | 0.217426 | 0.216671 | 0.214730 | 0.216287 | 0.218542 | 0.217038 | 0.215059 | 0.214676 | 0.216803 | 0.221370 | 0.227980 | 0.229356 | 0.228566 | 0.223026 | 0.216730 | 0.214629 | 0.216809 | 0.220814 | 0.221058 | 0.218338 | 0.216230 | 0.214949 | 0.214724 | 0.214996 | 0.217131 | 0.225767 | 0.236830 | 0.240978 | 0.230006 | 0.219306 | 0.214758 | 0.221458 | 0.233074 | 0.229982 | 0.220846 | 0.214626 | 0.220512 | 0.240444 | 0.257832 | 0.263926 | 0.256573 | 0.251339 | 0.240890 | 0.229763 | 0.220682 | 0.217555 | 0.215150 | 0.215895 | 0.225855 | 0.239719 | 0.249218 | 0.241146 | 0.218769 | 0.228074 | 0.339720 | 0.607632 | 0.983239 | 1.404935 | 1.487058 | 1.044931 | 0.544875 | 0.326159 | 0.245356 | 0.221868 | 0.218324 | 0.225290 | 0.249159 | 0.297295 | 0.357516 | 0.441189 | 0.574852 | 0.750981 | 0.928649 | 1.105756 | 1.156556 | 1.106283 | 1.018348 | 0.860040 | 0.691589 | 0.577647 | 0.499363 | 0.427352 | 0.367716 | 0.335373 | 0.313714 | 0.303922 | 0.289504 | 0.275846 | 0.263586 | 0.253335 | 0.237416 | 0.233745 | 0.233514 | 0.231499 | 0.228605 | 0.225404 | 0.220154 | 0.217368 | 0.215572 | 0.214866 | 0.217408 | 0.218050 | 0.216895 | 0.214904 | 0.215951 | 0.222846 | 0.232433 | 0.245056 | 0.254090 | 0.262579 | 0.275033 | 0.279674 | 0.268981 | 0.261642 | 0.253903 | 0.243426 | 0.237616 | 0.238349 | 0.245564 | 0.265938 | 0.301484 | 0.343744 | 0.396071 | 0.446571 | 0.461942 | 0.447217 | 0.429772 | 0.399629 | 0.362944 | 0.337458 | 0.319762 | 0.313940 | 0.318277 | 0.343355 | 0.393956 | 0.423515 | 0.467732 |
| right anterior | 1.197065 | 1.031406 | 0.685439 | 0.425070 | 0.297295 | 0.233166 | 0.219236 | 0.215343 | 0.214636 | 0.217672 | 0.249483 | 0.418717 | 0.720974 | 1.429903 | 1.847940 | 1.511654 | 0.998187 | 0.528156 | 0.266059 | 0.214783 | 0.237315 | 0.320687 | 0.467199 | 0.667995 | 0.959070 | 1.407470 | 1.678118 | 1.123596 | 0.515257 | 0.284093 | 0.215396 | 0.273147 | 0.427165 | 0.536088 | 0.501830 | 0.408105 | 0.307163 | 0.263721 | 0.228281 | 0.223126 | 0.231534 | 0.252839 | 0.280098 | 0.337931 | 0.354094 | 0.350982 | 0.302865 | 0.247091 | 0.217500 | 0.216513 | 0.229859 | 0.242467 | 0.255740 | 0.262442 | 0.275392 | 0.274209 | 0.271648 | 0.249254 | 0.233047 | 0.219406 | 0.215027 | 0.215526 | 0.214798 | 0.234255 | 0.350765 | 0.741874 | 1.789290 | 2.648375 | 2.663938 | 2.104748 | 1.691380 | 1.328699 | 1.049610 | 0.789888 | 0.626972 | 0.552521 | 0.497869 | 0.438603 | 0.388045 | 0.320989 | 0.260169 | 0.236716 | 0.234230 | 0.241718 | 0.264032 | 0.307378 | 0.371479 | 0.444262 | 0.579553 | 0.861666 | 1.478761 | 2.711911 | 5.574279 | 11.300574 | 20.329975 | 2.401210e+01 | 19.242826 | 13.005199 | 8.611905 | 4.767183 | 2.986326 | 2.000903 | 1.338675 | 0.978669 | 0.794254 | 0.684793 | 0.650614 | 0.673851 | 0.728163 | 0.828695 | 0.987378 | 1.217157 | 1.361020 | 1.553681 | 1.842915 | 2.010020 | 2.226079 | 2.480414 | 2.451521 | 2.560429 | 2.744698 | 2.627503 | 2.336415 | 1.825667 | 1.332317 | 1.012333 | 0.776355 | 0.602296 | 0.503074 | 0.449232 | 0.387048 | 0.334069 | 0.302102 | 0.295095 | 0.288933 | 0.280164 | 0.267213 | 0.249882 | 0.228057 | 0.215261 | 0.217891 | 0.232244 | 0.246034 | 0.240409 | 0.224101 | 0.214716 | 0.221626 | 0.241765 | 0.264889 | 0.302844 | 0.326851 | 0.340743 | 0.337353 | 0.314769 | 0.275290 | 0.246434 | 0.223017 | 0.214702 | 0.222198 | 0.250342 | 0.295398 | 0.333490 | 0.385520 | 0.449297 | 0.510197 | 0.576332 | 0.669974 | 0.750591 | 0.813512 | 0.830110 | 0.829460 | 0.743381 | 0.582010 | 0.453054 | 0.381814 | 0.338927 | 0.307502 | 0.297949 | 0.303864 | 0.317919 | 0.340129 | 0.412493 | 0.579792 | 0.834760 | 1.118035 | 1.226611 | 1.072554 | 0.810129 | 0.606321 | 0.461164 | 0.415414 | 0.386747 | 0.361368 | 0.344752 | 0.330393 | 0.299546 | 0.268742 | 0.245648 | 0.236674 | 0.242878 | 0.264476 | 0.305272 | 0.373188 | 0.434705 | 0.441299 | 0.388441 | 0.327864 | 0.283027 | 0.252573 | 0.232991 | 0.225796 | 0.224771 | 0.224307 | 0.223014 | 0.220183 | 0.219150 | 0.220041 | 0.218863 | 0.217882 | 0.218236 | 0.216560 | 0.214708 | 0.215089 | 0.218084 | 0.218410 | 0.215471 | 0.214760 | 0.216422 | 0.217706 | 0.214633 | 0.223079 | 0.262016 | 0.342517 | 0.455109 | 0.511836 | 0.490192 | 0.375472 | 0.278008 | 0.229502 | 0.218486 | 0.218412 | 0.228407 | 0.259264 | 0.308574 | 0.359176 | 0.384655 | 0.388279 | 0.346776 | 0.308188 | 0.269593 | 0.246057 | 0.231831 | 0.228320 | 0.225445 | 0.233367 | 0.254829 | 0.294258 | 0.344432 | 0.416189 | 0.472019 | 0.521101 | 0.559976 | 0.594166 | 0.606712 | 0.577688 | 0.507815 | 0.399004 | 0.291276 | 0.228602 | 0.215380 | 0.240170 | 0.278744 | 0.314092 | 0.337686 | 0.345284 | 0.346328 | 0.332749 | 0.318818 | 0.310463 | 0.292939 |
| left central | 0.283468 | 0.293445 | 0.306435 | 0.339086 | 0.363213 | 0.367958 | 0.329908 | 0.285600 | 0.259890 | 0.239246 | 0.223457 | 0.216313 | 0.216664 | 0.246087 | 0.338356 | 0.424688 | 0.481788 | 0.403519 | 0.284254 | 0.245142 | 0.226347 | 0.214654 | 0.228714 | 0.262256 | 0.358344 | 0.468316 | 0.587754 | 0.532753 | 0.390603 | 0.261939 | 0.217200 | 0.224377 | 0.240413 | 0.232356 | 0.215828 | 0.219384 | 0.243354 | 0.276400 | 0.309388 | 0.330403 | 0.313851 | 0.269963 | 0.228832 | 0.214632 | 0.224235 | 0.239608 | 0.247056 | 0.235629 | 0.224623 | 0.217844 | 0.215003 | 0.214663 | 0.214734 | 0.215553 | 0.215997 | 0.218659 | 0.223736 | 0.228775 | 0.227282 | 0.226105 | 0.221453 | 0.218901 | 0.217138 | 0.217941 | 0.226697 | 0.289876 | 0.565153 | 1.955803 | 8.600148 | 29.866616 | 66.239704 | 92.466291 | 80.678515 | 54.072005 | 31.728950 | 14.825734 | 7.203602 | 3.909720 | 2.330023 | 1.615323 | 1.285923 | 0.982406 | 0.840240 | 0.808341 | 0.780061 | 0.773624 | 0.772319 | 0.710184 | 0.724908 | 0.746116 | 0.986267 | 1.594205 | 2.785331 | 4.109874 | 5.922095 | 5.951043e+00 | 4.217976 | 2.197221 | 0.957318 | 0.462167 | 0.300570 | 0.244395 | 0.225356 | 0.223433 | 0.225557 | 0.233256 | 0.251356 | 0.282590 | 0.295887 | 0.288154 | 0.266238 | 0.247917 | 0.236850 | 0.234994 | 0.250217 | 0.291442 | 0.333994 | 0.384652 | 0.468395 | 0.571405 | 0.642341 | 0.672442 | 0.645453 | 0.622263 | 0.612950 | 0.646842 | 0.686201 | 0.873647 | 1.158485 | 1.431959 | 1.667238 | 1.908374 | 1.468645 | 0.917701 | 0.570802 | 0.390657 | 0.322273 | 0.289088 | 0.259457 | 0.237565 | 0.217610 | 0.223303 | 0.295807 | 0.515214 | 1.070198 | 1.810401 | 1.926184 | 1.492450 | 0.982604 | 0.624533 | 0.429037 | 0.336540 | 0.293000 | 0.278408 | 0.276703 | 0.284164 | 0.295362 | 0.292416 | 0.279979 | 0.262070 | 0.244458 | 0.236094 | 0.237902 | 0.250209 | 0.282737 | 0.325252 | 0.355917 | 0.385685 | 0.388581 | 0.375613 | 0.367740 | 0.383241 | 0.402812 | 0.446973 | 0.511441 | 0.629804 | 0.739855 | 0.826022 | 0.681963 | 0.536639 | 0.426802 | 0.351223 | 0.288303 | 0.263363 | 0.249135 | 0.253572 | 0.263006 | 0.286979 | 0.316502 | 0.361706 | 0.381804 | 0.413873 | 0.479834 | 0.565120 | 0.587640 | 0.539197 | 0.434729 | 0.309823 | 0.255784 | 0.232261 | 0.223350 | 0.220918 | 0.224395 | 0.236011 | 0.264301 | 0.308540 | 0.347379 | 0.353923 | 0.324948 | 0.287274 | 0.255930 | 0.240736 | 0.234589 | 0.230378 | 0.229476 | 0.230812 | 0.228794 | 0.227889 | 0.228138 | 0.224176 | 0.217958 | 0.214762 | 0.216448 | 0.222883 | 0.235404 | 0.257377 | 0.286713 | 0.331532 | 0.425602 | 0.618056 | 0.988538 | 1.387566 | 1.542330 | 1.401558 | 0.991077 | 0.596300 | 0.365021 | 0.266109 | 0.233721 | 0.223284 | 0.220436 | 0.221500 | 0.222986 | 0.224176 | 0.218930 | 0.214723 | 0.217262 | 0.229087 | 0.259486 | 0.311739 | 0.394179 | 0.481900 | 0.585501 | 0.709143 | 0.734519 | 0.637450 | 0.511834 | 0.396157 | 0.336035 | 0.303047 | 0.293224 | 0.306367 | 0.341827 | 0.392888 | 0.476069 | 0.596868 | 0.737648 | 0.896711 | 1.000696 | 0.972468 | 0.871674 | 0.806346 | 0.748922 | 0.793437 | 0.825869 | 0.868077 | 0.927761 | 1.041573 | 0.984202 | 1.009918 |
| right central | 3.776157 | 4.250900 | 5.018193 | 6.111748 | 5.766156 | 3.927363 | 2.757967 | 1.949593 | 1.585472 | 1.348886 | 0.994841 | 0.587938 | 0.338004 | 0.240895 | 0.216145 | 0.214628 | 0.216694 | 0.225827 | 0.249473 | 0.288678 | 0.346337 | 0.400327 | 0.432570 | 0.401039 | 0.324531 | 0.255775 | 0.217563 | 0.218790 | 0.232646 | 0.229111 | 0.216319 | 0.226844 | 0.283421 | 0.380844 | 0.462708 | 0.412726 | 0.281323 | 0.218607 | 0.230855 | 0.266355 | 0.270212 | 0.243424 | 0.219616 | 0.222586 | 0.281990 | 0.458938 | 0.707125 | 0.881141 | 0.690915 | 0.490493 | 0.327189 | 0.245619 | 0.215523 | 0.228094 | 0.321581 | 0.620518 | 1.478957 | 2.800097 | 2.633074 | 1.148355 | 0.497703 | 0.293189 | 0.232592 | 0.214895 | 0.219995 | 0.250497 | 0.327778 | 0.605320 | 1.861118 | 9.745993 | 46.696672 | 165.184804 | 257.895024 | 146.528880 | 52.595221 | 14.852929 | 4.593329 | 1.793068 | 0.850590 | 0.518301 | 0.495832 | 0.680811 | 1.786431 | 7.286883 | 29.464761 | 80.774119 | 107.784123 | 77.107964 | 61.778258 | 66.519161 | 121.848722 | 402.334608 | 2602.183169 | 32664.003080 | 576235.214903 | 1.677661e+06 | 811376.278998 | 104823.465525 | 12767.627011 | 2791.098969 | 1268.181853 | 1146.629430 | 1011.473337 | 749.263030 | 381.480780 | 163.877944 | 59.675445 | 30.823207 | 20.096329 | 17.494395 | 13.925626 | 13.379277 | 15.594229 | 23.383045 | 27.880768 | 30.983331 | 27.343856 | 25.230192 | 22.645788 | 24.256297 | 31.310722 | 59.160262 | 101.983328 | 186.389362 | 357.246188 | 508.977589 | 433.114089 | 304.625823 | 207.366369 | 180.369865 | 197.587165 | 269.894942 | 340.704995 | 327.861247 | 185.720610 | 90.168718 | 35.905595 | 12.254185 | 3.778034 | 1.600275 | 0.884237 | 0.634957 | 0.542983 | 0.512224 | 0.475173 | 0.436181 | 0.415136 | 0.434117 | 0.503260 | 0.590059 | 0.658477 | 0.701219 | 0.661794 | 0.632343 | 0.610538 | 0.680768 | 0.952888 | 1.610653 | 3.380949 | 8.639016 | 19.133397 | 23.381088 | 14.810448 | 5.510724 | 2.406584 | 1.294739 | 1.017242 | 0.980749 | 1.132558 | 1.128295 | 1.082940 | 0.881592 | 0.757004 | 0.633056 | 0.553967 | 0.473434 | 0.465921 | 0.473460 | 0.473637 | 0.457465 | 0.432971 | 0.397090 | 0.379009 | 0.368839 | 0.371190 | 0.394058 | 0.413620 | 0.434801 | 0.472713 | 0.504132 | 0.538739 | 0.595979 | 0.655016 | 0.684958 | 0.685217 | 0.606695 | 0.482343 | 0.396207 | 0.361171 | 0.353832 | 0.375970 | 0.418969 | 0.474030 | 0.524013 | 0.562045 | 0.493928 | 0.393690 | 0.298693 | 0.239731 | 0.217837 | 0.214991 | 0.216223 | 0.232490 | 0.279290 | 0.333593 | 0.419062 | 0.469499 | 0.433269 | 0.357512 | 0.303095 | 0.259691 | 0.246765 | 0.247070 | 0.272014 | 0.331065 | 0.431535 | 0.512412 | 0.573502 | 0.587109 | 0.536114 | 0.445529 | 0.355583 | 0.290222 | 0.250269 | 0.235544 | 0.229974 | 0.227358 | 0.223197 | 0.217702 | 0.215363 | 0.230956 | 0.275866 | 0.367771 | 0.506103 | 0.637653 | 0.715450 | 0.643856 | 0.487738 | 0.364071 | 0.275150 | 0.242878 | 0.231536 | 0.222279 | 0.217966 | 0.215858 | 0.214669 | 0.216348 | 0.227478 | 0.255350 | 0.284566 | 0.322121 | 0.367838 | 0.349725 | 0.324426 | 0.299189 | 0.274909 | 0.254213 | 0.234767 | 0.221043 | 0.215175 | 0.215941 | 0.227489 | 0.256383 | 0.310094 | 0.380231 | 0.486447 | 0.565007 | 0.564698 | 0.542796 | 0.512121 |
| left posterior | 1.231698 | 1.760161 | 2.586869 | 6.040886 | 11.590309 | 12.454009 | 5.487213 | 1.843559 | 0.553175 | 0.257740 | 0.214677 | 0.245445 | 0.300785 | 0.302511 | 0.255098 | 0.214720 | 0.378382 | 2.831455 | 41.470560 | 119.987579 | 31.255342 | 3.799864 | 0.499639 | 0.214684 | 0.480245 | 1.830879 | 3.046385 | 1.280186 | 0.510728 | 0.243399 | 0.219396 | 0.251647 | 0.230404 | 0.214913 | 0.231691 | 0.277646 | 0.338636 | 0.380355 | 0.394856 | 0.365355 | 0.314270 | 0.275164 | 0.268721 | 0.293134 | 0.337865 | 0.379742 | 0.382254 | 0.369660 | 0.339516 | 0.298467 | 0.259806 | 0.229696 | 0.214903 | 0.223690 | 0.255969 | 0.301623 | 0.342903 | 0.379048 | 0.419410 | 0.395357 | 0.293550 | 0.224480 | 0.229592 | 0.424759 | 1.622487 | 10.069562 | 70.392322 | 370.260673 | 1508.366676 | 5462.708875 | 15633.404059 | 36774.520757 | 64230.605029 | 84294.701690 | 73188.454244 | 49391.620297 | 20974.289795 | 9727.685153 | 5257.989269 | 2193.421900 | 506.955158 | 73.487578 | 13.863097 | 4.565308 | 2.349430 | 1.639872 | 1.386177 | 1.155458 | 1.050862 | 1.222813 | 1.656653 | 2.454742 | 4.135122 | 6.675412 | 10.290507 | 1.523418e+01 | 17.417572 | 13.850725 | 9.232109 | 5.857499 | 4.120591 | 3.254523 | 3.070948 | 3.469978 | 4.497760 | 6.598924 | 10.477370 | 16.463741 | 22.257687 | 23.318441 | 17.784938 | 11.754537 | 7.095293 | 4.797318 | 3.622289 | 3.267191 | 3.665251 | 5.147669 | 8.521070 | 15.921625 | 31.270307 | 55.915960 | 79.779093 | 87.635694 | 73.811167 | 39.534788 | 16.985236 | 6.031535 | 2.234905 | 1.008804 | 0.638749 | 0.491287 | 0.461737 | 0.470999 | 0.510035 | 0.573650 | 0.685044 | 0.820533 | 0.933375 | 1.034252 | 1.089187 | 1.054810 | 1.070261 | 1.121921 | 1.152863 | 1.054285 | 0.833955 | 0.601340 | 0.451979 | 0.350511 | 0.293573 | 0.260605 | 0.241073 | 0.231803 | 0.227641 | 0.228424 | 0.235583 | 0.250862 | 0.269448 | 0.294597 | 0.317497 | 0.343046 | 0.358064 | 0.357971 | 0.345138 | 0.328292 | 0.307990 | 0.291651 | 0.278341 | 0.263540 | 0.251193 | 0.244588 | 0.241782 | 0.245328 | 0.259665 | 0.282432 | 0.313346 | 0.359712 | 0.411251 | 0.462445 | 0.514567 | 0.554496 | 0.566392 | 0.588023 | 0.593155 | 0.566253 | 0.517133 | 0.464371 | 0.402844 | 0.353656 | 0.326505 | 0.307037 | 0.298034 | 0.301222 | 0.326391 | 0.373211 | 0.475574 | 0.642341 | 0.907351 | 1.198932 | 1.492186 | 1.581834 | 1.562676 | 1.405896 | 1.229067 | 1.072994 | 0.958437 | 0.888189 | 0.879716 | 0.978472 | 1.192667 | 1.642144 | 2.416851 | 3.911060 | 5.901700 | 8.806687 | 11.981536 | 14.012769 | 12.223204 | 9.518566 | 5.941321 | 3.590927 | 2.162592 | 1.408676 | 1.027945 | 0.889472 | 0.777696 | 0.724600 | 0.692786 | 0.677980 | 0.669058 | 0.755420 | 0.977790 | 1.542399 | 2.910239 | 6.349458 | 11.970354 | 20.435572 | 25.506697 | 23.173506 | 17.069661 | 10.584958 | 5.095025 | 2.825033 | 1.904599 | 1.649825 | 1.911550 | 2.561913 | 3.106231 | 3.284085 | 2.890655 | 2.235648 | 1.687125 | 1.283203 | 0.990322 | 0.836579 | 0.687182 | 0.591915 | 0.526578 | 0.458611 | 0.380791 | 0.320401 | 0.275113 | 0.248607 | 0.233301 | 0.225127 | 0.221382 | 0.219861 | 0.218772 | 0.217496 | 0.215355 | 0.214627 | 0.216805 | 0.221995 | 0.227261 | 0.227530 | 0.226504 | 0.223020 | 0.222776 | 0.220198 |
| right posterior | 1.459347 | 1.692970 | 2.036121 | 2.393837 | 2.445382 | 2.146409 | 1.620176 | 1.027284 | 0.549964 | 0.331198 | 0.256582 | 0.227542 | 0.217795 | 0.217639 | 0.232815 | 0.255551 | 0.294558 | 0.309694 | 0.284025 | 0.238129 | 0.216031 | 0.231561 | 0.264912 | 0.264115 | 0.238092 | 0.214645 | 0.254312 | 0.398528 | 0.762687 | 1.523842 | 2.595124 | 2.054203 | 0.968248 | 0.410734 | 0.228101 | 0.227902 | 0.312567 | 0.380160 | 0.386551 | 0.323670 | 0.283866 | 0.252503 | 0.242053 | 0.237753 | 0.238340 | 0.234368 | 0.221024 | 0.218932 | 0.310115 | 0.582752 | 0.978550 | 1.081519 | 0.897255 | 0.627347 | 0.403614 | 0.302050 | 0.271953 | 0.265005 | 0.265512 | 0.265677 | 0.237505 | 0.214757 | 0.295737 | 0.854889 | 4.097249 | 22.795969 | 107.925831 | 347.975826 | 747.277644 | 1374.174176 | 1678.655808 | 1408.531133 | 783.716920 | 276.713473 | 81.504735 | 27.135181 | 9.233195 | 3.394701 | 1.468487 | 0.667964 | 0.363780 | 0.269279 | 0.245935 | 0.246512 | 0.261430 | 0.299161 | 0.343812 | 0.411672 | 0.526880 | 0.741755 | 1.090725 | 1.838878 | 3.009558 | 4.601370 | 6.505598 | 7.669867e+00 | 7.437557 | 6.398006 | 5.261797 | 4.167243 | 3.790497 | 3.590429 | 3.606343 | 3.904364 | 4.221043 | 4.123319 | 4.066594 | 3.993723 | 3.563439 | 3.020373 | 2.475127 | 2.005138 | 1.803050 | 1.966235 | 2.331179 | 3.347719 | 6.075419 | 12.621559 | 30.060249 | 74.813028 | 158.387976 | 252.782945 | 261.763119 | 178.816528 | 93.585321 | 40.995192 | 16.537514 | 7.381693 | 3.744115 | 2.248947 | 1.566135 | 1.300039 | 1.202077 | 1.177756 | 1.243393 | 1.445608 | 1.665899 | 1.940578 | 2.204089 | 2.541838 | 2.876786 | 3.057211 | 3.290644 | 3.967259 | 4.932524 | 5.582038 | 5.247225 | 4.424898 | 3.396713 | 2.472757 | 1.775254 | 1.410032 | 1.194680 | 1.164016 | 1.325921 | 1.869862 | 2.951983 | 5.130308 | 8.474451 | 12.265848 | 13.725372 | 13.614640 | 11.377285 | 8.568833 | 5.984872 | 4.209738 | 2.919336 | 2.289926 | 1.856084 | 1.664547 | 1.672563 | 1.883154 | 2.155116 | 2.801663 | 3.554519 | 4.499189 | 4.865884 | 4.488188 | 3.533618 | 2.515582 | 1.625968 | 1.148919 | 0.897061 | 0.808380 | 0.881507 | 1.151919 | 1.778910 | 2.976426 | 4.572345 | 5.273490 | 4.774625 | 3.567263 | 2.828467 | 2.568041 | 2.539684 | 2.770960 | 3.017208 | 3.085419 | 3.249362 | 3.697756 | 4.524579 | 5.919559 | 7.800553 | 10.225194 | 12.500083 | 13.607742 | 13.665561 | 12.920451 | 11.759131 | 10.380770 | 9.493845 | 9.398393 | 10.727688 | 14.403487 | 21.576706 | 37.003898 | 61.531026 | 78.935220 | 82.207830 | 72.176362 | 45.844547 | 26.000045 | 15.243820 | 9.419333 | 6.839575 | 5.978061 | 5.643554 | 5.738435 | 6.332981 | 6.788679 | 7.509130 | 8.901169 | 11.781580 | 15.717456 | 19.516069 | 23.073548 | 25.060155 | 25.416471 | 24.071991 | 22.519219 | 17.857462 | 13.612293 | 10.271539 | 8.356903 | 7.592688 | 8.159414 | 9.879924 | 14.674736 | 20.903227 | 25.456082 | 25.889597 | 21.281387 | 13.720027 | 7.991143 | 4.745899 | 3.240595 | 2.585165 | 2.322783 | 2.358610 | 2.721295 | 3.168300 | 3.410939 | 3.322639 | 2.713066 | 2.018682 | 1.364221 | 0.924638 | 0.679076 | 0.572796 | 0.490585 | 0.460262 | 0.459430 | 0.473526 | 0.517848 | 0.637862 | 0.791361 | 1.037914 | 1.365473 | 1.528651 | 1.667790 |
| all electrodes | 0.432037 | 0.524746 | 0.532032 | 0.473049 | 0.404293 | 0.359464 | 0.266675 | 0.227874 | 0.216003 | 0.281874 | 0.427814 | 0.467046 | 0.483128 | 0.385399 | 0.304586 | 0.251492 | 0.223912 | 0.214730 | 0.223332 | 0.255706 | 0.268979 | 0.234340 | 0.215376 | 0.252947 | 0.415858 | 0.679080 | 1.260723 | 1.206256 | 0.908387 | 0.573141 | 0.430182 | 0.287375 | 0.254511 | 0.259956 | 0.304201 | 0.355968 | 0.423221 | 0.385599 | 0.313847 | 0.236874 | 0.217265 | 0.292403 | 0.549701 | 1.243745 | 1.661980 | 1.216915 | 0.587493 | 0.319532 | 0.260635 | 0.245542 | 0.246726 | 0.265098 | 0.318013 | 0.385047 | 0.501744 | 0.581929 | 0.552313 | 0.449145 | 0.373228 | 0.314551 | 0.257633 | 0.218729 | 0.227001 | 0.327337 | 0.737561 | 2.359021 | 12.031831 | 84.283934 | 633.078030 | 3310.637073 | 13300.911318 | 38839.791840 | 85292.344896 | 92916.544972 | 57358.788344 | 15143.797315 | 2165.387381 | 311.212336 | 66.491035 | 26.389547 | 16.948706 | 15.089302 | 12.909684 | 7.974573 | 3.664441 | 1.979379 | 1.205496 | 0.797138 | 0.622320 | 0.544560 | 0.585070 | 0.718362 | 1.126201 | 2.324441 | 6.525706 | 1.419825e+01 | 22.895131 | 16.240337 | 8.132781 | 3.525710 | 1.715932 | 0.897970 | 0.604551 | 0.498980 | 0.472542 | 0.519047 | 0.605369 | 0.806793 | 1.267479 | 2.456326 | 5.534012 | 14.680631 | 41.712644 | 92.724179 | 189.478407 | 248.688673 | 198.279762 | 155.607109 | 155.400154 | 150.304083 | 203.078131 | 310.990652 | 471.149722 | 487.928546 | 356.034553 | 190.179158 | 119.553543 | 70.043551 | 42.822885 | 24.974233 | 15.881796 | 12.427467 | 11.112268 | 9.765434 | 10.016828 | 11.457432 | 12.283967 | 11.797512 | 11.768187 | 10.391984 | 8.145040 | 5.490691 | 3.386948 | 2.467414 | 2.133550 | 1.985546 | 1.907441 | 2.049829 | 2.104026 | 2.094610 | 1.977720 | 1.760920 | 1.474889 | 1.122802 | 0.792512 | 0.556517 | 0.424869 | 0.349282 | 0.310617 | 0.273045 | 0.244431 | 0.224550 | 0.215396 | 0.216375 | 0.225615 | 0.246622 | 0.276958 | 0.323950 | 0.380572 | 0.428817 | 0.444407 | 0.463422 | 0.445315 | 0.431785 | 0.404536 | 0.348688 | 0.283179 | 0.239029 | 0.217696 | 0.216661 | 0.232088 | 0.253833 | 0.254092 | 0.242306 | 0.227369 | 0.219063 | 0.214685 | 0.217310 | 0.225749 | 0.230040 | 0.234679 | 0.232293 | 0.223678 | 0.217522 | 0.214964 | 0.215357 | 0.217591 | 0.220600 | 0.227278 | 0.234534 | 0.240281 | 0.249999 | 0.260183 | 0.266684 | 0.264574 | 0.255302 | 0.252464 | 0.265398 | 0.290198 | 0.330591 | 0.384894 | 0.446483 | 0.455694 | 0.410359 | 0.342994 | 0.298209 | 0.272430 | 0.273424 | 0.293660 | 0.323341 | 0.337465 | 0.329143 | 0.310056 | 0.298864 | 0.299344 | 0.315195 | 0.370244 | 0.450720 | 0.538479 | 0.637915 | 0.705368 | 0.634725 | 0.587091 | 0.562408 | 0.591165 | 0.686782 | 0.828148 | 1.032470 | 1.364989 | 1.619625 | 1.551556 | 1.277838 | 0.963997 | 0.714197 | 0.548090 | 0.493409 | 0.490357 | 0.484343 | 0.493093 | 0.525924 | 0.478148 | 0.413536 | 0.339846 | 0.272110 | 0.228676 | 0.214707 | 0.238560 | 0.281634 | 0.311666 | 0.312517 | 0.273580 | 0.230180 | 0.215369 | 0.216454 | 0.220434 | 0.220130 | 0.214994 | 0.221541 | 0.261168 | 0.337233 | 0.420601 | 0.509930 | 0.571297 | 0.632402 | 0.637220 | 0.662538 | 0.654805 | 0.640330 | 0.560811 | 0.558746 |

V) neural network features, neutral vs happy

  
|  | time window | peak latency | cluster *p* | peak Cohen's *d* |  | | | |
| **all electrodes** |  | | | |  | | | |
|  | | | | | | | | |

Model correlations, cluster permutation tests

|  | **left hemisphere** | | | | **right hemisphere** | | | |
|  | time window | peak latency | cluster *p* | peak Cohen's *d* | time window | peak latency | cluster *p* | peak Cohen's *d* |
| **anterior** |  | | | | 225 - 415 ms | 280 ms | 0.0049 | 1.0497 |
| **central** |  | | | |  | | | |
| **posterior** | 245 - 415 ms | 280 ms | 0.0126 | 0.8862 | 255 - 455 ms | 290 ms | 0.0117 | 0.7362 |
  | | | | 800 - 970 ms | 845 ms | 0.0275 | 0.6353 |  | | | | 995 - 1125 ms | 1050 ms | 0.0459 | 0.6343 |

  

Model correlations, Bayesian statistics

|  | -200 | -195 | -190 | -185 | -180 | -175 | -170 | -165 | -160 | -155 | -150 | -145 | -140 | -135 | -130 | -125 | -120 | -115 | -110 | -105 | -100 | -95 | -90 | -85 | -80 | -75 | -70 | -65 | -60 | -55 | -50 | -45 | -40 | -35 | -30 | -25 | -20 | -15 | -10 | -5 | 0 | 5 | 10 | 15 | 20 | 25 | 30 | 35 | 40 | 45 | 50 | 55 | 60 | 65 | 70 | 75 | 80 | 85 | 90 | 95 | 100 | 105 | 110 | 115 | 120 | 125 | 130 | 135 | 140 | 145 | 150 | 155 | 160 | 165 | 170 | 175 | 180 | 185 | 190 | 195 | 200 | 205 | 210 | 215 | 220 | 225 | 230 | 235 | 240 | 245 | 250 | 255 | 260 | 265 | 270 | 275 | 280 | 285 | 290 | 295 | 300 | 305 | 310 | 315 | 320 | 325 | 330 | 335 | 340 | 345 | 350 | 355 | 360 | 365 | 370 | 375 | 380 | 385 | 390 | 395 | 400 | 405 | 410 | 415 | 420 | 425 | 430 | 435 | 440 | 445 | 450 | 455 | 460 | 465 | 470 | 475 | 480 | 485 | 490 | 495 | 500 | 505 | 510 | 515 | 520 | 525 | 530 | 535 | 540 | 545 | 550 | 555 | 560 | 565 | 570 | 575 | 580 | 585 | 590 | 595 | 600 | 605 | 610 | 615 | 620 | 625 | 630 | 635 | 640 | 645 | 650 | 655 | 660 | 665 | 670 | 675 | 680 | 685 | 690 | 695 | 700 | 705 | 710 | 715 | 720 | 725 | 730 | 735 | 740 | 745 | 750 | 755 | 760 | 765 | 770 | 775 | 780 | 785 | 790 | 795 | 800 | 805 | 810 | 815 | 820 | 825 | 830 | 835 | 840 | 845 | 850 | 855 | 860 | 865 | 870 | 875 | 880 | 885 | 890 | 895 | 900 | 905 | 910 | 915 | 920 | 925 | 930 | 935 | 940 | 945 | 950 | 955 | 960 | 965 | 970 | 975 | 980 | 985 | 990 | 995 | 1000 | 1005 | 1010 | 1015 | 1020 | 1025 | 1030 | 1035 | 1040 | 1045 | 1050 | 1055 | 1060 | 1065 | 1070 | 1075 | 1080 | 1085 | 1090 | 1095 | 1100 | 1105 | 1110 | 1115 | 1120 | 1125 | 1130 | 1135 | 1140 | 1145 | 1150 | 1155 | 1160 | 1165 | 1170 | 1175 | 1180 | 1185 | 1190 | 1195 |
| --- | --- | --- | --- | --- | --- | --- | --- | --- | --- | --- | --- | --- | --- | --- | --- | --- | --- | --- | --- | --- | --- | --- | --- | --- | --- | --- | --- | --- | --- | --- | --- | --- | --- | --- | --- | --- | --- | --- | --- | --- | --- | --- | --- | --- | --- | --- | --- | --- | --- | --- | --- | --- | --- | --- | --- | --- | --- | --- | --- | --- | --- | --- | --- | --- | --- | --- | --- | --- | --- | --- | --- | --- | --- | --- | --- | --- | --- | --- | --- | --- | --- | --- | --- | --- | --- | --- | --- | --- | --- | --- | --- | --- | --- | --- | --- | --- | --- | --- | --- | --- | --- | --- | --- | --- | --- | --- | --- | --- | --- | --- | --- | --- | --- | --- | --- | --- | --- | --- | --- | --- | --- | --- | --- | --- | --- | --- | --- | --- | --- | --- | --- | --- | --- | --- | --- | --- | --- | --- | --- | --- | --- | --- | --- | --- | --- | --- | --- | --- | --- | --- | --- | --- | --- | --- | --- | --- | --- | --- | --- | --- | --- | --- | --- | --- | --- | --- | --- | --- | --- | --- | --- | --- | --- | --- | --- | --- | --- | --- | --- | --- | --- | --- | --- | --- | --- | --- | --- | --- | --- | --- | --- | --- | --- | --- | --- | --- | --- | --- | --- | --- | --- | --- | --- | --- | --- | --- | --- | --- | --- | --- | --- | --- | --- | --- | --- | --- | --- | --- | --- | --- | --- | --- | --- | --- | --- | --- | --- | --- | --- | --- | --- | --- | --- | --- | --- | --- | --- | --- | --- | --- | --- | --- | --- | --- | --- | --- | --- | --- | --- | --- | --- | --- | --- | --- | --- | --- | --- | --- | --- | --- | --- | --- | --- | --- | --- | --- | --- | --- | --- | --- | --- | --- | --- | --- | --- | --- | --- | --- | --- | --- |
| left anterior | 0.250252 | 0.256797 | 0.247358 | 0.235113 | 0.220832 | 0.214808 | 0.228490 | 0.248885 | 0.249095 | 0.233122 | 0.227490 | 0.218948 | 0.217389 | 0.221332 | 0.246780 | 0.297593 | 0.377474 | 0.388734 | 0.351921 | 0.304224 | 0.251375 | 0.215673 | 0.227829 | 0.242095 | 0.227572 | 0.214688 | 0.258254 | 0.358596 | 0.431207 | 0.434691 | 0.339338 | 0.219926 | 0.273987 | 0.740738 | 2.913007 | 4.708832 | 4.095507 | 2.230871 | 1.356779 | 0.889753 | 0.793947 | 0.643769 | 0.612408 | 0.540487 | 0.541103 | 0.528718 | 0.460339 | 0.324977 | 0.256182 | 0.226288 | 0.216564 | 0.215696 | 0.225592 | 0.263709 | 0.370158 | 0.582541 | 0.834713 | 0.825580 | 0.631069 | 0.386937 | 0.243724 | 0.216235 | 0.269965 | 0.474495 | 0.823900 | 1.071918 | 1.095597 | 0.972803 | 0.899491 | 0.817615 | 0.711094 | 0.570983 | 0.426677 | 0.285600 | 0.220692 | 0.221178 | 0.268790 | 0.353742 | 0.425945 | 0.479091 | 0.546501 | 0.518360 | 0.397287 | 0.298618 | 0.236147 | 0.216098 | 0.218302 | 0.242538 | 0.289347 | 0.342079 | 0.414688 | 0.468073 | 0.449762 | 0.403886 | 0.373325 | 0.327712 | 0.294415 | 0.269756 | 0.245970 | 0.227694 | 0.217751 | 0.215777 | 0.232190 | 0.269458 | 0.309894 | 0.326658 | 0.283250 | 0.238219 | 0.218452 | 0.214779 | 0.214681 | 0.214655 | 0.215424 | 0.216965 | 0.221440 | 0.225743 | 0.224035 | 0.221160 | 0.218585 | 0.215269 | 0.214664 | 0.214739 | 0.216100 | 0.221808 | 0.232443 | 0.238663 | 0.240257 | 0.237365 | 0.223375 | 0.214639 | 0.224843 | 0.260186 | 0.323864 | 0.407451 | 0.489216 | 0.531221 | 0.480478 | 0.423566 | 0.360480 | 0.319929 | 0.297949 | 0.302153 | 0.319423 | 0.365797 | 0.437000 | 0.491295 | 0.492983 | 0.408752 | 0.316205 | 0.249218 | 0.219238 | 0.218590 | 0.242794 | 0.272868 | 0.284467 | 0.278818 | 0.254565 | 0.229936 | 0.216797 | 0.215005 | 0.218805 | 0.221170 | 0.217292 | 0.214698 | 0.222449 | 0.245804 | 0.278974 | 0.313095 | 0.314522 | 0.293435 | 0.260400 | 0.240745 | 0.224418 | 0.219518 | 0.216753 | 0.215375 | 0.214722 | 0.215744 | 0.220164 | 0.224998 | 0.230744 | 0.228431 | 0.227961 | 0.232807 | 0.253728 | 0.286077 | 0.369314 | 0.509970 | 0.667438 | 0.739050 | 0.711829 | 0.576549 | 0.483415 | 0.408284 | 0.358133 | 0.366448 | 0.473631 | 0.699503 | 1.086804 | 1.502852 | 1.897521 | 2.071539 | 1.774606 | 1.204507 | 0.881378 | 0.686107 | 0.578460 | 0.544672 | 0.547523 | 0.600256 | 0.695118 | 0.794239 | 0.935743 | 1.256799 | 1.625312 | 1.714462 | 1.512850 | 1.098688 | 0.781340 | 0.572732 | 0.440179 | 0.357588 | 0.326511 | 0.313313 | 0.316883 | 0.330390 | 0.377998 | 0.433711 | 0.474903 | 0.472456 | 0.454218 | 0.426766 | 0.407906 | 0.388666 | 0.410937 | 0.476114 | 0.521187 | 0.542213 | 0.528678 | 0.435127 | 0.338861 | 0.267862 | 0.231017 | 0.219640 | 0.217382 | 0.218773 | 0.228235 | 0.242279 | 0.259525 | 0.272572 | 0.298897 | 0.331876 | 0.414027 | 0.578165 | 0.997791 | 1.374146 | 1.471459 | 1.120326 | 0.742716 | 0.455452 | 0.337199 | 0.290838 | 0.298964 | 0.353735 | 0.477648 | 0.653264 | 0.797083 | 0.810764 | 0.716037 | 0.627578 | 0.560414 | 0.491768 | 0.454882 | 0.446737 | 0.421834 | 0.398059 | 0.387229 | 0.375763 | 0.355664 | 0.352371 |
| right anterior | 3.413890 | 4.448856 | 4.970656 | 6.219743 | 3.593721 | 1.389311 | 0.843293 | 0.640547 | 0.505662 | 0.348935 | 0.243106 | 0.216483 | 0.280609 | 0.571225 | 0.846301 | 0.888964 | 0.746955 | 0.496462 | 0.294364 | 0.232844 | 0.216361 | 0.215743 | 0.217850 | 0.218967 | 0.219588 | 0.216411 | 0.219488 | 0.250798 | 0.285701 | 0.312377 | 0.314377 | 0.262019 | 0.227211 | 0.214627 | 0.221724 | 0.239979 | 0.239081 | 0.229591 | 0.215599 | 0.214931 | 0.216355 | 0.215583 | 0.214661 | 0.216927 | 0.221303 | 0.225542 | 0.223011 | 0.223081 | 0.220866 | 0.219008 | 0.216134 | 0.215216 | 0.214634 | 0.215746 | 0.219940 | 0.224526 | 0.225368 | 0.223329 | 0.221446 | 0.219589 | 0.219741 | 0.229994 | 0.298887 | 0.659351 | 1.787898 | 4.172699 | 9.768942 | 11.739717 | 7.112535 | 3.819204 | 2.116198 | 1.476930 | 1.190567 | 0.980316 | 0.900821 | 1.014424 | 1.018497 | 1.029710 | 1.077250 | 0.992652 | 0.769634 | 0.728939 | 0.747790 | 0.775327 | 0.963855 | 1.415354 | 2.320668 | 3.660289 | 6.986773 | 18.094408 | 57.403720 | 159.704768 | 397.412885 | 808.003442 | 1316.785714 | 1190.059515 | 726.900354 | 410.362552 | 239.621460 | 113.764124 | 56.048895 | 30.723622 | 17.381399 | 11.558605 | 9.224669 | 9.260452 | 10.503601 | 13.505614 | 15.798882 | 17.130571 | 16.713749 | 16.940831 | 15.609832 | 13.596673 | 11.717675 | 8.939511 | 7.237707 | 5.857077 | 4.268966 | 2.809213 | 2.307287 | 1.816837 | 1.540640 | 1.277825 | 1.150214 | 1.060742 | 1.056756 | 1.043066 | 1.041229 | 1.086099 | 0.985487 | 0.972584 | 1.072548 | 1.414993 | 1.980266 | 3.292047 | 4.973039 | 6.222614 | 5.123777 | 3.828231 | 2.800715 | 1.926452 | 1.340566 | 0.929736 | 0.604520 | 0.448453 | 0.379233 | 0.349493 | 0.346606 | 0.353603 | 0.332859 | 0.311320 | 0.301090 | 0.290209 | 0.281157 | 0.287317 | 0.303822 | 0.325869 | 0.347862 | 0.358099 | 0.366358 | 0.368793 | 0.348614 | 0.321331 | 0.299851 | 0.271526 | 0.245994 | 0.228373 | 0.218955 | 0.216001 | 0.215188 | 0.214729 | 0.215088 | 0.215386 | 0.216534 | 0.219298 | 0.221307 | 0.224025 | 0.228603 | 0.228395 | 0.228979 | 0.237156 | 0.246017 | 0.259971 | 0.281364 | 0.306297 | 0.326297 | 0.351484 | 0.360438 | 0.376910 | 0.402164 | 0.428902 | 0.445000 | 0.475918 | 0.545709 | 0.669677 | 0.837162 | 1.088369 | 1.286652 | 1.284759 | 1.143136 | 0.977799 | 0.792917 | 0.694735 | 0.602245 | 0.569390 | 0.556916 | 0.543386 | 0.497574 | 0.463813 | 0.392643 | 0.315936 | 0.263503 | 0.243831 | 0.243948 | 0.262930 | 0.302059 | 0.369360 | 0.481966 | 0.636773 | 0.840214 | 0.995201 | 0.986671 | 0.934623 | 0.734604 | 0.561371 | 0.539547 | 0.643308 | 0.795427 | 1.420914 | 3.226699 | 5.843900 | 8.635180 | 8.636983 | 6.337921 | 3.833499 | 2.006072 | 1.000742 | 0.600191 | 0.434656 | 0.383935 | 0.386159 | 0.434093 | 0.500506 | 0.579158 | 0.653974 | 0.767040 | 0.747742 | 0.704783 | 0.589129 | 0.498379 | 0.402601 | 0.352299 | 0.322510 | 0.332066 | 0.368370 | 0.454373 | 0.627419 | 0.928885 | 1.263621 | 1.481883 | 1.538904 | 1.451584 | 1.286163 | 1.100224 | 0.912491 | 0.666025 | 0.445663 | 0.290018 | 0.223487 | 0.219595 | 0.252953 | 0.301040 | 0.338575 | 0.343335 | 0.329384 | 0.297101 | 0.252395 | 0.239401 | 0.228190 |
| left central | 0.310965 | 0.346045 | 0.365953 | 0.416735 | 0.457829 | 0.439835 | 0.329726 | 0.264669 | 0.228733 | 0.214718 | 0.220801 | 0.224435 | 0.226907 | 0.234060 | 0.269569 | 0.332955 | 0.512332 | 0.811387 | 0.817496 | 0.579891 | 0.463882 | 0.359406 | 0.304335 | 0.256621 | 0.218804 | 0.221219 | 0.285501 | 0.447948 | 0.786771 | 1.207612 | 1.442163 | 1.297728 | 0.825584 | 0.386423 | 0.244371 | 0.216731 | 0.214835 | 0.214636 | 0.215340 | 0.218651 | 0.228776 | 0.234029 | 0.227286 | 0.230324 | 0.231636 | 0.231925 | 0.232081 | 0.232189 | 0.230935 | 0.227196 | 0.217964 | 0.214969 | 0.228264 | 0.247618 | 0.275704 | 0.303846 | 0.340134 | 0.384197 | 0.549511 | 0.704336 | 0.848240 | 0.677473 | 0.470417 | 0.301285 | 0.222189 | 0.226694 | 0.280137 | 0.373244 | 0.453960 | 0.397100 | 0.308485 | 0.245462 | 0.218043 | 0.214672 | 0.216191 | 0.222313 | 0.224091 | 0.221388 | 0.220926 | 0.220289 | 0.220199 | 0.222647 | 0.227908 | 0.239607 | 0.260002 | 0.284227 | 0.293903 | 0.285403 | 0.250932 | 0.220834 | 0.225824 | 0.359479 | 0.871974 | 2.624219 | 7.784987 | 17.855213 | 27.025006 | 31.207717 | 28.459036 | 17.085575 | 8.021167 | 2.774115 | 1.168978 | 0.693411 | 0.469049 | 0.335478 | 0.277774 | 0.243889 | 0.226119 | 0.219275 | 0.216921 | 0.215949 | 0.215561 | 0.214901 | 0.215834 | 0.219373 | 0.221098 | 0.219151 | 0.218846 | 0.216089 | 0.214647 | 0.216004 | 0.217021 | 0.217902 | 0.215554 | 0.214860 | 0.217797 | 0.227514 | 0.243632 | 0.259678 | 0.268238 | 0.274390 | 0.262651 | 0.251945 | 0.239459 | 0.231449 | 0.226388 | 0.225750 | 0.226662 | 0.229582 | 0.233094 | 0.232652 | 0.227580 | 0.219600 | 0.214655 | 0.220495 | 0.235041 | 0.259482 | 0.290951 | 0.343454 | 0.452139 | 0.639885 | 0.638939 | 0.596112 | 0.599346 | 0.509163 | 0.386640 | 0.302008 | 0.246503 | 0.216296 | 0.219181 | 0.235556 | 0.241520 | 0.236452 | 0.225188 | 0.220692 | 0.217486 | 0.215400 | 0.215386 | 0.215437 | 0.214819 | 0.219780 | 0.234008 | 0.260400 | 0.294712 | 0.329231 | 0.360698 | 0.372437 | 0.337977 | 0.297011 | 0.257721 | 0.230632 | 0.215327 | 0.216207 | 0.225580 | 0.231085 | 0.229125 | 0.221577 | 0.216976 | 0.214661 | 0.214702 | 0.214761 | 0.215024 | 0.215175 | 0.214871 | 0.214668 | 0.214635 | 0.215449 | 0.216907 | 0.220088 | 0.223970 | 0.228493 | 0.230585 | 0.229919 | 0.226579 | 0.222793 | 0.217851 | 0.215582 | 0.214915 | 0.214750 | 0.214744 | 0.215311 | 0.215862 | 0.216030 | 0.215887 | 0.215074 | 0.214641 | 0.216226 | 0.222013 | 0.231547 | 0.233208 | 0.228961 | 0.220562 | 0.215941 | 0.214658 | 0.214626 | 0.214642 | 0.214775 | 0.214639 | 0.214630 | 0.215043 | 0.215673 | 0.216435 | 0.216066 | 0.215008 | 0.215013 | 0.218640 | 0.222751 | 0.219779 | 0.215252 | 0.215883 | 0.223474 | 0.232712 | 0.239842 | 0.238074 | 0.229219 | 0.220758 | 0.215083 | 0.217452 | 0.231558 | 0.259341 | 0.286564 | 0.310028 | 0.328357 | 0.340314 | 0.330844 | 0.325798 | 0.326462 | 0.343606 | 0.357050 | 0.377693 | 0.412197 | 0.456968 | 0.532047 | 0.646648 | 0.809349 | 1.184859 | 1.596971 | 1.815764 | 1.914355 | 1.628248 | 1.220902 | 0.985793 | 0.816481 | 0.666006 | 0.570920 | 0.467798 | 0.406791 | 0.376584 | 0.346989 |
| right central | 0.219309 | 0.221125 | 0.228175 | 0.252154 | 0.271669 | 0.309212 | 0.307625 | 0.296722 | 0.258298 | 0.224196 | 0.216239 | 0.236621 | 0.284929 | 0.310186 | 0.285311 | 0.244816 | 0.219508 | 0.215613 | 0.215505 | 0.214658 | 0.217653 | 0.231973 | 0.259718 | 0.255062 | 0.242683 | 0.223799 | 0.215006 | 0.214817 | 0.214683 | 0.215054 | 0.219799 | 0.227502 | 0.222761 | 0.214784 | 0.239822 | 0.339267 | 0.524208 | 0.686013 | 0.680622 | 0.503435 | 0.326719 | 0.234320 | 0.214692 | 0.232144 | 0.270618 | 0.337296 | 0.429427 | 0.473472 | 0.432689 | 0.371436 | 0.290792 | 0.237110 | 0.215355 | 0.226359 | 0.286796 | 0.395856 | 0.566691 | 0.748051 | 0.964618 | 1.180951 | 1.490346 | 1.746266 | 1.659347 | 0.930598 | 0.468178 | 0.254094 | 0.215599 | 0.282066 | 0.528360 | 1.091556 | 1.474990 | 1.343002 | 0.842431 | 0.462614 | 0.284300 | 0.218913 | 0.231309 | 0.322758 | 0.548303 | 0.867301 | 0.813931 | 0.492666 | 0.292103 | 0.221691 | 0.219083 | 0.254134 | 0.308900 | 0.341360 | 0.420393 | 0.577511 | 0.956045 | 1.593212 | 2.738291 | 3.921909 | 5.379733 | 5.363251 | 5.515176 | 5.539067 | 4.952336 | 4.103091 | 3.701921 | 3.457367 | 2.824140 | 2.008734 | 1.448588 | 1.022361 | 0.679173 | 0.580387 | 0.602035 | 0.757793 | 0.933817 | 1.149964 | 1.401537 | 1.700468 | 1.708938 | 1.466370 | 0.920472 | 0.566358 | 0.378632 | 0.295487 | 0.265998 | 0.286475 | 0.318981 | 0.370172 | 0.412392 | 0.478186 | 0.525335 | 0.563638 | 0.575039 | 0.607990 | 0.610145 | 0.616412 | 0.576093 | 0.550228 | 0.523188 | 0.498231 | 0.460480 | 0.408362 | 0.331837 | 0.279934 | 0.241528 | 0.222053 | 0.215586 | 0.215813 | 0.219714 | 0.226755 | 0.241234 | 0.265266 | 0.269331 | 0.244721 | 0.221492 | 0.215970 | 0.244530 | 0.285053 | 0.311351 | 0.297602 | 0.258128 | 0.231238 | 0.216963 | 0.215344 | 0.220169 | 0.219431 | 0.216129 | 0.215089 | 0.219204 | 0.227116 | 0.228242 | 0.230776 | 0.234468 | 0.247443 | 0.258090 | 0.279250 | 0.296921 | 0.340546 | 0.379258 | 0.457023 | 0.498311 | 0.506666 | 0.461665 | 0.419430 | 0.362091 | 0.333993 | 0.301058 | 0.286683 | 0.284990 | 0.303071 | 0.339433 | 0.413226 | 0.499634 | 0.566912 | 0.530943 | 0.419180 | 0.351780 | 0.312088 | 0.294881 | 0.293933 | 0.298880 | 0.306564 | 0.309782 | 0.297748 | 0.275100 | 0.262121 | 0.244864 | 0.231129 | 0.221821 | 0.218267 | 0.217206 | 0.220907 | 0.229914 | 0.245281 | 0.268389 | 0.302590 | 0.302442 | 0.288676 | 0.281078 | 0.267999 | 0.263062 | 0.290689 | 0.343226 | 0.403358 | 0.454355 | 0.430202 | 0.359602 | 0.285922 | 0.232211 | 0.214689 | 0.228250 | 0.263282 | 0.291195 | 0.273819 | 0.239920 | 0.215124 | 0.227715 | 0.304825 | 0.480438 | 0.833552 | 1.384000 | 2.071917 | 1.993043 | 1.877894 | 1.324596 | 0.842181 | 0.548777 | 0.405731 | 0.330712 | 0.325080 | 0.360520 | 0.455673 | 0.640539 | 0.844860 | 0.996847 | 1.010074 | 0.791076 | 0.636956 | 0.533861 | 0.464255 | 0.419180 | 0.364970 | 0.318609 | 0.294004 | 0.262680 | 0.239890 | 0.227518 | 0.219384 | 0.216373 | 0.216220 | 0.216848 | 0.219120 | 0.224800 | 0.231962 | 0.243610 | 0.261885 | 0.332450 | 0.492774 | 0.690747 | 0.939983 | 1.114737 | 0.960896 | 0.873304 | 0.817333 |
| left posterior | 0.396891 | 0.488329 | 0.665594 | 1.159127 | 2.656899 | 3.837708 | 2.605781 | 0.906241 | 0.310848 | 0.214627 | 0.268982 | 0.421174 | 0.585591 | 0.598434 | 0.479947 | 0.296191 | 0.216275 | 0.257910 | 0.415844 | 0.573088 | 0.403220 | 0.237611 | 0.225468 | 0.334958 | 0.607932 | 1.140486 | 1.458408 | 1.039113 | 0.630052 | 0.352677 | 0.226696 | 0.227393 | 0.264877 | 0.267238 | 0.244425 | 0.228030 | 0.218693 | 0.215438 | 0.215049 | 0.223416 | 0.246106 | 0.297620 | 0.413837 | 0.639211 | 1.042250 | 1.517372 | 1.737167 | 1.893269 | 1.989416 | 2.316089 | 2.979438 | 2.828377 | 1.668469 | 0.728140 | 0.305864 | 0.215293 | 0.259337 | 0.428531 | 0.678331 | 0.627790 | 0.373499 | 0.216966 | 0.465730 | 4.187208 | 33.999293 | 163.563973 | 555.271479 | 1233.617651 | 1511.716678 | 842.523590 | 300.937375 | 71.288379 | 17.198540 | 5.152145 | 2.024880 | 1.031326 | 0.620762 | 0.425795 | 0.332973 | 0.290307 | 0.266098 | 0.258627 | 0.260068 | 0.272213 | 0.299726 | 0.337496 | 0.394183 | 0.477491 | 0.636035 | 1.314771 | 4.672961 | 20.543912 | 64.128873 | 119.224111 | 150.143828 | 162.574530 | 136.254431 | 86.838658 | 44.306289 | 22.602577 | 12.404850 | 7.954645 | 6.210623 | 6.553747 | 8.076287 | 10.296366 | 12.283960 | 14.005756 | 14.472435 | 14.289205 | 13.416501 | 11.921883 | 8.579983 | 5.704493 | 3.368097 | 2.118566 | 1.555576 | 1.351239 | 1.284737 | 1.407348 | 1.719796 | 1.926312 | 1.933782 | 1.712560 | 1.256536 | 0.753193 | 0.427259 | 0.269314 | 0.218236 | 0.217263 | 0.228630 | 0.232990 | 0.230286 | 0.222759 | 0.219508 | 0.215821 | 0.214675 | 0.219560 | 0.230910 | 0.254097 | 0.283692 | 0.326044 | 0.398143 | 0.476239 | 0.543929 | 0.584889 | 0.604678 | 0.547886 | 0.497179 | 0.443322 | 0.410063 | 0.375741 | 0.348138 | 0.328587 | 0.306271 | 0.281583 | 0.255848 | 0.240955 | 0.229272 | 0.228409 | 0.233546 | 0.251598 | 0.276795 | 0.302566 | 0.315667 | 0.317732 | 0.303960 | 0.277200 | 0.248191 | 0.224826 | 0.214671 | 0.218417 | 0.221948 | 0.220953 | 0.218136 | 0.214822 | 0.215932 | 0.222589 | 0.237927 | 0.262891 | 0.288229 | 0.301670 | 0.307720 | 0.305167 | 0.299838 | 0.293994 | 0.278258 | 0.254284 | 0.230316 | 0.217034 | 0.214861 | 0.218634 | 0.224364 | 0.222397 | 0.215985 | 0.215572 | 0.227460 | 0.255083 | 0.301629 | 0.367359 | 0.444359 | 0.531620 | 0.610085 | 0.648623 | 0.628035 | 0.592616 | 0.529764 | 0.471140 | 0.429060 | 0.435433 | 0.478702 | 0.582917 | 0.770951 | 1.121474 | 1.506242 | 1.761828 | 1.677158 | 1.459246 | 1.131594 | 0.889636 | 0.683736 | 0.604736 | 0.561691 | 0.523966 | 0.471920 | 0.418767 | 0.347219 | 0.294725 | 0.253561 | 0.235546 | 0.229139 | 0.231104 | 0.245514 | 0.286133 | 0.361979 | 0.499636 | 0.658363 | 0.774628 | 0.744594 | 0.616177 | 0.501071 | 0.423636 | 0.355082 | 0.322549 | 0.296569 | 0.287585 | 0.299273 | 0.322894 | 0.345196 | 0.368077 | 0.362569 | 0.336834 | 0.316016 | 0.289569 | 0.267092 | 0.252832 | 0.242647 | 0.234709 | 0.232741 | 0.228979 | 0.226323 | 0.222437 | 0.217631 | 0.215015 | 0.214830 | 0.219602 | 0.231442 | 0.249173 | 0.273503 | 0.304415 | 0.355701 | 0.472401 | 0.674368 | 0.980921 | 1.346367 | 1.582487 | 1.448840 | 1.236552 | 1.179263 | 1.126711 |
| right posterior | 0.254580 | 0.257162 | 0.272528 | 0.280896 | 0.307530 | 0.375088 | 0.414521 | 0.372944 | 0.276684 | 0.217346 | 0.224894 | 0.262335 | 0.320353 | 0.362753 | 0.344660 | 0.315618 | 0.262494 | 0.225035 | 0.214722 | 0.216417 | 0.219170 | 0.215881 | 0.216217 | 0.217673 | 0.216789 | 0.218970 | 0.224403 | 0.225547 | 0.229293 | 0.238734 | 0.252767 | 0.265257 | 0.246119 | 0.219579 | 0.222141 | 0.309518 | 0.599519 | 1.162153 | 2.329895 | 4.444456 | 8.188823 | 15.067827 | 26.747278 | 42.887469 | 72.657448 | 95.553023 | 51.001186 | 11.731200 | 1.657994 | 0.436710 | 0.242333 | 0.215372 | 0.217669 | 0.220823 | 0.222165 | 0.219915 | 0.220926 | 0.221591 | 0.221301 | 0.215160 | 0.234973 | 0.518916 | 4.281634 | 88.580538 | 1474.358591 | 8414.080510 | 13793.653991 | 7144.259693 | 1689.714723 | 267.330581 | 42.795510 | 8.516717 | 2.249405 | 0.702575 | 0.313074 | 0.217444 | 0.253211 | 0.411124 | 0.671023 | 0.997231 | 1.013645 | 0.799476 | 0.537142 | 0.349727 | 0.256209 | 0.220476 | 0.216661 | 0.243449 | 0.308113 | 0.442812 | 0.656913 | 1.155709 | 2.030198 | 3.822758 | 7.347988 | 13.684986 | 20.350304 | 25.653292 | 26.352457 | 23.551423 | 22.877815 | 26.182042 | 37.486497 | 68.536722 | 143.343463 | 215.956565 | 173.181245 | 81.014757 | 31.312357 | 13.685999 | 8.047494 | 5.278139 | 3.695636 | 3.101366 | 2.792648 | 2.968602 | 3.681309 | 4.876153 | 7.046061 | 11.370219 | 16.777410 | 20.842985 | 21.670454 | 18.742331 | 14.134869 | 9.727453 | 5.933205 | 3.590174 | 2.194094 | 1.621172 | 1.247874 | 1.116492 | 1.077476 | 1.056606 | 1.076769 | 1.289565 | 1.560685 | 1.763322 | 1.869811 | 1.769236 | 1.674913 | 1.533033 | 1.466872 | 1.703706 | 2.691599 | 4.376846 | 6.304257 | 6.943835 | 5.359490 | 3.722937 | 2.525154 | 1.727148 | 1.209307 | 0.947196 | 0.842032 | 0.881997 | 1.012142 | 1.315557 | 1.809001 | 2.492535 | 2.989759 | 3.669857 | 3.973789 | 3.822511 | 3.127671 | 2.569344 | 2.058286 | 1.858698 | 1.612992 | 1.455037 | 1.382102 | 1.493977 | 1.856091 | 2.930901 | 4.646285 | 6.803581 | 7.226207 | 5.756073 | 3.992290 | 2.614476 | 1.651850 | 1.065121 | 0.740051 | 0.568636 | 0.499831 | 0.520107 | 0.598392 | 0.705214 | 0.855788 | 1.017712 | 1.083373 | 1.050861 | 0.970074 | 0.907785 | 0.892323 | 0.954721 | 1.010137 | 1.020887 | 1.032991 | 1.120693 | 1.298956 | 1.647209 | 2.261521 | 3.248095 | 4.770169 | 6.993729 | 9.134018 | 10.681116 | 10.186633 | 7.924596 | 6.199196 | 5.235791 | 5.004875 | 6.100527 | 7.366919 | 8.094806 | 8.289811 | 7.392585 | 5.436552 | 3.834580 | 2.522735 | 1.774903 | 1.434959 | 1.321697 | 1.377885 | 1.653491 | 1.930568 | 2.211935 | 2.365038 | 2.359267 | 2.144785 | 1.893130 | 1.647526 | 1.424045 | 1.255987 | 1.145726 | 1.061358 | 1.059025 | 1.126552 | 1.269731 | 1.429528 | 1.630553 | 1.842366 | 2.178816 | 2.698950 | 3.757981 | 6.618271 | 12.031545 | 16.655515 | 14.400515 | 8.882589 | 5.404172 | 3.269084 | 2.282449 | 1.975747 | 2.208499 | 2.693346 | 3.601550 | 4.747290 | 6.077302 | 5.982386 | 4.905213 | 3.687393 | 2.575637 | 1.782261 | 1.187180 | 0.817204 | 0.627763 | 0.601303 | 0.607161 | 0.699295 | 0.912358 | 1.184723 | 1.351127 | 1.630388 | 1.618663 | 1.460811 | 1.398235 | 1.333688 | 1.268421 |
| all electrodes | 0.499700 | 0.480889 | 0.421614 | 0.278179 | 0.220365 | 0.216649 | 0.263357 | 0.379428 | 0.505155 | 0.649688 | 0.752906 | 0.642404 | 0.643929 | 0.537582 | 0.358125 | 0.259540 | 0.226310 | 0.215100 | 0.215132 | 0.220004 | 0.225343 | 0.225232 | 0.216023 | 0.214823 | 0.235066 | 0.318158 | 0.508394 | 0.650154 | 0.802457 | 0.702181 | 0.542921 | 0.379111 | 0.275157 | 0.237104 | 0.239902 | 0.241037 | 0.245329 | 0.254069 | 0.264344 | 0.252168 | 0.232866 | 0.226948 | 0.217233 | 0.214659 | 0.214648 | 0.214787 | 0.216336 | 0.215559 | 0.222386 | 0.230752 | 0.238289 | 0.245354 | 0.259864 | 0.274470 | 0.294124 | 0.313185 | 0.299522 | 0.282465 | 0.257866 | 0.231845 | 0.215598 | 0.270891 | 0.601783 | 2.337412 | 12.848576 | 65.842796 | 238.071190 | 212.003322 | 80.556346 | 26.153281 | 15.891926 | 13.964002 | 13.508111 | 9.000285 | 4.513657 | 1.526098 | 0.610027 | 0.331029 | 0.247493 | 0.227463 | 0.232687 | 0.254453 | 0.304319 | 0.344738 | 0.333094 | 0.315794 | 0.297297 | 0.271398 | 0.271618 | 0.304289 | 0.383083 | 0.560528 | 0.915439 | 1.492164 | 2.733712 | 4.325399 | 6.649646 | 8.903152 | 8.254858 | 4.759304 | 2.328060 | 0.834140 | 0.392205 | 0.285908 | 0.254079 | 0.260052 | 0.332171 | 0.478952 | 0.814329 | 1.415931 | 2.342345 | 2.709626 | 2.806282 | 2.018374 | 1.606174 | 1.214553 | 1.032758 | 0.830894 | 0.817613 | 0.890091 | 1.166987 | 1.639109 | 2.829214 | 3.586140 | 2.935581 | 1.766987 | 1.083670 | 0.649848 | 0.448787 | 0.345594 | 0.302064 | 0.294982 | 0.329474 | 0.386415 | 0.470638 | 0.590492 | 0.749820 | 0.940771 | 1.407609 | 2.064902 | 2.911412 | 3.259196 | 2.718388 | 2.514046 | 2.803916 | 2.948032 | 2.810003 | 2.669318 | 2.661984 | 2.724424 | 2.759516 | 2.621624 | 2.561984 | 1.925238 | 1.202341 | 0.738347 | 0.537898 | 0.429560 | 0.423388 | 0.407519 | 0.379038 | 0.324939 | 0.276237 | 0.238234 | 0.221407 | 0.214676 | 0.216615 | 0.219182 | 0.219373 | 0.218730 | 0.218894 | 0.220596 | 0.222837 | 0.228463 | 0.235262 | 0.230897 | 0.221320 | 0.214831 | 0.220582 | 0.248543 | 0.312569 | 0.440964 | 0.566077 | 0.684927 | 0.737551 | 0.694391 | 0.532328 | 0.374349 | 0.260074 | 0.221806 | 0.214819 | 0.216182 | 0.214866 | 0.216090 | 0.226629 | 0.250841 | 0.269428 | 0.270654 | 0.259568 | 0.249123 | 0.244665 | 0.249972 | 0.281336 | 0.362735 | 0.455713 | 0.573662 | 0.723931 | 0.869815 | 0.953052 | 1.057161 | 0.972683 | 0.814867 | 0.581280 | 0.414769 | 0.317014 | 0.284819 | 0.278783 | 0.305462 | 0.381919 | 0.474677 | 0.479137 | 0.406458 | 0.332816 | 0.278439 | 0.259750 | 0.269533 | 0.337918 | 0.479694 | 0.727855 | 1.055723 | 1.341191 | 1.106977 | 0.932733 | 0.772432 | 0.693172 | 0.657293 | 0.614164 | 0.532323 | 0.448241 | 0.370568 | 0.305953 | 0.269100 | 0.255433 | 0.258371 | 0.267441 | 0.295435 | 0.321562 | 0.330784 | 0.327621 | 0.302527 | 0.256945 | 0.226213 | 0.214779 | 0.220539 | 0.231950 | 0.251095 | 0.258199 | 0.240254 | 0.221197 | 0.214738 | 0.233867 | 0.298188 | 0.393448 | 0.524514 | 0.605282 | 0.585756 | 0.485228 | 0.356683 | 0.265749 | 0.226614 | 0.214848 | 0.220127 | 0.232753 | 0.246613 | 0.253692 | 0.267364 | 0.281929 | 0.299026 | 0.297828 | 0.318290 |

W) neural network features, neutral vs angry

  
|  | time window | peak latency | cluster *p* | peak Cohen's *d* |  | | | |
| **all electrodes** | 120 - 600 ms | 160 ms | 0.0002 | 1.3353 |  | | | |
|  | | | | | | | | |

Model correlations, cluster permutation tests

|  | **left hemisphere** | | | | **right hemisphere** | | | |
|  | time window | peak latency | cluster *p* | peak Cohen's *d* | time window | peak latency | cluster *p* | peak Cohen's *d* |
| **anterior** |  | | | |  | | | |
| **central** |  | | | | 120 - 475 ms | 160 ms | 0.0002 | 1.3133 |
| **posterior** | 120 - 235 ms | 160 ms | 0.0182 | 1.1275 | 95 - 1195 ms | 155 ms | 0.0002 | 1.2874 |
 250 - 540 ms | 415 ms | 0.0044 | 0.8293 |  | | | | 580 - 815 ms | 660 ms | 0.0111 | 0.572 |  | | | |

  

Model correlations, Bayesian statistics

|  | -200 | -195 | -190 | -185 | -180 | -175 | -170 | -165 | -160 | -155 | -150 | -145 | -140 | -135 | -130 | -125 | -120 | -115 | -110 | -105 | -100 | -95 | -90 | -85 | -80 | -75 | -70 | -65 | -60 | -55 | -50 | -45 | -40 | -35 | -30 | -25 | -20 | -15 | -10 | -5 | 0 | 5 | 10 | 15 | 20 | 25 | 30 | 35 | 40 | 45 | 50 | 55 | 60 | 65 | 70 | 75 | 80 | 85 | 90 | 95 | 100 | 105 | 110 | 115 | 120 | 125 | 130 | 135 | 140 | 145 | 150 | 155 | 160 | 165 | 170 | 175 | 180 | 185 | 190 | 195 | 200 | 205 | 210 | 215 | 220 | 225 | 230 | 235 | 240 | 245 | 250 | 255 | 260 | 265 | 270 | 275 | 280 | 285 | 290 | 295 | 300 | 305 | 310 | 315 | 320 | 325 | 330 | 335 | 340 | 345 | 350 | 355 | 360 | 365 | 370 | 375 | 380 | 385 | 390 | 395 | 400 | 405 | 410 | 415 | 420 | 425 | 430 | 435 | 440 | 445 | 450 | 455 | 460 | 465 | 470 | 475 | 480 | 485 | 490 | 495 | 500 | 505 | 510 | 515 | 520 | 525 | 530 | 535 | 540 | 545 | 550 | 555 | 560 | 565 | 570 | 575 | 580 | 585 | 590 | 595 | 600 | 605 | 610 | 615 | 620 | 625 | 630 | 635 | 640 | 645 | 650 | 655 | 660 | 665 | 670 | 675 | 680 | 685 | 690 | 695 | 700 | 705 | 710 | 715 | 720 | 725 | 730 | 735 | 740 | 745 | 750 | 755 | 760 | 765 | 770 | 775 | 780 | 785 | 790 | 795 | 800 | 805 | 810 | 815 | 820 | 825 | 830 | 835 | 840 | 845 | 850 | 855 | 860 | 865 | 870 | 875 | 880 | 885 | 890 | 895 | 900 | 905 | 910 | 915 | 920 | 925 | 930 | 935 | 940 | 945 | 950 | 955 | 960 | 965 | 970 | 975 | 980 | 985 | 990 | 995 | 1000 | 1005 | 1010 | 1015 | 1020 | 1025 | 1030 | 1035 | 1040 | 1045 | 1050 | 1055 | 1060 | 1065 | 1070 | 1075 | 1080 | 1085 | 1090 | 1095 | 1100 | 1105 | 1110 | 1115 | 1120 | 1125 | 1130 | 1135 | 1140 | 1145 | 1150 | 1155 | 1160 | 1165 | 1170 | 1175 | 1180 | 1185 | 1190 | 1195 |
| --- | --- | --- | --- | --- | --- | --- | --- | --- | --- | --- | --- | --- | --- | --- | --- | --- | --- | --- | --- | --- | --- | --- | --- | --- | --- | --- | --- | --- | --- | --- | --- | --- | --- | --- | --- | --- | --- | --- | --- | --- | --- | --- | --- | --- | --- | --- | --- | --- | --- | --- | --- | --- | --- | --- | --- | --- | --- | --- | --- | --- | --- | --- | --- | --- | --- | --- | --- | --- | --- | --- | --- | --- | --- | --- | --- | --- | --- | --- | --- | --- | --- | --- | --- | --- | --- | --- | --- | --- | --- | --- | --- | --- | --- | --- | --- | --- | --- | --- | --- | --- | --- | --- | --- | --- | --- | --- | --- | --- | --- | --- | --- | --- | --- | --- | --- | --- | --- | --- | --- | --- | --- | --- | --- | --- | --- | --- | --- | --- | --- | --- | --- | --- | --- | --- | --- | --- | --- | --- | --- | --- | --- | --- | --- | --- | --- | --- | --- | --- | --- | --- | --- | --- | --- | --- | --- | --- | --- | --- | --- | --- | --- | --- | --- | --- | --- | --- | --- | --- | --- | --- | --- | --- | --- | --- | --- | --- | --- | --- | --- | --- | --- | --- | --- | --- | --- | --- | --- | --- | --- | --- | --- | --- | --- | --- | --- | --- | --- | --- | --- | --- | --- | --- | --- | --- | --- | --- | --- | --- | --- | --- | --- | --- | --- | --- | --- | --- | --- | --- | --- | --- | --- | --- | --- | --- | --- | --- | --- | --- | --- | --- | --- | --- | --- | --- | --- | --- | --- | --- | --- | --- | --- | --- | --- | --- | --- | --- | --- | --- | --- | --- | --- | --- | --- | --- | --- | --- | --- | --- | --- | --- | --- | --- | --- | --- | --- | --- | --- | --- | --- | --- | --- | --- | --- | --- | --- | --- | --- | --- | --- | --- |
| left anterior | 0.316416 | 0.340148 | 0.381344 | 0.502081 | 0.727192 | 1.692073 | 3.775745 | 9.811128 | 33.733148 | 48.794350 | 17.486195 | 4.917883 | 1.532376 | 0.582721 | 0.326582 | 0.277232 | 0.294443 | 0.392203 | 0.741085 | 1.744269 | 3.716651 | 5.619668 | 3.478339 | 1.024832 | 0.383272 | 0.233793 | 0.224558 | 0.326347 | 0.542205 | 0.634129 | 0.576021 | 0.328315 | 0.230836 | 0.218393 | 0.250429 | 0.258820 | 0.243656 | 0.240118 | 0.240153 | 0.236756 | 0.238106 | 0.253116 | 0.302342 | 0.366329 | 0.457327 | 0.816901 | 1.703174 | 2.007458 | 1.685699 | 0.897104 | 0.474447 | 0.321573 | 0.248769 | 0.221898 | 0.215829 | 0.214657 | 0.218025 | 0.227453 | 0.238815 | 0.245492 | 0.258974 | 0.247931 | 0.237225 | 0.240375 | 0.285873 | 0.373985 | 0.607700 | 0.899096 | 1.220535 | 1.207972 | 1.067711 | 0.747783 | 0.515558 | 0.352569 | 0.250802 | 0.215461 | 0.272764 | 0.499139 | 1.034946 | 1.611236 | 1.553392 | 1.132659 | 0.754264 | 0.518190 | 0.341966 | 0.260069 | 0.227370 | 0.214804 | 0.222736 | 0.254958 | 0.324004 | 0.431078 | 0.561784 | 0.717290 | 7.272743e-01 | 6.367139e-01 | 4.959253e-01 | 3.648348e-01 | 2.707980e-01 | 0.228846 | 0.215146 | 0.216382 | 0.220677 | 0.221005 | 0.220636 | 0.218296 | 0.215390 | 0.214996 | 0.218324 | 0.225565 | 0.244882 | 0.272143 | 0.310380 | 0.344937 | 0.395973 | 0.449177 | 0.549056 | 0.798708 | 1.467190 | 2.912901 | 4.802565 | 6.462368 | 5.744466 | 4.943006 | 4.060075 | 3.033588 | 2.099863 | 1.416116 | 0.867267 | 0.534144 | 0.373753 | 0.291297 | 0.246256 | 0.225415 | 0.217506 | 0.214661 | 0.217614 | 0.216947 | 0.214732 | 0.218756 | 0.227934 | 0.235711 | 0.239278 | 0.242096 | 0.230382 | 0.221556 | 0.215526 | 0.218573 | 0.249560 | 0.314689 | 0.389972 | 0.449170 | 0.461887 | 0.460892 | 0.429913 | 0.361406 | 0.289284 | 0.251036 | 0.228578 | 0.216241 | 0.215412 | 0.218442 | 0.223395 | 0.231123 | 0.244119 | 0.271767 | 0.333300 | 0.439766 | 0.607275 | 0.774510 | 0.843789 | 0.707212 | 0.527474 | 0.368027 | 0.279313 | 0.240641 | 0.232848 | 0.238417 | 0.263649 | 0.292562 | 0.320515 | 0.331004 | 0.328682 | 0.330798 | 0.355624 | 0.397925 | 0.442866 | 0.506845 | 0.575053 | 0.590098 | 0.475435 | 0.377533 | 0.313896 | 0.276779 | 0.262275 | 0.267509 | 0.302934 | 0.358151 | 0.398497 | 0.391161 | 0.357596 | 0.302697 | 0.249384 | 0.216771 | 0.221816 | 0.247653 | 0.261514 | 0.239756 | 0.216380 | 0.226112 | 0.284548 | 0.363856 | 0.419150 | 0.450426 | 0.397329 | 0.313345 | 0.261873 | 0.235891 | 0.227375 | 0.226779 | 0.223162 | 0.216208 | 0.215113 | 0.225999 | 0.271467 | 0.382161 | 0.560786 | 0.749588 | 0.766418 | 0.663266 | 0.557876 | 0.544680 | 0.634313 | 0.912296 | 1.300437 | 1.722969 | 2.004245 | 1.574119 | 0.966919 | 0.580564 | 0.369626 | 0.268313 | 0.231723 | 0.218069 | 0.215083 | 0.214627 | 0.214630 | 0.214630 | 0.215328 | 0.217640 | 0.222016 | 0.223196 | 0.221244 | 0.217017 | 0.214821 | 0.214898 | 0.215748 | 0.217332 | 0.216915 | 0.216687 | 0.216823 | 0.216751 | 0.217278 | 0.220084 | 0.223637 | 0.230621 | 0.246799 | 0.261633 | 0.270776 | 0.284445 | 0.290726 | 0.292782 | 0.301818 | 0.317468 | 0.340511 | 0.387677 | 0.457163 | 0.553454 | 0.612523 | 0.710969 |
| right anterior | 0.219497 | 0.218638 | 0.214723 | 0.224402 | 0.252253 | 0.304186 | 0.358640 | 0.382051 | 0.410774 | 0.446653 | 0.549975 | 1.018135 | 1.306521 | 0.947579 | 0.704103 | 0.505673 | 0.375475 | 0.327352 | 0.293713 | 0.271415 | 0.267276 | 0.249528 | 0.246005 | 0.270146 | 0.276935 | 0.278923 | 0.282552 | 0.303860 | 0.323165 | 0.318594 | 0.298833 | 0.357465 | 0.377190 | 0.338867 | 0.289602 | 0.237633 | 0.214888 | 0.239601 | 0.321168 | 0.343699 | 0.298508 | 0.267344 | 0.228044 | 0.215524 | 0.231267 | 0.257602 | 0.268058 | 0.242835 | 0.215312 | 0.223005 | 0.252065 | 0.271799 | 0.293230 | 0.295745 | 0.314000 | 0.315366 | 0.327328 | 0.340131 | 0.349975 | 0.325773 | 0.302209 | 0.257730 | 0.237005 | 0.236291 | 0.261623 | 0.342385 | 0.617467 | 1.377460 | 3.303123 | 6.774023 | 11.794752 | 15.594721 | 13.077831 | 7.708350 | 3.764384 | 1.985601 | 1.149842 | 0.742516 | 0.524042 | 0.427533 | 0.341832 | 0.296942 | 0.270900 | 0.244053 | 0.227383 | 0.221939 | 0.221902 | 0.226402 | 0.248608 | 0.328530 | 0.563923 | 1.162852 | 2.520675 | 5.655820 | 1.088409e+01 | 1.585441e+01 | 1.692348e+01 | 1.500148e+01 | 1.096185e+01 | 7.158121 | 3.938388 | 2.061736 | 1.056058 | 0.687521 | 0.529637 | 0.481195 | 0.434481 | 0.395352 | 0.353369 | 0.313997 | 0.282345 | 0.271700 | 0.266581 | 0.284798 | 0.322905 | 0.366379 | 0.459778 | 0.635730 | 0.890943 | 1.408071 | 2.421974 | 4.230990 | 6.928167 | 9.120239 | 10.016018 | 10.416219 | 9.054078 | 7.588941 | 6.142472 | 4.815762 | 3.565153 | 2.615682 | 1.756235 | 1.264047 | 0.807071 | 0.536635 | 0.383792 | 0.291478 | 0.237809 | 0.217683 | 0.214926 | 0.218098 | 0.218691 | 0.219619 | 0.218505 | 0.217997 | 0.219773 | 0.220945 | 0.218624 | 0.214790 | 0.219005 | 0.240978 | 0.293320 | 0.400354 | 0.505816 | 0.490425 | 0.398958 | 0.290689 | 0.223253 | 0.222827 | 0.285389 | 0.397223 | 0.560961 | 0.688078 | 0.751266 | 0.777962 | 0.873062 | 0.895925 | 0.894171 | 0.867300 | 0.692203 | 0.482170 | 0.363438 | 0.306914 | 0.281095 | 0.281295 | 0.282133 | 0.288371 | 0.301738 | 0.304835 | 0.297373 | 0.308221 | 0.322676 | 0.329792 | 0.336830 | 0.350236 | 0.369356 | 0.380363 | 0.375962 | 0.355813 | 0.342018 | 0.309565 | 0.284959 | 0.269360 | 0.259232 | 0.251191 | 0.244670 | 0.239955 | 0.248225 | 0.273572 | 0.302136 | 0.334975 | 0.352859 | 0.350621 | 0.345949 | 0.345193 | 0.350240 | 0.360031 | 0.340554 | 0.304949 | 0.272140 | 0.250060 | 0.235866 | 0.231189 | 0.230679 | 0.235916 | 0.246949 | 0.267396 | 0.294817 | 0.340435 | 0.376923 | 0.404339 | 0.414802 | 0.446359 | 0.498152 | 0.608849 | 0.746777 | 0.846815 | 0.848254 | 0.750252 | 0.600926 | 0.502019 | 0.454997 | 0.418106 | 0.420337 | 0.437199 | 0.474246 | 0.514569 | 0.550794 | 0.531326 | 0.486707 | 0.398699 | 0.316620 | 0.267144 | 0.243366 | 0.232900 | 0.228992 | 0.235247 | 0.249004 | 0.266972 | 0.286574 | 0.320400 | 0.350054 | 0.383548 | 0.392524 | 0.380537 | 0.349860 | 0.313172 | 0.267377 | 0.237785 | 0.221810 | 0.215368 | 0.215129 | 0.216076 | 0.215357 | 0.214721 | 0.220865 | 0.242975 | 0.282606 | 0.358030 | 0.438749 | 0.487582 | 0.526813 | 0.530763 | 0.474178 | 0.431906 | 0.378487 | 0.345415 | 0.325639 | 0.308484 |
| left central | 0.218413 | 0.230737 | 0.239803 | 0.252839 | 0.260329 | 0.238589 | 0.218972 | 0.215230 | 0.225335 | 0.244115 | 0.265884 | 0.266473 | 0.241812 | 0.221561 | 0.215042 | 0.218977 | 0.225272 | 0.236422 | 0.248707 | 0.256159 | 0.254191 | 0.224815 | 0.214645 | 0.228365 | 0.304455 | 0.461347 | 0.619006 | 0.718927 | 0.564473 | 0.366090 | 0.272344 | 0.236644 | 0.243981 | 0.316528 | 0.594541 | 1.351802 | 3.520133 | 7.801098 | 22.038518 | 42.400526 | 50.000913 | 21.964571 | 8.231269 | 3.086595 | 1.357669 | 0.681171 | 0.453884 | 0.354005 | 0.288164 | 0.261116 | 0.260540 | 0.260151 | 0.265294 | 0.280765 | 0.292339 | 0.316915 | 0.376181 | 0.498668 | 0.708102 | 0.994989 | 1.031611 | 0.672422 | 0.338501 | 0.268627 | 0.285797 | 0.463280 | 1.285076 | 5.780345 | 23.080636 | 71.369684 | 164.946022 | 301.271413 | 300.169382 | 201.537623 | 97.611811 | 34.776855 | 9.313021 | 2.462528 | 0.785524 | 0.407196 | 0.280185 | 0.231694 | 0.221903 | 0.225470 | 0.226955 | 0.241479 | 0.270633 | 0.307512 | 0.368494 | 0.508964 | 0.873294 | 2.265467 | 8.380350 | 33.554384 | 1.011175e+02 | 1.936290e+02 | 2.268187e+02 | 1.374855e+02 | 5.592011e+01 | 19.077074 | 7.637242 | 3.699200 | 1.794085 | 0.912081 | 0.563791 | 0.426931 | 0.353660 | 0.320617 | 0.317596 | 0.320467 | 0.323572 | 0.321895 | 0.331102 | 0.344425 | 0.388263 | 0.445266 | 0.513387 | 0.593908 | 0.752894 | 0.941548 | 1.100284 | 1.117472 | 1.109727 | 1.023499 | 0.886168 | 0.739084 | 0.676997 | 0.665399 | 0.606827 | 0.493075 | 0.381855 | 0.292694 | 0.232444 | 0.215334 | 0.214891 | 0.215160 | 0.226704 | 0.256606 | 0.321833 | 0.434844 | 0.498326 | 0.416818 | 0.303570 | 0.225228 | 0.219965 | 0.259268 | 0.287115 | 0.271662 | 0.239674 | 0.216538 | 0.219604 | 0.251020 | 0.288125 | 0.312741 | 0.310443 | 0.295016 | 0.266351 | 0.249179 | 0.232601 | 0.221408 | 0.217516 | 0.215234 | 0.214681 | 0.215900 | 0.221144 | 0.228170 | 0.233131 | 0.240970 | 0.249166 | 0.250048 | 0.244134 | 0.238125 | 0.228185 | 0.217542 | 0.215167 | 0.220832 | 0.230661 | 0.238200 | 0.243697 | 0.245016 | 0.238891 | 0.232212 | 0.230814 | 0.227511 | 0.223358 | 0.220154 | 0.218650 | 0.217141 | 0.215791 | 0.214940 | 0.215270 | 0.216311 | 0.219600 | 0.230130 | 0.255774 | 0.303609 | 0.359273 | 0.383979 | 0.374913 | 0.339029 | 0.277437 | 0.229936 | 0.214691 | 0.231112 | 0.281797 | 0.343891 | 0.362370 | 0.324409 | 0.271581 | 0.235866 | 0.222463 | 0.220871 | 0.224154 | 0.230226 | 0.242071 | 0.249149 | 0.246593 | 0.240535 | 0.240711 | 0.243052 | 0.252916 | 0.261081 | 0.274363 | 0.284371 | 0.255525 | 0.229196 | 0.219693 | 0.215070 | 0.215636 | 0.221403 | 0.243182 | 0.277484 | 0.325792 | 0.389131 | 0.438367 | 0.417522 | 0.349640 | 0.272847 | 0.233092 | 0.214978 | 0.222358 | 0.243181 | 0.276441 | 0.328346 | 0.421037 | 0.592253 | 0.785907 | 0.991325 | 1.297459 | 1.710238 | 2.021587 | 2.090273 | 1.831671 | 1.318856 | 0.703915 | 0.387886 | 0.274721 | 0.234073 | 0.222628 | 0.221859 | 0.231991 | 0.259664 | 0.310100 | 0.369502 | 0.405578 | 0.421458 | 0.421925 | 0.376692 | 0.331680 | 0.284183 | 0.242795 | 0.220391 | 0.214713 | 0.215534 | 0.216231 | 0.215226 | 0.214796 | 0.219272 | 0.222834 | 0.227845 |
| right central | 0.503772 | 0.546585 | 0.611434 | 0.642017 | 0.667541 | 0.846151 | 1.339268 | 2.000414 | 3.490026 | 4.990478 | 4.157446 | 1.820909 | 0.720380 | 0.292709 | 0.221094 | 0.217660 | 0.246163 | 0.285854 | 0.310547 | 0.330545 | 0.313688 | 0.279047 | 0.242840 | 0.214753 | 0.244464 | 0.425460 | 1.089572 | 3.420976 | 7.204142 | 9.445952 | 3.584775 | 1.034666 | 0.423494 | 0.275070 | 0.230339 | 0.214637 | 0.231836 | 0.284488 | 0.404548 | 0.582506 | 0.656010 | 0.735667 | 0.766820 | 0.848335 | 0.835643 | 0.745392 | 0.525730 | 0.365802 | 0.263076 | 0.226007 | 0.215238 | 0.215136 | 0.220998 | 0.230660 | 0.238212 | 0.238298 | 0.223558 | 0.214779 | 0.230877 | 0.278308 | 0.358009 | 0.436209 | 0.536034 | 0.727942 | 1.181766 | 2.785169 | 9.931119 | 60.486748 | 466.710000 | 3019.710704 | 9587.601124 | 19169.082176 | 17540.145035 | 8201.697242 | 2460.936690 | 580.779191 | 133.943550 | 38.560351 | 11.575625 | 4.483935 | 2.612119 | 1.893336 | 1.727417 | 1.778805 | 2.068297 | 2.634095 | 3.952125 | 6.347343 | 13.047760 | 43.915702 | 188.310944 | 1330.605665 | 14877.658708 | 158603.666922 | 1.019904e+06 | 3.250743e+06 | 4.789854e+06 | 3.796097e+06 | 1.843117e+06 | 465192.354687 | 75478.507183 | 14096.705743 | 2773.279839 | 690.902672 | 235.278866 | 104.507065 | 50.179406 | 26.796606 | 17.391930 | 15.205259 | 14.914419 | 19.559244 | 39.300785 | 101.676182 | 251.414143 | 622.734418 | 1193.232830 | 1925.870001 | 2572.068740 | 3069.643788 | 3354.531474 | 3742.727446 | 2635.652852 | 1299.420012 | 636.548551 | 284.015999 | 116.423875 | 55.472800 | 31.377425 | 17.161319 | 10.183143 | 5.773316 | 3.076838 | 1.866802 | 1.201977 | 0.981437 | 0.973927 | 1.160349 | 1.312482 | 1.421950 | 1.322498 | 1.181182 | 1.112272 | 1.086372 | 1.009185 | 0.889288 | 0.633587 | 0.464971 | 0.370845 | 0.333240 | 0.321542 | 0.349698 | 0.414613 | 0.582777 | 1.040641 | 2.493469 | 6.619850 | 15.658629 | 23.632160 | 23.739763 | 16.654355 | 9.598316 | 3.811413 | 1.555573 | 0.823123 | 0.541222 | 0.430801 | 0.421095 | 0.491233 | 0.634895 | 0.887588 | 1.224131 | 1.665853 | 1.970811 | 2.097680 | 2.026013 | 2.154817 | 2.062407 | 1.932166 | 1.764091 | 1.594750 | 1.350565 | 1.109225 | 0.892196 | 0.728281 | 0.574266 | 0.456439 | 0.395523 | 0.361779 | 0.328200 | 0.329017 | 0.334288 | 0.341154 | 0.358632 | 0.405153 | 0.416219 | 0.433848 | 0.478274 | 0.585891 | 0.617819 | 0.585621 | 0.533674 | 0.481794 | 0.397720 | 0.332965 | 0.287266 | 0.263125 | 0.248237 | 0.241479 | 0.248256 | 0.284932 | 0.382013 | 0.636873 | 1.279234 | 2.412212 | 4.406905 | 5.546759 | 5.083441 | 3.765429 | 2.447838 | 1.357862 | 0.961462 | 0.807658 | 0.880056 | 1.166566 | 1.863114 | 3.011850 | 4.331529 | 4.204262 | 2.707045 | 1.455017 | 0.760261 | 0.464321 | 0.326899 | 0.288895 | 0.289911 | 0.312125 | 0.367432 | 0.440825 | 0.439249 | 0.359792 | 0.263880 | 0.216596 | 0.225920 | 0.263039 | 0.294488 | 0.275301 | 0.240611 | 0.216929 | 0.215925 | 0.219690 | 0.221708 | 0.227620 | 0.234881 | 0.256330 | 0.317758 | 0.418815 | 0.658501 | 1.088636 | 1.609024 | 1.944184 | 2.100598 | 1.668259 | 1.293261 | 0.889194 | 0.539078 | 0.370856 | 0.302013 | 0.266424 | 0.256628 | 0.269499 | 0.282179 | 0.305720 | 0.324409 | 0.353079 | 0.382627 | 0.444934 | 0.520101 | 0.579184 | 0.637614 |
| left posterior | 0.218977 | 0.214763 | 0.216078 | 0.231250 | 0.274041 | 0.355930 | 0.488416 | 0.589692 | 0.492628 | 0.322620 | 0.240460 | 0.217050 | 0.214681 | 0.215633 | 0.225159 | 0.291529 | 0.483744 | 0.847914 | 2.106098 | 4.683776 | 3.755260 | 1.057384 | 0.320915 | 0.215842 | 0.324938 | 0.687742 | 1.133377 | 1.063134 | 0.758874 | 0.517428 | 0.372075 | 0.335113 | 0.392830 | 0.619221 | 1.150683 | 1.877875 | 2.738756 | 3.222219 | 2.982444 | 2.499996 | 2.079558 | 1.712433 | 1.646705 | 1.706416 | 1.390654 | 0.949235 | 0.636435 | 0.418709 | 0.294607 | 0.236136 | 0.215294 | 0.221633 | 0.245130 | 0.277248 | 0.287041 | 0.281585 | 0.281462 | 0.283491 | 0.297269 | 0.328598 | 0.300398 | 0.235166 | 0.218997 | 0.372744 | 1.311170 | 5.947106 | 24.740704 | 81.466074 | 212.926546 | 461.628514 | 841.849249 | 1216.892497 | 1439.659124 | 1550.763625 | 1432.889131 | 1126.152443 | 809.645566 | 689.437776 | 657.813443 | 611.424619 | 416.684596 | 163.672785 | 42.646169 | 10.916205 | 3.653559 | 1.940227 | 1.428272 | 1.109791 | 0.976797 | 1.032562 | 1.300636 | 1.906153 | 3.578912 | 7.624803 | 1.840870e+01 | 4.108065e+01 | 5.820188e+01 | 4.685363e+01 | 2.995683e+01 | 16.489827 | 9.999312 | 6.555575 | 4.494815 | 3.149166 | 2.406620 | 1.900665 | 1.746638 | 1.795008 | 2.109888 | 2.612709 | 3.355338 | 3.852904 | 4.227718 | 4.273490 | 4.366584 | 4.472698 | 5.304536 | 6.436891 | 8.701156 | 12.106627 | 17.947952 | 27.100356 | 43.343827 | 66.688709 | 94.850054 | 102.353266 | 80.999734 | 46.323052 | 22.201864 | 9.504198 | 4.667561 | 2.646968 | 1.835523 | 1.483912 | 1.419952 | 1.561371 | 1.990723 | 2.738138 | 3.919828 | 5.912178 | 8.493812 | 11.688874 | 15.420173 | 19.271478 | 19.361331 | 14.032156 | 7.164619 | 3.239047 | 1.649441 | 1.010862 | 0.751205 | 0.646160 | 0.636025 | 0.700185 | 0.851964 | 1.119822 | 1.627273 | 2.512774 | 3.359133 | 3.949715 | 4.528656 | 4.936331 | 4.950137 | 4.551177 | 3.962668 | 3.385720 | 3.136096 | 3.210850 | 3.799096 | 4.538868 | 4.931301 | 4.868061 | 4.462003 | 4.191433 | 4.245659 | 3.964037 | 3.543246 | 3.440221 | 3.175495 | 2.980264 | 3.013194 | 3.034215 | 2.790834 | 2.717984 | 2.380765 | 2.204871 | 2.034039 | 1.890984 | 1.662726 | 1.536820 | 1.444298 | 1.388729 | 1.457684 | 1.698297 | 2.142583 | 2.613182 | 3.590790 | 4.876502 | 6.066612 | 5.950545 | 5.061155 | 3.408660 | 2.299410 | 1.530998 | 1.041497 | 0.725917 | 0.534782 | 0.429911 | 0.385352 | 0.379357 | 0.417247 | 0.523361 | 0.767446 | 1.243926 | 1.934055 | 2.551243 | 2.618183 | 2.045881 | 1.243902 | 0.760658 | 0.480245 | 0.332811 | 0.266397 | 0.242519 | 0.233306 | 0.236659 | 0.249857 | 0.266709 | 0.288685 | 0.320572 | 0.371356 | 0.439323 | 0.542778 | 0.717331 | 0.995599 | 1.277508 | 1.464983 | 1.592239 | 1.590274 | 1.390506 | 1.149552 | 0.970336 | 0.848220 | 0.801507 | 0.832393 | 1.032986 | 1.654622 | 2.945460 | 4.080934 | 4.220138 | 3.117835 | 1.933235 | 1.254685 | 0.862118 | 0.635796 | 0.557284 | 0.524089 | 0.522766 | 0.573321 | 0.640430 | 0.623836 | 0.540457 | 0.433947 | 0.332989 | 0.261077 | 0.225480 | 0.214665 | 0.222185 | 0.245216 | 0.279226 | 0.334116 | 0.387086 | 0.436899 | 0.429345 | 0.392110 | 0.341230 | 0.302323 | 0.275303 | 0.264156 | 0.250186 |
| right posterior | 1.003755 | 0.625485 | 0.411423 | 0.233793 | 0.223738 | 0.283928 | 0.367589 | 0.389138 | 0.303230 | 0.238814 | 0.219324 | 0.215451 | 0.214632 | 0.215317 | 0.214838 | 0.223392 | 0.251970 | 0.289460 | 0.305339 | 0.307101 | 0.264248 | 0.216275 | 0.226846 | 0.248578 | 0.273836 | 0.257010 | 0.228376 | 0.214627 | 0.232599 | 0.305476 | 0.375355 | 0.380670 | 0.302178 | 0.225870 | 0.234814 | 0.456189 | 1.363153 | 2.362499 | 1.367661 | 0.694858 | 0.486966 | 0.452030 | 0.512550 | 0.643529 | 0.732107 | 0.740891 | 0.582406 | 0.383552 | 0.239559 | 0.216311 | 0.242547 | 0.248700 | 0.232261 | 0.214627 | 0.258250 | 0.444156 | 0.757213 | 1.180591 | 1.486492 | 1.351477 | 1.273064 | 1.302698 | 1.551432 | 2.397407 | 5.210486 | 14.277810 | 43.707309 | 140.307172 | 523.007223 | 1882.950309 | 5159.361893 | 12065.139191 | 15969.418476 | 13315.750812 | 7242.963705 | 4020.729225 | 1880.233621 | 743.797021 | 257.148662 | 71.465175 | 18.236485 | 5.995407 | 2.890192 | 1.836334 | 1.585078 | 1.688319 | 1.912301 | 2.217542 | 2.751215 | 3.570574 | 4.354916 | 5.332014 | 6.485757 | 7.529342 | 8.290029e+00 | 8.909929e+00 | 8.932951e+00 | 8.468020e+00 | 7.648315e+00 | 6.373795 | 5.276395 | 4.440224 | 3.993382 | 3.682114 | 3.649776 | 3.663545 | 3.672621 | 3.532587 | 3.452115 | 3.311938 | 3.336530 | 3.583841 | 4.415371 | 6.317874 | 10.340327 | 17.788382 | 33.792622 | 73.286924 | 161.006037 | 294.155533 | 449.791339 | 591.301700 | 602.039067 | 451.765677 | 266.543121 | 142.483488 | 71.947549 | 39.847704 | 26.401196 | 19.313326 | 15.142011 | 13.669570 | 13.959156 | 15.099162 | 17.807968 | 21.581914 | 23.066179 | 23.520744 | 23.167282 | 21.715801 | 19.754568 | 18.903099 | 17.910857 | 19.521515 | 25.111979 | 36.310037 | 48.950970 | 58.867674 | 62.013311 | 56.041405 | 43.366558 | 34.178657 | 28.700670 | 26.790921 | 30.842375 | 43.522799 | 69.746514 | 125.556718 | 192.930960 | 217.873781 | 165.531482 | 95.328981 | 44.529358 | 20.560052 | 10.915489 | 7.390392 | 5.992866 | 6.299444 | 7.378166 | 9.573686 | 13.498831 | 19.622002 | 25.574806 | 39.547847 | 64.118724 | 101.909729 | 166.314297 | 241.811484 | 270.826294 | 226.565739 | 149.368701 | 79.373890 | 45.835745 | 26.735783 | 20.039547 | 20.898717 | 30.297733 | 47.153751 | 80.039963 | 136.993411 | 209.619079 | 274.722920 | 358.395859 | 446.792839 | 440.651758 | 382.538214 | 284.700289 | 215.265744 | 170.071563 | 161.580008 | 189.720731 | 258.114113 | 316.765064 | 334.654315 | 291.715432 | 208.186654 | 135.628627 | 95.009259 | 74.470123 | 61.880870 | 64.835477 | 78.615513 | 112.054844 | 179.011944 | 285.719309 | 440.467425 | 853.266934 | 1164.711911 | 1225.020772 | 979.504158 | 467.530649 | 134.030935 | 44.240597 | 18.864485 | 11.493294 | 9.727406 | 10.895488 | 15.336958 | 25.765505 | 41.726970 | 55.082857 | 66.494886 | 71.930164 | 77.664232 | 77.008265 | 70.081701 | 62.787835 | 60.631568 | 52.362802 | 49.126273 | 46.564734 | 44.554033 | 46.719923 | 49.552099 | 41.340242 | 33.784769 | 27.853566 | 25.537629 | 27.844214 | 30.385827 | 27.346326 | 21.372474 | 15.575035 | 11.238929 | 8.852332 | 8.235940 | 9.071070 | 11.221229 | 14.673848 | 20.131955 | 26.413477 | 34.666259 | 46.834408 | 62.011260 | 78.113608 | 76.755220 | 54.130826 | 27.660154 | 12.189296 | 5.166820 | 2.520306 | 1.572254 | 1.253546 | 1.264063 | 1.656024 | 2.614800 | 4.786162 | 9.136417 | 12.726874 | 16.930805 |
| all electrodes | 0.231319 | 0.238657 | 0.239395 | 0.241222 | 0.235079 | 0.235679 | 0.234284 | 0.241957 | 0.253451 | 0.267397 | 0.264638 | 0.305321 | 0.365808 | 0.503701 | 0.787298 | 1.400178 | 2.111840 | 3.341238 | 5.054049 | 6.440995 | 4.239399 | 1.260076 | 0.394840 | 0.231460 | 0.219555 | 0.220952 | 0.222354 | 0.344956 | 0.787983 | 1.599229 | 1.747216 | 1.538442 | 0.855628 | 0.389515 | 0.228048 | 0.223032 | 0.328958 | 0.478097 | 0.417539 | 0.305717 | 0.242586 | 0.214633 | 0.228781 | 0.252958 | 0.261257 | 0.262220 | 0.258398 | 0.271236 | 0.340595 | 0.453281 | 0.544975 | 0.593338 | 0.610136 | 0.458050 | 0.343326 | 0.260642 | 0.225034 | 0.217589 | 0.219225 | 0.226256 | 0.239091 | 0.225922 | 0.218819 | 0.306066 | 0.789658 | 3.934696 | 26.083707 | 150.349523 | 651.557406 | 1757.524599 | 4556.981001 | 8729.917268 | 15282.323418 | 23252.497008 | 29662.321596 | 19672.800306 | 7489.263211 | 2229.435557 | 755.379433 | 240.558768 | 72.834131 | 29.924362 | 14.427681 | 8.963554 | 6.454199 | 5.125483 | 5.034590 | 7.240202 | 10.917735 | 28.489080 | 129.871187 | 497.077032 | 1463.069575 | 4440.527717 | 6.656124e+03 | 6.673956e+03 | 5.025754e+03 | 1.745709e+03 | 6.568306e+02 | 314.422627 | 166.916326 | 91.796208 | 46.034246 | 18.936800 | 9.541564 | 5.198289 | 3.190040 | 2.724580 | 3.267677 | 5.057189 | 9.584136 | 21.141208 | 46.503343 | 110.025316 | 217.321998 | 289.684971 | 292.281385 | 289.501507 | 286.478770 | 256.598834 | 350.181837 | 596.270355 | 992.601157 | 1387.113858 | 1737.198790 | 1569.342728 | 1558.290222 | 1094.522286 | 683.165231 | 345.512862 | 156.204168 | 78.049989 | 51.424818 | 33.375492 | 30.388336 | 38.964936 | 64.209662 | 130.256819 | 315.930164 | 845.878207 | 1695.460994 | 1466.104279 | 632.572579 | 320.864870 | 171.966699 | 102.906600 | 75.756543 | 62.760575 | 62.999700 | 78.606408 | 84.232387 | 80.555515 | 78.175042 | 62.768950 | 49.167592 | 35.012521 | 23.607384 | 13.982666 | 6.563007 | 2.944199 | 1.514777 | 0.930580 | 0.651271 | 0.547636 | 0.446765 | 0.366290 | 0.322231 | 0.292886 | 0.268187 | 0.258429 | 0.251898 | 0.241890 | 0.243640 | 0.245524 | 0.252847 | 0.278164 | 0.337252 | 0.402965 | 0.517110 | 0.663394 | 0.779241 | 0.796582 | 0.779329 | 0.744509 | 0.703340 | 0.768249 | 0.801954 | 0.789946 | 0.750930 | 0.736161 | 0.680537 | 0.723118 | 0.802531 | 0.980181 | 1.135133 | 1.227156 | 1.206487 | 1.076399 | 0.808537 | 0.555906 | 0.398622 | 0.331233 | 0.304757 | 0.292618 | 0.296371 | 0.299744 | 0.318107 | 0.345873 | 0.369779 | 0.400753 | 0.449703 | 0.450357 | 0.424200 | 0.384923 | 0.361722 | 0.359630 | 0.400389 | 0.489289 | 0.649753 | 0.830447 | 0.914971 | 0.731756 | 0.543197 | 0.401707 | 0.311595 | 0.254240 | 0.233892 | 0.222765 | 0.220240 | 0.218321 | 0.216296 | 0.215029 | 0.217968 | 0.222204 | 0.221692 | 0.218931 | 0.215372 | 0.215561 | 0.227742 | 0.253700 | 0.279396 | 0.292642 | 0.280367 | 0.245432 | 0.218979 | 0.214870 | 0.222280 | 0.244398 | 0.273647 | 0.283686 | 0.307378 | 0.318715 | 0.328180 | 0.362617 | 0.430539 | 0.527301 | 0.645631 | 0.641195 | 0.499853 | 0.384128 | 0.281557 | 0.229526 | 0.216114 | 0.217034 | 0.239601 | 0.364759 | 0.855414 | 3.324402 | 9.973029 | 13.129770 | 7.856443 | 3.370474 | 1.632254 | 1.133185 | 0.873103 | 0.760941 | 0.694441 | 0.599299 | 0.519218 | 0.474071 |

X) neural network features, neutral vs sad

  
|  | time window | peak latency | cluster *p* | peak Cohen's *d* |  | | | |  |
| **all electrodes** | 125 - 185 ms | 155 ms | 0.0468 | 1.2828 |  | | | |  |
 935 - 1080 ms | 965 ms | 0.0232 | 0.5656 |  | | | ||  |  | | | | | | | | |

Model correlations, cluster permutation tests

|  | **left hemisphere** | | | | **right hemisphere** | | | |  |
|  | time window | peak latency | cluster *p* | peak Cohen's *d* | time window | peak latency | cluster *p* | peak Cohen's *d* |  |
| **anterior** |  | | | |  | | | |  |
| **central** | 135 - 245 ms | 160 ms | 0.0459 | 0.6496 | -200 - -135 ms | -195 ms | 0.0435 | -0.7561 |  |
| **posterior** |  | | | |  | | | |  |

  

Model correlations, Bayesian statistics

|  | -200 | -195 | -190 | -185 | -180 | -175 | -170 | -165 | -160 | -155 | -150 | -145 | -140 | -135 | -130 | -125 | -120 | -115 | -110 | -105 | -100 | -95 | -90 | -85 | -80 | -75 | -70 | -65 | -60 | -55 | -50 | -45 | -40 | -35 | -30 | -25 | -20 | -15 | -10 | -5 | 0 | 5 | 10 | 15 | 20 | 25 | 30 | 35 | 40 | 45 | 50 | 55 | 60 | 65 | 70 | 75 | 80 | 85 | 90 | 95 | 100 | 105 | 110 | 115 | 120 | 125 | 130 | 135 | 140 | 145 | 150 | 155 | 160 | 165 | 170 | 175 | 180 | 185 | 190 | 195 | 200 | 205 | 210 | 215 | 220 | 225 | 230 | 235 | 240 | 245 | 250 | 255 | 260 | 265 | 270 | 275 | 280 | 285 | 290 | 295 | 300 | 305 | 310 | 315 | 320 | 325 | 330 | 335 | 340 | 345 | 350 | 355 | 360 | 365 | 370 | 375 | 380 | 385 | 390 | 395 | 400 | 405 | 410 | 415 | 420 | 425 | 430 | 435 | 440 | 445 | 450 | 455 | 460 | 465 | 470 | 475 | 480 | 485 | 490 | 495 | 500 | 505 | 510 | 515 | 520 | 525 | 530 | 535 | 540 | 545 | 550 | 555 | 560 | 565 | 570 | 575 | 580 | 585 | 590 | 595 | 600 | 605 | 610 | 615 | 620 | 625 | 630 | 635 | 640 | 645 | 650 | 655 | 660 | 665 | 670 | 675 | 680 | 685 | 690 | 695 | 700 | 705 | 710 | 715 | 720 | 725 | 730 | 735 | 740 | 745 | 750 | 755 | 760 | 765 | 770 | 775 | 780 | 785 | 790 | 795 | 800 | 805 | 810 | 815 | 820 | 825 | 830 | 835 | 840 | 845 | 850 | 855 | 860 | 865 | 870 | 875 | 880 | 885 | 890 | 895 | 900 | 905 | 910 | 915 | 920 | 925 | 930 | 935 | 940 | 945 | 950 | 955 | 960 | 965 | 970 | 975 | 980 | 985 | 990 | 995 | 1000 | 1005 | 1010 | 1015 | 1020 | 1025 | 1030 | 1035 | 1040 | 1045 | 1050 | 1055 | 1060 | 1065 | 1070 | 1075 | 1080 | 1085 | 1090 | 1095 | 1100 | 1105 | 1110 | 1115 | 1120 | 1125 | 1130 | 1135 | 1140 | 1145 | 1150 | 1155 | 1160 | 1165 | 1170 | 1175 | 1180 | 1185 | 1190 | 1195 |
| --- | --- | --- | --- | --- | --- | --- | --- | --- | --- | --- | --- | --- | --- | --- | --- | --- | --- | --- | --- | --- | --- | --- | --- | --- | --- | --- | --- | --- | --- | --- | --- | --- | --- | --- | --- | --- | --- | --- | --- | --- | --- | --- | --- | --- | --- | --- | --- | --- | --- | --- | --- | --- | --- | --- | --- | --- | --- | --- | --- | --- | --- | --- | --- | --- | --- | --- | --- | --- | --- | --- | --- | --- | --- | --- | --- | --- | --- | --- | --- | --- | --- | --- | --- | --- | --- | --- | --- | --- | --- | --- | --- | --- | --- | --- | --- | --- | --- | --- | --- | --- | --- | --- | --- | --- | --- | --- | --- | --- | --- | --- | --- | --- | --- | --- | --- | --- | --- | --- | --- | --- | --- | --- | --- | --- | --- | --- | --- | --- | --- | --- | --- | --- | --- | --- | --- | --- | --- | --- | --- | --- | --- | --- | --- | --- | --- | --- | --- | --- | --- | --- | --- | --- | --- | --- | --- | --- | --- | --- | --- | --- | --- | --- | --- | --- | --- | --- | --- | --- | --- | --- | --- | --- | --- | --- | --- | --- | --- | --- | --- | --- | --- | --- | --- | --- | --- | --- | --- | --- | --- | --- | --- | --- | --- | --- | --- | --- | --- | --- | --- | --- | --- | --- | --- | --- | --- | --- | --- | --- | --- | --- | --- | --- | --- | --- | --- | --- | --- | --- | --- | --- | --- | --- | --- | --- | --- | --- | --- | --- | --- | --- | --- | --- | --- | --- | --- | --- | --- | --- | --- | --- | --- | --- | --- | --- | --- | --- | --- | --- | --- | --- | --- | --- | --- | --- | --- | --- | --- | --- | --- | --- | --- | --- | --- | --- | --- | --- | --- | --- | --- | --- | --- | --- | --- | --- | --- | --- | --- | --- | --- | --- |
| left anterior | 0.524944 | 0.466762 | 0.451973 | 0.348322 | 0.262598 | 0.233784 | 0.217761 | 0.214628 | 0.230708 | 0.242754 | 0.242821 | 0.230328 | 0.227055 | 0.215666 | 0.216780 | 0.240762 | 0.257902 | 0.252460 | 0.241711 | 0.239202 | 0.233245 | 0.233580 | 0.249832 | 0.288736 | 0.356618 | 0.393650 | 0.433596 | 0.433278 | 0.386531 | 0.277349 | 0.227022 | 0.216298 | 0.254937 | 0.432388 | 0.860745 | 1.303041 | 1.280523 | 1.444278 | 1.434746 | 1.598658 | 2.098527 | 4.472509 | 11.922995 | 36.107167 | 98.492391 | 125.124271 | 102.585276 | 54.354517 | 28.044169 | 13.181308 | 6.446926 | 2.047312 | 1.023413 | 0.578811 | 0.362633 | 0.265716 | 0.233274 | 0.218153 | 0.215363 | 0.215606 | 0.216090 | 0.215119 | 0.214629 | 0.220386 | 0.250022 | 0.322085 | 0.480411 | 0.740561 | 1.075954 | 1.409122 | 1.709276 | 1.658679 | 1.543480 | 1.288493 | 0.964042 | 0.595694 | 0.393265 | 0.278424 | 0.227165 | 0.214626 | 0.220510 | 0.223390 | 0.219414 | 0.215246 | 0.214849 | 0.220075 | 0.225678 | 0.226418 | 0.225348 | 0.227552 | 0.233421 | 0.248617 | 0.273001 | 0.313860 | 0.353487 | 0.360010 | 0.350306 | 0.347233 | 0.318311 | 0.283633 | 0.252925 | 0.233713 | 0.231610 | 0.238000 | 0.248205 | 0.270916 | 0.306160 | 0.323829 | 0.339637 | 0.333193 | 0.320262 | 0.294645 | 0.269429 | 0.250120 | 0.241000 | 0.233006 | 0.232541 | 0.231244 | 0.228379 | 0.224619 | 0.220742 | 0.217475 | 0.216310 | 0.215003 | 0.214626 | 0.214728 | 0.216252 | 0.226131 | 0.256495 | 0.333926 | 0.490133 | 0.711512 | 0.805157 | 0.681319 | 0.468136 | 0.360651 | 0.300363 | 0.265551 | 0.244645 | 0.234507 | 0.228296 | 0.223469 | 0.219607 | 0.216293 | 0.215282 | 0.215172 | 0.215520 | 0.215957 | 0.216970 | 0.215963 | 0.215308 | 0.226100 | 0.249577 | 0.291051 | 0.347435 | 0.422727 | 0.528903 | 0.617491 | 0.686506 | 0.782992 | 0.812814 | 0.848393 | 0.867915 | 0.856664 | 0.845776 | 0.873335 | 0.841541 | 0.870318 | 0.887237 | 0.851816 | 0.748522 | 0.621537 | 0.481042 | 0.384543 | 0.325853 | 0.302396 | 0.304972 | 0.338048 | 0.403383 | 0.519178 | 0.695085 | 0.845731 | 0.896526 | 0.783379 | 0.609641 | 0.435804 | 0.328453 | 0.262898 | 0.233972 | 0.221156 | 0.215257 | 0.214946 | 0.215802 | 0.215174 | 0.214627 | 0.218532 | 0.242598 | 0.310496 | 0.437406 | 0.644598 | 0.903449 | 1.011942 | 0.767123 | 0.449228 | 0.280362 | 0.229617 | 0.219153 | 0.222673 | 0.249946 | 0.331900 | 0.478552 | 0.686058 | 0.874535 | 0.997333 | 1.051827 | 1.071804 | 0.996715 | 0.950606 | 0.920781 | 0.937508 | 0.907160 | 0.860538 | 0.809846 | 0.753696 | 0.709359 | 0.706141 | 0.782256 | 0.893789 | 1.026244 | 0.953227 | 0.758369 | 0.559727 | 0.449374 | 0.371062 | 0.353358 | 0.375356 | 0.452540 | 0.513605 | 0.565689 | 0.581725 | 0.562636 | 0.490265 | 0.415044 | 0.336415 | 0.288055 | 0.249705 | 0.225444 | 0.218058 | 0.216353 | 0.216257 | 0.218266 | 0.219867 | 0.219306 | 0.216895 | 0.214690 | 0.217203 | 0.226120 | 0.241627 | 0.253183 | 0.258526 | 0.258295 | 0.248214 | 0.232302 | 0.222133 | 0.217416 | 0.215496 | 0.214998 | 0.214633 | 0.214854 | 0.216166 | 0.219664 | 0.224722 | 0.228013 | 0.230640 | 0.230896 | 0.230296 | 0.235399 | 0.250147 | 0.255300 | 0.267937 |
| right anterior | 0.229678 | 0.223438 | 0.215699 | 0.216314 | 0.218153 | 0.217008 | 0.214807 | 0.218807 | 0.251601 | 0.281766 | 0.269071 | 0.243719 | 0.229928 | 0.215173 | 0.224363 | 0.274823 | 0.394689 | 0.477954 | 0.449688 | 0.368595 | 0.282276 | 0.228820 | 0.214808 | 0.235791 | 0.290974 | 0.365212 | 0.402493 | 0.341476 | 0.269135 | 0.223136 | 0.216838 | 0.254996 | 0.343573 | 0.423227 | 0.365495 | 0.275394 | 0.228193 | 0.217831 | 0.214812 | 0.214638 | 0.215198 | 0.216909 | 0.217002 | 0.218137 | 0.217638 | 0.216707 | 0.217773 | 0.216066 | 0.214843 | 0.224362 | 0.264592 | 0.362817 | 0.544844 | 0.884479 | 1.424258 | 2.149233 | 3.373916 | 3.772681 | 3.202242 | 1.851524 | 0.937949 | 0.550872 | 0.445354 | 0.467425 | 0.700704 | 1.456162 | 3.505998 | 6.438268 | 9.277944 | 11.231088 | 11.666784 | 9.802998 | 6.846514 | 4.390461 | 3.077853 | 2.448281 | 2.086994 | 1.887908 | 1.674638 | 1.278706 | 0.889359 | 0.691982 | 0.636827 | 0.688260 | 0.859938 | 1.148261 | 1.559850 | 2.055422 | 2.349168 | 2.804527 | 3.241299 | 3.265982 | 3.372528 | 3.751154 | 4.062574 | 4.341102 | 5.000716 | 5.528065 | 5.524644 | 3.634729 | 2.206240 | 1.382583 | 0.912427 | 0.631720 | 0.579888 | 0.625613 | 0.786075 | 1.007191 | 1.257399 | 1.331441 | 1.313839 | 1.144464 | 0.946763 | 0.775704 | 0.642385 | 0.470251 | 0.348691 | 0.269319 | 0.230939 | 0.216695 | 0.214757 | 0.217883 | 0.223393 | 0.236470 | 0.257869 | 0.299179 | 0.374906 | 0.482194 | 0.556858 | 0.551289 | 0.492152 | 0.433900 | 0.363560 | 0.304986 | 0.272392 | 0.263605 | 0.269527 | 0.293095 | 0.344439 | 0.452049 | 0.629611 | 0.873648 | 1.216187 | 1.498436 | 1.472671 | 1.287750 | 1.077718 | 0.740346 | 0.506702 | 0.353012 | 0.271614 | 0.233567 | 0.217953 | 0.214879 | 0.220658 | 0.231855 | 0.237199 | 0.230313 | 0.219310 | 0.215803 | 0.239509 | 0.280805 | 0.338381 | 0.377649 | 0.366567 | 0.351454 | 0.325932 | 0.294970 | 0.282058 | 0.279890 | 0.285309 | 0.297374 | 0.299237 | 0.293463 | 0.288174 | 0.274567 | 0.259714 | 0.247391 | 0.244669 | 0.256120 | 0.274118 | 0.305112 | 0.356790 | 0.414905 | 0.472833 | 0.508818 | 0.543805 | 0.621917 | 0.697385 | 0.697199 | 0.706288 | 0.663779 | 0.579392 | 0.519201 | 0.441419 | 0.350165 | 0.300850 | 0.272327 | 0.262548 | 0.277141 | 0.321019 | 0.370019 | 0.428319 | 0.439946 | 0.412099 | 0.341322 | 0.284068 | 0.242882 | 0.221686 | 0.214627 | 0.217611 | 0.223467 | 0.232815 | 0.248993 | 0.268173 | 0.276091 | 0.284050 | 0.290585 | 0.289555 | 0.274159 | 0.275151 | 0.273348 | 0.269009 | 0.262406 | 0.253980 | 0.236470 | 0.223325 | 0.214985 | 0.216847 | 0.216164 | 0.214633 | 0.217327 | 0.226356 | 0.236164 | 0.224048 | 0.215989 | 0.216102 | 0.226584 | 0.244167 | 0.237667 | 0.222540 | 0.215199 | 0.240033 | 0.289638 | 0.316423 | 0.291324 | 0.248439 | 0.222035 | 0.214643 | 0.219420 | 0.224382 | 0.222999 | 0.217275 | 0.214836 | 0.215745 | 0.220752 | 0.229187 | 0.232602 | 0.241458 | 0.244956 | 0.245406 | 0.245679 | 0.251810 | 0.252198 | 0.251451 | 0.250785 | 0.248091 | 0.239170 | 0.227773 | 0.219518 | 0.215996 | 0.214680 | 0.217817 | 0.228972 | 0.246553 | 0.270053 | 0.287201 | 0.297679 | 0.296858 | 0.282306 |
| left central | 0.989342 | 0.975476 | 1.196193 | 1.332436 | 1.346359 | 1.015721 | 0.743917 | 0.446245 | 0.310352 | 0.230537 | 0.214629 | 0.228887 | 0.270457 | 0.314261 | 0.289388 | 0.249418 | 0.223074 | 0.214754 | 0.241321 | 0.284006 | 0.308297 | 0.281280 | 0.271222 | 0.253533 | 0.250085 | 0.234435 | 0.229239 | 0.224209 | 0.217949 | 0.215237 | 0.221365 | 0.234307 | 0.244166 | 0.256417 | 0.246030 | 0.234501 | 0.228444 | 0.226706 | 0.222557 | 0.224768 | 0.234819 | 0.256471 | 0.284938 | 0.334502 | 0.385709 | 0.415799 | 0.376134 | 0.315140 | 0.257714 | 0.223184 | 0.214678 | 0.222167 | 0.236182 | 0.262803 | 0.288356 | 0.329106 | 0.370531 | 0.396119 | 0.405416 | 0.412842 | 0.404531 | 0.400540 | 0.416368 | 0.433713 | 0.476074 | 0.590763 | 0.820394 | 1.325599 | 2.306333 | 3.708884 | 6.053393 | 10.422466 | 12.062002 | 10.668966 | 7.735797 | 4.652054 | 2.795272 | 1.923198 | 1.500612 | 1.513011 | 1.841191 | 2.172650 | 2.648597 | 3.554948 | 4.093669 | 4.022455 | 3.053257 | 2.142538 | 1.647654 | 1.201874 | 0.968082 | 0.823045 | 0.690636 | 0.592942 | 0.534696 | 0.470123 | 0.429850 | 0.380458 | 0.326364 | 0.288374 | 0.269671 | 0.271469 | 0.292819 | 0.326268 | 0.347163 | 0.372914 | 0.373573 | 0.334664 | 0.261055 | 0.225073 | 0.214993 | 0.226361 | 0.243797 | 0.245303 | 0.237663 | 0.224410 | 0.220976 | 0.219615 | 0.216875 | 0.214772 | 0.214705 | 0.215774 | 0.219784 | 0.229497 | 0.243923 | 0.253854 | 0.253520 | 0.239641 | 0.221115 | 0.215102 | 0.216585 | 0.223214 | 0.227689 | 0.226539 | 0.227264 | 0.229013 | 0.243098 | 0.252044 | 0.253200 | 0.259378 | 0.261979 | 0.248199 | 0.240363 | 0.231062 | 0.223762 | 0.219432 | 0.217467 | 0.214633 | 0.219396 | 0.237648 | 0.261536 | 0.282917 | 0.281812 | 0.269936 | 0.264021 | 0.258421 | 0.252197 | 0.232972 | 0.218377 | 0.215665 | 0.238991 | 0.321521 | 0.482161 | 0.761596 | 1.026757 | 1.192313 | 1.222612 | 1.067043 | 0.809354 | 0.612115 | 0.440360 | 0.293156 | 0.224164 | 0.225592 | 0.326003 | 0.531895 | 0.676061 | 0.770502 | 0.702442 | 0.692858 | 0.647537 | 0.528348 | 0.429367 | 0.387456 | 0.321479 | 0.300894 | 0.290437 | 0.277347 | 0.274067 | 0.292656 | 0.312467 | 0.391009 | 0.591123 | 0.955456 | 1.470721 | 1.800796 | 1.483839 | 0.999704 | 0.670910 | 0.471540 | 0.396301 | 0.384485 | 0.417764 | 0.516858 | 0.756728 | 1.165127 | 1.722243 | 2.055726 | 1.813562 | 1.320045 | 0.913755 | 0.697330 | 0.593084 | 0.522068 | 0.524301 | 0.576933 | 0.588053 | 0.624146 | 0.757002 | 0.774689 | 0.608345 | 0.404947 | 0.273328 | 0.226665 | 0.215121 | 0.217472 | 0.221447 | 0.218865 | 0.216393 | 0.214716 | 0.214729 | 0.214642 | 0.215503 | 0.221366 | 0.239501 | 0.239593 | 0.228357 | 0.221149 | 0.218640 | 0.216700 | 0.216001 | 0.216008 | 0.217334 | 0.218914 | 0.222580 | 0.233414 | 0.256453 | 0.294261 | 0.326932 | 0.335732 | 0.297908 | 0.246467 | 0.216642 | 0.223734 | 0.265606 | 0.321358 | 0.368856 | 0.391667 | 0.394796 | 0.400469 | 0.392429 | 0.384013 | 0.373677 | 0.390107 | 0.420031 | 0.458376 | 0.470587 | 0.483762 | 0.429327 | 0.353525 | 0.287819 | 0.250943 | 0.231951 | 0.230195 | 0.228546 | 0.231720 | 0.240806 | 0.252648 | 0.251580 | 0.257703 |
| right central | 18.299340 | 22.997676 | 30.581411 | 50.841453 | 83.863465 | 104.237017 | 110.714486 | 207.009452 | 873.454689 | 978.647511 | 349.716140 | 49.129954 | 7.605408 | 1.553947 | 0.541275 | 0.318461 | 0.254103 | 0.219822 | 0.214875 | 0.223866 | 0.238506 | 0.247234 | 0.239751 | 0.225463 | 0.216525 | 0.256024 | 0.389743 | 0.714402 | 1.061344 | 0.679283 | 0.314429 | 0.214747 | 0.246338 | 0.277596 | 0.265334 | 0.257673 | 0.235246 | 0.220249 | 0.214671 | 0.221512 | 0.244652 | 0.254045 | 0.256461 | 0.242446 | 0.223591 | 0.214668 | 0.220678 | 0.254631 | 0.307735 | 0.374760 | 0.443859 | 0.466089 | 0.405542 | 0.326448 | 0.249450 | 0.215486 | 0.219291 | 0.226385 | 0.226752 | 0.216988 | 0.214909 | 0.217919 | 0.221996 | 0.222948 | 0.223779 | 0.247346 | 0.323727 | 0.568123 | 1.534416 | 4.621596 | 9.634841 | 14.977690 | 16.445747 | 11.435372 | 6.511622 | 3.236613 | 1.607686 | 0.882461 | 0.530461 | 0.335660 | 0.268532 | 0.237487 | 0.236198 | 0.249144 | 0.271206 | 0.282932 | 0.298283 | 0.293396 | 0.294056 | 0.289849 | 0.286441 | 0.281623 | 0.281662 | 0.274075 | 0.275365 | 0.281693 | 0.285746 | 0.285779 | 0.296826 | 0.317300 | 0.365262 | 0.452652 | 0.498090 | 0.463894 | 0.342906 | 0.249900 | 0.214708 | 0.234932 | 0.297161 | 0.324996 | 0.319756 | 0.289617 | 0.269627 | 0.248768 | 0.241068 | 0.240909 | 0.246111 | 0.241730 | 0.229943 | 0.218781 | 0.214979 | 0.226338 | 0.242772 | 0.251138 | 0.257626 | 0.253277 | 0.235041 | 0.221290 | 0.217040 | 0.216591 | 0.217001 | 0.218719 | 0.220558 | 0.221421 | 0.219701 | 0.217223 | 0.215363 | 0.214933 | 0.214679 | 0.217063 | 0.223571 | 0.236315 | 0.265570 | 0.308653 | 0.373175 | 0.466031 | 0.531094 | 0.524354 | 0.447942 | 0.360949 | 0.290264 | 0.253397 | 0.241451 | 0.239332 | 0.248987 | 0.275276 | 0.296722 | 0.296118 | 0.275959 | 0.241541 | 0.219565 | 0.214683 | 0.218891 | 0.221036 | 0.222878 | 0.225742 | 0.228078 | 0.226348 | 0.224547 | 0.217113 | 0.216765 | 0.254526 | 0.351688 | 0.463564 | 0.544379 | 0.552621 | 0.460779 | 0.386098 | 0.357608 | 0.336072 | 0.328685 | 0.331208 | 0.330423 | 0.329957 | 0.317874 | 0.289734 | 0.262697 | 0.234232 | 0.216048 | 0.219540 | 0.239909 | 0.280683 | 0.318987 | 0.320104 | 0.290725 | 0.258423 | 0.230276 | 0.216679 | 0.215672 | 0.223237 | 0.229337 | 0.238835 | 0.256798 | 0.267305 | 0.260461 | 0.263829 | 0.262361 | 0.270115 | 0.286583 | 0.304962 | 0.300601 | 0.292363 | 0.263417 | 0.242078 | 0.234557 | 0.228559 | 0.224922 | 0.229517 | 0.235941 | 0.242065 | 0.249754 | 0.250027 | 0.236465 | 0.219399 | 0.215411 | 0.227500 | 0.244262 | 0.258524 | 0.265507 | 0.265402 | 0.260175 | 0.255345 | 0.250542 | 0.240947 | 0.225428 | 0.215663 | 0.216247 | 0.230843 | 0.269731 | 0.323505 | 0.378901 | 0.426958 | 0.466989 | 0.499654 | 0.537104 | 0.561909 | 0.634583 | 0.698881 | 0.779696 | 0.711146 | 0.711057 | 0.762695 | 0.836660 | 0.781096 | 0.638212 | 0.430615 | 0.290733 | 0.221232 | 0.227001 | 0.278692 | 0.331719 | 0.371694 | 0.320383 | 0.254434 | 0.218161 | 0.221143 | 0.258162 | 0.291537 | 0.324277 | 0.343608 | 0.363154 | 0.385599 | 0.419566 | 0.469576 | 0.535978 | 0.609337 | 0.660439 | 0.699633 | 0.724537 | 0.702416 |
| left posterior | 0.904070 | 1.140521 | 1.474382 | 2.433555 | 2.168993 | 1.690143 | 0.980626 | 0.595613 | 0.372096 | 0.246302 | 0.215044 | 0.225005 | 0.255923 | 0.274144 | 0.279558 | 0.269527 | 0.241974 | 0.223152 | 0.216165 | 0.215580 | 0.219747 | 0.230783 | 0.242594 | 0.239999 | 0.220187 | 0.214705 | 0.226162 | 0.300574 | 0.503680 | 1.070967 | 2.621118 | 4.399348 | 5.362470 | 3.818217 | 1.302519 | 0.527729 | 0.293925 | 0.219441 | 0.220978 | 0.230199 | 0.223080 | 0.214818 | 0.217305 | 0.223645 | 0.230234 | 0.234483 | 0.235307 | 0.239361 | 0.257410 | 0.305718 | 0.401175 | 0.639111 | 1.060279 | 1.793738 | 2.387762 | 3.316029 | 4.516831 | 5.153085 | 4.232570 | 3.141020 | 1.800913 | 0.845078 | 0.308813 | 0.215691 | 0.333327 | 0.712461 | 1.770932 | 4.572677 | 11.553984 | 26.667783 | 57.171897 | 94.585685 | 121.648864 | 107.945185 | 73.521076 | 44.312916 | 20.730246 | 8.100723 | 3.256270 | 1.432184 | 0.739486 | 0.470595 | 0.395346 | 0.420694 | 0.507978 | 0.618979 | 0.770415 | 0.856394 | 0.825127 | 0.786693 | 0.769296 | 0.748056 | 0.763648 | 0.785098 | 0.804155 | 0.788058 | 0.726729 | 0.651360 | 0.586232 | 0.542646 | 0.551305 | 0.610907 | 0.767572 | 1.074072 | 1.538698 | 2.051390 | 2.520140 | 2.769632 | 2.919337 | 3.077420 | 3.472822 | 4.100719 | 4.555451 | 5.026854 | 5.392166 | 5.977132 | 6.956433 | 8.633610 | 10.861574 | 13.240068 | 14.919313 | 16.874466 | 18.592115 | 17.189581 | 14.184844 | 9.186435 | 4.838593 | 2.422672 | 1.317359 | 0.775358 | 0.557164 | 0.449001 | 0.417674 | 0.399268 | 0.390997 | 0.381383 | 0.372280 | 0.349587 | 0.328752 | 0.310314 | 0.293064 | 0.278455 | 0.272043 | 0.271549 | 0.276021 | 0.277402 | 0.269466 | 0.258369 | 0.248660 | 0.240644 | 0.234715 | 0.230123 | 0.225907 | 0.223010 | 0.220149 | 0.221049 | 0.226232 | 0.239909 | 0.265723 | 0.317319 | 0.410878 | 0.584456 | 0.835663 | 1.101035 | 1.228913 | 1.112086 | 0.811356 | 0.552867 | 0.385909 | 0.287650 | 0.234237 | 0.215292 | 0.220376 | 0.236715 | 0.240466 | 0.228915 | 0.215575 | 0.224734 | 0.279671 | 0.376134 | 0.505893 | 0.606275 | 0.604920 | 0.554693 | 0.466404 | 0.375055 | 0.320355 | 0.290874 | 0.268025 | 0.255801 | 0.255290 | 0.252392 | 0.249297 | 0.246769 | 0.245493 | 0.243120 | 0.252397 | 0.274562 | 0.323109 | 0.409201 | 0.569965 | 0.776813 | 1.033759 | 1.207582 | 1.335248 | 1.444534 | 1.496677 | 1.409973 | 1.360921 | 1.366991 | 1.357946 | 1.386204 | 1.409912 | 1.414120 | 1.255234 | 1.065321 | 0.897421 | 0.762498 | 0.656578 | 0.621483 | 0.604986 | 0.620450 | 0.717191 | 0.895616 | 1.120576 | 1.386824 | 1.559324 | 1.567873 | 1.466307 | 1.349789 | 1.257199 | 1.326849 | 1.571931 | 2.135082 | 3.142417 | 4.915452 | 7.664217 | 10.254394 | 10.096723 | 8.013482 | 5.537616 | 3.471487 | 2.115727 | 1.387343 | 0.941756 | 0.673685 | 0.558443 | 0.506849 | 0.467443 | 0.442074 | 0.415488 | 0.384364 | 0.356190 | 0.336985 | 0.329517 | 0.329081 | 0.316709 | 0.308420 | 0.293558 | 0.274724 | 0.257801 | 0.245587 | 0.235368 | 0.231107 | 0.229570 | 0.233240 | 0.243289 | 0.263266 | 0.287274 | 0.302872 | 0.297548 | 0.288069 | 0.260948 | 0.237506 | 0.222735 | 0.217077 | 0.215262 | 0.215067 | 0.214799 | 0.215403 |
| right posterior | 0.343598 | 0.417027 | 0.607211 | 0.959781 | 1.329423 | 1.123714 | 0.825975 | 0.450385 | 0.312474 | 0.235088 | 0.217452 | 0.214819 | 0.217937 | 0.228510 | 0.229384 | 0.234895 | 0.233366 | 0.259158 | 0.350484 | 0.430863 | 0.374203 | 0.326489 | 0.253137 | 0.214822 | 0.251136 | 0.426807 | 0.871782 | 1.565764 | 2.059989 | 1.542361 | 0.718461 | 0.397724 | 0.346760 | 0.377386 | 0.406109 | 0.400966 | 0.397436 | 0.383074 | 0.404379 | 0.475473 | 0.585485 | 0.757837 | 0.979903 | 1.042526 | 0.875860 | 0.711602 | 0.604428 | 0.527467 | 0.540778 | 0.744810 | 1.160025 | 1.601138 | 1.651432 | 1.698594 | 1.220737 | 0.822957 | 0.533645 | 0.349858 | 0.263381 | 0.236258 | 0.218593 | 0.216057 | 0.247282 | 0.377952 | 0.742822 | 1.542741 | 3.359299 | 6.603928 | 11.070266 | 18.292824 | 26.623428 | 28.961746 | 22.428644 | 14.050833 | 7.815790 | 4.227147 | 2.129880 | 1.021047 | 0.527068 | 0.314452 | 0.232466 | 0.215301 | 0.214627 | 0.216634 | 0.229014 | 0.269422 | 0.341567 | 0.497097 | 0.808443 | 1.443932 | 2.731888 | 5.335032 | 7.692736 | 9.422152 | 9.837051 | 8.941799 | 7.858469 | 6.894126 | 5.488659 | 4.822013 | 4.182168 | 3.693999 | 3.026570 | 2.544933 | 2.062703 | 1.716950 | 1.398112 | 1.243319 | 1.122743 | 1.068385 | 1.049729 | 1.128964 | 1.281314 | 1.564208 | 1.904966 | 2.455431 | 3.055334 | 3.592557 | 4.461599 | 5.903686 | 7.628607 | 9.843312 | 11.929977 | 13.536520 | 14.781935 | 14.836125 | 12.340586 | 8.911541 | 5.506646 | 2.853345 | 1.493598 | 0.909841 | 0.674891 | 0.610310 | 0.662083 | 0.785026 | 0.919626 | 1.068411 | 1.080083 | 1.032318 | 0.944336 | 0.817290 | 0.751374 | 0.806664 | 0.913804 | 0.973889 | 0.995699 | 0.878294 | 0.674324 | 0.477325 | 0.358499 | 0.296578 | 0.269136 | 0.262200 | 0.271237 | 0.295912 | 0.349404 | 0.458592 | 0.665454 | 1.048655 | 1.660090 | 2.431720 | 3.031970 | 3.086456 | 2.454591 | 1.597318 | 0.876537 | 0.507507 | 0.343664 | 0.275937 | 0.247856 | 0.240052 | 0.240495 | 0.252142 | 0.275565 | 0.319599 | 0.383409 | 0.454176 | 0.501520 | 0.507696 | 0.459860 | 0.389294 | 0.330445 | 0.301421 | 0.290518 | 0.292533 | 0.313164 | 0.345726 | 0.374499 | 0.379838 | 0.376023 | 0.361464 | 0.358859 | 0.365427 | 0.396100 | 0.452251 | 0.521358 | 0.552988 | 0.549796 | 0.540851 | 0.579591 | 0.672879 | 0.835470 | 1.046133 | 1.159567 | 1.113838 | 0.949426 | 0.735388 | 0.568019 | 0.466555 | 0.403006 | 0.370257 | 0.359482 | 0.372407 | 0.411574 | 0.450747 | 0.491112 | 0.530427 | 0.536959 | 0.504392 | 0.442459 | 0.385071 | 0.342282 | 0.316030 | 0.313115 | 0.338702 | 0.389137 | 0.466077 | 0.584708 | 0.707643 | 0.819619 | 0.969898 | 1.105094 | 1.105446 | 1.017569 | 0.967647 | 0.894885 | 0.765437 | 0.602061 | 0.459664 | 0.333715 | 0.258866 | 0.229190 | 0.220742 | 0.219626 | 0.224205 | 0.238654 | 0.281348 | 0.357823 | 0.455433 | 0.579633 | 0.692021 | 0.701726 | 0.643261 | 0.561250 | 0.483722 | 0.431552 | 0.404308 | 0.397284 | 0.410233 | 0.421323 | 0.413150 | 0.351798 | 0.286678 | 0.241633 | 0.219207 | 0.215005 | 0.217608 | 0.216486 | 0.214649 | 0.218477 | 0.236187 | 0.261628 | 0.287317 | 0.336184 | 0.388606 | 0.469873 | 0.575427 | 0.669786 | 0.725796 |
| all electrodes | 0.216917 | 0.217690 | 0.215590 | 0.215065 | 0.221088 | 0.225608 | 0.257165 | 0.292691 | 0.378738 | 0.563417 | 0.871058 | 0.928186 | 1.250591 | 1.209163 | 1.018754 | 0.690223 | 0.445132 | 0.276012 | 0.227656 | 0.216604 | 0.215406 | 0.214626 | 0.214769 | 0.216705 | 0.226465 | 0.259021 | 0.392896 | 0.633751 | 0.727039 | 0.792802 | 0.732986 | 0.507425 | 0.383199 | 0.346214 | 0.314067 | 0.327997 | 0.313885 | 0.250757 | 0.216428 | 0.234744 | 0.371989 | 0.690751 | 1.243281 | 1.900635 | 1.653319 | 0.988349 | 0.450315 | 0.266442 | 0.227859 | 0.215842 | 0.214636 | 0.216369 | 0.219985 | 0.227218 | 0.223457 | 0.219858 | 0.215706 | 0.214644 | 0.216139 | 0.221323 | 0.222508 | 0.216058 | 0.224006 | 0.328549 | 0.881834 | 4.549504 | 39.464703 | 287.717799 | 1434.855633 | 4477.786193 | 8530.747566 | 10318.293989 | 6623.379573 | 1719.183261 | 267.927977 | 45.417463 | 7.828606 | 1.638561 | 0.604212 | 0.340366 | 0.266116 | 0.252136 | 0.262980 | 0.281918 | 0.311073 | 0.310933 | 0.288956 | 0.256174 | 0.240892 | 0.234423 | 0.243344 | 0.274037 | 0.370504 | 0.611874 | 1.198021 | 2.125862 | 3.235311 | 3.230005 | 2.333476 | 1.447690 | 0.982715 | 0.710602 | 0.600579 | 0.554567 | 0.598085 | 0.723254 | 0.850545 | 0.872926 | 0.901435 | 0.982695 | 1.198704 | 1.681460 | 2.529052 | 3.855524 | 6.495288 | 7.946226 | 7.835895 | 7.306740 | 5.886338 | 3.972910 | 3.092677 | 2.406945 | 2.004978 | 1.647327 | 1.319903 | 1.181972 | 1.366794 | 1.534856 | 1.769207 | 2.019764 | 1.910351 | 1.470826 | 1.178020 | 0.984918 | 0.895207 | 0.938026 | 0.940910 | 0.942824 | 0.930471 | 0.865227 | 0.707815 | 0.630742 | 0.590045 | 0.575757 | 0.533723 | 0.505587 | 0.464013 | 0.422046 | 0.394703 | 0.399511 | 0.439371 | 0.586905 | 0.934153 | 1.484372 | 2.103847 | 2.484413 | 1.994709 | 1.406896 | 0.963082 | 0.643212 | 0.456015 | 0.351326 | 0.281873 | 0.245377 | 0.227906 | 0.215372 | 0.218858 | 0.241704 | 0.269914 | 0.306786 | 0.310241 | 0.302382 | 0.272425 | 0.260050 | 0.238909 | 0.224022 | 0.214722 | 0.220154 | 0.239765 | 0.263879 | 0.300824 | 0.355539 | 0.380461 | 0.381584 | 0.370477 | 0.368729 | 0.346166 | 0.319255 | 0.309424 | 0.308469 | 0.310609 | 0.337382 | 0.385973 | 0.415559 | 0.425342 | 0.397102 | 0.331268 | 0.299410 | 0.303509 | 0.316922 | 0.319231 | 0.332832 | 0.332379 | 0.299467 | 0.265234 | 0.244987 | 0.232844 | 0.236400 | 0.262069 | 0.318256 | 0.444442 | 0.655313 | 0.819949 | 0.839383 | 0.669136 | 0.489680 | 0.365025 | 0.320936 | 0.306047 | 0.329153 | 0.358693 | 0.393989 | 0.414923 | 0.486731 | 0.611480 | 0.826814 | 1.151238 | 1.626883 | 2.320675 | 3.562065 | 4.890395 | 5.588362 | 4.425808 | 3.316754 | 3.065272 | 3.127770 | 2.822776 | 2.753285 | 2.531770 | 2.387658 | 2.254474 | 2.385602 | 2.730037 | 2.952033 | 1.825544 | 1.413875 | 1.397709 | 1.465558 | 1.686631 | 2.567978 | 3.318952 | 4.760178 | 6.412161 | 7.416434 | 6.759680 | 4.940828 | 1.969380 | 0.919270 | 0.537327 | 0.383442 | 0.318394 | 0.301505 | 0.299225 | 0.314652 | 0.351842 | 0.410482 | 0.483823 | 0.560465 | 0.654685 | 0.762971 | 1.005292 | 1.427322 | 2.028656 | 2.372207 | 2.628463 | 2.258649 | 1.644308 | 0.954947 | 0.806713 | 0.609699 |

Y) neural network features, happy vs angry

  
|  | time window | peak latency | cluster *p* | peak Cohen's *d* |  | | | |  |
| **all electrodes** | 365 - 475 ms | 405 ms | 0.0342 | 0.7293 |  | | | |  |
|  | | | | | | | | |  |

Model correlations, cluster permutation tests

|  | **left hemisphere** | | | | **right hemisphere** | | | |  |
|  | time window | peak latency | cluster *p* | peak Cohen's *d* | time window | peak latency | cluster *p* | peak Cohen's *d* |  |
| **anterior** |  | | | |  | | | |  |
| **central** |  | | | | 385 - 485 ms | 450 ms | 0.0472 | 0.739 |  |
  | | | | 785 - 895 ms | 820 ms | 0.0433 | 0.802 ||  | **posterior** | 130 - 215 ms | 170 ms | 0.0287 | 1.1229 | 860 - 1065 ms | 895 ms | 0.0212 | 0.5435 |
 265 - 365 ms | 340 ms | 0.0493 | 0.5903 |  | | | | 965 - 1075 ms | 990 ms | 0.0437 | 0.4957 |  | | | |

  

Model correlations, Bayesian statistics

|  | -200 | -195 | -190 | -185 | -180 | -175 | -170 | -165 | -160 | -155 | -150 | -145 | -140 | -135 | -130 | -125 | -120 | -115 | -110 | -105 | -100 | -95 | -90 | -85 | -80 | -75 | -70 | -65 | -60 | -55 | -50 | -45 | -40 | -35 | -30 | -25 | -20 | -15 | -10 | -5 | 0 | 5 | 10 | 15 | 20 | 25 | 30 | 35 | 40 | 45 | 50 | 55 | 60 | 65 | 70 | 75 | 80 | 85 | 90 | 95 | 100 | 105 | 110 | 115 | 120 | 125 | 130 | 135 | 140 | 145 | 150 | 155 | 160 | 165 | 170 | 175 | 180 | 185 | 190 | 195 | 200 | 205 | 210 | 215 | 220 | 225 | 230 | 235 | 240 | 245 | 250 | 255 | 260 | 265 | 270 | 275 | 280 | 285 | 290 | 295 | 300 | 305 | 310 | 315 | 320 | 325 | 330 | 335 | 340 | 345 | 350 | 355 | 360 | 365 | 370 | 375 | 380 | 385 | 390 | 395 | 400 | 405 | 410 | 415 | 420 | 425 | 430 | 435 | 440 | 445 | 450 | 455 | 460 | 465 | 470 | 475 | 480 | 485 | 490 | 495 | 500 | 505 | 510 | 515 | 520 | 525 | 530 | 535 | 540 | 545 | 550 | 555 | 560 | 565 | 570 | 575 | 580 | 585 | 590 | 595 | 600 | 605 | 610 | 615 | 620 | 625 | 630 | 635 | 640 | 645 | 650 | 655 | 660 | 665 | 670 | 675 | 680 | 685 | 690 | 695 | 700 | 705 | 710 | 715 | 720 | 725 | 730 | 735 | 740 | 745 | 750 | 755 | 760 | 765 | 770 | 775 | 780 | 785 | 790 | 795 | 800 | 805 | 810 | 815 | 820 | 825 | 830 | 835 | 840 | 845 | 850 | 855 | 860 | 865 | 870 | 875 | 880 | 885 | 890 | 895 | 900 | 905 | 910 | 915 | 920 | 925 | 930 | 935 | 940 | 945 | 950 | 955 | 960 | 965 | 970 | 975 | 980 | 985 | 990 | 995 | 1000 | 1005 | 1010 | 1015 | 1020 | 1025 | 1030 | 1035 | 1040 | 1045 | 1050 | 1055 | 1060 | 1065 | 1070 | 1075 | 1080 | 1085 | 1090 | 1095 | 1100 | 1105 | 1110 | 1115 | 1120 | 1125 | 1130 | 1135 | 1140 | 1145 | 1150 | 1155 | 1160 | 1165 | 1170 | 1175 | 1180 | 1185 | 1190 | 1195 |
| --- | --- | --- | --- | --- | --- | --- | --- | --- | --- | --- | --- | --- | --- | --- | --- | --- | --- | --- | --- | --- | --- | --- | --- | --- | --- | --- | --- | --- | --- | --- | --- | --- | --- | --- | --- | --- | --- | --- | --- | --- | --- | --- | --- | --- | --- | --- | --- | --- | --- | --- | --- | --- | --- | --- | --- | --- | --- | --- | --- | --- | --- | --- | --- | --- | --- | --- | --- | --- | --- | --- | --- | --- | --- | --- | --- | --- | --- | --- | --- | --- | --- | --- | --- | --- | --- | --- | --- | --- | --- | --- | --- | --- | --- | --- | --- | --- | --- | --- | --- | --- | --- | --- | --- | --- | --- | --- | --- | --- | --- | --- | --- | --- | --- | --- | --- | --- | --- | --- | --- | --- | --- | --- | --- | --- | --- | --- | --- | --- | --- | --- | --- | --- | --- | --- | --- | --- | --- | --- | --- | --- | --- | --- | --- | --- | --- | --- | --- | --- | --- | --- | --- | --- | --- | --- | --- | --- | --- | --- | --- | --- | --- | --- | --- | --- | --- | --- | --- | --- | --- | --- | --- | --- | --- | --- | --- | --- | --- | --- | --- | --- | --- | --- | --- | --- | --- | --- | --- | --- | --- | --- | --- | --- | --- | --- | --- | --- | --- | --- | --- | --- | --- | --- | --- | --- | --- | --- | --- | --- | --- | --- | --- | --- | --- | --- | --- | --- | --- | --- | --- | --- | --- | --- | --- | --- | --- | --- | --- | --- | --- | --- | --- | --- | --- | --- | --- | --- | --- | --- | --- | --- | --- | --- | --- | --- | --- | --- | --- | --- | --- | --- | --- | --- | --- | --- | --- | --- | --- | --- | --- | --- | --- | --- | --- | --- | --- | --- | --- | --- | --- | --- | --- | --- | --- | --- | --- | --- | --- | --- | --- | --- |
| left anterior | 0.441417 | 0.593081 | 0.730125 | 1.438108 | 1.913822 | 1.577886 | 0.757621 | 0.410216 | 0.286267 | 0.244823 | 0.225191 | 0.249572 | 0.363269 | 0.849558 | 1.876596 | 1.858837 | 1.435479 | 1.249498 | 0.966464 | 0.851843 | 0.715720 | 0.542991 | 0.393894 | 0.281758 | 0.218011 | 0.221163 | 0.254188 | 0.296093 | 0.310753 | 0.284130 | 0.239914 | 0.215111 | 0.233352 | 0.330287 | 0.672452 | 1.822789 | 4.777391 | 4.584883 | 2.123060 | 0.740370 | 0.357634 | 0.246856 | 0.226794 | 0.220991 | 0.236583 | 0.292062 | 0.428518 | 0.432198 | 0.460270 | 0.414606 | 0.429291 | 0.542883 | 1.077891 | 1.983751 | 3.510530 | 4.326895 | 3.491862 | 1.784949 | 1.025335 | 0.611504 | 0.380177 | 0.287968 | 0.243596 | 0.220449 | 0.214851 | 0.221954 | 0.236951 | 0.255442 | 0.304026 | 0.342475 | 0.387132 | 0.428925 | 0.504462 | 0.515217 | 0.468048 | 0.395534 | 0.322035 | 0.262327 | 0.223288 | 0.219496 | 0.318965 | 0.745245 | 1.744615 | 2.579950 | 2.248755 | 1.595527 | 1.121426 | 0.702349 | 0.506801 | 0.435404 | 0.437174 | 0.537461 | 1.009147 | 2.376727 | 6.866930 | 18.318237 | 46.401532 | 81.539042 | 89.701858 | 67.215129 | 40.036534 | 18.859083 | 10.918355 | 6.798971 | 4.215395 | 2.575994 | 1.618354 | 0.793815 | 0.472538 | 0.358662 | 0.304016 | 0.269804 | 0.247170 | 0.228898 | 0.219458 | 0.215425 | 0.214630 | 0.216001 | 0.216695 | 0.217237 | 0.215878 | 0.214773 | 0.216978 | 0.228791 | 0.259894 | 0.313381 | 0.399060 | 0.471917 | 0.485159 | 0.427591 | 0.344497 | 0.282012 | 0.248019 | 0.228819 | 0.218971 | 0.215786 | 0.214995 | 0.214968 | 0.217160 | 0.225820 | 0.252720 | 0.315830 | 0.459487 | 0.651650 | 0.822847 | 0.639257 | 0.424493 | 0.302653 | 0.248600 | 0.225767 | 0.219767 | 0.219751 | 0.229523 | 0.254041 | 0.298626 | 0.416496 | 0.638273 | 0.808213 | 1.191359 | 2.031210 | 2.943359 | 4.644625 | 6.850205 | 7.913458 | 8.602632 | 7.583341 | 4.656063 | 2.696259 | 1.406900 | 0.741488 | 0.455666 | 0.348832 | 0.301129 | 0.299729 | 0.322966 | 0.375056 | 0.428795 | 0.453776 | 0.421986 | 0.375771 | 0.318790 | 0.287570 | 0.274274 | 0.286847 | 0.327446 | 0.427600 | 0.623768 | 0.843600 | 0.763667 | 0.661698 | 0.557859 | 0.468442 | 0.389641 | 0.338517 | 0.281910 | 0.244580 | 0.218589 | 0.215265 | 0.220648 | 0.232488 | 0.255237 | 0.278024 | 0.301156 | 0.317141 | 0.332529 | 0.339249 | 0.342039 | 0.359026 | 0.413442 | 0.447331 | 0.478789 | 0.490679 | 0.550565 | 0.711770 | 0.965347 | 1.265212 | 1.747662 | 2.332265 | 2.827924 | 2.962453 | 2.894740 | 2.545153 | 2.341654 | 2.056600 | 1.630746 | 1.291979 | 1.044543 | 0.805050 | 0.655995 | 0.585814 | 0.518634 | 0.477499 | 0.433870 | 0.390358 | 0.373183 | 0.361931 | 0.314625 | 0.281269 | 0.251513 | 0.225732 | 0.216081 | 0.214660 | 0.216400 | 0.219060 | 0.217287 | 0.215078 | 0.215201 | 0.219933 | 0.224508 | 0.227304 | 0.228388 | 0.229809 | 0.239218 | 0.263035 | 0.300785 | 0.353923 | 0.430867 | 0.512712 | 0.550464 | 0.528368 | 0.509043 | 0.492208 | 0.517687 | 0.577163 | 0.657471 | 0.759216 | 0.866119 | 0.899494 | 0.899315 | 0.950547 | 0.980351 | 0.962852 | 0.903302 | 0.814611 | 0.766993 | 0.702707 | 0.630058 | 0.593048 | 0.587286 | 0.546606 |
| right anterior | 0.466804 | 0.367767 | 0.306841 | 0.240303 | 0.214628 | 0.245033 | 0.344859 | 0.543347 | 0.904477 | 1.224731 | 1.283629 | 1.136893 | 1.068230 | 0.905693 | 0.632750 | 0.428991 | 0.305714 | 0.234468 | 0.215288 | 0.232932 | 0.252307 | 0.273455 | 0.356562 | 0.460399 | 0.584436 | 0.639258 | 0.616084 | 0.456304 | 0.287299 | 0.218232 | 0.239135 | 0.432901 | 0.914972 | 1.675689 | 1.988251 | 1.446995 | 0.852401 | 0.562430 | 0.371038 | 0.313447 | 0.309337 | 0.336535 | 0.370841 | 0.445871 | 0.644489 | 1.118799 | 1.196438 | 0.976483 | 0.570251 | 0.371614 | 0.315514 | 0.308722 | 0.318501 | 0.366615 | 0.376351 | 0.397606 | 0.415767 | 0.484054 | 0.470184 | 0.446708 | 0.351185 | 0.297939 | 0.234787 | 0.215522 | 0.218566 | 0.224889 | 0.236578 | 0.235883 | 0.235834 | 0.248221 | 0.305611 | 0.366416 | 0.421599 | 0.483769 | 0.505910 | 0.491983 | 0.448173 | 0.367391 | 0.273516 | 0.219537 | 0.236836 | 0.345566 | 0.554744 | 0.780480 | 0.975259 | 0.868501 | 0.640162 | 0.502820 | 0.358655 | 0.260872 | 0.223030 | 0.215469 | 0.235264 | 0.272655 | 0.317323 | 0.344074 | 0.307931 | 0.279478 | 0.250471 | 0.231405 | 0.218626 | 0.214692 | 0.217839 | 0.221274 | 0.227014 | 0.226434 | 0.221430 | 0.214970 | 0.215639 | 0.223055 | 0.238396 | 0.270518 | 0.306100 | 0.372001 | 0.491389 | 0.716206 | 1.090590 | 1.820817 | 2.776718 | 4.459060 | 6.896945 | 9.321332 | 11.514178 | 12.090318 | 7.304853 | 2.984579 | 1.013578 | 0.467997 | 0.278085 | 0.225973 | 0.215806 | 0.233024 | 0.265477 | 0.271985 | 0.264491 | 0.245094 | 0.242624 | 0.256995 | 0.292803 | 0.355638 | 0.367593 | 0.336338 | 0.248588 | 0.219330 | 0.354506 | 0.776388 | 1.764269 | 3.227387 | 3.869699 | 4.118103 | 2.962667 | 2.027148 | 1.261149 | 0.666187 | 0.347570 | 0.251109 | 0.214932 | 0.232930 | 0.295008 | 0.382017 | 0.473727 | 0.538799 | 0.586417 | 0.596244 | 0.665800 | 0.694055 | 0.611027 | 0.499374 | 0.382495 | 0.285422 | 0.239522 | 0.218732 | 0.214630 | 0.214886 | 0.214973 | 0.222066 | 0.238932 | 0.268012 | 0.290289 | 0.279528 | 0.266490 | 0.284803 | 0.374841 | 0.700240 | 1.567193 | 2.686720 | 3.235533 | 2.559547 | 1.763278 | 1.299765 | 1.213181 | 1.356428 | 1.572619 | 1.868343 | 2.036158 | 1.736177 | 1.253646 | 0.846416 | 0.685974 | 0.680262 | 0.887938 | 1.299667 | 2.284488 | 2.424308 | 1.798951 | 0.930105 | 0.495572 | 0.313758 | 0.259271 | 0.236653 | 0.236326 | 0.240773 | 0.251374 | 0.259396 | 0.260981 | 0.256737 | 0.259108 | 0.252288 | 0.245007 | 0.238255 | 0.227028 | 0.215964 | 0.215613 | 0.229444 | 0.250623 | 0.266438 | 0.262962 | 0.258493 | 0.252110 | 0.260282 | 0.275803 | 0.308396 | 0.377824 | 0.481238 | 0.529822 | 0.495150 | 0.390978 | 0.279369 | 0.222366 | 0.216341 | 0.229372 | 0.233890 | 0.225830 | 0.217872 | 0.214842 | 0.224474 | 0.244944 | 0.258820 | 0.267560 | 0.256482 | 0.243454 | 0.238044 | 0.238291 | 0.242793 | 0.269791 | 0.325560 | 0.408052 | 0.532419 | 0.664947 | 0.749963 | 0.778615 | 0.757013 | 0.687507 | 0.635695 | 0.596688 | 0.555481 | 0.472636 | 0.371060 | 0.288299 | 0.236275 | 0.216526 | 0.214902 | 0.215753 | 0.215914 | 0.216748 | 0.219302 | 0.220753 | 0.222102 | 0.224263 | 0.223807 |
| left central | 0.554195 | 0.513542 | 0.529416 | 0.497117 | 0.419410 | 0.358088 | 0.365152 | 0.368682 | 0.369365 | 0.377234 | 0.415209 | 0.433533 | 0.501087 | 0.650345 | 1.015977 | 1.634695 | 2.409683 | 2.581889 | 2.143090 | 1.570865 | 1.189817 | 0.820921 | 0.589064 | 0.518808 | 0.450820 | 0.382275 | 0.355458 | 0.385993 | 0.401943 | 0.432951 | 0.565258 | 0.711510 | 0.690717 | 0.454387 | 0.314937 | 0.250814 | 0.226617 | 0.217345 | 0.215375 | 0.214854 | 0.215061 | 0.214630 | 0.215365 | 0.218415 | 0.227383 | 0.231815 | 0.228962 | 0.218863 | 0.214670 | 0.216493 | 0.222150 | 0.231113 | 0.238873 | 0.233194 | 0.226160 | 0.219477 | 0.217232 | 0.214643 | 0.216745 | 0.225671 | 0.234845 | 0.248727 | 0.262620 | 0.274936 | 0.263335 | 0.240513 | 0.214884 | 0.246819 | 0.451747 | 1.054502 | 1.966282 | 2.471344 | 2.828481 | 2.803892 | 2.399092 | 1.736062 | 1.129615 | 0.710533 | 0.447223 | 0.271302 | 0.214627 | 0.302749 | 0.693909 | 1.750876 | 3.704004 | 6.072567 | 6.423807 | 5.076651 | 2.939021 | 1.518344 | 0.776251 | 0.451923 | 0.340499 | 0.301522 | 0.281020 | 0.286777 | 0.343807 | 0.444515 | 0.650273 | 0.981768 | 1.363105 | 1.457815 | 1.230689 | 0.900966 | 0.616762 | 0.417268 | 0.303476 | 0.246979 | 0.222960 | 0.216270 | 0.214673 | 0.216757 | 0.220276 | 0.227713 | 0.242490 | 0.267938 | 0.302866 | 0.351632 | 0.397271 | 0.435466 | 0.451409 | 0.441770 | 0.393465 | 0.349049 | 0.309522 | 0.280626 | 0.266658 | 0.270045 | 0.278438 | 0.295382 | 0.317632 | 0.338519 | 0.327793 | 0.302925 | 0.263771 | 0.229563 | 0.214803 | 0.222342 | 0.259573 | 0.330061 | 0.480152 | 0.792455 | 1.322936 | 1.952635 | 3.284303 | 4.974352 | 5.417436 | 4.237076 | 2.393109 | 1.187276 | 0.679234 | 0.462676 | 0.378275 | 0.376924 | 0.389634 | 0.426761 | 0.471560 | 0.539066 | 0.526842 | 0.470265 | 0.389955 | 0.378416 | 0.383073 | 0.477267 | 0.694860 | 1.041971 | 1.217411 | 1.087779 | 0.745708 | 0.582864 | 0.473153 | 0.408881 | 0.357146 | 0.328446 | 0.298340 | 0.272432 | 0.242844 | 0.228388 | 0.217243 | 0.214901 | 0.219344 | 0.221529 | 0.222957 | 0.219261 | 0.216060 | 0.214801 | 0.217780 | 0.227109 | 0.231107 | 0.237420 | 0.238825 | 0.236054 | 0.222347 | 0.215518 | 0.220066 | 0.248292 | 0.299281 | 0.350830 | 0.377246 | 0.388107 | 0.415076 | 0.435272 | 0.413219 | 0.371376 | 0.322069 | 0.283229 | 0.267508 | 0.266716 | 0.283664 | 0.327423 | 0.399803 | 0.476809 | 0.553419 | 0.600258 | 0.627575 | 0.654537 | 0.711323 | 0.827417 | 1.031248 | 1.345214 | 1.774126 | 2.212920 | 2.158535 | 1.646337 | 1.160896 | 0.908898 | 0.893417 | 1.171242 | 2.178562 | 5.554706 | 12.624349 | 16.401456 | 13.389167 | 7.595494 | 3.479164 | 1.600642 | 0.808666 | 0.464785 | 0.336868 | 0.282796 | 0.260588 | 0.245122 | 0.235153 | 0.227490 | 0.216972 | 0.216346 | 0.228713 | 0.243666 | 0.256082 | 0.268597 | 0.277558 | 0.267476 | 0.250994 | 0.237047 | 0.222654 | 0.214946 | 0.217182 | 0.228039 | 0.248855 | 0.279506 | 0.308973 | 0.326046 | 0.328007 | 0.324491 | 0.308769 | 0.288897 | 0.270741 | 0.245410 | 0.223949 | 0.214929 | 0.217831 | 0.237239 | 0.271240 | 0.323176 | 0.378951 | 0.429945 | 0.456940 | 0.461897 | 0.448378 | 0.438698 |
| right central | 0.246500 | 0.272022 | 0.284817 | 0.335189 | 0.391810 | 0.417434 | 0.401573 | 0.389153 | 0.341438 | 0.327449 | 0.285052 | 0.238518 | 0.217431 | 0.214720 | 0.219699 | 0.221304 | 0.230125 | 0.239137 | 0.241746 | 0.241225 | 0.246137 | 0.243521 | 0.269904 | 0.345173 | 0.468950 | 0.572505 | 0.529700 | 0.465530 | 0.433686 | 0.427006 | 0.440864 | 0.591733 | 0.835095 | 1.154872 | 1.105384 | 0.573570 | 0.282672 | 0.214626 | 0.308892 | 0.724298 | 1.827775 | 4.220292 | 6.700846 | 3.747827 | 1.317413 | 0.411120 | 0.239612 | 0.214945 | 0.214717 | 0.214700 | 0.220146 | 0.244221 | 0.296521 | 0.385156 | 0.658169 | 1.251097 | 3.034671 | 5.058555 | 2.930940 | 1.062805 | 0.505288 | 0.294682 | 0.231978 | 0.214682 | 0.223163 | 0.235288 | 0.250839 | 0.303172 | 0.446647 | 1.050509 | 3.603159 | 10.774741 | 18.816888 | 19.308307 | 10.303627 | 4.211883 | 1.743271 | 0.771879 | 0.399281 | 0.270687 | 0.228746 | 0.220969 | 0.227678 | 0.243902 | 0.272360 | 0.336189 | 0.382708 | 0.412375 | 0.517237 | 0.764683 | 1.289517 | 2.452135 | 4.729298 | 7.548681 | 10.147698 | 8.611640 | 6.185981 | 3.265492 | 1.439134 | 0.678194 | 0.437972 | 0.363024 | 0.379575 | 0.488337 | 0.663634 | 1.000560 | 1.432014 | 1.528862 | 1.162504 | 0.769964 | 0.534657 | 0.421826 | 0.385072 | 0.401280 | 0.461513 | 0.598911 | 0.872649 | 1.315833 | 2.004511 | 2.831305 | 3.263394 | 3.344393 | 3.222862 | 3.221784 | 3.826259 | 4.988028 | 7.355778 | 12.076839 | 17.892074 | 23.385204 | 29.942337 | 25.670710 | 16.836167 | 10.214628 | 6.272105 | 3.779114 | 2.390252 | 1.415098 | 0.943247 | 0.710293 | 0.615219 | 0.611570 | 0.697202 | 0.755805 | 0.763703 | 0.707773 | 0.608223 | 0.511916 | 0.527572 | 0.555548 | 0.589302 | 0.623880 | 0.664132 | 0.639234 | 0.630309 | 0.666924 | 0.744158 | 0.844425 | 1.143369 | 1.716961 | 2.568120 | 3.374416 | 3.571882 | 2.627996 | 1.703729 | 1.037004 | 0.739146 | 0.572955 | 0.530953 | 0.576565 | 0.682201 | 0.794345 | 0.984097 | 1.106374 | 1.146062 | 1.126834 | 1.141841 | 1.283297 | 1.457020 | 1.538285 | 1.628093 | 1.662082 | 1.501933 | 1.191995 | 0.899097 | 0.673266 | 0.510522 | 0.392708 | 0.347020 | 0.332264 | 0.334430 | 0.367424 | 0.437937 | 0.580215 | 0.861661 | 1.221323 | 1.333949 | 1.250338 | 1.200238 | 1.324017 | 1.833147 | 3.597482 | 10.706851 | 37.106626 | 83.286972 | 86.914482 | 42.777680 | 12.583526 | 3.197950 | 1.275375 | 0.851920 | 0.886411 | 1.211282 | 2.013019 | 2.943628 | 3.775321 | 3.597048 | 2.891421 | 1.973862 | 1.362668 | 0.935005 | 0.819407 | 0.768981 | 0.827515 | 0.952290 | 1.064034 | 0.965243 | 0.881132 | 0.798783 | 0.726454 | 0.631749 | 0.552484 | 0.502816 | 0.518726 | 0.574964 | 0.659226 | 0.726175 | 0.752052 | 0.719728 | 0.576587 | 0.435398 | 0.332815 | 0.265127 | 0.227554 | 0.218792 | 0.217814 | 0.225040 | 0.246674 | 0.293192 | 0.367248 | 0.473194 | 0.587969 | 0.702686 | 0.715558 | 0.618811 | 0.484033 | 0.342719 | 0.273809 | 0.236759 | 0.218088 | 0.214627 | 0.214636 | 0.214743 | 0.215227 | 0.226710 | 0.261032 | 0.304438 | 0.328476 | 0.335955 | 0.328628 | 0.302179 | 0.267575 | 0.241925 | 0.223190 | 0.215547 | 0.215455 | 0.217304 | 0.215698 | 0.214784 | 0.214646 |
| left posterior | 0.449324 | 0.528718 | 0.629143 | 1.055777 | 1.374095 | 1.268309 | 0.843702 | 0.614346 | 0.416514 | 0.290146 | 0.246570 | 0.231714 | 0.237786 | 0.273983 | 0.504291 | 3.149894 | 112.888303 | 3745.516260 | 24223.759507 | 9328.520747 | 345.752476 | 14.720086 | 1.795852 | 0.399254 | 0.219424 | 0.225062 | 0.260422 | 0.283304 | 0.297782 | 0.243349 | 0.219264 | 0.215909 | 0.224258 | 0.228996 | 0.234415 | 0.233638 | 0.234784 | 0.222705 | 0.217957 | 0.216379 | 0.215904 | 0.216375 | 0.228377 | 0.259576 | 0.316157 | 0.411841 | 0.518376 | 0.606501 | 0.670924 | 0.737515 | 0.690033 | 0.602406 | 0.512466 | 0.460801 | 0.388874 | 0.315107 | 0.285113 | 0.287627 | 0.283382 | 0.279922 | 0.287635 | 0.290101 | 0.306426 | 0.352577 | 0.446386 | 0.719059 | 1.500697 | 3.756814 | 10.776539 | 32.291491 | 82.919130 | 199.586608 | 470.144184 | 1021.697052 | 1859.769762 | 2385.411480 | 1842.762626 | 1099.630047 | 455.125637 | 148.464255 | 50.705542 | 17.874125 | 4.963072 | 1.358796 | 0.539043 | 0.348143 | 0.281330 | 0.248568 | 0.247326 | 0.268586 | 0.315413 | 0.456362 | 0.949327 | 2.479051 | 7.590590 | 17.228014 | 22.209021 | 17.170516 | 10.950829 | 7.611332 | 5.300593 | 3.379679 | 2.465898 | 2.366691 | 2.592014 | 3.515612 | 4.570823 | 6.163329 | 6.916640 | 5.686606 | 3.693851 | 2.308069 | 1.433912 | 1.078025 | 0.958088 | 0.963258 | 1.132301 | 1.555947 | 2.221636 | 3.717457 | 6.946954 | 11.270284 | 12.585628 | 11.075172 | 7.158216 | 3.774685 | 1.945037 | 1.103124 | 0.699751 | 0.514652 | 0.429297 | 0.388798 | 0.379759 | 0.389677 | 0.406099 | 0.423291 | 0.468458 | 0.541653 | 0.594481 | 0.658847 | 0.754640 | 0.851644 | 0.954301 | 1.133166 | 1.242744 | 1.306020 | 1.083807 | 0.861419 | 0.742481 | 0.684334 | 0.649225 | 0.673828 | 0.673402 | 0.602156 | 0.503487 | 0.389370 | 0.313005 | 0.258277 | 0.227944 | 0.214844 | 0.221281 | 0.243225 | 0.261799 | 0.264317 | 0.254084 | 0.242987 | 0.222919 | 0.215707 | 0.247003 | 0.337151 | 0.545054 | 0.871519 | 1.183423 | 1.427798 | 1.530487 | 1.434838 | 1.297950 | 1.073685 | 0.844165 | 0.762150 | 0.782245 | 0.831722 | 0.955474 | 1.101918 | 1.151721 | 1.055212 | 0.884887 | 0.716123 | 0.590368 | 0.504982 | 0.461023 | 0.474810 | 0.553797 | 0.717313 | 1.078474 | 1.830758 | 2.973939 | 4.325127 | 5.453775 | 5.141365 | 3.983603 | 2.711351 | 1.807448 | 1.264419 | 0.999665 | 0.816037 | 0.745755 | 0.751190 | 0.850953 | 1.072621 | 1.730986 | 3.321320 | 6.795925 | 14.433781 | 29.637882 | 49.397093 | 71.957513 | 98.220296 | 86.262604 | 53.776424 | 26.994219 | 11.371345 | 4.527991 | 2.056929 | 1.143523 | 0.828105 | 0.706342 | 0.737046 | 0.875391 | 1.099418 | 1.157866 | 1.198886 | 1.137088 | 1.099144 | 1.146481 | 1.301261 | 1.499377 | 1.926871 | 2.345016 | 2.713500 | 3.293076 | 4.091353 | 4.785862 | 6.238654 | 8.012804 | 11.389538 | 14.926261 | 17.729605 | 16.276738 | 13.560441 | 8.769127 | 5.374177 | 3.451937 | 2.431407 | 1.725271 | 1.417594 | 1.183596 | 1.017182 | 0.872361 | 0.741209 | 0.586887 | 0.463824 | 0.382545 | 0.340123 | 0.317376 | 0.306513 | 0.310052 | 0.315623 | 0.324324 | 0.328184 | 0.322734 | 0.308751 | 0.285929 | 0.259467 | 0.241949 | 0.231825 | 0.225289 | 0.222533 | 0.220577 | 0.219527 |
| right posterior | 0.372702 | 0.338915 | 0.305172 | 0.276888 | 0.270872 | 0.297330 | 0.347250 | 0.504162 | 0.821406 | 1.141571 | 1.160888 | 0.805646 | 0.448704 | 0.333592 | 0.303492 | 0.291132 | 0.322715 | 0.417665 | 0.555753 | 0.573145 | 0.524667 | 0.373646 | 0.271051 | 0.222878 | 0.215240 | 0.216889 | 0.239264 | 0.287657 | 0.392032 | 0.661408 | 1.310542 | 1.630903 | 0.879602 | 0.398567 | 0.243750 | 0.214627 | 0.237528 | 0.270236 | 0.303466 | 0.295104 | 0.252690 | 0.215631 | 0.230820 | 0.316038 | 0.462900 | 0.624844 | 0.751575 | 0.978561 | 1.114097 | 1.127854 | 0.877934 | 0.574056 | 0.359271 | 0.258809 | 0.226785 | 0.218445 | 0.219003 | 0.228501 | 0.247417 | 0.277969 | 0.320818 | 0.324828 | 0.283265 | 0.249648 | 0.227196 | 0.214894 | 0.224272 | 0.288816 | 0.516950 | 1.283317 | 2.946304 | 5.909478 | 8.998516 | 9.388832 | 8.636012 | 6.987931 | 4.495364 | 2.612345 | 1.311353 | 0.583372 | 0.302200 | 0.224149 | 0.215376 | 0.223836 | 0.231580 | 0.230952 | 0.227920 | 0.223795 | 0.219969 | 0.216150 | 0.214811 | 0.214646 | 0.215049 | 0.217595 | 0.223076 | 0.231692 | 0.237054 | 0.233058 | 0.224848 | 0.219495 | 0.214734 | 0.220039 | 0.238919 | 0.281696 | 0.344816 | 0.401951 | 0.461645 | 0.507576 | 0.501718 | 0.500228 | 0.473924 | 0.419614 | 0.382432 | 0.363910 | 0.357044 | 0.378657 | 0.437597 | 0.561885 | 0.808320 | 1.264630 | 1.920693 | 2.941905 | 3.927741 | 4.092916 | 3.110356 | 2.187283 | 1.411597 | 0.993799 | 0.765423 | 0.647484 | 0.579175 | 0.560783 | 0.538337 | 0.527238 | 0.521253 | 0.524509 | 0.511489 | 0.505690 | 0.538156 | 0.651175 | 0.855052 | 1.181552 | 1.768043 | 2.464267 | 2.615340 | 2.263662 | 1.512595 | 0.933573 | 0.594486 | 0.437540 | 0.342884 | 0.300182 | 0.268674 | 0.255095 | 0.251507 | 0.268307 | 0.300562 | 0.370655 | 0.509207 | 0.711285 | 0.889249 | 1.105313 | 1.261376 | 1.311698 | 1.356685 | 1.417091 | 1.409539 | 1.474510 | 1.309108 | 1.060502 | 0.835479 | 0.689375 | 0.572139 | 0.500193 | 0.451804 | 0.440815 | 0.425440 | 0.415686 | 0.405564 | 0.398433 | 0.383527 | 0.391004 | 0.422447 | 0.511705 | 0.698011 | 1.103745 | 1.742612 | 2.434583 | 2.614056 | 2.288180 | 1.642401 | 1.176388 | 0.940569 | 0.864776 | 0.878510 | 1.006940 | 1.201915 | 1.396808 | 1.523189 | 1.556531 | 1.463681 | 1.352558 | 1.219697 | 1.161348 | 1.164813 | 1.178013 | 1.184962 | 1.246158 | 1.331347 | 1.303401 | 1.188495 | 1.093565 | 1.128424 | 1.227607 | 1.448601 | 2.011001 | 2.726784 | 3.256950 | 3.539385 | 3.711094 | 3.599401 | 3.498045 | 3.257467 | 2.873437 | 2.589270 | 2.369401 | 2.218372 | 2.167702 | 2.204565 | 2.383470 | 2.871744 | 3.583013 | 4.416625 | 5.950058 | 8.062867 | 9.693829 | 11.151802 | 12.469537 | 12.527633 | 11.788995 | 9.921209 | 8.200201 | 6.410288 | 4.923036 | 3.857704 | 3.111222 | 2.661828 | 2.481909 | 2.354394 | 2.316510 | 2.296738 | 2.166513 | 1.846155 | 1.424288 | 1.031083 | 0.781676 | 0.651728 | 0.601459 | 0.614193 | 0.678149 | 0.768557 | 0.863735 | 0.964562 | 1.033430 | 1.085739 | 1.046694 | 0.975981 | 0.863000 | 0.697444 | 0.499603 | 0.368115 | 0.279533 | 0.234964 | 0.219424 | 0.216106 | 0.215905 | 0.221694 | 0.238811 | 0.253494 | 0.266315 |
| all electrodes | 0.924641 | 2.022937 | 4.035714 | 13.179207 | 21.509433 | 13.224608 | 4.160156 | 1.883981 | 0.970112 | 0.544713 | 0.369770 | 0.311932 | 0.290141 | 0.323214 | 0.363625 | 0.403810 | 0.485460 | 0.613112 | 0.691530 | 1.347509 | 2.525973 | 2.518320 | 0.900122 | 0.356128 | 0.226338 | 0.214627 | 0.220125 | 0.215026 | 0.217890 | 0.235713 | 0.255727 | 0.300204 | 0.293147 | 0.269973 | 0.232847 | 0.219334 | 0.214821 | 0.218663 | 0.225326 | 0.220969 | 0.214726 | 0.243898 | 0.390254 | 1.136064 | 3.722726 | 6.545217 | 6.514788 | 4.216678 | 2.037997 | 1.227934 | 0.638864 | 0.437643 | 0.397385 | 0.449767 | 0.517866 | 0.528017 | 0.394616 | 0.270719 | 0.216713 | 0.224620 | 0.251355 | 0.278031 | 0.267029 | 0.238556 | 0.219164 | 0.216104 | 0.224928 | 0.217655 | 0.220044 | 0.292356 | 0.612579 | 1.947117 | 7.960324 | 28.835429 | 62.274412 | 66.955590 | 50.134325 | 29.740891 | 14.827944 | 7.698700 | 3.482736 | 1.132496 | 0.413656 | 0.231840 | 0.231069 | 0.326865 | 0.485080 | 0.624409 | 0.665788 | 0.788603 | 0.831903 | 0.799623 | 0.733131 | 0.659373 | 0.494499 | 0.377259 | 0.301779 | 0.266054 | 0.245842 | 0.244264 | 0.244090 | 0.247895 | 0.256381 | 0.253323 | 0.241900 | 0.228598 | 0.217927 | 0.215677 | 0.230171 | 0.271560 | 0.339480 | 0.485572 | 0.818115 | 1.339187 | 2.149285 | 3.365528 | 5.271009 | 7.509726 | 12.377114 | 15.814480 | 20.483822 | 21.910474 | 21.957134 | 18.622589 | 17.126494 | 13.294631 | 10.151867 | 7.310345 | 5.235907 | 3.827524 | 3.320707 | 3.748555 | 4.469489 | 4.146032 | 3.254045 | 1.980088 | 1.060415 | 0.580219 | 0.378246 | 0.295688 | 0.262826 | 0.244243 | 0.235234 | 0.235938 | 0.246612 | 0.255255 | 0.265328 | 0.276736 | 0.288311 | 0.279105 | 0.257655 | 0.228340 | 0.215491 | 0.220640 | 0.250914 | 0.314776 | 0.358144 | 0.391318 | 0.374317 | 0.353847 | 0.334466 | 0.332941 | 0.317979 | 0.311854 | 0.314729 | 0.308233 | 0.303869 | 0.319480 | 0.352536 | 0.372211 | 0.376936 | 0.365987 | 0.347944 | 0.310436 | 0.278257 | 0.252163 | 0.234257 | 0.220866 | 0.215328 | 0.215435 | 0.220422 | 0.225687 | 0.226961 | 0.222671 | 0.215766 | 0.216397 | 0.230976 | 0.273163 | 0.345966 | 0.419350 | 0.482319 | 0.480679 | 0.393483 | 0.320826 | 0.267190 | 0.233488 | 0.218875 | 0.214932 | 0.215399 | 0.219986 | 0.233897 | 0.260419 | 0.285202 | 0.309071 | 0.307180 | 0.293071 | 0.279378 | 0.286861 | 0.305157 | 0.353415 | 0.395495 | 0.474027 | 0.584093 | 0.713673 | 0.851538 | 0.891698 | 0.757456 | 0.576919 | 0.427721 | 0.322569 | 0.266085 | 0.229081 | 0.219257 | 0.219423 | 0.231844 | 0.266880 | 0.366653 | 0.450581 | 0.501712 | 0.497791 | 0.475326 | 0.408415 | 0.366834 | 0.330199 | 0.310147 | 0.315254 | 0.347832 | 0.397630 | 0.483499 | 0.576985 | 0.646886 | 0.660170 | 0.609426 | 0.540727 | 0.504431 | 0.484661 | 0.485256 | 0.501150 | 0.519546 | 0.548637 | 0.522160 | 0.485976 | 0.424462 | 0.350939 | 0.275243 | 0.233932 | 0.215452 | 0.220515 | 0.246417 | 0.278866 | 0.305135 | 0.287668 | 0.261061 | 0.245829 | 0.244292 | 0.249277 | 0.274822 | 0.332944 | 0.441298 | 0.583035 | 0.710013 | 0.902750 | 1.043222 | 1.079459 | 1.025542 | 0.994458 | 0.875450 | 0.718434 | 0.594367 | 0.519245 |

Z) neural network features, angry vs sad

  
|  | time window | peak latency | cluster *p* | peak Cohen's *d* |  | | | |  |
| **all electrodes** |  | | | |  | | | |  |
|  | | | | | | | | |  |

Model correlations, cluster permutation tests

|  | **left hemisphere** | | | | **right hemisphere** | | | |  |
|  | time window | peak latency | cluster *p* | peak Cohen's *d* | time window | peak latency | cluster *p* | peak Cohen's *d* |  |
| **anterior** |  | | | |  | | | |  |
| **central** | 185 - 280 ms | 240 ms | 0.0196 | 0.9875 | 215 - 495 ms | 240 ms | 0.0007 | 0.8433 |  |
 335 - 475 ms | 420 ms | 0.0145 | 0.6439 |  | | | ||  | **posterior** |  | | | |  | | | |

  

Model correlations, Bayesian statistics

|  | -200 | -195 | -190 | -185 | -180 | -175 | -170 | -165 | -160 | -155 | -150 | -145 | -140 | -135 | -130 | -125 | -120 | -115 | -110 | -105 | -100 | -95 | -90 | -85 | -80 | -75 | -70 | -65 | -60 | -55 | -50 | -45 | -40 | -35 | -30 | -25 | -20 | -15 | -10 | -5 | 0 | 5 | 10 | 15 | 20 | 25 | 30 | 35 | 40 | 45 | 50 | 55 | 60 | 65 | 70 | 75 | 80 | 85 | 90 | 95 | 100 | 105 | 110 | 115 | 120 | 125 | 130 | 135 | 140 | 145 | 150 | 155 | 160 | 165 | 170 | 175 | 180 | 185 | 190 | 195 | 200 | 205 | 210 | 215 | 220 | 225 | 230 | 235 | 240 | 245 | 250 | 255 | 260 | 265 | 270 | 275 | 280 | 285 | 290 | 295 | 300 | 305 | 310 | 315 | 320 | 325 | 330 | 335 | 340 | 345 | 350 | 355 | 360 | 365 | 370 | 375 | 380 | 385 | 390 | 395 | 400 | 405 | 410 | 415 | 420 | 425 | 430 | 435 | 440 | 445 | 450 | 455 | 460 | 465 | 470 | 475 | 480 | 485 | 490 | 495 | 500 | 505 | 510 | 515 | 520 | 525 | 530 | 535 | 540 | 545 | 550 | 555 | 560 | 565 | 570 | 575 | 580 | 585 | 590 | 595 | 600 | 605 | 610 | 615 | 620 | 625 | 630 | 635 | 640 | 645 | 650 | 655 | 660 | 665 | 670 | 675 | 680 | 685 | 690 | 695 | 700 | 705 | 710 | 715 | 720 | 725 | 730 | 735 | 740 | 745 | 750 | 755 | 760 | 765 | 770 | 775 | 780 | 785 | 790 | 795 | 800 | 805 | 810 | 815 | 820 | 825 | 830 | 835 | 840 | 845 | 850 | 855 | 860 | 865 | 870 | 875 | 880 | 885 | 890 | 895 | 900 | 905 | 910 | 915 | 920 | 925 | 930 | 935 | 940 | 945 | 950 | 955 | 960 | 965 | 970 | 975 | 980 | 985 | 990 | 995 | 1000 | 1005 | 1010 | 1015 | 1020 | 1025 | 1030 | 1035 | 1040 | 1045 | 1050 | 1055 | 1060 | 1065 | 1070 | 1075 | 1080 | 1085 | 1090 | 1095 | 1100 | 1105 | 1110 | 1115 | 1120 | 1125 | 1130 | 1135 | 1140 | 1145 | 1150 | 1155 | 1160 | 1165 | 1170 | 1175 | 1180 | 1185 | 1190 | 1195 |
| --- | --- | --- | --- | --- | --- | --- | --- | --- | --- | --- | --- | --- | --- | --- | --- | --- | --- | --- | --- | --- | --- | --- | --- | --- | --- | --- | --- | --- | --- | --- | --- | --- | --- | --- | --- | --- | --- | --- | --- | --- | --- | --- | --- | --- | --- | --- | --- | --- | --- | --- | --- | --- | --- | --- | --- | --- | --- | --- | --- | --- | --- | --- | --- | --- | --- | --- | --- | --- | --- | --- | --- | --- | --- | --- | --- | --- | --- | --- | --- | --- | --- | --- | --- | --- | --- | --- | --- | --- | --- | --- | --- | --- | --- | --- | --- | --- | --- | --- | --- | --- | --- | --- | --- | --- | --- | --- | --- | --- | --- | --- | --- | --- | --- | --- | --- | --- | --- | --- | --- | --- | --- | --- | --- | --- | --- | --- | --- | --- | --- | --- | --- | --- | --- | --- | --- | --- | --- | --- | --- | --- | --- | --- | --- | --- | --- | --- | --- | --- | --- | --- | --- | --- | --- | --- | --- | --- | --- | --- | --- | --- | --- | --- | --- | --- | --- | --- | --- | --- | --- | --- | --- | --- | --- | --- | --- | --- | --- | --- | --- | --- | --- | --- | --- | --- | --- | --- | --- | --- | --- | --- | --- | --- | --- | --- | --- | --- | --- | --- | --- | --- | --- | --- | --- | --- | --- | --- | --- | --- | --- | --- | --- | --- | --- | --- | --- | --- | --- | --- | --- | --- | --- | --- | --- | --- | --- | --- | --- | --- | --- | --- | --- | --- | --- | --- | --- | --- | --- | --- | --- | --- | --- | --- | --- | --- | --- | --- | --- | --- | --- | --- | --- | --- | --- | --- | --- | --- | --- | --- | --- | --- | --- | --- | --- | --- | --- | --- | --- | --- | --- | --- | --- | --- | --- | --- | --- | --- | --- | --- | --- | --- |
| left anterior | 0.234319 | 0.246382 | 0.247606 | 0.240484 | 0.221535 | 0.216337 | 0.242312 | 0.258824 | 0.241914 | 0.226886 | 0.215318 | 0.228739 | 0.301985 | 0.504903 | 0.920146 | 1.084944 | 0.998050 | 0.746639 | 0.436165 | 0.309915 | 0.252200 | 0.215879 | 0.217209 | 0.247207 | 0.310584 | 0.302000 | 0.327994 | 0.398191 | 0.490216 | 0.577393 | 0.503884 | 0.277279 | 0.214663 | 0.285906 | 0.432568 | 0.564215 | 0.596392 | 0.534122 | 0.415510 | 0.375396 | 0.332613 | 0.324795 | 0.345799 | 0.401845 | 0.422658 | 0.382730 | 0.277920 | 0.224022 | 0.220530 | 0.288367 | 0.335254 | 0.302539 | 0.225289 | 0.248646 | 0.593104 | 1.565556 | 3.440303 | 4.009456 | 4.252050 | 3.929521 | 2.475360 | 1.810659 | 1.780370 | 1.398007 | 0.853645 | 0.610868 | 0.424575 | 0.343919 | 0.321695 | 0.363044 | 0.488178 | 0.897497 | 1.580514 | 2.442159 | 2.121740 | 1.181265 | 0.581036 | 0.419557 | 0.401732 | 0.555658 | 1.047361 | 1.781931 | 2.064113 | 1.514552 | 0.861780 | 0.502056 | 0.380942 | 0.319807 | 0.277846 | 0.240985 | 0.233515 | 0.229312 | 0.226717 | 0.222221 | 0.222055 | 0.226629 | 0.239805 | 0.259116 | 0.305677 | 0.439673 | 0.697033 | 1.245105 | 2.311964 | 3.386057 | 3.177290 | 2.705290 | 1.539378 | 0.842442 | 0.516890 | 0.461713 | 0.437437 | 0.451057 | 0.436782 | 0.435958 | 0.337165 | 0.253482 | 0.215030 | 0.228899 | 0.277313 | 0.330225 | 0.357547 | 0.351830 | 0.334318 | 0.298700 | 0.267213 | 0.247415 | 0.240467 | 0.242709 | 0.246714 | 0.255365 | 0.261564 | 0.249132 | 0.222297 | 0.215032 | 0.239573 | 0.310566 | 0.437106 | 0.590073 | 0.617261 | 0.611155 | 0.580148 | 0.562467 | 0.572393 | 0.657955 | 0.698673 | 0.753894 | 0.676475 | 0.561267 | 0.445850 | 0.354986 | 0.278256 | 0.243554 | 0.225394 | 0.217343 | 0.215010 | 0.216271 | 0.216606 | 0.216492 | 0.217845 | 0.219159 | 0.217694 | 0.216270 | 0.215380 | 0.215025 | 0.214871 | 0.214651 | 0.214895 | 0.216002 | 0.219510 | 0.221277 | 0.216592 | 0.216219 | 0.239255 | 0.280699 | 0.323967 | 0.344843 | 0.313322 | 0.270391 | 0.239793 | 0.221258 | 0.214890 | 0.215472 | 0.217031 | 0.214897 | 0.216744 | 0.233625 | 0.299422 | 0.471154 | 0.635075 | 0.789682 | 1.024347 | 1.494594 | 2.391744 | 4.669984 | 8.541915 | 10.438388 | 6.559571 | 2.994833 | 1.262662 | 0.682247 | 0.451937 | 0.389553 | 0.417470 | 0.568444 | 0.789459 | 1.206039 | 1.722947 | 2.566586 | 3.906423 | 7.161594 | 16.179523 | 45.984202 | 79.588835 | 89.467041 | 56.000374 | 28.485127 | 12.643559 | 5.978617 | 3.000408 | 1.730012 | 1.177589 | 0.861882 | 0.713073 | 0.611285 | 0.608497 | 0.658657 | 0.821457 | 0.968227 | 1.089409 | 1.098853 | 1.093861 | 1.027720 | 0.977850 | 0.930144 | 0.928598 | 0.939372 | 0.965680 | 1.064936 | 1.093446 | 1.003226 | 0.857285 | 0.692318 | 0.545428 | 0.500532 | 0.475183 | 0.482840 | 0.526990 | 0.565299 | 0.582257 | 0.589372 | 0.607000 | 0.647697 | 0.763903 | 0.905287 | 1.199301 | 1.608848 | 2.071180 | 2.274225 | 2.134729 | 1.505921 | 0.997468 | 0.634447 | 0.461081 | 0.362586 | 0.326220 | 0.310914 | 0.314427 | 0.339142 | 0.390736 | 0.503949 | 0.714160 | 0.964644 | 1.166189 | 1.328476 | 1.146420 | 0.869551 | 0.605071 | 0.415678 | 0.344366 | 0.298008 |
| right anterior | 0.260778 | 0.238023 | 0.220652 | 0.214980 | 0.224385 | 0.247627 | 0.266139 | 0.272215 | 0.253720 | 0.239611 | 0.243690 | 0.258061 | 0.257130 | 0.287690 | 0.372798 | 0.424339 | 0.419950 | 0.352035 | 0.269354 | 0.222298 | 0.214627 | 0.235066 | 0.266934 | 0.315676 | 0.330980 | 0.358285 | 0.346416 | 0.295508 | 0.239133 | 0.221949 | 0.214783 | 0.226019 | 0.233331 | 0.234636 | 0.230426 | 0.219214 | 0.215017 | 0.214627 | 0.217072 | 0.216884 | 0.214748 | 0.215127 | 0.217419 | 0.220211 | 0.215047 | 0.218426 | 0.243674 | 0.324355 | 0.478239 | 0.625654 | 0.576081 | 0.411043 | 0.286269 | 0.230981 | 0.216740 | 0.215192 | 0.218124 | 0.233184 | 0.276813 | 0.407246 | 0.664685 | 1.380196 | 2.636286 | 5.201974 | 3.658683 | 1.679630 | 0.822945 | 0.547928 | 0.449103 | 0.574771 | 0.865883 | 1.614061 | 2.588731 | 3.729555 | 3.975682 | 4.753519 | 5.522927 | 7.793950 | 12.155198 | 14.143699 | 13.482602 | 11.564426 | 6.395365 | 2.564017 | 1.145519 | 0.563283 | 0.350581 | 0.280612 | 0.251258 | 0.242117 | 0.243002 | 0.235236 | 0.224315 | 0.219581 | 0.215419 | 0.214996 | 0.214965 | 0.214650 | 0.216608 | 0.229695 | 0.253659 | 0.281743 | 0.298871 | 0.273581 | 0.244946 | 0.237792 | 0.229545 | 0.226459 | 0.228462 | 0.228998 | 0.222264 | 0.214704 | 0.221046 | 0.245650 | 0.286746 | 0.339124 | 0.356456 | 0.342302 | 0.311142 | 0.302687 | 0.294434 | 0.290641 | 0.291663 | 0.291927 | 0.292521 | 0.292021 | 0.281553 | 0.277944 | 0.280348 | 0.278787 | 0.266665 | 0.242522 | 0.218769 | 0.214905 | 0.224719 | 0.248327 | 0.271578 | 0.282279 | 0.325194 | 0.415913 | 0.547342 | 0.760034 | 1.009161 | 0.920032 | 0.826572 | 0.664419 | 0.494961 | 0.412819 | 0.354325 | 0.285973 | 0.260134 | 0.248156 | 0.238435 | 0.236591 | 0.243110 | 0.259933 | 0.294911 | 0.375755 | 0.581887 | 0.963925 | 1.344797 | 1.895865 | 3.347031 | 7.011296 | 15.060864 | 20.698999 | 23.523502 | 25.498070 | 33.466773 | 26.289138 | 17.965953 | 7.580500 | 3.198882 | 1.399324 | 0.861379 | 0.619751 | 0.535946 | 0.531820 | 0.630338 | 0.784771 | 1.055408 | 1.637695 | 2.753746 | 3.645812 | 4.426969 | 3.596463 | 2.401223 | 1.577533 | 1.178045 | 0.850595 | 0.738664 | 0.636721 | 0.544368 | 0.458577 | 0.401435 | 0.349152 | 0.307994 | 0.291943 | 0.305695 | 0.362128 | 0.524372 | 1.023477 | 2.175484 | 4.095741 | 4.811320 | 4.215782 | 2.956912 | 1.970028 | 1.271911 | 0.776368 | 0.520762 | 0.388644 | 0.315012 | 0.280995 | 0.274373 | 0.290455 | 0.369499 | 0.568215 | 0.969342 | 1.338920 | 1.320097 | 0.903627 | 0.574580 | 0.389787 | 0.323899 | 0.320352 | 0.336201 | 0.361950 | 0.379860 | 0.375716 | 0.334177 | 0.279189 | 0.245903 | 0.225023 | 0.216450 | 0.214628 | 0.214939 | 0.214870 | 0.214739 | 0.219148 | 0.235355 | 0.273216 | 0.344363 | 0.461026 | 0.553226 | 0.567929 | 0.490065 | 0.374693 | 0.280710 | 0.228813 | 0.214737 | 0.216770 | 0.218998 | 0.221853 | 0.220586 | 0.218578 | 0.218237 | 0.218365 | 0.218749 | 0.222393 | 0.220543 | 0.217757 | 0.214826 | 0.214865 | 0.215307 | 0.214757 | 0.214940 | 0.218659 | 0.227657 | 0.242609 | 0.264677 | 0.295889 | 0.339466 | 0.375469 | 0.392454 | 0.393879 | 0.356309 | 0.330642 | 0.314895 | 0.297177 |
| left central | 0.232350 | 0.228997 | 0.226491 | 0.220875 | 0.219441 | 0.224713 | 0.238819 | 0.248431 | 0.284565 | 0.365671 | 0.422244 | 0.436890 | 0.428721 | 0.364086 | 0.314221 | 0.288566 | 0.285567 | 0.324637 | 0.388541 | 0.421563 | 0.549854 | 0.889805 | 1.807913 | 2.401151 | 2.775897 | 2.301027 | 1.716611 | 0.673625 | 0.296269 | 0.215086 | 0.267132 | 0.362058 | 0.370903 | 0.277251 | 0.220693 | 0.220261 | 0.244222 | 0.258798 | 0.261988 | 0.264715 | 0.236210 | 0.216960 | 0.224388 | 0.311213 | 0.609633 | 1.003004 | 1.114119 | 0.754748 | 0.561069 | 0.460782 | 0.381856 | 0.318425 | 0.258154 | 0.218542 | 0.218472 | 0.242483 | 0.283720 | 0.315381 | 0.327404 | 0.313238 | 0.286451 | 0.266890 | 0.261433 | 0.259399 | 0.255552 | 0.258174 | 0.238864 | 0.222009 | 0.216610 | 0.216217 | 0.217565 | 0.232441 | 0.253481 | 0.290739 | 0.354204 | 0.486000 | 0.849665 | 2.178731 | 7.530033 | 20.640077 | 29.822926 | 27.415803 | 24.483024 | 26.885090 | 38.598324 | 73.308298 | 147.821083 | 277.271808 | 306.999555 | 175.698804 | 98.271196 | 56.286542 | 37.328468 | 21.292088 | 11.128904 | 4.913057 | 2.043753 | 0.797538 | 0.414675 | 0.276342 | 0.230839 | 0.218992 | 0.217557 | 0.224027 | 0.264376 | 0.400406 | 0.781139 | 2.005551 | 4.185424 | 5.434348 | 5.970660 | 5.330510 | 4.080003 | 3.615726 | 3.205827 | 3.454583 | 3.906232 | 5.104603 | 6.180101 | 8.124390 | 8.948313 | 8.640831 | 7.964482 | 7.306372 | 6.741619 | 6.273585 | 5.149816 | 4.156700 | 3.409785 | 2.733507 | 2.995348 | 3.995129 | 5.489676 | 6.307793 | 4.464901 | 2.090341 | 1.173949 | 0.754217 | 0.488149 | 0.338492 | 0.249697 | 0.214683 | 0.245026 | 0.355504 | 0.547760 | 0.669298 | 0.509149 | 0.332813 | 0.235281 | 0.215343 | 0.247831 | 0.308136 | 0.357936 | 0.389327 | 0.424376 | 0.426604 | 0.407749 | 0.388890 | 0.350113 | 0.294338 | 0.243604 | 0.217157 | 0.221622 | 0.263779 | 0.334098 | 0.410226 | 0.466688 | 0.446453 | 0.350666 | 0.257771 | 0.215903 | 0.229105 | 0.275374 | 0.306998 | 0.280306 | 0.240618 | 0.219392 | 0.215263 | 0.222184 | 0.228214 | 0.228515 | 0.222550 | 0.216306 | 0.214786 | 0.214695 | 0.215710 | 0.220274 | 0.233784 | 0.250241 | 0.253802 | 0.243618 | 0.234189 | 0.230285 | 0.231391 | 0.234397 | 0.244196 | 0.271510 | 0.312431 | 0.367417 | 0.424018 | 0.476185 | 0.503483 | 0.528284 | 0.563373 | 0.615265 | 0.664935 | 0.667253 | 0.613444 | 0.529794 | 0.444274 | 0.360716 | 0.319346 | 0.297023 | 0.285553 | 0.269562 | 0.255621 | 0.236800 | 0.223051 | 0.214734 | 0.219903 | 0.237024 | 0.250104 | 0.256741 | 0.260701 | 0.266252 | 0.281278 | 0.323718 | 0.406528 | 0.535612 | 0.723976 | 0.931376 | 1.073056 | 1.071269 | 0.975205 | 0.816038 | 0.667767 | 0.549158 | 0.505714 | 0.499608 | 0.522748 | 0.570233 | 0.637234 | 0.650566 | 0.671972 | 0.598539 | 0.479864 | 0.371124 | 0.293734 | 0.241846 | 0.220792 | 0.215169 | 0.214853 | 0.215225 | 0.215271 | 0.215336 | 0.215150 | 0.215062 | 0.215258 | 0.215165 | 0.215051 | 0.214929 | 0.214899 | 0.214826 | 0.216747 | 0.223696 | 0.233659 | 0.248809 | 0.262897 | 0.266974 | 0.250080 | 0.237400 | 0.235931 | 0.231484 | 0.245200 | 0.258518 | 0.274576 | 0.296968 | 0.327219 | 0.314266 | 0.327506 |
| right central | 2.163037 | 1.789976 | 1.620016 | 1.503226 | 1.457592 | 1.387476 | 1.228378 | 1.239468 | 0.857397 | 0.829001 | 0.917819 | 0.767358 | 0.426662 | 0.294281 | 0.236568 | 0.225480 | 0.223451 | 0.217956 | 0.214627 | 0.220056 | 0.249867 | 0.340783 | 0.432212 | 0.499927 | 0.454015 | 0.419962 | 0.348305 | 0.291685 | 0.237854 | 0.216721 | 0.214733 | 0.221154 | 0.235768 | 0.263877 | 0.310176 | 0.346889 | 0.364950 | 0.363230 | 0.315840 | 0.287145 | 0.281398 | 0.278094 | 0.286494 | 0.325995 | 0.343652 | 0.332881 | 0.306537 | 0.295939 | 0.280124 | 0.272094 | 0.249436 | 0.222816 | 0.215828 | 0.241074 | 0.344000 | 0.683957 | 1.778494 | 5.060000 | 7.952747 | 2.331728 | 0.635583 | 0.299652 | 0.236152 | 0.219024 | 0.221623 | 0.233293 | 0.278417 | 0.362158 | 0.492967 | 0.604113 | 0.611820 | 0.565064 | 0.486946 | 0.415838 | 0.326849 | 0.285645 | 0.249111 | 0.235903 | 0.227202 | 0.219002 | 0.218112 | 0.294992 | 0.916174 | 5.026001 | 29.117978 | 135.097764 | 244.209828 | 175.422235 | 83.027657 | 34.936244 | 16.474089 | 9.857055 | 7.204051 | 5.526587 | 4.122953 | 3.002314 | 2.192909 | 1.503843 | 1.228889 | 1.107510 | 1.215096 | 1.629405 | 2.455347 | 3.468111 | 4.714729 | 5.440148 | 5.811487 | 6.910278 | 10.348224 | 19.118912 | 31.586456 | 44.670803 | 52.857080 | 69.583796 | 63.614282 | 50.265884 | 28.990130 | 18.471277 | 12.261328 | 11.476583 | 13.923967 | 23.013843 | 34.918908 | 47.805451 | 63.935083 | 90.092505 | 101.411707 | 97.066928 | 93.991137 | 94.272862 | 104.913648 | 199.799495 | 584.347988 | 1834.313183 | 3061.775844 | 1201.582337 | 193.576576 | 26.762249 | 5.555244 | 1.777011 | 0.662654 | 0.339454 | 0.258987 | 0.233236 | 0.226263 | 0.232556 | 0.254745 | 0.320491 | 0.461483 | 0.671547 | 0.908201 | 1.183587 | 1.145894 | 1.022408 | 0.924435 | 0.871821 | 0.942288 | 1.234569 | 1.711893 | 2.206157 | 2.130366 | 1.477210 | 0.877651 | 0.504168 | 0.348421 | 0.292299 | 0.284691 | 0.326222 | 0.465739 | 0.697730 | 1.028064 | 1.212390 | 1.165378 | 0.971830 | 0.800114 | 0.675228 | 0.692402 | 0.689357 | 0.650636 | 0.645450 | 0.588068 | 0.518991 | 0.507093 | 0.531390 | 0.589997 | 0.731541 | 0.836278 | 0.895445 | 0.907122 | 0.828994 | 0.632387 | 0.514485 | 0.430846 | 0.335677 | 0.275655 | 0.240436 | 0.216516 | 0.219183 | 0.235127 | 0.237295 | 0.218153 | 0.223444 | 0.283996 | 0.418343 | 0.577389 | 0.551512 | 0.387197 | 0.273629 | 0.222741 | 0.217778 | 0.247610 | 0.280728 | 0.286148 | 0.277114 | 0.269848 | 0.253143 | 0.233378 | 0.222816 | 0.217674 | 0.215115 | 0.214638 | 0.214694 | 0.214698 | 0.214869 | 0.216167 | 0.220122 | 0.226661 | 0.239160 | 0.256215 | 0.269218 | 0.274010 | 0.268315 | 0.255203 | 0.252463 | 0.261725 | 0.273689 | 0.295937 | 0.322843 | 0.333848 | 0.319878 | 0.297329 | 0.258349 | 0.226343 | 0.214645 | 0.226617 | 0.260393 | 0.291921 | 0.300338 | 0.275142 | 0.236919 | 0.221963 | 0.215594 | 0.214969 | 0.218099 | 0.223449 | 0.236711 | 0.250682 | 0.276183 | 0.305564 | 0.323751 | 0.344079 | 0.363574 | 0.335938 | 0.306090 | 0.293955 | 0.290235 | 0.282160 | 0.276638 | 0.266254 | 0.249539 | 0.235671 | 0.223520 | 0.216493 | 0.214814 | 0.220510 | 0.246099 | 0.282900 | 0.326664 | 0.353117 | 0.385405 |
| left posterior | 1.807774 | 1.979805 | 1.695037 | 1.219335 | 0.716451 | 0.490064 | 0.385734 | 0.350858 | 0.305230 | 0.263310 | 0.228703 | 0.215917 | 0.250340 | 0.292889 | 0.307693 | 0.266981 | 0.215209 | 0.271869 | 0.493957 | 0.691961 | 0.509203 | 0.282664 | 0.219419 | 0.540576 | 2.287205 | 4.069493 | 2.860063 | 1.102857 | 0.433279 | 0.271755 | 0.238247 | 0.238688 | 0.284016 | 0.400156 | 0.706922 | 1.328570 | 2.018439 | 2.350238 | 2.176329 | 1.217671 | 0.392957 | 0.217013 | 0.411928 | 0.810392 | 1.060194 | 1.089146 | 1.011525 | 0.846870 | 0.670877 | 0.570330 | 0.494043 | 0.445924 | 0.396614 | 0.352484 | 0.296810 | 0.257096 | 0.239159 | 0.250470 | 0.328377 | 0.590571 | 0.946129 | 1.073960 | 0.877892 | 0.573349 | 0.376299 | 0.279333 | 0.235278 | 0.219031 | 0.214839 | 0.214841 | 0.214892 | 0.214649 | 0.218639 | 0.239081 | 0.302204 | 0.414000 | 0.455502 | 0.352371 | 0.253670 | 0.216653 | 0.221229 | 0.237148 | 0.240718 | 0.229721 | 0.218272 | 0.215558 | 0.230473 | 0.267485 | 0.317519 | 0.360116 | 0.414625 | 0.475191 | 0.530653 | 0.597771 | 0.662546 | 0.656209 | 0.647428 | 0.602537 | 0.514113 | 0.459724 | 0.439975 | 0.460231 | 0.523989 | 0.597913 | 0.623947 | 0.604794 | 0.513701 | 0.417218 | 0.355639 | 0.302209 | 0.268412 | 0.245175 | 0.226967 | 0.216736 | 0.214626 | 0.215651 | 0.216039 | 0.216290 | 0.216844 | 0.216038 | 0.216481 | 0.217146 | 0.215535 | 0.214628 | 0.216361 | 0.235544 | 0.258114 | 0.268356 | 0.267375 | 0.248292 | 0.225752 | 0.216593 | 0.214813 | 0.216275 | 0.216362 | 0.215665 | 0.214651 | 0.216213 | 0.222433 | 0.235237 | 0.256840 | 0.295114 | 0.329808 | 0.345583 | 0.329797 | 0.317104 | 0.308454 | 0.318684 | 0.359035 | 0.455430 | 0.662657 | 1.046609 | 1.593663 | 2.136739 | 2.480437 | 2.083060 | 1.397076 | 0.916482 | 0.623017 | 0.478519 | 0.477764 | 0.632160 | 1.025093 | 1.932831 | 2.971794 | 3.308088 | 3.256842 | 2.730779 | 1.959765 | 1.517566 | 1.102855 | 0.795718 | 0.630198 | 0.534110 | 0.450887 | 0.425598 | 0.434212 | 0.482385 | 0.596786 | 0.824535 | 1.113448 | 1.527586 | 2.079619 | 2.601999 | 2.935808 | 3.031877 | 2.667639 | 2.121277 | 1.649973 | 1.498738 | 1.590796 | 2.130091 | 3.094617 | 4.819811 | 5.954270 | 4.882731 | 3.070954 | 1.771359 | 0.953787 | 0.598385 | 0.469415 | 0.425795 | 0.430554 | 0.443482 | 0.463336 | 0.454996 | 0.436433 | 0.394662 | 0.364864 | 0.343412 | 0.339462 | 0.321331 | 0.292809 | 0.262732 | 0.240432 | 0.217267 | 0.217784 | 0.238123 | 0.267064 | 0.296776 | 0.287737 | 0.260766 | 0.230073 | 0.215012 | 0.223891 | 0.260563 | 0.320115 | 0.383035 | 0.403361 | 0.386725 | 0.363245 | 0.338188 | 0.309440 | 0.285532 | 0.258421 | 0.234114 | 0.219732 | 0.214719 | 0.215341 | 0.216884 | 0.215335 | 0.214799 | 0.217667 | 0.219460 | 0.221978 | 0.221155 | 0.217494 | 0.214811 | 0.215086 | 0.216893 | 0.219477 | 0.222154 | 0.226767 | 0.232876 | 0.235704 | 0.238783 | 0.232953 | 0.224694 | 0.217955 | 0.214657 | 0.219496 | 0.230412 | 0.245723 | 0.263624 | 0.275777 | 0.284064 | 0.293124 | 0.303802 | 0.325639 | 0.375879 | 0.427931 | 0.470865 | 0.566430 | 0.633098 | 0.668966 | 0.681793 | 0.699048 | 0.686549 | 0.696127 | 0.653806 |
| right posterior | 8.035051 | 9.434785 | 10.446007 | 11.814621 | 7.399993 | 4.274915 | 2.112141 | 1.035925 | 0.530417 | 0.398955 | 0.377700 | 0.368339 | 0.411430 | 0.584325 | 0.797249 | 0.909308 | 0.841990 | 0.562067 | 0.369223 | 0.246807 | 0.216967 | 0.282987 | 0.377494 | 0.459132 | 0.448860 | 0.361759 | 0.263256 | 0.221008 | 0.214785 | 0.220839 | 0.230141 | 0.239987 | 0.244980 | 0.237727 | 0.234810 | 0.254202 | 0.285188 | 0.310506 | 0.358177 | 0.434581 | 0.527798 | 0.619487 | 0.758022 | 0.814138 | 0.906040 | 0.760156 | 0.654188 | 0.686741 | 0.924937 | 1.031576 | 0.880780 | 0.508147 | 0.361484 | 0.325422 | 0.346725 | 0.421082 | 0.624909 | 1.145390 | 2.318391 | 3.783416 | 4.427183 | 5.174352 | 4.315215 | 2.330965 | 0.998755 | 0.453099 | 0.266052 | 0.220284 | 0.214652 | 0.216826 | 0.215059 | 0.217381 | 0.226864 | 0.241801 | 0.271707 | 0.288338 | 0.274376 | 0.246072 | 0.219360 | 0.216480 | 0.228442 | 0.249614 | 0.265043 | 0.274622 | 0.278571 | 0.279967 | 0.264456 | 0.251721 | 0.246821 | 0.256109 | 0.276643 | 0.306833 | 0.337398 | 0.334803 | 0.308562 | 0.266262 | 0.234465 | 0.216343 | 0.215386 | 0.217370 | 0.215303 | 0.215033 | 0.222932 | 0.246307 | 0.282251 | 0.325690 | 0.376543 | 0.429163 | 0.457559 | 0.462405 | 0.452086 | 0.424509 | 0.395930 | 0.387592 | 0.374795 | 0.382430 | 0.411389 | 0.480735 | 0.575942 | 0.717338 | 0.823543 | 0.785463 | 0.576060 | 0.363131 | 0.249435 | 0.214890 | 0.222120 | 0.228412 | 0.221061 | 0.214633 | 0.223274 | 0.258902 | 0.316076 | 0.356670 | 0.382519 | 0.382783 | 0.354023 | 0.325126 | 0.295926 | 0.268952 | 0.245440 | 0.229182 | 0.221123 | 0.218457 | 0.215871 | 0.214676 | 0.215183 | 0.216282 | 0.216198 | 0.214745 | 0.215782 | 0.228175 | 0.268575 | 0.365387 | 0.559520 | 0.985099 | 1.717383 | 2.783466 | 4.063098 | 4.835329 | 4.200528 | 2.994707 | 1.672585 | 0.836977 | 0.501309 | 0.364329 | 0.302185 | 0.284547 | 0.297587 | 0.322789 | 0.358065 | 0.389477 | 0.418228 | 0.439523 | 0.432860 | 0.415079 | 0.379262 | 0.328341 | 0.277984 | 0.241983 | 0.219634 | 0.214680 | 0.221122 | 0.234976 | 0.245180 | 0.248725 | 0.244653 | 0.234654 | 0.227359 | 0.223313 | 0.223685 | 0.232752 | 0.243878 | 0.255705 | 0.255671 | 0.239096 | 0.218607 | 0.215311 | 0.230808 | 0.249360 | 0.263717 | 0.264696 | 0.256233 | 0.239430 | 0.227212 | 0.216611 | 0.214975 | 0.221479 | 0.238167 | 0.273871 | 0.343714 | 0.447579 | 0.513826 | 0.472759 | 0.369759 | 0.262277 | 0.215332 | 0.236432 | 0.309263 | 0.421075 | 0.489026 | 0.449052 | 0.349683 | 0.264093 | 0.224515 | 0.214634 | 0.220030 | 0.225913 | 0.225993 | 0.223461 | 0.218309 | 0.214688 | 0.222644 | 0.258952 | 0.337757 | 0.464004 | 0.594773 | 0.669412 | 0.629536 | 0.570263 | 0.501495 | 0.430201 | 0.360388 | 0.298672 | 0.249392 | 0.222637 | 0.214688 | 0.218226 | 0.217874 | 0.215102 | 0.215742 | 0.226167 | 0.246543 | 0.267178 | 0.278665 | 0.278680 | 0.282906 | 0.290298 | 0.310971 | 0.348607 | 0.392230 | 0.401871 | 0.395126 | 0.359179 | 0.332834 | 0.313882 | 0.303266 | 0.291449 | 0.286425 | 0.271370 | 0.257100 | 0.244451 | 0.230711 | 0.222361 | 0.217680 | 0.215175 | 0.214626 | 0.215283 | 0.216590 | 0.218076 |
| all electrodes | 0.215179 | 0.227553 | 0.261608 | 0.356070 | 0.470229 | 0.434728 | 0.338980 | 0.273885 | 0.227714 | 0.214720 | 0.217630 | 0.221480 | 0.225713 | 0.229277 | 0.230134 | 0.236403 | 0.249539 | 0.303093 | 0.374673 | 0.434789 | 0.552801 | 0.760440 | 0.687450 | 0.645442 | 0.603713 | 0.605366 | 0.744253 | 0.790236 | 0.664733 | 0.573875 | 0.482748 | 0.376391 | 0.372656 | 0.381580 | 0.394120 | 0.412980 | 0.430770 | 0.394371 | 0.358461 | 0.327590 | 0.278789 | 0.245886 | 0.229528 | 0.225213 | 0.224930 | 0.243643 | 0.289937 | 0.347616 | 0.325601 | 0.289674 | 0.242147 | 0.216492 | 0.228099 | 0.282451 | 0.377318 | 0.483477 | 0.712113 | 1.056134 | 1.635922 | 2.080301 | 2.213571 | 1.638970 | 1.295246 | 0.858792 | 0.531179 | 0.301674 | 0.219372 | 0.233226 | 0.305852 | 0.358320 | 0.331961 | 0.256333 | 0.214849 | 0.252793 | 0.369484 | 0.470055 | 0.452495 | 0.336638 | 0.245446 | 0.214701 | 0.236132 | 0.300612 | 0.407419 | 0.529979 | 0.664566 | 0.699469 | 0.654847 | 0.593407 | 0.528114 | 0.407437 | 0.357173 | 0.299049 | 0.256014 | 0.237975 | 0.239295 | 0.237525 | 0.241469 | 0.244906 | 0.249601 | 0.252211 | 0.245122 | 0.244387 | 0.246738 | 0.244777 | 0.241897 | 0.242909 | 0.241349 | 0.250345 | 0.263618 | 0.290543 | 0.357521 | 0.466703 | 0.603664 | 0.754493 | 0.918792 | 0.938037 | 0.888403 | 0.733822 | 0.627163 | 0.593565 | 0.591207 | 0.581830 | 0.665723 | 0.827781 | 1.067404 | 1.432131 | 1.815028 | 1.743774 | 1.507189 | 1.141167 | 0.835829 | 0.653051 | 0.492857 | 0.365618 | 0.301753 | 0.270410 | 0.242726 | 0.226453 | 0.217197 | 0.215064 | 0.229230 | 0.260912 | 0.319266 | 0.340482 | 0.339251 | 0.319844 | 0.295523 | 0.257836 | 0.238134 | 0.221181 | 0.214626 | 0.220688 | 0.233577 | 0.251463 | 0.253852 | 0.248151 | 0.237568 | 0.228463 | 0.221356 | 0.216315 | 0.214694 | 0.216922 | 0.223942 | 0.239062 | 0.266535 | 0.311025 | 0.392319 | 0.515561 | 0.616437 | 0.622756 | 0.574813 | 0.496842 | 0.415689 | 0.352305 | 0.309868 | 0.276762 | 0.251014 | 0.233126 | 0.224548 | 0.222216 | 0.226296 | 0.237659 | 0.273856 | 0.340770 | 0.434229 | 0.502215 | 0.555811 | 0.570057 | 0.565750 | 0.498158 | 0.463308 | 0.433938 | 0.399929 | 0.382362 | 0.393495 | 0.387882 | 0.398559 | 0.385748 | 0.336736 | 0.298585 | 0.273622 | 0.258999 | 0.259526 | 0.265627 | 0.277357 | 0.317794 | 0.357087 | 0.362319 | 0.353419 | 0.340358 | 0.310207 | 0.282806 | 0.271652 | 0.275194 | 0.274432 | 0.267401 | 0.258909 | 0.249498 | 0.233800 | 0.221089 | 0.216015 | 0.214777 | 0.214766 | 0.215269 | 0.216923 | 0.220770 | 0.227144 | 0.232282 | 0.242880 | 0.257314 | 0.261608 | 0.250902 | 0.229756 | 0.217392 | 0.214746 | 0.218802 | 0.226073 | 0.228938 | 0.230170 | 0.227669 | 0.221684 | 0.218043 | 0.215874 | 0.214760 | 0.214905 | 0.215405 | 0.215652 | 0.214946 | 0.214627 | 0.215713 | 0.217343 | 0.217582 | 0.214937 | 0.215931 | 0.228543 | 0.261085 | 0.330965 | 0.424745 | 0.523414 | 0.591767 | 0.596487 | 0.477495 | 0.395287 | 0.338798 | 0.309609 | 0.307016 | 0.356480 | 0.420739 | 0.585063 | 0.848651 | 1.027051 | 1.157244 | 1.413565 | 1.370261 | 1.380621 | 1.332550 | 1.195977 | 0.892122 | 0.782545 | 0.667827 |

AA) neural network features, happy vs sad

  
|  | time window | peak latency | cluster *p* | peak Cohen's *d* |  | | | |  |
| **all electrodes** |  | | | |  | | | |  |
|  | | | | | | | | |  |

Model correlations, cluster permutation tests

|  | **left hemisphere** | | | | **right hemisphere** | | | |  |
|  | time window | peak latency | cluster *p* | peak Cohen's *d* | time window | peak latency | cluster *p* | peak Cohen's *d* |  |
| **anterior** |  | | | |  | | | |  |
| **central** | 250 - 385 ms | 360 ms | 0.026 | -0.5521 |  | | | |  |
 520 - 735 ms | 685 ms | 0.0039 | -0.7995 |  | | | ||  | **posterior** |  | | | |  | | | |

  

Model correlations, Bayesian statistics

|  | -200 | -195 | -190 | -185 | -180 | -175 | -170 | -165 | -160 | -155 | -150 | -145 | -140 | -135 | -130 | -125 | -120 | -115 | -110 | -105 | -100 | -95 | -90 | -85 | -80 | -75 | -70 | -65 | -60 | -55 | -50 | -45 | -40 | -35 | -30 | -25 | -20 | -15 | -10 | -5 | 0 | 5 | 10 | 15 | 20 | 25 | 30 | 35 | 40 | 45 | 50 | 55 | 60 | 65 | 70 | 75 | 80 | 85 | 90 | 95 | 100 | 105 | 110 | 115 | 120 | 125 | 130 | 135 | 140 | 145 | 150 | 155 | 160 | 165 | 170 | 175 | 180 | 185 | 190 | 195 | 200 | 205 | 210 | 215 | 220 | 225 | 230 | 235 | 240 | 245 | 250 | 255 | 260 | 265 | 270 | 275 | 280 | 285 | 290 | 295 | 300 | 305 | 310 | 315 | 320 | 325 | 330 | 335 | 340 | 345 | 350 | 355 | 360 | 365 | 370 | 375 | 380 | 385 | 390 | 395 | 400 | 405 | 410 | 415 | 420 | 425 | 430 | 435 | 440 | 445 | 450 | 455 | 460 | 465 | 470 | 475 | 480 | 485 | 490 | 495 | 500 | 505 | 510 | 515 | 520 | 525 | 530 | 535 | 540 | 545 | 550 | 555 | 560 | 565 | 570 | 575 | 580 | 585 | 590 | 595 | 600 | 605 | 610 | 615 | 620 | 625 | 630 | 635 | 640 | 645 | 650 | 655 | 660 | 665 | 670 | 675 | 680 | 685 | 690 | 695 | 700 | 705 | 710 | 715 | 720 | 725 | 730 | 735 | 740 | 745 | 750 | 755 | 760 | 765 | 770 | 775 | 780 | 785 | 790 | 795 | 800 | 805 | 810 | 815 | 820 | 825 | 830 | 835 | 840 | 845 | 850 | 855 | 860 | 865 | 870 | 875 | 880 | 885 | 890 | 895 | 900 | 905 | 910 | 915 | 920 | 925 | 930 | 935 | 940 | 945 | 950 | 955 | 960 | 965 | 970 | 975 | 980 | 985 | 990 | 995 | 1000 | 1005 | 1010 | 1015 | 1020 | 1025 | 1030 | 1035 | 1040 | 1045 | 1050 | 1055 | 1060 | 1065 | 1070 | 1075 | 1080 | 1085 | 1090 | 1095 | 1100 | 1105 | 1110 | 1115 | 1120 | 1125 | 1130 | 1135 | 1140 | 1145 | 1150 | 1155 | 1160 | 1165 | 1170 | 1175 | 1180 | 1185 | 1190 | 1195 |
| --- | --- | --- | --- | --- | --- | --- | --- | --- | --- | --- | --- | --- | --- | --- | --- | --- | --- | --- | --- | --- | --- | --- | --- | --- | --- | --- | --- | --- | --- | --- | --- | --- | --- | --- | --- | --- | --- | --- | --- | --- | --- | --- | --- | --- | --- | --- | --- | --- | --- | --- | --- | --- | --- | --- | --- | --- | --- | --- | --- | --- | --- | --- | --- | --- | --- | --- | --- | --- | --- | --- | --- | --- | --- | --- | --- | --- | --- | --- | --- | --- | --- | --- | --- | --- | --- | --- | --- | --- | --- | --- | --- | --- | --- | --- | --- | --- | --- | --- | --- | --- | --- | --- | --- | --- | --- | --- | --- | --- | --- | --- | --- | --- | --- | --- | --- | --- | --- | --- | --- | --- | --- | --- | --- | --- | --- | --- | --- | --- | --- | --- | --- | --- | --- | --- | --- | --- | --- | --- | --- | --- | --- | --- | --- | --- | --- | --- | --- | --- | --- | --- | --- | --- | --- | --- | --- | --- | --- | --- | --- | --- | --- | --- | --- | --- | --- | --- | --- | --- | --- | --- | --- | --- | --- | --- | --- | --- | --- | --- | --- | --- | --- | --- | --- | --- | --- | --- | --- | --- | --- | --- | --- | --- | --- | --- | --- | --- | --- | --- | --- | --- | --- | --- | --- | --- | --- | --- | --- | --- | --- | --- | --- | --- | --- | --- | --- | --- | --- | --- | --- | --- | --- | --- | --- | --- | --- | --- | --- | --- | --- | --- | --- | --- | --- | --- | --- | --- | --- | --- | --- | --- | --- | --- | --- | --- | --- | --- | --- | --- | --- | --- | --- | --- | --- | --- | --- | --- | --- | --- | --- | --- | --- | --- | --- | --- | --- | --- | --- | --- | --- | --- | --- | --- | --- | --- | --- | --- | --- | --- | --- | --- |
| left anterior | 0.214700 | 0.214643 | 0.217977 | 0.242631 | 0.275896 | 0.292091 | 0.273451 | 0.243446 | 0.214661 | 0.239159 | 0.285795 | 0.313227 | 0.319362 | 0.267132 | 0.228002 | 0.215635 | 0.233198 | 0.251712 | 0.267021 | 0.291302 | 0.294862 | 0.299781 | 0.322645 | 0.427776 | 0.649964 | 1.011708 | 1.929474 | 3.701001 | 5.782817 | 9.265799 | 9.093386 | 4.161126 | 1.492843 | 0.425274 | 0.221750 | 0.230164 | 0.296880 | 0.301784 | 0.273018 | 0.282305 | 0.311655 | 0.344828 | 0.402520 | 0.393301 | 0.327943 | 0.238125 | 0.219458 | 0.321351 | 0.557895 | 0.993698 | 1.232388 | 1.196552 | 0.695803 | 0.351013 | 0.219908 | 0.227848 | 0.282963 | 0.314085 | 0.354026 | 0.381254 | 0.370708 | 0.301600 | 0.252475 | 0.218946 | 0.220821 | 0.253317 | 0.296735 | 0.369382 | 0.531148 | 0.667970 | 1.004804 | 1.771645 | 3.319650 | 5.319474 | 7.447505 | 5.494654 | 3.127194 | 1.571623 | 0.697261 | 0.376479 | 0.261776 | 0.223462 | 0.215804 | 0.217183 | 0.221351 | 0.232171 | 0.244952 | 0.256869 | 0.259940 | 0.260915 | 0.246933 | 0.228987 | 0.216024 | 0.214626 | 0.214790 | 0.216361 | 0.221403 | 0.231008 | 0.268157 | 0.344918 | 0.446074 | 0.531387 | 0.557998 | 0.487268 | 0.355845 | 0.273412 | 0.224479 | 0.214641 | 0.219633 | 0.222407 | 0.221289 | 0.215770 | 0.214630 | 0.214668 | 0.214684 | 0.218353 | 0.247436 | 0.309228 | 0.399589 | 0.479497 | 0.451128 | 0.334983 | 0.244532 | 0.216578 | 0.293368 | 0.480281 | 0.683451 | 0.690209 | 0.390116 | 0.236081 | 0.225087 | 0.285662 | 0.364634 | 0.446809 | 0.523766 | 0.531222 | 0.493499 | 0.403881 | 0.328927 | 0.278278 | 0.254929 | 0.249530 | 0.255366 | 0.259381 | 0.278112 | 0.318882 | 0.386809 | 0.447588 | 0.473622 | 0.447355 | 0.410780 | 0.353985 | 0.297291 | 0.251380 | 0.227469 | 0.216915 | 0.214732 | 0.214979 | 0.218165 | 0.227293 | 0.235178 | 0.228590 | 0.219660 | 0.215444 | 0.215095 | 0.220270 | 0.230293 | 0.236656 | 0.242685 | 0.259893 | 0.291423 | 0.333572 | 0.382639 | 0.420197 | 0.442122 | 0.442264 | 0.392452 | 0.355854 | 0.330398 | 0.296891 | 0.257310 | 0.237139 | 0.228643 | 0.228690 | 0.236194 | 0.257304 | 0.289124 | 0.340428 | 0.399462 | 0.473555 | 0.527785 | 0.619530 | 0.594090 | 0.475092 | 0.317661 | 0.227469 | 0.236432 | 0.405647 | 0.666018 | 0.779657 | 0.760970 | 0.590449 | 0.433137 | 0.333416 | 0.272225 | 0.235396 | 0.220803 | 0.215222 | 0.214797 | 0.215935 | 0.216151 | 0.215438 | 0.214841 | 0.222228 | 0.252197 | 0.326356 | 0.468770 | 0.649918 | 0.775019 | 0.719707 | 0.586389 | 0.489745 | 0.421396 | 0.373470 | 0.357962 | 0.345999 | 0.328385 | 0.300415 | 0.276901 | 0.258071 | 0.253156 | 0.253480 | 0.259331 | 0.269714 | 0.285848 | 0.279899 | 0.262037 | 0.246780 | 0.229215 | 0.217919 | 0.215246 | 0.214868 | 0.214662 | 0.215081 | 0.219801 | 0.230979 | 0.251370 | 0.282938 | 0.328579 | 0.367118 | 0.414043 | 0.434614 | 0.447123 | 0.448487 | 0.431309 | 0.384682 | 0.373667 | 0.366276 | 0.357485 | 0.353131 | 0.364713 | 0.381762 | 0.415023 | 0.453571 | 0.485352 | 0.514682 | 0.531777 | 0.522866 | 0.504881 | 0.474393 | 0.432830 | 0.379519 | 0.339094 | 0.307876 | 0.297827 | 0.292282 | 0.297903 | 0.313643 | 0.323209 | 0.326825 |
| right anterior | 0.484942 | 0.635127 | 0.606403 | 0.609746 | 0.587405 | 0.475892 | 0.426947 | 0.391168 | 0.361803 | 0.311317 | 0.234068 | 0.217732 | 0.245195 | 0.320397 | 0.374323 | 0.298919 | 0.223393 | 0.254910 | 0.849465 | 4.779226 | 24.729104 | 73.249742 | 75.019744 | 43.627976 | 17.328559 | 7.763383 | 3.856767 | 1.626638 | 0.706570 | 0.390791 | 0.235063 | 0.219086 | 0.250788 | 0.275978 | 0.282594 | 0.281866 | 0.297642 | 0.336082 | 0.375298 | 0.474302 | 0.636753 | 0.635109 | 0.504798 | 0.405670 | 0.305866 | 0.258451 | 0.231987 | 0.219394 | 0.214715 | 0.216666 | 0.223481 | 0.227523 | 0.227875 | 0.225581 | 0.218128 | 0.215085 | 0.214744 | 0.218195 | 0.226564 | 0.230321 | 0.230656 | 0.224321 | 0.215630 | 0.219480 | 0.236125 | 0.254288 | 0.274000 | 0.278609 | 0.259717 | 0.247255 | 0.242177 | 0.249367 | 0.268436 | 0.295009 | 0.326308 | 0.361505 | 0.374764 | 0.363208 | 0.342320 | 0.320663 | 0.311657 | 0.309799 | 0.334523 | 0.408341 | 0.544865 | 0.736956 | 0.913834 | 0.909351 | 0.825104 | 0.738569 | 0.660666 | 0.572303 | 0.524399 | 0.469630 | 0.420962 | 0.353397 | 0.305296 | 0.267779 | 0.255419 | 0.244975 | 0.251081 | 0.264013 | 0.285417 | 0.295908 | 0.282950 | 0.240971 | 0.217977 | 0.215416 | 0.222117 | 0.222821 | 0.216533 | 0.215643 | 0.229128 | 0.261798 | 0.301059 | 0.354301 | 0.410481 | 0.442807 | 0.411047 | 0.376285 | 0.324050 | 0.269306 | 0.227223 | 0.214748 | 0.217786 | 0.227827 | 0.243234 | 0.270319 | 0.291209 | 0.326204 | 0.362851 | 0.393448 | 0.338204 | 0.269044 | 0.222353 | 0.215781 | 0.232888 | 0.259451 | 0.286817 | 0.285105 | 0.278207 | 0.278212 | 0.281279 | 0.283254 | 0.296103 | 0.313346 | 0.336900 | 0.379164 | 0.408486 | 0.454637 | 0.481248 | 0.460848 | 0.357177 | 0.276666 | 0.225704 | 0.215124 | 0.233709 | 0.250333 | 0.253011 | 0.244974 | 0.231348 | 0.219849 | 0.216459 | 0.217627 | 0.223340 | 0.236747 | 0.267764 | 0.317674 | 0.391590 | 0.470801 | 0.525639 | 0.519940 | 0.426849 | 0.317865 | 0.257360 | 0.226457 | 0.215797 | 0.214676 | 0.214680 | 0.214696 | 0.218980 | 0.237419 | 0.302829 | 0.381293 | 0.423502 | 0.364329 | 0.295937 | 0.241847 | 0.220501 | 0.215642 | 0.216888 | 0.223749 | 0.241230 | 0.274351 | 0.330578 | 0.391405 | 0.398378 | 0.351721 | 0.304956 | 0.276544 | 0.251712 | 0.238776 | 0.236420 | 0.238918 | 0.227687 | 0.217930 | 0.214668 | 0.215169 | 0.217823 | 0.218321 | 0.215333 | 0.215489 | 0.220131 | 0.225103 | 0.226681 | 0.224579 | 0.221839 | 0.216582 | 0.214919 | 0.214628 | 0.214927 | 0.216259 | 0.217859 | 0.220752 | 0.220601 | 0.218210 | 0.216975 | 0.217350 | 0.223881 | 0.262868 | 0.403247 | 0.856774 | 2.007872 | 4.534571 | 7.964099 | 9.161934 | 6.738327 | 3.912328 | 1.945107 | 1.084831 | 0.702183 | 0.496413 | 0.387115 | 0.321504 | 0.287233 | 0.269020 | 0.265031 | 0.270109 | 0.288153 | 0.308215 | 0.334812 | 0.365362 | 0.410810 | 0.449177 | 0.454430 | 0.446876 | 0.444886 | 0.424436 | 0.379118 | 0.352886 | 0.331642 | 0.315364 | 0.303536 | 0.310854 | 0.326892 | 0.351737 | 0.362156 | 0.346685 | 0.311634 | 0.271654 | 0.235634 | 0.220744 | 0.216918 | 0.214885 | 0.215180 | 0.216329 | 0.218566 | 0.222997 | 0.225569 | 0.224254 |
| left central | 0.244291 | 0.267981 | 0.292978 | 0.343951 | 0.349539 | 0.368137 | 0.347956 | 0.303785 | 0.266920 | 0.249937 | 0.239175 | 0.240441 | 0.252587 | 0.267179 | 0.273584 | 0.261922 | 0.254354 | 0.254926 | 0.271077 | 0.264894 | 0.267816 | 0.262919 | 0.258879 | 0.252856 | 0.266979 | 0.257648 | 0.245778 | 0.227759 | 0.216969 | 0.218009 | 0.234595 | 0.306098 | 0.435722 | 0.577497 | 0.534041 | 0.432168 | 0.296073 | 0.246038 | 0.236005 | 0.237618 | 0.248706 | 0.277696 | 0.319276 | 0.346169 | 0.365223 | 0.360686 | 0.333535 | 0.289694 | 0.256713 | 0.238164 | 0.243163 | 0.272760 | 0.322869 | 0.386858 | 0.456761 | 0.481450 | 0.483485 | 0.420574 | 0.359378 | 0.311502 | 0.283439 | 0.266076 | 0.269486 | 0.281966 | 0.316830 | 0.346254 | 0.364308 | 0.368538 | 0.319207 | 0.246961 | 0.215286 | 0.231926 | 0.282076 | 0.327843 | 0.341133 | 0.311268 | 0.280278 | 0.274028 | 0.291107 | 0.340150 | 0.459368 | 0.566666 | 0.656149 | 0.622998 | 0.503824 | 0.324267 | 0.226445 | 0.235664 | 0.374828 | 0.843833 | 1.876024 | 4.193973 | 6.014985 | 7.044177 | 5.123875 | 2.853695 | 2.107488 | 1.860091 | 2.276760 | 3.206824 | 4.510242 | 6.213432 | 8.698707 | 7.595016 | 6.582637 | 5.581762 | 3.005928 | 2.027483 | 1.943494 | 1.824295 | 1.877616 | 2.261221 | 2.451061 | 4.008432 | 7.163334 | 8.427934 | 5.914714 | 2.558592 | 0.739471 | 0.340933 | 0.242890 | 0.216361 | 0.217854 | 0.240690 | 0.270334 | 0.311274 | 0.336091 | 0.362133 | 0.386034 | 0.399327 | 0.368356 | 0.361956 | 0.342700 | 0.314170 | 0.287754 | 0.259023 | 0.233308 | 0.219801 | 0.214828 | 0.228638 | 0.279329 | 0.390892 | 0.638690 | 1.078755 | 1.671194 | 2.024350 | 1.801899 | 1.447614 | 1.315173 | 1.349018 | 1.400641 | 1.900334 | 2.994780 | 4.733358 | 7.347847 | 12.724966 | 25.122434 | 39.429115 | 42.668677 | 30.965285 | 18.540093 | 11.192044 | 7.956895 | 5.310958 | 4.254235 | 3.899346 | 3.336489 | 3.102044 | 3.043123 | 3.342122 | 4.217778 | 5.476079 | 7.279406 | 10.765469 | 13.329275 | 15.184742 | 21.711122 | 33.733108 | 50.723026 | 62.991810 | 53.320741 | 31.102578 | 15.977410 | 8.519381 | 4.328418 | 2.543258 | 1.539335 | 1.084347 | 0.881883 | 0.770856 | 0.714340 | 0.718522 | 0.861528 | 1.118498 | 1.674008 | 2.883885 | 5.276853 | 7.255542 | 6.378236 | 4.163779 | 1.984384 | 0.948195 | 0.565074 | 0.444505 | 0.381152 | 0.359613 | 0.360861 | 0.354768 | 0.341774 | 0.333973 | 0.324018 | 0.315449 | 0.322069 | 0.318709 | 0.327839 | 0.349626 | 0.386861 | 0.435824 | 0.507822 | 0.526831 | 0.523469 | 0.465186 | 0.397356 | 0.331158 | 0.293323 | 0.266917 | 0.256793 | 0.251858 | 0.253564 | 0.253052 | 0.246385 | 0.239033 | 0.237487 | 0.239415 | 0.255221 | 0.307922 | 0.444077 | 0.675005 | 0.919152 | 1.042854 | 1.027748 | 0.879597 | 0.696958 | 0.523880 | 0.423670 | 0.366699 | 0.341254 | 0.344552 | 0.386192 | 0.464114 | 0.609400 | 0.860880 | 1.256757 | 1.746708 | 1.987732 | 1.800134 | 1.454911 | 0.995157 | 0.811355 | 0.750614 | 0.783806 | 0.930836 | 1.336569 | 1.934127 | 2.836212 | 3.612031 | 3.898937 | 4.624031 | 5.631330 | 7.986387 | 14.098423 | 28.607652 | 42.675820 | 52.098718 | 42.121178 | 25.235098 | 12.828527 | 6.496915 | 4.356052 | 3.235695 |
| right central | 0.463858 | 0.420985 | 0.367902 | 0.299341 | 0.250461 | 0.217802 | 0.216322 | 0.223099 | 0.224500 | 0.220345 | 0.216095 | 0.215238 | 0.215934 | 0.217781 | 0.220352 | 0.221326 | 0.230107 | 0.255263 | 0.295536 | 0.298711 | 0.302619 | 0.307105 | 0.305029 | 0.303322 | 0.304437 | 0.305862 | 0.281732 | 0.251556 | 0.227308 | 0.222950 | 0.221531 | 0.228712 | 0.227095 | 0.241567 | 0.253657 | 0.233896 | 0.215974 | 0.218846 | 0.269221 | 0.336549 | 0.385591 | 0.458276 | 0.401627 | 0.275940 | 0.220681 | 0.221096 | 0.271391 | 0.331901 | 0.380001 | 0.345623 | 0.294680 | 0.250996 | 0.226832 | 0.214637 | 0.232945 | 0.326358 | 0.567999 | 1.014605 | 1.360062 | 1.162075 | 0.828752 | 0.559001 | 0.432357 | 0.384355 | 0.418518 | 0.547171 | 0.877989 | 1.556231 | 2.218261 | 1.575695 | 0.610298 | 0.252361 | 0.223259 | 0.295855 | 0.386897 | 0.438066 | 0.458590 | 0.493009 | 0.545007 | 0.628759 | 0.848527 | 1.177202 | 1.987193 | 4.018439 | 8.126456 | 10.742292 | 7.656238 | 3.213368 | 1.516786 | 0.768409 | 0.512858 | 0.387663 | 0.312938 | 0.261888 | 0.248637 | 0.237758 | 0.229803 | 0.234188 | 0.252442 | 0.273609 | 0.324150 | 0.396106 | 0.432226 | 0.409197 | 0.328812 | 0.259951 | 0.228327 | 0.219633 | 0.219067 | 0.226814 | 0.237973 | 0.256980 | 0.263164 | 0.252164 | 0.227960 | 0.215104 | 0.221007 | 0.239515 | 0.247239 | 0.243873 | 0.229681 | 0.218238 | 0.214642 | 0.219020 | 0.230221 | 0.243775 | 0.259228 | 0.273178 | 0.265390 | 0.254915 | 0.244740 | 0.243984 | 0.236470 | 0.227901 | 0.223360 | 0.222495 | 0.217881 | 0.216744 | 0.216200 | 0.215644 | 0.215247 | 0.215801 | 0.216387 | 0.219449 | 0.224483 | 0.232438 | 0.240208 | 0.250175 | 0.265670 | 0.284392 | 0.293467 | 0.283401 | 0.252322 | 0.222218 | 0.217363 | 0.262283 | 0.352703 | 0.443642 | 0.446884 | 0.361831 | 0.261648 | 0.217608 | 0.226459 | 0.273275 | 0.335884 | 0.374941 | 0.393690 | 0.378561 | 0.329817 | 0.274064 | 0.238130 | 0.219339 | 0.214817 | 0.214850 | 0.214884 | 0.214925 | 0.214714 | 0.215710 | 0.218882 | 0.223763 | 0.225933 | 0.226144 | 0.218277 | 0.214740 | 0.217195 | 0.228461 | 0.250241 | 0.280910 | 0.319249 | 0.349705 | 0.368637 | 0.368022 | 0.341430 | 0.317025 | 0.283739 | 0.259947 | 0.247112 | 0.236735 | 0.231455 | 0.226650 | 0.220489 | 0.215640 | 0.214661 | 0.216339 | 0.218499 | 0.223478 | 0.230232 | 0.237452 | 0.238992 | 0.235598 | 0.225710 | 0.218003 | 0.215098 | 0.214665 | 0.215037 | 0.216316 | 0.222134 | 0.234249 | 0.258580 | 0.268281 | 0.268387 | 0.257964 | 0.246908 | 0.226816 | 0.218602 | 0.215180 | 0.214857 | 0.214694 | 0.215346 | 0.218045 | 0.225675 | 0.234922 | 0.255235 | 0.284182 | 0.321060 | 0.373406 | 0.444476 | 0.562431 | 0.769072 | 1.117851 | 1.480922 | 1.506116 | 1.339248 | 1.082300 | 0.848681 | 0.605857 | 0.433610 | 0.336929 | 0.288890 | 0.251579 | 0.238224 | 0.239809 | 0.239367 | 0.240411 | 0.244034 | 0.244222 | 0.238877 | 0.228177 | 0.218234 | 0.214884 | 0.215176 | 0.216966 | 0.216124 | 0.215310 | 0.214742 | 0.214648 | 0.215223 | 0.221566 | 0.244267 | 0.297187 | 0.418296 | 0.685547 | 1.127452 | 1.772946 | 2.548652 | 3.258715 | 3.524314 | 3.447118 | 3.131019 | 2.803641 |
| left posterior | 0.914541 | 1.129299 | 1.549244 | 2.545278 | 2.750208 | 1.846818 | 0.855521 | 0.385996 | 0.231102 | 0.218289 | 0.248125 | 0.297431 | 0.339452 | 0.317449 | 0.267288 | 0.217115 | 0.274275 | 1.139162 | 4.312980 | 9.147725 | 10.616826 | 9.646037 | 6.466912 | 4.986340 | 1.781496 | 0.756865 | 0.409467 | 0.310263 | 0.248134 | 0.230287 | 0.220750 | 0.219432 | 0.217553 | 0.221116 | 0.221730 | 0.222519 | 0.219210 | 0.218769 | 0.222635 | 0.236509 | 0.256970 | 0.280832 | 0.293832 | 0.249538 | 0.215548 | 0.230200 | 0.266058 | 0.326000 | 0.366186 | 0.349362 | 0.285862 | 0.234514 | 0.214977 | 0.237755 | 0.283266 | 0.270904 | 0.231203 | 0.216188 | 0.272528 | 0.443521 | 0.516909 | 0.533437 | 0.545372 | 0.502973 | 0.460401 | 0.424709 | 0.424172 | 0.445997 | 0.670282 | 1.618504 | 6.232273 | 32.461551 | 160.394717 | 474.219441 | 743.069282 | 645.364916 | 259.127057 | 94.896103 | 22.678386 | 4.531062 | 1.335485 | 0.616863 | 0.402342 | 0.329715 | 0.281012 | 0.247209 | 0.232942 | 0.221110 | 0.216720 | 0.217270 | 0.220304 | 0.226224 | 0.239423 | 0.249704 | 0.251003 | 0.253827 | 0.249959 | 0.239486 | 0.230408 | 0.229634 | 0.227963 | 0.230326 | 0.241364 | 0.270314 | 0.309875 | 0.346856 | 0.348251 | 0.316042 | 0.255217 | 0.217894 | 0.219927 | 0.253626 | 0.318476 | 0.395466 | 0.443510 | 0.422919 | 0.343174 | 0.265036 | 0.220149 | 0.220164 | 0.258627 | 0.318001 | 0.361879 | 0.362441 | 0.319493 | 0.260850 | 0.226015 | 0.214631 | 0.225309 | 0.249219 | 0.251575 | 0.237330 | 0.220685 | 0.214632 | 0.221520 | 0.234945 | 0.245906 | 0.246206 | 0.236266 | 0.222327 | 0.215271 | 0.215396 | 0.217257 | 0.217189 | 0.215597 | 0.214630 | 0.215192 | 0.214704 | 0.216705 | 0.236481 | 0.284812 | 0.380013 | 0.520059 | 0.638985 | 0.733957 | 0.777373 | 0.738187 | 0.631442 | 0.537504 | 0.445911 | 0.402326 | 0.370328 | 0.345099 | 0.326235 | 0.317472 | 0.308229 | 0.324517 | 0.357036 | 0.404832 | 0.448638 | 0.517471 | 0.546969 | 0.532964 | 0.453632 | 0.374028 | 0.308858 | 0.270056 | 0.242276 | 0.227656 | 0.219215 | 0.215127 | 0.215665 | 0.222704 | 0.237158 | 0.259272 | 0.283678 | 0.298285 | 0.305116 | 0.304113 | 0.294402 | 0.283786 | 0.270393 | 0.262894 | 0.262555 | 0.266823 | 0.277269 | 0.305102 | 0.336321 | 0.379309 | 0.447066 | 0.531185 | 0.605196 | 0.654004 | 0.671956 | 0.624672 | 0.557055 | 0.477092 | 0.433094 | 0.399298 | 0.383395 | 0.375208 | 0.382999 | 0.400424 | 0.450862 | 0.514657 | 0.632395 | 0.804727 | 0.947064 | 0.916307 | 0.795386 | 0.627426 | 0.541362 | 0.496801 | 0.463146 | 0.457298 | 0.470621 | 0.446709 | 0.422034 | 0.382984 | 0.344647 | 0.309616 | 0.294908 | 0.289585 | 0.308906 | 0.353967 | 0.437426 | 0.535036 | 0.586427 | 0.517880 | 0.404292 | 0.323727 | 0.283597 | 0.271029 | 0.279307 | 0.309325 | 0.371450 | 0.482476 | 0.668953 | 0.926132 | 1.176208 | 1.396536 | 1.488678 | 1.275714 | 0.941491 | 0.667345 | 0.466864 | 0.338103 | 0.283589 | 0.256381 | 0.240981 | 0.230640 | 0.226043 | 0.222875 | 0.222627 | 0.223760 | 0.228741 | 0.240292 | 0.259726 | 0.288005 | 0.327185 | 0.370615 | 0.414246 | 0.459095 | 0.533668 | 0.673592 | 0.880437 | 1.099638 | 1.372720 | 1.484768 | 1.547732 |
| right posterior | 5.589167 | 3.391620 | 2.055447 | 0.897722 | 0.413514 | 0.265528 | 0.218897 | 0.216167 | 0.238944 | 0.259600 | 0.259182 | 0.238211 | 0.217888 | 0.219462 | 0.281258 | 0.392810 | 0.479831 | 0.425914 | 0.333804 | 0.230826 | 0.218375 | 0.283112 | 0.361870 | 0.365236 | 0.251923 | 0.215406 | 0.293182 | 0.488877 | 0.559182 | 0.367641 | 0.257480 | 0.218099 | 0.216377 | 0.226956 | 0.249122 | 0.262051 | 0.251234 | 0.231423 | 0.221817 | 0.214854 | 0.214813 | 0.216052 | 0.215908 | 0.215527 | 0.214688 | 0.214629 | 0.214785 | 0.216525 | 0.237052 | 0.289155 | 0.375600 | 0.478156 | 0.523689 | 0.527670 | 0.474880 | 0.442295 | 0.444821 | 0.510992 | 0.546419 | 0.484491 | 0.346784 | 0.256545 | 0.219762 | 0.215458 | 0.219229 | 0.218203 | 0.215216 | 0.214903 | 0.217117 | 0.242594 | 0.343846 | 0.564408 | 0.863111 | 1.087344 | 1.041906 | 0.813096 | 0.571858 | 0.378837 | 0.264249 | 0.218069 | 0.219148 | 0.236018 | 0.254055 | 0.269055 | 0.282952 | 0.315233 | 0.403227 | 0.527881 | 0.625057 | 0.657905 | 0.532649 | 0.407462 | 0.342263 | 0.332388 | 0.335165 | 0.382114 | 0.456112 | 0.579874 | 0.710570 | 0.802659 | 0.768364 | 0.682842 | 0.553542 | 0.416916 | 0.337373 | 0.311789 | 0.326893 | 0.370541 | 0.495702 | 0.778792 | 1.313113 | 2.172104 | 2.753491 | 2.123474 | 1.428123 | 0.877638 | 0.521595 | 0.353671 | 0.266019 | 0.227653 | 0.215976 | 0.214641 | 0.215195 | 0.214677 | 0.214749 | 0.217603 | 0.228881 | 0.251456 | 0.308133 | 0.464164 | 0.860421 | 2.236034 | 8.633074 | 29.310406 | 78.899108 | 120.960825 | 99.843480 | 60.728019 | 28.911668 | 12.267128 | 5.498967 | 2.668642 | 1.402861 | 0.862680 | 0.577374 | 0.418967 | 0.348813 | 0.321125 | 0.301454 | 0.301501 | 0.309680 | 0.346737 | 0.397418 | 0.478481 | 0.569813 | 0.631940 | 0.513876 | 0.393851 | 0.298751 | 0.249132 | 0.231119 | 0.228134 | 0.231467 | 0.241000 | 0.252928 | 0.264769 | 0.277597 | 0.298763 | 0.357877 | 0.476349 | 0.662007 | 0.954417 | 1.216039 | 1.178700 | 0.985866 | 0.781959 | 0.617123 | 0.558051 | 0.543048 | 0.553818 | 0.594177 | 0.522405 | 0.409493 | 0.319464 | 0.259973 | 0.233471 | 0.224355 | 0.219430 | 0.222641 | 0.234399 | 0.246643 | 0.264782 | 0.278350 | 0.269300 | 0.254684 | 0.243021 | 0.230761 | 0.224479 | 0.218610 | 0.215089 | 0.215459 | 0.220623 | 0.231470 | 0.244013 | 0.261942 | 0.287144 | 0.332772 | 0.404700 | 0.518207 | 0.641099 | 0.701981 | 0.627801 | 0.477640 | 0.353411 | 0.273758 | 0.232065 | 0.216955 | 0.214627 | 0.214701 | 0.215868 | 0.227649 | 0.255387 | 0.286567 | 0.296146 | 0.284244 | 0.260103 | 0.234981 | 0.221478 | 0.215617 | 0.214941 | 0.216088 | 0.215230 | 0.215738 | 0.226760 | 0.254584 | 0.324488 | 0.456893 | 0.599421 | 0.745040 | 0.847859 | 0.841510 | 0.818139 | 0.766534 | 0.757335 | 0.831980 | 0.856066 | 0.923946 | 1.108348 | 1.156424 | 1.035275 | 0.826038 | 0.573094 | 0.392836 | 0.291459 | 0.241990 | 0.223733 | 0.216105 | 0.215031 | 0.219375 | 0.225952 | 0.233010 | 0.240545 | 0.247249 | 0.259502 | 0.276533 | 0.309992 | 0.360296 | 0.404303 | 0.416234 | 0.410906 | 0.365080 | 0.312334 | 0.268844 | 0.234537 | 0.217716 | 0.214627 | 0.216158 | 0.218268 | 0.218947 | 0.219031 |
| all electrodes | 0.256422 | 0.248999 | 0.230078 | 0.214644 | 0.221997 | 0.237004 | 0.277166 | 0.310003 | 0.367929 | 0.562799 | 0.937279 | 1.498764 | 1.993686 | 2.208460 | 1.796325 | 1.605176 | 1.447364 | 1.248680 | 1.061843 | 0.800492 | 0.530018 | 0.443968 | 0.461454 | 0.471101 | 0.595094 | 1.055434 | 2.350675 | 2.964746 | 2.941809 | 2.453378 | 2.143189 | 1.538729 | 1.104695 | 0.622769 | 0.407261 | 0.293832 | 0.245887 | 0.224731 | 0.215600 | 0.223102 | 0.268205 | 0.375830 | 0.577494 | 0.897903 | 0.856195 | 0.591683 | 0.342585 | 0.239977 | 0.215085 | 0.215775 | 0.215030 | 0.217451 | 0.256708 | 0.366206 | 0.550298 | 0.703850 | 0.630924 | 0.424599 | 0.305446 | 0.236499 | 0.215706 | 0.216959 | 0.224040 | 0.227324 | 0.223619 | 0.217487 | 0.216791 | 0.223801 | 0.260698 | 0.379644 | 0.785579 | 1.810828 | 3.531981 | 4.482366 | 4.073770 | 2.317219 | 1.234361 | 0.770974 | 0.555360 | 0.527658 | 0.641810 | 0.857488 | 1.054053 | 1.147102 | 1.142893 | 1.139424 | 0.947945 | 0.583784 | 0.342270 | 0.249043 | 0.218869 | 0.214861 | 0.222138 | 0.228338 | 0.234114 | 0.238869 | 0.256982 | 0.303661 | 0.398858 | 0.550395 | 0.781451 | 0.890236 | 0.751398 | 0.541835 | 0.396205 | 0.298106 | 0.251131 | 0.229517 | 0.220158 | 0.216934 | 0.216453 | 0.215353 | 0.214631 | 0.215029 | 0.224333 | 0.248575 | 0.285856 | 0.351273 | 0.511750 | 0.659330 | 0.866475 | 0.960039 | 0.874045 | 0.587957 | 0.398400 | 0.276982 | 0.225093 | 0.216026 | 0.233479 | 0.258902 | 0.259247 | 0.242279 | 0.224200 | 0.215973 | 0.215282 | 0.218075 | 0.218062 | 0.215679 | 0.214678 | 0.222438 | 0.248642 | 0.293505 | 0.367107 | 0.477329 | 0.573424 | 0.599943 | 0.516457 | 0.423946 | 0.367461 | 0.344822 | 0.361279 | 0.460437 | 0.728065 | 1.285383 | 1.979926 | 2.471249 | 2.436610 | 1.919173 | 1.364905 | 1.152568 | 1.076844 | 1.116381 | 1.102685 | 1.069617 | 0.944480 | 0.784365 | 0.647226 | 0.583639 | 0.598438 | 0.717563 | 1.054120 | 1.916251 | 4.406138 | 9.803767 | 17.316721 | 22.398172 | 22.818877 | 15.476420 | 10.731954 | 5.770173 | 2.624658 | 1.176773 | 0.781495 | 0.621377 | 0.586746 | 0.619792 | 0.752477 | 0.952250 | 1.134765 | 1.250897 | 1.349542 | 1.330177 | 1.142189 | 0.988573 | 0.786720 | 0.594568 | 0.474101 | 0.377645 | 0.296096 | 0.262316 | 0.248937 | 0.240220 | 0.232330 | 0.226630 | 0.223754 | 0.223863 | 0.222422 | 0.218221 | 0.215753 | 0.214651 | 0.214924 | 0.215041 | 0.214726 | 0.217937 | 0.229153 | 0.243671 | 0.255014 | 0.253087 | 0.242050 | 0.225319 | 0.217168 | 0.214676 | 0.216730 | 0.220716 | 0.226647 | 0.240702 | 0.255710 | 0.280288 | 0.300779 | 0.319951 | 0.342778 | 0.376526 | 0.420703 | 0.513494 | 0.644586 | 0.796944 | 0.884411 | 0.818682 | 0.728248 | 0.635103 | 0.565053 | 0.517166 | 0.492035 | 0.471066 | 0.471899 | 0.472931 | 0.497256 | 0.506900 | 0.502955 | 0.467896 | 0.396283 | 0.313441 | 0.258862 | 0.225061 | 0.214976 | 0.217873 | 0.237164 | 0.267420 | 0.307765 | 0.352689 | 0.378366 | 0.341841 | 0.287235 | 0.242387 | 0.220665 | 0.214628 | 0.216953 | 0.219819 | 0.218414 | 0.215329 | 0.215514 | 0.223147 | 0.239114 | 0.250374 | 0.243855 | 0.234362 | 0.225124 | 0.222751 | 0.221826 | 0.224257 |
