## Supplementary Table for "Shared Neural Dynamics of Facial Expression Processing": SupplementaryTable_2.html

  


### Supplementary Table 2.

  

Classification analyses

  

**Classification analyses.** The procedure followed a leave-one participant-out scheme. For identity classification, classifiers underwent iterative training on three emotion categories,
successively tested on the emotion category left out. In the case of sex and emotion classifiers, training was conducted on six distinct identities (three male and three female) and subsequently tested on one
identity left out. The classification of pairs of emotional expressions followed a similar logic, with the distinction that only two emotional expressions were included at a time in the classification procedure.
The training involved six identities (three male and three female) and was subsequently tested on one identity that was omitted during training. Classification accuracies were tested against chance levels.
  
Classification accuracies underwent two-sided, one-sample cluster permutation tests (10,000 iterations) against chance. The Bayesian tests employed a two-sided approach,
utilizing a non-directional whole-Cauchy prior with a medium width (r = 0.707) and excluding an interval from δ = 0.5 to +0.5.

  


**A** identity

**B** sex

**C** emotion

**D** neutral vs happy

**E** neutral vs angry

**F** neutral vs sad

**G** happy vs angry

**H** angry vs sad

**I** happy vs sad

A) identity

  
|  | time window | peak latency | cluster *p* | peak Cohen's *d* |  | | | |  |
| **all electrodes** | 95 - 510 ms | 115 ms | 0.0001 | 1.6927 |  | | | |  |
 550 - 675 ms | 575 ms | 0.0194 | 0.7775 |  | | | ||  |  | | | | | | | | |

Time-resolved classification, cluster permutation tests

|  | **left hemisphere** | | | | **right hemisphere** | | | |  |
|  | time window | peak latency | cluster *p* | peak Cohen's *d* | time window | peak latency | cluster *p* | peak Cohen's *d* |  |
| **anterior** |  | | | | 140 - 310 ms | 260 ms | 0.0023 | 1.0066 |  |
| **central** |  | | | | 180 - 270 ms | 205 ms | 0.0356 | 0.7062 |  |
| **posterior** | 185 - 355 ms | 210 ms | 0.0039 | 1.1439 | 60 - 890 ms | 110 ms | 0.0001 | 1.2808 |  |

  

Time-resolved classification, Bayesian statistics

|  | -200 | -195 | -190 | -185 | -180 | -175 | -170 | -165 | -160 | -155 | -150 | -145 | -140 | -135 | -130 | -125 | -120 | -115 | -110 | -105 | -100 | -95 | -90 | -85 | -80 | -75 | -70 | -65 | -60 | -55 | -50 | -45 | -40 | -35 | -30 | -25 | -20 | -15 | -10 | -5 | 0 | 5 | 10 | 15 | 20 | 25 | 30 | 35 | 40 | 45 | 50 | 55 | 60 | 65 | 70 | 75 | 80 | 85 | 90 | 95 | 100 | 105 | 110 | 115 | 120 | 125 | 130 | 135 | 140 | 145 | 150 | 155 | 160 | 165 | 170 | 175 | 180 | 185 | 190 | 195 | 200 | 205 | 210 | 215 | 220 | 225 | 230 | 235 | 240 | 245 | 250 | 255 | 260 | 265 | 270 | 275 | 280 | 285 | 290 | 295 | 300 | 305 | 310 | 315 | 320 | 325 | 330 | 335 | 340 | 345 | 350 | 355 | 360 | 365 | 370 | 375 | 380 | 385 | 390 | 395 | 400 | 405 | 410 | 415 | 420 | 425 | 430 | 435 | 440 | 445 | 450 | 455 | 460 | 465 | 470 | 475 | 480 | 485 | 490 | 495 | 500 | 505 | 510 | 515 | 520 | 525 | 530 | 535 | 540 | 545 | 550 | 555 | 560 | 565 | 570 | 575 | 580 | 585 | 590 | 595 | 600 | 605 | 610 | 615 | 620 | 625 | 630 | 635 | 640 | 645 | 650 | 655 | 660 | 665 | 670 | 675 | 680 | 685 | 690 | 695 | 700 | 705 | 710 | 715 | 720 | 725 | 730 | 735 | 740 | 745 | 750 | 755 | 760 | 765 | 770 | 775 | 780 | 785 | 790 | 795 | 800 | 805 | 810 | 815 | 820 | 825 | 830 | 835 | 840 | 845 | 850 | 855 | 860 | 865 | 870 | 875 | 880 | 885 | 890 | 895 | 900 | 905 | 910 | 915 | 920 | 925 | 930 | 935 | 940 | 945 | 950 | 955 | 960 | 965 | 970 | 975 | 980 | 985 | 990 | 995 | 1000 | 1005 | 1010 | 1015 | 1020 | 1025 | 1030 | 1035 | 1040 | 1045 | 1050 | 1055 | 1060 | 1065 | 1070 | 1075 | 1080 | 1085 | 1090 | 1095 | 1100 | 1105 | 1110 | 1115 | 1120 | 1125 | 1130 | 1135 | 1140 | 1145 | 1150 | 1155 | 1160 | 1165 | 1170 | 1175 | 1180 | 1185 | 1190 | 1195 |
| --- | --- | --- | --- | --- | --- | --- | --- | --- | --- | --- | --- | --- | --- | --- | --- | --- | --- | --- | --- | --- | --- | --- | --- | --- | --- | --- | --- | --- | --- | --- | --- | --- | --- | --- | --- | --- | --- | --- | --- | --- | --- | --- | --- | --- | --- | --- | --- | --- | --- | --- | --- | --- | --- | --- | --- | --- | --- | --- | --- | --- | --- | --- | --- | --- | --- | --- | --- | --- | --- | --- | --- | --- | --- | --- | --- | --- | --- | --- | --- | --- | --- | --- | --- | --- | --- | --- | --- | --- | --- | --- | --- | --- | --- | --- | --- | --- | --- | --- | --- | --- | --- | --- | --- | --- | --- | --- | --- | --- | --- | --- | --- | --- | --- | --- | --- | --- | --- | --- | --- | --- | --- | --- | --- | --- | --- | --- | --- | --- | --- | --- | --- | --- | --- | --- | --- | --- | --- | --- | --- | --- | --- | --- | --- | --- | --- | --- | --- | --- | --- | --- | --- | --- | --- | --- | --- | --- | --- | --- | --- | --- | --- | --- | --- | --- | --- | --- | --- | --- | --- | --- | --- | --- | --- | --- | --- | --- | --- | --- | --- | --- | --- | --- | --- | --- | --- | --- | --- | --- | --- | --- | --- | --- | --- | --- | --- | --- | --- | --- | --- | --- | --- | --- | --- | --- | --- | --- | --- | --- | --- | --- | --- | --- | --- | --- | --- | --- | --- | --- | --- | --- | --- | --- | --- | --- | --- | --- | --- | --- | --- | --- | --- | --- | --- | --- | --- | --- | --- | --- | --- | --- | --- | --- | --- | --- | --- | --- | --- | --- | --- | --- | --- | --- | --- | --- | --- | --- | --- | --- | --- | --- | --- | --- | --- | --- | --- | --- | --- | --- | --- | --- | --- | --- | --- | --- | --- | --- | --- | --- | --- | --- |
| left anterior | 0.422665 | 0.238230 | 0.231634 | 0.248304 | 0.249978 | 0.442883 | 1.828934 | 2.192938 | 3.324022 | 1.463758 | 1.356918 | 2.128532 | 0.630617 | 0.330205 | 0.292559 | 0.220815 | 0.221395 | 0.218541 | 0.224450 | 0.215374 | 0.262396 | 0.272161 | 0.228304 | 0.317704 | 0.765852 | 0.611639 | 0.702297 | 0.489678 | 0.343214 | 0.233719 | 0.343044 | 1.800626 | 16.446061 | 55.653136 | 202.217658 | 92.068456 | 15.959999 | 3.052434 | 1.386554 | 0.410140 | 0.275858 | 0.218130 | 0.247283 | 0.257593 | 0.273787 | 0.356717 | 0.308532 | 0.412432 | 0.394909 | 0.362581 | 0.373164 | 0.277457 | 0.221408 | 0.215358 | 0.228604 | 0.249889 | 0.218362 | 0.221379 | 0.216766 | 0.244204 | 0.318474 | 0.520544 | 1.197489 | 1.487378 | 1.364207 | 1.626067 | 0.901923 | 0.750148 | 0.373363 | 0.252579 | 0.229921 | 0.253291 | 0.252858 | 0.222813 | 0.215150 | 0.221676 | 0.216512 | 0.215424 | 0.228309 | 0.215177 | 0.284113 | 0.463568 | 0.636819 | 1.150219 | 1.476257 | 1.189142 | 1.341425 | 1.529611 | 2.077331 | 1.485395 | 0.819997 | 0.652359 | 0.694253 | 0.561840 | 0.413858 | 0.326130 | 0.311212 | 0.337288 | 0.412391 | 0.327482 | 0.261237 | 0.237829 | 0.241178 | 0.218068 | 0.214724 | 0.219219 | 0.216241 | 0.216225 | 0.225893 | 0.243801 | 0.241974 | 0.251269 | 0.220862 | 0.219089 | 0.229936 | 0.224711 | 0.224906 | 0.224436 | 0.229187 | 0.308906 | 0.332818 | 0.275982 | 0.262456 | 0.249109 | 0.239127 | 0.231417 | 0.217627 | 0.216281 | 0.222624 | 0.224431 | 0.227105 | 0.220031 | 0.215051 | 0.215319 | 0.251352 | 0.264154 | 0.253046 | 0.283381 | 0.309247 | 0.300316 | 0.275717 | 0.225381 | 0.223083 | 0.220145 | 0.216925 | 0.236454 | 0.254627 | 0.290237 | 0.429633 | 0.567830 | 0.462716 | 0.342603 | 0.287748 | 0.267041 | 0.229142 | 0.214850 | 0.215456 | 0.214660 | 0.217939 | 0.238977 | 0.253175 | 0.224859 | 0.229481 | 0.222337 | 0.216248 | 0.215320 | 0.217683 | 0.223837 | 0.215640 | 0.214637 | 0.214670 | 0.215765 | 0.215416 | 0.224381 | 0.217522 | 0.215198 | 0.214699 | 0.214699 | 0.215418 | 0.220212 | 0.219173 | 0.222078 | 0.231268 | 0.223457 | 0.214863 | 0.231015 | 0.241706 | 0.257613 | 0.238774 | 0.225358 | 0.221262 | 0.217580 | 0.215425 | 0.220162 | 0.218323 | 0.216811 | 0.218654 | 0.218370 | 0.214926 | 0.217763 | 0.219523 | 0.214675 | 0.231111 | 0.262864 | 0.258579 | 0.235408 | 0.222387 | 0.215149 | 0.226857 | 0.285004 | 0.325521 | 0.273980 | 0.225727 | 0.215928 | 0.215284 | 0.223375 | 0.235695 | 0.240775 | 0.216169 | 0.229273 | 0.294399 | 0.281250 | 0.288681 | 0.260224 | 0.256295 | 0.231033 | 0.214784 | 0.236781 | 0.292454 | 0.408957 | 0.419469 | 0.327332 | 0.258131 | 0.245741 | 0.230291 | 0.217687 | 0.214707 | 0.214635 | 0.214826 | 0.223415 | 0.220972 | 0.226564 | 0.240003 | 0.229042 | 0.248723 | 0.271088 | 0.279110 | 0.284530 | 0.235974 | 0.230086 | 0.235382 | 0.220352 | 0.222173 | 0.218642 | 0.235520 | 0.265439 | 0.279030 | 0.284881 | 0.261449 | 0.259697 | 0.264073 | 0.253106 | 0.244459 | 0.234783 | 0.233728 | 0.233401 | 0.215047 | 0.228058 | 0.285957 | 0.297730 | 0.313540 | 0.312566 | 0.263716 | 0.220497 | 0.214657 | 0.219041 | 0.219598 | 0.222640 | 0.221297 | 0.220340 |
| right anterior | 0.302608 | 0.433301 | 0.229021 | 0.234927 | 0.281663 | 0.255129 | 0.351029 | 0.919087 | 2.737477 | 1.077835 | 0.453017 | 0.219147 | 0.215687 | 0.222114 | 0.229468 | 0.270308 | 0.251386 | 0.214761 | 0.220385 | 0.319087 | 0.348328 | 0.314037 | 0.369958 | 0.445093 | 0.221108 | 0.308444 | 1.113252 | 3.383627 | 1.128854 | 0.433702 | 0.394945 | 0.325383 | 0.242272 | 0.215559 | 0.246761 | 0.252600 | 0.295636 | 0.297209 | 0.366480 | 0.546713 | 0.761147 | 0.404042 | 0.257831 | 0.224740 | 0.267237 | 0.337482 | 1.616558 | 29.120078 | 103.773888 | 16.813480 | 2.710090 | 2.030412 | 1.404023 | 0.922622 | 0.452497 | 0.337056 | 0.479770 | 0.744575 | 0.564123 | 0.880119 | 0.748977 | 0.719606 | 0.733007 | 0.571450 | 0.518058 | 0.662669 | 0.748306 | 1.187562 | 1.747797 | 3.063102 | 6.123651 | 7.887215 | 11.980098 | 24.952611 | 32.992445 | 33.878672 | 16.321073 | 50.696142 | 59.046848 | 37.562363 | 18.680791 | 14.891255 | 14.614746 | 12.342901 | 3.447165 | 4.372317 | 13.331758 | 55.227157 | 64.103242 | 64.125289 | 85.746739 | 120.767564 | 460.712360 | 373.986528 | 115.483001 | 84.535725 | 106.789002 | 117.791465 | 60.494978 | 12.258462 | 6.400642 | 2.953442 | 1.302614 | 0.842983 | 0.539849 | 0.469812 | 0.399017 | 0.449431 | 0.535607 | 0.628870 | 0.553884 | 0.444001 | 0.342665 | 0.375356 | 0.258635 | 0.216561 | 0.219849 | 0.221048 | 0.236883 | 0.225095 | 0.222085 | 0.218335 | 0.243816 | 0.255752 | 0.260202 | 0.298440 | 0.309181 | 0.276585 | 0.226854 | 0.214673 | 0.214804 | 0.214626 | 0.220939 | 0.219728 | 0.215182 | 0.218540 | 0.216649 | 0.217383 | 0.228363 | 0.231738 | 0.250931 | 0.248141 | 0.236698 | 0.239955 | 0.277104 | 0.306497 | 0.298736 | 0.274945 | 0.355295 | 0.505348 | 0.820574 | 0.671610 | 0.597077 | 0.492734 | 0.588501 | 0.574016 | 0.517593 | 0.433394 | 0.470375 | 0.458144 | 0.352974 | 0.299540 | 0.261876 | 0.298387 | 0.293992 | 0.310724 | 0.335218 | 0.364076 | 0.411609 | 0.456191 | 0.331096 | 0.289076 | 0.294141 | 0.257479 | 0.291187 | 0.294056 | 0.292754 | 0.258449 | 0.246872 | 0.231452 | 0.218970 | 0.214626 | 0.218211 | 0.214626 | 0.244391 | 0.288780 | 0.257751 | 0.324180 | 0.401524 | 0.430351 | 0.341068 | 0.280736 | 0.317561 | 0.439263 | 0.450419 | 0.444436 | 0.435584 | 0.532796 | 0.648922 | 0.454838 | 0.312415 | 0.251849 | 0.222909 | 0.217788 | 0.215026 | 0.214668 | 0.214714 | 0.214636 | 0.215558 | 0.222835 | 0.271953 | 0.346478 | 0.418866 | 0.431253 | 0.383209 | 0.439205 | 0.498198 | 0.417527 | 0.368769 | 0.402832 | 0.399900 | 0.678159 | 0.923438 | 0.990199 | 1.450258 | 0.843084 | 0.531771 | 0.623174 | 0.524464 | 0.406518 | 0.436189 | 0.386284 | 0.499058 | 0.500679 | 0.525721 | 0.639184 | 0.781588 | 1.076392 | 0.929007 | 0.977283 | 1.687024 | 2.642395 | 3.827281 | 3.082470 | 1.667914 | 1.247153 | 0.716179 | 0.423402 | 0.297798 | 0.237423 | 0.214698 | 0.225442 | 0.234374 | 0.224670 | 0.217072 | 0.214679 | 0.215007 | 0.226932 | 0.252020 | 0.284362 | 0.263256 | 0.221840 | 0.214636 | 0.214726 | 0.215984 | 0.217438 | 0.219005 | 0.239264 | 0.267775 | 0.256471 | 0.227947 | 0.220218 | 0.217651 | 0.236676 | 0.235261 | 0.228098 | 0.215832 | 0.214745 | 0.215306 | 0.226869 |
| left central | 0.731542 | 0.504865 | 0.481677 | 0.508616 | 0.580094 | 0.736616 | 0.506850 | 0.380477 | 0.437254 | 0.295778 | 0.234146 | 0.331948 | 0.457130 | 0.402481 | 0.419337 | 0.422148 | 0.644515 | 0.291241 | 0.214626 | 0.235636 | 0.226755 | 0.237989 | 0.224508 | 0.273147 | 0.238010 | 0.219154 | 0.284259 | 0.525236 | 1.779259 | 3.607146 | 3.679820 | 1.278777 | 1.049919 | 0.852907 | 0.316960 | 0.218979 | 0.247328 | 0.270689 | 0.321684 | 0.377654 | 0.291217 | 0.225027 | 0.214643 | 0.220718 | 0.216365 | 0.214962 | 0.218203 | 0.256193 | 0.265169 | 0.276945 | 0.254508 | 0.227787 | 0.217703 | 0.216134 | 0.263578 | 0.266788 | 0.331740 | 0.380782 | 0.358854 | 0.397139 | 0.331359 | 0.222606 | 0.226138 | 0.345792 | 0.526324 | 0.520446 | 0.469983 | 0.537848 | 0.484856 | 0.365036 | 0.287438 | 0.259274 | 0.323738 | 0.429957 | 0.332175 | 0.294938 | 0.353748 | 0.380929 | 0.400918 | 0.444546 | 0.546158 | 0.715435 | 0.623225 | 0.596026 | 0.634343 | 0.636399 | 0.636021 | 0.601305 | 0.601900 | 0.739054 | 0.727783 | 1.123142 | 2.067681 | 3.328841 | 1.868687 | 1.836909 | 1.633517 | 1.886529 | 1.730018 | 1.769286 | 1.675584 | 1.265723 | 0.803503 | 0.741266 | 0.471933 | 0.298601 | 0.232612 | 0.225868 | 0.229985 | 0.225418 | 0.215586 | 0.225726 | 0.222645 | 0.223603 | 0.243956 | 0.222203 | 0.216030 | 0.215650 | 0.219667 | 0.224351 | 0.228135 | 0.224870 | 0.214939 | 0.219557 | 0.217197 | 0.216662 | 0.216492 | 0.215603 | 0.261177 | 0.415547 | 0.677418 | 0.913766 | 0.870309 | 0.713536 | 0.411567 | 0.285527 | 0.231857 | 0.215363 | 0.221250 | 0.236171 | 0.233002 | 0.230606 | 0.250056 | 0.322451 | 0.467708 | 0.435034 | 0.399673 | 0.362617 | 0.371971 | 0.282792 | 0.235321 | 0.217248 | 0.224197 | 0.281917 | 0.325890 | 0.472032 | 0.428776 | 0.470767 | 0.678809 | 0.608893 | 0.481710 | 0.429378 | 0.357208 | 0.356502 | 0.393264 | 0.394828 | 0.292831 | 0.254785 | 0.233148 | 0.226633 | 0.219718 | 0.218973 | 0.247772 | 0.248095 | 0.226780 | 0.214639 | 0.217075 | 0.224150 | 0.223873 | 0.220757 | 0.216243 | 0.219294 | 0.238529 | 0.392959 | 1.022869 | 1.418135 | 1.210397 | 1.565959 | 5.470998 | 13.212394 | 14.039941 | 4.107433 | 1.792766 | 1.036570 | 0.942569 | 0.807884 | 0.819337 | 0.550191 | 0.473307 | 0.608787 | 1.053193 | 0.922245 | 0.696104 | 1.001169 | 1.085386 | 1.446717 | 1.190508 | 0.628137 | 0.507547 | 0.450648 | 0.353925 | 0.271544 | 0.239883 | 0.228223 | 0.252366 | 0.245154 | 0.250689 | 0.250112 | 0.234485 | 0.215862 | 0.215203 | 0.215583 | 0.226411 | 0.233959 | 0.234009 | 0.283073 | 0.468457 | 0.556640 | 0.434200 | 0.341447 | 0.304569 | 0.324261 | 0.286161 | 0.270161 | 0.306451 | 0.468063 | 0.711099 | 0.863332 | 0.621313 | 0.974405 | 1.315167 | 1.335189 | 0.617101 | 0.347411 | 0.276764 | 0.306949 | 0.345741 | 0.465201 | 0.497209 | 0.688657 | 0.871977 | 1.135443 | 0.738946 | 0.507119 | 0.417797 | 0.404318 | 0.366769 | 0.380738 | 0.439808 | 0.672724 | 0.680738 | 0.871008 | 0.724152 | 0.714362 | 0.687408 | 0.540032 | 0.379917 | 0.282339 | 0.230186 | 0.235487 | 0.239305 | 0.244285 | 0.235653 | 0.255066 | 0.345356 | 0.390973 | 0.397037 | 0.438035 | 0.490740 | 0.547467 |
| right central | 0.218533 | 0.252865 | 0.265108 | 0.255131 | 0.225955 | 0.217759 | 0.216258 | 0.244287 | 0.293036 | 0.555565 | 0.878422 | 6.514028 | 91.863565 | 32.459270 | 1.354358 | 0.551694 | 0.421983 | 0.432955 | 0.258303 | 0.218571 | 0.225381 | 0.214699 | 0.255023 | 0.337144 | 0.290918 | 0.470306 | 1.332372 | 17.406640 | 136.077229 | 262.231297 | 25.168572 | 37.417734 | 22.427941 | 1.312147 | 0.303923 | 0.220939 | 0.324091 | 0.641858 | 0.932527 | 0.636596 | 0.332864 | 0.242137 | 0.225650 | 0.214803 | 0.216113 | 0.216092 | 0.236279 | 0.349390 | 0.579900 | 0.760880 | 0.928723 | 0.449780 | 0.285936 | 0.245999 | 0.214767 | 0.234946 | 0.244245 | 0.226578 | 0.215102 | 0.231913 | 0.380571 | 1.012699 | 3.696726 | 7.769919 | 11.260947 | 17.552064 | 7.646451 | 4.479654 | 1.952373 | 1.018150 | 1.014675 | 1.559249 | 1.473513 | 0.942553 | 0.558085 | 1.011552 | 2.293904 | 2.801206 | 2.483287 | 6.310908 | 10.688323 | 18.156623 | 5.654863 | 2.741623 | 1.767973 | 1.762875 | 2.717232 | 9.875128 | 20.969413 | 11.979044 | 7.152025 | 4.403405 | 4.776144 | 5.824000 | 2.965419 | 1.255129 | 0.834859 | 0.694352 | 1.230395 | 1.807734 | 1.177889 | 0.594412 | 0.506801 | 0.381104 | 0.306786 | 0.238548 | 0.215620 | 0.232351 | 0.229808 | 0.224827 | 0.224332 | 0.214690 | 0.234262 | 0.271043 | 0.256390 | 0.236775 | 0.231727 | 0.234064 | 0.217005 | 0.217997 | 0.215290 | 0.216933 | 0.231810 | 0.233048 | 0.244068 | 0.273437 | 0.259412 | 0.235751 | 0.233316 | 0.233102 | 0.233668 | 0.230899 | 0.227808 | 0.256802 | 0.259818 | 0.230908 | 0.216136 | 0.214636 | 0.220916 | 0.217094 | 0.215907 | 0.216695 | 0.221775 | 0.230970 | 0.257508 | 0.290329 | 0.356918 | 0.371693 | 0.370025 | 0.267655 | 0.215182 | 0.234482 | 0.289254 | 0.292685 | 0.272053 | 0.261851 | 0.259860 | 0.242843 | 0.216191 | 0.215804 | 0.214626 | 0.214626 | 0.215207 | 0.215286 | 0.216103 | 0.246133 | 0.303562 | 0.343801 | 0.415560 | 0.427808 | 0.404038 | 0.307010 | 0.219816 | 0.219517 | 0.247630 | 0.278098 | 0.260791 | 0.250160 | 0.276560 | 0.263674 | 0.251430 | 0.250303 | 0.244571 | 0.221396 | 0.222283 | 0.319873 | 0.366046 | 0.374681 | 0.405226 | 0.452680 | 0.457644 | 0.361359 | 0.309480 | 0.255437 | 0.219588 | 0.216333 | 0.215404 | 0.216125 | 0.229632 | 0.245579 | 0.232792 | 0.219318 | 0.224803 | 0.217292 | 0.214811 | 0.229541 | 0.233313 | 0.238019 | 0.238394 | 0.226069 | 0.214832 | 0.226616 | 0.250213 | 0.264432 | 0.304075 | 0.305183 | 0.262731 | 0.225359 | 0.214671 | 0.215730 | 0.215668 | 0.218189 | 0.232730 | 0.247907 | 0.277237 | 0.334025 | 0.371781 | 0.393979 | 0.504427 | 0.460965 | 0.469185 | 0.490118 | 0.464289 | 0.353570 | 0.326875 | 0.256440 | 0.241161 | 0.257563 | 0.247190 | 0.247830 | 0.249967 | 0.256263 | 0.248554 | 0.236737 | 0.217592 | 0.214881 | 0.214626 | 0.214808 | 0.217279 | 0.231190 | 0.261337 | 0.274322 | 0.264434 | 0.259742 | 0.276003 | 0.263055 | 0.243647 | 0.220441 | 0.214728 | 0.224931 | 0.246562 | 0.337162 | 0.496655 | 0.666464 | 0.612541 | 0.583065 | 0.412643 | 0.270064 | 0.227050 | 0.216023 | 0.215123 | 0.219814 | 0.264178 | 0.339647 | 0.614966 | 1.881635 | 5.288924 | 7.780059 | 4.083991 | 1.308529 |
| left posterior | 0.384579 | 0.327123 | 0.368226 | 0.297165 | 0.287709 | 0.218961 | 0.217309 | 0.216163 | 0.266148 | 0.375281 | 0.409371 | 0.497767 | 0.516751 | 0.346452 | 0.219893 | 0.258355 | 0.403105 | 0.436899 | 0.393098 | 0.303130 | 0.330830 | 0.244764 | 0.214860 | 0.215595 | 0.224001 | 0.238171 | 0.300293 | 1.054949 | 3.370077 | 6.865160 | 5.028418 | 6.663753 | 2.621504 | 0.564058 | 0.216271 | 0.219555 | 0.219120 | 0.214832 | 0.216302 | 0.232930 | 0.217208 | 0.214676 | 0.215662 | 0.232537 | 0.253551 | 0.334097 | 0.375381 | 0.421492 | 0.320171 | 0.329370 | 0.263839 | 0.220858 | 0.226609 | 0.257133 | 0.276012 | 0.247502 | 0.229678 | 0.215476 | 0.246278 | 0.701438 | 5.875765 | 63.384966 | 773.416435 | 968.107982 | 934.680543 | 775.332981 | 929.819562 | 207.869744 | 18.669143 | 1.218887 | 0.469957 | 0.294600 | 0.224548 | 0.215321 | 0.230229 | 0.335076 | 1.205958 | 9.403703 | 94.100567 | 742.837872 | 4136.246721 | 2158.583590 | 2065.005564 | 1263.188688 | 746.802205 | 425.303647 | 141.814502 | 56.031479 | 55.954136 | 45.092722 | 42.298047 | 29.722953 | 17.605066 | 8.777458 | 7.394124 | 6.074251 | 5.519453 | 3.079160 | 2.616799 | 2.796860 | 6.793359 | 6.690561 | 10.600375 | 15.009409 | 19.310378 | 23.786041 | 10.062103 | 3.237816 | 3.082109 | 2.268254 | 1.573952 | 1.353622 | 0.923483 | 0.675581 | 0.495635 | 0.400336 | 0.474921 | 0.622629 | 0.559026 | 0.633951 | 0.875777 | 1.239331 | 2.154819 | 1.763651 | 0.932437 | 0.668559 | 0.491188 | 0.456992 | 0.446746 | 0.332796 | 0.340494 | 0.369418 | 0.449322 | 0.537338 | 0.576428 | 0.626794 | 0.896607 | 0.796467 | 0.873661 | 1.174995 | 0.732174 | 0.496103 | 0.393619 | 0.358604 | 0.351561 | 0.361334 | 0.280618 | 0.301771 | 0.325263 | 0.346946 | 0.365656 | 0.408246 | 0.521639 | 0.699249 | 0.660231 | 0.515340 | 0.553821 | 0.599520 | 0.682402 | 0.835492 | 0.805867 | 0.865280 | 0.727714 | 0.554231 | 0.448773 | 0.418739 | 0.375270 | 0.433297 | 0.486906 | 0.564589 | 0.515631 | 0.435312 | 0.357161 | 0.339077 | 0.274013 | 0.237329 | 0.236104 | 0.225919 | 0.236116 | 0.261092 | 0.304369 | 0.412801 | 0.607345 | 0.735984 | 0.853615 | 0.798007 | 0.753573 | 0.638695 | 0.661051 | 0.532036 | 0.561916 | 0.684953 | 0.672494 | 0.742039 | 0.710863 | 0.538643 | 0.492522 | 0.419226 | 0.534498 | 0.888968 | 1.264333 | 1.916258 | 2.086523 | 2.013730 | 2.520309 | 1.941023 | 1.278081 | 0.933792 | 0.781791 | 0.735051 | 0.646566 | 0.508028 | 0.458320 | 0.431120 | 0.430542 | 0.394024 | 0.323881 | 0.387042 | 0.389605 | 0.405911 | 0.362335 | 0.335557 | 0.331913 | 0.433164 | 0.514305 | 0.711961 | 0.924605 | 1.507703 | 3.706649 | 6.965663 | 6.447491 | 4.956058 | 4.545640 | 3.236177 | 2.135783 | 0.846605 | 0.419680 | 0.360331 | 0.318037 | 0.286065 | 0.266359 | 0.236358 | 0.240425 | 0.242849 | 0.234266 | 0.238524 | 0.279137 | 0.348797 | 0.543589 | 0.626889 | 0.758454 | 0.753203 | 0.598558 | 0.436387 | 0.331834 | 0.269334 | 0.256007 | 0.281737 | 0.331764 | 0.394042 | 0.406439 | 0.549681 | 0.748777 | 0.732567 | 0.548719 | 0.430181 | 0.343800 | 0.300503 | 0.238720 | 0.221240 | 0.216254 | 0.215712 | 0.215118 | 0.214906 | 0.215879 | 0.224594 | 0.234276 | 0.237484 | 0.261521 | 0.247085 |
| right posterior | 0.228277 | 0.215599 | 0.214843 | 0.218441 | 0.214797 | 0.247256 | 0.250922 | 0.248015 | 0.221300 | 0.365234 | 1.397811 | 1.046656 | 0.587975 | 0.629111 | 0.682940 | 1.752261 | 0.849405 | 0.482177 | 1.010754 | 1.159024 | 1.886264 | 1.988286 | 2.672147 | 9.600908 | 7.238013 | 0.411112 | 0.268893 | 0.229554 | 0.238901 | 0.227297 | 0.215897 | 0.214704 | 0.218067 | 0.216201 | 0.214643 | 0.216087 | 0.217870 | 0.230262 | 0.219701 | 0.219555 | 0.219922 | 0.214626 | 0.226483 | 0.236121 | 0.250376 | 0.248352 | 0.304506 | 0.376720 | 0.500292 | 0.536582 | 0.667197 | 0.884829 | 1.488502 | 1.597064 | 2.537116 | 5.692983 | 25.556755 | 43.392363 | 150.660675 | 340.785055 | 1696.881448 | 6197.939036 | 9009.479581 | 18511.486666 | 41330.710739 | 21462.014821 | 7333.759946 | 1008.095702 | 161.052985 | 25.979838 | 7.965359 | 4.578617 | 5.732258 | 8.646761 | 23.167632 | 68.771422 | 334.324793 | 2625.631696 | 13215.390074 | 17913.829414 | 40371.243051 | 128638.328876 | 168102.540412 | 90196.593480 | 65105.242653 | 32802.824418 | 10954.078611 | 4135.878925 | 1534.830880 | 1218.138396 | 869.406382 | 427.603807 | 302.258371 | 155.351624 | 130.826484 | 111.461369 | 69.341261 | 54.979551 | 74.821302 | 148.064584 | 130.944159 | 74.981601 | 77.201258 | 142.456386 | 464.043077 | 1983.195454 | 907.878388 | 976.127479 | 2442.411569 | 2078.564338 | 2873.155383 | 1338.510745 | 166.866136 | 49.807829 | 24.641440 | 14.390597 | 11.253697 | 6.695735 | 4.193966 | 4.609650 | 5.840695 | 9.141125 | 8.133158 | 6.839105 | 4.362011 | 6.918846 | 8.923960 | 11.250554 | 12.180060 | 31.784934 | 63.484421 | 132.454991 | 129.921676 | 136.377715 | 129.464974 | 78.475214 | 27.319642 | 13.328482 | 7.371350 | 9.005648 | 12.288525 | 11.687098 | 11.341122 | 14.182869 | 17.102673 | 14.433155 | 6.887990 | 4.026942 | 4.288747 | 4.782410 | 7.156745 | 11.173895 | 16.505857 | 27.580575 | 32.445027 | 25.255414 | 18.233289 | 11.671297 | 6.307713 | 3.720958 | 2.721507 | 2.277305 | 2.980464 | 3.200607 | 2.341524 | 2.144785 | 1.958481 | 2.098804 | 3.400727 | 3.904563 | 3.909668 | 3.867387 | 6.268237 | 10.412001 | 10.893709 | 6.322658 | 5.465937 | 15.319227 | 69.954083 | 143.599952 | 169.576242 | 124.090656 | 70.949584 | 93.119986 | 34.384135 | 11.149203 | 6.658907 | 6.487640 | 9.043165 | 11.369895 | 9.960798 | 19.792020 | 43.547476 | 93.864709 | 199.731726 | 425.502426 | 901.723561 | 856.062252 | 253.650921 | 83.874579 | 21.469020 | 6.290739 | 2.317006 | 2.183317 | 2.123844 | 1.959943 | 1.463891 | 1.665893 | 2.607784 | 3.377618 | 2.753022 | 1.796035 | 1.693952 | 2.073215 | 3.276899 | 3.102472 | 2.972911 | 2.319824 | 1.779941 | 1.211679 | 0.800266 | 0.536541 | 0.459341 | 0.425868 | 0.455378 | 0.501749 | 0.542576 | 0.559777 | 0.512153 | 0.398901 | 0.479033 | 0.598680 | 0.604832 | 0.789245 | 1.000735 | 1.495901 | 3.378801 | 4.808702 | 3.694746 | 2.807047 | 1.619927 | 1.181568 | 0.829390 | 0.576950 | 0.401416 | 0.444641 | 0.552960 | 0.826341 | 0.943606 | 1.136165 | 1.099641 | 1.198515 | 0.845118 | 0.675517 | 0.602239 | 0.587806 | 0.614641 | 0.758865 | 0.902474 | 2.044651 | 4.052471 | 4.857576 | 5.730478 | 8.234046 | 7.219642 | 6.019108 | 2.511168 | 1.154918 | 0.729861 | 0.561954 | 0.472833 | 0.410830 | 0.446388 | 0.660392 | 1.088281 | 2.075181 | 4.697444 | 5.614043 | 7.965529 | 9.410386 |
| all electrodes | 0.368531 | 0.221134 | 0.216469 | 0.263165 | 0.244162 | 0.235398 | 0.216414 | 0.246075 | 0.418499 | 0.297601 | 0.290842 | 0.662770 | 0.985227 | 0.631514 | 0.397081 | 0.382506 | 0.343734 | 0.254652 | 0.277235 | 0.245438 | 0.223342 | 0.248388 | 0.216935 | 0.250277 | 0.332462 | 0.599272 | 0.314941 | 0.252298 | 0.219877 | 0.233553 | 0.321668 | 0.320456 | 0.317963 | 0.217702 | 0.215422 | 0.230353 | 0.235154 | 0.225694 | 0.231928 | 0.217495 | 0.214704 | 0.216991 | 0.227005 | 0.214626 | 0.284676 | 0.439790 | 1.416550 | 1.961982 | 1.331736 | 2.201033 | 0.923577 | 0.432171 | 0.320990 | 0.260532 | 0.246577 | 0.240444 | 0.236059 | 0.373749 | 1.141038 | 9.595847 | 408.711762 | 15108.511868 | 277252.812273 | 589805.090937 | 49213.385417 | 9495.051852 | 2412.584325 | 174.848095 | 43.164644 | 14.663270 | 16.090590 | 19.050319 | 39.493144 | 105.172072 | 591.811031 | 2807.691930 | 37722.402269 | 83813.103737 | 132543.950793 | 282042.518537 | 95391.370019 | 83909.031540 | 93350.664332 | 104043.787211 | 41962.578195 | 29632.473736 | 15175.101321 | 12561.018848 | 1996.715811 | 1116.280535 | 1820.543926 | 1336.402767 | 834.879630 | 421.158322 | 392.959418 | 306.902787 | 347.224987 | 364.923368 | 363.074710 | 251.055836 | 146.439477 | 73.407120 | 23.803729 | 32.716228 | 14.401868 | 6.407260 | 5.533809 | 3.714913 | 4.423110 | 13.768038 | 20.305739 | 53.392649 | 95.830605 | 71.019467 | 83.416215 | 79.255668 | 39.632577 | 36.054796 | 17.409555 | 6.599404 | 6.571206 | 10.411939 | 11.977534 | 18.367443 | 15.515656 | 12.925660 | 17.966763 | 13.320970 | 10.820481 | 4.912332 | 5.562736 | 9.358801 | 26.151891 | 38.366938 | 36.533211 | 22.419561 | 16.999006 | 7.832245 | 4.540229 | 1.923501 | 2.052709 | 1.617736 | 1.536117 | 1.103458 | 0.832514 | 0.662430 | 0.779956 | 0.405141 | 0.424413 | 0.744030 | 1.474781 | 3.415190 | 7.744318 | 12.360056 | 27.675786 | 38.331398 | 18.170007 | 6.133644 | 3.961400 | 2.051475 | 1.886236 | 1.887731 | 1.973712 | 2.858627 | 3.884671 | 3.446674 | 5.439794 | 4.797462 | 4.337392 | 4.028936 | 3.252472 | 2.696884 | 5.485078 | 5.277887 | 2.696203 | 1.321258 | 0.939602 | 0.830129 | 0.963187 | 0.626362 | 0.494071 | 0.596410 | 0.699703 | 0.861620 | 0.503151 | 0.414874 | 0.368346 | 0.296833 | 0.275810 | 0.363438 | 0.653174 | 0.894380 | 1.107118 | 1.333319 | 1.667232 | 1.147996 | 0.786998 | 0.429560 | 0.245305 | 0.218991 | 0.215170 | 0.216426 | 0.216788 | 0.216200 | 0.218138 | 0.215193 | 0.219400 | 0.224196 | 0.218340 | 0.217503 | 0.214742 | 0.217213 | 0.218576 | 0.220331 | 0.225095 | 0.248693 | 0.240959 | 0.242033 | 0.214706 | 0.224288 | 0.246441 | 0.265996 | 0.303602 | 0.292154 | 0.279582 | 0.235670 | 0.215577 | 0.215223 | 0.216015 | 0.216471 | 0.216747 | 0.216862 | 0.228834 | 0.232690 | 0.244054 | 0.258796 | 0.281896 | 0.262745 | 0.271884 | 0.223232 | 0.215242 | 0.233728 | 0.226556 | 0.221352 | 0.215621 | 0.217013 | 0.217678 | 0.217948 | 0.214680 | 0.214626 | 0.220519 | 0.224259 | 0.237050 | 0.254251 | 0.250721 | 0.248556 | 0.223058 | 0.214661 | 0.218168 | 0.239630 | 0.245138 | 0.221120 | 0.222716 | 0.217731 | 0.223142 | 0.221216 | 0.215187 | 0.214938 | 0.215178 | 0.222089 | 0.228097 | 0.230237 | 0.234780 | 0.226260 | 0.240961 | 0.254906 | 0.290208 | 0.313702 | 0.396816 | 0.486204 |

Searchlight, spatiotemporal cluster permutation test

|  | start time | stop time | peak time | peak channel | cluster p | peak Cohen's d | direction |
| --- | --- | --- | --- | --- | --- | --- | --- |
| #1 | 75 | 1075 | 225 | PO8 | 0.0001 | 1.594533 | positive |

B) sex

  
|  | time window | peak latency | cluster *p* | peak Cohen's *d* |  | | | |  |
| **all electrodes** | 155 - 450 ms | 195 ms | 0.008 | 1.0624 |  | | | |  |
|  | | | | | | | | |  |

Time-resolved classification, cluster permutation tests

|  | **left hemisphere** | | | | **right hemisphere** | | | |  |
|  | time window | peak latency | cluster *p* | peak Cohen's *d* | time window | peak latency | cluster *p* | peak Cohen's *d* |  |
| **anterior** | 85 - 515 ms | 255 ms | 0.0012 | 1.2387 | 160 - 585 ms | 250 ms | 0.0022 | 1.1276 |  |
 1000 - 1180 ms | 1140 ms | 0.0219 | 0.67 |  | | | ||  | **central** | 135 - 355 ms | 245 ms | 0.0054 | 0.8322 | 165 - 340 ms | 240 ms | 0.0051 | 1.0577 |
 735 - 890 ms | 770 ms | 0.0141 | 0.9374 |  | | | | 955 - 1150 ms | 1080 ms | 0.009 | 0.7835 |  | | | || **posterior** |  | 80 - 140 ms | 110 ms | 0.044 | 1.2774 | 75 - 135 ms | 105 ms | 0.0351 | 1.5687 |
 160 - 280 ms | 180 ms | 0.0243 | 1.1957 | 165 - 325 ms | 195 ms | 0.0143 | 0.8523 |

  

Time-resolved classification, Bayesian statistics

|  | -200 | -195 | -190 | -185 | -180 | -175 | -170 | -165 | -160 | -155 | -150 | -145 | -140 | -135 | -130 | -125 | -120 | -115 | -110 | -105 | -100 | -95 | -90 | -85 | -80 | -75 | -70 | -65 | -60 | -55 | -50 | -45 | -40 | -35 | -30 | -25 | -20 | -15 | -10 | -5 | 0 | 5 | 10 | 15 | 20 | 25 | 30 | 35 | 40 | 45 | 50 | 55 | 60 | 65 | 70 | 75 | 80 | 85 | 90 | 95 | 100 | 105 | 110 | 115 | 120 | 125 | 130 | 135 | 140 | 145 | 150 | 155 | 160 | 165 | 170 | 175 | 180 | 185 | 190 | 195 | 200 | 205 | 210 | 215 | 220 | 225 | 230 | 235 | 240 | 245 | 250 | 255 | 260 | 265 | 270 | 275 | 280 | 285 | 290 | 295 | 300 | 305 | 310 | 315 | 320 | 325 | 330 | 335 | 340 | 345 | 350 | 355 | 360 | 365 | 370 | 375 | 380 | 385 | 390 | 395 | 400 | 405 | 410 | 415 | 420 | 425 | 430 | 435 | 440 | 445 | 450 | 455 | 460 | 465 | 470 | 475 | 480 | 485 | 490 | 495 | 500 | 505 | 510 | 515 | 520 | 525 | 530 | 535 | 540 | 545 | 550 | 555 | 560 | 565 | 570 | 575 | 580 | 585 | 590 | 595 | 600 | 605 | 610 | 615 | 620 | 625 | 630 | 635 | 640 | 645 | 650 | 655 | 660 | 665 | 670 | 675 | 680 | 685 | 690 | 695 | 700 | 705 | 710 | 715 | 720 | 725 | 730 | 735 | 740 | 745 | 750 | 755 | 760 | 765 | 770 | 775 | 780 | 785 | 790 | 795 | 800 | 805 | 810 | 815 | 820 | 825 | 830 | 835 | 840 | 845 | 850 | 855 | 860 | 865 | 870 | 875 | 880 | 885 | 890 | 895 | 900 | 905 | 910 | 915 | 920 | 925 | 930 | 935 | 940 | 945 | 950 | 955 | 960 | 965 | 970 | 975 | 980 | 985 | 990 | 995 | 1000 | 1005 | 1010 | 1015 | 1020 | 1025 | 1030 | 1035 | 1040 | 1045 | 1050 | 1055 | 1060 | 1065 | 1070 | 1075 | 1080 | 1085 | 1090 | 1095 | 1100 | 1105 | 1110 | 1115 | 1120 | 1125 | 1130 | 1135 | 1140 | 1145 | 1150 | 1155 | 1160 | 1165 | 1170 | 1175 | 1180 | 1185 | 1190 | 1195 |
| --- | --- | --- | --- | --- | --- | --- | --- | --- | --- | --- | --- | --- | --- | --- | --- | --- | --- | --- | --- | --- | --- | --- | --- | --- | --- | --- | --- | --- | --- | --- | --- | --- | --- | --- | --- | --- | --- | --- | --- | --- | --- | --- | --- | --- | --- | --- | --- | --- | --- | --- | --- | --- | --- | --- | --- | --- | --- | --- | --- | --- | --- | --- | --- | --- | --- | --- | --- | --- | --- | --- | --- | --- | --- | --- | --- | --- | --- | --- | --- | --- | --- | --- | --- | --- | --- | --- | --- | --- | --- | --- | --- | --- | --- | --- | --- | --- | --- | --- | --- | --- | --- | --- | --- | --- | --- | --- | --- | --- | --- | --- | --- | --- | --- | --- | --- | --- | --- | --- | --- | --- | --- | --- | --- | --- | --- | --- | --- | --- | --- | --- | --- | --- | --- | --- | --- | --- | --- | --- | --- | --- | --- | --- | --- | --- | --- | --- | --- | --- | --- | --- | --- | --- | --- | --- | --- | --- | --- | --- | --- | --- | --- | --- | --- | --- | --- | --- | --- | --- | --- | --- | --- | --- | --- | --- | --- | --- | --- | --- | --- | --- | --- | --- | --- | --- | --- | --- | --- | --- | --- | --- | --- | --- | --- | --- | --- | --- | --- | --- | --- | --- | --- | --- | --- | --- | --- | --- | --- | --- | --- | --- | --- | --- | --- | --- | --- | --- | --- | --- | --- | --- | --- | --- | --- | --- | --- | --- | --- | --- | --- | --- | --- | --- | --- | --- | --- | --- | --- | --- | --- | --- | --- | --- | --- | --- | --- | --- | --- | --- | --- | --- | --- | --- | --- | --- | --- | --- | --- | --- | --- | --- | --- | --- | --- | --- | --- | --- | --- | --- | --- | --- | --- | --- | --- | --- | --- | --- | --- | --- | --- | --- |
| left anterior | 0.765735 | 0.760880 | 0.690445 | 0.423779 | 0.492581 | 0.280204 | 0.215692 | 0.215512 | 0.214979 | 0.215657 | 0.214654 | 0.217774 | 0.218246 | 0.224799 | 0.215006 | 0.214634 | 0.221769 | 0.274059 | 0.250862 | 0.217894 | 0.230420 | 0.223022 | 0.215428 | 0.244364 | 0.291524 | 0.219423 | 0.214783 | 0.215567 | 0.219853 | 0.244093 | 0.258437 | 0.250430 | 0.250779 | 0.277614 | 0.218588 | 0.234280 | 0.262127 | 0.215223 | 0.237418 | 0.272154 | 0.309469 | 0.352174 | 0.451035 | 0.312567 | 0.239369 | 0.218733 | 0.220032 | 0.220181 | 0.259560 | 0.322624 | 0.397808 | 0.493183 | 0.484082 | 0.368890 | 0.498322 | 0.734949 | 0.950024 | 5.971011 | 5.184733 | 2.157933 | 1.856174 | 1.720941 | 1.622663 | 1.591403 | 1.510793 | 1.570652 | 2.312831 | 3.113404 | 3.534781 | 2.899400 | 3.803655 | 5.535365 | 7.674134 | 6.459006 | 10.175940 | 19.693717 | 28.138145 | 45.810614 | 43.067891 | 45.807633 | 40.834298 | 25.775873 | 15.284735 | 14.667177 | 7.708144 | 9.693379 | 17.491267 | 69.937685 | 349.886319 | 1997.609007 | 3669.664882 | 5746.791911 | 2091.088903 | 838.898274 | 287.752314 | 209.593093 | 205.357010 | 260.248756 | 174.123564 | 142.086972 | 138.283240 | 137.239139 | 73.210706 | 41.481694 | 26.030001 | 25.391268 | 34.563082 | 20.181804 | 10.979846 | 4.776937 | 3.355984 | 2.833400 | 2.671844 | 2.730783 | 3.979257 | 5.785246 | 16.237178 | 27.450132 | 32.652649 | 33.945874 | 28.181059 | 17.394183 | 14.607734 | 9.427909 | 10.154093 | 19.285362 | 26.994407 | 47.333434 | 65.451971 | 61.014987 | 72.840654 | 46.075135 | 14.432181 | 7.723052 | 3.130668 | 2.374712 | 2.254177 | 2.070551 | 1.380936 | 1.633818 | 1.742749 | 2.729105 | 2.833431 | 2.398493 | 1.239040 | 0.933311 | 0.654257 | 0.546654 | 0.503600 | 0.469351 | 0.544515 | 0.783017 | 1.214385 | 2.311816 | 2.328010 | 1.753400 | 1.072243 | 0.626576 | 0.462043 | 0.346878 | 0.306010 | 0.325246 | 0.386496 | 0.481484 | 0.698461 | 0.740175 | 0.861863 | 0.709983 | 0.514846 | 0.509560 | 0.550253 | 0.445937 | 0.420770 | 0.318491 | 0.264304 | 0.263233 | 0.266733 | 0.250066 | 0.259771 | 0.241306 | 0.239103 | 0.255136 | 0.245802 | 0.220547 | 0.221622 | 0.228128 | 0.237659 | 0.258448 | 0.267865 | 0.301697 | 0.365021 | 0.444626 | 0.525454 | 0.690024 | 0.720497 | 0.744729 | 0.720704 | 0.564047 | 0.478702 | 0.298192 | 0.243333 | 0.246469 | 0.267218 | 0.313161 | 0.376281 | 0.368938 | 0.392240 | 0.511625 | 0.496893 | 0.474662 | 0.423662 | 0.451673 | 0.643257 | 0.914398 | 0.842318 | 1.004345 | 1.127298 | 1.629717 | 1.648289 | 0.894599 | 1.218313 | 2.704404 | 2.778254 | 2.105919 | 1.228544 | 0.849685 | 1.261077 | 1.186805 | 0.863730 | 0.942383 | 1.185592 | 1.191735 | 1.215090 | 0.769267 | 0.466115 | 0.351513 | 0.329295 | 0.359609 | 0.498495 | 0.842778 | 1.529290 | 3.353488 | 5.408942 | 6.984057 | 4.456885 | 3.204995 | 3.033190 | 2.922236 | 2.897204 | 2.693171 | 1.812700 | 1.553495 | 1.999971 | 2.161954 | 1.847800 | 1.773548 | 1.983502 | 3.296891 | 8.225754 | 9.529862 | 10.050639 | 10.649038 | 17.980551 | 19.023870 | 19.509082 | 13.353518 | 10.593595 | 10.960298 | 12.542086 | 8.293966 | 9.834460 | 9.582095 | 6.596302 | 4.133007 | 2.903347 | 2.035485 | 1.841651 | 1.256598 | 0.990411 | 1.031855 |
| right anterior | 0.274940 | 0.507909 | 1.292291 | 2.072146 | 3.074779 | 3.018358 | 1.191017 | 0.790280 | 0.477543 | 0.287243 | 0.253121 | 0.268880 | 0.338623 | 0.526993 | 0.700150 | 0.535438 | 0.301632 | 0.234808 | 0.236272 | 0.302314 | 0.249327 | 0.227157 | 0.278144 | 0.348752 | 0.416804 | 0.225270 | 0.219404 | 0.222894 | 0.217728 | 0.304906 | 1.170947 | 5.859687 | 8.396331 | 5.186343 | 1.988588 | 0.555999 | 0.267909 | 0.221205 | 0.214810 | 0.218139 | 0.275659 | 0.359626 | 0.574989 | 0.652299 | 0.512573 | 0.325357 | 0.225278 | 0.214758 | 0.216324 | 0.234517 | 0.245503 | 0.234814 | 0.290956 | 0.353624 | 0.337075 | 0.282358 | 0.241422 | 0.217904 | 0.240909 | 0.284413 | 0.342336 | 0.477189 | 0.589899 | 0.931189 | 1.698974 | 1.575928 | 1.211283 | 1.139988 | 0.915728 | 0.984647 | 1.110288 | 1.243357 | 2.206269 | 4.756665 | 9.161612 | 23.979867 | 25.936197 | 29.164770 | 14.357662 | 4.642243 | 2.647526 | 1.715558 | 1.588336 | 2.138180 | 2.180946 | 5.177049 | 16.030854 | 69.104905 | 390.087986 | 978.359538 | 1729.929097 | 1161.955476 | 434.761949 | 281.395582 | 247.476988 | 214.028303 | 303.217985 | 692.762312 | 2062.356661 | 5636.323023 | 12215.200708 | 9350.988328 | 12251.829974 | 6355.306500 | 3018.254474 | 1707.262529 | 1010.929251 | 528.007881 | 134.426832 | 29.964840 | 20.490921 | 15.791944 | 13.887412 | 14.622202 | 14.632724 | 18.260231 | 39.657089 | 28.485818 | 28.989767 | 23.657724 | 14.485943 | 11.689907 | 23.135594 | 16.173765 | 24.976846 | 30.264821 | 33.567953 | 42.177686 | 20.925361 | 12.001361 | 9.334447 | 10.446977 | 11.253581 | 11.365339 | 12.551700 | 11.263806 | 10.680677 | 14.243938 | 8.242856 | 3.441056 | 2.646774 | 2.944675 | 6.620696 | 9.178689 | 11.266303 | 14.589521 | 22.925921 | 18.956465 | 10.425526 | 5.740721 | 4.685466 | 4.337573 | 3.927864 | 3.393879 | 3.550403 | 3.925116 | 2.827472 | 1.677952 | 0.889441 | 0.617382 | 0.492055 | 0.446761 | 0.460417 | 0.555029 | 0.621654 | 0.725024 | 0.979803 | 1.112593 | 1.027880 | 0.909992 | 0.946379 | 1.235082 | 1.514982 | 1.447252 | 1.214355 | 1.168814 | 1.011382 | 0.941924 | 0.884984 | 0.658274 | 0.581630 | 0.604568 | 0.600210 | 0.683262 | 0.690344 | 0.714850 | 1.047488 | 1.692717 | 1.983119 | 1.857456 | 1.885063 | 2.333654 | 2.772543 | 3.090289 | 2.755619 | 2.367031 | 1.794357 | 1.437788 | 0.858532 | 0.456281 | 0.305507 | 0.264316 | 0.308791 | 0.425916 | 0.508744 | 0.791677 | 0.854461 | 0.699470 | 0.550404 | 0.460643 | 0.452752 | 0.492227 | 0.590392 | 0.676661 | 0.786454 | 0.651440 | 0.577971 | 0.505961 | 0.406374 | 0.279617 | 0.246480 | 0.242004 | 0.271561 | 0.294090 | 0.376835 | 0.407978 | 0.422027 | 0.431939 | 0.385815 | 0.278346 | 0.247987 | 0.216863 | 0.217782 | 0.237864 | 0.249229 | 0.225142 | 0.219640 | 0.224491 | 0.224510 | 0.233533 | 0.292373 | 0.357035 | 0.394071 | 0.346347 | 0.314223 | 0.285289 | 0.254200 | 0.221721 | 0.214659 | 0.220108 | 0.231638 | 0.225638 | 0.226174 | 0.220966 | 0.220771 | 0.224154 | 0.221203 | 0.215984 | 0.214964 | 0.238605 | 0.328670 | 0.463210 | 0.622529 | 0.750900 | 1.232414 | 1.829331 | 1.461917 | 0.911798 | 0.853869 | 0.889376 | 0.715320 | 0.583553 | 0.458086 | 0.394449 | 0.398086 | 0.656228 | 0.912661 | 1.066967 | 1.519132 | 2.156555 |
| left central | 0.383998 | 0.324800 | 0.256748 | 0.231076 | 0.214714 | 0.275115 | 0.366354 | 0.445349 | 0.478583 | 0.507693 | 0.567104 | 0.690996 | 0.478335 | 0.246742 | 0.220052 | 0.219136 | 0.339296 | 0.783663 | 2.525263 | 2.276505 | 0.661751 | 0.334200 | 0.286837 | 0.215189 | 0.219371 | 0.220011 | 0.214935 | 0.216918 | 0.232135 | 0.221991 | 0.215244 | 0.218906 | 0.225927 | 0.263435 | 0.259466 | 0.313584 | 0.247224 | 0.462279 | 2.502432 | 13.093786 | 13.241621 | 17.013878 | 33.868749 | 26.069234 | 1.642053 | 0.245784 | 0.229592 | 0.230288 | 0.235371 | 0.216641 | 0.214631 | 0.236081 | 0.275590 | 0.272677 | 0.224154 | 0.216773 | 0.221145 | 0.228771 | 0.235342 | 0.235669 | 0.222248 | 0.216900 | 0.222317 | 0.257892 | 0.351209 | 0.619117 | 0.973579 | 5.892057 | 21.338719 | 24.757896 | 3.980534 | 1.937773 | 1.428843 | 2.295170 | 2.459903 | 4.020743 | 4.173704 | 7.117687 | 11.382719 | 27.583205 | 30.855345 | 22.594019 | 13.924658 | 14.975629 | 16.104233 | 22.099038 | 20.976004 | 26.465027 | 43.030910 | 68.828222 | 77.737720 | 73.511696 | 34.566809 | 26.486817 | 23.028982 | 24.089255 | 20.084629 | 18.100701 | 15.990694 | 16.928407 | 29.061595 | 36.565731 | 19.826227 | 9.196731 | 9.288819 | 10.265101 | 7.685473 | 3.490619 | 2.800896 | 2.695714 | 2.542483 | 1.380787 | 0.774926 | 0.732270 | 0.649958 | 0.494780 | 0.476584 | 0.568398 | 0.562238 | 0.761579 | 0.831444 | 0.963080 | 1.269260 | 1.769996 | 2.753187 | 4.125303 | 4.581681 | 4.511178 | 4.160671 | 3.301732 | 3.969418 | 3.137300 | 2.517685 | 1.928947 | 1.736905 | 1.856493 | 2.732097 | 1.811842 | 1.278096 | 1.458723 | 1.980559 | 3.041815 | 3.615616 | 3.300055 | 2.810155 | 3.189440 | 3.733257 | 3.787663 | 3.324537 | 2.846645 | 1.942899 | 2.267768 | 2.650086 | 2.521368 | 1.612125 | 1.081293 | 0.961161 | 0.756768 | 0.509567 | 0.297933 | 0.234110 | 0.220686 | 0.226645 | 0.229835 | 0.271453 | 0.359063 | 0.833780 | 2.559217 | 5.788773 | 7.321342 | 6.442519 | 4.280979 | 2.875643 | 1.492745 | 0.671035 | 0.704610 | 0.858000 | 1.046230 | 1.579183 | 1.546324 | 1.514975 | 2.503237 | 1.704319 | 0.932438 | 0.634067 | 0.528562 | 0.858233 | 1.907134 | 2.649553 | 4.423324 | 8.933130 | 16.081088 | 46.526833 | 91.677863 | 216.058254 | 416.835190 | 385.621674 | 189.076708 | 71.065674 | 26.272078 | 16.208230 | 10.756737 | 6.976296 | 5.865235 | 3.998430 | 4.137379 | 5.045612 | 4.019517 | 4.295083 | 5.785219 | 5.869706 | 8.903527 | 17.459839 | 11.273481 | 9.481419 | 5.897241 | 3.015699 | 2.453942 | 1.403853 | 0.990686 | 1.108244 | 0.901782 | 1.327358 | 1.439209 | 1.363085 | 1.099423 | 1.052957 | 0.918454 | 1.207986 | 1.131191 | 1.293333 | 1.391446 | 2.196215 | 3.952800 | 5.215324 | 4.105728 | 3.302508 | 4.791539 | 6.456041 | 6.931082 | 4.620718 | 4.289791 | 5.237772 | 5.652250 | 4.478804 | 6.037325 | 5.519390 | 8.241652 | 9.013204 | 7.230578 | 11.393326 | 18.995736 | 19.377414 | 19.083256 | 15.004085 | 16.694561 | 40.877215 | 50.709332 | 39.523922 | 26.271297 | 16.925793 | 16.769236 | 15.242985 | 18.268342 | 21.152967 | 18.645161 | 13.340078 | 14.254906 | 10.061709 | 4.739561 | 1.894852 | 1.036897 | 0.829886 | 0.814162 | 0.905309 | 0.901512 | 1.212494 | 1.459321 | 2.036119 | 2.824865 |
| right central | 0.516853 | 0.642476 | 0.785772 | 0.508367 | 0.386371 | 0.390446 | 0.348535 | 0.304235 | 0.315117 | 0.275567 | 0.273248 | 0.359009 | 0.571278 | 0.303119 | 0.223477 | 0.223600 | 0.267134 | 0.242424 | 0.391337 | 1.327783 | 2.716918 | 2.087534 | 4.063924 | 2.043336 | 1.169462 | 0.425021 | 0.228779 | 0.222239 | 0.282953 | 0.694245 | 1.671468 | 1.428574 | 1.235253 | 1.985936 | 2.013336 | 0.865637 | 0.452779 | 0.265675 | 0.239901 | 0.232675 | 0.214677 | 0.215704 | 0.253098 | 0.328315 | 0.550880 | 0.671636 | 0.508986 | 0.738503 | 0.822639 | 0.717815 | 0.710506 | 0.656138 | 0.888795 | 1.260761 | 1.783205 | 2.809632 | 3.087904 | 2.080536 | 1.141720 | 0.500751 | 0.321789 | 0.260402 | 0.298157 | 0.430899 | 0.473444 | 0.463245 | 0.458389 | 0.393469 | 0.411341 | 0.413052 | 0.349151 | 0.422393 | 0.784997 | 2.060199 | 8.971978 | 25.727911 | 32.523041 | 59.261656 | 131.537439 | 65.302167 | 23.342218 | 12.683494 | 13.367284 | 25.241270 | 40.676489 | 55.343935 | 120.953305 | 342.453747 | 806.650312 | 706.261125 | 271.512361 | 143.807454 | 161.815476 | 184.068273 | 119.940449 | 127.775997 | 151.087926 | 344.906079 | 526.726161 | 389.243387 | 363.279347 | 274.584275 | 123.053312 | 50.057461 | 27.488097 | 17.628679 | 7.829598 | 3.197644 | 1.303363 | 0.855959 | 0.813850 | 0.814867 | 0.690804 | 0.818093 | 0.985525 | 1.325019 | 1.127927 | 0.792191 | 0.502997 | 0.357324 | 0.298424 | 0.310217 | 0.315500 | 0.378696 | 0.446050 | 0.534592 | 0.671325 | 1.014629 | 1.182661 | 1.245015 | 1.255557 | 1.226522 | 1.505664 | 2.045406 | 2.974599 | 4.535213 | 7.129716 | 9.851316 | 10.362465 | 10.568125 | 8.747447 | 4.612914 | 4.249145 | 4.308040 | 5.105322 | 5.888465 | 4.968332 | 5.206506 | 5.083481 | 3.633025 | 2.547773 | 1.395088 | 1.227745 | 1.285233 | 1.003227 | 0.826065 | 0.782201 | 0.728131 | 0.521045 | 0.404537 | 0.320952 | 0.311922 | 0.333095 | 0.330744 | 0.361327 | 0.554916 | 0.655773 | 0.586267 | 0.411557 | 0.325871 | 0.279036 | 0.248919 | 0.224012 | 0.215794 | 0.215956 | 0.220472 | 0.227746 | 0.235115 | 0.233123 | 0.215232 | 0.215305 | 0.214882 | 0.214708 | 0.214964 | 0.214650 | 0.215645 | 0.257045 | 0.374188 | 0.453717 | 0.441514 | 0.456385 | 0.495832 | 0.498265 | 0.437855 | 0.364415 | 0.298428 | 0.280098 | 0.247103 | 0.231195 | 0.226954 | 0.227958 | 0.219804 | 0.220819 | 0.224605 | 0.239013 | 0.274194 | 0.318282 | 0.352230 | 0.363338 | 0.460049 | 0.525750 | 0.554930 | 0.348817 | 0.256212 | 0.226278 | 0.216902 | 0.216963 | 0.237437 | 0.270895 | 0.269777 | 0.287566 | 0.331962 | 0.364431 | 0.537952 | 0.605398 | 0.515903 | 0.371893 | 0.314675 | 0.282309 | 0.270606 | 0.247256 | 0.233802 | 0.225717 | 0.222189 | 0.214854 | 0.216741 | 0.223564 | 0.251170 | 0.273635 | 0.318257 | 0.368854 | 0.377813 | 0.409062 | 0.440200 | 0.353192 | 0.319113 | 0.245827 | 0.216325 | 0.214757 | 0.217480 | 0.215065 | 0.215663 | 0.216948 | 0.228977 | 0.275606 | 0.290865 | 0.314134 | 0.310871 | 0.330441 | 0.339259 | 0.311690 | 0.254862 | 0.240669 | 0.238123 | 0.237385 | 0.235580 | 0.253140 | 0.334592 | 0.516379 | 0.620809 | 0.535149 | 0.543776 | 0.504465 | 0.433271 | 0.341014 | 0.244632 | 0.220604 | 0.221205 | 0.215661 | 0.214645 |
| left posterior | 0.526596 | 0.547531 | 0.588544 | 0.414700 | 0.265832 | 0.219102 | 0.214757 | 0.259303 | 0.387390 | 0.516792 | 0.542972 | 0.486915 | 0.438518 | 0.813578 | 0.457923 | 0.724218 | 0.832729 | 1.012367 | 0.588327 | 0.298920 | 0.217007 | 0.217924 | 0.240006 | 0.223234 | 0.231092 | 0.236581 | 0.249475 | 0.224305 | 0.214632 | 0.214692 | 0.215616 | 0.226049 | 0.251854 | 0.315518 | 0.341817 | 0.366572 | 0.278197 | 0.222698 | 0.254635 | 0.386484 | 0.355989 | 0.277606 | 0.220438 | 0.225851 | 0.255017 | 0.437840 | 0.697168 | 0.724663 | 0.544555 | 0.322896 | 0.284400 | 0.328811 | 0.242343 | 0.214873 | 0.275269 | 0.667879 | 4.351473 | 32.255643 | 139.231223 | 222.039702 | 944.482753 | 3370.805080 | 8688.434463 | 12144.602115 | 5662.184223 | 1027.742909 | 123.987120 | 7.151235 | 1.721000 | 0.840818 | 0.792781 | 0.977824 | 2.924024 | 20.140644 | 386.137278 | 1628.588549 | 3620.449818 | 1253.336709 | 729.565559 | 229.074613 | 40.891908 | 9.880703 | 4.449609 | 2.371726 | 2.433473 | 2.570021 | 2.664207 | 3.801703 | 4.501643 | 3.741512 | 2.991108 | 2.410657 | 2.145275 | 1.957148 | 1.527077 | 1.363852 | 1.401750 | 1.199731 | 1.621139 | 1.480508 | 1.896183 | 2.264518 | 2.388691 | 1.924686 | 2.634885 | 2.889847 | 4.051588 | 2.873854 | 1.995015 | 1.298160 | 1.133322 | 0.865245 | 0.616791 | 0.480576 | 0.442197 | 0.433818 | 0.436597 | 0.465480 | 0.421111 | 0.424444 | 0.489138 | 0.541230 | 0.561126 | 0.677757 | 0.931213 | 1.258573 | 0.825779 | 0.480786 | 0.289601 | 0.223631 | 0.216834 | 0.236840 | 0.243998 | 0.231557 | 0.223522 | 0.219428 | 0.215748 | 0.215050 | 0.215251 | 0.222368 | 0.222070 | 0.221300 | 0.220878 | 0.219052 | 0.218035 | 0.225910 | 0.225882 | 0.233870 | 0.239057 | 0.232643 | 0.227859 | 0.222274 | 0.215665 | 0.216307 | 0.231964 | 0.272372 | 0.326287 | 0.336993 | 0.341776 | 0.389596 | 0.407979 | 0.357740 | 0.274417 | 0.239434 | 0.245356 | 0.259273 | 0.270655 | 0.287009 | 0.326042 | 0.447116 | 0.633764 | 0.796843 | 0.939583 | 0.983878 | 0.948789 | 0.995836 | 0.823858 | 0.592371 | 0.471903 | 0.375740 | 0.383644 | 0.393284 | 0.389899 | 0.381465 | 0.400218 | 0.426056 | 0.491440 | 0.466876 | 0.453053 | 0.426313 | 0.464416 | 0.560737 | 0.702454 | 0.791684 | 0.758256 | 0.846435 | 0.946911 | 1.102525 | 1.165118 | 0.918008 | 0.722494 | 0.713071 | 0.652208 | 0.676285 | 0.536929 | 0.518823 | 0.732948 | 1.442207 | 1.901997 | 2.395480 | 2.333732 | 2.981836 | 4.110824 | 2.876982 | 1.690971 | 1.007632 | 0.615490 | 0.436696 | 0.347038 | 0.274161 | 0.248223 | 0.226120 | 0.222899 | 0.228363 | 0.244394 | 0.244854 | 0.238125 | 0.224561 | 0.224077 | 0.222876 | 0.220805 | 0.218804 | 0.225793 | 0.250051 | 0.290313 | 0.318633 | 0.404673 | 0.562343 | 0.758303 | 0.831612 | 0.703696 | 0.566076 | 0.458637 | 0.327140 | 0.260370 | 0.242042 | 0.239718 | 0.243330 | 0.268173 | 0.302275 | 0.324536 | 0.344855 | 0.311773 | 0.333690 | 0.360617 | 0.354465 | 0.384085 | 0.509033 | 0.676940 | 1.006629 | 0.933764 | 0.899556 | 0.784383 | 0.789538 | 0.791008 | 0.798713 | 0.721490 | 0.935236 | 0.834578 | 0.900694 | 0.862252 | 0.776592 | 0.770036 | 0.836052 | 0.610772 | 0.576899 | 0.624806 | 0.561602 | 0.546966 | 0.447445 |
| right posterior | 0.478340 | 0.294102 | 0.233746 | 0.233625 | 0.221596 | 0.223212 | 0.258114 | 0.257321 | 0.228939 | 0.226205 | 0.223092 | 0.223244 | 0.227252 | 0.215777 | 0.221482 | 0.236113 | 0.238820 | 0.215333 | 0.214733 | 0.221657 | 0.221425 | 0.219511 | 0.237794 | 0.382165 | 0.835405 | 2.732428 | 9.727992 | 15.914896 | 5.809956 | 1.182546 | 0.427336 | 0.312692 | 0.228998 | 0.219548 | 0.314863 | 0.386753 | 0.358067 | 0.364225 | 0.673139 | 1.551950 | 3.567986 | 8.265121 | 21.687107 | 26.183053 | 53.027924 | 19.754963 | 14.287390 | 9.216862 | 5.084039 | 3.164488 | 2.239006 | 1.547131 | 1.238740 | 0.295666 | 0.303728 | 4.958361 | 163.653833 | 1698.435108 | 12076.142241 | 41059.121498 | 56094.024053 | 175280.309656 | 256449.188971 | 205918.374122 | 28339.062094 | 953.802828 | 31.208094 | 2.023232 | 0.489334 | 0.298794 | 0.289439 | 0.379674 | 0.668939 | 1.856049 | 6.047409 | 18.739778 | 49.645121 | 72.302367 | 85.830855 | 85.521121 | 50.202750 | 35.922889 | 18.317804 | 8.878275 | 4.783885 | 3.850506 | 2.980370 | 3.771065 | 5.200495 | 10.477036 | 18.043716 | 54.084627 | 50.590985 | 73.245220 | 92.551460 | 93.205668 | 77.984743 | 72.501868 | 34.424484 | 30.747423 | 25.206738 | 12.294238 | 6.724331 | 3.099494 | 1.916875 | 1.489734 | 1.200803 | 1.008085 | 1.066017 | 1.123403 | 1.533140 | 2.561040 | 3.157825 | 3.425647 | 3.924141 | 3.320989 | 4.490720 | 5.317646 | 3.678165 | 3.616940 | 2.553314 | 2.213949 | 2.178340 | 1.777748 | 1.408769 | 1.462306 | 1.648948 | 1.782527 | 1.529829 | 1.277407 | 1.108561 | 0.948274 | 0.778723 | 0.514318 | 0.396956 | 0.336795 | 0.304701 | 0.311025 | 0.312242 | 0.353373 | 0.393560 | 0.424814 | 0.400667 | 0.421250 | 0.437212 | 0.494415 | 0.493488 | 0.530372 | 0.605346 | 0.669111 | 0.591403 | 0.573037 | 0.450231 | 0.424703 | 0.398622 | 0.374107 | 0.413126 | 0.481981 | 0.443632 | 0.447606 | 0.415363 | 0.422472 | 0.452726 | 0.448617 | 0.420379 | 0.418099 | 0.405030 | 0.388167 | 0.361955 | 0.355831 | 0.391411 | 0.430519 | 0.446372 | 0.441580 | 0.412744 | 0.309645 | 0.267696 | 0.225795 | 0.214832 | 0.230199 | 0.244130 | 0.243550 | 0.230060 | 0.242261 | 0.232929 | 0.218633 | 0.215175 | 0.232057 | 0.257023 | 0.263016 | 0.309528 | 0.357256 | 0.385978 | 0.361111 | 0.334699 | 0.281187 | 0.285820 | 0.245358 | 0.219719 | 0.214629 | 0.222152 | 0.248530 | 0.259481 | 0.288403 | 0.268529 | 0.241703 | 0.247033 | 0.232571 | 0.223419 | 0.223473 | 0.227517 | 0.242680 | 0.278408 | 0.276982 | 0.270102 | 0.308361 | 0.388551 | 0.370664 | 0.412930 | 0.421160 | 0.526273 | 0.917099 | 1.093390 | 1.294580 | 1.613860 | 1.293307 | 1.505737 | 1.964806 | 2.302315 | 4.141077 | 5.329160 | 5.956857 | 8.090978 | 6.241789 | 3.106720 | 1.179881 | 0.400316 | 0.249268 | 0.224219 | 0.216945 | 0.218152 | 0.216310 | 0.222660 | 0.234120 | 0.261049 | 0.256615 | 0.246102 | 0.240088 | 0.252579 | 0.252750 | 0.264377 | 0.266345 | 0.284447 | 0.332985 | 0.295468 | 0.264678 | 0.239837 | 0.217432 | 0.214874 | 0.222382 | 0.256763 | 0.260631 | 0.251380 | 0.243925 | 0.236352 | 0.245052 | 0.234527 | 0.218354 | 0.214638 | 0.214629 | 0.216735 | 0.225036 | 0.237183 | 0.252246 | 0.255082 | 0.251521 | 0.232795 | 0.229981 | 0.218907 | 0.215363 |
| all electrodes | 0.285753 | 0.374450 | 0.744845 | 2.028599 | 5.705253 | 3.901599 | 2.371633 | 0.834460 | 0.629179 | 0.381086 | 0.215527 | 0.236235 | 0.271582 | 0.233057 | 0.218343 | 0.225766 | 0.222564 | 0.261851 | 0.257081 | 0.347895 | 0.689904 | 0.744481 | 0.537547 | 0.315005 | 0.253008 | 0.448808 | 0.336305 | 0.214642 | 0.214685 | 0.307682 | 1.767802 | 3.553639 | 1.240036 | 1.452508 | 1.491066 | 0.911050 | 0.334686 | 0.221101 | 0.232372 | 0.239306 | 0.240449 | 0.235790 | 0.250615 | 0.262991 | 0.222518 | 0.215467 | 0.221211 | 0.228114 | 0.231852 | 0.306757 | 0.456373 | 0.379715 | 0.330261 | 0.360033 | 0.525197 | 1.432263 | 4.329911 | 23.211246 | 105.883617 | 375.025831 | 983.786984 | 2449.089207 | 1831.766326 | 1162.372333 | 280.224142 | 26.012254 | 2.028441 | 0.533262 | 0.399625 | 0.501075 | 0.575822 | 1.352853 | 7.095332 | 88.705314 | 596.231191 | 913.002807 | 743.175768 | 702.217199 | 649.752469 | 849.079450 | 266.296925 | 60.954685 | 30.993560 | 43.138721 | 61.199128 | 45.448020 | 28.299828 | 50.725280 | 75.529450 | 92.289091 | 84.678138 | 51.955444 | 30.576194 | 50.402987 | 89.129159 | 179.570457 | 125.252237 | 81.176253 | 67.689749 | 50.188901 | 22.515135 | 9.276008 | 5.991886 | 4.395350 | 3.856714 | 4.049153 | 6.048590 | 4.420525 | 2.970359 | 2.260671 | 1.702538 | 1.667836 | 1.675563 | 1.706845 | 2.426155 | 2.191086 | 2.181289 | 1.908085 | 1.396478 | 1.456017 | 1.742887 | 1.729231 | 2.095086 | 2.408346 | 3.856398 | 5.367610 | 4.724695 | 3.465759 | 2.828794 | 1.928029 | 1.550681 | 1.196848 | 0.970465 | 0.822206 | 0.771755 | 0.764863 | 0.815860 | 0.640941 | 0.553767 | 0.667858 | 0.792077 | 0.662129 | 0.622209 | 0.584110 | 0.714966 | 0.623950 | 0.418627 | 0.313909 | 0.298469 | 0.313917 | 0.335351 | 0.345470 | 0.402669 | 0.473120 | 0.652121 | 0.871176 | 0.794582 | 0.855627 | 0.776625 | 0.690491 | 0.691218 | 0.680202 | 0.644624 | 0.576797 | 0.477723 | 0.483413 | 0.543931 | 0.511522 | 0.447371 | 0.511812 | 0.876476 | 1.363257 | 1.600452 | 1.273029 | 0.984693 | 0.935035 | 0.703513 | 0.502629 | 0.417344 | 0.367955 | 0.381365 | 0.401544 | 0.364312 | 0.335545 | 0.318352 | 0.306955 | 0.360244 | 0.523707 | 1.038952 | 2.042418 | 3.055028 | 4.163576 | 5.315732 | 4.770141 | 2.971433 | 1.647594 | 1.222087 | 1.050099 | 0.737077 | 0.488703 | 0.331363 | 0.281493 | 0.285777 | 0.305093 | 0.333013 | 0.408592 | 0.466469 | 0.523663 | 0.529119 | 0.453832 | 0.465859 | 0.521807 | 0.528139 | 0.587562 | 0.545929 | 0.619389 | 0.882232 | 0.759879 | 0.509949 | 0.485474 | 0.485720 | 0.478595 | 0.422782 | 0.372288 | 0.385905 | 0.436829 | 0.371604 | 0.365258 | 0.320461 | 0.296202 | 0.261739 | 0.238976 | 0.230175 | 0.234036 | 0.236638 | 0.263829 | 0.310310 | 0.355557 | 0.386316 | 0.433774 | 0.505343 | 0.534866 | 0.465544 | 0.405384 | 0.329644 | 0.267835 | 0.252201 | 0.250254 | 0.273910 | 0.315545 | 0.306523 | 0.323654 | 0.365942 | 0.369801 | 0.365319 | 0.366897 | 0.448634 | 0.590795 | 1.069875 | 2.006707 | 3.722815 | 4.745210 | 4.200524 | 3.168208 | 2.513456 | 1.464153 | 0.926352 | 0.798686 | 0.682160 | 0.647211 | 0.637539 | 0.809484 | 1.222022 | 2.012641 | 2.429151 | 2.474074 | 2.701608 | 2.807071 | 2.323126 | 1.924325 |

Searchlight, spatiotemporal cluster permutation test

|  | start time | stop time | peak time | peak channel | cluster p | peak Cohen's d | direction |
| --- | --- | --- | --- | --- | --- | --- | --- |
| #1 | 70 | 1195 | 105 | POz | 0.0001 | 1.892772 | positive |

C) emotion

  
|  | time window | peak latency | cluster *p* | peak Cohen's *d* |  | | | |  |
| **all electrodes** | 120 - 225 ms | 170 ms | 0.0149 | 1.1992 |  | | | |  |
 245 - 335 ms | 280 ms | 0.0326 | 0.9169 |  | | | | 345 - 430 ms | 410 ms | 0.0437 | 0.7775 |  | | | ||  |  | | | | | | | | |

Time-resolved classification, cluster permutation tests

|  |  | **left hemisphere** | | | | **right hemisphere** | | | |
|  | time window | peak latency | cluster *p* | peak Cohen's *d* | time window | peak latency | cluster *p* | peak Cohen's *d* |  |
| **anterior** |  | | | | 320 - 430 ms | 350 ms | 0.0324 | 0.7676 |  |
| **central** |  | | | | 125 - 470 ms | 270 ms | 0.0002 | 1.3182 |  |
| **posterior** | 110 - 535 ms | 150 ms | 0.0012 | 1.9544 | 90 - 665 ms | 270 ms | 0.0007 | 1.0842 |  |
 840 - 1025 ms | 890 ms | 0.022 | 0.61 | 685 - 1125 ms | 890 ms | 0.0029 | 0.9069 |

  

Time-resolved classification, Bayesian statistics

|  | -200 | -195 | -190 | -185 | -180 | -175 | -170 | -165 | -160 | -155 | -150 | -145 | -140 | -135 | -130 | -125 | -120 | -115 | -110 | -105 | -100 | -95 | -90 | -85 | -80 | -75 | -70 | -65 | -60 | -55 | -50 | -45 | -40 | -35 | -30 | -25 | -20 | -15 | -10 | -5 | 0 | 5 | 10 | 15 | 20 | 25 | 30 | 35 | 40 | 45 | 50 | 55 | 60 | 65 | 70 | 75 | 80 | 85 | 90 | 95 | 100 | 105 | 110 | 115 | 120 | 125 | 130 | 135 | 140 | 145 | 150 | 155 | 160 | 165 | 170 | 175 | 180 | 185 | 190 | 195 | 200 | 205 | 210 | 215 | 220 | 225 | 230 | 235 | 240 | 245 | 250 | 255 | 260 | 265 | 270 | 275 | 280 | 285 | 290 | 295 | 300 | 305 | 310 | 315 | 320 | 325 | 330 | 335 | 340 | 345 | 350 | 355 | 360 | 365 | 370 | 375 | 380 | 385 | 390 | 395 | 400 | 405 | 410 | 415 | 420 | 425 | 430 | 435 | 440 | 445 | 450 | 455 | 460 | 465 | 470 | 475 | 480 | 485 | 490 | 495 | 500 | 505 | 510 | 515 | 520 | 525 | 530 | 535 | 540 | 545 | 550 | 555 | 560 | 565 | 570 | 575 | 580 | 585 | 590 | 595 | 600 | 605 | 610 | 615 | 620 | 625 | 630 | 635 | 640 | 645 | 650 | 655 | 660 | 665 | 670 | 675 | 680 | 685 | 690 | 695 | 700 | 705 | 710 | 715 | 720 | 725 | 730 | 735 | 740 | 745 | 750 | 755 | 760 | 765 | 770 | 775 | 780 | 785 | 790 | 795 | 800 | 805 | 810 | 815 | 820 | 825 | 830 | 835 | 840 | 845 | 850 | 855 | 860 | 865 | 870 | 875 | 880 | 885 | 890 | 895 | 900 | 905 | 910 | 915 | 920 | 925 | 930 | 935 | 940 | 945 | 950 | 955 | 960 | 965 | 970 | 975 | 980 | 985 | 990 | 995 | 1000 | 1005 | 1010 | 1015 | 1020 | 1025 | 1030 | 1035 | 1040 | 1045 | 1050 | 1055 | 1060 | 1065 | 1070 | 1075 | 1080 | 1085 | 1090 | 1095 | 1100 | 1105 | 1110 | 1115 | 1120 | 1125 | 1130 | 1135 | 1140 | 1145 | 1150 | 1155 | 1160 | 1165 | 1170 | 1175 | 1180 | 1185 | 1190 | 1195 |
| --- | --- | --- | --- | --- | --- | --- | --- | --- | --- | --- | --- | --- | --- | --- | --- | --- | --- | --- | --- | --- | --- | --- | --- | --- | --- | --- | --- | --- | --- | --- | --- | --- | --- | --- | --- | --- | --- | --- | --- | --- | --- | --- | --- | --- | --- | --- | --- | --- | --- | --- | --- | --- | --- | --- | --- | --- | --- | --- | --- | --- | --- | --- | --- | --- | --- | --- | --- | --- | --- | --- | --- | --- | --- | --- | --- | --- | --- | --- | --- | --- | --- | --- | --- | --- | --- | --- | --- | --- | --- | --- | --- | --- | --- | --- | --- | --- | --- | --- | --- | --- | --- | --- | --- | --- | --- | --- | --- | --- | --- | --- | --- | --- | --- | --- | --- | --- | --- | --- | --- | --- | --- | --- | --- | --- | --- | --- | --- | --- | --- | --- | --- | --- | --- | --- | --- | --- | --- | --- | --- | --- | --- | --- | --- | --- | --- | --- | --- | --- | --- | --- | --- | --- | --- | --- | --- | --- | --- | --- | --- | --- | --- | --- | --- | --- | --- | --- | --- | --- | --- | --- | --- | --- | --- | --- | --- | --- | --- | --- | --- | --- | --- | --- | --- | --- | --- | --- | --- | --- | --- | --- | --- | --- | --- | --- | --- | --- | --- | --- | --- | --- | --- | --- | --- | --- | --- | --- | --- | --- | --- | --- | --- | --- | --- | --- | --- | --- | --- | --- | --- | --- | --- | --- | --- | --- | --- | --- | --- | --- | --- | --- | --- | --- | --- | --- | --- | --- | --- | --- | --- | --- | --- | --- | --- | --- | --- | --- | --- | --- | --- | --- | --- | --- | --- | --- | --- | --- | --- | --- | --- | --- | --- | --- | --- | --- | --- | --- | --- | --- | --- | --- | --- | --- | --- | --- | --- | --- | --- | --- | --- | --- |
| left anterior | 0.249447 | 0.250807 | 0.248401 | 0.267471 | 0.253214 | 0.316832 | 0.288002 | 0.226914 | 0.242739 | 0.231438 | 0.214626 | 0.220910 | 0.217228 | 0.224840 | 0.235645 | 0.246754 | 0.269874 | 0.251961 | 0.241724 | 0.215427 | 0.214665 | 0.241338 | 0.263789 | 0.352125 | 0.702053 | 2.640009 | 6.731508 | 10.020962 | 5.012675 | 3.831471 | 1.986756 | 0.712248 | 0.215614 | 0.356506 | 0.973740 | 2.528500 | 8.275667 | 13.836477 | 2.333343 | 0.591987 | 0.304299 | 0.514897 | 0.869452 | 1.157003 | 1.346315 | 1.936545 | 3.192563 | 19.851330 | 31.934163 | 18.832712 | 18.455357 | 7.011618 | 3.406243 | 2.303253 | 1.128724 | 0.477614 | 0.281194 | 0.227970 | 0.221514 | 0.224814 | 0.244587 | 0.307060 | 0.303432 | 0.260770 | 0.215252 | 0.218207 | 0.245388 | 2.791346e-01 | 4.323004e-01 | 5.298500e-01 | 5.044683e-01 | 3.506814e-01 | 0.323049 | 0.290468 | 0.316597 | 0.254217 | 0.217996 | 0.403869 | 0.928133 | 4.320102 | 12.882760 | 18.377680 | 10.730822 | 4.270157 | 1.202479 | 0.393879 | 0.236877 | 0.220183 | 0.307358 | 0.873577 | 3.569675 | 8.518581 | 6.458934 | 7.645720 | 9.012092 | 10.677854 | 5.099296 | 1.789357 | 1.541176 | 2.849167 | 2.556837 | 1.268992 | 0.663295 | 0.371743 | 0.308113 | 0.234269 | 0.214870 | 0.218870 | 0.216212 | 0.255301 | 0.424902 | 0.756040 | 1.161207 | 1.295045 | 1.094236 | 0.721977 | 0.488273 | 0.481521 | 0.411819 | 0.451306 | 0.459215 | 0.460482 | 0.412309 | 0.444729 | 0.370828 | 0.375111 | 0.345658 | 0.260414 | 0.265592 | 0.260345 | 0.242231 | 0.223130 | 0.214903 | 0.215862 | 0.216279 | 0.236117 | 0.273238 | 0.272162 | 0.273165 | 0.232244 | 0.216344 | 0.222916 | 0.245512 | 0.314134 | 0.314523 | 0.399226 | 0.332635 | 0.349014 | 0.345024 | 0.366963 | 0.294519 | 0.262834 | 0.231915 | 0.218630 | 0.227662 | 0.272394 | 0.311016 | 0.278792 | 0.235612 | 0.219128 | 0.216173 | 0.257922 | 0.309006 | 0.397833 | 0.450662 | 0.488138 | 0.575365 | 0.478598 | 0.378215 | 0.350357 | 0.278767 | 0.245506 | 0.232944 | 0.224317 | 0.227532 | 0.229022 | 0.232129 | 0.252075 | 0.296710 | 0.354332 | 0.470231 | 0.605377 | 0.645572 | 0.628342 | 0.723259 | 0.816786 | 0.814831 | 0.577141 | 0.394102 | 0.405321 | 0.410337 | 0.317938 | 0.311274 | 0.358307 | 0.497852 | 0.918798 | 1.040661 | 1.026774 | 1.214456 | 1.318834 | 0.925089 | 0.664522 | 0.453697 | 0.362053 | 0.318450 | 0.272660 | 0.237793 | 0.234070 | 0.238034 | 0.269893 | 0.311808 | 0.379527 | 0.476412 | 0.605633 | 0.712606 | 1.247625 | 1.661634 | 2.378683 | 3.412085 | 5.569663 | 6.219846 | 10.533098 | 5.970180 | 4.386873 | 3.055252 | 2.461215 | 1.330475 | 0.854978 | 0.838426 | 1.093312 | 1.063241 | 0.991478 | 0.760288 | 0.677259 | 0.652117 | 0.508326 | 0.521029 | 0.730430 | 0.763798 | 0.783159 | 0.791132 | 1.007223 | 1.616143 | 2.324154 | 1.810965 | 2.304710 | 2.900951 | 3.861002 | 3.925891 | 1.764132 | 1.194661 | 1.000365 | 1.085351 | 2.107275 | 3.588787 | 4.468265 | 3.736221 | 2.300389 | 1.869835 | 1.432092 | 0.920339 | 0.715297 | 0.637556 | 0.639028 | 0.600369 | 0.824252 | 0.734580 | 0.687120 | 0.928013 | 1.026262 | 0.787706 | 0.480916 | 0.347792 | 0.341742 | 0.355685 | 0.307228 | 0.258231 | 0.261123 | 0.333662 | 0.373024 |
| right anterior | 0.412154 | 0.407008 | 0.304850 | 0.296168 | 0.235989 | 0.231516 | 0.238225 | 0.216946 | 0.215054 | 0.223469 | 0.219170 | 0.233599 | 0.238923 | 0.381683 | 0.419601 | 0.561534 | 0.399430 | 0.303139 | 0.226679 | 0.215873 | 0.230641 | 0.281646 | 0.321018 | 0.500394 | 0.565283 | 0.610513 | 0.746082 | 2.435664 | 5.372147 | 2.131875 | 0.534810 | 0.284999 | 0.217420 | 0.264281 | 0.454816 | 1.430236 | 1.584340 | 0.691254 | 0.577099 | 0.538245 | 0.482238 | 0.773662 | 0.585674 | 0.545559 | 0.642945 | 0.754762 | 0.703273 | 0.701851 | 0.353591 | 0.295656 | 0.368805 | 0.442528 | 0.718294 | 0.974073 | 0.647772 | 0.914640 | 1.282901 | 0.677513 | 0.533147 | 0.249376 | 0.216224 | 0.216100 | 0.234126 | 0.287527 | 0.353358 | 0.550375 | 0.707928 | 1.107774e+00 | 8.776582e-01 | 9.549388e-01 | 8.133322e-01 | 9.826051e-01 | 1.041745 | 0.747626 | 0.511897 | 0.471894 | 0.324953 | 0.243559 | 0.217594 | 0.239026 | 0.255472 | 0.222625 | 0.214820 | 0.221613 | 0.265672 | 0.443324 | 0.817599 | 1.407234 | 1.888609 | 2.946020 | 4.452240 | 6.613554 | 7.579005 | 5.162498 | 3.847790 | 4.056054 | 3.002024 | 2.043803 | 1.240550 | 0.797158 | 0.779727 | 0.725581 | 0.688304 | 0.711178 | 1.559855 | 5.002813 | 12.411258 | 20.030928 | 31.340807 | 32.878376 | 34.518186 | 16.524372 | 7.073238 | 2.944436 | 2.302802 | 2.346148 | 2.465113 | 1.932016 | 2.060617 | 1.805866 | 2.888792 | 4.252568 | 4.827203 | 3.700885 | 2.342272 | 1.580289 | 1.467917 | 1.060563 | 1.086647 | 0.810101 | 0.968281 | 1.074188 | 0.833663 | 0.547688 | 0.420214 | 0.313281 | 0.284532 | 0.224828 | 0.224768 | 0.304278 | 0.347802 | 0.271140 | 0.286280 | 0.449530 | 0.589032 | 0.374752 | 0.269430 | 0.237917 | 0.301591 | 0.308651 | 0.237171 | 0.216249 | 0.218600 | 0.226230 | 0.289054 | 0.335829 | 0.398732 | 0.464923 | 0.543564 | 0.903134 | 0.998423 | 0.741796 | 0.542896 | 0.498913 | 0.509068 | 0.560070 | 0.411253 | 0.327678 | 0.257922 | 0.251425 | 0.334025 | 0.592981 | 0.755491 | 0.561563 | 0.536441 | 0.554078 | 0.483244 | 0.294161 | 0.238888 | 0.218015 | 0.218806 | 0.218049 | 0.216997 | 0.240111 | 0.313253 | 0.307176 | 0.335667 | 0.355079 | 0.330169 | 0.365686 | 0.283142 | 0.232990 | 0.223750 | 0.216617 | 0.217825 | 0.225542 | 0.275504 | 0.336069 | 0.354527 | 0.331306 | 0.250677 | 0.214648 | 0.224343 | 0.252877 | 0.254953 | 0.219165 | 0.214923 | 0.215338 | 0.222085 | 0.221690 | 0.218008 | 0.218604 | 0.215354 | 0.214775 | 0.215938 | 0.232097 | 0.283270 | 0.274982 | 0.224399 | 0.214753 | 0.260289 | 0.248313 | 0.236094 | 0.221999 | 0.215884 | 0.222662 | 0.222768 | 0.236165 | 0.225584 | 0.224970 | 0.263579 | 0.291726 | 0.398084 | 0.478764 | 0.511940 | 0.585010 | 0.473498 | 0.289991 | 0.275514 | 0.256289 | 0.256593 | 0.294185 | 0.329769 | 0.287023 | 0.262024 | 0.258974 | 0.274147 | 0.290638 | 0.318378 | 0.328155 | 0.471446 | 0.703681 | 0.770496 | 0.662893 | 0.681418 | 0.635176 | 0.726592 | 0.585595 | 0.533022 | 0.546713 | 0.612302 | 0.641331 | 0.617638 | 0.664248 | 0.824771 | 1.049812 | 0.974574 | 0.774371 | 0.472243 | 0.351563 | 0.240067 | 0.215303 | 0.223694 | 0.257875 | 0.332866 | 0.406449 | 0.452067 | 0.454425 | 0.455399 | 0.651395 |
| left central | 0.371929 | 0.394624 | 0.333935 | 0.451249 | 1.002993 | 2.881199 | 1.905098 | 0.759139 | 0.582863 | 1.067806 | 0.664774 | 0.396092 | 0.264166 | 0.355483 | 0.257562 | 0.217048 | 0.294886 | 0.370973 | 0.340255 | 0.291910 | 0.487888 | 0.357514 | 0.219357 | 0.240803 | 0.468367 | 0.612213 | 0.683179 | 0.684645 | 0.527319 | 0.247668 | 0.246149 | 0.673151 | 1.008980 | 0.819337 | 0.391231 | 0.262116 | 0.215759 | 0.224103 | 0.217578 | 0.229963 | 0.257710 | 0.245933 | 0.253355 | 0.270369 | 0.352176 | 0.780050 | 0.808415 | 0.492419 | 0.346331 | 0.255038 | 0.215902 | 0.227614 | 0.297293 | 0.347497 | 0.301848 | 0.267472 | 0.259947 | 0.234377 | 0.224958 | 0.215602 | 0.217706 | 0.226920 | 0.233960 | 0.245183 | 0.252167 | 0.319038 | 0.427370 | 5.955589e-01 | 7.523932e-01 | 1.114103e+00 | 2.234759e+00 | 1.403889e+01 | 61.205186 | 235.971824 | 525.022166 | 943.381702 | 1004.398518 | 721.659029 | 133.545898 | 32.451696 | 8.891761 | 2.389249 | 0.797910 | 0.534304 | 0.541449 | 0.880099 | 0.675762 | 0.554850 | 0.768989 | 1.340701 | 2.286563 | 2.372070 | 1.459393 | 1.674451 | 3.381191 | 4.945405 | 6.555176 | 2.516299 | 1.129310 | 0.728855 | 0.557584 | 0.322390 | 0.227842 | 0.216592 | 0.214900 | 0.214809 | 0.216929 | 0.218576 | 0.215731 | 0.231176 | 0.238295 | 0.278420 | 0.311403 | 0.369075 | 0.397349 | 0.282656 | 0.236128 | 0.222989 | 0.218520 | 0.243401 | 0.287920 | 0.347646 | 0.480269 | 0.579302 | 0.562826 | 0.466085 | 0.355500 | 0.266012 | 0.283233 | 0.318046 | 0.379372 | 0.457669 | 0.506187 | 0.511067 | 0.393050 | 0.258937 | 0.218447 | 0.215462 | 0.218139 | 0.221307 | 0.215135 | 0.217376 | 0.233424 | 0.266340 | 0.360297 | 0.657616 | 1.227275 | 0.807734 | 0.564151 | 0.435576 | 0.307440 | 0.231861 | 0.222484 | 0.220260 | 0.231013 | 0.250670 | 0.286862 | 0.343040 | 0.499496 | 0.446604 | 0.394176 | 0.357234 | 0.354913 | 0.314589 | 0.308761 | 0.314703 | 0.326307 | 0.349352 | 0.368482 | 0.324632 | 0.336392 | 0.369599 | 0.315774 | 0.312723 | 0.344899 | 0.394963 | 0.464471 | 0.549916 | 0.469290 | 0.549676 | 0.465301 | 0.308401 | 0.253906 | 0.230411 | 0.219551 | 0.216108 | 0.215596 | 0.214690 | 0.216707 | 0.216390 | 0.214626 | 0.214980 | 0.215007 | 0.217137 | 0.222133 | 0.230775 | 0.236304 | 0.228087 | 0.236146 | 0.242997 | 0.242963 | 0.224334 | 0.215751 | 0.215888 | 0.214701 | 0.216918 | 0.231072 | 0.240266 | 0.320780 | 0.542222 | 1.339969 | 2.514219 | 3.570984 | 4.334440 | 7.917663 | 7.373450 | 3.632806 | 1.525968 | 0.623517 | 0.422251 | 0.337397 | 0.275434 | 0.226668 | 0.216110 | 0.215668 | 0.217449 | 0.215665 | 0.214652 | 0.214686 | 0.214685 | 0.215724 | 0.214862 | 0.215713 | 0.222021 | 0.218973 | 0.214626 | 0.214842 | 0.214819 | 0.245075 | 0.267804 | 0.298984 | 0.285143 | 0.250005 | 0.251431 | 0.251489 | 0.231718 | 0.219973 | 0.225593 | 0.242153 | 0.248799 | 0.258374 | 0.247749 | 0.245378 | 0.234922 | 0.216095 | 0.214642 | 0.214841 | 0.214675 | 0.234932 | 0.359958 | 0.692485 | 1.103172 | 1.354470 | 1.154702 | 1.434676 | 1.529693 | 1.097369 | 0.592829 | 0.535367 | 0.462527 | 0.540436 | 0.536167 | 0.468981 | 0.472926 | 0.648815 | 0.830739 | 0.808267 | 0.798470 | 0.727770 | 0.695827 |
| right central | 0.425515 | 0.604123 | 0.814318 | 1.554150 | 2.134826 | 4.951757 | 15.664431 | 6.844277 | 11.834960 | 27.147633 | 20.669452 | 18.799101 | 4.578182 | 2.913228 | 2.415824 | 1.560095 | 1.280302 | 0.400260 | 0.216107 | 0.215545 | 0.218397 | 0.234721 | 0.232120 | 0.331670 | 0.342629 | 0.252953 | 0.226083 | 0.228907 | 0.220748 | 0.220828 | 0.262636 | 0.255572 | 0.259732 | 0.350783 | 0.276583 | 0.361560 | 0.214928 | 0.277325 | 0.418834 | 0.460779 | 0.497811 | 0.346671 | 0.285636 | 0.220031 | 0.216044 | 0.262473 | 0.256665 | 0.271393 | 0.243968 | 0.248083 | 0.247533 | 0.225291 | 0.215469 | 0.214626 | 0.216243 | 0.217131 | 0.231998 | 0.240523 | 0.251042 | 0.285423 | 0.221362 | 0.233639 | 0.340896 | 0.502680 | 0.957803 | 2.726929 | 7.033552 | 1.393574e+01 | 3.390177e+01 | 7.625893e+01 | 2.343083e+02 | 1.897242e+02 | 150.465989 | 48.191092 | 11.494188 | 4.970823 | 2.871380 | 1.983673 | 2.118559 | 4.653294 | 9.057422 | 17.796809 | 29.444867 | 47.989222 | 83.136650 | 122.412559 | 79.775285 | 137.104350 | 523.658714 | 2735.693452 | 10120.958046 | 10077.058388 | 20136.209072 | 30277.926564 | 13400.659470 | 3880.194572 | 743.613623 | 194.812883 | 110.555460 | 52.079240 | 19.839585 | 38.602387 | 60.322288 | 168.745088 | 218.256990 | 56.902060 | 63.245677 | 86.582050 | 48.978568 | 123.828687 | 122.210873 | 189.105493 | 270.605003 | 338.807577 | 314.215671 | 152.996827 | 99.991229 | 294.800533 | 356.597002 | 118.877276 | 63.606594 | 24.547561 | 10.330325 | 7.324240 | 5.395684 | 3.548266 | 2.666676 | 2.527240 | 1.750894 | 1.821979 | 1.529907 | 1.695958 | 1.586366 | 1.763922 | 1.535481 | 1.248160 | 0.704071 | 0.517808 | 0.388845 | 0.373577 | 0.286892 | 0.260644 | 0.231572 | 0.241809 | 0.250216 | 0.234714 | 0.225865 | 0.227465 | 0.225422 | 0.258607 | 0.347546 | 0.424293 | 0.478435 | 0.511820 | 0.542322 | 0.923005 | 0.851985 | 0.474943 | 0.363747 | 0.336599 | 0.373370 | 0.389562 | 0.280316 | 0.247795 | 0.219423 | 0.218316 | 0.271762 | 0.322173 | 0.374085 | 0.378899 | 0.300633 | 0.367670 | 0.343018 | 0.266893 | 0.235571 | 0.222417 | 0.233215 | 0.283954 | 0.356649 | 0.554484 | 0.998669 | 2.056490 | 2.341333 | 1.984107 | 1.248673 | 0.789312 | 0.645030 | 0.478306 | 0.348935 | 0.395756 | 0.450910 | 0.554363 | 0.879843 | 1.205078 | 1.381685 | 1.342705 | 0.857880 | 0.786625 | 0.859092 | 0.709794 | 0.782956 | 1.067817 | 1.382180 | 1.232013 | 0.783868 | 0.674774 | 0.440596 | 0.291445 | 0.240547 | 0.227843 | 0.238696 | 0.247366 | 0.247570 | 0.278553 | 0.305579 | 0.306979 | 0.295269 | 0.261704 | 0.227273 | 0.214626 | 0.227051 | 0.238557 | 0.228043 | 0.234348 | 0.227847 | 0.219004 | 0.217782 | 0.218207 | 0.222082 | 0.231239 | 0.222559 | 0.222528 | 0.214929 | 0.215084 | 0.219936 | 0.217528 | 0.214959 | 0.244809 | 0.274898 | 0.299384 | 0.268981 | 0.267565 | 0.227812 | 0.216666 | 0.215049 | 0.219747 | 0.232329 | 0.224754 | 0.217852 | 0.218121 | 0.229836 | 0.229256 | 0.223352 | 0.229619 | 0.219487 | 0.216733 | 0.225618 | 0.231983 | 0.238820 | 0.240636 | 0.247477 | 0.234730 | 0.214626 | 0.227306 | 0.256105 | 0.276260 | 0.281246 | 0.302591 | 0.313523 | 0.270220 | 0.249096 | 0.237092 | 0.254987 | 0.334557 | 0.376637 | 0.400341 | 0.419697 | 0.410376 | 0.411040 | 0.369947 |
| left posterior | 1.384911 | 0.922621 | 0.924255 | 1.844579 | 1.974061 | 1.529694 | 1.334873 | 0.594115 | 0.309858 | 0.226161 | 0.233367 | 0.308103 | 0.432762 | 0.428223 | 0.277338 | 0.214829 | 0.267722 | 1.260332 | 8.161993 | 53.091118 | 29.930599 | 4.620251 | 0.764114 | 0.287821 | 0.218808 | 0.317958 | 0.420879 | 0.307881 | 0.216333 | 0.216566 | 0.236758 | 0.306057 | 0.357911 | 0.277352 | 0.218711 | 0.250456 | 0.290700 | 0.318845 | 0.432458 | 0.579485 | 0.544543 | 0.704866 | 0.527381 | 0.301369 | 0.286463 | 0.268273 | 0.241409 | 0.237157 | 0.217608 | 0.214978 | 0.221387 | 0.266792 | 0.365116 | 0.683567 | 1.004305 | 0.981672 | 0.908114 | 0.442461 | 0.252322 | 0.215472 | 0.279757 | 0.463194 | 1.435817 | 8.833935 | 113.363800 | 5698.826677 | 165955.893156 | 1.955619e+06 | 1.047231e+07 | 9.525794e+06 | 6.649319e+06 | 3.872849e+06 | 920200.933408 | 238146.262074 | 28517.393736 | 7246.220709 | 3042.648273 | 965.407169 | 736.293823 | 2517.629443 | 12915.082553 | 7148.242370 | 577.214596 | 201.110120 | 101.272959 | 49.902761 | 38.830884 | 43.472609 | 54.944086 | 124.443979 | 158.679104 | 165.638772 | 228.440807 | 255.314388 | 182.999386 | 197.506448 | 181.915496 | 145.316886 | 113.540840 | 85.926616 | 50.754093 | 35.097679 | 30.944342 | 43.704746 | 52.977250 | 46.880771 | 34.381053 | 28.885656 | 28.923645 | 20.847433 | 12.661766 | 7.877352 | 6.563098 | 7.815452 | 8.746557 | 13.605124 | 20.091684 | 29.436170 | 58.598487 | 144.755804 | 245.865323 | 309.519695 | 234.649253 | 162.780699 | 98.683851 | 46.262279 | 29.402346 | 14.924412 | 8.065050 | 3.958396 | 3.017552 | 2.037227 | 1.743476 | 1.434508 | 1.719338 | 2.726572 | 3.594660 | 4.124740 | 5.448331 | 5.830003 | 4.954490 | 4.903706 | 3.025797 | 2.555493 | 2.285248 | 1.742586 | 1.555584 | 1.650893 | 1.297683 | 1.105106 | 1.087573 | 0.728796 | 0.578027 | 0.544621 | 0.515562 | 0.531808 | 0.575823 | 0.686012 | 1.121489 | 1.773044 | 2.079856 | 3.267687 | 4.183154 | 4.993367 | 3.528052 | 2.716883 | 1.561154 | 1.209753 | 0.952103 | 0.764684 | 0.576128 | 0.591700 | 0.561096 | 0.766149 | 0.929866 | 0.844135 | 0.941422 | 1.006744 | 1.006106 | 1.000428 | 0.849383 | 0.745612 | 0.743161 | 0.688796 | 0.674258 | 0.571497 | 0.574676 | 0.624390 | 0.607420 | 0.502523 | 0.441171 | 0.417418 | 0.527661 | 0.546061 | 0.522634 | 0.560384 | 0.661624 | 0.744675 | 0.922920 | 0.855474 | 0.941309 | 1.094620 | 1.108977 | 1.113138 | 1.027568 | 0.938202 | 0.952252 | 1.075580 | 1.341801 | 1.832168 | 2.227811 | 2.756394 | 3.150642 | 3.284502 | 3.615064 | 4.315107 | 4.626224 | 5.293329 | 6.907455 | 7.786472 | 6.847693 | 6.037876 | 5.380137 | 4.906621 | 5.166195 | 5.221986 | 5.274390 | 5.885793 | 5.860121 | 5.778020 | 5.245631 | 4.544862 | 3.819935 | 3.525328 | 3.025846 | 2.884951 | 2.343610 | 2.111941 | 1.901587 | 2.109215 | 2.645092 | 3.351914 | 3.128185 | 3.080919 | 2.013248 | 1.512539 | 1.240711 | 1.128336 | 1.337944 | 1.846590 | 1.845425 | 2.100107 | 2.666673 | 2.226065 | 1.853732 | 1.305329 | 0.817647 | 0.732458 | 0.806355 | 0.683568 | 0.674766 | 0.581033 | 0.466349 | 0.391814 | 0.347043 | 0.297593 | 0.261194 | 0.233196 | 0.222602 | 0.216886 | 0.214769 | 0.221108 | 0.239613 | 0.255825 | 0.259058 | 0.264977 | 0.268217 | 0.251523 | 0.250015 | 0.250987 |
| right posterior | 0.240519 | 0.274228 | 0.315589 | 0.420704 | 0.994840 | 1.695303 | 0.898482 | 0.423518 | 0.334316 | 0.302072 | 0.269272 | 0.222744 | 0.214986 | 0.216970 | 0.217636 | 0.233311 | 0.293007 | 0.369818 | 0.553285 | 0.517364 | 0.589684 | 0.495245 | 0.373225 | 0.230579 | 0.214723 | 0.214674 | 0.219283 | 0.233552 | 0.340542 | 0.269110 | 0.279701 | 0.363563 | 0.324147 | 0.279285 | 0.226280 | 0.265301 | 0.531495 | 1.296673 | 2.367576 | 2.936524 | 3.182697 | 2.689136 | 1.184911 | 1.153226 | 0.896528 | 0.985998 | 0.663041 | 0.285117 | 0.215042 | 0.218943 | 0.273681 | 0.287838 | 0.233103 | 0.214720 | 0.236022 | 0.363386 | 0.467612 | 0.894990 | 2.680527 | 2.536713 | 2.029634 | 2.723920 | 4.575246 | 12.106084 | 18.112414 | 32.965939 | 129.733083 | 4.001794e+02 | 1.500469e+03 | 2.956489e+03 | 5.496674e+03 | 1.604936e+04 | 19286.972786 | 9074.750033 | 1508.486018 | 164.352735 | 37.741692 | 9.255252 | 3.250725 | 1.696207 | 1.678835 | 3.608543 | 3.919113 | 5.215536 | 5.504903 | 5.707446 | 8.098835 | 13.071858 | 15.407688 | 41.538168 | 130.498883 | 348.422240 | 545.420908 | 652.207395 | 1077.366139 | 1060.675278 | 427.400064 | 163.137133 | 129.877092 | 97.428297 | 67.912750 | 31.481984 | 26.891273 | 31.738211 | 39.316061 | 56.249507 | 127.339788 | 393.140166 | 1023.211894 | 2745.618144 | 3694.150487 | 1429.111644 | 538.459732 | 284.984902 | 174.081842 | 140.415996 | 148.785998 | 233.608927 | 388.895747 | 931.589548 | 1194.179686 | 1461.000372 | 1432.586259 | 1038.420250 | 405.723330 | 307.845117 | 204.699606 | 123.809994 | 58.865271 | 27.432189 | 16.673966 | 16.808091 | 26.157964 | 49.852273 | 57.705917 | 40.798449 | 35.256654 | 30.104702 | 39.465402 | 14.914070 | 8.924480 | 8.723689 | 12.188767 | 13.604379 | 20.301463 | 15.391190 | 19.954928 | 12.861719 | 7.149079 | 4.262714 | 3.195554 | 2.645896 | 2.926156 | 2.394802 | 2.402054 | 2.966287 | 3.947394 | 5.623439 | 5.862750 | 4.710857 | 4.625004 | 5.008048 | 5.601725 | 6.059921 | 6.872051 | 7.675269 | 10.025091 | 12.128899 | 14.766359 | 11.684892 | 9.089440 | 6.440514 | 3.604821 | 2.095600 | 1.137194 | 0.991825 | 1.271842 | 2.413763 | 4.877992 | 11.062116 | 13.975334 | 14.418005 | 10.749560 | 6.781522 | 4.378483 | 3.424996 | 2.828972 | 3.276624 | 4.134726 | 4.581180 | 4.276686 | 5.344876 | 5.952293 | 6.202742 | 4.682361 | 4.950195 | 7.276649 | 9.002267 | 7.447678 | 6.557819 | 6.319025 | 8.741390 | 6.430122 | 5.042730 | 5.765495 | 4.313036 | 3.173010 | 2.598918 | 2.734313 | 3.024958 | 3.447457 | 3.639639 | 6.714832 | 11.099201 | 21.551692 | 28.807285 | 62.437849 | 109.064397 | 154.815620 | 65.637738 | 33.123931 | 23.109199 | 18.340213 | 15.443190 | 12.699795 | 9.347189 | 10.837092 | 12.855164 | 22.288032 | 24.814689 | 21.347313 | 19.965769 | 19.658249 | 16.788098 | 16.178863 | 14.538981 | 15.379023 | 19.844450 | 21.820706 | 21.884462 | 20.839197 | 20.572502 | 17.791765 | 20.265182 | 22.793328 | 24.271260 | 27.358859 | 22.384978 | 20.058187 | 15.023599 | 10.533641 | 4.946076 | 4.430022 | 3.401382 | 2.722958 | 2.473748 | 2.669310 | 2.988857 | 3.313850 | 2.701031 | 2.317786 | 2.681504 | 3.133600 | 2.986697 | 2.135783 | 1.715812 | 1.298120 | 1.101086 | 1.006291 | 0.742985 | 0.691008 | 0.812686 | 0.951546 | 1.418561 | 1.266388 | 1.049838 | 1.053439 | 1.055731 | 1.188042 | 1.244164 |
| all electrodes | 0.224083 | 0.219415 | 0.227812 | 0.273053 | 0.424627 | 0.409044 | 1.415435 | 6.882095 | 20.791558 | 2.868254 | 1.037271 | 0.504660 | 0.422723 | 0.308092 | 0.317195 | 0.351711 | 0.700328 | 1.009908 | 1.503226 | 1.397299 | 2.163940 | 1.412931 | 0.395147 | 0.214689 | 0.301631 | 0.496121 | 0.622626 | 2.198921 | 3.091124 | 0.935078 | 0.459694 | 0.263942 | 0.215002 | 0.216939 | 0.219020 | 0.221706 | 0.220885 | 0.217603 | 0.221872 | 0.237452 | 0.306977 | 0.538571 | 1.664523 | 6.367723 | 9.407828 | 10.797795 | 7.943380 | 3.006071 | 1.518219 | 0.645330 | 0.477899 | 0.385654 | 0.308486 | 0.285709 | 0.266802 | 0.243720 | 0.246743 | 0.222328 | 0.214889 | 0.223735 | 0.247990 | 0.314794 | 0.460421 | 0.750852 | 1.438890 | 3.983504 | 5.950701 | 1.048037e+01 | 3.871028e+01 | 8.568445e+01 | 2.342408e+02 | 6.167901e+02 | 2623.790953 | 2137.718770 | 3758.122138 | 5913.650459 | 9422.084915 | 15870.485217 | 7965.854998 | 3163.575674 | 1628.822934 | 109.014514 | 11.448689 | 4.005475 | 2.344078 | 1.466748 | 0.861516 | 0.543660 | 0.877050 | 2.413084 | 8.079460 | 22.302101 | 25.681140 | 41.085076 | 100.402962 | 108.576561 | 172.764646 | 171.495652 | 104.101404 | 67.474657 | 24.544441 | 13.704349 | 10.320072 | 3.447165 | 2.769553 | 1.751836 | 1.694054 | 1.441429 | 1.188344 | 2.169775 | 4.516261 | 4.007573 | 5.973729 | 5.823326 | 8.788075 | 7.765432 | 10.793130 | 13.764636 | 23.465892 | 34.436992 | 37.306613 | 34.282219 | 38.333360 | 23.295189 | 8.572661 | 3.008157 | 1.376146 | 0.902114 | 1.019172 | 1.145635 | 1.407840 | 1.831131 | 1.716040 | 1.687593 | 1.944640 | 1.025195 | 0.804712 | 0.847824 | 1.312930 | 2.695408 | 4.173129 | 5.912487 | 12.232143 | 17.283965 | 8.944024 | 2.970086 | 1.360922 | 0.711158 | 0.520550 | 0.404283 | 0.340615 | 0.279771 | 0.282359 | 0.323729 | 0.380838 | 0.405290 | 0.376087 | 0.374997 | 0.546162 | 0.732144 | 0.650519 | 0.547782 | 0.410360 | 0.422151 | 0.401673 | 0.356442 | 0.352496 | 0.356696 | 0.377473 | 0.392695 | 0.306811 | 0.233718 | 0.216852 | 0.216208 | 0.221863 | 0.217273 | 0.218877 | 0.238899 | 0.324834 | 0.381777 | 0.457483 | 0.434619 | 0.358630 | 0.298034 | 0.307748 | 0.312037 | 0.327531 | 0.306074 | 0.274041 | 0.249880 | 0.243931 | 0.250748 | 0.244079 | 0.252151 | 0.268269 | 0.367611 | 0.524912 | 0.901759 | 0.928651 | 0.969245 | 1.213769 | 1.025884 | 0.876456 | 0.934532 | 0.777119 | 0.987741 | 0.807548 | 0.534791 | 0.752252 | 0.949595 | 1.096378 | 1.094728 | 0.970470 | 1.474051 | 2.763541 | 1.900977 | 1.486676 | 0.863531 | 0.758266 | 0.771265 | 0.549284 | 0.386301 | 0.401661 | 0.475843 | 0.632173 | 0.666968 | 0.514652 | 0.449635 | 0.555898 | 0.689151 | 0.854185 | 0.932013 | 0.992046 | 1.314225 | 2.037652 | 2.349501 | 1.966698 | 1.664500 | 1.460163 | 1.558417 | 1.821431 | 1.502495 | 1.522527 | 1.477318 | 1.350717 | 1.060878 | 1.072899 | 0.843140 | 0.582839 | 0.438403 | 0.420974 | 0.435778 | 0.517730 | 0.493344 | 0.540013 | 0.786729 | 0.892343 | 0.671697 | 0.673581 | 0.805628 | 1.184652 | 1.186833 | 0.903584 | 0.718458 | 0.780748 | 0.584485 | 0.379490 | 0.249757 | 0.219185 | 0.214661 | 0.215084 | 0.220741 | 0.236159 | 0.246498 | 0.277727 | 0.360012 | 0.481868 | 0.717706 | 0.817040 | 0.796023 |

Searchlight, spatiotemporal cluster permutation test

|  | start time | stop time | peak time | peak channel | cluster p | peak Cohen's d | direction |
| --- | --- | --- | --- | --- | --- | --- | --- |
| #1 | 80 | 1145 | 150 | PO7 | 0.0001 | 1.64574 | positive |

D) neutral vs happy

  
|  | time window | peak latency | cluster *p* | peak Cohen's *d* |  | | | |  |
| **all electrodes** | 250 - 365 ms | 280 ms | 0.0271 | 0.8811 |  | | | |  |
 785 - 890 ms | 820 ms | 0.0427 | 0.5888 |  | | | ||  |  | | | | | | | | |

Time-resolved classification, cluster permutation tests

|  | **left hemisphere** | | | | **right hemisphere** | | | |  |
|  | time window | peak latency | cluster *p* | peak Cohen's *d* | time window | peak latency | cluster *p* | peak Cohen's *d* |  |
| **anterior** |  | | | | 235 - 315 ms | 280 ms | 0.0375 | 0.9373 |  |
| **central** |  | | | | 250 - 370 ms | 305 ms | 0.0209 | 0.6268 |  |
| **posterior** | 235 - 430 ms | 280 ms | 0.0084 | 1.1 | 105 - 165 ms | 135 ms | 0.0449 | 1.1806 |  |
  | | | | 245 - 495 ms | 280 ms | 0.0053 | 0.9535 |  | | | | 770 - 1005 ms | 875 ms | 0.0082 | 0.7442 |

  

Time-resolved classification, Bayesian statistics

|  | -200 | -195 | -190 | -185 | -180 | -175 | -170 | -165 | -160 | -155 | -150 | -145 | -140 | -135 | -130 | -125 | -120 | -115 | -110 | -105 | -100 | -95 | -90 | -85 | -80 | -75 | -70 | -65 | -60 | -55 | -50 | -45 | -40 | -35 | -30 | -25 | -20 | -15 | -10 | -5 | 0 | 5 | 10 | 15 | 20 | 25 | 30 | 35 | 40 | 45 | 50 | 55 | 60 | 65 | 70 | 75 | 80 | 85 | 90 | 95 | 100 | 105 | 110 | 115 | 120 | 125 | 130 | 135 | 140 | 145 | 150 | 155 | 160 | 165 | 170 | 175 | 180 | 185 | 190 | 195 | 200 | 205 | 210 | 215 | 220 | 225 | 230 | 235 | 240 | 245 | 250 | 255 | 260 | 265 | 270 | 275 | 280 | 285 | 290 | 295 | 300 | 305 | 310 | 315 | 320 | 325 | 330 | 335 | 340 | 345 | 350 | 355 | 360 | 365 | 370 | 375 | 380 | 385 | 390 | 395 | 400 | 405 | 410 | 415 | 420 | 425 | 430 | 435 | 440 | 445 | 450 | 455 | 460 | 465 | 470 | 475 | 480 | 485 | 490 | 495 | 500 | 505 | 510 | 515 | 520 | 525 | 530 | 535 | 540 | 545 | 550 | 555 | 560 | 565 | 570 | 575 | 580 | 585 | 590 | 595 | 600 | 605 | 610 | 615 | 620 | 625 | 630 | 635 | 640 | 645 | 650 | 655 | 660 | 665 | 670 | 675 | 680 | 685 | 690 | 695 | 700 | 705 | 710 | 715 | 720 | 725 | 730 | 735 | 740 | 745 | 750 | 755 | 760 | 765 | 770 | 775 | 780 | 785 | 790 | 795 | 800 | 805 | 810 | 815 | 820 | 825 | 830 | 835 | 840 | 845 | 850 | 855 | 860 | 865 | 870 | 875 | 880 | 885 | 890 | 895 | 900 | 905 | 910 | 915 | 920 | 925 | 930 | 935 | 940 | 945 | 950 | 955 | 960 | 965 | 970 | 975 | 980 | 985 | 990 | 995 | 1000 | 1005 | 1010 | 1015 | 1020 | 1025 | 1030 | 1035 | 1040 | 1045 | 1050 | 1055 | 1060 | 1065 | 1070 | 1075 | 1080 | 1085 | 1090 | 1095 | 1100 | 1105 | 1110 | 1115 | 1120 | 1125 | 1130 | 1135 | 1140 | 1145 | 1150 | 1155 | 1160 | 1165 | 1170 | 1175 | 1180 | 1185 | 1190 | 1195 |
| --- | --- | --- | --- | --- | --- | --- | --- | --- | --- | --- | --- | --- | --- | --- | --- | --- | --- | --- | --- | --- | --- | --- | --- | --- | --- | --- | --- | --- | --- | --- | --- | --- | --- | --- | --- | --- | --- | --- | --- | --- | --- | --- | --- | --- | --- | --- | --- | --- | --- | --- | --- | --- | --- | --- | --- | --- | --- | --- | --- | --- | --- | --- | --- | --- | --- | --- | --- | --- | --- | --- | --- | --- | --- | --- | --- | --- | --- | --- | --- | --- | --- | --- | --- | --- | --- | --- | --- | --- | --- | --- | --- | --- | --- | --- | --- | --- | --- | --- | --- | --- | --- | --- | --- | --- | --- | --- | --- | --- | --- | --- | --- | --- | --- | --- | --- | --- | --- | --- | --- | --- | --- | --- | --- | --- | --- | --- | --- | --- | --- | --- | --- | --- | --- | --- | --- | --- | --- | --- | --- | --- | --- | --- | --- | --- | --- | --- | --- | --- | --- | --- | --- | --- | --- | --- | --- | --- | --- | --- | --- | --- | --- | --- | --- | --- | --- | --- | --- | --- | --- | --- | --- | --- | --- | --- | --- | --- | --- | --- | --- | --- | --- | --- | --- | --- | --- | --- | --- | --- | --- | --- | --- | --- | --- | --- | --- | --- | --- | --- | --- | --- | --- | --- | --- | --- | --- | --- | --- | --- | --- | --- | --- | --- | --- | --- | --- | --- | --- | --- | --- | --- | --- | --- | --- | --- | --- | --- | --- | --- | --- | --- | --- | --- | --- | --- | --- | --- | --- | --- | --- | --- | --- | --- | --- | --- | --- | --- | --- | --- | --- | --- | --- | --- | --- | --- | --- | --- | --- | --- | --- | --- | --- | --- | --- | --- | --- | --- | --- | --- | --- | --- | --- | --- | --- | --- | --- | --- | --- | --- | --- | --- |
| left anterior | 0.216767 | 0.216048 | 0.214675 | 0.217082 | 0.217561 | 0.221085 | 0.228660 | 0.248464 | 0.277899 | 0.305419 | 0.299950 | 0.301791 | 0.324614 | 0.250937 | 0.429086 | 1.064167 | 0.571086 | 0.309689 | 0.232631 | 0.228170 | 0.214963 | 0.235057 | 0.503593 | 0.524926 | 0.268483 | 0.216058 | 0.345728 | 0.444030 | 0.622915 | 0.316967 | 0.220250 | 0.231331 | 0.761357 | 1.765477 | 4.409968 | 20.444874 | 12.622644 | 3.231314 | 1.183760 | 0.430640 | 0.435837 | 0.683222 | 0.657741 | 0.551650 | 1.090455 | 2.634807 | 5.470011 | 15.700521 | 72.366532 | 152.201998 | 281.293650 | 297.819619 | 49.018432 | 13.860173 | 2.630078 | 0.601707 | 0.508729 | 0.642211 | 0.680377 | 0.578257 | 0.465260 | 0.404872 | 0.289352 | 0.221269 | 0.221876 | 0.229914 | 0.217658 | 0.215585 | 0.232141 | 0.373036 | 0.723974 | 1.136720 | 1.674323 | 2.502587 | 5.077839 | 9.922415 | 7.344921 | 8.673078 | 16.873356 | 76.366892 | 178.096465 | 68.357381 | 13.727746 | 2.909200 | 0.763753 | 0.458889 | 0.236160 | 0.220425 | 0.312276 | 0.506499 | 1.083515 | 2.166115 | 4.732873 | 6.323379 | 8.278512 | 12.279212 | 6.802059 | 3.031392 | 1.679500 | 1.037071 | 0.727315 | 0.588931 | 0.341389 | 0.237568 | 0.221744 | 0.216352 | 0.214937 | 0.216896 | 0.219694 | 0.216611 | 0.327294 | 0.436830 | 0.399531 | 0.358325 | 0.310237 | 0.260595 | 0.232434 | 0.214701 | 0.220374 | 0.215876 | 0.215691 | 0.225025 | 0.238434 | 0.249774 | 0.283243 | 0.325742 | 0.361544 | 0.414093 | 0.501456 | 0.585338 | 0.730329 | 0.802422 | 0.810127 | 0.821328 | 0.797517 | 0.941041 | 0.802490 | 0.672214 | 0.622559 | 0.580270 | 0.435827 | 0.372602 | 0.313571 | 0.350083 | 0.348250 | 0.304705 | 0.249829 | 0.241893 | 0.221730 | 0.217485 | 0.214719 | 0.215009 | 0.215272 | 0.218425 | 0.232789 | 0.246336 | 0.238088 | 0.223420 | 0.222901 | 0.218797 | 0.216204 | 0.236022 | 0.309079 | 0.339304 | 0.350093 | 0.351839 | 0.370895 | 0.324889 | 0.302281 | 0.272749 | 0.265926 | 0.245227 | 0.233252 | 0.219976 | 0.222493 | 0.223083 | 0.222454 | 0.223517 | 0.248979 | 0.270487 | 0.297625 | 0.355351 | 0.491828 | 0.756502 | 1.171222 | 1.661094 | 3.192411 | 5.739475 | 6.184555 | 4.968234 | 3.927312 | 3.381093 | 2.088695 | 1.560371 | 1.076187 | 1.095513 | 0.941868 | 0.846464 | 0.841570 | 0.952378 | 0.772911 | 0.839308 | 0.849264 | 0.876619 | 0.925347 | 0.724794 | 0.657053 | 0.732689 | 0.788666 | 0.831421 | 0.960572 | 1.103929 | 2.007256 | 2.109300 | 2.315262 | 2.056502 | 1.395739 | 1.035159 | 0.827647 | 0.617627 | 0.620465 | 0.664601 | 0.768073 | 0.944719 | 0.947154 | 1.007570 | 1.000512 | 1.032421 | 0.767452 | 0.583394 | 0.418951 | 0.467685 | 0.376592 | 0.258843 | 0.221765 | 0.224810 | 0.240324 | 0.298989 | 0.265929 | 0.247353 | 0.276221 | 0.379707 | 0.452571 | 0.391120 | 0.281097 | 0.318148 | 0.410164 | 0.453876 | 0.454049 | 0.356610 | 0.412426 | 0.714206 | 0.961144 | 1.130113 | 1.232228 | 1.280600 | 1.556824 | 1.035590 | 0.873490 | 0.631439 | 0.598088 | 0.570266 | 0.549739 | 0.448761 | 0.459382 | 0.437961 | 0.434762 | 0.358517 | 0.326100 | 0.280538 | 0.246723 | 0.230295 | 0.221512 | 0.216990 | 0.217870 | 0.219232 | 0.217897 | 0.222879 | 0.230098 | 0.233854 |
| right anterior | 0.645642 | 0.655214 | 0.530476 | 0.526128 | 0.426506 | 0.294919 | 0.249451 | 0.216664 | 0.215169 | 0.218928 | 0.293683 | 0.481790 | 1.097306 | 1.995741 | 0.823910 | 0.336402 | 0.234858 | 0.215492 | 0.219846 | 0.224884 | 0.265146 | 0.226834 | 0.216063 | 0.216821 | 0.219664 | 0.284152 | 0.405284 | 0.646602 | 1.636684 | 2.745762 | 1.679914 | 0.459794 | 0.214643 | 0.245736 | 0.461797 | 0.799553 | 0.905588 | 0.568996 | 0.360259 | 0.225317 | 0.222579 | 0.238044 | 0.270971 | 0.374366 | 0.673314 | 0.704159 | 0.840024 | 0.682759 | 0.464073 | 0.519567 | 0.442122 | 0.251298 | 0.242731 | 0.229143 | 0.214689 | 0.222646 | 0.220265 | 0.215179 | 0.227636 | 0.238640 | 0.233120 | 0.233058 | 0.249518 | 0.216819 | 0.242389 | 0.474607 | 0.772161 | 1.071584 | 0.866827 | 0.816662 | 0.510605 | 0.395212 | 0.329320 | 0.267903 | 0.223643 | 0.225549 | 0.234500 | 0.218069 | 0.219583 | 0.255069 | 0.265451 | 0.242296 | 0.220848 | 0.214851 | 0.255504 | 0.474686 | 1.197196 | 2.899043 | 6.676461 | 19.624265 | 137.632762 | 952.353671 | 1146.460785 | 906.325433 | 359.176287 | 284.790791 | 215.843113 | 71.207978 | 16.204841 | 6.888845 | 4.152977 | 3.889742 | 2.168939 | 1.525231 | 1.264094 | 1.213244 | 1.283397 | 1.461198 | 1.233313 | 2.223732 | 2.832682 | 2.984848 | 3.465155 | 2.735954 | 1.697774 | 1.827263 | 1.269846 | 1.138509 | 1.287659 | 1.243982 | 1.297895 | 1.887013 | 1.747877 | 1.444328 | 1.321665 | 1.791382 | 1.462797 | 1.309239 | 0.987776 | 0.821529 | 1.052581 | 1.209333 | 0.889523 | 1.149753 | 1.590862 | 2.733644 | 3.486039 | 1.655674 | 0.901086 | 0.484191 | 0.358448 | 0.312337 | 0.273611 | 0.234970 | 0.229560 | 0.223706 | 0.234984 | 0.234910 | 0.223198 | 0.214963 | 0.214708 | 0.216729 | 0.225915 | 0.238497 | 0.235221 | 0.224224 | 0.218908 | 0.217684 | 0.215405 | 0.214652 | 0.220934 | 0.222612 | 0.222720 | 0.230717 | 0.251899 | 0.276755 | 0.310271 | 0.278628 | 0.287602 | 0.260025 | 0.239338 | 0.226913 | 0.222997 | 0.231248 | 0.243512 | 0.263182 | 0.287470 | 0.286851 | 0.266792 | 0.253640 | 0.247188 | 0.233397 | 0.215778 | 0.214653 | 0.214731 | 0.217191 | 0.230248 | 0.237473 | 0.324203 | 0.661826 | 1.209556 | 1.820212 | 0.916677 | 0.592183 | 0.742041 | 0.810611 | 1.065879 | 1.307281 | 1.664345 | 3.854752 | 4.829327 | 3.917699 | 2.281637 | 1.568151 | 1.562328 | 1.221333 | 1.024029 | 1.121180 | 0.889359 | 0.721647 | 0.538844 | 0.369759 | 0.323849 | 0.307267 | 0.293320 | 0.281182 | 0.309023 | 0.322619 | 0.369823 | 0.370719 | 0.314923 | 0.384090 | 0.453553 | 0.643833 | 1.211443 | 1.428823 | 1.528661 | 2.064870 | 1.478770 | 1.596783 | 1.269418 | 0.991386 | 0.969107 | 0.986723 | 0.964501 | 1.018844 | 0.703566 | 0.614071 | 0.532888 | 0.539358 | 0.667402 | 0.825023 | 1.042810 | 1.271872 | 1.508078 | 1.700239 | 2.047379 | 1.647497 | 1.510734 | 1.219781 | 1.503627 | 1.333636 | 1.211344 | 0.942525 | 1.072332 | 1.255013 | 1.665528 | 1.925483 | 2.294569 | 3.093329 | 3.722368 | 3.312551 | 3.170639 | 3.380194 | 3.180999 | 2.492295 | 1.592691 | 1.152390 | 0.829909 | 0.532146 | 0.349724 | 0.262896 | 0.237061 | 0.219996 | 0.221027 | 0.221696 | 0.222876 | 0.230933 | 0.239311 | 0.235760 |
| left central | 1.318564 | 1.114589 | 1.123223 | 0.881754 | 1.048795 | 0.408062 | 0.236186 | 0.220581 | 0.237965 | 0.248774 | 0.265538 | 0.315742 | 0.392078 | 0.289440 | 0.228706 | 0.291541 | 1.250734 | 5.357191 | 1.649258 | 0.695492 | 0.522706 | 0.593499 | 0.550629 | 0.241417 | 0.224901 | 0.233678 | 0.577488 | 0.699442 | 0.699111 | 0.933072 | 0.680394 | 0.291214 | 0.232258 | 0.524107 | 0.733747 | 0.538376 | 0.405867 | 0.445008 | 0.321378 | 0.215768 | 0.221256 | 0.223557 | 0.219344 | 0.220242 | 0.230325 | 0.226229 | 0.215015 | 0.217653 | 0.216410 | 0.220071 | 0.253471 | 0.265340 | 0.277076 | 0.236736 | 0.216049 | 0.214952 | 0.218746 | 0.224526 | 0.217091 | 0.217413 | 0.269727 | 0.309015 | 0.298877 | 0.229798 | 0.216603 | 0.284936 | 0.492679 | 1.168621 | 1.175304 | 0.804435 | 0.391951 | 0.272897 | 0.233981 | 0.226110 | 0.227846 | 0.250005 | 0.275512 | 0.342999 | 0.361219 | 0.387871 | 0.456177 | 0.405721 | 0.305705 | 0.280106 | 0.247149 | 0.272839 | 0.251281 | 0.228278 | 0.293310 | 0.407712 | 0.495168 | 0.487375 | 0.457448 | 0.590260 | 0.796023 | 0.934586 | 1.589724 | 1.948309 | 3.505103 | 5.610861 | 5.352896 | 2.863740 | 1.324148 | 0.726494 | 0.611005 | 0.334537 | 0.247095 | 0.214938 | 0.218049 | 0.219425 | 0.240238 | 0.233802 | 0.236351 | 0.220197 | 0.217305 | 0.214634 | 0.214933 | 0.214764 | 0.214890 | 0.215386 | 0.224769 | 0.238339 | 0.250465 | 0.246455 | 0.242116 | 0.227702 | 0.241286 | 0.266106 | 0.317124 | 0.379346 | 0.469931 | 0.462355 | 0.477194 | 0.390880 | 0.280603 | 0.234793 | 0.223238 | 0.221290 | 0.227483 | 0.229610 | 0.223548 | 0.237657 | 0.263604 | 0.280544 | 0.243403 | 0.235023 | 0.235941 | 0.253894 | 0.244783 | 0.240953 | 0.243537 | 0.251927 | 0.254777 | 0.252319 | 0.242321 | 0.242952 | 0.236704 | 0.232765 | 0.241198 | 0.244587 | 0.241542 | 0.244098 | 0.235739 | 0.229115 | 0.225603 | 0.221047 | 0.215154 | 0.216812 | 0.225633 | 0.237861 | 0.246604 | 0.310459 | 0.361793 | 0.322036 | 0.296337 | 0.281392 | 0.263960 | 0.255970 | 0.243410 | 0.237701 | 0.233605 | 0.215683 | 0.214633 | 0.215604 | 0.219244 | 0.226933 | 0.221969 | 0.224482 | 0.223634 | 0.228073 | 0.222839 | 0.216281 | 0.216184 | 0.233024 | 0.242316 | 0.257099 | 0.297675 | 0.405369 | 0.535659 | 0.614707 | 0.547220 | 0.614172 | 0.619883 | 0.568469 | 0.566832 | 0.595285 | 0.694729 | 0.848060 | 0.830126 | 0.735051 | 0.657048 | 0.568516 | 0.476088 | 0.375424 | 0.293090 | 0.261007 | 0.237659 | 0.228939 | 0.220005 | 0.217213 | 0.215329 | 0.214633 | 0.215502 | 0.215810 | 0.214723 | 0.218305 | 0.221038 | 0.214651 | 0.215602 | 0.214995 | 0.216474 | 0.216854 | 0.226082 | 0.260459 | 0.322018 | 0.337376 | 0.272746 | 0.219855 | 0.221583 | 0.302437 | 0.465650 | 0.528178 | 0.492190 | 0.423732 | 0.345409 | 0.325229 | 0.274851 | 0.232330 | 0.221427 | 0.218358 | 0.214636 | 0.220403 | 0.238403 | 0.255103 | 0.268841 | 0.301950 | 0.328343 | 0.378545 | 0.493301 | 0.664695 | 0.726794 | 0.665293 | 0.545979 | 0.530207 | 0.483760 | 0.340934 | 0.279261 | 0.253846 | 0.254778 | 0.263225 | 0.253395 | 0.224334 | 0.221589 | 0.247996 | 0.329816 | 0.353144 | 0.371899 | 0.405198 | 0.574226 | 1.079360 |
| right central | 0.285595 | 0.277041 | 0.226085 | 0.214883 | 0.233121 | 0.241867 | 0.224363 | 0.237430 | 0.266032 | 0.241584 | 0.217462 | 0.218372 | 0.239148 | 0.225357 | 0.215197 | 0.316347 | 0.534433 | 0.382652 | 0.300739 | 0.375834 | 0.369650 | 0.310805 | 0.236669 | 0.215084 | 0.216722 | 0.225569 | 0.227523 | 0.282597 | 0.433272 | 0.461292 | 0.391550 | 0.348275 | 0.230923 | 0.227179 | 0.219989 | 0.217904 | 0.224479 | 0.228884 | 0.221027 | 0.218567 | 0.214927 | 0.220291 | 0.252058 | 0.297163 | 0.349036 | 0.413459 | 0.576385 | 0.551862 | 0.397229 | 0.316927 | 0.256674 | 0.222497 | 0.215206 | 0.234851 | 0.253706 | 0.260530 | 0.291471 | 0.292768 | 0.298158 | 0.393331 | 0.503367 | 0.490964 | 0.356379 | 0.219292 | 0.238316 | 0.334527 | 0.659216 | 1.096005 | 1.757719 | 1.288528 | 0.524166 | 0.337690 | 0.284639 | 0.258362 | 0.235518 | 0.224964 | 0.218023 | 0.217382 | 0.223769 | 0.243913 | 0.274541 | 0.338982 | 0.472846 | 0.912854 | 1.151636 | 0.903604 | 0.593314 | 0.538523 | 0.775447 | 1.231674 | 1.906242 | 2.974987 | 4.164521 | 6.228066 | 5.929230 | 6.317544 | 4.268603 | 2.599250 | 2.619868 | 3.422141 | 5.199055 | 8.139022 | 6.555481 | 4.239122 | 4.105535 | 3.984956 | 3.118757 | 2.754313 | 2.043026 | 2.693261 | 4.304583 | 4.211563 | 2.977702 | 2.551656 | 1.827701 | 1.137830 | 0.713096 | 0.637886 | 0.524000 | 0.591363 | 0.591799 | 0.536631 | 0.630339 | 0.670233 | 0.583498 | 0.425279 | 0.277228 | 0.235346 | 0.227332 | 0.219710 | 0.225190 | 0.238902 | 0.255225 | 0.289491 | 0.303340 | 0.289275 | 0.293928 | 0.290166 | 0.256755 | 0.248528 | 0.237281 | 0.223970 | 0.221203 | 0.222427 | 0.218899 | 0.215606 | 0.214861 | 0.216301 | 0.218198 | 0.229175 | 0.228002 | 0.229649 | 0.219174 | 0.215684 | 0.214771 | 0.214711 | 0.219645 | 0.260784 | 0.324047 | 0.367548 | 0.345507 | 0.343472 | 0.395400 | 0.444615 | 0.328161 | 0.267660 | 0.222018 | 0.217358 | 0.217742 | 0.238898 | 0.292203 | 0.344290 | 0.421173 | 0.524517 | 0.565930 | 0.527344 | 0.393881 | 0.351168 | 0.347494 | 0.314375 | 0.241475 | 0.223833 | 0.214725 | 0.215268 | 0.220288 | 0.214867 | 0.218329 | 0.257460 | 0.329628 | 0.354741 | 0.297681 | 0.241193 | 0.216761 | 0.226627 | 0.228401 | 0.220354 | 0.214989 | 0.214709 | 0.217343 | 0.227392 | 0.244175 | 0.297872 | 0.369736 | 0.377219 | 0.394802 | 0.432530 | 0.353229 | 0.266039 | 0.219628 | 0.214730 | 0.218877 | 0.237252 | 0.285295 | 0.281510 | 0.248258 | 0.231014 | 0.238419 | 0.235472 | 0.227497 | 0.217379 | 0.214967 | 0.214800 | 0.217209 | 0.242059 | 0.285705 | 0.322088 | 0.302202 | 0.286979 | 0.263489 | 0.241522 | 0.231928 | 0.233706 | 0.240413 | 0.230854 | 0.218380 | 0.214786 | 0.222257 | 0.264351 | 0.357164 | 0.391460 | 0.342632 | 0.288065 | 0.230180 | 0.215288 | 0.221699 | 0.251248 | 0.252563 | 0.241448 | 0.242028 | 0.250496 | 0.265539 | 0.280060 | 0.255074 | 0.236109 | 0.227985 | 0.219916 | 0.214704 | 0.216065 | 0.214635 | 0.230769 | 0.297013 | 0.403416 | 0.428778 | 0.330469 | 0.304784 | 0.240062 | 0.214640 | 0.218326 | 0.221786 | 0.214784 | 0.221144 | 0.221011 | 0.215098 | 0.216033 | 0.252429 | 0.335622 | 0.531246 | 0.737556 | 0.850132 | 0.880434 |
| left posterior | 0.368959 | 0.333695 | 0.474280 | 0.859855 | 1.317804 | 2.449029 | 1.051951 | 0.616205 | 0.461977 | 0.236702 | 0.226149 | 0.308238 | 0.385528 | 0.355998 | 0.331183 | 0.276276 | 0.226004 | 0.216077 | 0.338665 | 0.525668 | 1.125577 | 1.045502 | 0.327895 | 0.221554 | 0.224895 | 0.324832 | 0.685619 | 1.449417 | 1.765422 | 1.080037 | 0.785075 | 0.520027 | 0.352678 | 0.314369 | 0.295969 | 0.367949 | 0.399793 | 0.310541 | 0.289715 | 0.436821 | 0.735871 | 1.342652 | 1.509190 | 1.521156 | 2.580500 | 4.265208 | 3.390632 | 1.982434 | 0.744277 | 0.345082 | 0.250623 | 0.217799 | 0.242456 | 0.329880 | 0.448948 | 0.517787 | 0.543102 | 0.443439 | 0.414846 | 0.362969 | 0.272451 | 0.217398 | 0.623133 | 12.871802 | 288.830232 | 1963.042420 | 6478.828177 | 9900.398024 | 10261.675715 | 4461.456655 | 547.999727 | 40.693809 | 6.169385 | 1.280018 | 0.529255 | 0.331501 | 0.240116 | 0.223965 | 0.220462 | 0.228385 | 0.224493 | 0.216871 | 0.214635 | 0.219077 | 0.252586 | 0.453222 | 1.245792 | 4.455287 | 14.244519 | 46.124308 | 138.254949 | 276.701739 | 510.823746 | 995.853322 | 1359.615943 | 1782.749851 | 1280.708093 | 596.440046 | 263.322892 | 137.918689 | 71.190460 | 59.450612 | 39.771798 | 41.996450 | 57.684719 | 58.094988 | 50.091277 | 74.644403 | 87.279350 | 166.574029 | 212.955175 | 110.105504 | 69.231584 | 27.573199 | 9.197374 | 4.701743 | 2.424326 | 1.557084 | 1.689973 | 1.489199 | 1.445557 | 1.619151 | 2.045167 | 2.395422 | 1.862382 | 1.499942 | 1.417973 | 1.113358 | 0.760617 | 0.682622 | 0.544067 | 0.409685 | 0.312020 | 0.261899 | 0.267699 | 0.273405 | 0.257088 | 0.251221 | 0.259367 | 0.294481 | 0.398592 | 0.460057 | 0.464822 | 0.519542 | 0.572066 | 0.752169 | 0.875210 | 0.799793 | 0.711238 | 0.849164 | 0.867193 | 0.737869 | 0.611931 | 0.598780 | 0.666128 | 0.841448 | 0.826318 | 0.870298 | 0.887917 | 0.872812 | 0.836423 | 0.767708 | 0.825258 | 0.813897 | 0.657364 | 0.605424 | 0.601445 | 0.532150 | 0.472686 | 0.385930 | 0.376236 | 0.377406 | 0.371484 | 0.367268 | 0.434824 | 0.639061 | 0.906341 | 1.314961 | 2.037290 | 2.483863 | 2.092227 | 1.687052 | 1.324893 | 1.117615 | 0.817631 | 0.625912 | 0.496863 | 0.518276 | 0.437509 | 0.378589 | 0.363448 | 0.381196 | 0.387507 | 0.427185 | 0.385807 | 0.422377 | 0.492813 | 0.546749 | 0.657570 | 0.650413 | 0.705892 | 0.686749 | 0.799278 | 0.844316 | 0.831143 | 0.700571 | 0.640197 | 0.584388 | 0.715787 | 0.712776 | 0.762639 | 0.772018 | 0.792645 | 1.211856 | 1.817778 | 1.815559 | 1.577659 | 1.169303 | 0.884853 | 0.678097 | 0.420163 | 0.330709 | 0.345235 | 0.374510 | 0.455421 | 0.649872 | 0.941574 | 1.235390 | 1.650826 | 1.812190 | 1.776659 | 1.524965 | 0.947393 | 0.773307 | 0.663242 | 0.505775 | 0.387207 | 0.333664 | 0.289865 | 0.338682 | 0.380810 | 0.386433 | 0.377661 | 0.368819 | 0.308840 | 0.286851 | 0.253067 | 0.232315 | 0.237485 | 0.238016 | 0.222939 | 0.220314 | 0.219474 | 0.219528 | 0.214894 | 0.216793 | 0.221576 | 0.218570 | 0.214656 | 0.215162 | 0.215201 | 0.224682 | 0.233317 | 0.232555 | 0.231197 | 0.220340 | 0.216558 | 0.214706 | 0.215014 | 0.216204 | 0.220563 | 0.225788 | 0.226336 | 0.224086 | 0.220711 | 0.218864 | 0.222046 | 0.218144 | 0.215575 | 0.216334 |
| right posterior | 0.272185 | 0.264554 | 0.241370 | 0.216965 | 0.237048 | 0.268617 | 0.316584 | 0.283575 | 0.227685 | 0.215168 | 0.251681 | 0.327718 | 0.297459 | 0.382706 | 0.431722 | 0.469617 | 0.385083 | 0.285547 | 0.299568 | 0.357830 | 0.219744 | 0.215138 | 0.218535 | 0.219790 | 0.246147 | 0.218121 | 0.282094 | 0.216720 | 0.234372 | 0.222047 | 0.219718 | 0.253821 | 0.252704 | 0.225848 | 0.231333 | 0.216567 | 0.302083 | 0.745315 | 2.170881 | 10.589218 | 29.373579 | 55.185243 | 48.549190 | 47.737560 | 78.988144 | 124.423674 | 105.028986 | 33.102945 | 4.056110 | 0.785024 | 0.342369 | 0.243395 | 0.228535 | 0.244332 | 0.232632 | 0.252769 | 0.254556 | 0.303530 | 0.371715 | 0.433696 | 0.548501 | 2.392759 | 23.511031 | 442.926210 | 2990.107005 | 7174.718551 | 8632.904415 | 3075.954492 | 1070.045069 | 359.773228 | 157.313545 | 43.679250 | 8.701344 | 2.408646 | 1.088240 | 0.542191 | 0.270933 | 0.216073 | 0.214668 | 0.214668 | 0.219055 | 0.232169 | 0.261677 | 0.286758 | 0.333325 | 0.365892 | 0.499846 | 0.708755 | 1.103937 | 2.114110 | 7.810922 | 18.618382 | 32.853893 | 59.799706 | 138.164158 | 226.974421 | 257.744495 | 206.742067 | 187.697916 | 163.970323 | 155.870337 | 126.510178 | 174.559650 | 225.943261 | 362.646475 | 187.797258 | 126.570769 | 124.847352 | 127.136409 | 107.962914 | 97.329176 | 61.141616 | 59.425951 | 65.346961 | 58.827545 | 45.226500 | 35.020712 | 43.041887 | 58.095806 | 59.099491 | 52.189285 | 54.846134 | 51.207157 | 25.430831 | 7.004355 | 3.685940 | 3.665744 | 4.158287 | 2.524063 | 2.375484 | 2.546350 | 3.511637 | 3.829570 | 3.068473 | 2.194338 | 1.922155 | 1.640252 | 1.457367 | 1.589464 | 1.387932 | 0.916333 | 1.024210 | 1.402935 | 1.761485 | 2.100804 | 2.637725 | 3.705926 | 4.878808 | 2.835113 | 1.692719 | 1.219847 | 1.151154 | 1.003053 | 0.758040 | 0.691801 | 0.905413 | 1.274614 | 2.189422 | 2.357722 | 1.522285 | 1.500910 | 1.478328 | 1.148041 | 0.834385 | 0.657828 | 0.478510 | 0.424249 | 0.380251 | 0.333033 | 0.327458 | 0.331345 | 0.278510 | 0.270419 | 0.262485 | 0.260843 | 0.272968 | 0.263524 | 0.267025 | 0.303009 | 0.355745 | 0.412626 | 0.464454 | 0.461926 | 0.566108 | 0.669395 | 0.687292 | 0.684741 | 0.805669 | 0.957336 | 0.964905 | 0.718405 | 0.752619 | 0.935501 | 1.048759 | 1.415588 | 1.997592 | 3.427377 | 5.827244 | 3.660500 | 2.766100 | 2.682076 | 3.211801 | 4.468154 | 5.904692 | 8.080226 | 14.110263 | 17.144961 | 16.615136 | 12.046873 | 9.678022 | 7.408610 | 7.465310 | 9.147708 | 11.040786 | 17.913267 | 26.986326 | 31.616579 | 40.341783 | 34.620180 | 20.369874 | 13.108273 | 11.980277 | 11.956380 | 11.941102 | 14.086749 | 18.125575 | 28.337529 | 34.539171 | 22.519482 | 17.321519 | 17.600462 | 13.158980 | 10.993193 | 7.091622 | 4.381396 | 4.111429 | 3.862155 | 3.273456 | 2.697266 | 2.334300 | 1.751801 | 1.580049 | 1.277717 | 1.119066 | 1.114361 | 1.141613 | 1.077958 | 1.325364 | 1.718032 | 3.311366 | 3.970825 | 4.017678 | 4.070857 | 5.063816 | 3.146375 | 1.870778 | 1.142289 | 0.923145 | 0.911708 | 0.939146 | 0.942611 | 1.073581 | 1.361565 | 1.348923 | 1.448740 | 1.063148 | 0.638210 | 0.444974 | 0.401107 | 0.326996 | 0.312186 | 0.269666 | 0.289286 | 0.376705 | 0.399102 | 0.357269 | 0.379534 | 0.345429 | 0.382704 | 0.383810 |
| all electrodes | 0.315174 | 0.716610 | 1.077069 | 0.610526 | 0.569301 | 1.317011 | 0.709663 | 0.399778 | 0.267448 | 0.260568 | 0.261968 | 0.239004 | 0.215889 | 0.223377 | 0.218414 | 0.233734 | 0.215138 | 0.215583 | 0.300936 | 0.492024 | 0.369762 | 0.256493 | 0.243243 | 0.361708 | 1.065030 | 0.915607 | 1.326110 | 3.303804 | 4.747596 | 2.793814 | 0.585175 | 0.226566 | 0.250658 | 0.393034 | 0.469506 | 0.668887 | 0.550582 | 0.331053 | 0.257253 | 0.220517 | 0.257552 | 0.247853 | 0.270028 | 0.221562 | 0.231697 | 0.234888 | 0.261463 | 0.340894 | 0.584303 | 0.805513 | 1.018254 | 1.151694 | 3.253419 | 3.148844 | 1.996832 | 0.738414 | 0.481823 | 0.355757 | 0.274094 | 0.237656 | 0.222304 | 0.216769 | 0.233445 | 0.393642 | 1.678261 | 4.974541 | 21.415569 | 37.714438 | 47.726420 | 24.028333 | 6.798832 | 1.227227 | 0.465859 | 0.233092 | 0.222696 | 0.282186 | 0.362668 | 0.405828 | 0.384128 | 0.295755 | 0.227913 | 0.220092 | 0.230620 | 0.293137 | 0.333694 | 0.293297 | 0.254855 | 0.224574 | 0.214935 | 0.313121 | 1.484920 | 9.629940 | 24.688679 | 47.592368 | 56.570845 | 102.730877 | 116.933251 | 115.872608 | 105.288244 | 50.166733 | 28.454592 | 23.814716 | 6.100209 | 2.455097 | 1.881158 | 1.569406 | 2.267433 | 2.676737 | 1.943465 | 2.740712 | 3.954214 | 5.236018 | 4.229265 | 2.005404 | 1.135214 | 0.786729 | 0.596271 | 0.619182 | 0.596108 | 0.732348 | 0.800019 | 0.713288 | 0.787643 | 0.657314 | 0.487122 | 0.329676 | 0.246374 | 0.221826 | 0.220838 | 0.224280 | 0.247660 | 0.287145 | 0.330166 | 0.363232 | 0.482342 | 0.465732 | 0.397439 | 0.342491 | 0.257232 | 0.264119 | 0.262026 | 0.264754 | 0.306216 | 0.367910 | 0.364589 | 0.465983 | 0.470017 | 0.405544 | 0.281364 | 0.236177 | 0.223010 | 0.215434 | 0.215044 | 0.217006 | 0.224865 | 0.249513 | 0.258206 | 0.249610 | 0.260773 | 0.255721 | 0.239211 | 0.217286 | 0.217079 | 0.218195 | 0.221351 | 0.220536 | 0.217871 | 0.215374 | 0.225091 | 0.234455 | 0.224119 | 0.216315 | 0.214936 | 0.223915 | 0.241118 | 0.262686 | 0.255140 | 0.261766 | 0.238990 | 0.215237 | 0.232962 | 0.256411 | 0.278175 | 0.310959 | 0.354164 | 0.378768 | 0.338595 | 0.323381 | 0.363124 | 0.368253 | 0.324302 | 0.330593 | 0.376042 | 0.451727 | 0.421820 | 0.435431 | 0.683256 | 1.441454 | 2.268993 | 2.556788 | 3.752869 | 5.461761 | 5.994730 | 6.165587 | 5.625145 | 5.841095 | 7.163827 | 4.785802 | 4.947039 | 4.161023 | 2.460857 | 2.352603 | 2.837207 | 2.344567 | 2.526492 | 2.687762 | 2.274407 | 1.885237 | 1.409379 | 0.935235 | 1.209946 | 1.342874 | 0.957728 | 1.073320 | 1.898655 | 3.504425 | 6.210922 | 4.284563 | 2.751769 | 2.705882 | 1.987268 | 1.222439 | 0.756178 | 0.522482 | 0.423525 | 0.538578 | 0.646431 | 0.739663 | 0.636853 | 0.605405 | 0.710355 | 0.990537 | 0.872208 | 0.692824 | 0.560370 | 0.664912 | 0.650148 | 0.523325 | 0.374743 | 0.328058 | 0.362582 | 0.547867 | 0.873579 | 0.912903 | 0.819428 | 0.752229 | 0.580708 | 0.397506 | 0.274147 | 0.235844 | 0.263572 | 0.289167 | 0.318888 | 0.351410 | 0.333537 | 0.277500 | 0.233285 | 0.218672 | 0.221842 | 0.219931 | 0.219553 | 0.228788 | 0.253968 | 0.256636 | 0.225348 | 0.215151 | 0.214856 | 0.216119 | 0.218914 | 0.220593 |

Searchlight, spatiotemporal cluster permutation test

|  | start time | stop time | peak time | peak channel | cluster p | peak Cohen's d | direction |
| --- | --- | --- | --- | --- | --- | --- | --- |
| #1 | 215 | 1145 | 290 | P8 | 0.0005 | 1.164651 | positive |

E) neutral vs angry

  
|  | time window | peak latency | cluster *p* | peak Cohen's *d* |  | | | |
| **all electrodes** | 100 - 665 ms | 160 ms | 0.0001 | 2.106 |  | | | |
|  | | | | | | | | |

Time-resolved classification, cluster permutation tests

|  | **left hemisphere** | | | | **right hemisphere** | | | |
|  | time window | peak latency | cluster *p* | peak Cohen's *d* | time window | peak latency | cluster *p* | peak Cohen's *d* |
| **anterior** |  | | | |  | | | |
| **central** | 240 - 310 ms | 280 ms | 0.0379 | 1.0941 | 105 - 465 ms | 285 ms | 0.0001 | 1.5973 |
| **posterior** | 110 - 740 ms | 155 ms | 0.0001 | 1.7768 | 80 - 1195 ms | 150 ms | 0.0001 | 1.7856 |
 860 - 1070 ms | 900 ms | 0.0158 | 0.5718 |  | | | |

  

Time-resolved classification, Bayesian statistics

|  | -200 | -195 | -190 | -185 | -180 | -175 | -170 | -165 | -160 | -155 | -150 | -145 | -140 | -135 | -130 | -125 | -120 | -115 | -110 | -105 | -100 | -95 | -90 | -85 | -80 | -75 | -70 | -65 | -60 | -55 | -50 | -45 | -40 | -35 | -30 | -25 | -20 | -15 | -10 | -5 | 0 | 5 | 10 | 15 | 20 | 25 | 30 | 35 | 40 | 45 | 50 | 55 | 60 | 65 | 70 | 75 | 80 | 85 | 90 | 95 | 100 | 105 | 110 | 115 | 120 | 125 | 130 | 135 | 140 | 145 | 150 | 155 | 160 | 165 | 170 | 175 | 180 | 185 | 190 | 195 | 200 | 205 | 210 | 215 | 220 | 225 | 230 | 235 | 240 | 245 | 250 | 255 | 260 | 265 | 270 | 275 | 280 | 285 | 290 | 295 | 300 | 305 | 310 | 315 | 320 | 325 | 330 | 335 | 340 | 345 | 350 | 355 | 360 | 365 | 370 | 375 | 380 | 385 | 390 | 395 | 400 | 405 | 410 | 415 | 420 | 425 | 430 | 435 | 440 | 445 | 450 | 455 | 460 | 465 | 470 | 475 | 480 | 485 | 490 | 495 | 500 | 505 | 510 | 515 | 520 | 525 | 530 | 535 | 540 | 545 | 550 | 555 | 560 | 565 | 570 | 575 | 580 | 585 | 590 | 595 | 600 | 605 | 610 | 615 | 620 | 625 | 630 | 635 | 640 | 645 | 650 | 655 | 660 | 665 | 670 | 675 | 680 | 685 | 690 | 695 | 700 | 705 | 710 | 715 | 720 | 725 | 730 | 735 | 740 | 745 | 750 | 755 | 760 | 765 | 770 | 775 | 780 | 785 | 790 | 795 | 800 | 805 | 810 | 815 | 820 | 825 | 830 | 835 | 840 | 845 | 850 | 855 | 860 | 865 | 870 | 875 | 880 | 885 | 890 | 895 | 900 | 905 | 910 | 915 | 920 | 925 | 930 | 935 | 940 | 945 | 950 | 955 | 960 | 965 | 970 | 975 | 980 | 985 | 990 | 995 | 1000 | 1005 | 1010 | 1015 | 1020 | 1025 | 1030 | 1035 | 1040 | 1045 | 1050 | 1055 | 1060 | 1065 | 1070 | 1075 | 1080 | 1085 | 1090 | 1095 | 1100 | 1105 | 1110 | 1115 | 1120 | 1125 | 1130 | 1135 | 1140 | 1145 | 1150 | 1155 | 1160 | 1165 | 1170 | 1175 | 1180 | 1185 | 1190 | 1195 |
| --- | --- | --- | --- | --- | --- | --- | --- | --- | --- | --- | --- | --- | --- | --- | --- | --- | --- | --- | --- | --- | --- | --- | --- | --- | --- | --- | --- | --- | --- | --- | --- | --- | --- | --- | --- | --- | --- | --- | --- | --- | --- | --- | --- | --- | --- | --- | --- | --- | --- | --- | --- | --- | --- | --- | --- | --- | --- | --- | --- | --- | --- | --- | --- | --- | --- | --- | --- | --- | --- | --- | --- | --- | --- | --- | --- | --- | --- | --- | --- | --- | --- | --- | --- | --- | --- | --- | --- | --- | --- | --- | --- | --- | --- | --- | --- | --- | --- | --- | --- | --- | --- | --- | --- | --- | --- | --- | --- | --- | --- | --- | --- | --- | --- | --- | --- | --- | --- | --- | --- | --- | --- | --- | --- | --- | --- | --- | --- | --- | --- | --- | --- | --- | --- | --- | --- | --- | --- | --- | --- | --- | --- | --- | --- | --- | --- | --- | --- | --- | --- | --- | --- | --- | --- | --- | --- | --- | --- | --- | --- | --- | --- | --- | --- | --- | --- | --- | --- | --- | --- | --- | --- | --- | --- | --- | --- | --- | --- | --- | --- | --- | --- | --- | --- | --- | --- | --- | --- | --- | --- | --- | --- | --- | --- | --- | --- | --- | --- | --- | --- | --- | --- | --- | --- | --- | --- | --- | --- | --- | --- | --- | --- | --- | --- | --- | --- | --- | --- | --- | --- | --- | --- | --- | --- | --- | --- | --- | --- | --- | --- | --- | --- | --- | --- | --- | --- | --- | --- | --- | --- | --- | --- | --- | --- | --- | --- | --- | --- | --- | --- | --- | --- | --- | --- | --- | --- | --- | --- | --- | --- | --- | --- | --- | --- | --- | --- | --- | --- | --- | --- | --- | --- | --- | --- | --- | --- | --- | --- | --- | --- | --- |
| left anterior | 0.255454 | 0.217079 | 0.218182 | 0.218262 | 0.218584 | 0.227324 | 0.215008 | 0.231912 | 0.318261 | 0.303188 | 0.265482 | 0.226003 | 0.214949 | 0.227135 | 0.217784 | 0.234196 | 0.267055 | 0.417045 | 0.512183 | 0.623495 | 0.583249 | 0.489244 | 0.744822 | 0.477345 | 0.400015 | 0.246296 | 0.218659 | 0.226232 | 0.329458 | 0.594459 | 0.967911 | 0.729970 | 0.235720 | 0.238789 | 0.243120 | 0.355682 | 0.564143 | 0.353176 | 0.249968 | 0.214686 | 0.215279 | 0.249370 | 0.276557 | 0.265815 | 0.885375 | 1.837772 | 4.582637 | 10.823001 | 3.060657 | 1.304322 | 0.916306 | 0.322283 | 0.228408 | 0.214859 | 0.243600 | 0.392101 | 0.573462 | 0.897926 | 0.977681 | 0.613238 | 0.407961 | 0.258043 | 0.215849 | 0.214672 | 0.218259 | 0.226274 | 0.330644 | 0.596455 | 1.804118 | 4.409456 | 5.066746e+00 | 2.526319e+00 | 1.284265e+00 | 4.345392e-01 | 2.710691e-01 | 2.153509e-01 | 2.909308e-01 | 0.596962 | 0.930338 | 1.603062 | 1.904747 | 1.803483 | 1.080982 | 0.929044 | 0.597451 | 0.414767 | 0.309332 | 0.224823 | 0.226094 | 0.352544 | 0.797494 | 1.920047 | 4.093204 | 7.662892 | 10.573414 | 8.111573 | 3.597022 | 2.548979 | 1.704488 | 1.129643 | 0.662035 | 0.441150 | 0.347600 | 0.282419 | 0.228795 | 0.220700 | 0.221163 | 0.221622 | 0.217376 | 0.216945 | 0.216760 | 0.226483 | 0.272548 | 0.337854 | 0.492321 | 0.626550 | 0.900663 | 1.769179 | 2.239752 | 2.380178 | 2.739096 | 5.652530 | 10.294981 | 16.558701 | 36.086990 | 49.026660 | 68.275407 | 45.153454 | 14.889843 | 6.846248 | 2.709578 | 1.014495 | 0.573553 | 0.374179 | 0.314931 | 0.273842 | 0.259276 | 0.248170 | 0.271865 | 0.290238 | 0.318156 | 0.503107 | 0.567744 | 0.551614 | 0.627840 | 0.514896 | 0.490803 | 0.416204 | 0.275107 | 0.246229 | 0.224691 | 0.217276 | 0.218762 | 0.224716 | 0.230414 | 0.230492 | 0.221014 | 0.214816 | 0.243290 | 0.320546 | 0.472525 | 0.806257 | 1.029968 | 1.302569 | 1.083199 | 0.864171 | 0.658709 | 0.673673 | 0.527481 | 0.395771 | 0.280507 | 0.255142 | 0.234274 | 0.221512 | 0.214762 | 0.214635 | 0.217168 | 0.240457 | 0.254342 | 0.307795 | 0.340173 | 0.408992 | 0.453171 | 0.535115 | 0.533376 | 0.512598 | 0.373957 | 0.349671 | 0.321753 | 0.380689 | 0.401034 | 0.366307 | 0.373361 | 0.467563 | 0.449256 | 0.418273 | 0.286479 | 0.242240 | 0.247745 | 0.265037 | 0.246984 | 0.237467 | 0.226978 | 0.220104 | 0.215834 | 0.214687 | 0.215365 | 0.214698 | 0.216671 | 0.219976 | 0.215019 | 0.214998 | 0.215672 | 0.214739 | 0.214690 | 0.219894 | 0.238716 | 0.234198 | 0.230286 | 0.234243 | 0.222474 | 0.218299 | 0.215123 | 0.217953 | 0.233070 | 0.234644 | 0.220652 | 0.214696 | 0.214917 | 0.215131 | 0.216392 | 0.223915 | 0.240003 | 0.247224 | 0.239835 | 0.235745 | 0.217826 | 0.214819 | 0.221732 | 0.282862 | 0.496362 | 0.751700 | 0.731709 | 0.657944 | 0.670555 | 0.549242 | 0.358322 | 0.288908 | 0.267862 | 0.307872 | 0.328612 | 0.324351 | 0.333324 | 0.415925 | 0.428904 | 0.563452 | 0.551190 | 0.435596 | 0.478628 | 0.587314 | 0.739139 | 0.935729 | 0.741598 | 0.591877 | 0.665226 | 0.592314 | 0.492399 | 0.450464 | 0.538635 | 0.761375 | 0.866563 | 0.640423 | 0.730154 | 0.841241 | 0.900640 | 0.776127 | 0.998017 | 1.108858 | 2.980148 | 3.273234 |
| right anterior | 0.268811 | 0.405814 | 0.283100 | 0.240153 | 0.215737 | 0.236256 | 0.245229 | 0.253524 | 0.327830 | 0.290454 | 0.236682 | 0.251634 | 0.302469 | 0.305882 | 0.252012 | 0.254838 | 0.329982 | 0.434268 | 0.312423 | 0.252635 | 0.242071 | 0.267512 | 0.231745 | 0.217402 | 0.214989 | 0.238774 | 0.232046 | 0.240631 | 0.233434 | 0.236143 | 0.242305 | 0.237467 | 0.236365 | 0.334415 | 0.510807 | 0.960432 | 1.307898 | 1.028519 | 0.399594 | 0.297833 | 0.260684 | 0.231354 | 0.224826 | 0.246608 | 0.446090 | 1.261710 | 1.824302 | 1.087075 | 0.550534 | 0.345119 | 0.278783 | 0.225329 | 0.219335 | 0.218830 | 0.216990 | 0.220013 | 0.214728 | 0.217975 | 0.214750 | 0.215126 | 0.214641 | 0.215598 | 0.214689 | 0.214641 | 0.215743 | 0.273584 | 0.520017 | 1.596796 | 5.701872 | 20.802488 | 2.227218e+01 | 2.422820e+01 | 2.449620e+01 | 1.207870e+01 | 7.078966e+00 | 2.966598e+00 | 1.597558e+00 | 1.279218 | 0.734957 | 0.404131 | 0.287798 | 0.280871 | 0.279160 | 0.265471 | 0.278616 | 0.282543 | 0.291751 | 0.317745 | 0.419193 | 0.569592 | 0.940229 | 1.393705 | 2.552037 | 4.224877 | 8.176006 | 10.944306 | 22.412261 | 16.740198 | 9.106510 | 5.522178 | 5.143339 | 3.642378 | 1.183443 | 0.601422 | 0.573741 | 0.635286 | 0.591941 | 0.368539 | 0.241424 | 0.237468 | 0.261726 | 0.270813 | 0.297185 | 0.267213 | 0.332104 | 0.568275 | 1.292589 | 1.672974 | 1.986459 | 3.181568 | 7.428148 | 16.115745 | 43.493004 | 47.285049 | 33.404806 | 29.993232 | 14.634476 | 6.579706 | 5.011565 | 3.986475 | 4.172854 | 4.270974 | 1.958828 | 0.763546 | 0.385399 | 0.266712 | 0.215745 | 0.233469 | 0.303156 | 0.348520 | 0.295321 | 0.233510 | 0.227410 | 0.235982 | 0.261641 | 0.282794 | 0.275049 | 0.271530 | 0.294918 | 0.274209 | 0.243513 | 0.235440 | 0.243714 | 0.314966 | 0.512066 | 1.020320 | 2.441082 | 4.709524 | 2.640049 | 1.112957 | 0.551891 | 0.363269 | 0.240354 | 0.214736 | 0.223430 | 0.240366 | 0.246118 | 0.276667 | 0.312499 | 0.291696 | 0.265542 | 0.243618 | 0.219595 | 0.214829 | 0.216642 | 0.221996 | 0.233294 | 0.240555 | 0.254404 | 0.247080 | 0.238484 | 0.237056 | 0.237934 | 0.234814 | 0.239574 | 0.228737 | 0.226688 | 0.222259 | 0.217812 | 0.217725 | 0.220709 | 0.219030 | 0.215729 | 0.214681 | 0.218114 | 0.221448 | 0.230493 | 0.231487 | 0.228098 | 0.230460 | 0.229923 | 0.222740 | 0.217674 | 0.214633 | 0.214652 | 0.214724 | 0.221050 | 0.249082 | 0.261399 | 0.249214 | 0.227159 | 0.216001 | 0.214631 | 0.216560 | 0.220536 | 0.217631 | 0.214701 | 0.215800 | 0.231953 | 0.268190 | 0.315511 | 0.339694 | 0.280352 | 0.258071 | 0.240252 | 0.224947 | 0.225513 | 0.221074 | 0.214744 | 0.214626 | 0.218616 | 0.218978 | 0.222503 | 0.227677 | 0.223046 | 0.214659 | 0.223871 | 0.233241 | 0.232290 | 0.234384 | 0.225892 | 0.214999 | 0.219125 | 0.234244 | 0.229599 | 0.234108 | 0.243280 | 0.248789 | 0.238064 | 0.225095 | 0.221711 | 0.223295 | 0.220370 | 0.218721 | 0.222054 | 0.231956 | 0.258118 | 0.279234 | 0.288681 | 0.287818 | 0.286929 | 0.301213 | 0.337016 | 0.324866 | 0.328463 | 0.294362 | 0.267094 | 0.230793 | 0.216925 | 0.216996 | 0.218823 | 0.231681 | 0.242518 | 0.263605 | 0.248946 | 0.242428 | 0.233643 | 0.252335 | 0.273241 | 0.295794 |
| left central | 0.281898 | 0.353672 | 0.305579 | 0.268737 | 0.339814 | 0.438547 | 0.732228 | 0.496545 | 0.388050 | 0.627678 | 0.789522 | 0.383773 | 0.241919 | 0.218714 | 0.217014 | 0.299753 | 0.879720 | 4.337636 | 4.014514 | 1.498483 | 0.746288 | 0.464678 | 0.237563 | 0.235419 | 0.364474 | 0.427061 | 0.430428 | 0.438177 | 0.402579 | 0.314696 | 0.216154 | 0.223035 | 0.232514 | 0.215042 | 0.218879 | 0.238152 | 0.280898 | 0.401110 | 0.997744 | 3.993762 | 7.497592 | 6.739417 | 4.669469 | 3.258431 | 6.033209 | 5.726258 | 2.343889 | 0.624272 | 0.301940 | 0.259029 | 0.245026 | 0.228764 | 0.232987 | 0.260880 | 0.334781 | 0.577425 | 0.918365 | 1.418317 | 1.902288 | 0.891613 | 0.372940 | 0.299970 | 0.299927 | 0.292576 | 0.311958 | 0.478120 | 0.934216 | 3.213884 | 11.668009 | 23.575847 | 7.650001e+01 | 1.268161e+02 | 1.407781e+02 | 1.699922e+02 | 6.501168e+01 | 2.130945e+01 | 6.901023e+00 | 3.126679 | 2.247291 | 1.004557 | 0.548924 | 0.437192 | 0.355256 | 0.350173 | 0.348290 | 0.326146 | 0.428615 | 0.714665 | 1.705606 | 4.460417 | 16.408738 | 46.739237 | 112.960925 | 385.732527 | 466.122527 | 577.765659 | 1200.554334 | 798.112119 | 615.439646 | 164.125354 | 38.183206 | 15.209090 | 4.096925 | 0.861429 | 0.370943 | 0.230838 | 0.214667 | 0.221742 | 0.221647 | 0.225557 | 0.218510 | 0.214844 | 0.214853 | 0.222185 | 0.235499 | 0.234179 | 0.239265 | 0.232884 | 0.246678 | 0.306570 | 0.352531 | 0.439181 | 0.591677 | 0.657854 | 0.902853 | 0.859558 | 0.656018 | 0.507979 | 0.480783 | 0.427767 | 0.330026 | 0.239609 | 0.226138 | 0.214626 | 0.221194 | 0.235036 | 0.242903 | 0.219133 | 0.227257 | 0.260099 | 0.354659 | 0.478922 | 0.383012 | 0.281375 | 0.225328 | 0.214676 | 0.224686 | 0.226748 | 0.229877 | 0.214634 | 0.228065 | 0.316685 | 0.371966 | 0.396782 | 0.338462 | 0.293430 | 0.243093 | 0.224093 | 0.214652 | 0.216631 | 0.217755 | 0.222561 | 0.226997 | 0.230273 | 0.223222 | 0.218365 | 0.216077 | 0.214897 | 0.215448 | 0.219627 | 0.219146 | 0.231691 | 0.224179 | 0.221162 | 0.216989 | 0.215425 | 0.238220 | 0.257002 | 0.321408 | 0.267452 | 0.251983 | 0.244684 | 0.236678 | 0.228236 | 0.231244 | 0.223031 | 0.236225 | 0.241815 | 0.226943 | 0.218646 | 0.215802 | 0.214912 | 0.216191 | 0.223894 | 0.233592 | 0.219802 | 0.214707 | 0.216211 | 0.223453 | 0.227675 | 0.226952 | 0.216573 | 0.224428 | 0.276785 | 0.323950 | 0.405879 | 0.485671 | 0.511314 | 0.483571 | 0.338361 | 0.298062 | 0.285748 | 0.248348 | 0.223936 | 0.214718 | 0.225721 | 0.230708 | 0.254822 | 0.236335 | 0.214904 | 0.232565 | 0.290749 | 0.408428 | 0.531448 | 0.813974 | 1.038908 | 1.605716 | 2.599056 | 3.625188 | 5.602719 | 7.082975 | 4.922767 | 2.688243 | 2.306790 | 1.493123 | 1.293503 | 1.108599 | 0.892194 | 0.594923 | 0.394222 | 0.248854 | 0.245623 | 0.234963 | 0.226190 | 0.231862 | 0.247694 | 0.249209 | 0.285078 | 0.342893 | 0.432028 | 0.544212 | 0.723313 | 1.118010 | 1.429470 | 0.997319 | 0.653966 | 0.516821 | 0.499843 | 0.418677 | 0.394565 | 0.473069 | 0.682917 | 0.815384 | 1.402085 | 1.684038 | 1.867168 | 1.688204 | 1.389955 | 1.309812 | 1.621072 | 1.634442 | 1.678524 | 1.396112 | 1.152783 | 0.973112 | 0.954909 | 0.853206 | 0.706375 | 0.575514 | 0.529673 |
| right central | 0.224454 | 0.224354 | 0.244150 | 0.283165 | 0.357065 | 0.444247 | 0.625814 | 1.097812 | 1.908188 | 2.360155 | 1.742563 | 0.941776 | 0.700287 | 0.458761 | 0.286589 | 0.224048 | 0.270621 | 0.268598 | 0.224867 | 0.215914 | 0.227066 | 0.234860 | 0.217994 | 0.215088 | 0.223885 | 0.257058 | 0.348412 | 0.766566 | 2.505410 | 1.809337 | 0.522188 | 0.253511 | 0.220468 | 0.224759 | 0.297572 | 0.429137 | 0.331191 | 0.242426 | 0.214640 | 0.214640 | 0.215233 | 0.223711 | 0.249109 | 0.372850 | 0.585222 | 0.949543 | 0.838898 | 0.502185 | 0.366900 | 0.289795 | 0.288047 | 0.255649 | 0.217127 | 0.214665 | 0.216350 | 0.219898 | 0.233792 | 0.255733 | 0.294300 | 0.427479 | 1.002069 | 2.513927 | 8.523328 | 12.663167 | 17.915739 | 39.372934 | 113.493753 | 213.036860 | 891.258981 | 2203.571882 | 1.100668e+04 | 4.459917e+04 | 5.670639e+04 | 3.547170e+04 | 1.269882e+04 | 1.361013e+03 | 1.849957e+02 | 50.360014 | 25.859473 | 24.133057 | 29.784976 | 40.025861 | 79.153975 | 214.305750 | 224.317175 | 327.191104 | 337.997437 | 288.546791 | 371.730457 | 1564.992640 | 2187.472257 | 9132.736564 | 11491.296883 | 20256.434726 | 73770.189927 | 146378.598393 | 135237.997703 | 232670.451962 | 198952.331576 | 106543.065007 | 51163.485300 | 20578.377674 | 8754.087838 | 2439.959306 | 532.859534 | 103.762678 | 45.719573 | 26.858435 | 14.377222 | 14.417020 | 27.163534 | 52.355088 | 119.784963 | 374.798282 | 545.550378 | 560.522239 | 326.274449 | 249.279962 | 165.542922 | 86.854155 | 96.406417 | 64.014941 | 34.836685 | 29.797291 | 18.993175 | 16.164651 | 22.040129 | 16.542843 | 10.083733 | 6.078324 | 4.952815 | 6.286106 | 3.221677 | 1.927140 | 1.073037 | 0.738040 | 0.495084 | 0.436743 | 0.435359 | 0.478684 | 0.592629 | 0.824460 | 1.395164 | 1.770911 | 2.852423 | 1.340440 | 0.889541 | 0.570665 | 0.375302 | 0.284384 | 0.295003 | 0.256545 | 0.276731 | 0.282651 | 0.289162 | 0.417824 | 0.651579 | 0.837260 | 1.248532 | 1.662599 | 2.869527 | 3.696374 | 2.616512 | 2.031928 | 2.003612 | 2.158607 | 3.514080 | 3.946938 | 4.157042 | 4.591414 | 7.577420 | 6.324112 | 3.991226 | 1.805898 | 0.775002 | 0.453453 | 0.394283 | 0.338693 | 0.353711 | 0.394100 | 0.550541 | 0.915802 | 1.528823 | 1.748414 | 2.402540 | 2.639771 | 2.552698 | 1.821536 | 1.463163 | 1.129862 | 1.109168 | 0.724207 | 0.472800 | 0.423829 | 0.381151 | 0.304675 | 0.284238 | 0.294500 | 0.344216 | 0.434692 | 0.459588 | 0.432925 | 0.388476 | 0.364195 | 0.325825 | 0.307142 | 0.277986 | 0.263416 | 0.273109 | 0.266382 | 0.242664 | 0.236439 | 0.221597 | 0.220453 | 0.218179 | 0.222693 | 0.242821 | 0.302595 | 0.340939 | 0.335758 | 0.294871 | 0.294372 | 0.273825 | 0.264517 | 0.262807 | 0.250364 | 0.232400 | 0.222036 | 0.215613 | 0.214687 | 0.214684 | 0.214827 | 0.216503 | 0.236444 | 0.247425 | 0.286209 | 0.336872 | 0.334281 | 0.338169 | 0.342926 | 0.302934 | 0.309164 | 0.268226 | 0.238181 | 0.231681 | 0.220267 | 0.216408 | 0.238404 | 0.292193 | 0.365893 | 0.363825 | 0.360376 | 0.347330 | 0.318321 | 0.285191 | 0.229214 | 0.215865 | 0.214812 | 0.214667 | 0.214873 | 0.216089 | 0.217030 | 0.218235 | 0.236370 | 0.272932 | 0.303071 | 0.388544 | 0.414211 | 0.405818 | 0.300733 | 0.256960 | 0.230122 | 0.214749 | 0.223071 | 0.238564 | 0.266005 | 0.269421 | 0.290772 | 0.302753 | 0.298167 |
| left posterior | 0.771006 | 0.477475 | 0.481650 | 0.446531 | 0.413387 | 0.299898 | 0.219513 | 0.324963 | 0.298162 | 0.327592 | 0.334591 | 0.284895 | 0.271960 | 0.233610 | 0.214840 | 0.217602 | 0.331930 | 0.692513 | 2.973616 | 8.621072 | 31.548759 | 21.873162 | 4.684011 | 0.341226 | 0.262438 | 0.700224 | 1.288507 | 1.268334 | 0.483988 | 0.256146 | 0.214626 | 0.217872 | 0.217723 | 0.214702 | 0.245915 | 0.391012 | 0.621700 | 0.668688 | 0.622627 | 0.732625 | 0.991367 | 2.454603 | 4.637980 | 4.036016 | 3.568537 | 2.916251 | 2.223995 | 1.250555 | 0.341287 | 0.227245 | 0.218822 | 0.258974 | 0.371234 | 0.452091 | 0.374005 | 0.239904 | 0.222499 | 0.214876 | 0.223526 | 0.283937 | 0.363535 | 0.640271 | 1.385365 | 3.723072 | 14.625793 | 108.914829 | 720.339281 | 8177.083137 | 41828.793881 | 190592.521978 | 5.291192e+05 | 1.310877e+06 | 1.020844e+06 | 5.506678e+05 | 1.085433e+05 | 1.619290e+04 | 5.156790e+03 | 3042.769305 | 1277.338479 | 708.164871 | 357.574904 | 139.797099 | 70.028111 | 42.357088 | 32.050225 | 41.357357 | 59.001981 | 73.337409 | 124.204245 | 172.482097 | 184.145667 | 172.942151 | 156.289327 | 202.217642 | 333.700027 | 545.610929 | 615.523472 | 349.730248 | 139.135741 | 120.950456 | 55.029139 | 27.531426 | 12.407444 | 8.629692 | 8.404424 | 9.688188 | 7.363170 | 7.276649 | 8.171563 | 15.553681 | 22.718803 | 23.273128 | 29.569961 | 49.435206 | 113.269759 | 220.609558 | 410.106849 | 852.378390 | 2181.204468 | 5467.123370 | 7696.464703 | 9105.066311 | 10487.596584 | 6372.890079 | 3631.639297 | 2207.901312 | 996.123060 | 400.206682 | 129.431458 | 41.385603 | 22.539559 | 15.739133 | 16.241071 | 16.471343 | 21.137931 | 31.495876 | 43.067678 | 47.015719 | 49.859565 | 65.631887 | 81.153974 | 90.681238 | 112.931223 | 153.495128 | 239.255185 | 159.236331 | 69.214629 | 30.497171 | 22.520002 | 15.054106 | 11.224248 | 7.619039 | 6.815786 | 6.633072 | 11.097319 | 14.518114 | 21.411534 | 44.705964 | 106.008478 | 124.838682 | 91.264720 | 51.473508 | 40.463353 | 38.164421 | 20.486433 | 8.982289 | 9.481938 | 8.260843 | 10.109377 | 9.309492 | 9.309492 | 9.466526 | 11.404923 | 9.029515 | 8.438633 | 5.989386 | 4.943075 | 4.292695 | 4.075144 | 3.695103 | 3.229163 | 3.338793 | 4.006764 | 3.461114 | 2.863251 | 2.586132 | 2.603138 | 3.052412 | 2.268105 | 1.127140 | 0.815841 | 0.765411 | 0.864089 | 0.763687 | 0.756771 | 0.849185 | 1.199078 | 1.940079 | 2.842754 | 2.832121 | 2.566589 | 2.340577 | 2.332141 | 2.192674 | 1.472011 | 1.033638 | 0.796822 | 0.885669 | 0.993324 | 0.910841 | 0.984947 | 1.144337 | 1.622160 | 2.022431 | 2.127535 | 2.194367 | 2.701616 | 3.507161 | 4.841858 | 5.143869 | 4.787447 | 3.798345 | 3.373468 | 3.199085 | 2.641254 | 1.857539 | 1.539860 | 1.433726 | 1.750059 | 1.653837 | 1.698837 | 2.253978 | 2.882144 | 3.058024 | 3.815157 | 2.745492 | 3.133100 | 2.887428 | 3.442789 | 4.529502 | 6.726612 | 9.872798 | 18.515909 | 15.453279 | 14.052195 | 7.373387 | 5.455800 | 3.642950 | 2.242317 | 1.889470 | 1.339825 | 1.352618 | 1.553654 | 1.594873 | 1.440865 | 1.241765 | 1.353569 | 1.905794 | 2.023139 | 2.163803 | 2.182772 | 1.795426 | 1.490056 | 1.032962 | 0.639065 | 0.409914 | 0.291465 | 0.237202 | 0.218958 | 0.215078 | 0.217092 | 0.214859 | 0.215655 | 0.226716 | 0.250536 | 0.305815 | 0.365754 | 0.411094 | 0.442210 | 0.498468 |
| right posterior | 0.245564 | 0.272596 | 0.293707 | 0.554282 | 1.753030 | 4.380208 | 3.176719 | 0.823700 | 0.441164 | 0.369170 | 0.280578 | 0.243673 | 0.215178 | 0.215171 | 0.224968 | 0.311740 | 0.377291 | 0.438167 | 0.500344 | 0.663139 | 0.465186 | 0.244254 | 0.228025 | 0.302985 | 0.465517 | 0.544126 | 0.462768 | 0.243497 | 0.259741 | 0.320829 | 0.374609 | 0.373524 | 0.365493 | 0.296748 | 0.231334 | 0.248001 | 0.336724 | 0.417129 | 0.441881 | 0.832694 | 1.228708 | 1.667881 | 1.350052 | 0.948380 | 0.747731 | 0.830291 | 0.781540 | 0.479356 | 0.274288 | 0.217507 | 0.216570 | 0.220782 | 0.216377 | 0.215027 | 0.259285 | 0.582120 | 1.885438 | 5.389997 | 9.363246 | 7.510958 | 6.250082 | 9.685433 | 15.051152 | 26.174932 | 51.509959 | 117.529718 | 583.292127 | 7314.125083 | 67392.531921 | 526007.236104 | 1.422956e+06 | 2.246230e+06 | 2.242892e+06 | 2.958337e+06 | 9.045002e+05 | 2.510705e+05 | 5.614510e+04 | 22315.343067 | 14517.740845 | 18359.058634 | 5938.912264 | 1283.314508 | 233.088328 | 102.745667 | 85.436465 | 73.968619 | 99.090262 | 218.228846 | 697.180876 | 1550.005652 | 1520.326185 | 1791.660798 | 2861.401896 | 3169.689782 | 4898.145940 | 7311.802066 | 8170.579699 | 7553.476050 | 7110.090926 | 6902.065862 | 7316.364991 | 3471.544801 | 1237.460898 | 416.073745 | 284.485350 | 308.003440 | 235.582800 | 163.923937 | 129.978224 | 104.107117 | 99.025198 | 95.139009 | 136.009048 | 224.591629 | 423.571680 | 819.177409 | 2925.443150 | 7017.323349 | 12909.108044 | 9200.282535 | 4519.442466 | 4464.578533 | 3567.139568 | 2627.470617 | 1949.768733 | 988.692131 | 372.778282 | 256.944720 | 126.155530 | 101.305145 | 93.817094 | 73.387693 | 95.802546 | 174.250448 | 255.364643 | 301.709992 | 308.211067 | 355.251513 | 767.363076 | 1031.604416 | 1693.884934 | 2584.721604 | 3710.728250 | 3738.317372 | 3805.251683 | 2673.527410 | 1494.359285 | 593.338055 | 196.521177 | 95.003787 | 78.883971 | 60.237496 | 84.677726 | 132.360122 | 267.598323 | 613.543757 | 817.225002 | 944.277661 | 923.872086 | 664.751081 | 411.068278 | 238.209791 | 164.482200 | 134.878163 | 140.071065 | 186.480398 | 202.983315 | 186.529651 | 190.489451 | 154.896083 | 189.398306 | 160.830258 | 147.282556 | 154.684361 | 155.702848 | 205.846209 | 390.152989 | 558.257722 | 761.360678 | 549.452154 | 403.440389 | 334.802586 | 275.189499 | 169.053694 | 117.414878 | 120.443175 | 169.355380 | 214.508408 | 345.781254 | 299.722243 | 360.867344 | 304.762592 | 250.710277 | 121.975152 | 51.647348 | 22.689041 | 12.542782 | 6.472661 | 5.388035 | 4.232740 | 5.637363 | 8.251038 | 10.050422 | 15.631774 | 26.503707 | 27.764339 | 25.518947 | 22.513247 | 23.958136 | 38.328449 | 61.647009 | 90.242841 | 162.883504 | 389.284408 | 541.642561 | 1169.544498 | 2148.402129 | 2440.437924 | 2881.360167 | 1303.258824 | 702.133315 | 456.576569 | 248.904552 | 136.990707 | 77.756743 | 48.828147 | 65.151086 | 62.464193 | 61.613941 | 65.998720 | 70.095186 | 101.447232 | 113.251211 | 123.380737 | 101.616284 | 84.504156 | 111.260407 | 153.704242 | 151.454690 | 155.349758 | 116.199368 | 93.202618 | 84.614127 | 75.823077 | 83.813934 | 107.215881 | 137.099008 | 171.719058 | 134.557741 | 107.911102 | 95.648239 | 62.870138 | 38.312367 | 25.858719 | 26.548274 | 38.621819 | 66.767609 | 104.509164 | 306.559057 | 834.466178 | 1503.541995 | 3269.381163 | 6932.503624 | 10313.121998 | 6466.193092 | 1164.782933 | 229.041117 | 88.697409 | 44.505575 | 30.857140 | 27.425697 | 18.247858 | 22.487581 | 20.089407 | 18.900132 | 23.513592 | 26.506031 | 24.894760 | 26.075495 | 20.199594 |
| all electrodes | 0.815922 | 1.174110 | 0.923803 | 0.441720 | 0.271620 | 0.218255 | 0.229170 | 0.252533 | 0.301253 | 0.395410 | 0.238866 | 0.224838 | 0.221271 | 0.252521 | 0.234919 | 0.216651 | 0.363766 | 2.078795 | 5.634517 | 5.863572 | 4.354474 | 3.706805 | 0.707738 | 0.244221 | 0.239747 | 0.378732 | 0.450037 | 0.298024 | 0.275570 | 0.216761 | 0.218440 | 0.222128 | 0.217168 | 0.244252 | 0.325907 | 0.644091 | 0.968278 | 0.816633 | 0.693014 | 0.440386 | 0.312503 | 0.284731 | 0.289133 | 0.265929 | 0.260920 | 0.248148 | 0.238063 | 0.218404 | 0.214637 | 0.225488 | 0.228632 | 0.233820 | 0.217223 | 0.214754 | 0.257881 | 0.344528 | 0.553520 | 0.969617 | 1.259204 | 1.241775 | 1.403337 | 1.338300 | 1.496555 | 2.297223 | 5.093497 | 22.995620 | 186.667450 | 1614.485041 | 10847.399426 | 65020.539629 | 7.351689e+05 | 7.849094e+06 | 2.488721e+07 | 2.521965e+07 | 1.463823e+07 | 3.940995e+06 | 2.251075e+06 | 502908.604901 | 92718.992549 | 49830.940743 | 25098.706818 | 6487.334064 | 2189.008585 | 275.972160 | 157.661866 | 110.278560 | 108.115724 | 79.429489 | 72.779872 | 52.966688 | 102.884857 | 92.418871 | 130.669298 | 477.648857 | 2281.607841 | 4267.364159 | 3853.717387 | 2941.410808 | 4164.313362 | 4219.396852 | 1349.790806 | 650.410564 | 720.225149 | 568.791911 | 459.976336 | 258.501057 | 241.506077 | 206.797735 | 173.795329 | 197.450746 | 466.822005 | 859.465052 | 1575.889706 | 1071.530473 | 944.556637 | 1074.487324 | 657.430717 | 740.322735 | 712.474455 | 678.706781 | 836.153171 | 1136.126797 | 2937.706814 | 5509.181982 | 3899.036378 | 2143.011626 | 1150.123582 | 1364.761461 | 1286.456071 | 681.114451 | 797.983972 | 1076.039879 | 1218.730869 | 1033.349651 | 322.162557 | 136.377715 | 99.361620 | 68.660357 | 95.821853 | 223.798936 | 582.788946 | 1573.986987 | 1137.146539 | 312.354472 | 126.778137 | 43.889089 | 15.240369 | 5.185776 | 4.006861 | 5.799200 | 14.512527 | 42.236484 | 111.659297 | 223.259159 | 244.269938 | 318.592486 | 206.802749 | 97.996855 | 39.694905 | 25.519870 | 12.585705 | 12.223441 | 9.409057 | 9.670978 | 6.800322 | 6.174160 | 6.324567 | 10.070599 | 8.675247 | 6.447491 | 3.716363 | 4.481314 | 3.214788 | 1.392880 | 0.884395 | 0.959165 | 1.111701 | 1.244884 | 1.031552 | 1.186825 | 1.653632 | 2.270576 | 1.468094 | 1.255782 | 1.496128 | 1.297468 | 1.191095 | 1.569714 | 1.286693 | 0.942935 | 0.670042 | 0.672716 | 0.891071 | 0.973101 | 0.723703 | 0.597045 | 0.755005 | 0.979611 | 0.915134 | 0.693256 | 0.538071 | 0.507525 | 0.567050 | 0.604344 | 0.542743 | 0.454588 | 0.513132 | 0.428396 | 0.415584 | 0.350776 | 0.289402 | 0.282761 | 0.299612 | 0.294837 | 0.359551 | 0.401273 | 0.488789 | 0.621066 | 0.548188 | 0.460442 | 0.352444 | 0.282089 | 0.281094 | 0.312716 | 0.326580 | 0.310939 | 0.262310 | 0.253191 | 0.260061 | 0.234161 | 0.214793 | 0.218525 | 0.215229 | 0.214781 | 0.218125 | 0.228610 | 0.254446 | 0.281713 | 0.309184 | 0.331391 | 0.411101 | 0.474374 | 0.477983 | 0.393781 | 0.295106 | 0.269974 | 0.255271 | 0.245079 | 0.217737 | 0.217714 | 0.225624 | 0.223050 | 0.226517 | 0.224686 | 0.221397 | 0.217856 | 0.217668 | 0.215206 | 0.221392 | 0.246773 | 0.287375 | 0.302717 | 0.356214 | 0.373143 | 0.385744 | 0.279032 | 0.228905 | 0.215599 | 0.216393 | 0.228232 | 0.224602 | 0.225647 | 0.218171 | 0.214945 | 0.215030 | 0.219893 | 0.235447 | 0.237944 | 0.240418 | 0.250392 |

Searchlight, spatiotemporal cluster permutation test

|  | start time | stop time | peak time | peak channel | cluster p | peak Cohen's d | direction |
| --- | --- | --- | --- | --- | --- | --- | --- |
| #1 | 80 | 1195 | 150 | POz | 0.0001 | 1.691478 | positive |

F) neutral vs sad

  
|  | time window | peak latency | cluster *p* | peak Cohen's *d* |  | | | |  |
| **all electrodes** | 115 - 205 ms | 160 ms | 0.0313 | 1.0432 |  | | | |  |
 840 - 1105 ms | 865 ms | 0.0062 | 0.7572 |  | | | ||  |  | | | | | | | | |

Time-resolved classification, cluster permutation tests

|  | **left hemisphere** | | | | **right hemisphere** | | | |  |
|  | time window | peak latency | cluster *p* | peak Cohen's *d* | time window | peak latency | cluster *p* | peak Cohen's *d* |  |
| **anterior** |  | | | | 210 - 295 ms | 250 ms | 0.0435 | 0.7589 |  |
| **central** | 90 - 195 ms | 165 ms | 0.0217 | 0.8846 | 205 - 320 ms | 255 ms | 0.0122 | 0.6973 |  |
| **posterior** | 120 - 450 ms | 155 ms | 0.0064 | 1.304 | 235 - 550 ms | 265 ms | 0.0029 | 0.8002 |  |
 805 - 1015 ms | 935 ms | 0.0268 | 0.6426 |  | | | |

  

Time-resolved classification, Bayesian statistics

|  | -200 | -195 | -190 | -185 | -180 | -175 | -170 | -165 | -160 | -155 | -150 | -145 | -140 | -135 | -130 | -125 | -120 | -115 | -110 | -105 | -100 | -95 | -90 | -85 | -80 | -75 | -70 | -65 | -60 | -55 | -50 | -45 | -40 | -35 | -30 | -25 | -20 | -15 | -10 | -5 | 0 | 5 | 10 | 15 | 20 | 25 | 30 | 35 | 40 | 45 | 50 | 55 | 60 | 65 | 70 | 75 | 80 | 85 | 90 | 95 | 100 | 105 | 110 | 115 | 120 | 125 | 130 | 135 | 140 | 145 | 150 | 155 | 160 | 165 | 170 | 175 | 180 | 185 | 190 | 195 | 200 | 205 | 210 | 215 | 220 | 225 | 230 | 235 | 240 | 245 | 250 | 255 | 260 | 265 | 270 | 275 | 280 | 285 | 290 | 295 | 300 | 305 | 310 | 315 | 320 | 325 | 330 | 335 | 340 | 345 | 350 | 355 | 360 | 365 | 370 | 375 | 380 | 385 | 390 | 395 | 400 | 405 | 410 | 415 | 420 | 425 | 430 | 435 | 440 | 445 | 450 | 455 | 460 | 465 | 470 | 475 | 480 | 485 | 490 | 495 | 500 | 505 | 510 | 515 | 520 | 525 | 530 | 535 | 540 | 545 | 550 | 555 | 560 | 565 | 570 | 575 | 580 | 585 | 590 | 595 | 600 | 605 | 610 | 615 | 620 | 625 | 630 | 635 | 640 | 645 | 650 | 655 | 660 | 665 | 670 | 675 | 680 | 685 | 690 | 695 | 700 | 705 | 710 | 715 | 720 | 725 | 730 | 735 | 740 | 745 | 750 | 755 | 760 | 765 | 770 | 775 | 780 | 785 | 790 | 795 | 800 | 805 | 810 | 815 | 820 | 825 | 830 | 835 | 840 | 845 | 850 | 855 | 860 | 865 | 870 | 875 | 880 | 885 | 890 | 895 | 900 | 905 | 910 | 915 | 920 | 925 | 930 | 935 | 940 | 945 | 950 | 955 | 960 | 965 | 970 | 975 | 980 | 985 | 990 | 995 | 1000 | 1005 | 1010 | 1015 | 1020 | 1025 | 1030 | 1035 | 1040 | 1045 | 1050 | 1055 | 1060 | 1065 | 1070 | 1075 | 1080 | 1085 | 1090 | 1095 | 1100 | 1105 | 1110 | 1115 | 1120 | 1125 | 1130 | 1135 | 1140 | 1145 | 1150 | 1155 | 1160 | 1165 | 1170 | 1175 | 1180 | 1185 | 1190 | 1195 |
| --- | --- | --- | --- | --- | --- | --- | --- | --- | --- | --- | --- | --- | --- | --- | --- | --- | --- | --- | --- | --- | --- | --- | --- | --- | --- | --- | --- | --- | --- | --- | --- | --- | --- | --- | --- | --- | --- | --- | --- | --- | --- | --- | --- | --- | --- | --- | --- | --- | --- | --- | --- | --- | --- | --- | --- | --- | --- | --- | --- | --- | --- | --- | --- | --- | --- | --- | --- | --- | --- | --- | --- | --- | --- | --- | --- | --- | --- | --- | --- | --- | --- | --- | --- | --- | --- | --- | --- | --- | --- | --- | --- | --- | --- | --- | --- | --- | --- | --- | --- | --- | --- | --- | --- | --- | --- | --- | --- | --- | --- | --- | --- | --- | --- | --- | --- | --- | --- | --- | --- | --- | --- | --- | --- | --- | --- | --- | --- | --- | --- | --- | --- | --- | --- | --- | --- | --- | --- | --- | --- | --- | --- | --- | --- | --- | --- | --- | --- | --- | --- | --- | --- | --- | --- | --- | --- | --- | --- | --- | --- | --- | --- | --- | --- | --- | --- | --- | --- | --- | --- | --- | --- | --- | --- | --- | --- | --- | --- | --- | --- | --- | --- | --- | --- | --- | --- | --- | --- | --- | --- | --- | --- | --- | --- | --- | --- | --- | --- | --- | --- | --- | --- | --- | --- | --- | --- | --- | --- | --- | --- | --- | --- | --- | --- | --- | --- | --- | --- | --- | --- | --- | --- | --- | --- | --- | --- | --- | --- | --- | --- | --- | --- | --- | --- | --- | --- | --- | --- | --- | --- | --- | --- | --- | --- | --- | --- | --- | --- | --- | --- | --- | --- | --- | --- | --- | --- | --- | --- | --- | --- | --- | --- | --- | --- | --- | --- | --- | --- | --- | --- | --- | --- | --- | --- | --- | --- | --- | --- | --- | --- | --- |
| left anterior | 0.280217 | 0.237386 | 0.229923 | 0.275436 | 0.326198 | 0.333391 | 0.216648 | 0.214692 | 0.220342 | 0.224654 | 0.216812 | 0.215302 | 0.245728 | 0.356752 | 0.319003 | 0.300233 | 0.309994 | 0.298510 | 0.256259 | 0.216925 | 0.214654 | 0.270394 | 0.329114 | 0.309925 | 0.443053 | 0.872232 | 0.694203 | 0.512184 | 0.379891 | 0.290770 | 0.243969 | 0.215784 | 0.231273 | 0.281962 | 0.369480 | 0.783934 | 2.481190 | 1.994389 | 8.069053 | 33.290597 | 47.002320 | 16.818522 | 9.485562 | 7.868650 | 15.399618 | 3.994753 | 2.679111 | 6.332094 | 10.553273 | 23.018940 | 27.034744 | 8.704303 | 5.970735 | 2.871355 | 1.632964 | 0.774385 | 0.378832 | 0.246613 | 0.227101 | 0.216538 | 0.217989 | 0.215229 | 0.224476 | 0.279624 | 0.320198 | 0.400628 | 0.610947 | 1.609406 | 5.365967 | 9.162666 | 8.940070 | 5.749931 | 3.430395 | 1.682589 | 0.746145 | 0.403465 | 0.232538 | 0.220858 | 0.267698 | 0.339835 | 0.331841 | 0.329355 | 0.326037 | 0.256009 | 0.215061 | 0.224522 | 0.261545 | 0.346270 | 0.590152 | 0.853884 | 1.680388 | 2.339972 | 3.927289 | 5.616948 | 5.434385 | 4.723541 | 5.460607 | 4.058161 | 4.403926 | 3.306548 | 1.923935 | 1.474380 | 1.330012 | 0.897874 | 0.891421 | 0.628355 | 0.476496 | 0.354453 | 0.290963 | 0.244355 | 0.237945 | 0.228590 | 0.236046 | 0.277804 | 0.364225 | 0.357638 | 0.353757 | 0.352877 | 0.329861 | 0.315983 | 0.255252 | 0.230155 | 0.233385 | 0.278610 | 0.317383 | 0.309723 | 0.262229 | 0.249927 | 0.233799 | 0.219971 | 0.226455 | 0.311055 | 0.417576 | 0.391129 | 0.348786 | 0.321485 | 0.316423 | 0.293016 | 0.253106 | 0.217819 | 0.214626 | 0.215051 | 0.214626 | 0.219788 | 0.249160 | 0.233944 | 0.215416 | 0.227393 | 0.249486 | 0.241634 | 0.224025 | 0.217029 | 0.228771 | 0.281413 | 0.350250 | 0.553337 | 0.805201 | 1.051219 | 1.219040 | 0.899448 | 0.989253 | 1.024866 | 1.069626 | 1.208755 | 1.042765 | 0.869940 | 0.834227 | 0.632174 | 0.513056 | 0.454125 | 0.354239 | 0.316033 | 0.286897 | 0.263639 | 0.252427 | 0.258664 | 0.273175 | 0.297291 | 0.338955 | 0.440355 | 0.683543 | 0.955686 | 0.802313 | 0.692646 | 0.715749 | 0.688395 | 0.982097 | 1.083955 | 1.269379 | 2.145374 | 1.292396 | 0.619692 | 0.385893 | 0.282029 | 0.308125 | 0.408270 | 0.480169 | 0.766092 | 1.438255 | 2.772857 | 3.696064 | 1.674335 | 0.743675 | 0.466417 | 0.323027 | 0.266668 | 0.250655 | 0.254986 | 0.348593 | 0.515571 | 0.693202 | 0.836675 | 1.171650 | 1.663779 | 1.460886 | 0.814882 | 0.906608 | 0.840919 | 0.820403 | 0.934007 | 0.812766 | 0.751455 | 0.678071 | 0.480672 | 0.431146 | 0.491593 | 0.493049 | 0.673558 | 0.909816 | 1.248876 | 1.389875 | 1.181493 | 0.952567 | 0.778695 | 0.751204 | 0.969810 | 1.109280 | 1.119283 | 1.121432 | 1.173209 | 1.309293 | 0.902184 | 0.616147 | 0.503043 | 0.463350 | 0.486424 | 0.434848 | 0.390063 | 0.413651 | 0.390909 | 0.395376 | 0.391504 | 0.381785 | 0.374695 | 0.368271 | 0.349235 | 0.346438 | 0.306751 | 0.287042 | 0.254745 | 0.260508 | 0.251134 | 0.233169 | 0.229122 | 0.244218 | 0.263478 | 0.297807 | 0.320321 | 0.368438 | 0.563948 | 0.794416 | 0.780134 | 0.695872 | 0.719923 | 0.776847 | 0.724615 | 0.589073 | 0.510934 | 0.562154 | 0.791704 |
| right anterior | 0.296820 | 0.256162 | 0.241051 | 0.235282 | 0.229147 | 0.259363 | 0.230563 | 0.234839 | 0.285486 | 0.449434 | 0.678890 | 0.515804 | 0.224787 | 0.217852 | 0.220130 | 0.227202 | 0.259338 | 0.332833 | 0.250531 | 0.217972 | 0.302320 | 0.449057 | 0.438495 | 0.535376 | 0.501151 | 0.548672 | 0.559303 | 0.552104 | 0.303008 | 0.247015 | 0.215793 | 0.215519 | 0.231547 | 0.239176 | 0.268187 | 0.227888 | 0.233058 | 0.233906 | 0.239746 | 0.246501 | 0.374943 | 0.376423 | 0.370163 | 0.262065 | 0.224625 | 0.225442 | 0.225129 | 0.214803 | 0.218086 | 0.221685 | 0.239062 | 0.238409 | 0.215497 | 0.215365 | 0.220249 | 0.252926 | 0.323367 | 0.439044 | 0.617703 | 0.699067 | 0.748403 | 0.594160 | 0.399914 | 0.388363 | 0.445900 | 0.636665 | 1.252731 | 3.046256 | 7.012693 | 17.331338 | 21.869755 | 20.039144 | 10.693399 | 3.013452 | 1.287118 | 0.897383 | 0.603074 | 0.515278 | 0.415895 | 0.421769 | 0.529561 | 0.900436 | 1.531274 | 4.338163 | 16.995360 | 82.127631 | 159.074025 | 171.667022 | 134.088989 | 69.374941 | 31.512632 | 16.131023 | 12.813863 | 10.567889 | 11.907465 | 10.839653 | 7.078016 | 4.389340 | 2.380338 | 1.347875 | 0.899891 | 0.635567 | 0.465417 | 0.548559 | 0.842839 | 1.519075 | 4.217506 | 10.694373 | 12.143725 | 12.148402 | 7.513514 | 4.283928 | 2.926227 | 1.454865 | 0.967357 | 0.846220 | 0.629525 | 0.446968 | 0.364652 | 0.327070 | 0.373277 | 0.439322 | 0.383405 | 0.302726 | 0.239192 | 0.215120 | 0.230289 | 0.283160 | 0.352767 | 0.281209 | 0.227734 | 0.221730 | 0.236976 | 0.232443 | 0.220833 | 0.217954 | 0.214861 | 0.248660 | 0.437455 | 0.642263 | 0.662723 | 0.869607 | 1.741835 | 3.663066 | 5.341013 | 3.910494 | 3.764737 | 8.764399 | 3.177925 | 0.834155 | 0.383437 | 0.247983 | 0.233864 | 0.270921 | 0.272854 | 0.262559 | 0.255004 | 0.290356 | 0.420940 | 0.573271 | 0.468294 | 0.318735 | 0.264016 | 0.248056 | 0.225733 | 0.214660 | 0.225124 | 0.257577 | 0.269831 | 0.267745 | 0.241279 | 0.219815 | 0.214739 | 0.215284 | 0.217117 | 0.219392 | 0.223172 | 0.221881 | 0.221767 | 0.219270 | 0.217552 | 0.215042 | 0.219447 | 0.219984 | 0.218793 | 0.217733 | 0.214928 | 0.215699 | 0.214633 | 0.216794 | 0.214976 | 0.234115 | 0.297122 | 0.359961 | 0.339836 | 0.299727 | 0.281407 | 0.253320 | 0.224940 | 0.214873 | 0.215866 | 0.220300 | 0.215565 | 0.216760 | 0.214844 | 0.214663 | 0.215736 | 0.214847 | 0.214626 | 0.226877 | 0.250181 | 0.266220 | 0.268299 | 0.250341 | 0.234922 | 0.238826 | 0.218491 | 0.217540 | 0.231690 | 0.239873 | 0.284013 | 0.279612 | 0.268240 | 0.261776 | 0.252524 | 0.278759 | 0.339617 | 0.361562 | 0.363567 | 0.285239 | 0.275350 | 0.235634 | 0.214636 | 0.225562 | 0.232924 | 0.217354 | 0.236545 | 0.246551 | 0.271749 | 0.335423 | 0.326109 | 0.246463 | 0.215146 | 0.232962 | 0.230410 | 0.215639 | 0.218350 | 0.215443 | 0.215594 | 0.216321 | 0.220680 | 0.225026 | 0.241388 | 0.231191 | 0.241216 | 0.234907 | 0.227224 | 0.219456 | 0.222032 | 0.224247 | 0.222964 | 0.220932 | 0.216549 | 0.214998 | 0.215972 | 0.217942 | 0.223619 | 0.241521 | 0.263470 | 0.271025 | 0.257258 | 0.259497 | 0.275036 | 0.281395 | 0.277444 | 0.269471 | 0.268606 | 0.325983 | 0.346927 | 0.303357 |
| left central | 0.349728 | 0.478791 | 0.507831 | 0.451507 | 0.987583 | 3.780547 | 4.194017 | 1.070827 | 0.556498 | 0.512942 | 0.296418 | 0.267957 | 0.218025 | 0.229000 | 0.234836 | 0.226486 | 0.239211 | 0.296665 | 0.264157 | 0.299624 | 0.313182 | 0.300700 | 0.394255 | 0.547707 | 1.288038 | 2.527277 | 2.979360 | 1.694836 | 0.965709 | 0.433759 | 0.232602 | 0.399488 | 0.823037 | 1.059276 | 0.530730 | 0.245262 | 0.228058 | 0.223991 | 0.215943 | 0.221136 | 0.225803 | 0.217405 | 0.221848 | 0.215038 | 0.215270 | 0.219391 | 0.221383 | 0.223711 | 0.256029 | 0.340813 | 0.319999 | 0.282530 | 0.246743 | 0.253403 | 0.242558 | 0.250579 | 0.307483 | 0.557709 | 1.566281 | 5.221706 | 9.070476 | 10.168542 | 9.091959 | 8.013484 | 6.219577 | 3.843920 | 2.944332 | 4.195899 | 17.342584 | 43.256837 | 59.923841 | 43.614997 | 51.417078 | 121.420320 | 169.538950 | 81.908504 | 33.583269 | 13.035433 | 5.614637 | 2.014321 | 1.198072 | 0.878160 | 0.750680 | 0.646406 | 0.726461 | 1.107005 | 1.206778 | 1.081325 | 1.085591 | 1.385836 | 1.687722 | 1.776694 | 1.536987 | 1.413152 | 1.255111 | 1.098019 | 0.784417 | 0.549382 | 0.423213 | 0.390517 | 0.421765 | 0.407080 | 0.394468 | 0.414345 | 0.474570 | 0.540741 | 0.417239 | 0.299903 | 0.265282 | 0.244795 | 0.232989 | 0.225677 | 0.214678 | 0.215714 | 0.230259 | 0.217556 | 0.229917 | 0.261834 | 0.269952 | 0.231339 | 0.219039 | 0.224309 | 0.227935 | 0.215572 | 0.215347 | 0.227658 | 0.260651 | 0.350386 | 0.393251 | 0.329880 | 0.283089 | 0.240699 | 0.230624 | 0.270441 | 0.304341 | 0.278469 | 0.320383 | 0.330792 | 0.380506 | 0.325818 | 0.237861 | 0.217213 | 0.222303 | 0.215219 | 0.214637 | 0.215159 | 0.221068 | 0.248473 | 0.318389 | 0.343857 | 0.370330 | 0.282460 | 0.236898 | 0.231996 | 0.221643 | 0.223705 | 0.238675 | 0.278972 | 0.383207 | 0.437405 | 0.321230 | 0.293742 | 0.241478 | 0.235766 | 0.231332 | 0.225260 | 0.218487 | 0.217357 | 0.216541 | 0.226718 | 0.249682 | 0.316406 | 0.410192 | 0.628755 | 1.097417 | 1.734773 | 2.158064 | 1.229976 | 0.509297 | 0.403300 | 0.338306 | 0.302257 | 0.262201 | 0.250615 | 0.250909 | 0.243876 | 0.237623 | 0.226685 | 0.214896 | 0.214626 | 0.215402 | 0.218996 | 0.266106 | 0.375903 | 0.480141 | 0.735221 | 1.010165 | 0.849410 | 0.554100 | 0.350868 | 0.298303 | 0.327928 | 0.402425 | 0.451814 | 0.499656 | 0.443016 | 0.396133 | 0.346678 | 0.272179 | 0.229952 | 0.214885 | 0.235510 | 0.301311 | 0.507871 | 1.093852 | 1.450640 | 1.464668 | 1.104575 | 0.912613 | 0.740030 | 0.549647 | 0.526948 | 0.507414 | 0.556131 | 0.671304 | 0.711123 | 0.587502 | 0.412838 | 0.275970 | 0.216505 | 0.217639 | 0.231888 | 0.244086 | 0.247419 | 0.241323 | 0.235888 | 0.215230 | 0.214887 | 0.227753 | 0.248669 | 0.269851 | 0.283306 | 0.256294 | 0.222770 | 0.219590 | 0.222706 | 0.232862 | 0.249956 | 0.259712 | 0.316245 | 0.413323 | 0.429949 | 0.369785 | 0.253143 | 0.224433 | 0.227374 | 0.233345 | 0.223663 | 0.215501 | 0.216686 | 0.215134 | 0.216071 | 0.225782 | 0.242412 | 0.225743 | 0.216247 | 0.216962 | 0.223460 | 0.226251 | 0.248404 | 0.282060 | 0.257309 | 0.249319 | 0.246082 | 0.233559 | 0.220314 | 0.216798 | 0.215125 | 0.221874 | 0.235403 |
| right central | 0.232822 | 0.222336 | 0.226837 | 0.431079 | 0.498696 | 0.960303 | 2.479289 | 3.321681 | 13.672045 | 13.489302 | 0.993524 | 1.590285 | 1.469755 | 1.002295 | 0.542548 | 0.328924 | 0.271496 | 0.350878 | 0.228455 | 0.239430 | 0.320139 | 0.439939 | 0.332970 | 0.297023 | 0.310533 | 0.245907 | 0.214904 | 0.256831 | 0.495434 | 1.226214 | 0.536662 | 0.273364 | 0.214903 | 0.216925 | 0.272665 | 0.245990 | 0.249370 | 0.261664 | 0.241266 | 0.243260 | 0.288543 | 0.239755 | 0.221998 | 0.220645 | 0.217779 | 0.236370 | 0.250466 | 0.257757 | 0.227669 | 0.217089 | 0.262758 | 0.288970 | 0.328183 | 0.282808 | 0.266283 | 0.247666 | 0.217676 | 0.216934 | 0.215100 | 0.215935 | 0.248124 | 0.349218 | 0.434498 | 0.629829 | 1.705538 | 4.474673 | 15.072882 | 30.139159 | 40.595161 | 80.124079 | 165.961981 | 164.220775 | 167.971599 | 118.854928 | 41.656424 | 11.780666 | 5.054394 | 1.928451 | 1.154640 | 0.921626 | 1.134832 | 2.150659 | 3.771676 | 4.011294 | 6.325744 | 8.317174 | 14.227365 | 14.238047 | 9.156149 | 8.600064 | 11.989619 | 16.566819 | 20.450578 | 35.729738 | 94.267625 | 254.395332 | 388.980764 | 201.560627 | 37.568636 | 41.865033 | 32.979182 | 17.059080 | 11.752499 | 6.244193 | 1.471825 | 1.060958 | 0.798929 | 0.865457 | 0.520020 | 0.346314 | 0.363755 | 0.520131 | 0.555284 | 0.461340 | 0.301341 | 0.261722 | 0.226605 | 0.223117 | 0.255744 | 0.251423 | 0.233336 | 0.220545 | 0.221026 | 0.214838 | 0.217462 | 0.215392 | 0.214720 | 0.216885 | 0.231535 | 0.227427 | 0.234478 | 0.224721 | 0.219429 | 0.214626 | 0.227970 | 0.242743 | 0.253706 | 0.256442 | 0.266001 | 0.309041 | 0.272908 | 0.242090 | 0.235856 | 0.226959 | 0.218354 | 0.215643 | 0.235591 | 0.257517 | 0.273386 | 0.254981 | 0.230105 | 0.218041 | 0.214740 | 0.214742 | 0.215007 | 0.215617 | 0.214765 | 0.214710 | 0.240254 | 0.280028 | 0.442051 | 1.014654 | 1.929440 | 4.038286 | 10.290596 | 8.906254 | 14.344735 | 19.413749 | 11.610095 | 2.343701 | 0.388412 | 0.214647 | 0.424385 | 1.100754 | 1.354831 | 0.944528 | 0.429787 | 0.261777 | 0.242608 | 0.216218 | 0.215056 | 0.215549 | 0.217386 | 0.235574 | 0.291657 | 0.275122 | 0.299026 | 0.355270 | 0.433209 | 0.318644 | 0.244129 | 0.217159 | 0.214859 | 0.216462 | 0.214694 | 0.219721 | 0.243463 | 0.321589 | 0.438910 | 0.435385 | 0.308787 | 0.237976 | 0.219677 | 0.218416 | 0.215513 | 0.216023 | 0.235983 | 0.293765 | 0.367078 | 0.411195 | 0.396130 | 0.373526 | 0.314853 | 0.270378 | 0.261499 | 0.246045 | 0.237258 | 0.244399 | 0.260310 | 0.313430 | 0.347385 | 0.404903 | 0.347624 | 0.278121 | 0.247403 | 0.247044 | 0.256622 | 0.362679 | 0.376134 | 0.618599 | 0.816516 | 0.792066 | 0.756228 | 0.396511 | 0.244512 | 0.227213 | 0.229175 | 0.253628 | 0.310866 | 0.389367 | 0.469648 | 0.462822 | 0.376142 | 0.352323 | 0.331135 | 0.324452 | 0.309189 | 0.332332 | 0.429499 | 0.662929 | 0.624995 | 0.621183 | 0.513718 | 0.466417 | 0.494848 | 0.439415 | 0.401812 | 0.423947 | 0.490623 | 0.555493 | 0.424986 | 0.440693 | 0.602558 | 0.790923 | 0.889278 | 0.679910 | 0.568500 | 0.601832 | 0.490828 | 0.386279 | 0.311415 | 0.299382 | 0.295940 | 0.298903 | 0.313847 | 0.385769 | 0.667348 | 0.859677 | 1.042467 | 1.324855 |
| left posterior | 0.340836 | 0.583696 | 1.000669 | 1.105780 | 1.534666 | 1.625647 | 0.778638 | 0.308587 | 0.217004 | 0.256126 | 0.385834 | 0.574157 | 0.634362 | 0.574000 | 0.426794 | 0.322325 | 0.228742 | 0.221763 | 0.288012 | 0.321890 | 0.343332 | 0.340768 | 0.274812 | 0.283518 | 0.282865 | 0.262115 | 0.428463 | 0.527302 | 2.400617 | 8.748947 | 18.033484 | 22.534098 | 29.259201 | 7.317767 | 1.871195 | 0.519544 | 0.304750 | 0.222990 | 0.221277 | 0.308278 | 0.412507 | 0.418118 | 0.443198 | 0.338433 | 0.259781 | 0.225169 | 0.216779 | 0.253127 | 0.376980 | 0.560825 | 1.039806 | 2.290266 | 10.144037 | 34.509683 | 153.038393 | 430.325762 | 610.551715 | 87.094532 | 13.366945 | 3.227631 | 1.683003 | 0.596992 | 0.215042 | 0.561258 | 4.181063 | 42.999236 | 130.164051 | 505.819042 | 1940.535954 | 5993.127561 | 8280.876312 | 11522.148237 | 7474.081633 | 5721.618841 | 6664.682016 | 1843.851587 | 526.335277 | 148.940501 | 41.077597 | 9.232650 | 3.072899 | 1.671962 | 1.508707 | 2.121304 | 2.474984 | 2.606588 | 3.823990 | 6.753024 | 7.813043 | 9.346308 | 9.035134 | 9.315689 | 10.479395 | 11.655950 | 11.227295 | 13.797846 | 11.014449 | 8.235184 | 6.457412 | 7.658770 | 9.306018 | 11.133214 | 14.096419 | 23.341375 | 29.764722 | 36.486437 | 25.498328 | 11.646586 | 6.907211 | 4.628231 | 4.508303 | 5.451851 | 5.326256 | 6.480582 | 12.914235 | 25.384285 | 50.127508 | 51.874385 | 54.633399 | 76.948836 | 77.111954 | 59.618268 | 44.456419 | 40.873707 | 44.247058 | 25.031277 | 11.320293 | 5.769684 | 4.233847 | 3.055844 | 1.704365 | 1.291950 | 1.453851 | 1.788934 | 2.289816 | 2.418223 | 2.221161 | 2.981070 | 4.185133 | 5.149721 | 5.857319 | 5.644886 | 6.113201 | 6.356005 | 4.598448 | 3.303677 | 2.221685 | 1.466650 | 1.134865 | 0.766048 | 0.607578 | 0.541276 | 0.465621 | 0.477049 | 0.533596 | 0.590869 | 0.758708 | 0.957074 | 1.317937 | 1.946291 | 2.952976 | 5.147837 | 8.448902 | 14.744253 | 17.223485 | 15.470616 | 14.925464 | 8.764556 | 5.197724 | 2.999782 | 1.879988 | 1.395827 | 0.918664 | 0.820846 | 0.924718 | 0.937167 | 0.960816 | 1.405852 | 1.949226 | 2.686872 | 4.738190 | 7.503516 | 8.660927 | 8.866892 | 4.920615 | 3.263418 | 2.337917 | 1.233616 | 0.775925 | 0.623655 | 0.551774 | 0.478948 | 0.442248 | 0.437086 | 0.500667 | 0.499429 | 0.464658 | 0.518640 | 0.641284 | 0.929634 | 1.152254 | 1.364439 | 1.827748 | 2.439978 | 2.998913 | 3.643092 | 3.028708 | 2.842569 | 2.621417 | 1.820649 | 1.817639 | 2.018726 | 1.886948 | 1.941092 | 2.224051 | 2.090135 | 2.330495 | 2.259626 | 1.620686 | 1.639790 | 1.634471 | 2.320088 | 3.356739 | 3.948279 | 4.421476 | 7.457415 | 8.615166 | 9.524291 | 7.679386 | 6.541988 | 5.545607 | 4.837868 | 4.045295 | 5.618595 | 6.298667 | 8.721454 | 8.841883 | 8.078948 | 7.236766 | 5.407198 | 3.603680 | 2.869116 | 1.835332 | 1.502986 | 1.266725 | 0.945232 | 0.901330 | 0.945768 | 1.144305 | 1.228677 | 1.152742 | 1.207958 | 1.521878 | 1.806545 | 1.619353 | 1.443886 | 1.970174 | 2.715106 | 3.866635 | 3.667690 | 2.920334 | 2.554912 | 2.331778 | 1.580432 | 1.365605 | 1.192820 | 1.111132 | 1.037745 | 1.083447 | 0.936333 | 0.878018 | 0.793817 | 0.724361 | 0.687140 | 0.739492 | 0.655026 | 0.605893 | 0.647320 | 0.692698 | 0.613750 |
| right posterior | 0.396125 | 0.379644 | 0.360093 | 0.464013 | 0.530163 | 0.377683 | 0.256513 | 0.216478 | 0.218708 | 0.226429 | 0.218710 | 0.253567 | 0.239435 | 0.248355 | 0.237885 | 0.294219 | 0.270312 | 0.216923 | 0.234416 | 0.214908 | 0.216019 | 0.219380 | 0.258943 | 0.233685 | 0.214826 | 0.224844 | 0.317485 | 0.384988 | 0.643049 | 0.853169 | 0.760753 | 0.639910 | 0.478029 | 0.285372 | 0.411797 | 0.363306 | 0.285460 | 0.308278 | 0.312892 | 0.255118 | 0.228240 | 0.214677 | 0.216536 | 0.224524 | 0.230233 | 0.220531 | 0.219981 | 0.214626 | 0.220975 | 0.217724 | 0.230540 | 0.232100 | 0.216053 | 0.215528 | 0.229235 | 0.271926 | 0.309633 | 0.444561 | 0.991102 | 2.200289 | 1.064167 | 0.640663 | 0.496253 | 0.539754 | 0.739794 | 1.423568 | 3.252859 | 11.106136 | 38.510615 | 150.212481 | 333.233699 | 547.061851 | 427.234380 | 415.119194 | 305.023874 | 255.176547 | 119.698421 | 57.694548 | 26.941349 | 7.142488 | 1.895046 | 1.281297 | 0.954372 | 0.659931 | 0.574075 | 0.473955 | 0.689262 | 1.332138 | 1.391416 | 2.345632 | 6.045751 | 17.239047 | 36.608655 | 48.866379 | 58.271106 | 62.459195 | 61.712615 | 41.284765 | 26.468161 | 20.191245 | 17.063810 | 10.771672 | 9.263522 | 7.271307 | 6.558178 | 4.441298 | 3.688844 | 3.272841 | 2.489263 | 2.566045 | 4.728627 | 5.443548 | 5.856799 | 6.157118 | 6.090375 | 9.469930 | 13.245324 | 12.189501 | 11.198233 | 7.095662 | 5.108470 | 6.636845 | 7.735413 | 8.020833 | 5.988790 | 6.022178 | 9.811215 | 16.322325 | 24.422162 | 23.764455 | 18.655960 | 16.519876 | 17.829990 | 20.224690 | 21.862898 | 12.334312 | 12.913809 | 19.023139 | 30.110558 | 23.338865 | 25.313903 | 24.093370 | 26.502201 | 27.638556 | 21.854818 | 15.882756 | 15.750320 | 9.467636 | 3.834005 | 2.092823 | 1.310751 | 0.940737 | 0.694451 | 0.509860 | 0.515174 | 0.574734 | 0.712375 | 0.994376 | 1.881668 | 2.842723 | 3.757854 | 3.575602 | 4.494258 | 5.821211 | 3.769395 | 2.068216 | 1.545292 | 1.542552 | 1.239405 | 0.813277 | 0.570331 | 0.467502 | 0.416699 | 0.338099 | 0.284719 | 0.320273 | 0.411747 | 0.466575 | 0.540500 | 0.541607 | 0.636425 | 0.763720 | 0.600518 | 0.558180 | 0.554117 | 0.564461 | 0.795833 | 1.045656 | 1.340970 | 1.200984 | 0.701793 | 0.438602 | 0.331474 | 0.297164 | 0.288812 | 0.269015 | 0.237480 | 0.223000 | 0.225706 | 0.244911 | 0.274229 | 0.313767 | 0.345727 | 0.447234 | 0.787150 | 1.051877 | 1.285139 | 1.222448 | 1.105527 | 0.922023 | 1.129057 | 1.234440 | 1.498676 | 1.295697 | 1.239823 | 1.181383 | 1.510130 | 1.286598 | 1.034369 | 0.837513 | 0.759226 | 0.644888 | 0.573897 | 0.437803 | 0.407415 | 0.388394 | 0.493746 | 0.534089 | 0.596641 | 0.554434 | 0.494779 | 0.476244 | 0.537736 | 0.496833 | 0.477801 | 0.398746 | 0.398586 | 0.463685 | 0.509373 | 0.412469 | 0.327846 | 0.279738 | 0.270747 | 0.256489 | 0.244552 | 0.251566 | 0.279409 | 0.323066 | 0.362207 | 0.418091 | 0.438921 | 0.539095 | 0.591905 | 0.621896 | 0.471799 | 0.457024 | 0.394834 | 0.375176 | 0.284705 | 0.236091 | 0.215819 | 0.214656 | 0.217489 | 0.240352 | 0.276222 | 0.306883 | 0.322139 | 0.280384 | 0.237401 | 0.220453 | 0.218853 | 0.242607 | 0.287794 | 0.336686 | 0.325995 | 0.356162 | 0.456600 | 0.454811 | 0.524152 | 0.599456 |
| all electrodes | 0.284030 | 0.239891 | 0.220515 | 0.218935 | 0.293475 | 0.303217 | 0.217302 | 0.215045 | 0.215518 | 0.238440 | 0.226942 | 0.232386 | 0.304448 | 0.290645 | 0.247271 | 0.218145 | 0.231113 | 0.261061 | 0.256849 | 0.214693 | 0.221033 | 0.215857 | 0.227771 | 0.233315 | 0.214848 | 0.224015 | 0.220221 | 0.216270 | 0.225002 | 0.319325 | 0.460710 | 0.473316 | 0.462082 | 0.554800 | 0.783394 | 0.768622 | 0.333147 | 0.278417 | 0.460212 | 0.710819 | 0.912700 | 0.831558 | 0.903514 | 1.248669 | 1.237385 | 1.275510 | 1.091560 | 1.014050 | 0.886735 | 0.642327 | 0.442032 | 0.331787 | 0.247016 | 0.224559 | 0.216483 | 0.215597 | 0.214928 | 0.219140 | 0.248829 | 0.316408 | 0.378688 | 0.474672 | 0.867344 | 1.632395 | 3.487179 | 8.278737 | 23.096423 | 96.661446 | 285.294632 | 445.608762 | 595.366124 | 870.619307 | 687.983651 | 524.654301 | 400.689569 | 411.321187 | 276.746951 | 102.479008 | 19.391307 | 5.082371 | 2.344372 | 1.359318 | 0.944476 | 0.843968 | 0.917519 | 1.200680 | 1.428074 | 2.068987 | 2.963754 | 5.063611 | 11.285323 | 28.929747 | 70.688042 | 217.128207 | 587.209835 | 576.477547 | 245.283130 | 59.851331 | 13.957206 | 3.814271 | 1.649927 | 0.815422 | 0.591527 | 0.465174 | 0.444408 | 0.459235 | 0.471611 | 0.375416 | 0.365201 | 0.440543 | 0.777193 | 1.270582 | 2.147361 | 3.438401 | 9.901817 | 14.812484 | 17.346116 | 12.547200 | 7.211020 | 6.688196 | 5.260933 | 2.654433 | 1.838944 | 1.656476 | 1.206275 | 0.900015 | 0.777781 | 0.679802 | 0.782357 | 0.921001 | 1.012652 | 1.064814 | 1.047586 | 0.835403 | 0.939573 | 0.923883 | 0.895488 | 0.758854 | 0.873623 | 1.219023 | 1.792850 | 1.708502 | 1.713912 | 1.735287 | 1.987746 | 1.606486 | 0.901215 | 0.528496 | 0.412005 | 0.365031 | 0.348072 | 0.323458 | 0.328334 | 0.382168 | 0.443726 | 0.509832 | 0.470027 | 0.543678 | 0.692598 | 0.691825 | 0.732096 | 0.805966 | 0.972475 | 1.220746 | 1.104516 | 0.766400 | 0.777728 | 0.809512 | 0.912517 | 0.786410 | 0.526111 | 0.362756 | 0.354212 | 0.386001 | 0.422099 | 0.456511 | 0.621088 | 1.015191 | 1.580762 | 1.612854 | 1.362203 | 1.442841 | 1.249146 | 0.850187 | 0.717699 | 0.807027 | 0.990882 | 1.258745 | 1.833456 | 2.879107 | 3.029120 | 2.394249 | 1.469473 | 1.136663 | 1.088733 | 1.055995 | 1.222208 | 1.703385 | 2.423727 | 2.845084 | 3.878551 | 3.050588 | 2.264764 | 1.673348 | 1.036923 | 0.875169 | 0.975176 | 1.036455 | 2.017380 | 4.119017 | 9.036535 | 20.240149 | 25.313722 | 30.924203 | 25.764543 | 11.289799 | 5.642569 | 4.627158 | 6.996970 | 10.149385 | 8.516214 | 7.991153 | 9.018689 | 9.669036 | 6.390867 | 3.206240 | 1.830712 | 1.879598 | 2.703271 | 3.202403 | 3.350778 | 4.561869 | 5.080399 | 4.698131 | 5.329602 | 5.245571 | 4.219547 | 3.914649 | 3.847301 | 4.445370 | 8.599808 | 18.762189 | 35.239279 | 45.119724 | 25.708504 | 22.912822 | 26.394438 | 22.760145 | 9.529558 | 8.101834 | 8.120664 | 12.926976 | 13.924972 | 12.608871 | 14.167176 | 19.689670 | 14.823721 | 9.462785 | 7.383060 | 5.920814 | 4.340194 | 1.643869 | 0.626029 | 0.391470 | 0.299794 | 0.300942 | 0.284573 | 0.257178 | 0.273104 | 0.378732 | 0.511691 | 0.570463 | 0.357530 | 0.305231 | 0.288249 | 0.292784 | 0.267099 | 0.240515 | 0.237962 | 0.278279 |

Searchlight, spatiotemporal cluster permutation test

|  | start time | stop time | peak time | peak channel | cluster p | peak Cohen's d | direction |
| --- | --- | --- | --- | --- | --- | --- | --- |
| #1 | 75 | 1195 | 155 | POz | 0.0003 | 1.695485 | positive |

G) happy vs angry

  
|  | time window | peak latency | cluster *p* | peak Cohen's *d* |  | | | |
| **all electrodes** | 145 - 215 ms | 175 ms | 0.0267 | 1.2307 |  | | | |
|  | | | | | | | | |

Time-resolved classification, cluster permutation tests

|  | **left hemisphere** | | | | **right hemisphere** | | | |
|  | time window | peak latency | cluster *p* | peak Cohen's *d* | time window | peak latency | cluster *p* | peak Cohen's *d* |
| **anterior** |  | | | |  | | | |
| **central** |  | | | |  | | | |
| **posterior** | 125 - 270 ms | 170 ms | 0.0106 | 1.0986 | 145 - 215 ms | 170 ms | 0.0475 | 1.1431 |
 320 - 500 ms | 405 ms | 0.013 | 0.7323 | 360 - 525 ms | 415 ms | 0.0284 | 0.8843 |  | | | | 650 - 1125 ms | 1005 ms | 0.0047 | 0.7603 |

  

Time-resolved classification, Bayesian statistics

|  | -200 | -195 | -190 | -185 | -180 | -175 | -170 | -165 | -160 | -155 | -150 | -145 | -140 | -135 | -130 | -125 | -120 | -115 | -110 | -105 | -100 | -95 | -90 | -85 | -80 | -75 | -70 | -65 | -60 | -55 | -50 | -45 | -40 | -35 | -30 | -25 | -20 | -15 | -10 | -5 | 0 | 5 | 10 | 15 | 20 | 25 | 30 | 35 | 40 | 45 | 50 | 55 | 60 | 65 | 70 | 75 | 80 | 85 | 90 | 95 | 100 | 105 | 110 | 115 | 120 | 125 | 130 | 135 | 140 | 145 | 150 | 155 | 160 | 165 | 170 | 175 | 180 | 185 | 190 | 195 | 200 | 205 | 210 | 215 | 220 | 225 | 230 | 235 | 240 | 245 | 250 | 255 | 260 | 265 | 270 | 275 | 280 | 285 | 290 | 295 | 300 | 305 | 310 | 315 | 320 | 325 | 330 | 335 | 340 | 345 | 350 | 355 | 360 | 365 | 370 | 375 | 380 | 385 | 390 | 395 | 400 | 405 | 410 | 415 | 420 | 425 | 430 | 435 | 440 | 445 | 450 | 455 | 460 | 465 | 470 | 475 | 480 | 485 | 490 | 495 | 500 | 505 | 510 | 515 | 520 | 525 | 530 | 535 | 540 | 545 | 550 | 555 | 560 | 565 | 570 | 575 | 580 | 585 | 590 | 595 | 600 | 605 | 610 | 615 | 620 | 625 | 630 | 635 | 640 | 645 | 650 | 655 | 660 | 665 | 670 | 675 | 680 | 685 | 690 | 695 | 700 | 705 | 710 | 715 | 720 | 725 | 730 | 735 | 740 | 745 | 750 | 755 | 760 | 765 | 770 | 775 | 780 | 785 | 790 | 795 | 800 | 805 | 810 | 815 | 820 | 825 | 830 | 835 | 840 | 845 | 850 | 855 | 860 | 865 | 870 | 875 | 880 | 885 | 890 | 895 | 900 | 905 | 910 | 915 | 920 | 925 | 930 | 935 | 940 | 945 | 950 | 955 | 960 | 965 | 970 | 975 | 980 | 985 | 990 | 995 | 1000 | 1005 | 1010 | 1015 | 1020 | 1025 | 1030 | 1035 | 1040 | 1045 | 1050 | 1055 | 1060 | 1065 | 1070 | 1075 | 1080 | 1085 | 1090 | 1095 | 1100 | 1105 | 1110 | 1115 | 1120 | 1125 | 1130 | 1135 | 1140 | 1145 | 1150 | 1155 | 1160 | 1165 | 1170 | 1175 | 1180 | 1185 | 1190 | 1195 |
| --- | --- | --- | --- | --- | --- | --- | --- | --- | --- | --- | --- | --- | --- | --- | --- | --- | --- | --- | --- | --- | --- | --- | --- | --- | --- | --- | --- | --- | --- | --- | --- | --- | --- | --- | --- | --- | --- | --- | --- | --- | --- | --- | --- | --- | --- | --- | --- | --- | --- | --- | --- | --- | --- | --- | --- | --- | --- | --- | --- | --- | --- | --- | --- | --- | --- | --- | --- | --- | --- | --- | --- | --- | --- | --- | --- | --- | --- | --- | --- | --- | --- | --- | --- | --- | --- | --- | --- | --- | --- | --- | --- | --- | --- | --- | --- | --- | --- | --- | --- | --- | --- | --- | --- | --- | --- | --- | --- | --- | --- | --- | --- | --- | --- | --- | --- | --- | --- | --- | --- | --- | --- | --- | --- | --- | --- | --- | --- | --- | --- | --- | --- | --- | --- | --- | --- | --- | --- | --- | --- | --- | --- | --- | --- | --- | --- | --- | --- | --- | --- | --- | --- | --- | --- | --- | --- | --- | --- | --- | --- | --- | --- | --- | --- | --- | --- | --- | --- | --- | --- | --- | --- | --- | --- | --- | --- | --- | --- | --- | --- | --- | --- | --- | --- | --- | --- | --- | --- | --- | --- | --- | --- | --- | --- | --- | --- | --- | --- | --- | --- | --- | --- | --- | --- | --- | --- | --- | --- | --- | --- | --- | --- | --- | --- | --- | --- | --- | --- | --- | --- | --- | --- | --- | --- | --- | --- | --- | --- | --- | --- | --- | --- | --- | --- | --- | --- | --- | --- | --- | --- | --- | --- | --- | --- | --- | --- | --- | --- | --- | --- | --- | --- | --- | --- | --- | --- | --- | --- | --- | --- | --- | --- | --- | --- | --- | --- | --- | --- | --- | --- | --- | --- | --- | --- | --- | --- | --- | --- | --- | --- | --- |
| left anterior | 0.216576 | 0.256940 | 0.528384 | 1.520630 | 2.025647 | 1.324653 | 0.482304 | 0.339386 | 0.235851 | 0.227339 | 0.270564 | 0.304879 | 0.320703 | 0.244329 | 0.216492 | 0.224831 | 0.224958 | 0.216176 | 0.229564 | 0.218382 | 0.231437 | 0.217977 | 0.216287 | 0.233798 | 0.257935 | 0.245846 | 0.511968 | 1.080099 | 1.155465 | 1.271121 | 1.381020 | 0.944026 | 0.432719 | 0.236826 | 0.217125 | 0.227111 | 0.333068 | 1.162770 | 1.907566 | 0.695936 | 0.289190 | 0.312455 | 0.396228 | 0.376873 | 0.338598 | 0.439433 | 0.430817 | 0.380169 | 0.252717 | 0.310105 | 0.352311 | 0.310642 | 0.294153 | 0.266081 | 0.353953 | 0.288949 | 0.214639 | 0.218125 | 0.222746 | 0.262107 | 0.237877 | 0.232276 | 0.214935 | 0.231064 | 0.246743 | 0.234997 | 0.251625 | 0.253767 | 0.239680 | 0.216378 | 0.215549 | 0.223403 | 0.230372 | 0.242654 | 0.286945 | 0.318156 | 0.302300 | 0.331231 | 0.324301 | 0.225918 | 0.233555 | 0.393186 | 0.675338 | 1.395096 | 4.691520 | 10.355621 | 9.531017 | 6.353362 | 5.138613 | 3.854222 | 1.369431 | 0.703304 | 0.903061 | 1.300979 | 1.812108 | 1.405626 | 1.056135 | 1.289786 | 1.767468 | 0.854584 | 0.554402 | 0.584445 | 0.470944 | 0.431574 | 0.452843 | 0.359400 | 0.401953 | 0.419176 | 0.344653 | 0.282417 | 0.224377 | 0.216377 | 0.225389 | 0.235993 | 0.269241 | 0.338155 | 0.507027 | 0.635796 | 0.643559 | 0.778257 | 0.918489 | 0.685492 | 0.452074 | 0.277014 | 0.221746 | 0.214980 | 0.214640 | 0.216344 | 0.236424 | 0.296286 | 0.294620 | 0.258574 | 0.281510 | 0.303676 | 0.324756 | 0.380270 | 0.495997 | 0.616907 | 0.788300 | 0.748468 | 0.841326 | 0.829566 | 0.492663 | 0.284283 | 0.233252 | 0.220047 | 0.215441 | 0.224769 | 0.260815 | 0.328318 | 0.343882 | 0.319702 | 0.254035 | 0.214987 | 0.279827 | 0.457766 | 0.940049 | 1.019436 | 1.131949 | 1.577889 | 2.376254 | 3.528365 | 4.505505 | 3.501552 | 4.876668 | 4.353357 | 1.876409 | 0.744404 | 0.501450 | 0.359006 | 0.287200 | 0.240902 | 0.243687 | 0.255002 | 0.275968 | 0.279010 | 0.297729 | 0.316417 | 0.319779 | 0.289382 | 0.255497 | 0.226775 | 0.217619 | 0.218642 | 0.230707 | 0.248261 | 0.262338 | 0.308827 | 0.515312 | 0.941624 | 1.237245 | 1.341827 | 1.112920 | 1.412414 | 0.969920 | 0.504704 | 0.314332 | 0.257725 | 0.254701 | 0.274228 | 0.271694 | 0.347538 | 0.351428 | 0.359014 | 0.358735 | 0.265560 | 0.240003 | 0.219745 | 0.214636 | 0.216223 | 0.223087 | 0.225336 | 0.222581 | 0.224964 | 0.241637 | 0.245512 | 0.215725 | 0.229264 | 0.262733 | 0.252400 | 0.240901 | 0.302267 | 0.303170 | 0.257059 | 0.277525 | 0.303540 | 0.379376 | 0.511634 | 0.485139 | 0.579154 | 0.627368 | 0.508522 | 0.422355 | 0.318323 | 0.268091 | 0.248156 | 0.248080 | 0.227893 | 0.218620 | 0.214716 | 0.215091 | 0.214771 | 0.215530 | 0.216431 | 0.216941 | 0.220855 | 0.240835 | 0.259064 | 0.276397 | 0.235706 | 0.214793 | 0.216906 | 0.217288 | 0.214744 | 0.214689 | 0.215456 | 0.215630 | 0.224280 | 0.232770 | 0.235755 | 0.222243 | 0.235896 | 0.250534 | 0.255029 | 0.271188 | 0.273294 | 0.247910 | 0.236013 | 0.221123 | 0.216618 | 0.215013 | 0.219396 | 0.226410 | 0.237312 | 0.233820 | 0.253278 | 0.258856 | 0.261077 | 0.253213 | 0.229055 |
| right anterior | 1.801621 | 2.205024 | 1.111110 | 0.412696 | 0.245552 | 0.218566 | 0.230847 | 0.378796 | 0.570143 | 1.169602 | 0.546999 | 0.353769 | 0.374733 | 0.462600 | 0.714085 | 0.618934 | 0.285112 | 0.347772 | 0.581122 | 0.393684 | 0.236209 | 0.219778 | 0.218179 | 0.228892 | 0.285382 | 0.380027 | 0.518154 | 0.724174 | 0.563217 | 0.340441 | 0.218771 | 0.235120 | 0.341919 | 0.645580 | 2.103285 | 4.196599 | 2.540627 | 1.605433 | 0.946930 | 0.669748 | 0.642858 | 0.595593 | 0.298768 | 0.308104 | 0.356124 | 0.560218 | 0.526683 | 0.528812 | 0.396721 | 0.559521 | 0.735354 | 0.787807 | 0.681708 | 1.167348 | 0.834429 | 0.562835 | 0.350575 | 0.337918 | 0.352464 | 0.275988 | 0.220032 | 0.215120 | 0.217487 | 0.229523 | 0.244106 | 0.246727 | 0.230347 | 0.228606 | 0.251664 | 0.260037 | 0.330175 | 0.535569 | 1.398772 | 3.209684 | 4.317153 | 1.237646 | 1.114794 | 0.453198 | 0.354873 | 0.310450 | 0.268232 | 0.230210 | 0.225852 | 0.230953 | 0.277656 | 0.450706 | 0.414588 | 0.419062 | 0.396216 | 0.486481 | 0.546018 | 0.725621 | 0.556592 | 0.741798 | 0.756323 | 0.852967 | 0.608818 | 0.470968 | 0.402867 | 0.490990 | 0.573782 | 0.672358 | 0.535507 | 0.491733 | 0.319882 | 0.256816 | 0.225160 | 0.214759 | 0.223080 | 0.256328 | 0.341491 | 0.349188 | 0.418600 | 0.479804 | 0.548458 | 0.851343 | 1.658888 | 2.253805 | 3.015780 | 1.945877 | 1.403150 | 0.947439 | 0.606038 | 0.431626 | 0.422407 | 0.461922 | 0.673509 | 0.653113 | 0.533649 | 0.340517 | 0.249829 | 0.214946 | 0.225802 | 0.250929 | 0.238118 | 0.228707 | 0.221476 | 0.226564 | 0.248478 | 0.274893 | 0.281334 | 0.264317 | 0.236441 | 0.219218 | 0.216214 | 0.218144 | 0.239482 | 0.246426 | 0.237636 | 0.243115 | 0.249215 | 0.291248 | 0.323708 | 0.263968 | 0.216356 | 0.238617 | 0.300614 | 0.365046 | 0.428962 | 0.648257 | 0.684591 | 0.478295 | 0.434879 | 0.455969 | 0.426667 | 0.587433 | 0.573311 | 0.711928 | 0.745951 | 0.464952 | 0.451135 | 0.616963 | 0.497977 | 0.446133 | 0.373918 | 0.417912 | 0.442971 | 0.386630 | 0.317645 | 0.286684 | 0.258223 | 0.221296 | 0.215235 | 0.217130 | 0.214890 | 0.214626 | 0.215089 | 0.215865 | 0.229567 | 0.246889 | 0.279768 | 0.283466 | 0.291908 | 0.279963 | 0.272169 | 0.257917 | 0.247238 | 0.217210 | 0.216809 | 0.222975 | 0.219544 | 0.214679 | 0.221156 | 0.225743 | 0.215575 | 0.225515 | 0.248030 | 0.277591 | 0.320951 | 0.352318 | 0.349386 | 0.318433 | 0.271365 | 0.283169 | 0.282705 | 0.284721 | 0.277894 | 0.289975 | 0.289111 | 0.291881 | 0.267795 | 0.269226 | 0.328874 | 0.475252 | 0.578441 | 0.712195 | 0.611867 | 0.516553 | 0.434746 | 0.336029 | 0.306855 | 0.311188 | 0.294583 | 0.321401 | 0.340636 | 0.345875 | 0.373873 | 0.314589 | 0.283695 | 0.269437 | 0.254116 | 0.246024 | 0.239985 | 0.231738 | 0.237406 | 0.231377 | 0.234765 | 0.236451 | 0.250096 | 0.286071 | 0.300046 | 0.291213 | 0.286999 | 0.274847 | 0.284148 | 0.308376 | 0.299426 | 0.271879 | 0.245879 | 0.251810 | 0.256746 | 0.257103 | 0.239327 | 0.230487 | 0.242821 | 0.277457 | 0.257971 | 0.222678 | 0.216775 | 0.259602 | 0.349225 | 0.478764 | 0.934575 | 1.075432 | 1.147414 | 0.912944 | 0.536043 | 0.448313 | 0.428460 | 0.348205 |
| left central | 0.319507 | 0.335482 | 0.600173 | 0.592295 | 0.594178 | 0.792158 | 0.432472 | 0.301600 | 0.273370 | 0.224988 | 0.222751 | 0.351656 | 1.131192 | 3.876953 | 24.194188 | 41.289116 | 77.275148 | 8.885235 | 1.258130 | 0.627702 | 0.770891 | 0.404473 | 0.291282 | 0.238915 | 0.278456 | 0.284861 | 0.256750 | 0.234180 | 0.303764 | 0.636779 | 3.505679 | 8.699433 | 5.003015 | 1.481153 | 0.580015 | 0.275373 | 0.214668 | 0.233704 | 0.288728 | 0.268587 | 0.235326 | 0.219959 | 0.225874 | 0.269224 | 0.250262 | 0.238893 | 0.228340 | 0.253345 | 0.270404 | 0.243795 | 0.239523 | 0.265534 | 0.286401 | 0.270938 | 0.293233 | 0.387308 | 0.955415 | 1.169655 | 1.227239 | 1.792432 | 4.184264 | 2.280682 | 1.107351 | 0.596817 | 0.684113 | 0.729341 | 0.726419 | 0.431054 | 0.263322 | 0.219337 | 0.218303 | 0.367909 | 0.801928 | 1.698649 | 3.282604 | 3.400126 | 1.715000 | 0.756491 | 0.330312 | 0.214634 | 0.291258 | 0.650042 | 1.299905 | 1.345208 | 0.814207 | 0.558878 | 0.465208 | 0.483416 | 0.349433 | 0.233469 | 0.215962 | 0.220048 | 0.262861 | 0.310659 | 0.293312 | 0.301388 | 0.325071 | 0.348184 | 0.351072 | 0.343211 | 0.378823 | 0.559066 | 0.953792 | 1.231703 | 1.641900 | 1.664125 | 0.806024 | 0.316421 | 0.215117 | 0.261689 | 0.325249 | 0.392038 | 0.484265 | 0.485559 | 0.443848 | 0.359936 | 0.335824 | 0.359703 | 0.438624 | 0.502773 | 0.576905 | 0.520497 | 0.464002 | 0.416750 | 0.336016 | 0.256780 | 0.220251 | 0.214775 | 0.215517 | 0.216358 | 0.218816 | 0.214659 | 0.223680 | 0.242880 | 0.226257 | 0.215090 | 0.236480 | 0.280197 | 0.332582 | 0.457797 | 0.622695 | 0.779627 | 1.107761 | 1.738142 | 3.383566 | 6.361668 | 6.154765 | 5.422302 | 2.415447 | 0.830179 | 0.381957 | 0.258918 | 0.243453 | 0.262412 | 0.247191 | 0.269242 | 0.359414 | 0.486794 | 0.475450 | 0.329807 | 0.290298 | 0.341022 | 0.426768 | 0.438566 | 0.484536 | 0.611400 | 0.859367 | 0.932723 | 0.791477 | 0.603430 | 0.496527 | 0.418665 | 0.366155 | 0.343104 | 0.325291 | 0.285987 | 0.252916 | 0.234984 | 0.220345 | 0.214893 | 0.221344 | 0.244790 | 0.276384 | 0.288373 | 0.303605 | 0.291170 | 0.288330 | 0.275336 | 0.247106 | 0.226379 | 0.225376 | 0.217574 | 0.215316 | 0.214848 | 0.225461 | 0.276059 | 0.383435 | 0.486155 | 0.659129 | 0.828409 | 1.027755 | 0.833166 | 0.651512 | 0.543819 | 0.462338 | 0.480405 | 0.424426 | 0.418422 | 0.369643 | 0.335191 | 0.354290 | 0.393833 | 0.404024 | 0.595448 | 0.859194 | 1.703205 | 4.014438 | 6.833346 | 7.167208 | 5.286507 | 3.649688 | 2.687106 | 1.493646 | 0.808651 | 0.591831 | 0.604884 | 0.680571 | 0.782155 | 0.805426 | 0.895902 | 1.111876 | 1.359709 | 1.447766 | 1.479741 | 1.253083 | 0.903386 | 0.710664 | 0.567721 | 0.458469 | 0.389731 | 0.328882 | 0.277848 | 0.264117 | 0.265030 | 0.263565 | 0.241537 | 0.220983 | 0.214657 | 0.215184 | 0.215508 | 0.216247 | 0.216778 | 0.214675 | 0.225473 | 0.274594 | 0.312744 | 0.305739 | 0.309978 | 0.312501 | 0.290953 | 0.257157 | 0.224083 | 0.215885 | 0.214634 | 0.215103 | 0.224645 | 0.235186 | 0.248877 | 0.232399 | 0.222432 | 0.219512 | 0.227158 | 0.239956 | 0.284283 | 0.334243 | 0.467534 | 0.515069 | 0.562602 | 0.619385 | 0.585883 |
| right central | 0.215889 | 0.217919 | 0.237183 | 0.249833 | 0.266361 | 0.241318 | 0.274902 | 0.424023 | 0.598792 | 0.606189 | 0.495993 | 0.779949 | 1.709943 | 3.244650 | 2.721369 | 1.360679 | 0.869405 | 0.642837 | 0.309825 | 0.218240 | 0.215038 | 0.214680 | 0.219974 | 0.233759 | 0.248855 | 0.285258 | 0.224310 | 0.248720 | 0.236131 | 0.225761 | 0.234602 | 0.215027 | 0.215659 | 0.219953 | 0.217169 | 0.215252 | 0.254069 | 0.515046 | 0.950725 | 7.568813 | 13.250295 | 18.225305 | 14.492952 | 3.314690 | 1.740137 | 1.248451 | 0.711211 | 0.576226 | 0.471679 | 0.369362 | 0.356709 | 0.373246 | 0.304133 | 0.351109 | 0.451823 | 1.450616 | 4.198254 | 5.956790 | 4.441438 | 1.569903 | 0.426518 | 0.240634 | 0.220664 | 0.245145 | 0.248631 | 0.269572 | 0.275502 | 0.319346 | 0.458327 | 1.087910 | 6.224856 | 25.838996 | 64.633395 | 76.160899 | 113.303976 | 105.771496 | 53.444595 | 27.584596 | 20.596578 | 12.714182 | 9.362729 | 2.757099 | 1.293146 | 1.143137 | 1.180724 | 1.121555 | 1.084589 | 1.868235 | 2.895468 | 3.471363 | 3.347043 | 5.326876 | 6.110481 | 6.797816 | 3.867553 | 2.855137 | 2.921665 | 2.914790 | 1.714420 | 1.097302 | 0.652491 | 0.710763 | 0.680554 | 0.722962 | 0.558875 | 0.485887 | 0.459899 | 0.415640 | 0.412685 | 0.533125 | 0.604395 | 0.768310 | 0.891912 | 1.142450 | 1.518968 | 2.030168 | 2.423517 | 4.026339 | 6.544027 | 7.167248 | 4.732492 | 2.887572 | 1.876336 | 1.014064 | 0.717699 | 0.548634 | 0.519291 | 0.572958 | 0.505353 | 0.460394 | 0.469067 | 0.410360 | 0.424408 | 0.454481 | 0.316374 | 0.308120 | 0.249929 | 0.244613 | 0.267194 | 0.286515 | 0.263660 | 0.282715 | 0.300635 | 0.406430 | 0.476728 | 0.392596 | 0.284713 | 0.272701 | 0.274546 | 0.273081 | 0.277886 | 0.288325 | 0.325627 | 0.373072 | 0.359278 | 0.398915 | 0.426390 | 0.637760 | 0.695582 | 0.778820 | 1.145411 | 1.916004 | 1.428188 | 0.813454 | 0.370927 | 0.262703 | 0.237644 | 0.234122 | 0.230875 | 0.263691 | 0.358783 | 0.525500 | 0.543797 | 0.458110 | 0.396332 | 0.404755 | 0.445327 | 0.408346 | 0.337742 | 0.396480 | 0.503643 | 0.575488 | 0.502818 | 0.379342 | 0.352525 | 0.376340 | 0.377079 | 0.360540 | 0.370833 | 0.379469 | 0.401270 | 0.507056 | 0.590424 | 0.657315 | 0.796689 | 0.719777 | 0.701044 | 0.794472 | 0.641135 | 0.576298 | 0.492599 | 0.492596 | 0.622985 | 0.808311 | 0.694382 | 0.680874 | 0.490769 | 0.430031 | 0.320196 | 0.268904 | 0.261088 | 0.292892 | 0.331846 | 0.439850 | 0.489127 | 0.554589 | 0.516666 | 0.479005 | 0.373793 | 0.298001 | 0.244937 | 0.217456 | 0.214689 | 0.216238 | 0.224579 | 0.225836 | 0.215553 | 0.215427 | 0.242196 | 0.268202 | 0.271807 | 0.287502 | 0.316079 | 0.296594 | 0.295673 | 0.284424 | 0.286029 | 0.295685 | 0.276979 | 0.249531 | 0.241043 | 0.238733 | 0.236857 | 0.232769 | 0.226235 | 0.233344 | 0.257737 | 0.300260 | 0.328272 | 0.375559 | 0.493435 | 0.756338 | 1.034918 | 1.035551 | 1.107921 | 0.972454 | 0.854419 | 0.568928 | 0.386862 | 0.307683 | 0.278852 | 0.251310 | 0.238033 | 0.219893 | 0.219732 | 0.220356 | 0.235264 | 0.244921 | 0.231822 | 0.225533 | 0.217697 | 0.214715 | 0.218105 | 0.253344 | 0.330692 | 0.355249 | 0.299896 | 0.277048 | 0.254888 | 0.244177 |
| left posterior | 25.211720 | 13.938797 | 6.859280 | 9.391477 | 5.727229 | 2.794293 | 1.327867 | 0.330611 | 0.232555 | 0.226209 | 0.214636 | 0.222288 | 0.237267 | 0.229864 | 0.263294 | 1.198874 | 2.109179 | 3.703065 | 7.873684 | 7.027637 | 22.128852 | 3.765717 | 0.275628 | 0.246860 | 0.356883 | 0.712710 | 0.660648 | 0.466139 | 0.428523 | 0.290131 | 0.214974 | 0.252662 | 0.346123 | 0.328345 | 0.255242 | 0.256105 | 0.277005 | 0.264243 | 0.281483 | 0.278958 | 0.308026 | 0.410840 | 0.594571 | 0.751506 | 0.528812 | 0.307014 | 0.237045 | 0.214664 | 0.235998 | 0.267680 | 0.270068 | 0.264377 | 0.262771 | 0.256178 | 0.233737 | 0.244428 | 0.291841 | 0.360811 | 0.362385 | 0.473866 | 0.685433 | 0.954487 | 1.062818 | 0.953981 | 1.193212 | 2.679395 | 5.124899 | 7.104403 | 15.373304 | 39.411639 | 165.327808 | 629.124025 | 715.281006 | 735.730622 | 1260.638469 | 787.916483 | 727.118393 | 518.188945 | 494.824780 | 672.051960 | 849.504674 | 524.355076 | 355.529786 | 252.372211 | 80.910433 | 21.065046 | 7.687678 | 6.495285 | 4.721406 | 6.619789 | 9.682095 | 14.716278 | 8.462313 | 5.353972 | 1.639549 | 0.960335 | 0.576963 | 0.360109 | 0.289564 | 0.273185 | 0.263231 | 0.342870 | 0.537097 | 0.985760 | 1.881901 | 3.044699 | 3.771793 | 5.415488 | 4.588530 | 3.812506 | 3.465807 | 3.881520 | 3.316057 | 3.466946 | 3.056842 | 2.779389 | 3.547648 | 6.070094 | 7.411948 | 10.334872 | 16.568539 | 23.815079 | 19.269199 | 16.344220 | 10.863717 | 9.959276 | 8.803520 | 6.210724 | 5.101429 | 5.143010 | 5.387952 | 6.113630 | 5.587491 | 4.807419 | 3.137536 | 3.067830 | 4.962979 | 5.012639 | 4.260118 | 3.545093 | 1.723547 | 1.182000 | 0.708865 | 0.465667 | 0.369507 | 0.322613 | 0.268873 | 0.259867 | 0.231147 | 0.219100 | 0.215093 | 0.218990 | 0.247770 | 0.240882 | 0.235934 | 0.217336 | 0.218997 | 0.233212 | 0.265370 | 0.332044 | 0.335665 | 0.379188 | 0.318945 | 0.281216 | 0.267614 | 0.248713 | 0.228488 | 0.227771 | 0.227545 | 0.259987 | 0.303134 | 0.375150 | 0.455836 | 0.602785 | 0.747699 | 0.865833 | 0.905369 | 0.879252 | 0.779166 | 0.810413 | 0.693511 | 0.678140 | 0.676316 | 0.714190 | 0.721299 | 0.718583 | 0.703952 | 0.786182 | 0.755353 | 0.687395 | 0.579927 | 0.527460 | 0.545547 | 0.655879 | 0.743704 | 0.983686 | 1.586168 | 2.481993 | 3.196036 | 4.141130 | 3.567347 | 3.355981 | 2.458793 | 1.459309 | 0.780342 | 0.650121 | 0.512251 | 0.416976 | 0.341895 | 0.354525 | 0.383303 | 0.510643 | 0.545418 | 0.806846 | 1.755052 | 3.727862 | 7.233095 | 15.863686 | 41.287873 | 101.469661 | 101.486979 | 44.847254 | 19.084275 | 5.439449 | 2.464825 | 0.865283 | 0.650609 | 0.589337 | 0.551499 | 0.587113 | 0.631442 | 0.667108 | 0.729186 | 0.670577 | 0.712324 | 0.884821 | 0.944060 | 1.208115 | 1.395367 | 1.683400 | 1.909000 | 1.928946 | 1.736782 | 1.410317 | 1.090849 | 0.731620 | 0.567697 | 0.421823 | 0.322625 | 0.280465 | 0.257707 | 0.237131 | 0.233648 | 0.225979 | 0.229554 | 0.251588 | 0.262186 | 0.273166 | 0.292407 | 0.334433 | 0.370870 | 0.371962 | 0.351660 | 0.326589 | 0.307710 | 0.285978 | 0.242812 | 0.230010 | 0.224531 | 0.217214 | 0.214851 | 0.214761 | 0.214940 | 0.214698 | 0.215960 | 0.219383 | 0.236783 | 0.251712 | 0.266322 | 0.326526 |
| right posterior | 0.216012 | 0.217490 | 0.215118 | 0.215376 | 0.215157 | 0.215725 | 0.225120 | 0.217054 | 0.215626 | 0.216075 | 0.230204 | 0.222530 | 0.214842 | 0.216196 | 0.214998 | 0.287907 | 0.668988 | 1.487691 | 2.454230 | 3.522202 | 3.138317 | 0.838097 | 0.244900 | 0.247017 | 0.360552 | 0.393390 | 0.304194 | 0.224933 | 0.256027 | 0.394416 | 1.226727 | 3.204340 | 2.128136 | 0.803440 | 0.272591 | 0.214892 | 0.235010 | 0.287751 | 0.326352 | 0.275217 | 0.224239 | 0.219426 | 0.252784 | 0.320691 | 0.704404 | 1.533180 | 2.370118 | 3.635419 | 5.115838 | 4.704093 | 4.465873 | 1.923092 | 0.818749 | 0.582744 | 0.302371 | 0.230506 | 0.215616 | 0.218378 | 0.251815 | 0.316271 | 0.373659 | 0.342266 | 0.335326 | 0.314035 | 0.292339 | 0.289052 | 0.306566 | 0.415522 | 1.031533 | 3.551782 | 17.557350 | 71.030418 | 338.012179 | 1364.491070 | 2048.813880 | 1645.148229 | 780.976642 | 344.161240 | 181.173989 | 84.988180 | 51.221674 | 27.245108 | 7.349858 | 2.463511 | 1.138275 | 1.197820 | 1.012279 | 1.188474 | 2.187197 | 4.385738 | 7.493107 | 9.500242 | 5.842106 | 4.836658 | 2.778388 | 1.227984 | 0.773275 | 0.618775 | 0.549727 | 0.490925 | 0.515534 | 0.701670 | 0.833340 | 1.087853 | 0.918132 | 1.032578 | 1.169541 | 1.253853 | 1.001939 | 1.062875 | 0.969685 | 1.180700 | 1.365714 | 1.584257 | 1.697628 | 2.079366 | 2.522233 | 3.708320 | 5.845389 | 10.924131 | 19.367118 | 35.999111 | 71.662301 | 121.065845 | 115.891867 | 81.781579 | 43.981590 | 17.841142 | 7.489769 | 3.172024 | 2.000899 | 1.495385 | 1.404327 | 1.355247 | 1.598529 | 1.904379 | 2.362341 | 2.722571 | 2.944986 | 2.943382 | 2.937034 | 2.936616 | 2.534348 | 2.137218 | 1.523944 | 1.442838 | 1.209778 | 1.217162 | 1.523295 | 1.546484 | 1.668050 | 2.287327 | 2.266185 | 1.978824 | 1.717392 | 1.313928 | 1.355900 | 1.543553 | 1.852575 | 2.394945 | 3.465265 | 4.493937 | 5.177860 | 4.775221 | 3.275519 | 1.943698 | 1.368242 | 1.134789 | 1.029131 | 1.174727 | 1.364629 | 1.588960 | 1.904612 | 1.809373 | 1.737668 | 2.182380 | 2.670034 | 3.748126 | 4.880997 | 6.383930 | 7.043469 | 6.879930 | 6.993467 | 5.114815 | 4.238988 | 5.450651 | 8.500932 | 11.092199 | 13.002848 | 8.540363 | 7.164924 | 6.383420 | 4.992894 | 3.524501 | 3.207369 | 2.973924 | 4.050911 | 4.844420 | 4.724420 | 3.881346 | 3.022812 | 3.039455 | 3.631105 | 3.372186 | 3.884529 | 3.989332 | 3.717688 | 4.515773 | 3.915092 | 3.107840 | 2.544624 | 1.943250 | 1.792658 | 2.107617 | 2.427262 | 3.119696 | 4.418636 | 5.937833 | 7.261617 | 7.212734 | 4.734320 | 3.515236 | 2.804576 | 2.606177 | 1.988731 | 1.383592 | 1.315945 | 1.543489 | 2.009914 | 2.233956 | 2.154183 | 2.378989 | 2.928804 | 3.418987 | 4.668021 | 6.101691 | 8.368810 | 12.484256 | 16.708836 | 23.711701 | 27.507043 | 31.956219 | 25.635044 | 25.358263 | 15.841906 | 10.312022 | 6.705590 | 4.835608 | 3.245813 | 3.111772 | 2.497473 | 2.404670 | 3.358431 | 5.260287 | 8.055230 | 12.612501 | 16.803899 | 20.145142 | 23.792245 | 25.874609 | 13.595942 | 8.968615 | 5.383672 | 3.021931 | 2.540639 | 1.598485 | 0.913147 | 0.841490 | 0.737011 | 0.607608 | 0.491881 | 0.349175 | 0.296032 | 0.262525 | 0.238649 | 0.230637 | 0.236045 | 0.234586 | 0.241302 | 0.257286 |
| all electrodes | 0.445093 | 0.454941 | 0.662132 | 0.973307 | 1.064153 | 1.024814 | 1.195497 | 2.091969 | 2.130054 | 1.160077 | 0.548315 | 0.388529 | 0.352282 | 0.326532 | 0.370416 | 0.621350 | 0.937591 | 0.883517 | 1.090541 | 2.499417 | 9.200026 | 17.401723 | 0.984298 | 0.263171 | 0.218487 | 0.270630 | 0.506717 | 0.950371 | 1.568082 | 0.510929 | 0.259298 | 0.227467 | 0.254327 | 0.257439 | 0.221249 | 0.215530 | 0.214710 | 0.214719 | 0.217999 | 0.215959 | 0.227538 | 0.334151 | 0.487204 | 0.836649 | 0.983921 | 2.341497 | 5.256516 | 6.995451 | 7.471269 | 3.937129 | 2.945156 | 1.391126 | 0.663686 | 0.495466 | 0.550576 | 0.466249 | 0.934046 | 0.724585 | 0.627523 | 0.579891 | 0.368562 | 0.223560 | 0.218702 | 0.216326 | 0.216658 | 0.216334 | 0.218505 | 0.257820 | 0.438701 | 1.405240 | 5.393450 | 25.564825 | 234.421819 | 1887.029389 | 5919.306847 | 5272.745857 | 6525.741166 | 3617.981963 | 2640.173193 | 982.012914 | 310.418659 | 58.316695 | 9.034660 | 1.364193 | 0.564884 | 0.413289 | 0.272079 | 0.251717 | 0.231870 | 0.236873 | 0.256568 | 0.248958 | 0.227346 | 0.218828 | 0.214691 | 0.219237 | 0.263825 | 0.274878 | 0.327572 | 0.425268 | 0.468896 | 0.452308 | 0.484573 | 0.376832 | 0.335848 | 0.242592 | 0.218526 | 0.214626 | 0.230991 | 0.328762 | 0.464852 | 0.376136 | 0.345646 | 0.426118 | 0.491177 | 0.503709 | 0.458923 | 0.401628 | 0.744724 | 0.878213 | 0.835813 | 0.961622 | 1.126582 | 1.174077 | 1.045258 | 0.660650 | 0.568136 | 0.378858 | 0.303187 | 0.278144 | 0.250227 | 0.257416 | 0.268659 | 0.261315 | 0.265547 | 0.240867 | 0.221946 | 0.220744 | 0.215715 | 0.214626 | 0.218228 | 0.214653 | 0.223905 | 0.239993 | 0.233333 | 0.215167 | 0.216600 | 0.215782 | 0.220399 | 0.258942 | 0.351197 | 0.423346 | 0.391513 | 0.433555 | 0.484189 | 0.588143 | 0.507901 | 0.404560 | 0.275942 | 0.225529 | 0.214727 | 0.214674 | 0.215886 | 0.217896 | 0.220460 | 0.272838 | 0.454530 | 0.648259 | 0.589380 | 0.476997 | 0.418642 | 0.332711 | 0.292372 | 0.245379 | 0.227322 | 0.217852 | 0.215653 | 0.239165 | 0.229623 | 0.236349 | 0.246585 | 0.251553 | 0.235094 | 0.216684 | 0.215191 | 0.214626 | 0.215148 | 0.223954 | 0.233545 | 0.232555 | 0.227006 | 0.246541 | 0.282714 | 0.258344 | 0.224133 | 0.216919 | 0.241269 | 0.262654 | 0.292339 | 0.311256 | 0.289609 | 0.273736 | 0.245062 | 0.241566 | 0.241838 | 0.237827 | 0.234257 | 0.245926 | 0.239197 | 0.221108 | 0.217441 | 0.217701 | 0.224284 | 0.247496 | 0.285766 | 0.354981 | 0.453199 | 0.513041 | 0.455808 | 0.460118 | 0.358008 | 0.289755 | 0.255947 | 0.288729 | 0.360526 | 0.423178 | 0.325773 | 0.299993 | 0.270667 | 0.284403 | 0.289653 | 0.286894 | 0.346349 | 0.599711 | 1.442079 | 3.629974 | 4.030143 | 4.531373 | 4.321519 | 2.280967 | 2.025842 | 1.533272 | 1.422743 | 1.961048 | 2.488257 | 2.842227 | 4.277921 | 2.936564 | 1.506568 | 0.701703 | 0.341358 | 0.246571 | 0.229062 | 0.217596 | 0.217942 | 0.229916 | 0.249601 | 0.250723 | 0.249966 | 0.236138 | 0.242150 | 0.251157 | 0.246764 | 0.230677 | 0.227656 | 0.224073 | 0.214980 | 0.215211 | 0.220927 | 0.226527 | 0.223613 | 0.230773 | 0.245053 | 0.251873 | 0.316770 | 0.372812 | 0.427915 | 0.539090 | 0.527783 | 0.521829 |

Searchlight, spatiotemporal cluster permutation test

|  | start time | stop time | peak time | peak channel | cluster p | peak Cohen's d | direction |
| --- | --- | --- | --- | --- | --- | --- | --- |
| #1 | 95 | 260 | 170 | PO7 | 0.0437 | 1.13891 | positive |
| #2 | 225 | 1155 | 425 | Oz | 0.0098 | 1.144394 | positive |

H) angry vs sad

  
|  | time window | peak latency | cluster *p* | peak Cohen's *d* |  | | | |  |
| **all electrodes** |  | | | |  | | | |  |
|  | | | | | | | | |  |

Time-resolved classification, cluster permutation tests

|  | **left hemisphere** | | | | **right hemisphere** | | | |  |
|  | time window | peak latency | cluster *p* | peak Cohen's *d* | time window | peak latency | cluster *p* | peak Cohen's *d* |  |
| **anterior** |  | | | |  | | | |  |
| **central** | 185 - 290 ms | 255 ms | 0.0179 | 0.9377 | 200 - 290 ms | 235 ms | 0.025 | 0.9198 |  |
 340 - 495 ms | 365 ms | 0.0061 | 0.8882 | 315 - 445 ms | 415 ms | 0.0079 | 0.9321 |  | | | | -200 - -115 ms | -115 ms | 0.0402 | -0.567 ||  | **posterior** |  | | | | 200 - 295 ms | 265 ms | 0.0107 | 1.1306 |
  | | | | 335 - 415 ms | 390 ms | 0.0414 | 0.7291 |

  

Time-resolved classification, Bayesian statistics

|  | -200 | -195 | -190 | -185 | -180 | -175 | -170 | -165 | -160 | -155 | -150 | -145 | -140 | -135 | -130 | -125 | -120 | -115 | -110 | -105 | -100 | -95 | -90 | -85 | -80 | -75 | -70 | -65 | -60 | -55 | -50 | -45 | -40 | -35 | -30 | -25 | -20 | -15 | -10 | -5 | 0 | 5 | 10 | 15 | 20 | 25 | 30 | 35 | 40 | 45 | 50 | 55 | 60 | 65 | 70 | 75 | 80 | 85 | 90 | 95 | 100 | 105 | 110 | 115 | 120 | 125 | 130 | 135 | 140 | 145 | 150 | 155 | 160 | 165 | 170 | 175 | 180 | 185 | 190 | 195 | 200 | 205 | 210 | 215 | 220 | 225 | 230 | 235 | 240 | 245 | 250 | 255 | 260 | 265 | 270 | 275 | 280 | 285 | 290 | 295 | 300 | 305 | 310 | 315 | 320 | 325 | 330 | 335 | 340 | 345 | 350 | 355 | 360 | 365 | 370 | 375 | 380 | 385 | 390 | 395 | 400 | 405 | 410 | 415 | 420 | 425 | 430 | 435 | 440 | 445 | 450 | 455 | 460 | 465 | 470 | 475 | 480 | 485 | 490 | 495 | 500 | 505 | 510 | 515 | 520 | 525 | 530 | 535 | 540 | 545 | 550 | 555 | 560 | 565 | 570 | 575 | 580 | 585 | 590 | 595 | 600 | 605 | 610 | 615 | 620 | 625 | 630 | 635 | 640 | 645 | 650 | 655 | 660 | 665 | 670 | 675 | 680 | 685 | 690 | 695 | 700 | 705 | 710 | 715 | 720 | 725 | 730 | 735 | 740 | 745 | 750 | 755 | 760 | 765 | 770 | 775 | 780 | 785 | 790 | 795 | 800 | 805 | 810 | 815 | 820 | 825 | 830 | 835 | 840 | 845 | 850 | 855 | 860 | 865 | 870 | 875 | 880 | 885 | 890 | 895 | 900 | 905 | 910 | 915 | 920 | 925 | 930 | 935 | 940 | 945 | 950 | 955 | 960 | 965 | 970 | 975 | 980 | 985 | 990 | 995 | 1000 | 1005 | 1010 | 1015 | 1020 | 1025 | 1030 | 1035 | 1040 | 1045 | 1050 | 1055 | 1060 | 1065 | 1070 | 1075 | 1080 | 1085 | 1090 | 1095 | 1100 | 1105 | 1110 | 1115 | 1120 | 1125 | 1130 | 1135 | 1140 | 1145 | 1150 | 1155 | 1160 | 1165 | 1170 | 1175 | 1180 | 1185 | 1190 | 1195 |
| --- | --- | --- | --- | --- | --- | --- | --- | --- | --- | --- | --- | --- | --- | --- | --- | --- | --- | --- | --- | --- | --- | --- | --- | --- | --- | --- | --- | --- | --- | --- | --- | --- | --- | --- | --- | --- | --- | --- | --- | --- | --- | --- | --- | --- | --- | --- | --- | --- | --- | --- | --- | --- | --- | --- | --- | --- | --- | --- | --- | --- | --- | --- | --- | --- | --- | --- | --- | --- | --- | --- | --- | --- | --- | --- | --- | --- | --- | --- | --- | --- | --- | --- | --- | --- | --- | --- | --- | --- | --- | --- | --- | --- | --- | --- | --- | --- | --- | --- | --- | --- | --- | --- | --- | --- | --- | --- | --- | --- | --- | --- | --- | --- | --- | --- | --- | --- | --- | --- | --- | --- | --- | --- | --- | --- | --- | --- | --- | --- | --- | --- | --- | --- | --- | --- | --- | --- | --- | --- | --- | --- | --- | --- | --- | --- | --- | --- | --- | --- | --- | --- | --- | --- | --- | --- | --- | --- | --- | --- | --- | --- | --- | --- | --- | --- | --- | --- | --- | --- | --- | --- | --- | --- | --- | --- | --- | --- | --- | --- | --- | --- | --- | --- | --- | --- | --- | --- | --- | --- | --- | --- | --- | --- | --- | --- | --- | --- | --- | --- | --- | --- | --- | --- | --- | --- | --- | --- | --- | --- | --- | --- | --- | --- | --- | --- | --- | --- | --- | --- | --- | --- | --- | --- | --- | --- | --- | --- | --- | --- | --- | --- | --- | --- | --- | --- | --- | --- | --- | --- | --- | --- | --- | --- | --- | --- | --- | --- | --- | --- | --- | --- | --- | --- | --- | --- | --- | --- | --- | --- | --- | --- | --- | --- | --- | --- | --- | --- | --- | --- | --- | --- | --- | --- | --- | --- | --- | --- | --- | --- | --- | --- |
| left anterior | 0.514354 | 1.548651 | 3.641402 | 2.943323 | 5.547288 | 3.683580 | 1.248408 | 0.571438 | 0.326779 | 0.214642 | 0.391438 | 2.074665 | 19.900674 | 64.158224 | 252.807578 | 869.067739 | 38.789110 | 3.073882 | 0.634800 | 0.240698 | 0.229289 | 0.597970 | 2.261057 | 2.193280 | 2.330149 | 1.902783 | 2.158478 | 1.173551 | 0.417795 | 0.267812 | 0.269931 | 0.246703 | 0.214841 | 0.238887 | 0.248906 | 0.307534 | 0.317727 | 0.416674 | 0.486906 | 0.450223 | 0.307534 | 0.308139 | 0.267737 | 0.279855 | 0.249682 | 0.231769 | 0.219000 | 0.243578 | 0.258917 | 0.245433 | 0.215877 | 0.216414 | 0.218089 | 0.224823 | 0.233189 | 0.231262 | 0.231658 | 0.314866 | 0.394390 | 0.495602 | 0.659142 | 0.642066 | 0.609215 | 0.643221 | 0.472737 | 0.464764 | 0.496025 | 0.922257 | 0.836617 | 0.743319 | 0.789110 | 1.782704 | 2.750190 | 3.887647 | 1.855323 | 2.047225 | 2.803127 | 2.748582 | 0.976893 | 0.753236 | 0.867892 | 1.153199 | 1.600336 | 0.987975 | 0.590774 | 0.571728 | 0.460956 | 0.303007 | 0.246560 | 0.214636 | 0.215217 | 0.215647 | 0.214626 | 0.219560 | 0.218213 | 0.218455 | 0.214698 | 0.224747 | 0.251615 | 0.401472 | 1.120184 | 1.948224 | 2.774975 | 3.241717 | 2.563755 | 1.212901 | 0.660083 | 0.428667 | 0.362062 | 0.284953 | 0.258814 | 0.220609 | 0.217611 | 0.260303 | 0.281698 | 0.312402 | 0.372473 | 0.372692 | 0.307650 | 0.248560 | 0.234425 | 0.258235 | 0.315643 | 0.346919 | 0.372497 | 0.463909 | 0.588327 | 0.655765 | 0.612160 | 0.381141 | 0.253287 | 0.215979 | 0.229180 | 0.276581 | 0.393571 | 0.759039 | 1.009106 | 1.139334 | 1.589655 | 1.282013 | 0.740076 | 0.392217 | 0.273785 | 0.265828 | 0.318390 | 0.297397 | 0.332291 | 0.305354 | 0.289411 | 0.234824 | 0.214906 | 0.215680 | 0.217034 | 0.214752 | 0.214975 | 0.218453 | 0.230880 | 0.248946 | 0.219891 | 0.214638 | 0.217432 | 0.215634 | 0.225775 | 0.264128 | 0.302905 | 0.306360 | 0.290291 | 0.277262 | 0.257355 | 0.248216 | 0.238862 | 0.239767 | 0.253731 | 0.312885 | 0.416392 | 0.532026 | 0.510727 | 0.465807 | 0.434052 | 0.428997 | 0.389097 | 0.391951 | 0.398084 | 0.462279 | 0.344819 | 0.258608 | 0.217307 | 0.222736 | 0.237134 | 0.229436 | 0.240434 | 0.232804 | 0.225428 | 0.215470 | 0.214640 | 0.214983 | 0.220124 | 0.219089 | 0.216018 | 0.219673 | 0.224594 | 0.214933 | 0.220872 | 0.256750 | 0.335031 | 0.506961 | 0.805345 | 1.117370 | 1.341604 | 2.029984 | 1.773770 | 0.758234 | 0.606682 | 0.354865 | 0.263490 | 0.218291 | 0.215025 | 0.237590 | 0.227029 | 0.249238 | 0.225586 | 0.217742 | 0.218169 | 0.222835 | 0.228389 | 0.274267 | 0.321199 | 0.381717 | 0.387709 | 0.348033 | 0.290080 | 0.240973 | 0.215687 | 0.217342 | 0.224980 | 0.228522 | 0.232992 | 0.218175 | 0.214747 | 0.220643 | 0.231698 | 0.256281 | 0.282158 | 0.386178 | 0.386705 | 0.419627 | 0.343310 | 0.300733 | 0.289887 | 0.281732 | 0.249570 | 0.249862 | 0.248797 | 0.254239 | 0.245424 | 0.225369 | 0.216618 | 0.215180 | 0.219415 | 0.221042 | 0.217314 | 0.214995 | 0.225631 | 0.267861 | 0.330937 | 0.381529 | 0.402869 | 0.400637 | 0.320862 | 0.257839 | 0.215190 | 0.226442 | 0.285656 | 0.295553 | 0.350348 | 0.337616 | 0.303335 | 0.261681 | 0.228590 | 0.217573 |
| right anterior | 0.215176 | 0.216653 | 0.228413 | 0.241736 | 0.321563 | 0.513678 | 0.358881 | 0.348656 | 0.279443 | 0.214777 | 0.228182 | 0.260275 | 0.234465 | 0.268283 | 0.707938 | 4.465869 | 128.662150 | 35.470948 | 4.278459 | 0.791003 | 0.216408 | 0.265179 | 0.602213 | 1.009462 | 0.679926 | 0.473057 | 0.345551 | 0.310516 | 0.420684 | 0.312868 | 0.265015 | 0.354946 | 0.287651 | 0.288722 | 0.224691 | 0.214691 | 0.216520 | 0.220311 | 0.215398 | 0.230319 | 0.304329 | 0.288341 | 0.254425 | 0.266520 | 0.214711 | 0.223166 | 0.315465 | 0.424430 | 0.314626 | 0.238911 | 0.221578 | 0.218771 | 0.254412 | 0.348727 | 0.388867 | 0.318946 | 0.290437 | 0.270849 | 0.283725 | 0.299304 | 0.301364 | 0.278319 | 0.280624 | 0.274156 | 0.368815 | 0.395546 | 0.361312 | 0.336081 | 0.335480 | 0.274731 | 0.332602 | 0.297919 | 0.290441 | 0.318562 | 0.371281 | 0.543851 | 1.222686 | 1.180981 | 1.803758 | 2.191733 | 1.851580 | 1.205694 | 0.648502 | 0.358601 | 0.268949 | 0.216117 | 0.235279 | 0.352233 | 0.367838 | 0.328666 | 0.298009 | 0.295078 | 0.236224 | 0.216603 | 0.219838 | 0.219010 | 0.214696 | 0.214971 | 0.222400 | 0.217873 | 0.215759 | 0.218353 | 0.220319 | 0.214688 | 0.222891 | 0.250390 | 0.258632 | 0.248774 | 0.258542 | 0.297694 | 0.327423 | 0.426651 | 0.732118 | 1.414648 | 2.814701 | 4.893546 | 5.017912 | 4.377918 | 2.984235 | 1.346566 | 1.848757 | 3.472950 | 5.876915 | 3.359450 | 1.661530 | 1.316320 | 1.603375 | 0.989574 | 0.699890 | 0.351348 | 0.289880 | 0.306053 | 0.307370 | 0.244474 | 0.216883 | 0.236052 | 0.263070 | 0.272162 | 0.302010 | 0.353745 | 0.380584 | 0.424553 | 0.476371 | 0.672262 | 1.410801 | 3.268826 | 9.210612 | 13.739707 | 7.602029 | 1.199419 | 0.458368 | 0.323423 | 0.435171 | 0.631982 | 0.850115 | 0.623891 | 0.738219 | 0.605673 | 0.527463 | 0.366564 | 0.294457 | 0.248834 | 0.234418 | 0.251048 | 0.326719 | 0.425031 | 0.417382 | 0.339074 | 0.318786 | 0.367587 | 0.346542 | 0.403765 | 0.374817 | 0.360763 | 0.469549 | 0.502475 | 0.433935 | 0.427450 | 0.353283 | 0.384039 | 0.399341 | 0.296877 | 0.261955 | 0.249597 | 0.243241 | 0.243307 | 0.216102 | 0.215504 | 0.218539 | 0.216838 | 0.215908 | 0.215186 | 0.243325 | 0.253015 | 0.316877 | 0.297485 | 0.305595 | 0.312187 | 0.263956 | 0.219760 | 0.226203 | 0.369231 | 0.412583 | 0.495437 | 0.422611 | 0.387635 | 0.420282 | 0.357796 | 0.347862 | 0.444769 | 0.441819 | 0.307512 | 0.247424 | 0.215010 | 0.230884 | 0.260013 | 0.318025 | 0.344289 | 0.266295 | 0.247404 | 0.231107 | 0.220005 | 0.214626 | 0.215341 | 0.215542 | 0.220910 | 0.230569 | 0.236134 | 0.228296 | 0.258643 | 0.266679 | 0.235486 | 0.214754 | 0.227222 | 0.230803 | 0.230008 | 0.230515 | 0.234789 | 0.238432 | 0.217754 | 0.216275 | 0.226129 | 0.271381 | 0.329108 | 0.342050 | 0.386563 | 0.400356 | 0.414332 | 0.386703 | 0.342061 | 0.289718 | 0.252024 | 0.244550 | 0.226240 | 0.215622 | 0.215419 | 0.216010 | 0.214759 | 0.216975 | 0.215214 | 0.221250 | 0.236304 | 0.332983 | 0.354250 | 0.343503 | 0.367864 | 0.428279 | 0.360797 | 0.327516 | 0.234586 | 0.214714 | 0.235850 | 0.293408 | 0.419365 | 0.961497 | 3.260805 | 10.570611 | 13.903520 | 8.737627 | 10.548578 |
| left central | 0.327054 | 0.306972 | 0.311596 | 0.281534 | 0.236713 | 0.241929 | 0.268787 | 0.378254 | 0.515527 | 0.410606 | 0.368719 | 0.460207 | 0.223990 | 0.214653 | 0.232867 | 0.260320 | 0.317330 | 0.420727 | 0.526123 | 0.221208 | 0.217786 | 0.215492 | 0.296034 | 0.834952 | 1.283951 | 1.565391 | 0.486506 | 0.232182 | 0.215642 | 0.264803 | 0.555323 | 1.073390 | 0.510451 | 0.333551 | 0.233430 | 0.307891 | 0.333074 | 0.294545 | 0.342517 | 0.234354 | 0.229339 | 0.214831 | 0.231983 | 0.272001 | 0.263673 | 0.247361 | 0.225859 | 0.233024 | 0.265716 | 0.279846 | 0.243222 | 0.215461 | 0.214835 | 0.218734 | 0.216380 | 0.215343 | 0.214986 | 0.231774 | 0.295987 | 0.307748 | 0.295953 | 0.239412 | 0.214674 | 0.215205 | 0.217744 | 0.218729 | 0.214922 | 0.223649 | 0.245226 | 0.371274 | 0.300394 | 0.274316 | 0.239458 | 0.217819 | 0.228870 | 0.317350 | 0.989617 | 3.396611 | 12.696556 | 15.939520 | 9.157921 | 5.261803 | 3.907085 | 5.469060 | 5.251754 | 5.765414 | 5.064508 | 6.864800 | 9.546046 | 21.596973 | 52.818073 | 216.814918 | 87.683679 | 61.302200 | 55.920262 | 23.682639 | 8.980301 | 5.238033 | 2.121257 | 1.285438 | 0.774112 | 0.408189 | 0.291606 | 0.290396 | 0.288619 | 0.277567 | 0.409403 | 1.182782 | 6.756257 | 58.983710 | 142.744743 | 315.341730 | 204.644817 | 126.236544 | 69.068867 | 39.494109 | 10.013555 | 5.780588 | 2.855277 | 3.968633 | 4.677244 | 3.826205 | 4.199329 | 3.809391 | 3.248652 | 2.844922 | 3.658505 | 4.158862 | 5.245631 | 4.949813 | 5.727796 | 12.621091 | 18.294205 | 21.498670 | 11.962788 | 6.133726 | 3.506047 | 3.739542 | 2.418062 | 1.633905 | 0.562133 | 0.422421 | 0.311614 | 0.240558 | 0.216046 | 0.226037 | 0.229240 | 0.215220 | 0.263064 | 0.513544 | 1.722984 | 3.537923 | 10.670681 | 17.202258 | 11.300282 | 9.614898 | 4.303068 | 1.892321 | 0.851334 | 0.439611 | 0.389128 | 0.303663 | 0.261234 | 0.277396 | 0.268374 | 0.275284 | 0.292655 | 0.311141 | 0.336552 | 0.367836 | 0.387775 | 0.420397 | 0.565170 | 0.556384 | 0.488691 | 0.426281 | 0.363638 | 0.308945 | 0.303761 | 0.258888 | 0.235962 | 0.220607 | 0.218615 | 0.276776 | 0.328363 | 0.407505 | 0.391043 | 0.398956 | 0.376700 | 0.264181 | 0.219984 | 0.218815 | 0.227452 | 0.255015 | 0.281047 | 0.277561 | 0.300754 | 0.292944 | 0.293351 | 0.250469 | 0.255981 | 0.288990 | 0.366898 | 0.386178 | 0.373990 | 0.299484 | 0.272196 | 0.219997 | 0.224939 | 0.252846 | 0.286369 | 0.279268 | 0.309490 | 0.355893 | 0.379647 | 0.360217 | 0.462064 | 0.614801 | 0.803925 | 0.690169 | 0.704215 | 0.729909 | 0.547211 | 0.443017 | 0.343915 | 0.359669 | 0.350935 | 0.344058 | 0.321431 | 0.315894 | 0.311603 | 0.314516 | 0.293969 | 0.308875 | 0.305184 | 0.300041 | 0.315214 | 0.337681 | 0.365111 | 0.361208 | 0.343011 | 0.324004 | 0.312124 | 0.272370 | 0.230045 | 0.222818 | 0.217561 | 0.216014 | 0.234948 | 0.252466 | 0.262994 | 0.249385 | 0.277906 | 0.298792 | 0.301984 | 0.340997 | 0.404746 | 0.525340 | 0.772296 | 0.900839 | 1.099688 | 1.015291 | 0.694253 | 0.481959 | 0.381955 | 0.332431 | 0.300416 | 0.243982 | 0.224822 | 0.226411 | 0.234047 | 0.242526 | 0.260849 | 0.308615 | 0.366987 | 0.573038 | 0.988150 | 1.280335 | 1.399034 | 1.257671 |
| right central | 2.260404 | 4.017851 | 2.828956 | 5.315430 | 12.608693 | 6.271697 | 7.134813 | 2.915838 | 2.301458 | 4.830671 | 6.011451 | 10.268513 | 30.113436 | 47.230339 | 32.226276 | 10.856677 | 7.121609 | 4.573623 | 1.125510 | 0.390752 | 0.233508 | 0.218632 | 0.217676 | 0.296538 | 0.610184 | 0.356147 | 0.243934 | 0.233546 | 0.388188 | 0.310733 | 0.333192 | 0.265025 | 0.223782 | 0.221213 | 0.237317 | 0.227416 | 0.224640 | 0.226525 | 0.214661 | 0.216474 | 0.226727 | 0.224052 | 0.214639 | 0.214770 | 0.216082 | 0.215461 | 0.216933 | 0.214812 | 0.215843 | 0.223887 | 0.220460 | 0.224401 | 0.215300 | 0.217870 | 0.219546 | 0.227267 | 0.288239 | 0.386725 | 0.569536 | 0.617244 | 0.388561 | 0.252748 | 0.218111 | 0.218898 | 0.262607 | 0.443160 | 0.606748 | 0.569240 | 0.298347 | 0.219649 | 0.217876 | 0.250075 | 0.273589 | 0.263923 | 0.264873 | 0.218273 | 0.230813 | 0.279717 | 0.543066 | 1.155672 | 1.936003 | 4.708702 | 9.018440 | 16.256369 | 47.970854 | 109.694270 | 215.637441 | 178.301131 | 92.472201 | 45.492974 | 23.018143 | 15.542684 | 14.569377 | 10.997587 | 7.876610 | 4.178425 | 2.219628 | 1.707676 | 1.335384 | 0.926618 | 0.740842 | 0.646488 | 1.089425 | 1.730278 | 2.065214 | 2.142991 | 2.647114 | 3.653084 | 4.879288 | 5.794344 | 3.825657 | 5.794743 | 7.818706 | 7.661552 | 12.537168 | 14.963283 | 19.310646 | 104.179402 | 96.660441 | 85.993309 | 85.820674 | 198.680262 | 309.647151 | 203.945521 | 63.841255 | 24.102075 | 15.270437 | 12.632669 | 4.410485 | 1.415167 | 0.955755 | 0.824333 | 0.711489 | 0.639984 | 0.462639 | 0.426269 | 0.499841 | 0.422872 | 0.279432 | 0.228362 | 0.215129 | 0.235039 | 0.283173 | 0.336544 | 0.299542 | 0.297698 | 0.270359 | 0.263933 | 0.240292 | 0.220340 | 0.214916 | 0.218357 | 0.252120 | 0.293438 | 0.367617 | 0.351119 | 0.330245 | 0.287870 | 0.258257 | 0.222257 | 0.216974 | 0.264880 | 0.403753 | 0.518168 | 0.896689 | 0.961785 | 0.696268 | 0.391800 | 0.262556 | 0.215559 | 0.222717 | 0.297677 | 0.411845 | 0.517652 | 0.495654 | 0.423444 | 0.367981 | 0.344886 | 0.280194 | 0.247593 | 0.230617 | 0.230033 | 0.251307 | 0.274380 | 0.288970 | 0.315836 | 0.336591 | 0.297050 | 0.245443 | 0.218540 | 0.217334 | 0.222267 | 0.247404 | 0.309070 | 0.424355 | 0.491377 | 0.408898 | 0.354680 | 0.252337 | 0.216039 | 0.228718 | 0.227305 | 0.218720 | 0.215606 | 0.220877 | 0.220347 | 0.220096 | 0.215833 | 0.215348 | 0.217594 | 0.216246 | 0.216384 | 0.215397 | 0.215076 | 0.221119 | 0.220548 | 0.218976 | 0.216947 | 0.216897 | 0.215591 | 0.215329 | 0.216325 | 0.220416 | 0.255133 | 0.292940 | 0.284245 | 0.263607 | 0.239035 | 0.218007 | 0.304108 | 0.626533 | 1.116916 | 1.274173 | 1.352785 | 0.984320 | 0.448414 | 0.294459 | 0.222148 | 0.219867 | 0.229980 | 0.232244 | 0.218015 | 0.217150 | 0.214897 | 0.215845 | 0.222822 | 0.223763 | 0.236561 | 0.262182 | 0.312296 | 0.365464 | 0.375032 | 0.383206 | 0.468816 | 0.435007 | 0.348678 | 0.360765 | 0.351720 | 0.465057 | 0.655808 | 0.561418 | 0.325445 | 0.286409 | 0.239122 | 0.252061 | 0.237814 | 0.222429 | 0.216557 | 0.249254 | 0.324700 | 0.476342 | 0.595995 | 0.960375 | 1.353837 | 1.420907 | 1.178217 | 0.774148 | 0.549901 | 0.438920 | 0.344765 |
| left posterior | 3.404823 | 2.065821 | 2.040336 | 1.996443 | 0.872257 | 0.576653 | 0.370980 | 0.284457 | 0.266495 | 0.241260 | 0.215597 | 0.277100 | 0.608360 | 0.566875 | 0.458427 | 0.267086 | 0.215425 | 0.214650 | 0.214626 | 0.215365 | 0.221299 | 0.227658 | 0.496151 | 2.279725 | 4.081081 | 2.904369 | 1.313072 | 0.716466 | 0.389854 | 0.252242 | 0.228170 | 0.249349 | 0.279789 | 0.328238 | 0.446414 | 0.828537 | 1.009084 | 0.881261 | 0.373720 | 0.214879 | 0.298551 | 0.450202 | 0.885040 | 1.518409 | 3.538240 | 2.957368 | 3.532364 | 1.480060 | 0.929713 | 0.587347 | 0.483013 | 0.508898 | 0.418613 | 0.408648 | 0.592902 | 0.472981 | 0.408124 | 0.273699 | 0.219658 | 0.220849 | 0.214790 | 0.236196 | 0.246779 | 0.258170 | 0.262932 | 0.285889 | 0.407348 | 0.621307 | 0.724489 | 0.761258 | 0.660604 | 0.520271 | 0.376634 | 0.253262 | 0.227395 | 0.238613 | 0.265351 | 0.348340 | 0.575152 | 1.556520 | 5.326442 | 5.493119 | 3.581035 | 2.857042 | 3.161779 | 3.356863 | 1.750986 | 1.069284 | 0.743689 | 0.594414 | 0.477535 | 0.371793 | 0.292800 | 0.296352 | 0.291412 | 0.348519 | 0.494752 | 0.668176 | 0.703146 | 0.705493 | 0.517710 | 0.385720 | 0.275656 | 0.233156 | 0.216238 | 0.218936 | 0.248490 | 0.250888 | 0.243371 | 0.216707 | 0.216903 | 0.241190 | 0.338678 | 0.482793 | 0.776489 | 1.300993 | 1.677677 | 1.950126 | 2.345042 | 1.861509 | 1.891751 | 1.297038 | 0.637539 | 0.418572 | 0.291584 | 0.228210 | 0.237573 | 0.248412 | 0.258627 | 0.266280 | 0.264495 | 0.268455 | 0.275376 | 0.243234 | 0.257441 | 0.276385 | 0.309989 | 0.317031 | 0.320537 | 0.291146 | 0.251378 | 0.224439 | 0.214668 | 0.217620 | 0.221556 | 0.241756 | 0.254174 | 0.236984 | 0.237172 | 0.227023 | 0.230884 | 0.252232 | 0.251889 | 0.251577 | 0.276041 | 0.292830 | 0.295817 | 0.289144 | 0.271119 | 0.276087 | 0.291359 | 0.292128 | 0.326302 | 0.410511 | 0.497001 | 0.572472 | 0.700549 | 0.542678 | 0.360312 | 0.290788 | 0.262218 | 0.250606 | 0.242472 | 0.226067 | 0.216507 | 0.218006 | 0.217749 | 0.223207 | 0.225058 | 0.228245 | 0.258015 | 0.414543 | 0.627173 | 1.167362 | 1.258281 | 1.303263 | 1.590898 | 1.286512 | 0.878097 | 0.876777 | 1.072876 | 1.054653 | 1.399664 | 2.223371 | 4.262987 | 6.353421 | 4.833114 | 2.808106 | 3.839572 | 3.872798 | 3.906378 | 2.765078 | 2.394244 | 2.077345 | 1.514029 | 1.286785 | 1.094086 | 0.777823 | 0.704209 | 0.608259 | 0.580675 | 0.723787 | 0.798842 | 0.775502 | 0.565115 | 0.383940 | 0.289751 | 0.245361 | 0.221359 | 0.214782 | 0.217766 | 0.218622 | 0.215408 | 0.218697 | 0.221481 | 0.222232 | 0.222599 | 0.216830 | 0.215987 | 0.214862 | 0.219295 | 0.231175 | 0.238705 | 0.239004 | 0.230744 | 0.226020 | 0.225259 | 0.220317 | 0.214649 | 0.216387 | 0.218617 | 0.219839 | 0.221638 | 0.234479 | 0.258662 | 0.275634 | 0.330447 | 0.517615 | 0.638856 | 0.647603 | 0.518958 | 0.443268 | 0.400894 | 0.388416 | 0.342257 | 0.403164 | 0.435419 | 0.493906 | 0.536307 | 0.635682 | 0.563375 | 0.538359 | 0.410939 | 0.398105 | 0.355102 | 0.364514 | 0.413923 | 0.490058 | 0.486202 | 0.546348 | 0.759339 | 2.059592 | 7.497794 | 17.864180 | 32.336479 | 90.527246 | 246.292184 | 289.188773 | 255.616166 | 167.956910 |
| right posterior | 5.367903 | 1.806557 | 1.397761 | 1.355958 | 1.415938 | 0.809186 | 0.657817 | 0.343839 | 0.373727 | 0.374780 | 0.487575 | 0.400215 | 0.246101 | 0.253144 | 0.263984 | 0.233830 | 0.224973 | 0.268272 | 0.347724 | 0.345621 | 0.248796 | 0.223007 | 0.263323 | 0.288481 | 0.263371 | 0.230525 | 0.221155 | 0.215630 | 0.220684 | 0.217530 | 0.222247 | 0.232206 | 0.227430 | 0.237105 | 0.263425 | 0.219570 | 0.218680 | 0.230264 | 0.226829 | 0.215332 | 0.226780 | 0.264050 | 0.254967 | 0.261271 | 0.380330 | 0.426354 | 0.283152 | 0.223287 | 0.216096 | 0.219878 | 0.226736 | 0.246774 | 0.265992 | 0.254652 | 0.219292 | 0.216499 | 0.221286 | 0.219393 | 0.221978 | 0.229761 | 0.253718 | 0.224865 | 0.219326 | 0.227959 | 0.252230 | 0.276055 | 0.322221 | 0.350067 | 0.372298 | 0.320531 | 0.256150 | 0.232783 | 0.223121 | 0.214635 | 0.216460 | 0.216341 | 0.216845 | 0.233857 | 0.280157 | 0.472712 | 1.678607 | 4.407377 | 15.713474 | 98.592228 | 224.059578 | 166.073653 | 310.193277 | 183.791459 | 240.916552 | 537.756676 | 2183.655761 | 6745.413608 | 4495.920026 | 1787.935001 | 434.770458 | 133.417407 | 29.977234 | 9.507732 | 5.672747 | 2.011037 | 0.581612 | 0.377221 | 0.368122 | 0.410902 | 0.523444 | 0.748418 | 1.268565 | 2.109559 | 2.537013 | 2.671941 | 3.828130 | 5.489415 | 6.183561 | 5.068255 | 5.659033 | 11.227186 | 17.108711 | 22.225879 | 23.033109 | 23.311882 | 19.806437 | 11.005961 | 3.944874 | 1.620345 | 0.745352 | 0.466842 | 0.380935 | 0.284397 | 0.313381 | 0.372962 | 0.464679 | 0.536926 | 0.612901 | 0.673085 | 0.765006 | 0.660788 | 0.557279 | 0.482855 | 0.398318 | 0.313078 | 0.263087 | 0.240671 | 0.222274 | 0.214774 | 0.224201 | 0.222488 | 0.215839 | 0.222347 | 0.257853 | 0.347000 | 0.457754 | 0.560892 | 0.770748 | 1.169782 | 1.304434 | 1.355870 | 1.047411 | 1.668899 | 3.009006 | 2.550167 | 1.414773 | 0.928685 | 0.784337 | 0.810317 | 0.582466 | 0.517479 | 0.711682 | 1.686760 | 3.043704 | 3.083948 | 2.324956 | 2.282208 | 2.248523 | 2.563301 | 1.965800 | 2.146440 | 3.349607 | 7.600681 | 15.139814 | 12.698172 | 6.151791 | 2.940650 | 1.238094 | 0.658842 | 0.332579 | 0.252068 | 0.222419 | 0.215214 | 0.220923 | 0.220482 | 0.222336 | 0.216272 | 0.216442 | 0.214755 | 0.220014 | 0.242187 | 0.281438 | 0.311946 | 0.424427 | 0.708218 | 0.996682 | 1.180354 | 0.944957 | 0.579319 | 0.501971 | 0.414110 | 0.295250 | 0.235510 | 0.215977 | 0.215415 | 0.216913 | 0.226064 | 0.249821 | 0.246139 | 0.230666 | 0.214679 | 0.270190 | 0.366843 | 0.512609 | 0.751441 | 0.712931 | 0.559915 | 0.423800 | 0.272646 | 0.246359 | 0.233876 | 0.227406 | 0.253852 | 0.303642 | 0.365844 | 0.483195 | 0.496332 | 0.514751 | 0.581197 | 0.572420 | 0.658731 | 0.734879 | 0.691763 | 0.830082 | 0.917735 | 0.601544 | 0.369136 | 0.256498 | 0.217798 | 0.215184 | 0.229088 | 0.239071 | 0.231046 | 0.223754 | 0.220900 | 0.214637 | 0.217260 | 0.225653 | 0.227959 | 0.225422 | 0.220166 | 0.216359 | 0.216058 | 0.234974 | 0.251044 | 0.278863 | 0.301472 | 0.312178 | 0.324085 | 0.292772 | 0.259792 | 0.248251 | 0.220807 | 0.217081 | 0.233001 | 0.242082 | 0.246409 | 0.251006 | 0.237752 | 0.214626 | 0.250142 | 0.422044 | 0.448274 | 0.568395 | 1.120785 |
| all electrodes | 0.912496 | 1.131293 | 1.961515 | 10.621556 | 45.639460 | 27.523664 | 12.499969 | 7.849056 | 2.782822 | 0.687988 | 0.255683 | 0.222804 | 0.242006 | 0.248825 | 0.239858 | 0.215506 | 0.249440 | 0.282724 | 0.322433 | 0.529805 | 0.630638 | 0.279061 | 0.214646 | 0.323481 | 0.497951 | 0.775472 | 1.192234 | 1.230557 | 0.496914 | 0.332711 | 0.240856 | 0.219665 | 0.214761 | 0.214626 | 0.220766 | 0.221769 | 0.214999 | 0.214643 | 0.214768 | 0.215467 | 0.220965 | 0.227901 | 0.236984 | 0.274819 | 0.308258 | 0.262818 | 0.255764 | 0.244944 | 0.218015 | 0.216655 | 0.214626 | 0.216087 | 0.214781 | 0.215296 | 0.214897 | 0.214731 | 0.224267 | 0.259796 | 0.314505 | 0.277425 | 0.296499 | 0.216083 | 0.220082 | 0.249541 | 0.313114 | 0.315514 | 0.285396 | 0.281121 | 0.222503 | 0.215559 | 0.248074 | 0.331876 | 0.357206 | 0.347869 | 0.276737 | 0.222736 | 0.217476 | 0.229907 | 0.357038 | 0.685695 | 1.306468 | 1.278093 | 0.822488 | 1.106890 | 1.324160 | 1.543306 | 1.354872 | 0.943154 | 0.822253 | 1.023643 | 1.162700 | 2.421455 | 2.575250 | 3.094155 | 2.249628 | 1.209554 | 0.900355 | 0.681609 | 0.568805 | 0.554594 | 0.587047 | 0.655456 | 0.880848 | 1.016906 | 1.093491 | 0.886899 | 0.805143 | 0.651345 | 0.491109 | 0.416507 | 0.391317 | 0.408694 | 0.389178 | 0.413121 | 0.424557 | 0.439576 | 0.449786 | 0.535396 | 0.428722 | 0.465622 | 0.504655 | 0.494075 | 0.363589 | 0.304666 | 0.264696 | 0.273744 | 0.292355 | 0.333875 | 0.423562 | 0.655425 | 0.818037 | 0.839568 | 0.742140 | 0.576388 | 0.372583 | 0.290721 | 0.273598 | 0.237363 | 0.222513 | 0.248963 | 0.256476 | 0.251209 | 0.251773 | 0.256573 | 0.234119 | 0.215456 | 0.217320 | 0.220315 | 0.249959 | 0.292162 | 0.316030 | 0.336519 | 0.415891 | 0.555708 | 0.958435 | 1.303126 | 1.882032 | 2.493708 | 3.486926 | 3.006231 | 1.985882 | 0.984548 | 0.658992 | 0.494265 | 0.394873 | 0.306595 | 0.339152 | 0.479884 | 0.715819 | 0.959389 | 1.236801 | 1.585558 | 1.337237 | 1.110402 | 0.749932 | 0.566268 | 0.591453 | 0.591707 | 0.406576 | 0.397322 | 0.292547 | 0.256773 | 0.235491 | 0.214686 | 0.222125 | 0.222084 | 0.224751 | 0.228773 | 0.226518 | 0.223978 | 0.215802 | 0.214805 | 0.222592 | 0.220639 | 0.216960 | 0.223663 | 0.240603 | 0.269062 | 0.293057 | 0.249942 | 0.241210 | 0.253629 | 0.259960 | 0.255890 | 0.243969 | 0.258917 | 0.375148 | 0.736216 | 0.765605 | 0.686177 | 0.722149 | 0.947882 | 0.706633 | 0.462519 | 0.281006 | 0.230598 | 0.215487 | 0.216749 | 0.220620 | 0.220983 | 0.225544 | 0.220198 | 0.216304 | 0.254572 | 0.340049 | 0.379622 | 0.374938 | 0.376143 | 0.347448 | 0.278359 | 0.231138 | 0.214654 | 0.216146 | 0.240786 | 0.309737 | 0.362819 | 0.377344 | 0.408398 | 0.374242 | 0.590781 | 0.802285 | 1.067094 | 1.118552 | 0.864862 | 0.736307 | 0.818685 | 0.624484 | 0.235572 | 0.217450 | 0.223023 | 0.218602 | 0.220358 | 0.225422 | 0.220019 | 0.217046 | 0.224482 | 0.218670 | 0.216524 | 0.214761 | 0.214701 | 0.214635 | 0.214704 | 0.218300 | 0.218086 | 0.217646 | 0.215971 | 0.224390 | 0.231866 | 0.237306 | 0.250332 | 0.259371 | 0.302659 | 0.369644 | 0.477355 | 0.700601 | 1.174933 | 2.170968 | 3.094263 | 4.320070 | 3.881237 |

Searchlight, spatiotemporal cluster permutation test

|  | start time | stop time | peak time | peak channel | cluster p | peak Cohen's d | direction |
| --- | --- | --- | --- | --- | --- | --- | --- |
| #1 | 140 | 500 | 260 | CP2 | 0.0002 | 1.146939 | positive |
| #2 | -200 | -80 | -120 | CP1 | 0.0278 | -0.94798 | negative |

I) happy vs sad

  
|  | time window | peak latency | cluster *p* | peak Cohen's *d* |  | | | |
| **all electrodes** |  | | | |  | | | |
|  | | | | | | | | |

Time-resolved classification, cluster permutation tests

|  | **left hemisphere** | | | | **right hemisphere** | | | |
|  | time window | peak latency | cluster *p* | peak Cohen's *d* | time window | peak latency | cluster *p* | peak Cohen's *d* |
| **anterior** |  | | | |  | | | |
| **central** | 500 - 715 ms | 500 ms | 0.0029 | -0.515 | 160 - 300 ms | 225 ms | 0.0177 | 0.6667 |
| **posterior** |  | | | |  | | | |

  

Time-resolved classification, Bayesian statistics

|  | -200 | -195 | -190 | -185 | -180 | -175 | -170 | -165 | -160 | -155 | -150 | -145 | -140 | -135 | -130 | -125 | -120 | -115 | -110 | -105 | -100 | -95 | -90 | -85 | -80 | -75 | -70 | -65 | -60 | -55 | -50 | -45 | -40 | -35 | -30 | -25 | -20 | -15 | -10 | -5 | 0 | 5 | 10 | 15 | 20 | 25 | 30 | 35 | 40 | 45 | 50 | 55 | 60 | 65 | 70 | 75 | 80 | 85 | 90 | 95 | 100 | 105 | 110 | 115 | 120 | 125 | 130 | 135 | 140 | 145 | 150 | 155 | 160 | 165 | 170 | 175 | 180 | 185 | 190 | 195 | 200 | 205 | 210 | 215 | 220 | 225 | 230 | 235 | 240 | 245 | 250 | 255 | 260 | 265 | 270 | 275 | 280 | 285 | 290 | 295 | 300 | 305 | 310 | 315 | 320 | 325 | 330 | 335 | 340 | 345 | 350 | 355 | 360 | 365 | 370 | 375 | 380 | 385 | 390 | 395 | 400 | 405 | 410 | 415 | 420 | 425 | 430 | 435 | 440 | 445 | 450 | 455 | 460 | 465 | 470 | 475 | 480 | 485 | 490 | 495 | 500 | 505 | 510 | 515 | 520 | 525 | 530 | 535 | 540 | 545 | 550 | 555 | 560 | 565 | 570 | 575 | 580 | 585 | 590 | 595 | 600 | 605 | 610 | 615 | 620 | 625 | 630 | 635 | 640 | 645 | 650 | 655 | 660 | 665 | 670 | 675 | 680 | 685 | 690 | 695 | 700 | 705 | 710 | 715 | 720 | 725 | 730 | 735 | 740 | 745 | 750 | 755 | 760 | 765 | 770 | 775 | 780 | 785 | 790 | 795 | 800 | 805 | 810 | 815 | 820 | 825 | 830 | 835 | 840 | 845 | 850 | 855 | 860 | 865 | 870 | 875 | 880 | 885 | 890 | 895 | 900 | 905 | 910 | 915 | 920 | 925 | 930 | 935 | 940 | 945 | 950 | 955 | 960 | 965 | 970 | 975 | 980 | 985 | 990 | 995 | 1000 | 1005 | 1010 | 1015 | 1020 | 1025 | 1030 | 1035 | 1040 | 1045 | 1050 | 1055 | 1060 | 1065 | 1070 | 1075 | 1080 | 1085 | 1090 | 1095 | 1100 | 1105 | 1110 | 1115 | 1120 | 1125 | 1130 | 1135 | 1140 | 1145 | 1150 | 1155 | 1160 | 1165 | 1170 | 1175 | 1180 | 1185 | 1190 | 1195 |
| --- | --- | --- | --- | --- | --- | --- | --- | --- | --- | --- | --- | --- | --- | --- | --- | --- | --- | --- | --- | --- | --- | --- | --- | --- | --- | --- | --- | --- | --- | --- | --- | --- | --- | --- | --- | --- | --- | --- | --- | --- | --- | --- | --- | --- | --- | --- | --- | --- | --- | --- | --- | --- | --- | --- | --- | --- | --- | --- | --- | --- | --- | --- | --- | --- | --- | --- | --- | --- | --- | --- | --- | --- | --- | --- | --- | --- | --- | --- | --- | --- | --- | --- | --- | --- | --- | --- | --- | --- | --- | --- | --- | --- | --- | --- | --- | --- | --- | --- | --- | --- | --- | --- | --- | --- | --- | --- | --- | --- | --- | --- | --- | --- | --- | --- | --- | --- | --- | --- | --- | --- | --- | --- | --- | --- | --- | --- | --- | --- | --- | --- | --- | --- | --- | --- | --- | --- | --- | --- | --- | --- | --- | --- | --- | --- | --- | --- | --- | --- | --- | --- | --- | --- | --- | --- | --- | --- | --- | --- | --- | --- | --- | --- | --- | --- | --- | --- | --- | --- | --- | --- | --- | --- | --- | --- | --- | --- | --- | --- | --- | --- | --- | --- | --- | --- | --- | --- | --- | --- | --- | --- | --- | --- | --- | --- | --- | --- | --- | --- | --- | --- | --- | --- | --- | --- | --- | --- | --- | --- | --- | --- | --- | --- | --- | --- | --- | --- | --- | --- | --- | --- | --- | --- | --- | --- | --- | --- | --- | --- | --- | --- | --- | --- | --- | --- | --- | --- | --- | --- | --- | --- | --- | --- | --- | --- | --- | --- | --- | --- | --- | --- | --- | --- | --- | --- | --- | --- | --- | --- | --- | --- | --- | --- | --- | --- | --- | --- | --- | --- | --- | --- | --- | --- | --- | --- | --- | --- | --- | --- | --- | --- |
| left anterior | 0.714527 | 0.648496 | 0.520998 | 0.443707 | 0.393079 | 0.305345 | 0.224262 | 0.219633 | 0.220690 | 0.239385 | 0.316163 | 0.322113 | 0.237917 | 0.214649 | 0.230585 | 0.215914 | 0.219228 | 0.266463 | 0.231774 | 0.214837 | 0.217839 | 0.237504 | 0.363481 | 0.915534 | 1.119068 | 1.803541 | 3.066387 | 6.140526 | 12.956286 | 9.888153 | 8.871872 | 4.418439 | 0.896413 | 0.276261 | 0.216513 | 0.232568 | 0.239425 | 0.354316 | 0.262333 | 0.214626 | 0.215456 | 0.232433 | 0.234525 | 0.257838 | 0.239961 | 0.261507 | 0.234037 | 0.233772 | 0.275881 | 0.328707 | 0.309303 | 0.282615 | 0.305316 | 0.561686 | 1.045437 | 0.632153 | 0.557626 | 0.654672 | 0.594295 | 0.499626 | 0.415227 | 0.303361 | 0.228930 | 0.215835 | 0.215646 | 0.215052 | 0.218849 | 0.237791 | 0.244205 | 0.246816 | 0.279546 | 0.440195 | 1.571171 | 4.918881 | 11.333612 | 22.596395 | 11.748655 | 4.335835 | 1.109773 | 0.380906 | 0.262419 | 0.217666 | 0.219160 | 0.227384 | 0.214748 | 0.223829 | 0.234271 | 0.236075 | 0.247439 | 0.239878 | 0.220054 | 0.217604 | 0.214753 | 0.218568 | 0.215120 | 0.215789 | 0.214638 | 0.217500 | 0.215253 | 0.214678 | 0.214626 | 0.215143 | 0.220120 | 0.243371 | 0.253607 | 0.242183 | 0.256363 | 0.248791 | 0.233101 | 0.215531 | 0.215543 | 0.220598 | 0.227141 | 0.228768 | 0.258805 | 0.337872 | 0.516558 | 0.757621 | 1.212626 | 0.946761 | 0.642986 | 0.368816 | 0.249123 | 0.217053 | 0.217321 | 0.239374 | 0.261421 | 0.300719 | 0.269724 | 0.229312 | 0.217697 | 0.219229 | 0.229121 | 0.253112 | 0.285916 | 0.297406 | 0.332527 | 0.337455 | 0.318954 | 0.288251 | 0.228755 | 0.215598 | 0.236996 | 0.337854 | 0.438963 | 0.563769 | 0.967604 | 1.273932 | 1.359178 | 1.049497 | 0.602099 | 0.410600 | 0.337324 | 0.268276 | 0.233142 | 0.218251 | 0.216850 | 0.223825 | 0.227957 | 0.241443 | 0.268341 | 0.275478 | 0.302502 | 0.345976 | 0.321456 | 0.375340 | 0.317076 | 0.245743 | 0.225659 | 0.220453 | 0.215646 | 0.214702 | 0.222429 | 0.214877 | 0.222698 | 0.235934 | 0.239847 | 0.241260 | 0.240600 | 0.257835 | 0.253254 | 0.241030 | 0.239245 | 0.222459 | 0.219677 | 0.244973 | 0.246313 | 0.234937 | 0.265752 | 0.322999 | 0.461251 | 0.584606 | 0.815702 | 2.363220 | 5.280026 | 3.778191 | 1.748908 | 0.962512 | 0.726111 | 0.501327 | 0.305026 | 0.257948 | 0.238859 | 0.240972 | 0.252001 | 0.233453 | 0.220757 | 0.217371 | 0.216509 | 0.214775 | 0.220065 | 0.223151 | 0.227359 | 0.231206 | 0.217768 | 0.217388 | 0.273795 | 0.366006 | 0.520466 | 0.600516 | 0.687453 | 0.621213 | 0.472712 | 0.387606 | 0.340253 | 0.281467 | 0.265033 | 0.303188 | 0.311998 | 0.287608 | 0.261949 | 0.275804 | 0.272945 | 0.244656 | 0.220289 | 0.215376 | 0.215843 | 0.225013 | 0.217416 | 0.215747 | 0.217363 | 0.229485 | 0.246247 | 0.261687 | 0.243918 | 0.260842 | 0.336859 | 0.426871 | 0.490102 | 0.458108 | 0.490896 | 0.621971 | 0.638222 | 0.556302 | 0.565402 | 0.529185 | 0.550526 | 0.508945 | 0.424924 | 0.431642 | 0.435044 | 0.466443 | 0.459364 | 0.492297 | 0.508677 | 0.522049 | 0.439155 | 0.369636 | 0.306409 | 0.278361 | 0.258152 | 0.246009 | 0.225545 | 0.216145 | 0.214693 | 0.216411 | 0.220249 | 0.227865 | 0.245434 | 0.239915 |
| right anterior | 0.223952 | 0.258812 | 0.317278 | 0.407640 | 0.435118 | 0.543348 | 0.587091 | 0.900574 | 0.682858 | 0.519141 | 0.284868 | 0.255880 | 0.241751 | 0.255049 | 0.289042 | 0.327410 | 0.302297 | 0.288648 | 0.336261 | 0.353739 | 0.440356 | 0.479138 | 0.594038 | 1.253103 | 6.686594 | 12.176674 | 25.297425 | 15.293490 | 7.555305 | 4.274267 | 0.945464 | 0.329849 | 0.234811 | 0.228058 | 0.239355 | 0.285364 | 0.312268 | 0.319597 | 0.246439 | 0.281898 | 0.249950 | 0.238656 | 0.225250 | 0.242390 | 0.242696 | 0.243875 | 0.224390 | 0.228461 | 0.229684 | 0.233719 | 0.237634 | 0.225746 | 0.232518 | 0.221097 | 0.221902 | 0.230394 | 0.232470 | 0.219716 | 0.240599 | 0.288051 | 0.458367 | 0.760000 | 1.062479 | 1.338324 | 1.253097 | 1.386876 | 1.250391 | 0.764597 | 0.512197 | 0.520421 | 0.629399 | 0.898270 | 0.809141 | 0.799533 | 0.754604 | 0.845544 | 0.538857 | 0.301281 | 0.257861 | 0.292241 | 0.345192 | 0.478630 | 0.611004 | 1.076520 | 4.288289 | 10.129892 | 11.246283 | 11.278940 | 6.095420 | 2.770715 | 1.315883 | 1.137188 | 1.025734 | 0.811395 | 0.448269 | 0.324173 | 0.312738 | 0.319288 | 0.276312 | 0.266950 | 0.287625 | 0.353847 | 0.461445 | 0.559490 | 0.613248 | 0.865694 | 0.820745 | 0.658466 | 0.576765 | 0.375115 | 0.412774 | 0.422124 | 0.351619 | 0.414625 | 0.558260 | 0.636784 | 0.969603 | 0.711251 | 0.835372 | 1.091729 | 0.886373 | 0.583347 | 0.483246 | 0.458977 | 0.732469 | 0.649465 | 0.608084 | 0.398465 | 0.329001 | 0.292531 | 0.247956 | 0.214634 | 0.232868 | 0.301657 | 0.344528 | 0.301058 | 0.295882 | 0.299757 | 0.284543 | 0.340379 | 0.341368 | 0.303969 | 0.306585 | 0.263239 | 0.235565 | 0.214776 | 0.228350 | 0.253238 | 0.332474 | 0.359889 | 0.428953 | 0.463016 | 0.385626 | 0.342144 | 0.265716 | 0.224628 | 0.217570 | 0.216921 | 0.216247 | 0.216533 | 0.215669 | 0.215649 | 0.218787 | 0.217309 | 0.214639 | 0.217844 | 0.217998 | 0.217010 | 0.216016 | 0.226257 | 0.260752 | 0.307772 | 0.477326 | 0.711305 | 0.975408 | 1.176213 | 1.181112 | 1.003964 | 0.786212 | 0.869345 | 1.258473 | 0.993689 | 0.703859 | 0.583934 | 0.401081 | 0.306457 | 0.248983 | 0.220418 | 0.229219 | 0.260475 | 0.332518 | 0.345812 | 0.316377 | 0.381077 | 0.522526 | 0.436489 | 0.434943 | 0.434671 | 0.360663 | 0.333199 | 0.249819 | 0.218411 | 0.224504 | 0.249120 | 0.242699 | 0.215089 | 0.222707 | 0.227912 | 0.224514 | 0.242589 | 0.290155 | 0.245119 | 0.224291 | 0.218774 | 0.246959 | 0.275677 | 0.356621 | 0.303174 | 0.365116 | 0.353999 | 0.344895 | 0.315387 | 0.343704 | 0.373263 | 0.564298 | 0.774234 | 0.671983 | 0.514784 | 0.427541 | 0.387287 | 0.429428 | 0.396539 | 0.537655 | 0.828686 | 0.750270 | 0.858609 | 0.607182 | 0.358541 | 0.303683 | 0.236176 | 0.223966 | 0.279431 | 0.386897 | 0.424830 | 0.473007 | 0.483748 | 0.561117 | 0.721500 | 0.867470 | 0.948593 | 1.659648 | 1.825452 | 2.064053 | 1.331401 | 1.291513 | 1.219768 | 0.969093 | 0.674339 | 0.607201 | 0.576739 | 0.802802 | 0.753642 | 0.637266 | 0.628012 | 0.748488 | 0.735335 | 0.695317 | 0.485574 | 0.377622 | 0.304955 | 0.256906 | 0.225543 | 0.220339 | 0.216560 | 0.217321 | 0.218347 | 0.216365 | 0.216084 | 0.216059 | 0.215162 |
| left central | 0.277295 | 0.359748 | 0.391111 | 0.435193 | 0.739204 | 0.779516 | 0.643400 | 0.649237 | 0.443841 | 0.392575 | 0.492217 | 0.400956 | 0.517428 | 0.539145 | 0.428292 | 0.436699 | 0.435771 | 0.412740 | 0.563929 | 1.271728 | 2.481317 | 2.186468 | 3.078105 | 3.816739 | 12.386590 | 11.706406 | 6.631711 | 1.591319 | 0.620161 | 0.239040 | 0.226817 | 0.274881 | 0.309097 | 0.427945 | 0.253997 | 0.215613 | 0.218602 | 0.232254 | 0.223696 | 0.219750 | 0.246586 | 0.215708 | 0.222946 | 0.221671 | 0.215754 | 0.215866 | 0.220946 | 0.239303 | 0.264822 | 0.276269 | 0.282518 | 0.406031 | 0.617302 | 0.682301 | 0.916504 | 1.499292 | 2.743906 | 4.279696 | 2.345832 | 1.006324 | 0.500421 | 0.471871 | 0.278573 | 0.243014 | 0.237730 | 0.244139 | 0.319633 | 0.591374 | 0.476032 | 0.321649 | 0.240825 | 0.215487 | 0.261285 | 0.355803 | 0.537164 | 0.757389 | 0.732784 | 0.473359 | 0.330302 | 0.294371 | 0.277176 | 0.309383 | 0.258004 | 0.244298 | 0.262458 | 0.240657 | 0.222512 | 0.214626 | 0.274250 | 0.350443 | 0.524869 | 0.611163 | 0.497154 | 0.570234 | 0.624677 | 0.485077 | 0.499315 | 0.557527 | 0.984393 | 1.468903 | 1.689988 | 1.636747 | 1.603498 | 2.411690 | 2.727249 | 1.679468 | 2.600148 | 2.697063 | 3.683339 | 4.301452 | 2.481773 | 1.530219 | 1.792193 | 1.486379 | 2.408809 | 3.335683 | 1.374489 | 0.639576 | 0.423024 | 0.292029 | 0.263719 | 0.220151 | 0.220049 | 0.232664 | 0.232997 | 0.221073 | 0.214680 | 0.218484 | 0.234140 | 0.254680 | 0.304514 | 0.255726 | 0.229983 | 0.221118 | 0.214896 | 0.232405 | 0.262468 | 0.323251 | 0.340112 | 0.647298 | 2.839704 | 14.310639 | 104.564640 | 557.439021 | 1148.377419 | 1792.108972 | 242.454513 | 20.294725 | 5.164472 | 2.030123 | 1.546797 | 2.889022 | 6.462275 | 27.690441 | 188.936994 | 350.543422 | 525.161182 | 697.655933 | 769.717836 | 648.622328 | 126.811349 | 21.490127 | 7.757068 | 4.732828 | 9.000902 | 20.803903 | 46.234591 | 106.365700 | 408.840304 | 454.546240 | 166.677812 | 31.102795 | 6.418630 | 4.410631 | 15.522835 | 32.932721 | 49.817234 | 135.080189 | 128.121910 | 65.663453 | 29.587124 | 5.491152 | 1.736672 | 1.411429 | 0.872763 | 0.630538 | 0.452339 | 0.290431 | 0.246738 | 0.221291 | 0.217871 | 0.224337 | 0.227448 | 0.222716 | 0.214661 | 0.219215 | 0.221128 | 0.225169 | 0.216352 | 0.214767 | 0.216294 | 0.219772 | 0.224014 | 0.221759 | 0.221673 | 0.219063 | 0.217410 | 0.218235 | 0.218914 | 0.219181 | 0.219562 | 0.219283 | 0.216655 | 0.218747 | 0.220694 | 0.218599 | 0.215994 | 0.215090 | 0.226597 | 0.250348 | 0.278645 | 0.294565 | 0.288869 | 0.360719 | 0.447235 | 0.576291 | 0.777358 | 0.999317 | 1.279973 | 1.817105 | 1.588290 | 1.182989 | 0.645148 | 0.457598 | 0.474772 | 0.533344 | 0.648381 | 0.778754 | 0.837050 | 1.120647 | 1.682333 | 2.504876 | 2.384998 | 2.027353 | 1.487439 | 1.353556 | 1.355988 | 1.265625 | 1.151990 | 1.200612 | 1.056795 | 1.026005 | 1.056033 | 1.100194 | 0.830667 | 0.548714 | 0.432425 | 0.443123 | 0.573980 | 0.851350 | 0.861686 | 0.996898 | 0.984854 | 0.812204 | 0.843345 | 0.909986 | 0.889170 | 0.884590 | 1.105325 | 1.816289 | 4.505382 | 10.164961 | 12.653539 | 11.038744 | 14.262612 | 10.872939 | 3.885330 | 2.806653 | 1.536854 | 0.797691 |
| right central | 0.355112 | 0.300239 | 0.270743 | 0.278062 | 0.262456 | 0.247274 | 0.284506 | 0.270945 | 0.286770 | 0.291366 | 0.264966 | 0.231107 | 0.214852 | 0.223100 | 0.215151 | 0.241284 | 0.258439 | 0.216896 | 0.214642 | 0.215882 | 0.237841 | 0.228720 | 0.217597 | 0.259744 | 0.271720 | 0.229841 | 0.216492 | 0.216345 | 0.237969 | 0.280923 | 0.241694 | 0.214742 | 0.218746 | 0.218178 | 0.216133 | 0.236388 | 0.313815 | 0.602043 | 1.457445 | 3.191898 | 2.372025 | 0.747376 | 0.529099 | 0.277623 | 0.214671 | 0.219739 | 0.228345 | 0.214717 | 0.226654 | 0.219517 | 0.222071 | 0.258779 | 0.269734 | 0.325273 | 0.296583 | 0.281935 | 0.293224 | 0.378202 | 0.369335 | 0.287559 | 0.214638 | 0.236960 | 0.261139 | 0.248680 | 0.217220 | 0.225362 | 0.249424 | 0.293656 | 0.258067 | 0.218554 | 0.244133 | 0.451124 | 1.352832 | 2.382266 | 3.129139 | 4.685493 | 3.211416 | 1.594873 | 2.033020 | 3.757666 | 6.128903 | 10.007781 | 4.728776 | 4.300368 | 6.766821 | 12.125811 | 12.278365 | 9.384404 | 5.582510 | 5.978508 | 7.854881 | 15.569040 | 17.882990 | 10.203178 | 2.937311 | 2.408570 | 1.798380 | 1.769286 | 1.629946 | 1.375722 | 1.310638 | 1.258474 | 1.345066 | 1.185658 | 0.921766 | 0.604550 | 0.530102 | 0.460048 | 0.503466 | 0.368109 | 0.276679 | 0.226234 | 0.218443 | 0.215960 | 0.218038 | 0.217007 | 0.226534 | 0.257461 | 0.330662 | 0.450014 | 0.440060 | 0.483994 | 0.503565 | 0.491898 | 0.489078 | 0.444766 | 0.365463 | 0.406224 | 0.339036 | 0.321134 | 0.284470 | 0.262749 | 0.244915 | 0.229927 | 0.214663 | 0.226154 | 0.228031 | 0.220490 | 0.215619 | 0.216538 | 0.229184 | 0.250372 | 0.302456 | 0.376404 | 0.476193 | 0.585774 | 0.684148 | 0.723653 | 0.888892 | 1.377861 | 1.761338 | 1.539296 | 1.553841 | 1.021530 | 0.816589 | 0.620830 | 0.386227 | 0.279178 | 0.250596 | 0.236479 | 0.236060 | 0.236333 | 0.267977 | 0.349065 | 0.401284 | 0.483320 | 0.670320 | 0.707098 | 0.588451 | 0.526856 | 0.360956 | 0.270989 | 0.233841 | 0.215322 | 0.230635 | 0.256492 | 0.293846 | 0.280402 | 0.257850 | 0.243797 | 0.229472 | 0.218270 | 0.214669 | 0.214639 | 0.214749 | 0.215673 | 0.214830 | 0.214961 | 0.224157 | 0.248594 | 0.327267 | 0.464239 | 0.704660 | 1.151967 | 1.367576 | 1.179910 | 1.304595 | 1.582198 | 1.347217 | 0.922795 | 0.643095 | 0.572743 | 0.631905 | 0.483501 | 0.337616 | 0.285503 | 0.247305 | 0.242434 | 0.230832 | 0.221659 | 0.233067 | 0.250401 | 0.248142 | 0.259922 | 0.242346 | 0.222477 | 0.217025 | 0.214833 | 0.214631 | 0.216665 | 0.215711 | 0.221842 | 0.234549 | 0.225144 | 0.217717 | 0.216336 | 0.226248 | 0.221730 | 0.230446 | 0.238416 | 0.224828 | 0.217980 | 0.217732 | 0.220786 | 0.246603 | 0.270469 | 0.313774 | 0.361827 | 0.339053 | 0.315074 | 0.293758 | 0.244158 | 0.217155 | 0.218157 | 0.225976 | 0.223508 | 0.228566 | 0.237155 | 0.226829 | 0.214933 | 0.216836 | 0.214948 | 0.216581 | 0.220200 | 0.222164 | 0.228679 | 0.239092 | 0.265763 | 0.268488 | 0.248376 | 0.229701 | 0.218937 | 0.215189 | 0.215878 | 0.224091 | 0.247173 | 0.244415 | 0.249565 | 0.272675 | 0.301280 | 0.301668 | 0.321983 | 0.338390 | 0.430912 | 0.460963 | 0.380008 | 0.347633 | 0.354408 | 0.345179 | 0.318206 |
| left posterior | 85.060240 | 18.827916 | 7.550530 | 4.802067 | 3.678903 | 1.370789 | 0.425205 | 0.221061 | 0.218275 | 0.328996 | 0.723092 | 2.445178 | 6.320592 | 8.374790 | 2.742733 | 0.868754 | 0.247410 | 0.256545 | 1.431160 | 11.879738 | 80.103695 | 74.305262 | 31.509129 | 5.083900 | 1.819788 | 0.369361 | 0.215615 | 0.216256 | 0.220913 | 0.236511 | 0.234474 | 0.245867 | 0.260410 | 0.255180 | 0.274414 | 0.281633 | 0.238286 | 0.222236 | 0.215823 | 0.216560 | 0.228325 | 0.226803 | 0.229759 | 0.221003 | 0.223146 | 0.214889 | 0.243661 | 0.298507 | 0.346150 | 0.329241 | 0.270846 | 0.252437 | 0.219572 | 0.233963 | 0.310628 | 0.358925 | 0.391065 | 0.264104 | 0.218996 | 0.249932 | 0.478488 | 0.783494 | 0.831613 | 0.988466 | 0.621228 | 0.464713 | 0.272054 | 0.215848 | 0.216876 | 0.214972 | 0.230374 | 0.446968 | 1.293121 | 4.670368 | 26.753655 | 51.810845 | 60.259854 | 36.488242 | 9.262197 | 3.525458 | 2.024631 | 1.198515 | 1.287478 | 1.143476 | 1.142022 | 1.023321 | 0.963058 | 0.866524 | 0.665467 | 0.476932 | 0.426178 | 0.324831 | 0.306451 | 0.282948 | 0.247770 | 0.235295 | 0.261831 | 0.296710 | 0.365683 | 0.375145 | 0.397702 | 0.477714 | 0.503085 | 0.413599 | 0.406932 | 0.431659 | 0.450928 | 0.344764 | 0.261516 | 0.226812 | 0.214686 | 0.217955 | 0.233025 | 0.232701 | 0.214650 | 0.241562 | 0.297770 | 0.387011 | 0.406871 | 0.422181 | 0.387198 | 0.369652 | 0.337307 | 0.320658 | 0.356121 | 0.396831 | 0.460522 | 0.456202 | 0.359248 | 0.274889 | 0.233332 | 0.218635 | 0.215172 | 0.214752 | 0.214626 | 0.214704 | 0.217221 | 0.228775 | 0.249128 | 0.262590 | 0.253476 | 0.257909 | 0.266973 | 0.246352 | 0.230843 | 0.214777 | 0.217017 | 0.220309 | 0.222844 | 0.222618 | 0.225877 | 0.237944 | 0.252672 | 0.273446 | 0.280404 | 0.267172 | 0.271999 | 0.241218 | 0.218902 | 0.216138 | 0.222507 | 0.219461 | 0.215963 | 0.215645 | 0.216325 | 0.215470 | 0.214730 | 0.219484 | 0.217473 | 0.215614 | 0.216092 | 0.222671 | 0.233213 | 0.231432 | 0.227171 | 0.228586 | 0.230209 | 0.230242 | 0.222122 | 0.216970 | 0.214709 | 0.217566 | 0.226172 | 0.244523 | 0.269746 | 0.292895 | 0.312773 | 0.299941 | 0.322809 | 0.359490 | 0.386369 | 0.385426 | 0.348597 | 0.319490 | 0.379946 | 0.419504 | 0.436279 | 0.403480 | 0.362674 | 0.385838 | 0.425930 | 0.407060 | 0.374572 | 0.377573 | 0.471447 | 0.662500 | 0.772758 | 0.962753 | 1.036177 | 1.133930 | 1.149943 | 1.024551 | 0.842287 | 0.897319 | 1.024458 | 1.134568 | 1.169132 | 1.154540 | 1.056191 | 0.859541 | 0.593854 | 0.453171 | 0.355539 | 0.330830 | 0.320204 | 0.347412 | 0.418229 | 0.409749 | 0.414785 | 0.404549 | 0.365827 | 0.334276 | 0.280026 | 0.259174 | 0.284247 | 0.278484 | 0.296990 | 0.327484 | 0.344678 | 0.410417 | 0.429441 | 0.441315 | 0.448838 | 0.482779 | 0.509906 | 0.789932 | 1.241819 | 1.923811 | 2.702151 | 3.051905 | 2.277206 | 1.529919 | 0.797363 | 0.452565 | 0.319038 | 0.251241 | 0.225023 | 0.224114 | 0.247262 | 0.290487 | 0.359493 | 0.467610 | 0.572531 | 0.780484 | 0.925388 | 0.943930 | 0.893022 | 0.919559 | 0.891319 | 0.833632 | 0.813298 | 0.854085 | 0.706147 | 0.559787 | 0.473151 | 0.512993 | 0.685471 | 0.650525 | 0.576454 | 0.580489 |
| right posterior | 0.255614 | 0.253413 | 0.249160 | 0.252076 | 0.241081 | 0.239524 | 0.216262 | 0.226886 | 0.249181 | 0.248606 | 0.269160 | 0.259427 | 0.253890 | 0.225998 | 0.215096 | 0.224609 | 0.230637 | 0.240294 | 0.255020 | 0.240637 | 0.217235 | 0.215890 | 0.225509 | 0.289664 | 0.422642 | 0.670220 | 2.385032 | 20.925438 | 33.560808 | 4.424439 | 0.564228 | 0.295622 | 0.225768 | 0.216251 | 0.279587 | 0.351529 | 0.337382 | 0.293255 | 0.235744 | 0.229316 | 0.232056 | 0.245682 | 0.254305 | 0.290658 | 0.273555 | 0.307086 | 0.292866 | 0.274136 | 0.236348 | 0.219109 | 0.215023 | 0.223281 | 0.233855 | 0.230997 | 0.221559 | 0.220757 | 0.214635 | 0.229794 | 0.453153 | 1.435729 | 3.964664 | 9.607694 | 7.920539 | 2.145412 | 0.445804 | 0.214668 | 0.498298 | 1.807633 | 3.083319 | 1.893710 | 0.504293 | 0.215913 | 0.350725 | 0.998698 | 2.407787 | 3.484627 | 2.960504 | 1.369735 | 0.680909 | 0.367910 | 0.295275 | 0.261920 | 0.238474 | 0.235171 | 0.232090 | 0.232537 | 0.221456 | 0.214626 | 0.231241 | 0.288070 | 0.324091 | 0.318304 | 0.282477 | 0.250456 | 0.242692 | 0.228994 | 0.223930 | 0.216097 | 0.214969 | 0.214841 | 0.215904 | 0.215730 | 0.216391 | 0.224984 | 0.222924 | 0.218684 | 0.217225 | 0.221919 | 0.235390 | 0.243512 | 0.239318 | 0.244297 | 0.230330 | 0.224610 | 0.216196 | 0.214849 | 0.217377 | 0.223336 | 0.224087 | 0.214631 | 0.225592 | 0.239508 | 0.283428 | 0.346705 | 0.547394 | 0.768016 | 0.727983 | 0.661252 | 0.591522 | 0.408324 | 0.319345 | 0.261372 | 0.219528 | 0.215040 | 0.231238 | 0.252765 | 0.288414 | 0.308207 | 0.338408 | 0.304724 | 0.267086 | 0.231485 | 0.217491 | 0.215088 | 0.226469 | 0.241379 | 0.258016 | 0.279939 | 0.289733 | 0.278347 | 0.261344 | 0.233181 | 0.217707 | 0.215107 | 0.234951 | 0.264173 | 0.311383 | 0.338394 | 0.290424 | 0.241382 | 0.232384 | 0.224138 | 0.217740 | 0.214697 | 0.220111 | 0.227218 | 0.233369 | 0.251352 | 0.291384 | 0.278223 | 0.240278 | 0.219659 | 0.266210 | 0.324536 | 0.497290 | 0.758322 | 0.975200 | 1.175482 | 0.747530 | 0.583079 | 0.450040 | 0.309882 | 0.246181 | 0.215920 | 0.217372 | 0.230926 | 0.236602 | 0.271969 | 0.291279 | 0.318797 | 0.322292 | 0.331690 | 0.315565 | 0.319635 | 0.263909 | 0.246675 | 0.237649 | 0.228234 | 0.224988 | 0.222760 | 0.228061 | 0.247121 | 0.260279 | 0.253770 | 0.255705 | 0.244617 | 0.244670 | 0.256185 | 0.269401 | 0.283921 | 0.326040 | 0.352125 | 0.381412 | 0.361262 | 0.327549 | 0.306168 | 0.315692 | 0.326765 | 0.333497 | 0.323849 | 0.376504 | 0.351808 | 0.305943 | 0.238903 | 0.217771 | 0.214891 | 0.214997 | 0.219193 | 0.218346 | 0.214701 | 0.229865 | 0.258874 | 0.271315 | 0.323784 | 0.378888 | 0.417793 | 0.369554 | 0.362177 | 0.321923 | 0.317388 | 0.287023 | 0.272576 | 0.290563 | 0.315395 | 0.313924 | 0.374995 | 0.428861 | 0.537839 | 0.698298 | 0.534727 | 0.487551 | 0.506222 | 0.455426 | 0.383311 | 0.295561 | 0.245483 | 0.231857 | 0.221928 | 0.216328 | 0.214820 | 0.215434 | 0.214676 | 0.214676 | 0.214764 | 0.216460 | 0.229063 | 0.238065 | 0.280419 | 0.348028 | 0.384866 | 0.447760 | 0.439531 | 0.404569 | 0.388050 | 0.344876 | 0.276630 | 0.238619 | 0.229781 | 0.230248 | 0.218430 |
| all electrodes | 0.483593 | 0.264701 | 0.233896 | 0.269132 | 0.230812 | 0.217757 | 0.248145 | 0.358736 | 0.691957 | 0.848663 | 0.512538 | 0.237394 | 0.220207 | 0.248928 | 0.314720 | 0.382862 | 0.340379 | 0.345002 | 0.235657 | 0.251456 | 0.263386 | 0.257168 | 0.242738 | 0.226201 | 0.303207 | 0.434261 | 0.791381 | 0.953179 | 1.979127 | 1.522203 | 0.708983 | 0.428554 | 0.334421 | 0.292671 | 0.222672 | 0.229554 | 0.227140 | 0.221968 | 0.216052 | 0.215587 | 0.215260 | 0.226945 | 0.321456 | 0.382879 | 0.563395 | 0.479410 | 0.371687 | 0.322869 | 0.332102 | 0.316366 | 0.271174 | 0.235813 | 0.234324 | 0.283265 | 0.316417 | 0.292083 | 0.324888 | 0.447772 | 0.364360 | 0.253639 | 0.217775 | 0.229183 | 0.242478 | 0.231663 | 0.216085 | 0.231512 | 0.391460 | 0.359422 | 0.320500 | 0.231400 | 0.238277 | 0.441035 | 1.340316 | 6.195905 | 14.166944 | 24.644775 | 19.607238 | 10.652408 | 8.117448 | 5.201108 | 3.591515 | 4.416456 | 3.227704 | 1.873519 | 1.437752 | 1.678788 | 1.219617 | 1.047904 | 0.625471 | 0.417206 | 0.361070 | 0.361196 | 0.339039 | 0.297051 | 0.263122 | 0.248207 | 0.287619 | 0.360379 | 0.319125 | 0.252422 | 0.250731 | 0.270572 | 0.314574 | 0.299593 | 0.278113 | 0.296609 | 0.341227 | 0.309137 | 0.278542 | 0.238367 | 0.228703 | 0.216099 | 0.215346 | 0.214877 | 0.221712 | 0.255970 | 0.448076 | 0.953555 | 2.601393 | 6.317903 | 5.820467 | 3.420584 | 2.132609 | 1.222724 | 0.712496 | 0.511986 | 0.361444 | 0.268792 | 0.228340 | 0.216737 | 0.214647 | 0.215115 | 0.215141 | 0.215669 | 0.215090 | 0.220302 | 0.237137 | 0.249373 | 0.267501 | 0.268448 | 0.240437 | 0.230402 | 0.230013 | 0.217058 | 0.215340 | 0.214850 | 0.219887 | 0.214771 | 0.214626 | 0.214626 | 0.214860 | 0.214698 | 0.217510 | 0.218153 | 0.219728 | 0.215483 | 0.219841 | 0.222780 | 0.215695 | 0.224202 | 0.232681 | 0.224588 | 0.216913 | 0.218435 | 0.216284 | 0.215323 | 0.215528 | 0.217027 | 0.216133 | 0.215919 | 0.219237 | 0.223118 | 0.250725 | 0.288539 | 0.258428 | 0.246419 | 0.234189 | 0.227454 | 0.218126 | 0.216530 | 0.229367 | 0.228532 | 0.233621 | 0.225180 | 0.252231 | 0.265018 | 0.273882 | 0.270813 | 0.278669 | 0.291352 | 0.334421 | 0.378002 | 0.392250 | 0.290789 | 0.286371 | 0.361756 | 0.451462 | 0.435423 | 0.272948 | 0.230733 | 0.226140 | 0.221132 | 0.214998 | 0.214828 | 0.216671 | 0.234408 | 0.259226 | 0.308487 | 0.385135 | 0.460987 | 0.513399 | 0.499967 | 0.482345 | 0.488785 | 0.539933 | 0.494265 | 0.482725 | 0.401155 | 0.384221 | 0.473005 | 0.621700 | 0.578050 | 0.578987 | 0.398990 | 0.393256 | 0.332667 | 0.248698 | 0.229102 | 0.233410 | 0.240969 | 0.284855 | 0.316956 | 0.316558 | 0.381650 | 0.305809 | 0.242289 | 0.220482 | 0.216167 | 0.214765 | 0.214635 | 0.216863 | 0.215986 | 0.229509 | 0.266068 | 0.284864 | 0.284632 | 0.342388 | 0.478511 | 0.518886 | 0.587730 | 0.583088 | 0.416315 | 0.341545 | 0.278435 | 0.275385 | 0.271546 | 0.261953 | 0.247001 | 0.247679 | 0.259179 | 0.295798 | 0.279862 | 0.259239 | 0.250360 | 0.243914 | 0.229312 | 0.221589 | 0.214857 | 0.222416 | 0.232584 | 0.251540 | 0.286623 | 0.281749 | 0.302803 | 0.272558 | 0.239534 | 0.220680 | 0.217117 | 0.214740 | 0.217966 |

Searchlight, spatiotemporal cluster permutation test

No significant clusters observed.
