## Supplementary Table for "Shared Neural Dynamics of Facial Expression Processing": SupplementaryTable_3.html

  


### Supplementary Table 3.

  

Classification analyses

  

**Identity and sex decoding.** Identity and sex decoding for emotional expressions vs neutral faces (classification accuracies for the directions emotion-to-neutral and neutral-to-emotion averaged).

  


**A** identity, happy

**B** identity, angry

**C** identity, sad

**G** sex, happy

**H** sex, angry

**I** sex, sad

**D** difference, identity, happy vs angry

**E** difference, identity, angry vs sad

**F** difference, identity, happy vs sad

**J** difference, sex, happy vs angry

**K** difference, sex, angry vs sad

**L** difference, sex, happy vs sad

A) identity, happy

  
|  | time window | peak latency | cluster *p* | peak Cohen's *d* |  | | | |
| **all electrodes** | 260 - 360 ms | 275 ms | 0.0193 | 0.8727 |  | | | |
|  | | | | | | | | |

Time-resolved classification, cluster permutation tests

|  | **left hemisphere** | | | | **right hemisphere** | | | |
|  | time window | peak latency | cluster *p* | peak Cohen's *d* | time window | peak latency | cluster *p* | peak Cohen's *d* |
| **anterior** |  | | | |  | | | |
| **central** |  | | | |  | | | |
| **posterior** |  | | | |  | | | |

  

Time-resolved classification, Bayesian statistics

|  | -200 | -195 | -190 | -185 | -180 | -175 | -170 | -165 | -160 | -155 | -150 | -145 | -140 | -135 | -130 | -125 | -120 | -115 | -110 | -105 | -100 | -95 | -90 | -85 | -80 | -75 | -70 | -65 | -60 | -55 | -50 | -45 | -40 | -35 | -30 | -25 | -20 | -15 | -10 | -5 | 0 | 5 | 10 | 15 | 20 | 25 | 30 | 35 | 40 | 45 | 50 | 55 | 60 | 65 | 70 | 75 | 80 | 85 | 90 | 95 | 100 | 105 | 110 | 115 | 120 | 125 | 130 | 135 | 140 | 145 | 150 | 155 | 160 | 165 | 170 | 175 | 180 | 185 | 190 | 195 | 200 | 205 | 210 | 215 | 220 | 225 | 230 | 235 | 240 | 245 | 250 | 255 | 260 | 265 | 270 | 275 | 280 | 285 | 290 | 295 | 300 | 305 | 310 | 315 | 320 | 325 | 330 | 335 | 340 | 345 | 350 | 355 | 360 | 365 | 370 | 375 | 380 | 385 | 390 | 395 | 400 | 405 | 410 | 415 | 420 | 425 | 430 | 435 | 440 | 445 | 450 | 455 | 460 | 465 | 470 | 475 | 480 | 485 | 490 | 495 | 500 | 505 | 510 | 515 | 520 | 525 | 530 | 535 | 540 | 545 | 550 | 555 | 560 | 565 | 570 | 575 | 580 | 585 | 590 | 595 | 600 | 605 | 610 | 615 | 620 | 625 | 630 | 635 | 640 | 645 | 650 | 655 | 660 | 665 | 670 | 675 | 680 | 685 | 690 | 695 | 700 | 705 | 710 | 715 | 720 | 725 | 730 | 735 | 740 | 745 | 750 | 755 | 760 | 765 | 770 | 775 | 780 | 785 | 790 | 795 | 800 | 805 | 810 | 815 | 820 | 825 | 830 | 835 | 840 | 845 | 850 | 855 | 860 | 865 | 870 | 875 | 880 | 885 | 890 | 895 | 900 | 905 | 910 | 915 | 920 | 925 | 930 | 935 | 940 | 945 | 950 | 955 | 960 | 965 | 970 | 975 | 980 | 985 | 990 | 995 | 1000 | 1005 | 1010 | 1015 | 1020 | 1025 | 1030 | 1035 | 1040 | 1045 | 1050 | 1055 | 1060 | 1065 | 1070 | 1075 | 1080 | 1085 | 1090 | 1095 | 1100 | 1105 | 1110 | 1115 | 1120 | 1125 | 1130 | 1135 | 1140 | 1145 | 1150 | 1155 | 1160 | 1165 | 1170 | 1175 | 1180 | 1185 | 1190 | 1195 |
| --- | --- | --- | --- | --- | --- | --- | --- | --- | --- | --- | --- | --- | --- | --- | --- | --- | --- | --- | --- | --- | --- | --- | --- | --- | --- | --- | --- | --- | --- | --- | --- | --- | --- | --- | --- | --- | --- | --- | --- | --- | --- | --- | --- | --- | --- | --- | --- | --- | --- | --- | --- | --- | --- | --- | --- | --- | --- | --- | --- | --- | --- | --- | --- | --- | --- | --- | --- | --- | --- | --- | --- | --- | --- | --- | --- | --- | --- | --- | --- | --- | --- | --- | --- | --- | --- | --- | --- | --- | --- | --- | --- | --- | --- | --- | --- | --- | --- | --- | --- | --- | --- | --- | --- | --- | --- | --- | --- | --- | --- | --- | --- | --- | --- | --- | --- | --- | --- | --- | --- | --- | --- | --- | --- | --- | --- | --- | --- | --- | --- | --- | --- | --- | --- | --- | --- | --- | --- | --- | --- | --- | --- | --- | --- | --- | --- | --- | --- | --- | --- | --- | --- | --- | --- | --- | --- | --- | --- | --- | --- | --- | --- | --- | --- | --- | --- | --- | --- | --- | --- | --- | --- | --- | --- | --- | --- | --- | --- | --- | --- | --- | --- | --- | --- | --- | --- | --- | --- | --- | --- | --- | --- | --- | --- | --- | --- | --- | --- | --- | --- | --- | --- | --- | --- | --- | --- | --- | --- | --- | --- | --- | --- | --- | --- | --- | --- | --- | --- | --- | --- | --- | --- | --- | --- | --- | --- | --- | --- | --- | --- | --- | --- | --- | --- | --- | --- | --- | --- | --- | --- | --- | --- | --- | --- | --- | --- | --- | --- | --- | --- | --- | --- | --- | --- | --- | --- | --- | --- | --- | --- | --- | --- | --- | --- | --- | --- | --- | --- | --- | --- | --- | --- | --- | --- | --- | --- | --- | --- | --- | --- | --- |
| left anterior | 2.058533 | 2.127543 | 2.139890 | 2.852714 | 2.021929 | 0.887698 | 0.276262 | 0.217881 | 0.358435 | 0.322141 | 0.363509 | 0.459407 | 0.416713 | 0.392107 | 0.392343 | 0.349079 | 0.242763 | 0.215118 | 0.214679 | 0.217949 | 0.221677 | 0.262106 | 0.336370 | 0.282445 | 0.333546 | 0.330272 | 0.285995 | 0.320541 | 0.317075 | 0.244496 | 0.222166 | 0.435462 | 1.235619 | 1.198213 | 1.274690 | 1.031989 | 0.452690 | 0.241212 | 0.214737 | 0.223914 | 0.260082 | 0.272268 | 0.277122 | 0.228348 | 0.229276 | 0.238045 | 0.223351 | 0.215223 | 0.258715 | 0.272201 | 0.246811 | 0.228837 | 0.216585 | 0.257548 | 0.304560 | 0.443212 | 0.415566 | 0.310737 | 0.239214 | 0.216579 | 0.215153 | 0.233142 | 0.260612 | 0.267832 | 0.229437 | 0.218951 | 0.222858 | 0.223015 | 0.316718 | 0.376857 | 0.453165 | 0.412019 | 0.314055 | 0.344214 | 0.410433 | 0.399935 | 0.519939 | 0.548676 | 0.748120 | 0.871565 | 0.941655 | 0.907088 | 0.535696 | 0.354948 | 0.305511 | 0.234339 | 0.223450 | 0.273485 | 0.346645 | 0.334931 | 0.455787 | 0.689209 | 0.560273 | 0.361045 | 0.349655 | 0.314604 | 0.319559 | 0.269010 | 0.229381 | 0.220616 | 0.221019 | 0.235799 | 0.251980 | 0.249622 | 0.235888 | 0.256308 | 0.361305 | 0.435013 | 0.356919 | 0.340225 | 0.351255 | 0.408599 | 0.468994 | 0.317118 | 0.237183 | 0.216189 | 0.214651 | 0.218723 | 0.277357 | 0.402238 | 0.588540 | 0.576071 | 0.579052 | 0.845288 | 0.534560 | 0.326658 | 0.232293 | 0.214850 | 0.214837 | 0.219095 | 0.239080 | 0.215447 | 0.215788 | 0.218571 | 0.216974 | 0.263433 | 0.302185 | 0.291017 | 0.252704 | 0.257164 | 0.232426 | 0.218560 | 0.217513 | 0.249927 | 0.297082 | 0.263843 | 0.264297 | 0.276693 | 0.309684 | 0.342554 | 0.326368 | 0.328228 | 0.460182 | 0.687638 | 0.579395 | 0.374066 | 0.284528 | 0.231573 | 0.215659 | 0.225643 | 0.243141 | 0.277143 | 0.243256 | 0.246762 | 0.276634 | 0.340946 | 0.344884 | 0.393640 | 0.342834 | 0.357850 | 0.307267 | 0.233965 | 0.216857 | 0.216470 | 0.215115 | 0.215439 | 0.222178 | 0.217352 | 0.214753 | 0.214920 | 0.215300 | 0.215230 | 0.220968 | 0.231387 | 0.237791 | 0.225217 | 0.221501 | 0.217666 | 0.214709 | 0.225067 | 0.231339 | 0.271981 | 0.284337 | 0.274571 | 0.226368 | 0.214659 | 0.214626 | 0.214768 | 0.217443 | 0.214755 | 0.226491 | 0.239084 | 0.222556 | 0.232235 | 0.219079 | 0.221953 | 0.215352 | 0.245066 | 0.302084 | 0.298732 | 0.287655 | 0.281425 | 0.311473 | 0.269131 | 0.239015 | 0.225499 | 0.228694 | 0.252680 | 0.265645 | 0.269196 | 0.256647 | 0.286525 | 0.345896 | 0.428624 | 0.403933 | 0.387923 | 0.334636 | 0.393912 | 0.386926 | 0.371100 | 0.372391 | 0.448872 | 0.615726 | 0.574833 | 0.407918 | 0.393114 | 0.335738 | 0.311136 | 0.309486 | 0.275495 | 0.284725 | 0.346987 | 0.293258 | 0.263503 | 0.241226 | 0.233207 | 0.228808 | 0.237495 | 0.216931 | 0.215147 | 0.217007 | 0.228257 | 0.231261 | 0.228581 | 0.225446 | 0.263157 | 0.295632 | 0.357456 | 0.363249 | 0.345092 | 0.376509 | 0.300703 | 0.249800 | 0.226350 | 0.214761 | 0.219757 | 0.232016 | 0.302855 | 0.303999 | 0.299319 | 0.284245 | 0.254486 | 0.221409 | 0.216088 | 0.231752 | 0.240487 | 0.240813 | 0.255465 | 0.255439 | 0.238465 |
| right anterior | 0.277396 | 0.391995 | 0.635085 | 0.517229 | 0.921074 | 0.651863 | 0.315686 | 0.317151 | 0.222508 | 0.228561 | 0.214626 | 0.214827 | 0.229326 | 0.233008 | 0.305395 | 0.412387 | 0.575840 | 0.629385 | 0.387675 | 0.254246 | 0.216081 | 0.214679 | 0.263133 | 0.390914 | 0.335136 | 0.299295 | 0.261389 | 0.352356 | 0.237731 | 0.216129 | 0.214778 | 0.214772 | 0.239260 | 0.259347 | 0.227774 | 0.214930 | 0.260644 | 0.306264 | 0.274911 | 0.321392 | 0.397601 | 0.498850 | 0.431958 | 0.283066 | 0.312415 | 0.251931 | 0.236334 | 0.218062 | 0.332831 | 0.429162 | 0.267389 | 0.254840 | 0.228407 | 0.222766 | 0.215269 | 0.215053 | 0.220317 | 0.214734 | 0.219775 | 0.215859 | 0.220900 | 0.252685 | 0.336474 | 0.369894 | 0.388483 | 0.392911 | 0.330982 | 0.371335 | 0.370010 | 0.345646 | 0.347146 | 0.326219 | 0.383274 | 0.449444 | 0.368367 | 0.284596 | 0.227862 | 0.223946 | 0.222983 | 0.219956 | 0.216170 | 0.216140 | 0.234078 | 0.278636 | 0.355928 | 0.460161 | 0.644324 | 1.703408 | 3.578987 | 3.741321 | 5.362183 | 6.390346 | 9.043329 | 3.544371 | 0.931851 | 0.457849 | 0.398107 | 0.323247 | 0.269047 | 0.231885 | 0.221544 | 0.218621 | 0.216560 | 0.215719 | 0.221448 | 0.245177 | 0.224705 | 0.215376 | 0.234722 | 0.267699 | 0.261876 | 0.233770 | 0.238387 | 0.214813 | 0.234990 | 0.250767 | 0.289413 | 0.294977 | 0.229698 | 0.217907 | 0.241985 | 0.309139 | 0.412212 | 0.813716 | 1.387651 | 1.718170 | 1.272553 | 0.825105 | 0.698555 | 0.337897 | 0.250444 | 0.219039 | 0.216289 | 0.218790 | 0.216109 | 0.217822 | 0.216075 | 0.220221 | 0.216529 | 0.217845 | 0.214714 | 0.226120 | 0.227017 | 0.286993 | 0.482115 | 0.684859 | 2.100879 | 1.867920 | 2.514722 | 2.416638 | 1.206441 | 0.599343 | 0.475762 | 0.256213 | 0.249432 | 0.216997 | 0.286961 | 0.343161 | 0.242679 | 0.214626 | 0.219810 | 0.219081 | 0.255088 | 0.272568 | 0.244591 | 0.214813 | 0.235428 | 0.247999 | 0.283856 | 0.350801 | 0.392629 | 0.354573 | 0.289484 | 0.249032 | 0.277795 | 0.277259 | 0.248467 | 0.216575 | 0.218020 | 0.225511 | 0.222301 | 0.222699 | 0.257113 | 0.252454 | 0.228607 | 0.221863 | 0.233564 | 0.222472 | 0.215230 | 0.229873 | 0.261264 | 0.260105 | 0.229682 | 0.220596 | 0.229351 | 0.255708 | 0.258103 | 0.254935 | 0.254962 | 0.298295 | 0.343414 | 0.260405 | 0.231827 | 0.215821 | 0.216914 | 0.215573 | 0.233965 | 0.262945 | 0.256176 | 0.256598 | 0.242574 | 0.227132 | 0.237931 | 0.232264 | 0.246405 | 0.227064 | 0.246456 | 0.268545 | 0.252892 | 0.224912 | 0.219210 | 0.214626 | 0.215938 | 0.230737 | 0.279894 | 0.266818 | 0.243028 | 0.258531 | 0.278151 | 0.279357 | 0.265127 | 0.251683 | 0.256111 | 0.266253 | 0.240816 | 0.257264 | 0.253042 | 0.338817 | 0.443761 | 0.666772 | 0.961921 | 1.570626 | 0.971369 | 0.668079 | 0.391306 | 0.291180 | 0.256574 | 0.233433 | 0.230492 | 0.219212 | 0.221492 | 0.218817 | 0.219252 | 0.216160 | 0.215179 | 0.214773 | 0.216364 | 0.225647 | 0.232263 | 0.232787 | 0.222993 | 0.234943 | 0.244022 | 0.265047 | 0.225790 | 0.215372 | 0.217165 | 0.232267 | 0.222484 | 0.215669 | 0.215810 | 0.214652 | 0.214735 | 0.214626 | 0.215601 | 0.234096 | 0.232745 | 0.225631 | 0.219242 | 0.236912 |
| left central | 0.218013 | 0.217260 | 0.214675 | 0.241403 | 0.350455 | 0.618267 | 2.086566 | 2.997039 | 2.094253 | 1.492313 | 0.611787 | 0.314528 | 0.234425 | 0.215423 | 0.229447 | 0.214734 | 0.227811 | 0.230510 | 0.214809 | 0.220471 | 0.218592 | 0.217195 | 0.224789 | 0.277830 | 0.357563 | 0.219209 | 0.222051 | 0.260688 | 0.432181 | 0.534455 | 0.554438 | 0.730668 | 0.367259 | 0.225581 | 0.215067 | 0.220980 | 0.214669 | 0.219697 | 0.222765 | 0.289699 | 0.494732 | 0.556248 | 0.782587 | 0.864025 | 1.064923 | 2.785846 | 2.054500 | 1.526855 | 0.864151 | 0.384128 | 0.279302 | 0.220382 | 0.235328 | 0.305646 | 0.463765 | 0.746975 | 0.779168 | 0.765869 | 0.296407 | 0.215524 | 0.215683 | 0.218475 | 0.277596 | 0.428617 | 0.670742 | 1.154848 | 1.214455 | 0.999346 | 0.804982 | 0.454220 | 0.326998 | 0.300149 | 0.279475 | 0.318596 | 0.515428 | 1.113776 | 3.073734 | 12.521691 | 61.696398 | 111.131095 | 251.587861 | 95.805903 | 20.898948 | 16.975831 | 8.102134 | 3.648601 | 2.567959 | 1.186327 | 0.879639 | 0.797635 | 0.832379 | 0.963319 | 1.099820 | 1.231051 | 1.980160 | 1.613212 | 1.877508 | 1.432310 | 1.036998 | 1.145234 | 1.097612 | 0.979754 | 0.830501 | 0.874766 | 0.722032 | 0.712356 | 0.411487 | 0.331046 | 0.267459 | 0.239220 | 0.220027 | 0.214780 | 0.215273 | 0.214939 | 0.221419 | 0.214626 | 0.217696 | 0.218397 | 0.232904 | 0.237992 | 0.245034 | 0.280512 | 0.257536 | 0.250054 | 0.257126 | 0.259760 | 0.241576 | 0.234148 | 0.225223 | 0.223513 | 0.247463 | 0.247694 | 0.249494 | 0.243797 | 0.267764 | 0.255617 | 0.275277 | 0.223918 | 0.226876 | 0.215254 | 0.214626 | 0.216237 | 0.214688 | 0.230717 | 0.274329 | 0.353975 | 0.372239 | 0.317545 | 0.308764 | 0.299484 | 0.238828 | 0.220793 | 0.239943 | 0.269341 | 0.249401 | 0.261475 | 0.229875 | 0.215723 | 0.231895 | 0.284494 | 0.310642 | 0.325351 | 0.397798 | 0.318261 | 0.236868 | 0.214723 | 0.221107 | 0.215666 | 0.219809 | 0.238623 | 0.225292 | 0.214718 | 0.218281 | 0.215600 | 0.225848 | 0.243331 | 0.249625 | 0.300350 | 0.329339 | 0.346195 | 0.286619 | 0.288150 | 0.236750 | 0.225671 | 0.215308 | 0.221838 | 0.217998 | 0.214640 | 0.235661 | 0.264588 | 0.260422 | 0.276038 | 0.325929 | 0.371183 | 0.411949 | 0.300033 | 0.243525 | 0.240634 | 0.218621 | 0.228197 | 0.225943 | 0.222380 | 0.220596 | 0.217368 | 0.215513 | 0.215828 | 0.215935 | 0.219318 | 0.226798 | 0.215982 | 0.214626 | 0.224668 | 0.224987 | 0.235315 | 0.252743 | 0.256445 | 0.232690 | 0.238091 | 0.251016 | 0.310362 | 0.449398 | 0.506772 | 0.573615 | 0.578910 | 0.440250 | 0.313849 | 0.247641 | 0.216783 | 0.215798 | 0.221412 | 0.219470 | 0.214626 | 0.218284 | 0.221890 | 0.223677 | 0.238015 | 0.273215 | 0.334044 | 0.347029 | 0.376535 | 0.450136 | 0.567615 | 0.545940 | 0.472934 | 0.436982 | 0.415657 | 0.428641 | 0.422452 | 0.396477 | 0.559385 | 0.834480 | 1.025818 | 1.374804 | 1.386500 | 2.618116 | 4.149420 | 3.126980 | 2.401244 | 1.271988 | 0.978057 | 0.885945 | 0.522674 | 0.415324 | 0.405973 | 0.336958 | 0.392030 | 0.444425 | 0.445464 | 0.451532 | 0.449747 | 0.493102 | 0.757706 | 1.016748 | 1.745049 | 3.235690 | 5.268106 | 7.416736 | 8.066188 | 8.021248 | 5.058271 |
| right central | 1.330135 | 1.056706 | 1.471093 | 1.325900 | 2.696650 | 3.054697 | 1.024458 | 0.662190 | 0.685491 | 0.391777 | 0.322295 | 0.223859 | 0.215863 | 0.269637 | 0.290525 | 0.243644 | 0.322503 | 0.429129 | 0.606554 | 0.428738 | 0.268462 | 0.242283 | 0.308053 | 0.273658 | 0.215144 | 0.214695 | 0.247397 | 0.271729 | 0.254002 | 0.214791 | 0.215548 | 0.214941 | 0.214987 | 0.214673 | 0.229269 | 0.233594 | 0.310225 | 0.334121 | 0.289816 | 0.307531 | 0.335212 | 0.449068 | 0.267442 | 0.214656 | 0.215919 | 0.216103 | 0.214794 | 0.214788 | 0.214818 | 0.214651 | 0.218241 | 0.215467 | 0.216890 | 0.229730 | 0.215437 | 0.221468 | 0.224099 | 0.224989 | 0.222798 | 0.219437 | 0.215588 | 0.268130 | 0.350120 | 0.514779 | 0.420131 | 0.343332 | 0.316076 | 0.281963 | 0.218796 | 0.214849 | 0.223738 | 0.215920 | 0.260482 | 0.276473 | 0.321546 | 0.380233 | 0.580205 | 0.924954 | 0.823685 | 0.396234 | 0.367936 | 0.320636 | 0.482288 | 0.497300 | 0.412875 | 0.368892 | 0.558335 | 1.135191 | 2.216780 | 1.316284 | 1.022600 | 0.715635 | 0.633503 | 0.534639 | 0.352147 | 0.259061 | 0.231552 | 0.219249 | 0.217830 | 0.223689 | 0.231768 | 0.229113 | 0.231791 | 0.254954 | 0.273433 | 0.258376 | 0.222736 | 0.221150 | 0.226072 | 0.271848 | 0.256941 | 0.242514 | 0.269001 | 0.319619 | 0.305594 | 0.321477 | 0.281865 | 0.272932 | 0.333916 | 0.383248 | 0.380290 | 0.356012 | 0.341127 | 0.310818 | 0.386574 | 0.359213 | 0.324753 | 0.349533 | 0.469368 | 0.694178 | 1.615747 | 2.469581 | 3.387958 | 4.486075 | 8.451936 | 7.680401 | 9.699637 | 8.806545 | 6.458099 | 4.252234 | 2.831292 | 1.821105 | 1.023757 | 0.759555 | 0.441718 | 0.309834 | 0.240398 | 0.223342 | 0.219034 | 0.227950 | 0.236835 | 0.277363 | 0.260535 | 0.268800 | 0.258334 | 0.272997 | 0.254281 | 0.225087 | 0.216201 | 0.234397 | 0.238195 | 0.269611 | 0.262272 | 0.232467 | 0.228640 | 0.234815 | 0.215069 | 0.214626 | 0.216930 | 0.216880 | 0.229801 | 0.295878 | 0.344761 | 0.412429 | 0.362970 | 0.329391 | 0.286900 | 0.240715 | 0.217680 | 0.214706 | 0.236669 | 0.216340 | 0.214626 | 0.215555 | 0.232912 | 0.253577 | 0.314380 | 0.252444 | 0.253440 | 0.248965 | 0.222751 | 0.227457 | 0.258309 | 0.343895 | 0.281728 | 0.214626 | 0.228227 | 0.226793 | 0.224640 | 0.217445 | 0.215141 | 0.215118 | 0.220454 | 0.220975 | 0.237199 | 0.228439 | 0.241718 | 0.234888 | 0.254323 | 0.289651 | 0.359398 | 0.406941 | 0.445928 | 0.407587 | 0.435098 | 0.571118 | 0.831791 | 0.815562 | 0.918444 | 1.135414 | 1.096659 | 0.881562 | 0.614855 | 0.452940 | 0.375879 | 0.333456 | 0.302429 | 0.302001 | 0.311531 | 0.355611 | 0.378147 | 0.321480 | 0.283910 | 0.285656 | 0.288854 | 0.325067 | 0.398223 | 0.425765 | 0.441668 | 0.440347 | 0.392825 | 0.350925 | 0.309546 | 0.244655 | 0.226008 | 0.231372 | 0.231573 | 0.244444 | 0.252789 | 0.273981 | 0.308041 | 0.322790 | 0.348078 | 0.490831 | 0.446859 | 0.495600 | 0.409919 | 0.323953 | 0.303854 | 0.271687 | 0.246153 | 0.230514 | 0.215539 | 0.215531 | 0.214644 | 0.215240 | 0.222191 | 0.216777 | 0.215439 | 0.214850 | 0.222880 | 0.243306 | 0.254220 | 0.230023 | 0.217967 | 0.216981 | 0.217572 | 0.232490 | 0.297656 | 0.532458 |
| left posterior | 0.978363 | 0.569876 | 0.270882 | 0.286999 | 0.253412 | 0.214950 | 0.238592 | 0.302020 | 0.258284 | 0.215203 | 0.217467 | 0.268992 | 0.242837 | 0.217739 | 0.221071 | 0.321539 | 0.432416 | 0.465499 | 0.460483 | 0.405783 | 0.338468 | 0.245804 | 0.223582 | 0.406059 | 0.809761 | 3.439768 | 4.089492 | 2.963502 | 5.298865 | 5.483978 | 1.739179 | 0.597087 | 0.360976 | 0.247712 | 0.217756 | 0.355103 | 0.775346 | 1.515482 | 1.331401 | 0.899760 | 0.393147 | 0.216230 | 0.244541 | 0.267269 | 0.353331 | 0.414321 | 0.390957 | 0.404983 | 0.310365 | 0.339542 | 0.434780 | 0.532096 | 0.842820 | 0.918555 | 1.065071 | 1.781934 | 1.146334 | 0.841546 | 0.924671 | 0.743727 | 1.018022 | 0.798937 | 0.396991 | 0.329899 | 0.258848 | 0.215111 | 0.215639 | 0.217782 | 0.214722 | 0.222655 | 0.249850 | 0.257559 | 0.286278 | 0.258603 | 0.243997 | 0.235180 | 0.220573 | 0.214626 | 0.217217 | 0.275418 | 0.335263 | 0.312053 | 0.297097 | 0.314032 | 0.317813 | 0.277321 | 0.220656 | 0.215525 | 0.214694 | 0.214783 | 0.225418 | 0.281654 | 0.548107 | 1.056323 | 0.978692 | 1.079291 | 1.199783 | 0.926783 | 0.740752 | 0.427623 | 0.292401 | 0.291950 | 0.257126 | 0.247197 | 0.239698 | 0.234033 | 0.239572 | 0.232258 | 0.216558 | 0.224110 | 0.214875 | 0.223851 | 0.278178 | 0.502239 | 0.466818 | 0.435621 | 0.536116 | 0.408022 | 0.364624 | 0.313092 | 0.240755 | 0.231298 | 0.224544 | 0.214645 | 0.214648 | 0.228996 | 0.231159 | 0.221761 | 0.215046 | 0.223279 | 0.253282 | 0.258521 | 0.323988 | 0.387562 | 0.381047 | 0.283306 | 0.217282 | 0.222484 | 0.267702 | 0.452641 | 0.605181 | 0.682858 | 0.594897 | 0.445751 | 0.358637 | 0.329505 | 0.316831 | 0.362640 | 0.352869 | 0.369680 | 0.404473 | 0.387815 | 0.376421 | 0.339633 | 0.281426 | 0.274056 | 0.241849 | 0.219544 | 0.222988 | 0.216808 | 0.214645 | 0.216526 | 0.224023 | 0.237079 | 0.268098 | 0.299608 | 0.334967 | 0.417035 | 0.439800 | 0.452192 | 0.491543 | 0.573075 | 0.622057 | 0.597221 | 0.484064 | 0.378593 | 0.302631 | 0.316619 | 0.361584 | 0.378622 | 0.378714 | 0.368921 | 0.328086 | 0.277440 | 0.221573 | 0.218000 | 0.258422 | 0.243526 | 0.220083 | 0.215334 | 0.225702 | 0.246191 | 0.252784 | 0.275437 | 0.267419 | 0.257381 | 0.240263 | 0.250522 | 0.276365 | 0.301776 | 0.329635 | 0.379911 | 0.449946 | 0.580959 | 0.760217 | 0.905530 | 1.274386 | 1.868480 | 2.492490 | 2.959234 | 2.189261 | 1.472630 | 0.982718 | 0.967671 | 0.935496 | 0.660185 | 0.816341 | 1.115155 | 1.202919 | 1.971438 | 2.400788 | 3.391625 | 5.120635 | 3.597813 | 2.694899 | 3.266642 | 3.117618 | 2.964271 | 3.019097 | 3.479878 | 3.540679 | 1.508115 | 1.081557 | 0.690734 | 0.403193 | 0.259720 | 0.229904 | 0.221696 | 0.219281 | 0.214752 | 0.215539 | 0.214846 | 0.214962 | 0.220292 | 0.225555 | 0.216980 | 0.222237 | 0.232271 | 0.271168 | 0.347851 | 0.518729 | 0.642705 | 0.651587 | 0.535742 | 0.441091 | 0.348425 | 0.302726 | 0.274432 | 0.272274 | 0.275857 | 0.251935 | 0.231622 | 0.225128 | 0.225766 | 0.224140 | 0.218781 | 0.223532 | 0.243327 | 0.286506 | 0.355324 | 0.354485 | 0.350571 | 0.336181 | 0.305031 | 0.273620 | 0.244338 | 0.229017 | 0.226070 | 0.227666 | 0.239731 |
| right posterior | 0.214698 | 0.215982 | 0.216705 | 0.223505 | 0.266719 | 0.317445 | 0.277580 | 0.221419 | 0.216366 | 0.215319 | 0.220787 | 0.215655 | 0.226659 | 0.233844 | 0.216275 | 0.218853 | 0.262254 | 0.412464 | 0.308614 | 0.244806 | 0.218203 | 0.230862 | 0.225957 | 0.219716 | 0.223349 | 0.230530 | 0.234436 | 0.218214 | 0.218368 | 0.215663 | 0.215896 | 0.236528 | 0.291952 | 0.426776 | 0.706708 | 0.444392 | 0.384097 | 0.770165 | 0.647704 | 0.870018 | 0.576413 | 0.336702 | 0.300391 | 0.311391 | 0.254897 | 0.260941 | 0.250667 | 0.236515 | 0.230369 | 0.323393 | 0.361427 | 0.413932 | 0.402403 | 0.323187 | 0.301824 | 0.344431 | 0.312177 | 0.284767 | 0.270350 | 0.278301 | 0.291126 | 0.300439 | 0.295172 | 0.255839 | 0.236837 | 0.255656 | 0.267738 | 0.273908 | 0.308867 | 0.349008 | 0.485783 | 0.823152 | 1.614831 | 1.993047 | 3.298611 | 4.105152 | 4.473925 | 2.567781 | 2.059322 | 2.035408 | 2.525823 | 1.957340 | 1.744384 | 2.810768 | 4.198969 | 3.705494 | 1.440190 | 0.962517 | 0.706974 | 1.058559 | 1.263201 | 0.791107 | 0.774568 | 1.113795 | 0.879246 | 0.868073 | 0.503173 | 0.406540 | 0.630345 | 0.735031 | 0.737133 | 1.038086 | 0.961303 | 1.393389 | 0.999726 | 0.666225 | 0.546585 | 0.428399 | 0.359606 | 0.356286 | 0.287289 | 0.248323 | 0.222091 | 0.214841 | 0.221853 | 0.240255 | 0.252182 | 0.215158 | 0.224409 | 0.249745 | 0.324923 | 0.467790 | 0.445026 | 0.365011 | 0.272141 | 0.259543 | 0.255298 | 0.248903 | 0.259893 | 0.353777 | 0.887721 | 2.702652 | 9.463358 | 28.175586 | 35.834553 | 22.976833 | 9.933571 | 2.406321 | 1.431474 | 1.183420 | 0.699345 | 0.463472 | 0.371289 | 0.317571 | 0.362458 | 0.289962 | 0.247983 | 0.235841 | 0.244103 | 0.245507 | 0.288672 | 0.382293 | 0.420064 | 0.386665 | 0.299573 | 0.283809 | 0.264149 | 0.242834 | 0.222337 | 0.227759 | 0.238853 | 0.296329 | 0.272641 | 0.264298 | 0.241078 | 0.237848 | 0.238370 | 0.236714 | 0.224022 | 0.230315 | 0.239622 | 0.272447 | 0.284585 | 0.266351 | 0.249362 | 0.257342 | 0.259946 | 0.246068 | 0.250665 | 0.235408 | 0.235241 | 0.234791 | 0.236543 | 0.229358 | 0.225461 | 0.221703 | 0.230997 | 0.230741 | 0.224673 | 0.216507 | 0.214752 | 0.215173 | 0.216230 | 0.217120 | 0.218528 | 0.228502 | 0.231475 | 0.251440 | 0.270618 | 0.253847 | 0.250305 | 0.232640 | 0.219472 | 0.236793 | 0.229631 | 0.225347 | 0.226468 | 0.219984 | 0.229872 | 0.238587 | 0.229884 | 0.228930 | 0.222283 | 0.225311 | 0.222195 | 0.214626 | 0.214990 | 0.215765 | 0.215329 | 0.216690 | 0.219468 | 0.226965 | 0.240822 | 0.231052 | 0.225564 | 0.229555 | 0.222614 | 0.214731 | 0.217050 | 0.219237 | 0.219862 | 0.218855 | 0.222610 | 0.215309 | 0.217495 | 0.226611 | 0.234228 | 0.247151 | 0.241582 | 0.251513 | 0.231872 | 0.226023 | 0.223581 | 0.224082 | 0.217375 | 0.214863 | 0.215234 | 0.217185 | 0.217181 | 0.223714 | 0.245868 | 0.277667 | 0.282230 | 0.282982 | 0.252797 | 0.304108 | 0.313914 | 0.283643 | 0.252715 | 0.237549 | 0.251609 | 0.293059 | 0.264258 | 0.234162 | 0.232967 | 0.237452 | 0.247491 | 0.243441 | 0.238409 | 0.237623 | 0.263823 | 0.269606 | 0.254214 | 0.243100 | 0.223564 | 0.214626 | 0.216497 | 0.219444 | 0.220236 | 0.219245 |
| all electrodes | 0.222963 | 0.214855 | 0.239469 | 0.274217 | 0.263820 | 0.391191 | 0.466563 | 0.631242 | 0.522739 | 0.263837 | 0.214626 | 0.302954 | 0.892217 | 0.812062 | 0.697242 | 0.770165 | 0.371333 | 0.245652 | 0.221473 | 0.270059 | 0.283392 | 0.236977 | 0.258804 | 0.233933 | 0.214817 | 0.280363 | 0.444805 | 0.920560 | 0.460132 | 0.432656 | 0.322889 | 0.222873 | 0.217085 | 0.232218 | 0.298809 | 0.281972 | 0.311391 | 0.330314 | 0.347227 | 0.376010 | 0.537851 | 0.375944 | 0.388294 | 0.411370 | 0.585757 | 0.476978 | 0.613668 | 0.564454 | 0.786342 | 0.537041 | 0.395198 | 0.291562 | 0.225693 | 0.218292 | 0.234168 | 0.252223 | 0.266736 | 0.274408 | 0.268562 | 0.322458 | 0.470124 | 0.517689 | 0.547819 | 1.201389 | 1.439110 | 1.117453 | 1.485373 | 0.619470 | 0.519053 | 0.431796 | 0.328944 | 0.454966 | 0.523008 | 0.546714 | 0.708636 | 0.999981 | 0.795533 | 1.323394 | 1.128661 | 1.585535 | 5.732628 | 15.392451 | 17.256421 | 30.201165 | 24.005333 | 7.692990 | 2.072643 | 0.649283 | 0.523594 | 0.597512 | 0.649676 | 0.998314 | 3.613930 | 23.050378 | 79.612185 | 106.731209 | 40.398408 | 30.745135 | 27.761529 | 17.649738 | 11.626888 | 8.734539 | 4.473206 | 3.036182 | 4.358347 | 4.818544 | 4.132876 | 1.947219 | 1.646104 | 1.840550 | 2.469162 | 2.532282 | 1.411813 | 0.854868 | 0.665606 | 0.624804 | 0.892364 | 1.171069 | 0.795380 | 1.163090 | 0.809265 | 0.923978 | 0.932135 | 1.094735 | 0.694299 | 0.617481 | 0.419575 | 0.492387 | 0.488617 | 0.876112 | 0.700741 | 0.653306 | 0.702882 | 1.139690 | 0.776126 | 0.653131 | 0.492255 | 0.384342 | 0.337161 | 0.407269 | 0.379487 | 0.325595 | 0.329008 | 0.243775 | 0.235078 | 0.222006 | 0.218673 | 0.220454 | 0.214800 | 0.216072 | 0.215664 | 0.218134 | 0.235521 | 0.290100 | 0.419675 | 0.726285 | 2.995292 | 5.130882 | 4.036062 | 3.146674 | 2.500840 | 1.190572 | 0.652127 | 0.520751 | 1.010212 | 2.536457 | 5.420841 | 5.171626 | 4.796541 | 6.565370 | 6.771421 | 4.533840 | 2.576510 | 1.748043 | 1.641850 | 1.209969 | 0.671999 | 0.444131 | 0.427912 | 0.447486 | 0.583065 | 0.372561 | 0.238518 | 0.215007 | 0.216114 | 0.227209 | 0.239144 | 0.286820 | 0.279993 | 0.221410 | 0.215877 | 0.222473 | 0.225241 | 0.241780 | 0.293638 | 0.321076 | 0.267812 | 0.255082 | 0.269698 | 0.292481 | 0.247323 | 0.225310 | 0.256051 | 0.319029 | 0.517304 | 0.686035 | 1.032298 | 1.526141 | 1.981609 | 1.195199 | 1.360352 | 0.947671 | 1.183795 | 0.697082 | 0.369398 | 0.291350 | 0.332289 | 0.382292 | 0.378626 | 0.299830 | 0.296400 | 0.366392 | 0.521662 | 0.633495 | 0.660013 | 0.865645 | 1.053598 | 1.265902 | 1.252419 | 0.774978 | 0.661507 | 0.420692 | 0.355899 | 0.321182 | 0.329084 | 0.336271 | 0.417832 | 0.416515 | 0.462977 | 0.457717 | 0.486658 | 0.468011 | 0.502221 | 0.451552 | 0.391310 | 0.388642 | 0.400736 | 0.394830 | 0.310843 | 0.231073 | 0.218235 | 0.223468 | 0.228627 | 0.222959 | 0.219816 | 0.228052 | 0.231010 | 0.222126 | 0.220186 | 0.217862 | 0.215969 | 0.216049 | 0.217844 | 0.226099 | 0.229420 | 0.219119 | 0.214649 | 0.214626 | 0.214626 | 0.217780 | 0.220803 | 0.222010 | 0.226487 | 0.250611 | 0.270909 | 0.269775 | 0.259987 | 0.282842 | 0.332632 | 0.331420 |

Searchlight, spatiotemporal cluster permutation test

|  | start time | stop time | peak time | peak channel | cluster p | peak Cohen's d | direction |
| --- | --- | --- | --- | --- | --- | --- | --- |
| #1 | 140 | 335 | 190 | P6 | 0.0044 | 0.828342 | positive |

B) identity, angry

  
|  | time window | peak latency | cluster *p* | peak Cohen's *d* |  | | | |  |
| **all electrodes** | 315 - 395 ms | 355 ms | 0.0315 | 0.9274 |  | | | |  |
 535 - 645 ms | 585 ms | 0.013 | 0.917 |  | | | ||  |  | | | | | | | | |

Time-resolved classification, cluster permutation tests

|  | **left hemisphere** | | | | **right hemisphere** | | | |  |
|  | time window | peak latency | cluster *p* | peak Cohen's *d* | time window | peak latency | cluster *p* | peak Cohen's *d* |  |
| **anterior** |  | | | |  | | | |  |
| **central** |  | | | |  | | | |  |
| **posterior** | 90 - 275 ms | 115 ms | 0.0047 | 1.1277 | 80 - 135 ms | 110 ms | 0.048 | 1.0941 |  |
 300 - 635 ms | 365 ms | 0.0006 | 0.6223 | 210 - 655 ms | 495 ms | 0.0002 | 1.0302 | 935 - 1165 ms | 1060 ms | 0.0031 | 0.6993 | 700 - 810 ms | 730 ms | 0.0191 | 0.8053 |

  

Time-resolved classification, Bayesian statistics

|  | -200 | -195 | -190 | -185 | -180 | -175 | -170 | -165 | -160 | -155 | -150 | -145 | -140 | -135 | -130 | -125 | -120 | -115 | -110 | -105 | -100 | -95 | -90 | -85 | -80 | -75 | -70 | -65 | -60 | -55 | -50 | -45 | -40 | -35 | -30 | -25 | -20 | -15 | -10 | -5 | 0 | 5 | 10 | 15 | 20 | 25 | 30 | 35 | 40 | 45 | 50 | 55 | 60 | 65 | 70 | 75 | 80 | 85 | 90 | 95 | 100 | 105 | 110 | 115 | 120 | 125 | 130 | 135 | 140 | 145 | 150 | 155 | 160 | 165 | 170 | 175 | 180 | 185 | 190 | 195 | 200 | 205 | 210 | 215 | 220 | 225 | 230 | 235 | 240 | 245 | 250 | 255 | 260 | 265 | 270 | 275 | 280 | 285 | 290 | 295 | 300 | 305 | 310 | 315 | 320 | 325 | 330 | 335 | 340 | 345 | 350 | 355 | 360 | 365 | 370 | 375 | 380 | 385 | 390 | 395 | 400 | 405 | 410 | 415 | 420 | 425 | 430 | 435 | 440 | 445 | 450 | 455 | 460 | 465 | 470 | 475 | 480 | 485 | 490 | 495 | 500 | 505 | 510 | 515 | 520 | 525 | 530 | 535 | 540 | 545 | 550 | 555 | 560 | 565 | 570 | 575 | 580 | 585 | 590 | 595 | 600 | 605 | 610 | 615 | 620 | 625 | 630 | 635 | 640 | 645 | 650 | 655 | 660 | 665 | 670 | 675 | 680 | 685 | 690 | 695 | 700 | 705 | 710 | 715 | 720 | 725 | 730 | 735 | 740 | 745 | 750 | 755 | 760 | 765 | 770 | 775 | 780 | 785 | 790 | 795 | 800 | 805 | 810 | 815 | 820 | 825 | 830 | 835 | 840 | 845 | 850 | 855 | 860 | 865 | 870 | 875 | 880 | 885 | 890 | 895 | 900 | 905 | 910 | 915 | 920 | 925 | 930 | 935 | 940 | 945 | 950 | 955 | 960 | 965 | 970 | 975 | 980 | 985 | 990 | 995 | 1000 | 1005 | 1010 | 1015 | 1020 | 1025 | 1030 | 1035 | 1040 | 1045 | 1050 | 1055 | 1060 | 1065 | 1070 | 1075 | 1080 | 1085 | 1090 | 1095 | 1100 | 1105 | 1110 | 1115 | 1120 | 1125 | 1130 | 1135 | 1140 | 1145 | 1150 | 1155 | 1160 | 1165 | 1170 | 1175 | 1180 | 1185 | 1190 | 1195 |
| --- | --- | --- | --- | --- | --- | --- | --- | --- | --- | --- | --- | --- | --- | --- | --- | --- | --- | --- | --- | --- | --- | --- | --- | --- | --- | --- | --- | --- | --- | --- | --- | --- | --- | --- | --- | --- | --- | --- | --- | --- | --- | --- | --- | --- | --- | --- | --- | --- | --- | --- | --- | --- | --- | --- | --- | --- | --- | --- | --- | --- | --- | --- | --- | --- | --- | --- | --- | --- | --- | --- | --- | --- | --- | --- | --- | --- | --- | --- | --- | --- | --- | --- | --- | --- | --- | --- | --- | --- | --- | --- | --- | --- | --- | --- | --- | --- | --- | --- | --- | --- | --- | --- | --- | --- | --- | --- | --- | --- | --- | --- | --- | --- | --- | --- | --- | --- | --- | --- | --- | --- | --- | --- | --- | --- | --- | --- | --- | --- | --- | --- | --- | --- | --- | --- | --- | --- | --- | --- | --- | --- | --- | --- | --- | --- | --- | --- | --- | --- | --- | --- | --- | --- | --- | --- | --- | --- | --- | --- | --- | --- | --- | --- | --- | --- | --- | --- | --- | --- | --- | --- | --- | --- | --- | --- | --- | --- | --- | --- | --- | --- | --- | --- | --- | --- | --- | --- | --- | --- | --- | --- | --- | --- | --- | --- | --- | --- | --- | --- | --- | --- | --- | --- | --- | --- | --- | --- | --- | --- | --- | --- | --- | --- | --- | --- | --- | --- | --- | --- | --- | --- | --- | --- | --- | --- | --- | --- | --- | --- | --- | --- | --- | --- | --- | --- | --- | --- | --- | --- | --- | --- | --- | --- | --- | --- | --- | --- | --- | --- | --- | --- | --- | --- | --- | --- | --- | --- | --- | --- | --- | --- | --- | --- | --- | --- | --- | --- | --- | --- | --- | --- | --- | --- | --- | --- | --- | --- | --- | --- | --- | --- |
| left anterior | 0.229437 | 0.216338 | 0.232380 | 0.263551 | 0.388978 | 0.411419 | 0.268143 | 0.215596 | 0.281352 | 0.523460 | 2.982280 | 17.658454 | 4.832480 | 0.660401 | 0.256968 | 0.221994 | 0.240449 | 0.241639 | 0.222383 | 0.261424 | 0.282481 | 0.232934 | 0.271729 | 0.310649 | 0.221895 | 0.253102 | 0.235839 | 0.230821 | 0.228016 | 0.388237 | 0.576836 | 0.505785 | 0.338765 | 0.280606 | 0.218352 | 0.214846 | 0.276155 | 0.420422 | 0.500070 | 0.842766 | 1.221258 | 1.098258 | 0.813173 | 0.479028 | 0.244661 | 0.218526 | 0.231235 | 0.414141 | 0.723091 | 0.848686 | 0.797660 | 0.472601 | 0.644409 | 1.121464 | 0.742275 | 0.463527 | 0.552139 | 0.733406 | 1.031565 | 0.578606 | 0.503696 | 0.725750 | 1.939941 | 0.801222 | 0.485478 | 0.248989 | 0.223198 | 0.219649 | 0.285157 | 0.436624 | 0.258595 | 0.221929 | 0.225964 | 0.242472 | 0.232555 | 0.226106 | 0.223764 | 0.214957 | 0.219844 | 0.235095 | 0.310704 | 0.502065 | 0.615301 | 0.973597 | 2.024049 | 3.127023 | 3.422573 | 2.503280 | 2.354885 | 2.926276 | 2.392404 | 1.748631 | 0.954605 | 0.627772 | 0.453080 | 0.263725 | 0.218539 | 0.214760 | 0.214655 | 0.220251 | 0.214995 | 0.218483 | 0.224308 | 0.224427 | 0.227926 | 0.221155 | 0.214653 | 0.215032 | 0.214962 | 0.215987 | 0.223086 | 0.231904 | 0.261170 | 0.303318 | 0.294404 | 0.375347 | 0.430283 | 0.386952 | 0.315534 | 0.283811 | 0.274868 | 0.256038 | 0.228455 | 0.215655 | 0.216711 | 0.241140 | 0.258467 | 0.237596 | 0.228073 | 0.224776 | 0.229107 | 0.235212 | 0.246726 | 0.249754 | 0.270218 | 0.276499 | 0.235818 | 0.225113 | 0.223801 | 0.219281 | 0.215229 | 0.216314 | 0.216706 | 0.216273 | 0.222229 | 0.218310 | 0.221124 | 0.242681 | 0.293064 | 0.370687 | 0.416833 | 0.474096 | 0.483614 | 0.364341 | 0.275385 | 0.236235 | 0.222960 | 0.231118 | 0.229665 | 0.258741 | 0.298292 | 0.350820 | 0.468601 | 0.518631 | 0.490679 | 0.417400 | 0.310856 | 0.286980 | 0.286143 | 0.229758 | 0.222243 | 0.226982 | 0.234458 | 0.255442 | 0.264593 | 0.251824 | 0.271231 | 0.287851 | 0.292194 | 0.265885 | 0.270018 | 0.258688 | 0.239767 | 0.250487 | 0.292692 | 0.293888 | 0.281302 | 0.267979 | 0.310802 | 0.398652 | 0.383134 | 0.258059 | 0.233971 | 0.245938 | 0.282709 | 0.272501 | 0.297913 | 0.324769 | 0.448970 | 0.561565 | 0.816913 | 0.803043 | 0.940317 | 0.785948 | 0.557927 | 0.394435 | 0.292461 | 0.220401 | 0.214801 | 0.222771 | 0.226318 | 0.218525 | 0.214812 | 0.214626 | 0.214986 | 0.214872 | 0.217997 | 0.220905 | 0.218400 | 0.226143 | 0.246620 | 0.256455 | 0.248652 | 0.257637 | 0.277271 | 0.307564 | 0.286985 | 0.239650 | 0.223433 | 0.214626 | 0.219144 | 0.226562 | 0.222185 | 0.218860 | 0.224013 | 0.238783 | 0.274907 | 0.361735 | 0.520076 | 0.727567 | 0.802254 | 0.741169 | 0.570048 | 0.383458 | 0.282606 | 0.238830 | 0.227362 | 0.227269 | 0.237112 | 0.246879 | 0.232777 | 0.237703 | 0.241523 | 0.225662 | 0.232239 | 0.297406 | 0.350630 | 0.402874 | 0.372028 | 0.464374 | 0.768891 | 0.697198 | 0.343570 | 0.256069 | 0.259519 | 0.369722 | 0.442959 | 0.399485 | 0.387684 | 0.504741 | 0.952434 | 1.214599 | 0.958772 | 0.869120 | 1.023097 | 1.512504 | 1.327972 | 0.941736 | 0.770309 | 0.592624 |
| right anterior | 0.307207 | 0.287607 | 0.239912 | 0.218456 | 0.291840 | 0.369358 | 0.431485 | 0.462884 | 0.450587 | 0.279913 | 0.216333 | 0.389131 | 0.605641 | 0.533849 | 0.367351 | 0.239802 | 0.214799 | 0.217296 | 0.218877 | 0.216497 | 0.214930 | 0.216588 | 0.218504 | 0.215177 | 0.221145 | 0.399531 | 0.932694 | 1.160741 | 0.542183 | 0.272268 | 0.233294 | 0.220816 | 0.229140 | 0.262978 | 0.554994 | 0.561190 | 0.842446 | 0.730142 | 0.928858 | 1.714663 | 2.952993 | 2.882201 | 3.079821 | 1.519023 | 1.076103 | 0.647865 | 0.587789 | 0.387867 | 0.243883 | 0.248913 | 0.231573 | 0.252624 | 0.283016 | 0.291797 | 0.345232 | 0.432548 | 0.784187 | 1.555771 | 0.880636 | 0.546773 | 0.361832 | 0.262734 | 0.223204 | 0.215087 | 0.239305 | 0.242648 | 0.275435 | 0.333850 | 0.547210 | 0.695839 | 1.208843 | 1.296258 | 1.788101 | 1.866824 | 1.191013 | 0.761207 | 0.468432 | 0.344786 | 0.318505 | 0.287142 | 0.316570 | 0.393252 | 0.432507 | 0.609150 | 0.849166 | 0.847209 | 0.904938 | 1.778346 | 1.519028 | 2.028003 | 1.615806 | 1.009394 | 0.694546 | 0.401153 | 0.241062 | 0.256012 | 0.224016 | 0.226250 | 0.218786 | 0.214746 | 0.219516 | 0.260744 | 0.412569 | 0.342200 | 0.362885 | 0.350347 | 0.354198 | 0.378215 | 0.231813 | 0.215401 | 0.215987 | 0.226474 | 0.214626 | 0.234059 | 0.232241 | 0.214747 | 0.220395 | 0.216066 | 0.227449 | 0.233275 | 0.262975 | 0.336677 | 0.368915 | 0.330874 | 0.357794 | 0.391760 | 0.441720 | 0.331437 | 0.284464 | 0.329041 | 0.464216 | 0.503056 | 0.404075 | 0.314318 | 0.268467 | 0.251130 | 0.228139 | 0.218036 | 0.242592 | 0.274374 | 0.309344 | 0.260539 | 0.236221 | 0.224454 | 0.221041 | 0.216678 | 0.214894 | 0.217220 | 0.220746 | 0.217462 | 0.215249 | 0.215043 | 0.222065 | 0.244612 | 0.263553 | 0.268230 | 0.277783 | 0.232299 | 0.214944 | 0.231084 | 0.251751 | 0.261792 | 0.246395 | 0.258872 | 0.242991 | 0.248755 | 0.257556 | 0.282573 | 0.317540 | 0.335651 | 0.440212 | 0.505039 | 0.443290 | 0.356254 | 0.276830 | 0.231313 | 0.220820 | 0.215536 | 0.229986 | 0.237557 | 0.222510 | 0.216907 | 0.214713 | 0.215115 | 0.214644 | 0.215070 | 0.215125 | 0.216479 | 0.220640 | 0.220342 | 0.217356 | 0.215311 | 0.216151 | 0.217649 | 0.217383 | 0.218910 | 0.223283 | 0.218749 | 0.224522 | 0.245491 | 0.248125 | 0.232044 | 0.221288 | 0.214967 | 0.214932 | 0.217076 | 0.221455 | 0.223028 | 0.214626 | 0.215477 | 0.224727 | 0.237706 | 0.248056 | 0.264860 | 0.235771 | 0.244212 | 0.229710 | 0.219475 | 0.218573 | 0.219795 | 0.214803 | 0.215798 | 0.216518 | 0.215866 | 0.215130 | 0.215806 | 0.231241 | 0.221077 | 0.217599 | 0.214959 | 0.214990 | 0.218060 | 0.223214 | 0.279690 | 0.280974 | 0.381201 | 0.474207 | 0.492575 | 0.449724 | 0.381081 | 0.347885 | 0.519415 | 0.653480 | 0.658682 | 0.453327 | 0.328716 | 0.323334 | 0.308097 | 0.254756 | 0.220572 | 0.215560 | 0.221748 | 0.275235 | 0.324156 | 0.291975 | 0.261857 | 0.270388 | 0.280284 | 0.293032 | 0.241748 | 0.233158 | 0.247497 | 0.288225 | 0.289429 | 0.270129 | 0.224787 | 0.221252 | 0.218971 | 0.215416 | 0.215000 | 0.242256 | 0.339084 | 0.337034 | 0.338346 | 0.328135 | 0.277856 | 0.240500 | 0.218160 | 0.222926 | 0.243539 |
| left central | 6.089851 | 10.860776 | 4.091044 | 2.395528 | 1.172917 | 0.542353 | 0.385978 | 0.236842 | 0.215381 | 0.228172 | 0.293071 | 0.531731 | 1.397161 | 4.298187 | 6.528855 | 93.287551 | 43.199929 | 3.327265 | 0.815899 | 0.328934 | 0.214914 | 0.231891 | 0.287196 | 0.484639 | 0.258549 | 0.271945 | 0.305907 | 0.339429 | 0.343528 | 0.434852 | 0.421603 | 1.086752 | 0.984796 | 0.544459 | 0.296102 | 0.230586 | 0.220502 | 0.214892 | 0.289887 | 0.556217 | 1.479770 | 3.477351 | 2.262358 | 2.126522 | 2.034896 | 0.740976 | 0.353980 | 0.257298 | 0.227495 | 0.226885 | 0.241805 | 0.219923 | 0.231901 | 0.272033 | 0.314726 | 0.283153 | 0.343864 | 0.455664 | 0.496048 | 0.421980 | 0.268832 | 0.227549 | 0.216183 | 0.260836 | 0.344670 | 0.309582 | 0.308926 | 0.280262 | 0.305203 | 0.302347 | 0.262336 | 0.245774 | 0.299295 | 0.411197 | 0.590223 | 0.354945 | 0.243165 | 0.225395 | 0.221787 | 0.223523 | 0.215634 | 0.215899 | 0.214647 | 0.216201 | 0.235089 | 0.299140 | 0.314267 | 0.386773 | 0.467076 | 0.598634 | 0.851808 | 0.592305 | 0.457515 | 0.438726 | 0.350264 | 0.284312 | 0.238045 | 0.215185 | 0.223082 | 0.217888 | 0.219818 | 0.219059 | 0.218814 | 0.217294 | 0.247192 | 0.217550 | 0.215469 | 0.215887 | 0.223075 | 0.223740 | 0.218425 | 0.228354 | 0.215770 | 0.214713 | 0.215120 | 0.221902 | 0.265585 | 0.285326 | 0.334452 | 0.422111 | 0.555565 | 0.646196 | 0.677849 | 0.464232 | 0.373970 | 0.346916 | 0.284928 | 0.251800 | 0.272455 | 0.315557 | 0.293222 | 0.280739 | 0.254920 | 0.294715 | 0.361184 | 0.386096 | 0.358087 | 0.504278 | 1.204822 | 2.759673 | 3.694003 | 2.962836 | 1.536624 | 1.290022 | 1.012109 | 0.717802 | 0.939452 | 1.003533 | 0.809455 | 0.697247 | 0.592643 | 0.487574 | 0.432585 | 0.280417 | 0.242094 | 0.238698 | 0.232954 | 0.244688 | 0.241503 | 0.235604 | 0.258056 | 0.302936 | 0.312567 | 0.294779 | 0.262599 | 0.288974 | 0.303553 | 0.333882 | 0.291167 | 0.320736 | 0.326394 | 0.346167 | 0.360870 | 0.440513 | 0.455671 | 0.731276 | 0.748168 | 1.148929 | 1.273448 | 0.965935 | 0.595298 | 0.575970 | 0.707130 | 1.269902 | 1.255136 | 0.924443 | 0.569100 | 0.534886 | 0.418814 | 0.256054 | 0.215516 | 0.234118 | 0.232746 | 0.225580 | 0.264357 | 0.355333 | 0.333920 | 0.313067 | 0.373830 | 0.317222 | 0.235180 | 0.214810 | 0.224992 | 0.215784 | 0.214626 | 0.215348 | 0.214626 | 0.224711 | 0.248481 | 0.273672 | 0.260159 | 0.242366 | 0.232302 | 0.233856 | 0.233637 | 0.218328 | 0.220276 | 0.227719 | 0.232868 | 0.219376 | 0.215027 | 0.217469 | 0.214819 | 0.217980 | 0.225788 | 0.221691 | 0.226395 | 0.302686 | 0.315688 | 0.316215 | 0.330945 | 0.412294 | 0.546387 | 0.453208 | 0.367037 | 0.351104 | 0.295447 | 0.276360 | 0.257917 | 0.240512 | 0.250528 | 0.240886 | 0.225117 | 0.229851 | 0.268984 | 0.295058 | 0.357777 | 0.377633 | 0.683613 | 2.148256 | 4.620967 | 5.301567 | 6.241694 | 4.510516 | 3.833510 | 2.387184 | 1.190434 | 0.750042 | 0.571640 | 0.489598 | 0.429999 | 0.381785 | 0.380837 | 0.441549 | 0.502065 | 0.571896 | 0.503009 | 0.402732 | 0.341057 | 0.308931 | 0.293548 | 0.290601 | 0.287360 | 0.318103 | 0.386127 | 0.427371 | 0.445758 | 0.430394 | 0.392378 | 0.356591 |
| right central | 0.990049 | 1.579538 | 1.466354 | 2.340684 | 1.628421 | 0.939585 | 0.526507 | 0.331415 | 0.316856 | 0.334589 | 0.243017 | 0.251803 | 0.238749 | 0.218615 | 0.239709 | 0.238918 | 0.241701 | 0.224803 | 0.231891 | 0.234776 | 0.250163 | 0.289634 | 0.269265 | 0.222840 | 0.214827 | 0.233303 | 0.309837 | 0.362427 | 0.563349 | 0.705937 | 0.546655 | 0.353114 | 0.250079 | 0.232818 | 0.258324 | 0.224891 | 0.215959 | 0.234410 | 0.272802 | 0.306277 | 0.510268 | 0.678541 | 1.411841 | 1.081608 | 0.402137 | 0.239582 | 0.215159 | 0.252703 | 0.354262 | 0.341665 | 0.246137 | 0.229230 | 0.323103 | 0.432663 | 0.825589 | 0.622513 | 0.263092 | 0.214744 | 0.252086 | 0.335142 | 0.475227 | 1.384911 | 1.320288 | 1.099286 | 0.948475 | 1.425220 | 1.674613 | 4.887900 | 3.434902 | 7.515371 | 15.700523 | 32.463021 | 13.027225 | 10.092219 | 2.371087 | 1.535365 | 0.670030 | 0.486821 | 0.488560 | 0.499024 | 0.439797 | 0.395901 | 0.493156 | 0.540411 | 0.564752 | 0.306348 | 0.224931 | 0.250797 | 0.400628 | 0.721109 | 1.236828 | 1.640293 | 2.488398 | 2.925966 | 1.215810 | 0.640021 | 0.347464 | 0.367034 | 0.436655 | 0.461180 | 0.573404 | 0.765900 | 1.775129 | 5.995647 | 14.333002 | 10.285928 | 6.072158 | 1.334022 | 0.825661 | 0.435743 | 0.289834 | 0.236618 | 0.226438 | 0.227919 | 0.268207 | 0.328093 | 0.334374 | 0.314091 | 0.282028 | 0.277002 | 0.264997 | 0.259089 | 0.262199 | 0.294139 | 0.259041 | 0.236376 | 0.233146 | 0.232384 | 0.223254 | 0.216326 | 0.216875 | 0.228852 | 0.265572 | 0.280844 | 0.332074 | 0.382497 | 0.494488 | 0.425571 | 0.488994 | 0.447815 | 0.417383 | 0.469859 | 0.506222 | 0.377941 | 0.300855 | 0.261475 | 0.222176 | 0.218297 | 0.217539 | 0.219430 | 0.215157 | 0.245632 | 0.248890 | 0.297651 | 0.343987 | 0.451120 | 0.487917 | 0.450383 | 0.349843 | 0.276025 | 0.240327 | 0.221344 | 0.233419 | 0.242146 | 0.249305 | 0.258401 | 0.325474 | 0.458637 | 0.591496 | 0.482185 | 0.554202 | 0.457254 | 0.325708 | 0.252265 | 0.234614 | 0.229485 | 0.227752 | 0.225134 | 0.238634 | 0.267172 | 0.298054 | 0.351890 | 0.415904 | 0.495893 | 0.548203 | 0.669574 | 0.732961 | 0.781370 | 0.588556 | 0.501188 | 0.467720 | 0.364922 | 0.295786 | 0.298583 | 0.290400 | 0.283601 | 0.281806 | 0.265226 | 0.256852 | 0.257953 | 0.255611 | 0.237673 | 0.221026 | 0.216829 | 0.219905 | 0.228157 | 0.237003 | 0.256186 | 0.275001 | 0.288353 | 0.295350 | 0.266975 | 0.268278 | 0.246360 | 0.229217 | 0.235496 | 0.256648 | 0.276854 | 0.345635 | 0.374739 | 0.422924 | 0.413481 | 0.365605 | 0.314468 | 0.267261 | 0.228416 | 0.215587 | 0.218022 | 0.215750 | 0.215104 | 0.226017 | 0.250227 | 0.271791 | 0.277891 | 0.359723 | 0.317607 | 0.232310 | 0.219962 | 0.245259 | 0.252809 | 0.232222 | 0.221819 | 0.215858 | 0.239080 | 0.281907 | 0.289864 | 0.270275 | 0.231912 | 0.220805 | 0.214641 | 0.216758 | 0.235558 | 0.271558 | 0.267723 | 0.241279 | 0.236092 | 0.218499 | 0.216142 | 0.217076 | 0.250740 | 0.254138 | 0.257917 | 0.245337 | 0.222373 | 0.217799 | 0.217667 | 0.215159 | 0.215206 | 0.217576 | 0.234664 | 0.242725 | 0.247991 | 0.274385 | 0.316624 | 0.478809 | 0.665178 | 0.923628 | 1.272977 | 1.497229 | 1.916963 |
| left posterior | 0.289341 | 0.402102 | 0.778578 | 0.874519 | 1.363526 | 1.752536 | 0.975794 | 0.487582 | 0.251952 | 0.226040 | 0.576676 | 2.337449 | 3.439788 | 5.116379 | 2.433663 | 0.681104 | 0.821756 | 0.854994 | 0.735885 | 0.484723 | 0.257046 | 0.223316 | 0.223990 | 0.229472 | 0.349609 | 0.489846 | 0.435771 | 0.316503 | 0.291902 | 0.241144 | 0.254256 | 0.318148 | 0.490957 | 0.709637 | 0.887100 | 0.714932 | 0.509054 | 0.325312 | 0.214669 | 0.298390 | 0.835583 | 0.941249 | 0.427450 | 0.251793 | 0.218083 | 0.241090 | 0.393126 | 0.801609 | 1.389659 | 1.018813 | 0.939083 | 0.657479 | 0.412092 | 0.369270 | 0.423568 | 0.398905 | 0.633759 | 1.110411 | 7.069193 | 56.947475 | 171.358716 | 582.211758 | 1675.317896 | 1731.187128 | 1914.837407 | 1206.037291 | 184.358702 | 77.688805 | 48.969953 | 48.361589 | 16.936335 | 4.911539 | 1.907324 | 1.467287 | 2.142537 | 2.664632 | 2.796066 | 4.035619 | 7.227885 | 11.129636 | 17.928294 | 22.438108 | 28.192835 | 31.984810 | 16.091517 | 15.197010 | 7.583841 | 3.771635 | 1.646104 | 2.313232 | 3.442313 | 3.757514 | 2.000536 | 2.866887 | 2.859619 | 1.632226 | 1.031023 | 0.839936 | 0.849824 | 1.036565 | 1.449340 | 1.857823 | 2.850259 | 4.209627 | 3.067741 | 2.302732 | 2.276106 | 2.160366 | 3.265590 | 3.892855 | 6.091120 | 6.481895 | 6.807313 | 7.788494 | 8.462317 | 6.554476 | 7.016001 | 8.044027 | 12.288996 | 16.846514 | 9.361841 | 10.083569 | 14.297458 | 14.083690 | 7.533338 | 4.000969 | 3.305765 | 3.616782 | 3.188391 | 2.039751 | 2.014311 | 4.337686 | 4.574186 | 3.827153 | 5.272416 | 4.762000 | 6.301267 | 5.523624 | 3.929500 | 4.573623 | 3.987410 | 2.387598 | 2.295665 | 2.063031 | 1.789020 | 2.317292 | 3.994375 | 5.280126 | 6.236271 | 6.459263 | 6.061859 | 5.132683 | 2.466404 | 1.676226 | 2.146271 | 3.634331 | 3.634938 | 3.037131 | 5.339979 | 13.898349 | 13.660136 | 7.715448 | 4.201584 | 3.646315 | 2.543015 | 2.524631 | 1.840411 | 1.471326 | 1.204435 | 1.197397 | 0.996100 | 1.281783 | 1.222017 | 1.083599 | 1.106945 | 1.024047 | 0.759328 | 0.729871 | 0.968890 | 1.026049 | 1.240742 | 1.377744 | 2.017637 | 3.511498 | 5.716666 | 5.762387 | 8.470226 | 9.683992 | 13.060183 | 11.463048 | 6.453394 | 4.421752 | 3.918449 | 2.147054 | 1.405902 | 1.201072 | 1.438880 | 1.834566 | 3.140577 | 5.985703 | 6.691590 | 10.231432 | 10.069994 | 6.634508 | 3.540725 | 1.984411 | 1.260244 | 1.106227 | 0.800434 | 0.756179 | 0.795084 | 0.858617 | 0.736272 | 0.665666 | 0.620689 | 0.567122 | 0.472625 | 0.474996 | 0.546134 | 0.684358 | 0.733798 | 0.737393 | 0.722254 | 0.902593 | 0.923637 | 1.032298 | 1.102393 | 1.429978 | 2.126807 | 3.343290 | 6.423860 | 7.479582 | 6.390126 | 8.039452 | 9.739961 | 15.938072 | 22.074415 | 11.320822 | 7.735026 | 4.186732 | 3.900814 | 3.847016 | 3.456900 | 3.422573 | 3.700062 | 4.020547 | 10.287574 | 11.197568 | 9.423220 | 7.165597 | 7.965968 | 11.758539 | 16.925028 | 16.215385 | 21.744356 | 39.116003 | 52.646044 | 36.490702 | 24.804301 | 13.319698 | 9.172698 | 4.773119 | 2.419542 | 2.026034 | 2.592053 | 2.845886 | 3.102004 | 4.759286 | 7.819627 | 6.538583 | 5.314532 | 2.784215 | 2.399239 | 2.314959 | 1.265223 | 0.920129 | 0.900865 | 0.793361 | 0.842342 | 0.608148 |
| right posterior | 0.932383 | 0.829361 | 1.201311 | 0.707544 | 0.324386 | 0.245897 | 0.216625 | 0.216107 | 0.218496 | 0.221243 | 0.263606 | 0.412684 | 0.741876 | 0.974966 | 1.351332 | 3.113141 | 3.225754 | 1.212446 | 0.527768 | 0.248343 | 0.214889 | 0.250469 | 0.227063 | 0.219550 | 0.245680 | 0.290061 | 0.318560 | 0.367523 | 0.901397 | 0.686180 | 0.467885 | 0.396825 | 0.250746 | 0.248744 | 0.249877 | 0.225855 | 0.223183 | 0.218121 | 0.224363 | 0.244544 | 0.287755 | 0.223564 | 0.214737 | 0.245389 | 0.286807 | 0.346084 | 0.483193 | 0.370095 | 0.462881 | 0.540418 | 0.334735 | 0.238099 | 0.242101 | 0.568668 | 0.619474 | 1.049292 | 4.530927 | 12.807443 | 25.041636 | 45.376616 | 150.323391 | 818.580192 | 1200.214072 | 215.905086 | 95.412242 | 64.218231 | 17.593416 | 5.283965 | 0.941511 | 0.594058 | 0.524877 | 0.776516 | 0.703824 | 0.777547 | 0.948203 | 0.986015 | 0.753748 | 0.622565 | 0.516142 | 0.611672 | 0.658643 | 0.940049 | 1.419510 | 4.238348 | 11.096272 | 14.189399 | 12.182368 | 10.714496 | 3.709124 | 3.557288 | 6.406654 | 8.750389 | 9.005088 | 4.964154 | 3.441411 | 9.927686 | 20.669747 | 32.256717 | 22.111844 | 11.519710 | 10.406804 | 9.481571 | 5.740273 | 3.936493 | 3.539529 | 4.419888 | 5.520502 | 5.812583 | 6.013395 | 6.990546 | 6.582737 | 4.013866 | 2.889627 | 3.406379 | 4.415998 | 5.121299 | 8.666371 | 24.258629 | 53.133769 | 58.890871 | 68.217137 | 53.412511 | 54.904654 | 40.460294 | 18.554231 | 20.242958 | 58.952413 | 517.844296 | 1377.747850 | 1264.601780 | 1010.926786 | 1658.127893 | 3649.846697 | 4689.390724 | 1377.532692 | 348.614343 | 351.431010 | 845.976202 | 820.881910 | 597.170133 | 448.563482 | 517.093433 | 718.263782 | 557.909968 | 303.937810 | 217.148777 | 187.878827 | 100.819967 | 31.672777 | 15.120671 | 8.799815 | 7.140773 | 7.368130 | 5.257847 | 9.136303 | 18.250523 | 36.671940 | 49.476792 | 47.449647 | 30.037394 | 20.247305 | 12.216805 | 5.522213 | 3.171723 | 2.835774 | 2.757435 | 2.667317 | 1.509252 | 1.351919 | 1.618182 | 2.031375 | 1.930786 | 1.250173 | 0.676646 | 0.595762 | 0.502104 | 0.455736 | 0.577286 | 0.722196 | 0.784601 | 1.959800 | 8.586084 | 44.457655 | 131.231744 | 152.969274 | 100.352752 | 51.606369 | 14.911305 | 5.680446 | 3.162559 | 2.549537 | 2.224710 | 2.739372 | 3.249311 | 3.905499 | 4.982504 | 6.452453 | 5.935308 | 4.907727 | 3.393761 | 2.202693 | 2.120823 | 1.651444 | 1.119400 | 1.585829 | 2.816389 | 2.619574 | 3.770990 | 9.044067 | 18.283986 | 36.655160 | 17.859211 | 5.272240 | 2.988054 | 1.603216 | 0.779625 | 0.530730 | 0.342673 | 0.299060 | 0.294659 | 0.321974 | 0.410278 | 0.496464 | 0.611967 | 0.894277 | 1.416690 | 2.341527 | 2.779807 | 2.165807 | 2.869495 | 3.051725 | 4.483272 | 5.372744 | 4.334793 | 6.092077 | 6.771112 | 2.246963 | 1.556263 | 1.200495 | 1.206651 | 1.434209 | 1.805852 | 1.734560 | 2.010684 | 2.938781 | 3.123414 | 2.957437 | 3.023261 | 1.886529 | 2.025195 | 3.182467 | 2.466648 | 2.188400 | 1.022056 | 0.521868 | 0.545586 | 0.513130 | 0.516953 | 0.611478 | 0.608110 | 0.750196 | 0.958075 | 1.021972 | 1.079520 | 1.505958 | 1.540027 | 1.298782 | 1.131559 | 0.680517 | 0.494571 | 0.334692 | 0.253316 | 0.217978 | 0.215709 | 0.216134 | 0.216262 | 0.217628 | 0.219983 | 0.225608 | 0.218171 |
| all electrodes | 1.020984 | 1.819598 | 7.331950 | 18.095742 | 487.688095 | 526.943413 | 40.894331 | 9.612296 | 4.358779 | 1.140581 | 0.332749 | 0.238965 | 0.424200 | 0.772087 | 0.660981 | 0.363880 | 0.282884 | 0.332110 | 0.215759 | 0.243365 | 0.282230 | 0.306887 | 0.274616 | 0.390949 | 1.000324 | 0.849877 | 1.185425 | 2.277555 | 0.776156 | 0.438970 | 0.310659 | 0.435485 | 0.714938 | 0.660331 | 0.451527 | 0.462533 | 0.965023 | 1.065150 | 0.372547 | 0.216008 | 0.219216 | 0.223799 | 0.229219 | 0.223505 | 0.220320 | 0.219547 | 0.250442 | 0.305532 | 0.385944 | 0.323936 | 0.236824 | 0.218826 | 0.216859 | 0.221728 | 0.214648 | 0.217550 | 0.223157 | 0.214738 | 0.231844 | 0.289513 | 0.389738 | 0.956233 | 1.348149 | 2.218869 | 1.681444 | 1.496993 | 1.354492 | 1.243479 | 0.807228 | 1.003433 | 1.087452 | 1.296426 | 1.931715 | 1.765904 | 1.312674 | 0.750100 | 0.559750 | 0.688087 | 1.170173 | 1.697132 | 3.279987 | 2.400215 | 3.136950 | 5.354538 | 3.891930 | 3.345555 | 2.489802 | 2.247705 | 2.559251 | 2.581889 | 1.032194 | 0.755607 | 0.776135 | 0.606696 | 0.492737 | 0.380363 | 0.270689 | 0.275857 | 0.296449 | 0.491415 | 0.605506 | 0.720135 | 0.916860 | 1.451484 | 1.943803 | 2.519564 | 3.917761 | 5.499780 | 5.152561 | 15.074103 | 43.037677 | 193.675413 | 266.103443 | 134.957339 | 77.421249 | 77.928253 | 21.996211 | 5.794877 | 2.094957 | 1.473702 | 1.155952 | 0.823438 | 0.421678 | 0.358617 | 0.351459 | 0.502104 | 0.579125 | 0.448340 | 0.448867 | 0.543049 | 0.679770 | 1.074977 | 1.206184 | 1.324118 | 2.099445 | 2.178716 | 2.333600 | 1.950797 | 2.102389 | 1.748735 | 1.083824 | 1.330179 | 1.850380 | 1.725603 | 1.135935 | 0.835872 | 1.027985 | 1.738332 | 1.461438 | 1.861824 | 2.202506 | 5.624184 | 8.030272 | 7.354725 | 9.344280 | 36.993559 | 83.538771 | 172.810962 | 106.298932 | 151.675390 | 132.595826 | 73.623637 | 17.659637 | 6.831627 | 8.409050 | 7.830887 | 7.322252 | 3.704977 | 1.935721 | 1.511346 | 1.005991 | 0.584375 | 0.518099 | 0.516891 | 0.973864 | 1.453620 | 2.427146 | 3.621744 | 3.810696 | 4.155821 | 3.336640 | 1.050330 | 1.135897 | 0.788446 | 1.478690 | 2.410148 | 2.828886 | 2.213618 | 1.904537 | 1.378310 | 1.120343 | 0.443373 | 0.380006 | 0.342109 | 0.283797 | 0.246258 | 0.232034 | 0.218061 | 0.222058 | 0.241523 | 0.305945 | 0.423385 | 0.551237 | 0.700641 | 0.881744 | 0.855966 | 0.506889 | 0.333718 | 0.335703 | 0.405439 | 0.481372 | 0.526489 | 0.739624 | 1.139922 | 1.132652 | 1.053204 | 1.195516 | 0.848409 | 0.688793 | 0.406538 | 0.316315 | 0.280485 | 0.262284 | 0.261464 | 0.296883 | 0.350380 | 0.466868 | 0.455874 | 0.493021 | 0.471538 | 0.372745 | 0.297543 | 0.263425 | 0.231630 | 0.231984 | 0.245036 | 0.268273 | 0.309960 | 0.401569 | 0.539556 | 1.076162 | 1.760265 | 2.762121 | 2.907416 | 2.287652 | 1.582322 | 1.070822 | 0.822323 | 0.803041 | 0.735079 | 0.472109 | 0.418909 | 0.363616 | 0.335720 | 0.362234 | 0.370085 | 0.373462 | 0.419400 | 0.412436 | 0.349760 | 0.372217 | 0.261235 | 0.232705 | 0.217036 | 0.216738 | 0.220874 | 0.230550 | 0.238513 | 0.276178 | 0.302996 | 0.337961 | 0.397491 | 0.442558 | 0.481575 | 0.335293 | 0.257979 | 0.219651 | 0.214626 | 0.225449 | 0.242503 |

Searchlight, spatiotemporal cluster permutation test

|  | start time | stop time | peak time | peak channel | cluster p | peak Cohen's d | direction |
| --- | --- | --- | --- | --- | --- | --- | --- |
| #1 | 85 | 675 | 110 | O1 | 0.0001 | 1.128508 | positive |

C) identity, sad

  
|  | time window | peak latency | cluster *p* | peak Cohen's *d* |  | | | |
| **all electrodes** | 870 - 950 ms | 870 ms | 0.0283 | -0.4318 |  | | | |
|  | | | | | | | | |

Time-resolved classification, cluster permutation tests

|  | **left hemisphere** | | | | **right hemisphere** | | | |
|  | time window | peak latency | cluster *p* | peak Cohen's *d* | time window | peak latency | cluster *p* | peak Cohen's *d* |
| **anterior** |  | | | |  | | | |
| **central** |  | | | | 940 - 1035 ms | 1030 ms | 0.0383 | -0.425 |
| **posterior** |  | | | |  | | | |

  

Time-resolved classification, Bayesian statistics

|  | -200 | -195 | -190 | -185 | -180 | -175 | -170 | -165 | -160 | -155 | -150 | -145 | -140 | -135 | -130 | -125 | -120 | -115 | -110 | -105 | -100 | -95 | -90 | -85 | -80 | -75 | -70 | -65 | -60 | -55 | -50 | -45 | -40 | -35 | -30 | -25 | -20 | -15 | -10 | -5 | 0 | 5 | 10 | 15 | 20 | 25 | 30 | 35 | 40 | 45 | 50 | 55 | 60 | 65 | 70 | 75 | 80 | 85 | 90 | 95 | 100 | 105 | 110 | 115 | 120 | 125 | 130 | 135 | 140 | 145 | 150 | 155 | 160 | 165 | 170 | 175 | 180 | 185 | 190 | 195 | 200 | 205 | 210 | 215 | 220 | 225 | 230 | 235 | 240 | 245 | 250 | 255 | 260 | 265 | 270 | 275 | 280 | 285 | 290 | 295 | 300 | 305 | 310 | 315 | 320 | 325 | 330 | 335 | 340 | 345 | 350 | 355 | 360 | 365 | 370 | 375 | 380 | 385 | 390 | 395 | 400 | 405 | 410 | 415 | 420 | 425 | 430 | 435 | 440 | 445 | 450 | 455 | 460 | 465 | 470 | 475 | 480 | 485 | 490 | 495 | 500 | 505 | 510 | 515 | 520 | 525 | 530 | 535 | 540 | 545 | 550 | 555 | 560 | 565 | 570 | 575 | 580 | 585 | 590 | 595 | 600 | 605 | 610 | 615 | 620 | 625 | 630 | 635 | 640 | 645 | 650 | 655 | 660 | 665 | 670 | 675 | 680 | 685 | 690 | 695 | 700 | 705 | 710 | 715 | 720 | 725 | 730 | 735 | 740 | 745 | 750 | 755 | 760 | 765 | 770 | 775 | 780 | 785 | 790 | 795 | 800 | 805 | 810 | 815 | 820 | 825 | 830 | 835 | 840 | 845 | 850 | 855 | 860 | 865 | 870 | 875 | 880 | 885 | 890 | 895 | 900 | 905 | 910 | 915 | 920 | 925 | 930 | 935 | 940 | 945 | 950 | 955 | 960 | 965 | 970 | 975 | 980 | 985 | 990 | 995 | 1000 | 1005 | 1010 | 1015 | 1020 | 1025 | 1030 | 1035 | 1040 | 1045 | 1050 | 1055 | 1060 | 1065 | 1070 | 1075 | 1080 | 1085 | 1090 | 1095 | 1100 | 1105 | 1110 | 1115 | 1120 | 1125 | 1130 | 1135 | 1140 | 1145 | 1150 | 1155 | 1160 | 1165 | 1170 | 1175 | 1180 | 1185 | 1190 | 1195 |
| --- | --- | --- | --- | --- | --- | --- | --- | --- | --- | --- | --- | --- | --- | --- | --- | --- | --- | --- | --- | --- | --- | --- | --- | --- | --- | --- | --- | --- | --- | --- | --- | --- | --- | --- | --- | --- | --- | --- | --- | --- | --- | --- | --- | --- | --- | --- | --- | --- | --- | --- | --- | --- | --- | --- | --- | --- | --- | --- | --- | --- | --- | --- | --- | --- | --- | --- | --- | --- | --- | --- | --- | --- | --- | --- | --- | --- | --- | --- | --- | --- | --- | --- | --- | --- | --- | --- | --- | --- | --- | --- | --- | --- | --- | --- | --- | --- | --- | --- | --- | --- | --- | --- | --- | --- | --- | --- | --- | --- | --- | --- | --- | --- | --- | --- | --- | --- | --- | --- | --- | --- | --- | --- | --- | --- | --- | --- | --- | --- | --- | --- | --- | --- | --- | --- | --- | --- | --- | --- | --- | --- | --- | --- | --- | --- | --- | --- | --- | --- | --- | --- | --- | --- | --- | --- | --- | --- | --- | --- | --- | --- | --- | --- | --- | --- | --- | --- | --- | --- | --- | --- | --- | --- | --- | --- | --- | --- | --- | --- | --- | --- | --- | --- | --- | --- | --- | --- | --- | --- | --- | --- | --- | --- | --- | --- | --- | --- | --- | --- | --- | --- | --- | --- | --- | --- | --- | --- | --- | --- | --- | --- | --- | --- | --- | --- | --- | --- | --- | --- | --- | --- | --- | --- | --- | --- | --- | --- | --- | --- | --- | --- | --- | --- | --- | --- | --- | --- | --- | --- | --- | --- | --- | --- | --- | --- | --- | --- | --- | --- | --- | --- | --- | --- | --- | --- | --- | --- | --- | --- | --- | --- | --- | --- | --- | --- | --- | --- | --- | --- | --- | --- | --- | --- | --- | --- | --- | --- | --- | --- | --- | --- |
| left anterior | 0.268772 | 0.216051 | 0.214789 | 0.215191 | 0.215967 | 0.218416 | 0.228002 | 0.331111 | 0.715294 | 0.638344 | 0.579703 | 0.310936 | 0.376945 | 0.219008 | 0.233965 | 0.275339 | 0.353693 | 0.357390 | 0.276189 | 0.376562 | 0.317521 | 0.229526 | 0.215661 | 0.269828 | 0.415736 | 0.517600 | 0.551092 | 0.473007 | 0.252666 | 0.241630 | 0.416945 | 0.555808 | 0.761974 | 1.210927 | 1.152746 | 0.789913 | 0.676575 | 1.590285 | 3.956371 | 10.192392 | 1.152248 | 0.276055 | 0.243165 | 0.235094 | 0.215014 | 0.222943 | 0.311504 | 0.305644 | 0.237640 | 0.238062 | 0.246895 | 0.240949 | 0.217402 | 0.228470 | 0.296232 | 0.568519 | 1.725846 | 3.037994 | 2.438200 | 0.977212 | 0.523986 | 0.320392 | 0.217558 | 0.218796 | 0.242149 | 0.227035 | 0.216459 | 0.220189 | 0.238766 | 0.249559 | 0.238843 | 0.235127 | 0.254044 | 0.278718 | 0.233474 | 0.216357 | 0.217361 | 0.214657 | 0.214897 | 0.220045 | 0.215023 | 0.226669 | 0.255267 | 0.314659 | 0.514036 | 0.592859 | 0.879236 | 0.957878 | 0.675119 | 0.594467 | 0.629272 | 0.520710 | 0.500342 | 0.311387 | 0.245454 | 0.224017 | 0.214961 | 0.221012 | 0.234960 | 0.248322 | 0.240984 | 0.231588 | 0.227226 | 0.217923 | 0.222455 | 0.214858 | 0.258344 | 0.403380 | 0.406099 | 0.504031 | 1.111017 | 2.717820 | 4.127870 | 3.451330 | 1.930859 | 3.157399 | 2.270627 | 0.953029 | 0.615171 | 0.567602 | 0.473648 | 0.781196 | 0.757621 | 0.851028 | 0.933213 | 1.646575 | 1.600382 | 1.020631 | 0.491293 | 0.291405 | 0.219045 | 0.214928 | 0.232930 | 0.228690 | 0.236643 | 0.222557 | 0.214665 | 0.218102 | 0.218619 | 0.223226 | 0.221182 | 0.219323 | 0.214734 | 0.217650 | 0.215908 | 0.214654 | 0.214652 | 0.222627 | 0.242906 | 0.286780 | 0.285192 | 0.220663 | 0.216163 | 0.243392 | 0.365381 | 0.805521 | 2.234754 | 3.620111 | 3.006439 | 1.668000 | 0.596470 | 0.350244 | 0.251681 | 0.272001 | 0.268080 | 0.304411 | 0.385541 | 0.628540 | 0.793877 | 1.293131 | 1.619277 | 4.787771 | 4.633804 | 2.599175 | 2.083942 | 1.762659 | 1.869215 | 1.526770 | 1.340840 | 1.399467 | 1.522343 | 1.300807 | 1.796081 | 1.426286 | 1.169937 | 0.665432 | 0.399336 | 0.412343 | 0.439403 | 0.350919 | 0.408651 | 0.499503 | 0.714049 | 0.688204 | 0.751324 | 0.674581 | 0.657693 | 0.724381 | 0.908290 | 0.871385 | 0.826020 | 0.557997 | 0.421736 | 0.373359 | 0.304744 | 0.250685 | 0.215739 | 0.215159 | 0.219690 | 0.215186 | 0.217564 | 0.214626 | 0.216715 | 0.215150 | 0.219264 | 0.219202 | 0.216804 | 0.217209 | 0.214626 | 0.215773 | 0.234281 | 0.264821 | 0.298621 | 0.287910 | 0.257451 | 0.257767 | 0.268283 | 0.243047 | 0.217018 | 0.214722 | 0.214834 | 0.216305 | 0.214802 | 0.214813 | 0.214720 | 0.215017 | 0.215464 | 0.215339 | 0.217869 | 0.216411 | 0.224706 | 0.243917 | 0.252247 | 0.220861 | 0.215107 | 0.223376 | 0.225028 | 0.256862 | 0.355579 | 0.664089 | 0.540220 | 0.413140 | 0.297221 | 0.248726 | 0.233770 | 0.217284 | 0.216174 | 0.215192 | 0.215936 | 0.215632 | 0.214817 | 0.220117 | 0.240072 | 0.272253 | 0.366166 | 0.337098 | 0.361269 | 0.420390 | 0.565289 | 0.598418 | 0.691893 | 0.658037 | 0.888773 | 0.901225 | 0.826380 | 0.799362 | 0.776239 | 0.667777 | 0.618244 | 0.608781 |
| right anterior | 0.373460 | 0.309372 | 0.248203 | 0.268351 | 0.276782 | 0.291798 | 0.319684 | 0.922955 | 4.388855 | 11.812741 | 29.643341 | 16.131081 | 1.971540 | 1.552097 | 0.802683 | 0.411885 | 0.334480 | 0.245040 | 0.357277 | 0.838521 | 0.532610 | 0.490722 | 0.615322 | 0.601259 | 0.697135 | 0.736070 | 0.791079 | 1.510222 | 1.311319 | 0.953443 | 1.050915 | 0.624900 | 0.349667 | 0.227683 | 0.227628 | 0.244299 | 0.251265 | 0.278744 | 0.271137 | 0.246400 | 0.219463 | 0.489924 | 1.316705 | 6.651744 | 15.260275 | 11.449850 | 5.695086 | 2.785349 | 1.021973 | 0.422773 | 0.268611 | 0.220293 | 0.214993 | 0.215170 | 0.214647 | 0.214646 | 0.214826 | 0.216792 | 0.244402 | 0.262912 | 0.229846 | 0.218383 | 0.243852 | 0.343374 | 0.349370 | 0.319282 | 0.338263 | 0.513314 | 0.387191 | 0.285862 | 0.224071 | 0.214710 | 0.220003 | 0.242224 | 0.302933 | 0.327227 | 0.261371 | 0.216919 | 0.217389 | 0.214802 | 0.214626 | 0.228151 | 0.247214 | 0.245310 | 0.220498 | 0.215721 | 0.230668 | 0.257930 | 0.226020 | 0.223944 | 0.215415 | 0.231462 | 0.232922 | 0.223237 | 0.242631 | 0.237857 | 0.241712 | 0.231728 | 0.232769 | 0.234975 | 0.269337 | 0.241453 | 0.228967 | 0.250028 | 0.290685 | 0.341704 | 0.459956 | 0.532106 | 0.828037 | 2.262829 | 1.336262 | 0.528532 | 0.283590 | 0.214822 | 0.261395 | 0.303456 | 0.385744 | 0.333318 | 0.230026 | 0.214990 | 0.252555 | 0.252556 | 0.276721 | 0.256494 | 0.258913 | 0.247097 | 0.251748 | 0.215939 | 0.238773 | 0.234023 | 0.229376 | 0.242346 | 0.227105 | 0.227543 | 0.231683 | 0.218569 | 0.214995 | 0.214626 | 0.214626 | 0.215956 | 0.215422 | 0.214839 | 0.214740 | 0.217738 | 0.235123 | 0.303893 | 0.390988 | 0.287326 | 0.293824 | 0.351910 | 0.409272 | 0.383520 | 0.268000 | 0.214665 | 0.215428 | 0.218613 | 0.238890 | 0.319707 | 0.603334 | 0.604439 | 0.686915 | 0.843802 | 1.031565 | 0.884245 | 0.615777 | 0.528779 | 0.546054 | 0.504217 | 0.446933 | 0.373860 | 0.388342 | 0.511667 | 0.393020 | 0.340681 | 0.285951 | 0.244699 | 0.219351 | 0.219212 | 0.250868 | 0.260743 | 0.271936 | 0.298328 | 0.305518 | 0.277936 | 0.292933 | 0.347875 | 0.474036 | 0.856235 | 1.095962 | 1.004864 | 0.615391 | 0.328138 | 0.374386 | 0.355673 | 0.352961 | 0.250788 | 0.216517 | 0.221781 | 0.275511 | 0.239938 | 0.227612 | 0.215368 | 0.222522 | 0.253274 | 0.276110 | 0.303275 | 0.292432 | 0.297639 | 0.237782 | 0.220509 | 0.224492 | 0.249702 | 0.258283 | 0.276593 | 0.285479 | 0.481307 | 0.570379 | 0.374670 | 0.251198 | 0.221907 | 0.222845 | 0.215790 | 0.214723 | 0.217367 | 0.224395 | 0.232092 | 0.226041 | 0.224860 | 0.223633 | 0.215241 | 0.216576 | 0.220672 | 0.215346 | 0.215350 | 0.229212 | 0.232411 | 0.234752 | 0.228446 | 0.214700 | 0.215499 | 0.224999 | 0.253557 | 0.276583 | 0.283101 | 0.239842 | 0.215587 | 0.238282 | 0.312780 | 0.415835 | 0.474003 | 0.714555 | 0.702534 | 0.469101 | 0.317394 | 0.254482 | 0.233872 | 0.218092 | 0.227269 | 0.244922 | 0.255037 | 0.275698 | 0.290473 | 0.390414 | 0.552494 | 0.599320 | 0.469959 | 0.425883 | 0.410208 | 0.423841 | 0.400474 | 0.428592 | 0.431882 | 0.463986 | 0.508103 | 0.452443 | 0.483378 | 0.553821 | 0.455416 | 0.387961 | 0.422872 |
| left central | 0.400971 | 0.378609 | 0.480983 | 0.381038 | 0.310086 | 0.464011 | 0.499778 | 0.407224 | 0.250949 | 0.215376 | 0.219050 | 0.280735 | 0.349728 | 0.374066 | 0.232203 | 0.238105 | 0.247425 | 0.291670 | 0.489013 | 0.432476 | 0.451867 | 0.225879 | 0.215540 | 0.279917 | 0.489871 | 1.650730 | 9.351365 | 10.617285 | 4.364494 | 1.609819 | 0.483369 | 0.336281 | 0.231356 | 0.215192 | 0.215533 | 0.218886 | 0.230120 | 0.279305 | 0.248378 | 0.300997 | 0.380075 | 0.471449 | 0.472467 | 0.516586 | 0.602799 | 0.672290 | 0.667449 | 0.398049 | 0.249023 | 0.226536 | 0.219162 | 0.216660 | 0.218627 | 0.243658 | 0.245939 | 0.232061 | 0.226352 | 0.214626 | 0.231278 | 0.255762 | 0.266976 | 0.315459 | 0.369276 | 0.331058 | 0.220607 | 0.225499 | 0.242989 | 0.243564 | 0.246332 | 0.238301 | 0.239211 | 0.263857 | 0.281968 | 0.387752 | 0.466583 | 0.697239 | 0.709685 | 0.697911 | 0.654944 | 0.618298 | 0.480567 | 0.456088 | 0.328812 | 0.492872 | 1.064641 | 1.763712 | 2.612802 | 1.729767 | 1.926988 | 2.138015 | 1.194079 | 0.295559 | 0.223257 | 0.215052 | 0.214649 | 0.233511 | 0.231418 | 0.214987 | 0.246021 | 0.271018 | 0.334455 | 0.675839 | 1.134363 | 1.596071 | 0.830556 | 0.634165 | 0.585482 | 0.715946 | 0.549356 | 0.456760 | 0.379769 | 0.377594 | 0.327419 | 0.313976 | 0.341318 | 0.351255 | 0.324682 | 0.327552 | 0.408579 | 0.464953 | 0.431622 | 0.332085 | 0.358476 | 0.437701 | 0.431159 | 0.522109 | 0.690323 | 1.251744 | 1.857152 | 1.332951 | 1.106941 | 1.044238 | 0.714897 | 0.900150 | 1.533178 | 4.438011 | 5.708671 | 3.203893 | 2.546419 | 2.751386 | 2.720057 | 2.767682 | 3.595135 | 7.788121 | 6.803006 | 4.053951 | 3.361369 | 2.177077 | 1.658834 | 0.882096 | 0.680202 | 0.881996 | 1.297010 | 1.174030 | 1.327895 | 1.149885 | 1.175652 | 0.835427 | 0.721761 | 0.519630 | 0.504266 | 0.399476 | 0.373377 | 0.365103 | 0.411908 | 0.311826 | 0.276790 | 0.263977 | 0.288971 | 0.307556 | 0.327691 | 0.327700 | 0.479535 | 0.851290 | 1.880892 | 3.867400 | 4.713780 | 6.149814 | 4.294085 | 4.995096 | 4.664990 | 1.885127 | 1.643719 | 1.148312 | 0.977612 | 0.846027 | 0.572312 | 0.431667 | 0.317350 | 0.242872 | 0.221865 | 0.216488 | 0.216987 | 0.216753 | 0.215887 | 0.217846 | 0.233939 | 0.250241 | 0.274902 | 0.304297 | 0.396287 | 0.658016 | 1.031103 | 1.033788 | 0.989237 | 0.927458 | 1.067473 | 1.399645 | 1.385637 | 1.077276 | 1.261400 | 1.879544 | 3.318017 | 3.476765 | 2.542483 | 1.465586 | 1.608558 | 1.186217 | 0.759150 | 0.515185 | 0.456061 | 0.406695 | 0.342077 | 0.302831 | 0.284961 | 0.299857 | 0.320729 | 0.315033 | 0.343503 | 0.385031 | 0.419234 | 0.393836 | 0.299139 | 0.309389 | 0.348623 | 0.345625 | 0.363348 | 0.303041 | 0.303164 | 0.388919 | 0.334728 | 0.303806 | 0.283114 | 0.282968 | 0.298258 | 0.293201 | 0.249449 | 0.223631 | 0.214696 | 0.218852 | 0.240185 | 0.299712 | 0.396246 | 0.397505 | 0.321544 | 0.289019 | 0.246791 | 0.237375 | 0.215886 | 0.215814 | 0.225292 | 0.218456 | 0.216136 | 0.215161 | 0.216376 | 0.214756 | 0.216129 | 0.228860 | 0.243728 | 0.261682 | 0.307859 | 0.580562 | 0.959024 | 1.067703 | 1.085306 | 0.820207 | 0.763799 | 0.693201 | 0.481323 | 0.345499 |
| right central | 0.219654 | 0.253262 | 0.247702 | 0.237295 | 0.227514 | 0.240015 | 0.222087 | 0.214702 | 0.260106 | 0.298551 | 0.520305 | 1.386441 | 3.320426 | 3.018877 | 4.923499 | 5.549040 | 8.430740 | 1.695335 | 0.585347 | 0.405804 | 0.292926 | 0.258661 | 0.215788 | 0.257458 | 0.346140 | 0.458943 | 0.298694 | 0.228193 | 0.267411 | 0.481862 | 0.583220 | 0.791496 | 1.201990 | 0.668111 | 0.383817 | 0.401790 | 0.473666 | 0.797784 | 0.729248 | 0.663728 | 0.630660 | 0.539385 | 0.368363 | 0.418977 | 0.419301 | 0.536998 | 0.477306 | 0.494058 | 0.406859 | 0.272901 | 0.214992 | 0.227499 | 0.317987 | 0.916907 | 2.772982 | 8.681709 | 7.641884 | 4.643065 | 7.280666 | 8.529757 | 0.817151 | 0.222152 | 0.285495 | 0.271541 | 0.263169 | 0.271478 | 0.217170 | 0.226655 | 0.273882 | 0.316718 | 0.246242 | 0.215034 | 0.216178 | 0.215915 | 0.240270 | 0.226365 | 0.271679 | 0.300519 | 0.366732 | 0.598326 | 0.951864 | 1.062348 | 1.330090 | 0.767183 | 0.817596 | 1.028215 | 1.129854 | 1.083284 | 2.571888 | 10.331889 | 47.037655 | 45.521288 | 42.400290 | 38.956626 | 46.673112 | 32.040285 | 6.130214 | 1.296624 | 0.728899 | 0.461630 | 0.337615 | 0.225513 | 0.223659 | 0.272278 | 0.332903 | 0.398676 | 0.490778 | 0.557411 | 0.386398 | 0.297189 | 0.283187 | 0.272776 | 0.318837 | 0.364774 | 0.449572 | 0.489949 | 0.418476 | 0.356457 | 0.396165 | 0.437566 | 0.419348 | 0.478434 | 0.603181 | 1.263375 | 2.894240 | 4.110212 | 2.584590 | 3.955261 | 1.307184 | 0.586596 | 0.280178 | 0.230094 | 0.216819 | 0.220262 | 0.281771 | 0.246430 | 0.222055 | 0.215088 | 0.226087 | 0.259087 | 0.276828 | 0.450196 | 0.659895 | 0.650964 | 0.408985 | 0.295063 | 0.253405 | 0.263812 | 0.242533 | 0.219372 | 0.221262 | 0.224415 | 0.215072 | 0.222555 | 0.229810 | 0.284864 | 0.406310 | 0.767318 | 0.874134 | 0.504055 | 0.374899 | 0.281928 | 0.241337 | 0.238835 | 0.233243 | 0.254343 | 0.391045 | 0.442423 | 0.431461 | 0.322192 | 0.242149 | 0.234834 | 0.231069 | 0.231563 | 0.261236 | 0.281341 | 0.327207 | 0.394970 | 0.398347 | 0.445314 | 0.463090 | 0.433056 | 0.433256 | 0.453171 | 0.475939 | 0.536268 | 0.673665 | 0.740739 | 0.648665 | 0.758399 | 0.590545 | 0.543856 | 0.753224 | 0.633085 | 0.481286 | 0.448717 | 0.446149 | 0.451727 | 0.296437 | 0.247460 | 0.277875 | 0.322293 | 0.394616 | 0.470802 | 0.735631 | 1.662643 | 2.615394 | 1.900562 | 1.255818 | 0.795432 | 0.965841 | 0.788914 | 0.630872 | 0.896413 | 1.130113 | 1.985471 | 4.531268 | 5.086383 | 6.662339 | 5.588818 | 1.698178 | 0.991761 | 0.654553 | 0.625855 | 0.457961 | 0.365053 | 0.420330 | 0.827123 | 2.041459 | 3.015726 | 3.520148 | 3.887042 | 5.654644 | 4.329988 | 3.296472 | 1.970260 | 1.846160 | 1.635959 | 2.189338 | 2.403483 | 2.290892 | 2.132767 | 2.151906 | 2.613312 | 1.946219 | 1.802864 | 1.329347 | 1.500158 | 1.238678 | 0.573462 | 0.300592 | 0.273051 | 0.228350 | 0.215432 | 0.215783 | 0.214626 | 0.220823 | 0.245643 | 0.244541 | 0.315742 | 0.492089 | 0.691034 | 0.995712 | 1.612949 | 2.295232 | 3.196716 | 2.556774 | 1.657185 | 0.785857 | 0.372265 | 0.257119 | 0.241467 | 0.243680 | 0.237290 | 0.235995 | 0.265142 | 0.278340 | 0.298985 | 0.282220 | 0.251125 |
| left posterior | 0.271510 | 0.282835 | 0.319599 | 0.333088 | 0.324726 | 0.339460 | 0.248732 | 0.222124 | 0.236687 | 0.337411 | 0.385687 | 0.394101 | 0.483954 | 0.356752 | 0.243029 | 0.239280 | 0.273416 | 0.285466 | 0.263140 | 0.226903 | 0.320343 | 0.331223 | 0.267606 | 0.265266 | 0.266688 | 0.253037 | 0.291901 | 0.217420 | 0.375048 | 0.446062 | 0.501538 | 0.759215 | 0.668821 | 0.881383 | 0.528281 | 0.266673 | 0.243787 | 0.214667 | 0.254948 | 0.286172 | 0.235372 | 0.227875 | 0.215156 | 0.221759 | 0.402321 | 1.092440 | 1.893827 | 0.749653 | 0.619975 | 0.601781 | 0.647193 | 0.426896 | 0.398168 | 0.403882 | 0.464141 | 0.561849 | 0.518118 | 0.348288 | 0.241314 | 0.228293 | 0.463984 | 1.029215 | 1.504628 | 1.649190 | 1.826413 | 1.188973 | 0.494260 | 0.252038 | 0.232818 | 0.258998 | 0.371183 | 0.463225 | 0.521791 | 0.504952 | 0.436611 | 0.275245 | 0.253976 | 0.281056 | 0.461725 | 1.149999 | 3.274611 | 12.008009 | 53.552310 | 53.848845 | 23.691024 | 10.943912 | 6.508808 | 5.817237 | 4.336031 | 1.830329 | 0.842062 | 0.703028 | 0.633822 | 0.648881 | 0.682113 | 0.706870 | 1.078963 | 1.535514 | 3.149542 | 5.855774 | 8.853136 | 7.395317 | 7.717337 | 4.125605 | 1.515290 | 0.628893 | 0.434403 | 0.338547 | 0.367173 | 0.388789 | 0.433061 | 0.585647 | 0.864090 | 0.872019 | 0.860575 | 0.654349 | 0.440044 | 0.330493 | 0.263898 | 0.256994 | 0.272756 | 0.328885 | 0.398188 | 0.585058 | 0.878005 | 1.372631 | 1.543896 | 1.495304 | 1.865099 | 2.255171 | 2.857215 | 2.796548 | 2.311270 | 2.149842 | 2.165174 | 0.790601 | 0.544012 | 0.427720 | 0.392422 | 0.351344 | 0.320163 | 0.335001 | 0.478972 | 0.594241 | 0.506061 | 0.471249 | 0.449017 | 0.404519 | 0.324594 | 0.299619 | 0.251495 | 0.244000 | 0.227123 | 0.226764 | 0.221010 | 0.214626 | 0.226779 | 0.221969 | 0.214657 | 0.220066 | 0.219267 | 0.221477 | 0.224052 | 0.241448 | 0.222858 | 0.217004 | 0.237711 | 0.260125 | 0.241104 | 0.228753 | 0.226207 | 0.214715 | 0.245520 | 0.373304 | 0.691567 | 0.648018 | 0.734447 | 0.745467 | 0.788476 | 0.852471 | 0.835901 | 0.702063 | 0.800388 | 0.945340 | 1.181404 | 1.993539 | 1.943679 | 1.687606 | 1.117803 | 1.030949 | 1.230102 | 1.703532 | 1.792871 | 1.523277 | 1.657652 | 1.074650 | 0.418752 | 0.264624 | 0.225589 | 0.222233 | 0.222663 | 0.216518 | 0.214652 | 0.215908 | 0.219134 | 0.228204 | 0.227547 | 0.236327 | 0.246479 | 0.278533 | 0.385816 | 0.489476 | 0.507268 | 0.462428 | 0.387068 | 0.328147 | 0.262031 | 0.220956 | 0.214883 | 0.225296 | 0.244746 | 0.245922 | 0.229192 | 0.216940 | 0.214917 | 0.216061 | 0.218102 | 0.233722 | 0.224971 | 0.223093 | 0.223994 | 0.221916 | 0.234979 | 0.246426 | 0.237694 | 0.258701 | 0.275590 | 0.301418 | 0.287596 | 0.278460 | 0.288979 | 0.310605 | 0.376543 | 0.462236 | 0.486889 | 0.597211 | 0.656074 | 0.848478 | 1.242354 | 1.551994 | 1.246045 | 1.411285 | 1.519714 | 1.410592 | 1.053827 | 0.677441 | 0.614611 | 0.801078 | 0.883964 | 0.853542 | 0.979716 | 0.735221 | 0.561302 | 0.480134 | 0.512052 | 0.664665 | 0.920183 | 1.169759 | 1.694357 | 2.240225 | 2.147740 | 1.600899 | 1.489739 | 1.829051 | 1.572811 | 1.519823 | 1.542344 | 1.260939 | 1.146176 | 1.131368 |
| right posterior | 0.226511 | 0.236551 | 0.215684 | 0.214661 | 0.243213 | 0.319490 | 0.281379 | 0.239824 | 0.232498 | 0.394305 | 0.462784 | 0.352961 | 0.235221 | 0.215279 | 0.227144 | 0.220229 | 0.256254 | 0.228958 | 0.217419 | 0.398800 | 0.755066 | 1.488874 | 1.042804 | 0.752229 | 0.464277 | 0.367865 | 0.223789 | 0.216296 | 0.214727 | 0.221939 | 0.239068 | 0.221485 | 0.216343 | 0.246526 | 0.225827 | 0.227000 | 0.215216 | 0.237864 | 0.243073 | 0.241060 | 0.309456 | 0.316665 | 0.414987 | 0.433225 | 0.338096 | 0.271538 | 0.299439 | 0.275431 | 0.227174 | 0.217836 | 0.222912 | 0.228831 | 0.267220 | 0.258204 | 0.271496 | 0.369745 | 0.527706 | 0.819985 | 1.646964 | 4.893546 | 12.717892 | 21.157555 | 42.795667 | 50.907177 | 22.592834 | 7.273354 | 1.645100 | 0.598335 | 0.404864 | 0.300183 | 0.229032 | 0.215211 | 0.215650 | 0.216499 | 0.214989 | 0.216778 | 0.228422 | 0.253892 | 0.321569 | 0.386146 | 0.548616 | 0.706376 | 1.222214 | 1.431221 | 1.711579 | 1.426822 | 1.029891 | 0.721546 | 0.471055 | 0.286764 | 0.260560 | 0.254612 | 0.245869 | 0.265625 | 0.256464 | 0.245490 | 0.245993 | 0.235609 | 0.234367 | 0.244105 | 0.232858 | 0.230048 | 0.256119 | 0.320952 | 0.386217 | 0.569154 | 0.637666 | 0.588649 | 0.639761 | 0.757017 | 0.663530 | 0.699147 | 0.587683 | 0.605687 | 0.895058 | 1.115353 | 1.237367 | 0.924056 | 0.742544 | 0.596851 | 0.405707 | 0.313213 | 0.237840 | 0.214771 | 0.216324 | 0.219305 | 0.225649 | 0.304203 | 0.412996 | 0.582416 | 0.824870 | 0.980200 | 0.791871 | 0.710147 | 0.449407 | 0.317895 | 0.318426 | 0.404776 | 0.451969 | 0.646625 | 0.871129 | 1.053424 | 1.646396 | 1.303849 | 0.760106 | 0.680936 | 0.634548 | 0.580208 | 0.529717 | 0.596826 | 0.877918 | 1.090918 | 0.936226 | 0.791742 | 0.602778 | 0.504217 | 0.332348 | 0.267021 | 0.238471 | 0.245226 | 0.258903 | 0.308969 | 0.429294 | 0.464725 | 0.326568 | 0.307512 | 0.252976 | 0.232075 | 0.235718 | 0.233328 | 0.238550 | 0.252483 | 0.275715 | 0.396379 | 0.474100 | 0.360554 | 0.318254 | 0.505044 | 0.868555 | 1.129854 | 1.231950 | 1.388276 | 1.111343 | 1.442986 | 1.124186 | 0.830895 | 0.828665 | 0.822112 | 0.751228 | 0.741768 | 0.588183 | 0.466106 | 0.492467 | 0.499545 | 0.492298 | 0.495516 | 0.592691 | 0.575593 | 0.408565 | 0.287131 | 0.229783 | 0.216834 | 0.215038 | 0.215050 | 0.214905 | 0.215985 | 0.218880 | 0.217686 | 0.230029 | 0.273117 | 0.325090 | 0.406310 | 0.448854 | 0.656035 | 0.775640 | 0.582634 | 0.474160 | 0.440369 | 0.450221 | 0.529776 | 0.484241 | 0.476388 | 0.449128 | 0.364218 | 0.315458 | 0.274835 | 0.256071 | 0.261238 | 0.256744 | 0.246883 | 0.258105 | 0.288820 | 0.336219 | 0.381160 | 0.423729 | 0.524821 | 0.761848 | 0.820914 | 0.839949 | 0.816117 | 0.741676 | 0.585577 | 0.524682 | 0.495509 | 0.510881 | 0.415735 | 0.341852 | 0.347107 | 0.304035 | 0.267782 | 0.260683 | 0.267277 | 0.312993 | 0.377988 | 0.377247 | 0.369679 | 0.367859 | 0.349863 | 0.315494 | 0.300264 | 0.310247 | 0.304098 | 0.347183 | 0.468232 | 0.601124 | 0.995061 | 1.366356 | 1.140487 | 1.143906 | 1.142990 | 1.214049 | 1.326078 | 1.298782 | 2.081174 | 4.199126 | 11.399668 | 15.119509 | 24.526443 | 31.073204 | 30.637043 |
| all electrodes | 6.946923 | 0.950186 | 0.277499 | 0.238320 | 0.215012 | 0.356615 | 1.663977 | 1.602237 | 0.451363 | 0.261792 | 0.259003 | 0.359278 | 0.586732 | 0.438235 | 0.324076 | 0.428741 | 0.369259 | 0.283732 | 0.287755 | 0.225436 | 0.221017 | 0.218976 | 0.227987 | 0.255173 | 0.286774 | 0.221219 | 0.215741 | 0.215507 | 0.218910 | 0.214626 | 0.215198 | 0.215833 | 0.242268 | 0.272839 | 0.319622 | 0.258586 | 0.280417 | 0.237937 | 0.231258 | 0.214797 | 0.239955 | 0.346006 | 0.394384 | 0.588999 | 0.603007 | 0.824656 | 0.823507 | 0.717815 | 0.315742 | 0.233750 | 0.226017 | 0.292788 | 0.614122 | 1.496579 | 1.776986 | 0.924488 | 1.043296 | 0.571006 | 0.233308 | 0.309026 | 1.046257 | 4.235499 | 5.470971 | 9.671196 | 9.458361 | 2.943134 | 0.626637 | 0.298636 | 0.214708 | 0.217688 | 0.225878 | 0.214774 | 0.217481 | 0.221985 | 0.229753 | 0.263551 | 0.472313 | 0.757296 | 0.593020 | 0.432631 | 0.377893 | 0.409012 | 0.416252 | 0.269439 | 0.229054 | 0.214919 | 0.216369 | 0.215344 | 0.254103 | 0.375711 | 0.370960 | 0.256627 | 0.220778 | 0.216121 | 0.219201 | 0.224179 | 0.232100 | 0.220454 | 0.214912 | 0.215319 | 0.220874 | 0.245488 | 0.269920 | 0.303019 | 0.275143 | 0.286348 | 0.578113 | 0.689846 | 0.581989 | 0.315797 | 0.237102 | 0.246930 | 0.240708 | 0.215224 | 0.244366 | 0.241443 | 0.237877 | 0.215886 | 0.216276 | 0.214726 | 0.223423 | 0.257502 | 0.291696 | 0.357548 | 0.326359 | 0.289269 | 0.241754 | 0.214626 | 0.254675 | 0.525158 | 1.903971 | 4.939723 | 8.411602 | 6.736565 | 5.011847 | 1.672876 | 0.816932 | 0.417336 | 0.288519 | 0.273968 | 0.264432 | 0.287695 | 0.335867 | 0.337438 | 0.359077 | 0.366654 | 0.308741 | 0.298828 | 0.338402 | 0.355842 | 0.374909 | 0.395115 | 0.293110 | 0.238346 | 0.243692 | 0.214626 | 0.229649 | 0.273223 | 0.303178 | 0.279550 | 0.232916 | 0.235619 | 0.222338 | 0.246741 | 0.307710 | 0.436242 | 0.634574 | 0.958983 | 1.416551 | 3.114864 | 3.817748 | 3.068228 | 3.502545 | 3.783010 | 3.672455 | 2.807944 | 1.464933 | 1.999136 | 2.019732 | 2.086296 | 3.744233 | 6.354393 | 3.809925 | 1.776488 | 0.817729 | 0.776171 | 0.532258 | 0.416796 | 0.340735 | 0.408204 | 0.478955 | 0.485101 | 0.560722 | 0.964274 | 1.366008 | 1.835618 | 3.895981 | 13.051290 | 17.455053 | 7.898212 | 18.106352 | 26.788904 | 51.353079 | 9.587072 | 2.842376 | 2.579459 | 2.331715 | 1.137952 | 0.860213 | 0.604054 | 1.157821 | 1.560262 | 1.285490 | 1.026854 | 1.402286 | 2.506010 | 3.873222 | 5.360595 | 4.096301 | 3.246191 | 5.921250 | 5.097159 | 5.856651 | 16.997041 | 12.833197 | 48.426526 | 57.945732 | 47.930836 | 29.210427 | 11.696426 | 2.065702 | 1.005514 | 0.509100 | 0.406498 | 0.294969 | 0.282963 | 0.285364 | 0.378626 | 0.571550 | 0.786920 | 1.005383 | 1.132734 | 0.979392 | 1.353032 | 1.351901 | 1.133180 | 0.834143 | 0.734802 | 0.595039 | 0.444231 | 0.253193 | 0.221623 | 0.214969 | 0.214934 | 0.214793 | 0.214968 | 0.219020 | 0.214892 | 0.240304 | 0.275184 | 0.329260 | 0.325250 | 0.295998 | 0.260978 | 0.270775 | 0.293040 | 0.254625 | 0.217811 | 0.217757 | 0.224101 | 0.227381 | 0.236267 | 0.295577 | 0.349162 | 0.328151 | 0.291685 | 0.271544 | 0.251721 | 0.273133 | 0.266386 |

Searchlight, spatiotemporal cluster permutation test

|  | start time | stop time | peak time | peak channel | cluster p | peak Cohen's d | direction |
| --- | --- | --- | --- | --- | --- | --- | --- |
| #1 | 80 | 325 | 275 | P9 | 0.0104 | 0.727709 | positive |

D) difference, identity, happy vs angry

  
|  | time window | peak latency | cluster *p* | peak Cohen's *d* |  | | | |
| **all electrodes** |  | | | |  | | | |
|  | | | | | | | | |

Time-resolved classification, cluster permutation tests

|  | **left hemisphere** | | | | **right hemisphere** | | | |
|  | time window | peak latency | cluster *p* | peak Cohen's *d* | time window | peak latency | cluster *p* | peak Cohen's *d* |
| **anterior** |  | | | |  | | | |
| **central** |  | | | |  | | | |
| **posterior** | 35 - 175 ms | 105 ms | 0.0084 | 1.0374 |  | | | |

  

Time-resolved classification, Bayesian statistics

|  | -200 | -195 | -190 | -185 | -180 | -175 | -170 | -165 | -160 | -155 | -150 | -145 | -140 | -135 | -130 | -125 | -120 | -115 | -110 | -105 | -100 | -95 | -90 | -85 | -80 | -75 | -70 | -65 | -60 | -55 | -50 | -45 | -40 | -35 | -30 | -25 | -20 | -15 | -10 | -5 | 0 | 5 | 10 | 15 | 20 | 25 | 30 | 35 | 40 | 45 | 50 | 55 | 60 | 65 | 70 | 75 | 80 | 85 | 90 | 95 | 100 | 105 | 110 | 115 | 120 | 125 | 130 | 135 | 140 | 145 | 150 | 155 | 160 | 165 | 170 | 175 | 180 | 185 | 190 | 195 | 200 | 205 | 210 | 215 | 220 | 225 | 230 | 235 | 240 | 245 | 250 | 255 | 260 | 265 | 270 | 275 | 280 | 285 | 290 | 295 | 300 | 305 | 310 | 315 | 320 | 325 | 330 | 335 | 340 | 345 | 350 | 355 | 360 | 365 | 370 | 375 | 380 | 385 | 390 | 395 | 400 | 405 | 410 | 415 | 420 | 425 | 430 | 435 | 440 | 445 | 450 | 455 | 460 | 465 | 470 | 475 | 480 | 485 | 490 | 495 | 500 | 505 | 510 | 515 | 520 | 525 | 530 | 535 | 540 | 545 | 550 | 555 | 560 | 565 | 570 | 575 | 580 | 585 | 590 | 595 | 600 | 605 | 610 | 615 | 620 | 625 | 630 | 635 | 640 | 645 | 650 | 655 | 660 | 665 | 670 | 675 | 680 | 685 | 690 | 695 | 700 | 705 | 710 | 715 | 720 | 725 | 730 | 735 | 740 | 745 | 750 | 755 | 760 | 765 | 770 | 775 | 780 | 785 | 790 | 795 | 800 | 805 | 810 | 815 | 820 | 825 | 830 | 835 | 840 | 845 | 850 | 855 | 860 | 865 | 870 | 875 | 880 | 885 | 890 | 895 | 900 | 905 | 910 | 915 | 920 | 925 | 930 | 935 | 940 | 945 | 950 | 955 | 960 | 965 | 970 | 975 | 980 | 985 | 990 | 995 | 1000 | 1005 | 1010 | 1015 | 1020 | 1025 | 1030 | 1035 | 1040 | 1045 | 1050 | 1055 | 1060 | 1065 | 1070 | 1075 | 1080 | 1085 | 1090 | 1095 | 1100 | 1105 | 1110 | 1115 | 1120 | 1125 | 1130 | 1135 | 1140 | 1145 | 1150 | 1155 | 1160 | 1165 | 1170 | 1175 | 1180 | 1185 | 1190 | 1195 |
| --- | --- | --- | --- | --- | --- | --- | --- | --- | --- | --- | --- | --- | --- | --- | --- | --- | --- | --- | --- | --- | --- | --- | --- | --- | --- | --- | --- | --- | --- | --- | --- | --- | --- | --- | --- | --- | --- | --- | --- | --- | --- | --- | --- | --- | --- | --- | --- | --- | --- | --- | --- | --- | --- | --- | --- | --- | --- | --- | --- | --- | --- | --- | --- | --- | --- | --- | --- | --- | --- | --- | --- | --- | --- | --- | --- | --- | --- | --- | --- | --- | --- | --- | --- | --- | --- | --- | --- | --- | --- | --- | --- | --- | --- | --- | --- | --- | --- | --- | --- | --- | --- | --- | --- | --- | --- | --- | --- | --- | --- | --- | --- | --- | --- | --- | --- | --- | --- | --- | --- | --- | --- | --- | --- | --- | --- | --- | --- | --- | --- | --- | --- | --- | --- | --- | --- | --- | --- | --- | --- | --- | --- | --- | --- | --- | --- | --- | --- | --- | --- | --- | --- | --- | --- | --- | --- | --- | --- | --- | --- | --- | --- | --- | --- | --- | --- | --- | --- | --- | --- | --- | --- | --- | --- | --- | --- | --- | --- | --- | --- | --- | --- | --- | --- | --- | --- | --- | --- | --- | --- | --- | --- | --- | --- | --- | --- | --- | --- | --- | --- | --- | --- | --- | --- | --- | --- | --- | --- | --- | --- | --- | --- | --- | --- | --- | --- | --- | --- | --- | --- | --- | --- | --- | --- | --- | --- | --- | --- | --- | --- | --- | --- | --- | --- | --- | --- | --- | --- | --- | --- | --- | --- | --- | --- | --- | --- | --- | --- | --- | --- | --- | --- | --- | --- | --- | --- | --- | --- | --- | --- | --- | --- | --- | --- | --- | --- | --- | --- | --- | --- | --- | --- | --- | --- | --- | --- | --- | --- | --- | --- | --- |
| left anterior | 0.741289 | 0.514781 | 0.413049 | 0.388585 | 0.269641 | 0.228132 | 0.215080 | 0.224038 | 0.221196 | 0.231391 | 0.277121 | 0.380852 | 0.362482 | 0.221540 | 0.231449 | 0.347832 | 0.286894 | 0.235660 | 0.219365 | 0.275980 | 0.231229 | 0.217588 | 0.430464 | 0.420684 | 0.298011 | 0.220996 | 0.218954 | 0.225479 | 0.222765 | 0.256052 | 0.471532 | 0.786080 | 0.858714 | 0.806024 | 0.592473 | 0.438595 | 0.231463 | 0.244215 | 0.345873 | 0.579657 | 0.945435 | 0.838236 | 0.657598 | 0.364925 | 0.215364 | 0.218242 | 0.236465 | 0.295936 | 0.319484 | 0.348153 | 0.402564 | 0.305111 | 0.496068 | 0.918509 | 0.796524 | 0.824553 | 0.980892 | 0.780601 | 0.639927 | 0.355867 | 0.306203 | 0.287703 | 0.387094 | 0.278793 | 0.256770 | 0.245297 | 0.230433 | 0.216262 | 0.229712 | 0.223726 | 0.266506 | 0.281430 | 0.326453 | 0.391475 | 0.428609 | 0.395312 | 0.423042 | 0.339759 | 0.430964 | 0.563479 | 0.954450 | 1.599231 | 1.264364 | 1.309410 | 2.459980 | 2.461686 | 1.273663 | 0.783841 | 0.698293 | 0.755097 | 0.409455 | 0.261624 | 0.231040 | 0.237890 | 0.221355 | 0.217692 | 0.243219 | 0.240892 | 0.221946 | 0.214661 | 0.216162 | 0.217808 | 0.218720 | 0.217976 | 0.215041 | 0.223239 | 0.284407 | 0.299772 | 0.278858 | 0.256079 | 0.247994 | 0.252764 | 0.237083 | 0.214635 | 0.222534 | 0.275950 | 0.326672 | 0.347033 | 0.414125 | 0.503024 | 0.614288 | 0.496438 | 0.405868 | 0.398685 | 0.328649 | 0.311298 | 0.267355 | 0.226857 | 0.222836 | 0.215425 | 0.214860 | 0.220826 | 0.233334 | 0.239070 | 0.228156 | 0.215913 | 0.217840 | 0.219938 | 0.216387 | 0.218088 | 0.217698 | 0.219183 | 0.214636 | 0.236340 | 0.273691 | 0.245260 | 0.249131 | 0.280040 | 0.354663 | 0.460356 | 0.490957 | 0.556379 | 0.711065 | 0.654440 | 0.466747 | 0.316517 | 0.253368 | 0.242254 | 0.225766 | 0.219581 | 0.218727 | 0.217981 | 0.242803 | 0.242723 | 0.225091 | 0.215108 | 0.215968 | 0.219663 | 0.215195 | 0.226036 | 0.224924 | 0.214626 | 0.220468 | 0.232026 | 0.241871 | 0.232486 | 0.230713 | 0.240468 | 0.246281 | 0.234176 | 0.231411 | 0.233795 | 0.233132 | 0.247516 | 0.276416 | 0.262748 | 0.252043 | 0.240222 | 0.245759 | 0.245371 | 0.238613 | 0.214635 | 0.216607 | 0.214702 | 0.229276 | 0.248776 | 0.267358 | 0.291065 | 0.411124 | 0.491021 | 0.915211 | 0.973987 | 0.747341 | 0.630841 | 0.417731 | 0.354723 | 0.244736 | 0.220026 | 0.269135 | 0.303538 | 0.298830 | 0.264943 | 0.256265 | 0.232753 | 0.219722 | 0.219442 | 0.227777 | 0.246568 | 0.248964 | 0.269017 | 0.296200 | 0.336002 | 0.363639 | 0.430436 | 0.459860 | 0.507356 | 0.429138 | 0.357418 | 0.308627 | 0.257001 | 0.238682 | 0.241531 | 0.279322 | 0.295732 | 0.258102 | 0.238532 | 0.217652 | 0.217932 | 0.239838 | 0.296764 | 0.305727 | 0.251769 | 0.238668 | 0.222976 | 0.216199 | 0.214633 | 0.214807 | 0.215651 | 0.218833 | 0.225723 | 0.217742 | 0.214985 | 0.214806 | 0.214874 | 0.214687 | 0.214783 | 0.214632 | 0.216063 | 0.217339 | 0.214744 | 0.216111 | 0.222681 | 0.217535 | 0.215917 | 0.227198 | 0.295446 | 0.364219 | 0.460638 | 0.442995 | 0.517158 | 0.706512 | 0.654597 | 0.419728 | 0.309629 | 0.270829 | 0.272786 | 0.258500 | 0.232441 | 0.229665 | 0.238840 |
| right anterior | 0.377739 | 0.446208 | 0.479516 | 0.292734 | 0.276865 | 0.240637 | 0.214860 | 0.215531 | 0.262816 | 0.295734 | 0.215806 | 0.357667 | 0.813883 | 0.731086 | 1.012231 | 0.526355 | 0.350137 | 0.307585 | 0.246038 | 0.219968 | 0.216040 | 0.215764 | 0.224686 | 0.285454 | 0.244019 | 0.231326 | 0.339060 | 0.319530 | 0.307340 | 0.235056 | 0.224036 | 0.216388 | 0.245434 | 0.286017 | 0.350394 | 0.290303 | 0.252250 | 0.236783 | 0.256389 | 0.283887 | 0.303896 | 0.314596 | 0.302937 | 0.364444 | 0.260805 | 0.266981 | 0.265011 | 0.326836 | 0.387360 | 0.502589 | 0.291282 | 0.300846 | 0.286248 | 0.272654 | 0.274524 | 0.289593 | 0.354708 | 0.647661 | 0.607255 | 0.417149 | 0.363746 | 0.341986 | 0.311694 | 0.265282 | 0.228701 | 0.228792 | 0.215941 | 0.214729 | 0.216275 | 0.219545 | 0.230087 | 0.238115 | 0.231082 | 0.220471 | 0.221999 | 0.229266 | 0.244355 | 0.232327 | 0.228565 | 0.225996 | 0.240653 | 0.250360 | 0.229908 | 0.230887 | 0.229361 | 0.221098 | 0.217610 | 0.214626 | 0.228070 | 0.227410 | 0.265602 | 0.318066 | 0.413872 | 0.379698 | 0.359258 | 0.258477 | 0.278089 | 0.242743 | 0.231499 | 0.224757 | 0.225021 | 0.242410 | 0.284493 | 0.245368 | 0.237557 | 0.220471 | 0.233754 | 0.273752 | 0.247682 | 0.253527 | 0.253873 | 0.245370 | 0.229961 | 0.222212 | 0.251377 | 0.234542 | 0.274198 | 0.261397 | 0.241149 | 0.234041 | 0.218850 | 0.219984 | 0.214640 | 0.257219 | 0.324812 | 0.341020 | 0.283998 | 0.271995 | 0.276492 | 0.214859 | 0.252489 | 0.318730 | 0.336343 | 0.294555 | 0.253607 | 0.244013 | 0.217589 | 0.222424 | 0.232581 | 0.250884 | 0.249309 | 0.219631 | 0.216294 | 0.221662 | 0.252294 | 0.284737 | 0.428648 | 0.521239 | 0.537338 | 0.529373 | 0.416533 | 0.311512 | 0.239329 | 0.216483 | 0.224541 | 0.269388 | 0.363298 | 0.302930 | 0.223156 | 0.229748 | 0.263195 | 0.279731 | 0.323837 | 0.385698 | 0.308139 | 0.247897 | 0.221972 | 0.228549 | 0.228745 | 0.221174 | 0.228784 | 0.249900 | 0.255665 | 0.239581 | 0.214752 | 0.223692 | 0.222334 | 0.218341 | 0.234155 | 0.257226 | 0.236406 | 0.216702 | 0.242024 | 0.233751 | 0.223775 | 0.221159 | 0.229296 | 0.216406 | 0.215532 | 0.217038 | 0.232678 | 0.237198 | 0.218696 | 0.214890 | 0.217770 | 0.268727 | 0.289940 | 0.266536 | 0.277862 | 0.354216 | 0.383895 | 0.276303 | 0.237018 | 0.216027 | 0.216860 | 0.217890 | 0.239733 | 0.266297 | 0.233517 | 0.240484 | 0.254649 | 0.258118 | 0.284575 | 0.297751 | 0.282662 | 0.273644 | 0.277162 | 0.270537 | 0.258353 | 0.235131 | 0.216736 | 0.215393 | 0.218439 | 0.230101 | 0.257635 | 0.249841 | 0.259133 | 0.259840 | 0.263319 | 0.252384 | 0.243469 | 0.226051 | 0.223829 | 0.215848 | 0.214934 | 0.220620 | 0.235075 | 0.220925 | 0.214729 | 0.227573 | 0.266180 | 0.284775 | 0.255002 | 0.237576 | 0.220578 | 0.215197 | 0.216593 | 0.222967 | 0.216796 | 0.214679 | 0.221839 | 0.226455 | 0.258791 | 0.261750 | 0.243697 | 0.225629 | 0.240319 | 0.264532 | 0.285809 | 0.255382 | 0.237617 | 0.264124 | 0.301211 | 0.320606 | 0.266403 | 0.221934 | 0.222793 | 0.233802 | 0.221417 | 0.214734 | 0.239167 | 0.286851 | 0.277681 | 0.285011 | 0.263903 | 0.222527 | 0.215082 | 0.216447 | 0.231239 | 0.288603 |
| left central | 2.055704 | 5.994374 | 3.628026 | 3.813899 | 2.944075 | 1.652026 | 1.593376 | 0.738318 | 0.388303 | 0.297248 | 0.224789 | 0.233508 | 0.389961 | 1.078985 | 1.421602 | 1.410826 | 0.968064 | 0.430220 | 0.361406 | 0.292527 | 0.217635 | 0.227510 | 0.221236 | 0.222960 | 0.232922 | 0.219208 | 0.273181 | 0.448097 | 0.823005 | 1.164747 | 1.240309 | 4.407850 | 1.830257 | 0.516246 | 0.298433 | 0.217931 | 0.218002 | 0.216065 | 0.231059 | 0.249493 | 0.233189 | 0.261764 | 0.224166 | 0.227905 | 0.231605 | 0.217890 | 0.247237 | 0.301607 | 0.332635 | 0.247444 | 0.218853 | 0.214642 | 0.249178 | 0.377563 | 0.573025 | 0.653738 | 1.087253 | 1.892532 | 0.703183 | 0.328669 | 0.236125 | 0.231966 | 0.227420 | 0.218865 | 0.215843 | 0.246319 | 0.271776 | 0.298415 | 0.283806 | 0.244419 | 0.225581 | 0.223572 | 0.214677 | 0.217076 | 0.215588 | 0.251894 | 0.454799 | 0.805939 | 0.974456 | 0.975956 | 1.636627 | 3.428139 | 1.977601 | 1.777402 | 0.885669 | 0.467062 | 0.415666 | 0.316280 | 0.262126 | 0.235477 | 0.221427 | 0.246104 | 0.297991 | 0.319036 | 0.432750 | 0.488679 | 0.679127 | 0.964851 | 0.745537 | 0.837792 | 0.577916 | 0.430685 | 0.398232 | 0.436803 | 0.337697 | 0.439252 | 0.330196 | 0.307682 | 0.270311 | 0.248014 | 0.224363 | 0.219217 | 0.214639 | 0.214680 | 0.221824 | 0.219761 | 0.232986 | 0.236925 | 0.228790 | 0.232131 | 0.240014 | 0.228211 | 0.246431 | 0.238700 | 0.229190 | 0.223143 | 0.217644 | 0.215304 | 0.220884 | 0.231601 | 0.218217 | 0.219051 | 0.217485 | 0.239941 | 0.265274 | 0.285256 | 0.247889 | 0.361172 | 0.522951 | 0.794495 | 0.736970 | 0.673619 | 0.430813 | 0.530157 | 0.651596 | 0.748276 | 0.961569 | 0.903069 | 0.788111 | 0.740348 | 0.441838 | 0.255413 | 0.227077 | 0.215245 | 0.215742 | 0.217298 | 0.214638 | 0.223259 | 0.259637 | 0.303846 | 0.360519 | 0.435360 | 0.559066 | 0.458751 | 0.303478 | 0.264539 | 0.248491 | 0.272053 | 0.237340 | 0.226837 | 0.234964 | 0.272253 | 0.293563 | 0.270403 | 0.247695 | 0.259303 | 0.260378 | 0.271996 | 0.297960 | 0.269337 | 0.257088 | 0.244993 | 0.292027 | 0.385913 | 0.448925 | 0.305990 | 0.269960 | 0.284528 | 0.353687 | 0.307941 | 0.233202 | 0.221785 | 0.228592 | 0.238074 | 0.223722 | 0.218274 | 0.225971 | 0.223514 | 0.260845 | 0.232656 | 0.215554 | 0.220143 | 0.235018 | 0.219078 | 0.215040 | 0.216512 | 0.215231 | 0.226892 | 0.250610 | 0.247366 | 0.234279 | 0.216530 | 0.214949 | 0.214808 | 0.218228 | 0.229551 | 0.218966 | 0.217803 | 0.219505 | 0.246124 | 0.321533 | 0.361807 | 0.324898 | 0.375723 | 0.366911 | 0.291182 | 0.215849 | 0.245636 | 0.270924 | 0.284186 | 0.285577 | 0.278627 | 0.280420 | 0.259174 | 0.240225 | 0.227602 | 0.215339 | 0.216489 | 0.219939 | 0.235685 | 0.246789 | 0.292586 | 0.335778 | 0.294243 | 0.246951 | 0.233369 | 0.223810 | 0.221066 | 0.216150 | 0.230507 | 0.240886 | 0.239031 | 0.238925 | 0.235669 | 0.221584 | 0.214948 | 0.221816 | 0.238096 | 0.238528 | 0.241039 | 0.247193 | 0.226349 | 0.219796 | 0.216700 | 0.215721 | 0.215235 | 0.214626 | 0.214980 | 0.216247 | 0.217531 | 0.218286 | 0.226970 | 0.239243 | 0.260608 | 0.272945 | 0.299065 | 0.336666 | 0.360191 | 0.386786 | 0.373048 |
| right central | 17.198598 | 14.974262 | 13.205051 | 14.889077 | 22.859274 | 10.595784 | 2.256055 | 0.826495 | 0.818046 | 0.572958 | 0.322625 | 0.245837 | 0.229456 | 0.248840 | 0.291845 | 0.259988 | 0.343225 | 0.418638 | 0.777839 | 0.589051 | 0.441508 | 0.472748 | 0.577387 | 0.285705 | 0.215218 | 0.223787 | 0.221694 | 0.219027 | 0.246329 | 0.353739 | 0.342616 | 0.289022 | 0.248278 | 0.233447 | 0.221131 | 0.244238 | 0.266793 | 0.327160 | 0.345755 | 0.430591 | 0.675282 | 1.116343 | 1.169012 | 0.478295 | 0.262248 | 0.218935 | 0.214637 | 0.225997 | 0.265700 | 0.269997 | 0.221228 | 0.229152 | 0.323767 | 0.580108 | 0.784269 | 0.642129 | 0.294171 | 0.223529 | 0.217723 | 0.242245 | 0.300248 | 0.299904 | 0.248666 | 0.218691 | 0.224035 | 0.262250 | 0.315919 | 0.516364 | 0.805084 | 1.344665 | 2.226326 | 1.612489 | 0.691134 | 0.564980 | 0.350331 | 0.271745 | 0.215168 | 0.225227 | 0.223003 | 0.219037 | 0.215349 | 0.214685 | 0.222683 | 0.219541 | 0.215264 | 0.225378 | 0.298809 | 0.329720 | 0.315234 | 0.252988 | 0.221958 | 0.214643 | 0.217875 | 0.222439 | 0.223728 | 0.231144 | 0.224498 | 0.241054 | 0.261118 | 0.259902 | 0.268418 | 0.284310 | 0.359856 | 0.492564 | 0.625416 | 0.523025 | 0.576767 | 0.393438 | 0.339846 | 0.228928 | 0.216575 | 0.214662 | 0.219973 | 0.228995 | 0.214975 | 0.216106 | 0.223124 | 0.220313 | 0.216248 | 0.223953 | 0.224801 | 0.225428 | 0.221258 | 0.214756 | 0.230589 | 0.235100 | 0.225715 | 0.227093 | 0.256583 | 0.344833 | 0.489664 | 0.437672 | 0.427321 | 0.472048 | 0.521616 | 0.461722 | 0.408492 | 0.447049 | 0.384628 | 0.359082 | 0.315533 | 0.263724 | 0.226662 | 0.230807 | 0.219333 | 0.215360 | 0.215669 | 0.214835 | 0.220197 | 0.227973 | 0.220489 | 0.217077 | 0.216369 | 0.214663 | 0.217424 | 0.221512 | 0.228288 | 0.246327 | 0.249270 | 0.217407 | 0.214705 | 0.228200 | 0.218850 | 0.215290 | 0.218891 | 0.220148 | 0.284238 | 0.362570 | 0.438853 | 0.363767 | 0.266370 | 0.217399 | 0.218822 | 0.250510 | 0.242448 | 0.233899 | 0.221730 | 0.215064 | 0.223428 | 0.255522 | 0.355307 | 0.356343 | 0.370722 | 0.429483 | 0.533019 | 0.668830 | 1.029273 | 0.783863 | 0.568310 | 0.481062 | 0.407541 | 0.264411 | 0.225768 | 0.217453 | 0.224865 | 0.270164 | 0.306581 | 0.286852 | 0.276772 | 0.264637 | 0.253836 | 0.228921 | 0.215238 | 0.214665 | 0.214968 | 0.215712 | 0.215682 | 0.220500 | 0.221963 | 0.220225 | 0.216194 | 0.214840 | 0.215019 | 0.215987 | 0.221070 | 0.221695 | 0.222032 | 0.221432 | 0.216974 | 0.220113 | 0.218011 | 0.215562 | 0.214665 | 0.214636 | 0.216035 | 0.222784 | 0.229882 | 0.253314 | 0.245268 | 0.235027 | 0.222464 | 0.214922 | 0.215968 | 0.215237 | 0.219761 | 0.214634 | 0.239545 | 0.333656 | 0.405727 | 0.384211 | 0.325189 | 0.289676 | 0.241395 | 0.214696 | 0.226121 | 0.226201 | 0.221557 | 0.214708 | 0.218390 | 0.239099 | 0.275480 | 0.364302 | 0.482685 | 0.610332 | 0.465197 | 0.487192 | 0.345256 | 0.287652 | 0.247511 | 0.216888 | 0.214710 | 0.217739 | 0.224763 | 0.221171 | 0.216322 | 0.215122 | 0.215913 | 0.214739 | 0.214980 | 0.221977 | 0.240895 | 0.263293 | 0.297068 | 0.302505 | 0.347012 | 0.408235 | 0.368990 | 0.368991 | 0.308160 | 0.259703 |
| left posterior | 0.716610 | 0.805136 | 0.712452 | 0.858714 | 1.091066 | 0.732176 | 0.348292 | 0.227537 | 0.214639 | 0.222920 | 0.424379 | 3.224130 | 5.974579 | 4.124284 | 1.610619 | 0.883432 | 1.232240 | 1.323145 | 1.046656 | 0.675536 | 0.363600 | 0.247548 | 0.214701 | 0.343859 | 0.857480 | 3.635438 | 3.140799 | 1.885763 | 1.950186 | 1.317134 | 1.081695 | 1.365614 | 1.716578 | 1.239504 | 0.581546 | 0.263624 | 0.217084 | 0.309738 | 0.718105 | 1.371369 | 1.634405 | 0.502122 | 0.241401 | 0.215305 | 0.250952 | 0.405973 | 0.715677 | 1.305370 | 1.554230 | 1.581215 | 2.344182 | 2.243785 | 2.680798 | 3.146991 | 4.690482 | 5.912473 | 5.612270 | 5.789415 | 18.752817 | 50.100759 | 245.671881 | 645.804617 | 801.232644 | 785.894351 | 539.116291 | 81.654458 | 11.192712 | 3.986608 | 2.706632 | 4.675065 | 6.066650 | 3.718647 | 2.769268 | 1.551816 | 1.710391 | 1.502371 | 1.169697 | 1.216655 | 2.064783 | 4.719472 | 8.512293 | 8.854362 | 9.897503 | 13.996461 | 11.443246 | 6.143844 | 1.416872 | 0.613337 | 0.465459 | 0.579027 | 0.526802 | 0.446514 | 0.292098 | 0.270240 | 0.268601 | 0.227206 | 0.214638 | 0.215141 | 0.215116 | 0.223089 | 0.269086 | 0.282261 | 0.374586 | 0.482418 | 0.464208 | 0.426959 | 0.404949 | 0.439968 | 0.637919 | 0.691311 | 1.101196 | 2.135248 | 4.296890 | 10.858201 | 15.067838 | 18.294834 | 49.072715 | 51.198002 | 45.270443 | 27.344970 | 5.470569 | 3.270566 | 2.755490 | 1.713373 | 1.421515 | 0.723429 | 0.607250 | 0.626080 | 0.678208 | 0.856734 | 1.047517 | 1.721871 | 3.548620 | 7.356931 | 13.099444 | 7.757864 | 3.631528 | 2.079201 | 0.650136 | 0.375594 | 0.281574 | 0.249936 | 0.259051 | 0.282833 | 0.288545 | 0.326815 | 0.384547 | 0.374226 | 0.367017 | 0.357671 | 0.335584 | 0.319117 | 0.278681 | 0.270738 | 0.328471 | 0.385743 | 0.435387 | 0.522690 | 0.612610 | 0.894565 | 1.070751 | 0.779928 | 0.537904 | 0.511337 | 0.388473 | 0.329233 | 0.276627 | 0.237670 | 0.224448 | 0.219330 | 0.214626 | 0.215394 | 0.215813 | 0.215291 | 0.215411 | 0.221492 | 0.229808 | 0.225757 | 0.226182 | 0.230068 | 0.253072 | 0.307331 | 0.530025 | 1.091367 | 3.101493 | 4.879827 | 15.149265 | 7.894524 | 3.714604 | 1.405247 | 0.701558 | 0.437907 | 0.418551 | 0.324280 | 0.314478 | 0.327210 | 0.420881 | 0.441172 | 0.520429 | 0.492226 | 0.390793 | 0.380826 | 0.340424 | 0.270786 | 0.229209 | 0.214914 | 0.220253 | 0.230796 | 0.262412 | 0.276854 | 0.261483 | 0.237247 | 0.225937 | 0.228453 | 0.224619 | 0.216885 | 0.226093 | 0.239274 | 0.235046 | 0.242971 | 0.252173 | 0.274838 | 0.295191 | 0.246338 | 0.239486 | 0.238372 | 0.231481 | 0.221597 | 0.216807 | 0.214637 | 0.219882 | 0.249269 | 0.268361 | 0.321732 | 0.473813 | 0.887353 | 1.183799 | 1.104817 | 1.074645 | 1.083011 | 1.205959 | 1.291508 | 1.372028 | 1.650512 | 1.993910 | 1.708432 | 1.520314 | 1.475311 | 1.184700 | 0.883581 | 0.778535 | 0.863137 | 1.169760 | 1.149474 | 0.952382 | 1.134829 | 1.015032 | 0.798797 | 0.632980 | 0.438835 | 0.431742 | 0.443539 | 0.408848 | 0.396105 | 0.466458 | 0.598112 | 0.672315 | 0.723404 | 0.658424 | 0.449434 | 0.481602 | 0.406544 | 0.403718 | 0.416470 | 0.351459 | 0.346876 | 0.370425 | 0.348986 | 0.348190 | 0.275922 |
| right posterior | 0.511310 | 0.432847 | 0.424224 | 0.315606 | 0.219253 | 0.218590 | 0.224650 | 0.214962 | 0.218639 | 0.215619 | 0.219685 | 0.278769 | 0.624318 | 1.147757 | 1.090158 | 1.890375 | 3.207473 | 2.791208 | 0.897586 | 0.296100 | 0.218699 | 0.216062 | 0.214740 | 0.227055 | 0.222286 | 0.234328 | 0.235352 | 0.267544 | 0.535974 | 0.397526 | 0.278202 | 0.389805 | 0.354880 | 0.458489 | 0.643217 | 0.323375 | 0.293439 | 0.396202 | 0.447830 | 0.657974 | 0.628611 | 0.309833 | 0.247964 | 0.345812 | 0.371672 | 0.435116 | 0.489373 | 0.353893 | 0.395774 | 0.865734 | 0.714129 | 0.479106 | 0.241789 | 0.233033 | 0.251623 | 0.250874 | 0.365117 | 0.584920 | 0.828630 | 1.278552 | 2.209997 | 7.855256 | 9.369393 | 10.960559 | 7.687639 | 3.185153 | 1.048758 | 0.471408 | 0.243776 | 0.215894 | 0.217981 | 0.217719 | 0.223534 | 0.225689 | 0.231502 | 0.247246 | 0.262352 | 0.250833 | 0.264965 | 0.256161 | 0.278183 | 0.262994 | 0.240002 | 0.232972 | 0.232113 | 0.226739 | 0.216508 | 0.247903 | 0.246502 | 0.230076 | 0.233922 | 0.254867 | 0.256568 | 0.229319 | 0.222445 | 0.252145 | 0.382380 | 0.513041 | 0.393423 | 0.317079 | 0.305982 | 0.287872 | 0.271219 | 0.238636 | 0.238045 | 0.255772 | 0.291536 | 0.314658 | 0.318974 | 0.307857 | 0.339471 | 0.362328 | 0.419185 | 0.603079 | 1.000998 | 1.757565 | 2.859514 | 3.193398 | 2.882132 | 2.186341 | 1.125401 | 0.672014 | 0.597333 | 0.590572 | 0.596478 | 0.722199 | 1.132057 | 3.323762 | 6.282108 | 6.714734 | 2.675894 | 1.743074 | 1.087515 | 0.716136 | 0.359100 | 0.265251 | 0.319785 | 0.701555 | 1.295422 | 2.199136 | 3.470887 | 6.193479 | 9.654334 | 8.736168 | 4.478003 | 4.641718 | 6.189708 | 4.845962 | 2.394916 | 1.556799 | 0.913173 | 0.600777 | 0.557517 | 0.509253 | 0.700710 | 0.725975 | 0.816932 | 0.933589 | 1.470960 | 1.506843 | 1.474691 | 0.745668 | 0.524537 | 0.399627 | 0.424536 | 0.446778 | 0.441955 | 0.360891 | 0.416591 | 0.443260 | 0.432904 | 0.356541 | 0.297173 | 0.263161 | 0.267883 | 0.247386 | 0.240405 | 0.269428 | 0.279012 | 0.298535 | 0.359936 | 0.509495 | 0.685046 | 0.904210 | 1.143679 | 1.617940 | 1.959317 | 1.761143 | 1.637064 | 1.661113 | 1.466031 | 1.072987 | 1.010758 | 1.005987 | 0.904697 | 0.725787 | 0.754977 | 0.636988 | 0.505107 | 0.511013 | 0.432952 | 0.422666 | 0.436537 | 0.312831 | 0.357023 | 0.440541 | 0.409985 | 0.456451 | 0.495409 | 0.504043 | 0.635530 | 0.508546 | 0.426363 | 0.371677 | 0.334298 | 0.323730 | 0.277608 | 0.240051 | 0.232650 | 0.228404 | 0.229779 | 0.234355 | 0.235989 | 0.263811 | 0.316902 | 0.372276 | 0.474549 | 0.572683 | 0.585094 | 0.653561 | 0.595667 | 0.574384 | 0.565431 | 0.434600 | 0.377835 | 0.341903 | 0.290709 | 0.263630 | 0.263147 | 0.258628 | 0.290307 | 0.329515 | 0.345313 | 0.337642 | 0.402593 | 0.454307 | 0.490000 | 0.526976 | 0.447134 | 0.475168 | 0.629096 | 0.684660 | 0.646755 | 0.510117 | 0.354140 | 0.447481 | 0.461857 | 0.430275 | 0.422003 | 0.411067 | 0.559883 | 0.985186 | 0.909658 | 0.805427 | 0.968301 | 0.960323 | 0.848968 | 0.656422 | 0.432651 | 0.362051 | 0.331484 | 0.279178 | 0.236494 | 0.228355 | 0.215581 | 0.215416 | 0.218594 | 0.222755 | 0.228023 | 0.221020 |
| all electrodes | 0.277130 | 0.404911 | 0.503381 | 0.649084 | 2.376518 | 4.466635 | 3.419758 | 0.818764 | 0.472830 | 0.391222 | 0.266243 | 0.227565 | 0.256180 | 0.230058 | 0.245073 | 0.280551 | 0.223895 | 0.227211 | 0.220586 | 0.214888 | 0.214715 | 0.224099 | 0.215376 | 0.260824 | 0.615292 | 0.987049 | 2.075557 | 4.309464 | 0.944427 | 0.593706 | 0.395276 | 0.344208 | 0.365524 | 0.300429 | 0.220593 | 0.219525 | 0.236062 | 0.243839 | 0.215305 | 0.245247 | 0.327643 | 0.313362 | 0.340213 | 0.352050 | 0.382409 | 0.279417 | 0.251854 | 0.228121 | 0.228923 | 0.227279 | 0.237410 | 0.230445 | 0.215622 | 0.215449 | 0.224207 | 0.259498 | 0.307021 | 0.269328 | 0.223723 | 0.216871 | 0.214640 | 0.245316 | 0.276975 | 0.287031 | 0.249439 | 0.227730 | 0.215691 | 0.219283 | 0.214984 | 0.223058 | 0.242817 | 0.244577 | 0.259066 | 0.240793 | 0.221865 | 0.216669 | 0.216627 | 0.226367 | 0.220483 | 0.228522 | 0.258889 | 0.338003 | 0.268945 | 0.245689 | 0.220609 | 0.215160 | 0.224242 | 0.256797 | 0.302784 | 0.293300 | 0.233782 | 0.214734 | 0.284150 | 0.573763 | 0.931655 | 1.503369 | 1.497191 | 0.979737 | 0.875386 | 0.513683 | 0.400173 | 0.315956 | 0.235811 | 0.215490 | 0.215337 | 0.219751 | 0.236106 | 0.293743 | 0.310643 | 0.355488 | 0.373184 | 0.421585 | 0.396786 | 0.360327 | 0.413343 | 0.551365 | 0.386461 | 0.279324 | 0.267888 | 0.248418 | 0.260665 | 0.226448 | 0.215019 | 0.220709 | 0.215087 | 0.218423 | 0.239889 | 0.221363 | 0.222416 | 0.218728 | 0.231204 | 0.256175 | 0.255880 | 0.242010 | 0.267517 | 0.266379 | 0.269918 | 0.268077 | 0.282612 | 0.258670 | 0.246588 | 0.275921 | 0.288964 | 0.390582 | 0.399158 | 0.389529 | 0.576266 | 0.790915 | 0.603378 | 0.702091 | 0.603418 | 0.955834 | 0.969245 | 0.827812 | 0.734285 | 0.943122 | 1.028147 | 1.253661 | 0.708304 | 0.799651 | 0.573168 | 0.528812 | 0.508263 | 0.442530 | 0.380206 | 0.273670 | 0.229642 | 0.214758 | 0.221533 | 0.235688 | 0.254352 | 0.260986 | 0.250158 | 0.229580 | 0.216435 | 0.214851 | 0.235176 | 0.309599 | 0.299406 | 0.297662 | 0.266875 | 0.260741 | 0.345183 | 0.393168 | 0.612059 | 1.024320 | 1.346453 | 2.054839 | 2.111788 | 0.971041 | 0.497366 | 0.268190 | 0.248133 | 0.226616 | 0.216417 | 0.227467 | 0.219069 | 0.223392 | 0.222314 | 0.217242 | 0.219556 | 0.249696 | 0.234838 | 0.221468 | 0.214626 | 0.215834 | 0.242286 | 0.297686 | 0.310121 | 0.240680 | 0.230900 | 0.216490 | 0.214900 | 0.227476 | 0.274354 | 0.303404 | 0.315921 | 0.284758 | 0.266037 | 0.240929 | 0.222960 | 0.214928 | 0.218381 | 0.227925 | 0.222837 | 0.221276 | 0.216395 | 0.223124 | 0.223554 | 0.215809 | 0.215797 | 0.215517 | 0.216865 | 0.223733 | 0.226128 | 0.221092 | 0.220712 | 0.217645 | 0.214925 | 0.217400 | 0.243748 | 0.273770 | 0.283438 | 0.286022 | 0.280036 | 0.254134 | 0.236611 | 0.230251 | 0.258006 | 0.340634 | 0.315164 | 0.285711 | 0.266543 | 0.257378 | 0.266941 | 0.250927 | 0.245107 | 0.265498 | 0.264143 | 0.256820 | 0.276194 | 0.232609 | 0.218881 | 0.215292 | 0.215723 | 0.215010 | 0.221413 | 0.225078 | 0.240049 | 0.235607 | 0.235072 | 0.241657 | 0.234868 | 0.224260 | 0.215079 | 0.215866 | 0.222571 | 0.238834 | 0.287005 | 0.312410 |

Searchlight, spatiotemporal cluster permutation test

No significant clusters observed.

E) difference, identity, angry vs sad

  
|  | time window | peak latency | cluster *p* | peak Cohen's *d* |  | | | |
| **all electrodes** | 560 - 830 ms | 645 ms | 0.001 | 0.6286 |  | | | |
|  | | | | | | | | |

Time-resolved classification, cluster permutation tests

|  | **left hemisphere** | | | | **right hemisphere** | | | |
|  | time window | peak latency | cluster *p* | peak Cohen's *d* | time window | peak latency | cluster *p* | peak Cohen's *d* |
| **anterior** |  | | | |  | | | |
| **central** | 660 - 735 ms | 690 ms | 0.0462 | 0.6975 |  | | | |
| **posterior** |  | | | |  | | | |

  

Time-resolved classification, Bayesian statistics

|  | -200 | -195 | -190 | -185 | -180 | -175 | -170 | -165 | -160 | -155 | -150 | -145 | -140 | -135 | -130 | -125 | -120 | -115 | -110 | -105 | -100 | -95 | -90 | -85 | -80 | -75 | -70 | -65 | -60 | -55 | -50 | -45 | -40 | -35 | -30 | -25 | -20 | -15 | -10 | -5 | 0 | 5 | 10 | 15 | 20 | 25 | 30 | 35 | 40 | 45 | 50 | 55 | 60 | 65 | 70 | 75 | 80 | 85 | 90 | 95 | 100 | 105 | 110 | 115 | 120 | 125 | 130 | 135 | 140 | 145 | 150 | 155 | 160 | 165 | 170 | 175 | 180 | 185 | 190 | 195 | 200 | 205 | 210 | 215 | 220 | 225 | 230 | 235 | 240 | 245 | 250 | 255 | 260 | 265 | 270 | 275 | 280 | 285 | 290 | 295 | 300 | 305 | 310 | 315 | 320 | 325 | 330 | 335 | 340 | 345 | 350 | 355 | 360 | 365 | 370 | 375 | 380 | 385 | 390 | 395 | 400 | 405 | 410 | 415 | 420 | 425 | 430 | 435 | 440 | 445 | 450 | 455 | 460 | 465 | 470 | 475 | 480 | 485 | 490 | 495 | 500 | 505 | 510 | 515 | 520 | 525 | 530 | 535 | 540 | 545 | 550 | 555 | 560 | 565 | 570 | 575 | 580 | 585 | 590 | 595 | 600 | 605 | 610 | 615 | 620 | 625 | 630 | 635 | 640 | 645 | 650 | 655 | 660 | 665 | 670 | 675 | 680 | 685 | 690 | 695 | 700 | 705 | 710 | 715 | 720 | 725 | 730 | 735 | 740 | 745 | 750 | 755 | 760 | 765 | 770 | 775 | 780 | 785 | 790 | 795 | 800 | 805 | 810 | 815 | 820 | 825 | 830 | 835 | 840 | 845 | 850 | 855 | 860 | 865 | 870 | 875 | 880 | 885 | 890 | 895 | 900 | 905 | 910 | 915 | 920 | 925 | 930 | 935 | 940 | 945 | 950 | 955 | 960 | 965 | 970 | 975 | 980 | 985 | 990 | 995 | 1000 | 1005 | 1010 | 1015 | 1020 | 1025 | 1030 | 1035 | 1040 | 1045 | 1050 | 1055 | 1060 | 1065 | 1070 | 1075 | 1080 | 1085 | 1090 | 1095 | 1100 | 1105 | 1110 | 1115 | 1120 | 1125 | 1130 | 1135 | 1140 | 1145 | 1150 | 1155 | 1160 | 1165 | 1170 | 1175 | 1180 | 1185 | 1190 | 1195 |
| --- | --- | --- | --- | --- | --- | --- | --- | --- | --- | --- | --- | --- | --- | --- | --- | --- | --- | --- | --- | --- | --- | --- | --- | --- | --- | --- | --- | --- | --- | --- | --- | --- | --- | --- | --- | --- | --- | --- | --- | --- | --- | --- | --- | --- | --- | --- | --- | --- | --- | --- | --- | --- | --- | --- | --- | --- | --- | --- | --- | --- | --- | --- | --- | --- | --- | --- | --- | --- | --- | --- | --- | --- | --- | --- | --- | --- | --- | --- | --- | --- | --- | --- | --- | --- | --- | --- | --- | --- | --- | --- | --- | --- | --- | --- | --- | --- | --- | --- | --- | --- | --- | --- | --- | --- | --- | --- | --- | --- | --- | --- | --- | --- | --- | --- | --- | --- | --- | --- | --- | --- | --- | --- | --- | --- | --- | --- | --- | --- | --- | --- | --- | --- | --- | --- | --- | --- | --- | --- | --- | --- | --- | --- | --- | --- | --- | --- | --- | --- | --- | --- | --- | --- | --- | --- | --- | --- | --- | --- | --- | --- | --- | --- | --- | --- | --- | --- | --- | --- | --- | --- | --- | --- | --- | --- | --- | --- | --- | --- | --- | --- | --- | --- | --- | --- | --- | --- | --- | --- | --- | --- | --- | --- | --- | --- | --- | --- | --- | --- | --- | --- | --- | --- | --- | --- | --- | --- | --- | --- | --- | --- | --- | --- | --- | --- | --- | --- | --- | --- | --- | --- | --- | --- | --- | --- | --- | --- | --- | --- | --- | --- | --- | --- | --- | --- | --- | --- | --- | --- | --- | --- | --- | --- | --- | --- | --- | --- | --- | --- | --- | --- | --- | --- | --- | --- | --- | --- | --- | --- | --- | --- | --- | --- | --- | --- | --- | --- | --- | --- | --- | --- | --- | --- | --- | --- | --- | --- | --- | --- | --- | --- |
| left anterior | 0.266163 | 0.214646 | 0.220027 | 0.240443 | 0.275413 | 0.266030 | 0.217422 | 0.272132 | 0.783954 | 1.512903 | 2.746219 | 1.853698 | 1.920904 | 0.375853 | 0.214979 | 0.290612 | 0.450632 | 0.465205 | 0.294451 | 0.574841 | 0.521486 | 0.256044 | 0.255307 | 0.408247 | 0.387714 | 0.263276 | 0.260166 | 0.239265 | 0.214643 | 0.463081 | 0.888102 | 0.898969 | 0.731088 | 0.864385 | 0.510592 | 0.328761 | 0.224170 | 0.215743 | 0.219981 | 0.221888 | 0.225625 | 0.294958 | 0.299484 | 0.244170 | 0.228404 | 0.226794 | 0.226981 | 0.222578 | 0.308973 | 0.329904 | 0.311420 | 0.275858 | 0.352741 | 0.622620 | 0.826003 | 1.251369 | 4.733666 | 12.356038 | 14.026432 | 2.938334 | 1.615483 | 1.231663 | 0.890219 | 0.343704 | 0.228060 | 0.214626 | 0.214689 | 0.223537 | 0.282520 | 0.417831 | 0.299493 | 0.245378 | 0.218487 | 0.219551 | 0.214691 | 0.217220 | 0.226632 | 0.214745 | 0.216693 | 0.218597 | 0.277043 | 0.474504 | 0.623636 | 1.092499 | 3.339087 | 6.390613 | 9.163355 | 6.844401 | 4.319929 | 4.991058 | 3.583216 | 2.209993 | 1.465266 | 0.737485 | 0.443438 | 0.267577 | 0.217981 | 0.221783 | 0.231699 | 0.225076 | 0.230067 | 0.219138 | 0.215243 | 0.215250 | 0.214967 | 0.220068 | 0.232428 | 0.293484 | 0.267395 | 0.321362 | 0.487915 | 0.783570 | 1.047550 | 1.368206 | 1.161450 | 1.866852 | 1.526531 | 0.851319 | 0.552018 | 0.470233 | 0.403031 | 0.440909 | 0.380256 | 0.374704 | 0.462977 | 0.905156 | 0.877576 | 0.535344 | 0.335993 | 0.262528 | 0.229937 | 0.223272 | 0.216659 | 0.217422 | 0.220792 | 0.230659 | 0.227138 | 0.226546 | 0.226837 | 0.226657 | 0.219330 | 0.214727 | 0.216524 | 0.214637 | 0.216940 | 0.217384 | 0.218956 | 0.223338 | 0.242118 | 0.261928 | 0.287959 | 0.385122 | 0.447820 | 0.450557 | 0.454082 | 0.572045 | 0.764555 | 1.086968 | 1.012962 | 0.979293 | 0.648521 | 0.531464 | 0.460129 | 0.505241 | 0.488954 | 0.496172 | 0.463673 | 0.626447 | 0.772810 | 0.613582 | 0.625590 | 1.080314 | 1.315758 | 1.244197 | 1.079462 | 0.886251 | 1.158250 | 1.153961 | 1.114290 | 0.933266 | 0.965855 | 0.850861 | 0.905237 | 0.815321 | 0.874169 | 0.612871 | 0.419934 | 0.386850 | 0.468558 | 0.541926 | 0.626619 | 0.498790 | 0.581789 | 0.655505 | 0.971532 | 0.805101 | 0.959872 | 1.111358 | 1.661964 | 1.711915 | 2.392857 | 1.744003 | 1.684051 | 1.233249 | 0.721045 | 0.447937 | 0.269977 | 0.216713 | 0.217287 | 0.221408 | 0.227624 | 0.217099 | 0.215614 | 0.214726 | 0.216448 | 0.214890 | 0.215189 | 0.215896 | 0.216590 | 0.218401 | 0.218055 | 0.215914 | 0.214661 | 0.215996 | 0.223827 | 0.228242 | 0.221317 | 0.215843 | 0.216402 | 0.214658 | 0.216067 | 0.224630 | 0.219071 | 0.217278 | 0.217985 | 0.223368 | 0.236509 | 0.273972 | 0.330608 | 0.407525 | 0.379402 | 0.314472 | 0.270282 | 0.265833 | 0.255175 | 0.241624 | 0.233105 | 0.249145 | 0.301247 | 0.408724 | 0.342633 | 0.316144 | 0.285111 | 0.247648 | 0.246313 | 0.273126 | 0.272860 | 0.302570 | 0.277418 | 0.297710 | 0.366008 | 0.309068 | 0.225250 | 0.215214 | 0.221269 | 0.217857 | 0.224564 | 0.216053 | 0.216582 | 0.214626 | 0.219172 | 0.221904 | 0.214695 | 0.215436 | 0.214661 | 0.216360 | 0.215069 | 0.214643 | 0.214626 | 0.214656 |
| right anterior | 0.549351 | 0.458402 | 0.282999 | 0.229205 | 0.214626 | 0.215913 | 0.221130 | 0.215854 | 0.255873 | 0.582629 | 3.109733 | 7.408837 | 3.129068 | 2.537603 | 1.133846 | 0.451438 | 0.293041 | 0.228170 | 0.284741 | 0.501845 | 0.370873 | 0.325496 | 0.318184 | 0.335044 | 0.358234 | 0.240558 | 0.216386 | 0.219883 | 0.235288 | 0.291060 | 0.330486 | 0.302284 | 0.339046 | 0.281034 | 0.281924 | 0.266104 | 0.294475 | 0.251846 | 0.281093 | 0.380521 | 0.993841 | 3.583552 | 8.866243 | 15.422982 | 32.167119 | 30.456800 | 11.456375 | 3.255432 | 0.729099 | 0.422574 | 0.281010 | 0.247938 | 0.243666 | 0.236005 | 0.258793 | 0.275343 | 0.361015 | 0.666413 | 0.663471 | 0.592587 | 0.383391 | 0.267853 | 0.256404 | 0.253682 | 0.219966 | 0.216604 | 0.214689 | 0.216989 | 0.219718 | 0.241370 | 0.329562 | 0.391239 | 0.522193 | 0.624961 | 0.712764 | 0.665386 | 0.406193 | 0.275701 | 0.245353 | 0.250419 | 0.264467 | 0.254073 | 0.239498 | 0.293428 | 0.415923 | 0.465530 | 0.449908 | 0.559497 | 0.514217 | 0.571627 | 0.763500 | 0.828266 | 0.686282 | 0.395747 | 0.287659 | 0.291493 | 0.253433 | 0.246821 | 0.238397 | 0.227305 | 0.235750 | 0.214638 | 0.232682 | 0.216702 | 0.214626 | 0.217705 | 0.230283 | 0.239271 | 0.404216 | 0.857682 | 0.541392 | 0.290394 | 0.247131 | 0.226817 | 0.215231 | 0.244489 | 0.237543 | 0.239457 | 0.215857 | 0.233508 | 0.429640 | 0.609919 | 0.651301 | 0.419653 | 0.419115 | 0.430686 | 0.489427 | 0.289425 | 0.333916 | 0.392889 | 0.572389 | 0.625729 | 0.450361 | 0.355949 | 0.307748 | 0.254725 | 0.225529 | 0.216638 | 0.230563 | 0.233496 | 0.270420 | 0.239623 | 0.228342 | 0.226588 | 0.235318 | 0.252682 | 0.274357 | 0.232993 | 0.230506 | 0.249391 | 0.281707 | 0.266187 | 0.271479 | 0.243215 | 0.250469 | 0.241797 | 0.225667 | 0.220017 | 0.287044 | 0.383667 | 0.456320 | 0.497211 | 0.498583 | 0.568781 | 0.457579 | 0.456817 | 0.479875 | 0.542083 | 0.556783 | 0.521600 | 0.661852 | 0.912657 | 0.674866 | 0.503689 | 0.350441 | 0.259847 | 0.225499 | 0.219323 | 0.273385 | 0.300506 | 0.284815 | 0.282902 | 0.259821 | 0.247653 | 0.241842 | 0.248966 | 0.272378 | 0.363766 | 0.330933 | 0.312623 | 0.285520 | 0.242839 | 0.248921 | 0.240577 | 0.241240 | 0.241392 | 0.223512 | 0.224210 | 0.261562 | 0.263725 | 0.254046 | 0.226949 | 0.228086 | 0.233550 | 0.233304 | 0.234831 | 0.224597 | 0.223958 | 0.222767 | 0.215395 | 0.214849 | 0.214692 | 0.215152 | 0.214832 | 0.218341 | 0.247166 | 0.280719 | 0.248966 | 0.219517 | 0.214626 | 0.220345 | 0.217233 | 0.216206 | 0.218500 | 0.221993 | 0.228096 | 0.241995 | 0.230020 | 0.224570 | 0.215455 | 0.214887 | 0.223906 | 0.220723 | 0.243433 | 0.277975 | 0.349454 | 0.403913 | 0.390062 | 0.290156 | 0.255002 | 0.228920 | 0.226228 | 0.225914 | 0.223128 | 0.229563 | 0.267235 | 0.344724 | 0.470169 | 0.464537 | 0.388198 | 0.386675 | 0.360581 | 0.247660 | 0.215716 | 0.214645 | 0.214706 | 0.223641 | 0.304660 | 0.382086 | 0.319619 | 0.330261 | 0.359058 | 0.551213 | 0.715116 | 0.643605 | 0.387781 | 0.344311 | 0.323318 | 0.302086 | 0.265969 | 0.231055 | 0.215788 | 0.219817 | 0.224628 | 0.226421 | 0.239822 | 0.269105 | 0.281677 | 0.312574 | 0.356842 |
| left central | 0.366505 | 0.491949 | 0.399603 | 0.474037 | 0.412024 | 0.221467 | 0.215825 | 0.235939 | 0.235947 | 0.227389 | 0.246484 | 0.258320 | 0.341503 | 0.572111 | 0.968226 | 4.962980 | 3.083892 | 1.660129 | 2.609302 | 0.916524 | 0.363156 | 0.214844 | 0.252207 | 0.227539 | 0.255467 | 0.448965 | 0.714409 | 0.500344 | 0.368747 | 0.264114 | 0.215098 | 0.266428 | 0.385088 | 0.362742 | 0.250207 | 0.215716 | 0.215645 | 0.240340 | 0.294569 | 0.504037 | 0.923504 | 1.672232 | 1.443671 | 1.658318 | 1.783389 | 1.207946 | 0.775877 | 0.391702 | 0.252899 | 0.233857 | 0.235584 | 0.214760 | 0.216814 | 0.218031 | 0.227752 | 0.225491 | 0.248203 | 0.342760 | 0.525469 | 0.534464 | 0.338955 | 0.300239 | 0.241332 | 0.214867 | 0.271687 | 0.306830 | 0.336574 | 0.315484 | 0.359539 | 0.346361 | 0.299970 | 0.319872 | 0.504224 | 1.538232 | 3.701735 | 3.333596 | 1.208048 | 0.884259 | 0.783273 | 0.682893 | 0.385619 | 0.314047 | 0.270020 | 0.338653 | 0.614936 | 1.125324 | 1.315744 | 1.968546 | 3.337166 | 5.871559 | 6.013198 | 1.037088 | 0.448365 | 0.334601 | 0.278882 | 0.219217 | 0.214626 | 0.214626 | 0.262110 | 0.281787 | 0.343291 | 0.466835 | 0.598531 | 0.713788 | 0.688863 | 0.448461 | 0.456708 | 0.522020 | 0.356550 | 0.308073 | 0.281371 | 0.247016 | 0.252568 | 0.254184 | 0.279807 | 0.334898 | 0.488686 | 0.517036 | 0.766361 | 0.987634 | 0.947705 | 0.698114 | 0.795373 | 0.843809 | 0.786936 | 0.887528 | 0.847916 | 0.908231 | 1.010434 | 0.945855 | 0.811776 | 0.732954 | 0.491828 | 0.625592 | 1.138608 | 2.492321 | 4.043352 | 5.395934 | 21.395951 | 42.935192 | 76.127023 | 37.707319 | 14.058517 | 12.556791 | 9.266693 | 4.980931 | 5.600142 | 4.158768 | 2.856309 | 1.579063 | 1.070114 | 1.203434 | 1.503988 | 0.951823 | 0.991367 | 1.019330 | 1.231634 | 1.380676 | 1.182968 | 0.715829 | 0.929882 | 0.912350 | 0.858945 | 0.693575 | 0.587034 | 0.499077 | 0.452617 | 0.457100 | 0.439906 | 0.552829 | 0.685416 | 0.822819 | 1.396806 | 2.813352 | 4.673183 | 10.917338 | 10.195124 | 18.219653 | 16.608196 | 16.892645 | 10.395500 | 7.131947 | 10.627562 | 23.301309 | 22.363692 | 8.513249 | 2.454507 | 1.499734 | 0.720726 | 0.300198 | 0.217048 | 0.218887 | 0.218789 | 0.216524 | 0.232502 | 0.253571 | 0.221830 | 0.215121 | 0.215191 | 0.217291 | 0.276861 | 0.586144 | 1.101428 | 0.665380 | 0.539868 | 0.607042 | 0.613481 | 0.515569 | 0.373779 | 0.288441 | 0.312131 | 0.416842 | 0.572371 | 0.667161 | 0.617751 | 0.681841 | 0.618922 | 0.452518 | 0.357900 | 0.315413 | 0.350398 | 0.345741 | 0.275670 | 0.294483 | 0.306611 | 0.305833 | 0.239946 | 0.215869 | 0.216549 | 0.219671 | 0.221296 | 0.215230 | 0.229591 | 0.224439 | 0.216791 | 0.216127 | 0.214876 | 0.214663 | 0.215643 | 0.227402 | 0.219443 | 0.220365 | 0.225321 | 0.223701 | 0.216002 | 0.214635 | 0.222426 | 0.242221 | 0.359803 | 0.669173 | 1.130460 | 1.690278 | 2.411360 | 2.176178 | 1.652819 | 1.286638 | 0.719821 | 0.536508 | 0.358084 | 0.296140 | 0.256021 | 0.259740 | 0.263635 | 0.321211 | 0.302839 | 0.349305 | 0.389506 | 0.447882 | 0.474287 | 0.496342 | 0.623453 | 1.386556 | 2.022281 | 2.361237 | 2.553987 | 2.160999 | 1.967999 | 1.495016 | 0.969634 | 0.687044 |
| right central | 0.566428 | 1.070115 | 0.768653 | 0.715465 | 0.525810 | 0.486465 | 0.363302 | 0.277603 | 0.224492 | 0.217242 | 0.272906 | 0.446264 | 0.677169 | 1.142338 | 1.409777 | 1.353369 | 2.028158 | 0.783934 | 0.353733 | 0.277285 | 0.230043 | 0.214878 | 0.243994 | 0.298726 | 0.316347 | 0.254876 | 0.215266 | 0.324565 | 0.477694 | 0.932394 | 0.998390 | 0.878880 | 0.775029 | 0.531998 | 0.444846 | 0.373093 | 0.395138 | 0.772535 | 0.991019 | 0.997269 | 1.733306 | 1.790567 | 1.784610 | 1.766713 | 0.787325 | 0.520592 | 0.363467 | 0.243395 | 0.215902 | 0.215064 | 0.222940 | 0.214686 | 0.215435 | 0.252697 | 0.301864 | 0.428998 | 0.529718 | 0.659945 | 1.240550 | 1.895065 | 1.100849 | 0.967738 | 0.442795 | 0.445297 | 0.438418 | 0.605808 | 1.032160 | 3.085886 | 3.545222 | 10.847900 | 8.797063 | 5.607414 | 5.153458 | 2.464234 | 0.617781 | 0.522151 | 0.265926 | 0.231524 | 0.221199 | 0.214716 | 0.218393 | 0.229092 | 0.241658 | 0.222094 | 0.228566 | 0.301245 | 0.403813 | 0.325978 | 0.307014 | 0.375507 | 0.458489 | 0.371286 | 0.272548 | 0.259390 | 0.333634 | 0.344111 | 0.327673 | 0.238241 | 0.219421 | 0.214632 | 0.222854 | 0.280399 | 0.549375 | 1.650484 | 4.100864 | 4.462833 | 3.606066 | 1.711543 | 0.908505 | 0.455556 | 0.331937 | 0.275526 | 0.287166 | 0.308251 | 0.426417 | 0.551167 | 0.517547 | 0.425486 | 0.411459 | 0.436792 | 0.415131 | 0.438967 | 0.508961 | 0.825894 | 0.988073 | 0.795066 | 0.576294 | 0.571775 | 0.350454 | 0.263137 | 0.232671 | 0.237100 | 0.239083 | 0.241238 | 0.227872 | 0.257785 | 0.348136 | 0.367571 | 0.483953 | 0.522120 | 0.525251 | 0.927399 | 1.503096 | 1.049946 | 0.524276 | 0.332348 | 0.249374 | 0.250020 | 0.220263 | 0.214626 | 0.215960 | 0.218892 | 0.236639 | 0.282199 | 0.310677 | 0.444100 | 0.625856 | 0.931967 | 0.839820 | 0.449825 | 0.322589 | 0.250933 | 0.245580 | 0.251561 | 0.252705 | 0.273158 | 0.415846 | 0.623387 | 0.844092 | 0.591827 | 0.488129 | 0.419163 | 0.329413 | 0.274867 | 0.281681 | 0.283153 | 0.305739 | 0.325171 | 0.351491 | 0.421227 | 0.483083 | 0.547686 | 0.656810 | 0.799408 | 0.937637 | 1.374391 | 2.107290 | 2.894106 | 2.171032 | 2.055606 | 1.621646 | 1.100962 | 0.943267 | 0.741796 | 0.547420 | 0.519017 | 0.491006 | 0.447934 | 0.339255 | 0.302059 | 0.339637 | 0.353267 | 0.356624 | 0.377416 | 0.486757 | 0.787343 | 0.867588 | 0.835170 | 0.748406 | 0.688730 | 0.740258 | 0.597480 | 0.560282 | 0.620113 | 0.588285 | 0.808649 | 1.268536 | 1.870619 | 3.937409 | 6.250900 | 4.669349 | 3.413261 | 1.630032 | 1.069139 | 0.587585 | 0.357028 | 0.294791 | 0.301459 | 0.428699 | 0.617764 | 1.113045 | 1.854671 | 1.887468 | 1.630320 | 2.619799 | 1.722027 | 0.986264 | 0.473868 | 0.439225 | 0.487591 | 0.648822 | 0.675732 | 0.961348 | 1.313881 | 1.219559 | 1.046236 | 0.798426 | 0.615062 | 0.459933 | 0.292351 | 0.228732 | 0.215311 | 0.224162 | 0.234423 | 0.233284 | 0.224068 | 0.214726 | 0.222987 | 0.237051 | 0.334525 | 0.419046 | 0.473972 | 0.516017 | 0.495808 | 0.558361 | 0.692305 | 0.636849 | 0.608331 | 0.350247 | 0.226407 | 0.214636 | 0.214752 | 0.215613 | 0.219131 | 0.232617 | 0.229517 | 0.228818 | 0.229429 | 0.239539 | 0.262284 |
| left posterior | 0.215178 | 0.217906 | 0.233432 | 0.245784 | 0.285363 | 0.299865 | 0.342464 | 0.314033 | 0.291021 | 0.235828 | 0.216886 | 0.315186 | 0.383104 | 0.522367 | 0.598072 | 0.676173 | 1.113827 | 1.219838 | 1.071354 | 0.546134 | 0.449675 | 0.351243 | 0.277661 | 0.224072 | 0.215735 | 0.232592 | 0.225803 | 0.280287 | 0.415460 | 0.366134 | 0.472433 | 0.828048 | 1.174337 | 2.332411 | 2.792227 | 0.908170 | 0.583794 | 0.276969 | 0.241266 | 0.469594 | 0.690645 | 0.766427 | 0.359734 | 0.224873 | 0.255349 | 0.729996 | 2.339254 | 2.677850 | 3.032039 | 2.062450 | 2.175007 | 1.427719 | 0.958995 | 1.004339 | 1.662387 | 2.209505 | 2.985976 | 1.759826 | 2.216527 | 1.993837 | 1.948058 | 1.930802 | 3.289905 | 5.511700 | 9.127857 | 9.371520 | 5.738148 | 5.174533 | 3.011494 | 1.128503 | 0.497989 | 0.306487 | 0.251423 | 0.238330 | 0.274174 | 0.498902 | 0.620442 | 0.667956 | 0.642964 | 0.486846 | 0.421595 | 0.311344 | 0.214795 | 0.230859 | 0.236069 | 0.219577 | 0.222393 | 0.222589 | 0.226088 | 0.220628 | 0.247530 | 0.262620 | 0.250054 | 0.258015 | 0.252811 | 0.229277 | 0.214694 | 0.222765 | 0.247635 | 0.273935 | 0.262180 | 0.237683 | 0.223026 | 0.215356 | 0.222182 | 0.245386 | 0.273779 | 0.301811 | 0.309039 | 0.326749 | 0.351335 | 0.320279 | 0.287129 | 0.310800 | 0.335514 | 0.325306 | 0.354742 | 0.404746 | 0.586515 | 0.697956 | 0.465810 | 0.333517 | 0.347929 | 0.301212 | 0.247837 | 0.217151 | 0.216332 | 0.215540 | 0.214786 | 0.215260 | 0.215391 | 0.217247 | 0.219280 | 0.221027 | 0.246856 | 0.288090 | 0.333194 | 0.379125 | 0.357726 | 0.407528 | 0.402165 | 0.309114 | 0.262562 | 0.249658 | 0.252148 | 0.275298 | 0.330076 | 0.396595 | 0.505209 | 0.619529 | 0.768950 | 0.731960 | 0.618825 | 0.537308 | 0.747411 | 1.354881 | 2.612651 | 1.940789 | 1.453620 | 1.150989 | 1.044132 | 0.702853 | 0.558315 | 0.497303 | 0.556984 | 0.880371 | 1.215467 | 1.384148 | 1.080096 | 0.932659 | 0.677802 | 0.529902 | 0.336656 | 0.260359 | 0.223554 | 0.218876 | 0.214634 | 0.214989 | 0.214692 | 0.214959 | 0.214845 | 0.219455 | 0.236659 | 0.272011 | 0.325597 | 0.309932 | 0.344857 | 0.329566 | 0.330599 | 0.276448 | 0.232353 | 0.222226 | 0.232899 | 0.241181 | 0.239530 | 0.259195 | 0.363031 | 0.562914 | 0.930951 | 1.141614 | 0.988781 | 1.387270 | 1.868803 | 1.402125 | 0.968304 | 0.620395 | 0.511915 | 0.430915 | 0.342284 | 0.281766 | 0.237029 | 0.224037 | 0.219051 | 0.218832 | 0.222048 | 0.230712 | 0.242496 | 0.290789 | 0.384836 | 0.627853 | 0.880971 | 0.867405 | 0.624274 | 0.562747 | 0.445110 | 0.433295 | 0.413720 | 0.378790 | 0.489936 | 0.627369 | 0.769874 | 0.883790 | 0.731088 | 0.616864 | 0.696449 | 0.732762 | 0.868136 | 0.698997 | 0.807951 | 0.693902 | 0.775451 | 0.735983 | 0.530129 | 0.428850 | 0.451470 | 0.410506 | 0.620313 | 0.597130 | 0.521967 | 0.569622 | 0.787997 | 1.217512 | 1.954300 | 2.209332 | 2.967452 | 6.270222 | 12.349805 | 4.814842 | 1.909737 | 0.698952 | 0.463860 | 0.478660 | 0.467296 | 0.488552 | 0.545344 | 0.525389 | 0.457976 | 0.425696 | 0.336313 | 0.249123 | 0.229144 | 0.220526 | 0.221348 | 0.218204 | 0.215825 | 0.215636 | 0.217697 | 0.217037 | 0.219046 | 0.215089 |
| right posterior | 0.381631 | 0.332063 | 0.485922 | 0.453580 | 0.228621 | 0.221213 | 0.240648 | 0.222071 | 0.216400 | 0.254683 | 0.248725 | 0.215873 | 0.306463 | 0.515832 | 0.701870 | 0.698683 | 0.824596 | 0.560996 | 0.292154 | 0.265695 | 0.495654 | 1.037522 | 0.602557 | 0.313506 | 0.230619 | 0.215910 | 0.227105 | 0.242269 | 0.328444 | 0.252564 | 0.225002 | 0.228477 | 0.220311 | 0.215043 | 0.216190 | 0.214645 | 0.216969 | 0.234106 | 0.250964 | 0.273081 | 0.426758 | 0.346424 | 0.385448 | 0.689167 | 0.664038 | 0.533842 | 0.713124 | 0.497496 | 0.397930 | 0.391200 | 0.306940 | 0.252238 | 0.217923 | 0.241813 | 0.243451 | 0.231607 | 0.252212 | 0.255525 | 0.227530 | 0.214644 | 0.216014 | 0.229923 | 0.231929 | 0.223376 | 0.223810 | 0.226347 | 0.236997 | 0.248790 | 0.218046 | 0.216014 | 0.243803 | 0.345947 | 0.407478 | 0.469856 | 0.537461 | 0.545336 | 0.451780 | 0.346431 | 0.269181 | 0.261033 | 0.222350 | 0.214973 | 0.221148 | 0.215524 | 0.216551 | 0.220901 | 0.235470 | 0.272552 | 0.280571 | 0.440749 | 0.666418 | 0.932436 | 1.469703 | 0.942379 | 0.845559 | 1.945143 | 2.473179 | 2.783057 | 2.281670 | 1.238572 | 1.508103 | 1.767206 | 0.971796 | 0.583880 | 0.443548 | 0.324105 | 0.304534 | 0.284280 | 0.271440 | 0.261639 | 0.268077 | 0.245469 | 0.267373 | 0.311709 | 0.305401 | 0.294010 | 0.351891 | 0.537014 | 0.770557 | 0.867451 | 0.917626 | 0.865850 | 1.359060 | 2.510539 | 1.385801 | 1.227735 | 1.617192 | 1.565174 | 1.728031 | 1.226486 | 0.578641 | 0.680688 | 0.818450 | 1.006287 | 0.980607 | 0.911407 | 1.036903 | 1.271006 | 1.476828 | 1.275779 | 0.904997 | 0.957194 | 0.752032 | 0.649914 | 0.621568 | 0.573857 | 0.569961 | 0.510037 | 0.483247 | 0.386248 | 0.316001 | 0.292073 | 0.312825 | 0.310369 | 0.378142 | 0.518450 | 1.088446 | 1.835066 | 2.827448 | 2.571566 | 1.835477 | 0.955009 | 0.402322 | 0.299056 | 0.337020 | 0.354519 | 0.443919 | 0.456337 | 0.475877 | 0.604127 | 0.659963 | 0.594257 | 0.417824 | 0.252059 | 0.237684 | 0.242248 | 0.241421 | 0.235541 | 0.223652 | 0.216800 | 0.226609 | 0.270635 | 0.435690 | 0.615051 | 0.664808 | 0.640251 | 0.542122 | 0.416905 | 0.360393 | 0.326743 | 0.356992 | 0.380875 | 0.372261 | 0.354951 | 0.321264 | 0.306759 | 0.285774 | 0.286550 | 0.311065 | 0.351601 | 0.400321 | 0.432760 | 0.422794 | 0.408508 | 0.459564 | 0.535927 | 0.517513 | 0.621332 | 0.826887 | 0.758178 | 0.713558 | 0.480836 | 0.339351 | 0.272029 | 0.231385 | 0.219871 | 0.216962 | 0.214740 | 0.217401 | 0.223405 | 0.218948 | 0.215241 | 0.214906 | 0.223718 | 0.246974 | 0.294962 | 0.360862 | 0.339534 | 0.312277 | 0.320378 | 0.290983 | 0.270901 | 0.249984 | 0.231140 | 0.238055 | 0.242687 | 0.223849 | 0.223279 | 0.217271 | 0.216234 | 0.222258 | 0.243591 | 0.246976 | 0.243091 | 0.252637 | 0.289919 | 0.326760 | 0.328659 | 0.309250 | 0.331713 | 0.366068 | 0.322128 | 0.263982 | 0.225329 | 0.214844 | 0.214695 | 0.214634 | 0.214860 | 0.218746 | 0.224056 | 0.231595 | 0.258970 | 0.264946 | 0.263371 | 0.263601 | 0.234314 | 0.217264 | 0.216172 | 0.217318 | 0.229193 | 0.257184 | 0.306323 | 0.412907 | 0.571702 | 1.000825 | 1.641913 | 2.140571 | 3.151870 | 3.542919 | 2.078275 |
| all electrodes | 0.462252 | 0.218909 | 0.472480 | 1.359804 | 15.850230 | 82.641021 | 178.657507 | 45.170817 | 4.327298 | 0.726260 | 0.346197 | 0.235390 | 0.218400 | 0.225082 | 0.219579 | 0.222581 | 0.220915 | 0.215741 | 0.239080 | 0.260036 | 0.290486 | 0.294497 | 0.283306 | 0.459655 | 1.371475 | 0.704982 | 0.549641 | 0.647316 | 0.521360 | 0.299850 | 0.256602 | 0.329644 | 0.695617 | 0.907853 | 0.802727 | 0.542846 | 1.005669 | 0.694546 | 0.321983 | 0.215614 | 0.233569 | 0.299015 | 0.343697 | 0.455073 | 0.456215 | 0.419415 | 0.354014 | 0.273711 | 0.216891 | 0.229678 | 0.251083 | 0.276290 | 0.431335 | 1.119266 | 1.017525 | 0.528959 | 0.506352 | 0.434833 | 0.273449 | 0.214809 | 0.233424 | 0.226231 | 0.232386 | 0.227658 | 0.224600 | 0.215131 | 0.228672 | 0.263289 | 0.352370 | 0.469995 | 0.564594 | 0.488632 | 0.580638 | 0.564679 | 0.487743 | 0.319302 | 0.249274 | 0.247278 | 0.296076 | 0.353266 | 0.610621 | 0.454523 | 0.517278 | 0.943077 | 1.245916 | 1.591481 | 1.406815 | 1.268125 | 2.059914 | 2.885395 | 1.614933 | 0.931039 | 0.771443 | 0.493345 | 0.319877 | 0.255894 | 0.219810 | 0.230784 | 0.262907 | 0.373070 | 0.514015 | 0.775277 | 1.081505 | 1.635965 | 1.472218 | 2.101463 | 6.995585 | 10.605741 | 8.641110 | 8.568588 | 7.925019 | 18.243031 | 22.524723 | 5.756393 | 1.823275 | 1.863891 | 1.301070 | 1.014093 | 0.637307 | 0.561717 | 0.617718 | 0.683937 | 0.470045 | 0.444581 | 0.402439 | 0.482786 | 0.456585 | 0.313846 | 0.257744 | 0.229382 | 0.216214 | 0.215196 | 0.214810 | 0.215235 | 0.221664 | 0.234677 | 0.258774 | 0.297828 | 0.408954 | 0.426704 | 0.349185 | 0.340762 | 0.324391 | 0.349002 | 0.304272 | 0.272337 | 0.339043 | 0.535146 | 0.536690 | 0.580038 | 0.557052 | 0.994570 | 1.332831 | 1.552592 | 1.429996 | 5.540408 | 13.219002 | 28.765186 | 23.361281 | 17.541938 | 8.718113 | 4.905015 | 1.819363 | 1.567521 | 2.448492 | 3.282069 | 4.515494 | 4.529436 | 4.728337 | 8.291315 | 9.360538 | 4.670617 | 4.175159 | 4.431351 | 7.163742 | 9.122077 | 7.250581 | 11.959699 | 14.819947 | 11.021387 | 14.199280 | 9.482732 | 12.512638 | 8.056191 | 10.265835 | 13.677460 | 7.496598 | 4.350347 | 3.399905 | 3.650541 | 4.259337 | 1.637598 | 1.629784 | 2.648389 | 2.130958 | 1.547583 | 1.917247 | 1.603375 | 1.577142 | 1.306711 | 2.392042 | 4.095768 | 6.924207 | 5.109075 | 4.710415 | 5.527522 | 2.864403 | 0.945616 | 0.814573 | 0.817211 | 1.305902 | 1.732811 | 2.289226 | 3.516054 | 4.130228 | 5.126793 | 9.319325 | 8.522972 | 4.976185 | 1.610617 | 1.161761 | 1.053575 | 1.111388 | 1.622163 | 2.007639 | 4.986850 | 12.530704 | 13.090230 | 10.060780 | 4.486690 | 1.338673 | 0.710213 | 0.437676 | 0.319638 | 0.273270 | 0.279453 | 0.300708 | 0.404553 | 0.624402 | 0.975428 | 2.191054 | 4.759523 | 8.708196 | 10.718636 | 6.189775 | 3.691224 | 2.065105 | 1.460776 | 1.119262 | 0.862864 | 0.408339 | 0.328893 | 0.287424 | 0.259698 | 0.267465 | 0.273133 | 0.265952 | 0.318850 | 0.410243 | 0.419838 | 0.543686 | 0.394834 | 0.309954 | 0.249025 | 0.256325 | 0.285838 | 0.278781 | 0.242078 | 0.233619 | 0.232691 | 0.234848 | 0.233895 | 0.215574 | 0.214664 | 0.216966 | 0.221118 | 0.230753 | 0.234185 | 0.278731 | 0.296531 |

Searchlight, spatiotemporal cluster permutation test

No significant clusters observed.

F) difference, identity, happy vs sad

  
|  | time window | peak latency | cluster *p* | peak Cohen's *d* |  | | | |  |
| **all electrodes** | 260 - 345 ms | 300 ms | 0.0345 | 0.5879 |  | | | |  |
 625 - 710 ms | 650 ms | 0.0215 | 0.7639 |  | | | | 775 - 865 ms | 830 ms | 0.0286 | 0.7068 |  | | | | 880 - 955 ms | 940 ms | 0.0162 | 1.0248 |  | | | ||  |  | | | | | | | | |

Time-resolved classification, cluster permutation tests

|  |  | **left hemisphere** | | | | **right hemisphere** | | | |
|  |  | time window | peak latency | cluster *p* | peak Cohen's *d* | time window | peak latency | cluster *p* | peak Cohen's *d* |
| **anterior** |  | | | |  | | | |  |
| **central** | 170 - 270 ms | 190 ms | 0.0253 | 0.775 |  | | | |  |
| **posterior** |  | | | |  | | | |  |

  

Time-resolved classification, Bayesian statistics

|  | -200 | -195 | -190 | -185 | -180 | -175 | -170 | -165 | -160 | -155 | -150 | -145 | -140 | -135 | -130 | -125 | -120 | -115 | -110 | -105 | -100 | -95 | -90 | -85 | -80 | -75 | -70 | -65 | -60 | -55 | -50 | -45 | -40 | -35 | -30 | -25 | -20 | -15 | -10 | -5 | 0 | 5 | 10 | 15 | 20 | 25 | 30 | 35 | 40 | 45 | 50 | 55 | 60 | 65 | 70 | 75 | 80 | 85 | 90 | 95 | 100 | 105 | 110 | 115 | 120 | 125 | 130 | 135 | 140 | 145 | 150 | 155 | 160 | 165 | 170 | 175 | 180 | 185 | 190 | 195 | 200 | 205 | 210 | 215 | 220 | 225 | 230 | 235 | 240 | 245 | 250 | 255 | 260 | 265 | 270 | 275 | 280 | 285 | 290 | 295 | 300 | 305 | 310 | 315 | 320 | 325 | 330 | 335 | 340 | 345 | 350 | 355 | 360 | 365 | 370 | 375 | 380 | 385 | 390 | 395 | 400 | 405 | 410 | 415 | 420 | 425 | 430 | 435 | 440 | 445 | 450 | 455 | 460 | 465 | 470 | 475 | 480 | 485 | 490 | 495 | 500 | 505 | 510 | 515 | 520 | 525 | 530 | 535 | 540 | 545 | 550 | 555 | 560 | 565 | 570 | 575 | 580 | 585 | 590 | 595 | 600 | 605 | 610 | 615 | 620 | 625 | 630 | 635 | 640 | 645 | 650 | 655 | 660 | 665 | 670 | 675 | 680 | 685 | 690 | 695 | 700 | 705 | 710 | 715 | 720 | 725 | 730 | 735 | 740 | 745 | 750 | 755 | 760 | 765 | 770 | 775 | 780 | 785 | 790 | 795 | 800 | 805 | 810 | 815 | 820 | 825 | 830 | 835 | 840 | 845 | 850 | 855 | 860 | 865 | 870 | 875 | 880 | 885 | 890 | 895 | 900 | 905 | 910 | 915 | 920 | 925 | 930 | 935 | 940 | 945 | 950 | 955 | 960 | 965 | 970 | 975 | 980 | 985 | 990 | 995 | 1000 | 1005 | 1010 | 1015 | 1020 | 1025 | 1030 | 1035 | 1040 | 1045 | 1050 | 1055 | 1060 | 1065 | 1070 | 1075 | 1080 | 1085 | 1090 | 1095 | 1100 | 1105 | 1110 | 1115 | 1120 | 1125 | 1130 | 1135 | 1140 | 1145 | 1150 | 1155 | 1160 | 1165 | 1170 | 1175 | 1180 | 1185 | 1190 | 1195 |
| --- | --- | --- | --- | --- | --- | --- | --- | --- | --- | --- | --- | --- | --- | --- | --- | --- | --- | --- | --- | --- | --- | --- | --- | --- | --- | --- | --- | --- | --- | --- | --- | --- | --- | --- | --- | --- | --- | --- | --- | --- | --- | --- | --- | --- | --- | --- | --- | --- | --- | --- | --- | --- | --- | --- | --- | --- | --- | --- | --- | --- | --- | --- | --- | --- | --- | --- | --- | --- | --- | --- | --- | --- | --- | --- | --- | --- | --- | --- | --- | --- | --- | --- | --- | --- | --- | --- | --- | --- | --- | --- | --- | --- | --- | --- | --- | --- | --- | --- | --- | --- | --- | --- | --- | --- | --- | --- | --- | --- | --- | --- | --- | --- | --- | --- | --- | --- | --- | --- | --- | --- | --- | --- | --- | --- | --- | --- | --- | --- | --- | --- | --- | --- | --- | --- | --- | --- | --- | --- | --- | --- | --- | --- | --- | --- | --- | --- | --- | --- | --- | --- | --- | --- | --- | --- | --- | --- | --- | --- | --- | --- | --- | --- | --- | --- | --- | --- | --- | --- | --- | --- | --- | --- | --- | --- | --- | --- | --- | --- | --- | --- | --- | --- | --- | --- | --- | --- | --- | --- | --- | --- | --- | --- | --- | --- | --- | --- | --- | --- | --- | --- | --- | --- | --- | --- | --- | --- | --- | --- | --- | --- | --- | --- | --- | --- | --- | --- | --- | --- | --- | --- | --- | --- | --- | --- | --- | --- | --- | --- | --- | --- | --- | --- | --- | --- | --- | --- | --- | --- | --- | --- | --- | --- | --- | --- | --- | --- | --- | --- | --- | --- | --- | --- | --- | --- | --- | --- | --- | --- | --- | --- | --- | --- | --- | --- | --- | --- | --- | --- | --- | --- | --- | --- | --- | --- | --- | --- | --- | --- | --- | --- |
| left anterior | 0.440927 | 0.695666 | 0.607640 | 0.806521 | 0.511365 | 0.318156 | 0.218172 | 0.284025 | 0.747631 | 0.595877 | 0.614776 | 0.453238 | 0.661530 | 0.313569 | 0.230380 | 0.214840 | 0.244876 | 0.268953 | 0.243961 | 0.260168 | 0.311133 | 0.276731 | 0.256775 | 0.215225 | 0.222726 | 0.233040 | 0.236225 | 0.216565 | 0.229934 | 0.268027 | 0.262734 | 0.220390 | 0.215185 | 0.215135 | 0.217075 | 0.233080 | 0.217348 | 0.261616 | 0.446554 | 0.870017 | 0.723973 | 0.329489 | 0.302361 | 0.250007 | 0.223119 | 0.242975 | 0.296776 | 0.248205 | 0.215109 | 0.215672 | 0.215481 | 0.216266 | 0.221117 | 0.216678 | 0.214868 | 0.215023 | 0.227759 | 0.264616 | 0.321084 | 0.307953 | 0.296385 | 0.310338 | 0.256989 | 0.228251 | 0.216854 | 0.233604 | 0.224739 | 0.230900 | 0.347467 | 0.425140 | 0.461968 | 0.428624 | 0.382745 | 0.464266 | 0.442530 | 0.361982 | 0.351442 | 0.417750 | 0.562454 | 0.734634 | 0.768982 | 0.522511 | 0.290352 | 0.216987 | 0.227598 | 0.271650 | 0.513276 | 0.815179 | 0.927605 | 1.046904 | 1.840881 | 1.551360 | 1.222436 | 0.505613 | 0.359882 | 0.282835 | 0.253783 | 0.222324 | 0.214846 | 0.220130 | 0.218622 | 0.214626 | 0.216111 | 0.220009 | 0.215999 | 0.243521 | 0.506151 | 1.293811 | 1.105997 | 1.598813 | 3.626723 | 9.087553 | 21.611184 | 5.268780 | 1.137630 | 0.869464 | 0.460029 | 0.286813 | 0.216312 | 0.226248 | 0.257716 | 0.228670 | 0.228230 | 0.231340 | 0.215127 | 0.248203 | 0.298330 | 0.325811 | 0.265940 | 0.251751 | 0.237632 | 0.214748 | 0.231118 | 0.238149 | 0.219426 | 0.231952 | 0.303680 | 0.319191 | 0.264469 | 0.270577 | 0.240461 | 0.223402 | 0.216532 | 0.239733 | 0.255821 | 0.232997 | 0.232161 | 0.261773 | 0.317840 | 0.393567 | 0.391190 | 0.309012 | 0.294657 | 0.252992 | 0.214650 | 0.295833 | 0.728508 | 1.099153 | 0.801522 | 0.828048 | 0.544053 | 0.511876 | 0.309356 | 0.356803 | 0.469458 | 0.890526 | 1.139141 | 2.136239 | 1.820426 | 2.793806 | 2.019892 | 1.570747 | 1.127479 | 0.861933 | 0.777737 | 0.763670 | 0.848370 | 0.649924 | 0.545024 | 0.612214 | 0.609363 | 0.472832 | 0.461977 | 0.380585 | 0.321124 | 0.280943 | 0.242505 | 0.252420 | 0.277444 | 0.301300 | 0.354778 | 0.539124 | 0.708847 | 0.665281 | 0.528090 | 0.399903 | 0.385773 | 0.369639 | 0.381119 | 0.411235 | 0.338746 | 0.263615 | 0.258245 | 0.238665 | 0.236298 | 0.218218 | 0.216268 | 0.228545 | 0.247331 | 0.255676 | 0.240753 | 0.246050 | 0.246968 | 0.245660 | 0.237937 | 0.228789 | 0.228200 | 0.247995 | 0.246085 | 0.261651 | 0.289889 | 0.394916 | 0.562735 | 0.666783 | 0.514870 | 0.490805 | 0.414725 | 0.387269 | 0.299005 | 0.270429 | 0.271109 | 0.329166 | 0.352246 | 0.359176 | 0.298634 | 0.288121 | 0.260030 | 0.247286 | 0.237861 | 0.228821 | 0.221482 | 0.224153 | 0.217811 | 0.226026 | 0.233347 | 0.239043 | 0.236502 | 0.267173 | 0.268776 | 0.344515 | 0.338339 | 0.353901 | 0.304228 | 0.264934 | 0.247029 | 0.261638 | 0.255973 | 0.296478 | 0.284789 | 0.277298 | 0.291910 | 0.241618 | 0.216081 | 0.219469 | 0.267817 | 0.291493 | 0.344505 | 0.603552 | 0.721813 | 0.761833 | 0.752469 | 0.626719 | 0.568601 | 0.385925 | 0.290340 | 0.271550 | 0.268374 | 0.240248 | 0.237413 | 0.255274 |
| right anterior | 0.218501 | 0.221927 | 0.303875 | 0.257102 | 0.316191 | 0.272598 | 0.218679 | 0.221653 | 0.715439 | 3.123843 | 3.173430 | 1.808933 | 0.433640 | 0.356803 | 0.249735 | 0.216192 | 0.214864 | 0.228343 | 0.216635 | 0.314905 | 0.363669 | 0.342325 | 0.263513 | 0.232218 | 0.272706 | 0.312047 | 0.327224 | 0.328500 | 0.408965 | 0.416856 | 0.417963 | 0.321651 | 0.226328 | 0.217211 | 0.237085 | 0.224300 | 0.215367 | 0.215725 | 0.215657 | 0.224889 | 0.324615 | 0.934718 | 1.849840 | 2.335364 | 4.366724 | 2.704874 | 2.177672 | 0.670715 | 0.268588 | 0.214643 | 0.214779 | 0.225932 | 0.220855 | 0.222781 | 0.215239 | 0.215037 | 0.216497 | 0.215394 | 0.219769 | 0.227865 | 0.215207 | 0.222320 | 0.226833 | 0.215935 | 0.217738 | 0.221723 | 0.215501 | 0.215856 | 0.215319 | 0.221895 | 0.251993 | 0.280537 | 0.354661 | 0.499458 | 0.593333 | 0.447297 | 0.263780 | 0.226251 | 0.217223 | 0.219195 | 0.215833 | 0.215567 | 0.214746 | 0.225406 | 0.279560 | 0.356682 | 0.380415 | 0.500462 | 0.804891 | 0.817010 | 1.590538 | 2.238566 | 2.480282 | 1.076213 | 0.663887 | 0.411284 | 0.380480 | 0.309605 | 0.271090 | 0.245886 | 0.257638 | 0.239099 | 0.227621 | 0.229814 | 0.238967 | 0.234422 | 0.301980 | 0.408521 | 0.702241 | 1.670397 | 1.066474 | 0.453806 | 0.302351 | 0.214642 | 0.278811 | 0.342984 | 0.512704 | 0.465721 | 0.254778 | 0.218487 | 0.299591 | 0.355412 | 0.495059 | 0.925851 | 1.599232 | 2.135079 | 2.136076 | 0.713616 | 0.903096 | 0.433005 | 0.278264 | 0.238501 | 0.215646 | 0.214852 | 0.216746 | 0.214626 | 0.215849 | 0.216787 | 0.215380 | 0.218038 | 0.215189 | 0.220971 | 0.221109 | 0.260812 | 0.392483 | 0.588308 | 1.401730 | 0.810240 | 0.945253 | 1.771671 | 1.560388 | 0.788921 | 0.495276 | 0.234282 | 0.223876 | 0.221589 | 0.311392 | 0.536515 | 0.728886 | 0.433966 | 0.343763 | 0.318295 | 0.264835 | 0.257033 | 0.253151 | 0.302209 | 0.452162 | 0.523486 | 0.610169 | 0.699515 | 0.847733 | 0.884812 | 0.493762 | 0.358888 | 0.333350 | 0.284806 | 0.237376 | 0.214692 | 0.217691 | 0.216132 | 0.218913 | 0.263170 | 0.322405 | 0.301550 | 0.276726 | 0.287165 | 0.369869 | 0.414394 | 0.354016 | 0.283987 | 0.235571 | 0.216579 | 0.239391 | 0.250814 | 0.234726 | 0.216786 | 0.238123 | 0.226392 | 0.214909 | 0.224743 | 0.256476 | 0.248500 | 0.218672 | 0.232964 | 0.283444 | 0.292132 | 0.364212 | 0.423956 | 0.299406 | 0.260609 | 0.254230 | 0.259588 | 0.287480 | 0.299068 | 0.326376 | 0.409589 | 0.567768 | 0.500957 | 0.317560 | 0.237219 | 0.230261 | 0.215255 | 0.215201 | 0.219211 | 0.233260 | 0.223286 | 0.218373 | 0.224034 | 0.232576 | 0.248237 | 0.255936 | 0.255069 | 0.242791 | 0.247860 | 0.256702 | 0.276201 | 0.275152 | 0.337339 | 0.314962 | 0.373032 | 0.426250 | 0.420998 | 0.338670 | 0.289807 | 0.260996 | 0.287103 | 0.311124 | 0.353747 | 0.466966 | 0.466735 | 0.611778 | 0.515874 | 0.387276 | 0.277802 | 0.240266 | 0.222021 | 0.220537 | 0.214626 | 0.215049 | 0.217052 | 0.230229 | 0.225664 | 0.256503 | 0.289807 | 0.360328 | 0.341160 | 0.298953 | 0.255875 | 0.268542 | 0.271307 | 0.304738 | 0.283507 | 0.300372 | 0.337734 | 0.306895 | 0.272989 | 0.290772 | 0.277409 | 0.272246 | 0.255013 |
| left central | 0.305001 | 0.297991 | 0.388398 | 0.484086 | 0.689956 | 1.624044 | 2.562938 | 1.712276 | 0.741049 | 0.480781 | 0.300703 | 0.216704 | 0.222993 | 0.290197 | 0.256675 | 0.227207 | 0.267570 | 0.310311 | 0.353744 | 0.265854 | 0.317334 | 0.224381 | 0.216769 | 0.214648 | 0.216724 | 0.756330 | 3.947465 | 2.843222 | 2.198044 | 1.698727 | 0.852726 | 0.934718 | 0.373767 | 0.217181 | 0.214770 | 0.214641 | 0.221644 | 0.258182 | 0.246114 | 0.367723 | 0.691704 | 0.941846 | 1.401024 | 1.942838 | 2.774347 | 6.468673 | 4.197923 | 1.685009 | 0.689440 | 0.351129 | 0.262891 | 0.214872 | 0.237956 | 0.343417 | 0.443771 | 0.478239 | 0.472570 | 0.363050 | 0.218154 | 0.235105 | 0.250196 | 0.244092 | 0.217572 | 0.228505 | 0.422115 | 1.394922 | 2.533379 | 2.628430 | 2.245535 | 0.965039 | 0.559415 | 0.647989 | 0.552151 | 0.921737 | 2.142579 | 7.195660 | 10.383641 | 21.721762 | 37.328647 | 39.316769 | 41.448550 | 39.643484 | 7.945497 | 15.354229 | 24.461964 | 20.620844 | 17.379421 | 6.972795 | 4.959408 | 4.334678 | 4.463821 | 1.973579 | 1.404217 | 1.310323 | 2.014948 | 1.228313 | 2.069521 | 2.837959 | 3.127469 | 2.828609 | 2.494956 | 3.384853 | 3.378451 | 4.107048 | 1.783638 | 1.613294 | 0.921030 | 0.960427 | 0.623029 | 0.444583 | 0.320999 | 0.272151 | 0.256119 | 0.259365 | 0.254890 | 0.339769 | 0.428225 | 0.474963 | 0.964983 | 1.372880 | 1.030205 | 0.676931 | 0.588812 | 0.629172 | 0.565884 | 0.584015 | 0.543234 | 0.658945 | 0.650562 | 0.519049 | 0.668078 | 0.639910 | 0.512546 | 0.529228 | 0.787620 | 1.539389 | 2.943195 | 1.146595 | 1.106200 | 0.733945 | 0.590514 | 0.498699 | 0.549990 | 0.455471 | 0.362129 | 0.314425 | 0.346540 | 0.369665 | 0.351734 | 0.302259 | 0.329353 | 0.634703 | 0.922669 | 1.086910 | 0.964980 | 0.980069 | 0.722595 | 0.502525 | 0.325465 | 0.239306 | 0.226193 | 0.216570 | 0.214784 | 0.216932 | 0.246065 | 0.256445 | 0.267144 | 0.249378 | 0.277296 | 0.328773 | 0.337232 | 0.287307 | 0.324934 | 0.530608 | 1.067099 | 1.658238 | 1.707006 | 2.253041 | 2.127217 | 3.032039 | 3.027756 | 1.750360 | 0.858900 | 0.580569 | 0.427588 | 0.454472 | 0.348299 | 0.272608 | 0.221711 | 0.216091 | 0.223466 | 0.243503 | 0.277599 | 0.297899 | 0.313095 | 0.243070 | 0.214902 | 0.221377 | 0.266351 | 0.264076 | 0.324931 | 0.536577 | 0.687193 | 0.648961 | 0.724858 | 0.528800 | 0.596673 | 0.704301 | 0.762812 | 0.563197 | 0.564534 | 0.485253 | 0.550173 | 0.437649 | 0.316153 | 0.272427 | 0.301672 | 0.276305 | 0.252920 | 0.219655 | 0.214640 | 0.214863 | 0.219366 | 0.223733 | 0.218301 | 0.215269 | 0.228855 | 0.253626 | 0.286728 | 0.324770 | 0.327060 | 0.285100 | 0.236487 | 0.231979 | 0.235463 | 0.224459 | 0.217161 | 0.216404 | 0.216238 | 0.215100 | 0.223529 | 0.240660 | 0.246694 | 0.238644 | 0.231935 | 0.230543 | 0.242806 | 0.264015 | 0.282945 | 0.357329 | 0.539721 | 0.934501 | 1.926940 | 2.184619 | 2.897042 | 4.795604 | 2.571390 | 1.623023 | 0.644708 | 0.465973 | 0.398430 | 0.327971 | 0.293630 | 0.322683 | 0.256550 | 0.280670 | 0.309637 | 0.348548 | 0.374210 | 0.417999 | 0.491895 | 1.100258 | 2.267846 | 3.856730 | 6.395303 | 8.070793 | 10.488243 | 14.129637 | 11.812673 | 5.191395 |
| right central | 0.446607 | 0.348107 | 0.464256 | 0.455135 | 0.809434 | 0.747364 | 0.454525 | 0.414575 | 0.664222 | 0.445092 | 0.552359 | 0.654553 | 1.022547 | 2.060373 | 3.217492 | 3.747760 | 11.863619 | 5.467025 | 3.720207 | 1.471245 | 0.535074 | 0.358557 | 0.324808 | 0.214935 | 0.258402 | 0.318220 | 0.220170 | 0.272357 | 0.302369 | 0.310996 | 0.322769 | 0.386405 | 0.442753 | 0.383520 | 0.376543 | 0.250403 | 0.223083 | 0.240231 | 0.255902 | 0.250401 | 0.239238 | 0.217698 | 0.225088 | 0.305687 | 0.307534 | 0.335025 | 0.303492 | 0.283296 | 0.280033 | 0.245664 | 0.215323 | 0.226816 | 0.296034 | 0.907153 | 1.484453 | 3.100598 | 2.301901 | 1.444276 | 1.289228 | 0.803805 | 0.350591 | 0.291462 | 0.236585 | 0.284840 | 0.270732 | 0.253123 | 0.282957 | 0.298101 | 0.248493 | 0.236714 | 0.215471 | 0.215860 | 0.240918 | 0.234195 | 0.228339 | 0.258070 | 0.261223 | 0.282037 | 0.250839 | 0.217195 | 0.224070 | 0.231341 | 0.220171 | 0.214795 | 0.219683 | 0.231052 | 0.218577 | 0.214856 | 0.214626 | 0.242385 | 0.301097 | 0.326645 | 0.292133 | 0.305237 | 0.441512 | 0.533814 | 0.504423 | 0.358122 | 0.303940 | 0.255302 | 0.228270 | 0.214735 | 0.237986 | 0.300416 | 0.373471 | 0.393684 | 0.330379 | 0.338188 | 0.301612 | 0.335599 | 0.308525 | 0.285479 | 0.357320 | 0.490134 | 0.562735 | 0.644478 | 0.473652 | 0.407135 | 0.549458 | 0.738101 | 0.811789 | 0.884667 | 1.139641 | 1.893915 | 3.640406 | 3.196528 | 1.696786 | 2.279042 | 1.976675 | 1.835268 | 1.933919 | 1.851826 | 1.302565 | 1.392594 | 0.976703 | 1.134755 | 1.382014 | 1.565634 | 1.599655 | 1.598374 | 1.340426 | 1.579473 | 1.368273 | 1.163540 | 0.631097 | 0.375443 | 0.267038 | 0.254220 | 0.238738 | 0.230102 | 0.217438 | 0.230972 | 0.249017 | 0.272847 | 0.272854 | 0.351031 | 0.398563 | 0.431741 | 0.392559 | 0.380443 | 0.345759 | 0.344795 | 0.290164 | 0.259370 | 0.249696 | 0.271565 | 0.266649 | 0.262596 | 0.242293 | 0.228207 | 0.243328 | 0.286244 | 0.309794 | 0.340192 | 0.355813 | 0.355235 | 0.347145 | 0.326249 | 0.286508 | 0.280658 | 0.245745 | 0.267243 | 0.279890 | 0.282529 | 0.258555 | 0.252839 | 0.245733 | 0.303761 | 0.304866 | 0.323273 | 0.314267 | 0.425502 | 0.601838 | 0.656218 | 0.442168 | 0.290302 | 0.253528 | 0.257048 | 0.229755 | 0.222126 | 0.234200 | 0.256740 | 0.296470 | 0.327169 | 0.477140 | 0.731530 | 1.330248 | 1.160000 | 1.151429 | 0.987308 | 1.651328 | 1.333383 | 1.049410 | 1.234867 | 1.561610 | 2.821713 | 7.742817 | 9.117369 | 12.458638 | 8.300069 | 3.495549 | 1.954331 | 1.097230 | 0.786252 | 0.541328 | 0.407606 | 0.418187 | 0.603489 | 0.996068 | 1.390600 | 1.860268 | 1.667414 | 1.744806 | 1.425998 | 1.292250 | 1.438246 | 2.028249 | 2.097928 | 3.827049 | 5.213495 | 4.350555 | 2.743422 | 1.722282 | 1.057221 | 0.664953 | 0.632464 | 0.560400 | 0.713164 | 0.815682 | 0.708369 | 0.525729 | 0.574035 | 0.486751 | 0.539190 | 0.389427 | 0.416641 | 0.391025 | 0.367879 | 0.331941 | 0.370916 | 0.408408 | 0.417774 | 0.405486 | 0.425111 | 0.511013 | 0.626080 | 0.712608 | 0.593512 | 0.352915 | 0.266625 | 0.256558 | 0.268163 | 0.276858 | 0.249240 | 0.235492 | 0.256534 | 0.242114 | 0.237485 | 0.218652 | 0.220120 |
| left posterior | 0.863202 | 0.695246 | 0.423970 | 0.453968 | 0.384943 | 0.291702 | 0.214899 | 0.243717 | 0.291415 | 0.288387 | 0.455673 | 0.894567 | 1.220533 | 0.418503 | 0.264010 | 0.223560 | 0.232094 | 0.232770 | 0.231804 | 0.238860 | 0.215080 | 0.228380 | 0.267813 | 0.434732 | 0.707596 | 1.759385 | 2.799802 | 0.673155 | 0.318785 | 0.298733 | 0.233485 | 0.218019 | 0.243556 | 0.334260 | 0.385105 | 0.397827 | 0.641730 | 0.716903 | 0.386428 | 0.262767 | 0.245049 | 0.219017 | 0.253836 | 0.238082 | 0.216613 | 0.221642 | 0.242848 | 0.216796 | 0.223592 | 0.219355 | 0.214868 | 0.228527 | 0.262481 | 0.282832 | 0.261307 | 0.262789 | 0.235048 | 0.248340 | 0.297503 | 0.456869 | 1.422311 | 3.170900 | 2.996911 | 2.675867 | 1.545987 | 0.526564 | 0.302934 | 0.223166 | 0.223403 | 0.280152 | 0.604978 | 0.754344 | 1.047851 | 0.744740 | 0.545055 | 0.307190 | 0.262091 | 0.255547 | 0.390177 | 1.277851 | 4.065744 | 10.211423 | 39.377347 | 49.230475 | 22.615919 | 5.927812 | 1.716038 | 0.843866 | 0.750201 | 0.542426 | 0.347909 | 0.263041 | 0.217965 | 0.215184 | 0.215164 | 0.216425 | 0.214797 | 0.220787 | 0.243434 | 0.370530 | 0.540633 | 0.457931 | 0.563989 | 0.494207 | 0.380760 | 0.286981 | 0.246705 | 0.236386 | 0.262767 | 0.263083 | 0.307964 | 0.459133 | 0.831197 | 1.410962 | 1.448118 | 1.161936 | 0.839375 | 0.543770 | 0.396428 | 0.359468 | 0.317605 | 0.384350 | 0.454637 | 0.498535 | 0.734307 | 0.734264 | 0.713146 | 0.892981 | 1.233170 | 2.055109 | 2.943201 | 2.454766 | 2.963398 | 2.849798 | 2.237899 | 0.780521 | 0.386893 | 0.266676 | 0.222778 | 0.219893 | 0.241675 | 0.238057 | 0.216481 | 0.217165 | 0.219255 | 0.220409 | 0.219693 | 0.215029 | 0.215095 | 0.218047 | 0.233960 | 0.235065 | 0.245975 | 0.238129 | 0.230595 | 0.246883 | 0.258617 | 0.229489 | 0.221025 | 0.214762 | 0.218165 | 0.215594 | 0.214641 | 0.214735 | 0.226170 | 0.287859 | 0.378030 | 0.484121 | 0.443823 | 0.419581 | 0.437810 | 0.418231 | 0.319717 | 0.262024 | 0.217892 | 0.214991 | 0.226448 | 0.226929 | 0.224573 | 0.230562 | 0.237027 | 0.241662 | 0.267683 | 0.313963 | 0.449199 | 0.743060 | 0.999973 | 0.846297 | 0.546295 | 0.437711 | 0.398282 | 0.345975 | 0.310799 | 0.255990 | 0.251828 | 0.238419 | 0.222071 | 0.215969 | 0.234525 | 0.247948 | 0.257830 | 0.292159 | 0.344464 | 0.373153 | 0.408693 | 0.428131 | 0.511578 | 0.557472 | 0.631563 | 0.559096 | 0.359382 | 0.273495 | 0.244110 | 0.251330 | 0.257623 | 0.249282 | 0.300913 | 0.419996 | 0.519078 | 0.807171 | 1.080480 | 1.309739 | 1.269826 | 0.887799 | 0.716263 | 0.784338 | 0.719211 | 0.541291 | 0.617792 | 0.642322 | 0.573704 | 0.409895 | 0.339050 | 0.275358 | 0.240692 | 0.214736 | 0.217665 | 0.226394 | 0.226226 | 0.236450 | 0.234835 | 0.245289 | 0.279584 | 0.363682 | 0.416804 | 0.421916 | 0.305152 | 0.312711 | 0.310216 | 0.263713 | 0.217283 | 0.214772 | 0.214769 | 0.214858 | 0.215185 | 0.215446 | 0.216528 | 0.225924 | 0.232161 | 0.229247 | 0.249078 | 0.251512 | 0.245047 | 0.234324 | 0.240610 | 0.269306 | 0.282095 | 0.279048 | 0.278227 | 0.270760 | 0.315221 | 0.314833 | 0.315079 | 0.351235 | 0.344158 | 0.363557 | 0.377216 | 0.352662 | 0.329788 | 0.284451 |
| right posterior | 0.218218 | 0.219219 | 0.214703 | 0.218332 | 0.215809 | 0.214626 | 0.215101 | 0.218164 | 0.229375 | 0.272578 | 0.288124 | 0.280380 | 0.265864 | 0.222863 | 0.217896 | 0.215080 | 0.215808 | 0.255372 | 0.324096 | 0.477499 | 0.441001 | 0.846022 | 0.613147 | 0.403876 | 0.261283 | 0.241908 | 0.214642 | 0.214626 | 0.216493 | 0.222831 | 0.222855 | 0.241415 | 0.269643 | 0.453224 | 0.541417 | 0.422832 | 0.304690 | 0.308949 | 0.292662 | 0.297638 | 0.215439 | 0.219391 | 0.256377 | 0.256875 | 0.237965 | 0.217637 | 0.224466 | 0.222119 | 0.214626 | 0.244138 | 0.237145 | 0.237970 | 0.218809 | 0.216482 | 0.214640 | 0.216767 | 0.237427 | 0.304038 | 0.536286 | 1.323365 | 3.202780 | 5.983252 | 5.319874 | 5.411195 | 2.466681 | 1.276836 | 0.486731 | 0.276336 | 0.229128 | 0.214668 | 0.252657 | 0.372466 | 0.661488 | 1.135490 | 2.346014 | 7.840731 | 37.401201 | 8.480293 | 1.919452 | 0.915912 | 0.530688 | 0.317160 | 0.225658 | 0.228652 | 0.248896 | 0.254203 | 0.221135 | 0.216024 | 0.218523 | 0.293203 | 0.332562 | 0.289941 | 0.294242 | 0.310431 | 0.305473 | 0.320875 | 0.263019 | 0.248533 | 0.292784 | 0.286479 | 0.316073 | 0.353585 | 0.293360 | 0.272369 | 0.235758 | 0.214793 | 0.217280 | 0.222845 | 0.229816 | 0.231203 | 0.237402 | 0.259438 | 0.271369 | 0.313127 | 0.430863 | 0.703426 | 1.002157 | 0.541878 | 0.361438 | 0.288050 | 0.223329 | 0.216437 | 0.239156 | 0.265854 | 0.227004 | 0.219607 | 0.216010 | 0.231678 | 0.263584 | 0.276910 | 0.255828 | 0.228126 | 0.214822 | 0.230047 | 0.304524 | 0.427446 | 0.391955 | 0.259770 | 0.228406 | 0.214626 | 0.231489 | 0.286723 | 0.417967 | 0.392250 | 0.290665 | 0.306630 | 0.327461 | 0.330961 | 0.290046 | 0.290251 | 0.280997 | 0.251495 | 0.233278 | 0.228877 | 0.231290 | 0.224241 | 0.215741 | 0.214702 | 0.215388 | 0.215086 | 0.215155 | 0.214658 | 0.223171 | 0.225761 | 0.220411 | 0.219367 | 0.214664 | 0.215883 | 0.215025 | 0.214626 | 0.214708 | 0.215941 | 0.215590 | 0.218432 | 0.226677 | 0.216021 | 0.214671 | 0.224222 | 0.242118 | 0.273566 | 0.282815 | 0.293280 | 0.267281 | 0.280901 | 0.274146 | 0.272125 | 0.263095 | 0.270336 | 0.290064 | 0.342633 | 0.357303 | 0.332296 | 0.346538 | 0.365638 | 0.362219 | 0.303663 | 0.319661 | 0.273069 | 0.228260 | 0.217272 | 0.215892 | 0.217742 | 0.215711 | 0.228732 | 0.225069 | 0.217591 | 0.216656 | 0.215257 | 0.215116 | 0.215276 | 0.227186 | 0.242628 | 0.276754 | 0.309048 | 0.362557 | 0.430701 | 0.355589 | 0.320939 | 0.341947 | 0.398983 | 0.379476 | 0.316274 | 0.266793 | 0.253497 | 0.243157 | 0.226135 | 0.225215 | 0.250498 | 0.272873 | 0.277469 | 0.305205 | 0.354453 | 0.449931 | 0.417848 | 0.314550 | 0.282511 | 0.280060 | 0.251321 | 0.264973 | 0.262282 | 0.279811 | 0.276009 | 0.277812 | 0.276042 | 0.311803 | 0.294585 | 0.283344 | 0.298466 | 0.273711 | 0.275321 | 0.314268 | 0.388831 | 0.489326 | 0.561444 | 0.460463 | 0.660837 | 0.721446 | 0.619790 | 0.435341 | 0.355475 | 0.405278 | 0.461706 | 0.434506 | 0.414987 | 0.481040 | 0.688457 | 1.040762 | 1.157200 | 1.480167 | 1.854569 | 2.718991 | 3.216485 | 2.417953 | 2.625114 | 2.602368 | 3.424997 | 3.482870 | 3.916148 | 4.049219 | 2.945572 |
| all electrodes | 0.722585 | 0.418814 | 0.218979 | 0.217896 | 0.238559 | 0.539351 | 3.252801 | 4.413438 | 0.902707 | 0.292395 | 0.232968 | 0.216051 | 0.230498 | 0.253140 | 0.248770 | 0.226241 | 0.214626 | 0.218930 | 0.270433 | 0.277747 | 0.310715 | 0.248277 | 0.293124 | 0.285271 | 0.239712 | 0.234077 | 0.314710 | 0.399759 | 0.281610 | 0.304630 | 0.248836 | 0.215892 | 0.229552 | 0.261482 | 0.347102 | 0.289175 | 0.340225 | 0.323337 | 0.331288 | 0.279704 | 0.232132 | 0.220839 | 0.227410 | 0.258043 | 0.231435 | 0.249817 | 0.238728 | 0.224662 | 0.248675 | 0.267180 | 0.337799 | 0.375841 | 0.504139 | 0.924732 | 1.631151 | 1.088905 | 1.229380 | 0.752255 | 0.300740 | 0.218151 | 0.229637 | 0.317919 | 0.412500 | 0.423351 | 0.333882 | 0.239715 | 0.225848 | 0.243455 | 0.378615 | 0.391270 | 0.366632 | 0.363468 | 0.345507 | 0.350309 | 0.393499 | 0.405025 | 0.291609 | 0.341548 | 0.327635 | 0.394909 | 0.816117 | 1.055862 | 0.863900 | 1.605410 | 1.426617 | 1.309802 | 0.850171 | 0.465057 | 0.592275 | 1.052370 | 1.327460 | 1.232029 | 2.216348 | 4.989949 | 4.097406 | 3.959275 | 2.123014 | 2.904750 | 8.047604 | 6.628130 | 5.575551 | 4.604437 | 3.030156 | 2.669674 | 2.629374 | 2.308607 | 5.177499 | 3.582918 | 3.028809 | 1.420149 | 1.054094 | 1.339417 | 1.042699 | 0.498255 | 0.309891 | 0.307775 | 0.377417 | 0.512746 | 0.377163 | 0.484338 | 0.625845 | 1.550403 | 1.974128 | 2.784539 | 1.476640 | 0.966136 | 0.454641 | 0.326491 | 0.242269 | 0.221205 | 0.229599 | 0.273940 | 0.307338 | 0.256294 | 0.251133 | 0.224904 | 0.216110 | 0.215452 | 0.222241 | 0.238542 | 0.234244 | 0.218834 | 0.215346 | 0.220649 | 0.229847 | 0.243824 | 0.274341 | 0.274095 | 0.255823 | 0.269719 | 0.244966 | 0.240162 | 0.217764 | 0.219975 | 0.233493 | 0.377178 | 1.284216 | 3.829904 | 5.100652 | 3.182838 | 1.656080 | 0.902372 | 0.477216 | 0.480744 | 1.000365 | 2.742281 | 8.159411 | 13.318011 | 15.633469 | 29.204682 | 33.196988 | 20.431026 | 12.564562 | 15.502560 | 16.652033 | 10.969885 | 3.865729 | 3.023569 | 2.981746 | 3.679683 | 17.332137 | 13.961343 | 2.206256 | 0.772159 | 0.416279 | 0.364885 | 0.276940 | 0.225507 | 0.217126 | 0.250871 | 0.314941 | 0.345133 | 0.379622 | 0.599143 | 1.115530 | 1.596549 | 1.844418 | 2.703989 | 3.254343 | 2.377402 | 2.129198 | 1.612140 | 4.752102 | 5.315683 | 7.118625 | 13.222690 | 18.281939 | 11.295683 | 13.643540 | 3.050842 | 3.667658 | 2.729322 | 2.890010 | 1.761045 | 1.250715 | 1.276471 | 2.953710 | 5.017130 | 4.589447 | 2.456058 | 4.779306 | 17.479792 | 96.239968 | 207.005022 | 257.183292 | 1237.282925 | 1922.544504 | 1596.801556 | 562.426839 | 81.780570 | 10.609761 | 2.844332 | 1.191532 | 0.926293 | 0.607322 | 0.576060 | 0.746586 | 1.101396 | 1.997804 | 2.385379 | 2.974408 | 2.672274 | 2.355649 | 2.570783 | 2.374236 | 2.438435 | 2.358683 | 2.258624 | 1.484918 | 0.606716 | 0.286961 | 0.254931 | 0.234970 | 0.218634 | 0.217188 | 0.222604 | 0.218245 | 0.225287 | 0.264486 | 0.301909 | 0.365235 | 0.396662 | 0.364071 | 0.316480 | 0.329021 | 0.313892 | 0.243687 | 0.216646 | 0.216752 | 0.216057 | 0.215528 | 0.217885 | 0.241365 | 0.238517 | 0.223502 | 0.217321 | 0.216057 | 0.215542 | 0.216324 | 0.217205 |

Searchlight, spatiotemporal cluster permutation test

No significant clusters observed.

G) sex, happy

  
|  | time window | peak latency | cluster *p* | peak Cohen's *d* |  | | | |
| **all electrodes** | 235 - 330 ms | 305 ms | 0.0229 | 0.796 |  | | | |
|  | | | | | | | | |

Time-resolved classification, cluster permutation tests

|  | **left hemisphere** | | | | **right hemisphere** | | | |
|  | time window | peak latency | cluster *p* | peak Cohen's *d* | time window | peak latency | cluster *p* | peak Cohen's *d* |
| **anterior** |  | | | | 290 - 445 ms | 420 ms | 0.0172 | 0.823 |
| **central** | 180 - 330 ms | 250 ms | 0.0147 | 0.8393 | 195 - 280 ms | 260 ms | 0.0376 | 1.1078 |
| **posterior** |  | | | | 160 - 235 ms | 190 ms | 0.0413 | 0.9122 |

  

Time-resolved classification, Bayesian statistics

|  | -200 | -195 | -190 | -185 | -180 | -175 | -170 | -165 | -160 | -155 | -150 | -145 | -140 | -135 | -130 | -125 | -120 | -115 | -110 | -105 | -100 | -95 | -90 | -85 | -80 | -75 | -70 | -65 | -60 | -55 | -50 | -45 | -40 | -35 | -30 | -25 | -20 | -15 | -10 | -5 | 0 | 5 | 10 | 15 | 20 | 25 | 30 | 35 | 40 | 45 | 50 | 55 | 60 | 65 | 70 | 75 | 80 | 85 | 90 | 95 | 100 | 105 | 110 | 115 | 120 | 125 | 130 | 135 | 140 | 145 | 150 | 155 | 160 | 165 | 170 | 175 | 180 | 185 | 190 | 195 | 200 | 205 | 210 | 215 | 220 | 225 | 230 | 235 | 240 | 245 | 250 | 255 | 260 | 265 | 270 | 275 | 280 | 285 | 290 | 295 | 300 | 305 | 310 | 315 | 320 | 325 | 330 | 335 | 340 | 345 | 350 | 355 | 360 | 365 | 370 | 375 | 380 | 385 | 390 | 395 | 400 | 405 | 410 | 415 | 420 | 425 | 430 | 435 | 440 | 445 | 450 | 455 | 460 | 465 | 470 | 475 | 480 | 485 | 490 | 495 | 500 | 505 | 510 | 515 | 520 | 525 | 530 | 535 | 540 | 545 | 550 | 555 | 560 | 565 | 570 | 575 | 580 | 585 | 590 | 595 | 600 | 605 | 610 | 615 | 620 | 625 | 630 | 635 | 640 | 645 | 650 | 655 | 660 | 665 | 670 | 675 | 680 | 685 | 690 | 695 | 700 | 705 | 710 | 715 | 720 | 725 | 730 | 735 | 740 | 745 | 750 | 755 | 760 | 765 | 770 | 775 | 780 | 785 | 790 | 795 | 800 | 805 | 810 | 815 | 820 | 825 | 830 | 835 | 840 | 845 | 850 | 855 | 860 | 865 | 870 | 875 | 880 | 885 | 890 | 895 | 900 | 905 | 910 | 915 | 920 | 925 | 930 | 935 | 940 | 945 | 950 | 955 | 960 | 965 | 970 | 975 | 980 | 985 | 990 | 995 | 1000 | 1005 | 1010 | 1015 | 1020 | 1025 | 1030 | 1035 | 1040 | 1045 | 1050 | 1055 | 1060 | 1065 | 1070 | 1075 | 1080 | 1085 | 1090 | 1095 | 1100 | 1105 | 1110 | 1115 | 1120 | 1125 | 1130 | 1135 | 1140 | 1145 | 1150 | 1155 | 1160 | 1165 | 1170 | 1175 | 1180 | 1185 | 1190 | 1195 |
| --- | --- | --- | --- | --- | --- | --- | --- | --- | --- | --- | --- | --- | --- | --- | --- | --- | --- | --- | --- | --- | --- | --- | --- | --- | --- | --- | --- | --- | --- | --- | --- | --- | --- | --- | --- | --- | --- | --- | --- | --- | --- | --- | --- | --- | --- | --- | --- | --- | --- | --- | --- | --- | --- | --- | --- | --- | --- | --- | --- | --- | --- | --- | --- | --- | --- | --- | --- | --- | --- | --- | --- | --- | --- | --- | --- | --- | --- | --- | --- | --- | --- | --- | --- | --- | --- | --- | --- | --- | --- | --- | --- | --- | --- | --- | --- | --- | --- | --- | --- | --- | --- | --- | --- | --- | --- | --- | --- | --- | --- | --- | --- | --- | --- | --- | --- | --- | --- | --- | --- | --- | --- | --- | --- | --- | --- | --- | --- | --- | --- | --- | --- | --- | --- | --- | --- | --- | --- | --- | --- | --- | --- | --- | --- | --- | --- | --- | --- | --- | --- | --- | --- | --- | --- | --- | --- | --- | --- | --- | --- | --- | --- | --- | --- | --- | --- | --- | --- | --- | --- | --- | --- | --- | --- | --- | --- | --- | --- | --- | --- | --- | --- | --- | --- | --- | --- | --- | --- | --- | --- | --- | --- | --- | --- | --- | --- | --- | --- | --- | --- | --- | --- | --- | --- | --- | --- | --- | --- | --- | --- | --- | --- | --- | --- | --- | --- | --- | --- | --- | --- | --- | --- | --- | --- | --- | --- | --- | --- | --- | --- | --- | --- | --- | --- | --- | --- | --- | --- | --- | --- | --- | --- | --- | --- | --- | --- | --- | --- | --- | --- | --- | --- | --- | --- | --- | --- | --- | --- | --- | --- | --- | --- | --- | --- | --- | --- | --- | --- | --- | --- | --- | --- | --- | --- | --- | --- | --- | --- | --- | --- | --- |
| left anterior | 0.970229 | 1.585176 | 1.883659 | 2.166823 | 2.135855 | 0.530089 | 0.272155 | 0.215197 | 0.253997 | 0.463246 | 1.000992 | 0.789286 | 0.579632 | 0.613641 | 0.351238 | 0.215896 | 0.345851 | 1.786332 | 1.462338 | 0.754537 | 0.734394 | 0.288175 | 0.222530 | 0.438920 | 1.724224 | 3.382361 | 4.508390 | 13.854422 | 31.659513 | 51.728513 | 24.392981 | 8.232860 | 8.272815 | 3.200317 | 0.974631 | 0.289398 | 0.220726 | 0.216420 | 0.232542 | 0.353865 | 0.567034 | 0.749351 | 0.613013 | 0.299782 | 0.233452 | 0.227473 | 0.245698 | 0.214672 | 0.272685 | 0.569065 | 0.456214 | 0.433465 | 0.750911 | 1.009538 | 0.498818 | 0.284490 | 0.216849 | 0.226509 | 0.268353 | 0.308095 | 0.304758 | 0.260812 | 0.227408 | 0.217606 | 0.218612 | 0.229022 | 0.217266 | 0.227135 | 0.271765 | 0.311556 | 0.309442 | 0.304313 | 0.362129 | 0.294986 | 0.219624 | 0.224223 | 0.229817 | 0.218831 | 0.226585 | 0.247677 | 0.314195 | 0.350819 | 0.334773 | 0.274177 | 0.239133 | 0.233123 | 0.258294 | 0.344539 | 0.536689 | 1.153860 | 2.806550 | 3.833661 | 4.435616 | 1.873479 | 1.061462 | 1.004873 | 1.127130 | 1.193550 | 1.570288 | 1.393186 | 2.168063 | 2.637395 | 1.275230 | 1.080962 | 0.531102 | 0.336760 | 0.382041 | 0.325801 | 0.284756 | 0.305873 | 0.260791 | 0.277944 | 0.279871 | 0.237444 | 0.249256 | 0.269633 | 0.278043 | 0.268407 | 0.272719 | 0.277222 | 0.275085 | 0.245540 | 0.224642 | 0.225241 | 0.238380 | 0.239839 | 0.235371 | 0.236030 | 0.233369 | 0.257861 | 0.274940 | 0.290910 | 0.364589 | 0.511820 | 0.741796 | 0.987710 | 0.625282 | 0.329531 | 0.235672 | 0.217821 | 0.218009 | 0.214732 | 0.215724 | 0.222166 | 0.241250 | 0.268875 | 0.343720 | 0.377439 | 0.325871 | 0.336887 | 0.359506 | 0.468203 | 0.673627 | 0.669018 | 0.477255 | 0.353478 | 0.264700 | 0.240765 | 0.219832 | 0.214818 | 0.214659 | 0.218895 | 0.244075 | 0.291533 | 0.307534 | 0.310294 | 0.305404 | 0.270129 | 0.242328 | 0.219291 | 0.215707 | 0.220223 | 0.222993 | 0.241434 | 0.246140 | 0.241549 | 0.216271 | 0.214726 | 0.215241 | 0.219631 | 0.214682 | 0.218111 | 0.223522 | 0.232295 | 0.261321 | 0.260303 | 0.307551 | 0.320909 | 0.337541 | 0.361059 | 0.341784 | 0.417611 | 0.791049 | 0.804428 | 0.776300 | 0.687566 | 0.533129 | 0.562154 | 0.447352 | 0.356464 | 0.336505 | 0.311674 | 0.321599 | 0.307553 | 0.244988 | 0.245764 | 0.247791 | 0.244763 | 0.260061 | 0.237925 | 0.229461 | 0.240879 | 0.244530 | 0.268958 | 0.303464 | 0.363763 | 0.608365 | 1.260935 | 1.757835 | 1.773015 | 1.937328 | 2.777350 | 2.749617 | 1.675173 | 0.650086 | 0.494871 | 0.499055 | 0.526081 | 0.413657 | 0.355462 | 0.322871 | 0.369203 | 0.336614 | 0.331734 | 0.311127 | 0.316403 | 0.307441 | 0.286039 | 0.298829 | 0.423708 | 0.643564 | 0.741603 | 0.903758 | 0.868563 | 1.080367 | 0.942552 | 0.725003 | 0.578092 | 0.470376 | 0.336994 | 0.331545 | 0.309007 | 0.320100 | 0.308816 | 0.289555 | 0.296083 | 0.296703 | 0.305589 | 0.332955 | 0.350103 | 0.308709 | 0.261126 | 0.226557 | 0.230141 | 0.222722 | 0.243140 | 0.270064 | 0.295554 | 0.290294 | 0.279525 | 0.241483 | 0.241634 | 0.219884 | 0.214626 | 0.215847 | 0.216216 | 0.215482 | 0.215241 | 0.216005 | 0.215602 |
| right anterior | 1.354672 | 1.803765 | 1.550442 | 1.213893 | 1.129407 | 0.257707 | 0.234970 | 0.649431 | 4.343812 | 14.315626 | 5.891193 | 1.138584 | 0.297646 | 0.287039 | 0.223352 | 1.390904 | 7.656672 | 16.285425 | 11.098916 | 1.528024 | 1.393408 | 1.326527 | 0.343718 | 0.242369 | 0.359068 | 0.403875 | 0.336350 | 0.526886 | 0.461017 | 0.558050 | 0.333453 | 0.260973 | 0.235903 | 0.215169 | 0.215264 | 0.218179 | 0.236430 | 0.222317 | 0.269105 | 0.473930 | 0.861527 | 2.998894 | 4.012514 | 2.376386 | 1.131656 | 0.479443 | 0.279122 | 0.265581 | 0.331879 | 0.398865 | 0.369413 | 0.351551 | 0.341586 | 0.242882 | 0.220334 | 0.285616 | 0.320506 | 0.246087 | 0.216682 | 0.234339 | 0.448459 | 0.713386 | 0.919344 | 1.243813 | 1.263295 | 1.342219 | 1.057418 | 1.019329 | 2.510557 | 4.614797 | 4.823073 | 4.327298 | 1.839645 | 1.750418 | 1.794018 | 1.990294 | 2.059496 | 2.128852 | 1.794245 | 2.421510 | 1.933709 | 1.885598 | 1.371632 | 1.398514 | 1.338984 | 1.199833 | 1.569358 | 3.388112 | 5.167759 | 8.936642 | 4.318402 | 2.338570 | 1.416124 | 0.913305 | 0.690764 | 0.724449 | 0.672630 | 0.885625 | 1.352915 | 1.723203 | 3.145574 | 3.830369 | 3.904868 | 4.583982 | 5.687758 | 3.008901 | 3.194286 | 3.181211 | 4.305545 | 3.932394 | 3.426426 | 3.863524 | 4.186421 | 3.108489 | 1.911785 | 1.589483 | 1.592616 | 1.999240 | 2.964751 | 6.100537 | 17.644218 | 32.135910 | 29.337223 | 47.305392 | 62.337682 | 46.479140 | 22.906817 | 8.093189 | 4.714727 | 2.603348 | 0.876489 | 0.491586 | 0.390201 | 0.506609 | 0.566673 | 0.456148 | 0.501538 | 0.667510 | 0.554545 | 0.423559 | 0.245657 | 0.214679 | 0.216295 | 0.225630 | 0.245860 | 0.305309 | 0.292724 | 0.300960 | 0.373441 | 0.474602 | 0.381184 | 0.342603 | 0.406605 | 0.640455 | 1.125242 | 1.662610 | 1.875107 | 2.144687 | 1.628690 | 1.263054 | 0.984802 | 1.844045 | 2.769183 | 2.092049 | 2.073055 | 2.229703 | 1.923803 | 1.168229 | 0.534331 | 0.488668 | 0.437776 | 0.390941 | 0.454135 | 0.466133 | 0.616283 | 0.530409 | 0.333385 | 0.329669 | 0.372375 | 0.354779 | 0.365461 | 0.325953 | 0.333991 | 0.345494 | 0.322331 | 0.291204 | 0.255493 | 0.288358 | 0.442545 | 0.816959 | 1.359401 | 1.274415 | 1.112344 | 1.349048 | 1.156838 | 0.855970 | 0.624702 | 0.589740 | 0.822589 | 1.242757 | 1.369336 | 1.289065 | 0.959471 | 0.785066 | 0.660155 | 0.558593 | 0.413962 | 0.388621 | 0.475301 | 0.616883 | 0.765247 | 0.696441 | 0.646314 | 0.619709 | 0.607957 | 0.549032 | 0.544139 | 0.613547 | 0.690505 | 0.806455 | 1.021073 | 1.008584 | 0.830093 | 0.681168 | 0.423498 | 0.324960 | 0.293186 | 0.277403 | 0.322744 | 0.366137 | 0.344798 | 0.429948 | 0.590245 | 0.802394 | 1.028861 | 0.863662 | 0.784929 | 1.014795 | 0.936943 | 1.371632 | 2.021439 | 2.387291 | 2.947546 | 3.037887 | 3.089856 | 2.630844 | 1.877481 | 0.958560 | 0.530717 | 0.446388 | 0.379323 | 0.327960 | 0.329567 | 0.318143 | 0.333049 | 0.365308 | 0.331738 | 0.274886 | 0.259153 | 0.241765 | 0.229072 | 0.238314 | 0.251683 | 0.272481 | 0.326509 | 0.298529 | 0.343419 | 0.351358 | 0.318888 | 0.311333 | 0.292137 | 0.272264 | 0.249516 | 0.231202 | 0.240334 | 0.283519 | 0.292139 | 0.294151 | 0.316828 | 0.519045 |
| left central | 0.525443 | 0.361681 | 0.318425 | 0.287723 | 0.244714 | 0.215735 | 0.219851 | 0.227413 | 0.217838 | 0.239028 | 0.309692 | 0.360848 | 0.291148 | 0.355179 | 0.344661 | 0.238783 | 0.260023 | 0.375571 | 0.539685 | 0.906316 | 1.350153 | 2.101979 | 1.284497 | 0.498582 | 0.348586 | 0.244979 | 0.270365 | 0.227382 | 0.228607 | 0.353365 | 0.624475 | 0.773782 | 0.643254 | 1.609621 | 1.052608 | 0.425538 | 0.282248 | 0.251763 | 0.315819 | 0.378260 | 0.550192 | 0.920782 | 1.478026 | 0.842667 | 0.757056 | 0.641512 | 0.602852 | 0.425803 | 0.389032 | 0.465343 | 0.778224 | 0.831661 | 0.703483 | 0.562521 | 0.416076 | 0.234427 | 0.218385 | 0.251932 | 0.241261 | 0.241091 | 0.222842 | 0.220384 | 0.216777 | 0.218481 | 0.214635 | 0.219553 | 0.248303 | 0.295722 | 0.385100 | 0.519101 | 0.577789 | 0.393532 | 0.242649 | 0.216846 | 0.301910 | 0.713703 | 1.633509 | 6.560633 | 28.191183 | 77.598977 | 92.688121 | 127.459001 | 64.762310 | 29.027859 | 15.129121 | 5.651473 | 4.658675 | 4.872870 | 9.884875 | 19.469582 | 74.353770 | 42.392070 | 49.878845 | 37.405966 | 43.380358 | 42.843385 | 18.229668 | 6.178383 | 8.883408 | 8.975517 | 8.390470 | 7.336295 | 4.245534 | 3.354870 | 3.500968 | 3.296671 | 1.752971 | 1.294819 | 1.717715 | 1.597763 | 1.128517 | 0.698555 | 0.385478 | 0.495433 | 0.692784 | 0.676931 | 0.740456 | 0.849492 | 0.773502 | 1.020362 | 2.026034 | 2.659795 | 3.192045 | 3.219851 | 6.728327 | 4.519626 | 1.708973 | 0.591032 | 0.410256 | 0.358667 | 0.321767 | 0.270043 | 0.273203 | 0.278833 | 0.288208 | 0.297461 | 0.233424 | 0.215347 | 0.259249 | 0.335923 | 0.398507 | 0.374704 | 0.489843 | 0.354469 | 0.267807 | 0.226399 | 0.217995 | 0.216972 | 0.230648 | 0.259718 | 0.271932 | 0.257514 | 0.244118 | 0.260429 | 0.221619 | 0.216171 | 0.215695 | 0.214637 | 0.216392 | 0.229369 | 0.227716 | 0.261027 | 0.265622 | 0.280457 | 0.376109 | 0.522794 | 0.822811 | 1.275158 | 1.341030 | 1.063873 | 0.636529 | 0.472770 | 0.392222 | 0.301420 | 0.241156 | 0.217700 | 0.215162 | 0.218075 | 0.226099 | 0.226706 | 0.223023 | 0.238658 | 0.282128 | 0.370563 | 0.442471 | 0.482911 | 0.420498 | 0.318127 | 0.245931 | 0.223687 | 0.217899 | 0.218381 | 0.214835 | 0.215056 | 0.217713 | 0.220701 | 0.221276 | 0.216543 | 0.214626 | 0.217198 | 0.216309 | 0.216171 | 0.236251 | 0.253171 | 0.306671 | 0.325524 | 0.397081 | 0.403383 | 0.361839 | 0.317458 | 0.315146 | 0.308762 | 0.327341 | 0.342614 | 0.344740 | 0.380146 | 0.504518 | 0.748136 | 0.928378 | 0.938281 | 0.846201 | 0.589467 | 0.495562 | 0.362270 | 0.310319 | 0.293286 | 0.297949 | 0.325801 | 0.386056 | 0.376583 | 0.436404 | 0.465926 | 0.444013 | 0.470485 | 0.422621 | 0.372812 | 0.384894 | 0.402996 | 0.445963 | 0.487338 | 0.535871 | 0.672551 | 0.856927 | 0.756962 | 0.589997 | 0.382077 | 0.334181 | 0.297521 | 0.269107 | 0.256126 | 0.263410 | 0.276555 | 0.369617 | 0.457414 | 0.524656 | 0.629222 | 0.705289 | 0.578257 | 0.445398 | 0.334112 | 0.267544 | 0.240122 | 0.233257 | 0.220638 | 0.228404 | 0.227636 | 0.230010 | 0.234048 | 0.245961 | 0.266062 | 0.333994 | 0.437517 | 0.650357 | 0.750441 | 0.708367 | 0.593531 | 0.450397 | 0.347523 | 0.263174 | 0.217812 |
| right central | 0.249829 | 0.459725 | 0.681289 | 0.666700 | 0.763911 | 0.854868 | 1.193740 | 0.993552 | 0.276046 | 0.223169 | 0.217479 | 0.247371 | 0.295825 | 0.415938 | 0.551469 | 0.268980 | 0.363931 | 0.537707 | 0.315837 | 0.376475 | 0.621729 | 1.539502 | 4.680153 | 1.344925 | 0.506842 | 0.656069 | 1.096865 | 1.870619 | 1.048394 | 0.591929 | 0.455872 | 0.511021 | 0.424219 | 0.312335 | 0.219524 | 0.214636 | 0.226577 | 0.217145 | 0.217487 | 0.228797 | 0.266599 | 0.299033 | 0.415616 | 0.434551 | 0.505601 | 0.632434 | 0.964993 | 0.973442 | 1.308465 | 0.917237 | 0.511517 | 0.442340 | 0.268824 | 0.228955 | 0.221864 | 0.216523 | 0.227320 | 0.310224 | 0.308746 | 0.338957 | 0.272413 | 0.247855 | 0.268917 | 0.250521 | 0.230702 | 0.217337 | 0.217390 | 0.226861 | 0.227782 | 0.232745 | 0.231621 | 0.221686 | 0.216249 | 0.256025 | 0.399276 | 0.698147 | 1.099837 | 1.111998 | 1.155324 | 1.412355 | 1.757532 | 3.668745 | 4.467254 | 4.158827 | 5.270836 | 4.277561 | 4.801691 | 7.271895 | 10.083588 | 48.917316 | 604.307886 | 2102.198944 | 1395.105399 | 139.360637 | 23.765559 | 6.306948 | 1.643952 | 0.901081 | 0.662553 | 0.604429 | 0.970549 | 1.230679 | 1.049528 | 0.887797 | 0.517921 | 0.363810 | 0.316984 | 0.272423 | 0.274629 | 0.312590 | 0.398659 | 0.439352 | 0.379176 | 0.313947 | 0.255918 | 0.217644 | 0.216138 | 0.227250 | 0.216476 | 0.217924 | 0.236405 | 0.250901 | 0.256657 | 0.246176 | 0.253409 | 0.260401 | 0.270198 | 0.249016 | 0.258005 | 0.315939 | 0.364258 | 0.298683 | 0.231514 | 0.227248 | 0.256304 | 0.298728 | 0.286142 | 0.311488 | 0.333192 | 0.407983 | 0.440246 | 0.353458 | 0.274740 | 0.215726 | 0.257440 | 0.265509 | 0.271995 | 0.226089 | 0.226408 | 0.312099 | 0.755713 | 1.715259 | 1.768281 | 1.936937 | 1.224605 | 0.743315 | 0.537704 | 0.407667 | 0.334623 | 0.332023 | 0.392137 | 0.426235 | 0.380455 | 0.334069 | 0.290693 | 0.340937 | 0.337063 | 0.272791 | 0.222570 | 0.215301 | 0.228197 | 0.245524 | 0.280767 | 0.287258 | 0.253218 | 0.223347 | 0.220672 | 0.222505 | 0.228239 | 0.241830 | 0.262485 | 0.273456 | 0.309680 | 0.314074 | 0.319943 | 0.319877 | 0.306421 | 0.278412 | 0.263283 | 0.240797 | 0.230646 | 0.220438 | 0.223001 | 0.232372 | 0.254946 | 0.277039 | 0.266573 | 0.270080 | 0.299404 | 0.343808 | 0.448825 | 0.592723 | 0.618780 | 0.801129 | 0.811425 | 0.706283 | 0.572923 | 0.410850 | 0.333326 | 0.368443 | 0.415232 | 0.560724 | 0.643440 | 0.782790 | 1.026283 | 1.404480 | 2.595544 | 5.779207 | 6.549017 | 4.815690 | 3.543309 | 2.233539 | 1.928227 | 1.299259 | 0.741595 | 0.617532 | 0.763271 | 0.987169 | 1.266965 | 0.849868 | 0.576484 | 0.558948 | 0.604667 | 0.566043 | 0.419672 | 0.265328 | 0.231865 | 0.223140 | 0.217968 | 0.214675 | 0.215396 | 0.220387 | 0.232883 | 0.248267 | 0.246287 | 0.231718 | 0.219231 | 0.215133 | 0.215364 | 0.225834 | 0.261379 | 0.274103 | 0.329087 | 0.321960 | 0.327767 | 0.360774 | 0.307774 | 0.245981 | 0.225370 | 0.214737 | 0.217151 | 0.227626 | 0.256150 | 0.295415 | 0.304847 | 0.286550 | 0.254571 | 0.239908 | 0.227491 | 0.223641 | 0.215306 | 0.214697 | 0.217008 | 0.229089 | 0.247359 | 0.252742 | 0.269876 | 0.294999 | 0.303242 | 0.283716 |
| left posterior | 0.330874 | 0.418771 | 0.415480 | 0.369964 | 0.422676 | 0.346316 | 0.221492 | 0.219897 | 0.247581 | 0.281713 | 0.265004 | 0.226865 | 0.214756 | 0.222156 | 0.215891 | 0.214823 | 0.215876 | 0.215096 | 0.219653 | 0.214673 | 0.226225 | 0.293222 | 0.520299 | 0.658452 | 0.980782 | 1.705353 | 1.099966 | 0.613644 | 0.497742 | 0.404423 | 0.385551 | 0.295753 | 0.219056 | 0.328268 | 0.511661 | 0.686240 | 0.656165 | 0.727118 | 0.642945 | 0.344381 | 0.230133 | 0.222410 | 0.214669 | 0.225444 | 0.240461 | 0.234422 | 0.231070 | 0.242375 | 0.271019 | 0.293943 | 0.296148 | 0.334829 | 0.381628 | 0.455709 | 0.415085 | 0.354964 | 0.330811 | 0.277359 | 0.228927 | 0.214789 | 0.224151 | 0.276722 | 0.389504 | 0.770956 | 1.420591 | 2.618834 | 2.957695 | 3.882244 | 1.829682 | 0.815256 | 0.419548 | 0.323398 | 0.325447 | 0.378743 | 0.323495 | 0.438623 | 0.672523 | 1.182575 | 1.500635 | 1.144975 | 0.938993 | 0.830504 | 0.445456 | 0.296297 | 0.222579 | 0.221058 | 0.230443 | 0.241984 | 0.235466 | 0.220470 | 0.214739 | 0.228444 | 0.261985 | 0.303552 | 0.348133 | 0.295428 | 0.243926 | 0.220184 | 0.214694 | 0.214692 | 0.215464 | 0.214851 | 0.217522 | 0.231353 | 0.228252 | 0.231672 | 0.227751 | 0.227490 | 0.216862 | 0.214873 | 0.216405 | 0.221083 | 0.225737 | 0.224679 | 0.227710 | 0.220066 | 0.220197 | 0.214635 | 0.220650 | 0.227513 | 0.220563 | 0.217424 | 0.214706 | 0.214770 | 0.221871 | 0.235615 | 0.244636 | 0.231752 | 0.215694 | 0.215140 | 0.214702 | 0.215231 | 0.214778 | 0.214926 | 0.224122 | 0.269155 | 0.326453 | 0.359836 | 0.366411 | 0.398287 | 0.361332 | 0.320019 | 0.328509 | 0.287401 | 0.247978 | 0.262562 | 0.262855 | 0.278529 | 0.260186 | 0.228647 | 0.227506 | 0.222949 | 0.218925 | 0.223823 | 0.250077 | 0.323481 | 0.394065 | 0.363271 | 0.390591 | 0.297362 | 0.226131 | 0.218462 | 0.234729 | 0.243721 | 0.246804 | 0.243629 | 0.246238 | 0.240093 | 0.236504 | 0.242195 | 0.259457 | 0.295349 | 0.388386 | 0.531590 | 0.744421 | 1.368890 | 2.133172 | 2.173538 | 1.538849 | 1.392520 | 1.001023 | 0.764295 | 0.417120 | 0.281326 | 0.229749 | 0.217075 | 0.218907 | 0.236264 | 0.250060 | 0.240288 | 0.219324 | 0.214981 | 0.217673 | 0.216333 | 0.218742 | 0.221871 | 0.217294 | 0.214715 | 0.214674 | 0.215548 | 0.217797 | 0.217700 | 0.215931 | 0.215351 | 0.228355 | 0.248921 | 0.283963 | 0.384843 | 0.409350 | 0.394415 | 0.343644 | 0.300074 | 0.300923 | 0.316933 | 0.333427 | 0.340730 | 0.354112 | 0.354804 | 0.371106 | 0.296979 | 0.254412 | 0.219290 | 0.215006 | 0.218231 | 0.215090 | 0.214626 | 0.220085 | 0.247043 | 0.335758 | 0.544608 | 0.765016 | 1.031314 | 1.468465 | 1.820742 | 1.470815 | 1.140388 | 0.821572 | 0.810090 | 0.719401 | 0.564455 | 0.465319 | 0.452041 | 0.436702 | 0.457632 | 0.450771 | 0.470257 | 0.486008 | 0.514606 | 0.528785 | 0.581732 | 0.765760 | 0.965661 | 0.971837 | 0.925280 | 0.679622 | 0.551984 | 0.572008 | 0.491037 | 0.424509 | 0.470634 | 0.477946 | 0.530844 | 0.507885 | 0.437534 | 0.427502 | 0.425930 | 0.425131 | 0.454911 | 0.432632 | 0.425889 | 0.445808 | 0.454458 | 0.509100 | 0.536900 | 0.479851 | 0.500194 | 0.599305 | 0.614580 | 0.575115 | 0.475110 |
| right posterior | 0.306809 | 0.302236 | 0.245603 | 0.273702 | 0.218597 | 0.217025 | 0.223488 | 0.287266 | 0.428152 | 0.706380 | 0.485021 | 0.834923 | 2.060358 | 3.482483 | 4.789651 | 3.628419 | 6.977989 | 12.082603 | 1.871843 | 0.474223 | 0.247957 | 0.217857 | 0.273443 | 0.275895 | 0.247753 | 0.222905 | 0.225363 | 0.322476 | 0.432034 | 0.374740 | 0.252780 | 0.227733 | 0.256078 | 0.237342 | 0.215590 | 0.214890 | 0.220166 | 0.222748 | 0.262493 | 0.243727 | 0.220780 | 0.215124 | 0.217499 | 0.222714 | 0.230371 | 0.251803 | 0.328729 | 0.382311 | 0.464753 | 0.386637 | 0.297624 | 0.284285 | 0.268063 | 0.235610 | 0.229513 | 0.214668 | 0.228241 | 0.240543 | 0.301509 | 0.405929 | 0.615747 | 1.106604 | 1.473832 | 1.464863 | 1.635871 | 1.308407 | 1.529986 | 1.210976 | 0.638369 | 0.392141 | 0.341721 | 0.497757 | 1.694515 | 9.379512 | 51.255624 | 157.520541 | 138.893353 | 197.875032 | 164.091587 | 189.249062 | 218.870874 | 155.205143 | 92.569543 | 39.320362 | 20.800012 | 12.676554 | 4.979053 | 2.364307 | 1.266867 | 1.252153 | 1.853601 | 2.164852 | 2.374759 | 2.941641 | 3.917225 | 4.601328 | 3.231078 | 1.879054 | 1.473166 | 1.633126 | 1.821212 | 1.532076 | 1.142425 | 0.856855 | 0.589099 | 0.389777 | 0.283869 | 0.240624 | 0.229079 | 0.239826 | 0.281125 | 0.333289 | 0.416877 | 0.436823 | 0.465091 | 0.495074 | 0.544623 | 0.557961 | 0.591604 | 0.522300 | 0.518100 | 0.474548 | 0.360209 | 0.303400 | 0.287313 | 0.288929 | 0.347528 | 0.415930 | 0.470066 | 0.633377 | 0.936870 | 1.013153 | 1.034948 | 0.811417 | 0.630999 | 0.532558 | 0.459517 | 0.373210 | 0.339691 | 0.321568 | 0.307717 | 0.264786 | 0.242636 | 0.236878 | 0.241076 | 0.231760 | 0.216176 | 0.214631 | 0.216816 | 0.228053 | 0.262288 | 0.270204 | 0.281250 | 0.303452 | 0.324697 | 0.331554 | 0.318891 | 0.287064 | 0.277871 | 0.281219 | 0.306515 | 0.317949 | 0.304124 | 0.296262 | 0.309051 | 0.326142 | 0.375730 | 0.431775 | 0.542672 | 0.762684 | 1.189605 | 1.179662 | 1.160530 | 1.070758 | 0.974086 | 0.862582 | 0.912237 | 0.915913 | 0.942421 | 0.924677 | 0.827569 | 0.520990 | 0.365802 | 0.260395 | 0.223443 | 0.214744 | 0.214793 | 0.214633 | 0.227581 | 0.272176 | 0.375635 | 0.420936 | 0.552293 | 0.647844 | 0.554561 | 0.402142 | 0.312925 | 0.251566 | 0.239081 | 0.228096 | 0.227612 | 0.221414 | 0.236496 | 0.258731 | 0.312179 | 0.390527 | 0.463009 | 0.541364 | 0.797358 | 1.004048 | 1.077477 | 1.042834 | 0.988386 | 1.016736 | 1.138917 | 1.055765 | 0.841112 | 0.806631 | 0.680663 | 0.425295 | 0.271108 | 0.215096 | 0.216717 | 0.223604 | 0.247497 | 0.255316 | 0.250653 | 0.228593 | 0.214995 | 0.216758 | 0.222994 | 0.225453 | 0.221049 | 0.216288 | 0.214626 | 0.219684 | 0.254144 | 0.331887 | 0.356427 | 0.347738 | 0.327139 | 0.300348 | 0.251802 | 0.223073 | 0.216009 | 0.217955 | 0.218791 | 0.232599 | 0.251801 | 0.293878 | 0.316359 | 0.323311 | 0.298251 | 0.268276 | 0.221898 | 0.217269 | 0.249018 | 0.290758 | 0.286233 | 0.256416 | 0.227514 | 0.215201 | 0.221134 | 0.221861 | 0.218368 | 0.214713 | 0.214663 | 0.223681 | 0.254144 | 0.275333 | 0.261032 | 0.237637 | 0.226350 | 0.224869 | 0.215875 | 0.215679 | 0.231704 | 0.247618 | 0.266617 | 0.277914 |
| all electrodes | 0.218209 | 0.214649 | 0.246066 | 0.369958 | 0.648296 | 0.394838 | 0.286099 | 0.427082 | 0.425443 | 0.272655 | 0.214726 | 0.220236 | 0.229745 | 0.298964 | 0.292745 | 0.658973 | 1.543489 | 2.308891 | 4.233245 | 2.049068 | 2.421525 | 1.018955 | 0.238058 | 0.228830 | 0.337541 | 1.252337 | 1.542080 | 1.758355 | 1.461983 | 0.723899 | 0.422381 | 0.276014 | 0.214776 | 0.252533 | 0.281498 | 0.334504 | 0.408007 | 0.548874 | 0.574823 | 0.601096 | 0.370482 | 0.307636 | 0.255649 | 0.228143 | 0.217319 | 0.231570 | 0.246581 | 0.243896 | 0.226579 | 0.221931 | 0.224502 | 0.222263 | 0.224390 | 0.295748 | 0.293263 | 0.232221 | 0.216155 | 0.263154 | 0.295123 | 0.395195 | 0.492418 | 0.784952 | 1.296954 | 1.970452 | 1.747573 | 1.842024 | 1.960052 | 2.689326 | 1.988983 | 1.067396 | 0.709592 | 0.621623 | 0.745037 | 1.071469 | 1.824617 | 5.150371 | 10.014880 | 28.015731 | 61.457371 | 144.097748 | 585.139631 | 568.820101 | 65.236198 | 9.463627 | 3.053253 | 1.394766 | 1.197026 | 1.318376 | 1.894198 | 5.502212 | 21.201356 | 41.328750 | 61.024658 | 49.463785 | 28.127259 | 16.114461 | 14.254721 | 25.107702 | 48.861758 | 63.719098 | 63.897013 | 46.680925 | 23.057027 | 16.718344 | 12.105635 | 5.194612 | 1.608999 | 0.889768 | 0.672404 | 0.836694 | 0.883314 | 1.331384 | 2.827591 | 6.853718 | 3.441195 | 3.185202 | 3.206717 | 4.348107 | 2.287905 | 4.100688 | 6.807153 | 8.122790 | 5.915999 | 3.905119 | 3.372752 | 3.022740 | 0.904970 | 0.411897 | 0.305026 | 0.252439 | 0.219950 | 0.220072 | 0.239857 | 0.232422 | 0.223776 | 0.216482 | 0.214708 | 0.228775 | 0.235544 | 0.248355 | 0.261666 | 0.230368 | 0.216309 | 0.217840 | 0.239029 | 0.238961 | 0.225662 | 0.219804 | 0.217742 | 0.223603 | 0.239708 | 0.222325 | 0.230876 | 0.230588 | 0.220941 | 0.217888 | 0.228893 | 0.251021 | 0.314548 | 0.347770 | 0.319672 | 0.411129 | 0.441080 | 0.404478 | 0.311398 | 0.340287 | 0.492615 | 1.016156 | 0.914714 | 0.600382 | 0.426924 | 0.491177 | 0.563708 | 0.394587 | 0.295923 | 0.326982 | 0.347460 | 0.325588 | 0.316672 | 0.265514 | 0.241288 | 0.279302 | 0.278872 | 0.301052 | 0.314522 | 0.241152 | 0.229347 | 0.275345 | 0.236523 | 0.221645 | 0.221558 | 0.223562 | 0.228496 | 0.218428 | 0.215794 | 0.228251 | 0.271444 | 0.371640 | 0.386952 | 0.352257 | 0.346750 | 0.341104 | 0.372514 | 0.365956 | 0.301522 | 0.307427 | 0.252399 | 0.220798 | 0.215205 | 0.216468 | 0.229578 | 0.277143 | 0.448335 | 0.481829 | 0.548484 | 0.425821 | 0.393708 | 0.344489 | 0.252412 | 0.220164 | 0.217762 | 0.261751 | 0.299827 | 0.317493 | 0.299499 | 0.271443 | 0.290357 | 0.284377 | 0.237225 | 0.214865 | 0.215806 | 0.218978 | 0.224517 | 0.242654 | 0.270231 | 0.261768 | 0.240214 | 0.236879 | 0.251541 | 0.280400 | 0.371479 | 0.368028 | 0.309431 | 0.276983 | 0.237722 | 0.226051 | 0.225445 | 0.216151 | 0.237066 | 0.232978 | 0.220661 | 0.221287 | 0.234460 | 0.226056 | 0.247250 | 0.261337 | 0.260897 | 0.228874 | 0.214653 | 0.223667 | 0.233605 | 0.300534 | 0.366925 | 0.434613 | 0.406599 | 0.361571 | 0.335718 | 0.311580 | 0.257447 | 0.226476 | 0.221446 | 0.214668 | 0.221164 | 0.238131 | 0.257348 | 0.329164 | 0.357523 | 0.441262 | 0.513382 | 0.594350 |

Searchlight, spatiotemporal cluster permutation test

|  | start time | stop time | peak time | peak channel | cluster p | peak Cohen's d | direction |
| --- | --- | --- | --- | --- | --- | --- | --- |
| #1 | 75 | 905 | 205 | P4 | 0.0005 | 0.949451 | positive |

H) sex, angry

  
|  | time window | peak latency | cluster *p* | peak Cohen's *d* |  | | | |  |
| **all electrodes** | 210 - 365 ms | 340 ms | 0.0211 | 0.7002 |  | | | |  |
|  | | | | | | | | |  |

Time-resolved classification, cluster permutation tests

|  | **left hemisphere** | | | | **right hemisphere** | | | |  |
|  | time window | peak latency | cluster *p* | peak Cohen's *d* | time window | peak latency | cluster *p* | peak Cohen's *d* |  |
| **anterior** |  | | | |  | | | |  |
| **central** | 125 - 250 ms | 230 ms | 0.0329 | 0.5487 | 215 - 335 ms | 300 ms | 0.0226 | 0.8209 |  |
 285 - 365 ms | 335 ms | 0.044 | 1.0003 | 415 - 585 ms | 510 ms | 0.0154 | 0.7193 ||  | **posterior** |  | | | | 155 - 555 ms | 280 ms | 0.0004 | 1.2214 |

  

Time-resolved classification, Bayesian statistics

|  | -200 | -195 | -190 | -185 | -180 | -175 | -170 | -165 | -160 | -155 | -150 | -145 | -140 | -135 | -130 | -125 | -120 | -115 | -110 | -105 | -100 | -95 | -90 | -85 | -80 | -75 | -70 | -65 | -60 | -55 | -50 | -45 | -40 | -35 | -30 | -25 | -20 | -15 | -10 | -5 | 0 | 5 | 10 | 15 | 20 | 25 | 30 | 35 | 40 | 45 | 50 | 55 | 60 | 65 | 70 | 75 | 80 | 85 | 90 | 95 | 100 | 105 | 110 | 115 | 120 | 125 | 130 | 135 | 140 | 145 | 150 | 155 | 160 | 165 | 170 | 175 | 180 | 185 | 190 | 195 | 200 | 205 | 210 | 215 | 220 | 225 | 230 | 235 | 240 | 245 | 250 | 255 | 260 | 265 | 270 | 275 | 280 | 285 | 290 | 295 | 300 | 305 | 310 | 315 | 320 | 325 | 330 | 335 | 340 | 345 | 350 | 355 | 360 | 365 | 370 | 375 | 380 | 385 | 390 | 395 | 400 | 405 | 410 | 415 | 420 | 425 | 430 | 435 | 440 | 445 | 450 | 455 | 460 | 465 | 470 | 475 | 480 | 485 | 490 | 495 | 500 | 505 | 510 | 515 | 520 | 525 | 530 | 535 | 540 | 545 | 550 | 555 | 560 | 565 | 570 | 575 | 580 | 585 | 590 | 595 | 600 | 605 | 610 | 615 | 620 | 625 | 630 | 635 | 640 | 645 | 650 | 655 | 660 | 665 | 670 | 675 | 680 | 685 | 690 | 695 | 700 | 705 | 710 | 715 | 720 | 725 | 730 | 735 | 740 | 745 | 750 | 755 | 760 | 765 | 770 | 775 | 780 | 785 | 790 | 795 | 800 | 805 | 810 | 815 | 820 | 825 | 830 | 835 | 840 | 845 | 850 | 855 | 860 | 865 | 870 | 875 | 880 | 885 | 890 | 895 | 900 | 905 | 910 | 915 | 920 | 925 | 930 | 935 | 940 | 945 | 950 | 955 | 960 | 965 | 970 | 975 | 980 | 985 | 990 | 995 | 1000 | 1005 | 1010 | 1015 | 1020 | 1025 | 1030 | 1035 | 1040 | 1045 | 1050 | 1055 | 1060 | 1065 | 1070 | 1075 | 1080 | 1085 | 1090 | 1095 | 1100 | 1105 | 1110 | 1115 | 1120 | 1125 | 1130 | 1135 | 1140 | 1145 | 1150 | 1155 | 1160 | 1165 | 1170 | 1175 | 1180 | 1185 | 1190 | 1195 |
| --- | --- | --- | --- | --- | --- | --- | --- | --- | --- | --- | --- | --- | --- | --- | --- | --- | --- | --- | --- | --- | --- | --- | --- | --- | --- | --- | --- | --- | --- | --- | --- | --- | --- | --- | --- | --- | --- | --- | --- | --- | --- | --- | --- | --- | --- | --- | --- | --- | --- | --- | --- | --- | --- | --- | --- | --- | --- | --- | --- | --- | --- | --- | --- | --- | --- | --- | --- | --- | --- | --- | --- | --- | --- | --- | --- | --- | --- | --- | --- | --- | --- | --- | --- | --- | --- | --- | --- | --- | --- | --- | --- | --- | --- | --- | --- | --- | --- | --- | --- | --- | --- | --- | --- | --- | --- | --- | --- | --- | --- | --- | --- | --- | --- | --- | --- | --- | --- | --- | --- | --- | --- | --- | --- | --- | --- | --- | --- | --- | --- | --- | --- | --- | --- | --- | --- | --- | --- | --- | --- | --- | --- | --- | --- | --- | --- | --- | --- | --- | --- | --- | --- | --- | --- | --- | --- | --- | --- | --- | --- | --- | --- | --- | --- | --- | --- | --- | --- | --- | --- | --- | --- | --- | --- | --- | --- | --- | --- | --- | --- | --- | --- | --- | --- | --- | --- | --- | --- | --- | --- | --- | --- | --- | --- | --- | --- | --- | --- | --- | --- | --- | --- | --- | --- | --- | --- | --- | --- | --- | --- | --- | --- | --- | --- | --- | --- | --- | --- | --- | --- | --- | --- | --- | --- | --- | --- | --- | --- | --- | --- | --- | --- | --- | --- | --- | --- | --- | --- | --- | --- | --- | --- | --- | --- | --- | --- | --- | --- | --- | --- | --- | --- | --- | --- | --- | --- | --- | --- | --- | --- | --- | --- | --- | --- | --- | --- | --- | --- | --- | --- | --- | --- | --- | --- | --- | --- | --- | --- | --- | --- | --- |
| left anterior | 0.226145 | 0.262576 | 0.292328 | 0.311342 | 0.314363 | 0.295310 | 0.252024 | 0.215494 | 0.240763 | 0.249483 | 0.286895 | 0.249362 | 0.237534 | 0.215957 | 0.230631 | 0.246228 | 0.219110 | 0.215814 | 0.214674 | 0.227854 | 0.220407 | 0.228449 | 0.222956 | 0.215037 | 0.214681 | 0.217109 | 0.214642 | 0.215357 | 0.237518 | 0.280975 | 0.299319 | 0.413319 | 0.365060 | 0.258068 | 0.216263 | 0.222156 | 0.249817 | 0.236231 | 0.219169 | 0.223822 | 0.230580 | 0.228681 | 0.219414 | 0.221344 | 0.216564 | 0.216595 | 0.237577 | 0.226900 | 0.215962 | 0.262733 | 0.359336 | 0.444369 | 0.706696 | 0.785666 | 0.703055 | 0.541245 | 0.482985 | 0.349051 | 0.251062 | 0.223800 | 0.232555 | 0.271315 | 0.478120 | 0.609083 | 0.528171 | 0.534046 | 0.381506 | 0.305145 | 0.302781 | 0.251402 | 0.274875 | 0.419085 | 0.696666 | 1.059457 | 1.488088 | 2.008894 | 2.359213 | 2.330104 | 2.122496 | 1.887266 | 2.528424 | 1.538371 | 0.752677 | 0.648167 | 0.628852 | 0.524843 | 0.570010 | 0.721988 | 1.051976 | 1.638562 | 1.776671 | 1.588937 | 2.500574 | 2.310200 | 2.856988 | 2.984610 | 6.185417 | 5.394576 | 5.648086 | 3.673597 | 4.169258 | 3.851086 | 4.914337 | 2.838640 | 6.233689 | 22.668367 | 17.573525 | 8.328250 | 2.234206 | 0.712900 | 0.377936 | 0.234359 | 0.219810 | 0.226744 | 0.235569 | 0.230950 | 0.215878 | 0.216347 | 0.252302 | 0.376319 | 0.596246 | 0.990989 | 1.528636 | 1.636175 | 1.571772 | 1.034232 | 0.882870 | 0.736902 | 0.555719 | 0.353732 | 0.285530 | 0.260608 | 0.252585 | 0.224824 | 0.215655 | 0.222206 | 0.257001 | 0.305937 | 0.343051 | 0.335716 | 0.389938 | 0.449557 | 0.478209 | 0.402310 | 0.336776 | 0.302630 | 0.382353 | 0.477850 | 0.539172 | 0.528050 | 0.501936 | 0.551563 | 0.527929 | 0.368872 | 0.254329 | 0.222309 | 0.214647 | 0.215372 | 0.216912 | 0.217101 | 0.218058 | 0.215155 | 0.217290 | 0.229153 | 0.275960 | 0.379014 | 0.579532 | 0.780553 | 0.842482 | 0.745719 | 0.725609 | 0.581095 | 0.475283 | 0.427191 | 0.534339 | 0.739244 | 0.718977 | 0.659922 | 0.592851 | 0.603644 | 0.675958 | 0.694816 | 0.723674 | 0.659032 | 0.691685 | 0.792100 | 0.709608 | 0.658665 | 0.628570 | 0.592822 | 0.946382 | 0.990714 | 0.718040 | 0.575118 | 0.457054 | 0.325854 | 0.262375 | 0.218110 | 0.214939 | 0.219060 | 0.216649 | 0.215052 | 0.234774 | 0.294532 | 0.336833 | 0.321527 | 0.288200 | 0.247130 | 0.226752 | 0.220955 | 0.216741 | 0.214636 | 0.215239 | 0.215543 | 0.214966 | 0.234525 | 0.317372 | 0.443795 | 0.453507 | 0.570861 | 0.605964 | 0.577647 | 0.431347 | 0.370351 | 0.316592 | 0.299636 | 0.247290 | 0.224104 | 0.216741 | 0.241129 | 0.258063 | 0.265581 | 0.243600 | 0.220915 | 0.216463 | 0.215112 | 0.214924 | 0.217953 | 0.233334 | 0.283070 | 0.337273 | 0.320974 | 0.289664 | 0.223630 | 0.221581 | 0.269341 | 0.305820 | 0.297037 | 0.253698 | 0.244306 | 0.220650 | 0.215105 | 0.225394 | 0.227554 | 0.230907 | 0.263797 | 0.452596 | 0.678416 | 0.859207 | 0.814167 | 0.725255 | 0.721923 | 0.483133 | 0.356506 | 0.330461 | 0.391854 | 0.534462 | 0.792952 | 1.042323 | 1.032461 | 0.786012 | 0.758753 | 0.621469 | 0.572552 | 0.462616 | 0.451025 | 0.541744 | 0.623509 | 0.647847 | 0.782024 |
| right anterior | 0.678739 | 0.993865 | 2.549758 | 5.412034 | 4.492897 | 0.800553 | 0.466272 | 0.513421 | 0.274501 | 0.229950 | 0.224999 | 0.265269 | 0.216523 | 0.214647 | 0.216374 | 0.215118 | 0.262660 | 0.219749 | 0.216304 | 0.229252 | 0.246903 | 0.216673 | 0.214626 | 0.233385 | 0.215787 | 0.234980 | 0.219034 | 0.223070 | 0.233248 | 0.214756 | 0.219004 | 0.217497 | 0.235005 | 0.214626 | 0.254511 | 0.248920 | 0.287141 | 0.344462 | 0.344737 | 0.260830 | 0.324448 | 0.350285 | 0.366164 | 0.537987 | 0.672358 | 0.894331 | 0.939569 | 0.637095 | 0.557136 | 0.645842 | 0.789505 | 0.929719 | 2.826966 | 18.072868 | 64.528468 | 102.051093 | 373.043704 | 444.281224 | 367.284541 | 70.740730 | 22.148817 | 11.543678 | 3.676768 | 0.832331 | 0.267423 | 0.235876 | 0.241809 | 0.215245 | 0.226803 | 0.224387 | 0.227445 | 0.219912 | 0.227051 | 0.232729 | 0.225892 | 0.215100 | 0.220223 | 0.222309 | 0.231608 | 0.218356 | 0.218901 | 0.247441 | 0.280487 | 0.367531 | 0.478679 | 0.766616 | 1.062169 | 1.188812 | 1.678197 | 1.663220 | 1.014983 | 0.496611 | 0.288074 | 0.223914 | 0.214633 | 0.215375 | 0.223718 | 0.264201 | 0.313457 | 0.374825 | 0.455926 | 0.426899 | 0.388615 | 0.362941 | 0.361080 | 0.392345 | 0.440443 | 0.582443 | 0.728957 | 0.611273 | 0.537400 | 0.458996 | 0.432425 | 0.561285 | 0.541202 | 0.582162 | 0.697186 | 0.761204 | 0.946605 | 1.220947 | 1.281038 | 1.664837 | 2.367947 | 2.710260 | 3.029213 | 2.999092 | 2.644698 | 1.644505 | 1.689702 | 1.615125 | 1.786802 | 1.579649 | 1.158739 | 1.279067 | 1.074221 | 0.835083 | 0.843317 | 0.640265 | 0.615155 | 0.631822 | 0.644674 | 1.208391 | 1.717464 | 2.231567 | 3.642360 | 3.810690 | 3.538570 | 2.523995 | 1.613874 | 1.284344 | 1.002141 | 0.689279 | 0.529988 | 0.541897 | 0.556965 | 0.524922 | 0.384389 | 0.301786 | 0.299949 | 0.383695 | 0.457796 | 0.546473 | 0.778762 | 1.603223 | 3.681457 | 5.224354 | 4.787698 | 4.888035 | 3.493749 | 2.630375 | 2.120237 | 2.150415 | 2.182772 | 2.962746 | 3.021372 | 3.574505 | 3.936095 | 4.530462 | 3.015988 | 3.041255 | 1.861191 | 1.830626 | 1.308599 | 0.774359 | 0.640920 | 0.719160 | 0.523007 | 0.373775 | 0.346733 | 0.328110 | 0.298101 | 0.330305 | 0.433614 | 0.535811 | 0.787903 | 0.791079 | 0.816272 | 0.842191 | 0.725874 | 0.572478 | 0.445931 | 0.445889 | 0.399031 | 0.466524 | 0.591428 | 0.613622 | 0.584787 | 0.537707 | 0.383754 | 0.388507 | 0.439593 | 0.473384 | 0.480810 | 0.458602 | 0.492225 | 0.487094 | 0.439588 | 0.337518 | 0.309219 | 0.280003 | 0.290159 | 0.282349 | 0.294320 | 0.364278 | 0.417503 | 0.393820 | 0.415617 | 0.348101 | 0.314786 | 0.292239 | 0.247322 | 0.252492 | 0.270508 | 0.271223 | 0.300673 | 0.364175 | 0.492670 | 0.700128 | 0.569628 | 0.434912 | 0.430240 | 0.439729 | 0.394346 | 0.310171 | 0.266953 | 0.296919 | 0.359600 | 0.378785 | 0.368075 | 0.388012 | 0.514801 | 0.661236 | 0.581377 | 0.425911 | 0.390303 | 0.366824 | 0.372852 | 0.346542 | 0.317844 | 0.331139 | 0.399831 | 0.415656 | 0.490895 | 0.551812 | 0.576875 | 0.659272 | 0.602477 | 0.708015 | 1.028192 | 1.429940 | 1.660614 | 2.033408 | 2.183164 | 2.584534 | 2.090389 | 1.407517 | 1.125758 | 0.927733 | 0.805231 | 0.749600 |
| left central | 0.270655 | 0.324890 | 0.395246 | 0.333632 | 0.225041 | 0.216778 | 0.221605 | 0.215830 | 0.216607 | 0.215348 | 0.214697 | 0.215335 | 0.215672 | 0.219450 | 0.237195 | 0.258308 | 0.402923 | 1.761339 | 1.701460 | 7.975379 | 19.878139 | 5.853386 | 1.384734 | 0.985976 | 0.698293 | 0.765534 | 1.123903 | 1.228514 | 1.839834 | 4.115238 | 5.277482 | 4.149705 | 5.586911 | 5.863315 | 5.470011 | 1.995925 | 0.880784 | 0.390117 | 0.280917 | 0.245485 | 0.216471 | 0.235021 | 0.281629 | 0.321200 | 0.301822 | 0.245937 | 0.226911 | 0.217256 | 0.216007 | 0.229566 | 0.231444 | 0.222113 | 0.224576 | 0.224742 | 0.232779 | 0.227532 | 0.218180 | 0.215137 | 0.215497 | 0.215356 | 0.227581 | 0.256067 | 0.356527 | 0.512254 | 0.765068 | 1.715521 | 1.682883 | 2.183320 | 3.076654 | 5.564732 | 5.816936 | 7.225052 | 6.475060 | 8.839446 | 6.708081 | 4.559607 | 3.601645 | 3.074998 | 2.875680 | 5.159528 | 5.261760 | 3.737750 | 2.431954 | 2.443176 | 3.572681 | 4.816796 | 3.854692 | 3.690408 | 3.479907 | 2.494768 | 1.398128 | 0.733162 | 0.481967 | 0.403349 | 0.537716 | 0.632970 | 0.858869 | 1.902493 | 2.991014 | 3.381697 | 5.371343 | 8.817186 | 19.229538 | 56.323594 | 76.863445 | 304.663064 | 551.041247 | 430.293019 | 126.604029 | 97.812650 | 21.621200 | 7.407046 | 2.636123 | 1.675728 | 1.034721 | 0.867726 | 1.128688 | 1.533500 | 1.521294 | 1.276750 | 0.956566 | 0.859032 | 0.823919 | 0.776565 | 0.853896 | 0.830348 | 0.972877 | 1.083765 | 1.620825 | 1.994389 | 1.746107 | 1.921932 | 1.952249 | 1.876914 | 2.206203 | 1.864690 | 2.225088 | 2.258013 | 2.230488 | 2.485557 | 1.540986 | 1.304221 | 1.463993 | 1.318100 | 1.213916 | 0.941484 | 0.900700 | 1.699904 | 2.671880 | 3.366359 | 3.774836 | 2.430056 | 2.689656 | 2.482302 | 1.883572 | 1.135299 | 0.705177 | 0.462673 | 0.384921 | 0.276351 | 0.238275 | 0.228904 | 0.226089 | 0.222201 | 0.220278 | 0.219863 | 0.224988 | 0.222443 | 0.217666 | 0.217202 | 0.218001 | 0.220538 | 0.225643 | 0.223093 | 0.231361 | 0.259809 | 0.246243 | 0.252554 | 0.280758 | 0.356638 | 0.482027 | 0.515610 | 0.563137 | 0.779579 | 0.890075 | 1.268121 | 0.834100 | 0.660401 | 0.513724 | 0.376306 | 0.323345 | 0.340104 | 0.289389 | 0.312769 | 0.297744 | 0.257682 | 0.231130 | 0.217334 | 0.216507 | 0.224721 | 0.237184 | 0.236439 | 0.220476 | 0.216874 | 0.214661 | 0.214707 | 0.227506 | 0.258736 | 0.265571 | 0.262230 | 0.281692 | 0.292146 | 0.341450 | 0.343399 | 0.357918 | 0.418217 | 0.415334 | 0.440371 | 0.511565 | 0.485042 | 0.356845 | 0.268051 | 0.235776 | 0.232161 | 0.234005 | 0.229821 | 0.226607 | 0.245459 | 0.259938 | 0.276988 | 0.262152 | 0.242896 | 0.237319 | 0.240740 | 0.225395 | 0.230674 | 0.228549 | 0.244721 | 0.252665 | 0.268989 | 0.252816 | 0.246543 | 0.246574 | 0.246828 | 0.238912 | 0.224448 | 0.216780 | 0.235951 | 0.263809 | 0.315996 | 0.340553 | 0.352209 | 0.291611 | 0.227747 | 0.214652 | 0.227262 | 0.249952 | 0.279596 | 0.291763 | 0.276066 | 0.241562 | 0.222402 | 0.217378 | 0.229696 | 0.236854 | 0.241358 | 0.237347 | 0.240380 | 0.248721 | 0.238134 | 0.216233 | 0.220246 | 0.245752 | 0.279128 | 0.292758 | 0.276002 | 0.256417 | 0.261514 | 0.249165 | 0.232115 |
| right central | 0.215592 | 0.218785 | 0.215092 | 0.228530 | 0.337157 | 0.529817 | 0.814074 | 0.832670 | 1.098382 | 0.742406 | 0.645580 | 0.411453 | 0.336874 | 0.302866 | 0.250972 | 0.215117 | 0.240755 | 0.280515 | 0.393382 | 0.344673 | 0.324176 | 0.262980 | 0.238142 | 0.226280 | 0.225814 | 0.216287 | 0.233667 | 0.294950 | 0.363964 | 0.506031 | 0.861908 | 2.012815 | 5.299232 | 4.966312 | 4.022767 | 3.048567 | 2.820435 | 1.553781 | 0.725902 | 0.398401 | 0.276684 | 0.222332 | 0.216817 | 0.270743 | 0.337532 | 0.333146 | 0.287863 | 0.273643 | 0.219131 | 0.215541 | 0.217626 | 0.217647 | 0.214673 | 0.217187 | 0.218506 | 0.226859 | 0.214895 | 0.237760 | 0.372807 | 0.741584 | 0.791521 | 1.133293 | 1.521056 | 1.728788 | 1.278033 | 0.949657 | 0.644598 | 0.786035 | 0.620762 | 0.417785 | 0.300812 | 0.330328 | 0.380102 | 0.613242 | 0.871413 | 1.211648 | 0.657740 | 0.457144 | 0.335662 | 0.279810 | 0.253495 | 0.306964 | 0.512383 | 2.357696 | 9.117757 | 11.412162 | 19.742217 | 35.766666 | 51.977791 | 37.575725 | 19.307149 | 23.380352 | 30.248088 | 29.093861 | 40.950356 | 54.661042 | 86.618582 | 92.667724 | 57.701646 | 50.957255 | 60.947372 | 47.317718 | 31.916126 | 17.264857 | 15.853385 | 10.213260 | 3.595568 | 2.118860 | 1.051486 | 0.860209 | 0.687196 | 0.362081 | 0.286835 | 0.375813 | 0.556361 | 1.008545 | 1.242177 | 1.681675 | 2.712819 | 3.394665 | 2.822177 | 1.895090 | 1.299156 | 1.693747 | 2.035321 | 1.650980 | 1.700703 | 2.668544 | 4.076722 | 7.283005 | 8.280353 | 12.239610 | 17.956909 | 25.020754 | 18.251588 | 11.957243 | 11.709057 | 12.826126 | 12.119149 | 10.472600 | 10.255364 | 13.041514 | 20.812298 | 19.491394 | 12.717956 | 10.031687 | 9.733343 | 8.326991 | 6.562040 | 5.332809 | 7.320392 | 9.422601 | 10.711311 | 12.079261 | 20.110039 | 14.936632 | 5.860166 | 1.504323 | 1.012509 | 0.713397 | 0.844660 | 0.807354 | 1.091780 | 1.741808 | 2.974019 | 3.176663 | 3.675841 | 2.548445 | 1.844409 | 1.695724 | 1.602364 | 1.353166 | 0.840702 | 0.529318 | 0.415119 | 0.318227 | 0.266106 | 0.259080 | 0.270957 | 0.277285 | 0.268784 | 0.296611 | 0.365911 | 0.441622 | 0.493897 | 0.762262 | 1.402706 | 3.141879 | 3.804771 | 4.016476 | 2.190604 | 1.754039 | 1.417929 | 1.738728 | 2.409429 | 3.664326 | 3.845598 | 3.791255 | 1.889444 | 1.328784 | 0.655267 | 0.449180 | 0.340126 | 0.289080 | 0.316378 | 0.408052 | 0.445764 | 0.533031 | 0.477368 | 0.395699 | 0.395004 | 0.343542 | 0.381071 | 0.413151 | 0.351503 | 0.303846 | 0.293696 | 0.256090 | 0.255883 | 0.222550 | 0.216705 | 0.214653 | 0.216011 | 0.219831 | 0.216873 | 0.215013 | 0.217537 | 0.218009 | 0.226801 | 0.245797 | 0.286445 | 0.320225 | 0.307630 | 0.266001 | 0.281039 | 0.300674 | 0.301252 | 0.272308 | 0.244963 | 0.257204 | 0.290062 | 0.326763 | 0.338159 | 0.331175 | 0.311290 | 0.287033 | 0.257699 | 0.227597 | 0.215079 | 0.244926 | 0.294259 | 0.307587 | 0.326158 | 0.404007 | 0.418016 | 0.339736 | 0.255115 | 0.236873 | 0.232978 | 0.228917 | 0.214694 | 0.214807 | 0.214871 | 0.219740 | 0.225600 | 0.235002 | 0.233605 | 0.247452 | 0.241216 | 0.243047 | 0.247435 | 0.259580 | 0.241672 | 0.252788 | 0.339615 | 0.565647 | 0.949074 | 1.201710 | 1.299488 | 2.480770 |
| left posterior | 0.218726 | 0.241757 | 0.220983 | 0.220743 | 0.275568 | 0.241952 | 0.291393 | 0.467861 | 0.654687 | 0.784337 | 0.998655 | 0.585255 | 0.906749 | 0.917773 | 0.638690 | 1.086088 | 0.307988 | 0.234975 | 0.277087 | 0.335705 | 0.327214 | 0.313568 | 0.306642 | 0.244840 | 0.225241 | 0.217604 | 0.219182 | 0.232011 | 0.226428 | 0.221309 | 0.216316 | 0.222963 | 0.241842 | 0.228138 | 0.222513 | 0.220986 | 0.226180 | 0.250942 | 0.238082 | 0.225573 | 0.219272 | 0.215593 | 0.232558 | 0.288108 | 0.511630 | 1.290588 | 1.459950 | 0.964880 | 0.431759 | 0.277784 | 0.242252 | 0.223141 | 0.215230 | 0.218379 | 0.232513 | 0.287216 | 0.378988 | 0.615353 | 1.626384 | 4.083523 | 11.757887 | 18.463526 | 14.845643 | 7.771463 | 2.743789 | 0.707466 | 0.268259 | 0.218260 | 0.282801 | 0.333532 | 0.346398 | 0.331524 | 0.288244 | 0.227150 | 0.214694 | 0.226490 | 0.248774 | 0.292799 | 0.377244 | 0.505235 | 0.597940 | 0.841491 | 0.843017 | 0.823078 | 1.019129 | 1.662873 | 2.141159 | 1.994133 | 2.272282 | 3.331939 | 4.197550 | 3.926766 | 3.303276 | 2.809749 | 2.405043 | 1.729410 | 1.510559 | 1.301921 | 1.042840 | 0.880373 | 0.993842 | 1.323085 | 1.734689 | 1.782885 | 2.561801 | 5.316958 | 10.087220 | 8.017615 | 5.449296 | 3.558108 | 2.371283 | 1.471113 | 1.113389 | 0.807349 | 0.622989 | 0.534606 | 0.558434 | 0.700392 | 0.868966 | 0.925179 | 0.925583 | 1.065082 | 1.281202 | 1.118153 | 0.814370 | 0.839567 | 0.719701 | 0.476864 | 0.322465 | 0.266633 | 0.260246 | 0.266677 | 0.259232 | 0.258517 | 0.306976 | 0.391200 | 0.436708 | 0.442089 | 0.435557 | 0.426075 | 0.372094 | 0.341362 | 0.351901 | 0.353280 | 0.328472 | 0.324060 | 0.319038 | 0.319143 | 0.300103 | 0.265754 | 0.238147 | 0.237839 | 0.217125 | 0.214877 | 0.216132 | 0.223249 | 0.248786 | 0.279012 | 0.316415 | 0.286564 | 0.261984 | 0.253502 | 0.241505 | 0.230471 | 0.215852 | 0.219634 | 0.232272 | 0.245940 | 0.267640 | 0.291915 | 0.307288 | 0.257561 | 0.227493 | 0.219393 | 0.215474 | 0.217545 | 0.223444 | 0.226384 | 0.217145 | 0.214911 | 0.219420 | 0.237448 | 0.291604 | 0.407570 | 0.521845 | 0.706191 | 0.870714 | 0.911212 | 0.763735 | 0.668914 | 0.516065 | 0.522610 | 0.428564 | 0.372083 | 0.310299 | 0.267837 | 0.256159 | 0.256271 | 0.244925 | 0.232212 | 0.225630 | 0.237325 | 0.288681 | 0.359641 | 0.502960 | 0.728618 | 1.332824 | 1.975890 | 3.130154 | 3.599797 | 2.292783 | 1.035966 | 0.519420 | 0.348147 | 0.267018 | 0.227337 | 0.217586 | 0.216528 | 0.223490 | 0.241621 | 0.249880 | 0.277550 | 0.299857 | 0.305829 | 0.336262 | 0.333301 | 0.291264 | 0.266362 | 0.232606 | 0.223524 | 0.226096 | 0.230975 | 0.244427 | 0.268477 | 0.318374 | 0.495102 | 0.959760 | 1.673800 | 2.682167 | 3.079892 | 3.987652 | 3.572332 | 2.878412 | 1.919472 | 1.245720 | 0.658515 | 0.410358 | 0.312098 | 0.257962 | 0.222196 | 0.219860 | 0.252799 | 0.284866 | 0.290788 | 0.295419 | 0.265590 | 0.240981 | 0.215240 | 0.220476 | 0.263405 | 0.300557 | 0.297092 | 0.274370 | 0.247068 | 0.217894 | 0.215439 | 0.229890 | 0.243539 | 0.230676 | 0.232986 | 0.238099 | 0.252166 | 0.268837 | 0.309479 | 0.394299 | 0.547847 | 0.558506 | 0.614457 | 0.513473 | 0.454724 |
| right posterior | 0.514517 | 0.814510 | 1.525234 | 3.090675 | 2.855345 | 1.889789 | 3.107674 | 5.624687 | 2.374967 | 1.617271 | 1.036885 | 1.283010 | 2.534821 | 1.792394 | 1.372246 | 1.883345 | 2.428396 | 1.892216 | 0.688863 | 0.420150 | 0.459413 | 0.308056 | 0.261327 | 0.225099 | 0.215375 | 0.216596 | 0.214626 | 0.220779 | 0.233366 | 0.216232 | 0.219210 | 0.231938 | 0.283629 | 0.422899 | 0.570802 | 0.765900 | 0.872386 | 0.574496 | 0.413402 | 0.340777 | 0.434150 | 0.343169 | 0.280158 | 0.247237 | 0.273750 | 0.326156 | 0.377693 | 0.373217 | 0.493701 | 0.488243 | 0.326209 | 0.236688 | 0.215226 | 0.230993 | 0.338606 | 0.672516 | 2.356923 | 5.196910 | 11.823730 | 28.996773 | 81.678236 | 223.488919 | 1057.881776 | 592.199882 | 110.706703 | 19.126376 | 4.935999 | 2.060176 | 0.824525 | 0.525175 | 0.996931 | 3.852116 | 14.986760 | 35.418688 | 40.003929 | 72.434907 | 112.699916 | 97.659918 | 124.582209 | 94.902836 | 106.799856 | 207.193005 | 190.865318 | 221.109838 | 249.169328 | 219.906905 | 145.104225 | 62.859803 | 40.728981 | 40.118245 | 62.887503 | 93.964862 | 146.423153 | 298.314775 | 734.364718 | 1948.626973 | 4771.930110 | 5726.053115 | 3168.906572 | 1602.339446 | 1905.640496 | 1436.451963 | 1431.619827 | 522.794274 | 268.820095 | 153.033505 | 114.917035 | 106.312380 | 165.289406 | 155.220874 | 175.807833 | 190.909098 | 232.763780 | 267.268843 | 285.907351 | 429.869513 | 532.873190 | 382.980764 | 360.028180 | 199.898854 | 117.160116 | 40.559529 | 13.225170 | 7.102105 | 3.383848 | 2.561203 | 2.774129 | 2.191554 | 2.521299 | 3.036720 | 2.740111 | 3.669998 | 4.986323 | 5.512297 | 5.857227 | 5.339099 | 5.599170 | 8.488846 | 16.619213 | 31.238503 | 46.625020 | 84.833484 | 126.120526 | 149.688722 | 81.954820 | 38.756364 | 13.996723 | 7.197456 | 3.912626 | 2.178688 | 1.476833 | 1.342157 | 1.170488 | 1.105557 | 1.021256 | 1.014838 | 1.011183 | 0.885124 | 0.771191 | 0.664678 | 0.668434 | 0.710379 | 0.714033 | 0.636834 | 0.743619 | 0.782058 | 0.754430 | 0.619192 | 0.453023 | 0.334482 | 0.266464 | 0.232188 | 0.217878 | 0.214854 | 0.214679 | 0.220265 | 0.252025 | 0.350919 | 0.476349 | 0.701037 | 1.174469 | 2.086224 | 3.637358 | 3.683005 | 3.225087 | 3.237207 | 3.526714 | 3.806865 | 3.615872 | 2.124280 | 1.700720 | 1.601377 | 1.366283 | 1.156281 | 0.706631 | 0.556393 | 0.505301 | 0.460497 | 0.441740 | 0.414054 | 0.341709 | 0.302602 | 0.285089 | 0.322845 | 0.367997 | 0.420248 | 0.456571 | 0.549173 | 0.596007 | 0.554270 | 0.370268 | 0.295046 | 0.261277 | 0.283719 | 0.294045 | 0.303432 | 0.276450 | 0.258680 | 0.258123 | 0.284742 | 0.265228 | 0.229290 | 0.215877 | 0.220642 | 0.226686 | 0.238339 | 0.234315 | 0.232992 | 0.238592 | 0.270250 | 0.279792 | 0.347755 | 0.362840 | 0.375782 | 0.386176 | 0.571851 | 1.060310 | 3.311429 | 5.751629 | 9.648918 | 16.578353 | 30.251599 | 21.408325 | 8.591108 | 3.311418 | 2.287128 | 1.303537 | 0.655041 | 0.337268 | 0.250857 | 0.216357 | 0.214633 | 0.214779 | 0.218997 | 0.258666 | 0.478409 | 0.917090 | 1.507887 | 1.616534 | 1.582904 | 0.916802 | 0.504589 | 0.339782 | 0.303492 | 0.331642 | 0.367614 | 0.418571 | 0.561588 | 0.770088 | 0.961480 | 1.018326 | 0.672817 | 0.426220 | 0.264798 | 0.224487 | 0.215020 | 0.225726 | 0.245482 | 0.244429 | 0.243197 |
| all electrodes | 0.217852 | 0.261344 | 0.442122 | 0.665351 | 0.634223 | 0.730741 | 0.767441 | 0.299253 | 0.214757 | 0.375814 | 0.627580 | 0.683325 | 0.502946 | 0.459969 | 0.234775 | 0.251151 | 0.215012 | 0.215010 | 0.227382 | 0.593387 | 2.050275 | 0.815362 | 0.774543 | 0.928858 | 1.215374 | 0.805739 | 0.521195 | 0.467101 | 0.498945 | 0.444049 | 0.283205 | 0.270136 | 0.283097 | 0.282612 | 0.306413 | 0.303075 | 0.260748 | 0.249461 | 0.218063 | 0.216589 | 0.231739 | 0.444263 | 0.947324 | 1.122752 | 1.886588 | 1.891110 | 2.493989 | 1.770310 | 0.689619 | 0.481920 | 0.422550 | 0.367310 | 0.303036 | 0.261645 | 0.244999 | 0.223190 | 0.223604 | 0.334241 | 0.929363 | 2.153162 | 4.623341 | 17.261097 | 28.117447 | 11.470619 | 2.642291 | 0.484899 | 0.240252 | 0.217297 | 0.216222 | 0.214749 | 0.215618 | 0.254748 | 0.734248 | 2.694579 | 4.394726 | 3.041686 | 2.606507 | 1.885686 | 1.883480 | 1.578537 | 1.266833 | 1.086857 | 1.432582 | 1.480232 | 2.337092 | 3.052510 | 3.718143 | 5.625293 | 7.338237 | 10.802446 | 29.808835 | 42.053910 | 21.821769 | 13.984518 | 10.078884 | 11.229821 | 9.184665 | 4.113768 | 2.494699 | 2.408737 | 1.656282 | 2.106181 | 2.611124 | 4.302541 | 8.721219 | 11.078124 | 8.757190 | 15.365356 | 17.083570 | 13.932684 | 8.925944 | 4.153743 | 2.719115 | 2.193485 | 1.289121 | 0.821247 | 0.818822 | 0.809948 | 1.168102 | 1.508661 | 2.694026 | 3.819499 | 4.813497 | 2.989224 | 1.676173 | 1.635121 | 1.393409 | 1.020619 | 0.880586 | 0.596208 | 0.513694 | 0.475473 | 0.366979 | 0.289371 | 0.245899 | 0.232905 | 0.231136 | 0.239423 | 0.295660 | 0.326621 | 0.416332 | 0.713124 | 1.531579 | 3.597853 | 4.620473 | 2.762469 | 2.938993 | 3.180411 | 2.021759 | 1.200664 | 0.780044 | 0.575261 | 0.620840 | 0.509615 | 0.363920 | 0.262460 | 0.220842 | 0.215554 | 0.219879 | 0.226919 | 0.225972 | 0.219881 | 0.215024 | 0.249534 | 0.472702 | 1.129940 | 2.234388 | 2.846072 | 2.006422 | 1.345109 | 1.033236 | 0.600465 | 0.380777 | 0.314929 | 0.266153 | 0.278837 | 0.318344 | 0.315931 | 0.321088 | 0.411837 | 0.499894 | 0.676610 | 0.766303 | 0.834628 | 0.986274 | 1.900504 | 2.739797 | 4.698333 | 8.106032 | 7.592145 | 4.737801 | 3.142090 | 2.171191 | 1.054002 | 0.623738 | 0.447306 | 0.415965 | 0.439643 | 0.421284 | 0.390180 | 0.360604 | 0.364004 | 0.398645 | 0.436853 | 0.503863 | 0.572849 | 0.633148 | 0.985306 | 0.943911 | 0.662920 | 0.496794 | 0.382812 | 0.432283 | 0.584297 | 0.682919 | 0.707127 | 0.690486 | 0.516392 | 0.421225 | 0.332196 | 0.244773 | 0.218170 | 0.214768 | 0.214889 | 0.234166 | 0.301937 | 0.357775 | 0.450758 | 0.665214 | 0.869326 | 0.816567 | 0.631337 | 0.438215 | 0.410001 | 0.384578 | 0.339071 | 0.338656 | 0.326634 | 0.310932 | 0.361915 | 0.402664 | 0.465631 | 0.596692 | 0.573946 | 0.693817 | 0.680758 | 0.584657 | 0.513384 | 0.539101 | 0.427904 | 0.391609 | 0.331047 | 0.339182 | 0.344932 | 0.374974 | 0.390426 | 0.433355 | 0.398509 | 0.358128 | 0.361271 | 0.339930 | 0.299356 | 0.292468 | 0.278315 | 0.291148 | 0.313916 | 0.329312 | 0.408140 | 0.444352 | 0.427283 | 0.417319 | 0.453405 | 0.457135 | 0.472090 | 0.367433 | 0.321153 | 0.280096 | 0.278658 | 0.247946 | 0.232047 |

Searchlight, spatiotemporal cluster permutation test

|  | start time | stop time | peak time | peak channel | cluster p | peak Cohen's d | direction |
| --- | --- | --- | --- | --- | --- | --- | --- |
| #1 | 75 | 650 | 390 | P4 | 0.0002 | 0.952955 | positive |

I) sex, sad

  
|  | time window | peak latency | cluster *p* | peak Cohen's *d* |  | | | |
| **all electrodes** |  | | | |  | | | |
|  | | | | | | | | |

Time-resolved classification, cluster permutation tests

|  | **left hemisphere** | | | | **right hemisphere** | | | |
|  | time window | peak latency | cluster *p* | peak Cohen's *d* | time window | peak latency | cluster *p* | peak Cohen's *d* |
| **anterior** |  | | | |  | | | |
| **central** |  | | | |  | | | |
| **posterior** | 1100 - 1195 ms | 1165 ms | 0.026 | 0.7253 |  | | | |

  

Time-resolved classification, Bayesian statistics

|  | -200 | -195 | -190 | -185 | -180 | -175 | -170 | -165 | -160 | -155 | -150 | -145 | -140 | -135 | -130 | -125 | -120 | -115 | -110 | -105 | -100 | -95 | -90 | -85 | -80 | -75 | -70 | -65 | -60 | -55 | -50 | -45 | -40 | -35 | -30 | -25 | -20 | -15 | -10 | -5 | 0 | 5 | 10 | 15 | 20 | 25 | 30 | 35 | 40 | 45 | 50 | 55 | 60 | 65 | 70 | 75 | 80 | 85 | 90 | 95 | 100 | 105 | 110 | 115 | 120 | 125 | 130 | 135 | 140 | 145 | 150 | 155 | 160 | 165 | 170 | 175 | 180 | 185 | 190 | 195 | 200 | 205 | 210 | 215 | 220 | 225 | 230 | 235 | 240 | 245 | 250 | 255 | 260 | 265 | 270 | 275 | 280 | 285 | 290 | 295 | 300 | 305 | 310 | 315 | 320 | 325 | 330 | 335 | 340 | 345 | 350 | 355 | 360 | 365 | 370 | 375 | 380 | 385 | 390 | 395 | 400 | 405 | 410 | 415 | 420 | 425 | 430 | 435 | 440 | 445 | 450 | 455 | 460 | 465 | 470 | 475 | 480 | 485 | 490 | 495 | 500 | 505 | 510 | 515 | 520 | 525 | 530 | 535 | 540 | 545 | 550 | 555 | 560 | 565 | 570 | 575 | 580 | 585 | 590 | 595 | 600 | 605 | 610 | 615 | 620 | 625 | 630 | 635 | 640 | 645 | 650 | 655 | 660 | 665 | 670 | 675 | 680 | 685 | 690 | 695 | 700 | 705 | 710 | 715 | 720 | 725 | 730 | 735 | 740 | 745 | 750 | 755 | 760 | 765 | 770 | 775 | 780 | 785 | 790 | 795 | 800 | 805 | 810 | 815 | 820 | 825 | 830 | 835 | 840 | 845 | 850 | 855 | 860 | 865 | 870 | 875 | 880 | 885 | 890 | 895 | 900 | 905 | 910 | 915 | 920 | 925 | 930 | 935 | 940 | 945 | 950 | 955 | 960 | 965 | 970 | 975 | 980 | 985 | 990 | 995 | 1000 | 1005 | 1010 | 1015 | 1020 | 1025 | 1030 | 1035 | 1040 | 1045 | 1050 | 1055 | 1060 | 1065 | 1070 | 1075 | 1080 | 1085 | 1090 | 1095 | 1100 | 1105 | 1110 | 1115 | 1120 | 1125 | 1130 | 1135 | 1140 | 1145 | 1150 | 1155 | 1160 | 1165 | 1170 | 1175 | 1180 | 1185 | 1190 | 1195 |
| --- | --- | --- | --- | --- | --- | --- | --- | --- | --- | --- | --- | --- | --- | --- | --- | --- | --- | --- | --- | --- | --- | --- | --- | --- | --- | --- | --- | --- | --- | --- | --- | --- | --- | --- | --- | --- | --- | --- | --- | --- | --- | --- | --- | --- | --- | --- | --- | --- | --- | --- | --- | --- | --- | --- | --- | --- | --- | --- | --- | --- | --- | --- | --- | --- | --- | --- | --- | --- | --- | --- | --- | --- | --- | --- | --- | --- | --- | --- | --- | --- | --- | --- | --- | --- | --- | --- | --- | --- | --- | --- | --- | --- | --- | --- | --- | --- | --- | --- | --- | --- | --- | --- | --- | --- | --- | --- | --- | --- | --- | --- | --- | --- | --- | --- | --- | --- | --- | --- | --- | --- | --- | --- | --- | --- | --- | --- | --- | --- | --- | --- | --- | --- | --- | --- | --- | --- | --- | --- | --- | --- | --- | --- | --- | --- | --- | --- | --- | --- | --- | --- | --- | --- | --- | --- | --- | --- | --- | --- | --- | --- | --- | --- | --- | --- | --- | --- | --- | --- | --- | --- | --- | --- | --- | --- | --- | --- | --- | --- | --- | --- | --- | --- | --- | --- | --- | --- | --- | --- | --- | --- | --- | --- | --- | --- | --- | --- | --- | --- | --- | --- | --- | --- | --- | --- | --- | --- | --- | --- | --- | --- | --- | --- | --- | --- | --- | --- | --- | --- | --- | --- | --- | --- | --- | --- | --- | --- | --- | --- | --- | --- | --- | --- | --- | --- | --- | --- | --- | --- | --- | --- | --- | --- | --- | --- | --- | --- | --- | --- | --- | --- | --- | --- | --- | --- | --- | --- | --- | --- | --- | --- | --- | --- | --- | --- | --- | --- | --- | --- | --- | --- | --- | --- | --- | --- | --- | --- | --- | --- | --- | --- |
| left anterior | 0.214660 | 0.224811 | 0.235660 | 0.225679 | 0.221026 | 0.224297 | 0.224244 | 0.236094 | 0.253055 | 0.242400 | 0.280632 | 0.474989 | 0.995293 | 0.975327 | 0.508958 | 0.328996 | 0.406331 | 0.369119 | 0.911621 | 1.427579 | 1.135321 | 0.866310 | 0.484477 | 0.239572 | 0.223469 | 0.326967 | 0.555098 | 0.825049 | 0.775375 | 0.387318 | 0.260901 | 0.220949 | 0.394905 | 1.128517 | 4.685779 | 3.380435 | 2.151516 | 1.707076 | 1.679682 | 0.605243 | 0.307517 | 0.214897 | 0.240275 | 0.215317 | 0.280349 | 0.264829 | 0.282540 | 0.265046 | 0.272311 | 0.282800 | 0.224893 | 0.215643 | 0.216964 | 0.214821 | 0.240087 | 0.358378 | 0.589977 | 0.669096 | 0.836986 | 0.835576 | 0.649651 | 0.379471 | 0.239595 | 0.216125 | 0.218548 | 0.245029 | 0.267456 | 0.335926 | 0.330298 | 0.324594 | 0.352910 | 0.575913 | 0.518177 | 0.477671 | 0.432182 | 0.429509 | 0.461932 | 0.376061 | 0.282862 | 0.237110 | 0.217871 | 0.239224 | 0.337147 | 0.690484 | 1.324611 | 3.174642 | 4.846468 | 5.794704 | 3.034032 | 1.358331 | 0.557370 | 0.245973 | 0.236607 | 0.349760 | 0.397633 | 0.444011 | 0.638112 | 1.134840 | 0.981829 | 0.547392 | 0.443639 | 0.596860 | 0.597405 | 0.502556 | 0.341362 | 0.344290 | 0.369908 | 0.413825 | 0.376339 | 0.326813 | 0.309610 | 0.362546 | 0.388131 | 0.471498 | 0.404837 | 0.411331 | 0.416111 | 0.368320 | 0.277094 | 0.228681 | 0.218364 | 0.234888 | 0.267019 | 0.283319 | 0.329776 | 0.354903 | 0.324245 | 0.254171 | 0.225517 | 0.215248 | 0.215875 | 0.217068 | 0.219511 | 0.230238 | 0.258600 | 0.319501 | 0.417841 | 0.407427 | 0.346254 | 0.268024 | 0.242601 | 0.246153 | 0.228537 | 0.216461 | 0.215989 | 0.223516 | 0.250066 | 0.267063 | 0.254122 | 0.224302 | 0.223205 | 0.225054 | 0.225779 | 0.227259 | 0.236560 | 0.269193 | 0.415874 | 0.566820 | 0.559906 | 0.387381 | 0.270078 | 0.250723 | 0.228525 | 0.218386 | 0.216277 | 0.215941 | 0.215810 | 0.224452 | 0.219579 | 0.220138 | 0.216425 | 0.225540 | 0.264588 | 0.389765 | 0.594058 | 1.024735 | 1.791110 | 4.535329 | 5.305932 | 5.572283 | 5.090184 | 4.482400 | 4.394777 | 5.128646 | 5.972589 | 8.626765 | 8.467801 | 6.759387 | 5.930501 | 3.503275 | 1.680920 | 0.902385 | 0.458056 | 0.323190 | 0.265308 | 0.242742 | 0.240283 | 0.227179 | 0.223196 | 0.239169 | 0.258639 | 0.295593 | 0.308867 | 0.306759 | 0.308606 | 0.265876 | 0.223317 | 0.216721 | 0.214635 | 0.222386 | 0.221574 | 0.217021 | 0.214999 | 0.215054 | 0.216873 | 0.217742 | 0.218579 | 0.233854 | 0.263192 | 0.309647 | 0.347255 | 0.331385 | 0.375964 | 0.356813 | 0.270935 | 0.235414 | 0.215679 | 0.219243 | 0.232287 | 0.253888 | 0.267062 | 0.254603 | 0.252174 | 0.251748 | 0.236023 | 0.216558 | 0.215347 | 0.218322 | 0.227927 | 0.218793 | 0.220490 | 0.218621 | 0.215290 | 0.215534 | 0.222156 | 0.245680 | 0.249795 | 0.270280 | 0.307982 | 0.307936 | 0.283945 | 0.242748 | 0.216951 | 0.219266 | 0.224919 | 0.239206 | 0.242251 | 0.225444 | 0.217168 | 0.216138 | 0.233721 | 0.240375 | 0.282272 | 0.372154 | 0.490990 | 1.039677 | 2.466294 | 5.664513 | 8.288454 | 8.510842 | 7.011221 | 3.392170 | 1.561671 | 0.802447 | 0.579414 | 0.464285 | 0.410857 | 0.364444 | 0.335192 | 0.311560 |
| right anterior | 1.857725 | 0.843473 | 0.806256 | 1.322512 | 1.144782 | 0.769717 | 0.516728 | 0.878311 | 1.590310 | 3.869548 | 8.018705 | 8.790532 | 2.094897 | 2.169068 | 1.910886 | 2.393812 | 1.170601 | 0.255113 | 0.236075 | 0.225222 | 0.229315 | 0.234025 | 0.299579 | 0.780276 | 1.682142 | 0.715339 | 0.300352 | 0.346812 | 0.265815 | 0.234171 | 0.726316 | 3.727533 | 9.431044 | 17.817600 | 12.133355 | 1.402328 | 0.299613 | 0.214938 | 0.224872 | 0.216923 | 0.415840 | 1.432851 | 6.674244 | 4.260411 | 5.237360 | 12.360546 | 6.865936 | 3.288926 | 4.998876 | 5.342742 | 3.487781 | 3.390585 | 1.902632 | 0.881676 | 0.384925 | 0.245296 | 0.214636 | 0.219383 | 0.228555 | 0.233214 | 0.229601 | 0.215846 | 0.241281 | 0.280332 | 0.373888 | 0.411643 | 0.529912 | 0.708930 | 0.727938 | 0.945359 | 0.881153 | 0.987114 | 0.897701 | 0.526324 | 0.302797 | 0.222092 | 0.225600 | 0.288111 | 0.295152 | 0.268464 | 0.259571 | 0.237279 | 0.242993 | 0.263538 | 0.278640 | 0.284542 | 0.273619 | 0.230186 | 0.214991 | 0.220474 | 0.245548 | 0.392894 | 0.771447 | 0.918877 | 0.825031 | 0.712879 | 0.854575 | 1.088668 | 0.925402 | 0.538705 | 0.499262 | 0.433409 | 0.342976 | 0.309165 | 0.272592 | 0.280025 | 0.290189 | 0.306552 | 0.281993 | 0.310678 | 0.364611 | 0.574635 | 0.684785 | 0.732961 | 0.701538 | 0.905341 | 0.943932 | 0.577606 | 0.310069 | 0.224999 | 0.214635 | 0.225488 | 0.220356 | 0.216623 | 0.215315 | 0.219732 | 0.248099 | 0.260914 | 0.263707 | 0.232196 | 0.229364 | 0.215835 | 0.214877 | 0.215924 | 0.214950 | 0.214706 | 0.214948 | 0.221507 | 0.226656 | 0.245295 | 0.255757 | 0.293833 | 0.291738 | 0.287343 | 0.287103 | 0.250616 | 0.224594 | 0.215507 | 0.214681 | 0.216421 | 0.215236 | 0.216103 | 0.218605 | 0.220203 | 0.214706 | 0.214745 | 0.217770 | 0.220526 | 0.224867 | 0.230576 | 0.232242 | 0.226772 | 0.236308 | 0.237416 | 0.221780 | 0.215203 | 0.223231 | 0.243732 | 0.265036 | 0.300055 | 0.293221 | 0.275915 | 0.246396 | 0.219877 | 0.214755 | 0.214813 | 0.217142 | 0.214753 | 0.214672 | 0.214704 | 0.216802 | 0.217183 | 0.214671 | 0.216345 | 0.214791 | 0.215394 | 0.221488 | 0.225580 | 0.238389 | 0.226466 | 0.232771 | 0.252550 | 0.244250 | 0.226882 | 0.220680 | 0.216497 | 0.217404 | 0.214822 | 0.218214 | 0.217168 | 0.215529 | 0.217652 | 0.221148 | 0.224156 | 0.220775 | 0.215310 | 0.214700 | 0.215019 | 0.217713 | 0.228768 | 0.249738 | 0.243854 | 0.220839 | 0.215490 | 0.224043 | 0.226836 | 0.233804 | 0.237103 | 0.234377 | 0.220121 | 0.214825 | 0.223284 | 0.220253 | 0.220313 | 0.216962 | 0.214631 | 0.218094 | 0.225980 | 0.244916 | 0.241024 | 0.237136 | 0.229590 | 0.220816 | 0.218161 | 0.217663 | 0.214934 | 0.214661 | 0.214693 | 0.214631 | 0.214631 | 0.217645 | 0.214828 | 0.217415 | 0.224702 | 0.227516 | 0.228471 | 0.222776 | 0.230416 | 0.221621 | 0.218979 | 0.219538 | 0.218410 | 0.219143 | 0.223022 | 0.225398 | 0.224863 | 0.220048 | 0.217290 | 0.223829 | 0.238971 | 0.269468 | 0.311964 | 0.346423 | 0.349451 | 0.348244 | 0.336111 | 0.334497 | 0.334591 | 0.332833 | 0.339531 | 0.402034 | 0.527071 | 0.717947 | 0.818344 | 0.799983 | 0.895908 | 0.949487 | 0.969338 | 0.929085 | 0.731542 |
| left central | 0.226870 | 0.214853 | 0.214674 | 0.215982 | 0.215232 | 0.221027 | 0.221521 | 0.220229 | 0.217959 | 0.216917 | 0.218614 | 0.219728 | 0.215762 | 0.216934 | 0.217764 | 0.235993 | 0.227458 | 0.215708 | 0.214641 | 0.242527 | 0.296045 | 0.509653 | 0.744270 | 0.600651 | 0.232973 | 0.220364 | 0.378287 | 1.257014 | 5.591563 | 10.674019 | 1.492931 | 0.307783 | 0.214967 | 0.272883 | 0.451701 | 0.700682 | 1.032594 | 0.577492 | 0.477521 | 0.377125 | 0.291581 | 0.258575 | 0.225639 | 0.217487 | 0.261189 | 0.251744 | 0.241878 | 0.215379 | 0.214626 | 0.223467 | 0.246365 | 0.343837 | 0.401229 | 0.521110 | 0.591389 | 0.589757 | 0.470896 | 0.302837 | 0.226502 | 0.215219 | 0.216376 | 0.215008 | 0.215049 | 0.218674 | 0.262050 | 0.315638 | 0.241833 | 0.216196 | 0.261978 | 0.365585 | 0.396578 | 0.325210 | 0.271152 | 0.215558 | 0.245895 | 0.433953 | 1.060814 | 1.626997 | 1.134491 | 0.702961 | 0.361427 | 0.266128 | 0.224667 | 0.220245 | 0.226832 | 0.229655 | 0.217868 | 0.216523 | 0.224222 | 0.220722 | 0.217935 | 0.221885 | 0.230397 | 0.227440 | 0.217827 | 0.214947 | 0.214881 | 0.214626 | 0.216097 | 0.223151 | 0.233671 | 0.242829 | 0.255253 | 0.297194 | 0.302108 | 0.291578 | 0.262968 | 0.267873 | 0.305249 | 0.283312 | 0.250190 | 0.233497 | 0.224617 | 0.234153 | 0.239833 | 0.224509 | 0.225550 | 0.229084 | 0.233125 | 0.257314 | 0.269462 | 0.294686 | 0.393819 | 0.412143 | 0.399628 | 0.436387 | 0.400067 | 0.385439 | 0.368322 | 0.280094 | 0.261259 | 0.245105 | 0.237223 | 0.239845 | 0.265894 | 0.316132 | 0.496578 | 0.648133 | 0.703552 | 0.548158 | 0.486662 | 0.360306 | 0.288348 | 0.264913 | 0.271269 | 0.279408 | 0.279997 | 0.274697 | 0.268139 | 0.262776 | 0.246797 | 0.247811 | 0.245479 | 0.254863 | 0.271744 | 0.284163 | 0.277387 | 0.272579 | 0.251169 | 0.242954 | 0.243417 | 0.252365 | 0.252316 | 0.289562 | 0.312198 | 0.363158 | 0.406073 | 0.492131 | 0.494791 | 0.655996 | 0.549727 | 0.572533 | 0.502566 | 0.610808 | 0.649277 | 0.546454 | 0.417976 | 0.410852 | 0.504236 | 0.828079 | 1.119682 | 1.710968 | 4.387619 | 16.725085 | 43.461075 | 46.299207 | 25.940598 | 11.805073 | 3.051979 | 0.986361 | 0.469334 | 0.317722 | 0.250774 | 0.221983 | 0.214631 | 0.214632 | 0.215486 | 0.217856 | 0.221226 | 0.245291 | 0.307650 | 0.431515 | 0.531172 | 0.644077 | 1.086942 | 2.157520 | 2.813651 | 2.698064 | 3.024022 | 3.324460 | 2.669826 | 1.935279 | 1.284815 | 1.232311 | 1.412629 | 1.557558 | 1.266608 | 1.142056 | 0.836640 | 0.812641 | 0.747951 | 0.580778 | 0.404484 | 0.380129 | 0.349384 | 0.413319 | 0.475783 | 0.573686 | 0.648454 | 0.497974 | 0.374091 | 0.316643 | 0.248491 | 0.220596 | 0.214825 | 0.214756 | 0.217680 | 0.222048 | 0.225499 | 0.226079 | 0.222479 | 0.218635 | 0.214644 | 0.215182 | 0.216445 | 0.223968 | 0.222559 | 0.219464 | 0.223691 | 0.219182 | 0.224869 | 0.233402 | 0.229042 | 0.234297 | 0.231955 | 0.218826 | 0.215966 | 0.219612 | 0.253629 | 0.272166 | 0.266409 | 0.249537 | 0.227548 | 0.217130 | 0.215841 | 0.222862 | 0.240004 | 0.251960 | 0.263665 | 0.288759 | 0.275113 | 0.259957 | 0.306921 | 0.290146 | 0.267902 | 0.230686 | 0.215743 | 0.215887 | 0.221534 | 0.214660 |
| right central | 0.253863 | 0.337673 | 0.607514 | 0.802005 | 1.092156 | 1.075378 | 0.735904 | 0.384030 | 0.290355 | 0.215367 | 0.253778 | 0.232362 | 0.214738 | 0.214660 | 0.226441 | 0.225550 | 0.215993 | 0.233462 | 0.355326 | 0.465483 | 0.405834 | 0.443447 | 0.747591 | 0.569889 | 0.261982 | 0.214769 | 0.219326 | 0.222224 | 0.325772 | 0.757482 | 1.185803 | 0.880853 | 0.896158 | 0.986660 | 0.708616 | 0.407954 | 0.255330 | 0.230602 | 0.214704 | 0.214715 | 0.218294 | 0.214639 | 0.217331 | 0.217343 | 0.257515 | 0.414892 | 0.307669 | 0.237388 | 0.234541 | 0.234767 | 0.216971 | 0.216544 | 0.236342 | 0.217554 | 0.215032 | 0.221259 | 0.227291 | 0.225544 | 0.215833 | 0.215086 | 0.215937 | 0.215744 | 0.215370 | 0.218741 | 0.223814 | 0.235022 | 0.237142 | 0.233243 | 0.243821 | 0.256726 | 0.243476 | 0.229628 | 0.220800 | 0.214626 | 0.214972 | 0.217046 | 0.230877 | 0.215392 | 0.220434 | 0.264222 | 0.486237 | 0.646199 | 0.765130 | 0.962264 | 0.687506 | 0.672052 | 0.938923 | 0.858945 | 1.036617 | 0.953474 | 1.293260 | 1.965340 | 3.394886 | 4.753103 | 5.549440 | 3.893467 | 2.689939 | 1.804671 | 0.978701 | 0.468525 | 0.307491 | 0.233452 | 0.216143 | 0.214780 | 0.216275 | 0.214803 | 0.215178 | 0.216912 | 0.217139 | 0.215483 | 0.250913 | 0.260674 | 0.240039 | 0.243889 | 0.224379 | 0.215287 | 0.218744 | 0.246102 | 0.253115 | 0.255701 | 0.262892 | 0.240724 | 0.233052 | 0.227942 | 0.229158 | 0.222637 | 0.215113 | 0.218248 | 0.240343 | 0.261635 | 0.239659 | 0.215393 | 0.227936 | 0.250197 | 0.260852 | 0.287845 | 0.324122 | 0.297274 | 0.274494 | 0.296508 | 0.400302 | 0.707220 | 1.307724 | 0.969800 | 0.840930 | 0.531979 | 0.332863 | 0.265517 | 0.234665 | 0.219553 | 0.217789 | 0.216942 | 0.218609 | 0.227617 | 0.239437 | 0.285607 | 0.396962 | 0.662715 | 0.856967 | 0.955658 | 0.819760 | 0.796736 | 0.824247 | 0.679509 | 0.610526 | 0.531632 | 0.641268 | 1.270138 | 2.166611 | 2.704510 | 3.940009 | 5.927696 | 8.727532 | 6.786152 | 5.924508 | 7.893987 | 10.374496 | 13.428690 | 5.519497 | 3.907066 | 2.925472 | 2.173823 | 1.272673 | 0.820242 | 0.554072 | 0.628689 | 0.648877 | 0.628353 | 0.527204 | 0.439737 | 0.361894 | 0.416078 | 0.363370 | 0.304872 | 0.295263 | 0.305314 | 0.335006 | 0.427252 | 0.416201 | 0.387753 | 0.388608 | 0.480018 | 0.669106 | 0.941539 | 1.294054 | 1.245738 | 1.448776 | 1.711538 | 1.926953 | 2.276193 | 2.631482 | 3.162061 | 5.201915 | 8.006255 | 9.657378 | 9.919599 | 11.243399 | 9.434586 | 5.055035 | 2.047187 | 1.293267 | 0.944885 | 0.733393 | 0.733605 | 0.715032 | 0.738289 | 0.834968 | 1.063636 | 1.269027 | 1.233984 | 0.901984 | 0.684990 | 0.407244 | 0.324991 | 0.265585 | 0.239314 | 0.228867 | 0.233683 | 0.259832 | 0.344554 | 0.427295 | 0.438030 | 0.476468 | 0.576471 | 0.909280 | 1.121194 | 2.545881 | 3.148148 | 3.366375 | 3.818737 | 3.408655 | 1.919485 | 1.548924 | 0.989617 | 0.775042 | 0.861677 | 0.887835 | 1.316951 | 1.237458 | 1.081801 | 1.139259 | 1.218247 | 1.124115 | 1.252928 | 1.019254 | 0.868645 | 0.910380 | 0.920985 | 0.677408 | 0.545664 | 0.492566 | 0.510496 | 0.619241 | 0.610587 | 0.492208 | 0.696391 | 1.256068 | 1.401172 | 1.421624 | 1.215101 |
| left posterior | 0.352961 | 0.304375 | 0.335740 | 0.311220 | 0.265848 | 0.216707 | 0.216709 | 0.250782 | 0.253253 | 0.273005 | 0.235462 | 0.214788 | 0.247957 | 0.380804 | 0.564679 | 0.825953 | 0.890559 | 0.805845 | 0.690379 | 0.657598 | 0.808720 | 2.925352 | 2.317838 | 0.825324 | 0.579474 | 0.416738 | 0.293229 | 0.217481 | 0.267427 | 0.270062 | 0.220311 | 0.221622 | 0.253774 | 0.342236 | 0.358821 | 0.291008 | 0.280338 | 0.255025 | 0.236174 | 0.216481 | 0.227040 | 0.299261 | 0.454944 | 0.827310 | 0.694922 | 0.692835 | 0.499026 | 0.367587 | 0.318534 | 0.258860 | 0.227365 | 0.214658 | 0.226472 | 0.215306 | 0.226582 | 0.300010 | 0.685468 | 1.215781 | 5.104492 | 57.759823 | 140.171130 | 207.348400 | 212.955175 | 105.659537 | 34.175104 | 6.658801 | 1.343106 | 0.526834 | 0.291040 | 0.236743 | 0.222256 | 0.266132 | 0.302847 | 0.259360 | 0.267311 | 0.354373 | 0.500595 | 0.690120 | 0.601881 | 0.686204 | 1.163157 | 1.506898 | 1.278328 | 0.829503 | 0.793936 | 0.643178 | 0.555083 | 0.414270 | 0.337214 | 0.335365 | 0.382433 | 0.399026 | 0.489429 | 0.593791 | 0.748337 | 0.937977 | 0.760457 | 0.574236 | 0.476858 | 0.440451 | 0.412645 | 0.354081 | 0.377789 | 0.484126 | 0.886405 | 0.957524 | 0.839820 | 0.809179 | 0.801007 | 0.632950 | 0.443240 | 0.313567 | 0.264799 | 0.234432 | 0.222704 | 0.222309 | 0.219733 | 0.221644 | 0.215928 | 0.216789 | 0.224330 | 0.223417 | 0.218090 | 0.214658 | 0.217497 | 0.243209 | 0.302828 | 0.454194 | 0.719023 | 1.115901 | 1.390102 | 1.409241 | 1.268318 | 1.114939 | 0.853134 | 0.454578 | 0.294495 | 0.247677 | 0.223911 | 0.216410 | 0.216105 | 0.214636 | 0.230998 | 0.265647 | 0.332264 | 0.528871 | 0.721388 | 0.794111 | 0.693520 | 0.424683 | 0.311665 | 0.266853 | 0.230333 | 0.218640 | 0.215295 | 0.216614 | 0.214752 | 0.224804 | 0.228696 | 0.235970 | 0.263515 | 0.318083 | 0.493815 | 0.641022 | 0.810899 | 1.173443 | 1.656813 | 1.813676 | 2.638259 | 2.605226 | 2.633327 | 1.943264 | 1.227913 | 0.760199 | 0.545112 | 0.417214 | 0.310057 | 0.294622 | 0.295620 | 0.347710 | 0.463356 | 0.652660 | 1.115539 | 1.636773 | 1.685044 | 1.823843 | 1.553169 | 1.173134 | 0.850343 | 0.453247 | 0.328523 | 0.286422 | 0.268267 | 0.253376 | 0.257808 | 0.266250 | 0.273575 | 0.310364 | 0.313421 | 0.301702 | 0.336235 | 0.346297 | 0.394458 | 0.473777 | 0.449883 | 0.392045 | 0.384638 | 0.351188 | 0.396667 | 0.398499 | 0.365540 | 0.320316 | 0.312248 | 0.301376 | 0.248924 | 0.216975 | 0.214693 | 0.214634 | 0.218418 | 0.235365 | 0.251921 | 0.326368 | 0.445882 | 0.614334 | 0.745785 | 0.613651 | 0.476114 | 0.355412 | 0.304985 | 0.272846 | 0.237446 | 0.221012 | 0.218196 | 0.216048 | 0.214779 | 0.217227 | 0.218222 | 0.222032 | 0.221166 | 0.217360 | 0.215542 | 0.215783 | 0.214923 | 0.214849 | 0.218770 | 0.223826 | 0.228653 | 0.249522 | 0.337849 | 0.426716 | 0.415036 | 0.400356 | 0.404024 | 0.474648 | 0.452241 | 0.391951 | 0.380895 | 0.461366 | 0.755466 | 1.130687 | 1.629324 | 2.407645 | 4.348305 | 5.176028 | 6.610018 | 7.290610 | 10.899668 | 10.815170 | 11.561626 | 9.522011 | 11.448711 | 11.724178 | 14.276082 | 22.148175 | 20.628303 | 11.346143 | 7.064830 | 6.140163 | 4.315337 | 2.433143 |
| right posterior | 1.420251 | 0.641243 | 0.416641 | 0.394074 | 0.282293 | 0.219675 | 0.215051 | 0.214642 | 0.214782 | 0.214936 | 0.219034 | 0.234845 | 0.261682 | 0.217073 | 0.222267 | 0.262883 | 0.233875 | 0.214852 | 0.216147 | 0.229631 | 0.217993 | 0.235842 | 0.271125 | 0.456760 | 1.221992 | 1.285083 | 1.512448 | 0.674715 | 0.354603 | 0.283677 | 0.220014 | 0.216565 | 0.220661 | 0.217148 | 0.226286 | 0.222919 | 0.215307 | 0.228819 | 0.241554 | 0.239188 | 0.225446 | 0.236358 | 0.276352 | 0.344140 | 0.406031 | 0.423862 | 0.460550 | 0.383315 | 0.304691 | 0.252586 | 0.225277 | 0.214889 | 0.215237 | 0.215631 | 0.287840 | 0.624400 | 2.268429 | 11.816542 | 52.417919 | 150.853958 | 405.537060 | 320.115810 | 183.261560 | 63.048620 | 20.474683 | 3.009551 | 0.464505 | 0.219191 | 0.241630 | 0.371277 | 0.513686 | 0.667394 | 0.819379 | 0.763993 | 0.422979 | 0.264233 | 0.214686 | 0.263440 | 0.421440 | 0.942174 | 1.707520 | 2.502577 | 3.183870 | 2.667384 | 1.694888 | 0.964302 | 0.482077 | 0.249360 | 0.237923 | 0.384135 | 0.563258 | 0.472202 | 0.298781 | 0.227603 | 0.221175 | 0.257025 | 0.269230 | 0.282412 | 0.248322 | 0.219009 | 0.214745 | 0.222736 | 0.225020 | 0.239050 | 0.293847 | 0.306517 | 0.281581 | 0.298462 | 0.345690 | 0.386310 | 0.368284 | 0.255633 | 0.226141 | 0.216960 | 0.216011 | 0.225399 | 0.256424 | 0.282287 | 0.275856 | 0.259045 | 0.242838 | 0.231563 | 0.219340 | 0.214666 | 0.218636 | 0.223656 | 0.222667 | 0.222609 | 0.227794 | 0.228172 | 0.235704 | 0.258871 | 0.357712 | 0.583331 | 0.959737 | 1.460695 | 2.371969 | 3.770560 | 3.622813 | 1.735290 | 0.845052 | 0.542827 | 0.378471 | 0.290899 | 0.232842 | 0.218469 | 0.217662 | 0.231899 | 0.245557 | 0.242156 | 0.247644 | 0.263752 | 0.283935 | 0.355914 | 0.400266 | 0.533651 | 1.707474 | 4.677664 | 7.201158 | 7.999837 | 8.183489 | 6.915404 | 2.621023 | 0.604435 | 0.284262 | 0.224663 | 0.214726 | 0.219064 | 0.237269 | 0.237771 | 0.225949 | 0.215489 | 0.260106 | 0.418373 | 0.664171 | 1.071391 | 1.296251 | 2.436229 | 3.493270 | 2.857454 | 2.438429 | 1.753111 | 1.132513 | 0.859033 | 0.490326 | 0.329005 | 0.279475 | 0.229315 | 0.217358 | 0.214626 | 0.214703 | 0.215279 | 0.215382 | 0.215199 | 0.215446 | 0.215983 | 0.221721 | 0.234527 | 0.263179 | 0.355979 | 0.488287 | 0.734552 | 1.112618 | 2.077266 | 3.334241 | 5.190623 | 4.469513 | 2.927986 | 2.750412 | 3.453550 | 5.275177 | 6.201002 | 7.361988 | 12.640513 | 19.719167 | 7.461134 | 2.335426 | 1.062400 | 0.505283 | 0.318800 | 0.257697 | 0.234077 | 0.225211 | 0.218695 | 0.218220 | 0.233251 | 0.252088 | 0.283625 | 0.310599 | 0.322082 | 0.361799 | 0.354758 | 0.254140 | 0.235411 | 0.225390 | 0.225771 | 0.240670 | 0.245329 | 0.249257 | 0.278971 | 0.298020 | 0.317280 | 0.353911 | 0.389230 | 0.396250 | 0.363793 | 0.333188 | 0.260109 | 0.224948 | 0.216575 | 0.216400 | 0.222777 | 0.215439 | 0.214635 | 0.228740 | 0.273194 | 0.363093 | 0.482087 | 0.558637 | 0.323948 | 0.235838 | 0.215959 | 0.218545 | 0.270290 | 0.388063 | 0.643715 | 0.873627 | 1.039919 | 1.225031 | 1.116112 | 0.869740 | 0.910245 | 1.244079 | 2.159210 | 2.904878 | 3.207731 | 4.465566 | 6.643363 | 6.513313 | 2.647858 |
| all electrodes | 0.416665 | 0.267927 | 0.272447 | 0.320880 | 0.263022 | 0.253454 | 0.223890 | 0.256746 | 0.356338 | 0.349956 | 0.333269 | 0.422026 | 0.343148 | 0.313372 | 0.260486 | 0.233896 | 0.215265 | 0.216411 | 0.424296 | 0.526779 | 0.458570 | 0.326751 | 0.361363 | 0.311932 | 0.220125 | 0.252013 | 0.320707 | 0.307615 | 0.331830 | 0.576797 | 0.873433 | 0.413103 | 0.361354 | 0.219381 | 0.242919 | 0.411968 | 1.321243 | 6.960673 | 7.605895 | 7.045959 | 1.191242 | 0.376336 | 0.242178 | 0.222752 | 0.218353 | 0.256118 | 0.290352 | 0.302024 | 0.381090 | 0.368928 | 0.370541 | 0.457993 | 0.329404 | 0.325924 | 0.365030 | 0.397581 | 0.542806 | 0.835890 | 1.004791 | 4.012913 | 8.419614 | 10.146454 | 11.247235 | 8.051095 | 5.289167 | 5.388005 | 1.162076 | 0.374505 | 0.269110 | 0.229862 | 0.222716 | 0.227118 | 0.214811 | 0.218512 | 0.216294 | 0.220168 | 0.232851 | 0.251405 | 0.299857 | 0.592731 | 0.920439 | 0.858869 | 0.384104 | 0.268864 | 0.278380 | 0.303470 | 0.310838 | 0.333932 | 0.339473 | 0.298426 | 0.279672 | 0.286432 | 0.315995 | 0.345260 | 0.322317 | 0.277057 | 0.302000 | 0.329712 | 0.288014 | 0.242515 | 0.223690 | 0.215969 | 0.222062 | 0.220305 | 0.221943 | 0.229284 | 0.254672 | 0.308602 | 0.352637 | 0.380641 | 0.616591 | 1.103673 | 1.597595 | 1.491593 | 1.110574 | 1.143163 | 0.664842 | 0.372159 | 0.267393 | 0.225784 | 0.216867 | 0.214929 | 0.214857 | 0.215561 | 0.226948 | 0.229924 | 0.259667 | 0.297799 | 0.353801 | 0.338456 | 0.334394 | 0.332385 | 0.312949 | 0.287423 | 0.234673 | 0.218547 | 0.258400 | 0.290932 | 0.334171 | 0.305152 | 0.302357 | 0.291620 | 0.259845 | 0.220721 | 0.215077 | 0.217759 | 0.221182 | 0.218697 | 0.219327 | 0.223207 | 0.223840 | 0.216378 | 0.215377 | 0.215005 | 0.215216 | 0.216538 | 0.214770 | 0.216803 | 0.214709 | 0.214815 | 0.216175 | 0.233218 | 0.249327 | 0.242881 | 0.233977 | 0.230943 | 0.232441 | 0.219286 | 0.218347 | 0.235345 | 0.221260 | 0.214648 | 0.214709 | 0.215132 | 0.214706 | 0.216289 | 0.215233 | 0.215680 | 0.223600 | 0.249733 | 0.253622 | 0.271278 | 0.306123 | 0.307420 | 0.280688 | 0.281549 | 0.260731 | 0.256159 | 0.256658 | 0.270552 | 0.298402 | 0.351559 | 0.377085 | 0.423948 | 0.469002 | 0.483738 | 0.395799 | 0.352825 | 0.303635 | 0.296736 | 0.309998 | 0.361467 | 0.476668 | 0.853911 | 1.609932 | 2.584725 | 3.314375 | 3.371496 | 3.889089 | 3.986126 | 4.057458 | 5.522218 | 10.028670 | 11.713537 | 23.151270 | 37.421852 | 44.699799 | 23.854309 | 8.711702 | 3.600690 | 1.828584 | 1.007177 | 0.575808 | 0.337878 | 0.279982 | 0.293219 | 0.368199 | 0.519611 | 0.529764 | 0.566070 | 0.800135 | 0.954851 | 0.886008 | 0.549475 | 0.428199 | 0.435152 | 0.559204 | 0.777401 | 0.946660 | 1.581604 | 2.488510 | 3.441528 | 3.270087 | 1.712490 | 1.170146 | 0.943745 | 0.512094 | 0.399346 | 0.350387 | 0.323250 | 0.340308 | 0.307384 | 0.305714 | 0.286598 | 0.245886 | 0.228106 | 0.229383 | 0.229015 | 0.218331 | 0.216036 | 0.219314 | 0.220414 | 0.220924 | 0.224245 | 0.254427 | 0.259906 | 0.250926 | 0.240825 | 0.234463 | 0.235669 | 0.254446 | 0.256735 | 0.292284 | 0.326975 | 0.413938 | 0.497846 | 0.568392 | 0.601462 | 0.555517 | 0.434192 |

Searchlight, spatiotemporal cluster permutation test

No significant clusters observed.

J) difference, sex, happy vs angry

  
|  | time window | peak latency | cluster *p* | peak Cohen's *d* |  | | | |
| **all electrodes** |  | | | |  | | | |
|  | | | | | | | | |

Time-resolved classification, cluster permutation tests

|  | **left hemisphere** | | | | **right hemisphere** | | | |
|  | time window | peak latency | cluster *p* | peak Cohen's *d* | time window | peak latency | cluster *p* | peak Cohen's *d* |
| **anterior** |  | | | |  | | | |
| **central** |  | | | |  | | | |
| **posterior** |  | | | |  | | | |

  

Time-resolved classification, Bayesian statistics

|  | -200 | -195 | -190 | -185 | -180 | -175 | -170 | -165 | -160 | -155 | -150 | -145 | -140 | -135 | -130 | -125 | -120 | -115 | -110 | -105 | -100 | -95 | -90 | -85 | -80 | -75 | -70 | -65 | -60 | -55 | -50 | -45 | -40 | -35 | -30 | -25 | -20 | -15 | -10 | -5 | 0 | 5 | 10 | 15 | 20 | 25 | 30 | 35 | 40 | 45 | 50 | 55 | 60 | 65 | 70 | 75 | 80 | 85 | 90 | 95 | 100 | 105 | 110 | 115 | 120 | 125 | 130 | 135 | 140 | 145 | 150 | 155 | 160 | 165 | 170 | 175 | 180 | 185 | 190 | 195 | 200 | 205 | 210 | 215 | 220 | 225 | 230 | 235 | 240 | 245 | 250 | 255 | 260 | 265 | 270 | 275 | 280 | 285 | 290 | 295 | 300 | 305 | 310 | 315 | 320 | 325 | 330 | 335 | 340 | 345 | 350 | 355 | 360 | 365 | 370 | 375 | 380 | 385 | 390 | 395 | 400 | 405 | 410 | 415 | 420 | 425 | 430 | 435 | 440 | 445 | 450 | 455 | 460 | 465 | 470 | 475 | 480 | 485 | 490 | 495 | 500 | 505 | 510 | 515 | 520 | 525 | 530 | 535 | 540 | 545 | 550 | 555 | 560 | 565 | 570 | 575 | 580 | 585 | 590 | 595 | 600 | 605 | 610 | 615 | 620 | 625 | 630 | 635 | 640 | 645 | 650 | 655 | 660 | 665 | 670 | 675 | 680 | 685 | 690 | 695 | 700 | 705 | 710 | 715 | 720 | 725 | 730 | 735 | 740 | 745 | 750 | 755 | 760 | 765 | 770 | 775 | 780 | 785 | 790 | 795 | 800 | 805 | 810 | 815 | 820 | 825 | 830 | 835 | 840 | 845 | 850 | 855 | 860 | 865 | 870 | 875 | 880 | 885 | 890 | 895 | 900 | 905 | 910 | 915 | 920 | 925 | 930 | 935 | 940 | 945 | 950 | 955 | 960 | 965 | 970 | 975 | 980 | 985 | 990 | 995 | 1000 | 1005 | 1010 | 1015 | 1020 | 1025 | 1030 | 1035 | 1040 | 1045 | 1050 | 1055 | 1060 | 1065 | 1070 | 1075 | 1080 | 1085 | 1090 | 1095 | 1100 | 1105 | 1110 | 1115 | 1120 | 1125 | 1130 | 1135 | 1140 | 1145 | 1150 | 1155 | 1160 | 1165 | 1170 | 1175 | 1180 | 1185 | 1190 | 1195 |
| --- | --- | --- | --- | --- | --- | --- | --- | --- | --- | --- | --- | --- | --- | --- | --- | --- | --- | --- | --- | --- | --- | --- | --- | --- | --- | --- | --- | --- | --- | --- | --- | --- | --- | --- | --- | --- | --- | --- | --- | --- | --- | --- | --- | --- | --- | --- | --- | --- | --- | --- | --- | --- | --- | --- | --- | --- | --- | --- | --- | --- | --- | --- | --- | --- | --- | --- | --- | --- | --- | --- | --- | --- | --- | --- | --- | --- | --- | --- | --- | --- | --- | --- | --- | --- | --- | --- | --- | --- | --- | --- | --- | --- | --- | --- | --- | --- | --- | --- | --- | --- | --- | --- | --- | --- | --- | --- | --- | --- | --- | --- | --- | --- | --- | --- | --- | --- | --- | --- | --- | --- | --- | --- | --- | --- | --- | --- | --- | --- | --- | --- | --- | --- | --- | --- | --- | --- | --- | --- | --- | --- | --- | --- | --- | --- | --- | --- | --- | --- | --- | --- | --- | --- | --- | --- | --- | --- | --- | --- | --- | --- | --- | --- | --- | --- | --- | --- | --- | --- | --- | --- | --- | --- | --- | --- | --- | --- | --- | --- | --- | --- | --- | --- | --- | --- | --- | --- | --- | --- | --- | --- | --- | --- | --- | --- | --- | --- | --- | --- | --- | --- | --- | --- | --- | --- | --- | --- | --- | --- | --- | --- | --- | --- | --- | --- | --- | --- | --- | --- | --- | --- | --- | --- | --- | --- | --- | --- | --- | --- | --- | --- | --- | --- | --- | --- | --- | --- | --- | --- | --- | --- | --- | --- | --- | --- | --- | --- | --- | --- | --- | --- | --- | --- | --- | --- | --- | --- | --- | --- | --- | --- | --- | --- | --- | --- | --- | --- | --- | --- | --- | --- | --- | --- | --- | --- | --- | --- | --- | --- | --- | --- |
| left anterior | 0.422998 | 0.444582 | 0.433232 | 0.407507 | 0.318901 | 0.227537 | 0.215032 | 0.214661 | 0.214751 | 0.236364 | 0.271466 | 0.278154 | 0.257105 | 0.334235 | 0.360701 | 0.241891 | 0.237100 | 0.404408 | 0.420845 | 0.273960 | 0.320483 | 0.308141 | 0.214850 | 0.288606 | 0.766759 | 1.451452 | 1.200480 | 1.216732 | 4.842804 | 6.210160 | 3.000275 | 3.053249 | 3.742611 | 1.841108 | 0.590788 | 0.235811 | 0.219221 | 0.220702 | 0.236989 | 0.263537 | 0.310619 | 0.358715 | 0.345139 | 0.234408 | 0.218832 | 0.226248 | 0.267324 | 0.219798 | 0.248640 | 0.525374 | 0.558428 | 0.598986 | 1.211303 | 1.585179 | 1.003124 | 0.566185 | 0.366488 | 0.247099 | 0.214647 | 0.230181 | 0.222991 | 0.215353 | 0.272614 | 0.331086 | 0.394205 | 0.428244 | 0.302848 | 0.224493 | 0.215001 | 0.221493 | 0.217527 | 0.221454 | 0.248559 | 0.342743 | 0.625964 | 1.052958 | 1.204210 | 0.987347 | 0.523515 | 0.411385 | 0.348815 | 0.288230 | 0.256871 | 0.276891 | 0.302032 | 0.271938 | 0.246378 | 0.227023 | 0.219458 | 0.214869 | 0.243745 | 0.296042 | 0.265787 | 0.236926 | 0.216769 | 0.215080 | 0.216509 | 0.218185 | 0.216949 | 0.219054 | 0.227813 | 0.228853 | 0.302772 | 0.289224 | 0.653817 | 1.800569 | 0.845183 | 0.478672 | 0.320686 | 0.231400 | 0.218789 | 0.222993 | 0.270635 | 0.253107 | 0.274060 | 0.280802 | 0.248787 | 0.226688 | 0.214836 | 0.221891 | 0.246748 | 0.332387 | 0.498105 | 0.548338 | 0.497731 | 0.398869 | 0.397722 | 0.356752 | 0.296875 | 0.223412 | 0.214626 | 0.217848 | 0.231900 | 0.279509 | 0.340442 | 0.333689 | 0.243878 | 0.214831 | 0.228627 | 0.277928 | 0.304722 | 0.307454 | 0.302164 | 0.264758 | 0.236458 | 0.220892 | 0.222020 | 0.230308 | 0.242322 | 0.237574 | 0.226545 | 0.218145 | 0.215012 | 0.224047 | 0.237319 | 0.240665 | 0.236628 | 0.232266 | 0.222603 | 0.215358 | 0.216536 | 0.217750 | 0.218702 | 0.219575 | 0.215012 | 0.217005 | 0.229204 | 0.255323 | 0.285154 | 0.323222 | 0.423484 | 0.430448 | 0.395230 | 0.430855 | 0.503412 | 0.589803 | 0.447946 | 0.413732 | 0.439076 | 0.533532 | 0.479771 | 0.427561 | 0.399575 | 0.337331 | 0.303262 | 0.321393 | 0.282995 | 0.267229 | 0.253224 | 0.236826 | 0.269684 | 0.257771 | 0.220920 | 0.217627 | 0.214789 | 0.221712 | 0.236669 | 0.326063 | 0.337702 | 0.327689 | 0.305918 | 0.258241 | 0.228671 | 0.214846 | 0.227131 | 0.224344 | 0.218293 | 0.214786 | 0.218639 | 0.216621 | 0.216929 | 0.225418 | 0.224195 | 0.231594 | 0.255796 | 0.342931 | 0.728538 | 1.680162 | 2.164785 | 2.605085 | 2.559013 | 3.298612 | 2.348790 | 1.542880 | 0.721546 | 0.596927 | 0.475115 | 0.412313 | 0.274460 | 0.225518 | 0.216356 | 0.219758 | 0.223263 | 0.236703 | 0.239626 | 0.261354 | 0.259426 | 0.232310 | 0.223786 | 0.227028 | 0.240465 | 0.252101 | 0.275542 | 0.352449 | 0.613895 | 0.840172 | 0.817272 | 0.655532 | 0.454841 | 0.335904 | 0.279382 | 0.242794 | 0.229493 | 0.225686 | 0.220625 | 0.215608 | 0.221306 | 0.232172 | 0.235984 | 0.231001 | 0.237334 | 0.253299 | 0.255039 | 0.233407 | 0.234015 | 0.233107 | 0.246619 | 0.273893 | 0.318533 | 0.337850 | 0.336322 | 0.322582 | 0.329810 | 0.357624 | 0.331963 | 0.326063 | 0.352942 | 0.344036 | 0.347032 | 0.376621 |
| right anterior | 0.286417 | 0.244913 | 0.215157 | 0.284105 | 0.410067 | 0.462772 | 0.560705 | 1.269369 | 1.387920 | 1.287445 | 0.481386 | 0.257383 | 0.230324 | 0.245191 | 0.224129 | 0.620146 | 7.028634 | 9.366153 | 3.434247 | 0.528004 | 0.409691 | 0.588331 | 0.296362 | 0.290236 | 0.301907 | 0.262135 | 0.285004 | 0.378858 | 0.328295 | 0.408527 | 0.269609 | 0.249468 | 0.245856 | 0.214988 | 0.228551 | 0.235514 | 0.289024 | 0.289312 | 0.384624 | 0.541050 | 1.456871 | 4.030501 | 2.930991 | 3.305616 | 3.016669 | 1.689903 | 0.884436 | 0.576835 | 0.642057 | 0.893696 | 1.097392 | 1.281952 | 2.997451 | 6.150954 | 4.500020 | 2.841273 | 6.267908 | 15.304428 | 25.487044 | 17.786964 | 24.698966 | 25.193244 | 14.662399 | 5.243712 | 1.183465 | 0.936352 | 0.761407 | 0.470116 | 0.500936 | 0.674463 | 0.597197 | 0.587287 | 0.416621 | 0.425663 | 0.501026 | 0.650228 | 0.769040 | 0.794251 | 0.848723 | 0.772565 | 0.433364 | 0.303704 | 0.253767 | 0.227860 | 0.218096 | 0.214630 | 0.214630 | 0.218651 | 0.224887 | 0.248506 | 0.254835 | 0.301980 | 0.349918 | 0.420040 | 0.463446 | 0.406834 | 0.324714 | 0.278101 | 0.269019 | 0.263679 | 0.266477 | 0.283047 | 0.280775 | 0.274298 | 0.274252 | 0.246071 | 0.243522 | 0.234353 | 0.235353 | 0.251530 | 0.261219 | 0.266409 | 0.265594 | 0.227746 | 0.216902 | 0.214701 | 0.216189 | 0.215427 | 0.214667 | 0.215212 | 0.225967 | 0.230151 | 0.219595 | 0.220087 | 0.219063 | 0.215682 | 0.215783 | 0.219797 | 0.233457 | 0.262146 | 0.373163 | 0.416912 | 0.341671 | 0.294888 | 0.262137 | 0.247793 | 0.233393 | 0.216436 | 0.217931 | 0.226968 | 0.277322 | 0.471101 | 0.573874 | 0.668263 | 0.991216 | 0.909571 | 0.867261 | 0.618814 | 0.374761 | 0.303170 | 0.285194 | 0.252656 | 0.225040 | 0.214768 | 0.217095 | 0.227576 | 0.251526 | 0.289950 | 0.268673 | 0.229921 | 0.217576 | 0.224112 | 0.220835 | 0.216516 | 0.224221 | 0.230049 | 0.248447 | 0.311140 | 0.433966 | 0.465764 | 0.435309 | 0.495129 | 0.457921 | 0.490508 | 0.395731 | 0.406149 | 0.497139 | 0.497004 | 0.387511 | 0.388628 | 0.310246 | 0.310174 | 0.257985 | 0.225250 | 0.220817 | 0.226566 | 0.226476 | 0.214720 | 0.231350 | 0.291052 | 0.410792 | 0.365025 | 0.281377 | 0.267218 | 0.229877 | 0.216134 | 0.216081 | 0.221063 | 0.215214 | 0.216621 | 0.223159 | 0.223275 | 0.227654 | 0.222919 | 0.215749 | 0.215216 | 0.218751 | 0.216605 | 0.215977 | 0.219032 | 0.218352 | 0.215515 | 0.214899 | 0.214893 | 0.214651 | 0.215640 | 0.215193 | 0.216800 | 0.221207 | 0.237423 | 0.253977 | 0.264324 | 0.251262 | 0.225255 | 0.214822 | 0.218074 | 0.225166 | 0.221679 | 0.215154 | 0.216012 | 0.224398 | 0.236520 | 0.251563 | 0.276426 | 0.274369 | 0.235518 | 0.216582 | 0.214631 | 0.214985 | 0.228445 | 0.239179 | 0.244929 | 0.256884 | 0.282596 | 0.323094 | 0.283801 | 0.245331 | 0.224823 | 0.215742 | 0.214728 | 0.222048 | 0.242490 | 0.233905 | 0.223187 | 0.218959 | 0.216077 | 0.217809 | 0.220453 | 0.218425 | 0.223633 | 0.239364 | 0.235567 | 0.236840 | 0.235817 | 0.226340 | 0.248196 | 0.236695 | 0.249665 | 0.293599 | 0.346697 | 0.375375 | 0.452828 | 0.496534 | 0.595757 | 0.499423 | 0.361547 | 0.327635 | 0.309847 | 0.274258 | 0.229509 |
| left central | 0.236997 | 0.220007 | 0.215149 | 0.216296 | 0.221265 | 0.214636 | 0.214666 | 0.219874 | 0.221105 | 0.235869 | 0.266690 | 0.300413 | 0.257928 | 0.293124 | 0.347233 | 0.303662 | 0.225396 | 0.224853 | 0.214844 | 0.214731 | 0.217149 | 0.216289 | 0.216013 | 0.261542 | 0.254027 | 0.363081 | 0.475028 | 0.508745 | 0.985940 | 3.440944 | 6.453037 | 7.949199 | 6.658691 | 16.448186 | 10.800002 | 2.326397 | 0.794490 | 0.415943 | 0.394568 | 0.371878 | 0.366473 | 0.340001 | 0.335307 | 0.271912 | 0.286897 | 0.324607 | 0.331555 | 0.271838 | 0.289325 | 0.368320 | 0.469491 | 0.421624 | 0.429100 | 0.411773 | 0.397260 | 0.254464 | 0.214656 | 0.241282 | 0.236786 | 0.224478 | 0.214865 | 0.221448 | 0.264614 | 0.301241 | 0.417066 | 0.845633 | 1.412773 | 2.637317 | 4.794305 | 7.458753 | 7.650161 | 6.339901 | 1.941171 | 0.796054 | 0.362509 | 0.229321 | 0.223866 | 0.347377 | 0.524296 | 0.659609 | 0.824982 | 0.777789 | 0.450585 | 0.332908 | 0.233482 | 0.215648 | 0.215986 | 0.214626 | 0.226104 | 0.301592 | 0.517805 | 0.655590 | 0.815277 | 0.834307 | 0.570961 | 0.418182 | 0.255905 | 0.217125 | 0.217763 | 0.215692 | 0.217779 | 0.219411 | 0.230460 | 0.251900 | 0.275871 | 0.417472 | 0.946328 | 1.430819 | 1.155870 | 1.498399 | 0.872463 | 0.525433 | 0.448591 | 0.356966 | 0.270597 | 0.253120 | 0.268113 | 0.281654 | 0.287176 | 0.244731 | 0.219385 | 0.216212 | 0.217161 | 0.217346 | 0.214998 | 0.217366 | 0.247961 | 0.334879 | 0.450968 | 0.434797 | 0.383758 | 0.423934 | 0.396847 | 0.372010 | 0.384439 | 0.350453 | 0.542853 | 1.052503 | 2.260627 | 3.497615 | 2.869374 | 2.285298 | 4.278756 | 2.763245 | 1.529682 | 0.778629 | 0.591083 | 0.533575 | 0.459997 | 0.362641 | 0.321459 | 0.328736 | 0.385323 | 0.388365 | 0.554528 | 0.504506 | 0.382546 | 0.333332 | 0.284349 | 0.222716 | 0.215264 | 0.220898 | 0.224217 | 0.233195 | 0.281313 | 0.337961 | 0.401618 | 0.538403 | 0.625321 | 0.544901 | 0.405875 | 0.307115 | 0.256865 | 0.230684 | 0.214748 | 0.229991 | 0.230480 | 0.227356 | 0.227571 | 0.245390 | 0.298894 | 0.281382 | 0.259158 | 0.254105 | 0.246837 | 0.261154 | 0.238987 | 0.242127 | 0.259364 | 0.252646 | 0.245218 | 0.248221 | 0.243618 | 0.251154 | 0.236767 | 0.222046 | 0.215753 | 0.214658 | 0.215411 | 0.215564 | 0.219990 | 0.231976 | 0.236738 | 0.240484 | 0.251343 | 0.263855 | 0.254328 | 0.234271 | 0.226481 | 0.220886 | 0.217416 | 0.215595 | 0.214642 | 0.214630 | 0.215331 | 0.218015 | 0.215038 | 0.214854 | 0.215003 | 0.215559 | 0.223666 | 0.239683 | 0.259095 | 0.240908 | 0.227443 | 0.226054 | 0.230882 | 0.226849 | 0.230821 | 0.223726 | 0.236256 | 0.249121 | 0.248089 | 0.248948 | 0.257870 | 0.240085 | 0.242155 | 0.231923 | 0.234394 | 0.230956 | 0.242579 | 0.263260 | 0.288922 | 0.277865 | 0.267845 | 0.246339 | 0.294448 | 0.325888 | 0.353940 | 0.401433 | 0.440506 | 0.454895 | 0.489698 | 0.376307 | 0.319118 | 0.272547 | 0.256296 | 0.232809 | 0.221540 | 0.216566 | 0.215900 | 0.217420 | 0.219244 | 0.215399 | 0.214752 | 0.215377 | 0.214773 | 0.214653 | 0.214629 | 0.217387 | 0.253315 | 0.351880 | 0.608077 | 0.878084 | 0.919610 | 0.764046 | 0.564473 | 0.480824 | 0.334577 | 0.240720 |
| right central | 0.228472 | 0.282289 | 0.394356 | 0.469222 | 0.891350 | 1.266425 | 2.218078 | 1.739217 | 0.723973 | 0.466524 | 0.414742 | 0.244513 | 0.219340 | 0.214733 | 0.228037 | 0.238354 | 0.386720 | 0.638188 | 0.530708 | 0.527639 | 0.710794 | 0.999946 | 1.478408 | 0.589937 | 0.380837 | 0.409867 | 0.979736 | 4.161979 | 3.938534 | 4.639405 | 3.720763 | 6.342321 | 9.057403 | 4.646351 | 1.246500 | 0.702960 | 0.486803 | 0.419985 | 0.317584 | 0.244675 | 0.215158 | 0.227158 | 0.302608 | 0.450903 | 0.711026 | 0.911135 | 0.978149 | 0.936406 | 0.512559 | 0.315111 | 0.262848 | 0.258016 | 0.235623 | 0.231687 | 0.227776 | 0.228386 | 0.223699 | 0.224569 | 0.216826 | 0.236340 | 0.265372 | 0.315368 | 0.297888 | 0.331190 | 0.364361 | 0.408436 | 0.463333 | 0.589172 | 0.422653 | 0.336781 | 0.287032 | 0.285755 | 0.263557 | 0.254403 | 0.222762 | 0.215305 | 0.258676 | 0.314475 | 0.430896 | 0.567045 | 0.722092 | 0.899039 | 0.602857 | 0.276457 | 0.225546 | 0.214630 | 0.226538 | 0.243090 | 0.303431 | 0.280951 | 0.242162 | 0.254474 | 0.268866 | 0.278759 | 0.310641 | 0.324303 | 0.581004 | 0.751273 | 0.697908 | 0.929033 | 0.946961 | 0.716094 | 0.684887 | 0.562242 | 0.719449 | 0.681802 | 0.406328 | 0.343580 | 0.275501 | 0.236601 | 0.216876 | 0.219531 | 0.219294 | 0.220197 | 0.281808 | 0.503263 | 0.725326 | 1.142166 | 1.380851 | 1.347205 | 0.895805 | 0.554603 | 0.418502 | 0.495193 | 0.520452 | 0.512860 | 0.545236 | 0.894469 | 1.161482 | 1.113649 | 1.104260 | 1.650360 | 2.685119 | 2.817899 | 1.529003 | 1.028882 | 1.299500 | 1.215721 | 1.032511 | 0.885532 | 0.867273 | 1.520451 | 3.167369 | 5.539849 | 8.401523 | 7.614808 | 9.072152 | 5.765556 | 1.993876 | 0.969304 | 0.478121 | 0.315799 | 0.276291 | 0.271819 | 0.333326 | 0.308605 | 0.264188 | 0.226789 | 0.228020 | 0.225465 | 0.222423 | 0.219222 | 0.238724 | 0.305788 | 0.478636 | 0.439307 | 0.435549 | 0.440180 | 0.512447 | 0.660639 | 0.763095 | 0.790711 | 0.767361 | 0.601504 | 0.427958 | 0.290782 | 0.253037 | 0.252299 | 0.269873 | 0.293055 | 0.305855 | 0.342636 | 0.468079 | 0.566282 | 0.661122 | 0.990484 | 1.606211 | 2.768953 | 2.521810 | 1.946884 | 1.140091 | 0.788174 | 0.672065 | 0.766910 | 0.955923 | 1.230358 | 1.147958 | 1.219404 | 1.047770 | 1.097373 | 0.923078 | 0.856106 | 0.704433 | 0.668347 | 0.727805 | 0.921605 | 0.866784 | 0.774704 | 0.608810 | 0.580091 | 0.649501 | 0.820675 | 1.095736 | 1.542378 | 1.746393 | 1.670404 | 2.134019 | 2.152355 | 1.818409 | 0.998698 | 0.721245 | 0.493196 | 0.434142 | 0.350393 | 0.323170 | 0.331298 | 0.469949 | 0.582067 | 0.974218 | 0.931275 | 0.865557 | 0.945854 | 0.903799 | 0.640282 | 0.507352 | 0.368819 | 0.314069 | 0.275039 | 0.243598 | 0.239771 | 0.268465 | 0.336258 | 0.407290 | 0.454908 | 0.426458 | 0.350203 | 0.277295 | 0.228562 | 0.215820 | 0.257201 | 0.348260 | 0.366207 | 0.440057 | 0.503619 | 0.540893 | 0.513284 | 0.349772 | 0.269568 | 0.245686 | 0.220802 | 0.216493 | 0.222701 | 0.231381 | 0.233233 | 0.227414 | 0.218880 | 0.215605 | 0.215043 | 0.216150 | 0.217957 | 0.227998 | 0.235845 | 0.220304 | 0.215961 | 0.221591 | 0.243408 | 0.263532 | 0.254525 | 0.245285 | 0.298194 |
| left posterior | 0.288093 | 0.452915 | 0.404174 | 0.253103 | 0.216008 | 0.217731 | 0.235770 | 0.332181 | 0.434823 | 0.582350 | 0.639689 | 0.404711 | 0.439975 | 0.363219 | 0.332698 | 0.315430 | 0.222610 | 0.220848 | 0.217129 | 0.247264 | 0.294388 | 0.386485 | 0.533031 | 0.519941 | 0.638527 | 0.999853 | 0.872061 | 0.623175 | 0.488375 | 0.398248 | 0.348467 | 0.305382 | 0.219142 | 0.247869 | 0.326225 | 0.375897 | 0.332357 | 0.271740 | 0.255875 | 0.230398 | 0.215662 | 0.221266 | 0.222070 | 0.221591 | 0.237793 | 0.296751 | 0.302114 | 0.273021 | 0.224517 | 0.215152 | 0.219168 | 0.236072 | 0.281668 | 0.309867 | 0.333535 | 0.387867 | 0.469557 | 0.584962 | 0.807194 | 1.071353 | 1.568475 | 1.474292 | 0.851540 | 0.341648 | 0.221069 | 0.244579 | 0.391154 | 0.832802 | 0.949833 | 0.758522 | 0.559781 | 0.447116 | 0.388982 | 0.316392 | 0.251040 | 0.239497 | 0.255454 | 0.274964 | 0.261345 | 0.227075 | 0.216189 | 0.216015 | 0.227461 | 0.264100 | 0.405456 | 0.966241 | 1.342813 | 1.710046 | 1.695143 | 1.658150 | 1.272034 | 0.890096 | 0.635019 | 0.616694 | 0.594829 | 0.741909 | 0.989986 | 1.216189 | 1.119919 | 0.910001 | 0.978991 | 0.983580 | 0.669466 | 0.459687 | 0.494739 | 0.558031 | 0.716965 | 0.708250 | 0.775998 | 0.785136 | 0.790836 | 0.748702 | 0.752979 | 0.637624 | 0.570404 | 0.495489 | 0.524440 | 0.542426 | 0.567207 | 0.567691 | 0.620918 | 0.669883 | 0.795976 | 0.813284 | 0.895062 | 1.201340 | 1.120447 | 0.601034 | 0.309819 | 0.243429 | 0.248353 | 0.257066 | 0.240458 | 0.246599 | 0.320725 | 0.617904 | 1.185592 | 1.729468 | 2.000603 | 2.259589 | 1.267648 | 0.874054 | 1.026911 | 0.835890 | 0.537230 | 0.569388 | 0.523008 | 0.543357 | 0.436704 | 0.296457 | 0.261425 | 0.250060 | 0.223211 | 0.220046 | 0.233079 | 0.252802 | 0.240715 | 0.219570 | 0.215646 | 0.214725 | 0.221573 | 0.254684 | 0.269068 | 0.261034 | 0.235801 | 0.219113 | 0.215641 | 0.214654 | 0.216725 | 0.218449 | 0.216508 | 0.217823 | 0.246509 | 0.284641 | 0.338570 | 0.520586 | 0.662648 | 0.667439 | 0.519387 | 0.453258 | 0.375915 | 0.291034 | 0.219629 | 0.231573 | 0.296238 | 0.377558 | 0.531364 | 0.628587 | 0.592399 | 0.529789 | 0.374307 | 0.329343 | 0.284820 | 0.278968 | 0.249556 | 0.228075 | 0.229663 | 0.239131 | 0.235534 | 0.231381 | 0.231414 | 0.241973 | 0.273051 | 0.281838 | 0.291070 | 0.300239 | 0.311994 | 0.267601 | 0.259833 | 0.249153 | 0.232858 | 0.221035 | 0.214675 | 0.217928 | 0.229297 | 0.244429 | 0.257741 | 0.256273 | 0.241781 | 0.218663 | 0.214630 | 0.230105 | 0.265977 | 0.294998 | 0.297486 | 0.285676 | 0.238876 | 0.215870 | 0.232994 | 0.295843 | 0.343498 | 0.396003 | 0.473664 | 0.535069 | 0.419019 | 0.301706 | 0.225579 | 0.215257 | 0.223043 | 0.248855 | 0.296557 | 0.280253 | 0.261197 | 0.240943 | 0.228647 | 0.215095 | 0.218999 | 0.238564 | 0.260768 | 0.326520 | 0.567793 | 0.922330 | 0.992456 | 0.896152 | 0.733515 | 0.586859 | 0.522970 | 0.347856 | 0.279889 | 0.246866 | 0.231632 | 0.240000 | 0.246437 | 0.251955 | 0.298010 | 0.343555 | 0.411672 | 0.479388 | 0.416789 | 0.430389 | 0.470731 | 0.540823 | 0.699863 | 0.983715 | 1.149385 | 1.573155 | 1.876256 | 2.092517 | 1.651784 | 1.123953 |
| right posterior | 0.713010 | 0.997474 | 1.265411 | 2.673123 | 1.315559 | 0.569479 | 0.830250 | 1.558526 | 1.582053 | 2.290642 | 1.574023 | 3.585868 | 12.483271 | 12.540128 | 9.326457 | 11.343265 | 26.155060 | 45.824001 | 7.803504 | 1.740583 | 0.573049 | 0.234994 | 0.215742 | 0.222264 | 0.231668 | 0.222351 | 0.220817 | 0.310587 | 0.458500 | 0.368541 | 0.285478 | 0.272063 | 0.428156 | 0.488705 | 0.379962 | 0.371297 | 0.337663 | 0.381021 | 0.402661 | 0.344254 | 0.325123 | 0.266170 | 0.225493 | 0.215755 | 0.216554 | 0.216834 | 0.215773 | 0.226491 | 0.224070 | 0.215413 | 0.214732 | 0.218172 | 0.228173 | 0.248893 | 0.341528 | 0.454446 | 0.777341 | 1.136924 | 1.106808 | 1.158002 | 1.059752 | 0.715705 | 0.677996 | 0.445158 | 0.263180 | 0.228170 | 0.214912 | 0.215223 | 0.218508 | 0.217035 | 0.221001 | 0.264609 | 0.306736 | 0.354871 | 0.260223 | 0.226107 | 0.216469 | 0.259807 | 0.261842 | 0.296301 | 0.331513 | 0.262448 | 0.229890 | 0.214722 | 0.225707 | 0.240523 | 0.254911 | 0.250937 | 0.255499 | 0.258984 | 0.253493 | 0.257898 | 0.260377 | 0.279796 | 0.351144 | 0.405004 | 0.524900 | 0.619293 | 0.507283 | 0.371263 | 0.338280 | 0.325655 | 0.380860 | 0.497118 | 0.735354 | 1.448155 | 3.956724 | 6.797039 | 10.564839 | 8.340997 | 4.525979 | 2.849386 | 1.969137 | 1.643363 | 1.834818 | 2.154873 | 2.482586 | 2.600905 | 2.216841 | 1.848706 | 1.440805 | 1.007629 | 0.846311 | 0.743117 | 0.517731 | 0.406656 | 0.328503 | 0.275817 | 0.267328 | 0.248796 | 0.222916 | 0.224799 | 0.230095 | 0.244585 | 0.272107 | 0.294646 | 0.345794 | 0.517198 | 0.868834 | 1.212026 | 1.592943 | 2.607701 | 4.336555 | 6.389965 | 5.958939 | 6.759609 | 6.870219 | 6.072339 | 2.897995 | 1.115040 | 0.516871 | 0.389512 | 0.313300 | 0.273023 | 0.252724 | 0.250328 | 0.262091 | 0.270581 | 0.272807 | 0.255956 | 0.243737 | 0.241079 | 0.242590 | 0.233745 | 0.237493 | 0.235696 | 0.223990 | 0.215210 | 0.223163 | 0.279710 | 0.424112 | 0.584366 | 0.762737 | 0.791779 | 0.659578 | 0.455282 | 0.330654 | 0.246511 | 0.221184 | 0.215300 | 0.232150 | 0.300045 | 0.483985 | 0.829053 | 1.243141 | 2.297703 | 3.174006 | 3.688688 | 2.595027 | 0.959675 | 0.430369 | 0.330934 | 0.251034 | 0.225976 | 0.218211 | 0.221894 | 0.229718 | 0.249081 | 0.255640 | 0.259246 | 0.242282 | 0.239489 | 0.223714 | 0.226071 | 0.221097 | 0.217351 | 0.216898 | 0.219295 | 0.215298 | 0.214707 | 0.227122 | 0.245949 | 0.262250 | 0.258764 | 0.271808 | 0.272507 | 0.279905 | 0.294230 | 0.280055 | 0.229170 | 0.215648 | 0.217900 | 0.217803 | 0.228777 | 0.254232 | 0.278604 | 0.268776 | 0.247847 | 0.234496 | 0.264848 | 0.290620 | 0.378553 | 0.407863 | 0.400890 | 0.383012 | 0.461172 | 0.493012 | 0.596534 | 0.769709 | 1.056652 | 1.353680 | 1.982218 | 1.703935 | 1.253277 | 0.815527 | 0.551468 | 0.387019 | 0.268733 | 0.219933 | 0.219489 | 0.252696 | 0.264124 | 0.248679 | 0.221599 | 0.223452 | 0.355420 | 0.744916 | 1.736323 | 2.270695 | 2.200876 | 1.051448 | 0.408472 | 0.266864 | 0.246144 | 0.264812 | 0.297525 | 0.330179 | 0.492661 | 0.870048 | 1.417052 | 1.745891 | 0.910919 | 0.477485 | 0.286994 | 0.224954 | 0.215972 | 0.241316 | 0.276267 | 0.289414 | 0.299347 |
| all electrodes | 0.214649 | 0.252537 | 0.253414 | 0.233588 | 0.220685 | 0.228610 | 0.230303 | 0.234611 | 0.293756 | 0.384735 | 0.363200 | 0.349155 | 0.510792 | 1.102650 | 0.353384 | 0.609352 | 0.509208 | 0.476418 | 0.533034 | 0.251519 | 0.219267 | 0.235864 | 0.296762 | 0.624614 | 1.247528 | 3.202481 | 2.742675 | 2.931869 | 2.528953 | 1.417505 | 0.652309 | 0.389464 | 0.263589 | 0.215208 | 0.214878 | 0.221132 | 0.251582 | 0.275044 | 0.338452 | 0.370634 | 0.408988 | 0.808607 | 0.937730 | 0.793604 | 1.177235 | 1.619740 | 2.511725 | 1.755621 | 0.676538 | 0.451428 | 0.376019 | 0.329578 | 0.241833 | 0.215129 | 0.214642 | 0.214630 | 0.217724 | 0.224139 | 0.288493 | 0.310128 | 0.331333 | 0.317813 | 0.245302 | 0.216829 | 0.217815 | 0.302286 | 0.476592 | 0.606595 | 0.685801 | 0.458540 | 0.346749 | 0.249652 | 0.216153 | 0.234373 | 0.232160 | 0.214679 | 0.218252 | 0.246893 | 0.253393 | 0.281041 | 0.335113 | 0.305018 | 0.220577 | 0.216168 | 0.255194 | 0.286541 | 0.280932 | 0.317623 | 0.295136 | 0.241528 | 0.219569 | 0.214656 | 0.226806 | 0.223311 | 0.215946 | 0.214821 | 0.214885 | 0.220010 | 0.231991 | 0.250450 | 0.290494 | 0.312029 | 0.264946 | 0.230745 | 0.215749 | 0.250449 | 0.328145 | 0.448856 | 0.525530 | 0.464894 | 0.379106 | 0.288284 | 0.251478 | 0.237264 | 0.227912 | 0.214632 | 0.214964 | 0.215130 | 0.226112 | 0.215947 | 0.214696 | 0.214694 | 0.222357 | 0.226780 | 0.219092 | 0.223304 | 0.276778 | 0.334476 | 0.367151 | 0.319625 | 0.335121 | 0.399180 | 0.384688 | 0.300682 | 0.252686 | 0.231798 | 0.227009 | 0.217771 | 0.239168 | 0.251415 | 0.292731 | 0.590878 | 1.327402 | 3.188261 | 6.474344 | 4.366170 | 3.450940 | 3.036257 | 1.126794 | 0.605669 | 0.386148 | 0.346890 | 0.319955 | 0.284041 | 0.255260 | 0.227118 | 0.215414 | 0.237383 | 0.280161 | 0.315609 | 0.300662 | 0.324521 | 0.288841 | 0.233963 | 0.220183 | 0.258638 | 0.262332 | 0.245853 | 0.255797 | 0.275181 | 0.286259 | 0.234897 | 0.214630 | 0.214630 | 0.214644 | 0.214994 | 0.214645 | 0.214800 | 0.215729 | 0.238375 | 0.288906 | 0.317369 | 0.350938 | 0.363244 | 0.413030 | 0.794529 | 1.095028 | 1.042102 | 1.256109 | 1.115777 | 0.757079 | 0.570920 | 0.468586 | 0.413034 | 0.396295 | 0.380197 | 0.452605 | 0.671347 | 0.718154 | 0.643669 | 0.576087 | 0.573858 | 0.661574 | 0.728702 | 0.719012 | 0.847583 | 0.692029 | 0.754944 | 0.582932 | 0.395459 | 0.308811 | 0.246363 | 0.234455 | 0.258648 | 0.270034 | 0.304470 | 0.304952 | 0.270992 | 0.296841 | 0.293776 | 0.251769 | 0.238331 | 0.233781 | 0.239848 | 0.296233 | 0.382646 | 0.471051 | 0.576570 | 0.661397 | 0.565772 | 0.471432 | 0.383865 | 0.297984 | 0.264829 | 0.245479 | 0.241037 | 0.253576 | 0.251911 | 0.239404 | 0.250516 | 0.248244 | 0.269320 | 0.335460 | 0.362000 | 0.478832 | 0.508688 | 0.445644 | 0.487555 | 0.600557 | 0.456196 | 0.373418 | 0.262631 | 0.245721 | 0.247127 | 0.233896 | 0.227002 | 0.231430 | 0.245641 | 0.280023 | 0.318533 | 0.322266 | 0.381065 | 0.429249 | 0.432397 | 0.426999 | 0.430055 | 0.433110 | 0.505833 | 0.458802 | 0.384785 | 0.372727 | 0.344699 | 0.307736 | 0.276015 | 0.238766 | 0.218350 | 0.214626 | 0.217612 | 0.231502 | 0.249636 |

Searchlight, spatiotemporal cluster permutation test

No significant clusters observed.

K) difference, sex, angry vs sad

  
|  | time window | peak latency | cluster *p* | peak Cohen's *d* |  | | | |  |
| **all electrodes** | 815 - 895 ms | 880 ms | 0.038 | 0.8669 |  | | | |  |
|  | | | | | | | | |  |

Time-resolved classification, cluster permutation tests

|  | **left hemisphere** | | | | **right hemisphere** | | | |  |
|  | time window | peak latency | cluster *p* | peak Cohen's *d* | time window | peak latency | cluster *p* | peak Cohen's *d* |  |
| **anterior** | 670 - 765 ms | 725 ms | 0.036 | 0.7059 | 5 - 110 ms | 65 ms | 0.0453 | -0.7235 |  |
| **central** | 470 - 585 ms | 485 ms | 0.0193 | 0.8108 | 445 - 795 ms | 510 ms | 0.0033 | 0.873 |  |
 690 - 755 ms | 725 ms | 0.0426 | 1.074 |  | | | ||  | **posterior** |  | | | | 225 - 620 ms | 490 ms | 0.0003 | 1.1628 |

  

Time-resolved classification, Bayesian statistics

|  | -200 | -195 | -190 | -185 | -180 | -175 | -170 | -165 | -160 | -155 | -150 | -145 | -140 | -135 | -130 | -125 | -120 | -115 | -110 | -105 | -100 | -95 | -90 | -85 | -80 | -75 | -70 | -65 | -60 | -55 | -50 | -45 | -40 | -35 | -30 | -25 | -20 | -15 | -10 | -5 | 0 | 5 | 10 | 15 | 20 | 25 | 30 | 35 | 40 | 45 | 50 | 55 | 60 | 65 | 70 | 75 | 80 | 85 | 90 | 95 | 100 | 105 | 110 | 115 | 120 | 125 | 130 | 135 | 140 | 145 | 150 | 155 | 160 | 165 | 170 | 175 | 180 | 185 | 190 | 195 | 200 | 205 | 210 | 215 | 220 | 225 | 230 | 235 | 240 | 245 | 250 | 255 | 260 | 265 | 270 | 275 | 280 | 285 | 290 | 295 | 300 | 305 | 310 | 315 | 320 | 325 | 330 | 335 | 340 | 345 | 350 | 355 | 360 | 365 | 370 | 375 | 380 | 385 | 390 | 395 | 400 | 405 | 410 | 415 | 420 | 425 | 430 | 435 | 440 | 445 | 450 | 455 | 460 | 465 | 470 | 475 | 480 | 485 | 490 | 495 | 500 | 505 | 510 | 515 | 520 | 525 | 530 | 535 | 540 | 545 | 550 | 555 | 560 | 565 | 570 | 575 | 580 | 585 | 590 | 595 | 600 | 605 | 610 | 615 | 620 | 625 | 630 | 635 | 640 | 645 | 650 | 655 | 660 | 665 | 670 | 675 | 680 | 685 | 690 | 695 | 700 | 705 | 710 | 715 | 720 | 725 | 730 | 735 | 740 | 745 | 750 | 755 | 760 | 765 | 770 | 775 | 780 | 785 | 790 | 795 | 800 | 805 | 810 | 815 | 820 | 825 | 830 | 835 | 840 | 845 | 850 | 855 | 860 | 865 | 870 | 875 | 880 | 885 | 890 | 895 | 900 | 905 | 910 | 915 | 920 | 925 | 930 | 935 | 940 | 945 | 950 | 955 | 960 | 965 | 970 | 975 | 980 | 985 | 990 | 995 | 1000 | 1005 | 1010 | 1015 | 1020 | 1025 | 1030 | 1035 | 1040 | 1045 | 1050 | 1055 | 1060 | 1065 | 1070 | 1075 | 1080 | 1085 | 1090 | 1095 | 1100 | 1105 | 1110 | 1115 | 1120 | 1125 | 1130 | 1135 | 1140 | 1145 | 1150 | 1155 | 1160 | 1165 | 1170 | 1175 | 1180 | 1185 | 1190 | 1195 |
| --- | --- | --- | --- | --- | --- | --- | --- | --- | --- | --- | --- | --- | --- | --- | --- | --- | --- | --- | --- | --- | --- | --- | --- | --- | --- | --- | --- | --- | --- | --- | --- | --- | --- | --- | --- | --- | --- | --- | --- | --- | --- | --- | --- | --- | --- | --- | --- | --- | --- | --- | --- | --- | --- | --- | --- | --- | --- | --- | --- | --- | --- | --- | --- | --- | --- | --- | --- | --- | --- | --- | --- | --- | --- | --- | --- | --- | --- | --- | --- | --- | --- | --- | --- | --- | --- | --- | --- | --- | --- | --- | --- | --- | --- | --- | --- | --- | --- | --- | --- | --- | --- | --- | --- | --- | --- | --- | --- | --- | --- | --- | --- | --- | --- | --- | --- | --- | --- | --- | --- | --- | --- | --- | --- | --- | --- | --- | --- | --- | --- | --- | --- | --- | --- | --- | --- | --- | --- | --- | --- | --- | --- | --- | --- | --- | --- | --- | --- | --- | --- | --- | --- | --- | --- | --- | --- | --- | --- | --- | --- | --- | --- | --- | --- | --- | --- | --- | --- | --- | --- | --- | --- | --- | --- | --- | --- | --- | --- | --- | --- | --- | --- | --- | --- | --- | --- | --- | --- | --- | --- | --- | --- | --- | --- | --- | --- | --- | --- | --- | --- | --- | --- | --- | --- | --- | --- | --- | --- | --- | --- | --- | --- | --- | --- | --- | --- | --- | --- | --- | --- | --- | --- | --- | --- | --- | --- | --- | --- | --- | --- | --- | --- | --- | --- | --- | --- | --- | --- | --- | --- | --- | --- | --- | --- | --- | --- | --- | --- | --- | --- | --- | --- | --- | --- | --- | --- | --- | --- | --- | --- | --- | --- | --- | --- | --- | --- | --- | --- | --- | --- | --- | --- | --- | --- | --- | --- | --- | --- | --- | --- | --- |
| left anterior | 0.221893 | 0.282195 | 0.352709 | 0.364978 | 0.371773 | 0.384904 | 0.304530 | 0.245565 | 0.216693 | 0.214626 | 0.214634 | 0.236136 | 0.290358 | 0.428404 | 0.487336 | 0.410524 | 0.333472 | 0.283874 | 0.362306 | 0.634467 | 0.543224 | 0.327905 | 0.252061 | 0.219042 | 0.217163 | 0.243073 | 0.335761 | 0.396301 | 0.277018 | 0.217077 | 0.219216 | 0.316461 | 0.483253 | 0.722549 | 0.744848 | 0.628744 | 0.465544 | 0.558732 | 0.993000 | 0.567378 | 0.313731 | 0.222490 | 0.217594 | 0.220503 | 0.253617 | 0.223871 | 0.216517 | 0.218867 | 0.254734 | 0.362306 | 0.372600 | 0.330270 | 0.460910 | 0.569319 | 0.770088 | 0.930971 | 1.226883 | 1.011417 | 0.717425 | 0.531517 | 0.482716 | 0.432784 | 0.476079 | 0.360176 | 0.317312 | 0.260859 | 0.222797 | 0.215471 | 0.215176 | 0.220585 | 0.218061 | 0.214952 | 0.223740 | 0.255161 | 0.310396 | 0.368495 | 0.369990 | 0.391607 | 0.446361 | 0.555736 | 0.830030 | 1.430042 | 1.875678 | 3.140101 | 4.924827 | 8.068323 | 9.889126 | 10.287848 | 6.649747 | 5.830957 | 2.878306 | 0.826875 | 0.384873 | 0.256935 | 0.240065 | 0.240939 | 0.250603 | 0.230230 | 0.245109 | 0.289921 | 0.359860 | 0.290818 | 0.285690 | 0.259111 | 0.338252 | 0.413220 | 0.361271 | 0.272703 | 0.238694 | 0.219613 | 0.214696 | 0.250675 | 0.365844 | 0.480572 | 0.482525 | 0.513881 | 0.407098 | 0.295161 | 0.215822 | 0.257281 | 0.484201 | 0.847332 | 1.413278 | 1.601695 | 2.051519 | 1.656127 | 1.335832 | 0.817251 | 0.544714 | 0.337765 | 0.301350 | 0.291696 | 0.298070 | 0.274333 | 0.287443 | 0.460830 | 0.986029 | 1.241074 | 1.019391 | 0.627458 | 0.603219 | 0.665106 | 0.527627 | 0.372689 | 0.331214 | 0.354620 | 0.630369 | 0.986516 | 1.161353 | 0.881240 | 0.755545 | 0.723619 | 0.604610 | 0.406790 | 0.293728 | 0.271613 | 0.295440 | 0.307662 | 0.300384 | 0.266519 | 0.228991 | 0.230255 | 0.231922 | 0.232644 | 0.251917 | 0.297243 | 0.365083 | 0.532825 | 0.519013 | 0.480356 | 0.468374 | 0.512816 | 0.601637 | 0.920031 | 1.881489 | 4.762434 | 8.887622 | 18.438777 | 17.328229 | 13.967581 | 12.190330 | 9.984234 | 10.559551 | 10.705390 | 12.005675 | 18.109384 | 14.812821 | 12.102007 | 11.147635 | 7.190846 | 8.775738 | 6.171023 | 2.342118 | 1.332386 | 0.849553 | 0.508444 | 0.358643 | 0.239570 | 0.219309 | 0.221769 | 0.235892 | 0.281395 | 0.357437 | 0.478224 | 0.517268 | 0.400987 | 0.280889 | 0.238528 | 0.219791 | 0.214691 | 0.215458 | 0.215701 | 0.214630 | 0.214664 | 0.216570 | 0.233729 | 0.244631 | 0.262676 | 0.241652 | 0.241537 | 0.229116 | 0.233599 | 0.219010 | 0.216802 | 0.220900 | 0.225759 | 0.225199 | 0.228225 | 0.217572 | 0.214959 | 0.214733 | 0.214759 | 0.214786 | 0.219010 | 0.219003 | 0.216559 | 0.214670 | 0.221193 | 0.246382 | 0.261870 | 0.291702 | 0.280582 | 0.255384 | 0.216554 | 0.227682 | 0.289452 | 0.318307 | 0.326798 | 0.313999 | 0.307659 | 0.261572 | 0.222957 | 0.216244 | 0.230175 | 0.240761 | 0.288620 | 0.403363 | 0.409207 | 0.389837 | 0.324694 | 0.270857 | 0.267788 | 0.228884 | 0.214729 | 0.217434 | 0.227222 | 0.228427 | 0.221403 | 0.218963 | 0.217952 | 0.219772 | 0.215321 | 0.214708 | 0.215488 | 0.215330 | 0.218170 | 0.228522 | 0.242231 | 0.253840 | 0.282436 |
| right anterior | 0.275445 | 0.214853 | 0.224380 | 0.231542 | 0.249241 | 0.214975 | 0.215734 | 0.241743 | 0.458763 | 0.838555 | 1.935110 | 2.535897 | 0.631035 | 0.555762 | 0.609948 | 0.603832 | 0.705944 | 0.245028 | 0.232553 | 0.241269 | 0.268126 | 0.237261 | 0.257977 | 0.679326 | 0.414295 | 0.249931 | 0.239721 | 0.269377 | 0.227370 | 0.238797 | 0.958674 | 1.936867 | 2.130851 | 3.901644 | 2.536275 | 0.653396 | 0.334687 | 0.264806 | 0.236595 | 0.243198 | 0.521824 | 1.304096 | 2.658130 | 3.594171 | 5.648318 | 13.298248 | 17.566601 | 8.105139 | 8.884850 | 14.235388 | 12.650525 | 12.602909 | 18.620857 | 21.741973 | 14.210552 | 8.384860 | 8.533317 | 6.599218 | 4.362456 | 1.813176 | 1.384509 | 1.932447 | 1.922582 | 1.118654 | 0.620063 | 0.589622 | 0.839519 | 0.703109 | 0.447954 | 0.458318 | 0.404961 | 0.398078 | 0.316362 | 0.244056 | 0.221110 | 0.215835 | 0.214671 | 0.223908 | 0.219288 | 0.232061 | 0.318509 | 0.361335 | 0.457515 | 0.731509 | 0.974396 | 1.498546 | 1.630813 | 1.077658 | 0.946295 | 0.715435 | 0.444134 | 0.218233 | 0.331775 | 0.762295 | 1.418291 | 0.806632 | 0.604851 | 0.402982 | 0.271518 | 0.215531 | 0.217916 | 0.218277 | 0.220757 | 0.220155 | 0.223168 | 0.222622 | 0.222604 | 0.228817 | 0.252488 | 0.231943 | 0.217476 | 0.217329 | 0.221990 | 0.214784 | 0.214734 | 0.214654 | 0.219493 | 0.250062 | 0.393740 | 0.835875 | 1.240838 | 2.128326 | 2.260342 | 1.829135 | 1.407008 | 1.253137 | 0.811034 | 0.549327 | 0.551482 | 0.662682 | 0.703054 | 0.777082 | 0.607314 | 0.675740 | 0.525218 | 0.431279 | 0.415801 | 0.409242 | 0.415136 | 0.509273 | 0.573630 | 1.106476 | 1.504286 | 2.014756 | 3.879656 | 3.265358 | 1.865080 | 1.210942 | 0.821020 | 0.621819 | 0.559387 | 0.404598 | 0.318240 | 0.296475 | 0.339220 | 0.345588 | 0.325840 | 0.289123 | 0.299317 | 0.364237 | 0.416263 | 0.446506 | 0.582328 | 0.775720 | 0.811724 | 0.652023 | 0.565592 | 0.486050 | 0.395081 | 0.334639 | 0.332245 | 0.378236 | 0.486169 | 0.856037 | 1.351072 | 1.357388 | 1.370552 | 0.944111 | 0.659792 | 0.657899 | 0.469145 | 0.406666 | 0.375090 | 0.284788 | 0.280873 | 0.301988 | 0.303960 | 0.286472 | 0.303615 | 0.275328 | 0.275646 | 0.318047 | 0.350029 | 0.341309 | 0.369511 | 0.350244 | 0.371376 | 0.352040 | 0.285318 | 0.267948 | 0.255199 | 0.244057 | 0.227842 | 0.227083 | 0.240366 | 0.268703 | 0.280588 | 0.269428 | 0.280388 | 0.321553 | 0.401666 | 0.432175 | 0.367819 | 0.290670 | 0.264062 | 0.253573 | 0.237077 | 0.221466 | 0.219029 | 0.222345 | 0.243118 | 0.269633 | 0.266010 | 0.292829 | 0.299769 | 0.268690 | 0.252437 | 0.230979 | 0.218205 | 0.216586 | 0.214629 | 0.214904 | 0.218456 | 0.220997 | 0.225786 | 0.244997 | 0.270395 | 0.296990 | 0.283467 | 0.262982 | 0.286245 | 0.278320 | 0.251751 | 0.228019 | 0.218285 | 0.221896 | 0.237173 | 0.229582 | 0.234667 | 0.241860 | 0.260353 | 0.292335 | 0.278586 | 0.242096 | 0.232911 | 0.229081 | 0.234537 | 0.232894 | 0.220320 | 0.216402 | 0.215130 | 0.214854 | 0.215166 | 0.214722 | 0.214693 | 0.218867 | 0.219655 | 0.228493 | 0.245237 | 0.264474 | 0.253714 | 0.238359 | 0.222847 | 0.222752 | 0.219651 | 0.214791 | 0.216046 | 0.218475 | 0.220575 | 0.217287 |
| left central | 0.215949 | 0.240321 | 0.265544 | 0.263639 | 0.220932 | 0.215561 | 0.225274 | 0.219524 | 0.214704 | 0.214834 | 0.215905 | 0.215976 | 0.214633 | 0.220563 | 0.218853 | 0.216323 | 0.241197 | 0.438285 | 0.496231 | 1.273512 | 3.376538 | 5.452251 | 4.177476 | 2.429471 | 0.617969 | 0.364639 | 0.257838 | 0.221293 | 0.225471 | 0.276233 | 0.415646 | 0.765075 | 1.812104 | 3.569537 | 5.428144 | 3.367373 | 2.130456 | 0.701387 | 0.451313 | 0.341859 | 0.253173 | 0.215648 | 0.225112 | 0.299586 | 0.414109 | 0.313815 | 0.266463 | 0.218480 | 0.215464 | 0.243889 | 0.272446 | 0.328958 | 0.365222 | 0.405991 | 0.438270 | 0.435441 | 0.357304 | 0.254227 | 0.218436 | 0.215728 | 0.222754 | 0.228857 | 0.265393 | 0.353047 | 0.813044 | 3.839369 | 2.644199 | 1.243453 | 0.600478 | 0.357360 | 0.291709 | 0.328576 | 0.384476 | 1.028945 | 2.117640 | 3.757809 | 6.758787 | 7.942247 | 6.921419 | 7.210588 | 2.718049 | 1.450951 | 0.795414 | 0.817246 | 1.458284 | 1.839511 | 1.395574 | 1.054240 | 0.728407 | 0.593851 | 0.429152 | 0.291573 | 0.243931 | 0.237851 | 0.278037 | 0.321344 | 0.390347 | 0.681334 | 1.449164 | 2.708650 | 4.289635 | 4.192022 | 4.493244 | 12.400660 | 13.199147 | 20.066457 | 18.712616 | 24.022871 | 23.579554 | 12.815184 | 3.790350 | 1.995024 | 1.135697 | 1.076407 | 0.854246 | 0.702317 | 1.038004 | 1.489532 | 1.641403 | 2.650231 | 3.182169 | 2.812850 | 3.757053 | 3.789476 | 3.624761 | 4.164482 | 4.401090 | 4.721053 | 6.076323 | 2.694897 | 1.438644 | 1.149371 | 0.966513 | 1.015731 | 1.806484 | 2.967830 | 16.544177 | 54.695185 | 100.947143 | 83.553592 | 30.476044 | 9.585749 | 4.928603 | 2.687335 | 2.131129 | 1.966038 | 2.333590 | 4.287988 | 6.704957 | 7.178941 | 9.647354 | 11.594816 | 9.016499 | 9.984515 | 9.326033 | 7.848436 | 4.389365 | 1.760248 | 0.799098 | 0.452789 | 0.336094 | 0.335481 | 0.332283 | 0.398228 | 0.431964 | 0.524786 | 0.752857 | 0.936914 | 0.763158 | 0.930553 | 0.784188 | 0.888265 | 0.878261 | 0.983972 | 1.360791 | 1.721459 | 1.007333 | 0.979607 | 1.641613 | 5.157501 | 17.523274 | 24.895901 | 47.433366 | 188.810957 | 556.114719 | 963.595519 | 576.146381 | 350.533558 | 70.123011 | 13.360157 | 2.896987 | 1.391148 | 0.614432 | 0.440856 | 0.298243 | 0.256513 | 0.238721 | 0.226340 | 0.216692 | 0.220633 | 0.232277 | 0.271261 | 0.382776 | 0.508650 | 0.867872 | 1.281123 | 2.584912 | 3.461141 | 3.695500 | 3.288466 | 3.455278 | 2.661681 | 2.459531 | 2.103777 | 2.288879 | 3.398889 | 3.086687 | 3.570950 | 4.343289 | 4.444661 | 2.723906 | 1.212138 | 0.587128 | 0.528313 | 0.464168 | 0.469115 | 0.494965 | 0.707191 | 1.015861 | 1.028426 | 0.684817 | 0.489116 | 0.337729 | 0.272324 | 0.227568 | 0.231129 | 0.241378 | 0.274069 | 0.291593 | 0.309143 | 0.278203 | 0.256746 | 0.230168 | 0.225271 | 0.220169 | 0.214741 | 0.227994 | 0.243472 | 0.272863 | 0.296840 | 0.339003 | 0.373871 | 0.308842 | 0.252844 | 0.224365 | 0.215966 | 0.227341 | 0.274500 | 0.362050 | 0.371510 | 0.313336 | 0.262359 | 0.231171 | 0.230132 | 0.220018 | 0.216361 | 0.214895 | 0.215745 | 0.215512 | 0.220510 | 0.233471 | 0.258378 | 0.338579 | 0.372809 | 0.374686 | 0.307595 | 0.257815 | 0.262393 | 0.265527 | 0.230415 |
| right central | 0.278687 | 0.361740 | 1.282139 | 3.510838 | 16.636586 | 15.277338 | 5.261839 | 1.567072 | 1.299636 | 0.518038 | 0.308768 | 0.272719 | 0.278839 | 0.265570 | 0.272724 | 0.217767 | 0.244554 | 0.297900 | 0.587337 | 0.726979 | 0.563727 | 0.451816 | 0.521746 | 0.411089 | 0.266113 | 0.216341 | 0.219727 | 0.244700 | 0.222029 | 0.214949 | 0.214663 | 0.229958 | 0.284764 | 0.292914 | 0.333566 | 0.431797 | 0.696601 | 0.613265 | 0.542508 | 0.330964 | 0.237552 | 0.219446 | 0.214646 | 0.256069 | 0.350019 | 0.473153 | 0.362104 | 0.286007 | 0.235162 | 0.221209 | 0.214630 | 0.219324 | 0.229438 | 0.214778 | 0.218767 | 0.243687 | 0.227177 | 0.215959 | 0.290778 | 0.430996 | 0.535446 | 0.708227 | 0.663394 | 0.543797 | 0.408844 | 0.304315 | 0.253271 | 0.262668 | 0.234439 | 0.221238 | 0.217541 | 0.232171 | 0.264908 | 0.404421 | 0.525525 | 0.646085 | 0.610229 | 0.359596 | 0.248569 | 0.214637 | 0.296637 | 0.301708 | 0.256403 | 0.227067 | 0.562521 | 0.862260 | 1.001863 | 1.379585 | 1.088874 | 0.879253 | 0.509345 | 0.442956 | 0.365959 | 0.291523 | 0.270721 | 0.305495 | 0.508910 | 0.985816 | 1.328343 | 2.453759 | 4.942401 | 8.786918 | 7.203845 | 4.107885 | 4.365594 | 2.257501 | 1.064259 | 0.877742 | 0.608574 | 0.423945 | 0.281643 | 0.224511 | 0.221048 | 0.241537 | 0.327683 | 0.569033 | 0.833308 | 1.485749 | 2.350197 | 3.044698 | 2.718926 | 1.709500 | 1.037632 | 0.997642 | 1.243491 | 1.118604 | 1.012928 | 1.155999 | 1.279805 | 1.578641 | 2.675103 | 8.221072 | 23.774745 | 25.497246 | 18.266042 | 11.417211 | 17.229966 | 12.627484 | 7.302865 | 7.711649 | 12.593301 | 34.643661 | 107.087785 | 89.423573 | 45.280649 | 21.609812 | 9.043404 | 4.652153 | 2.385781 | 1.339226 | 1.357380 | 1.421533 | 1.668298 | 2.362496 | 3.357227 | 4.417917 | 5.884737 | 4.706863 | 5.327293 | 4.657551 | 4.488478 | 4.672240 | 6.165625 | 6.071493 | 10.292753 | 9.889196 | 12.564574 | 16.671494 | 19.348150 | 22.603035 | 37.121104 | 38.719079 | 27.189393 | 13.691087 | 10.124949 | 7.570194 | 6.101661 | 6.532751 | 3.898542 | 3.437851 | 2.679128 | 2.645249 | 2.389691 | 1.981579 | 1.392756 | 2.186906 | 3.973518 | 10.596875 | 18.032802 | 29.452661 | 14.158638 | 14.365759 | 10.869738 | 11.235205 | 13.911616 | 16.279546 | 11.957367 | 11.413219 | 4.778643 | 2.680448 | 1.278002 | 1.075383 | 0.940633 | 0.876775 | 1.094249 | 1.312123 | 1.473736 | 1.866582 | 1.813877 | 1.915257 | 2.538296 | 2.662228 | 3.975058 | 5.662085 | 4.675857 | 3.605813 | 3.610449 | 2.363784 | 1.658998 | 0.775867 | 0.522610 | 0.389361 | 0.325568 | 0.296367 | 0.299814 | 0.318235 | 0.386994 | 0.449860 | 0.594657 | 0.814062 | 0.967632 | 0.941229 | 0.567792 | 0.397438 | 0.353489 | 0.324289 | 0.300495 | 0.281475 | 0.283212 | 0.363635 | 0.450820 | 0.498630 | 0.539726 | 0.582158 | 0.696538 | 0.690440 | 0.820591 | 0.732629 | 0.520643 | 0.369913 | 0.291745 | 0.260688 | 0.247106 | 0.220306 | 0.216018 | 0.226923 | 0.255510 | 0.309268 | 0.318902 | 0.327201 | 0.428719 | 0.438006 | 0.411372 | 0.410125 | 0.355098 | 0.313203 | 0.322524 | 0.313359 | 0.303021 | 0.274150 | 0.256963 | 0.249827 | 0.280920 | 0.272848 | 0.227327 | 0.224945 | 0.228349 | 0.225957 | 0.229809 | 0.216702 |
| left posterior | 0.277342 | 0.234365 | 0.287452 | 0.392737 | 0.612479 | 0.273483 | 0.274636 | 0.278142 | 0.332542 | 0.319484 | 0.536359 | 0.421360 | 0.994122 | 1.340642 | 1.716774 | 2.621301 | 1.116008 | 0.377364 | 0.299164 | 0.249786 | 0.257032 | 0.393675 | 0.329181 | 0.297352 | 0.295453 | 0.291955 | 0.247227 | 0.216339 | 0.315648 | 0.294360 | 0.224970 | 0.237225 | 0.319947 | 0.375599 | 0.346784 | 0.289442 | 0.305544 | 0.333055 | 0.278356 | 0.227251 | 0.215540 | 0.257568 | 0.344536 | 0.536150 | 0.819535 | 1.691642 | 1.433358 | 0.912236 | 0.528576 | 0.325999 | 0.258808 | 0.222139 | 0.227854 | 0.222532 | 0.218374 | 0.217838 | 0.216166 | 0.215027 | 0.216032 | 0.261360 | 0.254572 | 0.261750 | 0.303441 | 0.329485 | 0.369472 | 0.393480 | 0.401796 | 0.461610 | 0.435935 | 0.365247 | 0.325372 | 0.368367 | 0.341304 | 0.248551 | 0.228368 | 0.225316 | 0.232194 | 0.232723 | 0.216182 | 0.214689 | 0.217643 | 0.215303 | 0.216359 | 0.214706 | 0.215036 | 0.225669 | 0.239893 | 0.260316 | 0.297272 | 0.362460 | 0.390755 | 0.456487 | 0.393071 | 0.354683 | 0.313978 | 0.294282 | 0.317086 | 0.334950 | 0.339857 | 0.335197 | 0.383509 | 0.460949 | 0.478427 | 0.391257 | 0.339598 | 0.388331 | 0.607069 | 0.657303 | 0.492655 | 0.475727 | 0.530887 | 0.590112 | 0.686675 | 0.693331 | 0.597001 | 0.504646 | 0.589410 | 0.727111 | 0.897257 | 0.976158 | 0.933140 | 0.946704 | 0.923652 | 0.594887 | 0.402670 | 0.303215 | 0.242225 | 0.214785 | 0.245611 | 0.308217 | 0.335211 | 0.324083 | 0.307951 | 0.295882 | 0.252224 | 0.215825 | 0.221793 | 0.234523 | 0.259385 | 0.281028 | 0.272444 | 0.281482 | 0.254724 | 0.235157 | 0.217359 | 0.219931 | 0.235994 | 0.252533 | 0.263815 | 0.247494 | 0.235821 | 0.221233 | 0.219105 | 0.218731 | 0.214683 | 0.215978 | 0.242379 | 0.321763 | 0.388681 | 0.380907 | 0.433525 | 0.519075 | 0.711882 | 0.698458 | 0.527566 | 0.429417 | 0.410523 | 0.390698 | 0.407150 | 0.397930 | 0.400052 | 0.444660 | 0.456457 | 0.381416 | 0.341603 | 0.355466 | 0.306108 | 0.302484 | 0.278672 | 0.276633 | 0.294171 | 0.299302 | 0.303776 | 0.287307 | 0.266731 | 0.246319 | 0.225513 | 0.216570 | 0.214848 | 0.221632 | 0.227163 | 0.241484 | 0.239039 | 0.240337 | 0.223265 | 0.215161 | 0.214645 | 0.216254 | 0.218514 | 0.222703 | 0.238131 | 0.229925 | 0.217140 | 0.214645 | 0.226993 | 0.260573 | 0.324738 | 0.396322 | 0.410481 | 0.356930 | 0.286532 | 0.242574 | 0.219123 | 0.214644 | 0.214664 | 0.216040 | 0.216386 | 0.215410 | 0.215168 | 0.214884 | 0.214639 | 0.216239 | 0.223076 | 0.235066 | 0.238138 | 0.232468 | 0.229336 | 0.224921 | 0.227523 | 0.224656 | 0.216360 | 0.214782 | 0.216814 | 0.222457 | 0.236886 | 0.291861 | 0.374611 | 0.485479 | 0.581382 | 0.581956 | 0.641663 | 0.659420 | 0.629958 | 0.543670 | 0.429172 | 0.310887 | 0.252275 | 0.220813 | 0.219042 | 0.250424 | 0.309366 | 0.384642 | 0.457838 | 0.546568 | 0.549168 | 0.439733 | 0.383845 | 0.325258 | 0.333581 | 0.304935 | 0.317412 | 0.363807 | 0.478831 | 0.586060 | 0.925133 | 1.218235 | 1.939749 | 2.117358 | 1.893827 | 1.732720 | 2.035364 | 2.206585 | 2.893524 | 4.891610 | 7.576601 | 8.618723 | 7.168484 | 7.264573 | 5.656169 | 4.088448 |
| right posterior | 1.885130 | 1.792751 | 1.853794 | 2.901505 | 1.928755 | 0.861165 | 1.012688 | 1.385365 | 1.132759 | 1.134144 | 1.023296 | 1.271310 | 3.212430 | 1.294478 | 0.805150 | 1.055124 | 1.102867 | 0.729999 | 0.365117 | 0.252027 | 0.321615 | 0.297697 | 0.319461 | 0.342547 | 0.380039 | 0.435559 | 0.572684 | 0.503664 | 0.402738 | 0.266018 | 0.231107 | 0.223976 | 0.241331 | 0.359246 | 0.477942 | 0.574141 | 0.457278 | 0.521265 | 0.430330 | 0.342028 | 0.346832 | 0.328220 | 0.349988 | 0.371359 | 0.535423 | 0.763158 | 0.968561 | 0.718232 | 0.684595 | 0.518691 | 0.326680 | 0.226561 | 0.214670 | 0.222565 | 0.227911 | 0.230034 | 0.239693 | 0.229913 | 0.217468 | 0.214647 | 0.216369 | 0.214654 | 0.222181 | 0.230307 | 0.220242 | 0.247481 | 0.386096 | 0.682920 | 0.752697 | 0.992850 | 2.163536 | 5.619740 | 15.445941 | 43.166338 | 66.279216 | 81.468846 | 53.929867 | 13.265388 | 5.135608 | 1.667357 | 0.874980 | 0.841686 | 0.588189 | 0.641466 | 0.964923 | 1.456502 | 2.722230 | 3.974075 | 11.001893 | 31.596060 | 89.739249 | 136.349985 | 111.252181 | 106.130793 | 105.158138 | 116.383667 | 114.246204 | 136.760378 | 136.955687 | 115.612849 | 168.107886 | 116.833499 | 74.889975 | 45.346744 | 54.958457 | 39.408072 | 27.824262 | 30.160426 | 72.723254 | 154.918652 | 325.436222 | 129.138895 | 88.220867 | 59.047814 | 47.690807 | 57.453755 | 42.105971 | 34.366214 | 30.465123 | 23.432058 | 25.912330 | 11.547250 | 5.986732 | 4.841998 | 3.609844 | 3.699532 | 5.209401 | 5.204166 | 11.526992 | 12.684953 | 8.558272 | 14.278955 | 40.709556 | 86.486015 | 135.671382 | 205.421035 | 345.587320 | 921.551689 | 2536.178443 | 2137.759252 | 1330.174295 | 1073.130599 | 519.695336 | 266.836150 | 66.088735 | 18.176124 | 7.870820 | 7.656405 | 6.182773 | 3.068797 | 1.915547 | 2.159052 | 2.535740 | 4.977298 | 7.343496 | 15.348732 | 94.089114 | 98.755829 | 62.315789 | 34.434166 | 25.150809 | 21.526768 | 11.430511 | 2.748073 | 1.362656 | 0.824034 | 0.538769 | 0.390007 | 0.274156 | 0.237907 | 0.225617 | 0.232812 | 0.254017 | 0.299351 | 0.355775 | 0.533362 | 0.897830 | 2.130931 | 3.556408 | 4.401747 | 6.611020 | 9.310712 | 10.278270 | 8.556245 | 5.477573 | 4.446646 | 3.689463 | 2.255462 | 1.628647 | 0.963113 | 0.815817 | 0.805941 | 0.700784 | 0.649711 | 0.482432 | 0.419618 | 0.460335 | 0.497627 | 0.589198 | 0.784943 | 0.749479 | 0.836274 | 1.003758 | 1.569484 | 2.225480 | 3.139434 | 3.153614 | 3.544022 | 4.352739 | 5.334950 | 4.208191 | 2.922875 | 2.465903 | 3.648473 | 4.545796 | 3.524979 | 1.634808 | 0.892107 | 0.542565 | 0.409287 | 0.306813 | 0.246803 | 0.224394 | 0.223823 | 0.227806 | 0.254715 | 0.265159 | 0.284537 | 0.310403 | 0.376610 | 0.417714 | 0.478759 | 0.375920 | 0.348372 | 0.326380 | 0.412753 | 0.695466 | 1.600369 | 2.541773 | 4.877667 | 6.446540 | 8.676932 | 6.005615 | 3.618996 | 2.213209 | 1.681154 | 0.957993 | 0.499087 | 0.292458 | 0.237307 | 0.214626 | 0.218361 | 0.215356 | 0.217232 | 0.266318 | 0.510959 | 1.105977 | 2.137539 | 2.876386 | 1.994723 | 0.841556 | 0.403449 | 0.264483 | 0.220574 | 0.214787 | 0.216199 | 0.215508 | 0.216330 | 0.226859 | 0.243175 | 0.247212 | 0.216042 | 0.227104 | 0.341380 | 0.502533 | 0.716658 | 1.015725 | 1.606309 | 1.320543 | 0.906039 |
| all electrodes | 0.352186 | 0.214640 | 0.236650 | 0.243705 | 0.274025 | 0.275766 | 0.298761 | 0.214640 | 0.380866 | 0.871782 | 0.947390 | 1.081449 | 0.634355 | 0.601901 | 0.283469 | 0.268421 | 0.215533 | 0.216204 | 0.344768 | 1.057585 | 2.051371 | 1.154757 | 2.381822 | 2.583139 | 1.000157 | 0.326724 | 0.234835 | 0.226133 | 0.228060 | 0.214711 | 0.262258 | 0.223722 | 0.214662 | 0.246723 | 0.411639 | 0.701716 | 0.943104 | 1.691131 | 1.120935 | 1.031044 | 0.302257 | 0.218584 | 0.296135 | 0.366578 | 0.763034 | 1.117072 | 1.981534 | 2.153005 | 1.750629 | 1.133683 | 0.889909 | 0.835150 | 0.505609 | 0.400273 | 0.385988 | 0.331906 | 0.254045 | 0.223673 | 0.219522 | 0.216088 | 0.221756 | 0.216999 | 0.214688 | 0.215065 | 0.222658 | 0.302630 | 0.314929 | 0.247882 | 0.241106 | 0.220439 | 0.215674 | 0.216534 | 0.318101 | 0.561922 | 0.749900 | 0.580655 | 0.704026 | 0.646770 | 0.631943 | 0.369112 | 0.279078 | 0.276288 | 0.451555 | 0.611680 | 0.802197 | 0.723416 | 0.629331 | 0.629621 | 0.563423 | 0.599105 | 0.664588 | 0.592603 | 0.469473 | 0.454111 | 0.493028 | 0.642533 | 0.528286 | 0.376467 | 0.387599 | 0.510197 | 0.580118 | 0.760925 | 0.698037 | 1.020822 | 2.001441 | 2.186133 | 1.374873 | 1.263001 | 1.257797 | 1.098490 | 0.627309 | 0.345811 | 0.253968 | 0.245628 | 0.238529 | 0.219698 | 0.231917 | 0.273902 | 0.389009 | 0.630267 | 1.279435 | 2.306335 | 2.928708 | 1.428544 | 0.658895 | 0.614831 | 0.451676 | 0.335316 | 0.269181 | 0.238718 | 0.230981 | 0.229469 | 0.222741 | 0.216054 | 0.216468 | 0.236057 | 0.273412 | 0.311194 | 0.444908 | 0.475080 | 0.643775 | 1.148104 | 2.415600 | 4.420217 | 4.534512 | 2.712543 | 2.574136 | 2.767217 | 1.508913 | 0.727791 | 0.466500 | 0.403051 | 0.415904 | 0.365259 | 0.302659 | 0.238183 | 0.218370 | 0.214994 | 0.220622 | 0.229669 | 0.234520 | 0.257374 | 0.242179 | 0.215343 | 0.238802 | 0.305703 | 0.358958 | 0.497579 | 0.790632 | 1.077613 | 0.799541 | 0.464295 | 0.338451 | 0.313234 | 0.253853 | 0.247297 | 0.279004 | 0.316444 | 0.391413 | 0.677820 | 0.837006 | 1.172170 | 1.434725 | 1.358575 | 1.348091 | 2.055580 | 2.363294 | 3.712261 | 6.482752 | 11.045943 | 16.422766 | 18.212669 | 16.823656 | 9.144058 | 5.713685 | 3.168420 | 1.872236 | 1.412428 | 0.942908 | 0.784181 | 0.769454 | 0.888057 | 1.246015 | 2.531533 | 6.586004 | 16.673105 | 27.722043 | 47.047247 | 37.640537 | 17.035284 | 9.348971 | 6.585411 | 10.057502 | 20.398488 | 39.231918 | 68.496901 | 100.169210 | 39.705865 | 10.516338 | 3.572966 | 1.109433 | 0.507464 | 0.325829 | 0.264907 | 0.297107 | 0.440359 | 0.674563 | 1.086934 | 1.556228 | 1.876566 | 2.119047 | 1.771466 | 1.161398 | 0.795549 | 0.654258 | 0.576455 | 0.702515 | 0.794836 | 0.778115 | 1.080930 | 1.455792 | 2.057476 | 2.932552 | 2.174518 | 2.339003 | 2.153142 | 1.390387 | 1.041868 | 0.988859 | 0.747130 | 0.712412 | 0.516201 | 0.509410 | 0.452338 | 0.413170 | 0.366024 | 0.384403 | 0.354847 | 0.305039 | 0.267180 | 0.248771 | 0.236069 | 0.233950 | 0.227947 | 0.219670 | 0.223822 | 0.233381 | 0.267398 | 0.283954 | 0.272843 | 0.245976 | 0.244052 | 0.224453 | 0.217792 | 0.216705 | 0.225227 | 0.239915 | 0.242485 | 0.252728 | 0.246321 |

Searchlight, spatiotemporal cluster permutation test

|  | start time | stop time | peak time | peak channel | cluster p | peak Cohen's d | direction |
| --- | --- | --- | --- | --- | --- | --- | --- |
| #1 | 225 | 340 | 300 | O2 | 0.0167 | 0.910399 | positive |

L) difference, sex, happy vs sad

  
|  | time window | peak latency | cluster *p* | peak Cohen's *d* |  | | | |
| **all electrodes** |  | | | |  | | | |
|  | | | | | | | | |

Time-resolved classification, cluster permutation tests

|  | **left hemisphere** | | | | **right hemisphere** | | | |
|  | time window | peak latency | cluster *p* | peak Cohen's *d* | time window | peak latency | cluster *p* | peak Cohen's *d* |
| **anterior** |  | | | |  | | | |
| **central** | 180 - 280 ms | 190 ms | 0.0333 | 0.9166 |  | | | |
| **posterior** |  | | | |  | | | |

  

Time-resolved classification, Bayesian statistics

|  | -200 | -195 | -190 | -185 | -180 | -175 | -170 | -165 | -160 | -155 | -150 | -145 | -140 | -135 | -130 | -125 | -120 | -115 | -110 | -105 | -100 | -95 | -90 | -85 | -80 | -75 | -70 | -65 | -60 | -55 | -50 | -45 | -40 | -35 | -30 | -25 | -20 | -15 | -10 | -5 | 0 | 5 | 10 | 15 | 20 | 25 | 30 | 35 | 40 | 45 | 50 | 55 | 60 | 65 | 70 | 75 | 80 | 85 | 90 | 95 | 100 | 105 | 110 | 115 | 120 | 125 | 130 | 135 | 140 | 145 | 150 | 155 | 160 | 165 | 170 | 175 | 180 | 185 | 190 | 195 | 200 | 205 | 210 | 215 | 220 | 225 | 230 | 235 | 240 | 245 | 250 | 255 | 260 | 265 | 270 | 275 | 280 | 285 | 290 | 295 | 300 | 305 | 310 | 315 | 320 | 325 | 330 | 335 | 340 | 345 | 350 | 355 | 360 | 365 | 370 | 375 | 380 | 385 | 390 | 395 | 400 | 405 | 410 | 415 | 420 | 425 | 430 | 435 | 440 | 445 | 450 | 455 | 460 | 465 | 470 | 475 | 480 | 485 | 490 | 495 | 500 | 505 | 510 | 515 | 520 | 525 | 530 | 535 | 540 | 545 | 550 | 555 | 560 | 565 | 570 | 575 | 580 | 585 | 590 | 595 | 600 | 605 | 610 | 615 | 620 | 625 | 630 | 635 | 640 | 645 | 650 | 655 | 660 | 665 | 670 | 675 | 680 | 685 | 690 | 695 | 700 | 705 | 710 | 715 | 720 | 725 | 730 | 735 | 740 | 745 | 750 | 755 | 760 | 765 | 770 | 775 | 780 | 785 | 790 | 795 | 800 | 805 | 810 | 815 | 820 | 825 | 830 | 835 | 840 | 845 | 850 | 855 | 860 | 865 | 870 | 875 | 880 | 885 | 890 | 895 | 900 | 905 | 910 | 915 | 920 | 925 | 930 | 935 | 940 | 945 | 950 | 955 | 960 | 965 | 970 | 975 | 980 | 985 | 990 | 995 | 1000 | 1005 | 1010 | 1015 | 1020 | 1025 | 1030 | 1035 | 1040 | 1045 | 1050 | 1055 | 1060 | 1065 | 1070 | 1075 | 1080 | 1085 | 1090 | 1095 | 1100 | 1105 | 1110 | 1115 | 1120 | 1125 | 1130 | 1135 | 1140 | 1145 | 1150 | 1155 | 1160 | 1165 | 1170 | 1175 | 1180 | 1185 | 1190 | 1195 |
| --- | --- | --- | --- | --- | --- | --- | --- | --- | --- | --- | --- | --- | --- | --- | --- | --- | --- | --- | --- | --- | --- | --- | --- | --- | --- | --- | --- | --- | --- | --- | --- | --- | --- | --- | --- | --- | --- | --- | --- | --- | --- | --- | --- | --- | --- | --- | --- | --- | --- | --- | --- | --- | --- | --- | --- | --- | --- | --- | --- | --- | --- | --- | --- | --- | --- | --- | --- | --- | --- | --- | --- | --- | --- | --- | --- | --- | --- | --- | --- | --- | --- | --- | --- | --- | --- | --- | --- | --- | --- | --- | --- | --- | --- | --- | --- | --- | --- | --- | --- | --- | --- | --- | --- | --- | --- | --- | --- | --- | --- | --- | --- | --- | --- | --- | --- | --- | --- | --- | --- | --- | --- | --- | --- | --- | --- | --- | --- | --- | --- | --- | --- | --- | --- | --- | --- | --- | --- | --- | --- | --- | --- | --- | --- | --- | --- | --- | --- | --- | --- | --- | --- | --- | --- | --- | --- | --- | --- | --- | --- | --- | --- | --- | --- | --- | --- | --- | --- | --- | --- | --- | --- | --- | --- | --- | --- | --- | --- | --- | --- | --- | --- | --- | --- | --- | --- | --- | --- | --- | --- | --- | --- | --- | --- | --- | --- | --- | --- | --- | --- | --- | --- | --- | --- | --- | --- | --- | --- | --- | --- | --- | --- | --- | --- | --- | --- | --- | --- | --- | --- | --- | --- | --- | --- | --- | --- | --- | --- | --- | --- | --- | --- | --- | --- | --- | --- | --- | --- | --- | --- | --- | --- | --- | --- | --- | --- | --- | --- | --- | --- | --- | --- | --- | --- | --- | --- | --- | --- | --- | --- | --- | --- | --- | --- | --- | --- | --- | --- | --- | --- | --- | --- | --- | --- | --- | --- | --- | --- | --- | --- | --- |
| left anterior | 0.670785 | 1.184440 | 1.462271 | 1.486799 | 1.434050 | 0.556051 | 0.306879 | 0.245686 | 0.216083 | 0.259506 | 0.371202 | 0.236432 | 0.214731 | 0.214677 | 0.215143 | 0.247049 | 0.693522 | 2.514288 | 11.517435 | 17.093783 | 12.029358 | 4.194798 | 0.350219 | 0.283473 | 1.825274 | 4.340424 | 8.361457 | 33.256710 | 80.516838 | 45.428817 | 12.056413 | 1.992242 | 0.922383 | 0.299935 | 0.218475 | 0.331642 | 0.401310 | 0.454144 | 0.649820 | 0.629129 | 0.539521 | 0.369051 | 0.278730 | 0.258670 | 0.269988 | 0.256437 | 0.284421 | 0.230444 | 0.215745 | 0.272994 | 0.300795 | 0.375706 | 0.508713 | 0.497400 | 0.266379 | 0.218378 | 0.347262 | 0.559074 | 0.936397 | 1.204411 | 0.937967 | 0.436704 | 0.248224 | 0.214918 | 0.221513 | 0.251472 | 0.248210 | 0.229015 | 0.216100 | 0.214658 | 0.214630 | 0.222918 | 0.217861 | 0.223150 | 0.252571 | 0.325193 | 0.375794 | 0.320685 | 0.223354 | 0.215618 | 0.256903 | 0.424730 | 0.598941 | 0.929596 | 1.061434 | 1.601733 | 2.320965 | 4.114264 | 5.449357 | 8.831433 | 10.280891 | 3.273308 | 0.789621 | 0.332377 | 0.251917 | 0.247520 | 0.239294 | 0.221686 | 0.237869 | 0.279639 | 0.322573 | 0.266262 | 0.225233 | 0.219891 | 0.216488 | 0.215940 | 0.215331 | 0.220396 | 0.221065 | 0.216059 | 0.218983 | 0.222659 | 0.231476 | 0.279081 | 0.250785 | 0.240643 | 0.233984 | 0.226756 | 0.214857 | 0.222334 | 0.262438 | 0.271056 | 0.279265 | 0.304346 | 0.414819 | 0.471952 | 0.395280 | 0.301143 | 0.256705 | 0.257128 | 0.271444 | 0.290890 | 0.372052 | 0.562389 | 0.844025 | 1.518627 | 1.551199 | 0.821016 | 0.395476 | 0.234129 | 0.222486 | 0.231531 | 0.227301 | 0.221516 | 0.229156 | 0.267882 | 0.399792 | 0.517524 | 0.460020 | 0.390504 | 0.411389 | 0.505715 | 0.675666 | 0.682250 | 0.611955 | 0.533389 | 0.577166 | 0.602176 | 0.438277 | 0.282269 | 0.240924 | 0.244510 | 0.247085 | 0.249097 | 0.245542 | 0.246520 | 0.247337 | 0.258839 | 0.240756 | 0.226181 | 0.214713 | 0.215165 | 0.225096 | 0.244965 | 0.289044 | 0.377102 | 0.625480 | 1.072641 | 1.469212 | 1.520100 | 1.783276 | 1.832529 | 1.992919 | 2.231113 | 2.663744 | 2.872333 | 3.752903 | 3.443605 | 3.120285 | 2.175746 | 1.339204 | 1.275128 | 1.276047 | 0.840449 | 0.632815 | 0.548044 | 0.502066 | 0.466683 | 0.384262 | 0.390777 | 0.451267 | 0.504377 | 0.548158 | 0.527680 | 0.398832 | 0.349579 | 0.276104 | 0.248298 | 0.239571 | 0.216165 | 0.214881 | 0.219372 | 0.226202 | 0.241601 | 0.258054 | 0.293784 | 0.721747 | 2.018596 | 3.928067 | 4.875549 | 6.237464 | 6.837571 | 8.637324 | 4.491164 | 1.028653 | 0.550612 | 0.391578 | 0.318240 | 0.254516 | 0.223398 | 0.215720 | 0.224490 | 0.224593 | 0.225356 | 0.228731 | 0.276292 | 0.327474 | 0.324388 | 0.377309 | 0.426031 | 0.580788 | 0.629160 | 0.662735 | 0.523018 | 0.455255 | 0.329067 | 0.293395 | 0.252469 | 0.225412 | 0.215167 | 0.215909 | 0.220817 | 0.243172 | 0.273120 | 0.277943 | 0.306061 | 0.313323 | 0.285282 | 0.277069 | 0.248504 | 0.221305 | 0.215349 | 0.224849 | 0.242206 | 0.277927 | 0.318676 | 0.390373 | 0.467340 | 0.607833 | 0.656770 | 0.738110 | 0.530420 | 0.483742 | 0.424264 | 0.374095 | 0.324264 | 0.296051 | 0.258006 | 0.244968 | 0.238102 |
| right anterior | 0.216880 | 0.233770 | 0.220087 | 0.219983 | 0.228884 | 0.341208 | 0.502934 | 2.434489 | 23.546171 | 98.748600 | 108.418632 | 58.784808 | 2.676931 | 2.655160 | 0.572823 | 0.214626 | 0.423513 | 1.650421 | 7.567711 | 0.947250 | 0.728538 | 0.745141 | 0.474868 | 0.311898 | 0.242027 | 0.214770 | 0.228822 | 0.265307 | 0.307389 | 0.638531 | 0.832045 | 1.197031 | 1.411058 | 0.872295 | 0.610593 | 0.360959 | 0.220522 | 0.220588 | 0.325159 | 0.352290 | 0.269216 | 0.266525 | 0.217734 | 0.215174 | 0.216458 | 0.302637 | 0.436385 | 0.407240 | 0.385733 | 0.370246 | 0.365811 | 0.409380 | 0.348420 | 0.352457 | 0.334542 | 0.325039 | 0.265506 | 0.219244 | 0.219714 | 0.268581 | 0.409482 | 0.359948 | 0.303360 | 0.301825 | 0.267128 | 0.261769 | 0.234473 | 0.221583 | 0.243035 | 0.282469 | 0.277551 | 0.273512 | 0.257767 | 0.316365 | 0.438486 | 0.702925 | 1.087528 | 1.601867 | 1.606787 | 2.205814 | 1.647168 | 0.893627 | 0.678401 | 0.690897 | 0.759159 | 0.740684 | 0.844797 | 0.887559 | 0.792550 | 0.766642 | 0.519303 | 0.302009 | 0.224752 | 0.215051 | 0.215813 | 0.214632 | 0.216749 | 0.215730 | 0.218585 | 0.257000 | 0.307202 | 0.338879 | 0.355757 | 0.353532 | 0.372640 | 0.303452 | 0.296942 | 0.303734 | 0.360089 | 0.332262 | 0.278175 | 0.242034 | 0.233871 | 0.225728 | 0.218007 | 0.214649 | 0.215116 | 0.238705 | 0.395027 | 1.116375 | 2.725205 | 4.719381 | 2.738257 | 2.266293 | 1.727995 | 1.324326 | 0.660352 | 0.448557 | 0.374680 | 0.360695 | 0.265894 | 0.252886 | 0.243999 | 0.282118 | 0.276034 | 0.259361 | 0.273482 | 0.335471 | 0.332836 | 0.349732 | 0.283782 | 0.269393 | 0.285836 | 0.310467 | 0.342678 | 0.347732 | 0.280970 | 0.257013 | 0.269858 | 0.275367 | 0.263422 | 0.248708 | 0.254268 | 0.289935 | 0.384743 | 0.438982 | 0.495821 | 0.526871 | 0.506569 | 0.484900 | 0.446150 | 0.533194 | 0.680556 | 0.653636 | 0.555796 | 0.418747 | 0.330584 | 0.253266 | 0.216283 | 0.216237 | 0.217313 | 0.217273 | 0.215647 | 0.233992 | 0.292648 | 0.284903 | 0.268407 | 0.250722 | 0.256923 | 0.256713 | 0.245616 | 0.235542 | 0.255603 | 0.246359 | 0.253194 | 0.257068 | 0.259631 | 0.294996 | 0.453601 | 0.563651 | 0.795071 | 0.974227 | 0.854159 | 0.762515 | 0.590341 | 0.419408 | 0.369059 | 0.325974 | 0.299079 | 0.327096 | 0.343395 | 0.317993 | 0.282512 | 0.267218 | 0.273318 | 0.295686 | 0.283439 | 0.278973 | 0.405459 | 0.686064 | 1.065000 | 0.854291 | 0.509433 | 0.329795 | 0.268144 | 0.243677 | 0.230975 | 0.229728 | 0.232926 | 0.261157 | 0.345161 | 0.476179 | 0.442191 | 0.395612 | 0.302070 | 0.251365 | 0.226658 | 0.218042 | 0.216734 | 0.222232 | 0.222781 | 0.239285 | 0.281505 | 0.334997 | 0.353933 | 0.352563 | 0.344683 | 0.384538 | 0.396833 | 0.468328 | 0.696651 | 0.748948 | 0.645978 | 0.506463 | 0.438631 | 0.374071 | 0.356129 | 0.272627 | 0.249086 | 0.242733 | 0.235097 | 0.230393 | 0.230107 | 0.222181 | 0.221238 | 0.224342 | 0.226891 | 0.222194 | 0.215862 | 0.215831 | 0.228780 | 0.238457 | 0.243312 | 0.236648 | 0.222927 | 0.223935 | 0.218132 | 0.216535 | 0.218932 | 0.220548 | 0.233098 | 0.278266 | 0.363089 | 0.449074 | 0.433496 | 0.383313 | 0.383752 | 0.389610 | 0.342182 | 0.246905 |
| left central | 0.243935 | 0.255065 | 0.252904 | 0.253730 | 0.231800 | 0.215602 | 0.226011 | 0.230790 | 0.220782 | 0.237041 | 0.277158 | 0.284621 | 0.248841 | 0.254388 | 0.300203 | 0.261884 | 0.220046 | 0.307535 | 0.435371 | 0.835286 | 1.237846 | 2.610073 | 2.455465 | 0.843380 | 0.308815 | 0.215675 | 0.240201 | 0.350113 | 0.678252 | 1.409677 | 1.309731 | 0.829827 | 0.572162 | 0.555403 | 0.258318 | 0.214657 | 0.242600 | 0.236181 | 0.224329 | 0.215593 | 0.227124 | 0.289169 | 0.378211 | 0.422305 | 0.582865 | 0.502604 | 0.454296 | 0.291963 | 0.266143 | 0.250000 | 0.261821 | 0.226387 | 0.218065 | 0.216323 | 0.229121 | 0.296179 | 0.368094 | 0.333601 | 0.244952 | 0.226942 | 0.221966 | 0.218460 | 0.216839 | 0.223805 | 0.239955 | 0.235737 | 0.216690 | 0.294662 | 0.488903 | 0.758801 | 0.854869 | 0.533262 | 0.292984 | 0.214892 | 0.325158 | 1.146974 | 5.067367 | 31.070685 | 172.183572 | 699.391422 | 260.760772 | 92.944402 | 22.133358 | 14.558977 | 13.093990 | 5.975106 | 3.984956 | 3.455053 | 4.671451 | 8.935528 | 10.874193 | 6.822176 | 5.818689 | 4.325216 | 3.579914 | 2.831773 | 1.777731 | 0.951033 | 1.257114 | 1.399400 | 1.383451 | 1.309498 | 1.184741 | 1.497678 | 1.686398 | 1.693628 | 1.015409 | 0.921270 | 1.275658 | 1.025405 | 0.703068 | 0.530522 | 0.328441 | 0.393451 | 0.463925 | 0.389473 | 0.398312 | 0.409661 | 0.406155 | 0.607054 | 1.036402 | 1.506317 | 2.955899 | 3.991607 | 6.533410 | 6.338026 | 3.304800 | 1.321616 | 0.829318 | 0.487151 | 0.382176 | 0.305612 | 0.290161 | 0.288395 | 0.321508 | 0.389434 | 0.415220 | 0.362398 | 0.298101 | 0.243317 | 0.234698 | 0.220095 | 0.216306 | 0.215283 | 0.216732 | 0.230282 | 0.242202 | 0.275863 | 0.316187 | 0.363200 | 0.343522 | 0.317563 | 0.289433 | 0.315620 | 0.282956 | 0.275890 | 0.265563 | 0.253615 | 0.254001 | 0.272291 | 0.270909 | 0.337251 | 0.344409 | 0.438156 | 0.665559 | 1.136611 | 2.010159 | 3.930611 | 3.678658 | 3.796210 | 1.887364 | 1.424023 | 0.939194 | 0.806357 | 0.592859 | 0.413721 | 0.315582 | 0.337244 | 0.435677 | 0.605284 | 0.741681 | 1.318843 | 4.285051 | 22.062730 | 57.377040 | 56.386263 | 24.527450 | 8.438884 | 1.883066 | 0.743882 | 0.402226 | 0.301533 | 0.240876 | 0.221622 | 0.215991 | 0.216957 | 0.220354 | 0.219742 | 0.218132 | 0.222189 | 0.249846 | 0.352045 | 0.540418 | 0.698487 | 1.339494 | 2.416450 | 3.564077 | 3.683568 | 3.907737 | 3.457542 | 3.304771 | 2.566194 | 2.116763 | 2.001778 | 2.274676 | 2.828314 | 2.959379 | 3.192227 | 2.766561 | 3.059832 | 3.050624 | 1.707368 | 0.964594 | 0.654010 | 0.497772 | 0.507292 | 0.549708 | 0.721847 | 1.081370 | 0.902357 | 0.821428 | 0.740216 | 0.535795 | 0.430812 | 0.337494 | 0.308739 | 0.340569 | 0.363883 | 0.389493 | 0.384437 | 0.373097 | 0.367910 | 0.344442 | 0.331808 | 0.303823 | 0.241372 | 0.236527 | 0.233573 | 0.222953 | 0.223442 | 0.220707 | 0.219300 | 0.240836 | 0.254354 | 0.267028 | 0.311455 | 0.346446 | 0.387459 | 0.465185 | 0.415854 | 0.335544 | 0.287576 | 0.253070 | 0.224876 | 0.218561 | 0.214848 | 0.215523 | 0.216096 | 0.215386 | 0.214965 | 0.219899 | 0.241710 | 0.251643 | 0.270751 | 0.278084 | 0.307088 | 0.314858 | 0.270156 | 0.227911 | 0.216976 |
| right central | 0.215715 | 0.214639 | 0.221110 | 0.235929 | 0.238728 | 0.220715 | 0.214737 | 0.231523 | 0.214694 | 0.217144 | 0.240856 | 0.216243 | 0.246336 | 0.276145 | 0.386852 | 0.270218 | 0.274878 | 0.257463 | 0.215439 | 0.218953 | 0.220992 | 0.259320 | 0.280680 | 0.227463 | 0.242263 | 0.376043 | 0.623150 | 0.874640 | 1.018498 | 1.294236 | 1.686242 | 1.615644 | 1.351353 | 1.012744 | 0.392391 | 0.271638 | 0.218226 | 0.217301 | 0.215402 | 0.220046 | 0.229319 | 0.267491 | 0.397062 | 0.291142 | 0.237207 | 0.215422 | 0.235056 | 0.265639 | 0.262865 | 0.238532 | 0.241232 | 0.267594 | 0.264265 | 0.226272 | 0.216187 | 0.215603 | 0.214633 | 0.233708 | 0.255158 | 0.273206 | 0.257662 | 0.239601 | 0.236161 | 0.221798 | 0.214768 | 0.219797 | 0.237644 | 0.248781 | 0.259717 | 0.270043 | 0.259811 | 0.235569 | 0.215443 | 0.232005 | 0.296459 | 0.422877 | 0.684093 | 0.567224 | 0.509660 | 0.453931 | 0.325359 | 0.365827 | 0.389294 | 0.375633 | 0.536091 | 0.483328 | 0.381400 | 0.403282 | 0.295340 | 0.321669 | 0.310426 | 0.269206 | 0.234808 | 0.215649 | 0.217611 | 0.217747 | 0.231860 | 0.228599 | 0.217078 | 0.216848 | 0.255321 | 0.335114 | 0.366380 | 0.379120 | 0.320183 | 0.263577 | 0.257215 | 0.250090 | 0.256340 | 0.255520 | 0.243808 | 0.247488 | 0.243472 | 0.222232 | 0.217978 | 0.214885 | 0.215156 | 0.216759 | 0.224740 | 0.246883 | 0.286276 | 0.279754 | 0.269038 | 0.249218 | 0.251091 | 0.242513 | 0.228693 | 0.215668 | 0.214953 | 0.214776 | 0.222484 | 0.233496 | 0.237339 | 0.248659 | 0.282521 | 0.344929 | 0.380783 | 0.396891 | 0.405215 | 0.502610 | 0.772551 | 0.966408 | 1.139709 | 0.543990 | 0.338225 | 0.266513 | 0.228213 | 0.226415 | 0.240074 | 0.254471 | 0.327336 | 0.394557 | 0.430288 | 0.507123 | 0.478989 | 0.530385 | 0.621456 | 0.700203 | 0.664531 | 0.678969 | 0.704936 | 0.725127 | 0.676282 | 0.551086 | 0.477147 | 0.521027 | 0.568966 | 0.590138 | 0.505752 | 0.437992 | 0.419956 | 0.445283 | 0.448005 | 0.424393 | 0.478200 | 0.661417 | 0.746760 | 0.813967 | 0.565116 | 0.472422 | 0.402248 | 0.358221 | 0.289086 | 0.257431 | 0.233290 | 0.240382 | 0.247439 | 0.253594 | 0.237951 | 0.231958 | 0.224143 | 0.238323 | 0.230804 | 0.218514 | 0.214926 | 0.214685 | 0.217329 | 0.228730 | 0.223957 | 0.216835 | 0.214629 | 0.214640 | 0.217192 | 0.218285 | 0.228750 | 0.232097 | 0.263882 | 0.328752 | 0.392075 | 0.388790 | 0.383836 | 0.314150 | 0.320063 | 0.290732 | 0.259076 | 0.235588 | 0.221423 | 0.214793 | 0.215736 | 0.217599 | 0.218039 | 0.215163 | 0.215111 | 0.214813 | 0.216814 | 0.220304 | 0.219110 | 0.221750 | 0.220189 | 0.227471 | 0.225850 | 0.217149 | 0.218759 | 0.224633 | 0.222690 | 0.215092 | 0.214660 | 0.216693 | 0.229216 | 0.279154 | 0.313210 | 0.295157 | 0.275216 | 0.280236 | 0.343614 | 0.412067 | 0.722009 | 0.930233 | 1.108980 | 1.696994 | 2.103751 | 1.394896 | 1.753718 | 1.190359 | 1.009955 | 1.215567 | 0.926628 | 0.786847 | 0.657237 | 0.473120 | 0.477324 | 0.474044 | 0.384422 | 0.373519 | 0.333538 | 0.334023 | 0.379023 | 0.419494 | 0.384909 | 0.342154 | 0.356665 | 0.369343 | 0.359205 | 0.304172 | 0.255292 | 0.284325 | 0.322900 | 0.302769 | 0.293836 | 0.298418 |
| left posterior | 0.866235 | 0.874038 | 1.133160 | 1.088532 | 0.770776 | 0.299423 | 0.214938 | 0.240473 | 0.262477 | 0.295909 | 0.260434 | 0.220294 | 0.223696 | 0.302094 | 0.357127 | 0.380517 | 0.405836 | 0.342698 | 0.289921 | 0.335898 | 0.474479 | 1.529393 | 2.441444 | 1.645826 | 1.817487 | 1.888499 | 0.987608 | 0.390673 | 0.236524 | 0.228811 | 0.261824 | 0.228708 | 0.247701 | 0.465400 | 0.789427 | 0.784430 | 0.657790 | 0.562451 | 0.454535 | 0.271170 | 0.214632 | 0.230285 | 0.298171 | 0.501841 | 0.514685 | 0.469396 | 0.371798 | 0.335055 | 0.338563 | 0.305648 | 0.276159 | 0.264201 | 0.244786 | 0.288681 | 0.348067 | 0.417752 | 0.664705 | 0.804370 | 1.294770 | 4.530322 | 8.343968 | 6.787139 | 3.612280 | 0.958272 | 0.397252 | 0.229125 | 0.219698 | 0.283894 | 0.325988 | 0.316905 | 0.282395 | 0.224712 | 0.216592 | 0.230468 | 0.219611 | 0.216192 | 0.216763 | 0.222025 | 0.238017 | 0.223237 | 0.214780 | 0.218216 | 0.235502 | 0.258393 | 0.361885 | 0.509601 | 0.553391 | 0.502394 | 0.404469 | 0.339489 | 0.303314 | 0.259519 | 0.247645 | 0.243582 | 0.253197 | 0.316509 | 0.369759 | 0.429777 | 0.392595 | 0.325643 | 0.306747 | 0.266054 | 0.242108 | 0.233997 | 0.267524 | 0.274086 | 0.272765 | 0.267282 | 0.298274 | 0.295511 | 0.287925 | 0.267783 | 0.257317 | 0.240673 | 0.235136 | 0.227728 | 0.225019 | 0.217814 | 0.215369 | 0.225930 | 0.228475 | 0.223862 | 0.216576 | 0.214763 | 0.222991 | 0.258064 | 0.316263 | 0.389066 | 0.437947 | 0.554747 | 0.760226 | 0.866129 | 0.716379 | 0.631292 | 0.627768 | 0.546742 | 0.442269 | 0.390618 | 0.341245 | 0.319215 | 0.294509 | 0.260379 | 0.318132 | 0.345387 | 0.370077 | 0.623185 | 0.854373 | 1.194125 | 1.132080 | 0.524252 | 0.384708 | 0.295892 | 0.241101 | 0.233630 | 0.234141 | 0.264777 | 0.321096 | 0.343733 | 0.355670 | 0.307070 | 0.270269 | 0.242265 | 0.253997 | 0.266886 | 0.277770 | 0.315442 | 0.354414 | 0.393920 | 0.484787 | 0.473876 | 0.422761 | 0.342849 | 0.269532 | 0.227952 | 0.215353 | 0.219752 | 0.243045 | 0.248486 | 0.241990 | 0.234267 | 0.218582 | 0.215389 | 0.265866 | 0.427771 | 0.654642 | 0.850501 | 1.356096 | 1.712049 | 1.522406 | 0.613912 | 0.326209 | 0.249631 | 0.230305 | 0.226397 | 0.224418 | 0.222618 | 0.228963 | 0.248243 | 0.251792 | 0.259945 | 0.294584 | 0.302977 | 0.303291 | 0.281699 | 0.238209 | 0.219270 | 0.214658 | 0.224708 | 0.226086 | 0.223590 | 0.220720 | 0.217541 | 0.217149 | 0.218935 | 0.234430 | 0.277597 | 0.324712 | 0.334288 | 0.293657 | 0.227200 | 0.214632 | 0.260238 | 0.377023 | 0.521509 | 0.530967 | 0.448524 | 0.317769 | 0.239236 | 0.215263 | 0.227952 | 0.279060 | 0.375201 | 0.525744 | 0.726512 | 0.801073 | 0.829725 | 0.608060 | 0.623575 | 0.527821 | 0.411304 | 0.348233 | 0.341482 | 0.322787 | 0.309439 | 0.284084 | 0.274198 | 0.265933 | 0.253563 | 0.228321 | 0.222834 | 0.235805 | 0.250378 | 0.252916 | 0.245805 | 0.231629 | 0.229624 | 0.228349 | 0.216359 | 0.220550 | 0.232033 | 0.262575 | 0.286496 | 0.368561 | 0.452695 | 0.593502 | 0.650804 | 0.766658 | 0.713377 | 0.760118 | 0.620661 | 0.592012 | 0.521871 | 0.464494 | 0.480767 | 0.484056 | 0.386704 | 0.331255 | 0.320365 | 0.307337 | 0.289973 |
| right posterior | 0.273800 | 0.239814 | 0.259520 | 0.238568 | 0.252415 | 0.228961 | 0.220382 | 0.288962 | 0.421264 | 0.512034 | 0.374563 | 0.520008 | 1.003829 | 2.320306 | 2.330480 | 0.772106 | 0.942421 | 1.470030 | 0.560559 | 0.346265 | 0.220459 | 0.227972 | 0.302914 | 0.408327 | 0.590569 | 0.553170 | 0.342689 | 0.215529 | 0.235971 | 0.242387 | 0.229168 | 0.230032 | 0.266255 | 0.221862 | 0.217912 | 0.221393 | 0.218734 | 0.215992 | 0.215051 | 0.215122 | 0.214626 | 0.219223 | 0.248209 | 0.288801 | 0.353958 | 0.453307 | 0.862054 | 1.103575 | 1.118859 | 0.689737 | 0.393966 | 0.279134 | 0.249276 | 0.230758 | 0.283621 | 0.325991 | 0.419115 | 0.696514 | 0.991877 | 1.477861 | 1.770204 | 1.036207 | 0.761372 | 0.417608 | 0.251703 | 0.215218 | 0.316471 | 0.457384 | 0.490115 | 0.506635 | 0.536670 | 0.829957 | 2.509846 | 11.678973 | 92.538745 | 362.097572 | 255.017245 | 187.254750 | 51.553712 | 17.708871 | 7.585190 | 2.825100 | 1.038312 | 0.617905 | 0.534369 | 0.618410 | 0.763776 | 1.130423 | 2.012596 | 3.613383 | 6.434796 | 5.762306 | 3.876211 | 2.966350 | 2.133899 | 1.644631 | 1.200223 | 0.780211 | 0.775346 | 1.182467 | 1.548611 | 1.430009 | 1.039747 | 0.881362 | 0.786954 | 0.550389 | 0.357195 | 0.307915 | 0.311579 | 0.377141 | 0.522896 | 0.505323 | 0.566642 | 0.527082 | 0.458582 | 0.402212 | 0.322359 | 0.283881 | 0.280313 | 0.269988 | 0.284964 | 0.280510 | 0.264302 | 0.263260 | 0.271942 | 0.284357 | 0.321049 | 0.359651 | 0.423603 | 0.548204 | 0.827282 | 1.128974 | 1.920274 | 2.158660 | 2.108971 | 2.039126 | 2.228005 | 2.217490 | 1.840438 | 1.054398 | 0.700038 | 0.449143 | 0.339305 | 0.291695 | 0.261903 | 0.236483 | 0.219568 | 0.222510 | 0.239344 | 0.257100 | 0.305231 | 0.329470 | 0.360598 | 0.462462 | 0.538355 | 0.665005 | 1.205498 | 1.546884 | 1.760962 | 1.833961 | 2.109592 | 2.094769 | 1.356318 | 0.637198 | 0.426283 | 0.339266 | 0.307437 | 0.307267 | 0.312872 | 0.402471 | 0.604454 | 0.911876 | 1.599928 | 2.752989 | 3.935302 | 5.909757 | 7.912595 | 10.624304 | 11.191666 | 8.982400 | 6.614203 | 2.833236 | 1.239216 | 0.619562 | 0.354773 | 0.255972 | 0.233001 | 0.219997 | 0.228475 | 0.242269 | 0.291943 | 0.328139 | 0.412203 | 0.473705 | 0.426225 | 0.342286 | 0.317086 | 0.286813 | 0.304678 | 0.358408 | 0.456250 | 0.556557 | 0.790022 | 1.213274 | 1.838727 | 2.737779 | 2.779512 | 2.634902 | 3.814808 | 5.504044 | 6.776140 | 8.122424 | 11.166856 | 25.255584 | 45.452146 | 28.707548 | 10.620024 | 6.038401 | 2.289606 | 0.720163 | 0.341965 | 0.231173 | 0.217569 | 0.214739 | 0.219481 | 0.215010 | 0.215492 | 0.229955 | 0.261278 | 0.251702 | 0.248571 | 0.239533 | 0.220708 | 0.219560 | 0.220114 | 0.232739 | 0.296364 | 0.372245 | 0.413305 | 0.523495 | 0.552582 | 0.502226 | 0.409824 | 0.341381 | 0.300675 | 0.291451 | 0.276769 | 0.262446 | 0.247918 | 0.251781 | 0.238253 | 0.230385 | 0.234343 | 0.232449 | 0.229921 | 0.225614 | 0.221387 | 0.219522 | 0.222120 | 0.215435 | 0.214647 | 0.216380 | 0.214865 | 0.225718 | 0.277386 | 0.400351 | 0.448495 | 0.597150 | 0.928409 | 1.198954 | 0.992219 | 0.857024 | 0.839642 | 1.050656 | 0.864501 | 0.622784 | 0.528721 | 0.486721 | 0.406572 | 0.322668 |
| all electrodes | 0.324427 | 0.242628 | 0.216300 | 0.215322 | 0.237869 | 0.222402 | 0.223976 | 0.229849 | 0.214763 | 0.227055 | 0.278512 | 0.349388 | 0.252409 | 0.220578 | 0.214862 | 0.278930 | 0.698254 | 1.051557 | 8.148150 | 4.089363 | 2.984586 | 1.358218 | 0.414447 | 0.230015 | 0.253750 | 0.921466 | 1.510245 | 1.687974 | 1.606217 | 1.181828 | 0.915841 | 0.399717 | 0.251278 | 0.226292 | 0.286746 | 0.434774 | 0.890143 | 2.543722 | 2.899907 | 2.810032 | 0.879441 | 0.408654 | 0.267233 | 0.232061 | 0.214653 | 0.216307 | 0.217089 | 0.217624 | 0.240802 | 0.254781 | 0.248309 | 0.277609 | 0.339814 | 0.520428 | 0.544596 | 0.374643 | 0.288894 | 0.246344 | 0.245676 | 0.298488 | 0.341863 | 0.265798 | 0.233386 | 0.217420 | 0.214880 | 0.215837 | 0.255402 | 0.470502 | 0.621511 | 0.523086 | 0.364326 | 0.285464 | 0.373085 | 0.526837 | 0.691084 | 0.947735 | 1.878754 | 4.629830 | 6.852614 | 4.724339 | 2.979381 | 1.815516 | 1.333868 | 0.791132 | 0.420646 | 0.291444 | 0.275283 | 0.255982 | 0.270904 | 0.373482 | 0.527468 | 0.646684 | 0.709279 | 0.724014 | 0.864162 | 1.121559 | 0.814298 | 0.722624 | 1.044340 | 1.742002 | 2.316656 | 2.349759 | 1.418704 | 1.282553 | 1.303590 | 0.836387 | 0.366813 | 0.250764 | 0.223760 | 0.226883 | 0.216984 | 0.214736 | 0.215178 | 0.214816 | 0.215666 | 0.220607 | 0.241919 | 0.309456 | 0.333600 | 0.743135 | 1.425845 | 1.701988 | 1.387270 | 0.851455 | 0.562928 | 0.504715 | 0.264051 | 0.215429 | 0.225829 | 0.237972 | 0.266887 | 0.335821 | 0.374106 | 0.322849 | 0.246761 | 0.214784 | 0.232537 | 0.284744 | 0.312045 | 0.308293 | 0.320804 | 0.287881 | 0.245669 | 0.215379 | 0.224163 | 0.232924 | 0.232866 | 0.224665 | 0.215084 | 0.214852 | 0.215409 | 0.215759 | 0.221911 | 0.223597 | 0.217210 | 0.214767 | 0.224865 | 0.264149 | 0.277705 | 0.285386 | 0.256493 | 0.238740 | 0.227637 | 0.226678 | 0.220458 | 0.224348 | 0.240492 | 0.329536 | 0.480306 | 0.486879 | 0.339452 | 0.299701 | 0.315938 | 0.292507 | 0.244896 | 0.239123 | 0.246944 | 0.259099 | 0.282253 | 0.306971 | 0.293097 | 0.356092 | 0.414663 | 0.440554 | 0.401499 | 0.323115 | 0.279422 | 0.302711 | 0.273409 | 0.263213 | 0.273165 | 0.301158 | 0.326043 | 0.319394 | 0.292014 | 0.266846 | 0.229975 | 0.216319 | 0.214657 | 0.214900 | 0.215126 | 0.219895 | 0.235056 | 0.296375 | 0.481221 | 0.641901 | 0.955443 | 1.154472 | 1.342964 | 1.345955 | 1.563363 | 2.476562 | 5.962804 | 7.881977 | 18.627995 | 22.216810 | 22.928201 | 13.332664 | 4.568497 | 1.956940 | 0.818670 | 0.396346 | 0.274437 | 0.224248 | 0.215803 | 0.218829 | 0.224564 | 0.250194 | 0.281248 | 0.358206 | 0.506625 | 0.603433 | 0.594658 | 0.485308 | 0.462787 | 0.451079 | 0.451452 | 0.495447 | 0.586681 | 0.871566 | 1.683690 | 2.188160 | 1.826909 | 1.203375 | 0.904734 | 0.866906 | 0.560535 | 0.309648 | 0.241511 | 0.234299 | 0.253644 | 0.292523 | 0.317992 | 0.280161 | 0.272721 | 0.263295 | 0.265108 | 0.239163 | 0.215735 | 0.222483 | 0.233510 | 0.274267 | 0.306586 | 0.354946 | 0.471244 | 0.461219 | 0.404178 | 0.347775 | 0.281204 | 0.252283 | 0.264909 | 0.242592 | 0.244877 | 0.242121 | 0.255146 | 0.244194 | 0.242960 | 0.227539 | 0.219813 | 0.214667 |

Searchlight, spatiotemporal cluster permutation test

No significant clusters observed.
